# Deducing ensemble dynamics and information flow from the whole-brain imaging data

**DOI:** 10.1101/2022.11.18.517011

**Authors:** Yu Toyoshima, Hirofumi Sato, Daiki Nagata, Manami Kanamori, Moon Sun Jang, Koyo Kuze, Suzu Oe, Takayuki Teramoto, Yuishi Iwasaki, Ryo Yoshida, Takeshi Ishihara, Yuichi Iino

## Abstract

Recent development of large-scale activity imaging of neuronal ensembles provides opportunities for understanding how activity patterns are generated in the brain and how information is transmitted between neurons or neuronal ensembles. However, methodologies for extracting the component properties that generate overall dynamics are still limited. In this study, the results of time-lapse 3D imaging (4D imaging) of head neurons of the nematode *C. elegans* were analyzed by hitherto unemployed methodologies.

By combining time-delay embedding with independent component analysis, the whole-brain activities were decomposed to a small number of component dynamics. Results from multiple samples, where different subsets of neurons were observed, were further combined by matrix factorization, revealing common dynamics from neuronal activities that are apparently divergent across sampled animals. By this analysis, we could identify components that show common relationships across different samples and those that show relationships distinct between individual samples.

We also constructed a network model building on time-lagged prediction models of synaptic communications. This was achieved by dimension reduction of 4D imaging data using the general framework gKDR (gradient kernel dimension reduction). The model is able to decompose basal dynamics of the network. We further extended the model by incorporating probabilistic distribution, resulting in models that we call gKDR-GMM and gKDR-GP. The models capture the overall relationships of neural activities and reproduce the stochastic but coordinated dynamics in the neural network simulation. By virtual manipulation of individual neurons and synaptic contacts in this model, information flow could be estimated from whole-brain imaging results.

## Introduction

How brain achieves various integrative and instructive functions is a major question in neurobiology. Needless to say, brain is composed of interconnected web of neurons, and transmission of activity across neurons is fundamental for the functions. Activity of each neuron is transmitted to other neurons through chemical and electrical synapses, whose signs, efficiency and dynamic properties determine the activity of receiver neurons. Also, each neuron typically receives inputs from multiple presynaptic neurons and interactions between these inputs are also important for determining the response of the postsynaptic neuron. Therefore, the network shape and synaptic properties are the major determinants of information flow.

Although mammalian cerebral cortex is as complex as being composed of tens of billions of neurons, recent advance of technologies is starting to reveal connectomes in various levels, from micro-connectome at synaptic resolution to macroscopic connectome between brain regions (Elam et al., 2021; Kasthuri et al., 2015; Oh et al., 2014). In parallel, a number of techniques to monitor neuronal activities are being actively utilized, such as multi-unit electrical recording, electroencephalogram (EEG), Magnetoencephalography (MEG), functional magnetic resonance imaging (fMRI) and calcium imaging (Breakspear, 2017). A difficulty in all these analyses is that cerebral cortex is layered and therefore the structure is essentially three-dimensional, to which deep brain nuclei add further complexity. Along with the huge number of neurons, the structural complexity hampers full resolution understanding of the whole neural circuit.

In smaller animals, quasi-whole-brain activity measurements at single-cell resolution have been made possible. One of the most successful subjects is zebrafish larvae, where the brain of agar-embedded larval animals have been observed during fictive swimming, for example by light-sheet microscopy, revealing various activity groups and their relationships in behavioral contexts (Ahrens et al., 2012; Naumann et al., 2016).

Another model animal well-suited for these kinds of analyses is the nematode *C. elegans. C. elegans* is a small experimental animal whose body length is about 1 mm in young adults, which allowed reconstruction of the whole nervous system, which include exactly 302 neurons in an adult hermaphrodite, by electron microscopy, resulting in the full connectome data (Cook et al., 2019; White et al., 1986; Witvliet et al., 2021). A number of laboratories have performed whole-brain imaging in *C. elegans* (Kato et al., 2015; Nichols et al., 2017; Skora et al., 2018; Uzel et al., 2022). Global neuronal activities were shown to form a manifold in the state space, where typical behaviors such as forward, backward and turn are represented (Kato et al., 2015). Also, whole-brain imaging in freely moving animals have been performed, and activities that are related to specific behaviors were found (Nguyen et al., 2016; Venkatachalam et al., 2016). Furthermore, researchers have succeeded in predicting behavior based on whole brain activities (Hallinen et al., 2021; Susoy et al., 2021).

All these attempts so far described the whole brain states and described the transition of states, and found ensemble activities related to sensory input, behavior, or preparation of behavior. However, the question of HOW these activity patterns are generated have not been directly addressed. Namely, still lacking are the methodologies to understand how network dynamics are assembled and how network structure and synaptic connections generate the dynamics observed. Because whole-brain imaging in *C. elegans* provides high-dimensional data (time-series data of around 100 neurons), it is essential to extract relevant information while reducing the dimensionality of the data. In addition, due to difficulty in observing and annotating all neurons in the brain, methods for assembling the partially observed data across different individual animals is desired.

Here we present approaches to decompose and reconstruct the whole nervous system dynamics, based on *C. elegans* 4D imaging results. These approaches successfully extracted common dynamics across animals, as well as individual differences. Roles of individual neurons in the information flow could also be estimated by synapse-based models.

## Results

### Correlation between neuronal activities

In this study, a *C. elegans* adult hermaphrodite that express the calcium reporter Yellow Cameleon (nuclear localized YC2.60) in all neurons was placed in a narrow channel in the microfluidic device called olfactory chip (Chronis et al., 2007) and stimulated with a periodic switch between two different concentrations of NaCl under a fluorescence microscope. The focal plane was scanned up and down to obtain 3D fluorescence images (hereafter called 4D imaging) (Hirose et al., 2018; Tokunaga et al., 2014; Toyoshima et al., 2020, 2016). A total of 24 samples of whole-brain activity were obtained (Materials and Methods). As shown in Figure 1A and Supplementary File 1, there are several prominent characteristics in the activities. First, as the animals were given sensory stimuli (changes in NaCl concentration applied to the nose tip), there were small groups of neurons that apparently responded to the sensory stimuli. Other groups of neurons showed synchronized activation and inactivation. These groups did not show obvious synchrony with the sensory stimuli, and therefore considered as spontaneous activities. These observations are consistent with previous reports (Kato et al., 2015).

**Figure 1.**
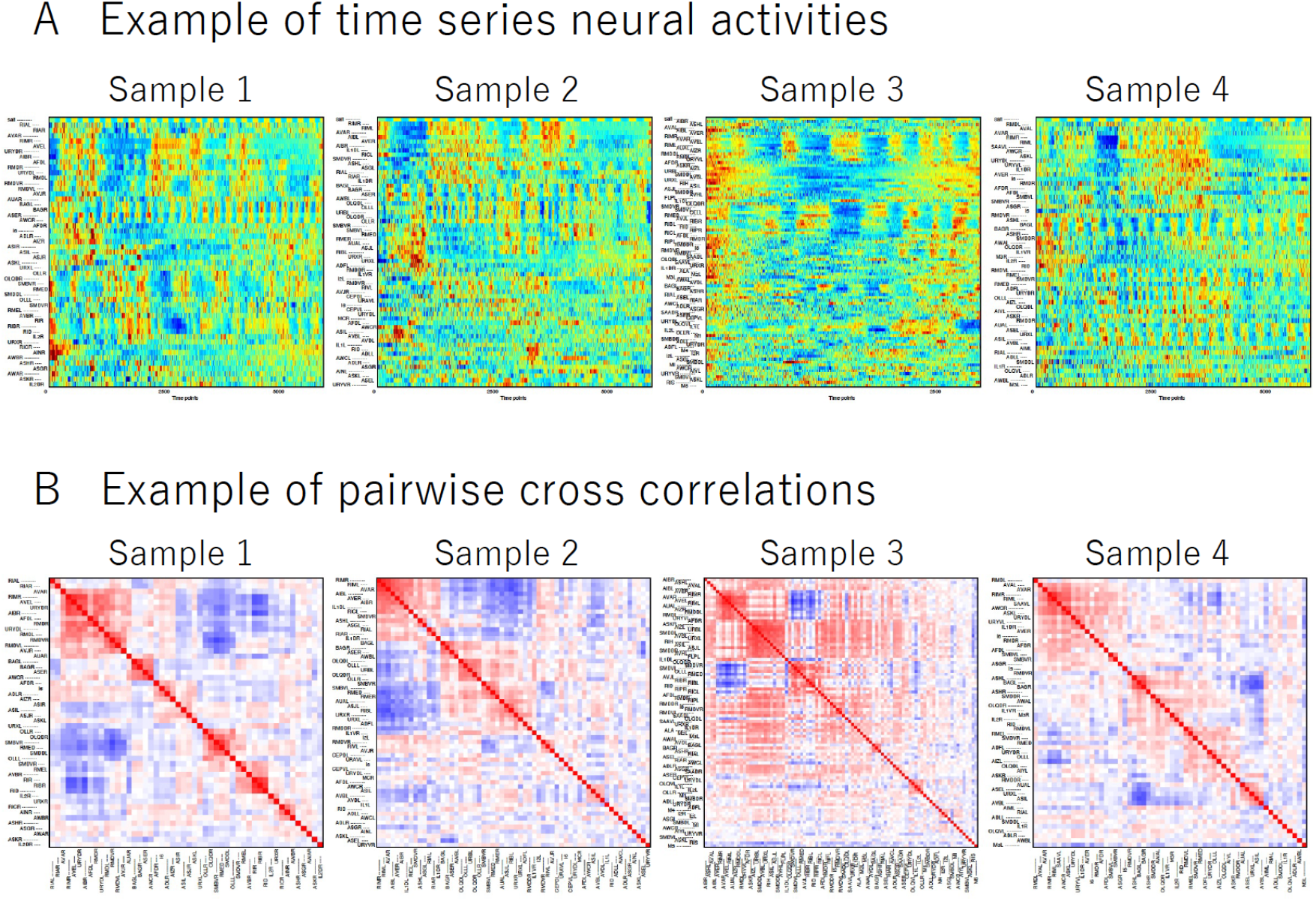
Data obtained by 4D imaging. (A) Activity time series of head neurons obtained by 4D imaging. Activity of each neuron in the scaled fluorescence ratio of YFP over CFP is shown in pseudocolor. Top row shows salt concentration changes applied to the nose tip of the animals. Following rows indicate neuronal activity profiles. Each row represents one neuron, whose order was determined by hierarchical clustering based on activity cross-correlations. Note that only a subset of head neurons, which differ between samples, are shown in each panel, because some of the neurons were unobserved (*e*.*g*. too dim) or unannotated. (B) Pairwise cross correlation of head neuron activities obtained by 4D imaging. Red color shows positive correlation and blue color shows negative correlation. Examples of four samples are shown. For all samples, see Supplementary File 1.

To further clarify this, cross-correlations in neuronal activities were determined in all pairwise combinations of neurons (Figure 1B). It was noted that the size of each correlated group differed considerably between individual samples. Here, it is also apparent that in many cases, in each sample there is at least one major group of neurons with correlated activity, and there is often another correlated group which is negatively correlated with the first group. Examination of member neurons reveals that these groups correspond to the well-known groups of neurons related to reversal and forward movements of the animals, namely, AVA, AVE and RIM neurons etc. in the first group (called group A here), and RIB, RID and RME neurons etc. in the second group (called group B). Other groups that were correlated across samples included [OLLL, OLLR, OLQDL, OLQDR] and [I2L, I2R, MCL], and left and right members of BAG, RIA, RMDV and SMDD classes (Figure 1-Figure supplement 1).

**Figure 1–figure supplement 1.**
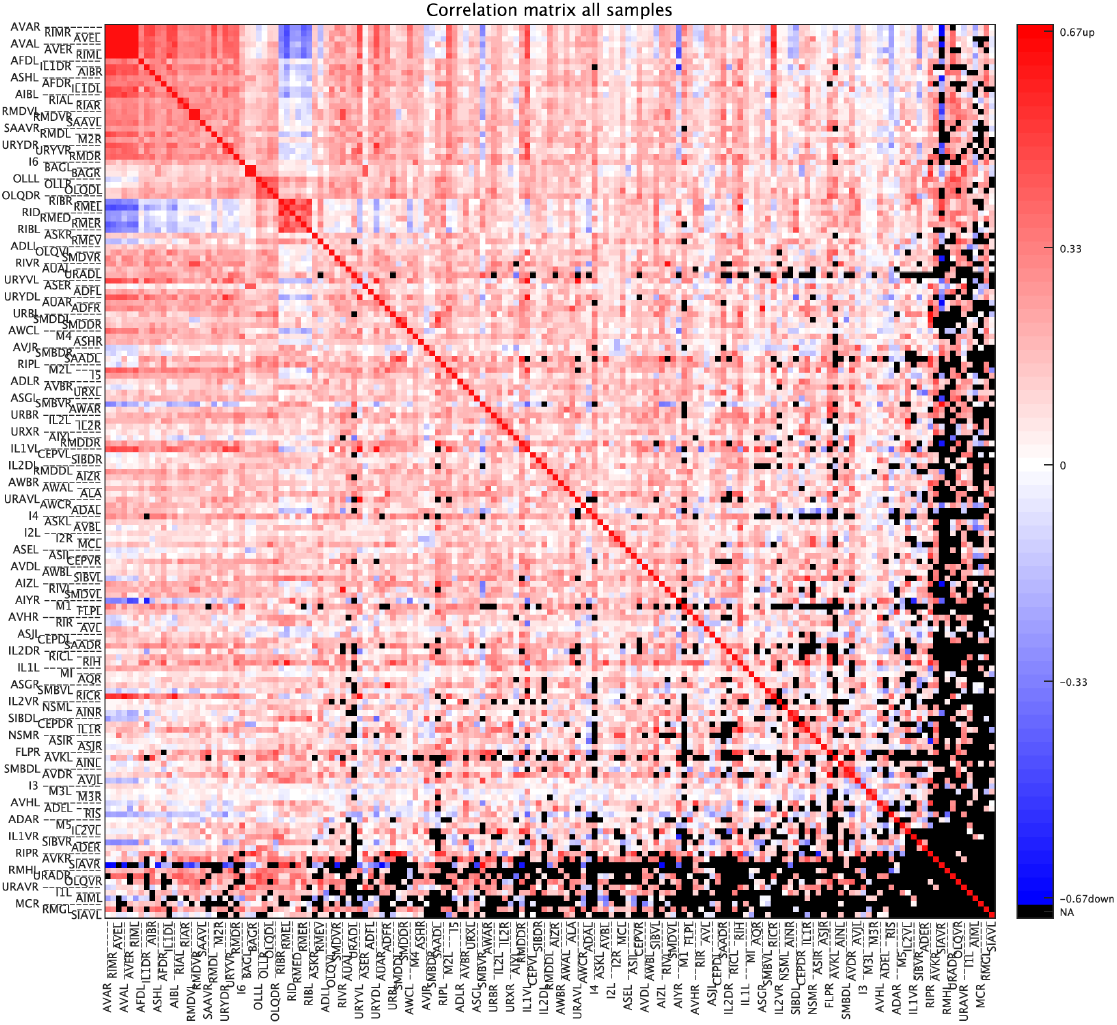
Commonly correlated neurons. Pairwise cross-correlation averaged across samples are shown. Hierarchical clustering was performed to arrange the neurons based on p-values. Red colors show positive and blue colors show negative correlations. Several large and small correlated groups are seen, two of which show prominent negative correlation with each other. Pairs of neurons that were never co-observed in any sample were filled black.

### Common and individual dynamics are revealed by time-delay embedding and independent component analysis

Next, we focus on the overall dynamics of the nervous system captured by the whole-brain activity data. In the previous studies, principal component analysis (PCA) was applied to the whole-brain activity data of individual samples (Hallinen et al., 2021; Kato et al., 2015). This approach is straightforward but has some drawbacks.

The first problem is that the principal component axes obtained from a sample cannot be applied to that of the other samples directly, suggesting that the observed dynamics in the PCA space cannot be compared between samples. This problem can be addressed by applying the analysis to the whole-brain activity data from multiple samples at once.

The second problem is that PCA requires the orthogonality between principal component axes. This requirement is derived from mathematical procedures and lacks biological validity, so the obtained components might be distorted. This requirement can be relaxed by using Independent Component Analysis (ICA) instead of PCA. A similar approach is applied for the descriptor of the posture of the worm (Gyenes and Brown, 2016).

The third problem is that the method is useful for decomposing the whole-brain dynamics into the sum of the instantaneous components, but is not useful for extracting “temporal motifs” of the neural activities from the dynamics. A temporal motif is a latent temporal pattern that appears repeatedly in neural activities. Time delay embedding (TDE) methods have been applied to model complex ecological systems (Sugihara and May, 1990) and neural dynamics (Tajima et al., 2015). The method combining TDE with ICA was applied to extract behavioral motifs from time series of worms’ posture (Ahamed et al., 2021). A similar approach should be useful to extract neural motifs from whole-brain activity data.

Here we introduced a method we called TDE-RICA (Reconstruction ICA with Time-Delay Embedding) and applied the method to the whole-brain activity data from multiple samples simultaneously. RICA is one of the variations of ICA methods and penalized with reconstruction cost so that the captured components and weights can reproduce the original data (Le et al., 2011).

Because RICA cannot handle missing values, 94 neurons of 10 samples (with no missing values) were selected from the whole dataset of 177 neurons of 24 samples (with missing values). Since our method of annotation for neuron identity is imperfect (Toyoshima et al., 2020), several neurons cannot be identified in a sample, resulting in missing values.

We set the time-delay for embedding as 300 time steps (approximately 60 seconds) so that the motifs of the neural activity capture the large-scale dynamics such as switching between forward and backward command neurons (Kato et al., 2015). In addition, the number of time steps corresponds well with the approximately 60-second period of sodium chloride stimulation used in the experiment. The whole-brain activity is measured during 6000 time steps (approximately 1200 seconds), and the embedded data is a matrix *X* that contains (94 [neurons] × 300 [delay time steps]) × (5701 [time steps] × 10 [samples]) elements (see Methods). We also set the number of components as 14, which was the minimum number to capture the neural response to the sodium chloride stimulation. Applying TDE-RICA to the selected dataset returns the independent components *M* (*M* = *W*^*T*^*X*^*T*^, matrix *M* contains 14 [components] × (94 [neurons] × 300 delay time steps]) elements) and their weights *W* (containing (5701 [time steps] × 10 [samples]) × 14 [components] elements) that represent the original data well (Figure 2, for all 10 selected samples, see Supplementary File 2).

**Figure 2.**
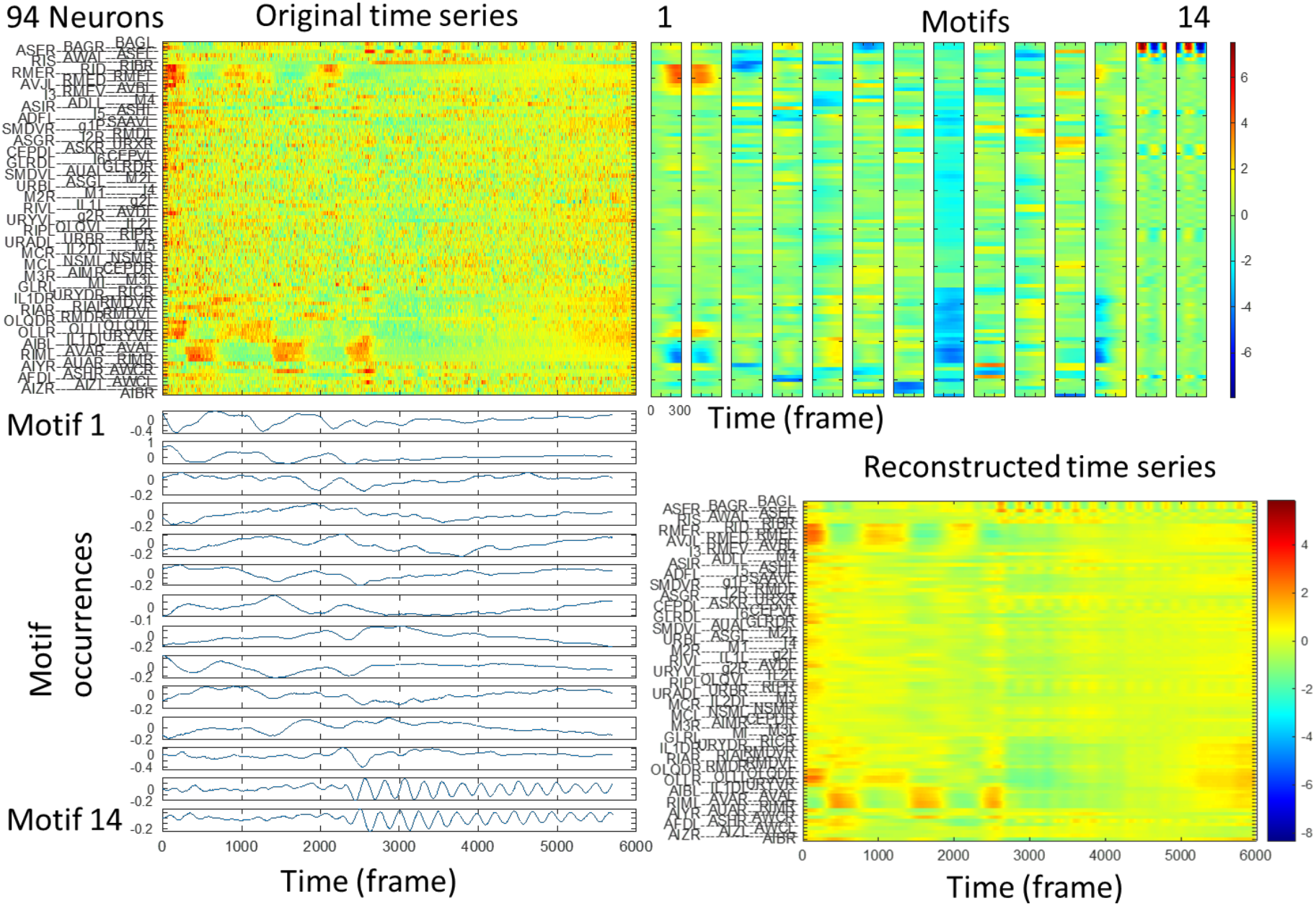
TDE-RICA captures the motifs of neural activities (Top left) Example of activity time series of selected 94 neurons, as shown in Figure 1A. TDE-RICA decomposes the time series into the product of the motifs (top right) and the motif occurrences (bottom left). The product of the motifs and its occurrences reproduce the captured features of the original time course (bottom right). (Top right) The motifs of neural activities obtained by TDE-RICA. Each motif consists of the activity of 94 neurons over 300 time points. The color indicates the relative intensity of individual neural activity in each motif. The motifs are common between samples. (Bottom left) The motif occurrences obtained by TDE-RICA. The occurrences are different between samples. (Bottom right) The reconstructed time series. The color range is the same as that of the upper left panel.

The independent components are common between samples and were regarded as motifs of neural activities. The weights are different between samples and were regarded as the occurrences of the motifs in the sample.

In order to complete the missing values, we extended the results obtained by TDE-RICA with a matrix factorization. Matrix factorization can handle missing values and complete the values based on the existing values. We already have the matrices of motifs and their occurrences obtained from the partial dataset without missing values. We can extend the matrix of motifs to all 177 neurons and the matrix of motif occurrences to all 24 samples so that the product of the motifs and their occurrences reproduces the observed neural activities. Thus we obtain the full set of motifs (*M*^*all*^containing 14 [components] × (177 [neurons] × 300 [delay time steps]) elements) and their occurrences (*W*^*all*^, containing (5701 [time steps] × 24 [samples]) × 14 [components] elements) (Figure 2 —figure supplement 1).

The motifs and their occurrences obtained by TDE-RICA are preserved in those obtained by matrix factorization. The analyzed results of the full set of motifs and the occurrences (see Supplementary File 3) were basically the same as that of the motifs and the occurrences obtained by TDE-RICA (see Supplementary File 2). Therefore we describe the results of the full set of motifs and occurrences.

**Figure 2–figure supplement 1.**
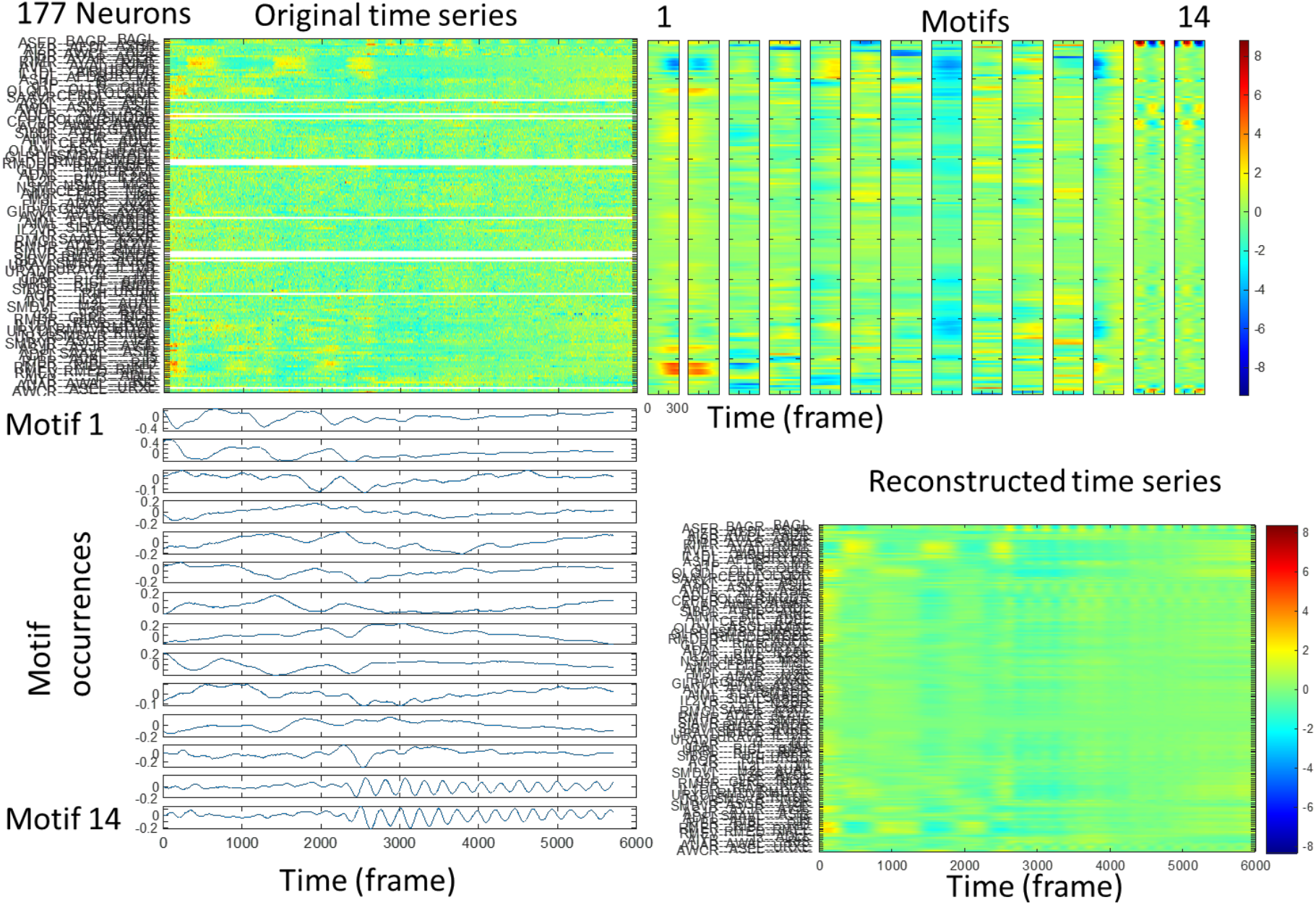
Completing missing value by TDE-RICA with matrix factorization. (Top left) Activity time series of all 177 neurons in the same sample as in Figure 1A. The activity of several neurons are missing and shown in white (top left). The missing values are completed in the reconstructed time series by TDE-RICA with matrix factorization (bottom right). (Top right) The full set of motifs of neural activities obtained by TDE-RICA with matrix factorization. (Bottom left) The full set of motif occurrences. (Bottom right) The reconstructed time series. The missing values in the top left panel are completed well.

We found that the last two motifs (13th and 14th) in the 14 motifs have large weights on neurons that respond to the sensory stimulation including ASE and BAG sensory neurons (Figure 3, Figure 3— figure supplement 1). The other neurons including downstream interneurons were less weighted, suggesting that the activities of sensory neurons might affect the downstream neurons more indirectly and independently than we have expected (see below).

**Figure 3.**
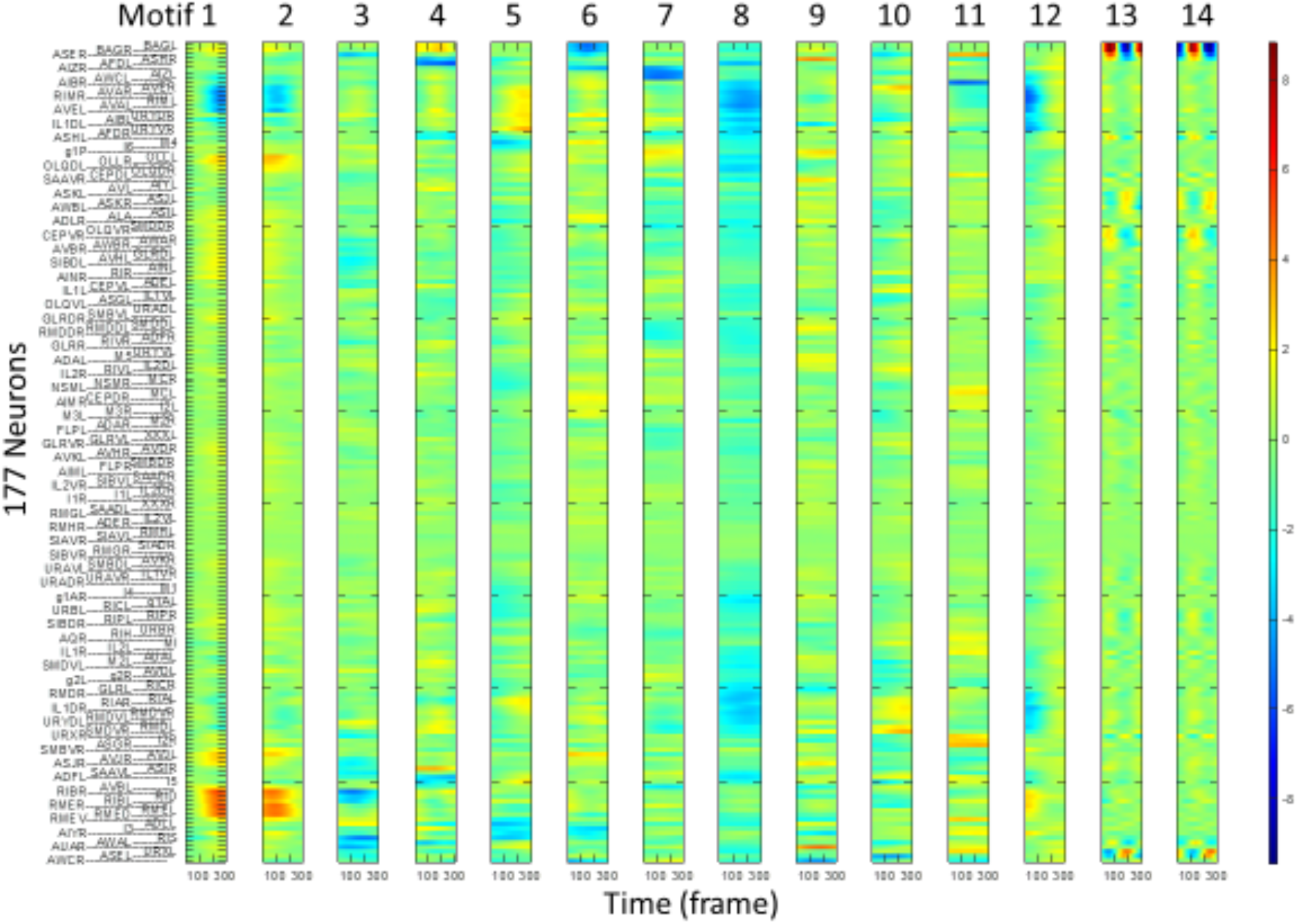
Temporal motifs of neural activities obtained by TDE-RICA with matrix factorization. Fourteen motifs are shown. Each motif consists of the activity of 177 neurons over 300 time points. The color indicates the relative intensity of individual neural activity in each motif.

We also found that the first and second motifs have large weights on neurons that govern the forward and backward movement of worms including interneurons AVA, AIB, RIM, AVB, and motor neurons RME (Figure 3, Figure 3—figure supplement 2). The activities of neurons governing forward movement (AVB and RME, group B) and backward movement (AVA, AIB, RIM, group A) were inversely correlated, suggesting mutual inhibition between these two groups (Kawano et al., 2011). The mechanosensory neurons including OLQ and OLL were positively correlated to the neurons governing forward movement, suggesting that the worms might have moved forward and hit the wall of the microfluidic device in which they were held.

In the 8th motif, the neurons for mechanosensation and backward movement had large negative weights, and the neurons for forward movement were less weighted (Figure 3, Figure 3—figure supplement 3). This result indicates that the relationships between these three groups of neurons are variable and can change in a context-dependent manner. In the 8th motif, the thermosensory neurons AFD and their downstream interneurons RIA and RMD were positively correlated to the neurons for backward movements (Mori and Ohshima, 1995; Ohnishi et al., 2011). In addition, the other thermosensory neurons AWC and downstream interneuron AIZ were positively correlated in the 7th motif. These results may indicate that these motifs capture the unintended activities of the thermotaxis circuit.

We further analyzed the relationships between motifs using the occurrences of the motifs. We calculated the cross-correlations of the occurrences in each sample and averaged them across samples (Figure 4).

**Figure 4.**
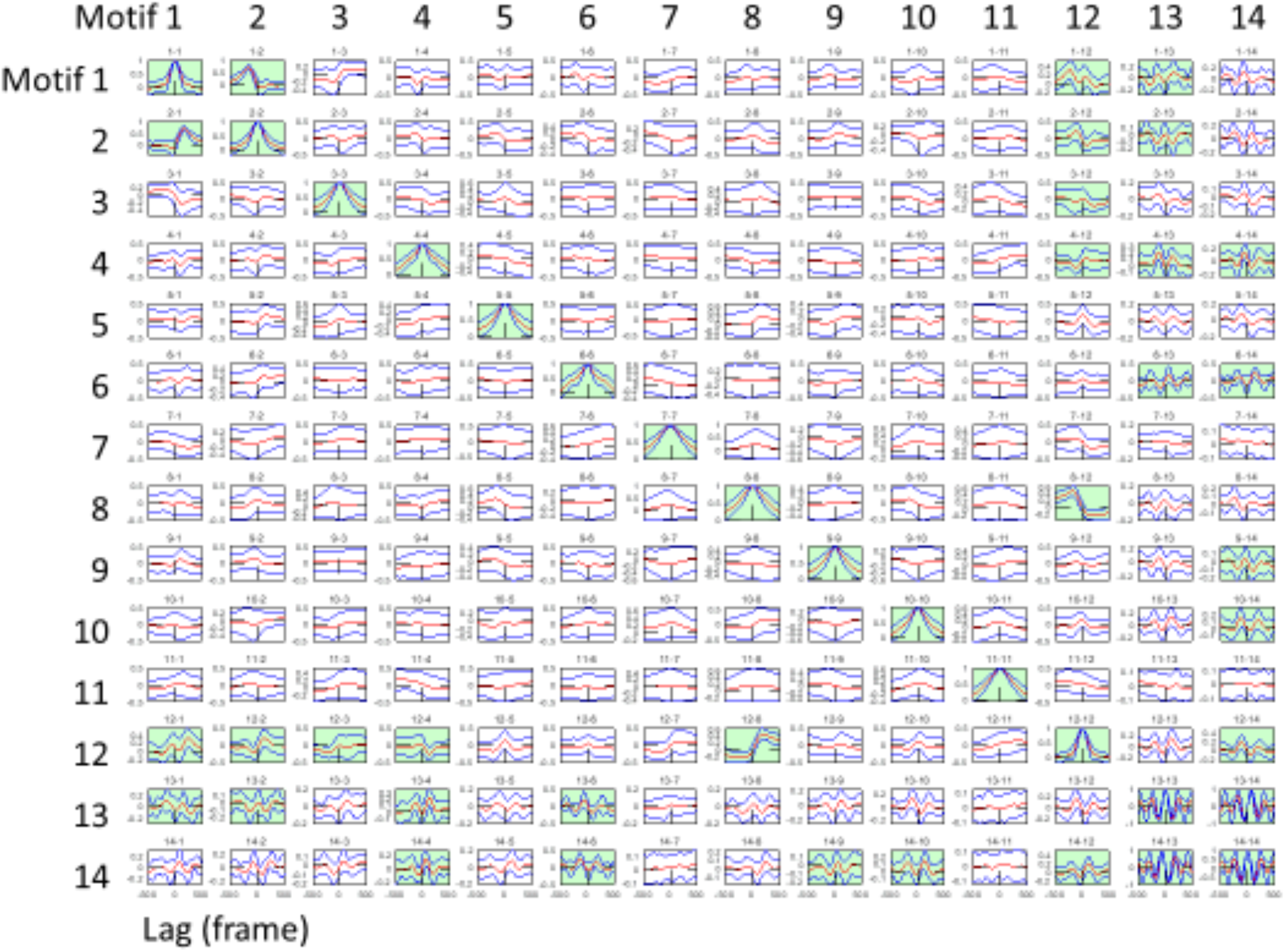
Cross-correlation function of motif occurrences. Cross-correlation functions of motif occurrence are averaged across samples. The red line indicates the mean and the blue lines indicate the mean ± standard deviation. Pairs are highlighted in green if the mean + standard deviation is smaller than 0 for any lag or if the mean - standard deviation is larger than 0 for any lag.

We found a prominent cross-correlation between the 13th and 14th motifs that represent the sensory responses. In order to compare the neural dynamics between different samples on the latent space of the motifs, we plot the occurrences of the 13th motif with that of the 14th motif (Figure 5). The 14th motif always preceded the 13th motif. In the phase diagram, their trajectories formed a circle during the period with sensory stimulation, which degenerated to the origin during the period without stimulation. These results suggest that the sensory motifs successfully represent the presence or absence of stimuli and capture the response to the periodic stimulus. These trajectories were common between samples, indicating that these sensory dynamics are common across animals.

**Figure 5.**
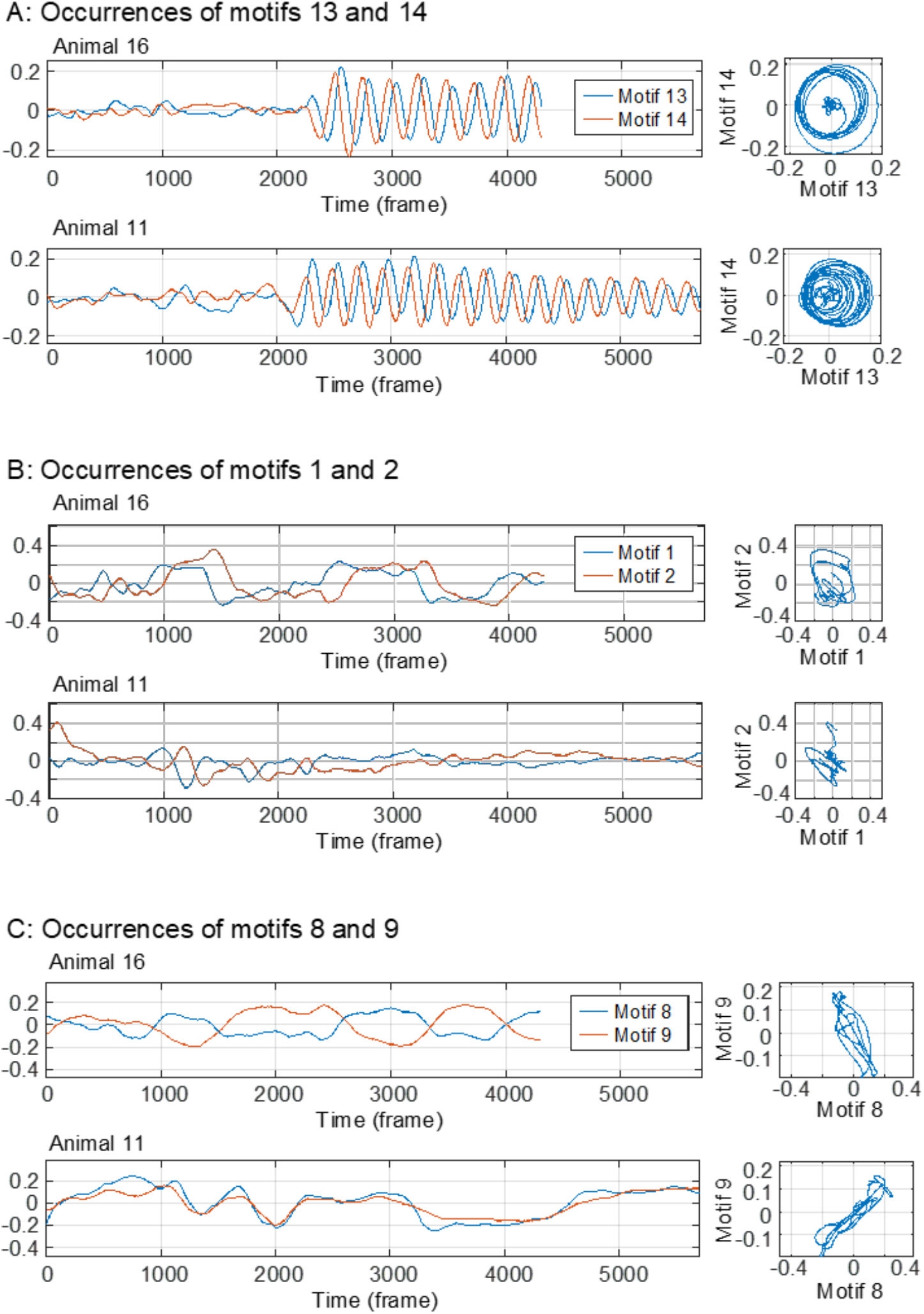
Common features and individual differences of motif occurrences in the latent space. (A) Left: The occurrence of motif 13 (red) and motif 14 (blue) in sample 16 (upper) and sample 11 (lower). Right: The phase diagram of motif 13 and motif 14 in sample 16 (upper) and sample 11 (lower). (B) Left: The occurrence of motif 1 (red) and motif 2 (blue) in the same samples as (A). Right: the phase diagram of motif 1 (red) and motif 2 (blue) in the same samples as (A). (C) Left: The occurrence of motif 8 (red) and motif 9 (blue) in the same samples as (A). Right: the phase diagram of motif 8 (red) and motif 9 (blue) in the same samples as (A).

We found another prominent cross-correlation between the first and the second motifs that represent the forward and backward movements (Figure 5). The first motif always preceded the second motif. In the phase diagram, the trajectories formed distorted circles. These features are common between samples, but the shapes of trajectories were different between samples. This may indicate the individual differences in quantitative dynamics of command interneurons and motor neurons between animals.

We also found an interesting relationship between the 8th and 9th motifs. These motifs were positively correlated in a sample, but they negatively correlated in another sample (Figure 5). These relationships were held throughout the recording of these samples. This result may suggest the individual differences in qualitative relationships of neurons between animals, although it has been believed in general that there are few individual differences in neural activity in *C. elegans* because of their uniform genetic background and stereotypic developmental process.

**Figure 3–figure supplement 1.**
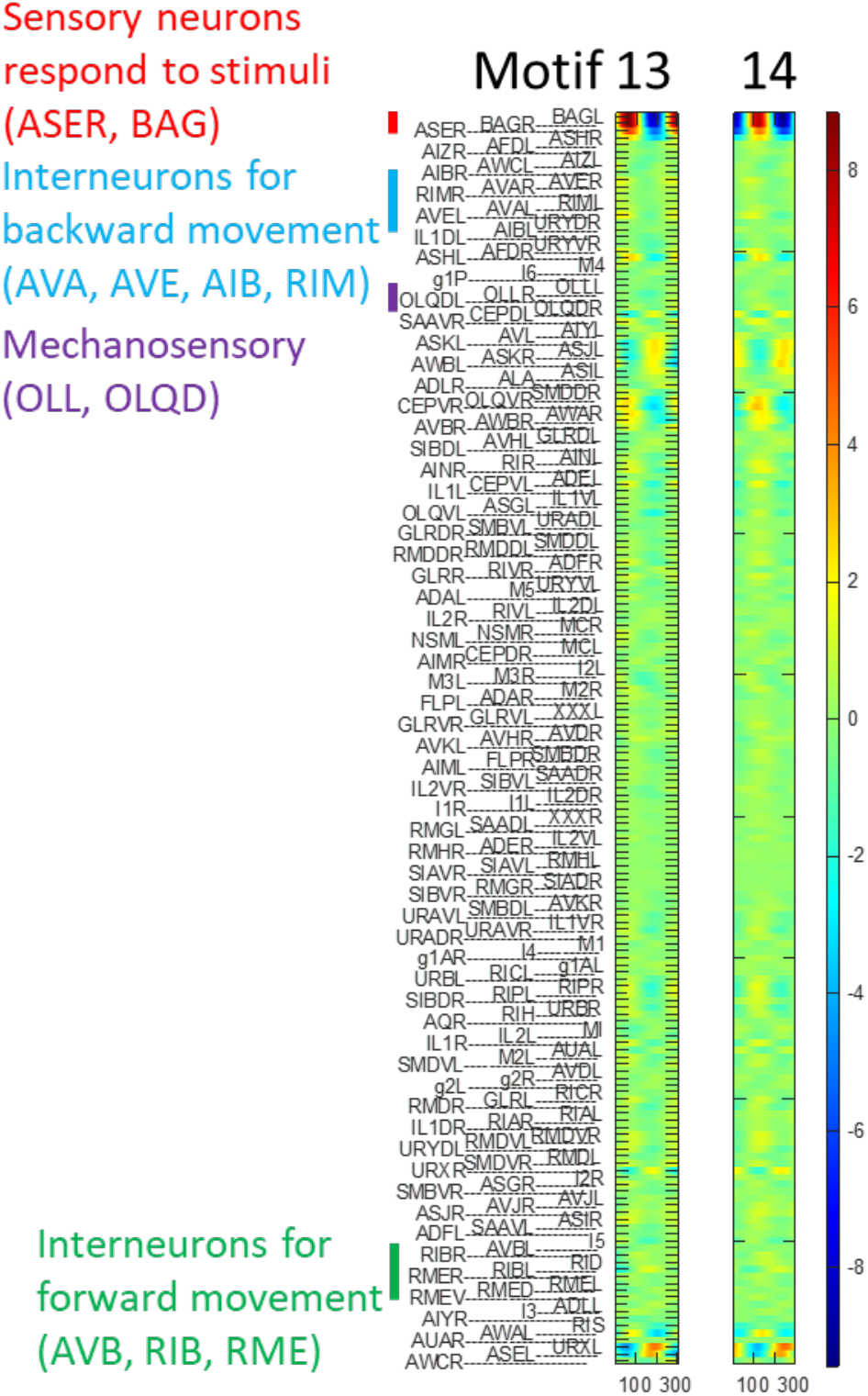
Motifs that correspond to the sensory responses.

**Figure 3–figure supplement 2.**
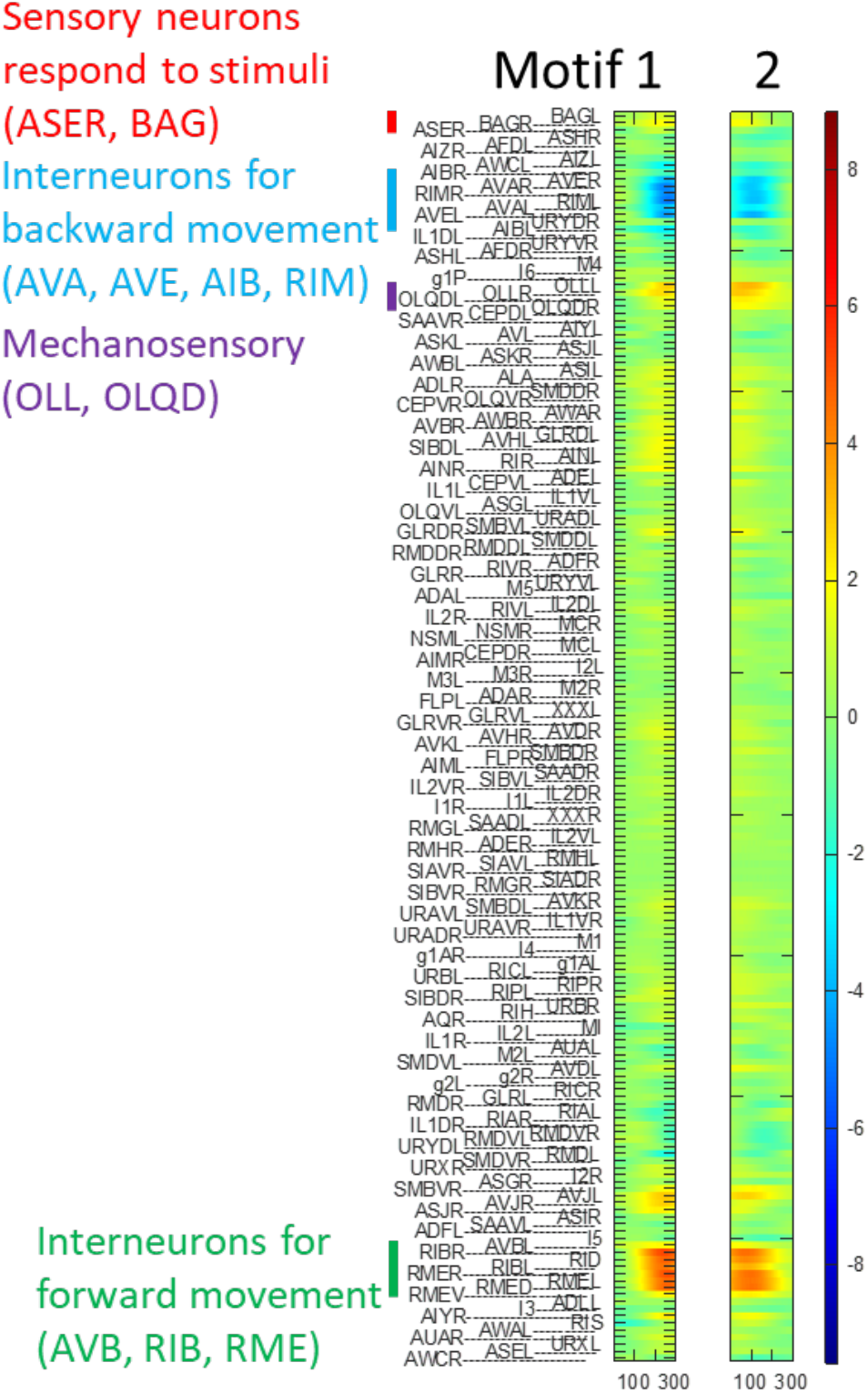
Motifs that correspond to the forward and backward movements.

**Figure 3–figure supplement 3.**
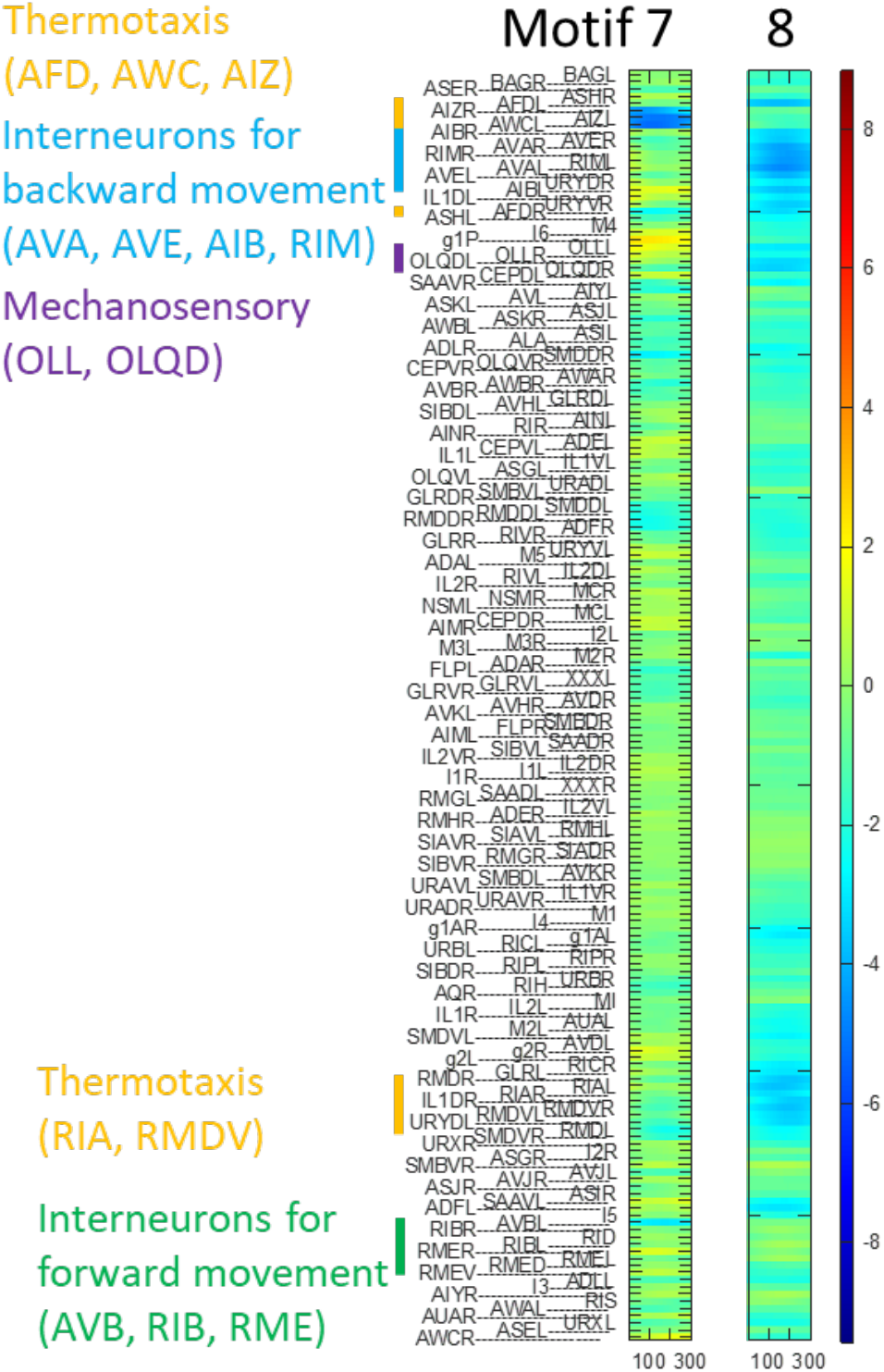
Motifs that correspond to the thermotaxis circuit.

### Synapse-based model reproduces overall dynamics of brain network

The TDE-RICA described above successfully decomposed the overall dynamics of the nervous system into separate dynamics of subsets of neurons as motifs. We next sought to understand the synaptic basis of the dynamics. To do this, we utilized the connectome information in addition to the observed neuronal activities. As described in Introduction, synaptic connections between neuron pairs throughout the nervous system (connectome) have been fully described. Based on this set of information, we constructed models in a bottom-up manner.

In these models, we consider only annotated neurons in which non-random time-series activity was observed, and unlike TDE-RICA each sample was treated separately (see Materials and Methods). First, one of the neurons in the selected sample (sample *j*) is set as a “target” neuron (neuron *i*). We then select neurons which send synaptic inputs to the target neuron via chemical or electrical synaptic connections (“explanatory neurons for target *i* “, 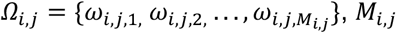 depicts the number of neurons presynaptic to target *i* in sample *j*). Our assumption is that the activity of the target neuron is determined by inputs from presynaptic neurons, and therefore we take into consideration that chemical synapses are directed but electrical synapses are not. Time-delay embedding is applied to the explanatory neurons to utilize the past time series of the neurons to predict target neuron activity ahead of time (*y*_*i,j*_(*t* + *Δt*)) (Figure 6, see Methods for details). Note that previous activity of the target neuron itself was also included in the explanatory variables (Figure 6).

**Figure 6.**
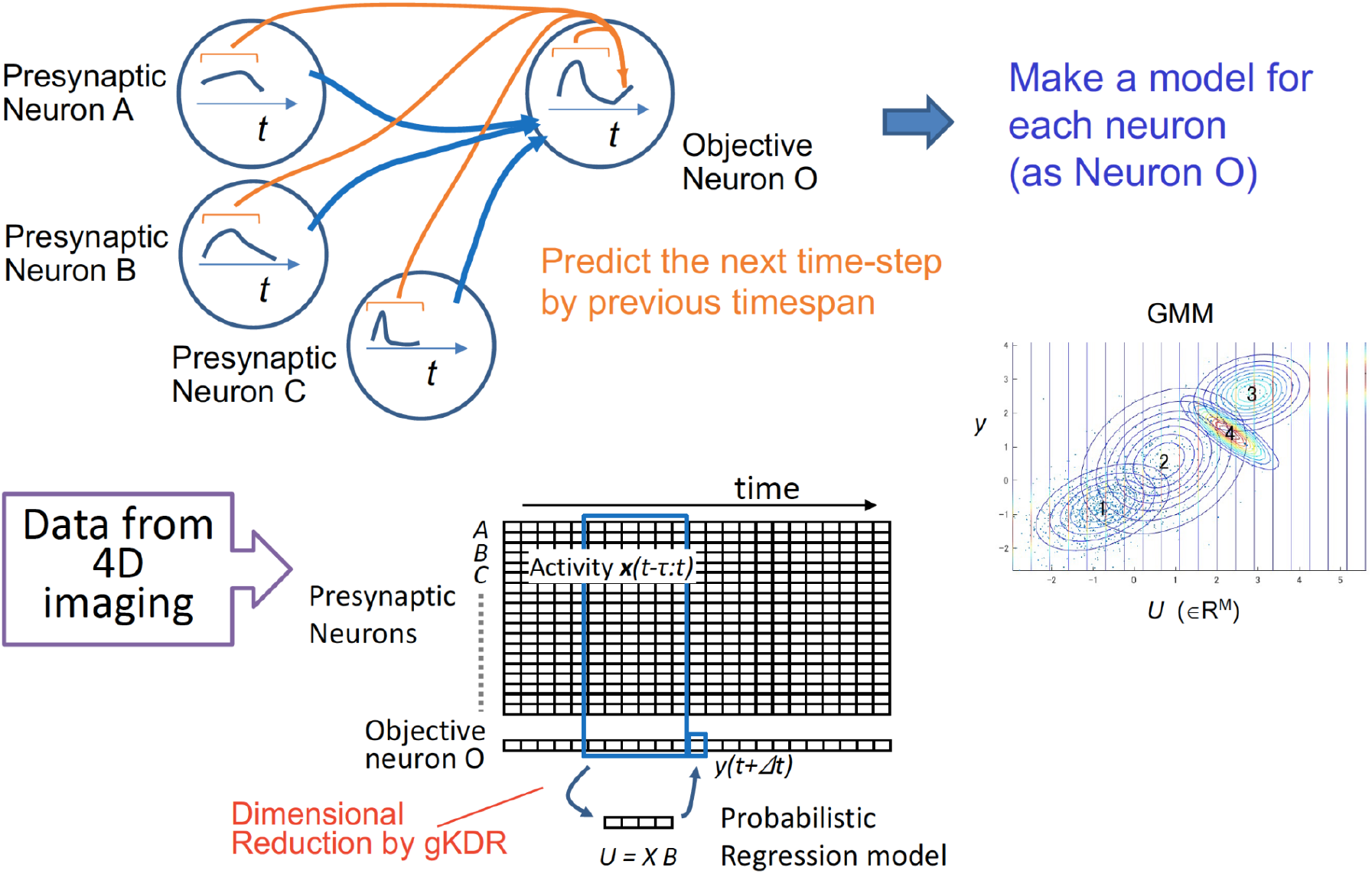
Overview of gKDR-GMM. The model learns to predict activity of the target neuron ahead of time (*y*(*t* + *Δt*)) from previous activity of presynaptic neurons (*X*) (blue lines represent physical connections). For this sake, gKDR reduces the dimension of presynaptic activities *X* and generates *K*-dimensional values *U* representing *X*, making sure that sufficient information for predicting *y* is included in *U*. GMM models the joint probability of (*U, y*) as a weighted sum of Gaussian distribution which is fitted to the real data. Conditional probability of *y* is determined from the GMM model and used for prediction. See Methods and Figure 13 for details.

We first employed dimensional reduction to reveal “important explanatory neurons and time-points” using gradient kernel dimension reduction (gKDR) (Fukumizu et al., 2004; Fukumizu and Leng, 2014)). gKDR determines the *K*-dimensional subspace (ℜ^*^) of the space spanned by the explanatory variables (activity patterns of presynaptic neurons). The subspace is defined by a dimension reduction matrix *B*_*i,j*_, so that *U*_*i,j*_(*t*) = *<u>X</u>*_*i,j*_(*t*)*B*_*i,j*_, *<u>X</u>*_*i,j*_(*t*) being embedded activities of explanatory neurons (see Methods). The aim of gKDR is to find *B*_*i,j*_ so that *U*_*ij*_(*t*) ∈ ℜ^*^ includes sufficient information for predicting *y*_*i,j*_(*t* + *Δt*) (see Methods for details). One prominent characteristic of gKDR is that the best *B*_*i,j*_ is reached by solving a convex optimization problem and can be obtained by eigenvalue calculation. Therefore, once hyperparameters *K* and embedding rank etc., have been determined, the solution is unique for a given data set and there is no worry of reaching a non-optimal locally minimal solution.

As described in Figure 1 and Figure 1-figure supplement 1, *C. elegans* nervous system includes several groups of neurons that show synchronized activities, showing positive cross-correlations among their members. However, most of the activity changes of these groups are not periodic and are irregular, except for responses to regular sensory stimuli that were applied by the experimenter. This characteristic of the neuronal ensemble corresponds to the stochastic nature of the behaviors. Namely, “when” a group of neurons are activated is unpredictable, but once the neurons are activated, they do so “at the same time”, driving the robust behavior. To reproduce this pattern, we employed a probabilistic model. In this model, called gKDR-GMM, the relationship between the *K*-dimensional explanatory variables obtained by gKDR and the target variable were described by the joint distribution and modeled as Gaussian mixture probabilistic distribution. Here, we have a GMM model (GMM_*i,j*_) for each target neuron *i* in each sample *j*. For simulation, we determine the conditional distribution of *y*_*ij*_(*t* + *Δt*) on *U*_*ij*_(*t*), namely *P*(*y*_*ij*_(*t* + *Δt*)|*U*_*ij*_(*t*)), from joint probability *P*(*U*_*ij*_(*t*), *y*_*ij*_(*t* + *Δt*)) modeled in GMM_*i,j*_, and select *y*_*ij*_(*t* + *Δt*) (Figure 6).

The gKDR-GMM model is in fact a collection of these models (*B*_*i,j*_ and GMM_*i,j*_, *i* = 1,2, …, *M*_*j*_ ; *M*_*j*_ depicts the number of observed neurons in sample *j*), each of which describes the synaptic input to the target neuron *i*. The meta-model composed of *M*_*j*_ models describe the dynamics of the whole nervous system, because it represents the activation rule of all neurons in the system. The model is then run for simulation (here called freerun simulation). Whole nervous system simulation is performed simply by repeating prediction of activity of *y*_*i,j*_(*t* + *Δt*) from activity data of neurons up to time *t* and repeating the process for *i* = 1 to *M* to generate 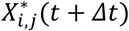 (* stands for estimated values) and repeating the process to proceed through time (*t* + *Δt, t* + 2*Δt*, ….).

Optimal hyperparameters for the gKDR-GMM model were searched for (Figure 7—figure supplement 1, Experimental Methods). First, presynaptic neurons were chosen as those that send direct chemical synaptic output to or have a gap junction with the target neuron (called direct links). However the reproduction of ensemble neural dynamics was not very good. This was expected because, as noted earlier, we cannot observe and annotate all neurons in the head of *C. elegans*, and typically, half of the neurons are missing from the data as described above (maximal number of annotated non-random neurons was 110 out of ∼190 head neurons). Therefore in another option, only for missing presynaptic neurons, neurons with another step of synaptic connections were included, namely presynaptic neurons for the missing neuron that was presynaptic to the target (called indirect links). The results of modeling including indirect links were generally better (Figure 7—figure supplement 1C-H; average numbers of “presynaptic” neurons in the connectome data, direct link option and indirect link option were 15.1, 6.8 and 21.1, respectively; Figure 7—figure supplement 2). Next, embedding ranks and time steps was searched for and *k*′ = 30 and *τ*′ = 10 was adopted (approximately 60 secs of time span was used for prediction of *y*_*i,j*_(*t* + *Δt*), Figure 7—figure supplement 1A-B). For *K*, namely the reduced number of dimensions after gKDR, the results were improved by increasing the dimension up to *K* = 3 or 4 but did not improve further (Figure 7—figure supplement 1C-E). Therefore, *K*=3 to 5 was adopted. Because slightly different results were obtained with different *K*, we hereafter created models with these different values. The number of gaussians were also varied and two Gaussians (*κ* = 2) showed the best performance in general, which we used hereafter (Figure 7—figure supplement 1F-H).

**Figure 7.**
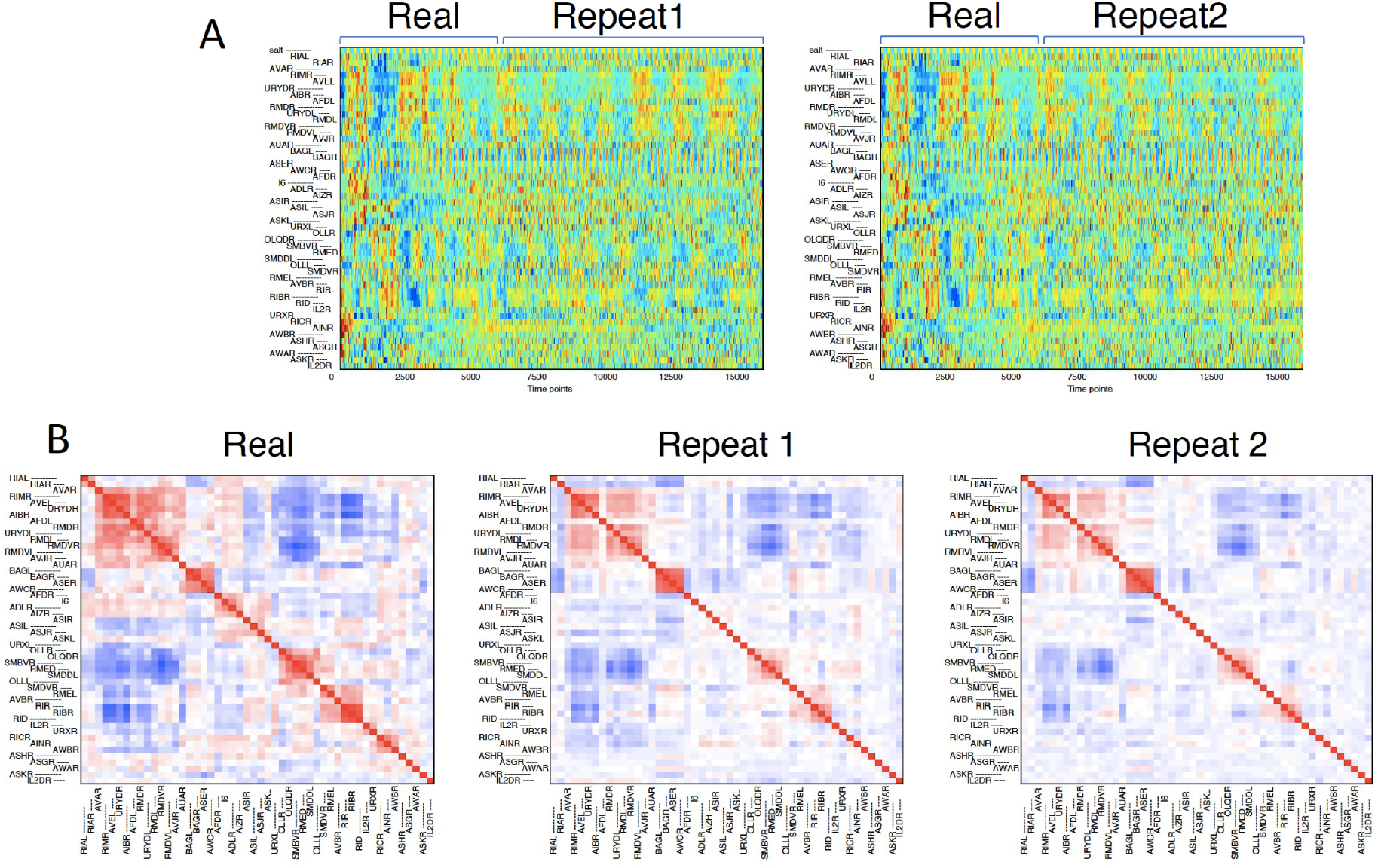
Prediction results by gKDR-GMM. (A) Neural activities obtained from freerun simulation by gKDR-GMM. The time-span depicted as “real” is actual activity data which is identical to Figure 1A, while the following time span depicted as “Repeat 1/2” displays results of simulation. Results of two simulation runs are shown. The order of neurons is the same as Figure 1A. (B) Cross-correlation of neural activities. Left panel shows the cross-correlation of actual activities identical to Figure 1B, shown for comparison. Middle and right panels are cross correlation of simulation results shown in A. Color coded as red for positive and blue for negative correlations. See Supplementary Files 4A-C for results of all samples.

Another popular and powerful approach for multidimensional nonlinear regression is Gaussian process. We also tested modeling the relationships between *U* and *y* by Gaussian process instead of GMM (called gKDR-GP). The resulting score of its performance was similar to gKDR-GMM (Figure 7—figure supplement 1I-K, Supplementary Files 4A-C, 5A-C). These results demonstrate the robustness of our approach. For the sake of simplicity and interpretability, we mainly used gKDR-GMM hereafter.

Figure 7A shows example results of the simulation by the gKDR-GMM model, and all results are shown in Supplementary Files 4A-C. Results of two independent repeats of simulations are shown from the same model for the same sample. Because gKDR-GMM is a probabilistic model, the results are different each time. Still, the overall ensemble patterns are well reproduced.

**Figure 7–figure supplement 1.**
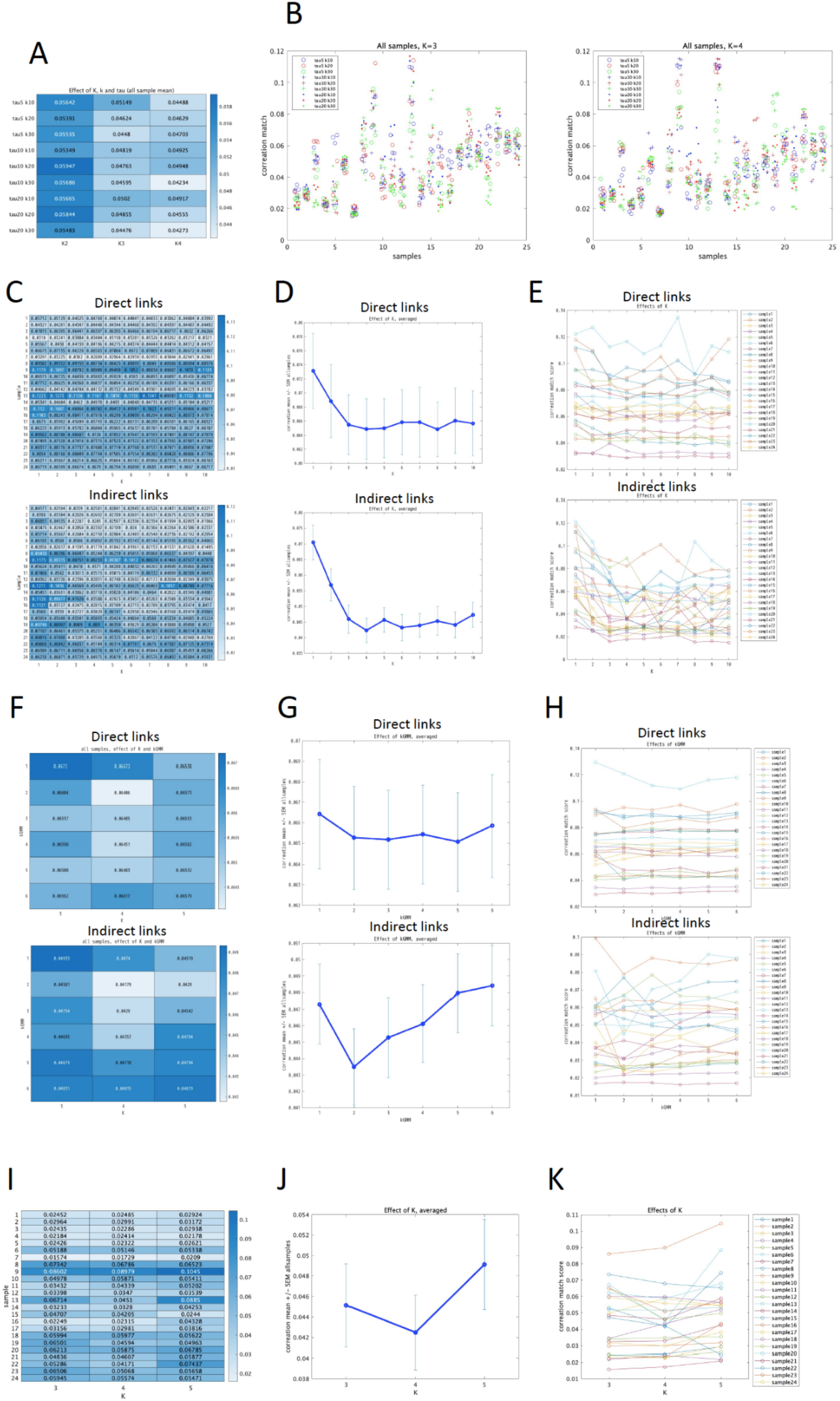
Determination of hyperparameters. **(A-B)** Determination of *k*′ and *τ*′ for gKDR-GMM. **(A)** gKDR-GMM models were generated for each sample with different embedding parameters *k*′ and *τ*′, and reduction dimension *K*, and freerun simulation was run for three times. Simulation results were evaluated by mean square difference between cross-correlation matrix of neuron activities between real data and simulation results. Averages of all samples and all repeats are shown in this figure. *κ* = 3 for all tests. **(B)** Same results in A are shown for each sample separately. **(C-E)** Determination of *K* for gKDR-GMM. **(C)** gKDR-GMM models were generated for each sample with different reduction dimension *K*, and freerun simulation was run for three times. *k*′ = 30, *τ*′ = 10, *κ* = 3 for all tests. In “direct links” only neurons that send direct synaptic input to the target neuron were considered “presynaptic”, while in “indirect links”, for missing presynaptic neuron, neurons presynaptic to the missing neuron were also included in “presynaptic neuron”. **(D)** Same results as in (C), but all samples were averaged and SEM are shown. **(E)** Line plot representation of (D). **(F-H)** Determination of hyperparameters *κ* for gKDR-GMM. **(F)** gKDR-GMM models were generated for each sample with different number of Gaussians, *κ*, for GMM as well as reduction dimension *K* for kGMM. “direct links” and “indirect links” are the same as in (C). **(G)** Same results as (F) but average of all samples and all *K* are shown as well as SEM. **(H)** Line plot representation of (G). **(I-K)** Determination of hyperparameters *K* for gKDR-GP. **(I)** gKDR-GP models were generated for each sample with different reduction dimension *K*, and freerun simulation was run for three times. *k*′ = 30, *τ*′ = 10 for all tests. Only results for “indirect links” are shown. **(J)** Same results as (I) but average of all samples and all *K* are shown as well as SEM. **(K)** Line plot representation of (J).

**Figure 7–figure supplement 2.**
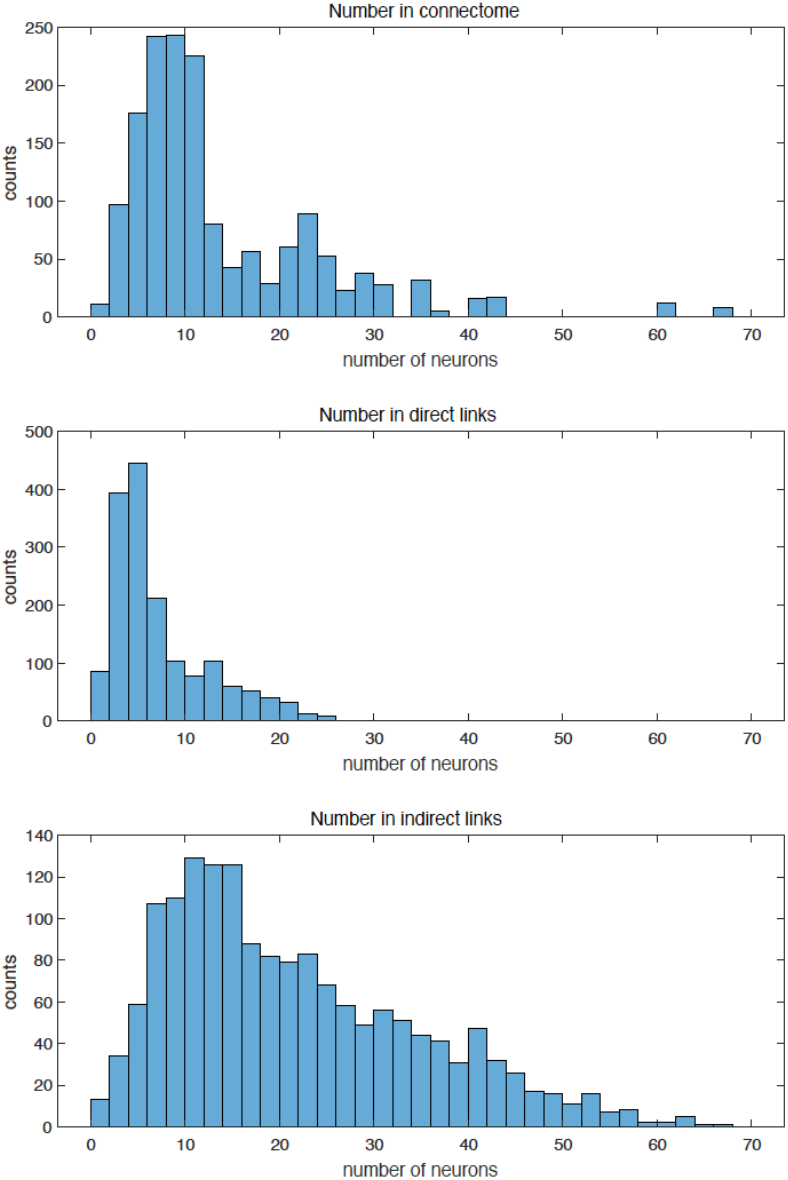
Numbers of “presynaptic neurons” used for prediction models. **(A)** Numbers of (directly connected) presynaptic neurons in the connectome data. **(B)** Numbers of directly connected presynaptic neurons that were observed and annotated in 4D imaging and included in the models. **(C)** Numbers of “indirectly linked” presynaptic neurons. These include directly linked presynaptic neurons and those presynaptic to unobserved or unannotated presynaptic neurons. All target neurons in all samples were included in (A)-(C).

The reproduction capacity of the model was further evaluated. In common practice, the reproduction capacity is often assessed by cross validation, where the difference (or similarity) between predicted data and real data is assessed. However, as described above, our basic assumption is that the nervous system is stochastic in nature - the changes in the activities are irregular and stochastic. Therefore, we do not necessarily expect the activities of model neurons change at the same time as the real neurons in long-term simulations. Rather, we expect that the relationships between neurons observed in the real nervous system are maintained in the simulation based on the model, and that sensory information is properly transmitted in the model nervous system.

Therefore, correlation of activity between neurons was first evaluated after the simulation by the model. An example is shown in Figure 7B and all results are also shown in Supplementary Files 4A-C. As described earlier, there are correlated groups of neurons in the real imaging data. Some of the relationship was maintained in the simulation, though the absolute value of correlation was generally lower and some correlated groups are missing in the model. Second, to evaluate not only the correlation but also temporal relationships between neurons, lagged cross-correlation of all pairwise combinations of neurons were evaluated. Figure 8A,B and Supplementary File 6A,B show that the lags that provided best correlation between each pair were significantly maintained in the simulation in most samples. Further, TDE-RICA was performed as described earlier, by using the same motif matrix, to compare the dynamic relationship of major groups (Supplementary File 7). The prominent relationship between motifs 13 and 14 were maintained in most samples, and characteristic behavior between motifs 8 and 9 as shown in Figure 5 were also reproduced in the freerun simulation. Finally, because the animals were stimulated by periodic change in salt concentrations during the 4D imaging, we evaluated the periodic signals in each neuron by fourier analysis and periodic shift of the time series data (Figure 8C and Supplementary Files 8). Fourier power of all neurons were compared between real data and simulation in Figure8-Figure supplement 1. Figure 8C indicates that, though not appreciated by direct inspection, the periodicity of sensory stimulus is in fact propagated to many neurons in the nervous system, which is also reflected in small but non-zero contribution of many neurons to motifs 13 and 14 in TDE-RICA (Figure 3), and this characteristic is reproduced in freerun simulation by gKDR-GMM (Figure 8C and Supplementary Files 8).

**Figure 8.**
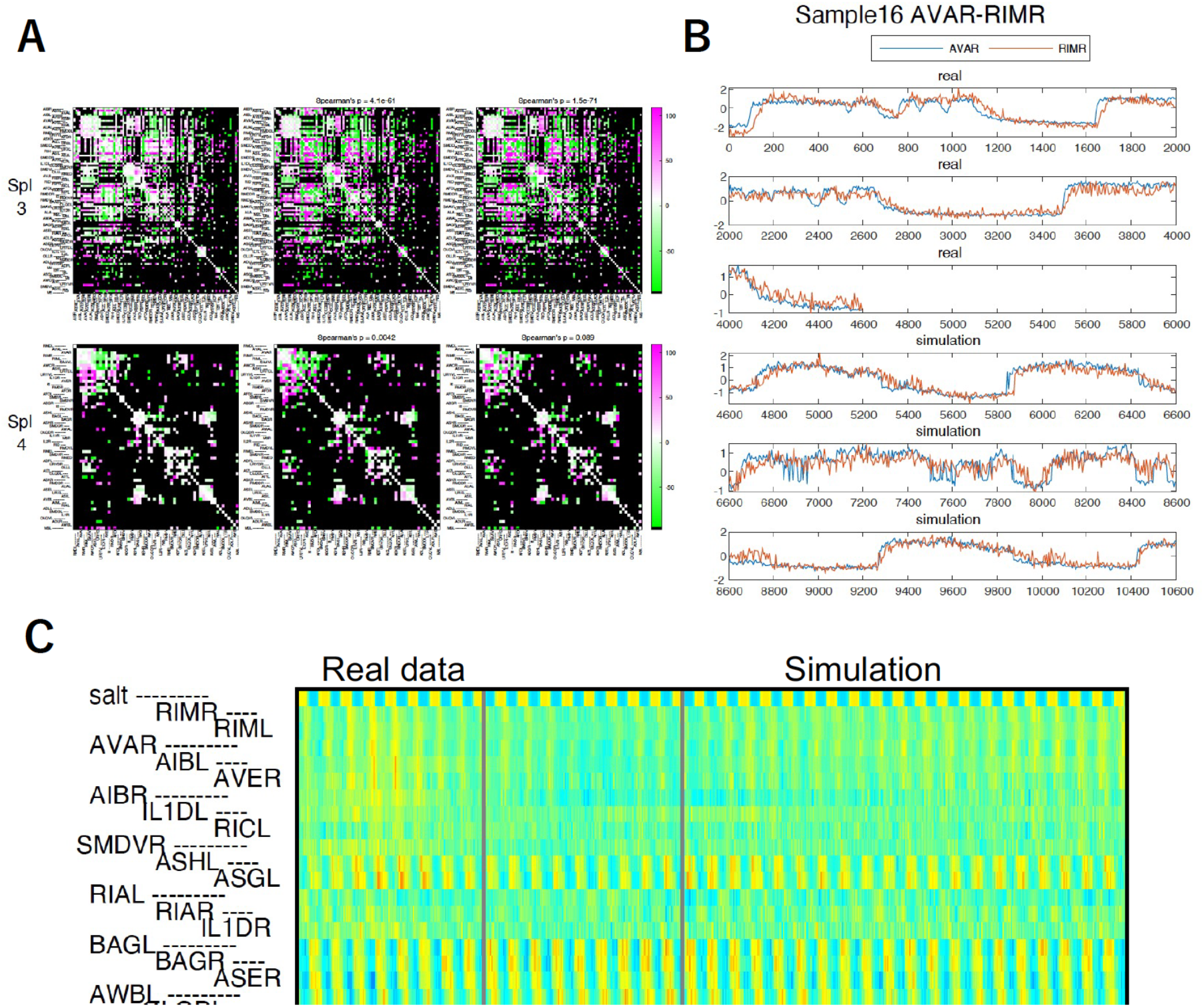
Comparison of real and simulation results. **(A)** Lagged cross-correlation of all combinations of neurons were determined and the lags that provided the best absolute value of cross-correlation are shown color-coded. Magenta are positive and green are negative lags. **(B)** Example of time series plot of a pair of neurons, AVA and RIM, which showed lagged correlation in a majority of samples. **(C)** Periodicity of neural activities. Top row shows salt stimulus (concentration change between 50 mM and 25 mM), which has a regular periodicity. Real and simulation results as Figure 7A was shifted by multiples (1-fold to 10-fold) of salt stimulus period and overlayed to visualize the periodicity of activity of each neuron.

**Figure 8 - Figure supplement 1.**
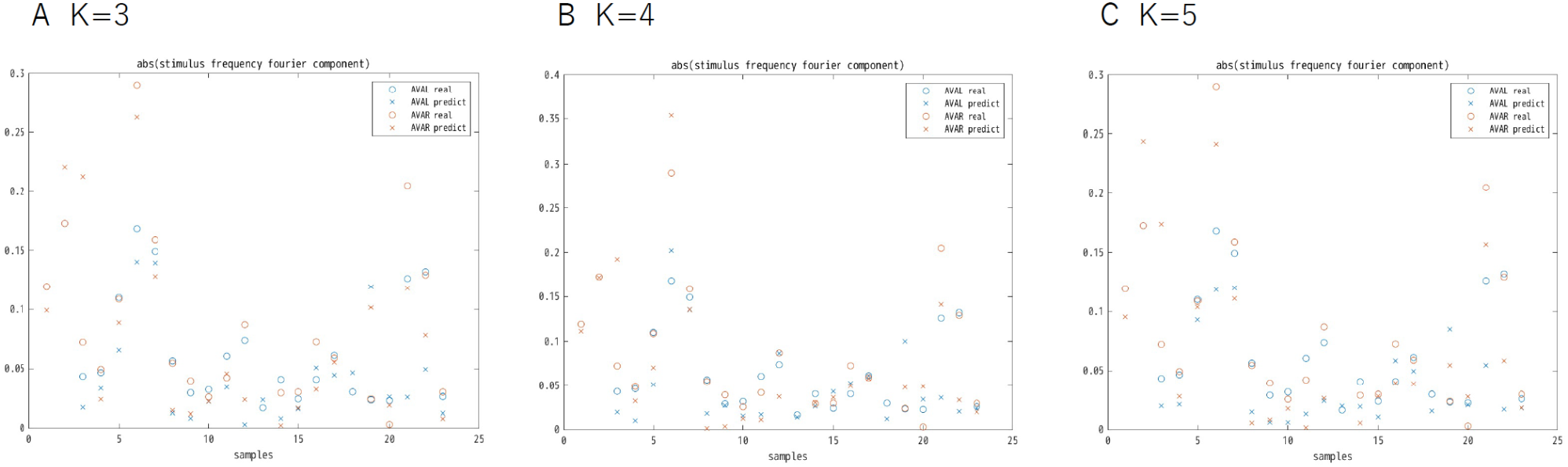
Fourier power of AVAL/R activity at the frequency of sensory stimuli. Fourier power of real AVAL/R activity (circles) and gKDR-GMM simulation results (crosses) with different model parameters **(A)** *K*=3, **(B)** *K*=4, **(C)** *K*=5 are shown. Only some of the samples show large periodic components.

These results indicate that our method reproduces major dynamics in a probabilistic manner. We note that this is valid for some but not all samples of 4D imaging. Through a survey of all results, it seems that there need to be clear correlated activities in groups of neurons for the dynamics to be reproduced. However, in cases where a majority of neurons showed grobally correlated (epilepsy-like) activity changes (for example sample 6, 19, 20 in Supplementary File 4B), the ensemble dynamics was difficult to reproduce.

### Noise-driven probabilistic behavior is essential for reproducing *C. elegans* brain activity

In the gKDR-GMM model, we intentionally incorporated a probabilistic model. However, because gKDR in principle conserves the presynaptic-postsynaptic relationship of the neural activity data, we can also generate a deterministic model based on the output of gKDR (lower dimension variables *U*_*i,j*_(*t*) = *X*_*i,j*_(*t*)*B*_*i,j*_). We therefore tested how a deterministic model would behave. It can be simply achieved by using the same GMM model, but instead of prediction by a conditional probabilistic distribution, we can use the expectation *E*(*y*_*i,j*_(*t* + *Δt*)|*U*_*i,j*_(*t*)) for the simulation. This is practically a non-linear regression model.

We first tested prediction of the near future based on real data. Time series data from calcium imaging were split into two halves in the time axis, then the gKDR-GMM model was created using half of the data as training data, and the correlation between prediction and real data was tested in the other half. As shown in Figure 9, the deterministic model was successful for predicting 5-20 timepoints ahead for some neurons. However, it was almost impossible to predict more than 200 time points ahead, except for neurons with periodic activities such as salt-sensing neurons. Deterministic model performed slightly better than the probabilistic model for prediction of the near future.

**Figure 9.**
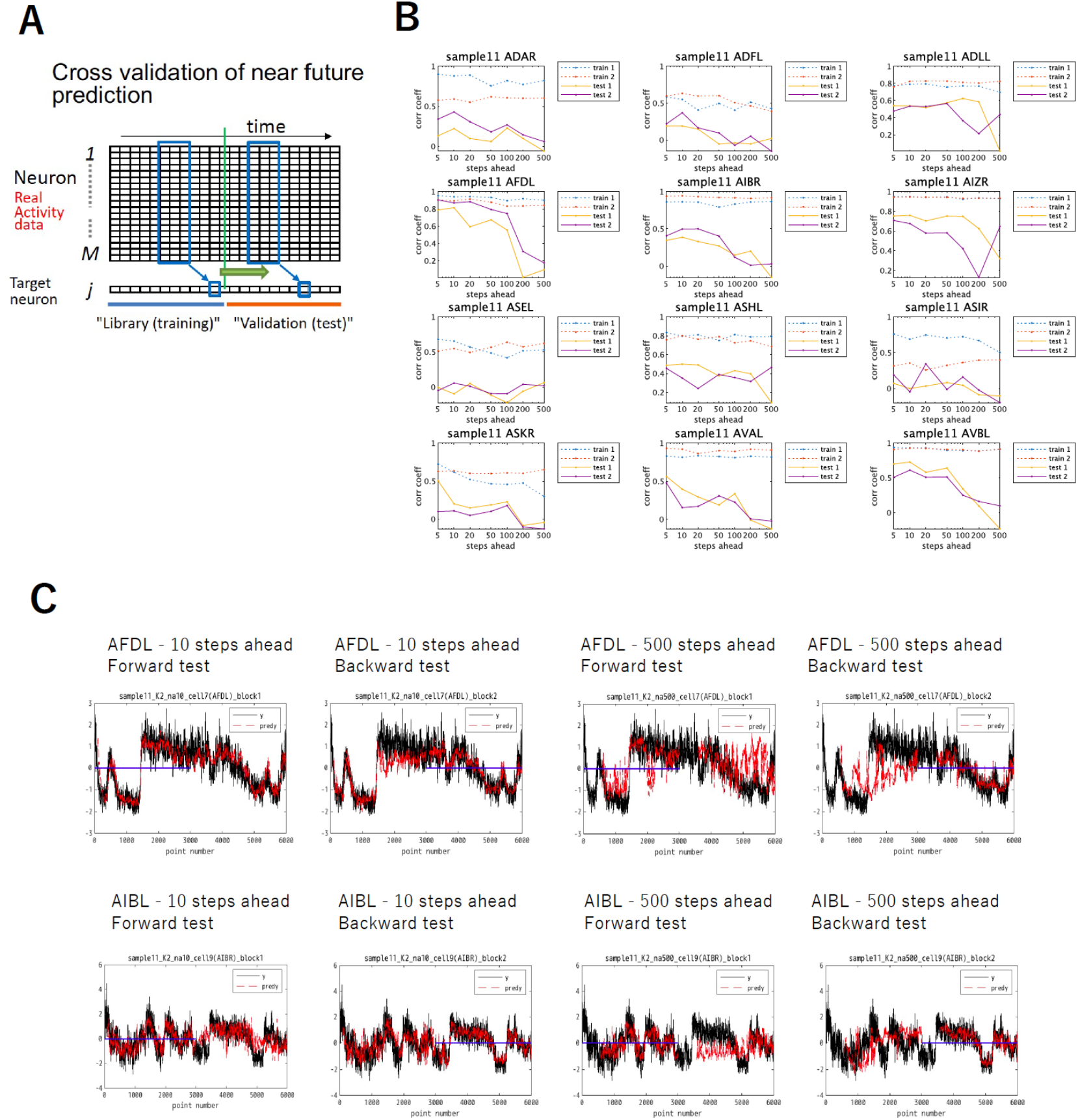
Prediction of near-future activity. **(A)** overview of the analysis. As in Figure 6, activity of target neuron was predicted, and ahead time span (*Δt* in Methods) was varied. Model was built using half of the data and the prediction was performed with the same data or the other half of the data the latter of which was not used for model fitting. **(B)** Near-future prediction results. Correlation between predicted data and real data are shown with different ahead time span. Blue for forward (training using the first half of data and testing in the second half), or red for reverse (opposite). Dotted lines for testing using the half data used for training, and filled lines for testing using testing data. Only a few neurons are shown as an example. **(C)** Comparison of predicted and real data. Blue line indicates the time span used for training. Black for real data and red for predicted results.

Figure 10A and Supplementary File 9A shows an example result of long-term simulation with the deterministic model (middle row). In this case, the neuronal activities either became constant or changed slowly and did not mimic real activity profiles, nor did correlation matrix mimic real activities (Figure 10B, Supplementary File 9B). If we omit the periodic sensory stimuli in the model, the simulated activities further degenerated because activities synchronized to sensory input were eliminated (middle row, right). Therefore, at least in our setting of the model, deterministic models cannot reproduce realistic activity patterns of the real neural network.

**Figure 10.**
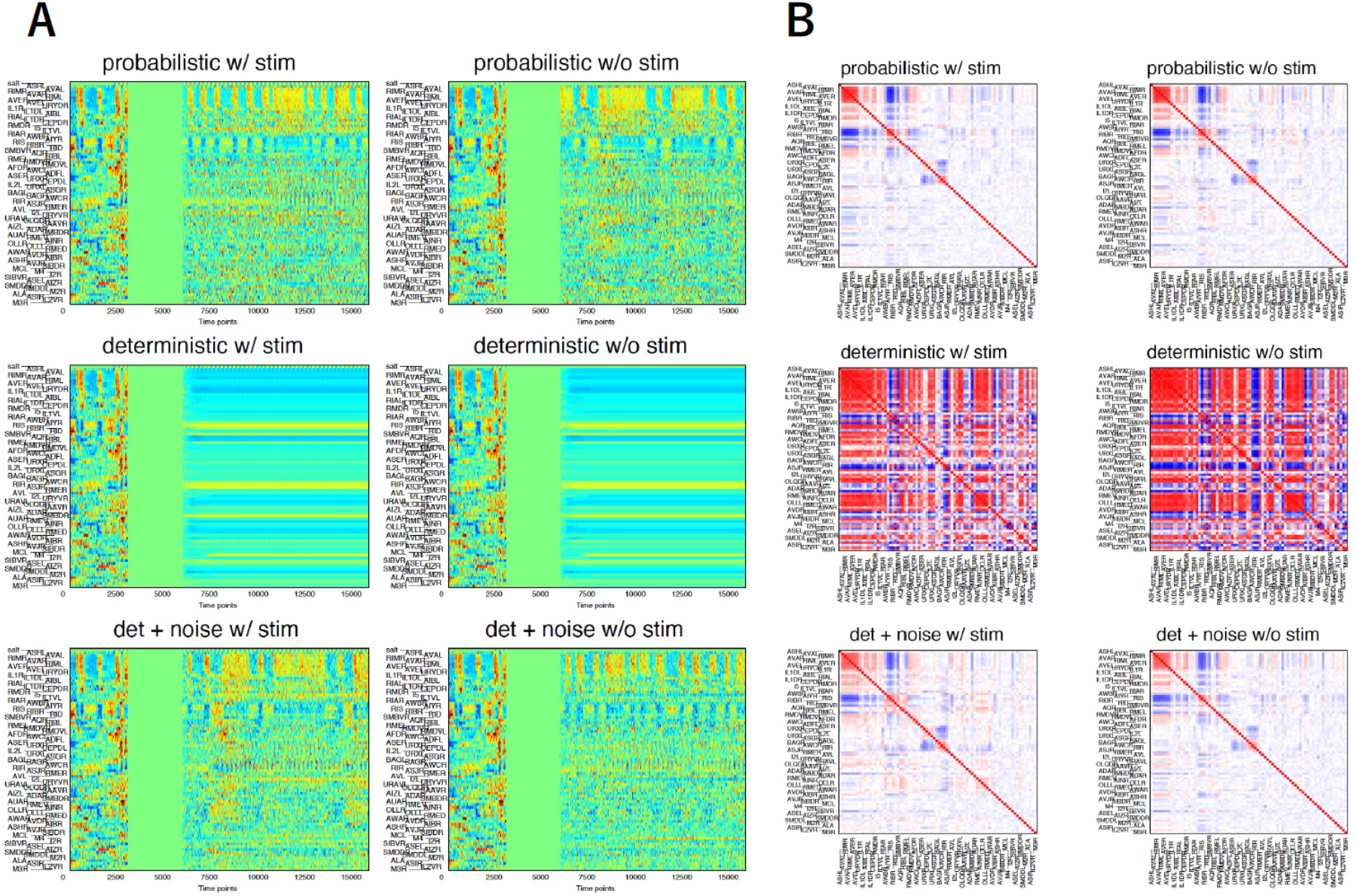
Freerun simulation by deterministic prediction. **(A)** Left part in each panel (0-1250 time points) shows real data. Simulated results follow zero-activity portion (1250-2500 time points), which was used as initial conditions for freerun simulation. “w/stim” indicates that periodic salt sensory input was added during the simulation, while “w/o stim” indicates that the sensory input was omitted. “probabilistic” indicates regular gKDR-GMM (Figure 7), while “deterministic” represents prediction by expectation value (which is unique) for GMM. “det + noise” represents deterministic prediction with random independent noise added. **(B)** Pairwise cross correlation of the simulation part of (A).

We reasoned that noise that is intrinsic to the neurons may be a major source of the dynamic activity of interconnected neurons. Based on this consideration, we next added Gaussian white noise with constant size (*i. i. d*. noise). This, in fact, caused generation of stochastic synchronized activities similar to that observed in real animals (Figure 10A, B, Supplementary File 9A, B, bottom row). Similar results were obtained by using gKDR-GP models (Supplementary File 10). These results suggest that irregular activities generated in each neuron are important for building a model that simulates the network properties of real animals.

### Principles of Network Dynamics

We next asked how the integrity of the neural network, such as correlation of activities among neurons, is maintained in the model. Although neural connectome is known in *C. elegans*, electron microscopy only provides information on physical connections but not the strength of the connections, nor whether the synapses are excitatory or inhibitory. Our gKDR-GMM model includes the estimate of the nature of synaptic transmission. The model describes a complicated relationship between multiple presynaptic neurons at previous time points and activity of the target neuron as a probabilistic distribution composed of multiple Gaussians. To simplify the relationship, we take an average of the gradient, მE(y_i_)/მx_j_. If this value is positive, it means that the more active the *j* input is, the more active is the target neuron at the next time step (excitatory transmission), and if the gradient is negative, the relationship is reversed. The gradient for all neurons in all samples are shown in figures and tables as Supplementary Files 11A-F.

One prominent feature of the whole brain dynamics is the backward (group A) and forward (group B) neurons, and how these correlated/anticorrelated activities are generated is an important and totally open question, with only reciprocal inhibition so far suggested (Kawano et al., 2011). In figure 11A (also Supplementary Files 12), gradients of synaptic contact are drawn as a network chart for A and B groups. As these examples show, multiple positive influences are present among A type neurons, and among them, there are a few hub-like neurons that send strong synaptic connections to other members. It is also true for group B neurons, and several negative connections are found between hub neurons in group A and group B. AIB and RIM neurons in A group and RIB neuron in B group tended to be hub neurons.

**Figure 11.**
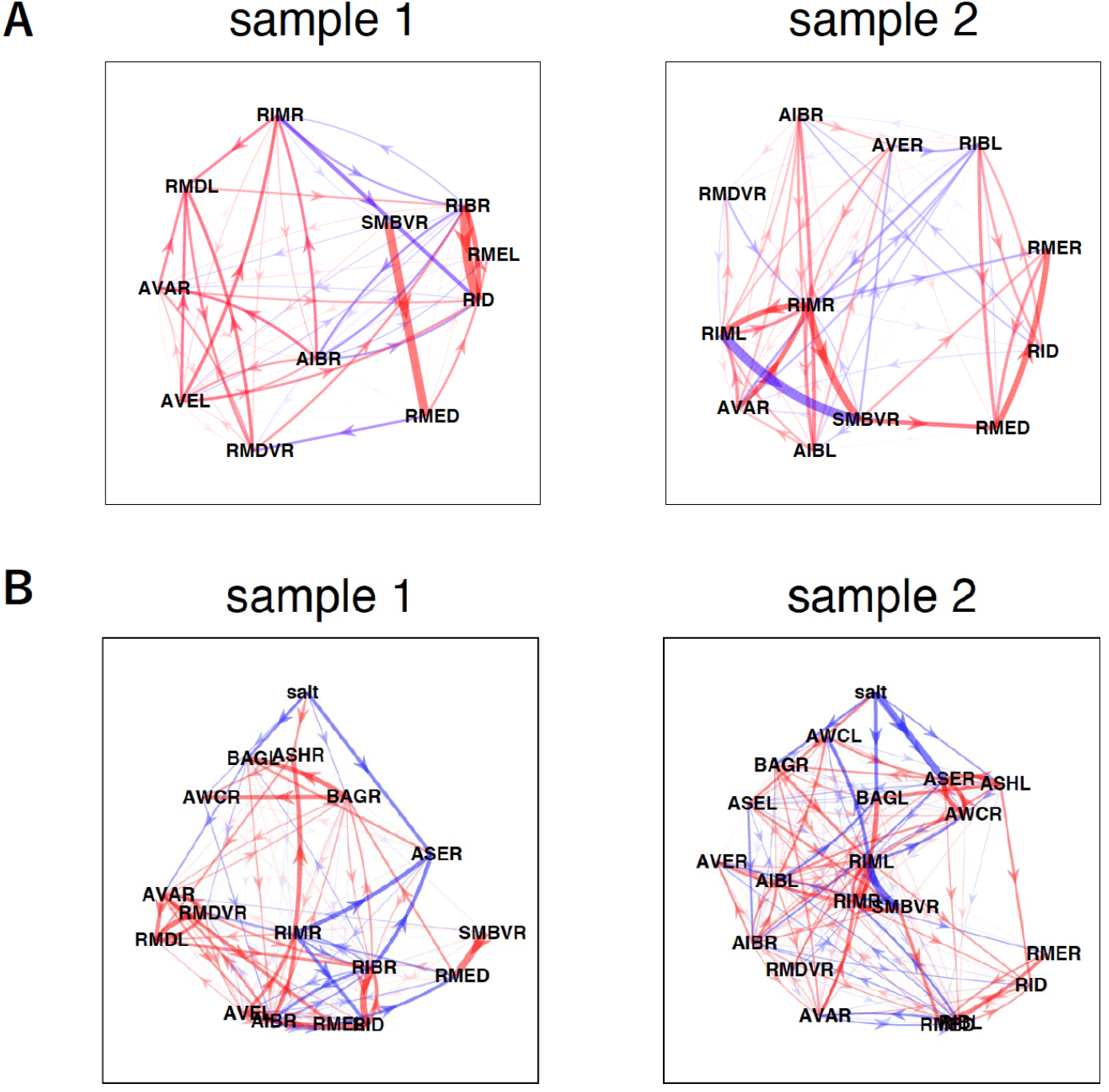
Predicted network structure for group A (backward) and group B (forward) neurons. **(A)** Mean gradient was used for estimation of synaptic weight for each neuron pair (see Methods). Red arrowed lines show positive (excitatory) influences, while blue shows negative (inhibitory) influences. Thickness and color strength indicated the magnitude of each gradient. **(B)** Predicted network structure for salt-sensing neurons and group A (backward) and group B (forward) neurons are shown. Similar to (A), synaptic strengths in the model are shown based on mean gradient (see Methods), but in this figure, salt sensory input and salt-sensing neurons were included.

We further evaluated the functional importance of each neuron. Because our model is based on synaptic transmission between each neuron pair, we can assess the role of each synaptic communication in generating the whole brain dynamics, by simply changing the synaptic transmission weight (corresponding to components of *B*_*i,j*_ matrix) to zero in the model. For simplicity, all the outputs of one neuron were abolished, and simulation was performed with modified *B*_*i,j*_. We then focused on each of highly correlated or anticorrelated neuron pairs. Table 1 (all data in Supplementary File 13) shows how killing each neuron affects the correlation. A general principle in these results is that, for a pair of neurons, killing output of either member of the pair does not necessarily have a strong effect, and instead, killing output of a third neuron can cause severe reduction. There are often hub neurons that affect multiple correlated pairs. Importantly, different models with different *K* lead to similar results, while the results tended to vary more between different samples.

**Table 1.**
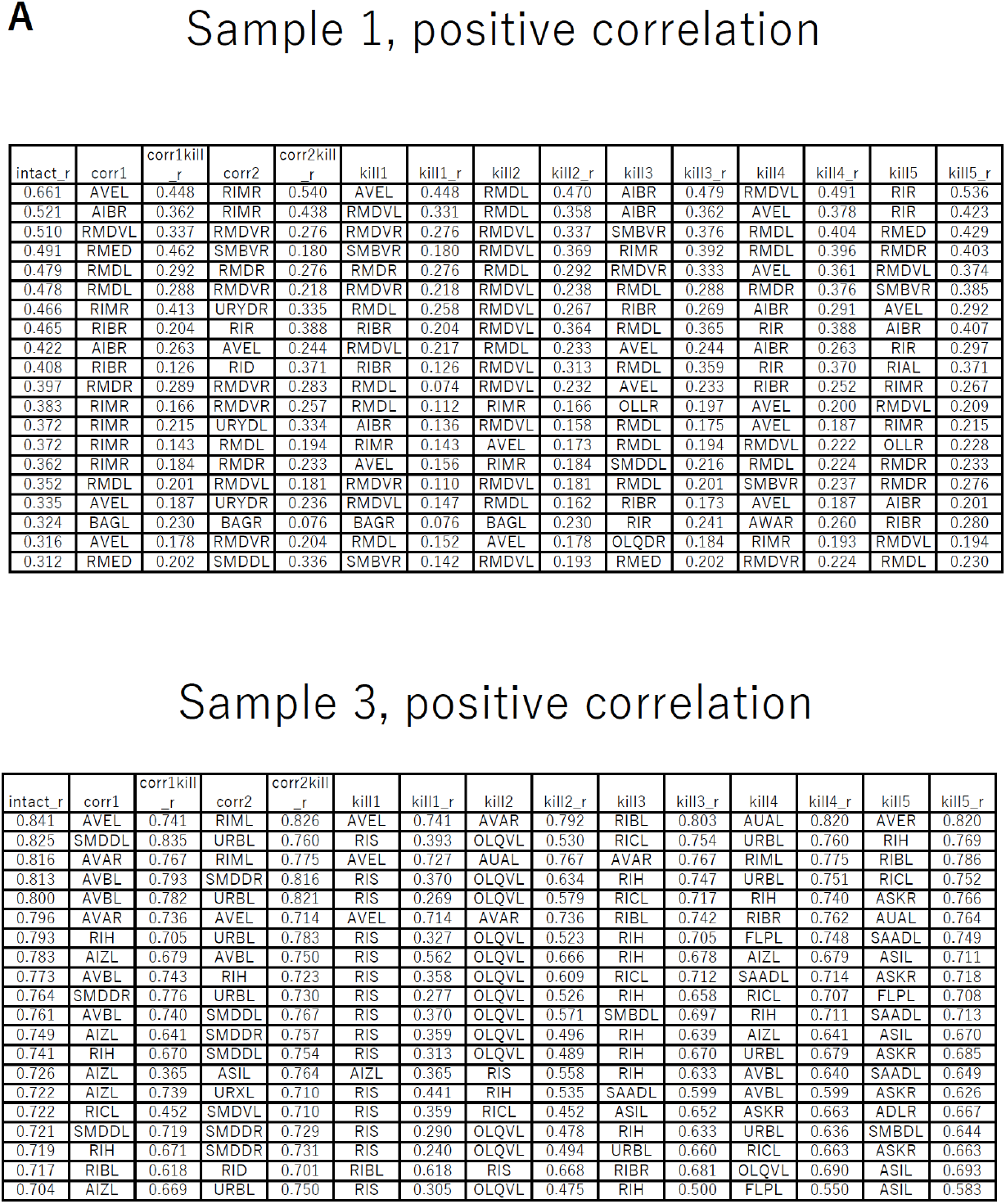
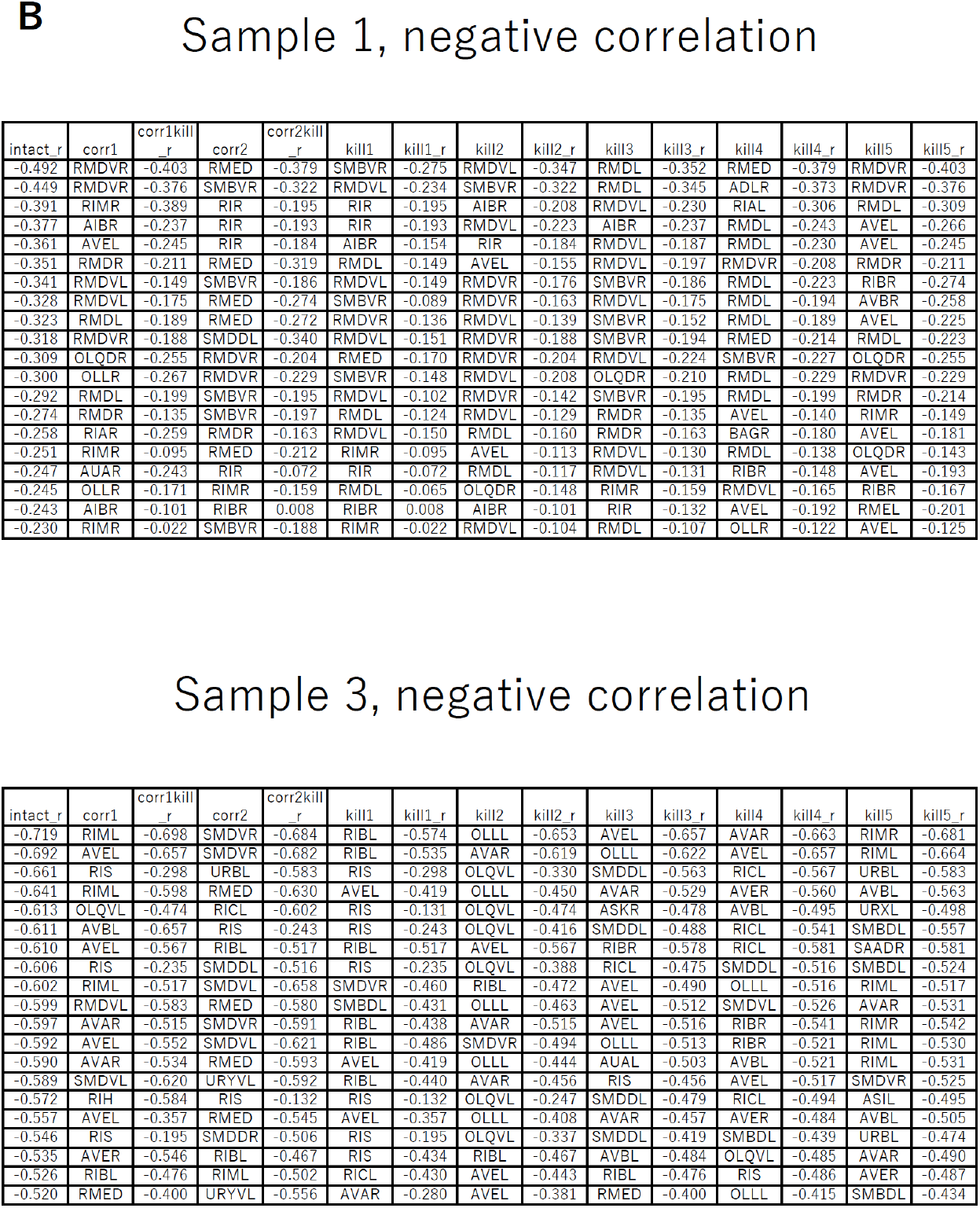
Effect of virtual ablation of neuronal outputs. **(A)** Highly correlated neuron pairs (“corr1” and “corr2”) are listed in the order of positive correlation, and changes in the correlation value of each pair was evaluated after removing outputs of each neuron in the model. “corr1 kill_r” shows the correlation between corr1 neuron and corr2 neuron when all output of the corr1 neuron was removed during the simulation. “corr2 kill_r” in a similar manner. “kill1” and “kill1_r” through “kill5” and “kill5_r” show neurons that caused the most severe reduction, in the order of effects, in the correlation between corr1 neuron and corr2 neuron. Examples of two samples are shown. **(B)** Similar to A except that most negatively correlated neuron pairs are shown and neurons that had the most severe effects on the negative correlation were listed. See Supplementary File 13 for all samples.

As described earlier, worms show either attraction or avoidance of NaCl, and the major mechanism of chemotaxis is known as biased random walk, in which the probability of switching from forward to backward movement is modulated depending on the salt concentration change that the sensory neurons sense (Pierce-Shimomura et al., 1999). How is this information transmitted within the nervous system? As noted earlier, many neurons including the backward command neuron AVAL/R show activity changes at the frequency of salt input. If we omit sensory input in the model, this periodicity of AVAL/R is greatly reduced. If we look at synaptic transmission from salt-sensing neurons (defined in the model) to group A/B circuits using the gradient-based analyses as described above, we see multiple routes from salt-sensing neurons to AVA (Figure 11B, Supplementary Files 14A-C).

To see the significance of each connection, we again picked each neuron and ablated synaptic input from all salt sensors to this neuron. As a result, in most cases effect of blocking input to a single neuron had only minor effects, with the exception of sample 1, where blocking direct input to AVA had a strong effect (Figure 12A, Supplementary File 15). In all other samples that had significant periodicity in AVA, blocking input to multiple neurons at the same time, if we choose appropriate neurons, gradually reduced the periodicity of AVA, suggesting there are multiple routes from salt sensor neurons to the motor command circuit (Figure 12B). Those neurons that were often found important were AIB and RIB, in addition to direct input to AVA. In these cases, the positive and negative correlations within and between A group and B group neurons were considerably maintained. We therefore conclude that motor command structure is intrinsic, and can be driven by random noise, but on top of it, multiple routes from sensory neurons stimulate members of the circuit to synchronize the motor circuit with the sensory input, eventually generating the stochastic sensori-motor responses of the animal.

**Figure 12.**
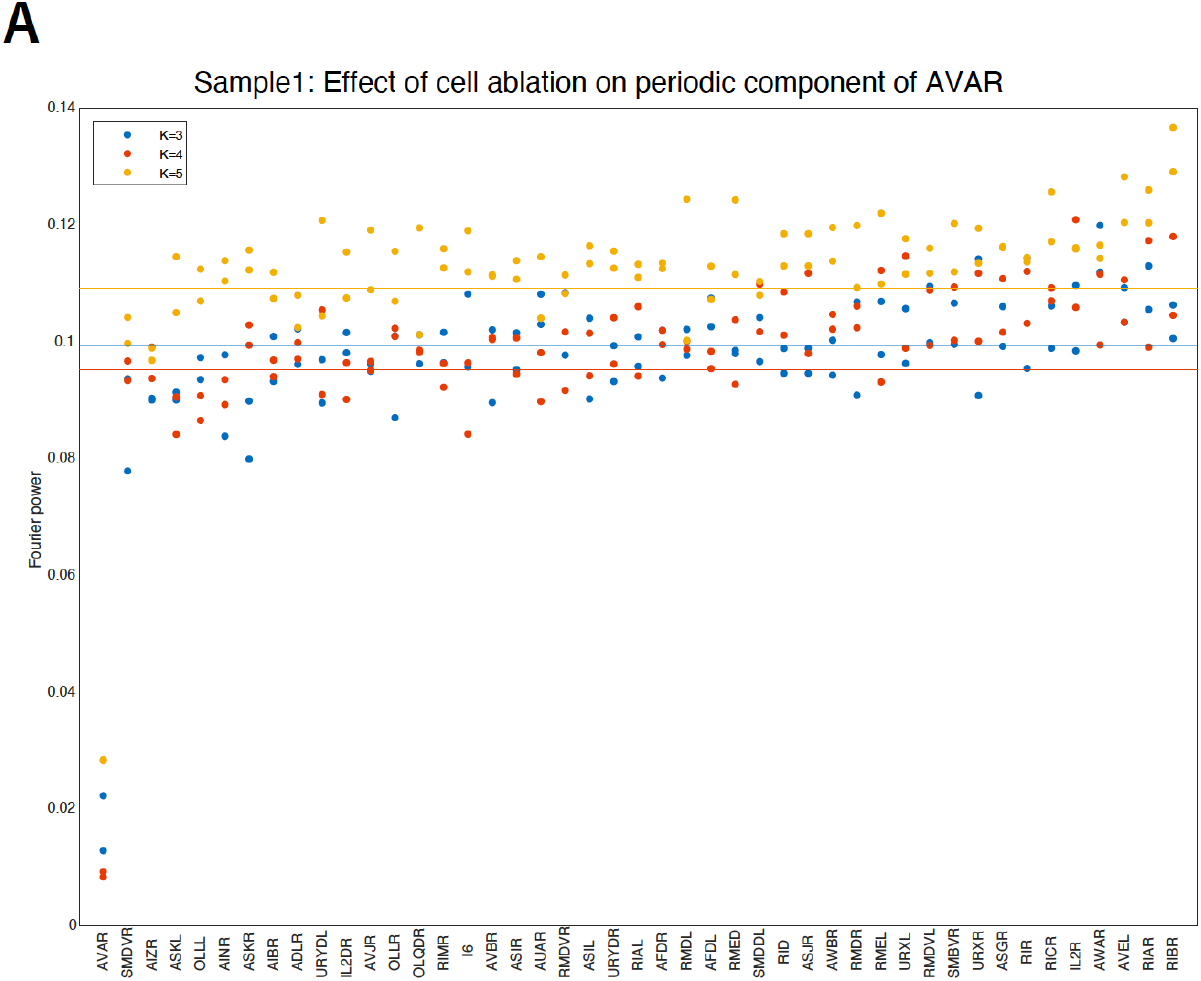
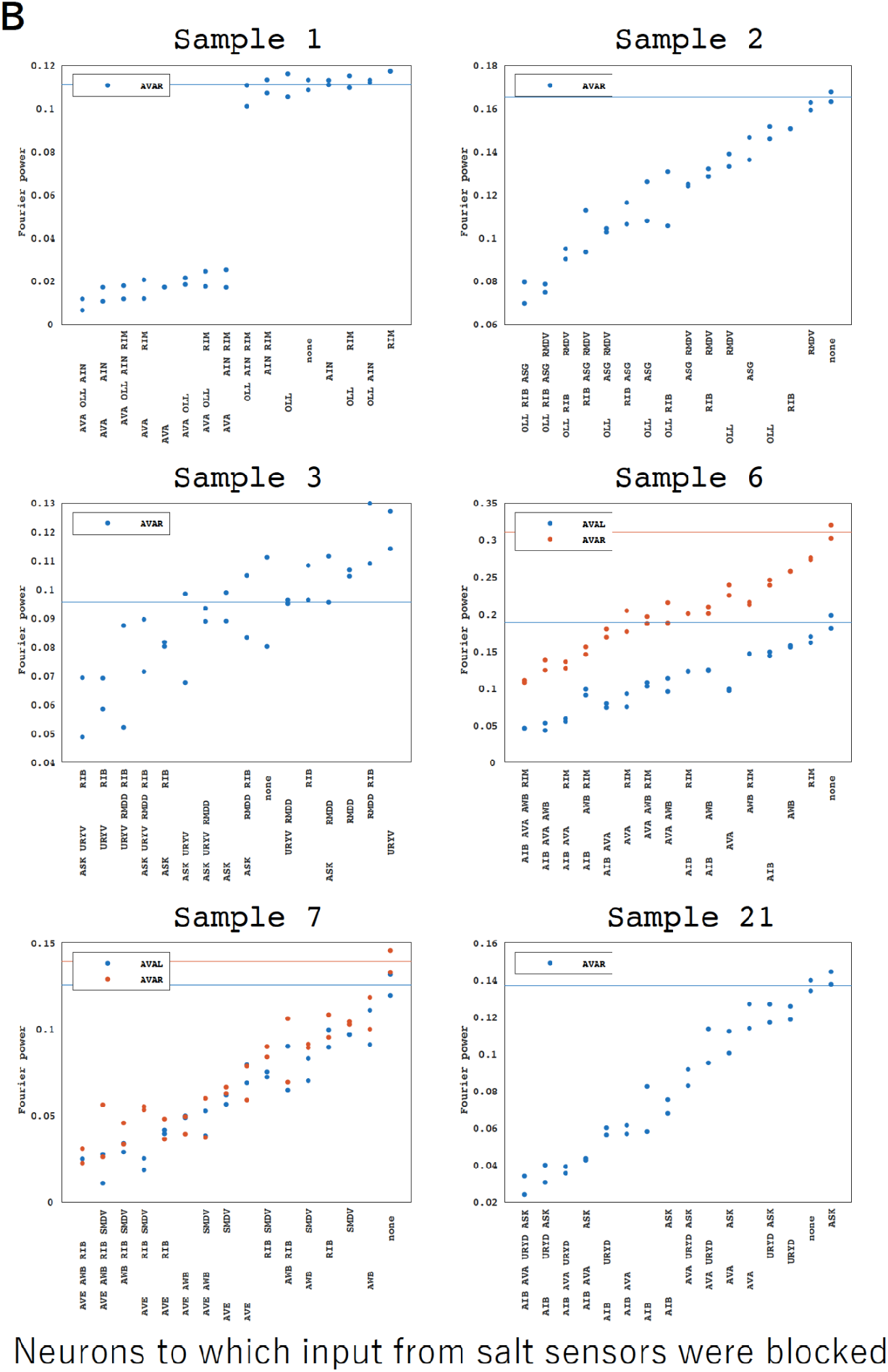
Effect of neuron ablation on transmission of periodic sensory information. **(A)** Single neurons were ablated individually. Periodic component in the frequency of sensory stimulus in the activity of backward command neurons AVAL and AVAR were estimated by Fourier power. Input from all salt sensory neurons were blocked for each neuron shown in abscissa during simulation, and the ablated neurons were ordered by the strength of the effect. Tests were performed for gKDR models with dimension reduction *K*=3,4,5. Only a few examples are shown. **(B)** Inputs from all sensory neurons were blocked for a combination of neurons indicated. Groups of neurons were ordered by the strength of their effects. All samples that showed above-threshold periodicity in AVAL/R are shown.

## Discussion

*C. elegans* has been an attractive experimental system for neuroscience, because apart from the advantage of genetics and ease of molecular manipulation, the compact nervous system is observable at single cell resolution, all neurons have been named and the whole-brain connectome has been determined. With this background, in this study, we have conducted calcium imaging of the *C. elegans* head region where calcium probe GCaMP2.6 was expressed in all neuronal nuclei. Through 3D segmentation, tracking and annotation, we successfully obtained “whole brain” neural activity data for 24 samples, in which neural activity data with neuronal identity has been obtained. To our knowledge, this is the largest annotated *C. elegans* neural activity data set so far reported.

By annotating neurons, one can for the first time compare the nervous system dynamics between different animals. Although our recording was performed in animals restrained in the same chamber and similarly stimulated by the chemoattractant NaCl, there is considerable difference in the activity patterns between individual samples. We currently do not know whether the difference stems from individual difference in the connectome structure that occurred during embryonic or post-embryonic development (which has been reported recently (Witvliet et al., 2021)), individual difference due to difference in the experience due to subtle difference in the environmental stimuli. Alternatively, it may not reflect individual differences but is rather fluctuation through time of the network dynamics (similar to dwelling and roaming in the animal locomotion) (Ji et al., 2021). Part of the reason for the apparent difference is due to the fact that only a fraction of all neurons are annotated in our dataset.

To overcome the above limitations and extract activity profiles common among different samples, we employed independent component analysis (ICA). ICA is used for separation of signals, but by combining ICA with time-delayed embedding, we succeeded in extracting common dynamic patterns out of the whole brain network. In addition, by employing matrix factorization, we could cover the missing data and could extract global patterns. The results revealed multiple sets of neurons that are engaged in major dynamic patterns, which included sensory response dynamics (motifs 13 and 14) and spontaneous motor dynamics (motifs 1 and 2). Thus, although the TDE-RICA method succeeded in extracting the stimulus-dependent and the spontaneous motor signals as separate components, it could not clearly extract how stimulus-dependent signal affects the motor signal. While this coarse-grained method might be suitable for examining global dynamics across all individuals, the effect of stimulus-dependent signals might have been too weak to be captured by this method. A more fine-grained method that can capture slight variations was needed to achieve this goal.

In this study, we estimated the synaptic weights along the physical network structure of *C. elegans* nervous system. We aimed at building data-driven phenomenological models rather than physical models that are described by a set of differential equations. We did not employ a differential equation approach which requires assumption of particular transmission function, but rather employed time-delay embedding and nonlinear regression models, which can be considered nonparametric models. Namely, activity values of presynaptic neurons at several different delay times were used as explanatory variables which are mapped to the response variable by a nonlinear function.

Given that synaptic transmission is intrinsically a nonlinear phenomenon, using nonlinear models is essential for reproducing neuronal events. Moreover, because there are on average 10-20 presynaptic neurons (Fig. 1 figure supplement 2), dimensionality of the explanatory variables is quite large. Therefore, choice of proper dimensionality reduction method is crucial. There are many commonly used dimensionality reduction methods such as principal component analysis (Kato et al., 2015), independent component analysis, kernel principal component analysis, t-SNE (Lin et al., 2020; Maaten and Hinton, 2008), UMAP etc. However, all these methods fail to take into account the relationship between explanatory variables (activity presynaptic neurons in our case) and response variables (postsynaptic neuron activity ahead of time). This could cause removal of (or giving too small weights to) variables (neurons and time points) that strongly influence the response of the target neuron. Rather, dimensionality reduction that takes into account of the response is preferred. Examples of linear approaches for this purpose are canonical correlation analysis (CCA) (Fung et al., 2002) and partial least square (PLS) (Helland, 1988; Höskuldsson, 1988) among others, which seek a low dimensional projection (*U*) of explanatory variables (*X*), so that *U* has a high correlation coefficient or high covariance with the response variable, *Y*. However, as mentioned earlier, *X* and *Y* are likely to have non-linear relationship, which leads us to the choice of KDR which is based on the kernel method and assumes no linearity.

In KDR, *K*-dimensional projection, *U*, of explanatory variables, *X*, is looked for so that variance of *Y* conditioned on *U* is minimal. This has the same effect as maximizing mutual information between *U* and *Y* given *U* is a subspace projection of *X*. Hence KDR is special in achieving the same purpose as CCA and PLS even if *X* and *Y* are not linearly related. While KDR is solved by numerical optimization, gKDR is an extension of KDR where the objective function is based on the gradient of kernel functions. This makes the problem convex and therefore the solution is unique (there is no risk of local minima), which can be determined by simple matrix-based calculation. To deal with nonlinear relationships, kernelization approaches have been employed and applied to CCA (kCCA) (Bach and Jordan 2002) and Support Vector Regression (Boser et al. 1992), both of which aim at direct regression but do not achieve dimensional reduction. This means that these methods cannot evaluate the relative importance of each explanatory variable for regression. KDR/gKDR is unique in achieving this goal by evaluating statistical conditional independence between subspaces of explanatory variables and the response variable. Ather potential means to achieve the same goal are deep neural networks, which will be interesting to pursue in the future based on our results.

By using dimensionality reduction by gKDR and probabilistic regression models GMM or GP, we could reproduce overall dynamics of the *C. elegans* brain reasonably well. We need to be cautious that the set of synaptic weights may not necessarily be the only solution that reflects the actual synaptic weights in the samples. In fact, we occasionally observed different relationships in different models for the same sample (for example synaptic weight between RIBR and RMER in *K*=4 vs *K*=3/5 in sample 8 in Supplementary Files 12A-C). However, we consider these as rare exceptions and the fact that models built with different numbers of *K*, which affects both gKDR and regression model, showed similar results in most cases, suggest that the obtained results are robust and likely be a good guess of how real neural network is working. Our results that the reproduction of the overall network dynamics was compromised in *K* = 1 than higher *K* and also *κ* = 1 than higher *κ* suggests that non-linear interaction of multiple presynaptic neurons is important for carrying out the actual brain functions.

The advantage of the synapse-based model is that it allows us to ask which neurons and synapses are responsible for each aspect of the whole-brain dynamics. By making use of the advantage, we tested the effect of each neuron on pairwise correlation and sensory information transfer. A general rule found through these analyses is that correlation of the activity in a pair of neurons is not necessarily formed by the neurons themselves, but rather there are hub neurons that govern correlated behaviors of a set of neurons. On the other hand, sensory transmission is a distributed system and in only a few cases a particular pathway has a major role, but rather information from sensory neurons are transmitted through the network in a percolation-like manner which collectively drives the sensory-driven control of behavior.

In neuroscience, recent advance in massive imaging technologies demands analytical methods. In such studies, most commonly used analysis is cross correlation, which suggest functional connectivity between brain areas (Needham et al., 2022; Ota et al., 2021). Another popular method is Granger causality (Vezoli et al., 2021). While these rely on assumption of linear relationships, nonlinear counterparts include mutual information and transfer entropy. All these methods are descriptive of the relationship between neurons or brain areas, and the results cannot be used for long-term prediction or simulation. Therefore, new methods are required for truly quantitative understanding of the ensemble behaviors. Our approach provides one example of such endeavor. This approach could be used for large brains, where connectome data are beginning to be obtained (Oh et al. 2014) and wide-field calcium imaging, fMRI, EEG and MEG data can be available (Breakspear, 2017). Further, though our current analyses utilized the connectome data of *C. elegans*, the same method may be applicable to neuronal activity data where no connectome information is available, by assuming all-to-all connections.

A major drawback of KDR, which is common in kernel approaches, is the limitation of the number of data points whose square is proportional to the memory requirement for computation. In our current implementation, 50-70 Gbyte memory was required for processing 1000 time points and computation time was roughly proportional to *O*(*N*^*2*^*M*) where *N* is the time point number and *M* is the number of variables, namely the number of presynaptic neurons. As a rule-of-thumb, for making a gKDR-GMM model, it took 0.61 sec/presynaptic neuron for one target neuron or around 10-20 min for one animal using a 3110 TFLOPS supercomputer.

## Materials and Methods

### Strains and culture

Animals were raised at 20 °C under standard conditions on nematode growth medium (NGM) plates with *E. coli* OP50. For 4D imaging, the following strain was used (Toyoshima et al., 2020): JN3038 *qjIs11[glr-1p::svnls2::TagBFPsyn, ser-2(prom2)p::svnls2::TagBFPsyn]; peIs3042[eat-4p::svnls2::TagRFP675syn, lin-44p::GFP]; peIs2100[H20p::nls4::mCherry]; qjIs14[H20p::nls::YC2*.*60]*. Transgenic strains were generated by germ-line transformations in which we co-injected the gene of interest with a visible transformation marker (*lin-44p::gfp* etc.) into the animals. The transgene was then integrated into a chromosome by UV irradiation.

### Microscopic setup

The microscope system was developed as previously described (Toyoshima et al., 2020). Briefly, the system consists of a confocal microscope and three cameras (one CMOS camera and two EM-CCD cameras). A piezo actuator is attached to the objective lens of the confocal microscope to enable high-speed 3d imaging. The CMOS camera is used to capture images for cell annotation, and the two EM-CCD cameras are used to measure neuronal activity (Yellow Cameleon imaging).

### Image acquisition

The detailed microscope settings for imaging were previously described (Toyoshima et al., 2020).

Animals were raised on standard NGM plates until young adults, and further incubated overnight on pre-imaging NGM plates with 50 mM of NaCl. Osmolarity of pre-imaging plates was adjusted to 350 mOsm with glycerol. Animals were introduced into a PDMS microfluidic chamber and stimulated with a change in salt concentration from 50 mM to 25 mM or from 25 mM to 50 mM every 30 seconds (the period was 60 seconds). We used a modified version of the olfactory chip (Chronis et al., 2007) for the microfluidic device. The stimuli were delivered to the animals by switching the imaging solutions (25 mM potassium phosphate (pH 6.0), 1 mM CaCl2, 1 mM MgSO4, 0.02% gelatin, NaCl at the indicated concentration and glycerol to adjust their osmolarity to 350 mOsm).

For the 3D images for neuronal annotation, 50 slices per volume were taken to cover the entire body of animals (0.72 - 1.00 μm / slice). The size of each image is 1024*150 pixels. A total of 8 volumes were taken, and the best one was used for neuronal annotation. To acquire images for measuring neural activity, 22 slices per volume were taken for the same area (1.62 - 2.29 μm / slice). The size of each image is 256*64 pixels. A total of 6,000 volumes were taken at a rate of about 4 volumes per second (about 25 minutes in total).

### Cell detection, annotation and tracking

All the nuclei in a volume in the 3D images for neuronal annotation (called the annotation movie) were detected by our image analysis pipeline roiedit3D (Toyoshima et al., 2016) and corrected manually. We detected 201.9 ± 15.5 (mean ± standard deviation) nuclei on average from 24 samples.

The cells were annotated with neuronal identity based on the expression patterns of cell-specific promoters as previously described (Toyoshima et al., 2020). We annotated 146.8 ± 23.1 (mean ± standard deviation) cells on average.

For cell tracking, our cell tracking pipeline CAT was used. CAT selects the volume most similar to the volume in which the cells were identified from the images for measuring neural activity (called the activity movie). The volume in the annotation movie was registered to the volume in the activity movie by B-spline transform implemented in elastix (Klein et al., 2010). The ROI information including position, size, and annotation of nuclei was copied from the annotation movie to the activity movie. CAT also constructs a shortest path tree, where the volumes of the activity movie correspond to the nodes of the tree, and the similarities between the volumes correspond to the edges of the tree. The volumes connected by the edge were registered by the B-spline transform, and ROI information was copied. CAT run on the Shirokane3 and Shriokane5 supercomputing systems of the Human Genome Center (the University of Tokyo), and processed an activity movie within about 8 hr. The tracking results were checked and corrected manually by using roiedit3d. The fluorescence intensities of CFP, YFP, and mCherry were obtained by least square fitting of the tri-variate Gaussian mixtures, which correspond to the tracked ROIs, to the nuclei in the volume (Toyoshima et al., 2016). If the worm moved outside the field of view, the error periods were removed from the dataset.

### Pre-treatment

To remove noise, a median filter with a 5-point window was applied to the time series of CFP and YFP intensities. If the filtered CFP intensity is smaller than 1/10 of the median of the filtered time series of CFP, the time point was regarded as an outlier. CFP and YFP intensities at the outliers were set as NaN (missing), and were completed by a median filter with a 4-point window. The missing values can be completed if the missing values are sparsely distributed. The outlier neurons were removed from the dataset if the number of time points of the outlier is larger than 400 or any missing values remained in the completed time series. We also removed non-neuronal cells including hypodermal cells. The ratio of YFP over CFP was calculated, and the linear trend of the ratio was removed to compensate for the photobleaching. The time series of the ratio was subtracted by its mean and divided by its standard deviation for scaling. The scaled ratio of YFP over CFP was regarded as the neural activity. The obtained whole-brain activity dataset contains the neural activity of 139.6 ± 24.9 (mean ± standard deviation) cells on average from 24 samples, and covers a total of 177 out of 196 cells in the head region of animals. For gKDR, noisy neurons were removed from the analyses. Specifically, only neurons with autocorrelation of greater than 0.3 at a lag of 20 time points were included and those that did not meet this criteria were considered missing.

### time-delay embedding and Reconstruction ICA (TDE-RICA)

Let *x*_*i,j*_(*t*) be an activity of neuron *i* (from 1 to 177) of sample *j* (from 1 to 24) at time point *t* (from 1 to 6000). The embedding of neuron *i* in sample *j* at time point *t, X*_*i,j*_(*t*), is *X*_*i,j*_(*t*) = [*x*_*i,j*_(*t* − (*k* − 1)*τ*), *x*_*i,j*_(*t* − (*k* − 2)*τ*), …, *x*_*i,j*_(*t*)] ∈ ℜ^1×k^, where *k* = 300 and *τ* = 1. We choose these values so that the motifs obtained by the following analysis can capture the large-scale dynamics of neural activities. The embedded time series of neuron *i* in sample *j* is *X*_*i,j*_ = [*X*_*i,j*_^⊤^((*k* − 1)*τ* + 1), *X*_*i,j*_ ^⊤^ ((*k* − 1)*τ* + 2), …, *X*_*i,j*_ ^⊤^ (6000)] ∈ ℜ^/00×1201^.

We used Reconstruction ICA (Le et al., 2011) implemented in Matlab so that the captured components and weights can reproduce the original data. Because the RICA cannot handle missing values, 94 neurons of 10 samples (with no missing values) were selected from the whole dataset of 177 neurons of 24 samples (with missing values).

The embedded time series of the 94 selected neurons in sample *j* is *X*_,*j*_ = [*X*_1,*j*_^⊤^, *X*_2,*j*_^⊤^, …, *X*_94,*j*_^⊤^]^⊤^ ∈ ℜ^(94×300)×5701^, and the whole embedded time series of selected neurons in the 10 selected samples is *X* = [*X*_,1_, *X*_,2_, …, *X*_,10_] ∈ ℜ^(94×300)×(5701×10)^.

The cost function of reconstruction ICA consists of the terms of the reconstruction cost and the independence (non-gaussianity). Reconstruction ICA searches a matrix *W* that minimize the cost function 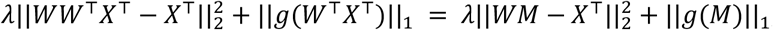, where *λ* is the weight for the reconstruction penalty, || · || is the entrywise norm, and *g* is the entrywise contrast function 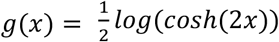. *M* is the independent components *M* = *W*^⊤^*X*^⊤^ ∈ ℜ^*n*×(94×300)^, and is regarded as the motifs of the neural activities. *n* is the number of components and we set *n* as 14, which was the minimum number to capture the neural response to the sodium chloride stimulation. *W* is the weight matrix *W* ∈ ℜ^(5701×10)×*n*^, and is regarded as the occurrences of the motifs.

For completing the missing values, the matrix factorization was combined with TDE-RICA. Matrix factorization does not take into account *M* = *W*^⊤^*X*^⊤^ and searches *M* and *W* that minimizes the reconstruction cost 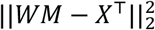. Therefore matrix factorization can be applied even if *X* contains missing values. Here we extend the matrices included in the reconstruction cost. *X*^*all*^ is the embedded time series of all 177 neurons in all 24 samples *X*^*all*^ ∈ ℜ^(177×300)×(5701×24)^. *M*^*all*^ is the extended matrix of motifs *M*^*all*^ = [*M*, M^*new*^] ∈ ℜ^*n*×(177×300)^. *W*^*all*^ is the extended matrix of occurrences *W*^*all*^ = [*W*, W^*new*^] ∈ ℜ^(5701×24)×*n*^. Then we search *M*^*new*^ and *W*^*new*^ to minimize the extended reconstruction cost 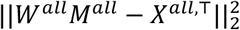 by using the L-BFGS optimization method implemented in matlab.

### Synapse-based regression models

#### gKDR

To construct a model that reproduces the dynamics of the whole neural network, we adopted several approaches that model synaptic inputs to each neuron. First, we collect the presynaptic-postsynaptic relationships in the physical network. Connectivity data was adopted from the electron microscopy reconstruction data (White et al., 1986) which had been digitized by Oshio et al. (http://ims.dse.ibaraki.ac.jp/ccep/) (Oshio et al., 1998).

Chemical synapses were treated as directional and gap junctions were treated as bidirectional synapses. Because in our 4D imaging data for each sample, there are many missing neurons, because 1) part of the nervous system can be obscure or too packed to obtain clear-cut signals for each neuron, 2) some neurons are difficult to name, for example because the positional relationship with surrounding neurons is atypical. Therefore, in one option, we included neurons directly connected to the target neuron (called “direct link”) to make the models, and in another option, we included neurons connected to the target neurons via two synapses in case the directly connected neuron was not observed (called “indirect link”). For the sake of simplicity, the neurons thus collected are denoted as “presynaptic neurons” for a given target neuron, even if the neurons are two synapses away in the case of employing indirect links.

To model dynamic relationships between the activity time-series of presynaptic neurons and that of target neurons, we generated regression models using time-delay embedding. As for TDE-RICA, *x*_*i,j*_(*t*) depicts the activity of neuron *i* at time *t* in sample *j*. Each neuron was set as a target neuron one by one. For example, assume that neuron *i* is set as a target neuron (*i* = 1,2, …, *M*_*j*_; *M*_*j*_ is the number of neurons in sample *j* data set). Activity of target neuron *i, y*_*i,j*_(*t* + *Δt*) ≡ *x*_*i,j*_(*t* + *Δt*), is predicted by activities of presynaptic neurons. Here, similar to TDE-RICA, time-delay embedding was adopted to utilize the past time series of the presynaptic neurons to predict target neuron activity. Namely, *y*_*i,j*_(*t* + *Δt*) is predicted from 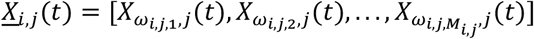, where *X*_*m,j*_(*t*) stands for a row vector representing time-delay-embedded activity of neuron *m*, 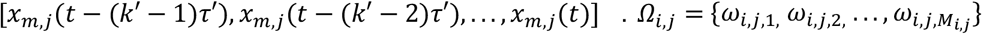 is a set of indices of neurons presynaptic to target *i, M*_*i,j*_ being total number of neurons presynaptic to neuron *i*) (Figures 6B and 13B). Note that previous activity of the target neuron itself, *X*_*i,j*_(*t*) = [*x*_*i,j*_(*t* − (*k*′ − 1)*τ*′), *x*_*i,j*_(*t* − (*k*′ − 2)*τ*′), …, *x*_*i,j*_(*t*)] (identical to [*y*_*i,j*_(*t* − (*k*′ − 1)*τ*′), *y*_*i,j*_(*t* − (*k*′ − 2)*τ*′), …, *y*_*i,j*_(*t*)]) was also included in the explanatory vectors 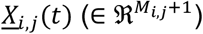.

Mean number of presynaptic neurons was 6.8 and 21.1, respectively, for direct link and indirect link (Figure 7 - Figure supplement 2). This causes a widely recognized challenge of the curse of dimensionality. Considering that synaptic transmission is an intrinsically nonlinear process, we employed the gradient kernel dimension reduction (gKDR) (Fukumizu and Leng, 2014) for dimension reduction. The basic principle of KDR is explained in Figure 13C. We shall model a function *y*_*i,j*_ = *f*_*i,j*_(*<u>X</u>*_*i,j*_) ≃ *g*_*i,j*_(*U*_*ij*_) (here *y*_*i,j*_, *<u>X</u>*_*i,j*_ and *U*_*ij*_ are considered random variables and *y*_*i,j*_(*t*), *X*_*i,j*_(*t*) and *U*_*ij*_(*t*) are samples from them), as a reasonably simple model, which is achieved by selecting a low dimensional subspace of explanatory variables as 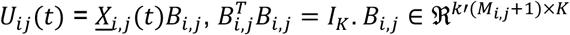 is selected aiming at making *U*_*ij*_ (∈ ℜ^1×*K*^) sufficiently informative for predicting *y*_*i,j*_. More precisely, gKDR evaluates statistical independence between *<u>X</u>*_*i,j*_(*t*) and *y*_*i,j*_(*t* + *Δt*) conditioned on *U*_*ij*_(*t*), using Reproducing Kernel Hilbert Space (RKHS), and maximizes it. It thereby finds a subspace including *U*_*i,j*_ that is most informative for estimating *y*_*i,j*_. In the toy example in Figure 13C, the value of *y* is shown in pseudo-colors. Although *y* is a function of *x*_1_, *x*_2_ and *x*_3_, *y* depends only on the values of *u*_1_ and *u*_(_ and is independent of *u*_3_ (an axis perpendicular to *u*_1_ and *u*_2_). Therefore, gKDR selects a *B* that maps (*x*_1_, *x*_2_, *x*_3_) to (*u*_1_, *u*_2_). The dimension of the subspace, *K* needs to be pre-determined and in this work *K* was optimized by a grid search as well as other hyperparameters *k*′, *τ*′ and *Δt* as described in the main text. gKDR was performed by Matlab codes distributed by Kenji Fukumizu on his web site (https://www.ism.ac.jp/~fukumizu/index_j.html).

**Figure 13.**
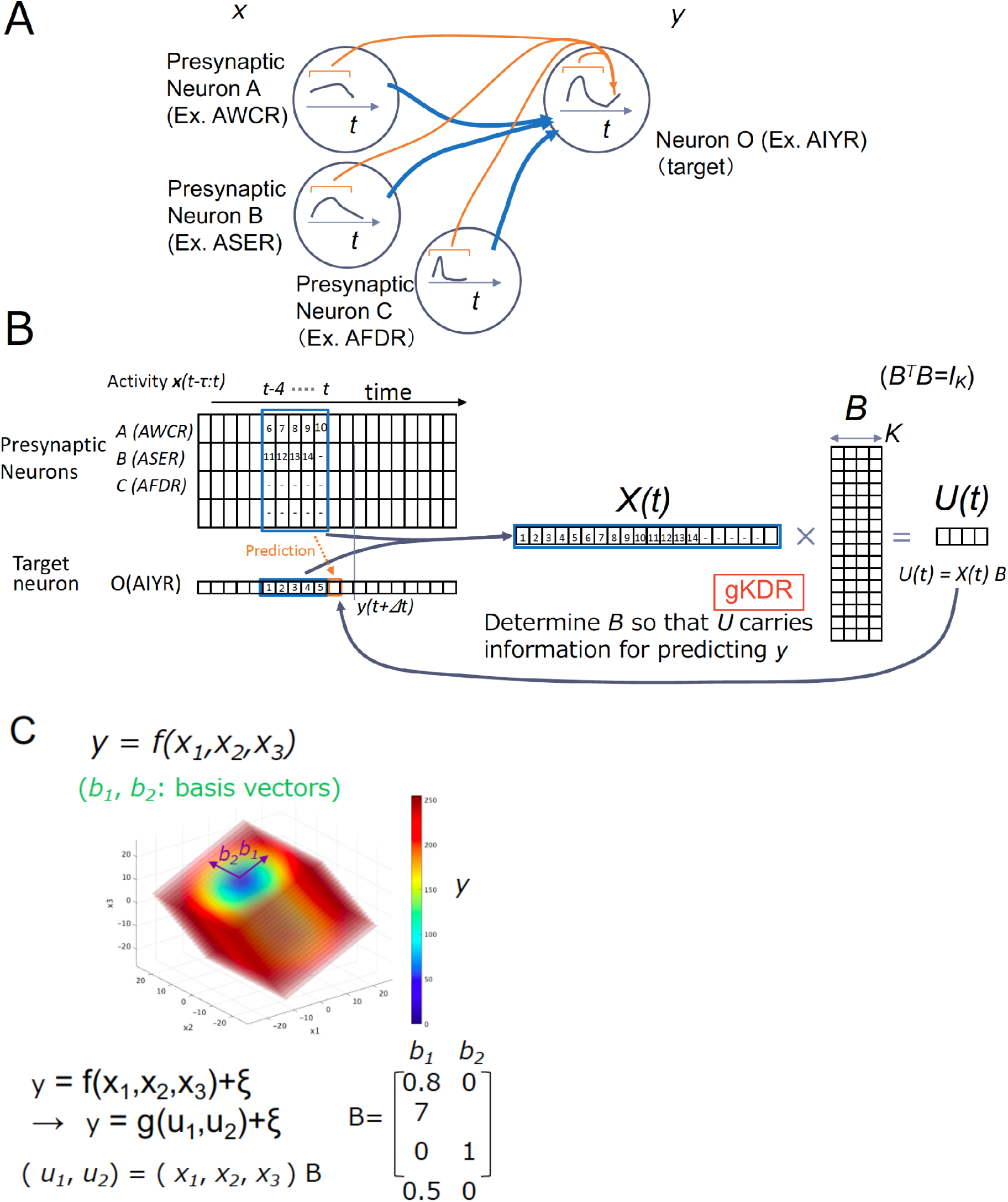
Outline of gKDR-GMM. (A, B) For a given sample *j*, the model learns to predict activity of the target neuron (neuron *i* = neuron O in this figure) ahead of time (*y*_*i,j*_(*t* + *Δt*)) from previous activity of presynaptic neurons. Activity of presynaptic neurons was time-delay embedded. *k*′ steps of previous activity in intervals *τ*′ were taken and made a row vercor *X*_*i*,*j*_(*t*). These vectors are concatenated for all presynaptic neurons for neuron *i* (*<u>X</u>*_*i,j*_(*t*)) (= *X*(*t*) in this figure). Then gKDR reduces the dimension of presynaptic activities *<u>X</u>*_*i,j*_(*t*) to *K* dimensional value *U*_*i,j*_(*t*) (= *U*(*t*)). GMM models the joint probability of (*U*_*ij*_(*t*), *y*_*ij*_(*t* + *Δt*)) as a weighted sum of Gaussian distribution which is fitted to the real data. Conditional probability of *y*_*ij*_(*t* + *Δt*) is determined from the GMM model and used for prediction. **(C)** Toy example explaining gKDR. This example represents an imaginary function *y* = *f*(*x*_1_, *x*_2_, *x*_3_). *y* value is color-coded in the 3D model. If given *K* = 2 (reduction dimension), from the dataset of *n* samples of (*x*_1_, *x*_*2*_, *x*_3_, *y*), gKDR determines the basis vectors (*b*_1_, *b*_2_) because *y* is independent of *b*_1_, which defined the rest of the dimensions spanned by *x*_1_, *x*_2_, *x*_1_, while *y* depends on *b*_1_ and *b*_2_. *b*_3_ and *b*_2_ constitutes column vectors of *B*.

#### Sensory input

As described above, we stimulated the animals with regular changes of salt concentrations while 4D imaging. In the gKDR-GMM and gKDR-GP models, salt sensor neurons were defined as ASEL/R, AWCL/R, BAGL/R and ASHL/R. ASEL and ASER are known to be salt-sensing neurons that are most important for salt chemotaxis. AWC and BAG were included because they are ciliated sensory neurons and consistently showed prominent activities synchronized to salt stimulus. ASH neurons were included because they have been shown to sense salt and occasionally showed salt response in our data set (Thiele et al. 2009). In the model, for these neurons, salt concentration was simply treated as an additional presynaptic neuron.

#### gKDR-GMM

Determination of the dimension reduction matrix *B*_*i,j*_ was achieved by gKDR as described in the previous section. In the case of gKDR-GMM, the mapping function *g*_*i,j*_(*U*_*ij*_) is considered a probabilistic distribution. Here, rather than considering *g*_*i,j*_(*U*_*ij*_) itself, the joint probability of *Z*_*ij*_ = (*U*_*ij*_(*t*), *y*_*ij*_(*t* + *Δt*)) was modeled by using a Gaussian Mixture model. Namely, 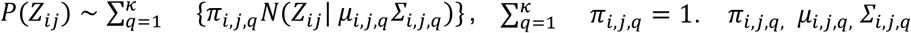 were determined by using { *Z*_*ij*_(*t*) } from the training data. The optimal number of gaussians (*κ*) was also searched for and as a result we employed two Gaussians as described in the main text. fitgmm function of Matlab was used. Prediction by the gKDR-GMM was performed by random extraction from the conditional distribution *P*(*y*_*ij*_(*t* + *Δt*)|*U*_*ij*_(*t*)).

#### gKDR-GP

Determination of the dimension reduction matrix *B*_*i,j*_ was the same as above. Probabilistic distribution *y*_*i,j*_(*t* + *Δt*) ∼ *g*_*i,j*_(*U*_*ij*_(*t*)) was modeled using Gaussian Process. The fitrgp function of Matlab was used for this modeling.

#### Cross validation and simulation based on the models

Two-way cross validation was performed by splitting the time series into two parts, and a gKDR-GMM model was built on either half, in turn, and the model prediction 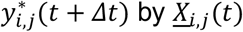 was compared to real data *y*_*i,j*_(*t* + *Δt*) in the latter half of the time series data. Pearson’s correlation constant was used for evaluation.

#### Freerun simulation

For simulation (also called freerun simulation), 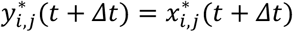 was predicted from *X*_*i,j*_(*t*), and the prediction was repeated for all *i* ‘s, which allows for embedding to generate 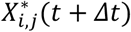 and 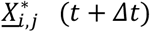 for all *i*, then 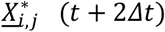 was predicted from 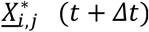 and this process was repeated using newly predicted values for further prediction.

#### Hyperparameters

Hyperparameter search was done by scoring similarity. Correlations of activity between neurons were represented by correlation matrices, and we scored similarity by calculating Mean Squared Error of correlation matrices of real and simulation results. Although the data length is typically 6000 time points, data for every five time points were used for gKDR analyses because for gKDR and GP, an inverse of *n* × *n* matrix needs to be calculated, *n* being the data number, and using *n*=6000 data points is impractical. For other hyperparameters, embedding steps and spans, *k*′ and *τ*′, the number of dimensionality reductions, *K*, in gKDR, and number of Gaussians in GMM (*κ*) were systematically searched for, and *k*′ = 30, *τ*′ = 10 (per original data with typical length 6000), and *K* = 3,4,5 were chosen. The number of gaussians were also varied and two Gaussians (*κ* = 2) showed the best performance in general, which we used (Figure 7 - Figure supplement 1F-H).

The total span of embedding is about the same as TDE-RICA, with wider spacing for gKDR. For hyperparameter search, freerun simulation with a length of 2000 steps was repeated five times for each set of hyperparameters.

## Supplementary Files

**Supplementary File 1**

Result of 4D imaging in 24 samples. Neuronal activities and their cross-correlations are shown.

SF1_4D imaging results.pdf

**Supplementary File 2**

Results of TDE-RICA for 94 selected neurons of 10 selected samples.

SF2_TDE-RICA_ver0.pdf

**Supplementary File 3**

Results of TDE-RICA with matrix factorization for all 177 neurons of all 24 samples, including neural activity data for all 24 samples obtained by the whole-brain imaging experiments.

Page 1-24: The experimental neural activity data and the results of TDE-RICA with matrix factorization in each sample.

Page 25-48: The phase diagram of all pairs of the motif occurrences in each sample.

Page 49: The motifs obtained by TDE-RICA with matrix factorization.

Page 50: Cross-correlation functions of motif occurrence, averaged across samples.

Page 51-141: Time course and phase diagram of all pairs of the motif occurrences in each sample.

SF3_TDE-RICA-MF_ver0.pdf

**Supplementary Files 4A-C (related to Figure 7)**

Simulation results of gKDR-GMM

SF4A/B/C_simulation_results_by_gKDR-GMM_K=3/4/5.pdf

**Supplementary Files 5A-C**

Simulation results by gKDR-GP

SF5A/B/C_simulation_results_by_gKDR-GP_K=3/4/5.pdf

**Supplementary File 6A (related to Figure 8)**

Lagged cross-correlation of all combinations of neurons for all samples

SF6A_lagged_cross_correlation_real_and_gKDR-GMM.pdf

**SupplementaryAdditional File 6B (related to Figure 8A, B)**

Activity profile of example neuron pairs with lagged correlation

SF6B_example_neuron_pairs_with_lagged_correlation.pdf

**Supplementary File 7**

Simulation results by gKDR-GMM, evaluated by TDE-RICA

SF7_test2022105_compare_real_and_simulation_by_tderica.pdf

**Supplementary File 8 (related to Figure 8C)**

Shifted overlay to visualize same periodicity as sensory input

SF8_periodic_activity_of_real_and_simulated.pdf

**Supplementary File 9 (related to Figure 10)**

Probabilistic and deterministic models A: Simulation result using probabilistic model, deterministic model and deterministic model with random noise

SF9A_probabilistic_vs_deterministic_hearmap.pdf

B: Cross correlation of results in A

SF9B_probabilistic_vs_deterministic_correlation_matrix.pdf

**Supplementary File 10 (related to Figure 10)**

Probabilistic and deterministic models in the gKDR-GP model

SF10_gKDR_GP_detprobabilistic_vs_deterministic.pdf

**Supplementary File 11A-F**

A-C: Synaptic connection estimated from gradients of the input-output function shown as a matrix

SF11A/B/C_connection_weight_matrix_K3/4/5.pdf

D-F: Synaptic connection weights estimated from gradients of the input-output function shown as tables

SF11D/E/F_connection_weight_matrix_K3/4/5.xls

**Supplementary File 12**

Synaptic weight map for the forward-backward circuit

SF12A/B/C_Forward_backward_network_K3/4/5.pdf

**Supplementary File 13**

Effect of cell ablation on top positive and negative correlated pairs

SF13A/B_pospairs_nosalt_indirect_table_K3/5.xlsx

SF13C/D_negpairs_nosalt_indirect_table_K3/5.xlsx

**Supplementary File 14**

Synaptic weight map from the salt sensory input to forward-backward circuit

SF14A/B/C_Sensory_and_forward_backward_network_K3/4/5.pdf

**Supplementary File 15**

Effect of blocking sensory input to individual cells SF15_effect_of_killing_on_AVA_periodicity.pdf

**Supplementary File 16**

Main Figures 1, 7-12 and their supplements are included.

SF16_Main_Figures_and_Figure_Supplements_1_7-12.pdf

**Supplementary File 17**

Main Figures 2-6 and 13 and their supplements are included.

SF17_Main_Figures_and_Figure_Supplements_2-6_13.pdf

## Acknowledgement

The calculations were mainly performed in the Shirokane supercomputer at the Human Genome Center at the Institute of Medical Science, the University of Tokyo.

## Notes

### Competing Interest Statement

The authors have declared no competing interest.

