## Supplementary figures and images for "Deducing ensemble dynamics and information flow from the whole-brain imaging data"

### Supplementary File 1

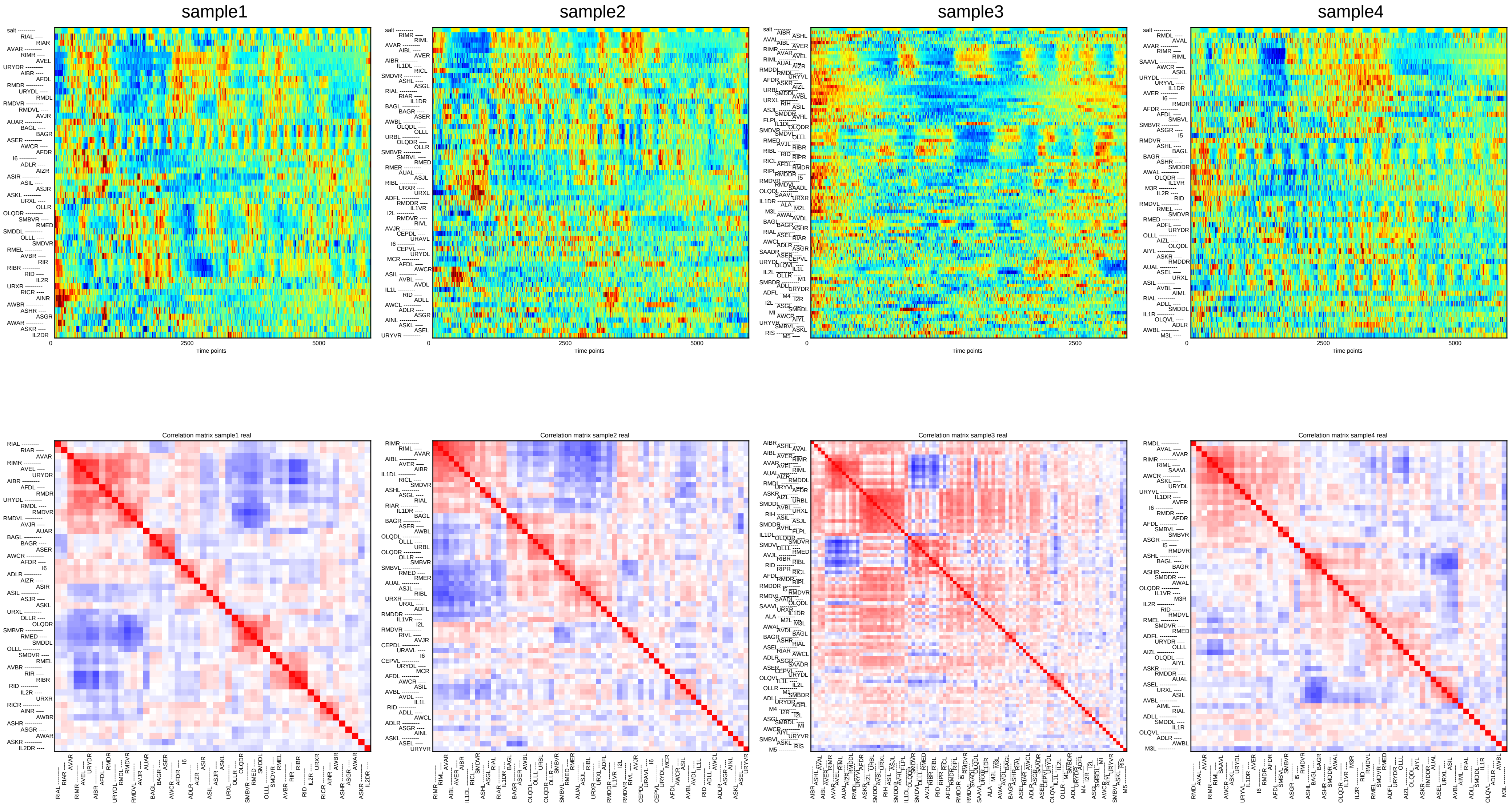

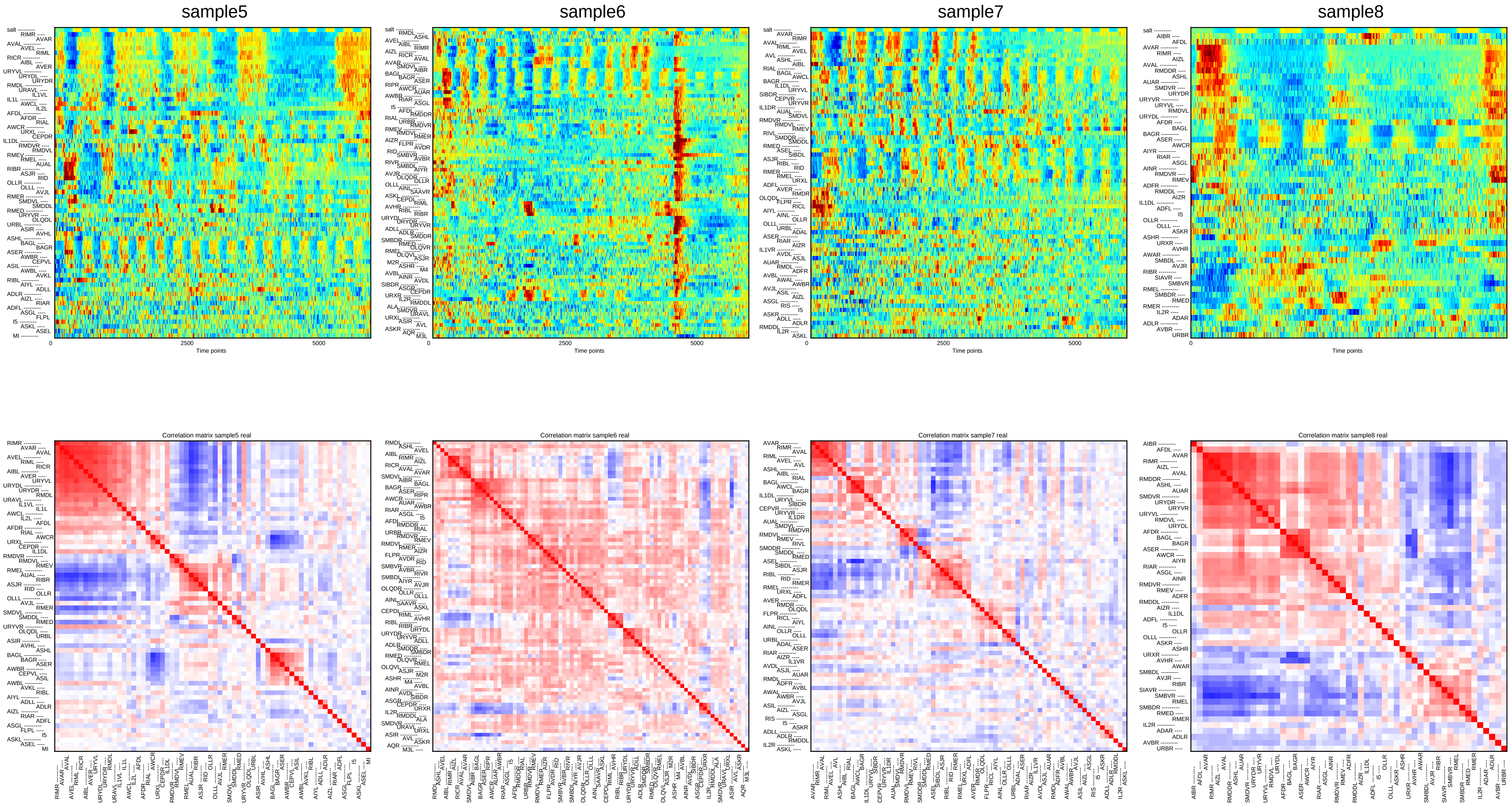

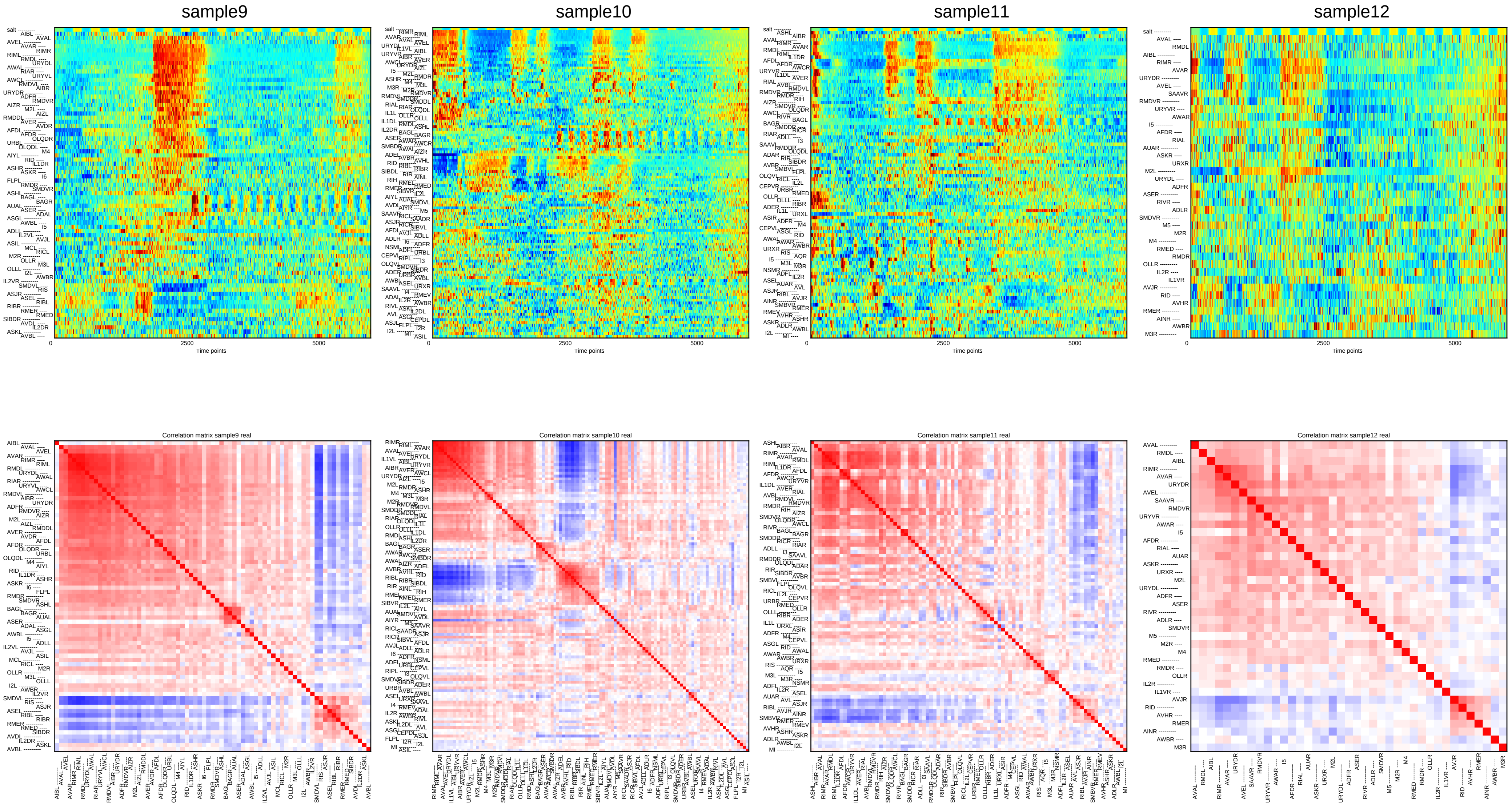

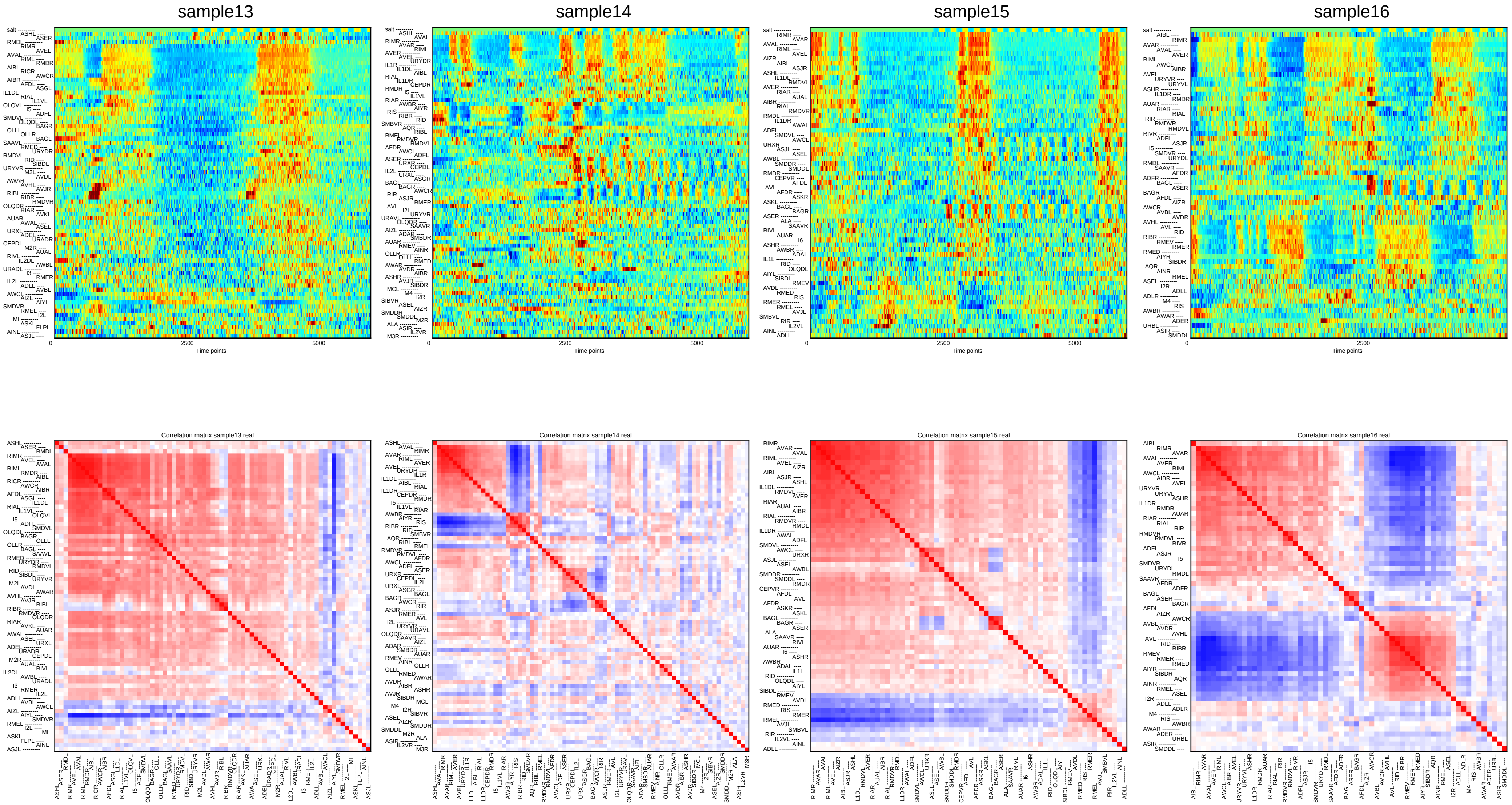

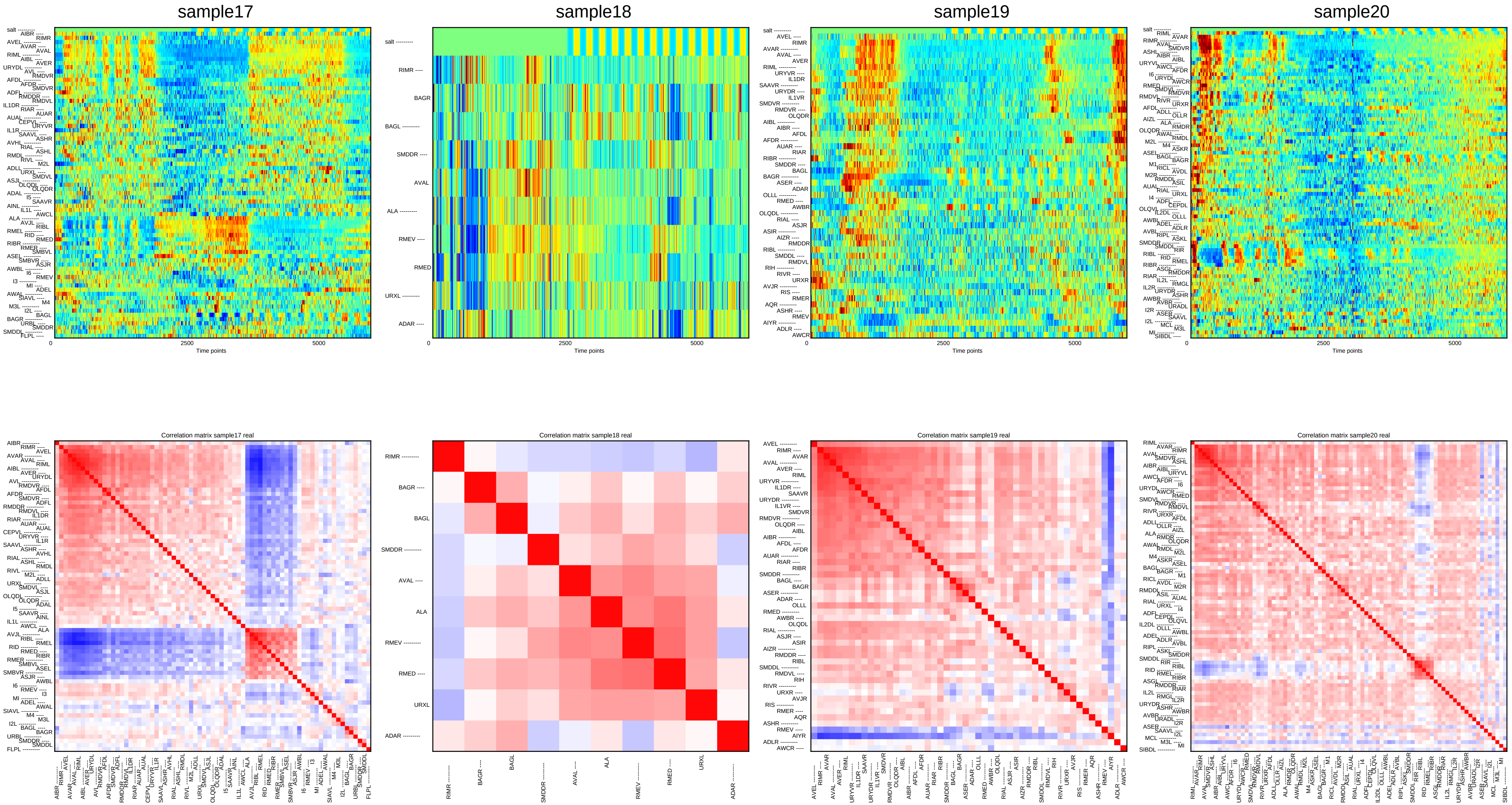

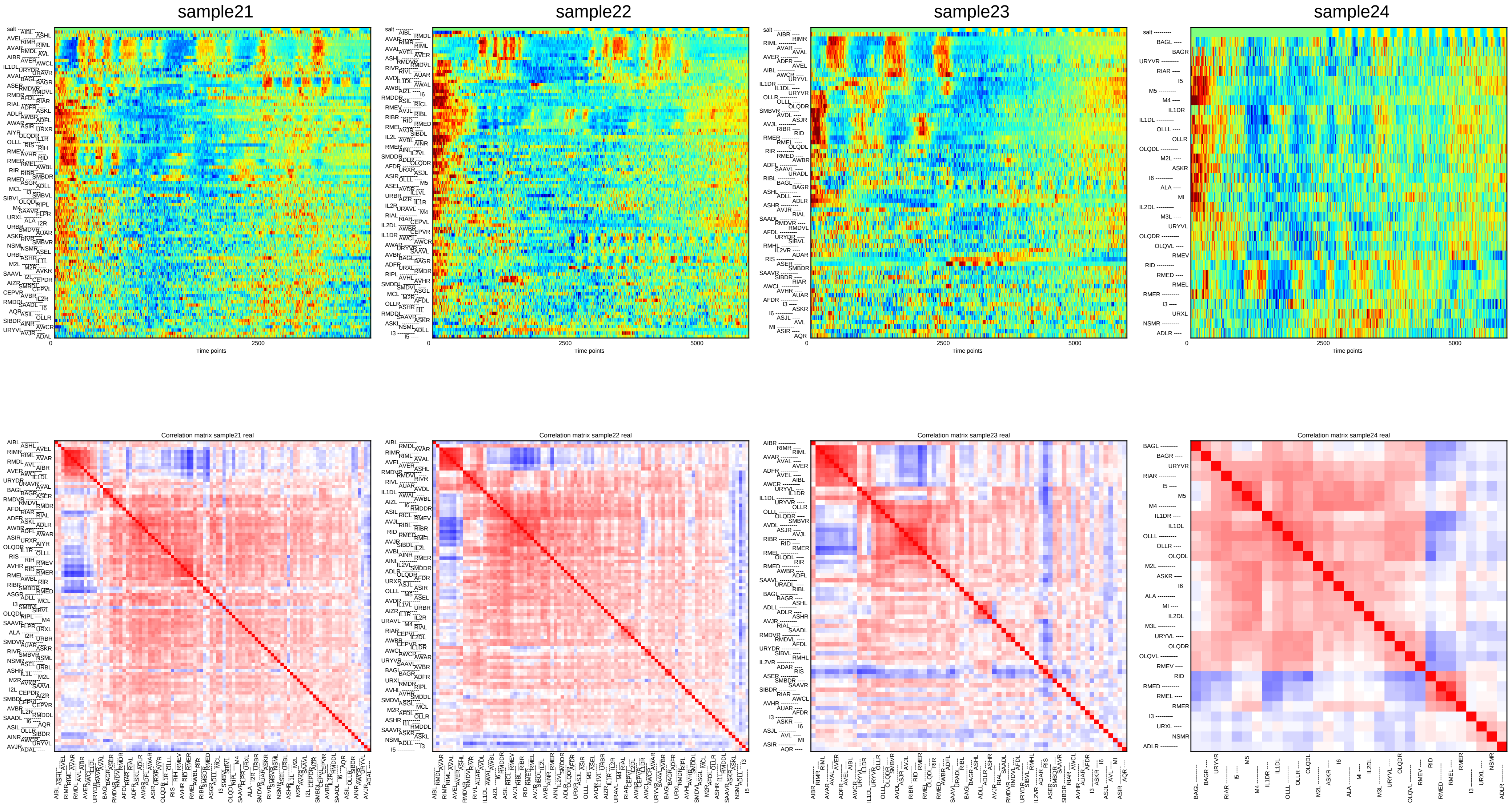

### Supplementary File 6A

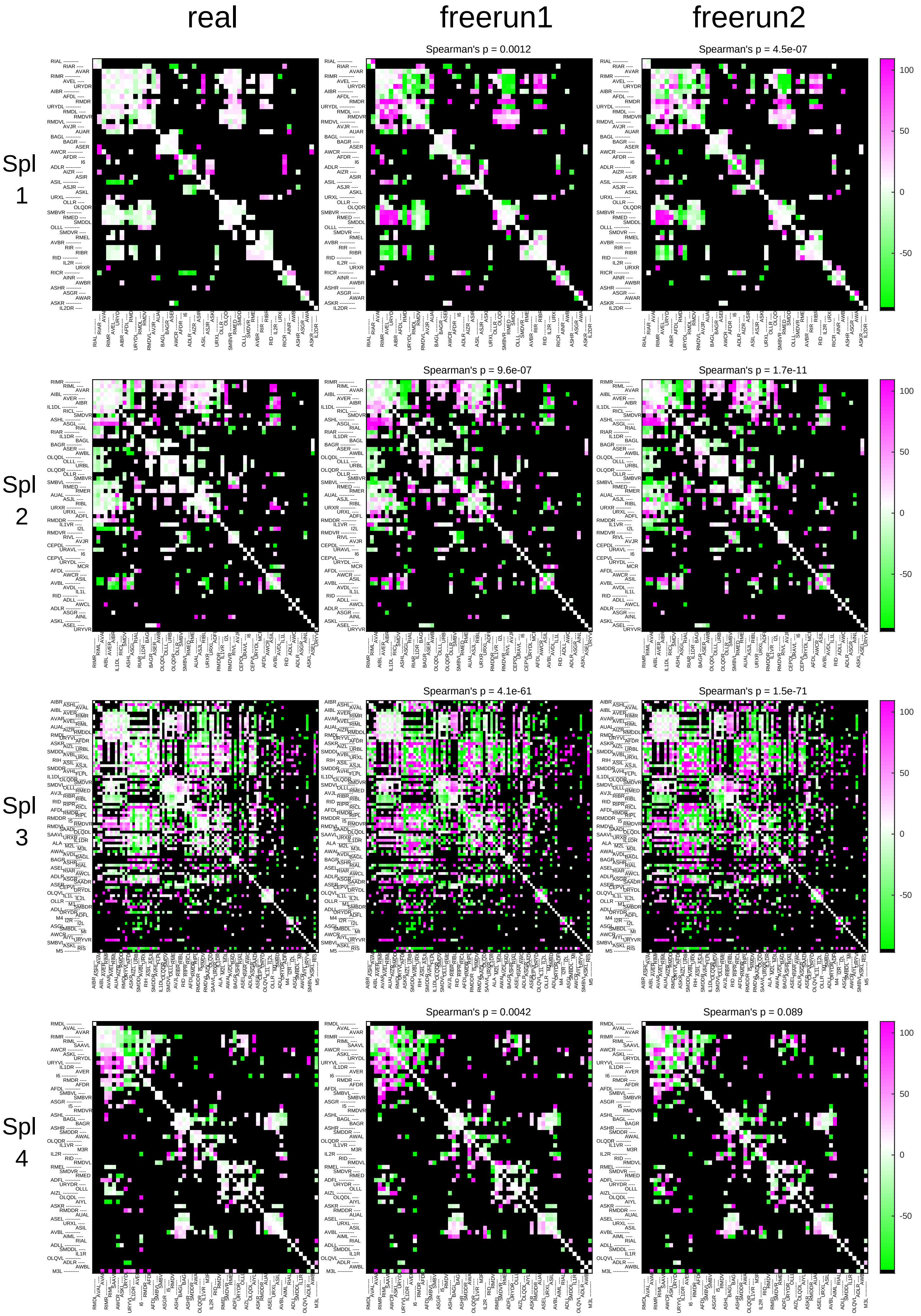

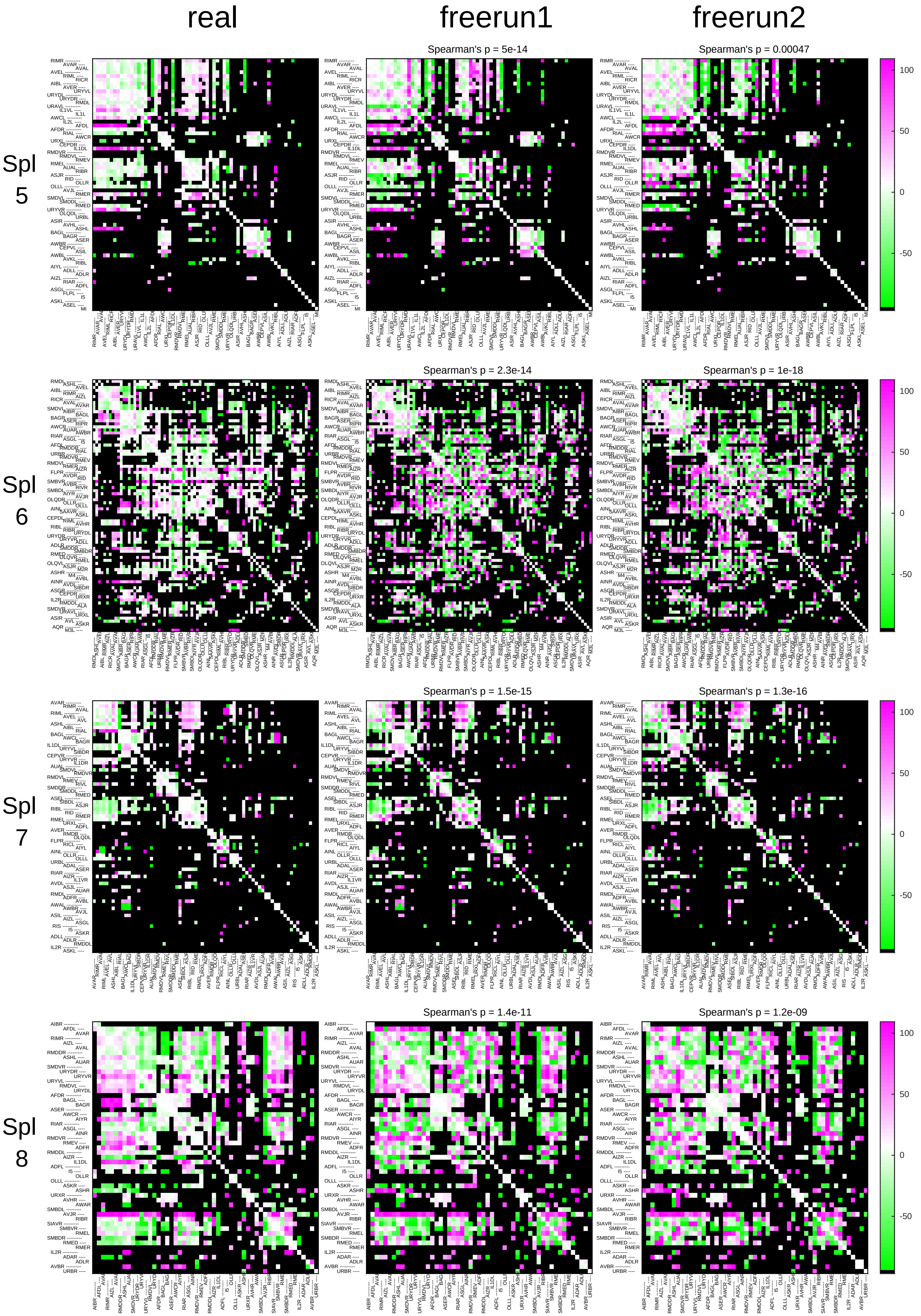

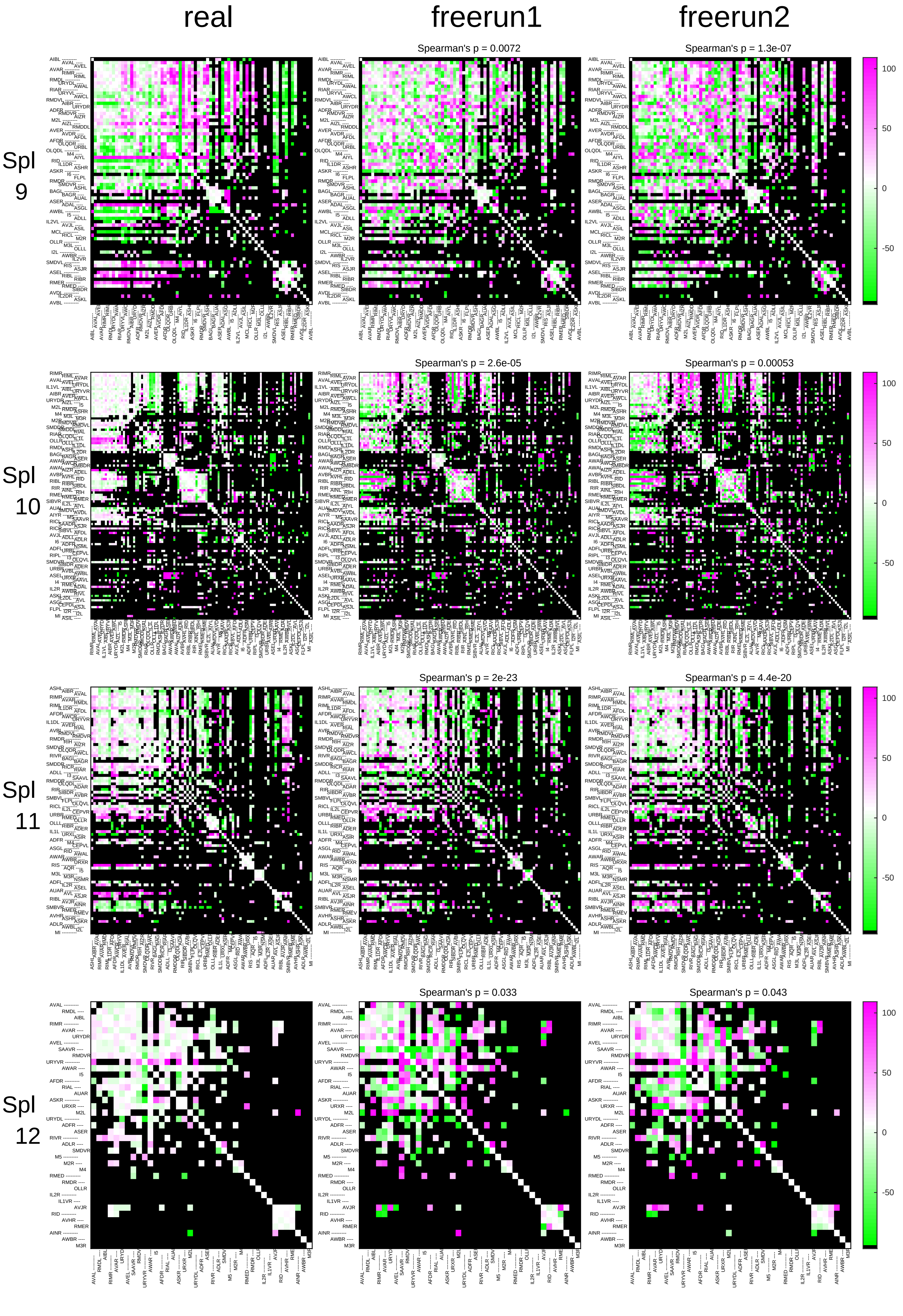

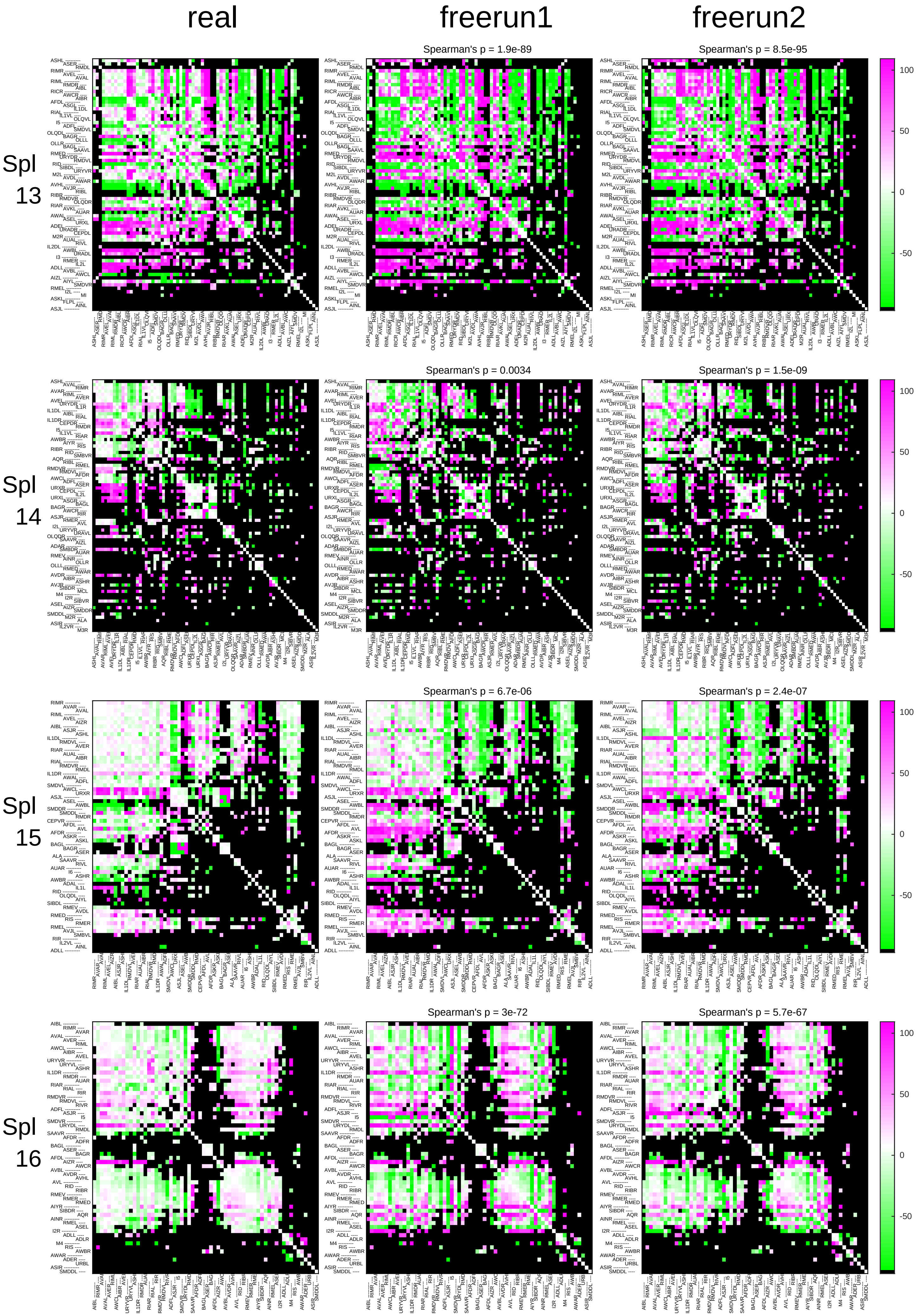

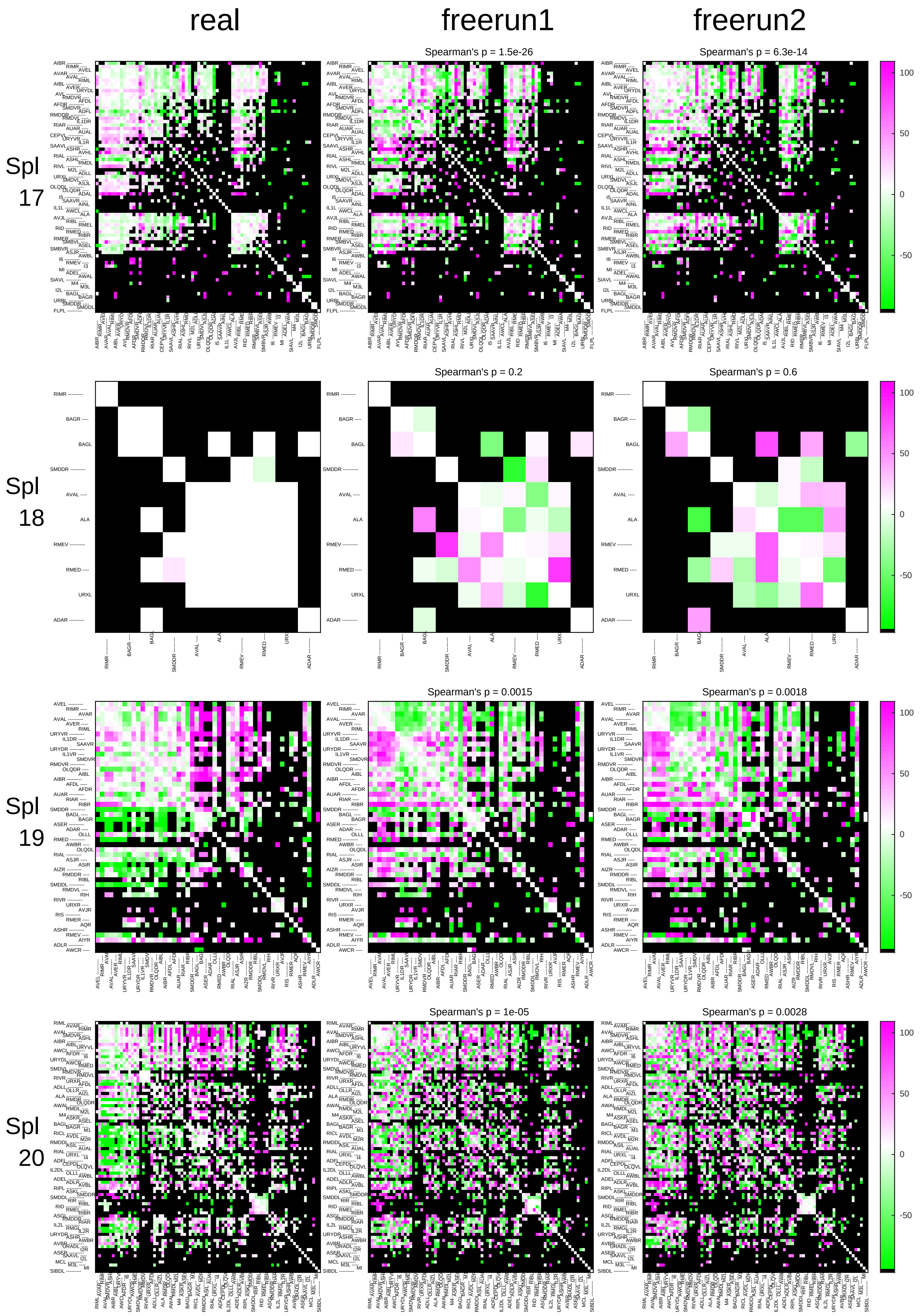

### Supplementary File 8

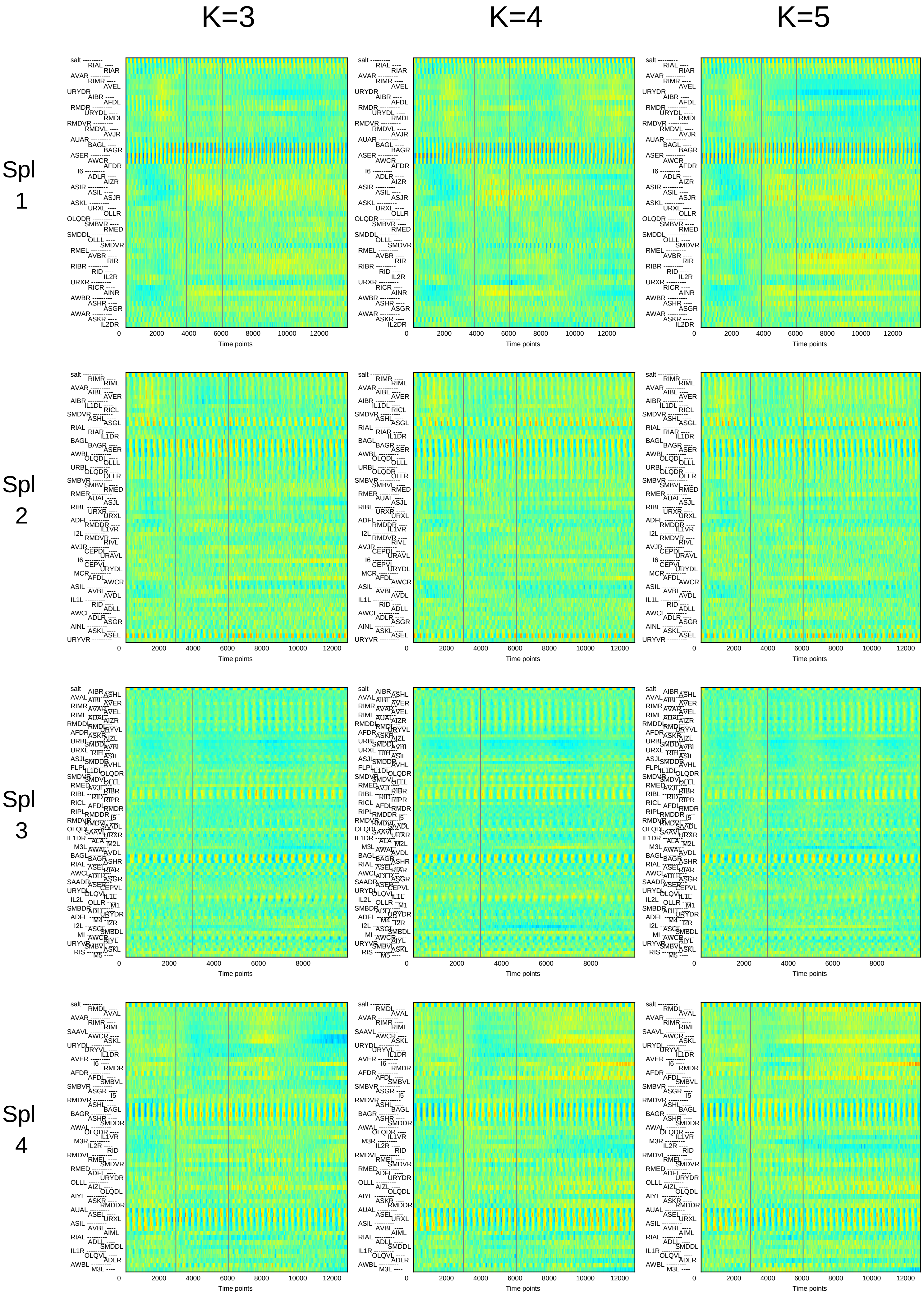

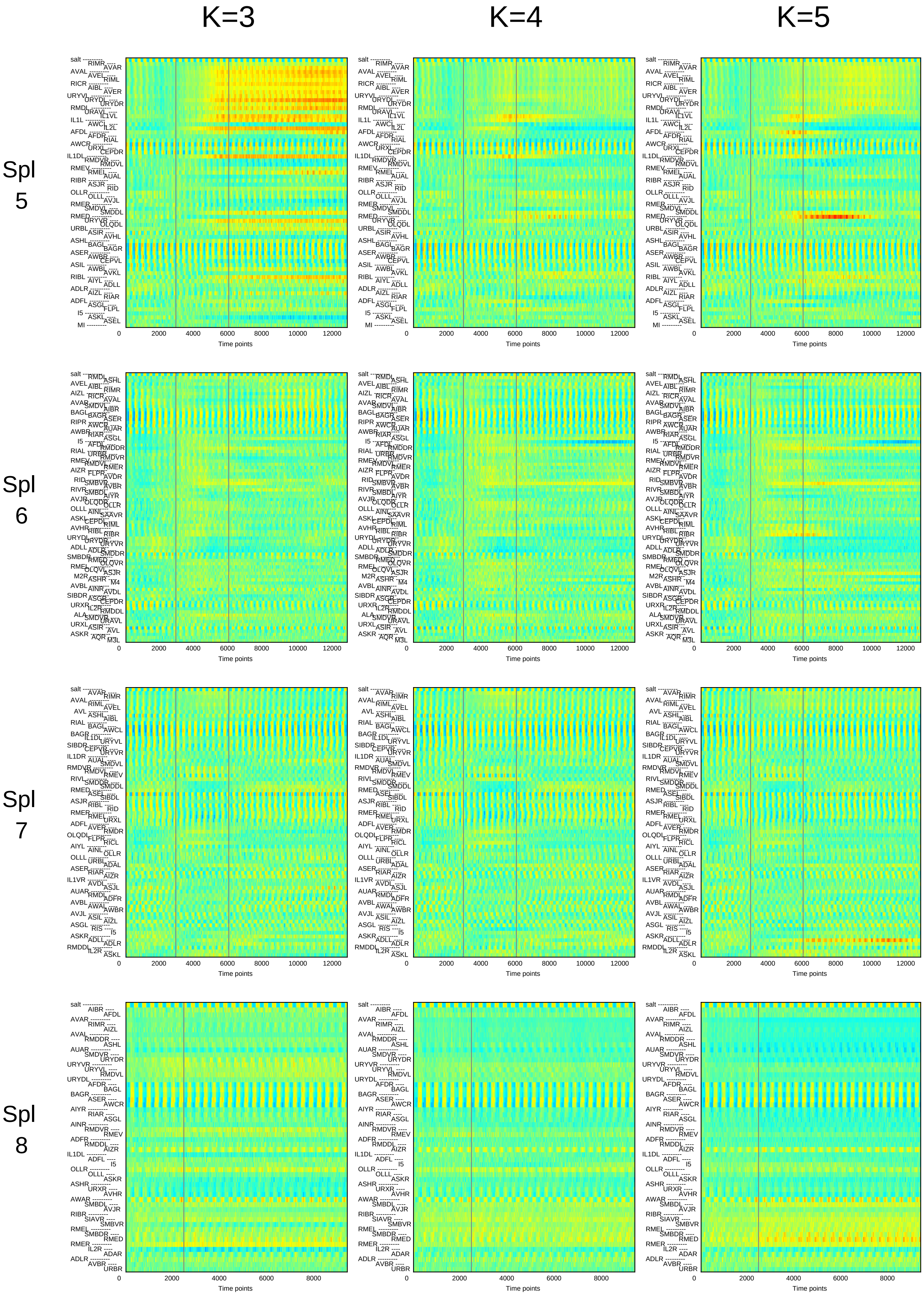

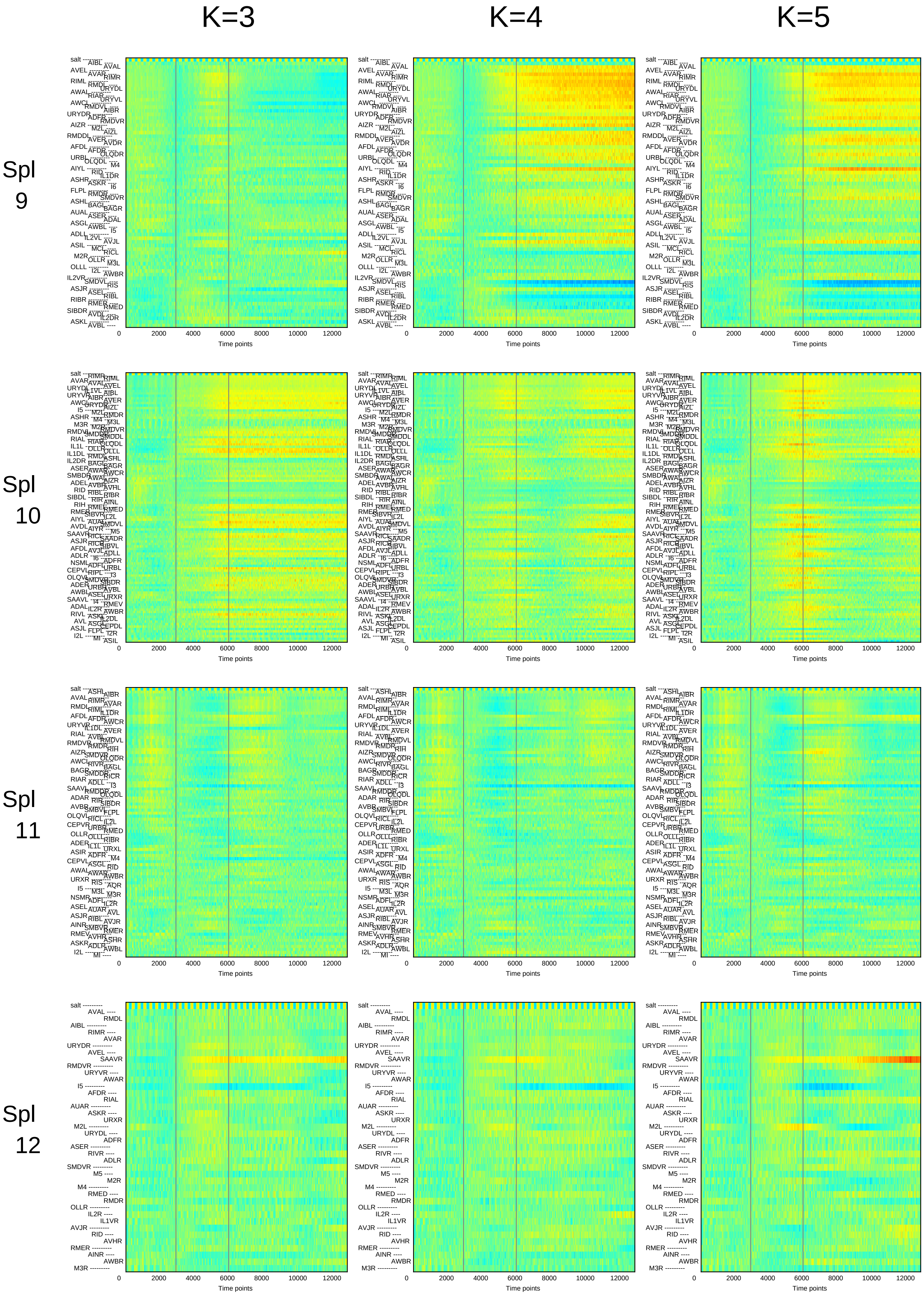

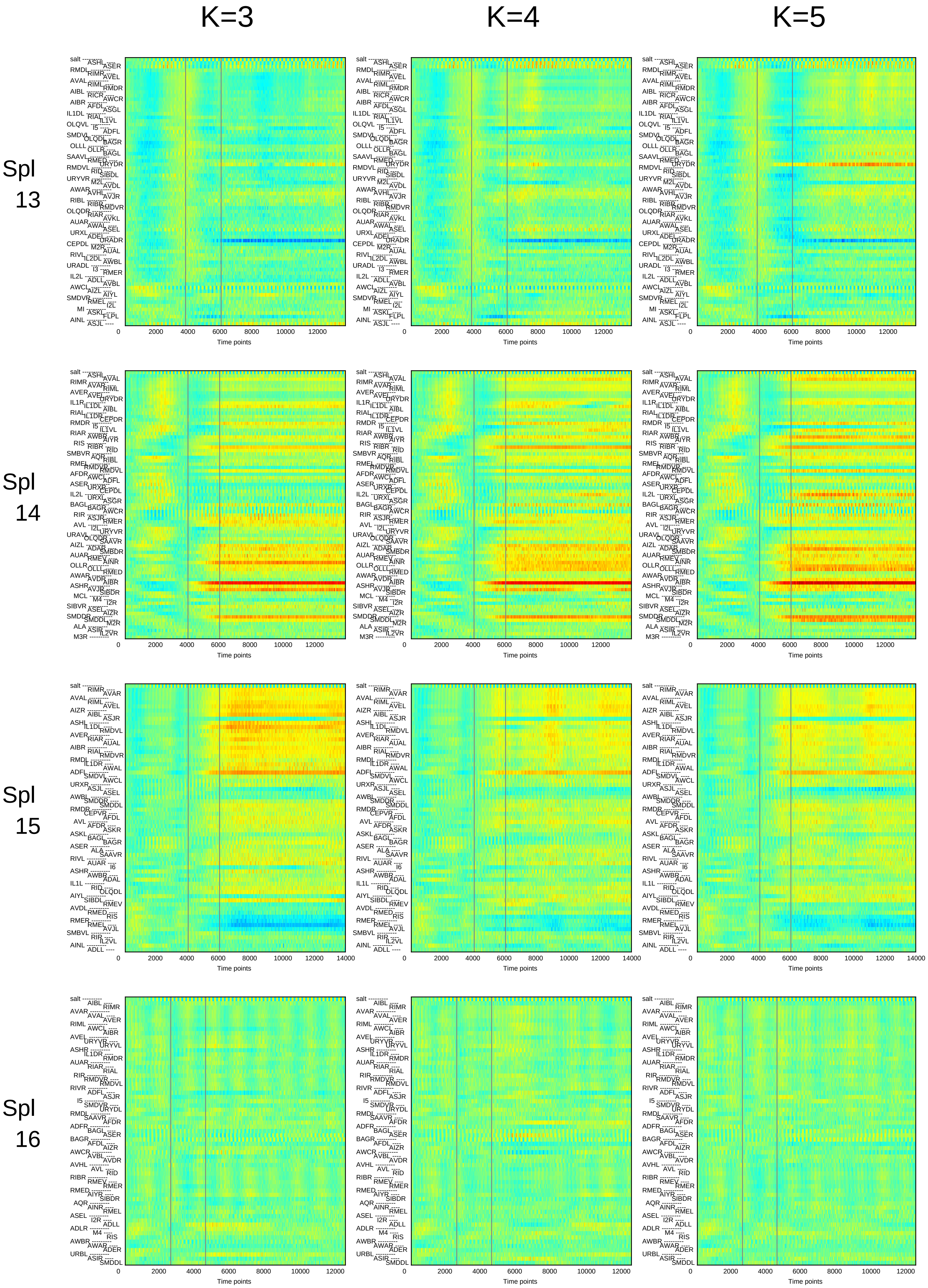

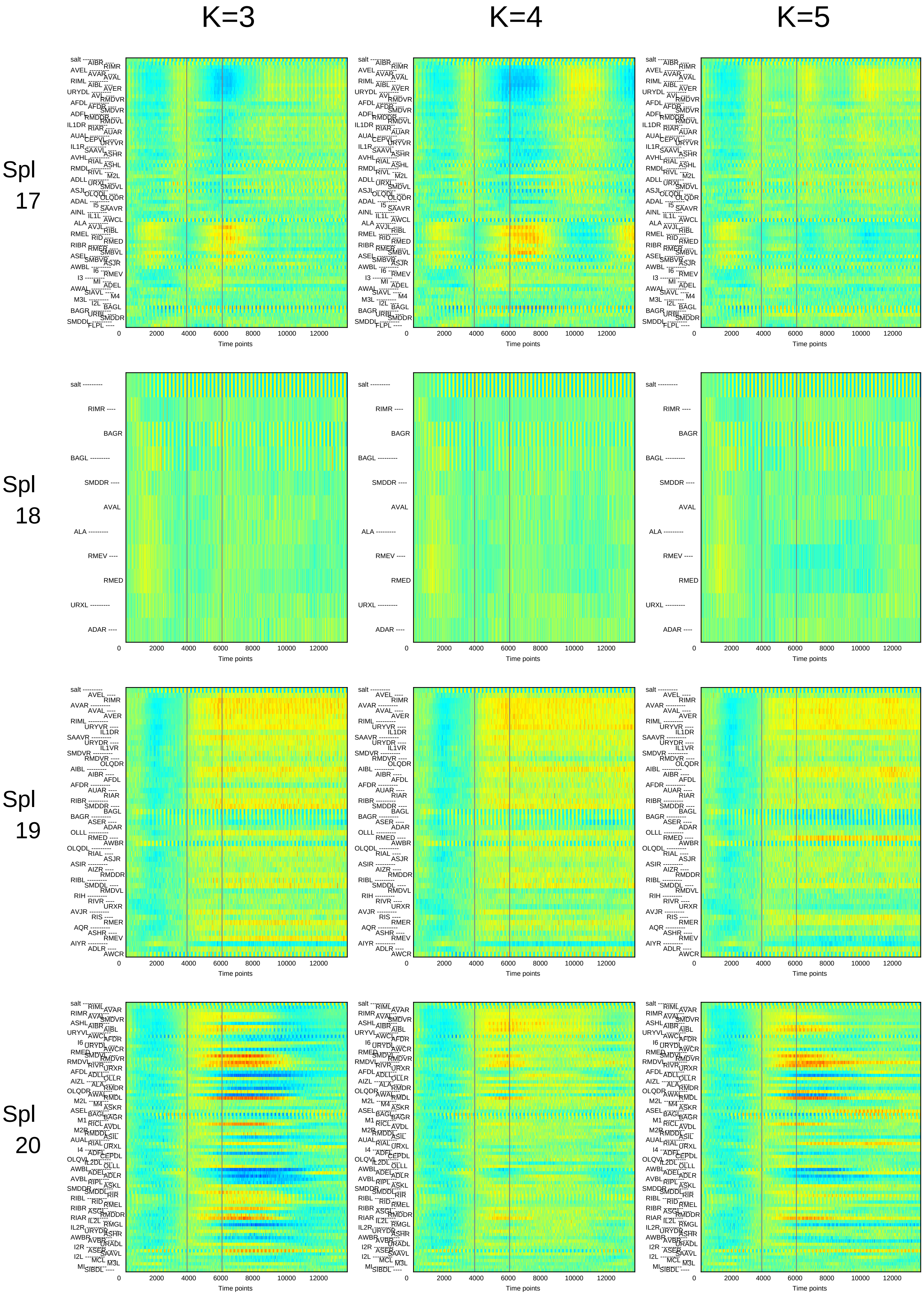

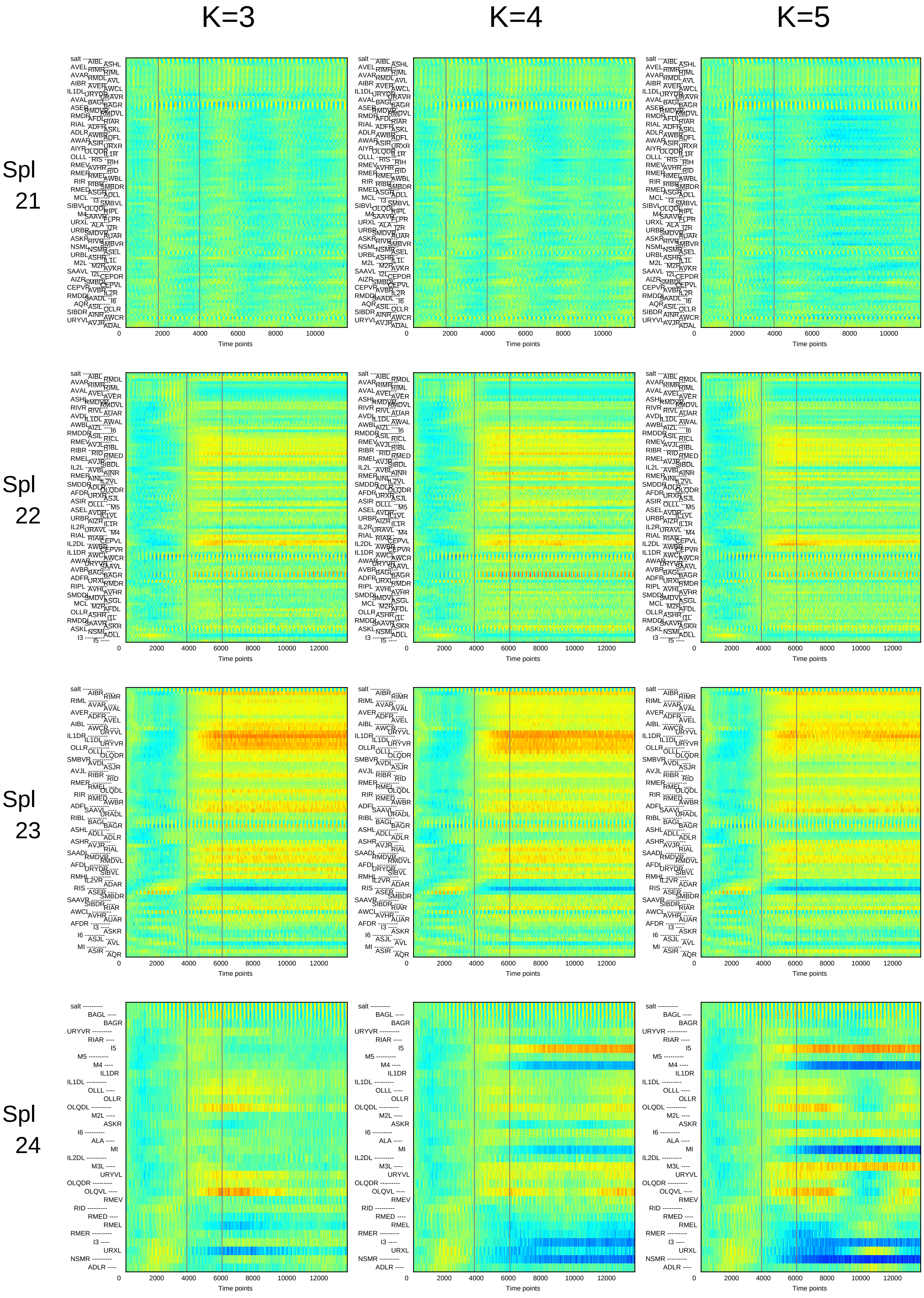

### Supplementary File 12A

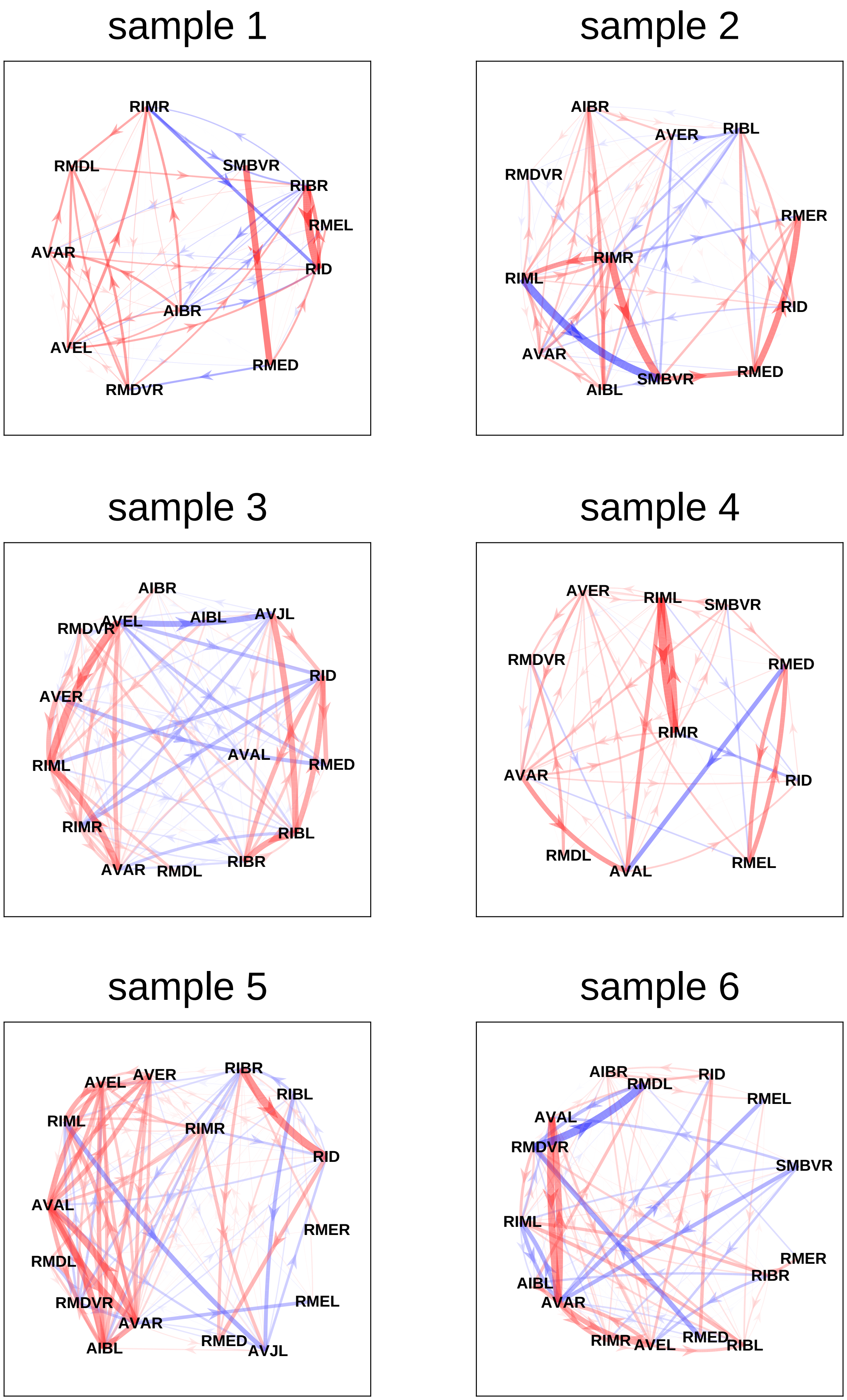

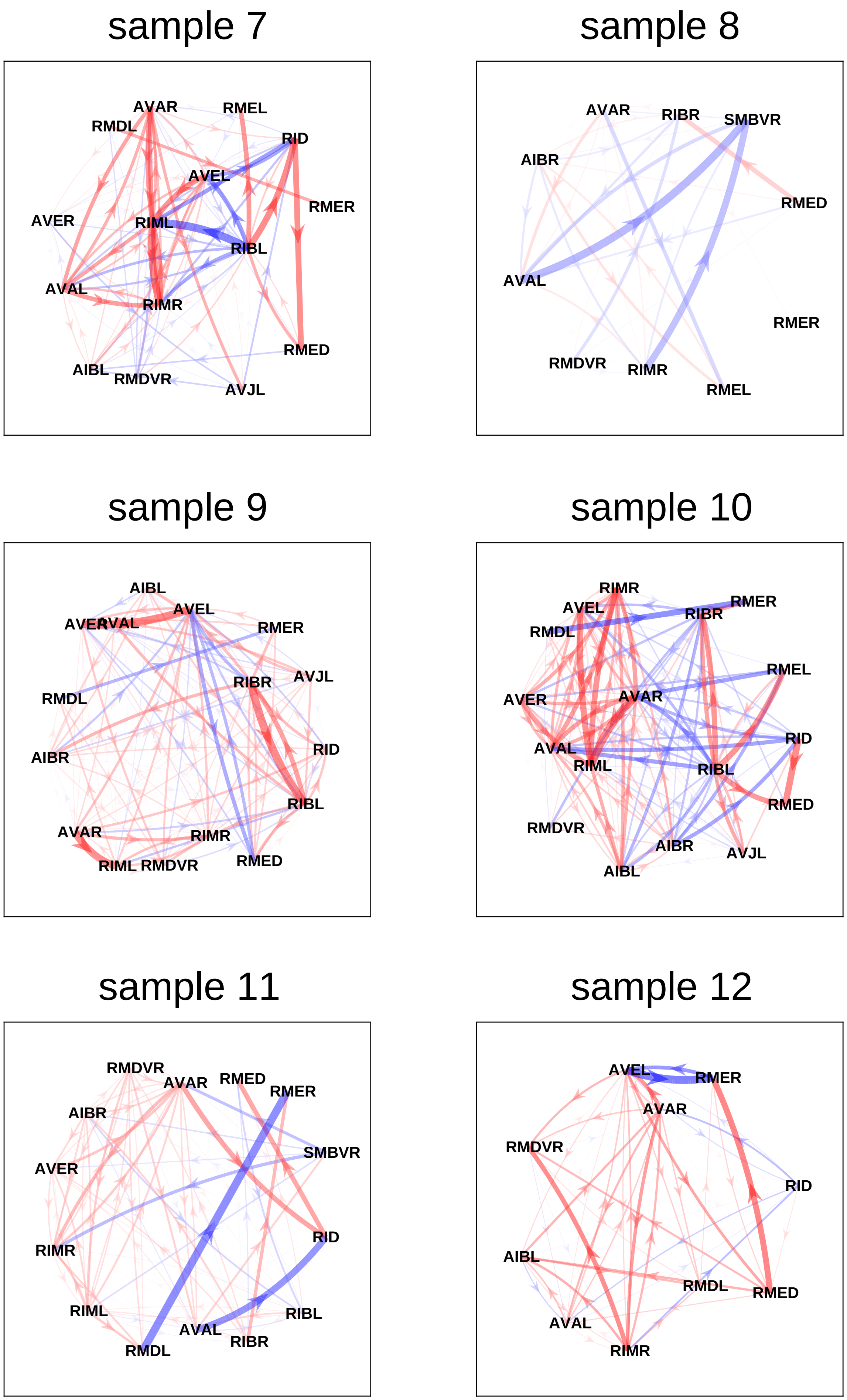

sample 13

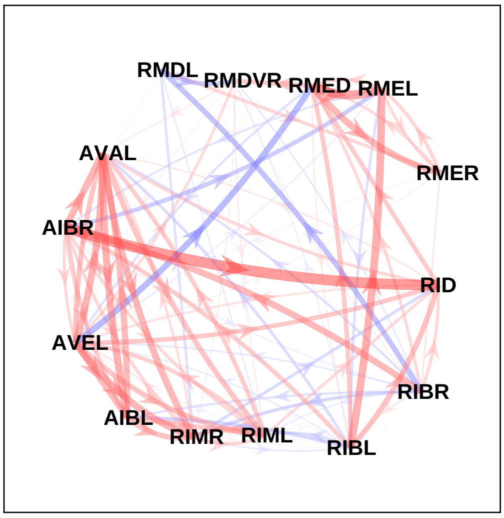


sample 14

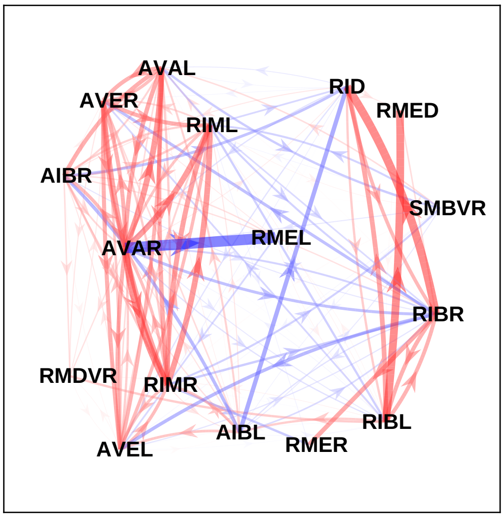


sample 15

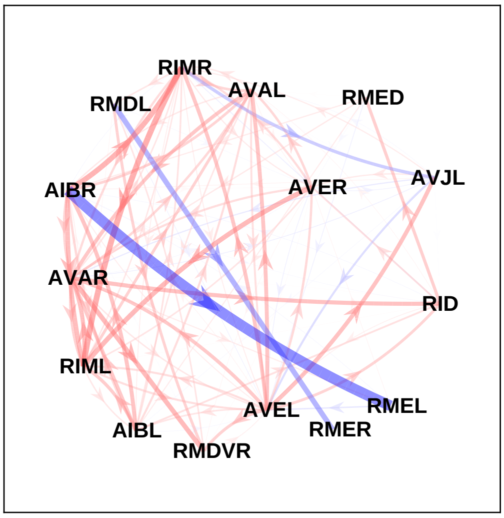


sample 16

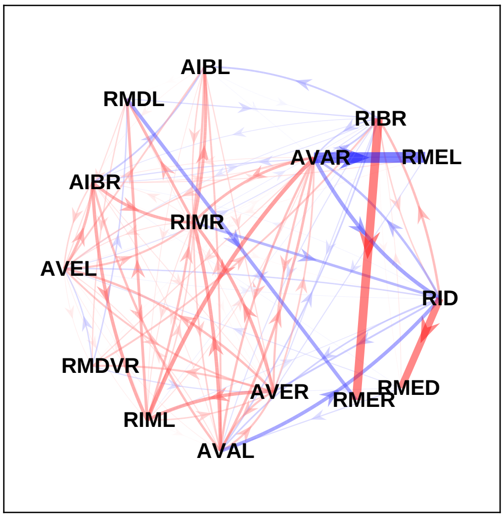


sample 17

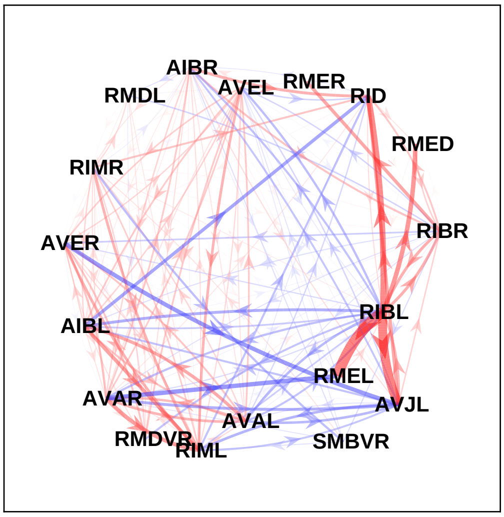


sample 18

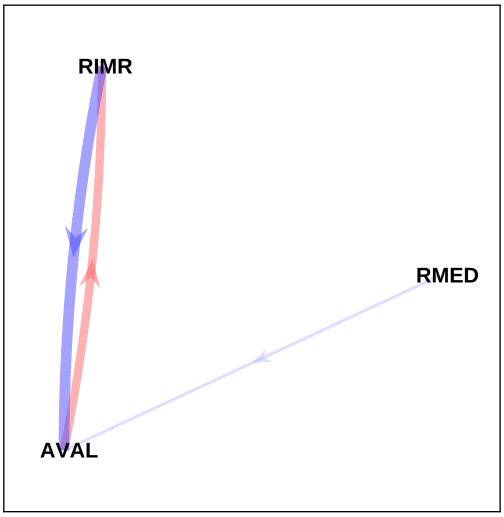

#### sample 19

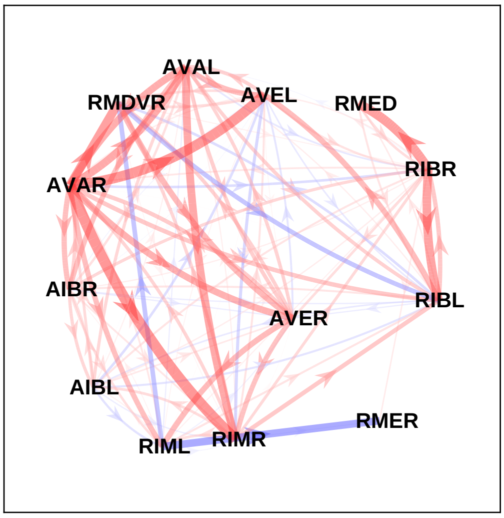


#### sample 20

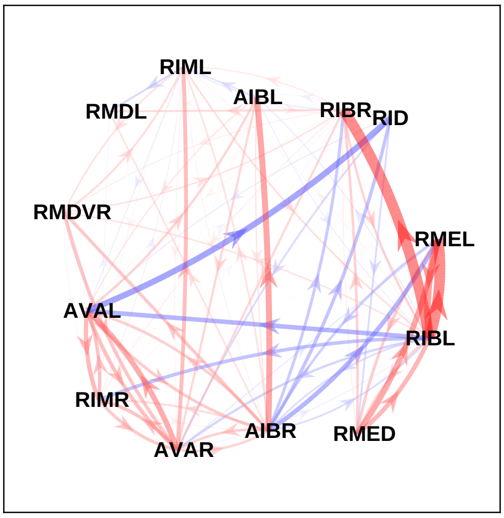


#### sample 21

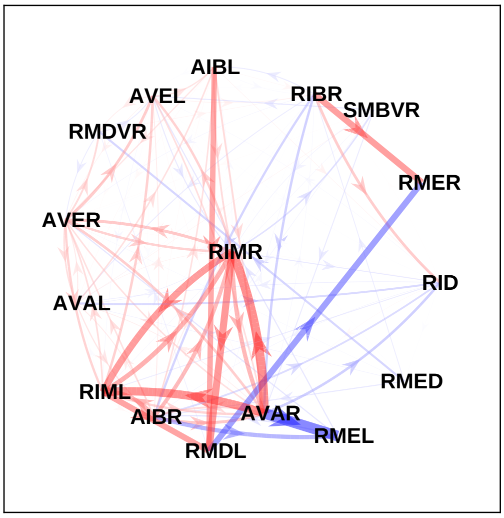


#### sample 22

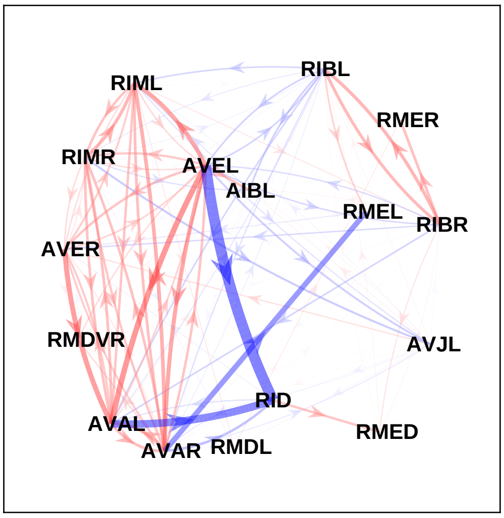


#### sample 23

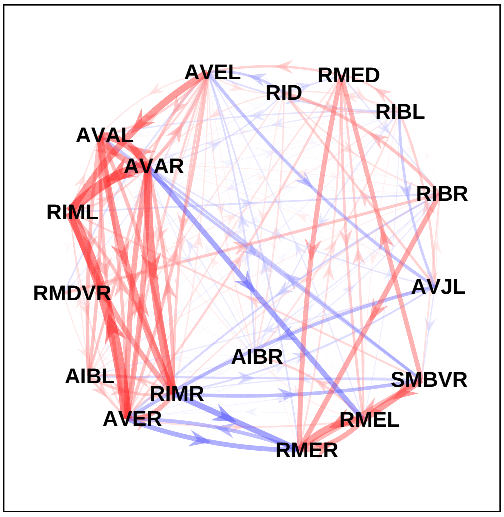


#### sample 24

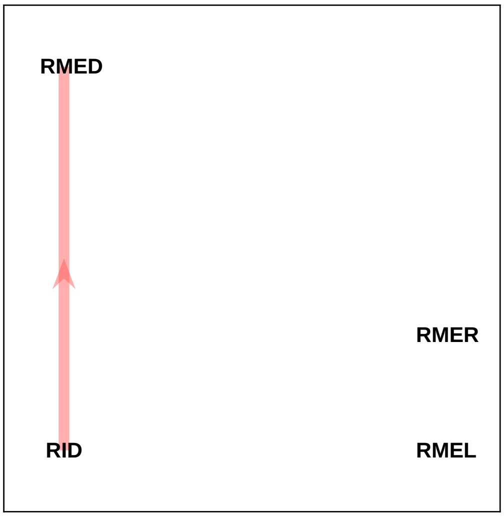

### Supplementary File 12C

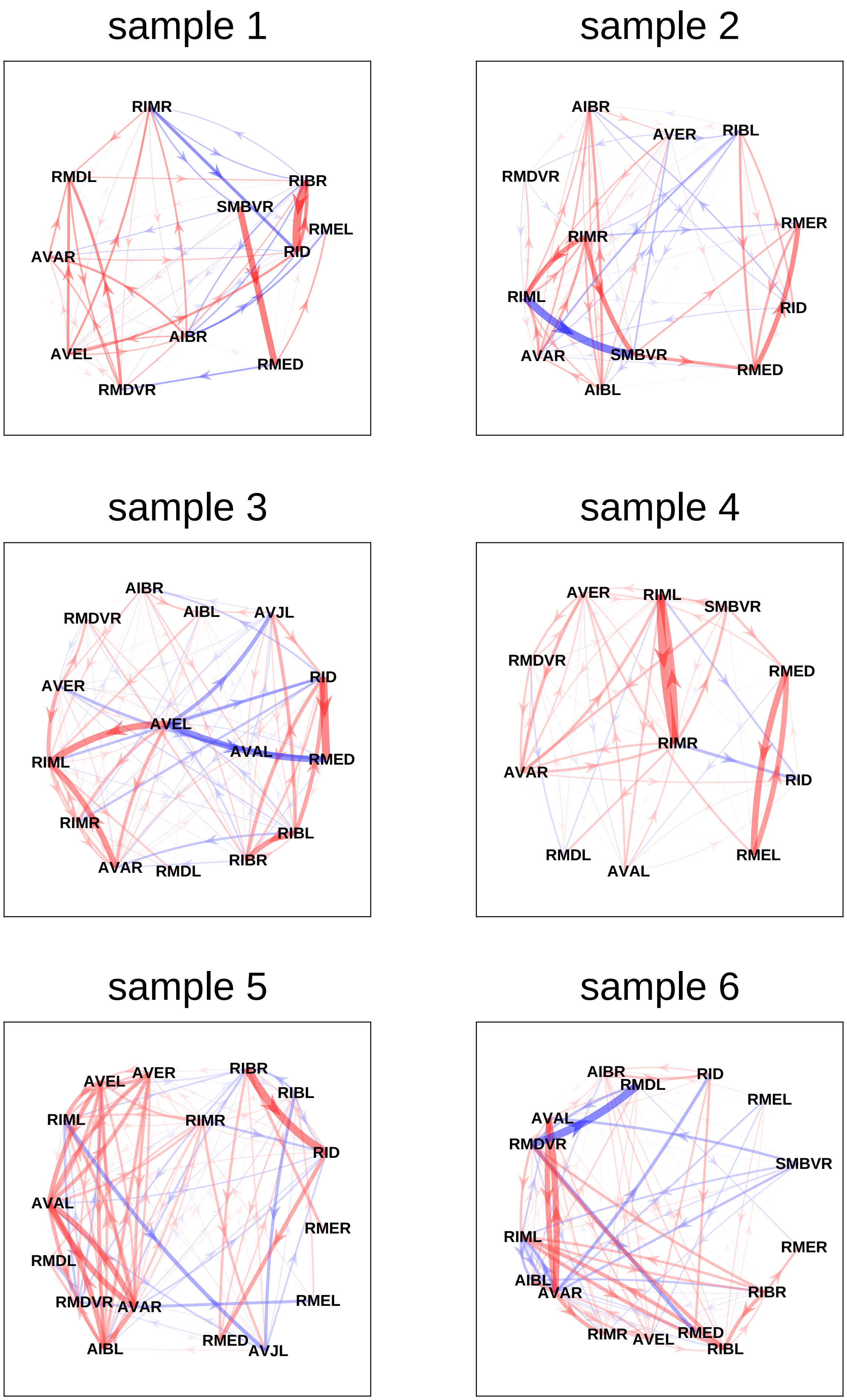

sample 7

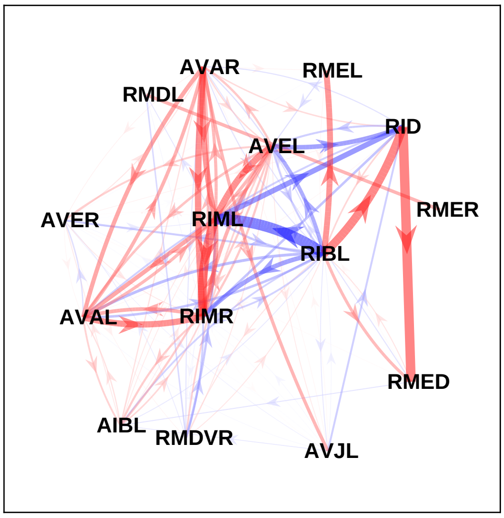


sample 8

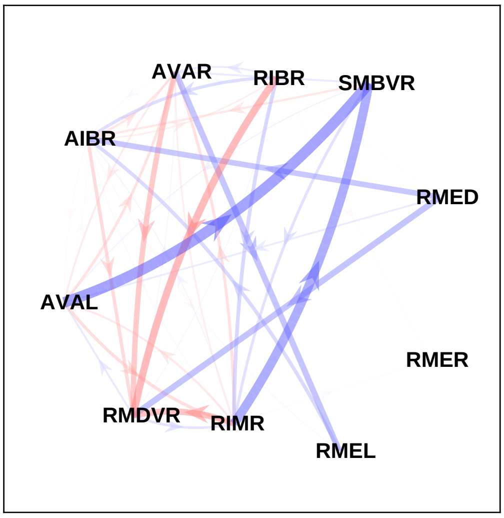


sample 9

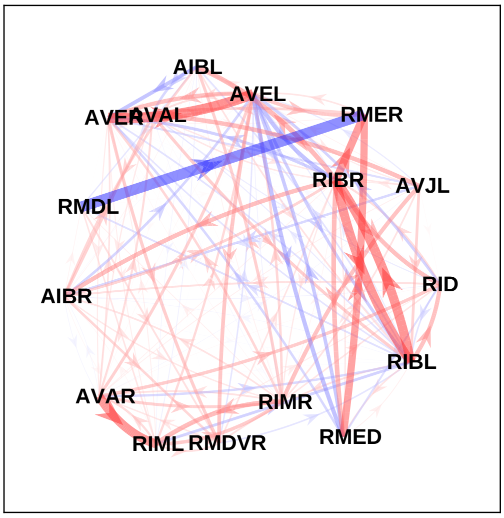


sample 10

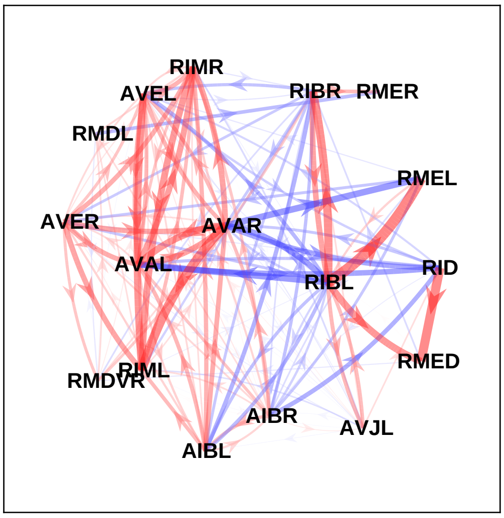


sample 11

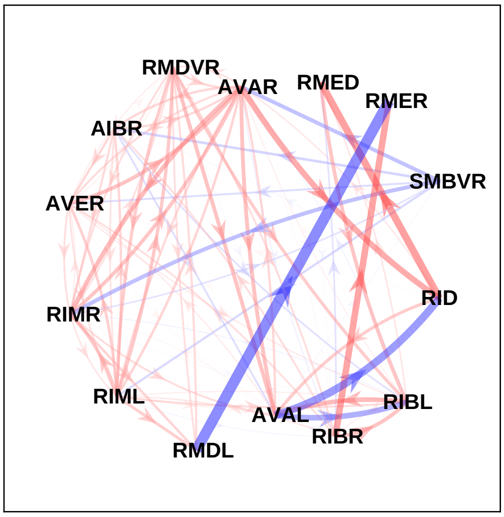


sample 12

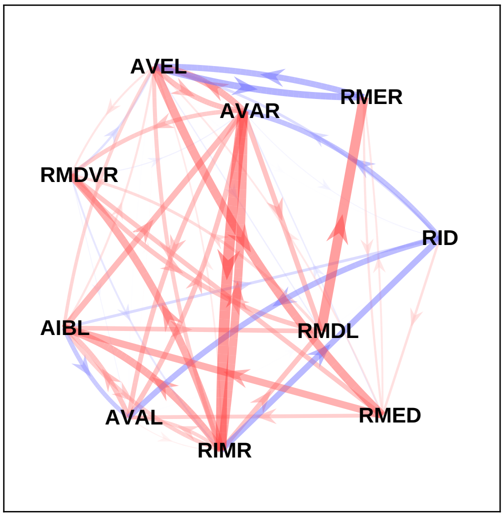

sample 13

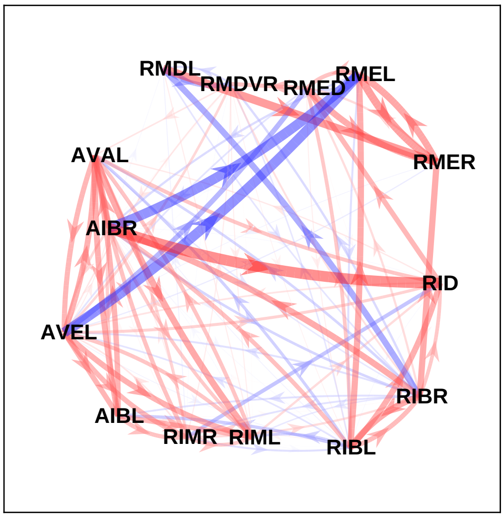


sample 14

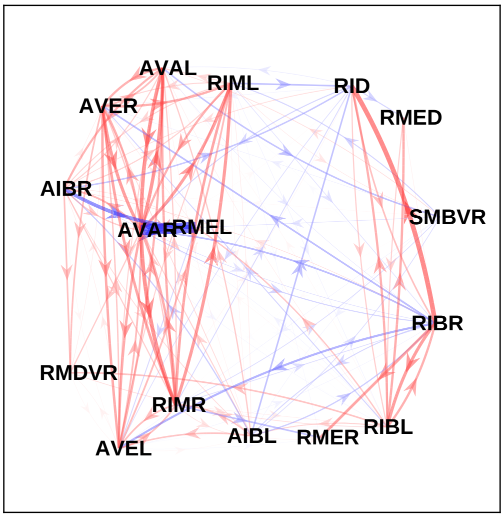


sample 15

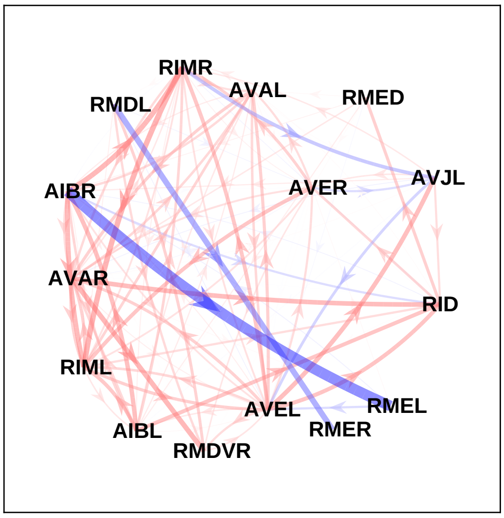


sample 16

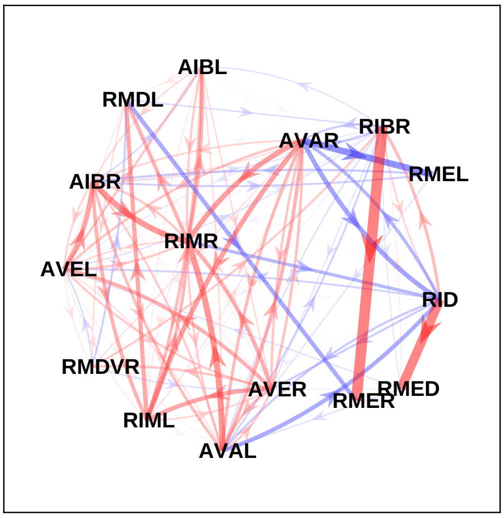


sample 17

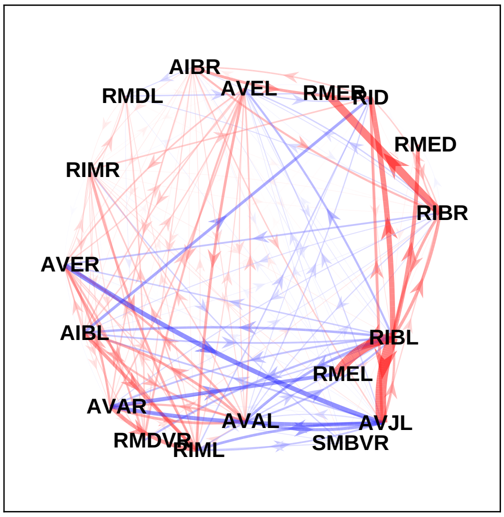


sample 18

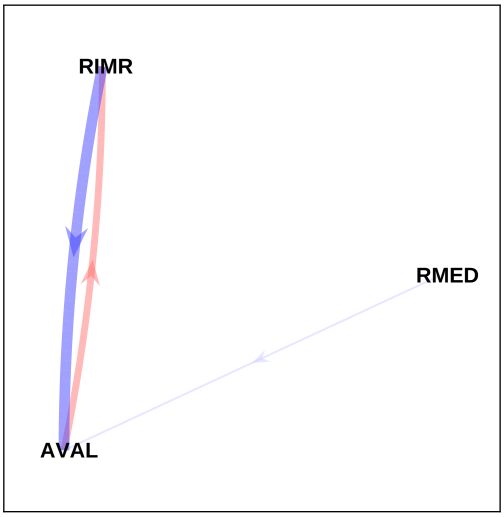

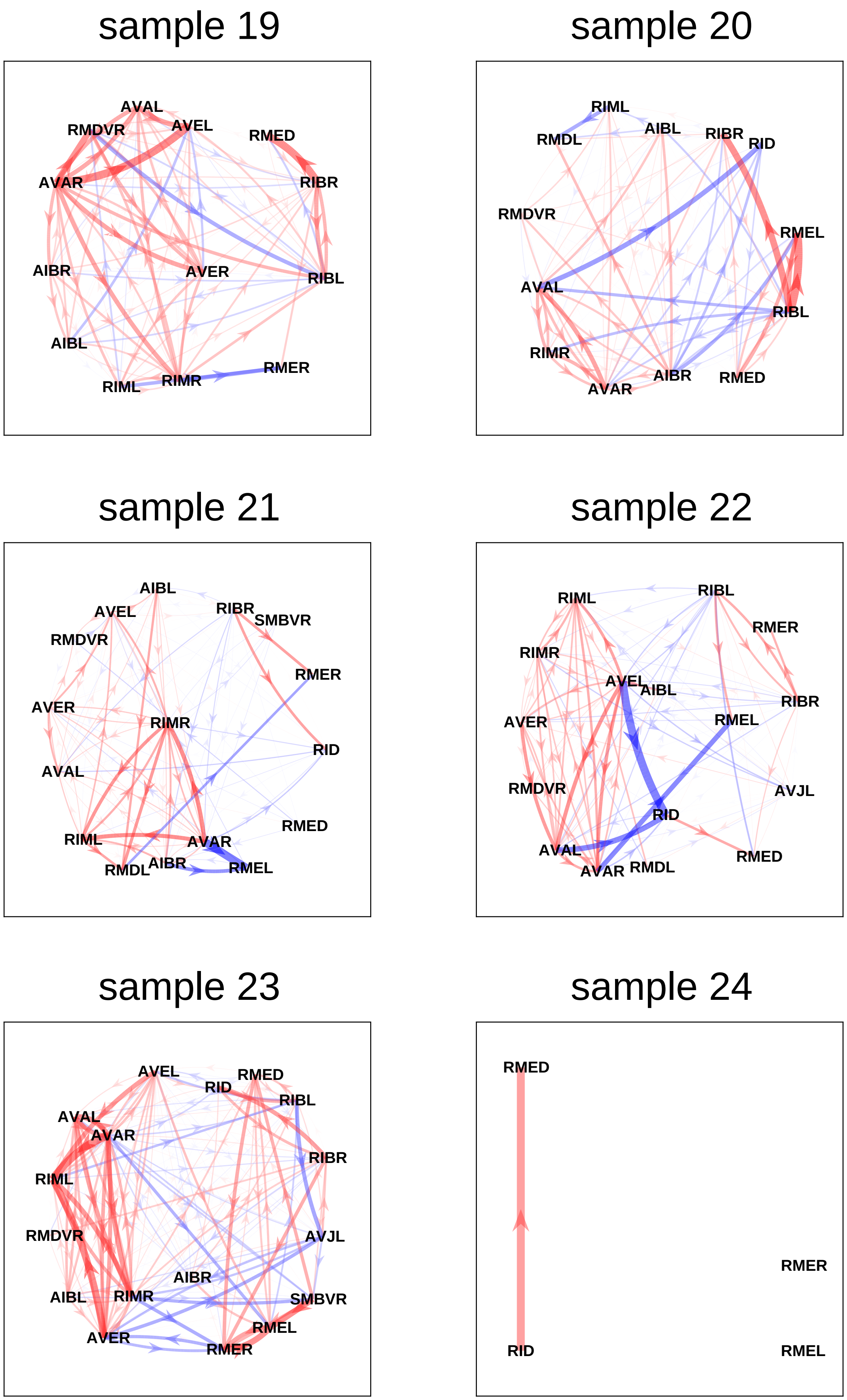
