## Supplementary File 2 for "Deducing ensemble dynamics and information flow from the whole-brain imaging data"

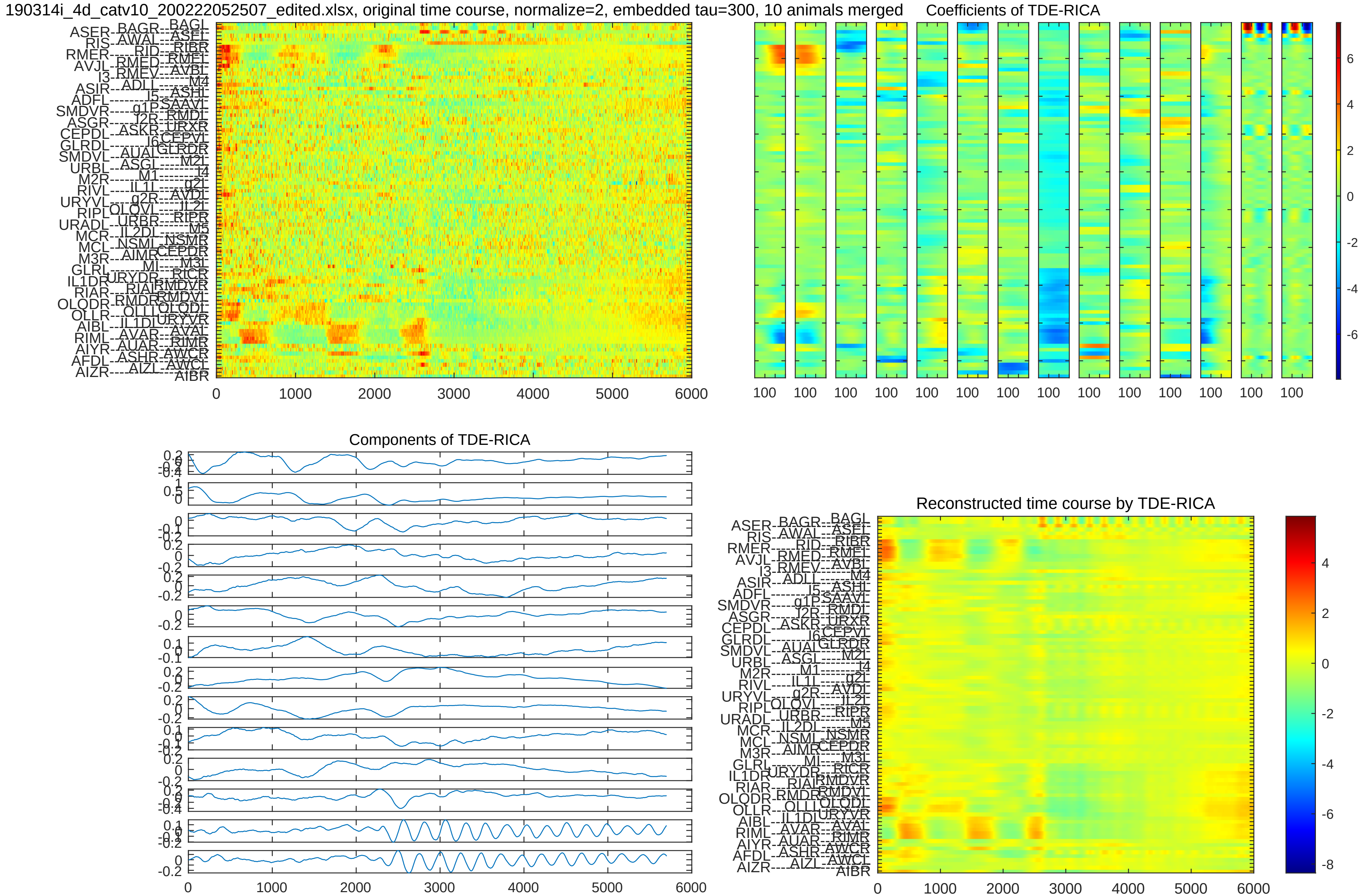
190314i_4d_catv10_200222052507_edited.xlsx, original time course, normalize=2, embedded tau=300, 10 animals merged
Coefficients of TDE-RICA
Components of TDE-RICA
Reconstructed time course by TDE-RICA

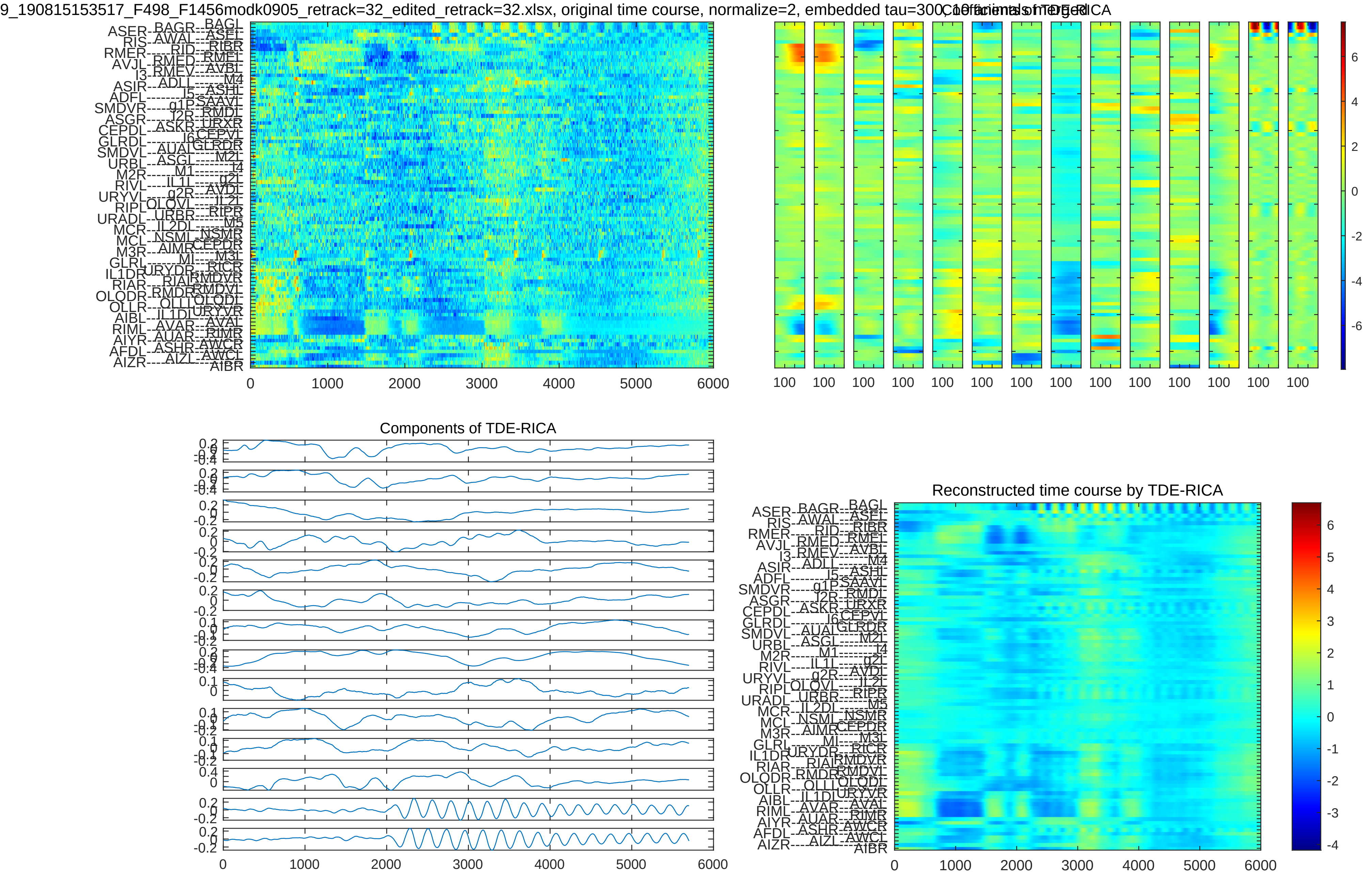

9_190815153517_F498_F1456modk0905_retrack=32_edited_retrack=32.xlsx, original time course, normalize=2, embedded tau=300, 10 trials of merged
Coefficients of TDE-RICA
Components of TDE-RICA
Reconstructed time course by TDE-RICA
0
1000
2000
3000
4000
5000
6000

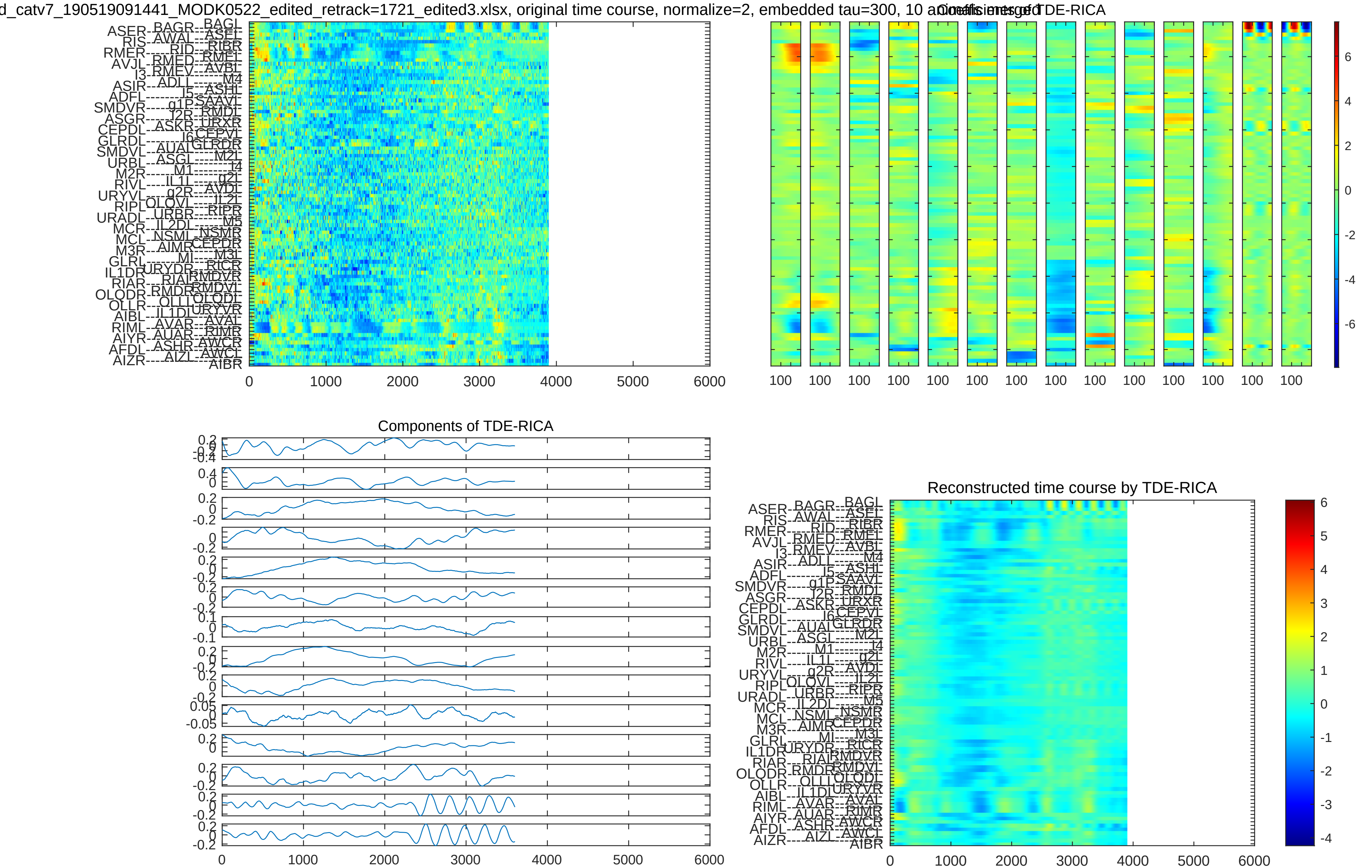
d_catv7_190519091441_MODK0522_edited_retrack=1721_edited3.xlsx, original time course, normalize=2, embedded tau=300, 10 animals merged
Coefficients of TDE-RICA
0 1000 2000 3000 4000 5000 6000
100 100 100 100 100 100 100 100 100 100 100 100 100 100
6 4 2 0 -2 -4 -6
Components of TDE-RICA
0 1000 2000 3000 4000 5000 6000
Reconstructed time course by TDE-RICA
0 1000 2000 3000 4000 5000 6000
6 5 4 3 2 1 0 -1 -2 -3 -4

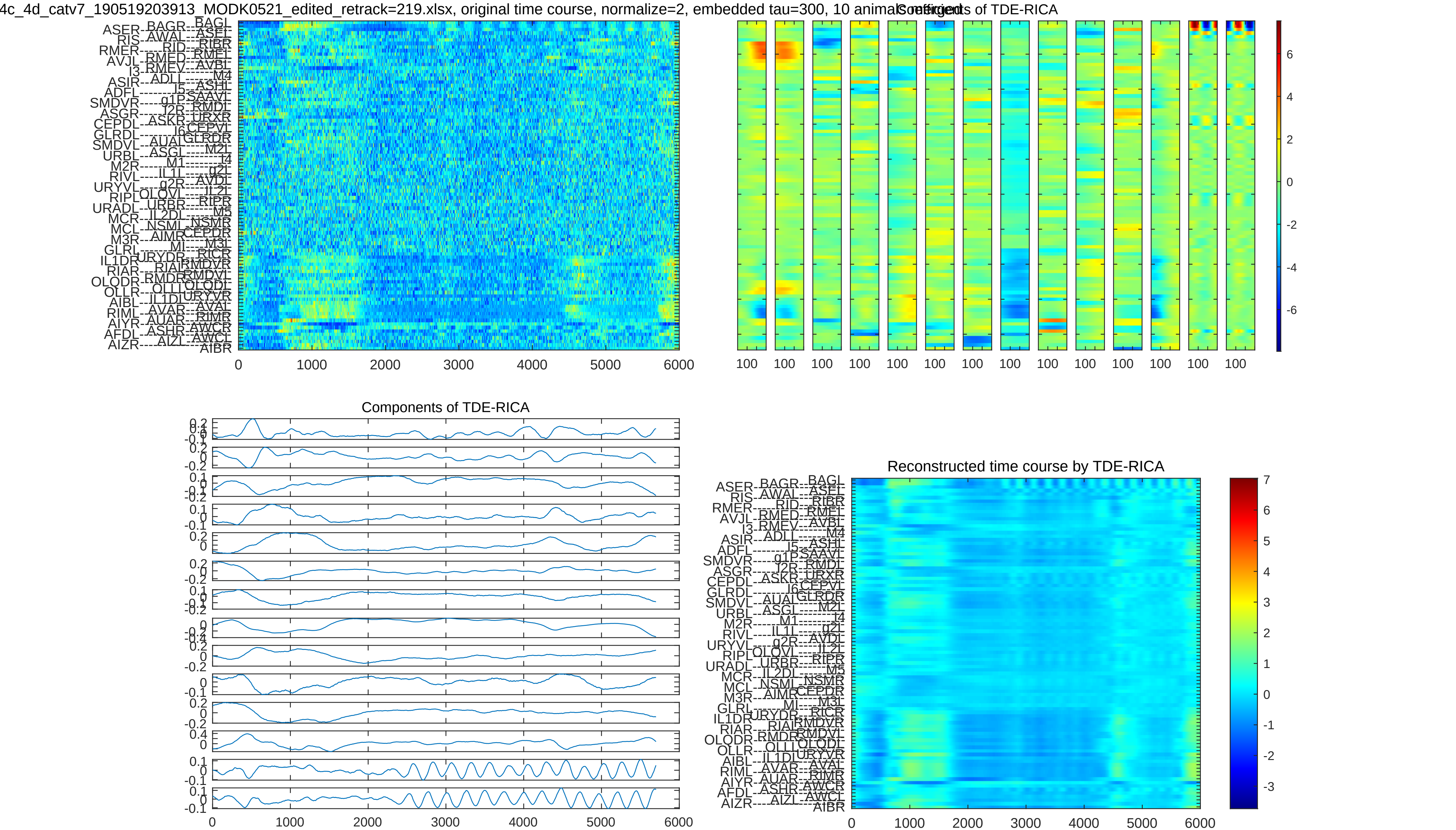
4c_4d_catv7_190519203913_MODK0521_edited_retrack=219.xlsx, original time course, normalize=2, embedded tau=300, 10 animals
Coefficients of TDE-RICA
Components of TDE-RICA
Reconstructed time course by TDE-RICA
0 1000 2000 3000 4000 5000 6000
100
6 4 2 0 -2 -4 -6
7 6 5 4 3 2 1 0 -1 -2 -3

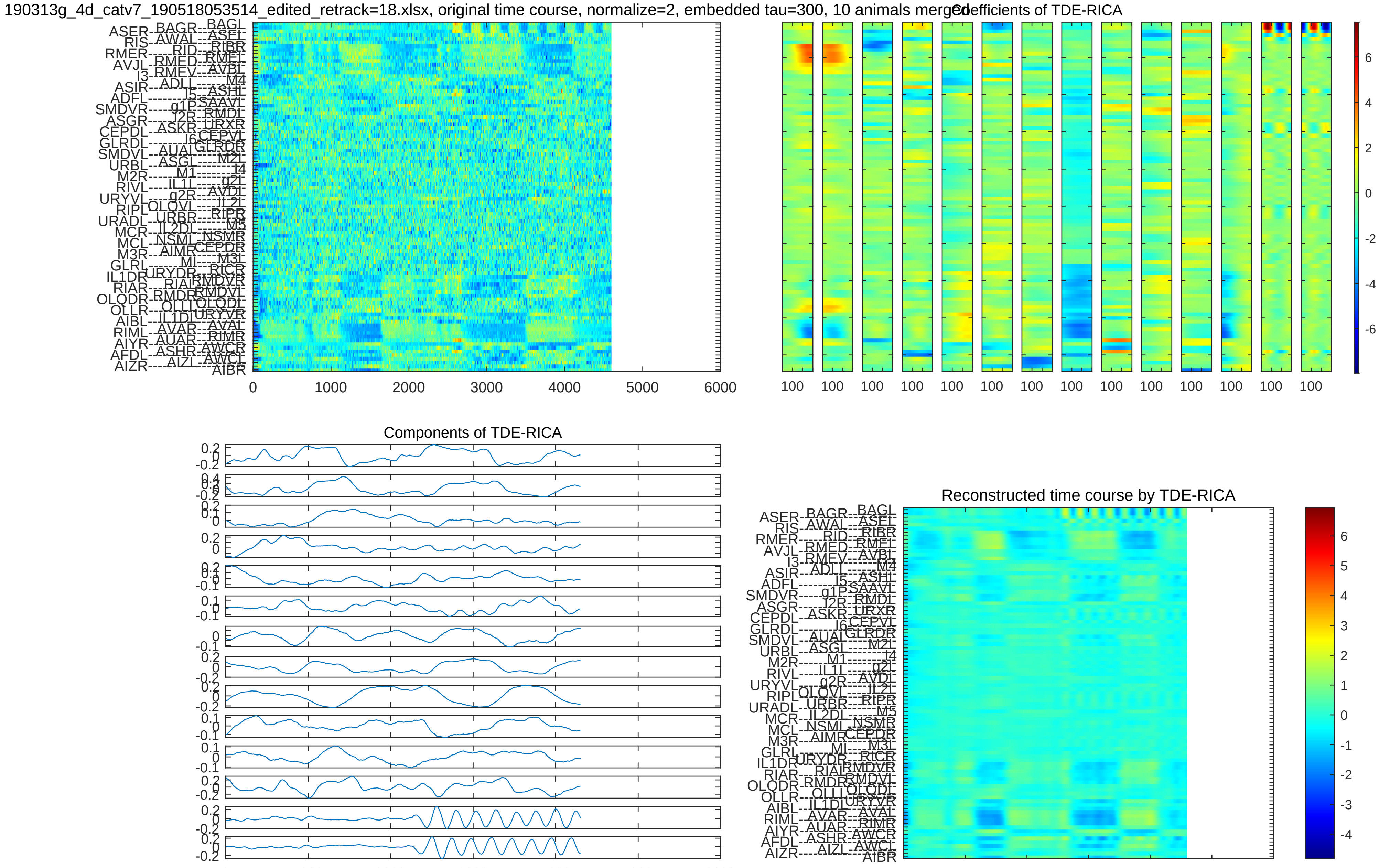
190313g_4d_catv7_190518053514_edited_retrack=18.xlsx, original time course, normalize=2, embedded tau=300, 10 animals merged
Coefficients of TDE-RICA
Components of TDE-RICA
Reconstructed time course by TDE-RICA

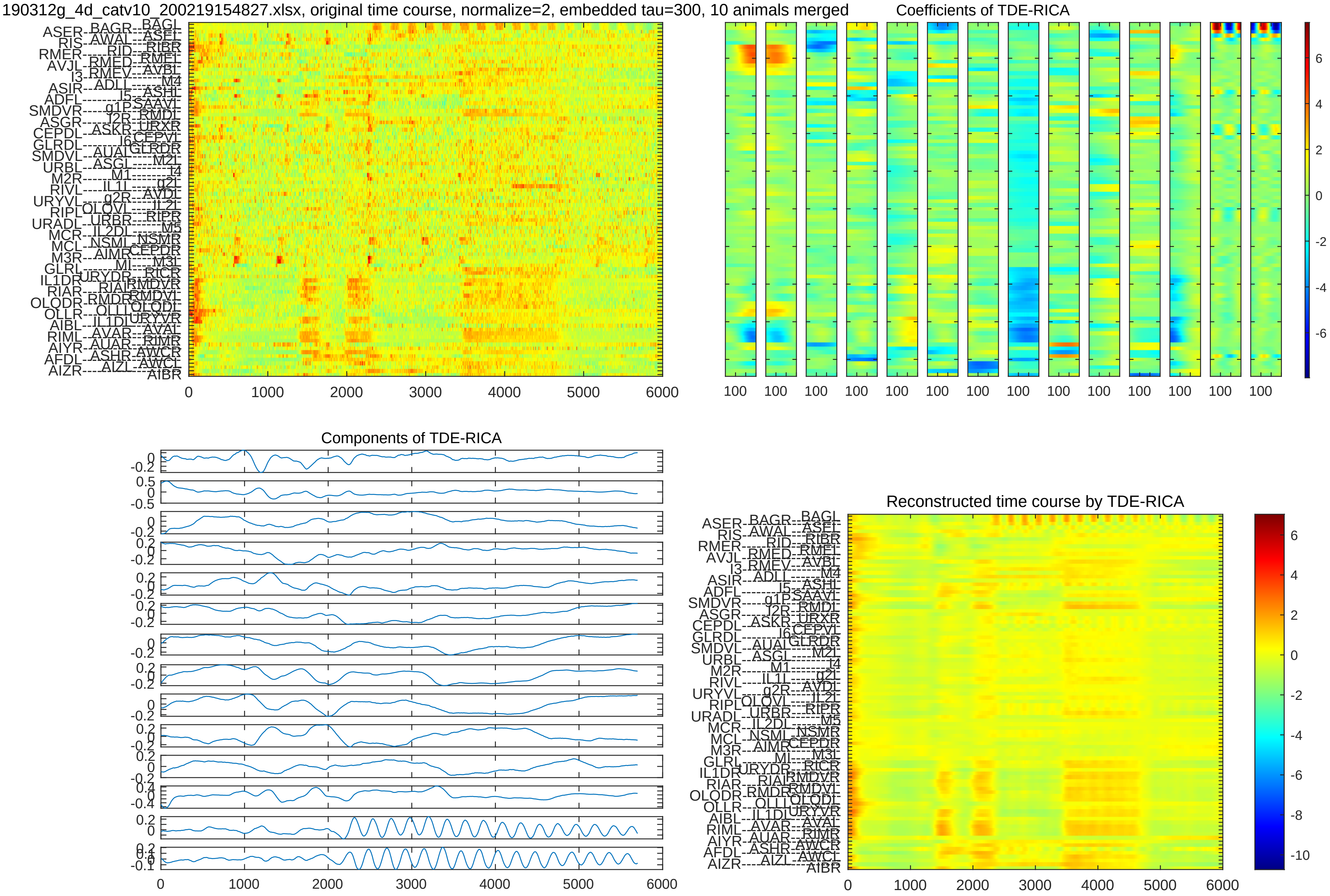
190312g_4d_catv10_200219154827.xlsx, original time course, normalize=2, embedded tau=300, 10 animals merged
Coefficients of TDE-RICA
Components of TDE-RICA
Reconstructed time course by TDE-RICA

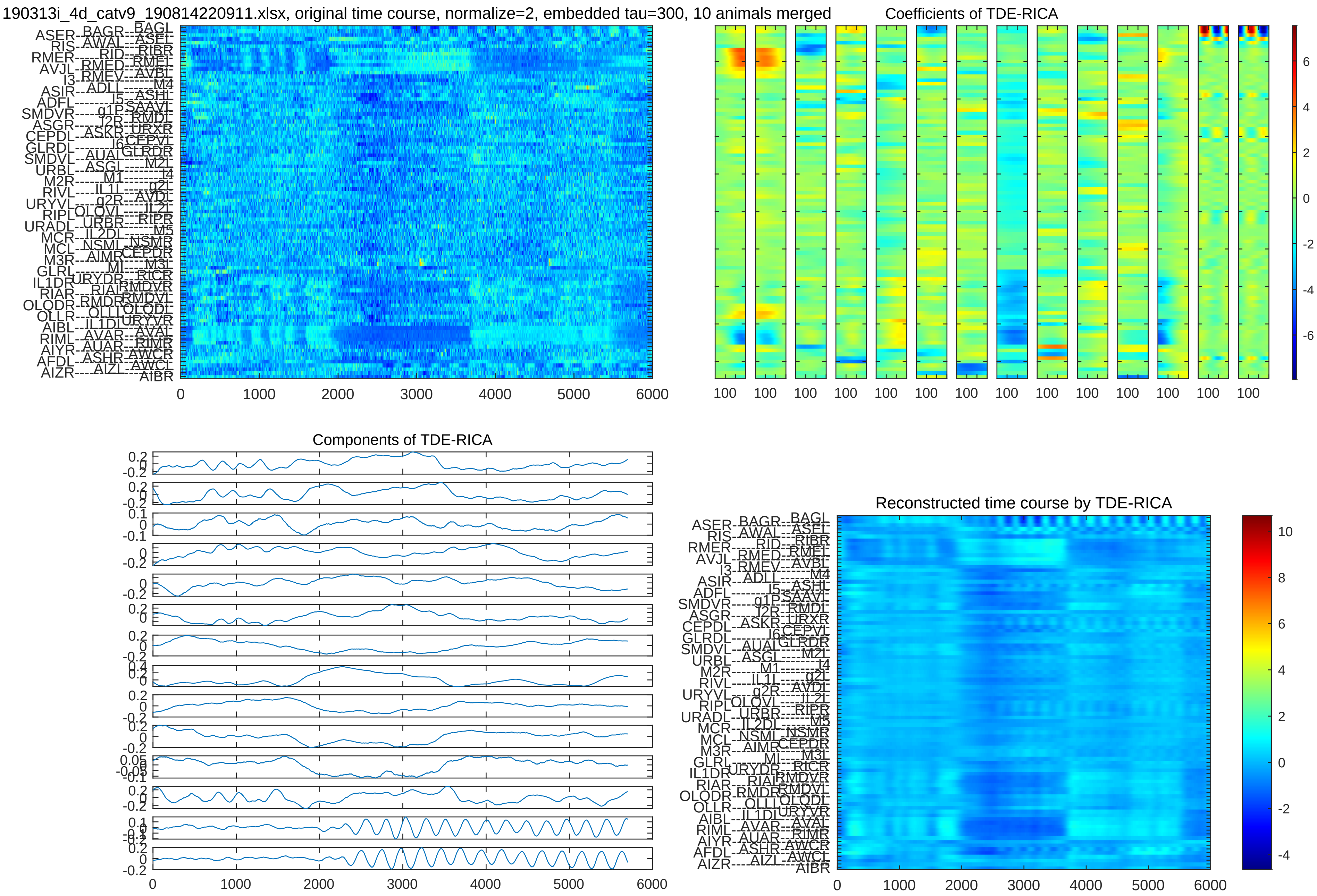
190313i_4d_catv9_190814220911.xlsx, original time course, normalize=2, embedded tau=300, 10 animals merged
Coefficients of TDE-RICA
Components of TDE-RICA
Reconstructed time course by TDE-RICA

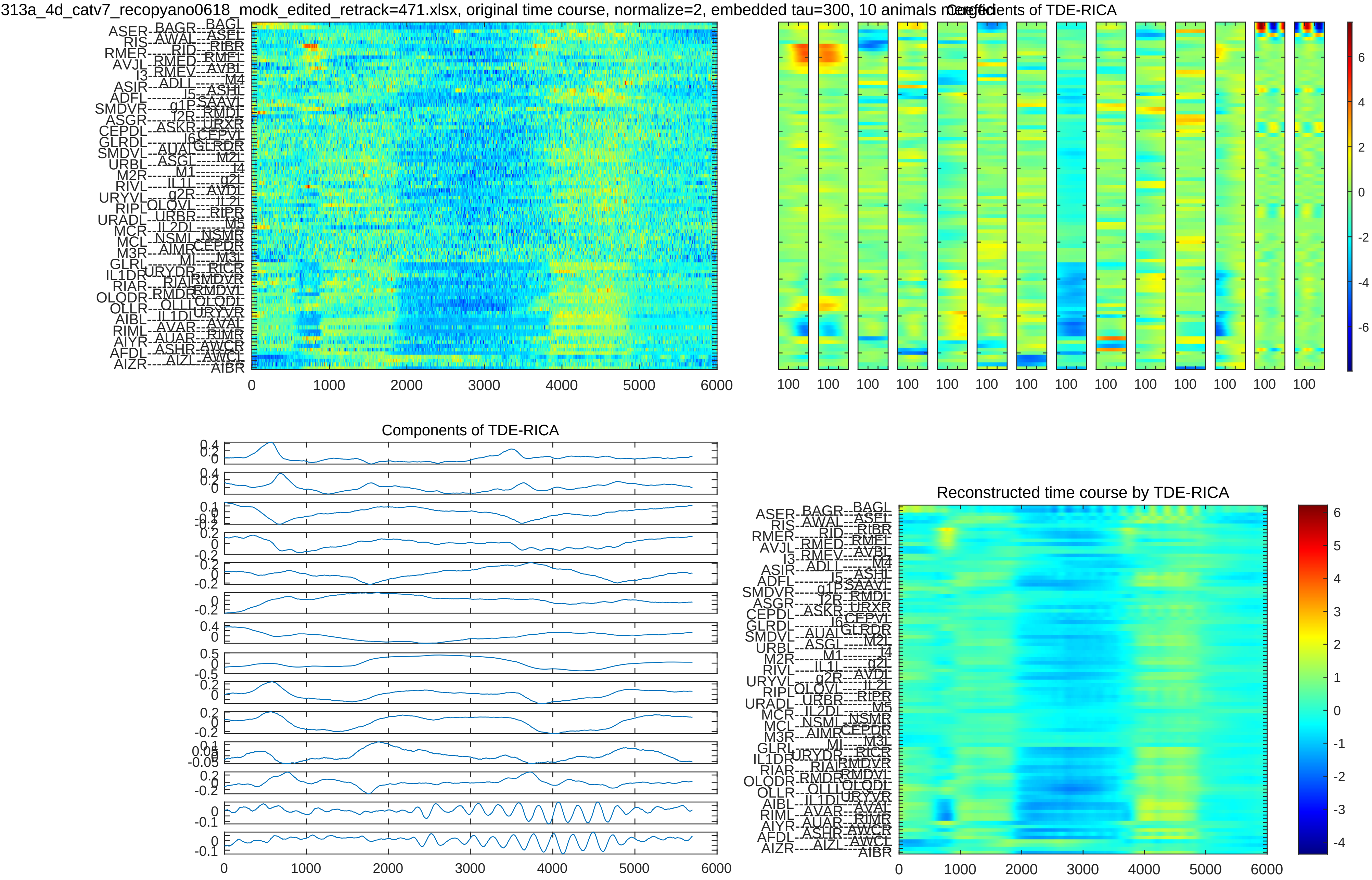
313a_4d_catv7_recopyano0618_modk_edited_retrack=471.xlsx, original time course, normalize=2, embedded tau=300, 10 animals merged
Coefficients of TDE-RICA
Components of TDE-RICA
Reconstructed time course by TDE-RICA
0
1000
2000
3000
4000
5000
6000
100

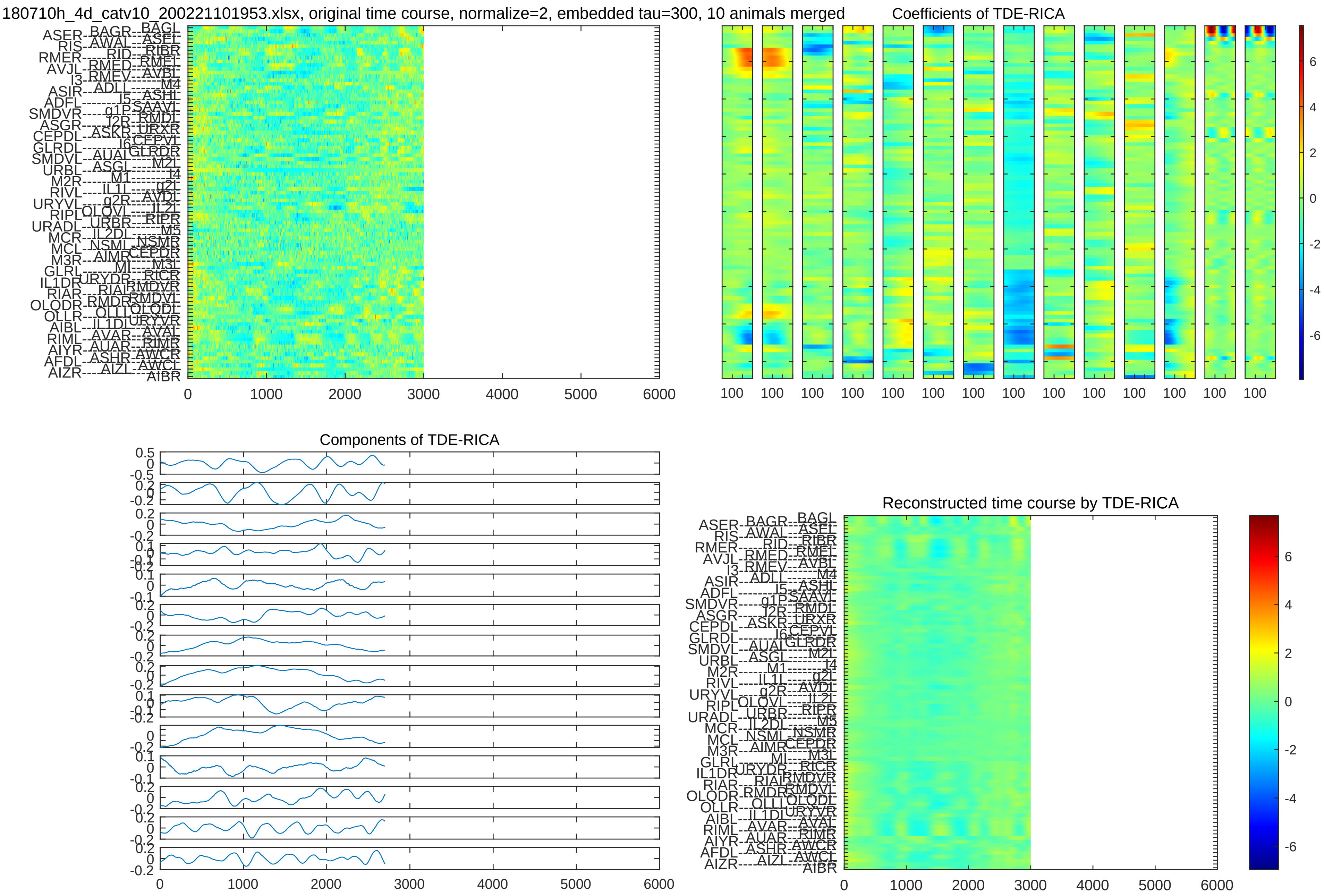

180710h_4d_catv10_200221101953.xlsx, original time course, normalize=2, embedded tau=300, 10 animals merged
Coefficients of TDE-RICA
Components of TDE-RICA
Reconstructed time course by TDE-RICA
0
1000
2000
3000
4000
5000
6000
100
6
4
2
0
-2
-4
-6

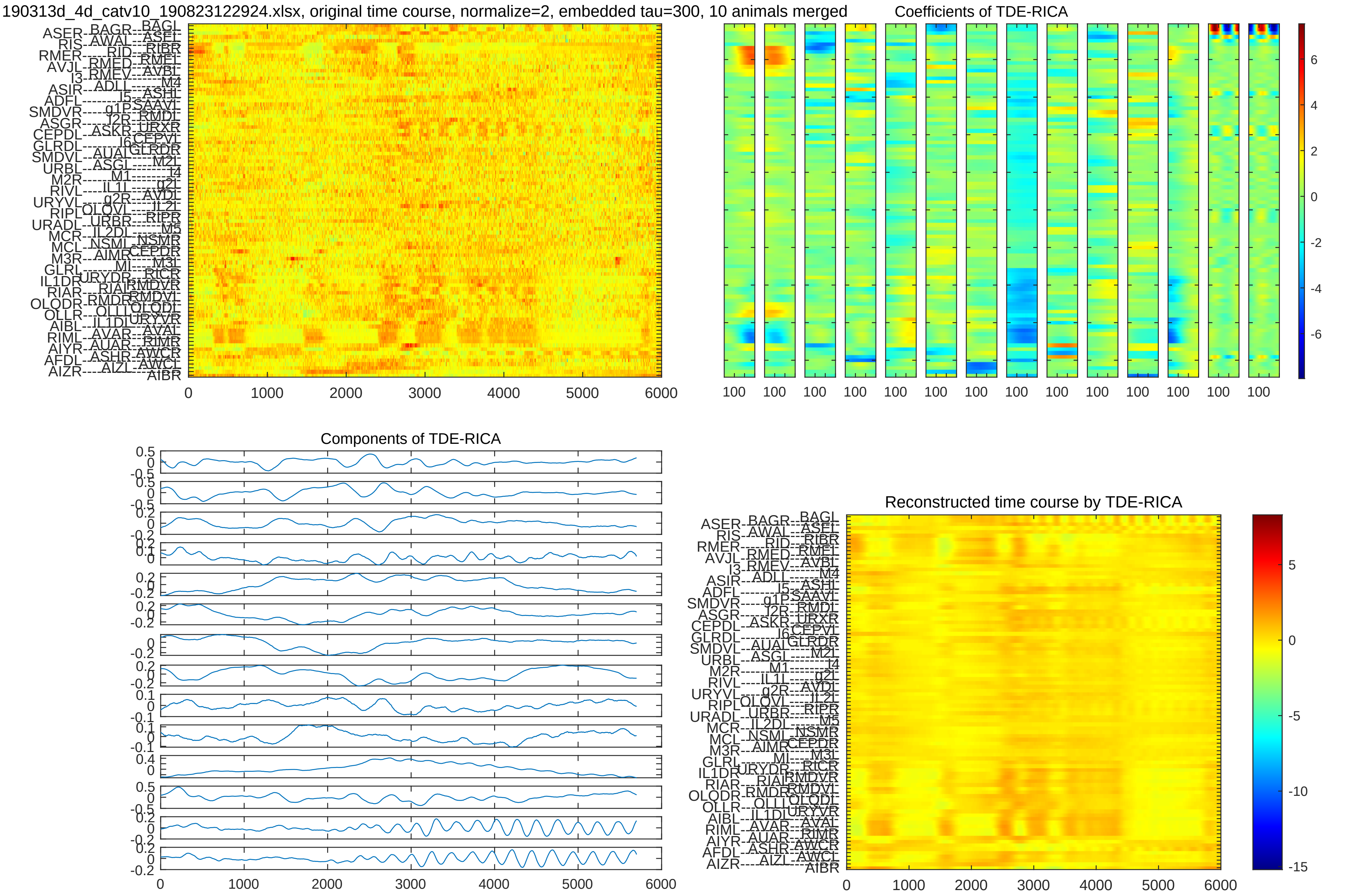

190313d_4d_catv10_190823122924.xlsx, original time course, normalize=2, embedded tau=300, 10 animals merged
Coefficients of TDE-RICA
Components of TDE-RICA
Reconstructed time course by TDE-RICA
0 1000 2000 3000 4000 5000 6000

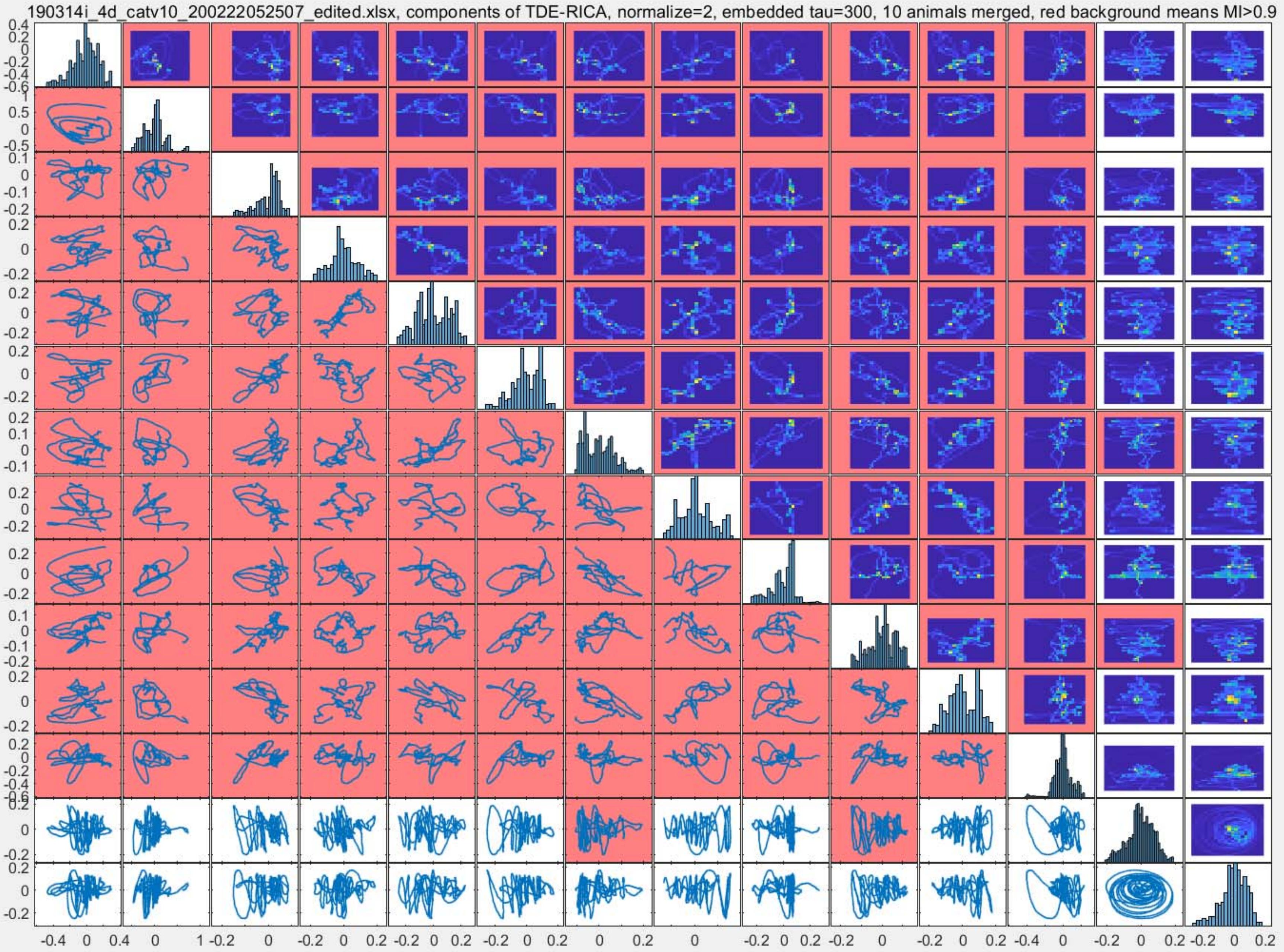
190314i_4d_catv10_200222052507_edited.xlsx, components of TDE-RICA, normalize=2, embedded tau=300, 10 animals merged, red background means MI>0.9

190312f_4d_catv9_190815153517_F498_F1456modk0905_retrack=32_edited_retrack=32.xlsx, components of TDE-RICA, normalize=2, embedded tau=300, 10 animals merged, red background means MI

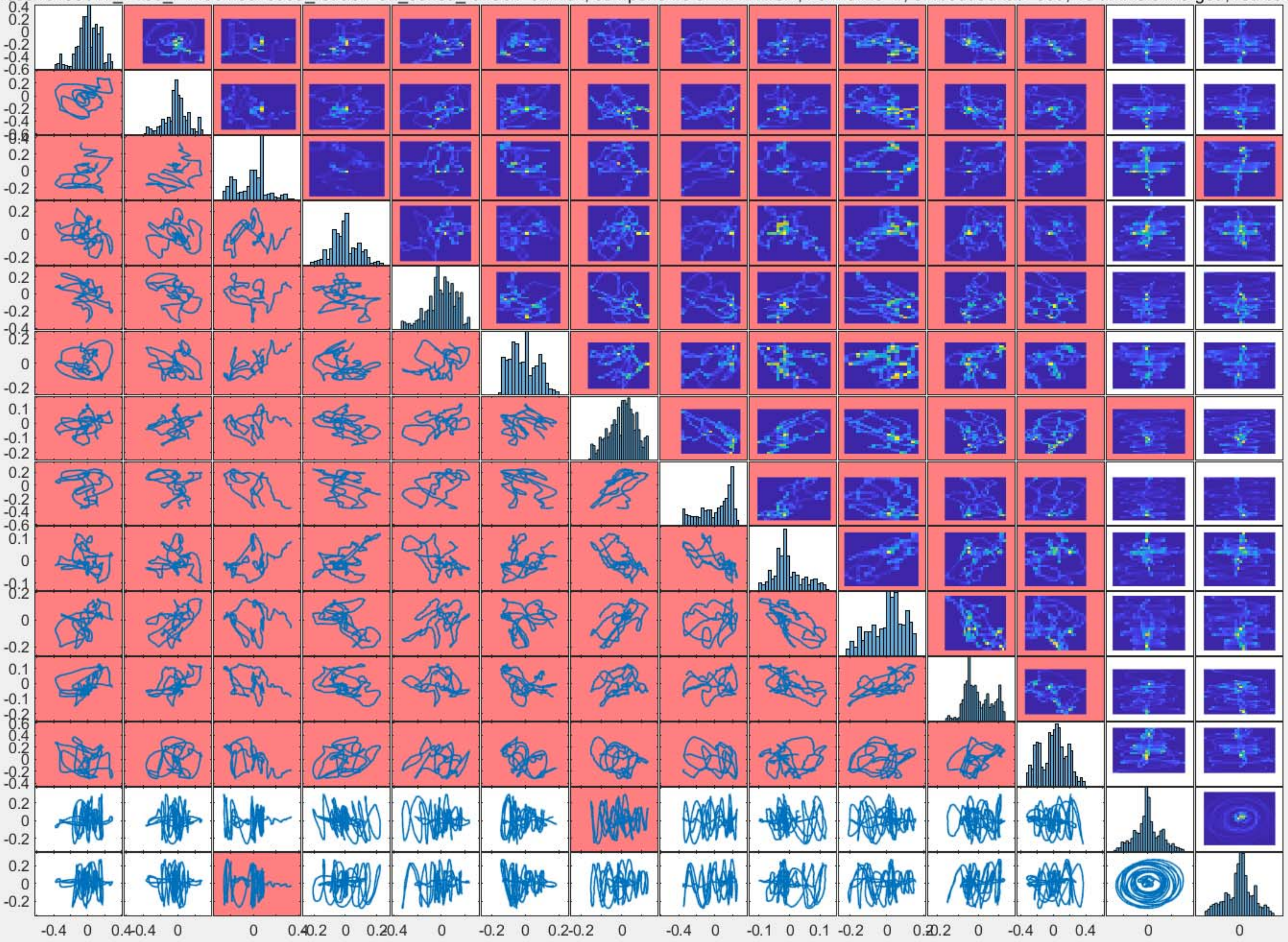

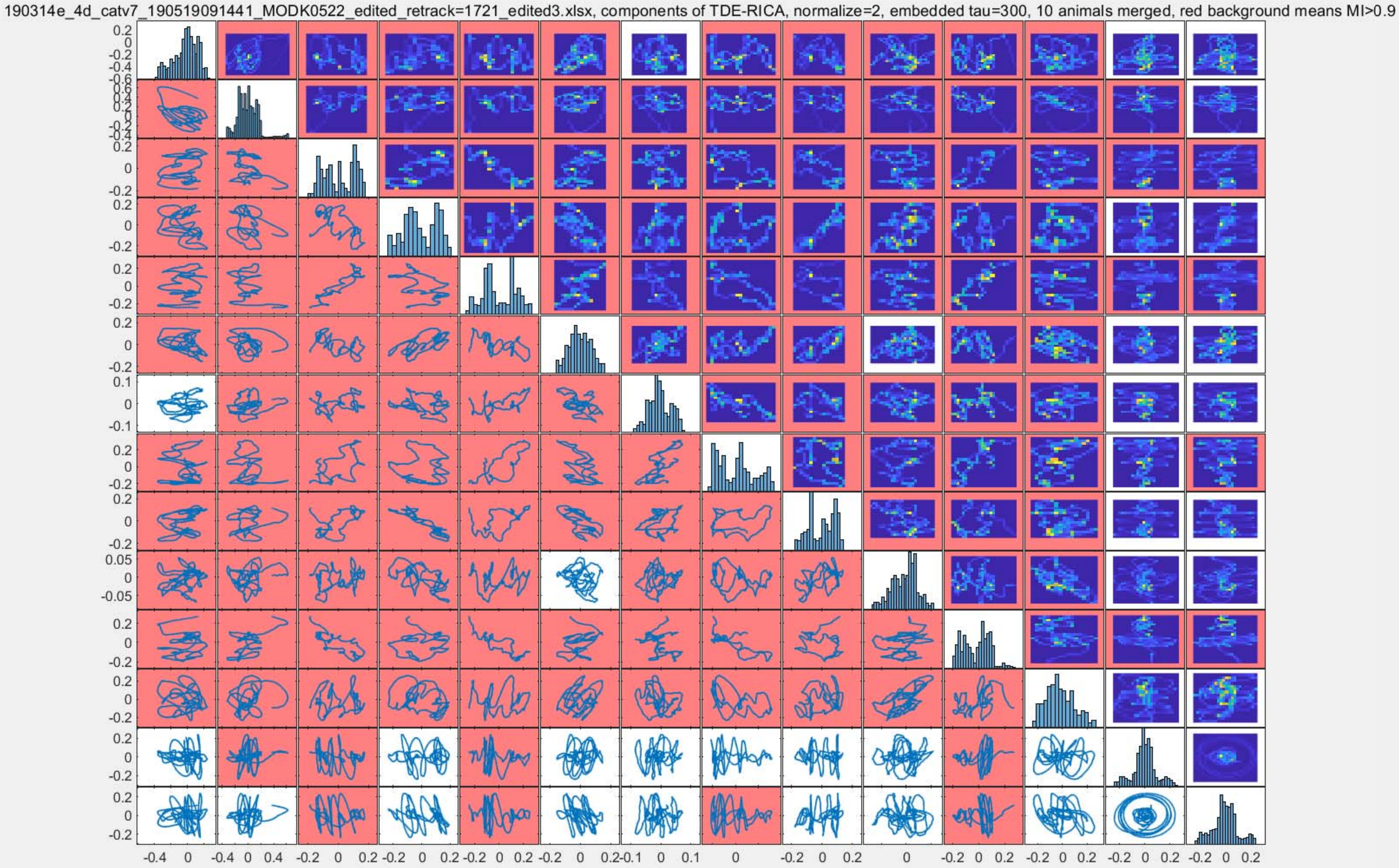
190314e_4d_catv7_190519091441_MODK0522_edited_retrack=1721_edited3.xlsx, components of TDE-RICA, normalize=2, embedded tau=300, 10 animals merged, red background means MI>0.9

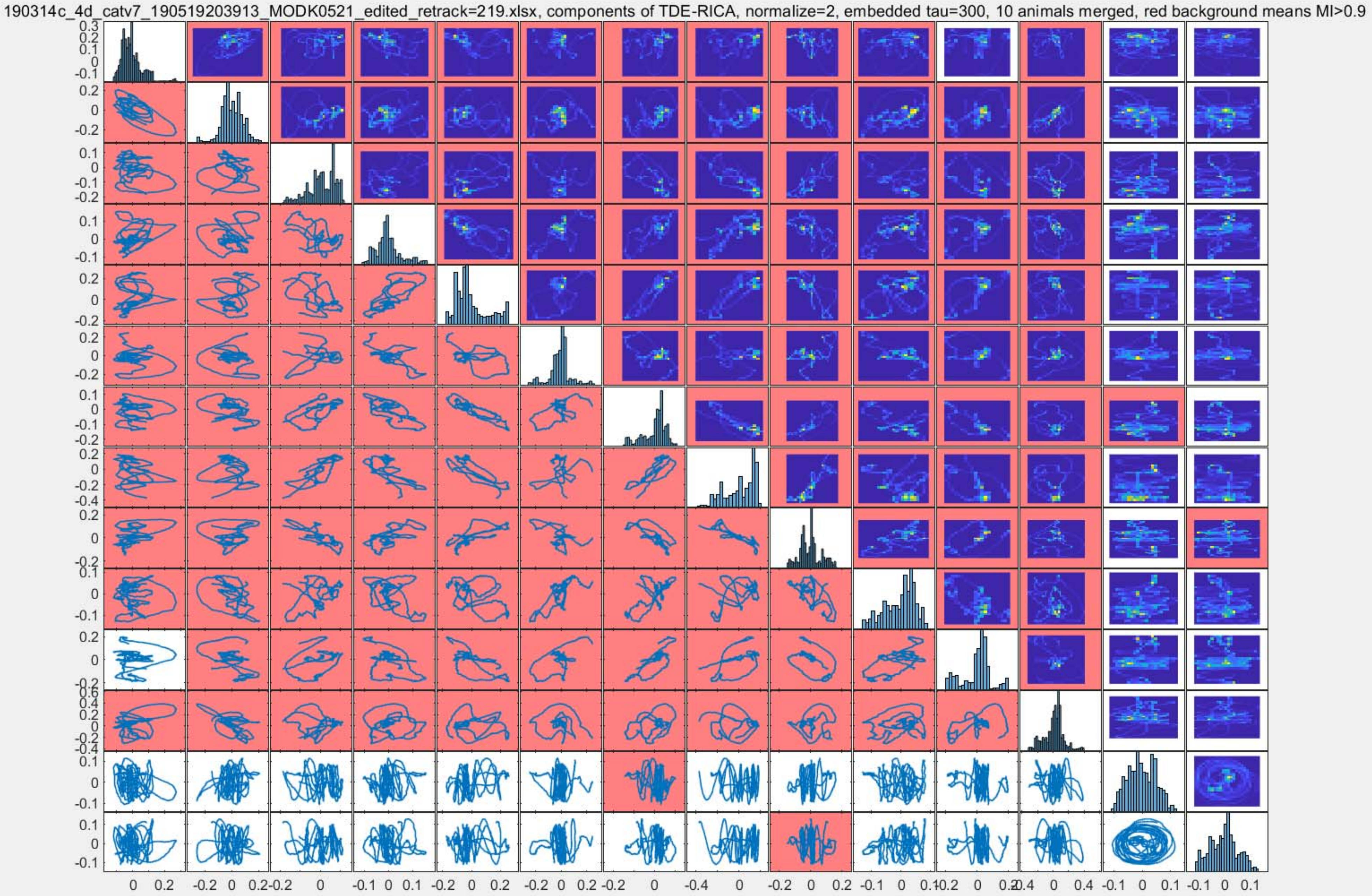
190314c_4d_catv7_190519203913_MODK0521_edited_retrack=219.xlsx, components of TDE-RICA, normalize=2, embedded tau=300, 10 animals merged, red background means MI>0.9

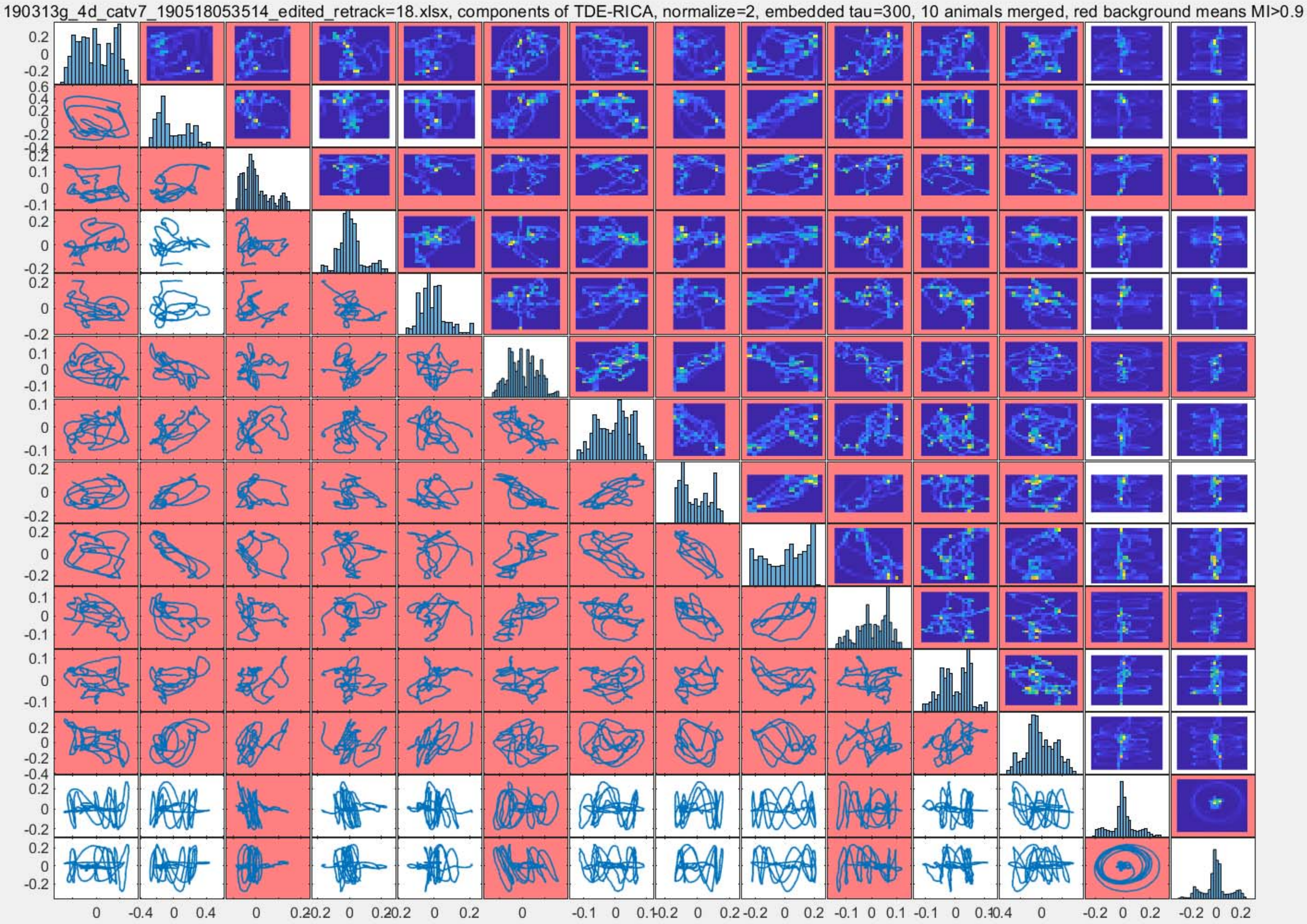
190313g_4d_catv7_190518053514_edited_retrack=18.xlsx, components of TDE-RICA, normalize=2, embedded tau=300, 10 animals merged, red background means MI>0.9

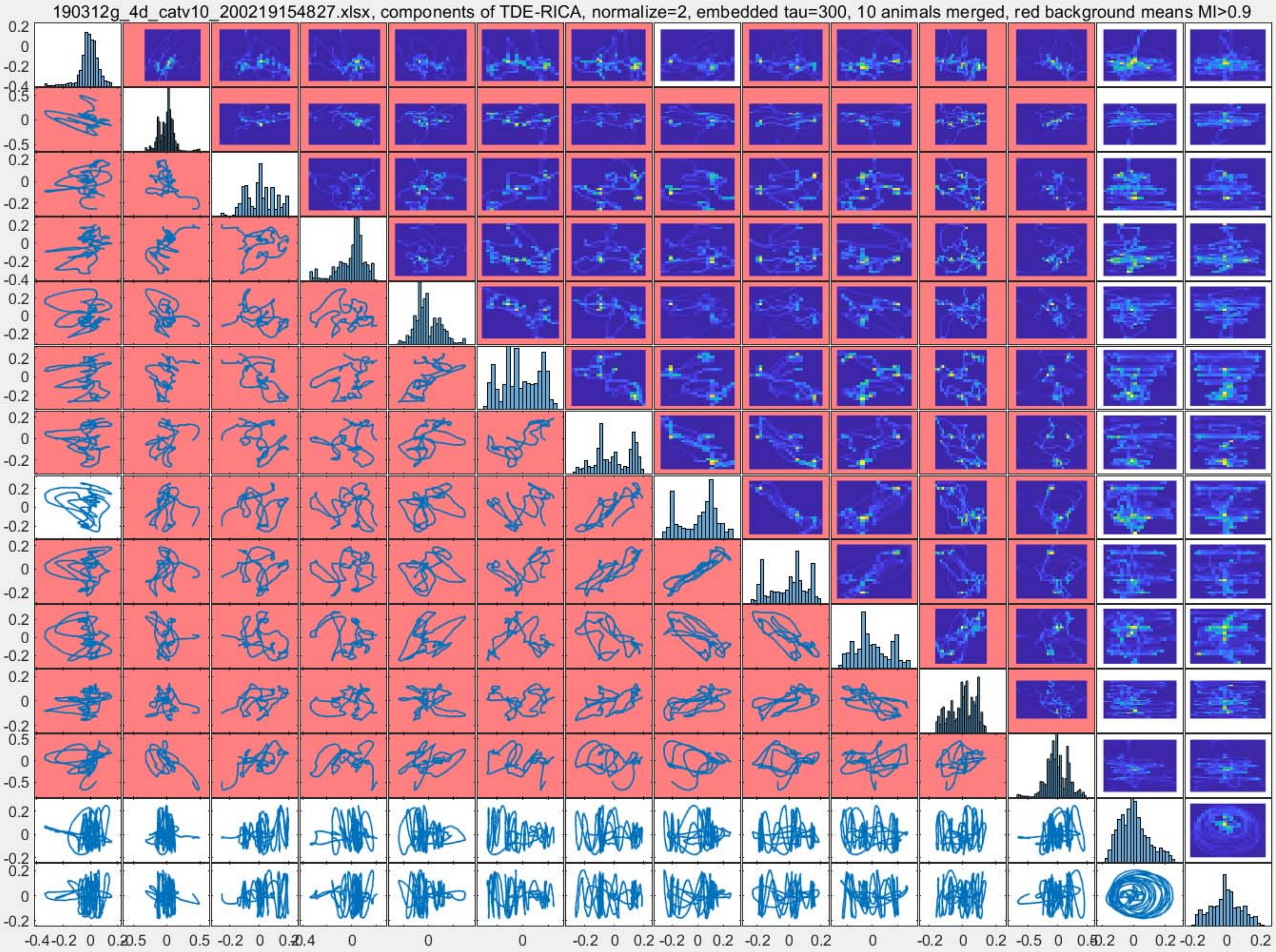
190312g_4d_catv10_200219154827.xlsx, components of TDE-RICA, normalize=2, embedded tau=300, 10 animals merged, red background means MI>0.9

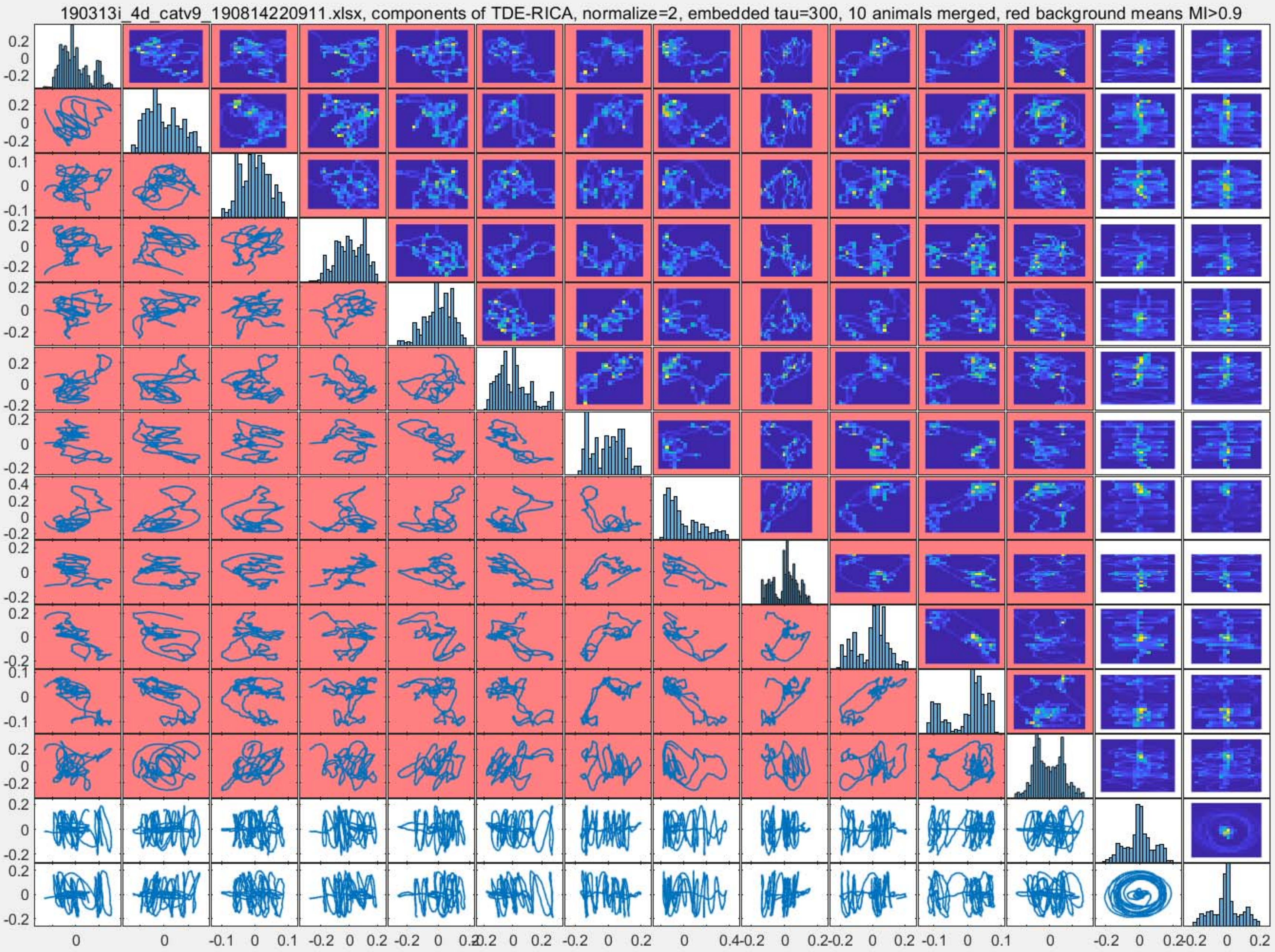
190313i_4d_catv9_190814220911.xlsx, components of TDE-RICA, normalize=2, embedded tau=300, 10 animals merged, red background means MI>0.9

190313a_4d_catv7_recopyano0618_modk_edited_retrack=471.xlsx, components of TDE-RICA, normalize=2, embedded tau=300, 10 animals merged, red background means MI>0.9

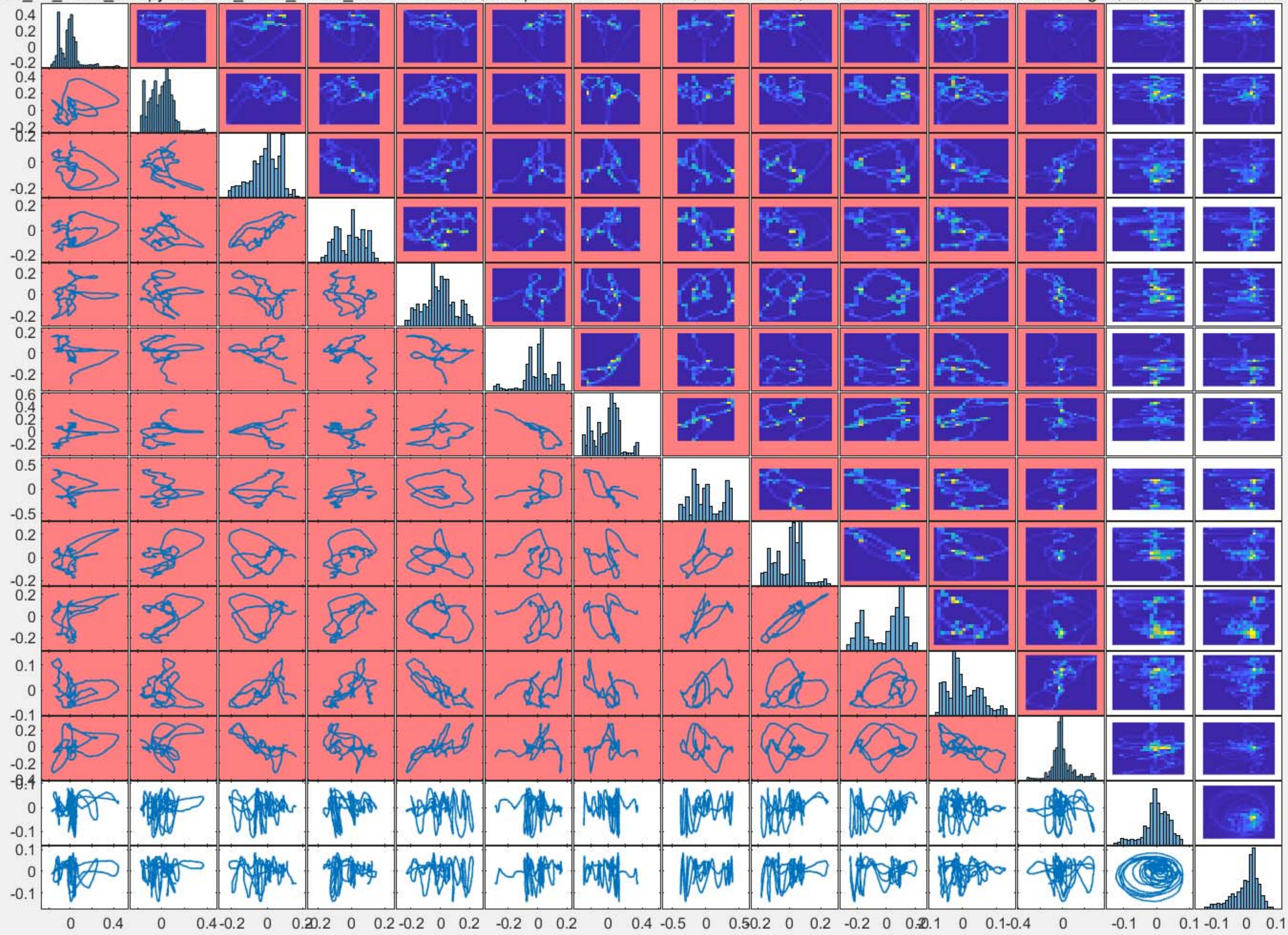

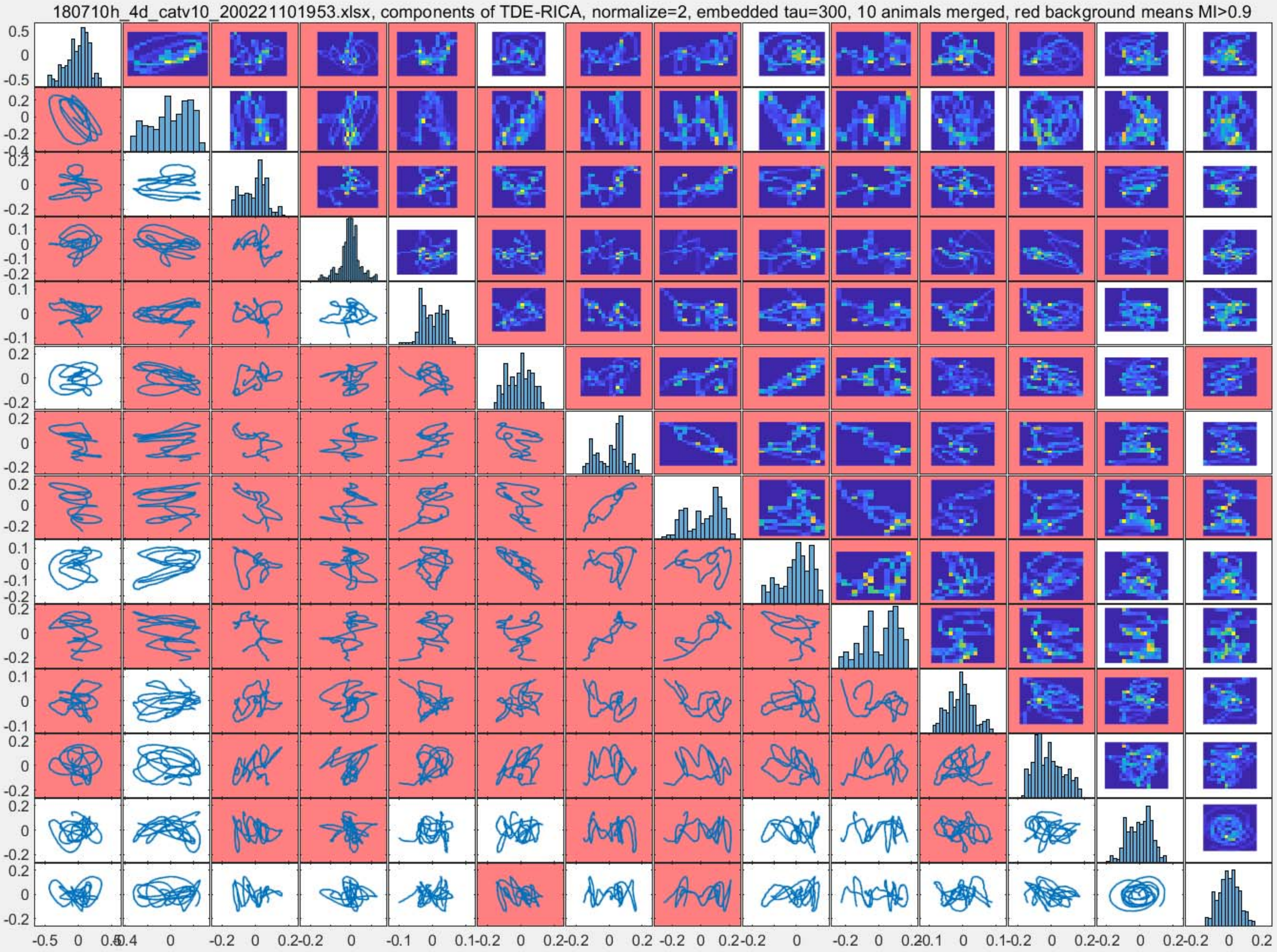
180710h_4d_catv10_200221101953.xlsx, components of TDE-RICA, normalize=2, embedded tau=300, 10 animals merged, red background means MI>0.9

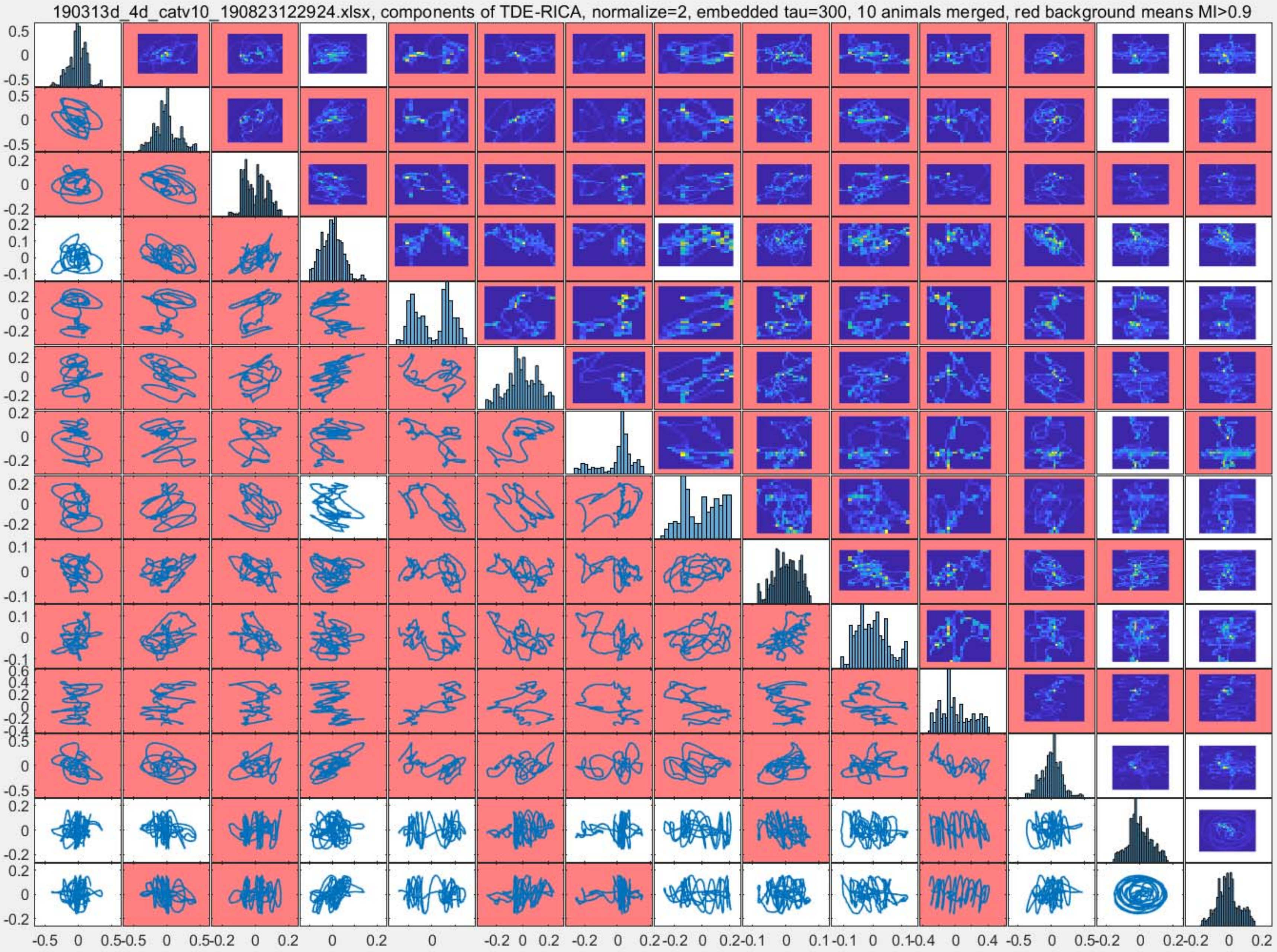
190313d_4d_catv10_190823122924.xlsx, components of TDE-RICA, normalize=2, embedded tau=300, 10 animals merged, red background means MI>0.9

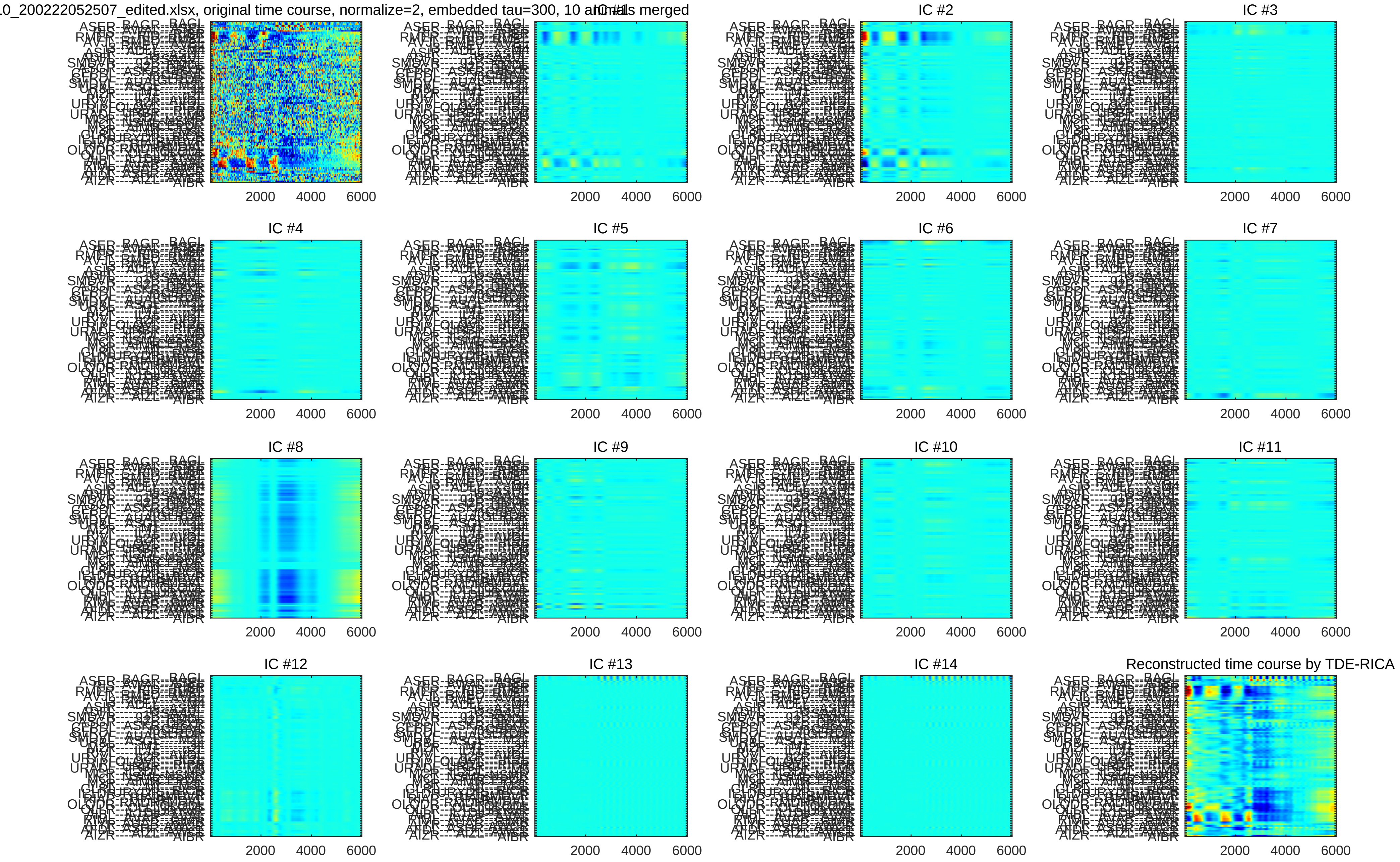
0_200222052507_edited.xlsx, original time course, normalize=2, embedded tau=300, 10 animals merged
IC #1
IC #2
IC #3
IC #4
IC #5
IC #6
IC #7
IC #8
IC #9
IC #10
IC #11
IC #12
IC #13
IC #14
Reconstructed time course by TDE-RICA
2000
4000
6000

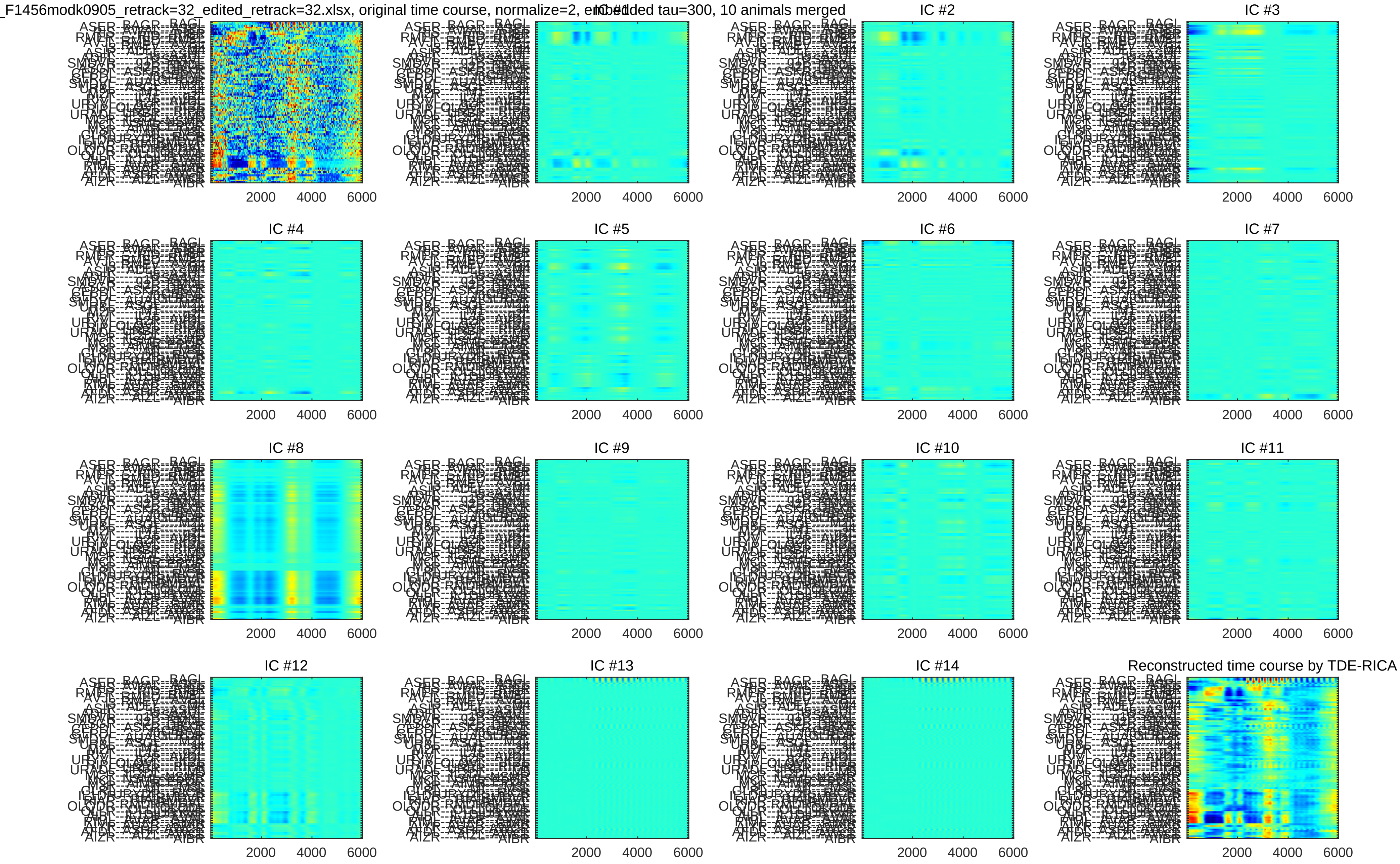
_F1456modk0905_retrack=32_edited_retrack=32.xlsx, original time course, normalize=2, embedded tau=300, 10 animals merged
IC #1
IC #2
IC #3
IC #4
IC #5
IC #6
IC #7
IC #8
IC #9
IC #10
IC #11
IC #12
IC #13
IC #14
Reconstructed time course by TDE-RICA
2000
4000
6000
ASER
BAGR
BAGL
AIZR
AIZL
AIBR

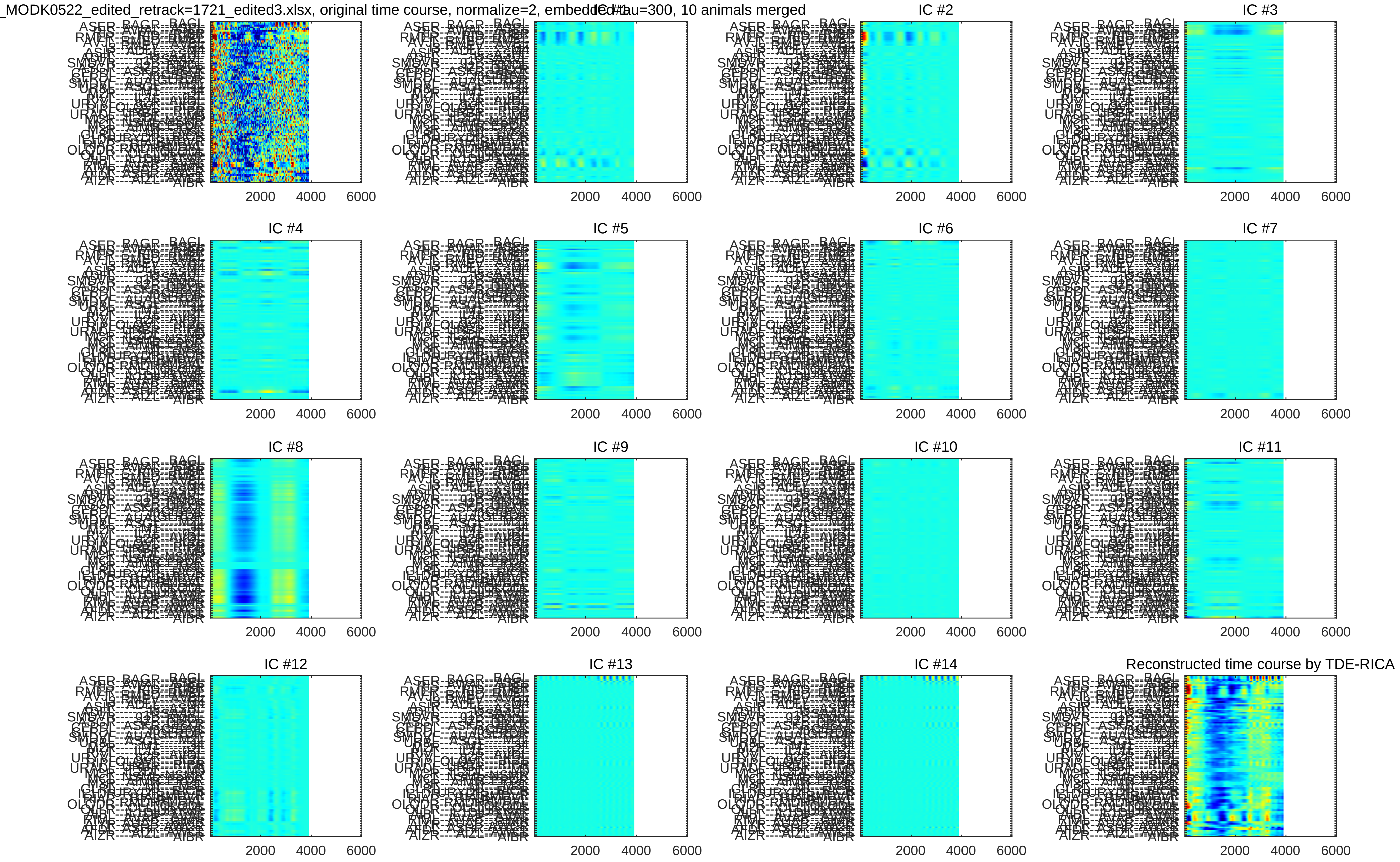
_MODK0522_edited_retrack=1721_edited3.xlsx, original time course, normalize=2, embedded tau=300, 10 animals merged
IC #1
IC #2
IC #3
IC #4
IC #5
IC #6
IC #7
IC #8
IC #9
IC #10
IC #11
IC #12
IC #13
IC #14
Reconstructed time course by TDE-RICA
2000
4000
6000

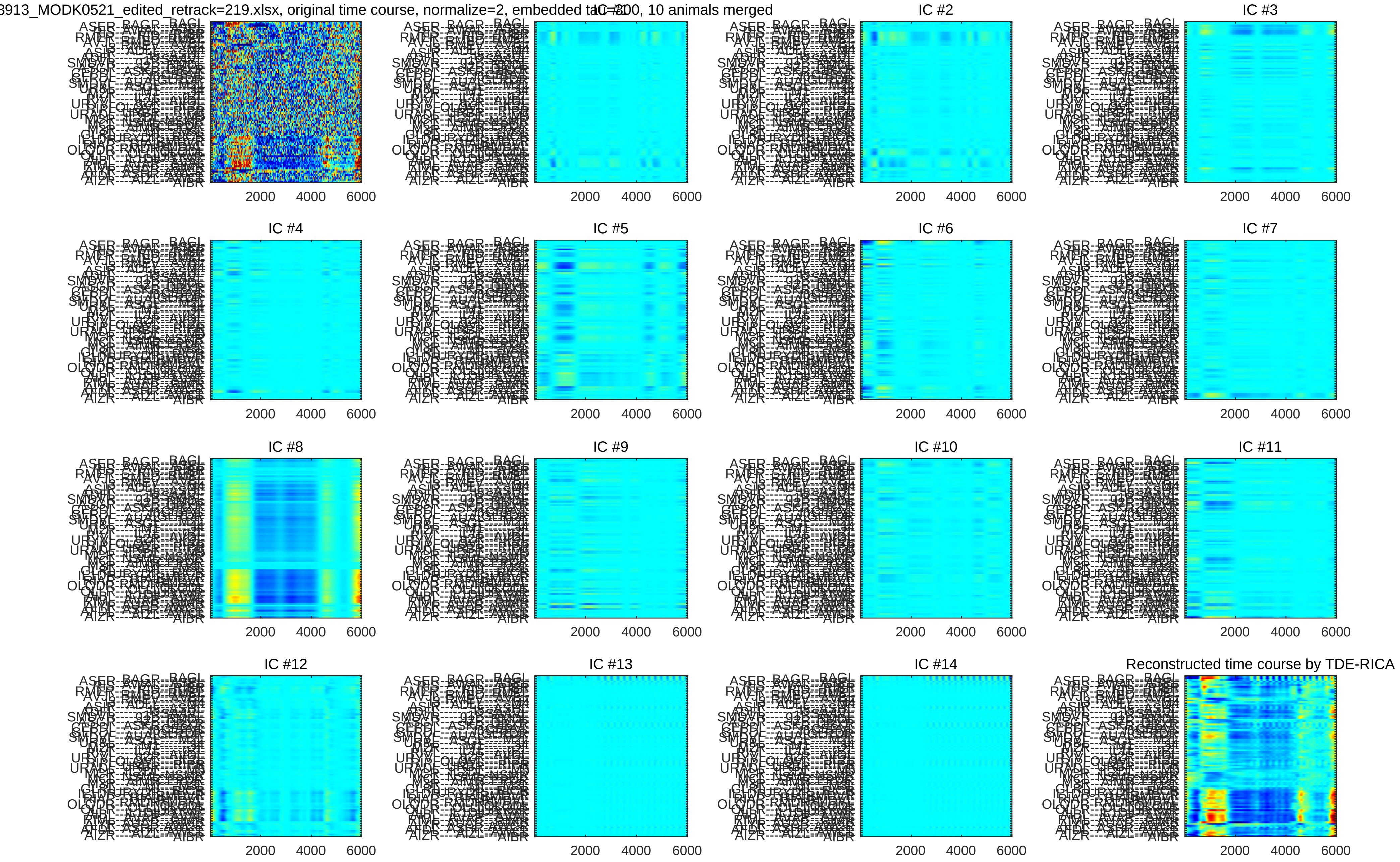
3913_MODK0521_edited_retrack=219.xlsx, original time course, normalize=2, embedded tau=100, 10 animals merged
IC #1
IC #2
IC #3
IC #4
IC #5
IC #6
IC #7
IC #8
IC #9
IC #10
IC #11
IC #12
IC #13
IC #14
Reconstructed time course by TDE-RICA
2000
4000
6000

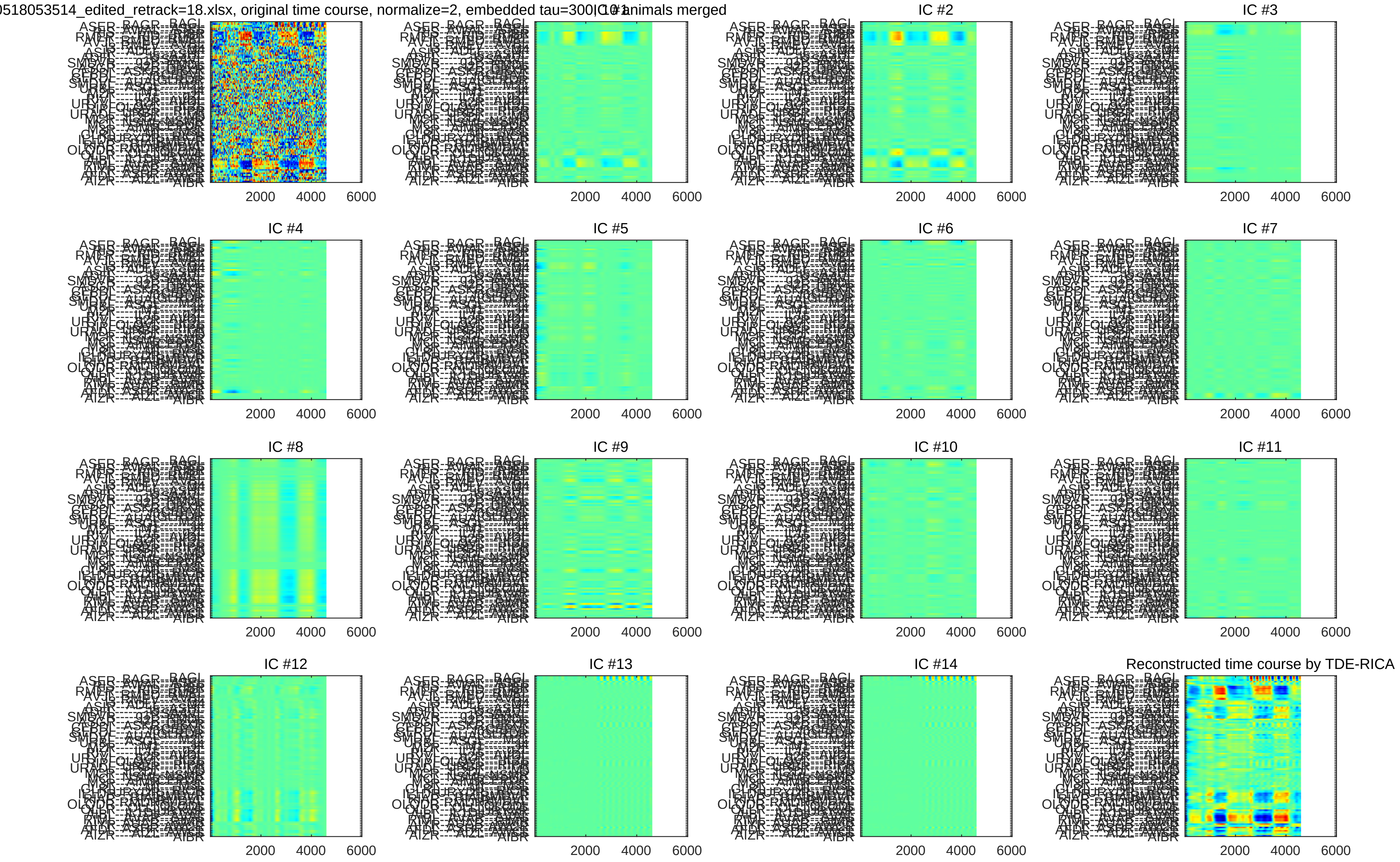

0518053514_edited_retrack=18.xlsx, original time course, normalize=2, embedded tau=300
animals merged
IC #2
IC #3
IC #4
IC #5
IC #6
IC #7
IC #8
IC #9
IC #10
IC #11
IC #12
IC #13
IC #14
Reconstructed time course by TDE-RICA
2000
4000
6000

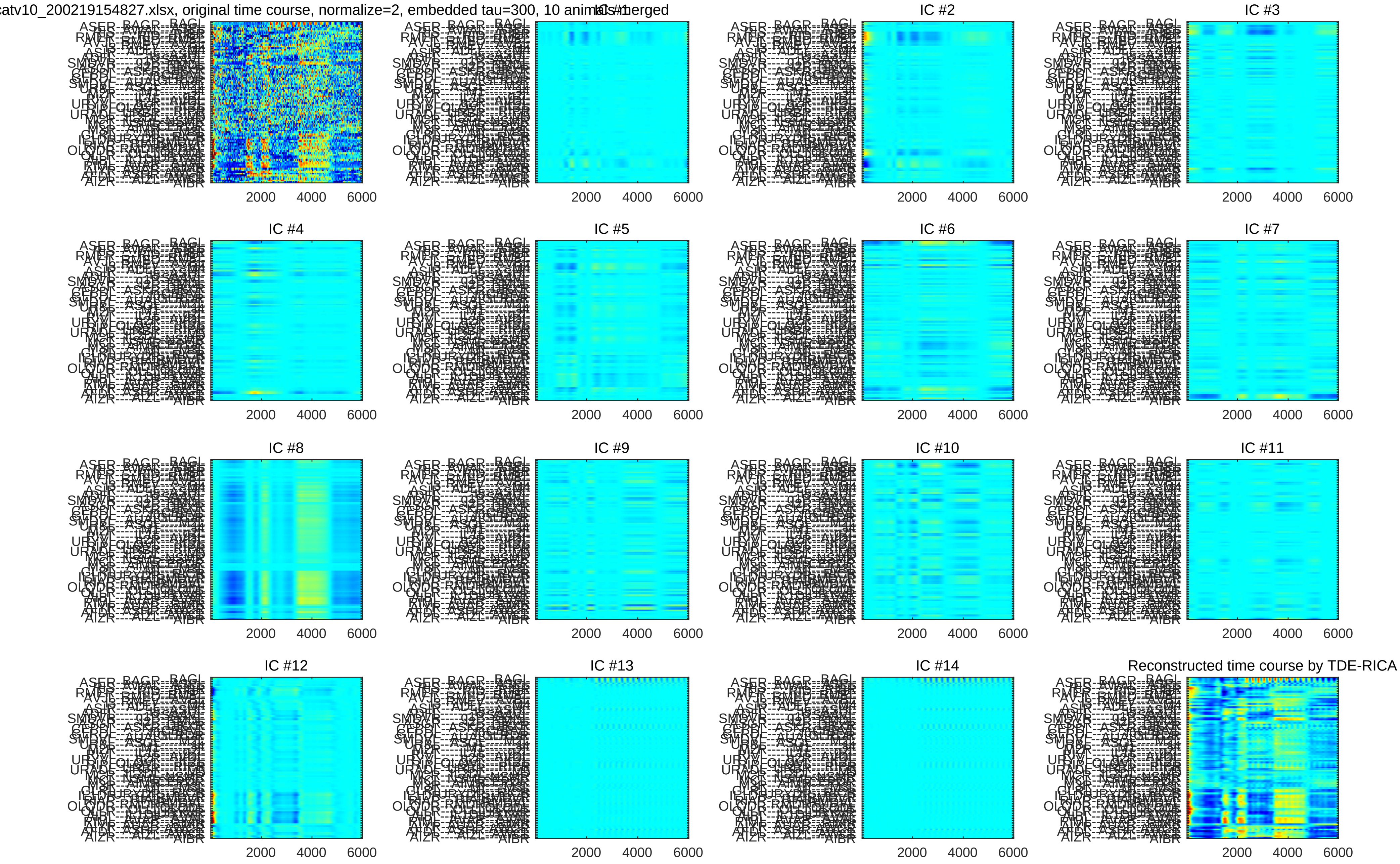
catv10_200219154827.xlsx, original time course, normalize=2, embedded tau=300, 10 animals merged
IC #1
IC #2
IC #3
IC #4
IC #5
IC #6
IC #7
IC #8
IC #9
IC #10
IC #11
IC #12
IC #13
IC #14
Reconstructed time course by TDE-RICA
2000
4000
6000

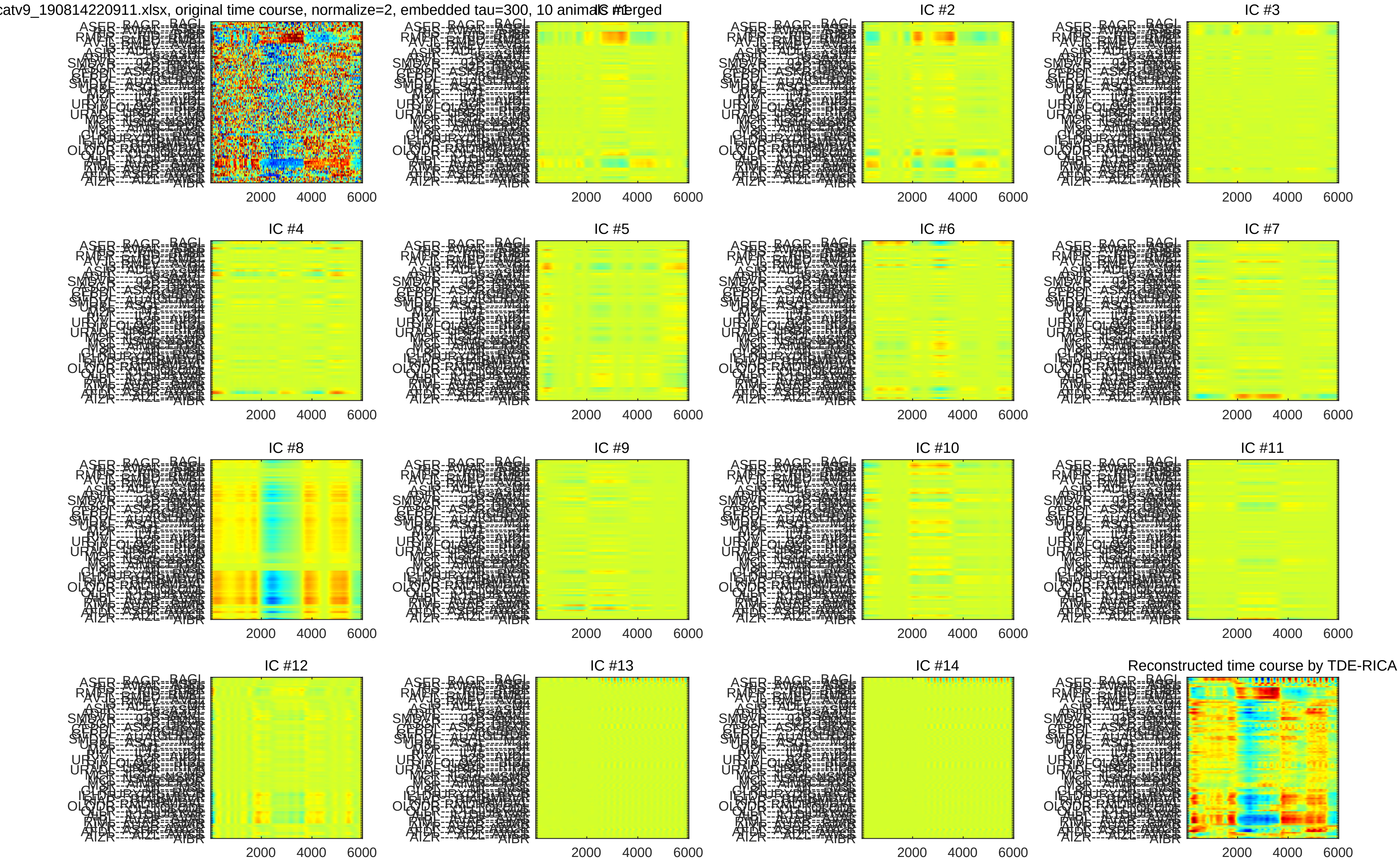
catv9_190814220911.xlsx, original time course, normalize=2, embedded tau=300, 10 animals merged
IC #1
IC #2
IC #3
IC #4
IC #5
IC #6
IC #7
IC #8
IC #9
IC #10
IC #11
IC #12
IC #13
IC #14
Reconstructed time course by TDE-RICA
2000
4000
6000

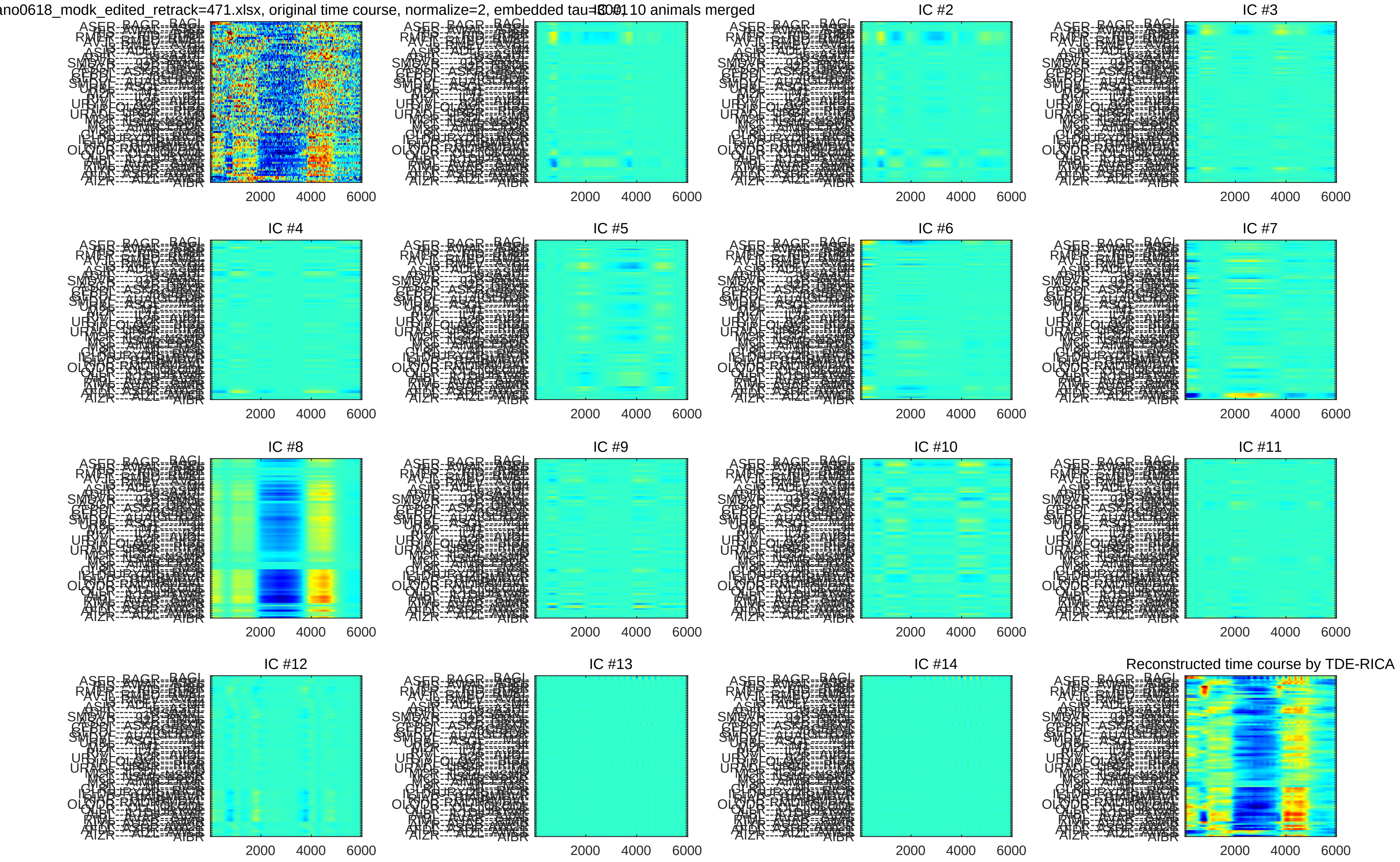
ano0618_modk_edited_retrack=471.xlsx, original time course, normalize=2, embedded tau=10, 10 animals merged
IC #1
IC #2
IC #3
IC #4
IC #5
IC #6
IC #7
IC #8
IC #9
IC #10
IC #11
IC #12
IC #13
IC #14
Reconstructed time course by TDE-RICA
2000
4000
6000

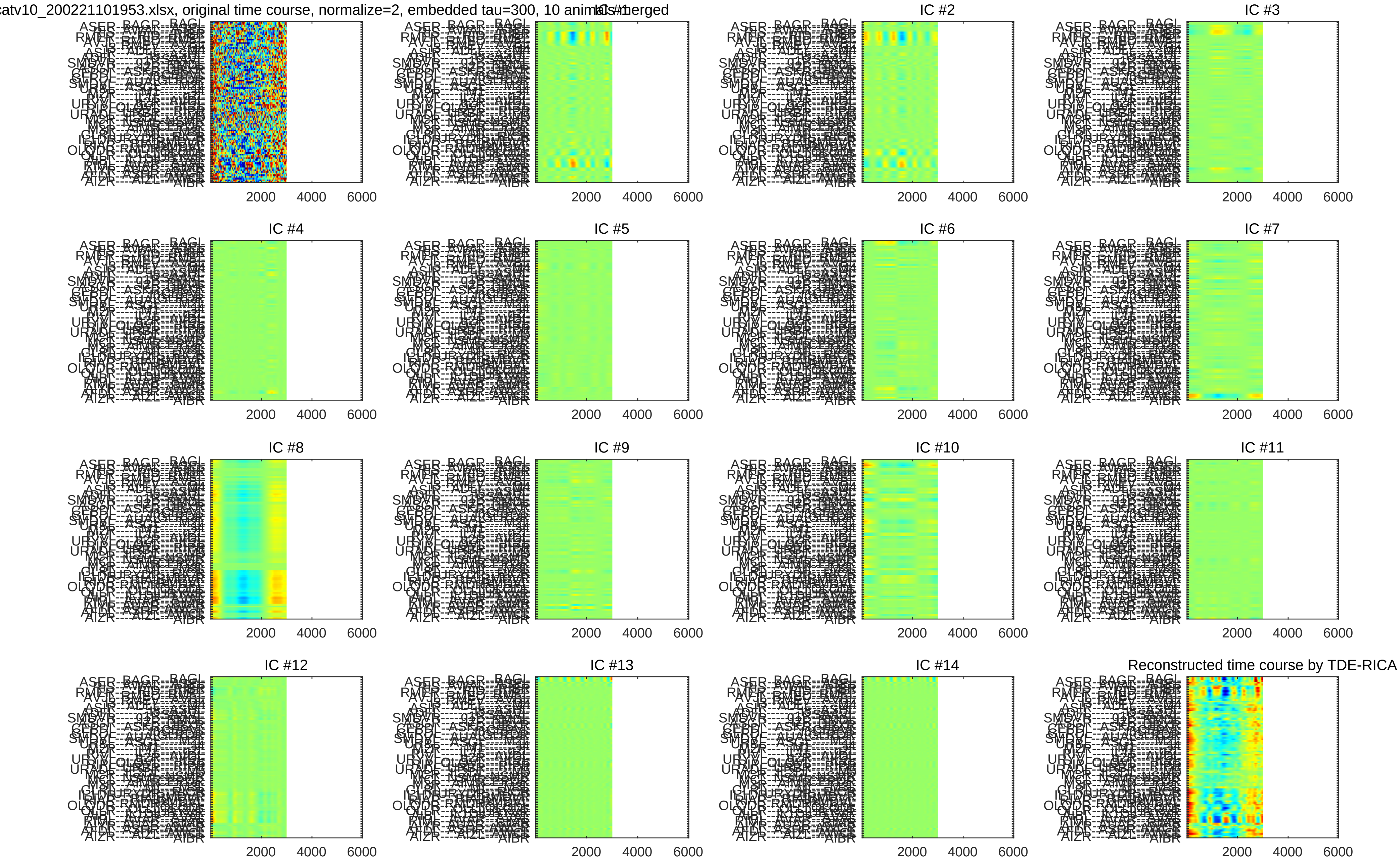
catv10_200221101953.xlsx, original time course, normalize=2, embedded tau=300, 10 animals merged
IC #1
IC #2
IC #3
IC #4
IC #5
IC #6
IC #7
IC #8
IC #9
IC #10
IC #11
IC #12
IC #13
IC #14
Reconstructed time course by TDE-RICA
2000
4000
6000

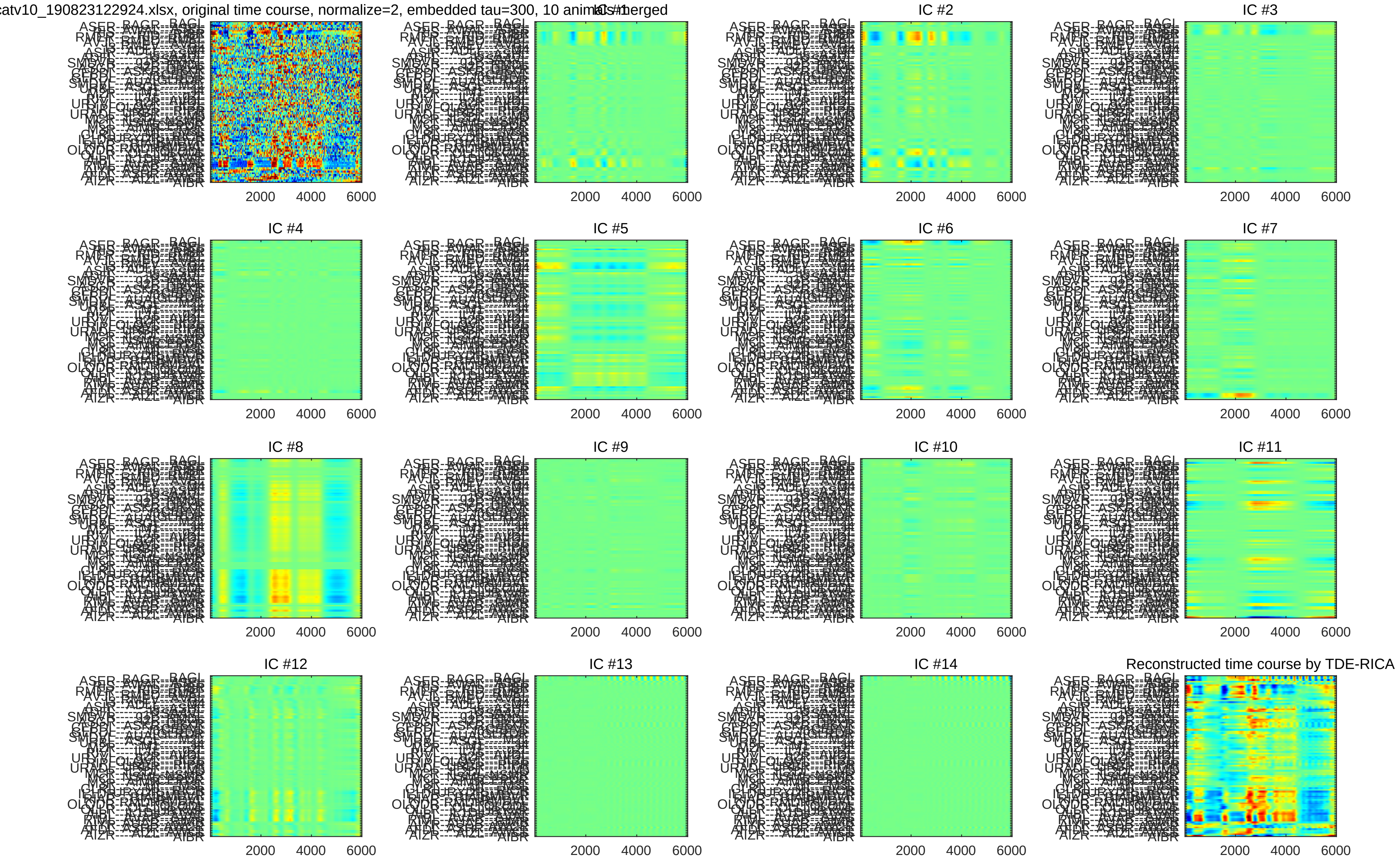
catv10_190823122924.xlsx, original time course, normalize=2, embedded tau=300, 10 animals merged
IC #1
IC #2
IC #3
IC #4
IC #5
IC #6
IC #7
IC #8
IC #9
IC #10
IC #11
IC #12
IC #13
IC #14
Reconstructed time course by TDE-RICA
2000
4000
6000
