## Supplementary File 3 for "Deducing ensemble dynamics and information flow from the whole-brain imaging data"

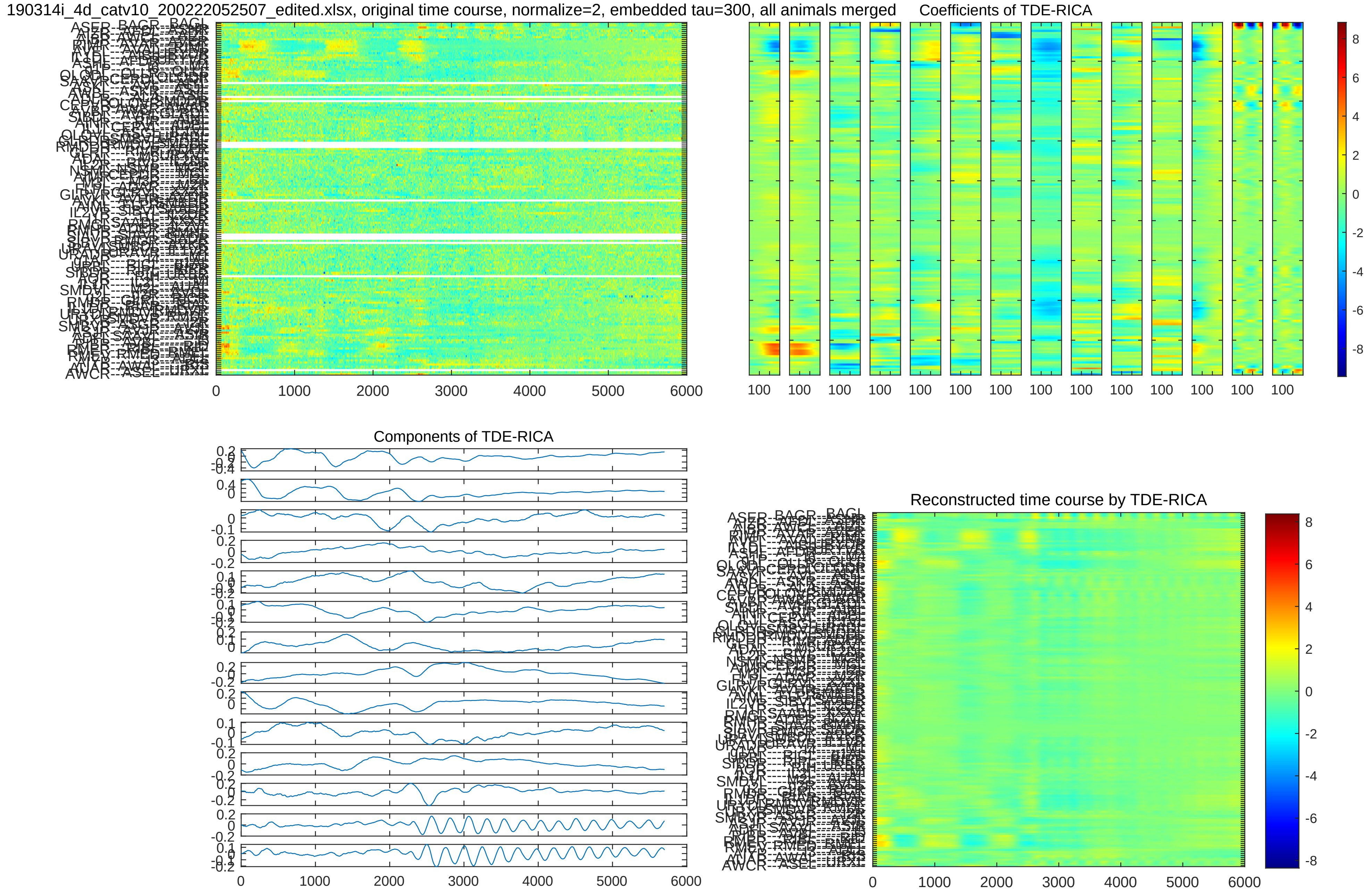
190314i_4d_catv10_200222052507_edited.xlsx, original time course, normalize=2, embedded tau=300, all animals merged
Coefficients of TDE-RICA
Components of TDE-RICA
Reconstructed time course by TDE-RICA
0
1000
2000
3000
4000
5000
6000
100

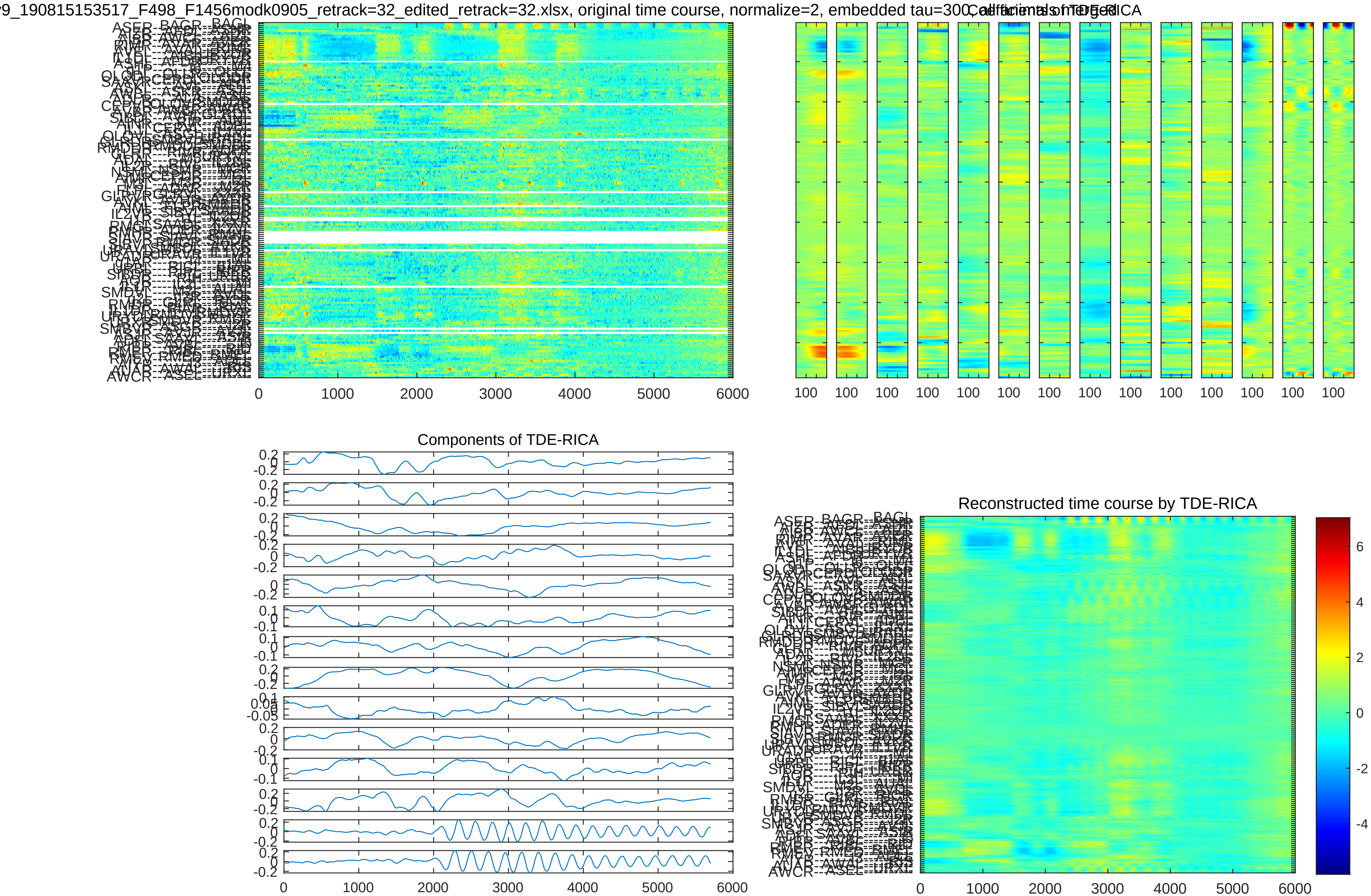
9_190815153517_F498_F1456modk0905_retrack=32_edited_retrack=32.xlsx, original time course, normalize=2, embedded tau=300
Coefficients of TDE-RICA
Components of TDE-RICA
Reconstructed time course by TDE-RICA

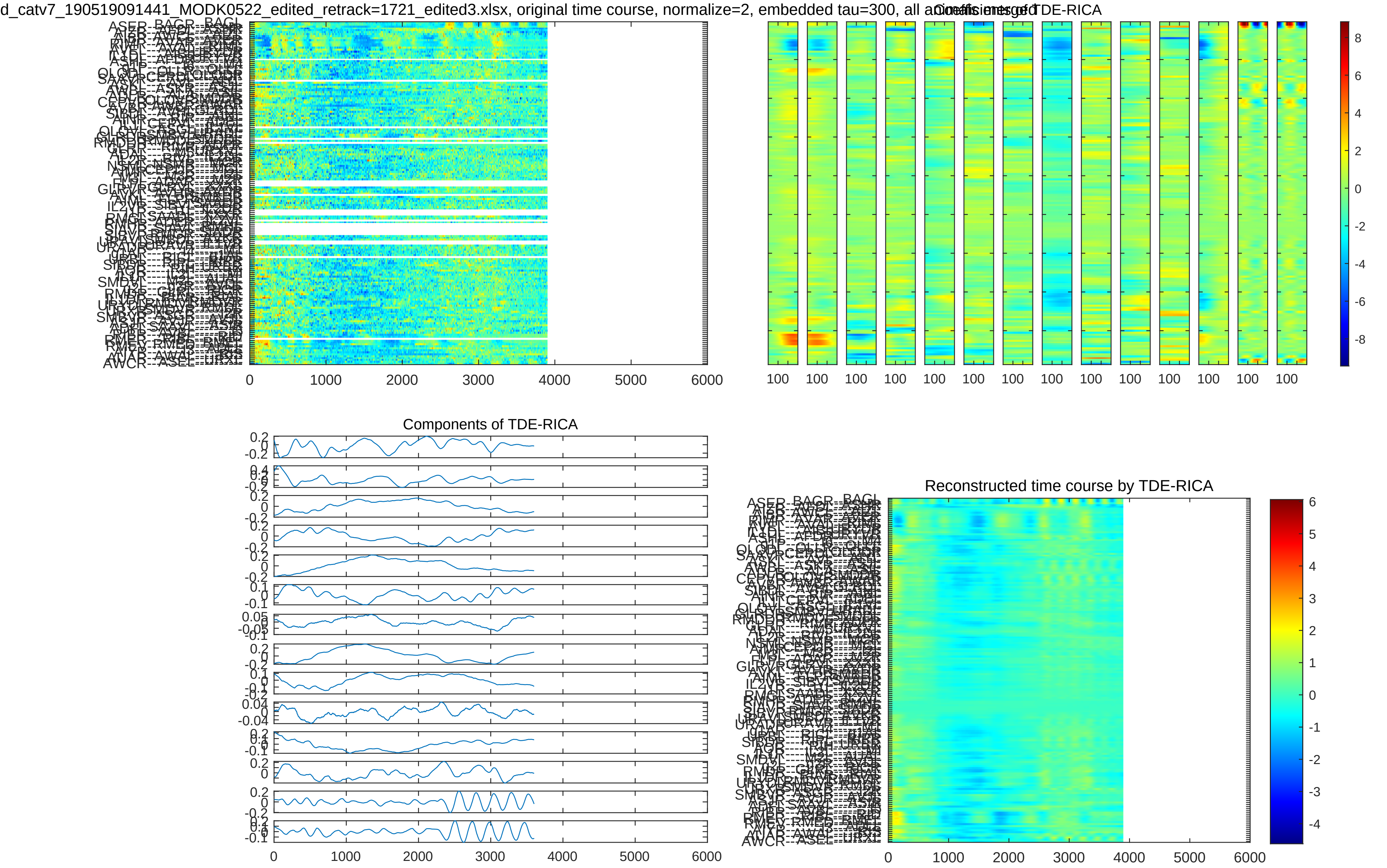
d_catv7_190519091441_MODK0522_edited_retrack=1721_edited3.xlsx, original time course, normalize=2, embedded tau=300, all animals merged
Coefficients of TDE-RICA
Components of TDE-RICA
Reconstructed time course by TDE-RICA

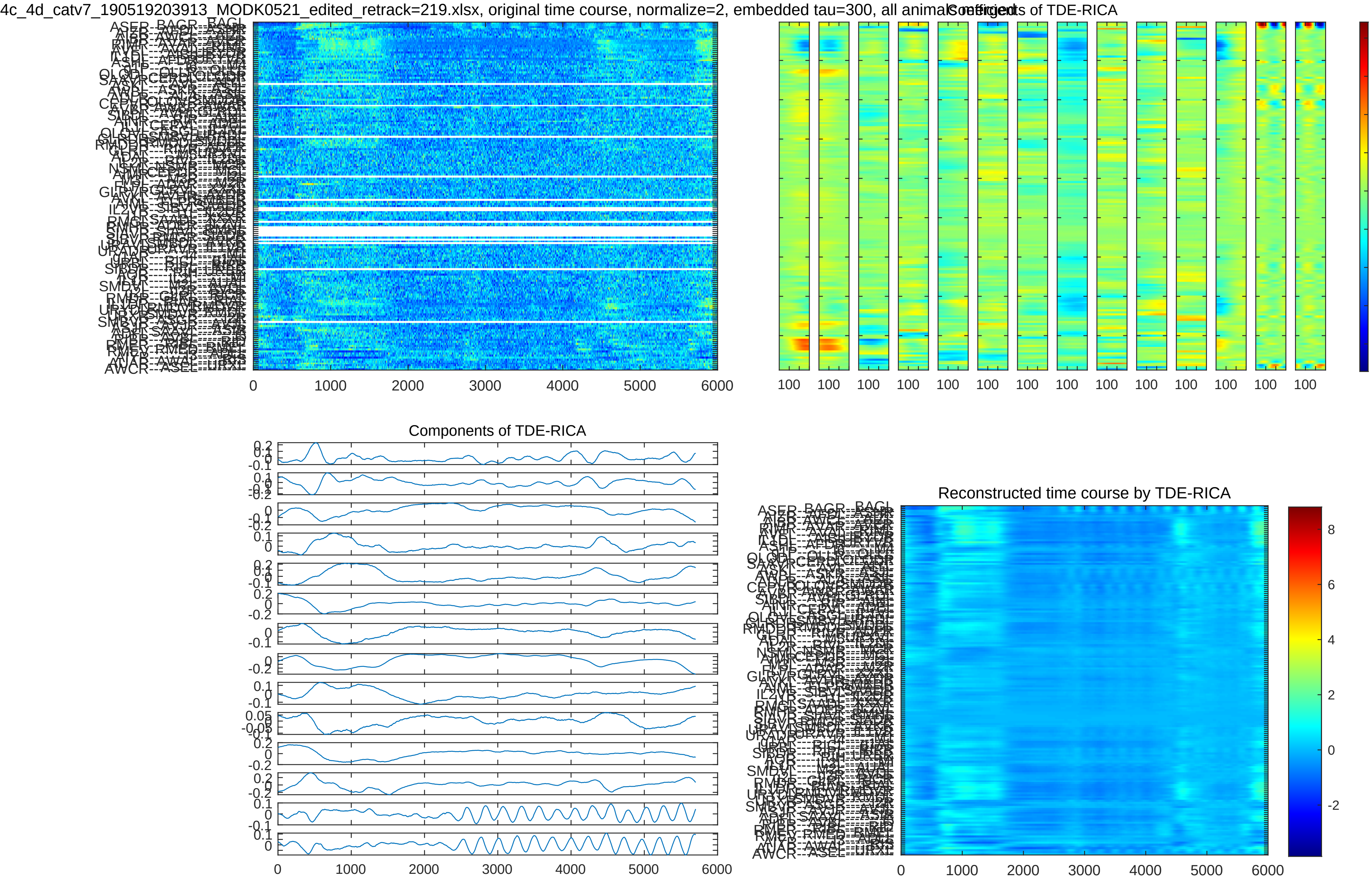
4c_4d_catv7_190519203913_MODK0521_edited_retrack=219.xlsx, original time course, normalize=2, embedded tau=300, all animals
Coefficients of TDE-RICA
Components of TDE-RICA
Reconstructed time course by TDE-RICA

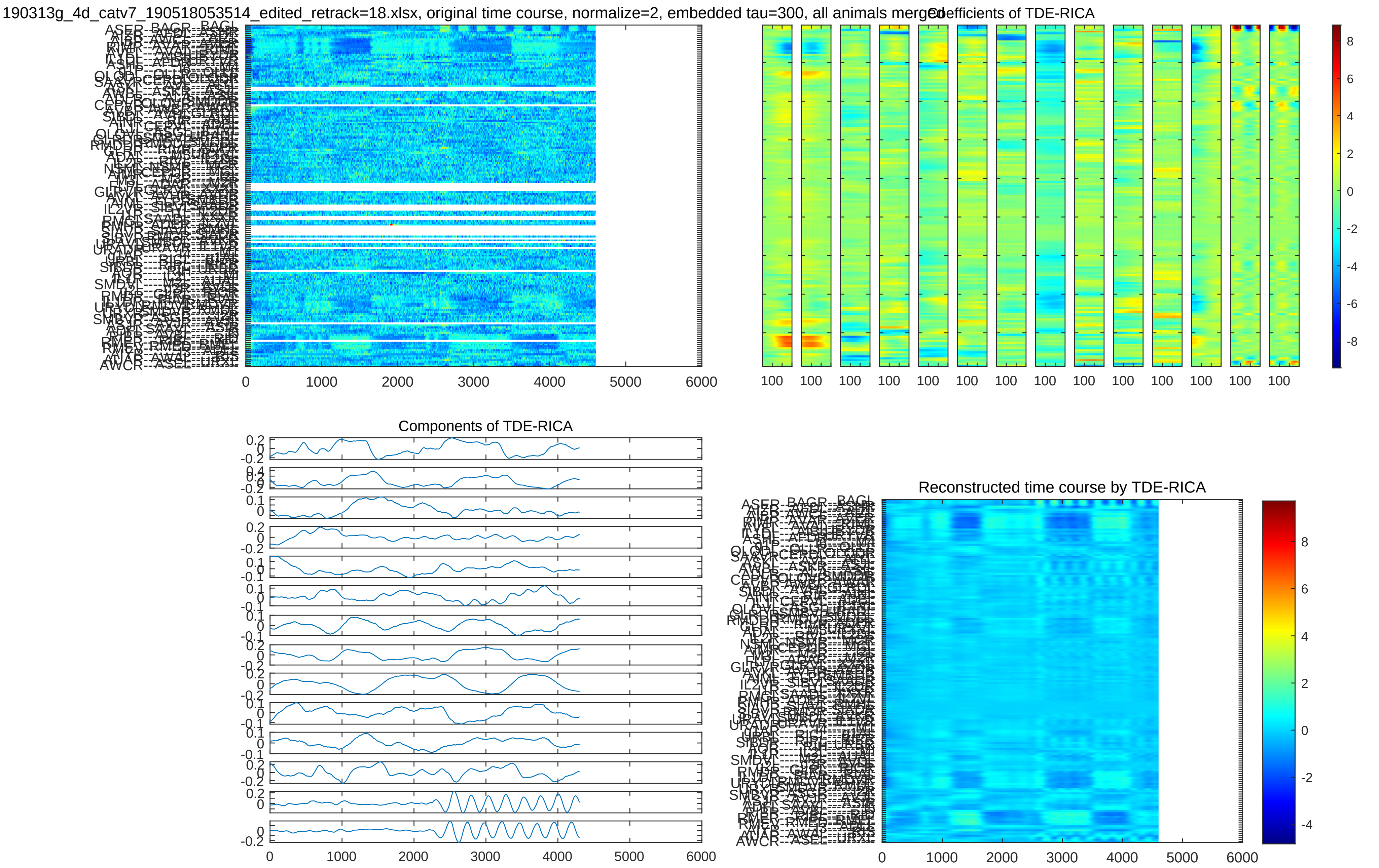

190313g_4d_catv7_190518053514_edited_retrack=18.xlsx, original time course, normalize=2, embedded tau=300, all animals merged
Coefficients of TDE-RICA
Components of TDE-RICA
Reconstructed time course by TDE-RICA
0
1000
2000
3000
4000
5000
6000

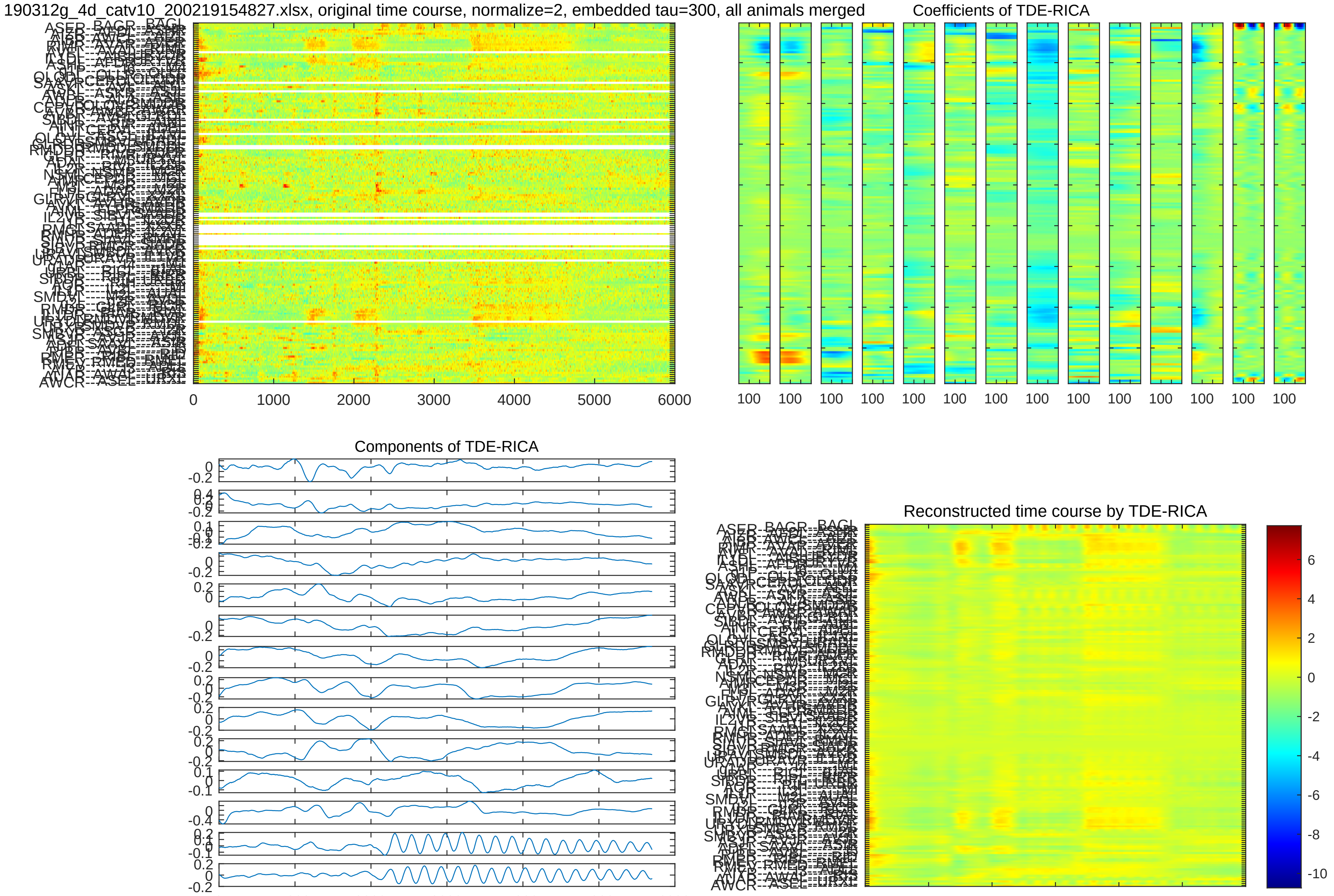
190312g_4d_catv10_200219154827.xlsx, original time course, normalize=2, embedded tau=300, all animals merged
Coefficients of TDE-RICA
Components of TDE-RICA
Reconstructed time course by TDE-RICA

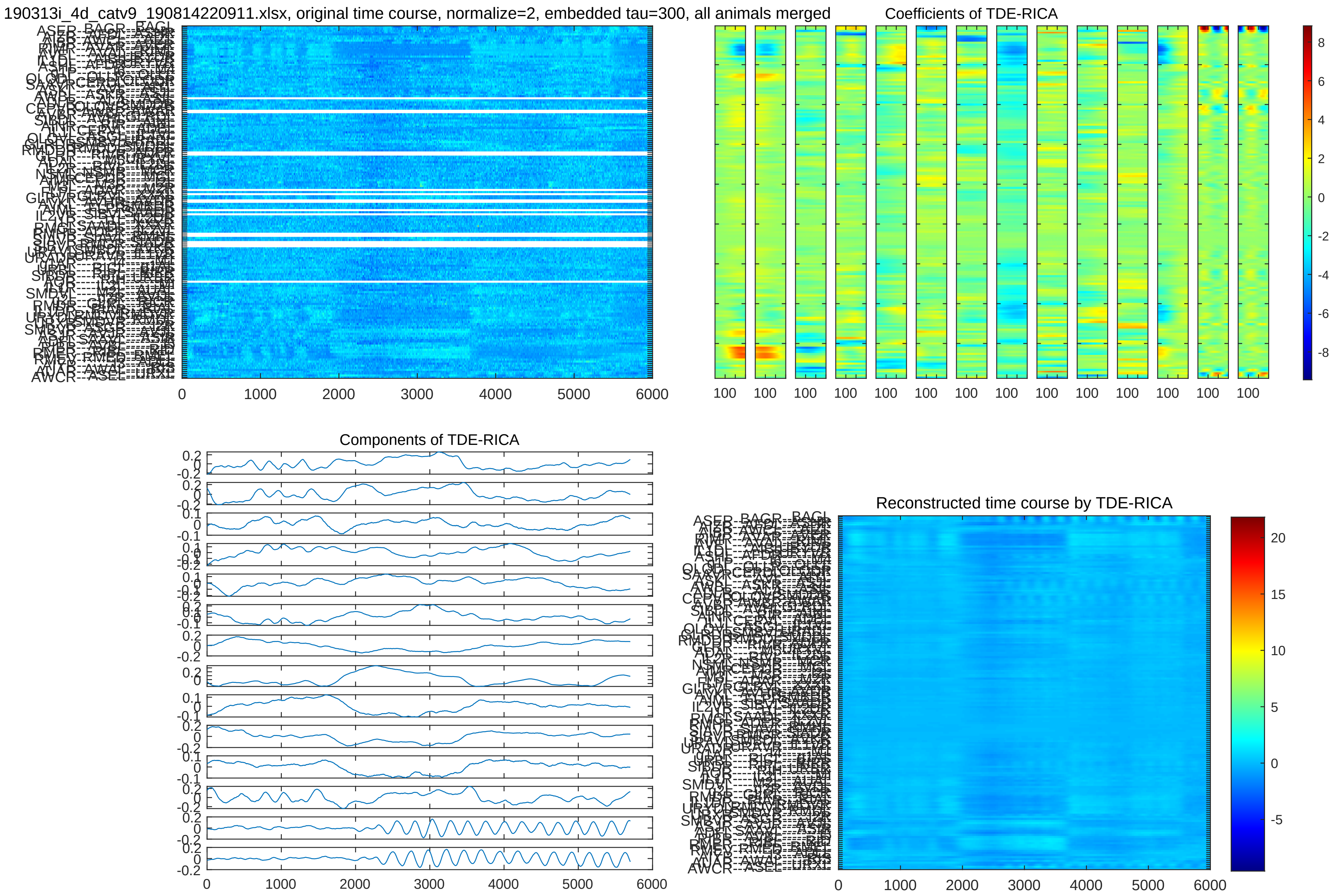

190313i_4d_catv9_190814220911.xlsx, original time course, normalize=2, embedded tau=300, all animals merged
0
1000
2000
3000
4000
5000
6000
Coefficients of TDE-RICA
100
8
6
4
2
0
-2
-4
-6
-8
Components of TDE-RICA
Reconstructed time course by TDE-RICA
20
15
10
5
0
-5

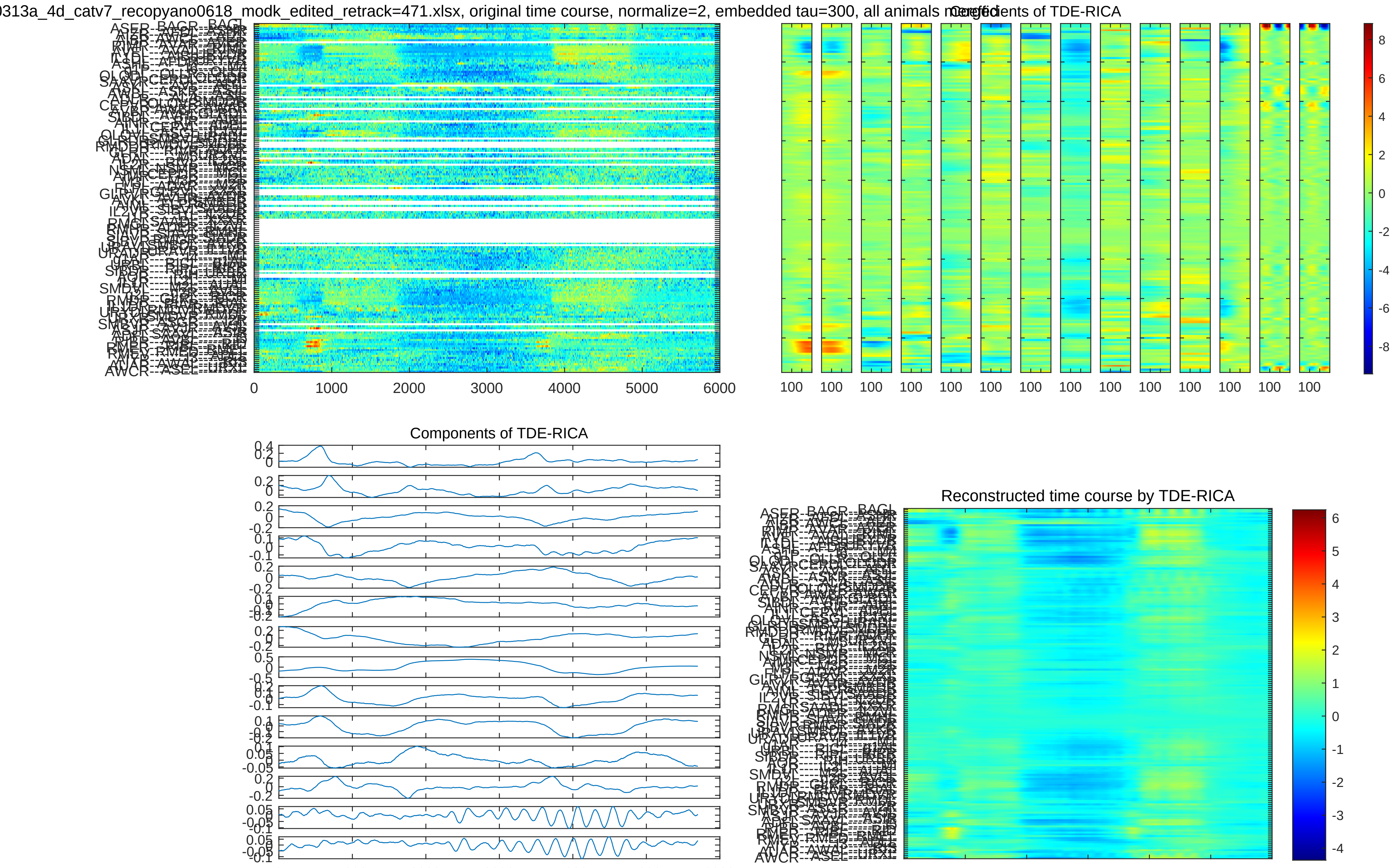

0313a_4d_catv7_recopyano0618_modk_edited_retrack=471.xlsx, original time course, normalize=2, embedded tau=300, all animals merged
Coefficients of TDE-RICA
0 1000 2000 3000 4000 5000 6000
100 100 100 100 100 100 100 100 100 100 100 100 100 100
8 6 4 2 0 -2 -4 -6 -8
Components of TDE-RICA
Reconstructed time course by TDE-RICA
6 5 4 3 2 1 0 -1 -2 -3 -4

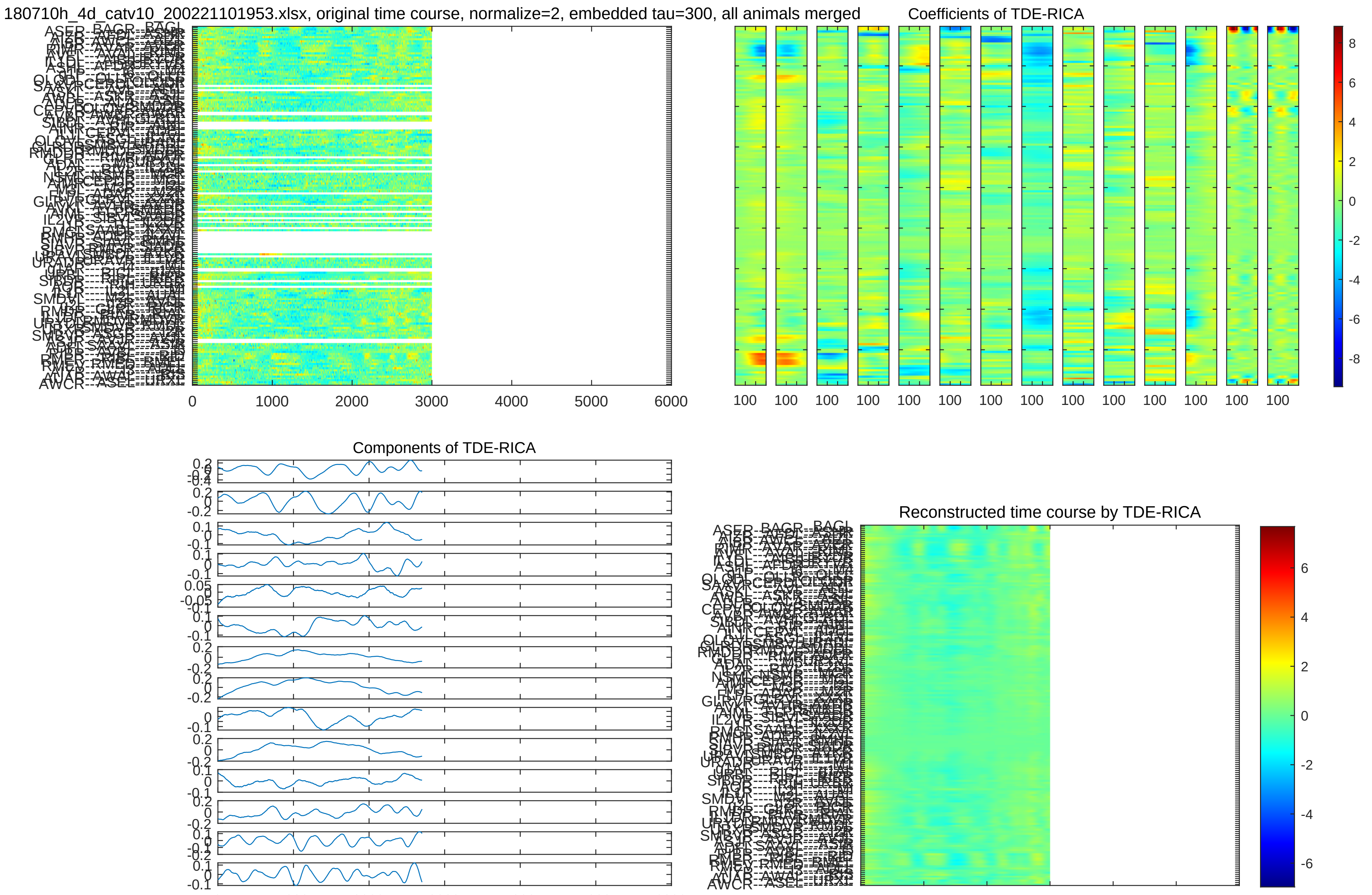

180710h_4d_catv10_200221101953.xlsx, original time course, normalize=2, embedded tau=300, all animals merged
Coefficients of TDE-RICA
Components of TDE-RICA
Reconstructed time course by TDE-RICA
0
1000
2000
3000
4000
5000
6000

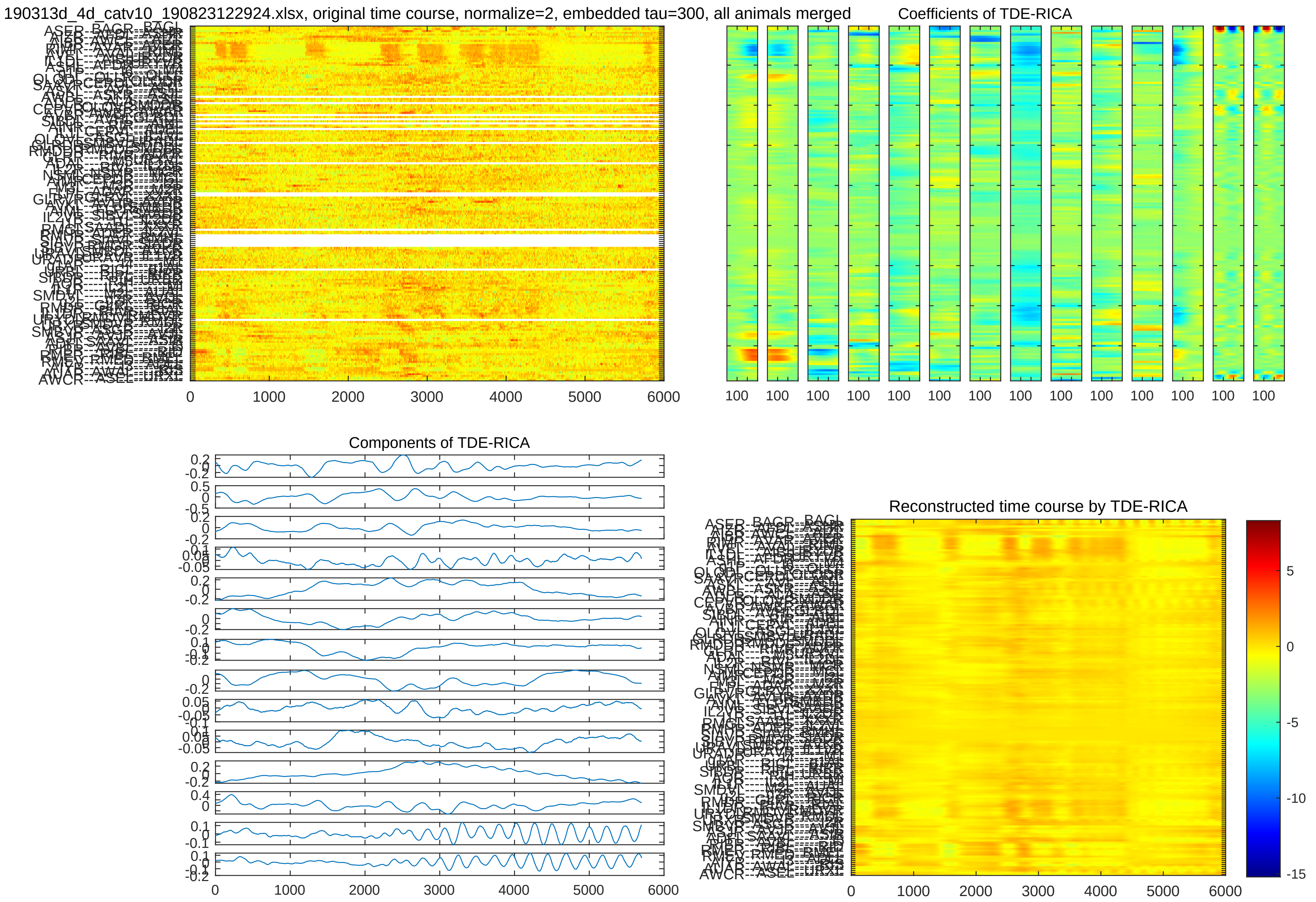
190313d_4d_catv10_190823122924.xlsx, original time course, normalize=2, embedded tau=300, all animals merged
Coefficients of TDE-RICA
Components of TDE-RICA
Reconstructed time course by TDE-RICA
0
1000
2000
3000
4000
5000
6000
100

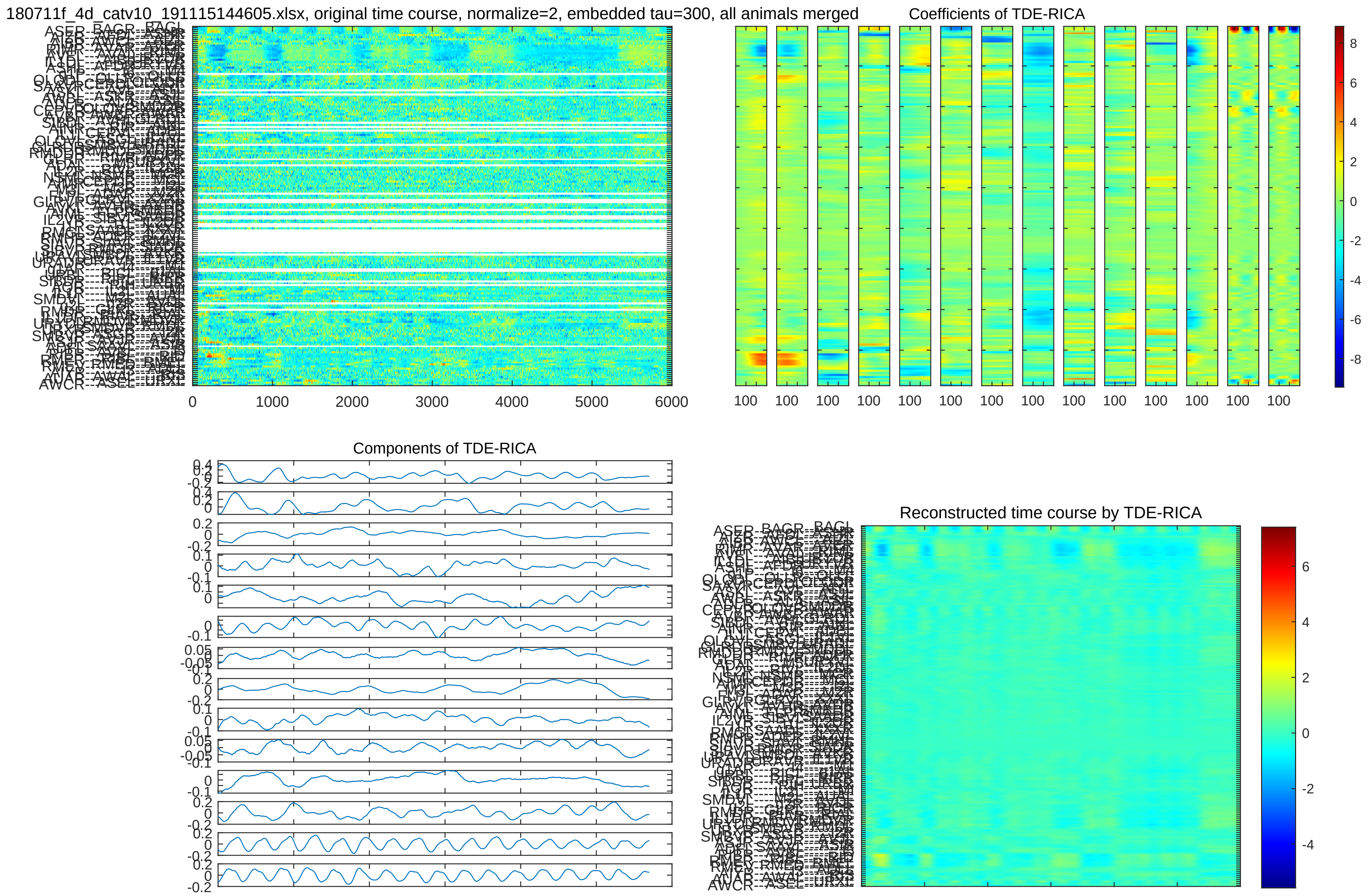

180711f_4d_catv10_191115144605.xlsx, original time course, normalize=2, embedded tau=300, all animals merged
Coefficients of TDE-RICA
Components of TDE-RICA
Reconstructed time course by TDE-RICA
0
1000
2000
3000
4000
5000
6000
100
8
6
4
2
0
-2
-4
-6
-8

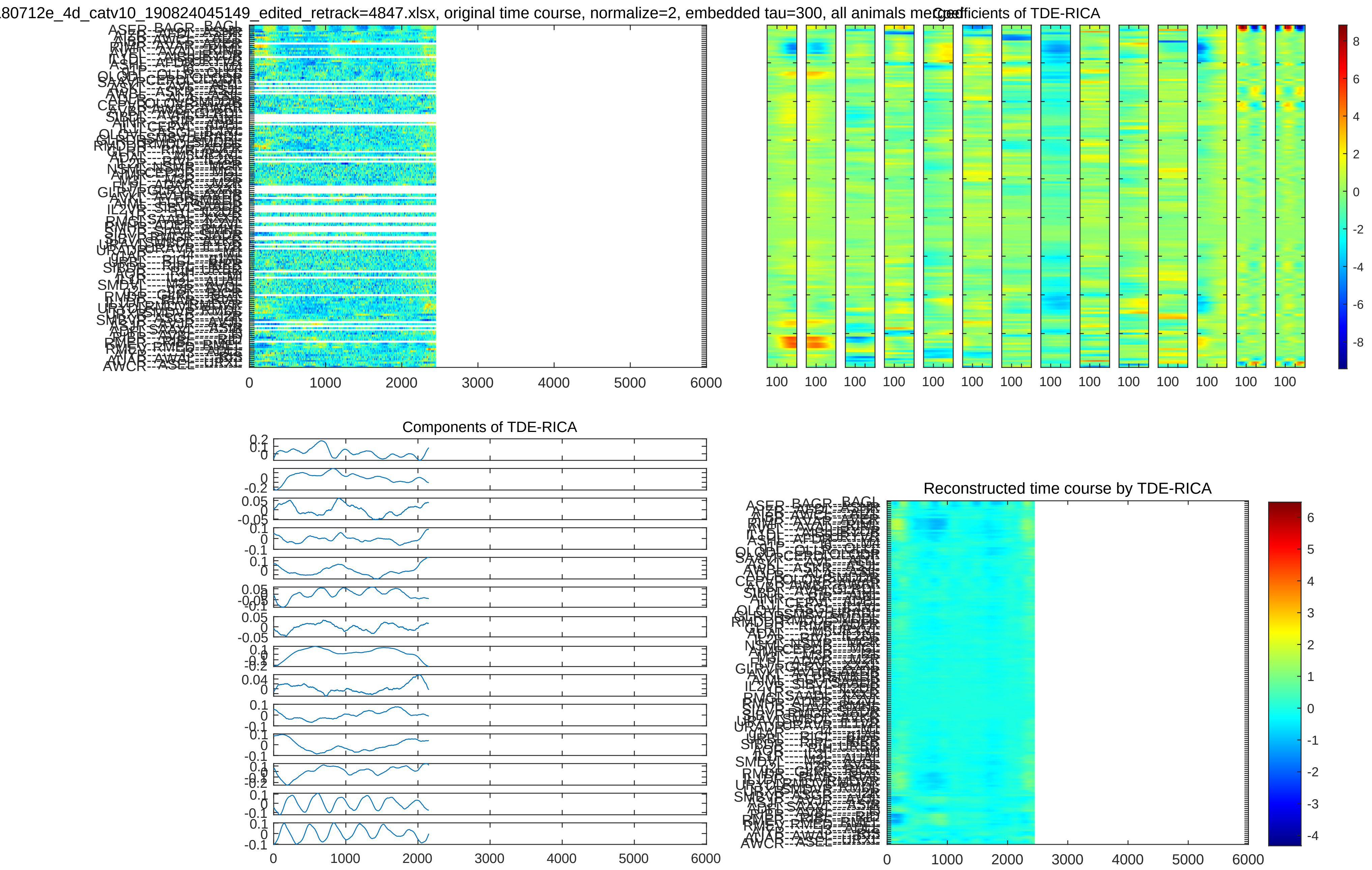

180712e_4d_catv10_190824045149_edited_retrack=4847.xlsx, original time course, normalize=2, embedded tau=300, all animals merged
Coefficients of TDE-RICA
Components of TDE-RICA
Reconstructed time course by TDE-RICA
0 1000 2000 3000 4000 5000 6000

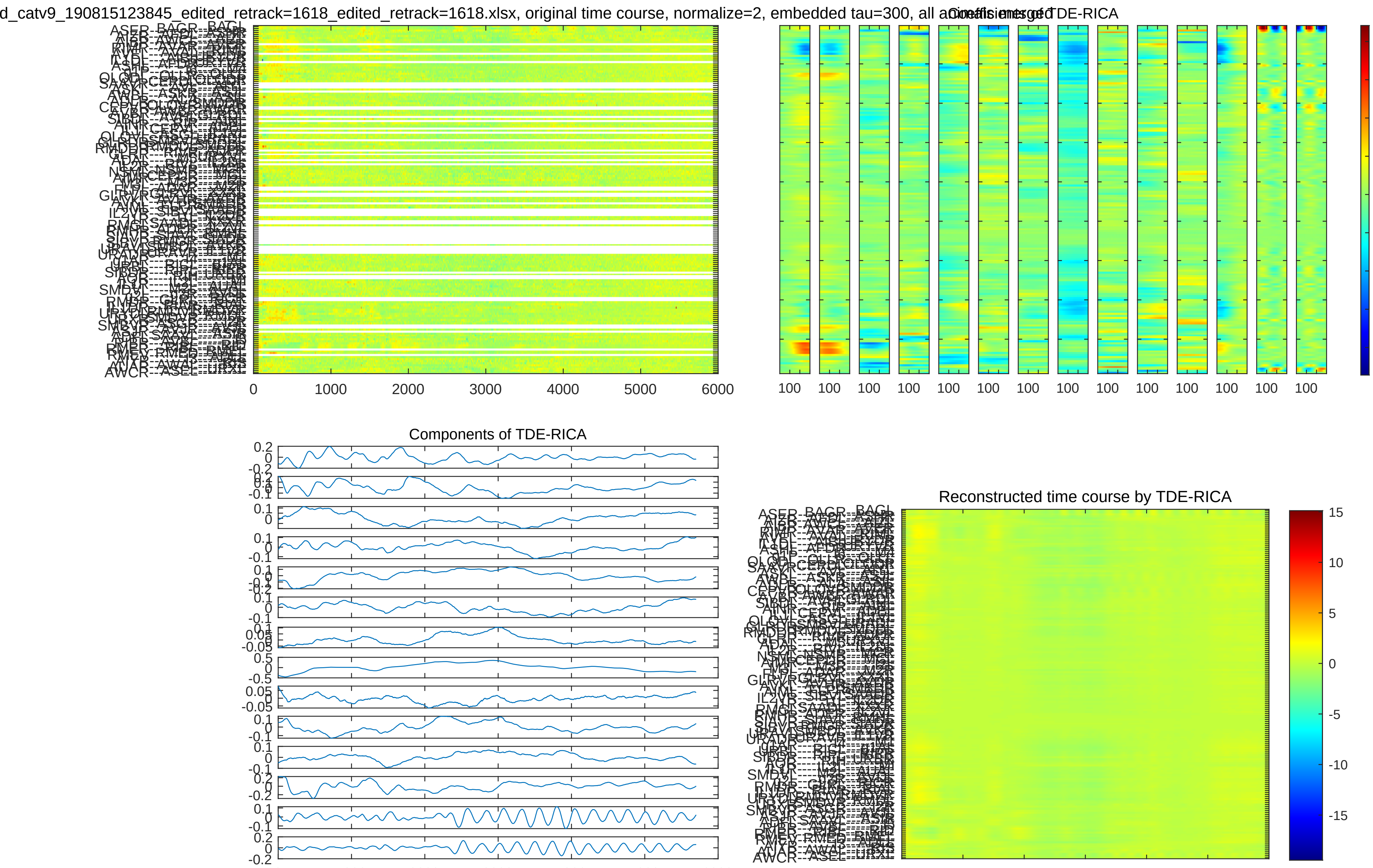
d_catv9_190815123845_edited_retrack=1618_edited_retrack=1618.xlsx, original time course, normalize=2, embedded tau=300, all animals merged
Coefficients of TDE-RICA
Components of TDE-RICA
Reconstructed time course by TDE-RICA

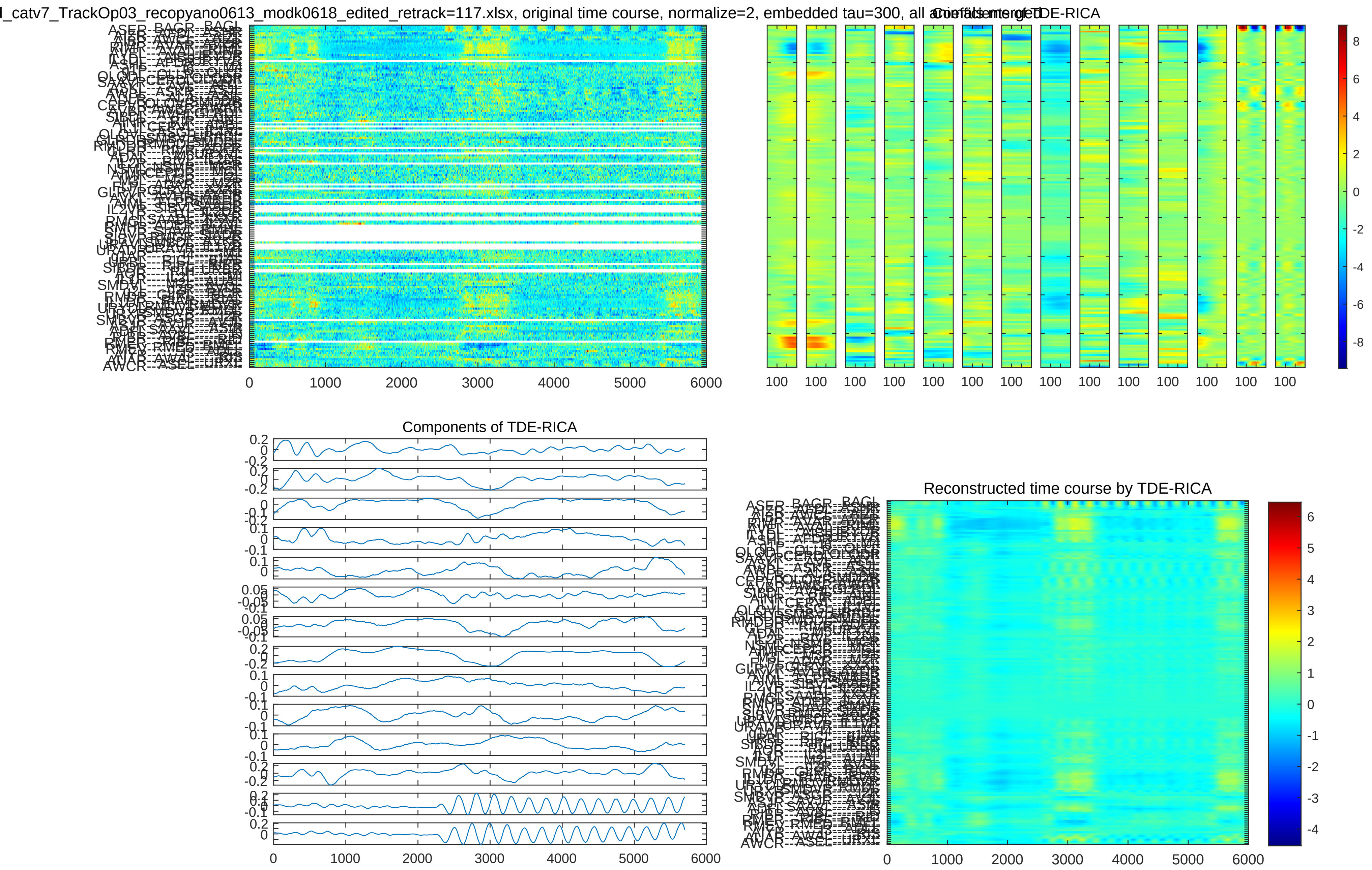

_catv7_TrackOp03_recopyano0613_modk0618_edited_retrack=117.xlsx, original time course, normalize=2, embedded tau=300, all animals merged
Coefficients of TDE-RICA
Components of TDE-RICA
Reconstructed time course by TDE-RICA

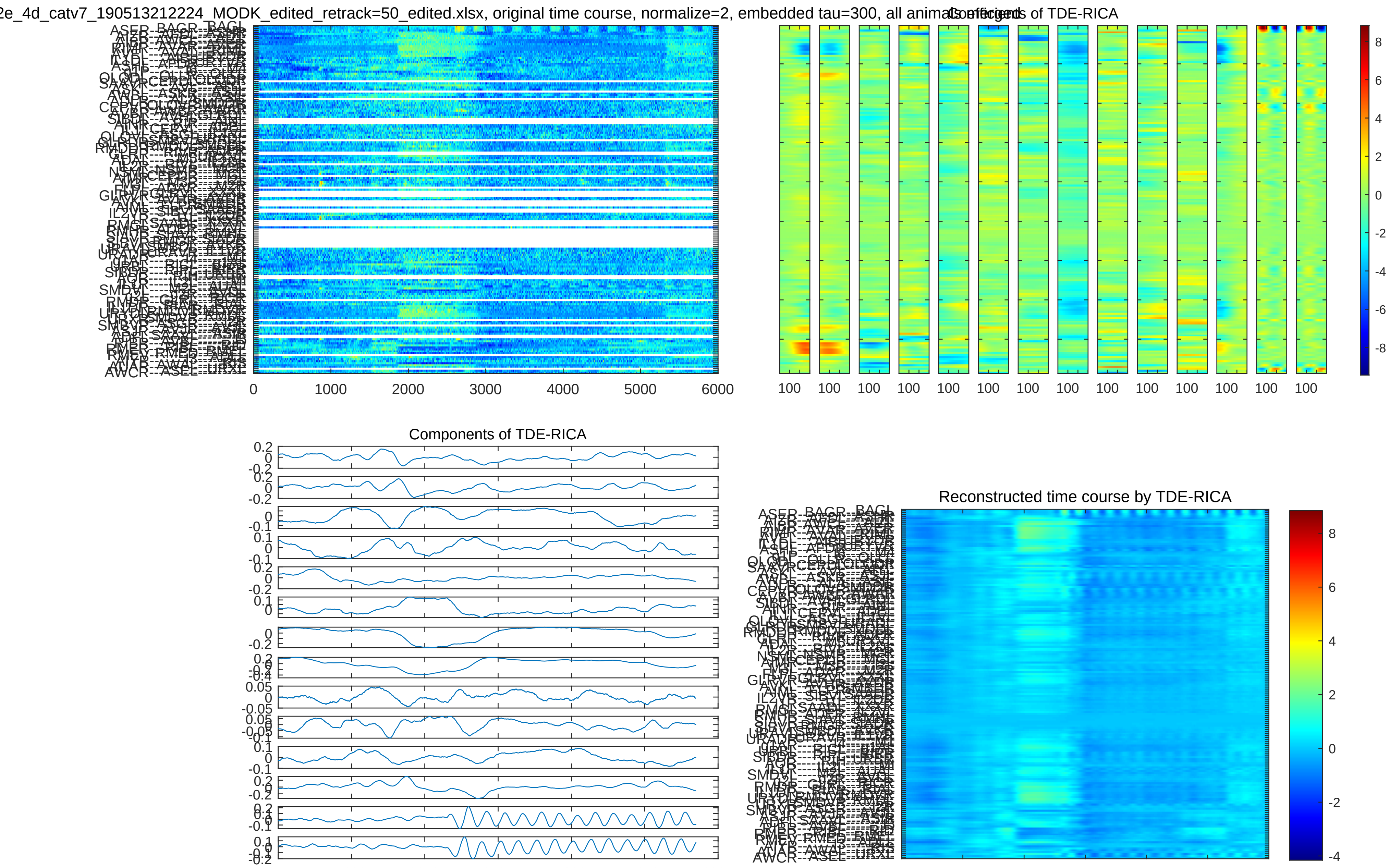
2e_4d_catv7_190513212224_MODK_edited_retrack=50_edited.xlsx, original time course, normalize=2, embedded tau=300, all animals
Coefficients of TDE-RICA
Components of TDE-RICA
Reconstructed time course by TDE-RICA
0 1000 2000 3000 4000 5000 6000
100 100 100 100 100 100 100 100 100 100 100 100 100 100

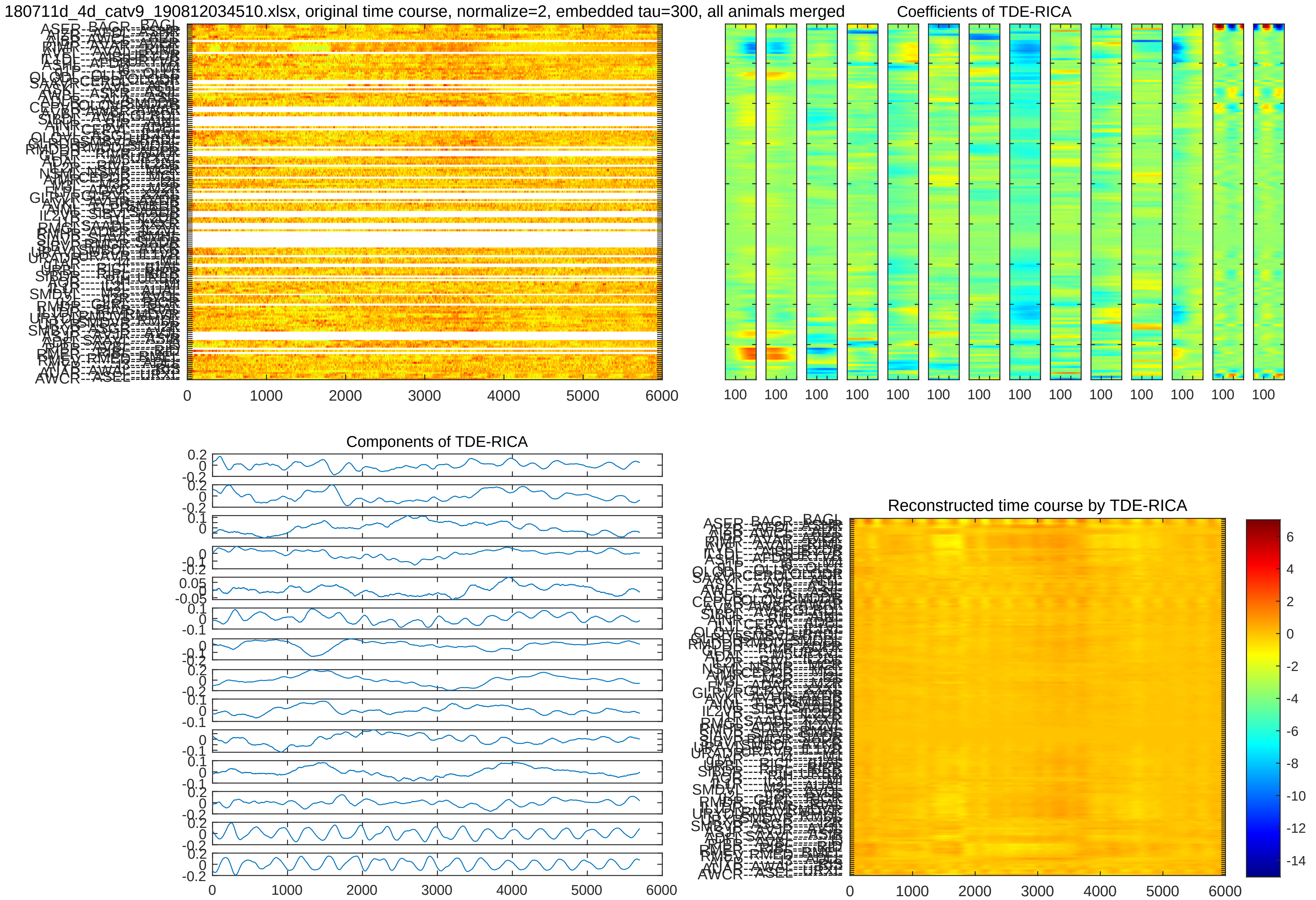

180711d_4d_catv9_190812034510.xlsx, original time course, normalize=2, embedded tau=300, all animals merged
Coefficients of TDE-RICA
Components of TDE-RICA
Reconstructed time course by TDE-RICA

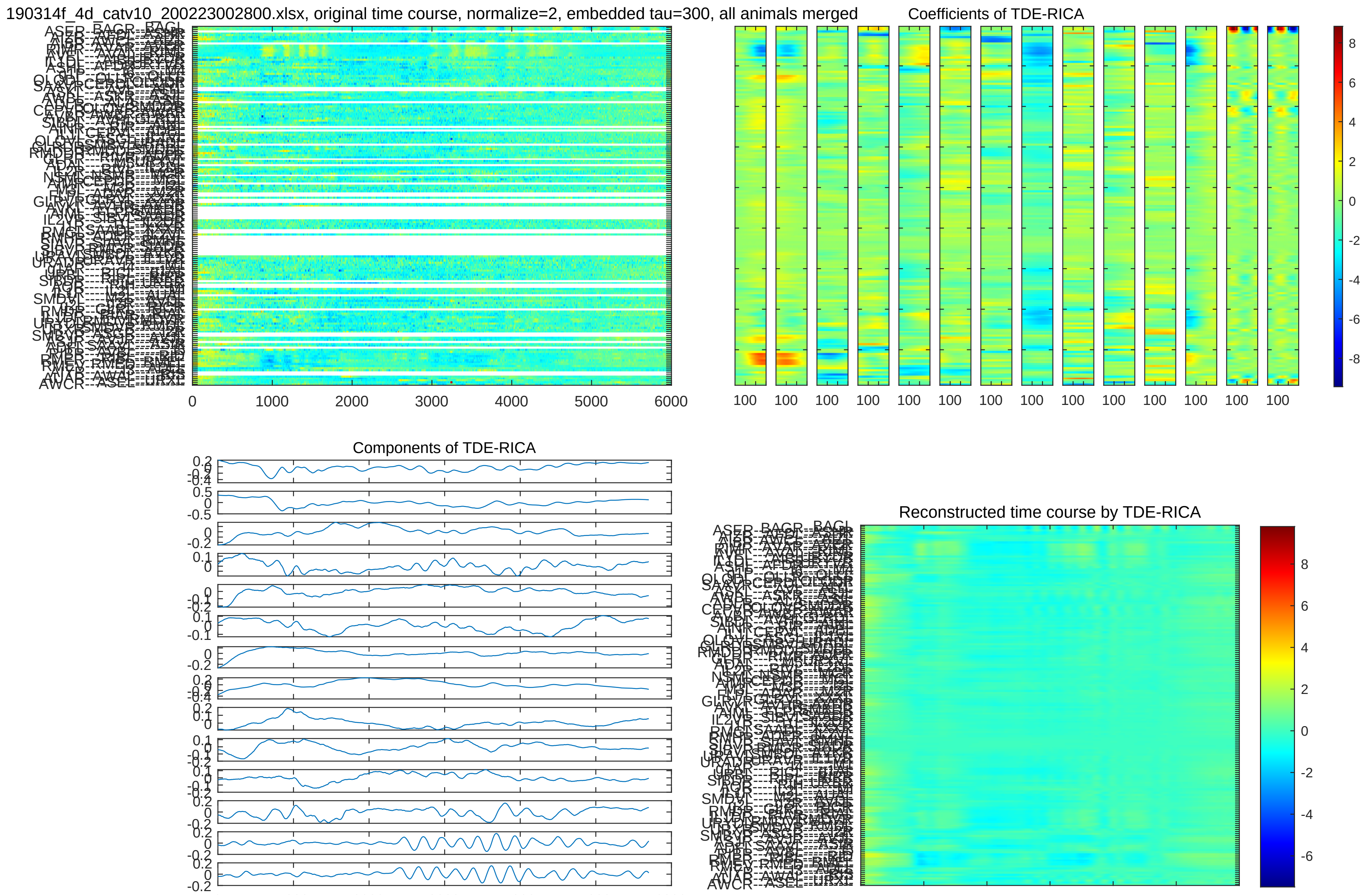
190314f_4d_catv10_200223002800.xlsx, original time course, normalize=2, embedded tau=300, all animals merged
Coefficients of TDE-RICA
Components of TDE-RICA
Reconstructed time course by TDE-RICA

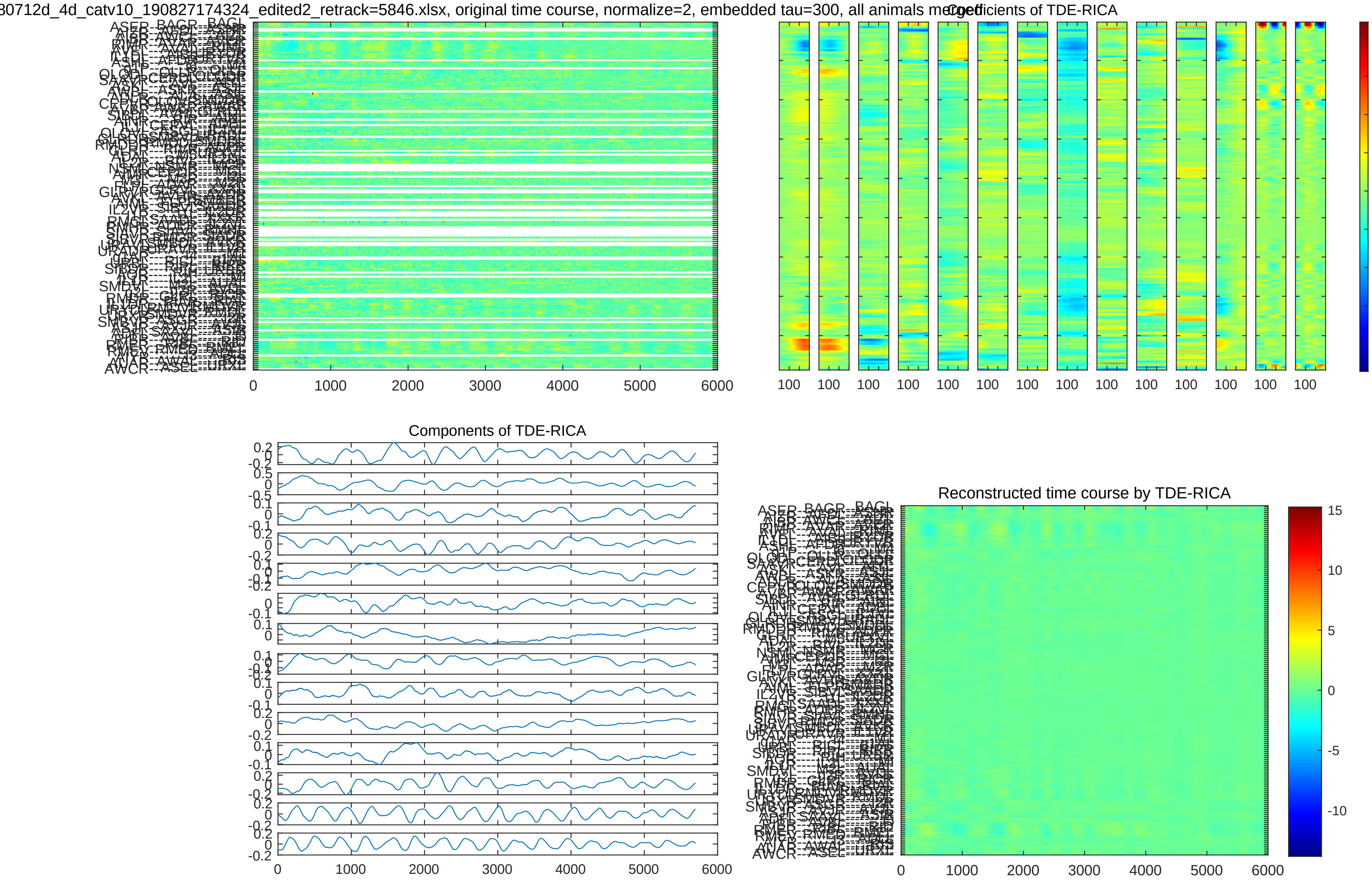
30712d_4d_catv10_190827174324_edited2_retrack=5846.xlsx, original time course, normalize=2, embedded tau=300, all animals merged
Coefficients of TDE-RICA
Components of TDE-RICA
Reconstructed time course by TDE-RICA

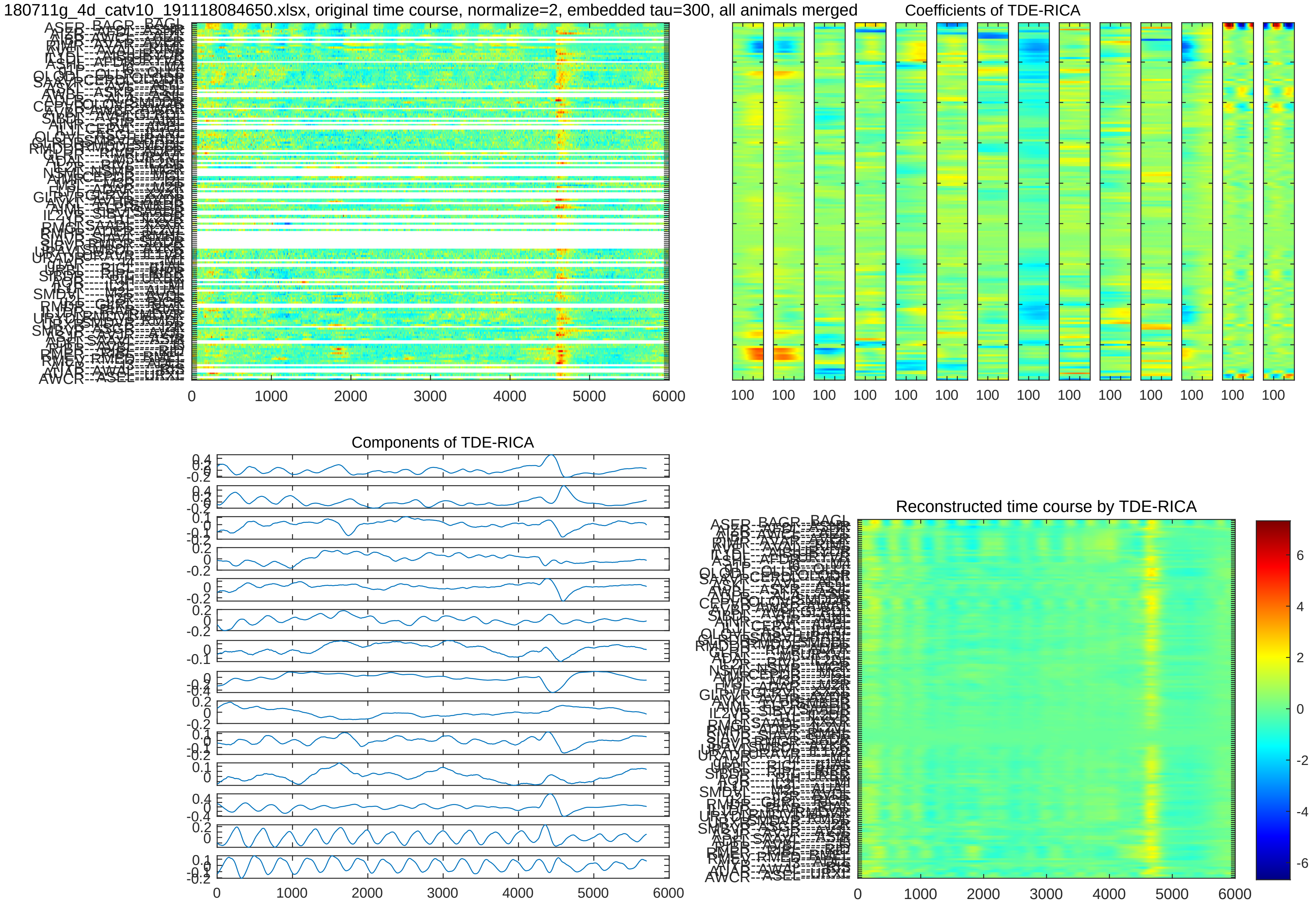
180711g_4d_catv10_191118084650.xlsx, original time course, normalize=2, embedded tau=300, all animals merged
Coefficients of TDE-RICA
Components of TDE-RICA
Reconstructed time course by TDE-RICA

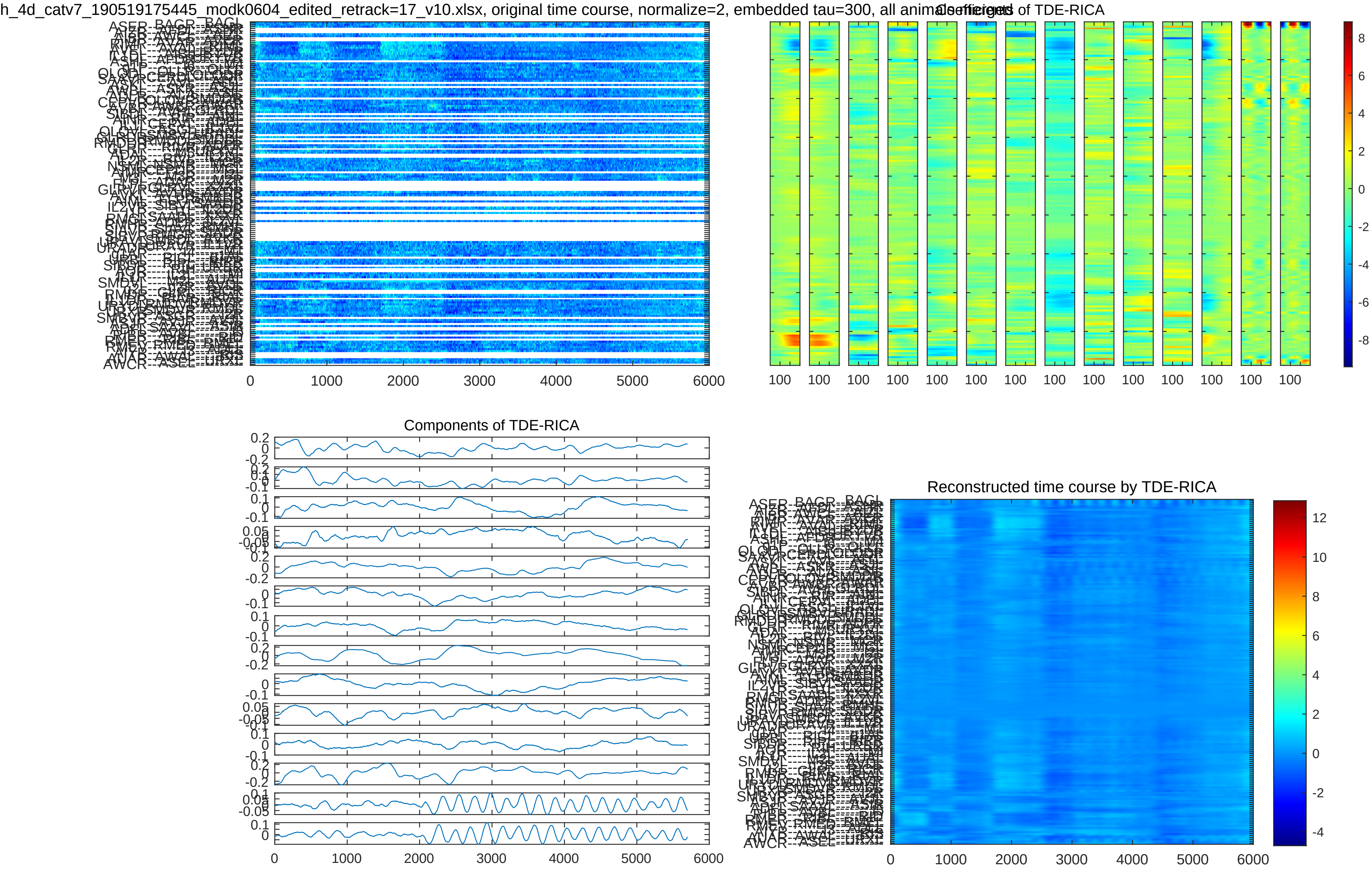
h_4d_catv7_190519175445_modk0604_edited_retrack=17_v10.xlsx, original time course, normalize=2, embedded tau=300, all animals
Coefficients of TDE-RICA
Components of TDE-RICA
Reconstructed time course by TDE-RICA

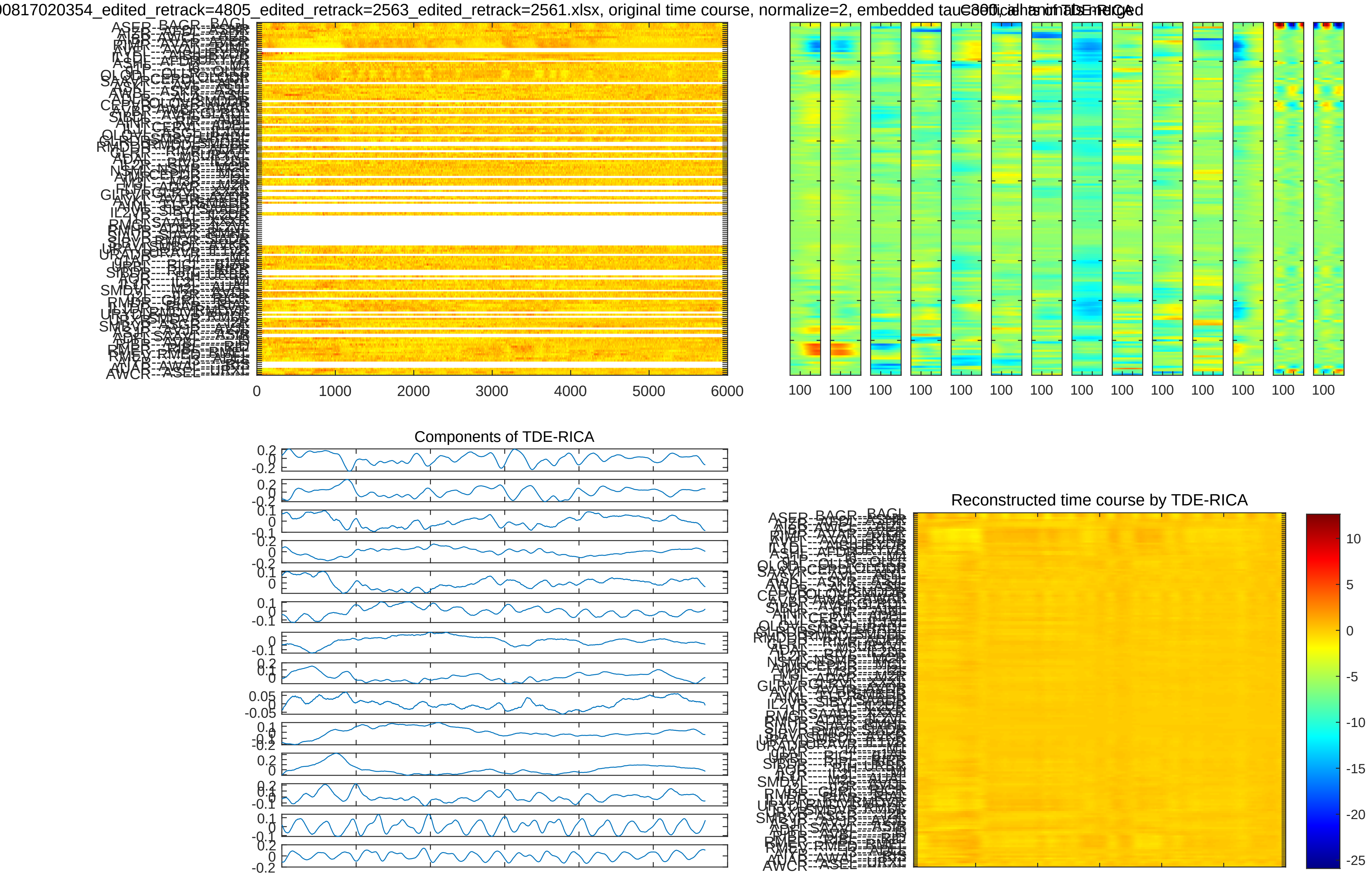
00817020354_edited_retrack=4805_edited_retrack=2563_edited_retrack=2561.xlsx, original time course, normalize=2, embedded tau
Components of TDE-RICA
Reconstructed time course by TDE-RICA

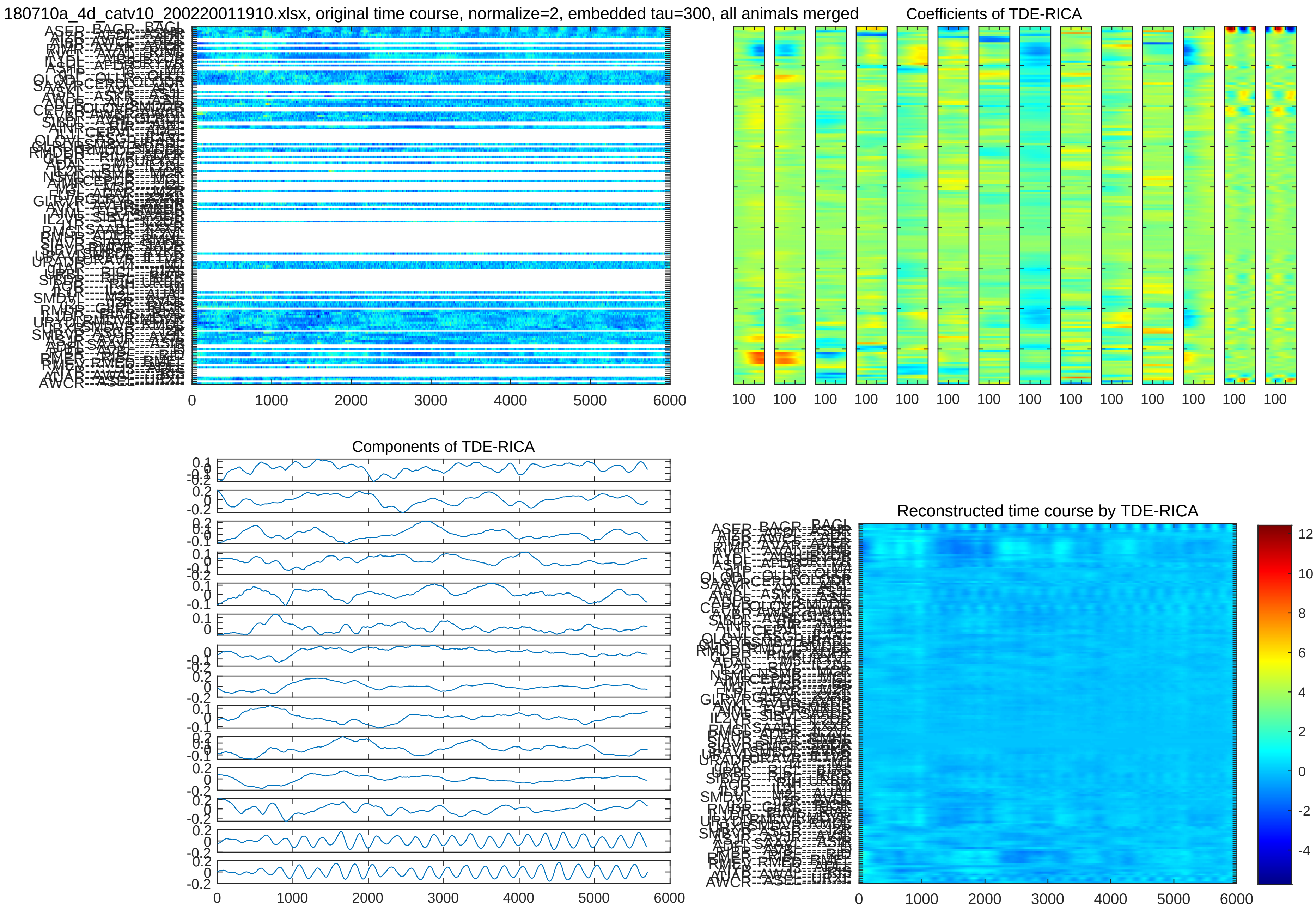
180710a_4d_catv10_200220011910.xlsx, original time course, normalize=2, embedded tau=300, all animals merged
Coefficients of TDE-RICA
Components of TDE-RICA
Reconstructed time course by TDE-RICA
0 1000 2000 3000 4000 5000 6000
100 100 100 100 100 100 100 100 100 100 100 100 100 100

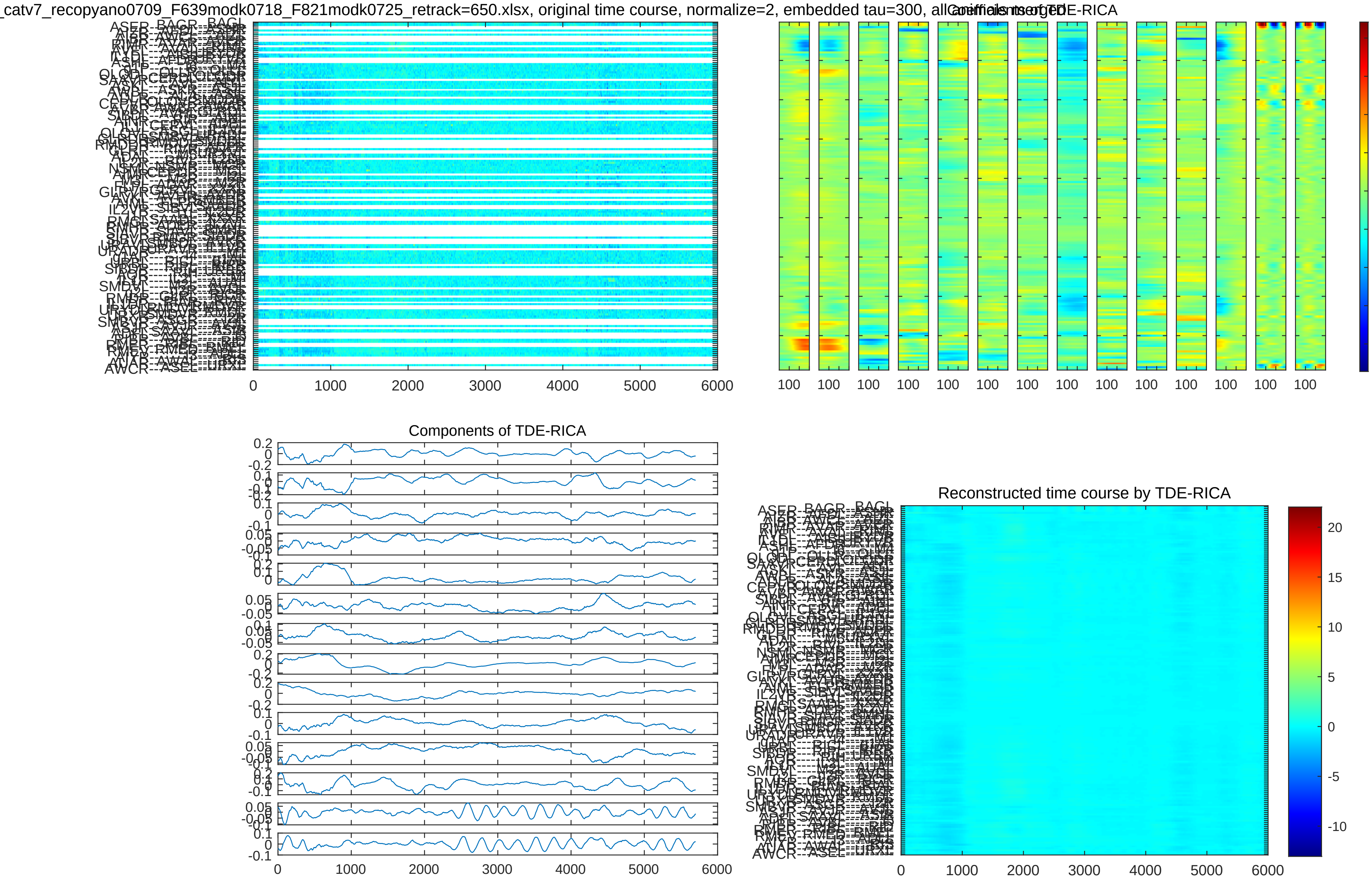
_catv7_recopyano0709_F639modk0718_F821modk0725_retrack=650.xlsx, original time course, normalize=2, embedded tau=300, all
Coefficients of TDE-RICA
Components of TDE-RICA
Reconstructed time course by TDE-RICA

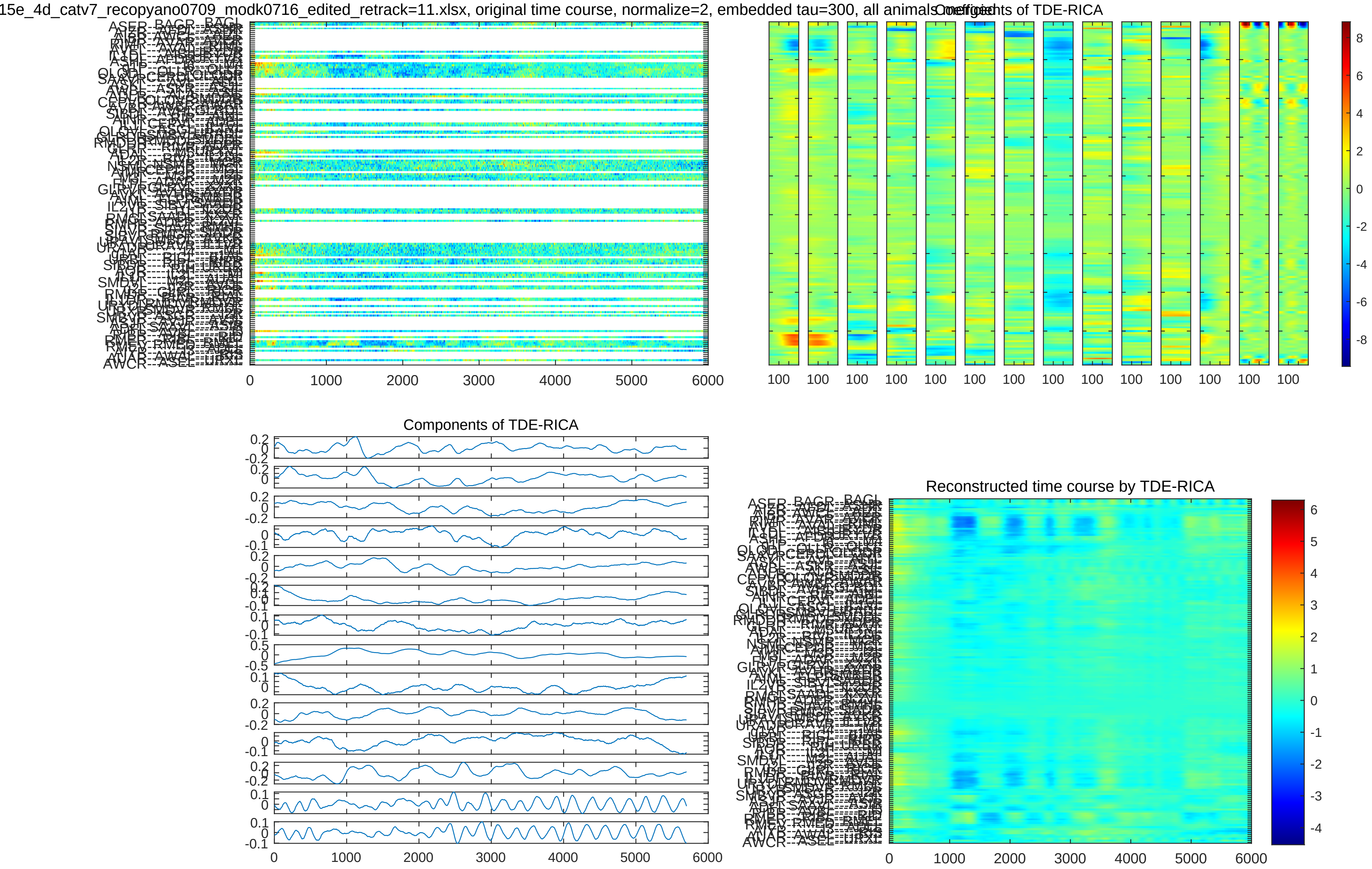
15e_4d_catv7_recopyano0709_modk0716_edited_retrack=11.xlsx, original time course, normalize=2, embedded tau=300, all animals
Coefficients of TDE-RICA
Components of TDE-RICA
Reconstructed time course by TDE-RICA

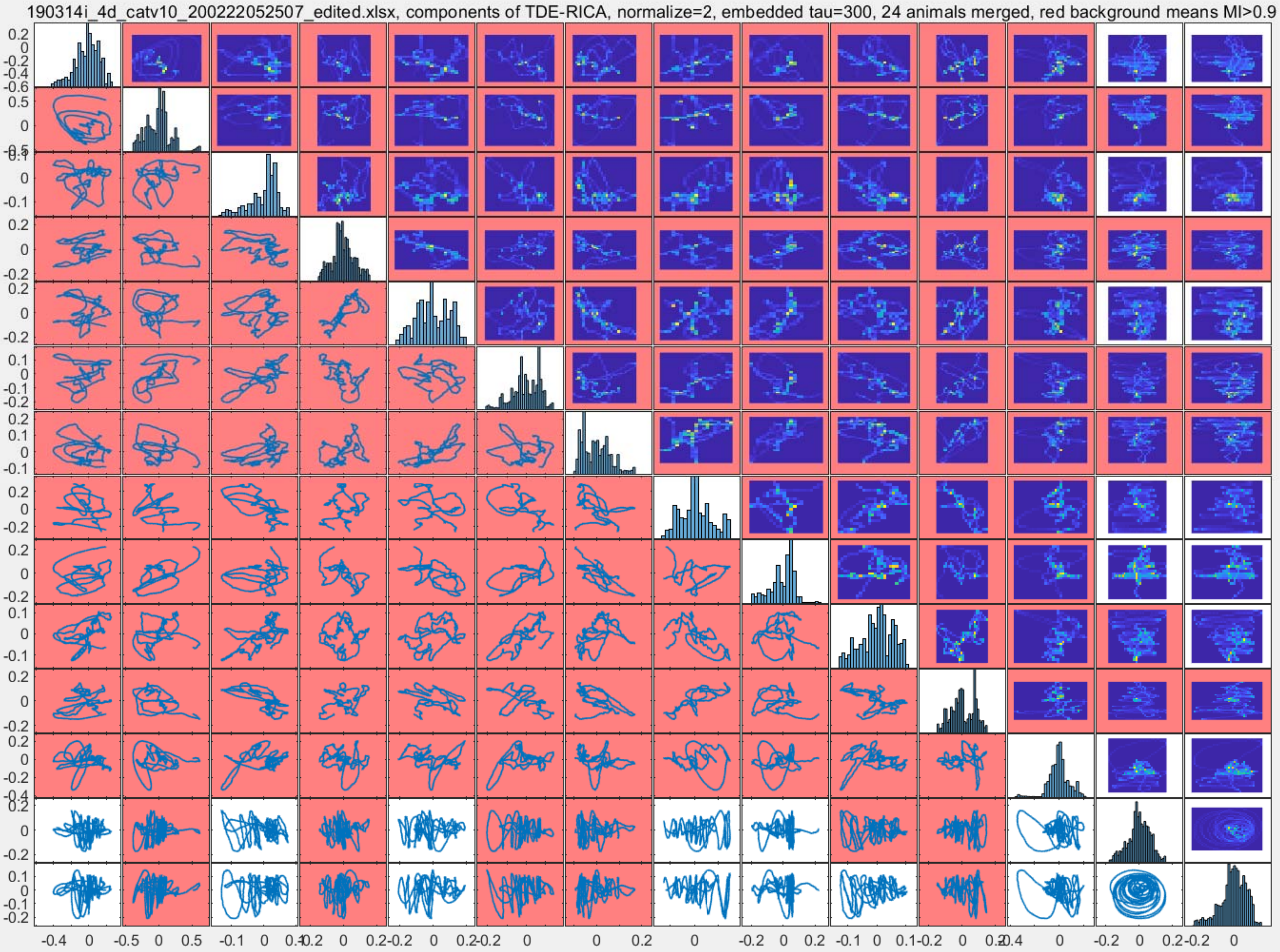
190314i_4d_catv10_200222052507_edited.xlsx, components of TDE-RICA, normalize=2, embedded tau=300, 24 animals merged, red background means MI>0.9

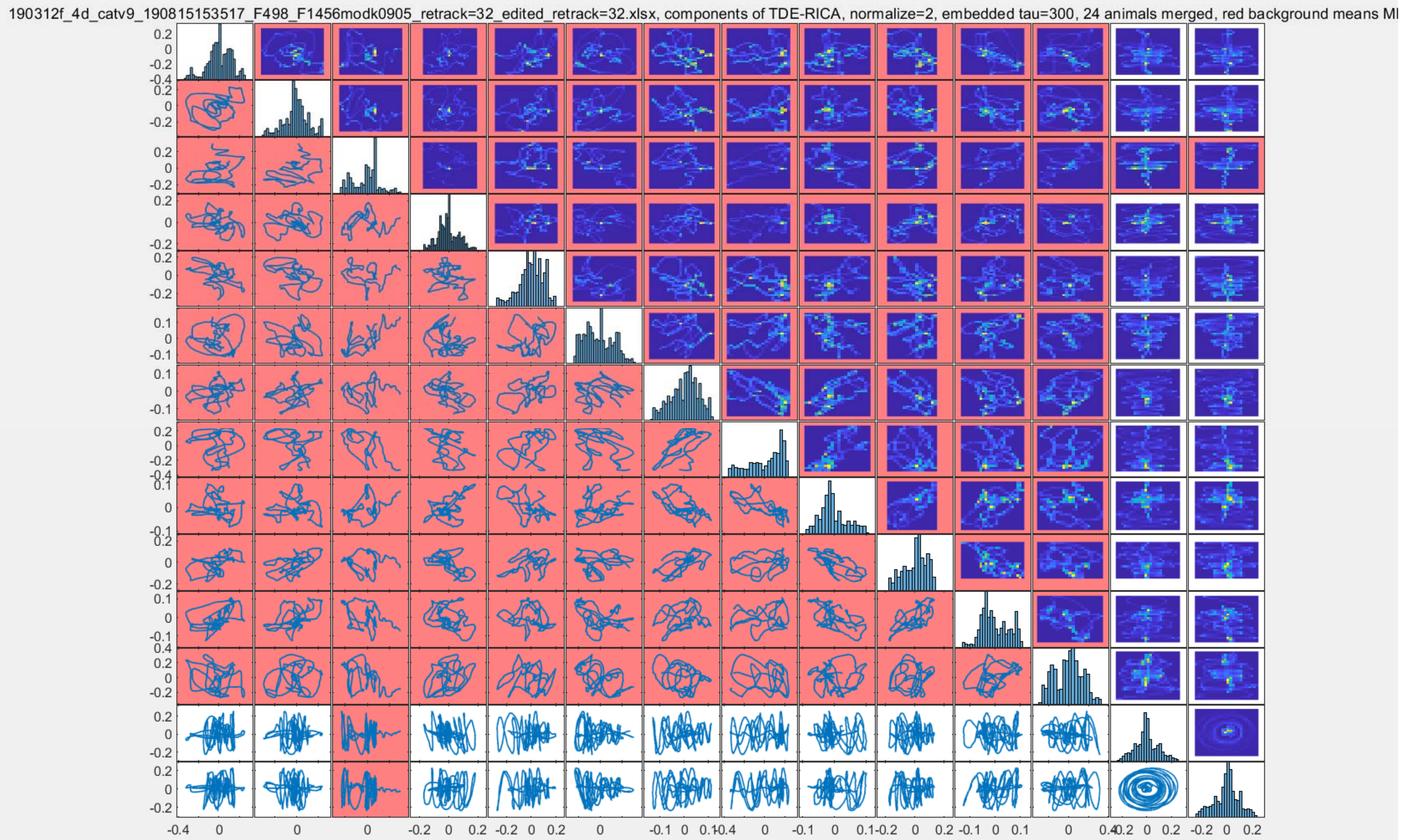
190312f_4d_catv9_190815153517_F498_F1456modk0905_retrack=32_edited_retrack=32.xlsx, components of TDE-RICA, normalize=2, embedded tau=300, 24 animals merged, red background means MI

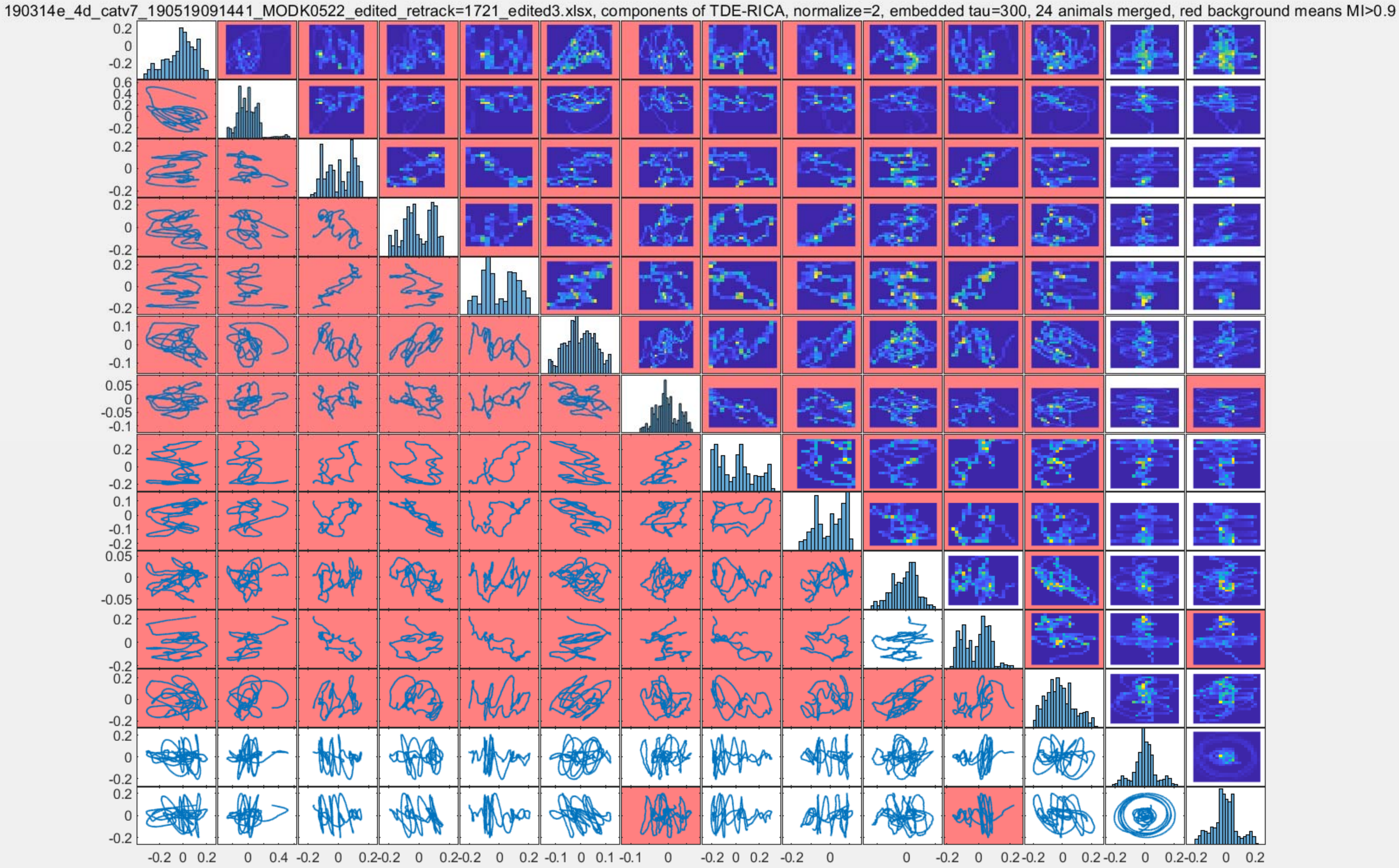
190314e_4d_catv7_190519091441_MODK0522_edited_retrack=1721_edited3.xlsx, components of TDE-RICA, normalize=2, embedded tau=300, 24 animals merged, red background means MI>0.9

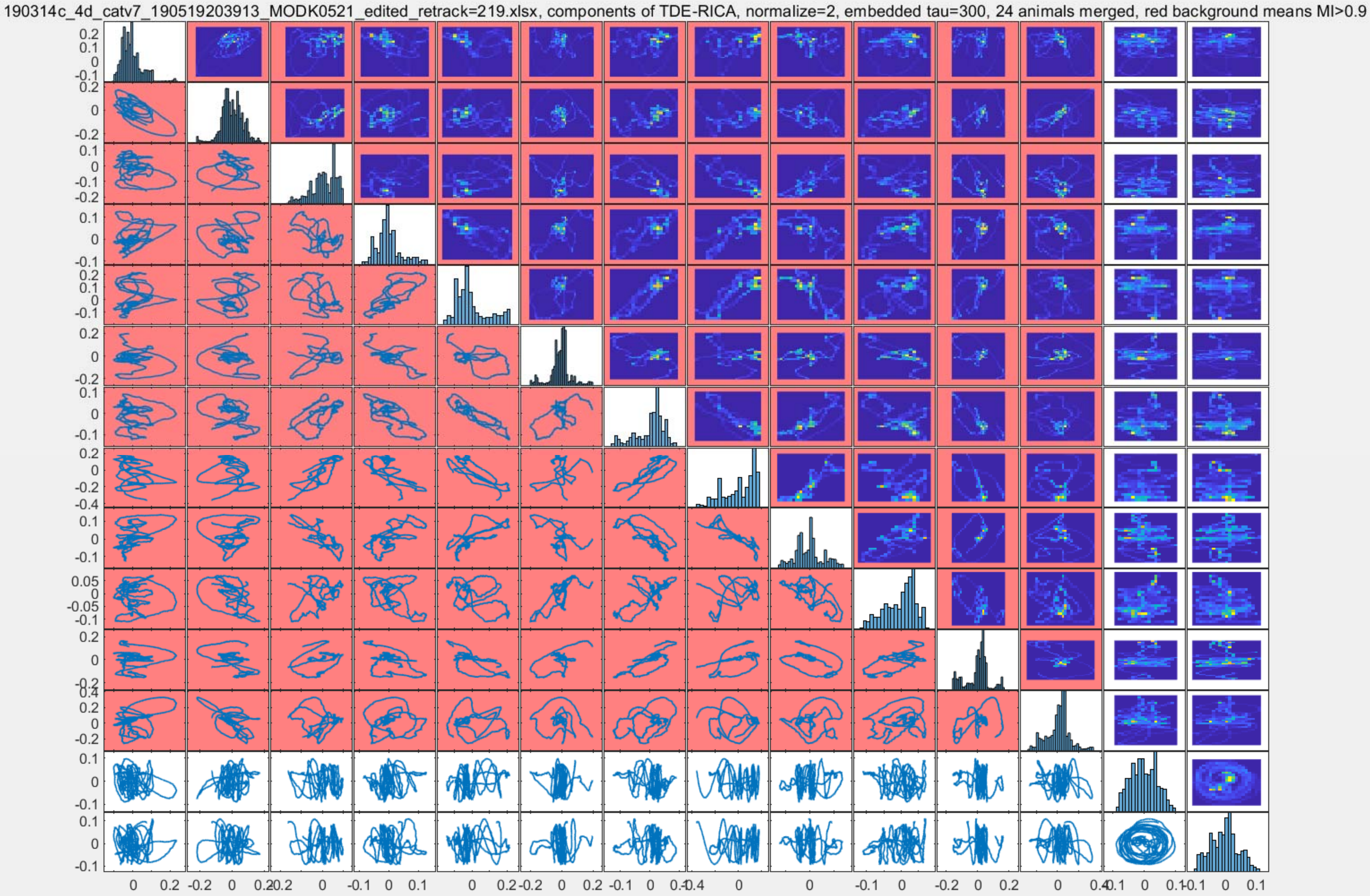
190314c_4d_catv7_190519203913_MODK0521_edited_retrack=219.xlsx, components of TDE-RICA, normalize=2, embedded tau=300, 24 animals merged, red background means MI>0.9

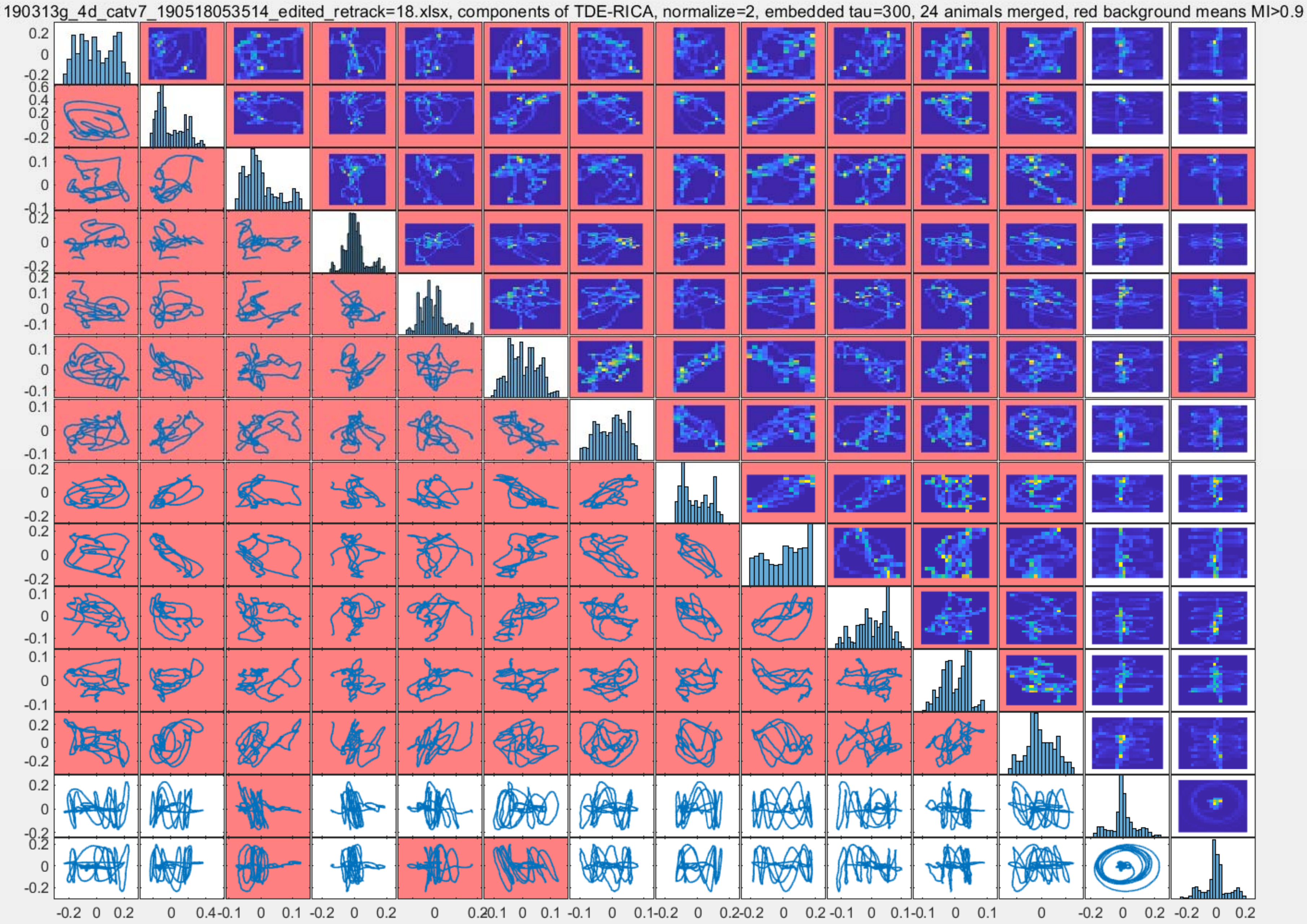
190313g_4d_catv7_190518053514_edited_retrack=18.xlsx, components of TDE-RICA, normalize=2, embedded tau=300, 24 animals merged, red background means MI>0.9

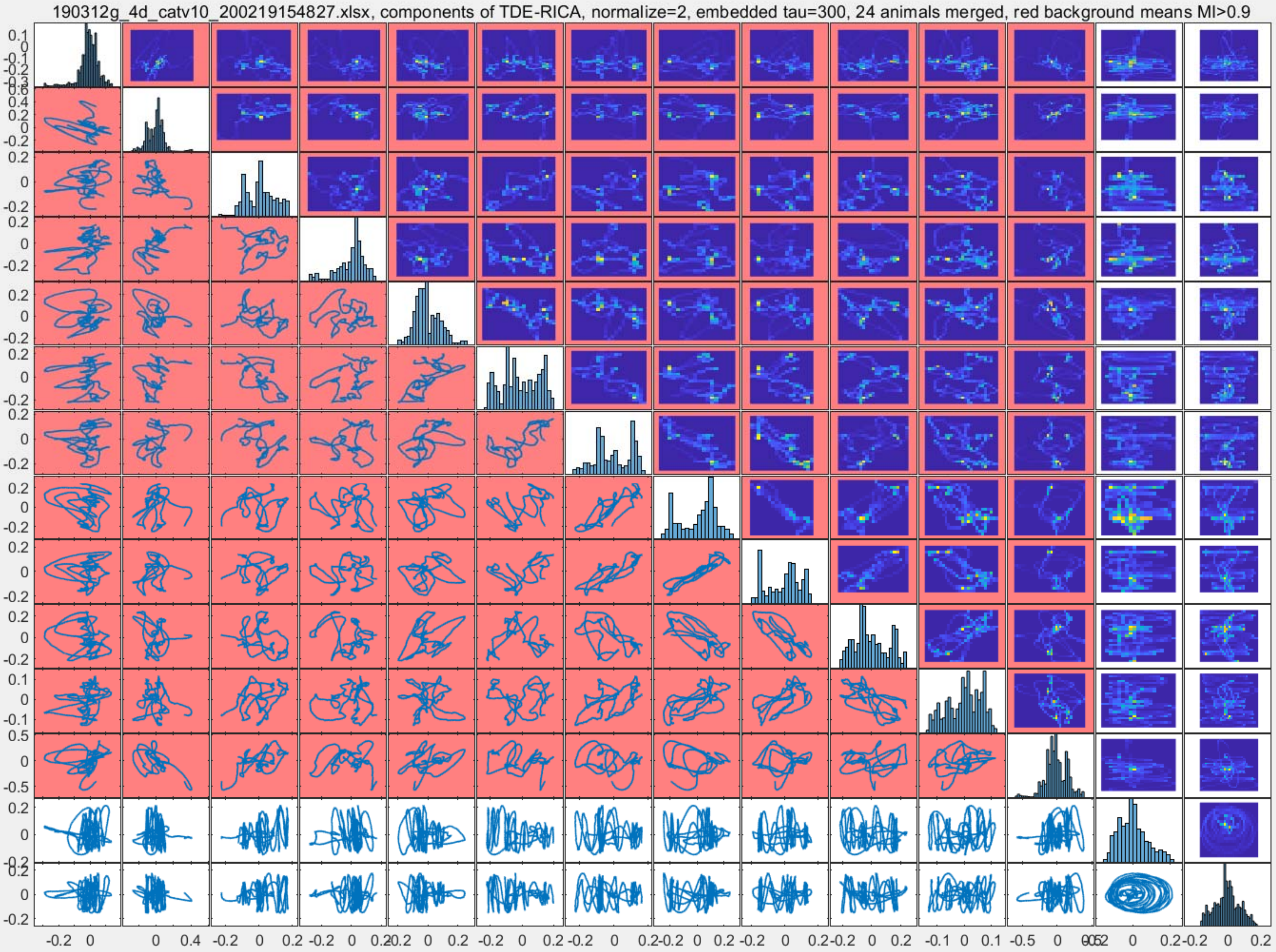

190312g_4d_catv10_200219154827.xlsx, components of TDE-RICA, normalize=2, embedded tau=300, 24 animals merged, red background means MI>0.9

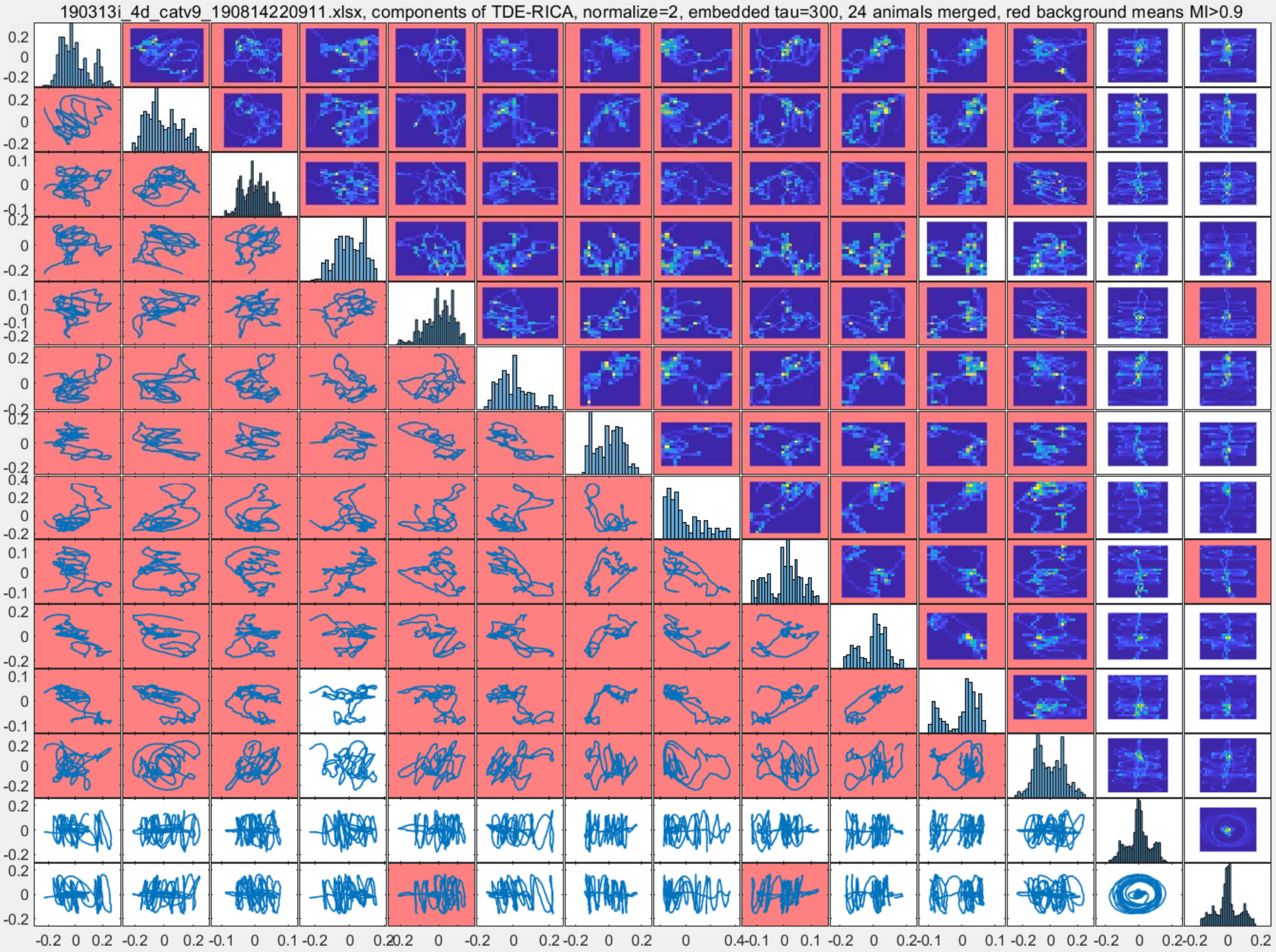
190313i_4d_catv9_190814220911.xlsx, components of TDE-RICA, normalize=2, embedded tau=300, 24 animals merged, red background means MI>0.9

190313a_4d_catv7_recopyano0618_modk_edited_retrack=471.xlsx, components of TDE-RICA, normalize=2, embedded tau=300, 24 animals merged, red background means MI>0.9

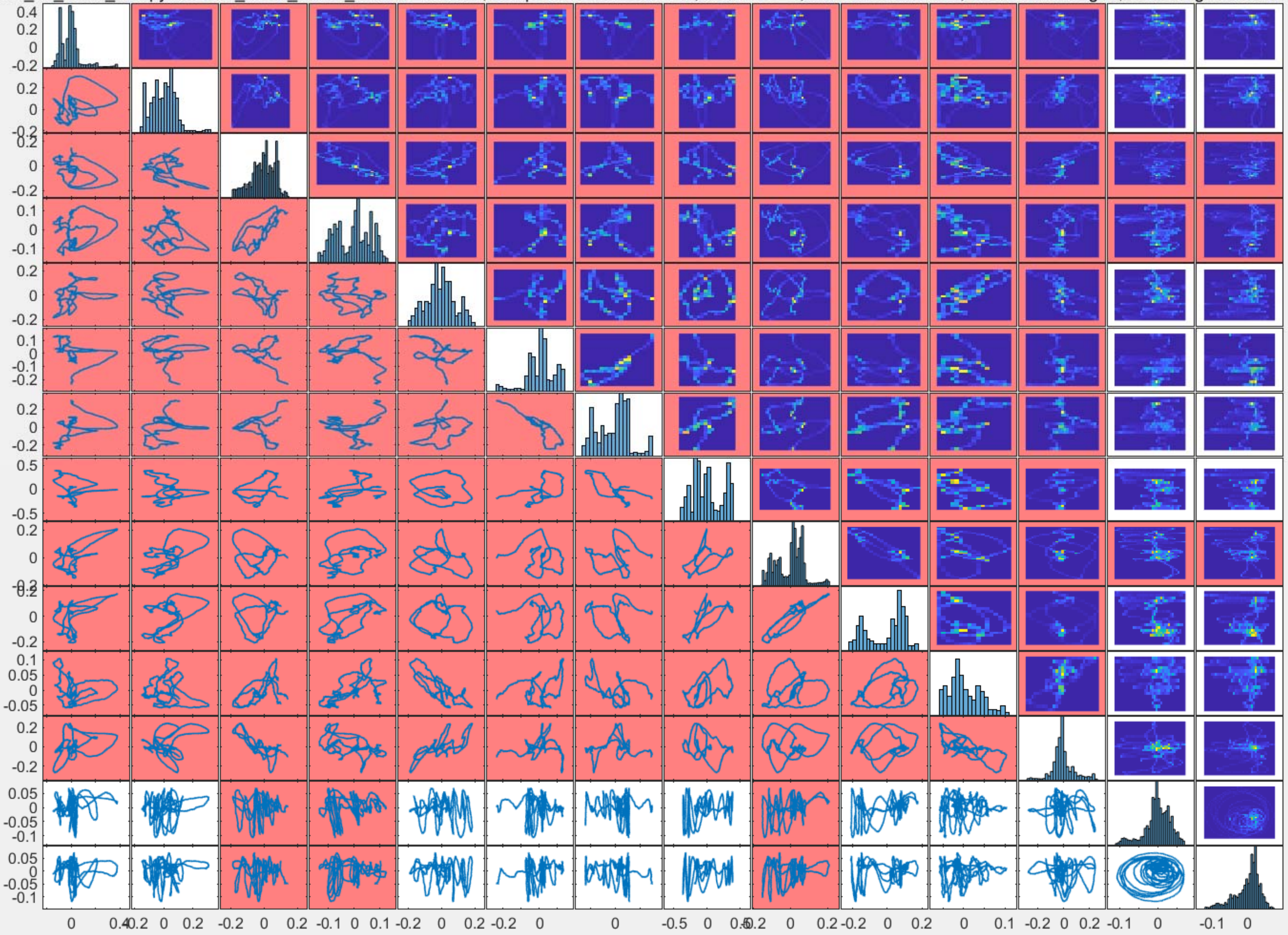

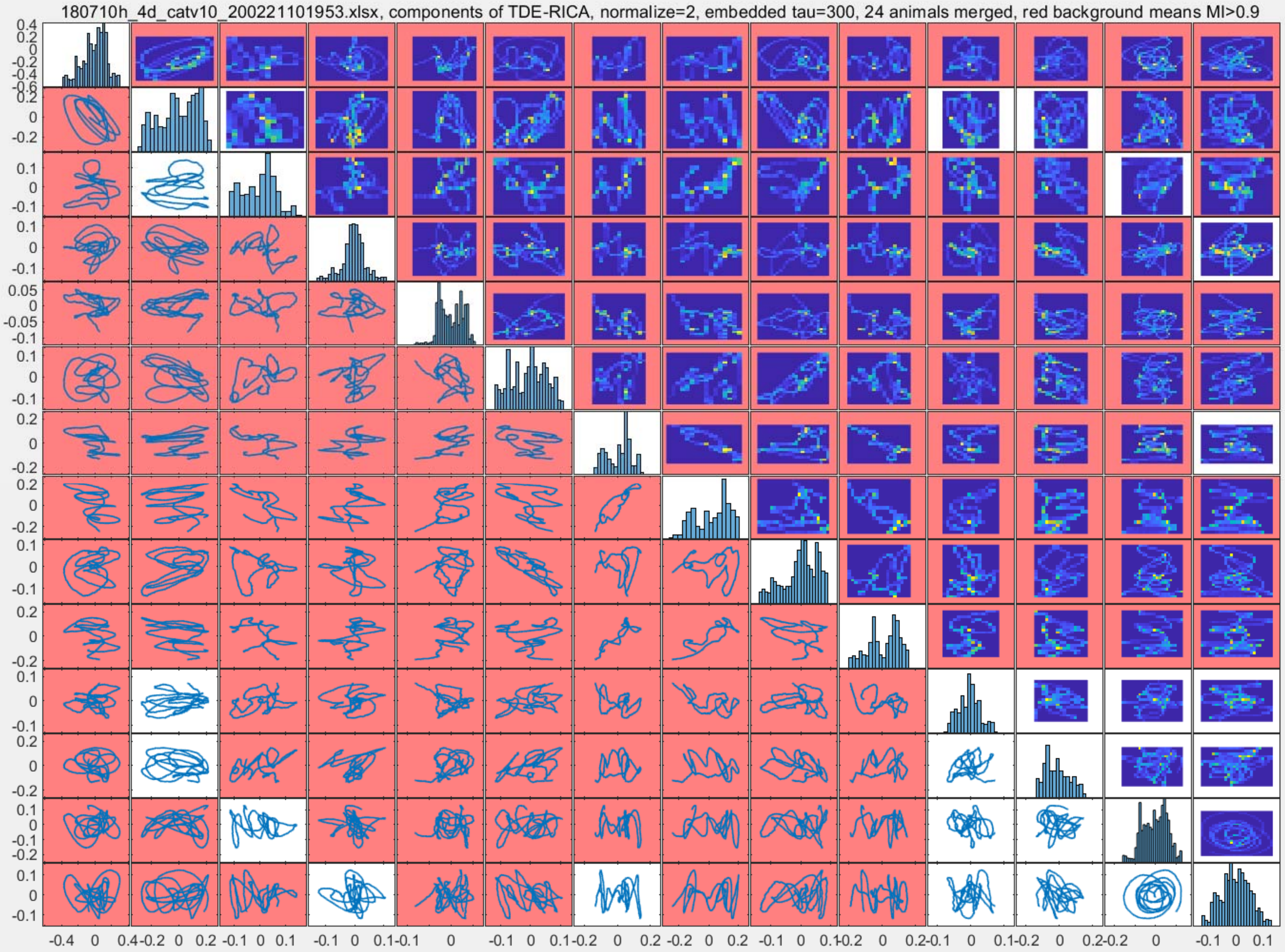

180710h_4d_catv10_200221101953.xlsx, components of TDE-RICA, normalize=2, embedded tau=300, 24 animals merged, red background means MI>0.9

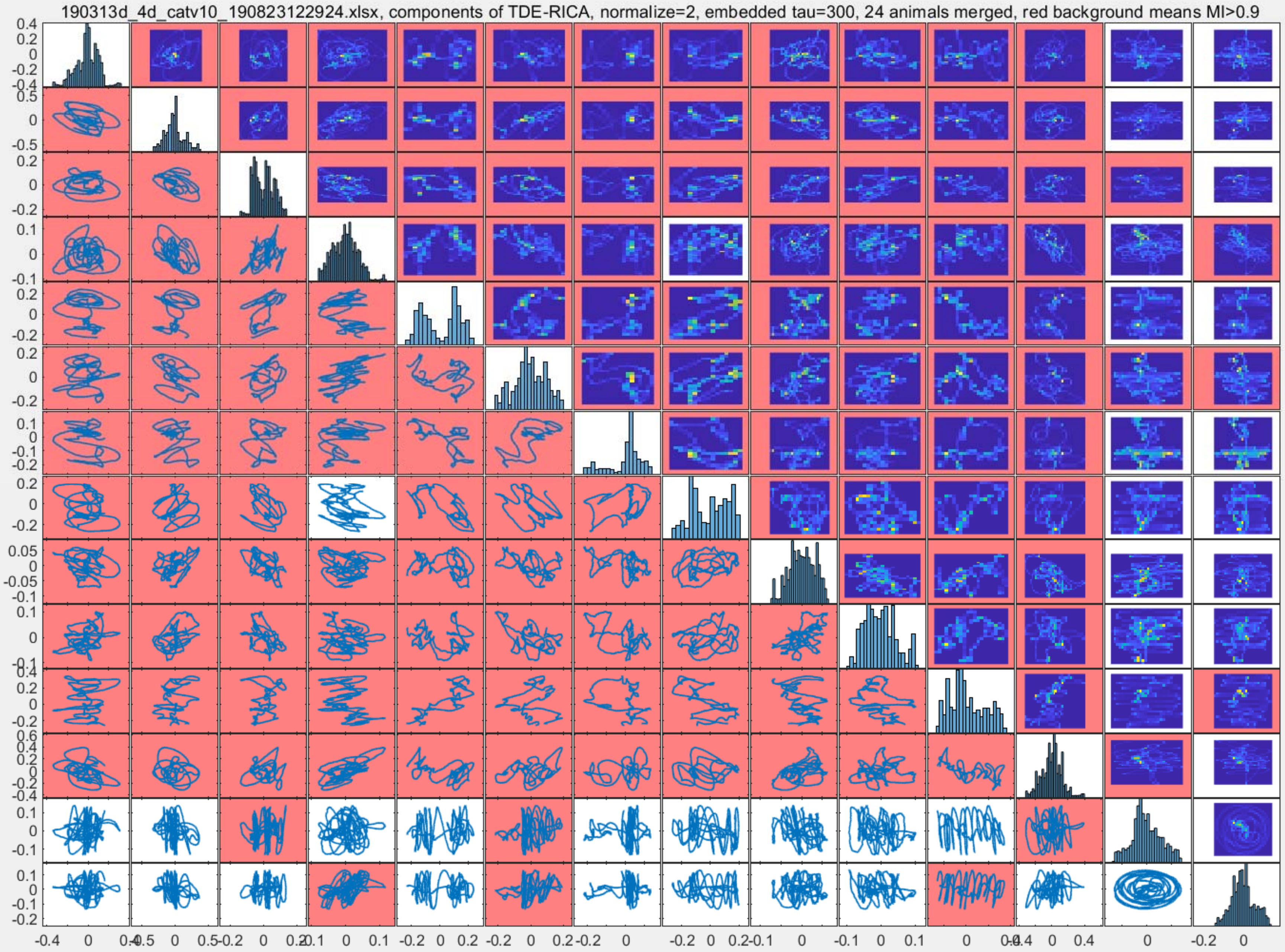
190313d_4d_catv10_190823122924.xlsx, components of TDE-RICA, normalize=2, embedded tau=300, 24 animals merged, red background means MI>0.9

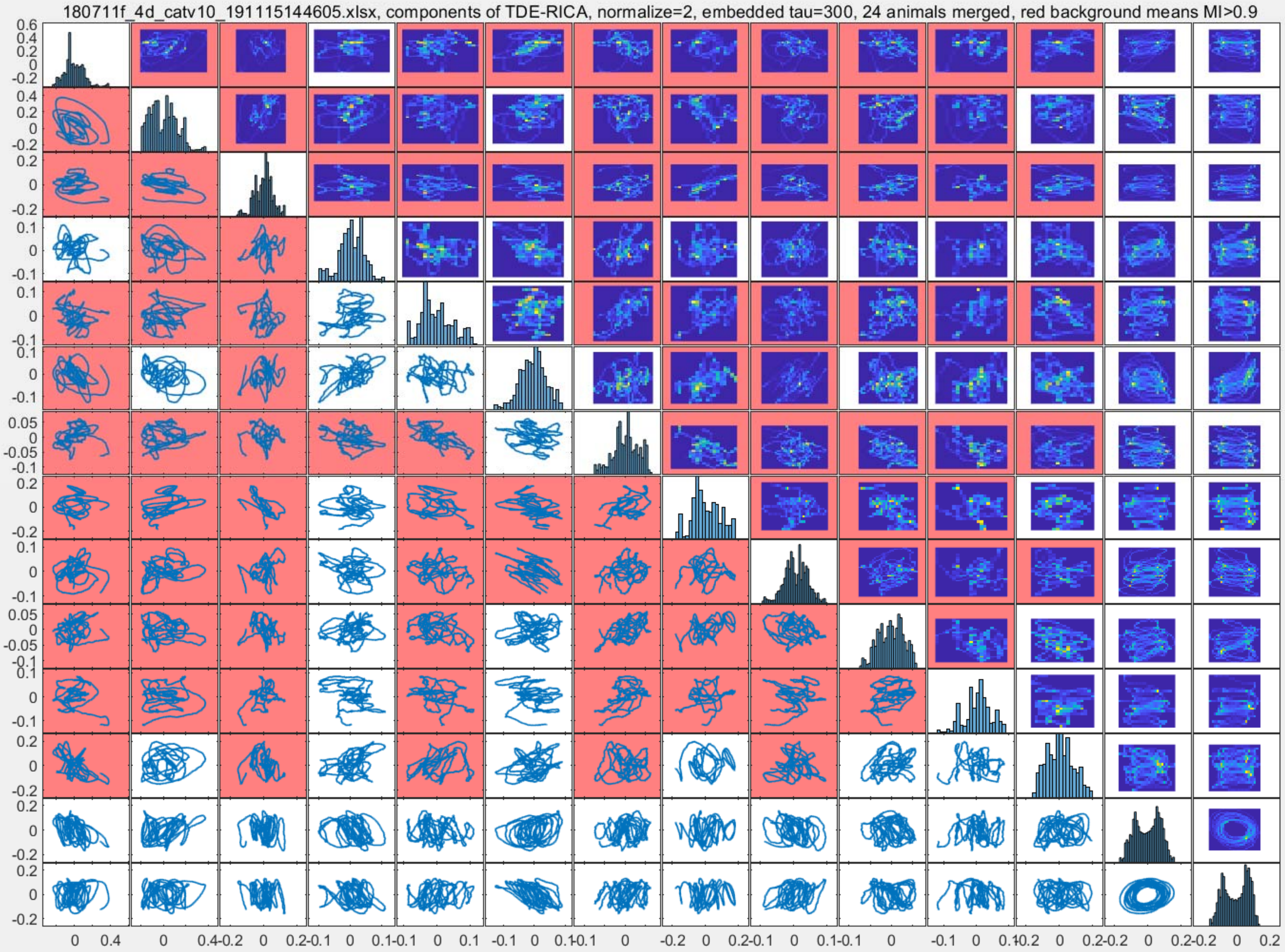

180711f_4d_catv10_191115144605.xlsx, components of TDE-RICA, normalize=2, embedded tau=300, 24 animals merged, red background means MI>0.9

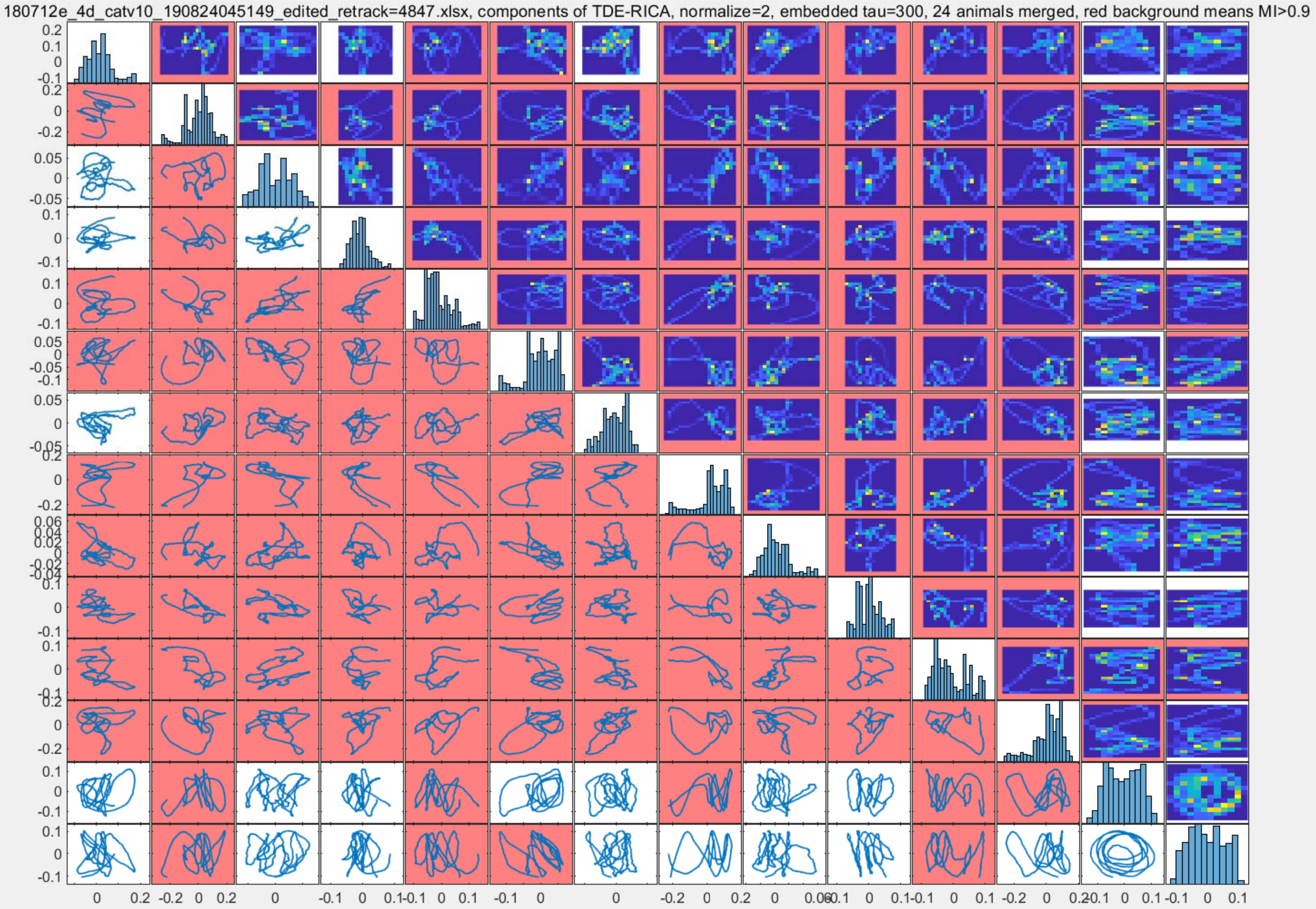

180712e_4d_catv10_190824045149_edited_retrack=4847.xlsx, components of TDE-RICA, normalize=2, embedded tau=300, 24 animals merged, red background means MI>0.9

190314d_4d_catv9_190815123845_edited_retrack=1618_edited_retrack=1618.xlsx, components of TDE-RICA, normalize=2, embedded tau=300, 24 animals merged, red background means MI>0.9

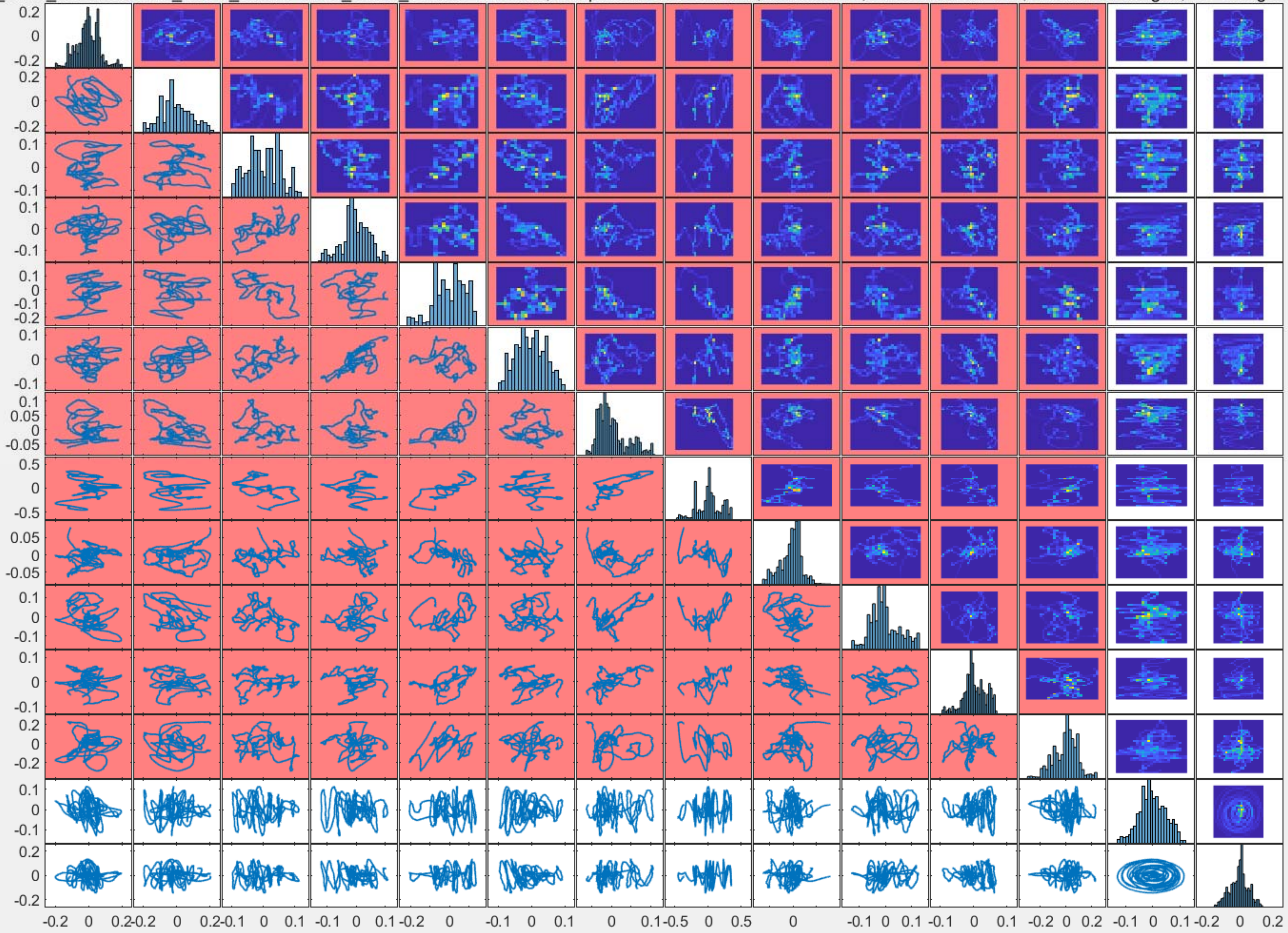

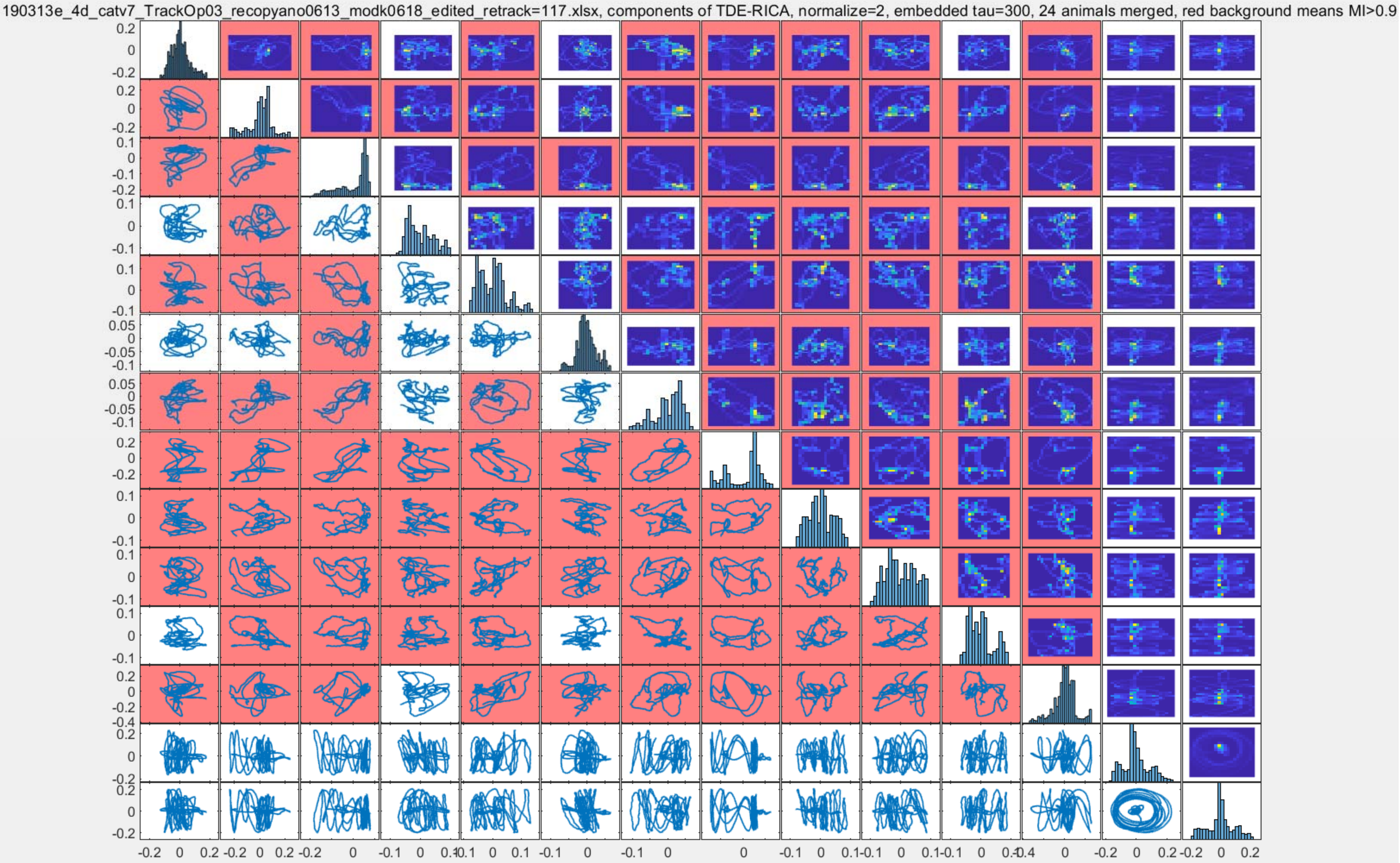
190313e_4d_catv7_TrackOp03_recopyano0613_modk0618_edited_retrack=117.xlsx, components of TDE-RICA, normalize=2, embedded tau=300, 24 animals merged, red background means MI>0.9

190312e_4d_catv7_190513212224_MODK_edited_retrack=50_edited.xlsx, components of TDE-RICA, normalize=2, embedded tau=300, 24 animals merged, red background means MI>0.9

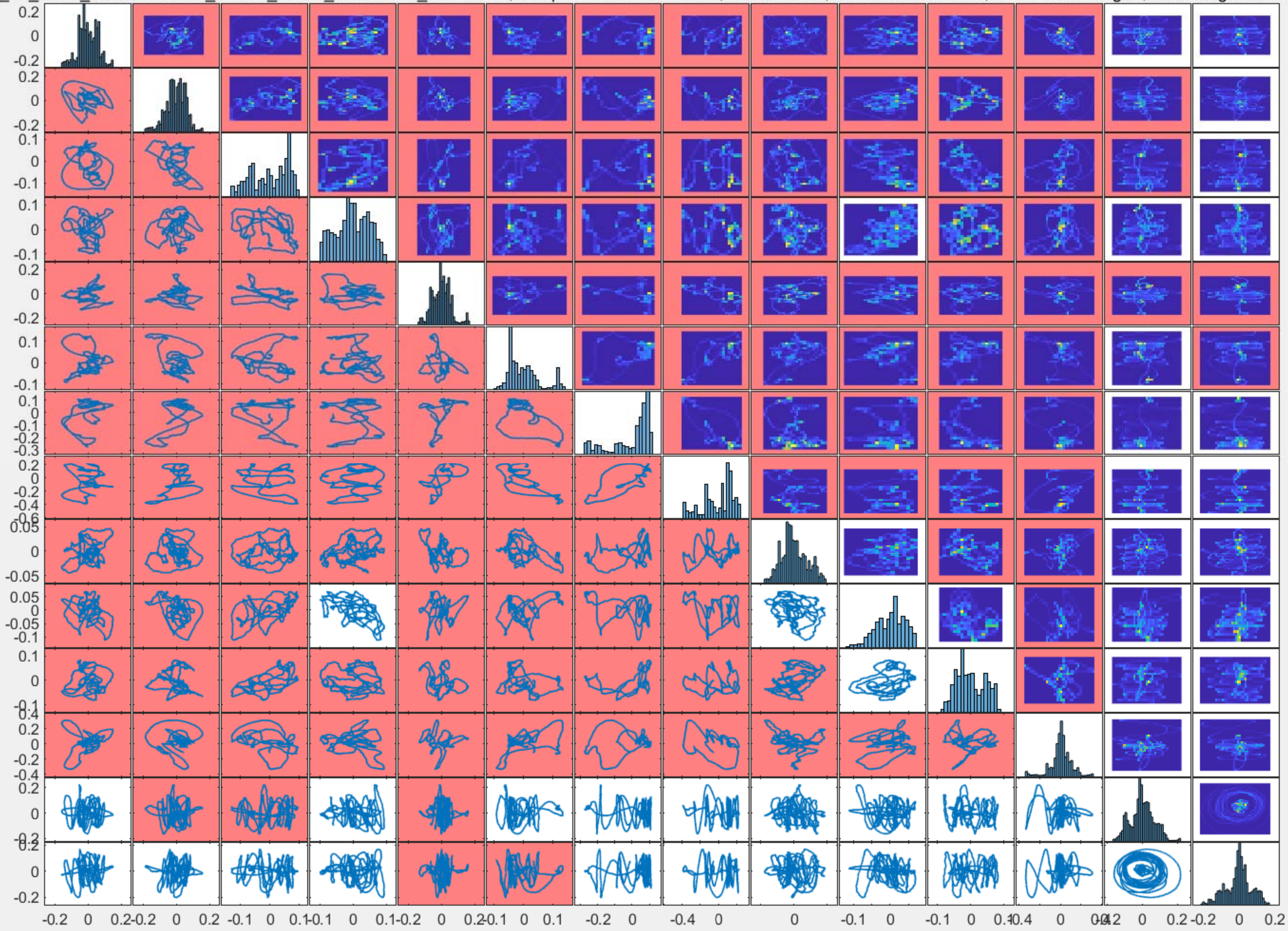

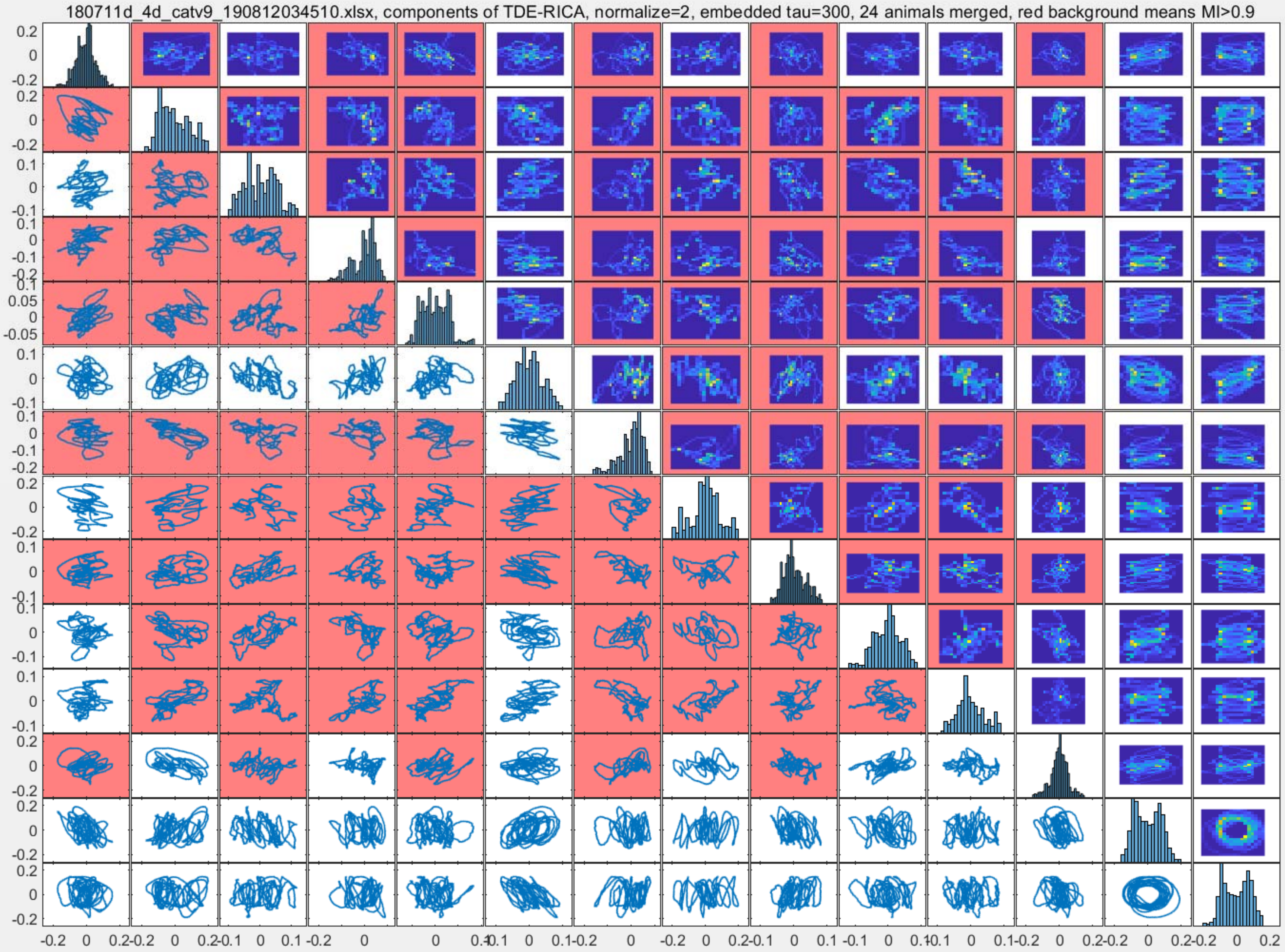

180711d_4d_catv9_190812034510.xlsx, components of TDE-RICA, normalize=2, embedded tau=300, 24 animals merged, red background means MI>0.9

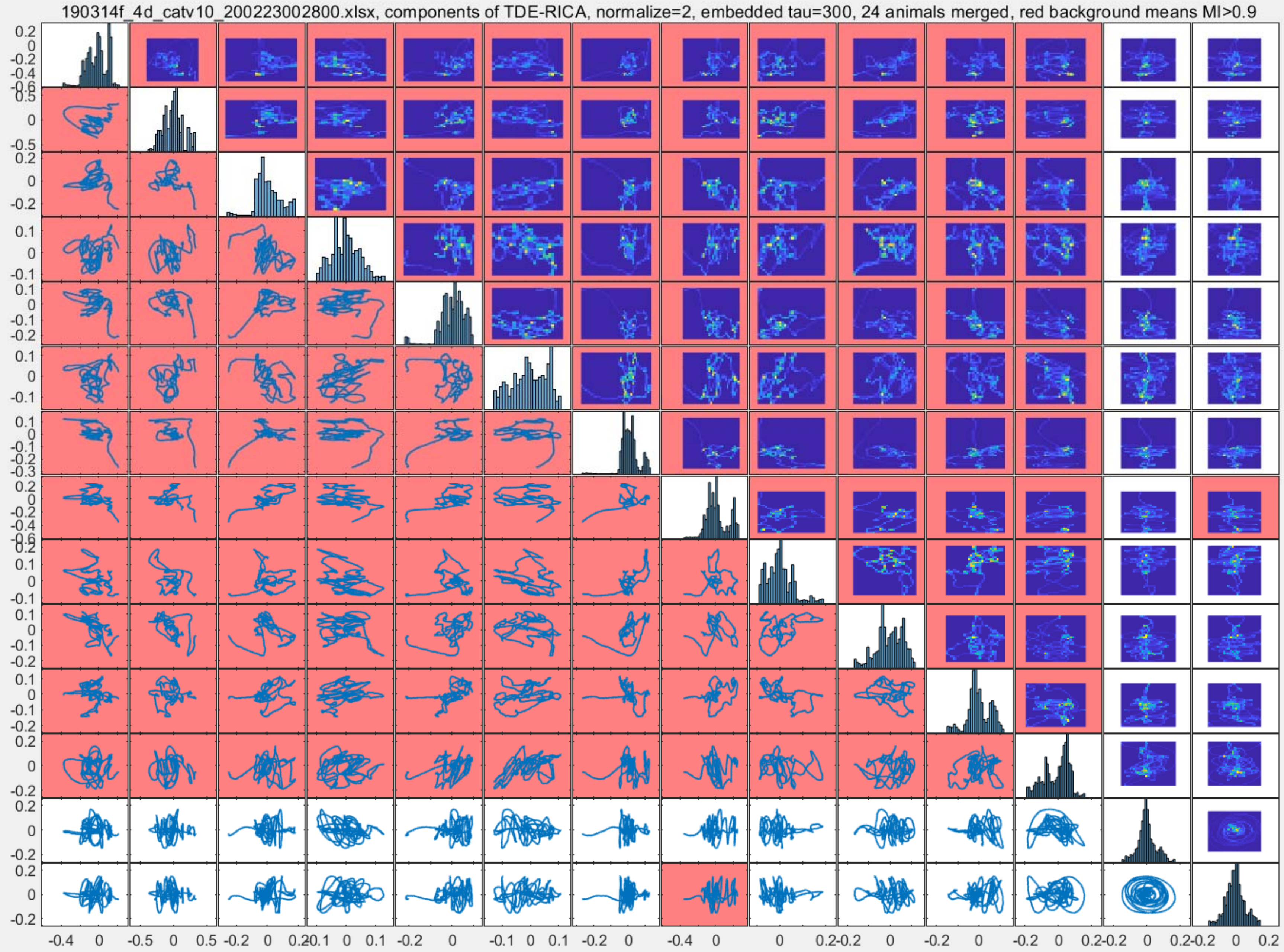

190314f_4d_catv10_200223002800.xlsx, components of TDE-RICA, normalize=2, embedded tau=300, 24 animals merged, red background means MI>0.9

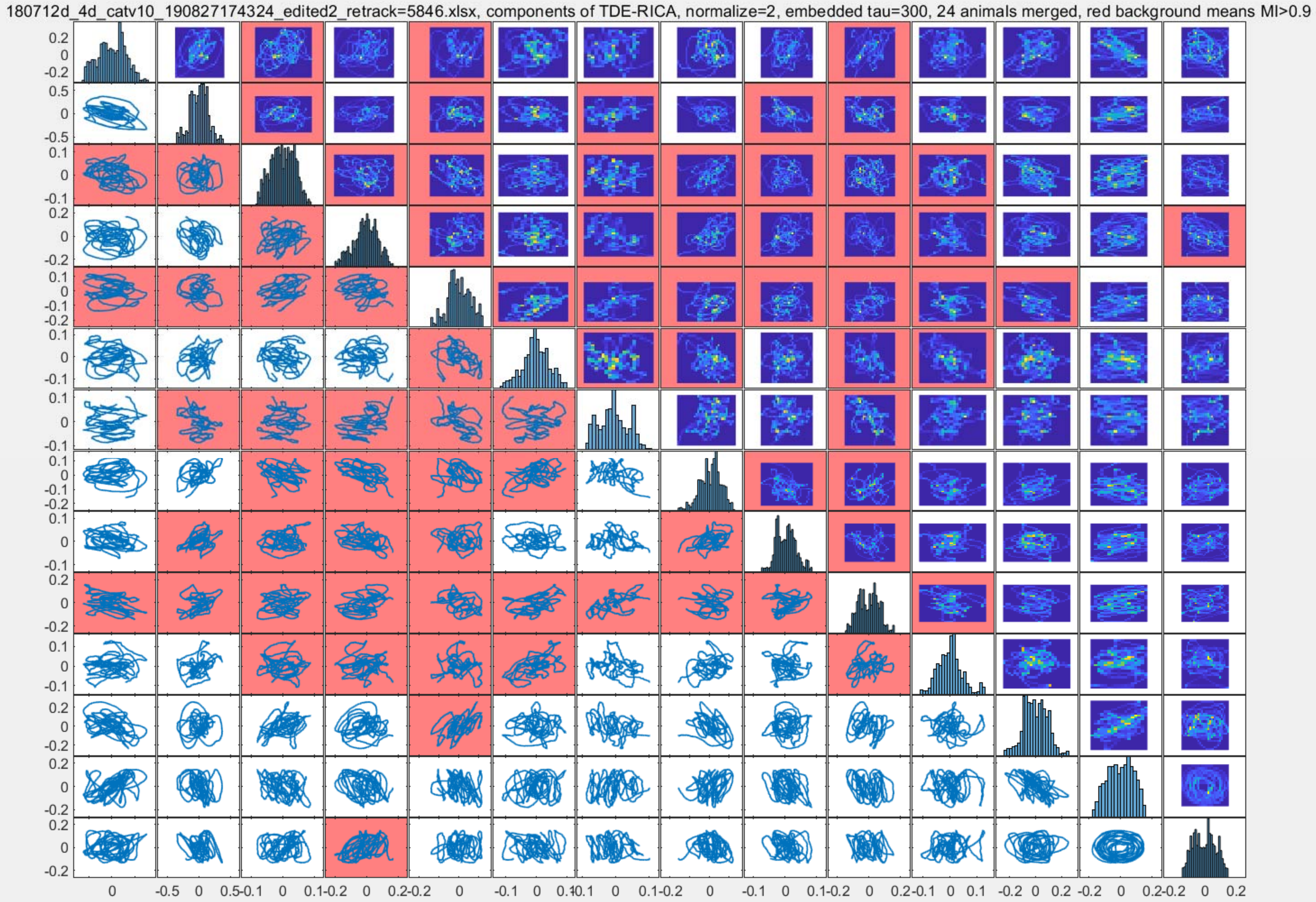
180712d_4d_catv10_190827174324_edited2_retrack=5846.xlsx, components of TDE-RICA, normalize=2, embedded tau=300, 24 animals merged, red background means MI>0.9

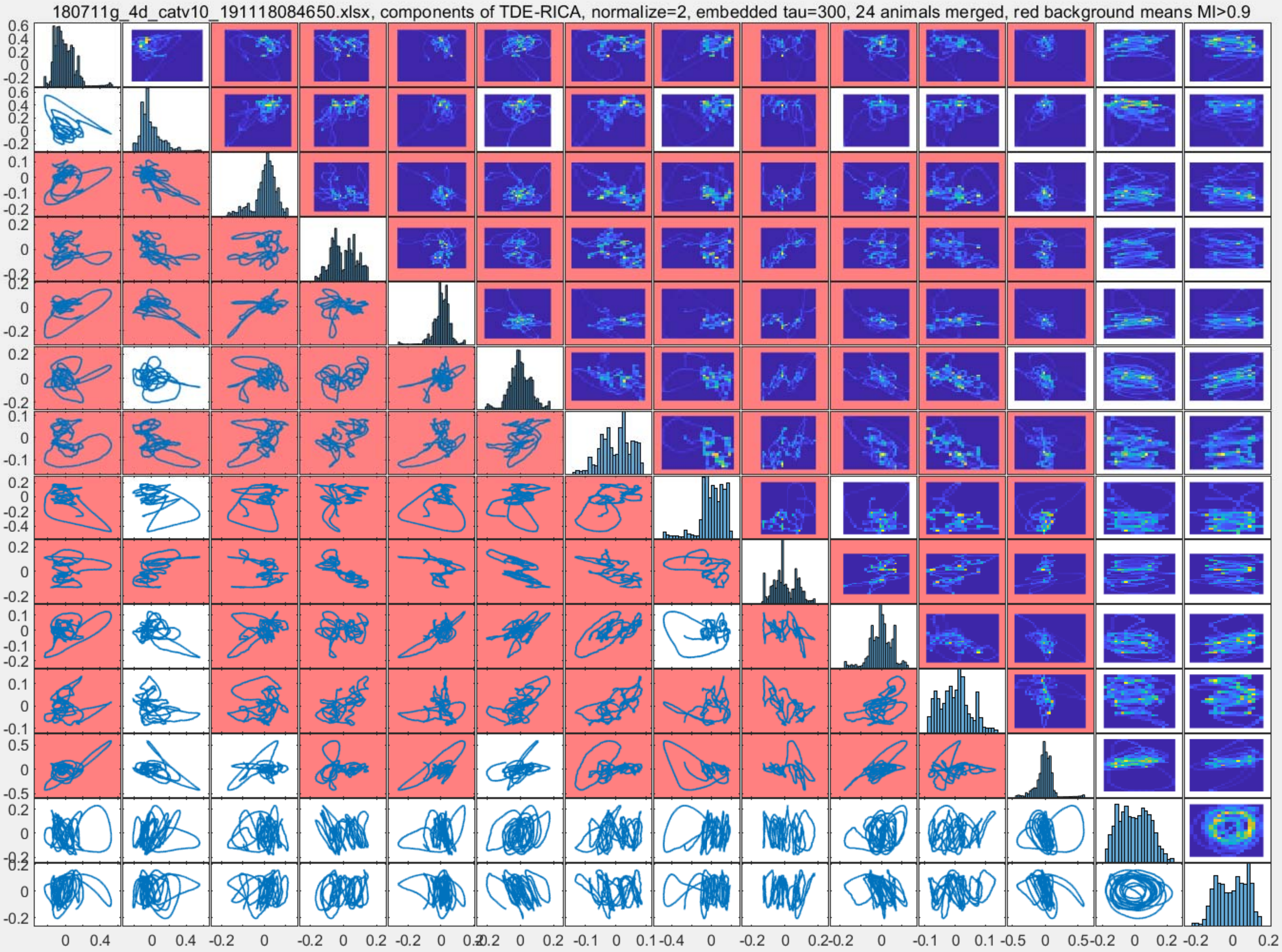

180711g_4d_catv10_191118084650.xlsx, components of TDE-RICA, normalize=2, embedded tau=300, 24 animals merged, red background means MI>0.9

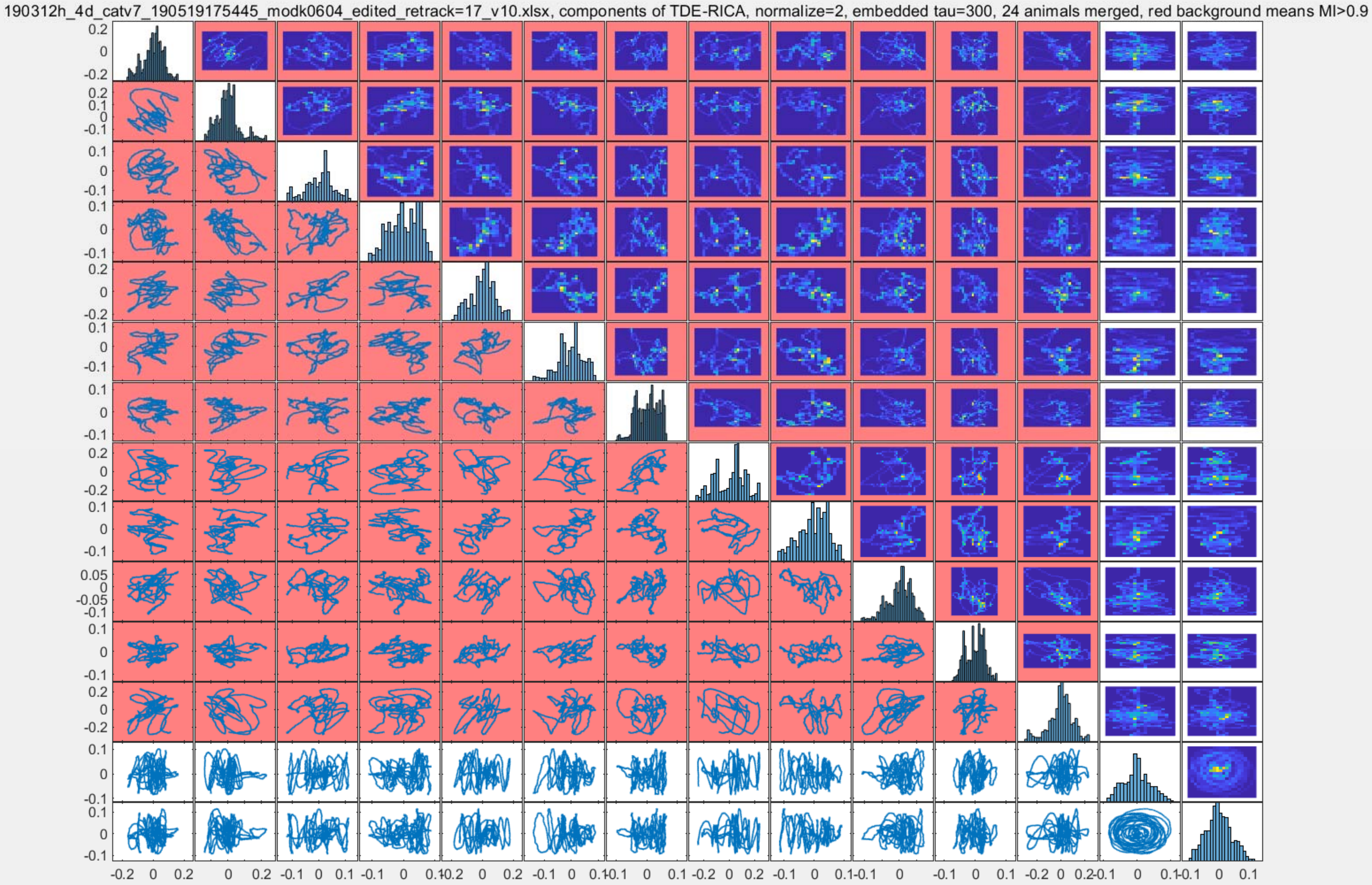
190312h_4d_catv7_190519175445_modk0604_edited_retrack=17_v10.xlsx, components of TDE-RICA, normalize=2, embedded tau=300, 24 animals merged, red background means MI>0.9

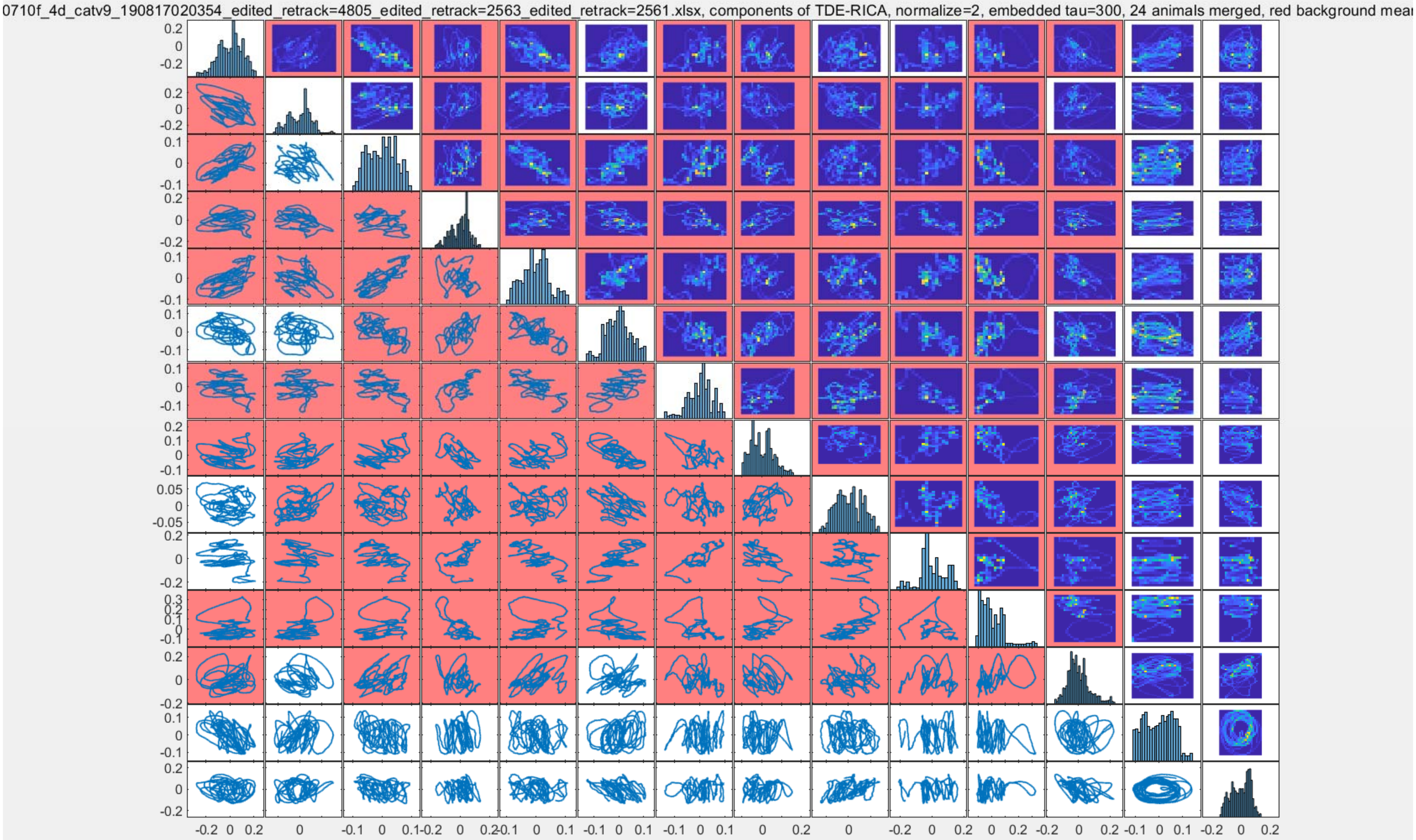
0710f_4d_catv9_190817020354_edited_retrack=4805_edited_retrack=2563_edited_retrack=2561.xlsx, components of TDE-RICA, normalize=2, embedded tau=300, 24 animals merged, red background mea

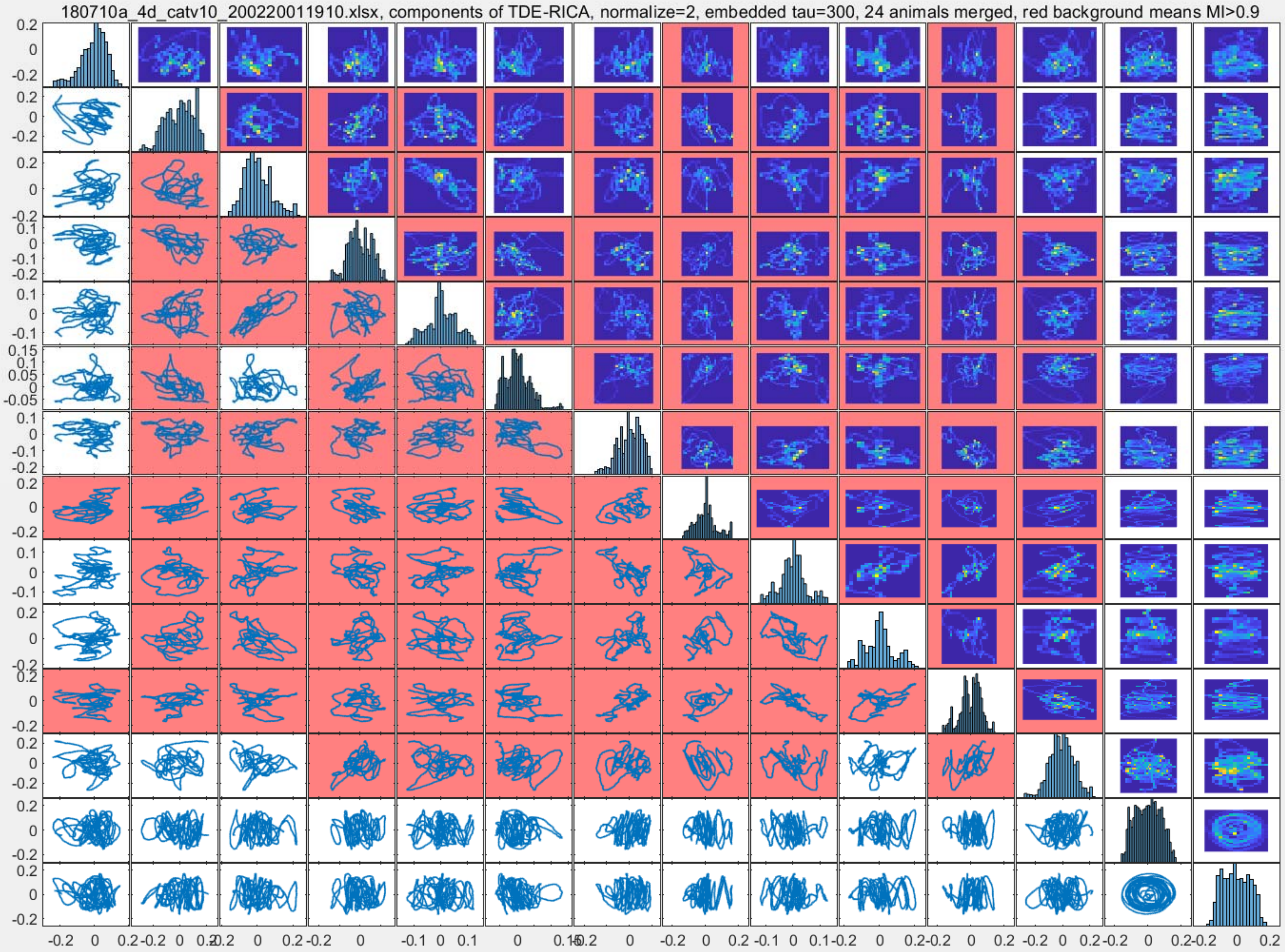
180710a_4d_catv10_200220011910.xlsx, components of TDE-RICA, normalize=2, embedded tau=300, 24 animals merged, red background means MI>0.9

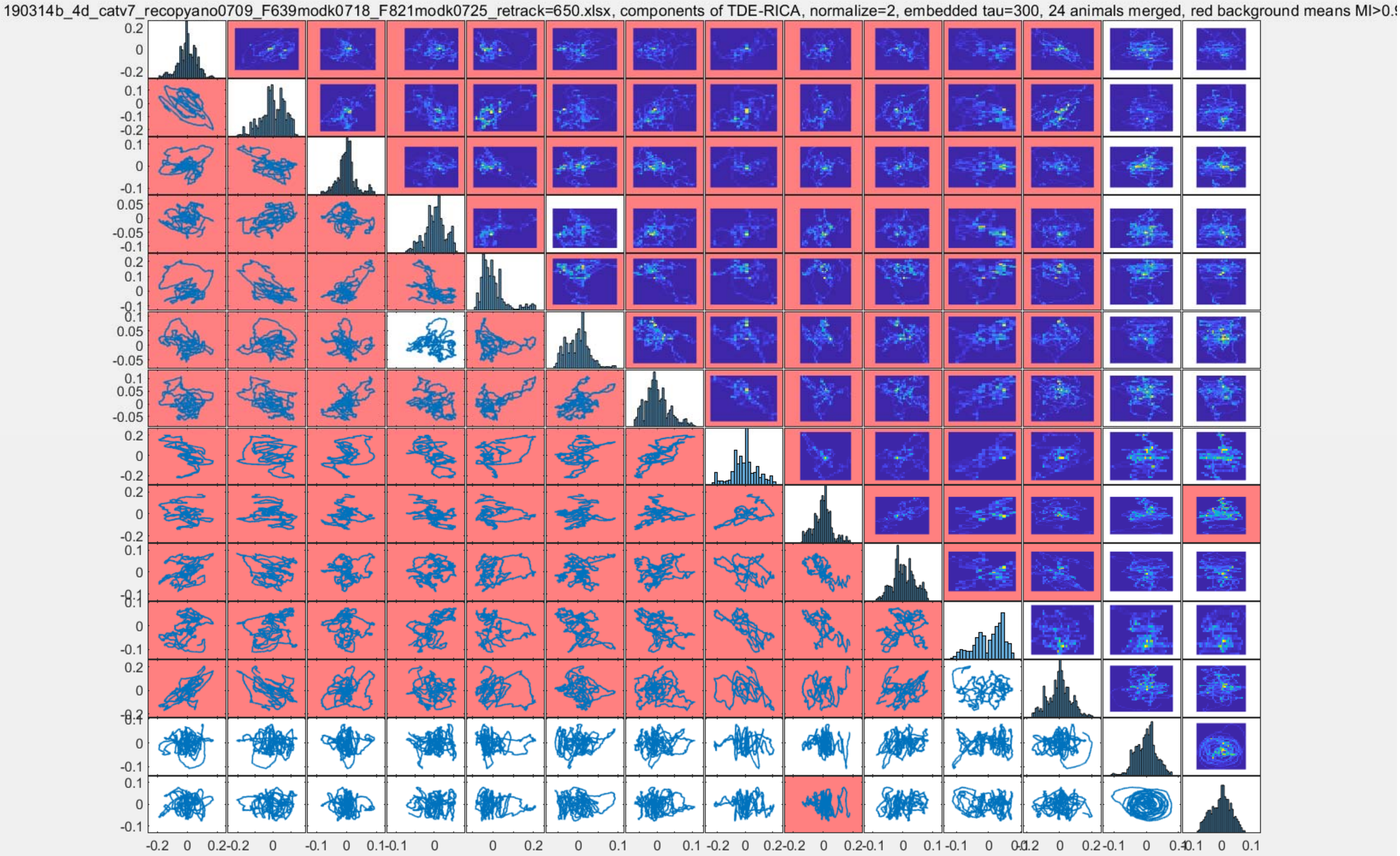
190314b_4d_catv7_recopyano0709_F639modk0718_F821modk0725_retrack=650.xlsx, components of TDE-RICA, normalize=2, embedded tau=300, 24 animals merged, red background means MI>0.

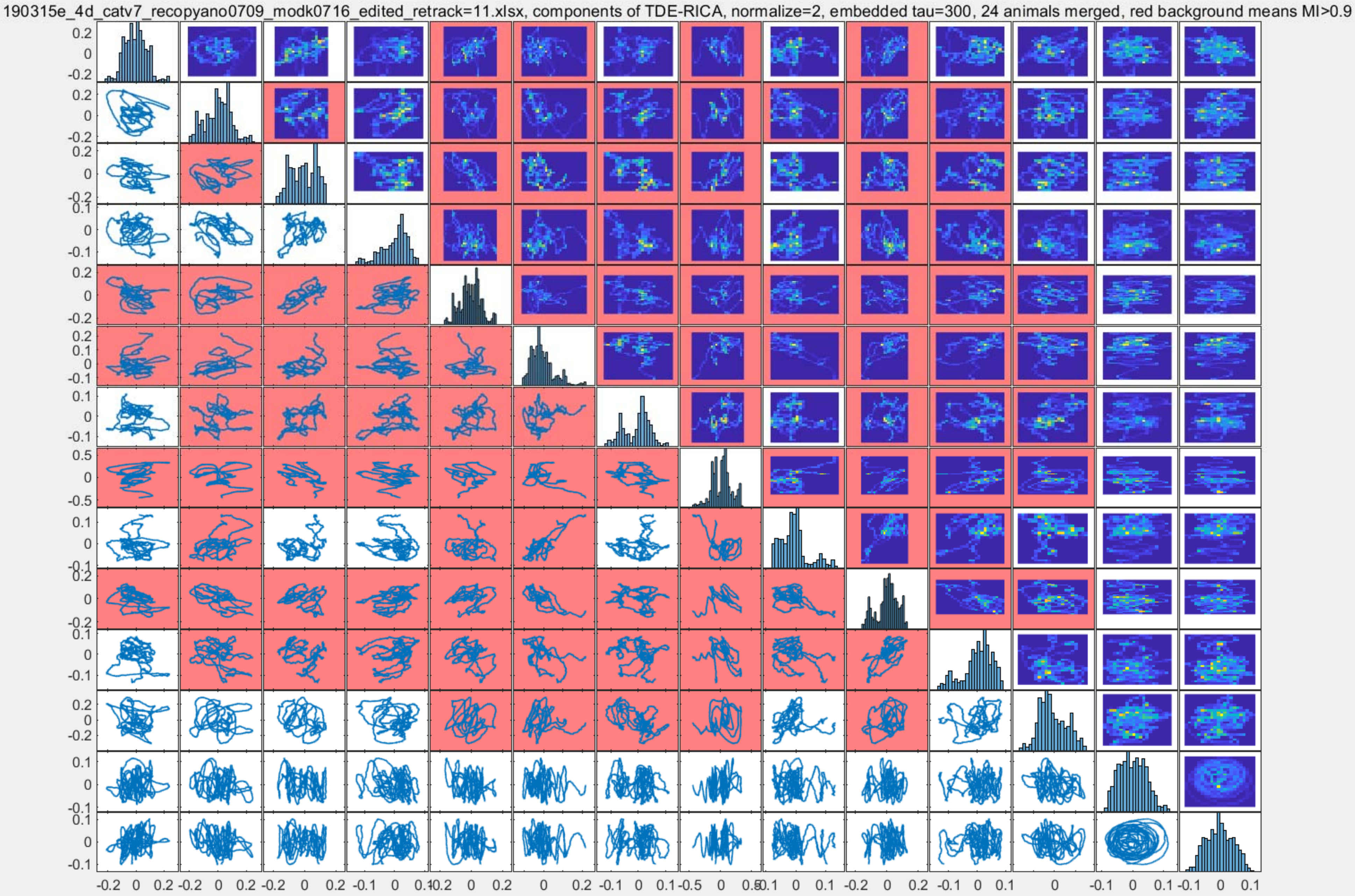
190315e_4d_catv7_recopyano0709_modk0716_edited_retrack=11.xlsx, components of TDE-RICA, normalize=2, embedded tau=300, 24 animals merged, red background means MI>0.9

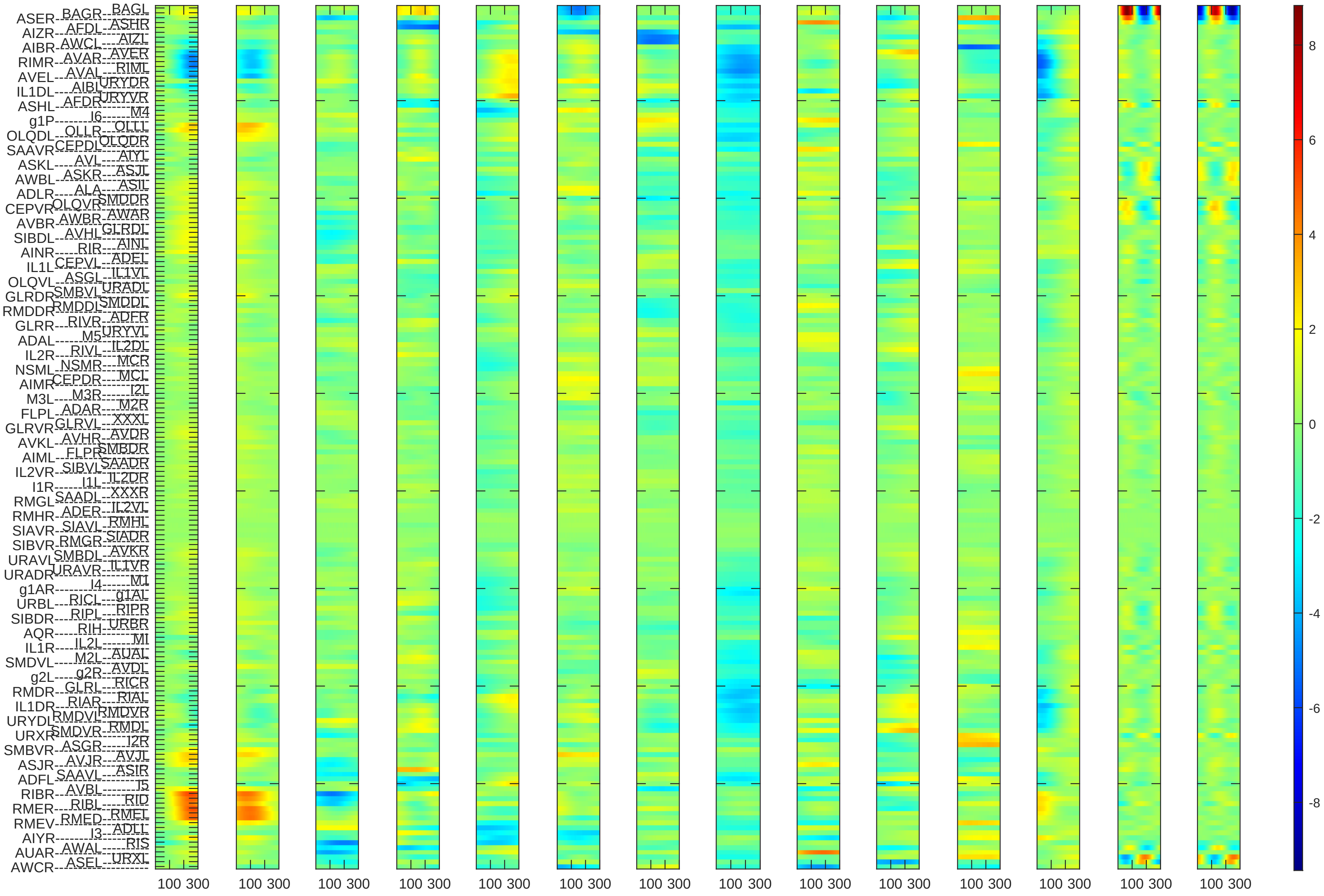

100 300
8
6
4
2
0
-2
-4
-6
-8

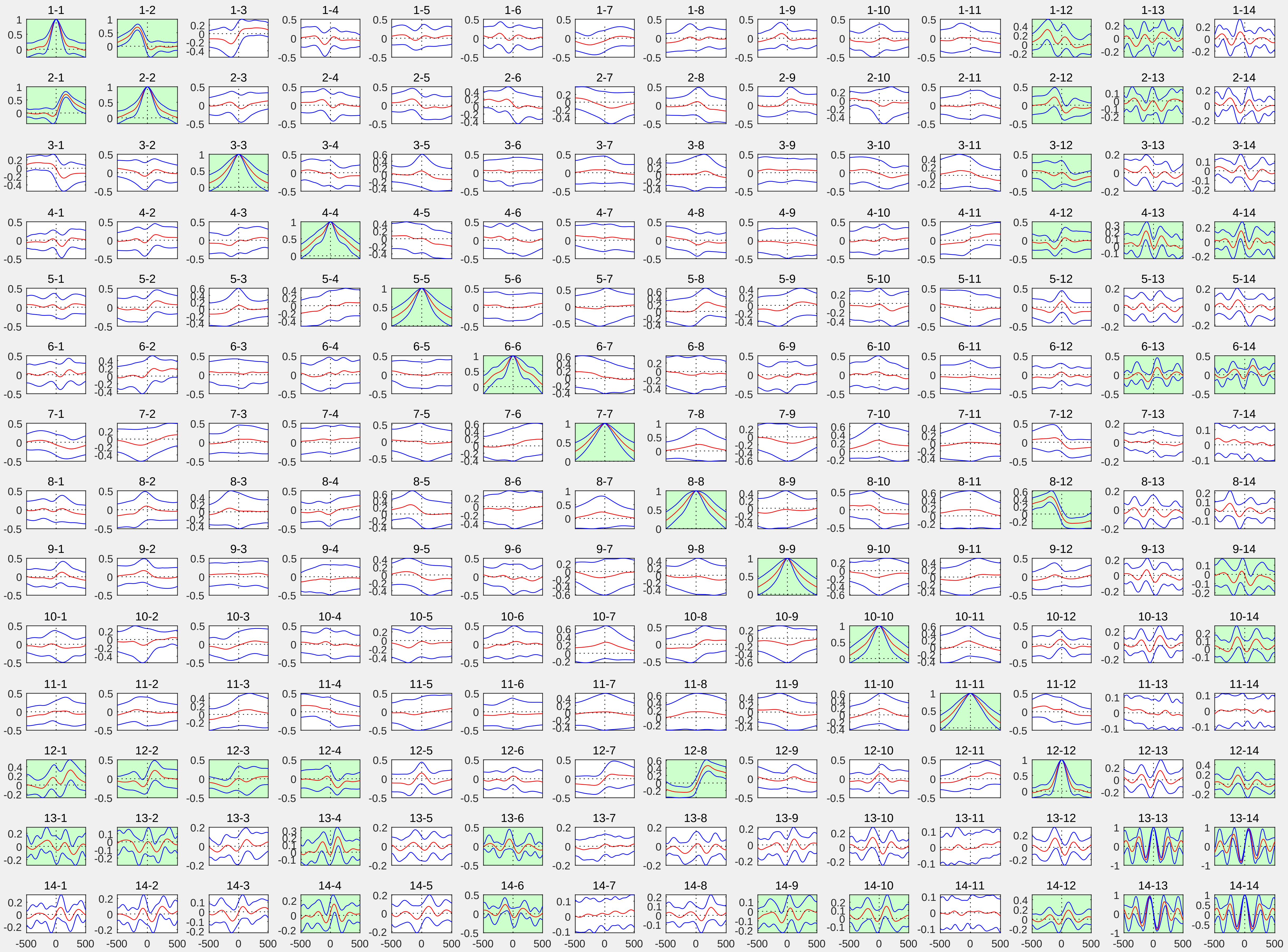

1-1 1-2 1-3 1-4 1-5 1-6 1-7 1-8 1-9 1-10 1-11 1-12 1-13 1-14
2-1 2-2 2-3 2-4 2-5 2-6 2-7 2-8 2-9 2-10 2-11 2-12 2-13 2-14
3-1 3-2 3-3 3-4 3-5 3-6 3-7 3-8 3-9 3-10 3-11 3-12 3-13 3-14
4-1 4-2 4-3 4-4 4-5 4-6 4-7 4-8 4-9 4-10 4-11 4-12 4-13 4-14
5-1 5-2 5-3 5-4 5-5 5-6 5-7 5-8 5-9 5-10 5-11 5-12 5-13 5-14
6-1 6-2 6-3 6-4 6-5 6-6 6-7 6-8 6-9 6-10 6-11 6-12 6-13 6-14
7-1 7-2 7-3 7-4 7-5 7-6 7-7 7-8 7-9 7-10 7-11 7-12 7-13 7-14
8-1 8-2 8-3 8-4 8-5 8-6 8-7 8-8 8-9 8-10 8-11 8-12 8-13 8-14
9-1 9-2 9-3 9-4 9-5 9-6 9-7 9-8 9-9 9-10 9-11 9-12 9-13 9-14
10-1 10-2 10-3 10-4 10-5 10-6 10-7 10-8 10-9 10-10 10-11 10-12 10-13 10-14
11-1 11-2 11-3 11-4 11-5 11-6 11-7 11-8 11-9 11-10 11-11 11-12 11-13 11-14
12-1 12-2 12-3 12-4 12-5 12-6 12-7 12-8 12-9 12-10 12-11 12-12 12-13 12-14
13-1 13-2 13-3 13-4 13-5 13-6 13-7 13-8 13-9 13-10 13-11 13-12 13-13 13-14
14-1 14-2 14-3 14-4 14-5 14-6 14-7 14-8 14-9 14-10 14-11 14-12 14-13 14-14
-500 0 500

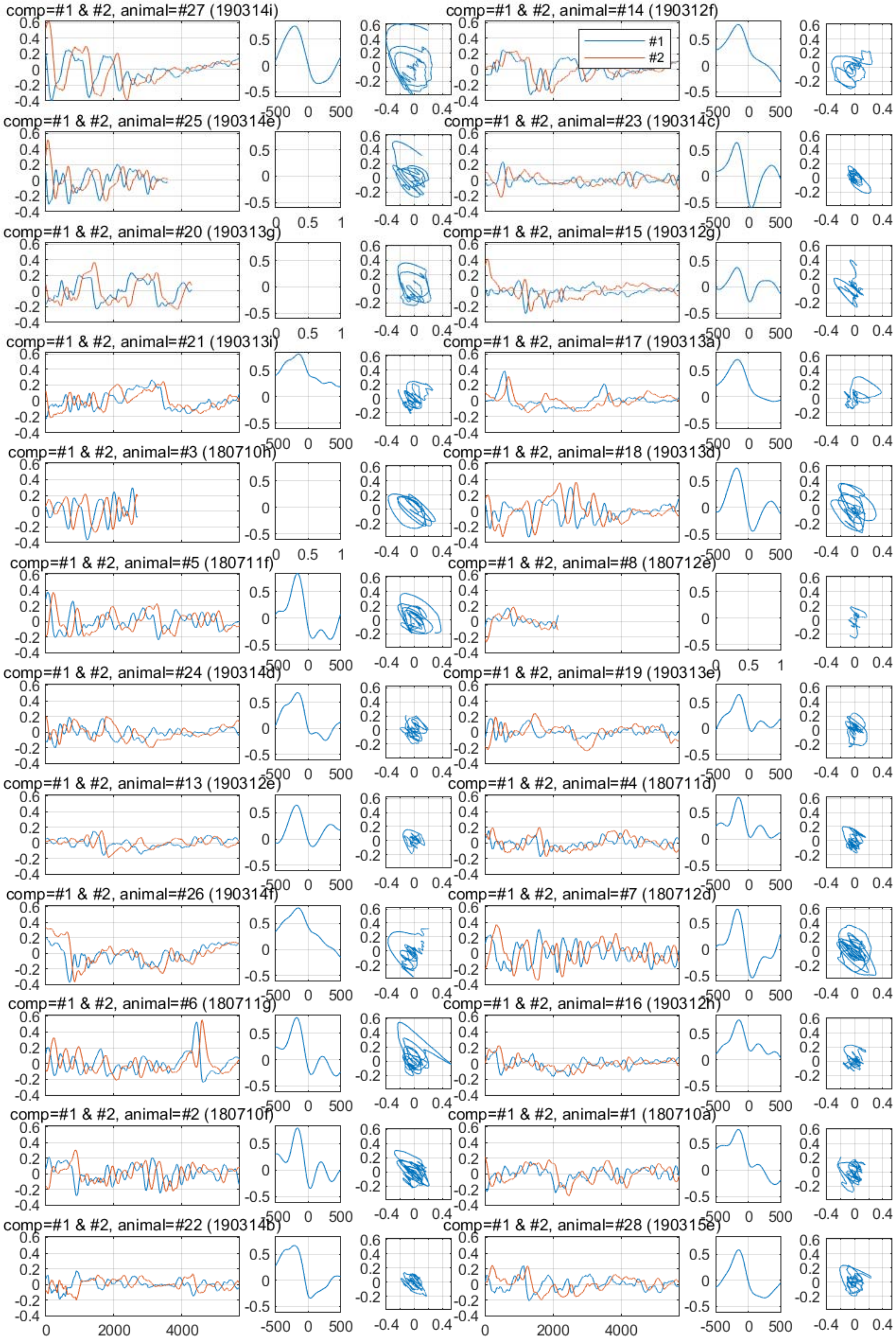

comp=#1 & #2, animal=#27 (190314i)
comp=#1 & #2, animal=#14 (190312f)
#1
#2
comp=#1 & #2, animal=#25 (190314e)
comp=#1 & #2, animal=#23 (190314c)
comp=#1 & #2, animal=#20 (190313g)
comp=#1 & #2, animal=#15 (190312g)
comp=#1 & #2, animal=#21 (190313i)
comp=#1 & #2, animal=#17 (190313a)
comp=#1 & #2, animal=#3 (180710h)
comp=#1 & #2, animal=#18 (190313d)
comp=#1 & #2, animal=#5 (180711f)
comp=#1 & #2, animal=#8 (180712e)
comp=#1 & #2, animal=#24 (190314d)
comp=#1 & #2, animal=#19 (190313e)
comp=#1 & #2, animal=#13 (190312e)
comp=#1 & #2, animal=#4 (180711d)
comp=#1 & #2, animal=#26 (190314f)
comp=#1 & #2, animal=#7 (180712d)
comp=#1 & #2, animal=#6 (180711g)
comp=#1 & #2, animal=#16 (190312h)
comp=#1 & #2, animal=#2 (180710f)
comp=#1 & #2, animal=#1 (180710a)
comp=#1 & #2, animal=#22 (190314b)
comp=#1 & #2, animal=#28 (190315e)

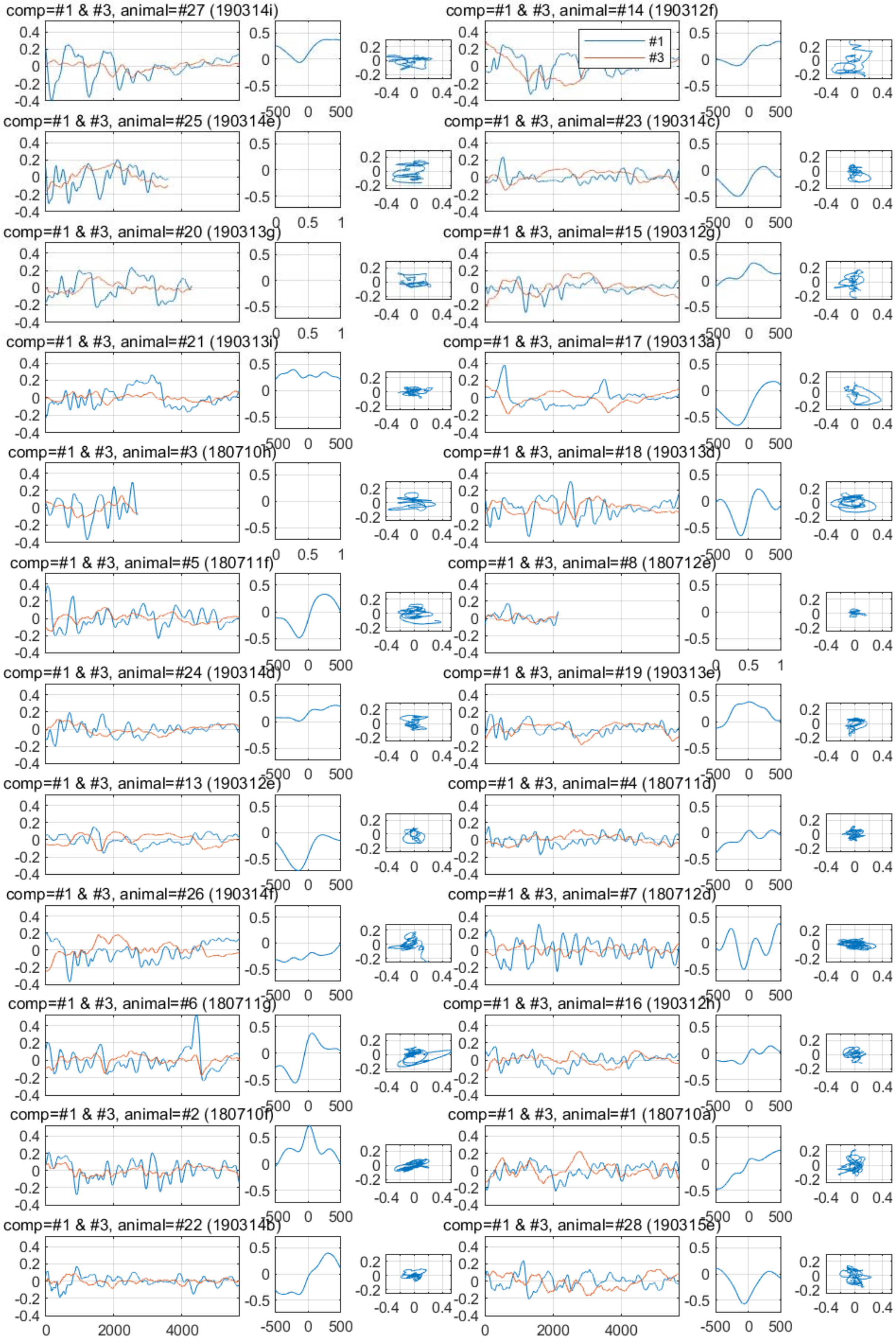
comp=#1 & #3, animal=#27 (190314i)
comp=#1 & #3, animal=#14 (190312f)
#1
#3
comp=#1 & #3, animal=#25 (190314e)
comp=#1 & #3, animal=#23 (190314c)
comp=#1 & #3, animal=#20 (190313g)
comp=#1 & #3, animal=#15 (190312g)
comp=#1 & #3, animal=#21 (190313i)
comp=#1 & #3, animal=#17 (190313a)
comp=#1 & #3, animal=#3 (180710h)
comp=#1 & #3, animal=#18 (190313d)
comp=#1 & #3, animal=#5 (180711f)
comp=#1 & #3, animal=#8 (180712e)
comp=#1 & #3, animal=#24 (190314d)
comp=#1 & #3, animal=#19 (190313e)
comp=#1 & #3, animal=#13 (190312e)
comp=#1 & #3, animal=#4 (180711d)
comp=#1 & #3, animal=#26 (190314f)
comp=#1 & #3, animal=#7 (180712d)
comp=#1 & #3, animal=#6 (180711g)
comp=#1 & #3, animal=#16 (190312h)
comp=#1 & #3, animal=#2 (180710f)
comp=#1 & #3, animal=#1 (180710a)
comp=#1 & #3, animal=#22 (190314b)
comp=#1 & #3, animal=#28 (190315e)

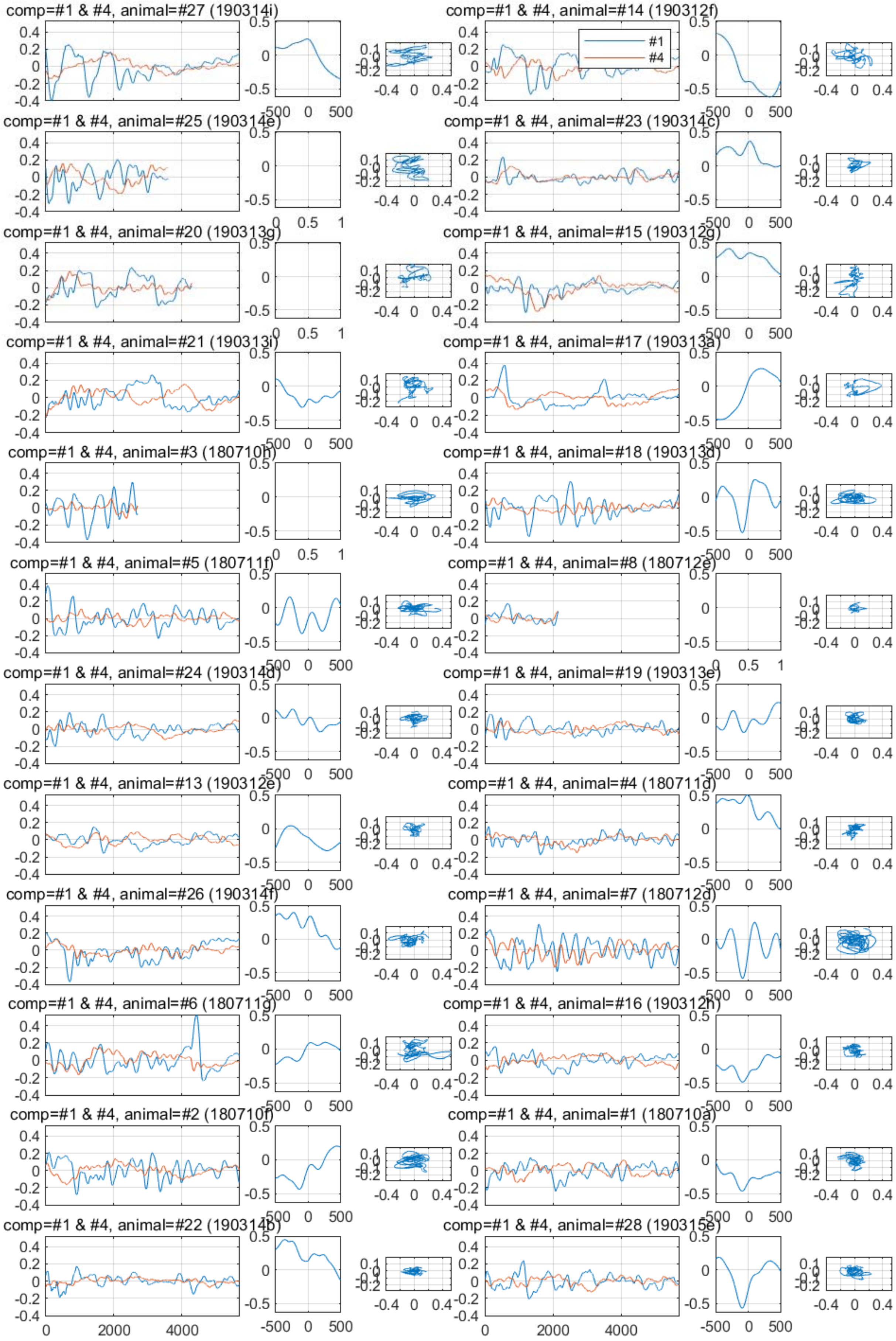
comp=#1 & #4, animal=#27 (190314i)
comp=#1 & #4, animal=#14 (190312f)
#1
#4
comp=#1 & #4, animal=#25 (190314e)
comp=#1 & #4, animal=#23 (190314c)
comp=#1 & #4, animal=#20 (190313g)
comp=#1 & #4, animal=#15 (190312g)
comp=#1 & #4, animal=#21 (190313i)
comp=#1 & #4, animal=#17 (190313a)
comp=#1 & #4, animal=#3 (180710h)
comp=#1 & #4, animal=#18 (190313d)
comp=#1 & #4, animal=#5 (180711f)
comp=#1 & #4, animal=#8 (180712e)
comp=#1 & #4, animal=#24 (190314d)
comp=#1 & #4, animal=#19 (190313e)
comp=#1 & #4, animal=#13 (190312e)
comp=#1 & #4, animal=#4 (180711d)
comp=#1 & #4, animal=#26 (190314f)
comp=#1 & #4, animal=#7 (180712d)
comp=#1 & #4, animal=#6 (180711g)
comp=#1 & #4, animal=#16 (190312h)
comp=#1 & #4, animal=#2 (180710f)
comp=#1 & #4, animal=#1 (180710a)
comp=#1 & #4, animal=#22 (190314b)
comp=#1 & #4, animal=#28 (190315e)
0
2000
4000
-500
0
500
-0.4
0
0.4

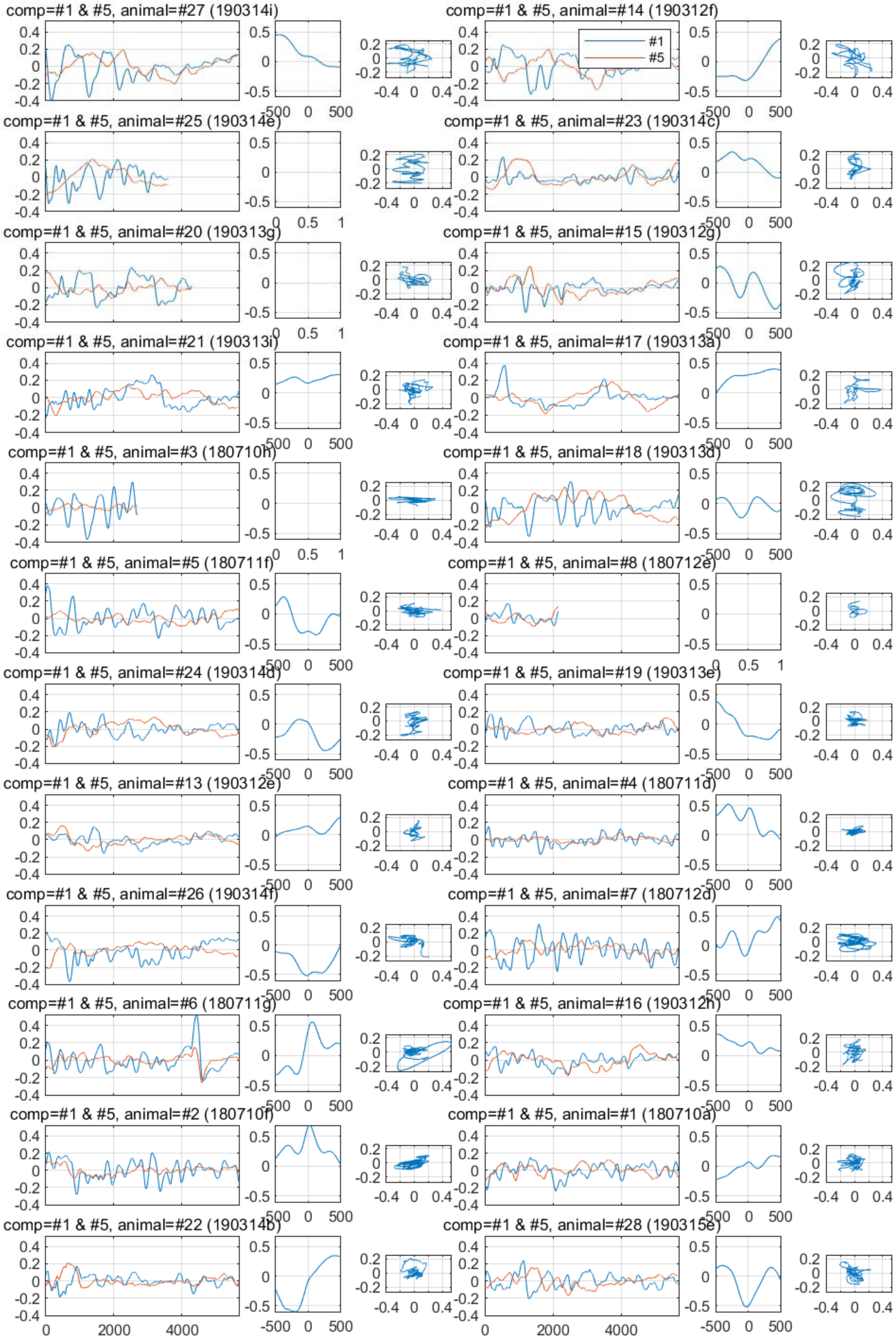

comp=#1 & #5, animal=#27 (190314i)
comp=#1 & #5, animal=#14 (190312f)
#1
#5
comp=#1 & #5, animal=#25 (190314e)
comp=#1 & #5, animal=#23 (190314c)
comp=#1 & #5, animal=#20 (190313g)
comp=#1 & #5, animal=#15 (190312g)
comp=#1 & #5, animal=#21 (190313i)
comp=#1 & #5, animal=#17 (190313a)
comp=#1 & #5, animal=#3 (180710h)
comp=#1 & #5, animal=#18 (190313d)
comp=#1 & #5, animal=#5 (180711f)
comp=#1 & #5, animal=#8 (180712e)
comp=#1 & #5, animal=#24 (190314d)
comp=#1 & #5, animal=#19 (190313e)
comp=#1 & #5, animal=#13 (190312e)
comp=#1 & #5, animal=#4 (180711d)
comp=#1 & #5, animal=#26 (190314f)
comp=#1 & #5, animal=#7 (180712d)
comp=#1 & #5, animal=#6 (180711g)
comp=#1 & #5, animal=#16 (190312h)
comp=#1 & #5, animal=#2 (180710f)
comp=#1 & #5, animal=#1 (180710a)
comp=#1 & #5, animal=#22 (190314b)
comp=#1 & #5, animal=#28 (190315e)

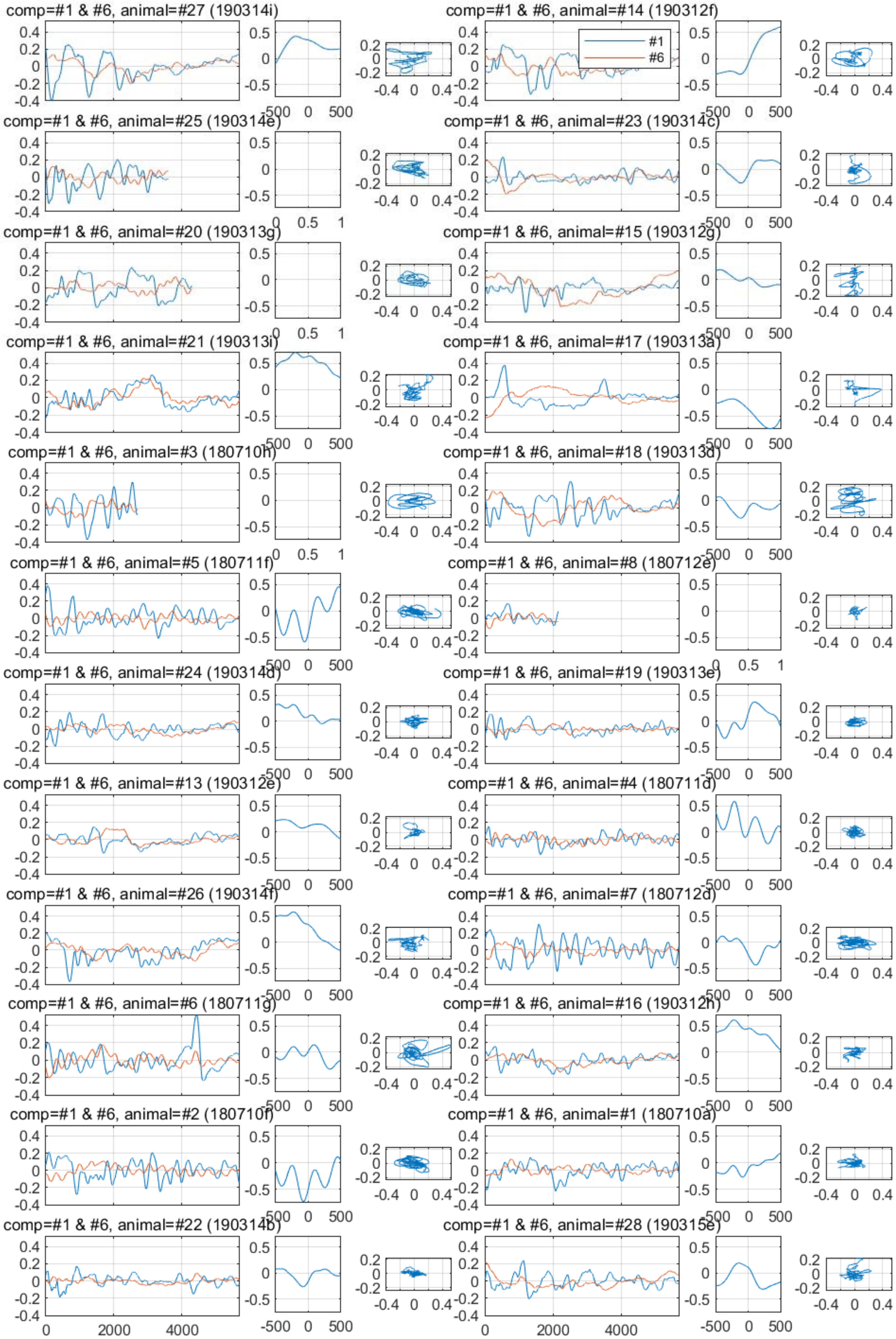
comp=#1 & #6, animal=#27 (190314i)
comp=#1 & #6, animal=#14 (190312f)
#1
#6
comp=#1 & #6, animal=#25 (190314e)
comp=#1 & #6, animal=#23 (190314c)
comp=#1 & #6, animal=#20 (190313g)
comp=#1 & #6, animal=#15 (190312g)
comp=#1 & #6, animal=#21 (190313i)
comp=#1 & #6, animal=#17 (190313a)
comp=#1 & #6, animal=#3 (180710h)
comp=#1 & #6, animal=#18 (190313d)
comp=#1 & #6, animal=#5 (180711f)
comp=#1 & #6, animal=#8 (180712e)
comp=#1 & #6, animal=#24 (190314d)
comp=#1 & #6, animal=#19 (190313e)
comp=#1 & #6, animal=#13 (190312e)
comp=#1 & #6, animal=#4 (180711d)
comp=#1 & #6, animal=#26 (190314f)
comp=#1 & #6, animal=#7 (180712d)
comp=#1 & #6, animal=#6 (180711g)
comp=#1 & #6, animal=#16 (190312h)
comp=#1 & #6, animal=#2 (180710f)
comp=#1 & #6, animal=#1 (180710a)
comp=#1 & #6, animal=#22 (190314b)
comp=#1 & #6, animal=#28 (190315e)
0
2000
4000
-500
0
500

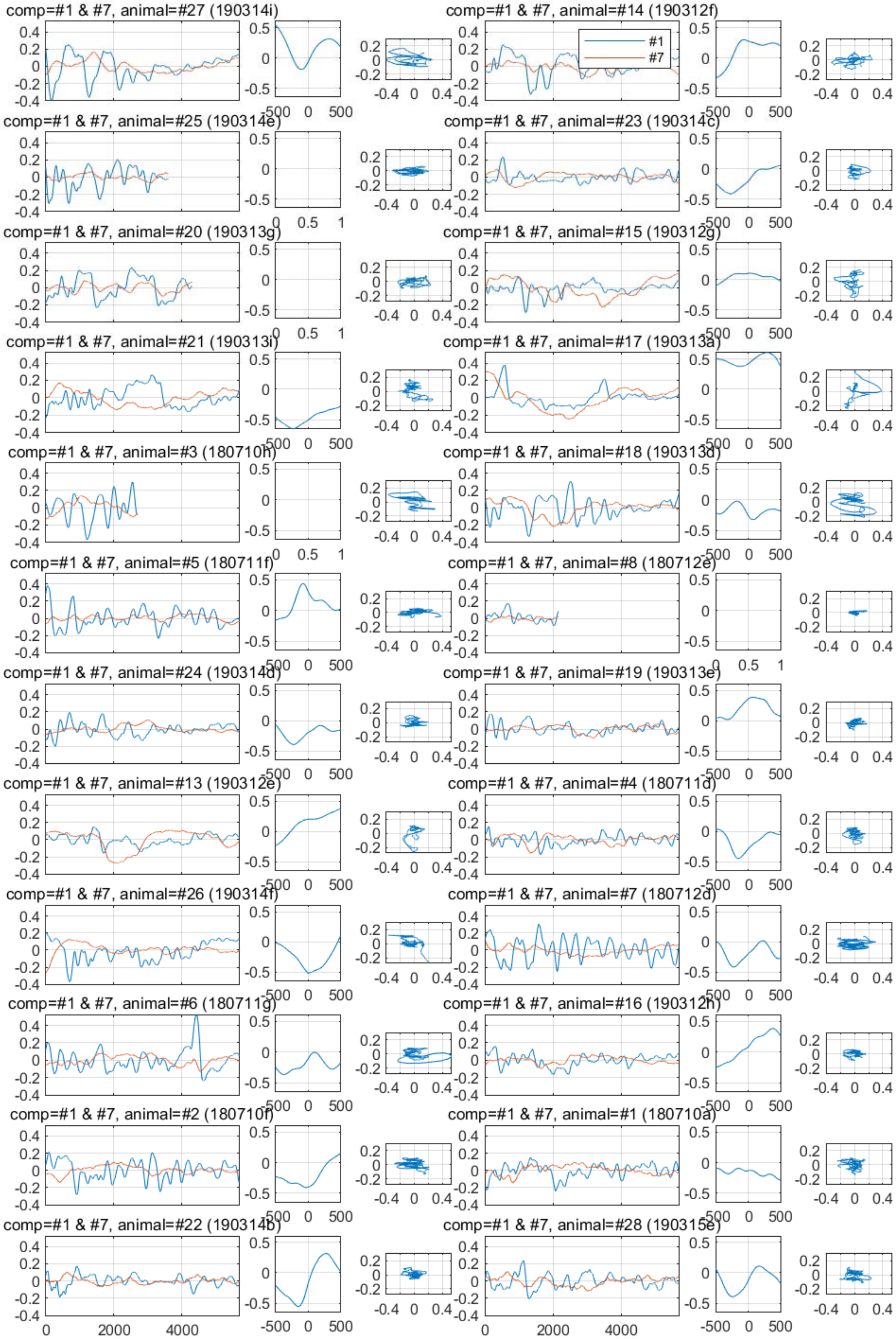

comp=#1 & #7, animal=#27 (190314i)
comp=#1 & #7, animal=#14 (190312f)
#1
#7
comp=#1 & #7, animal=#25 (190314e)
comp=#1 & #7, animal=#23 (190314c)
comp=#1 & #7, animal=#20 (190313g)
comp=#1 & #7, animal=#15 (190312g)
comp=#1 & #7, animal=#21 (190313i)
comp=#1 & #7, animal=#17 (190313a)
comp=#1 & #7, animal=#3 (180710h)
comp=#1 & #7, animal=#18 (190313d)
comp=#1 & #7, animal=#5 (180711f)
comp=#1 & #7, animal=#8 (180712e)
comp=#1 & #7, animal=#24 (190314d)
comp=#1 & #7, animal=#19 (190313e)
comp=#1 & #7, animal=#13 (190312e)
comp=#1 & #7, animal=#4 (180711d)
comp=#1 & #7, animal=#26 (190314f)
comp=#1 & #7, animal=#7 (180712d)
comp=#1 & #7, animal=#6 (180711g)
comp=#1 & #7, animal=#16 (190312h)
comp=#1 & #7, animal=#2 (180710f)
comp=#1 & #7, animal=#1 (180710a)
comp=#1 & #7, animal=#22 (190314b)
comp=#1 & #7, animal=#28 (190315e)

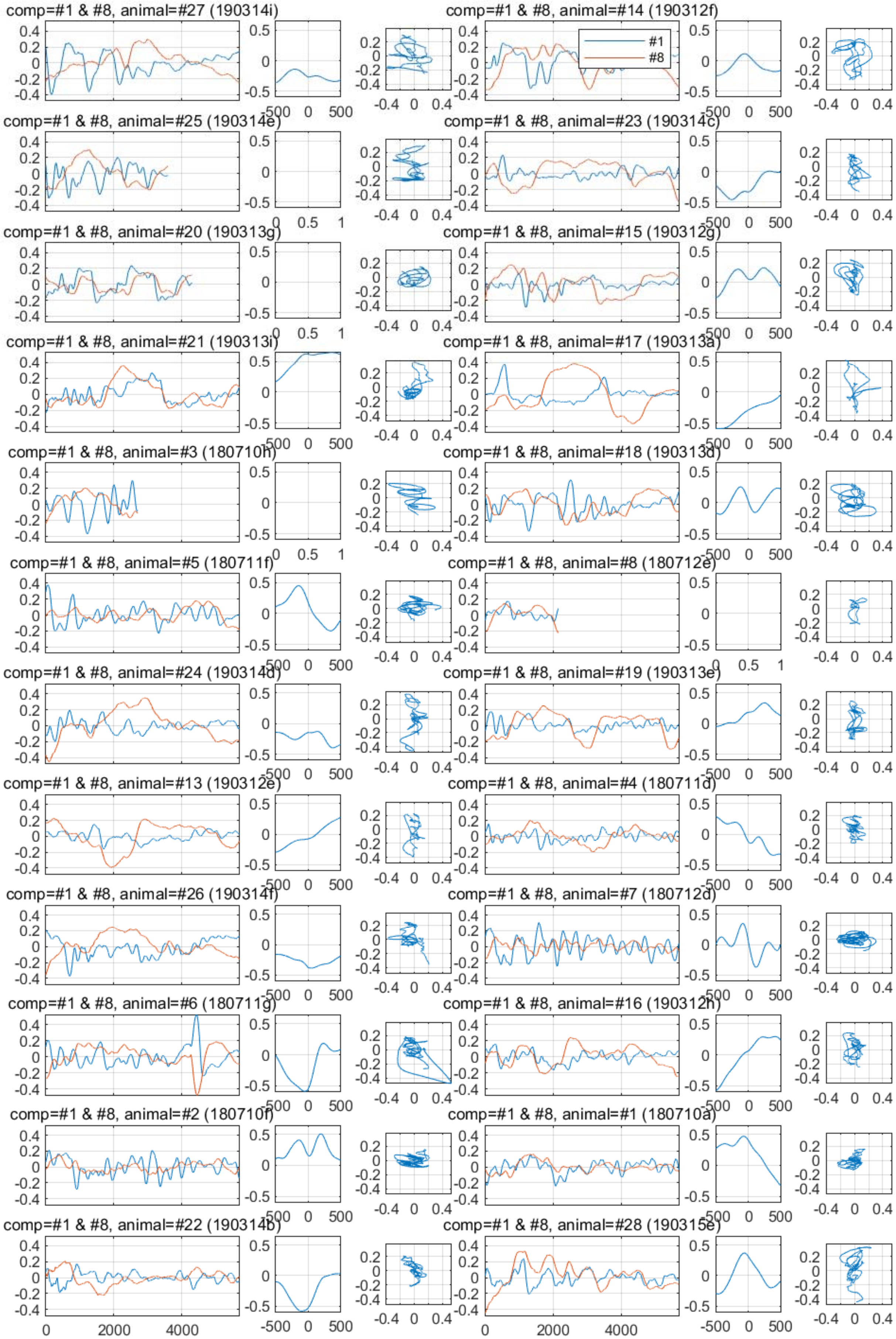

comp=#1 & #8, animal=#27 (190314i)
comp=#1 & #8, animal=#14 (190312f)
#1
#8
comp=#1 & #8, animal=#25 (190314e)
comp=#1 & #8, animal=#23 (190314c)
comp=#1 & #8, animal=#20 (190313g)
comp=#1 & #8, animal=#15 (190312g)
comp=#1 & #8, animal=#21 (190313i)
comp=#1 & #8, animal=#17 (190313a)
comp=#1 & #8, animal=#3 (180710h)
comp=#1 & #8, animal=#18 (190313d)
comp=#1 & #8, animal=#5 (180711f)
comp=#1 & #8, animal=#8 (180712e)
comp=#1 & #8, animal=#24 (190314d)
comp=#1 & #8, animal=#19 (190313e)
comp=#1 & #8, animal=#13 (190312e)
comp=#1 & #8, animal=#4 (180711d)
comp=#1 & #8, animal=#26 (190314f)
comp=#1 & #8, animal=#7 (180712d)
comp=#1 & #8, animal=#6 (180711g)
comp=#1 & #8, animal=#16 (190312h)
comp=#1 & #8, animal=#2 (180710f)
comp=#1 & #8, animal=#1 (180710a)
comp=#1 & #8, animal=#22 (190314b)
comp=#1 & #8, animal=#28 (190315e)
0
2000
4000
-500
0
500
-0.4
0
0.4

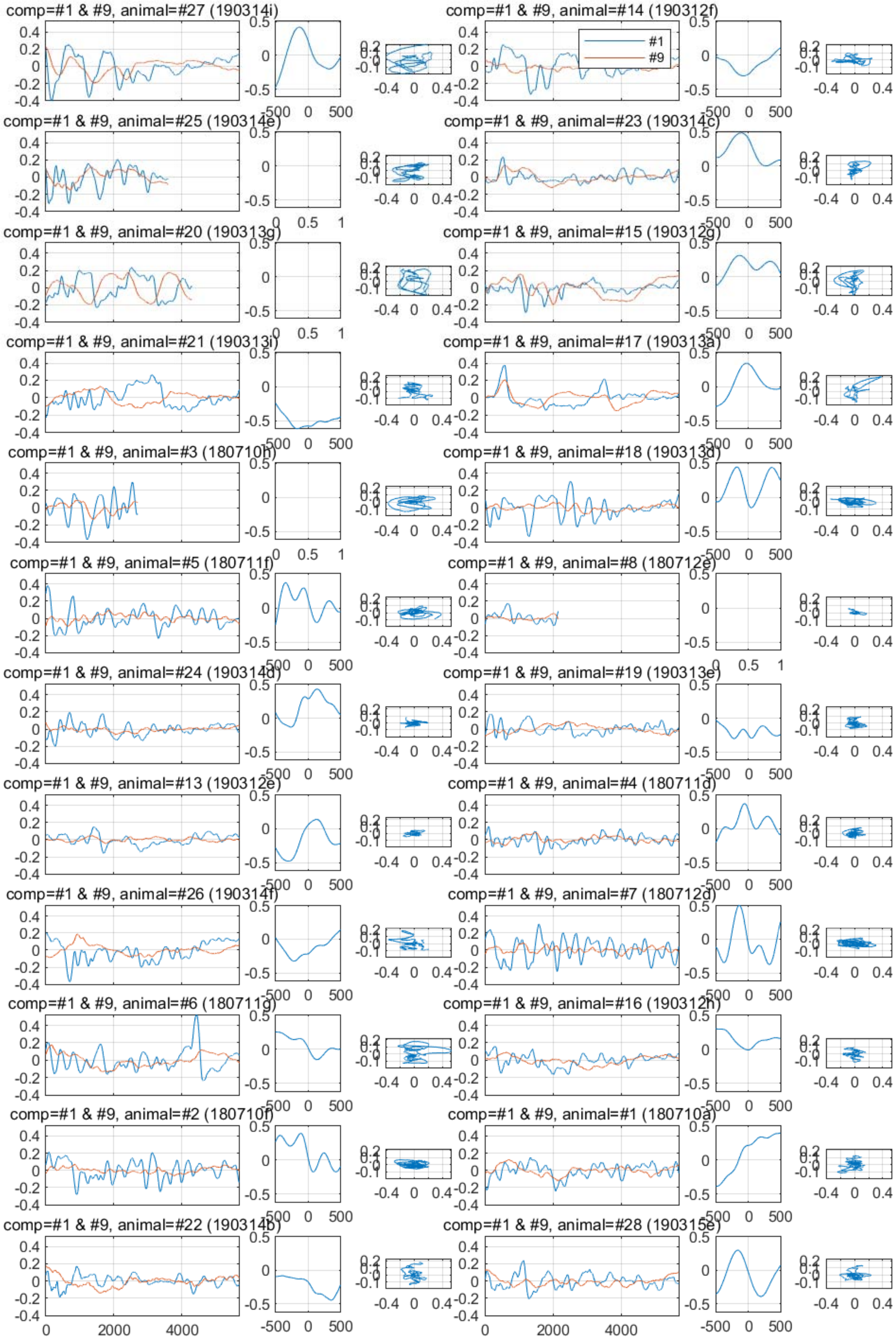
comp=#1 & #9, animal=#27 (190314i)
comp=#1 & #9, animal=#14 (190312f)
#1
#9
comp=#1 & #9, animal=#25 (190314e)
comp=#1 & #9, animal=#23 (190314c)
comp=#1 & #9, animal=#20 (190313g)
comp=#1 & #9, animal=#15 (190312g)
comp=#1 & #9, animal=#21 (190313i)
comp=#1 & #9, animal=#17 (190313a)
comp=#1 & #9, animal=#3 (180710h)
comp=#1 & #9, animal=#18 (190313d)
comp=#1 & #9, animal=#5 (180711f)
comp=#1 & #9, animal=#8 (180712e)
comp=#1 & #9, animal=#24 (190314d)
comp=#1 & #9, animal=#19 (190313e)
comp=#1 & #9, animal=#13 (190312e)
comp=#1 & #9, animal=#4 (180711d)
comp=#1 & #9, animal=#26 (190314f)
comp=#1 & #9, animal=#7 (180712d)
comp=#1 & #9, animal=#6 (180711g)
comp=#1 & #9, animal=#16 (190312h)
comp=#1 & #9, animal=#2 (180710f)
comp=#1 & #9, animal=#1 (180710a)
comp=#1 & #9, animal=#22 (190314b)
comp=#1 & #9, animal=#28 (190315e)

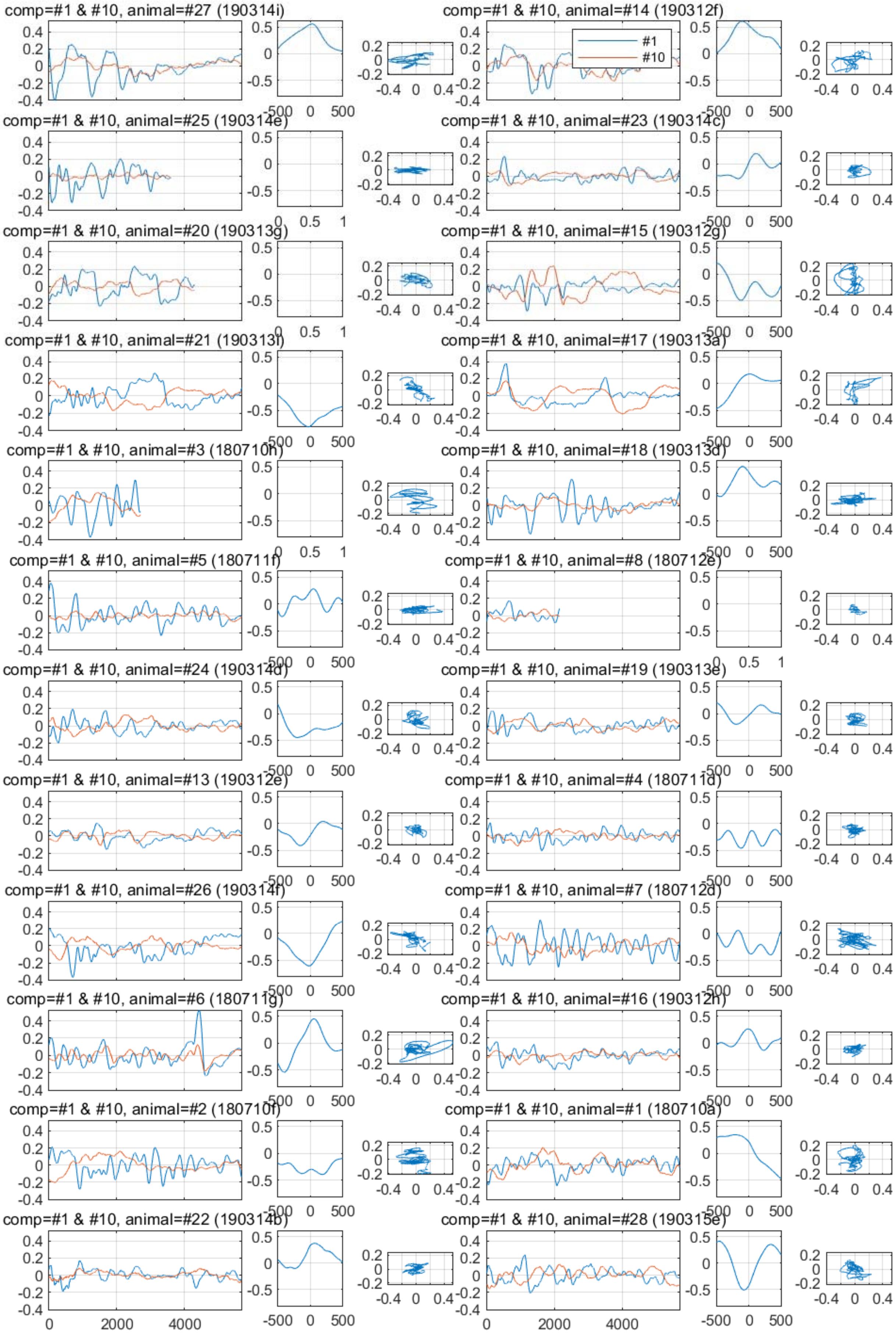
comp=#1 & #10, animal=#27 (190314i)
comp=#1 & #10, animal=#14 (190312f)
#1
#10
comp=#1 & #10, animal=#25 (190314e)
comp=#1 & #10, animal=#23 (190314c)
comp=#1 & #10, animal=#20 (190313g)
comp=#1 & #10, animal=#15 (190312g)
comp=#1 & #10, animal=#21 (190313i)
comp=#1 & #10, animal=#17 (190313a)
comp=#1 & #10, animal=#3 (180710h)
comp=#1 & #10, animal=#18 (190313d)
comp=#1 & #10, animal=#5 (180711f)
comp=#1 & #10, animal=#8 (180712e)
comp=#1 & #10, animal=#24 (190314d)
comp=#1 & #10, animal=#19 (190313e)
comp=#1 & #10, animal=#13 (190312e)
comp=#1 & #10, animal=#4 (180711d)
comp=#1 & #10, animal=#26 (190314f)
comp=#1 & #10, animal=#7 (180712d)
comp=#1 & #10, animal=#6 (180711g)
comp=#1 & #10, animal=#16 (190312h)
comp=#1 & #10, animal=#2 (180710f)
comp=#1 & #10, animal=#1 (180710a)
comp=#1 & #10, animal=#22 (190314b)
comp=#1 & #10, animal=#28 (190315e)

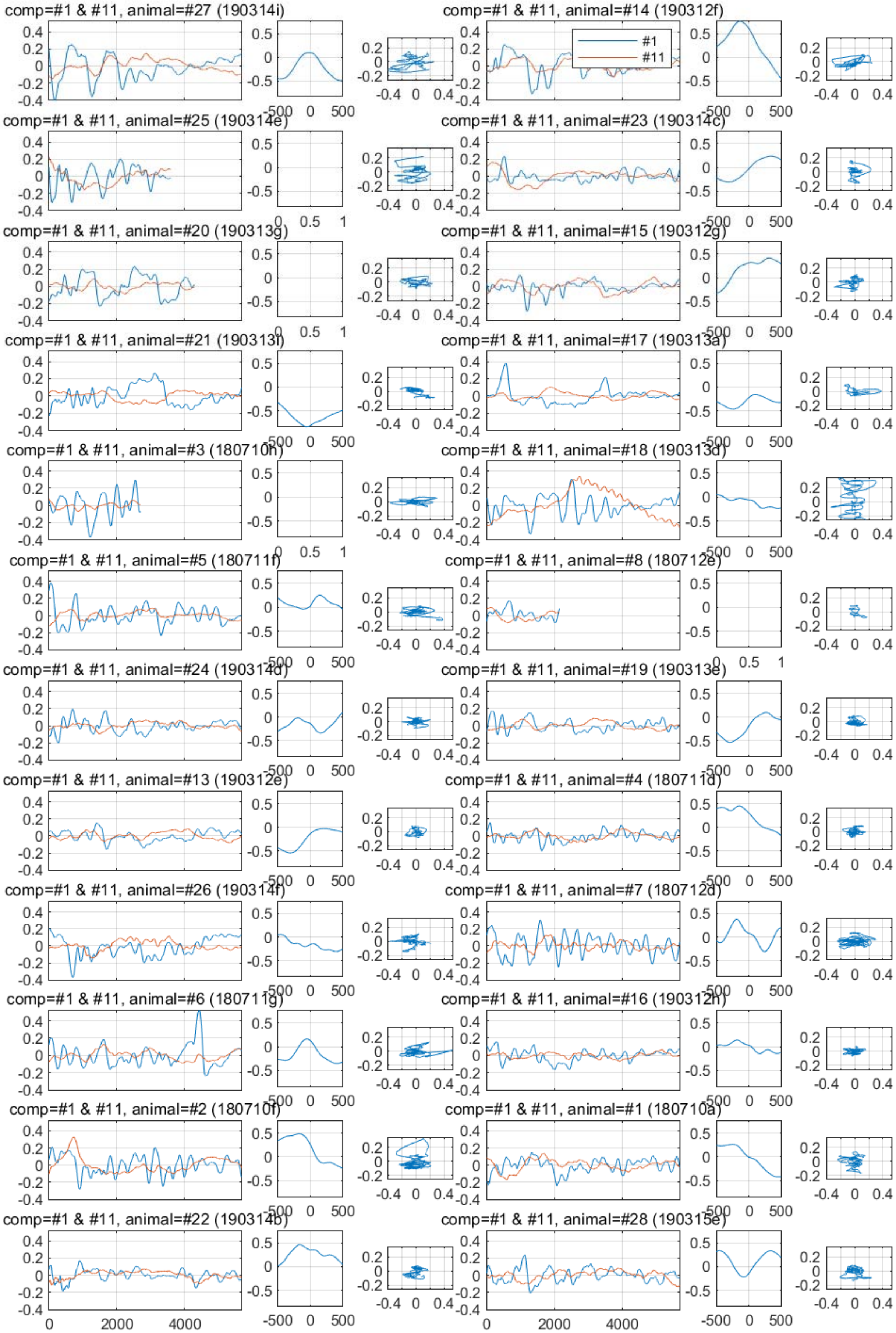

comp=#1 & #11, animal=#27 (190314i)
comp=#1 & #11, animal=#14 (190312f)
#1
#11
comp=#1 & #11, animal=#25 (190314e)
comp=#1 & #11, animal=#23 (190314c)
comp=#1 & #11, animal=#20 (190313g)
comp=#1 & #11, animal=#15 (190312g)
comp=#1 & #11, animal=#21 (190313i)
comp=#1 & #11, animal=#17 (190313a)
comp=#1 & #11, animal=#3 (180710h)
comp=#1 & #11, animal=#18 (190313d)
comp=#1 & #11, animal=#5 (180711f)
comp=#1 & #11, animal=#8 (180712e)
comp=#1 & #11, animal=#24 (190314d)
comp=#1 & #11, animal=#19 (190313e)
comp=#1 & #11, animal=#13 (190312e)
comp=#1 & #11, animal=#4 (180711d)
comp=#1 & #11, animal=#26 (190314f)
comp=#1 & #11, animal=#7 (180712d)
comp=#1 & #11, animal=#6 (180711g)
comp=#1 & #11, animal=#16 (190312h)
comp=#1 & #11, animal=#2 (180710f)
comp=#1 & #11, animal=#1 (180710a)
comp=#1 & #11, animal=#22 (190314b)
comp=#1 & #11, animal=#28 (190315e)

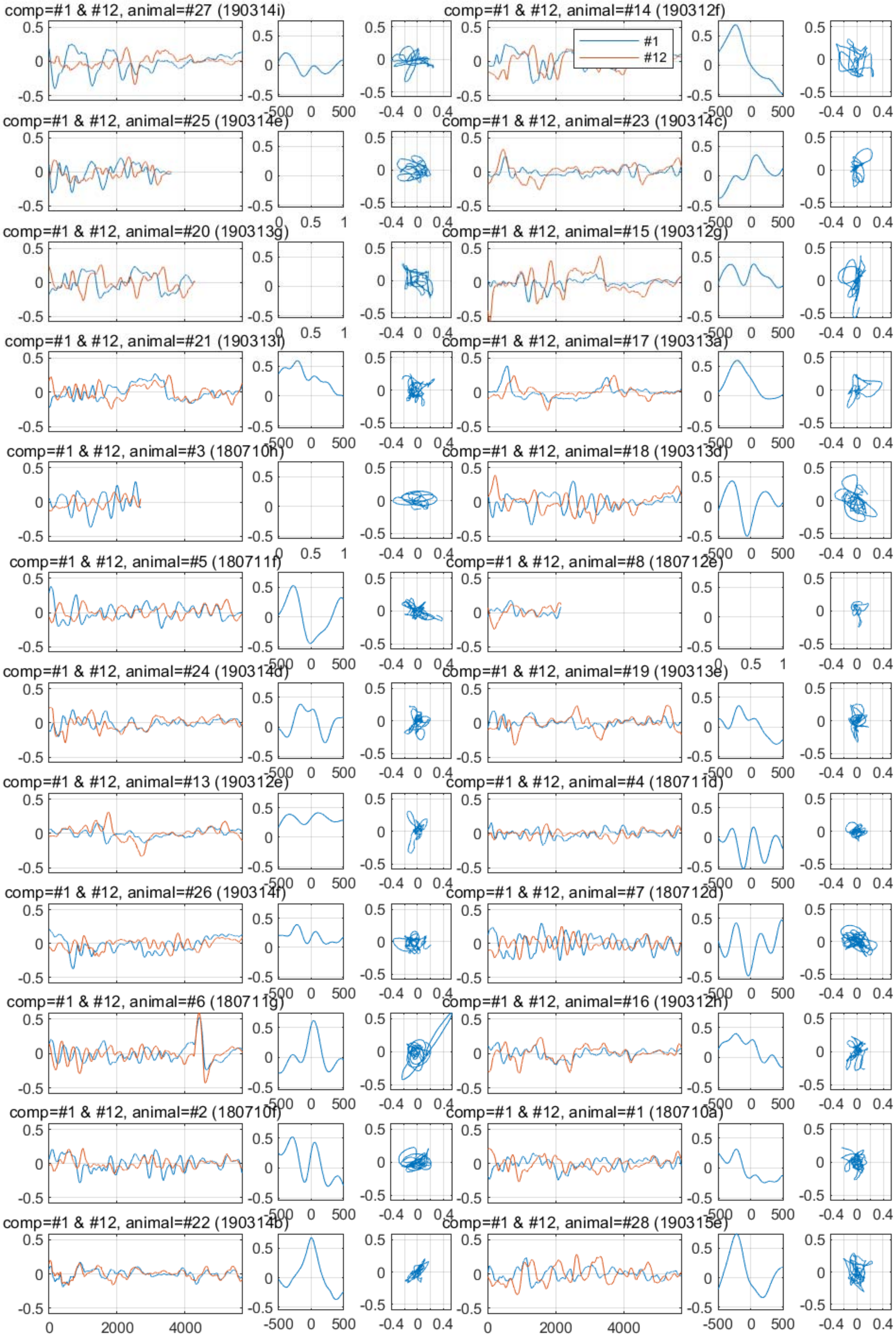
comp=#1 & #12, animal=#27 (190314i)
comp=#1 & #12, animal=#14 (190312f)
#1
#12
comp=#1 & #12, animal=#25 (190314e)
comp=#1 & #12, animal=#23 (190314c)
comp=#1 & #12, animal=#20 (190313g)
comp=#1 & #12, animal=#15 (190312g)
comp=#1 & #12, animal=#21 (190313f)
comp=#1 & #12, animal=#17 (190313a)
comp=#1 & #12, animal=#3 (180710h)
comp=#1 & #12, animal=#18 (190313d)
comp=#1 & #12, animal=#5 (180711f)
comp=#1 & #12, animal=#8 (180712e)
comp=#1 & #12, animal=#24 (190314d)
comp=#1 & #12, animal=#19 (190313e)
comp=#1 & #12, animal=#13 (190312e)
comp=#1 & #12, animal=#4 (180711d)
comp=#1 & #12, animal=#26 (190314f)
comp=#1 & #12, animal=#7 (180712d)
comp=#1 & #12, animal=#6 (180711g)
comp=#1 & #12, animal=#16 (190312h)
comp=#1 & #12, animal=#2 (180710f)
comp=#1 & #12, animal=#1 (180710a)
comp=#1 & #12, animal=#22 (190314b)
comp=#1 & #12, animal=#28 (190315e)

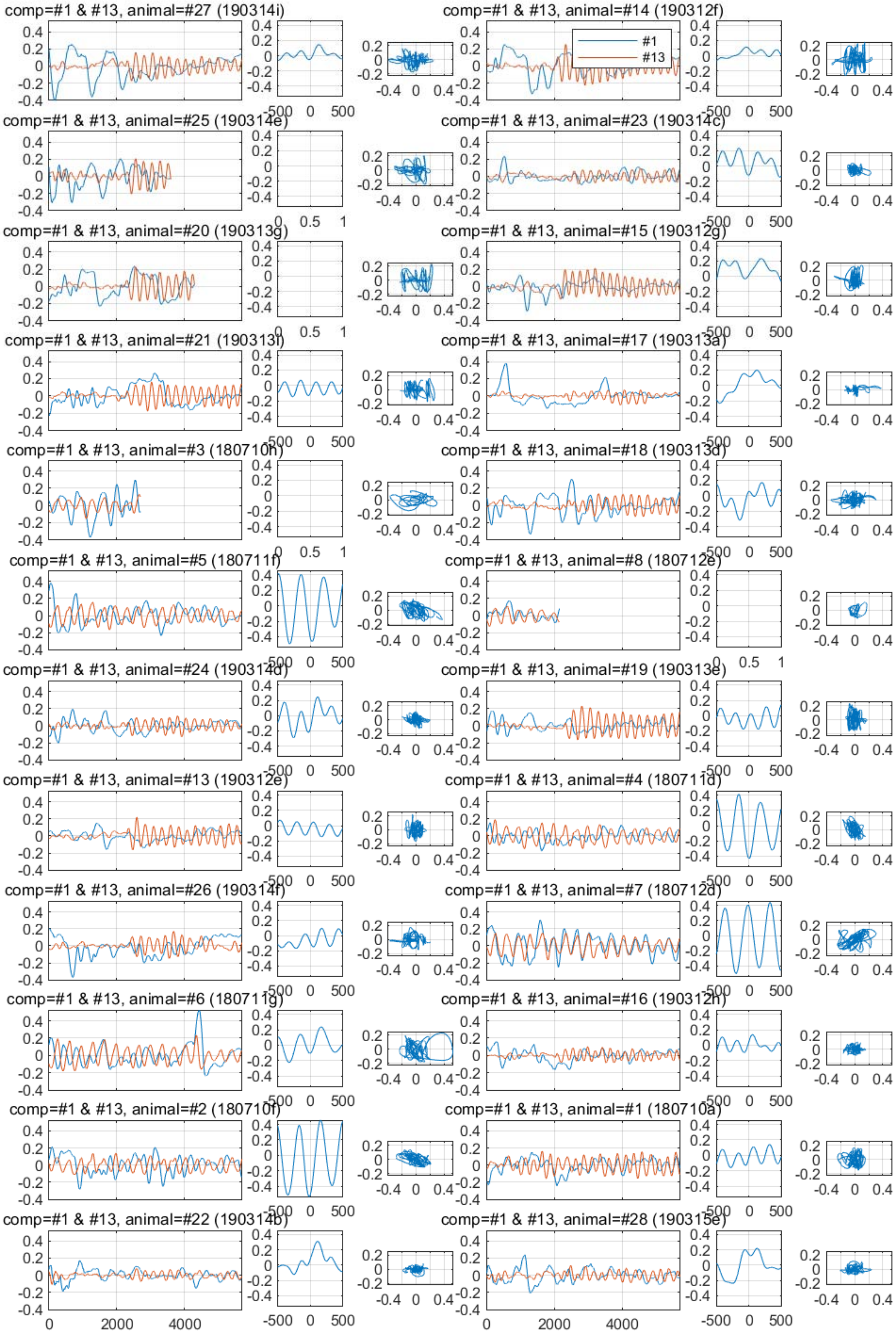
comp=#1 & #13, animal=#27 (190314i)
comp=#1 & #13, animal=#14 (190312f)
#1
#13
comp=#1 & #13, animal=#25 (190314e)
comp=#1 & #13, animal=#23 (190314c)
comp=#1 & #13, animal=#20 (190313g)
comp=#1 & #13, animal=#15 (190312g)
comp=#1 & #13, animal=#21 (190313i)
comp=#1 & #13, animal=#17 (190313a)
comp=#1 & #13, animal=#3 (180710h)
comp=#1 & #13, animal=#18 (190313d)
comp=#1 & #13, animal=#5 (180711f)
comp=#1 & #13, animal=#8 (180712e)
comp=#1 & #13, animal=#24 (190314d)
comp=#1 & #13, animal=#19 (190313e)
comp=#1 & #13, animal=#13 (190312e)
comp=#1 & #13, animal=#4 (180711d)
comp=#1 & #13, animal=#26 (190314f)
comp=#1 & #13, animal=#7 (180712d)
comp=#1 & #13, animal=#6 (180711g)
comp=#1 & #13, animal=#16 (190312h)
comp=#1 & #13, animal=#2 (180710f)
comp=#1 & #13, animal=#1 (180710a)
comp=#1 & #13, animal=#22 (190314b)
comp=#1 & #13, animal=#28 (190315e)

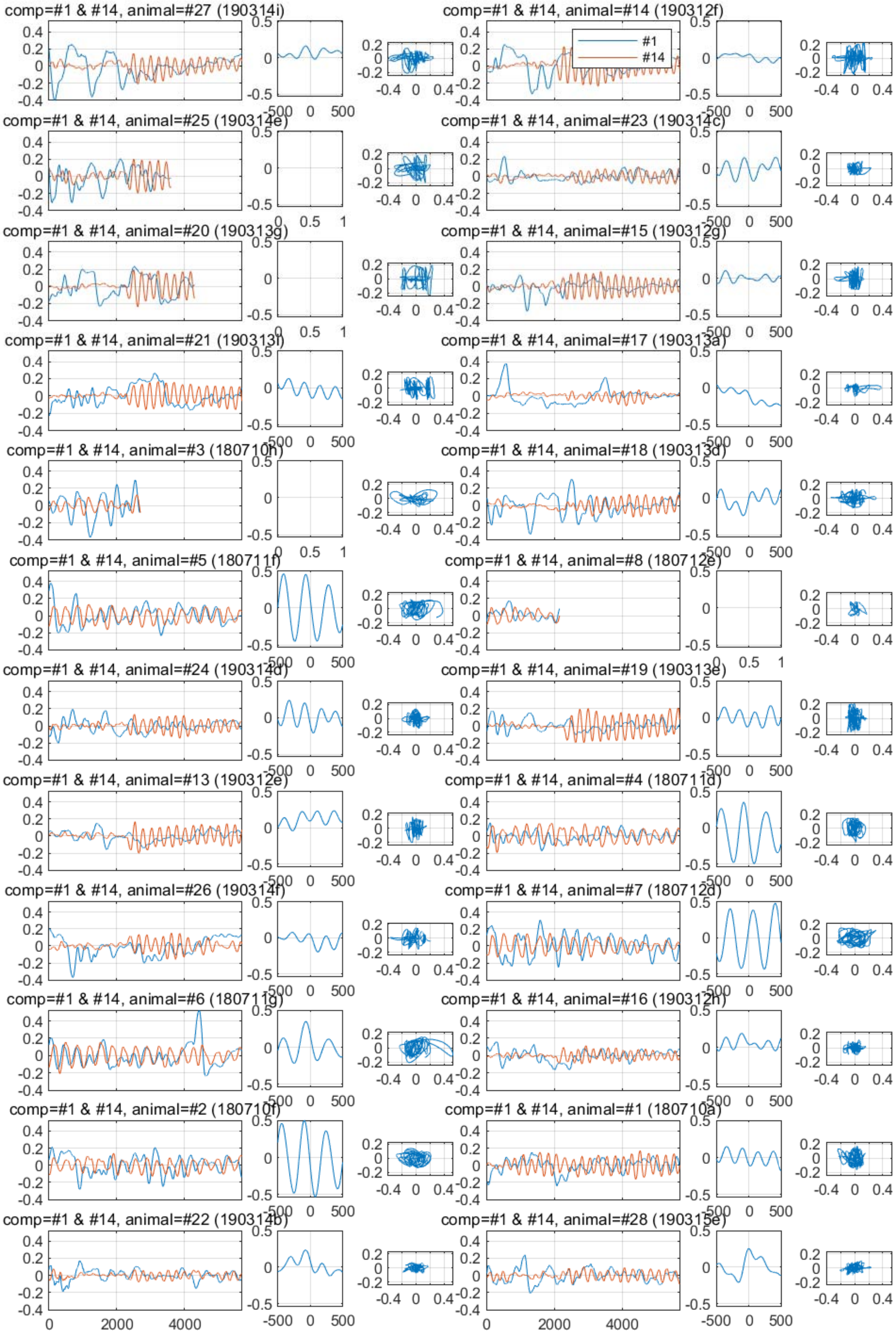
comp=#1 & #14, animal=#27 (190314i)
comp=#1 & #14, animal=#14 (190312f)
#1
#14
comp=#1 & #14, animal=#25 (190314e)
comp=#1 & #14, animal=#23 (190314c)
comp=#1 & #14, animal=#20 (190313g)
comp=#1 & #14, animal=#15 (190312g)
comp=#1 & #14, animal=#21 (190313i)
comp=#1 & #14, animal=#17 (190313a)
comp=#1 & #14, animal=#3 (180710h)
comp=#1 & #14, animal=#18 (190313d)
comp=#1 & #14, animal=#5 (180711f)
comp=#1 & #14, animal=#8 (180712e)
comp=#1 & #14, animal=#24 (190314d)
comp=#1 & #14, animal=#19 (190313e)
comp=#1 & #14, animal=#13 (190312e)
comp=#1 & #14, animal=#4 (180711d)
comp=#1 & #14, animal=#26 (190314f)
comp=#1 & #14, animal=#7 (180712d)
comp=#1 & #14, animal=#6 (180711g)
comp=#1 & #14, animal=#16 (190312h)
comp=#1 & #14, animal=#2 (180710f)
comp=#1 & #14, animal=#1 (180710a)
comp=#1 & #14, animal=#22 (190314b)
comp=#1 & #14, animal=#28 (190315e)

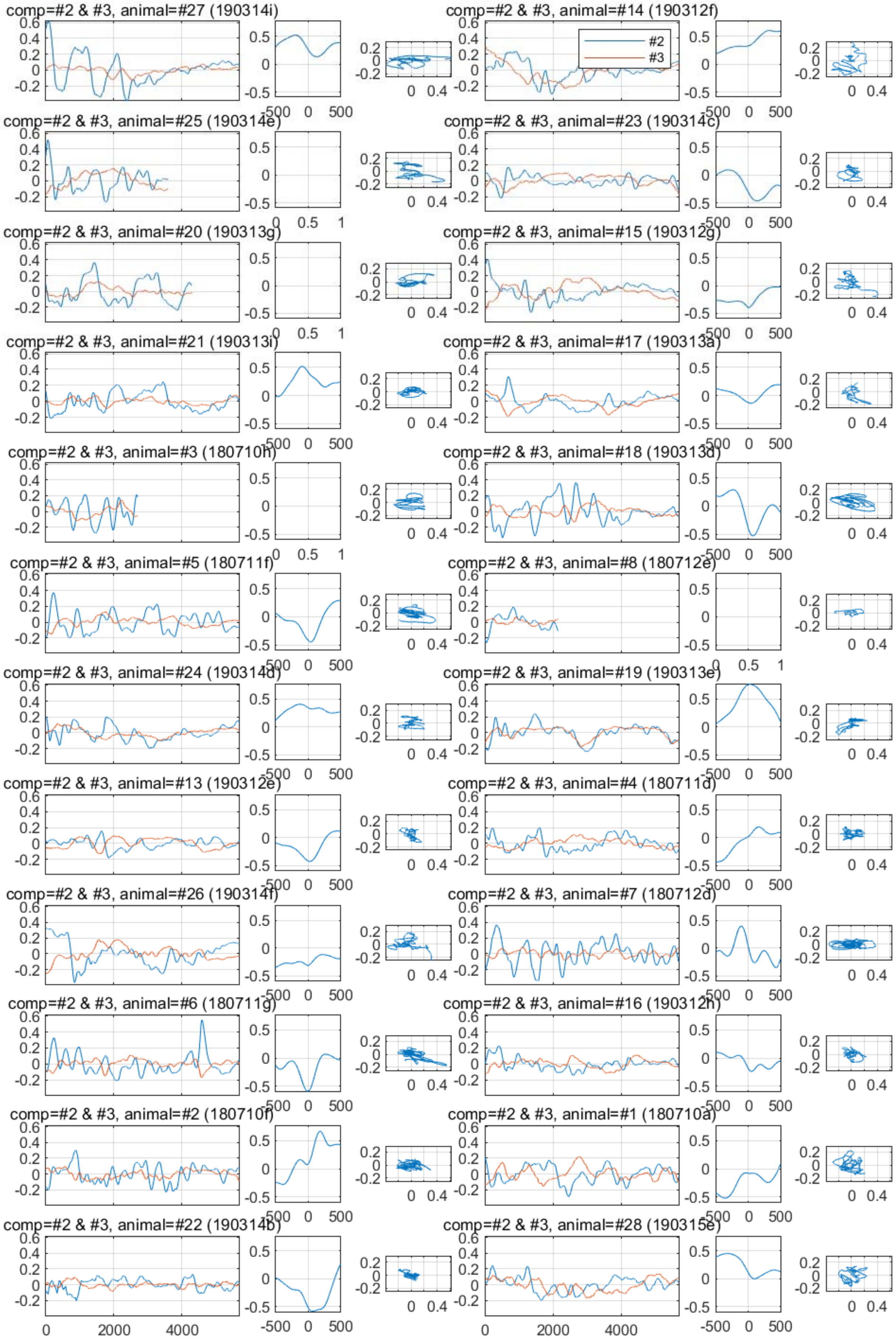
comp=#2 & #3, animal=#27 (190314i)
comp=#2 & #3, animal=#14 (190312f)
#2
#3
comp=#2 & #3, animal=#25 (190314e)
comp=#2 & #3, animal=#23 (190314c)
comp=#2 & #3, animal=#20 (190313g)
comp=#2 & #3, animal=#15 (190312g)
comp=#2 & #3, animal=#21 (190313i)
comp=#2 & #3, animal=#17 (190313a)
comp=#2 & #3, animal=#3 (180710h)
comp=#2 & #3, animal=#18 (190313d)
comp=#2 & #3, animal=#5 (180711f)
comp=#2 & #3, animal=#8 (180712e)
comp=#2 & #3, animal=#24 (190314d)
comp=#2 & #3, animal=#19 (190313e)
comp=#2 & #3, animal=#13 (190312e)
comp=#2 & #3, animal=#4 (180711d)
comp=#2 & #3, animal=#26 (190314f)
comp=#2 & #3, animal=#7 (180712d)
comp=#2 & #3, animal=#6 (180711g)
comp=#2 & #3, animal=#16 (190312h)
comp=#2 & #3, animal=#2 (180710f)
comp=#2 & #3, animal=#1 (180710a)
comp=#2 & #3, animal=#22 (190314b)
comp=#2 & #3, animal=#28 (190315e)

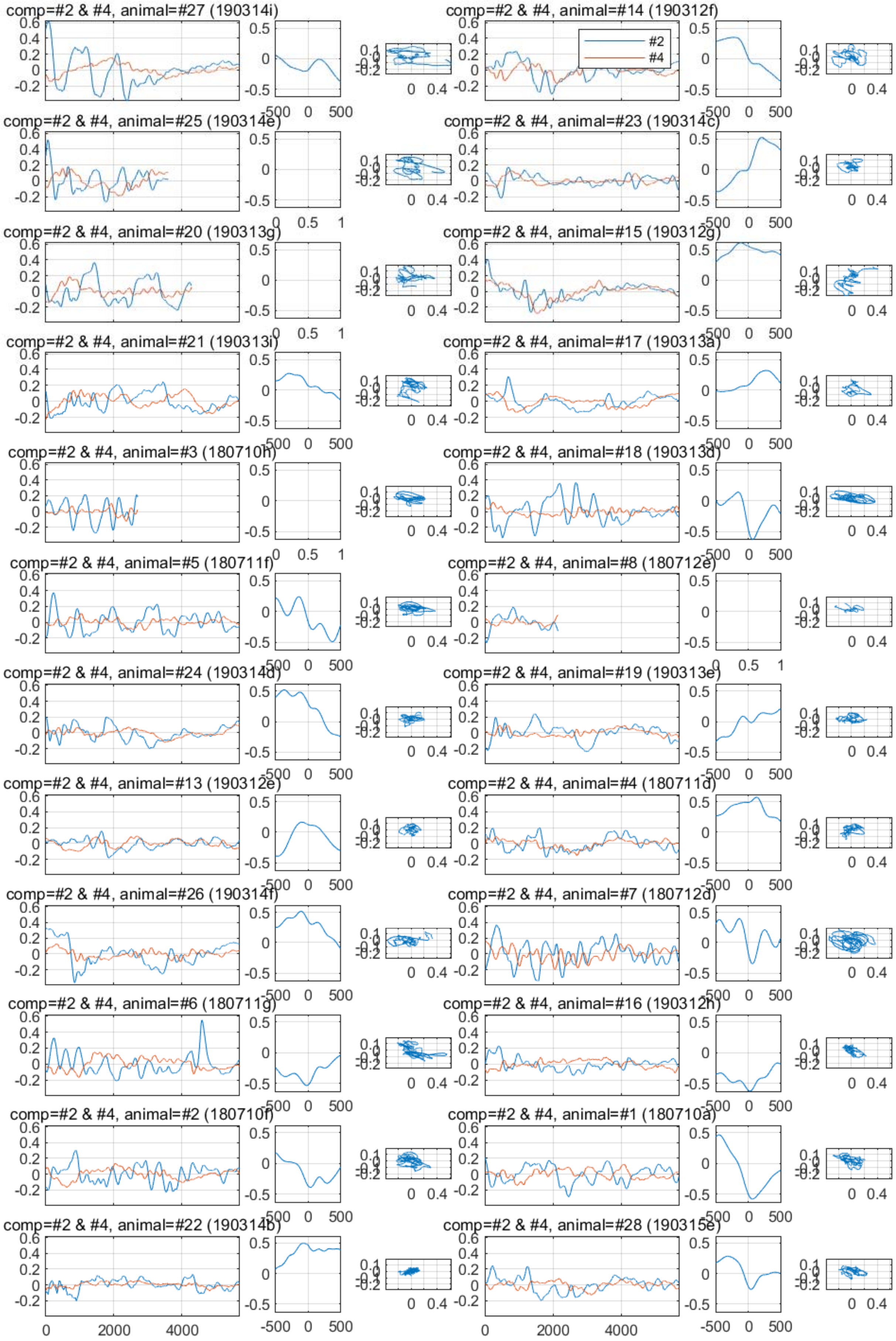
comp=#2 & #4, animal=#27 (190314i)
comp=#2 & #4, animal=#14 (190312f)
#2
#4
comp=#2 & #4, animal=#25 (190314e)
comp=#2 & #4, animal=#23 (190314c)
comp=#2 & #4, animal=#20 (190313g)
comp=#2 & #4, animal=#15 (190312g)
comp=#2 & #4, animal=#21 (190313i)
comp=#2 & #4, animal=#17 (190313a)
comp=#2 & #4, animal=#3 (180710h)
comp=#2 & #4, animal=#18 (190313d)
comp=#2 & #4, animal=#5 (180711f)
comp=#2 & #4, animal=#8 (180712e)
comp=#2 & #4, animal=#24 (190314d)
comp=#2 & #4, animal=#19 (190313e)
comp=#2 & #4, animal=#13 (190312e)
comp=#2 & #4, animal=#4 (180711d)
comp=#2 & #4, animal=#26 (190314f)
comp=#2 & #4, animal=#7 (180712d)
comp=#2 & #4, animal=#6 (180711g)
comp=#2 & #4, animal=#16 (190312h)
comp=#2 & #4, animal=#2 (180710f)
comp=#2 & #4, animal=#1 (180710a)
comp=#2 & #4, animal=#22 (190314b)
comp=#2 & #4, animal=#28 (190315e)

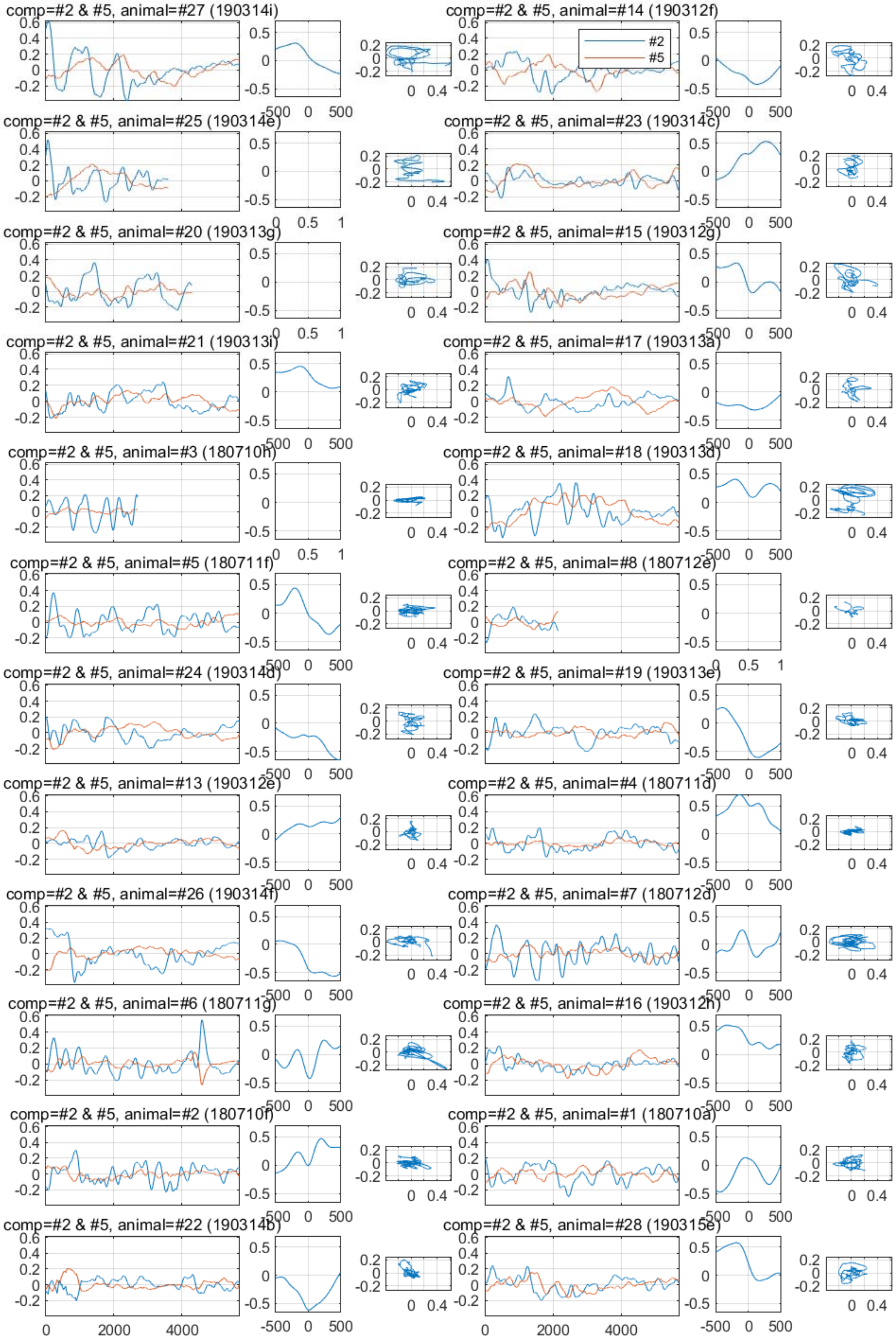
comp=#2 & #5, animal=#27 (190314i)
comp=#2 & #5, animal=#14 (190312f)
#2
#5
comp=#2 & #5, animal=#25 (190314e)
comp=#2 & #5, animal=#23 (190314c)
comp=#2 & #5, animal=#20 (190313g)
comp=#2 & #5, animal=#15 (190312g)
comp=#2 & #5, animal=#21 (190313i)
comp=#2 & #5, animal=#17 (190313a)
comp=#2 & #5, animal=#3 (180710h)
comp=#2 & #5, animal=#18 (190313d)
comp=#2 & #5, animal=#5 (180711f)
comp=#2 & #5, animal=#8 (180712e)
comp=#2 & #5, animal=#24 (190314d)
comp=#2 & #5, animal=#19 (190313e)
comp=#2 & #5, animal=#13 (190312e)
comp=#2 & #5, animal=#4 (180711d)
comp=#2 & #5, animal=#26 (190314f)
comp=#2 & #5, animal=#7 (180712d)
comp=#2 & #5, animal=#6 (180711g)
comp=#2 & #5, animal=#16 (190312h)
comp=#2 & #5, animal=#2 (180710f)
comp=#2 & #5, animal=#1 (180710a)
comp=#2 & #5, animal=#22 (190314b)
comp=#2 & #5, animal=#28 (190315e)
0
2000
4000
-500
0
500

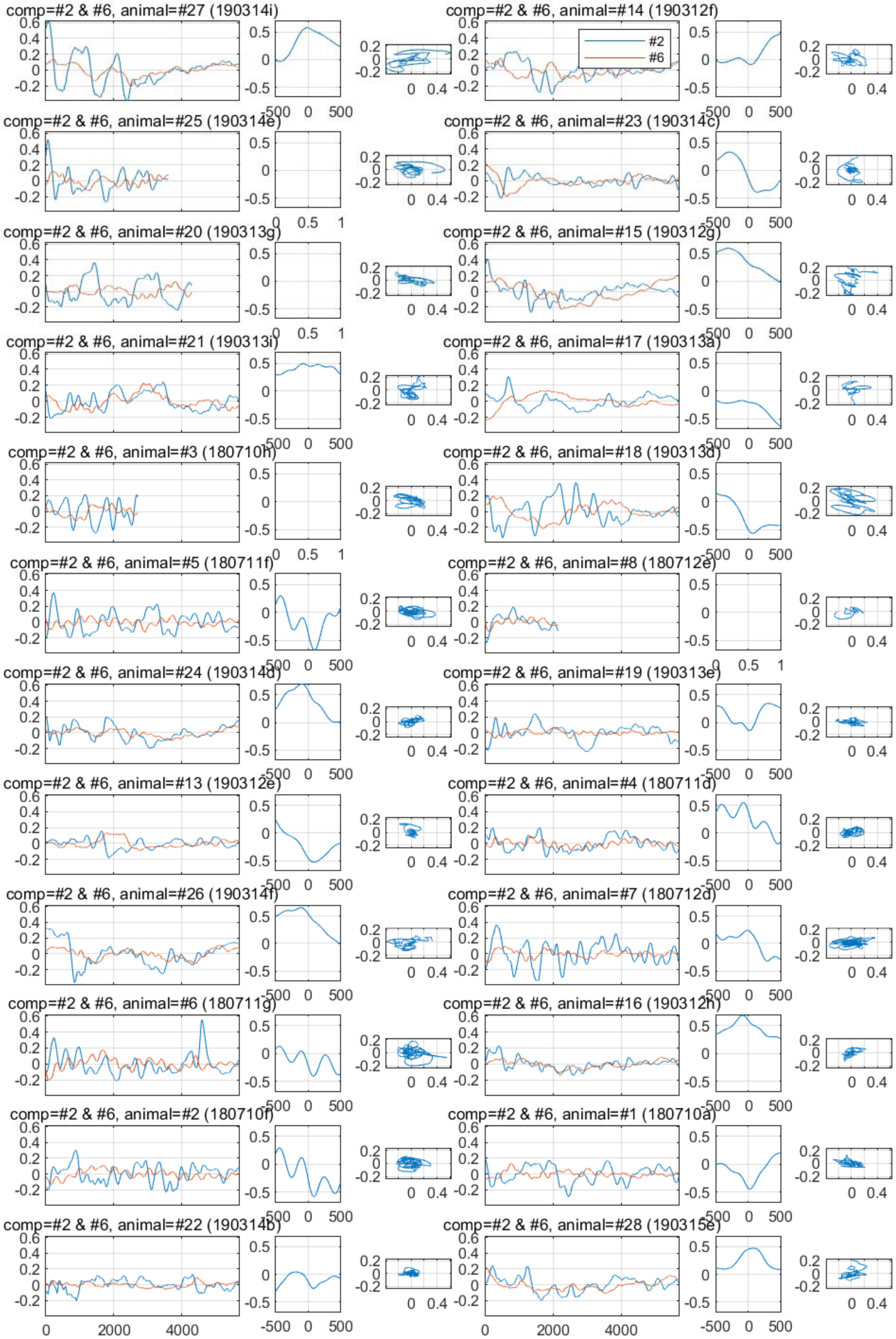
comp=#2 & #6, animal=#27 (190314i)
comp=#2 & #6, animal=#14 (190312f)
#2
#6
comp=#2 & #6, animal=#25 (190314e)
comp=#2 & #6, animal=#23 (190314c)
comp=#2 & #6, animal=#20 (190313g)
comp=#2 & #6, animal=#15 (190312g)
comp=#2 & #6, animal=#21 (190313i)
comp=#2 & #6, animal=#17 (190313a)
comp=#2 & #6, animal=#3 (180710h)
comp=#2 & #6, animal=#18 (190313d)
comp=#2 & #6, animal=#5 (180711f)
comp=#2 & #6, animal=#8 (180712e)
comp=#2 & #6, animal=#24 (190314d)
comp=#2 & #6, animal=#19 (190313e)
comp=#2 & #6, animal=#13 (190312e)
comp=#2 & #6, animal=#4 (180711d)
comp=#2 & #6, animal=#26 (190314f)
comp=#2 & #6, animal=#7 (180712d)
comp=#2 & #6, animal=#6 (180711g)
comp=#2 & #6, animal=#16 (190312h)
comp=#2 & #6, animal=#2 (180710f)
comp=#2 & #6, animal=#1 (180710a)
comp=#2 & #6, animal=#22 (190314b)
comp=#2 & #6, animal=#28 (190315e)
0
2000
4000
-500
0
500

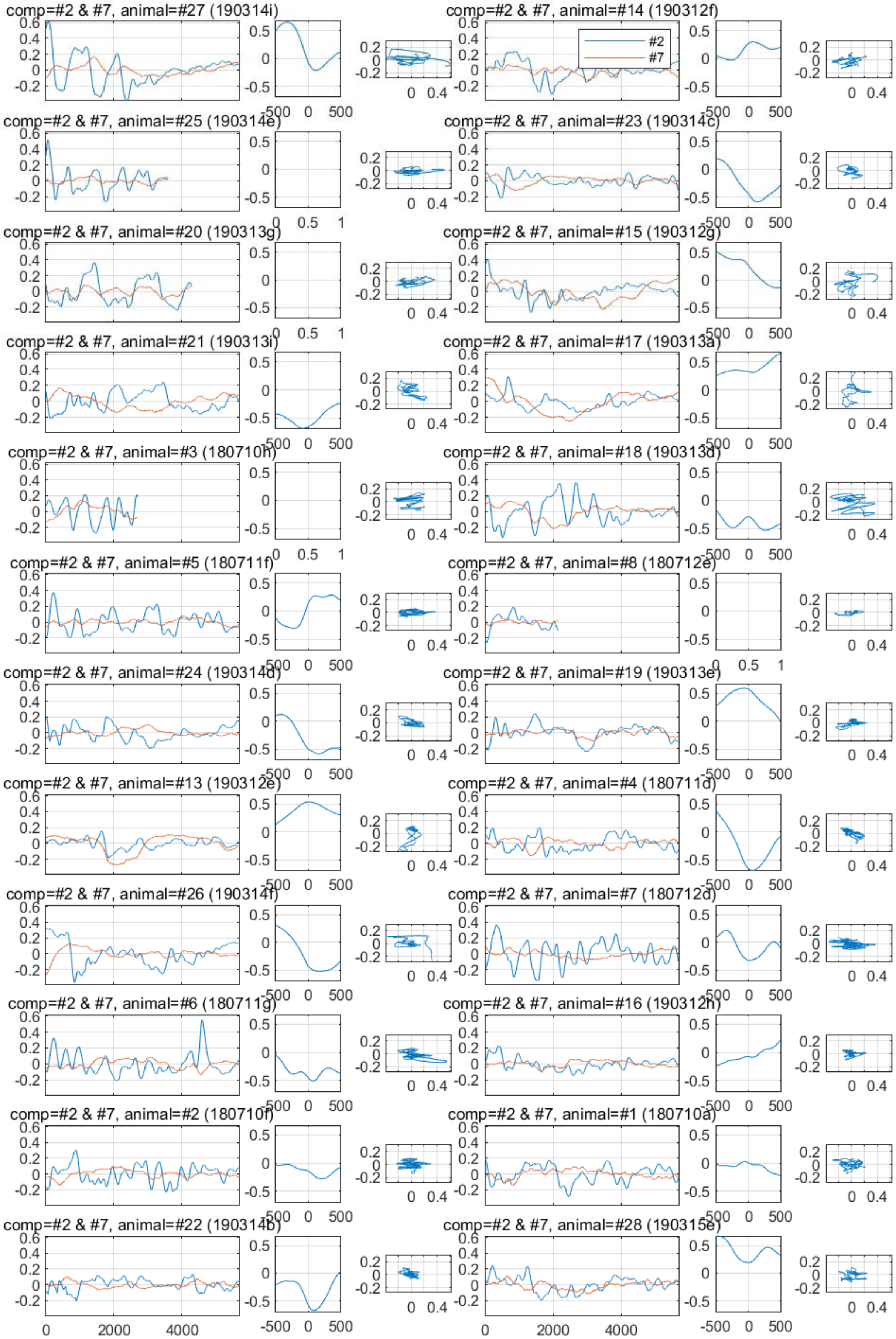
comp=#2 & #7, animal=#27 (190314i)
comp=#2 & #7, animal=#14 (190312f)
#2
#7
comp=#2 & #7, animal=#25 (190314e)
comp=#2 & #7, animal=#23 (190314c)
comp=#2 & #7, animal=#20 (190313g)
comp=#2 & #7, animal=#15 (190312g)
comp=#2 & #7, animal=#21 (190313i)
comp=#2 & #7, animal=#17 (190313a)
comp=#2 & #7, animal=#3 (180710h)
comp=#2 & #7, animal=#18 (190313d)
comp=#2 & #7, animal=#5 (180711f)
comp=#2 & #7, animal=#8 (180712e)
comp=#2 & #7, animal=#24 (190314d)
comp=#2 & #7, animal=#19 (190313e)
comp=#2 & #7, animal=#13 (190312e)
comp=#2 & #7, animal=#4 (180711d)
comp=#2 & #7, animal=#26 (190314f)
comp=#2 & #7, animal=#7 (180712d)
comp=#2 & #7, animal=#6 (180711g)
comp=#2 & #7, animal=#16 (190312h)
comp=#2 & #7, animal=#2 (180710f)
comp=#2 & #7, animal=#1 (180710a)
comp=#2 & #7, animal=#22 (190314b)
comp=#2 & #7, animal=#28 (190315e)

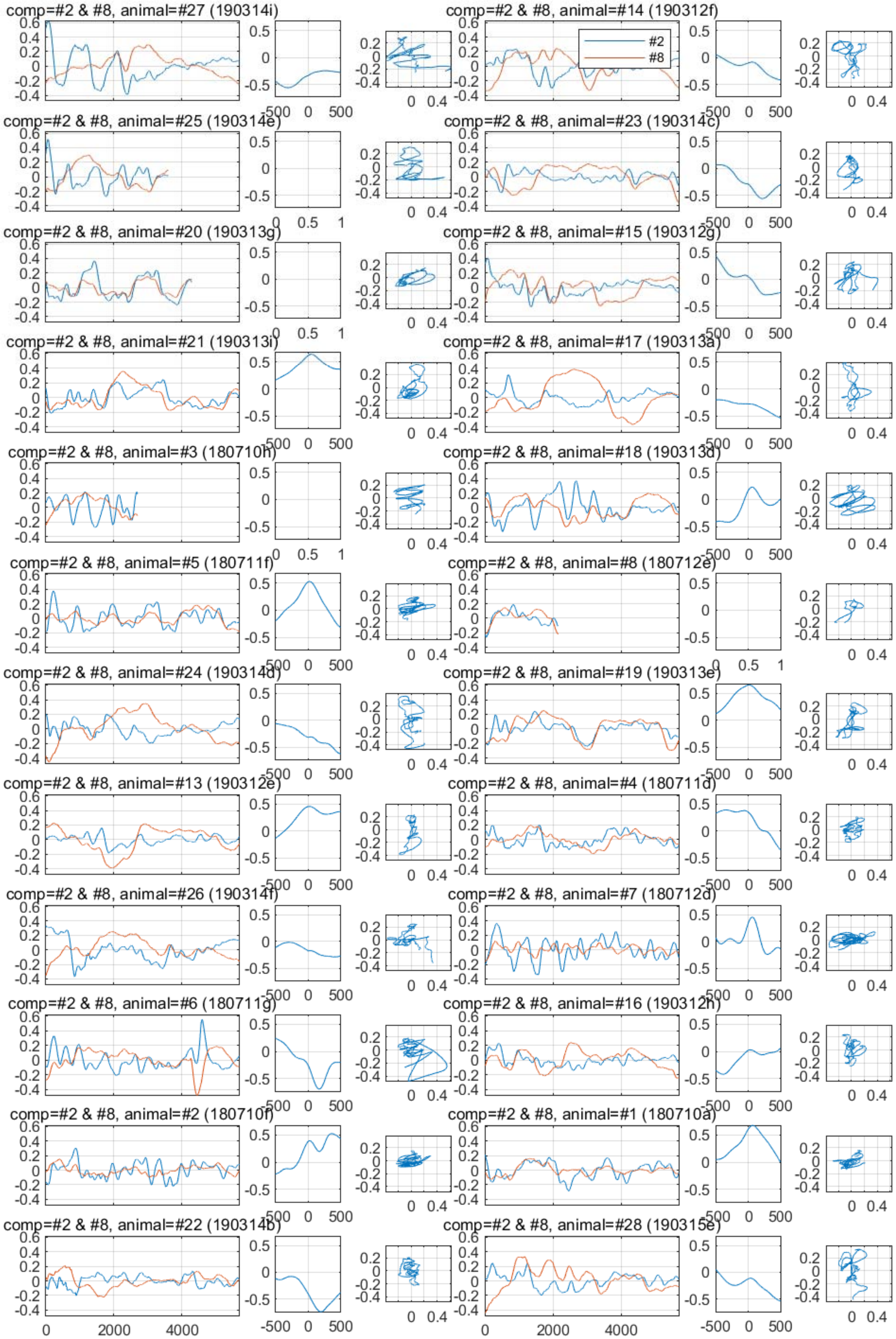

comp=#2 & #8, animal=#27 (190314i)
comp=#2 & #8, animal=#14 (190312f)
#2
#8
comp=#2 & #8, animal=#25 (190314e)
comp=#2 & #8, animal=#23 (190314c)
comp=#2 & #8, animal=#20 (190313g)
comp=#2 & #8, animal=#15 (190312g)
comp=#2 & #8, animal=#21 (190313i)
comp=#2 & #8, animal=#17 (190313a)
comp=#2 & #8, animal=#3 (180710h)
comp=#2 & #8, animal=#18 (190313d)
comp=#2 & #8, animal=#5 (180711f)
comp=#2 & #8, animal=#8 (180712e)
comp=#2 & #8, animal=#24 (190314d)
comp=#2 & #8, animal=#19 (190313e)
comp=#2 & #8, animal=#13 (190312e)
comp=#2 & #8, animal=#4 (180711d)
comp=#2 & #8, animal=#26 (190314f)
comp=#2 & #8, animal=#7 (180712d)
comp=#2 & #8, animal=#6 (180711g)
comp=#2 & #8, animal=#16 (190312h)
comp=#2 & #8, animal=#2 (180710f)
comp=#2 & #8, animal=#1 (180710a)
comp=#2 & #8, animal=#22 (190314b)
comp=#2 & #8, animal=#28 (190315e)

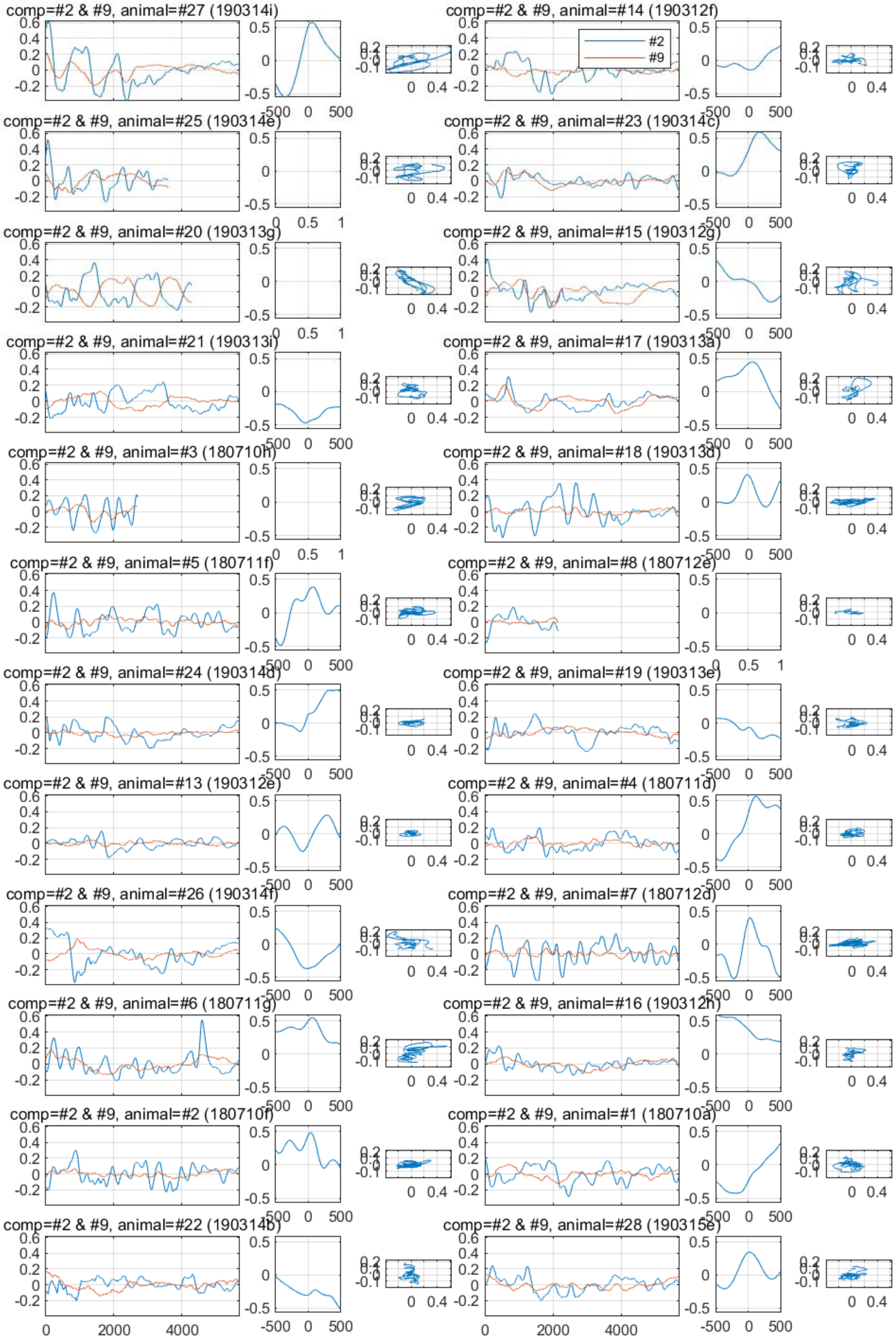
comp=#2 & #9, animal=#27 (190314i)
comp=#2 & #9, animal=#14 (190312f)
#2
#9
comp=#2 & #9, animal=#25 (190314e)
comp=#2 & #9, animal=#23 (190314c)
comp=#2 & #9, animal=#20 (190313g)
comp=#2 & #9, animal=#15 (190312g)
comp=#2 & #9, animal=#21 (190313i)
comp=#2 & #9, animal=#17 (190313a)
comp=#2 & #9, animal=#3 (180710h)
comp=#2 & #9, animal=#18 (190313d)
comp=#2 & #9, animal=#5 (180711f)
comp=#2 & #9, animal=#8 (180712e)
comp=#2 & #9, animal=#24 (190314d)
comp=#2 & #9, animal=#19 (190313e)
comp=#2 & #9, animal=#13 (190312e)
comp=#2 & #9, animal=#4 (180711d)
comp=#2 & #9, animal=#26 (190314f)
comp=#2 & #9, animal=#7 (180712d)
comp=#2 & #9, animal=#6 (180711g)
comp=#2 & #9, animal=#16 (190312h)
comp=#2 & #9, animal=#2 (180710f)
comp=#2 & #9, animal=#1 (180710a)
comp=#2 & #9, animal=#22 (190314b)
comp=#2 & #9, animal=#28 (190315e)
0
2000
4000
-500
0
500

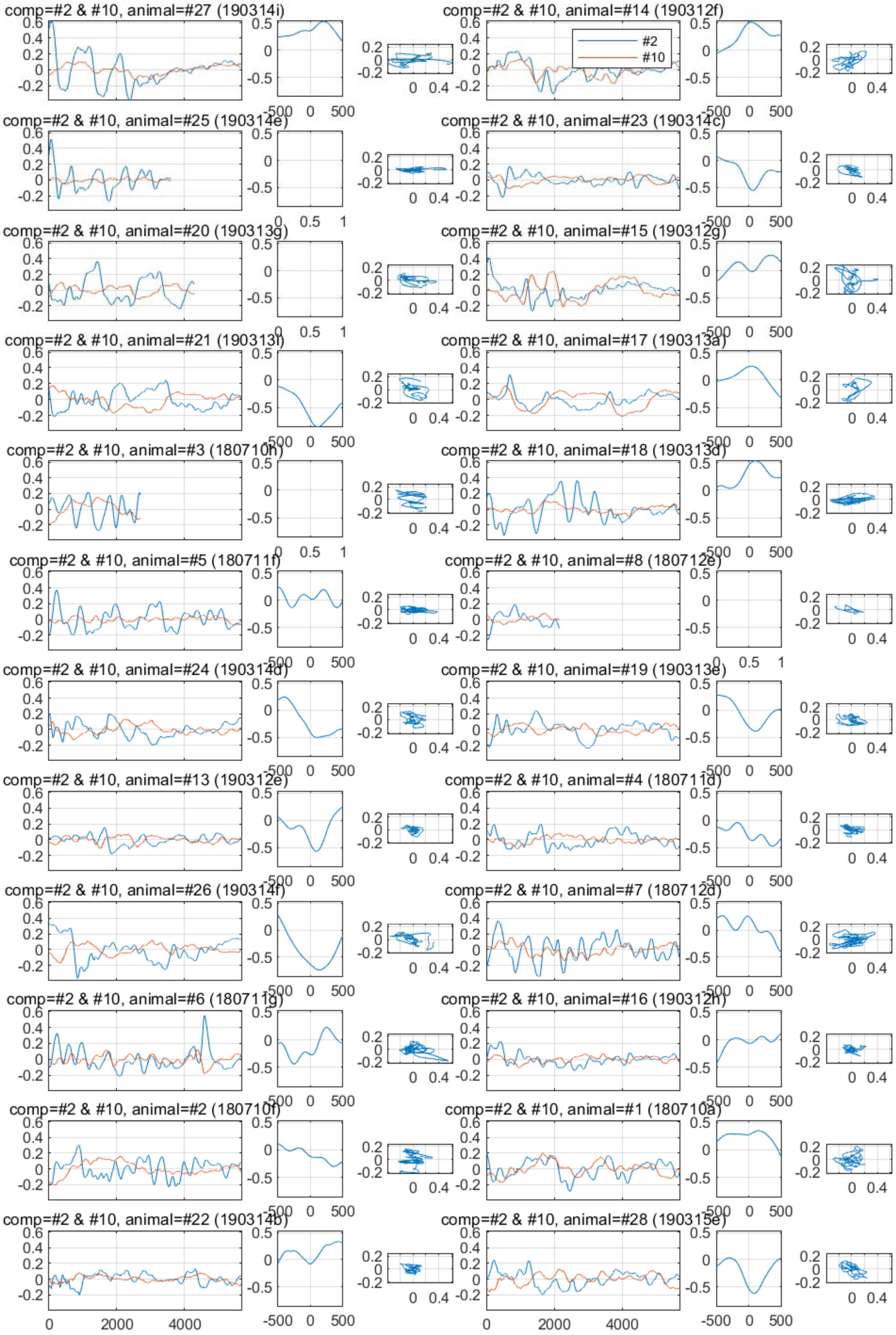
comp=#2 & #10, animal=#27 (190314i)
comp=#2 & #10, animal=#14 (190312f)
#2
#10
comp=#2 & #10, animal=#25 (190314e)
comp=#2 & #10, animal=#23 (190314c)
comp=#2 & #10, animal=#20 (190313g)
comp=#2 & #10, animal=#15 (190312g)
comp=#2 & #10, animal=#21 (190313i)
comp=#2 & #10, animal=#17 (190313a)
comp=#2 & #10, animal=#3 (180710h)
comp=#2 & #10, animal=#18 (190313d)
comp=#2 & #10, animal=#5 (180711f)
comp=#2 & #10, animal=#8 (180712e)
comp=#2 & #10, animal=#24 (190314d)
comp=#2 & #10, animal=#19 (190313e)
comp=#2 & #10, animal=#13 (190312e)
comp=#2 & #10, animal=#4 (180711d)
comp=#2 & #10, animal=#26 (190314f)
comp=#2 & #10, animal=#7 (180712d)
comp=#2 & #10, animal=#6 (180711g)
comp=#2 & #10, animal=#16 (190312h)
comp=#2 & #10, animal=#2 (180710f)
comp=#2 & #10, animal=#1 (180710a)
comp=#2 & #10, animal=#22 (190314b)
comp=#2 & #10, animal=#28 (190315e)
0
2000
4000
-500
0
500

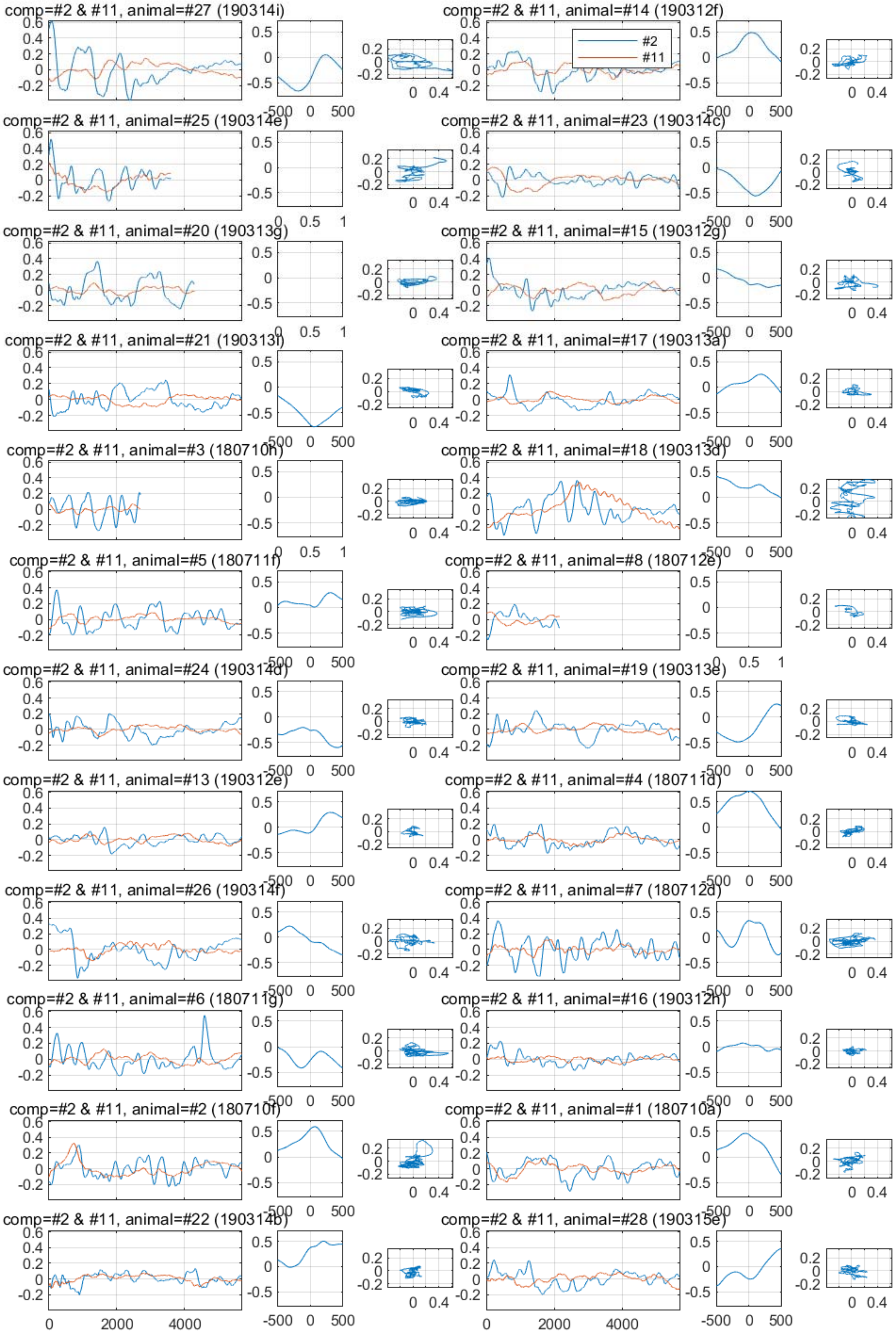
comp=#2 & #11, animal=#27 (190314i)
comp=#2 & #11, animal=#14 (190312f)
#2
#11
comp=#2 & #11, animal=#25 (190314e)
comp=#2 & #11, animal=#23 (190314c)
comp=#2 & #11, animal=#20 (190313g)
comp=#2 & #11, animal=#15 (190312g)
comp=#2 & #11, animal=#21 (190313i)
comp=#2 & #11, animal=#17 (190313a)
comp=#2 & #11, animal=#3 (180710h)
comp=#2 & #11, animal=#18 (190313d)
comp=#2 & #11, animal=#5 (180711f)
comp=#2 & #11, animal=#8 (180712e)
comp=#2 & #11, animal=#24 (190314d)
comp=#2 & #11, animal=#19 (190313e)
comp=#2 & #11, animal=#13 (190312e)
comp=#2 & #11, animal=#4 (180711d)
comp=#2 & #11, animal=#26 (190314f)
comp=#2 & #11, animal=#7 (180712d)
comp=#2 & #11, animal=#6 (180711g)
comp=#2 & #11, animal=#16 (190312h)
comp=#2 & #11, animal=#2 (180710f)
comp=#2 & #11, animal=#1 (180710a)
comp=#2 & #11, animal=#22 (190314b)
comp=#2 & #11, animal=#28 (190315e)

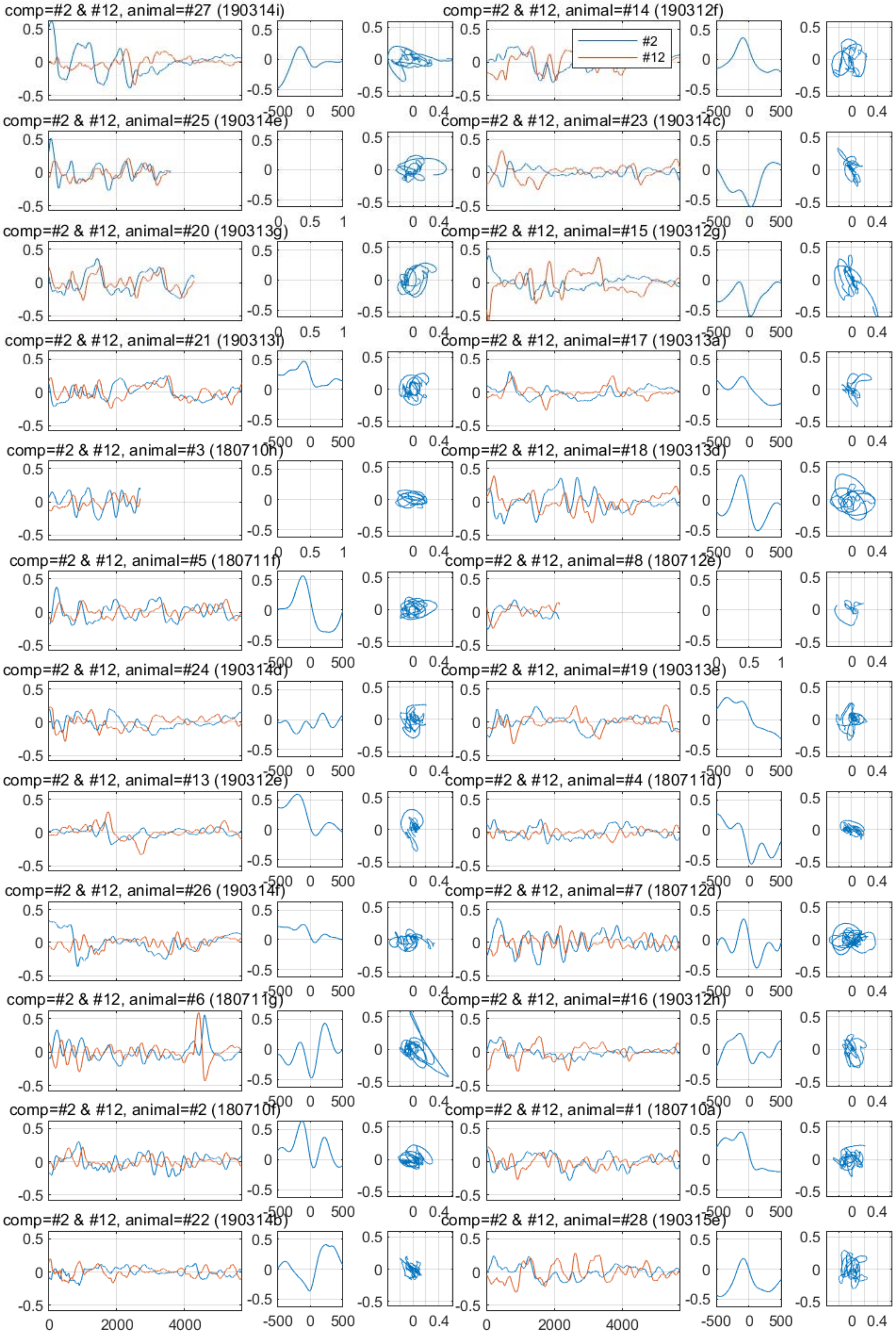
comp=#2 & #12, animal=#27 (190314i)
comp=#2 & #12, animal=#14 (190312f)
#2
#12
comp=#2 & #12, animal=#25 (190314e)
comp=#2 & #12, animal=#23 (190314c)
comp=#2 & #12, animal=#20 (190313g)
comp=#2 & #12, animal=#15 (190312g)
comp=#2 & #12, animal=#21 (190313i)
comp=#2 & #12, animal=#17 (190313a)
comp=#2 & #12, animal=#3 (180710h)
comp=#2 & #12, animal=#18 (190313d)
comp=#2 & #12, animal=#5 (180711f)
comp=#2 & #12, animal=#8 (180712e)
comp=#2 & #12, animal=#24 (190314d)
comp=#2 & #12, animal=#19 (190313e)
comp=#2 & #12, animal=#13 (190312e)
comp=#2 & #12, animal=#4 (180711d)
comp=#2 & #12, animal=#26 (190314f)
comp=#2 & #12, animal=#7 (180712d)
comp=#2 & #12, animal=#6 (180711g)
comp=#2 & #12, animal=#16 (190312h)
comp=#2 & #12, animal=#2 (180710f)
comp=#2 & #12, animal=#1 (180710a)
comp=#2 & #12, animal=#22 (190314b)
comp=#2 & #12, animal=#28 (190315e)

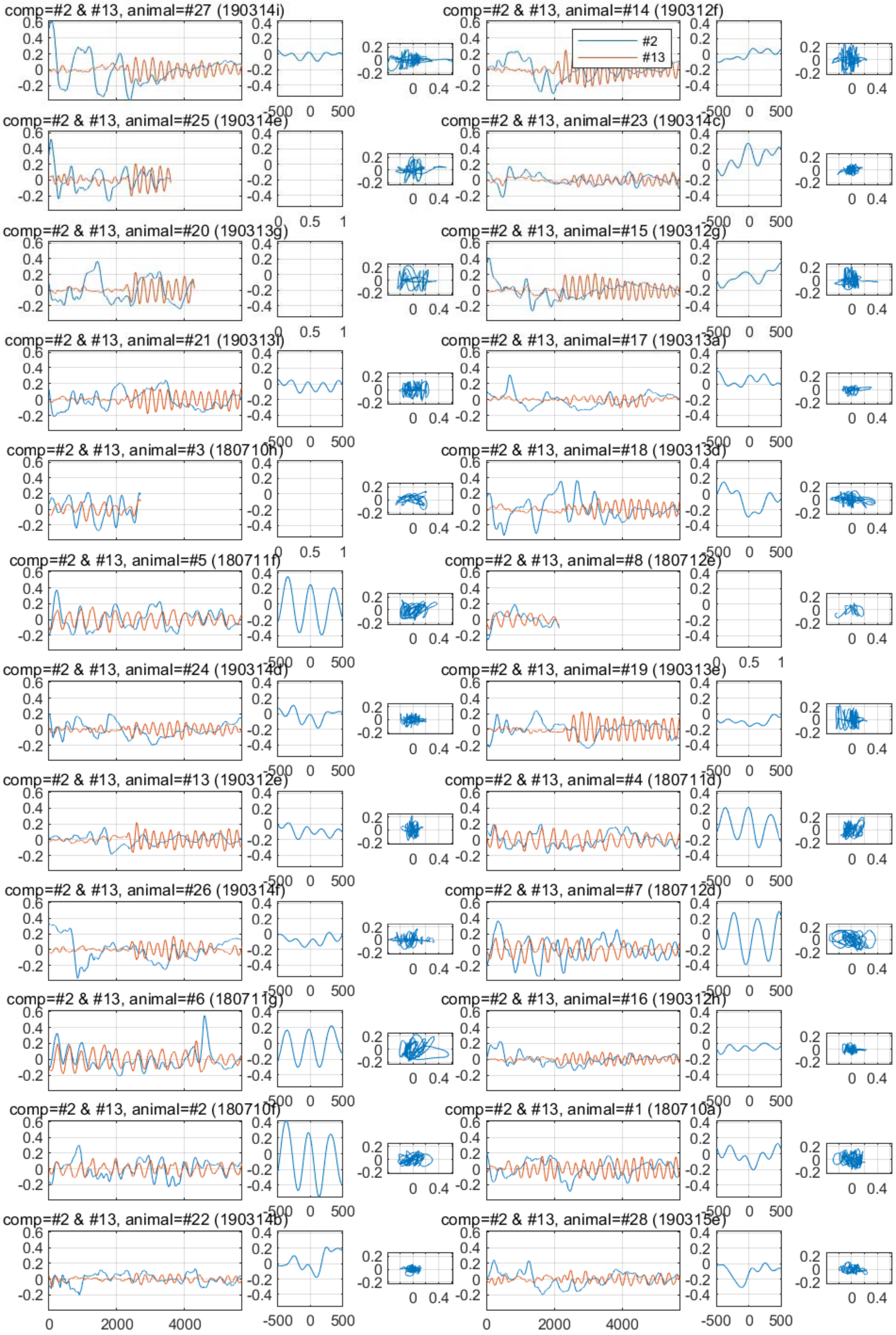

#2
#13
comp=#2 & #13, animal=#27 (190314i)
comp=#2 & #13, animal=#25 (190314e)
comp=#2 & #13, animal=#20 (190313g)
comp=#2 & #13, animal=#21 (190313i)
comp=#2 & #13, animal=#3 (180710h)
comp=#2 & #13, animal=#5 (180711f)
comp=#2 & #13, animal=#24 (190314d)
comp=#2 & #13, animal=#13 (190312e)
comp=#2 & #13, animal=#26 (190314f)
comp=#2 & #13, animal=#6 (180711g)
comp=#2 & #13, animal=#2 (180710f)
comp=#2 & #13, animal=#22 (190314b)
comp=#2 & #13, animal=#14 (190312f)
comp=#2 & #13, animal=#23 (190314c)
comp=#2 & #13, animal=#15 (190312g)
comp=#2 & #13, animal=#17 (190313a)
comp=#2 & #13, animal=#18 (190313d)
comp=#2 & #13, animal=#8 (180712e)
comp=#2 & #13, animal=#19 (190313e)
comp=#2 & #13, animal=#4 (180711d)
comp=#2 & #13, animal=#7 (180712d)
comp=#2 & #13, animal=#16 (190312h)
comp=#2 & #13, animal=#1 (180710a)
comp=#2 & #13, animal=#28 (190315e)

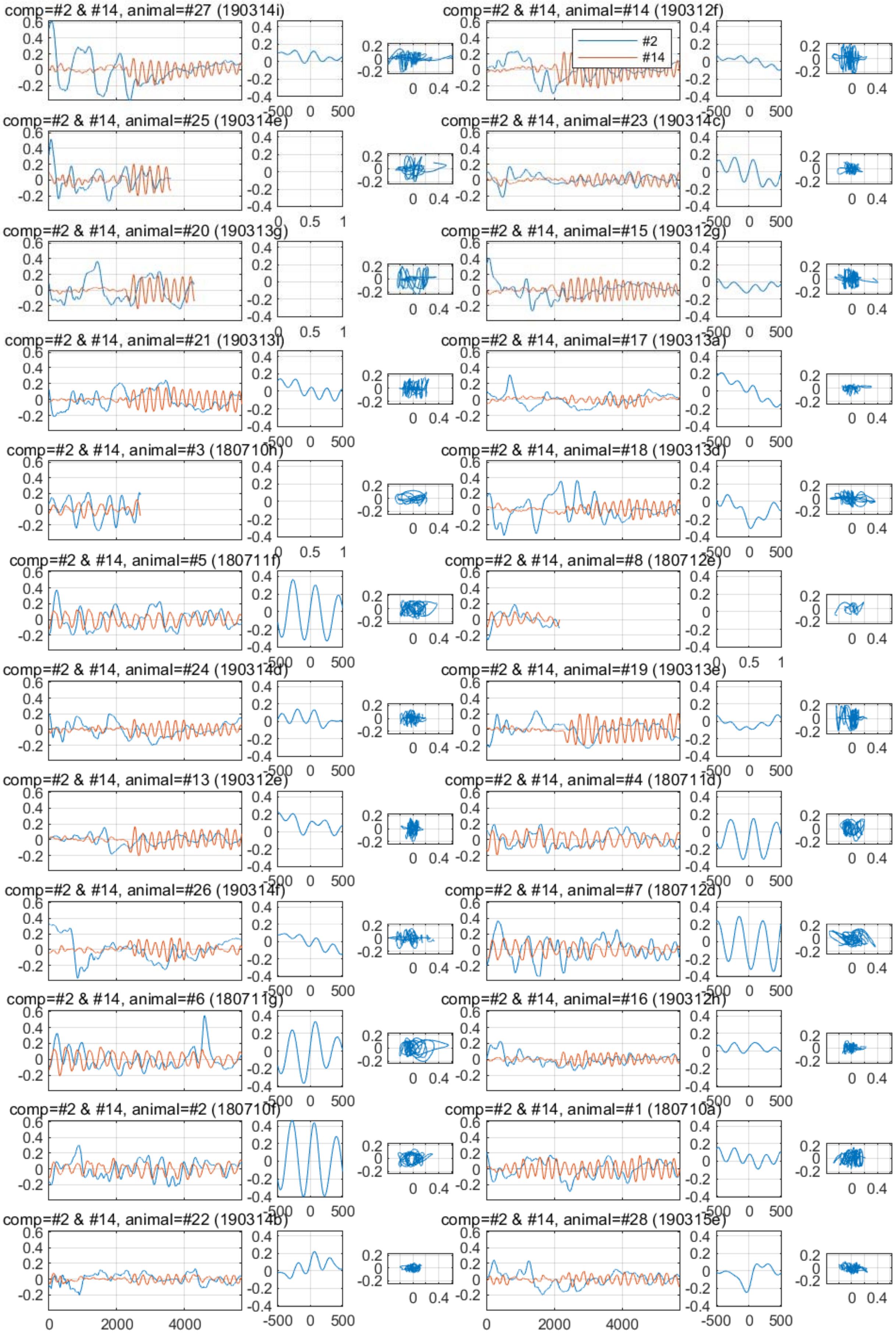
comp=#2 & #14, animal=#27 (190314i)
comp=#2 & #14, animal=#14 (190312f)
#2
#14
comp=#2 & #14, animal=#25 (190314e)
comp=#2 & #14, animal=#23 (190314c)
comp=#2 & #14, animal=#20 (190313g)
comp=#2 & #14, animal=#15 (190312g)
comp=#2 & #14, animal=#21 (190313i)
comp=#2 & #14, animal=#17 (190313a)
comp=#2 & #14, animal=#3 (180710h)
comp=#2 & #14, animal=#18 (190313d)
comp=#2 & #14, animal=#5 (180711f)
comp=#2 & #14, animal=#8 (180712e)
comp=#2 & #14, animal=#24 (190314d)
comp=#2 & #14, animal=#19 (190313e)
comp=#2 & #14, animal=#13 (190312e)
comp=#2 & #14, animal=#4 (180711d)
comp=#2 & #14, animal=#26 (190314f)
comp=#2 & #14, animal=#7 (180712d)
comp=#2 & #14, animal=#6 (180711g)
comp=#2 & #14, animal=#16 (190312h)
comp=#2 & #14, animal=#2 (180710f)
comp=#2 & #14, animal=#1 (180710a)
comp=#2 & #14, animal=#22 (190314b)
comp=#2 & #14, animal=#28 (190315e)

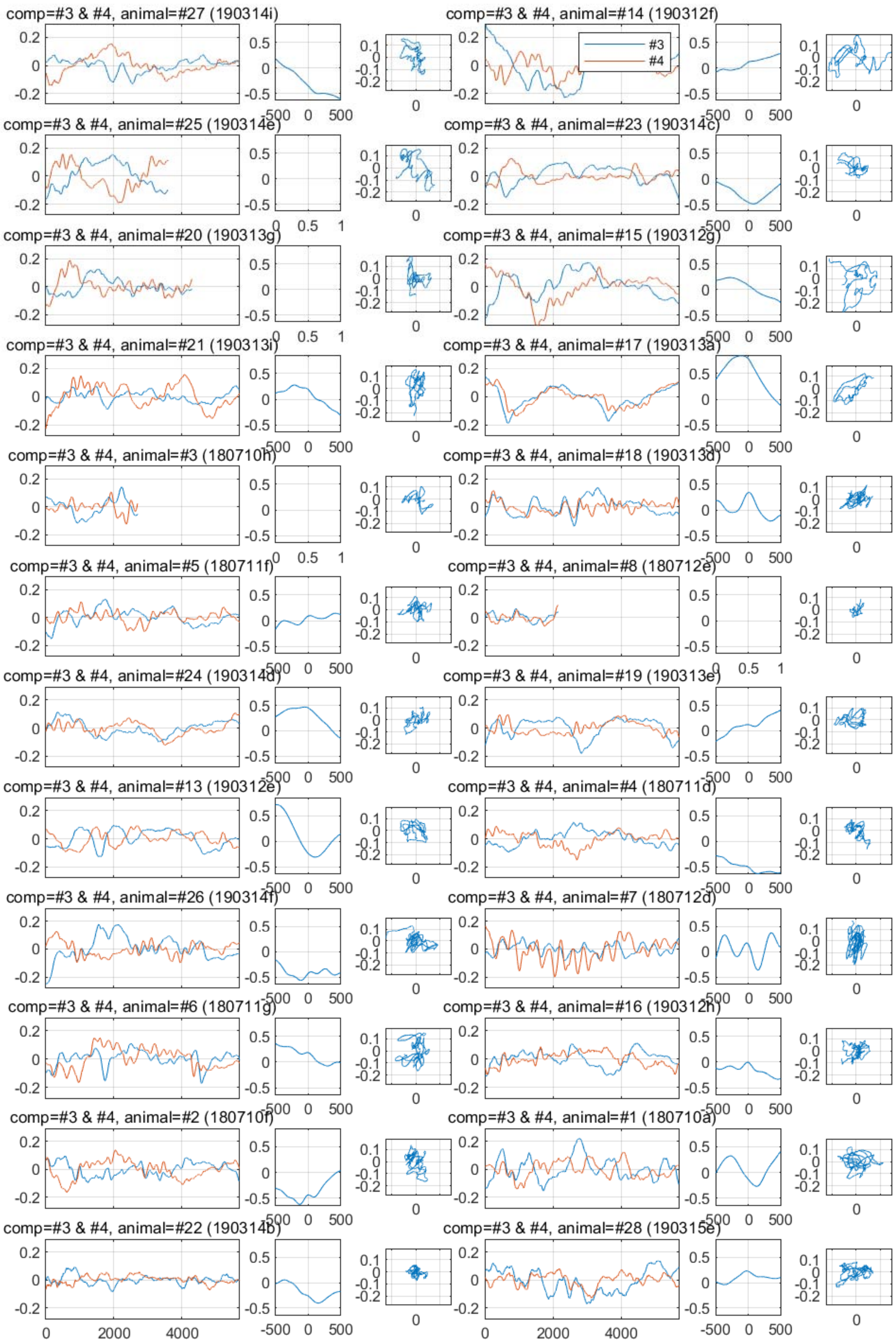
comp=#3 & #4, animal=#27 (190314i)
comp=#3 & #4, animal=#14 (190312f)
#3
#4
comp=#3 & #4, animal=#25 (190314e)
comp=#3 & #4, animal=#23 (190314c)
comp=#3 & #4, animal=#20 (190313g)
comp=#3 & #4, animal=#15 (190312g)
comp=#3 & #4, animal=#21 (190313i)
comp=#3 & #4, animal=#17 (190313a)
comp=#3 & #4, animal=#3 (180710h)
comp=#3 & #4, animal=#18 (190313d)
comp=#3 & #4, animal=#5 (180711f)
comp=#3 & #4, animal=#8 (180712e)
comp=#3 & #4, animal=#24 (190314d)
comp=#3 & #4, animal=#19 (190313e)
comp=#3 & #4, animal=#13 (190312e)
comp=#3 & #4, animal=#4 (180711d)
comp=#3 & #4, animal=#26 (190314f)
comp=#3 & #4, animal=#7 (180712d)
comp=#3 & #4, animal=#6 (180711g)
comp=#3 & #4, animal=#16 (190312h)
comp=#3 & #4, animal=#2 (180710f)
comp=#3 & #4, animal=#1 (180710a)
comp=#3 & #4, animal=#22 (190314b)
comp=#3 & #4, animal=#28 (190315e)
0
2000
4000
-500
0
500

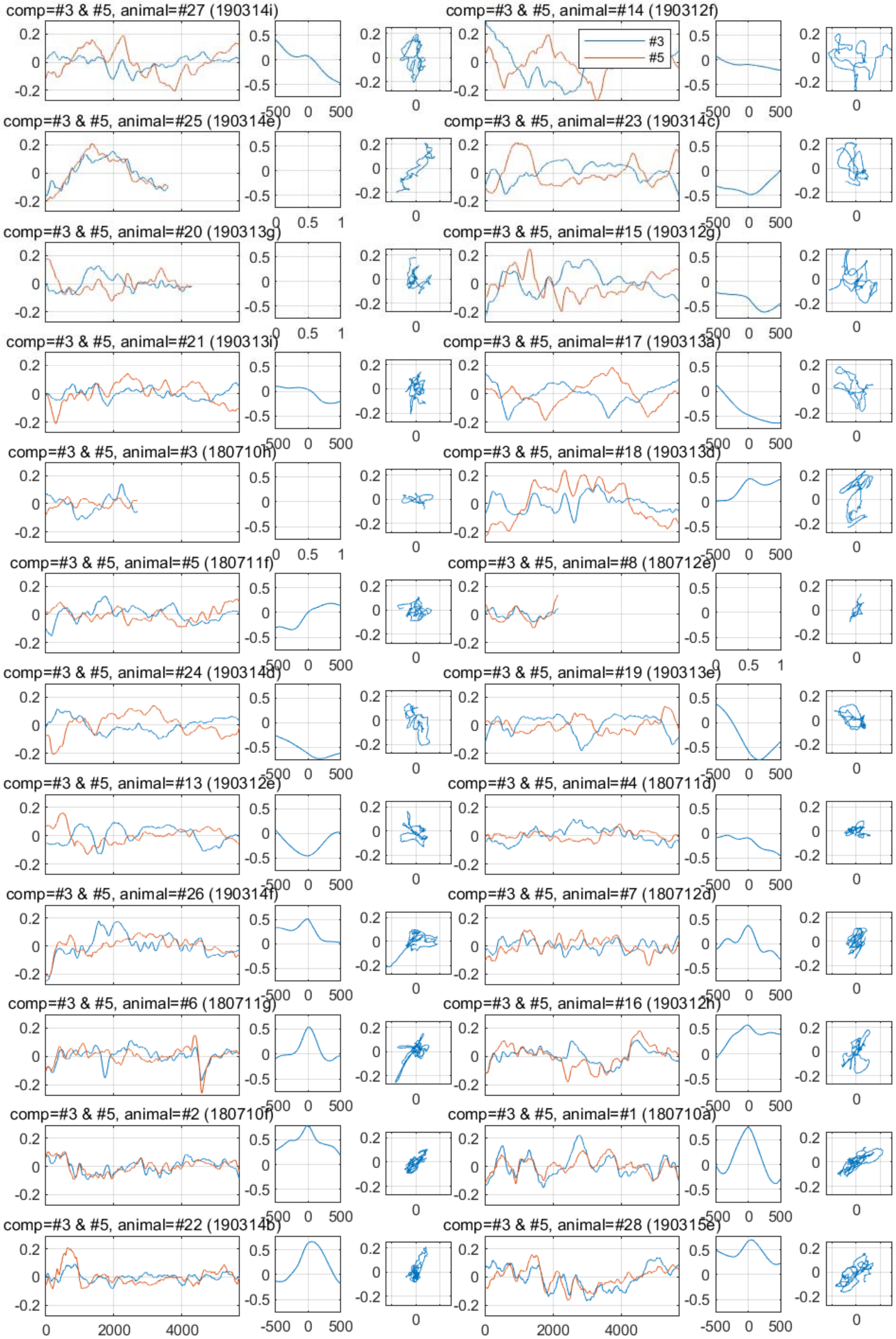
comp=#3 & #5, animal=#27 (190314i)
comp=#3 & #5, animal=#14 (190312f)
#3
#5
comp=#3 & #5, animal=#25 (190314e)
comp=#3 & #5, animal=#23 (190314c)
comp=#3 & #5, animal=#20 (190313g)
comp=#3 & #5, animal=#15 (190312g)
comp=#3 & #5, animal=#21 (190313i)
comp=#3 & #5, animal=#17 (190313a)
comp=#3 & #5, animal=#3 (180710h)
comp=#3 & #5, animal=#18 (190313d)
comp=#3 & #5, animal=#5 (180711f)
comp=#3 & #5, animal=#8 (180712e)
comp=#3 & #5, animal=#24 (190314d)
comp=#3 & #5, animal=#19 (190313e)
comp=#3 & #5, animal=#13 (190312e)
comp=#3 & #5, animal=#4 (180711d)
comp=#3 & #5, animal=#26 (190314f)
comp=#3 & #5, animal=#7 (180712d)
comp=#3 & #5, animal=#6 (180711g)
comp=#3 & #5, animal=#16 (190312h)
comp=#3 & #5, animal=#2 (180710f)
comp=#3 & #5, animal=#1 (180710a)
comp=#3 & #5, animal=#22 (190314b)
comp=#3 & #5, animal=#28 (190315e)

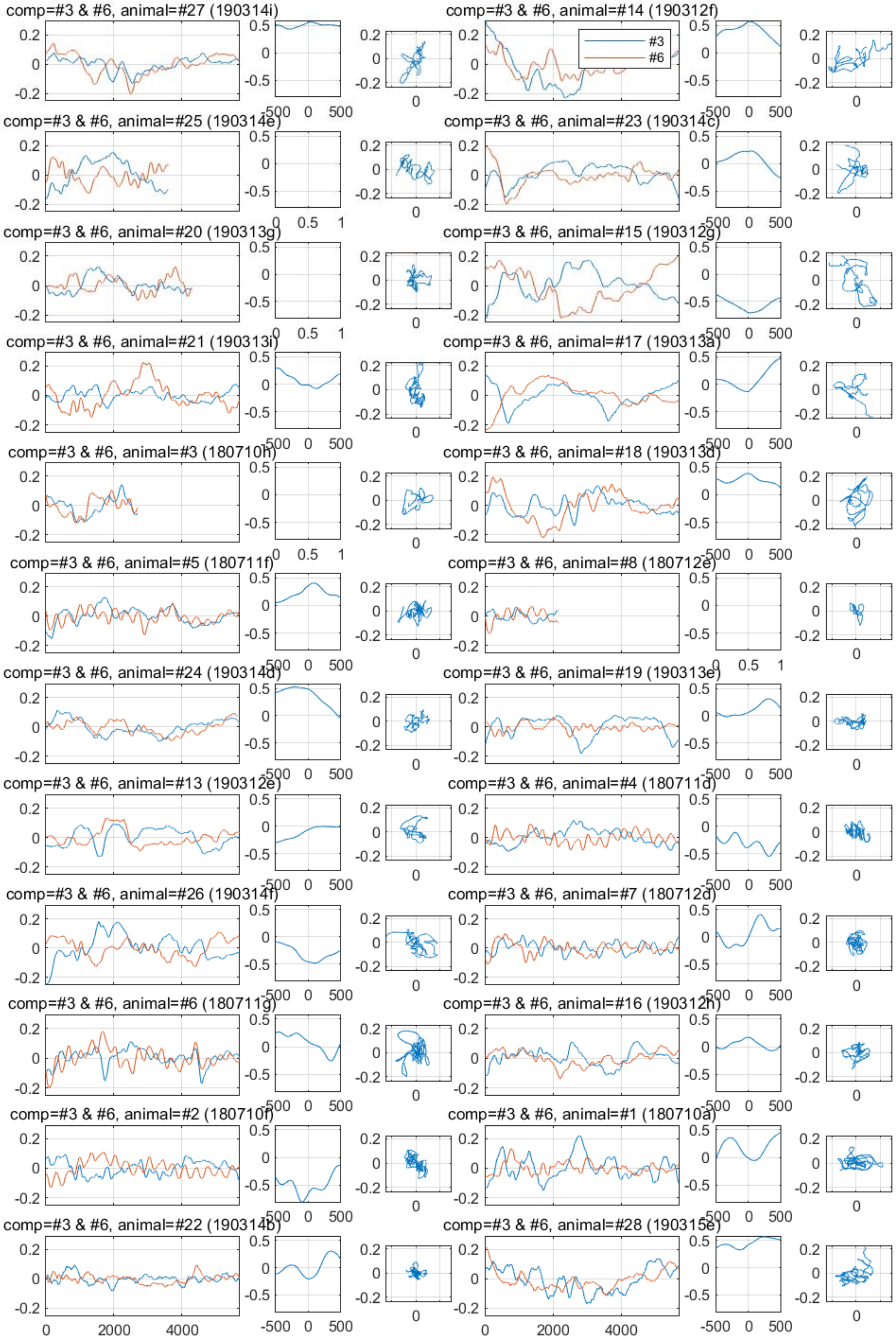
comp=#3 & #6, animal=#27 (190314i)
comp=#3 & #6, animal=#14 (190312f)
#3
#6
comp=#3 & #6, animal=#25 (190314e)
comp=#3 & #6, animal=#23 (190314c)
comp=#3 & #6, animal=#20 (190313g)
comp=#3 & #6, animal=#15 (190312g)
comp=#3 & #6, animal=#21 (190313i)
comp=#3 & #6, animal=#17 (190313a)
comp=#3 & #6, animal=#3 (180710h)
comp=#3 & #6, animal=#18 (190313d)
comp=#3 & #6, animal=#5 (180711f)
comp=#3 & #6, animal=#8 (180712e)
comp=#3 & #6, animal=#24 (190314d)
comp=#3 & #6, animal=#19 (190313e)
comp=#3 & #6, animal=#13 (190312e)
comp=#3 & #6, animal=#4 (180711d)
comp=#3 & #6, animal=#26 (190314f)
comp=#3 & #6, animal=#7 (180712d)
comp=#3 & #6, animal=#6 (180711g)
comp=#3 & #6, animal=#16 (190312h)
comp=#3 & #6, animal=#2 (180710f)
comp=#3 & #6, animal=#1 (180710a)
comp=#3 & #6, animal=#22 (190314b)
comp=#3 & #6, animal=#28 (190315e)
0
2000
4000
-500
500

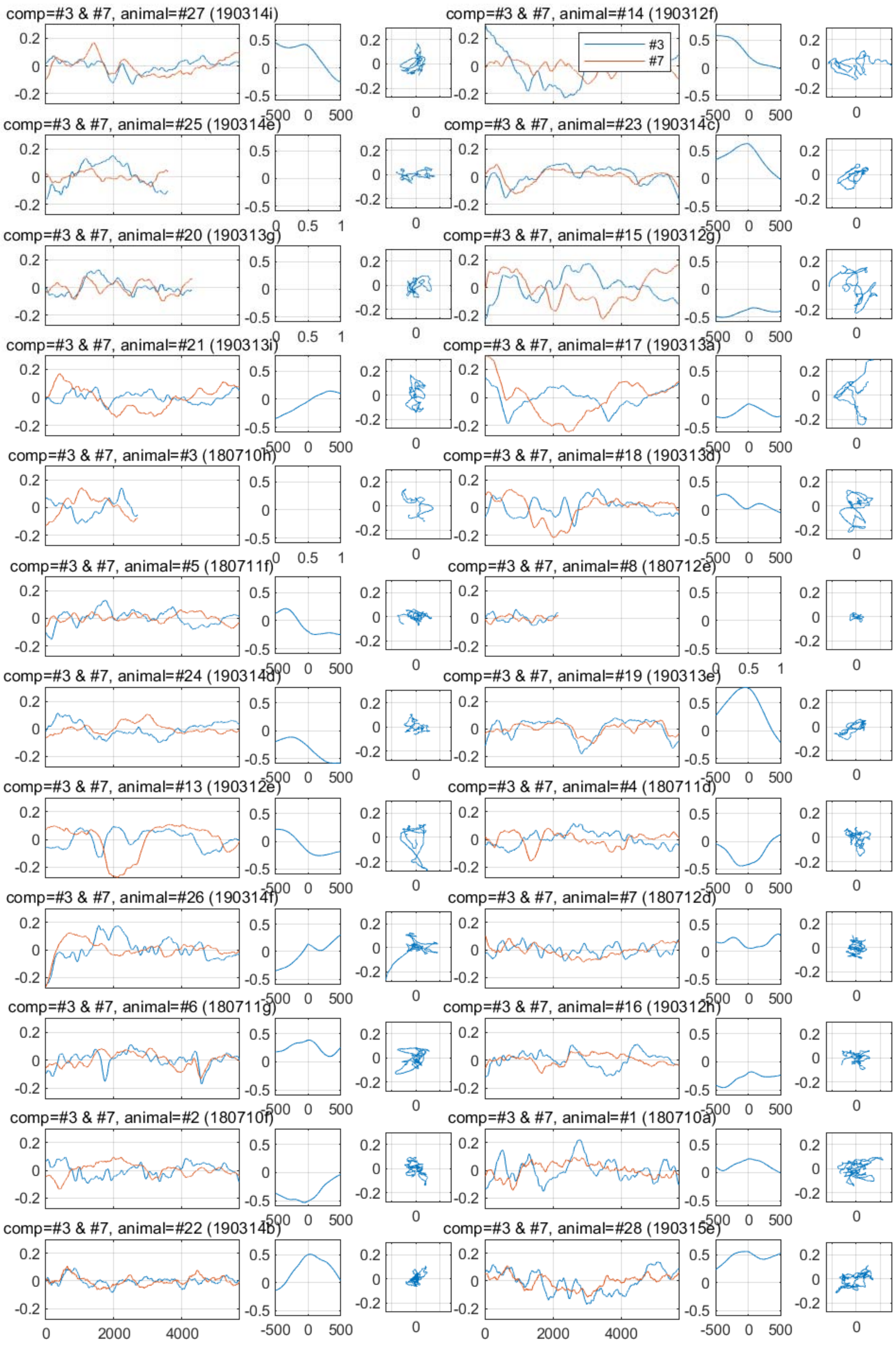
comp=#3 & #7, animal=#27 (190314i)
comp=#3 & #7, animal=#14 (190312f)
#3
#7
comp=#3 & #7, animal=#25 (190314e)
comp=#3 & #7, animal=#23 (190314c)
comp=#3 & #7, animal=#20 (190313g)
comp=#3 & #7, animal=#15 (190312g)
comp=#3 & #7, animal=#21 (190313i)
comp=#3 & #7, animal=#17 (190313a)
comp=#3 & #7, animal=#3 (180710h)
comp=#3 & #7, animal=#18 (190313d)
comp=#3 & #7, animal=#5 (180711f)
comp=#3 & #7, animal=#8 (180712e)
comp=#3 & #7, animal=#24 (190314d)
comp=#3 & #7, animal=#19 (190313e)
comp=#3 & #7, animal=#13 (190312e)
comp=#3 & #7, animal=#4 (180711d)
comp=#3 & #7, animal=#26 (190314f)
comp=#3 & #7, animal=#7 (180712d)
comp=#3 & #7, animal=#6 (180711g)
comp=#3 & #7, animal=#16 (190312h)
comp=#3 & #7, animal=#2 (180710f)
comp=#3 & #7, animal=#1 (180710a)
comp=#3 & #7, animal=#22 (190314b)
comp=#3 & #7, animal=#28 (190315e)

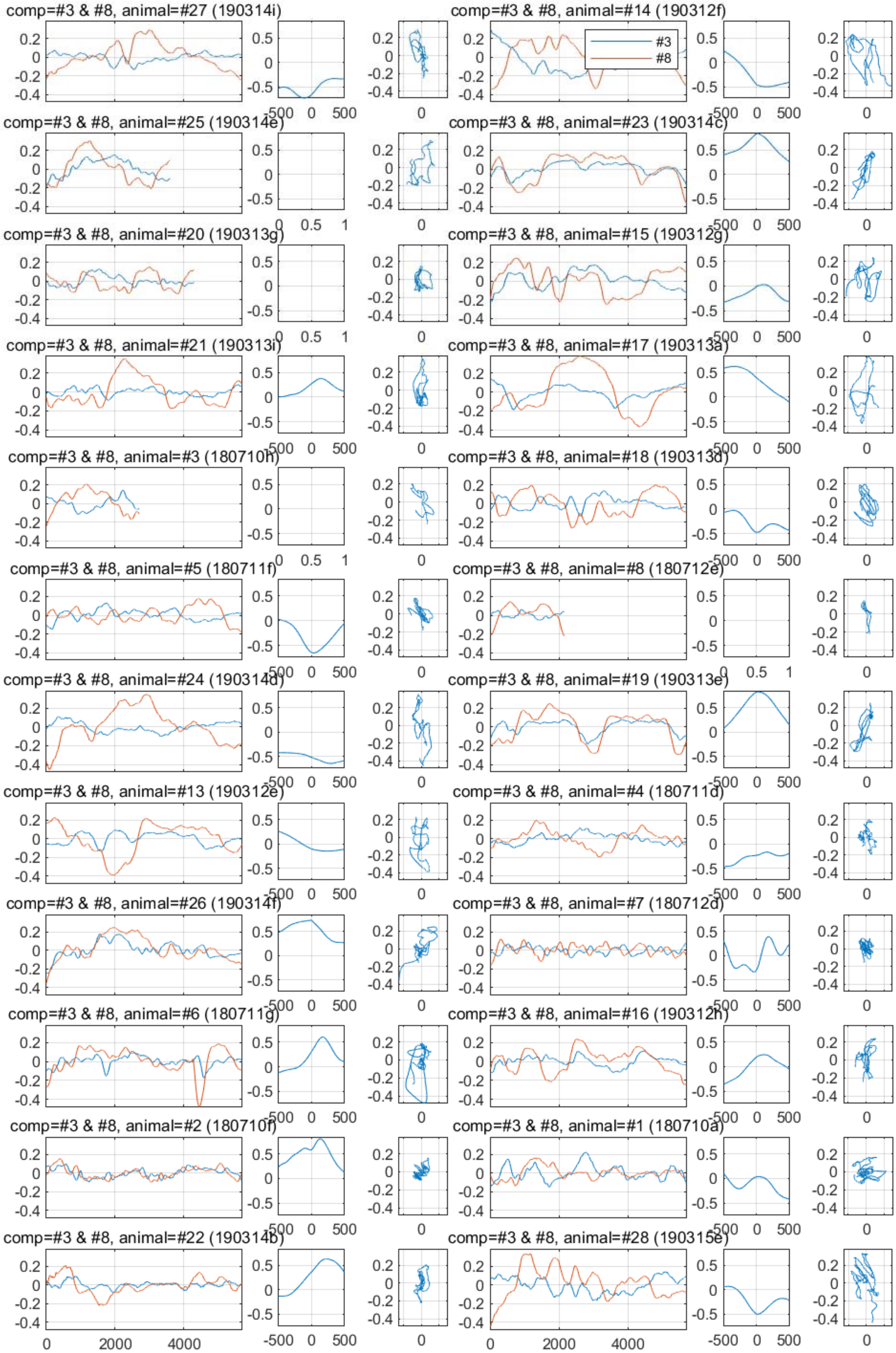

comp=#3 & #8, animal=#27 (190314i)
comp=#3 & #8, animal=#14 (190312f)
#3
#8
comp=#3 & #8, animal=#25 (190314e)
comp=#3 & #8, animal=#23 (190314c)
comp=#3 & #8, animal=#20 (190313g)
comp=#3 & #8, animal=#15 (190312g)
comp=#3 & #8, animal=#21 (190313i)
comp=#3 & #8, animal=#17 (190313a)
comp=#3 & #8, animal=#3 (180710h)
comp=#3 & #8, animal=#18 (190313d)
comp=#3 & #8, animal=#5 (180711f)
comp=#3 & #8, animal=#8 (180712e)
comp=#3 & #8, animal=#24 (190314d)
comp=#3 & #8, animal=#19 (190313e)
comp=#3 & #8, animal=#13 (190312e)
comp=#3 & #8, animal=#4 (180711d)
comp=#3 & #8, animal=#26 (190314f)
comp=#3 & #8, animal=#7 (180712d)
comp=#3 & #8, animal=#6 (180711g)
comp=#3 & #8, animal=#16 (190312h)
comp=#3 & #8, animal=#2 (180710f)
comp=#3 & #8, animal=#1 (180710a)
comp=#3 & #8, animal=#22 (190314b)
comp=#3 & #8, animal=#28 (190315e)

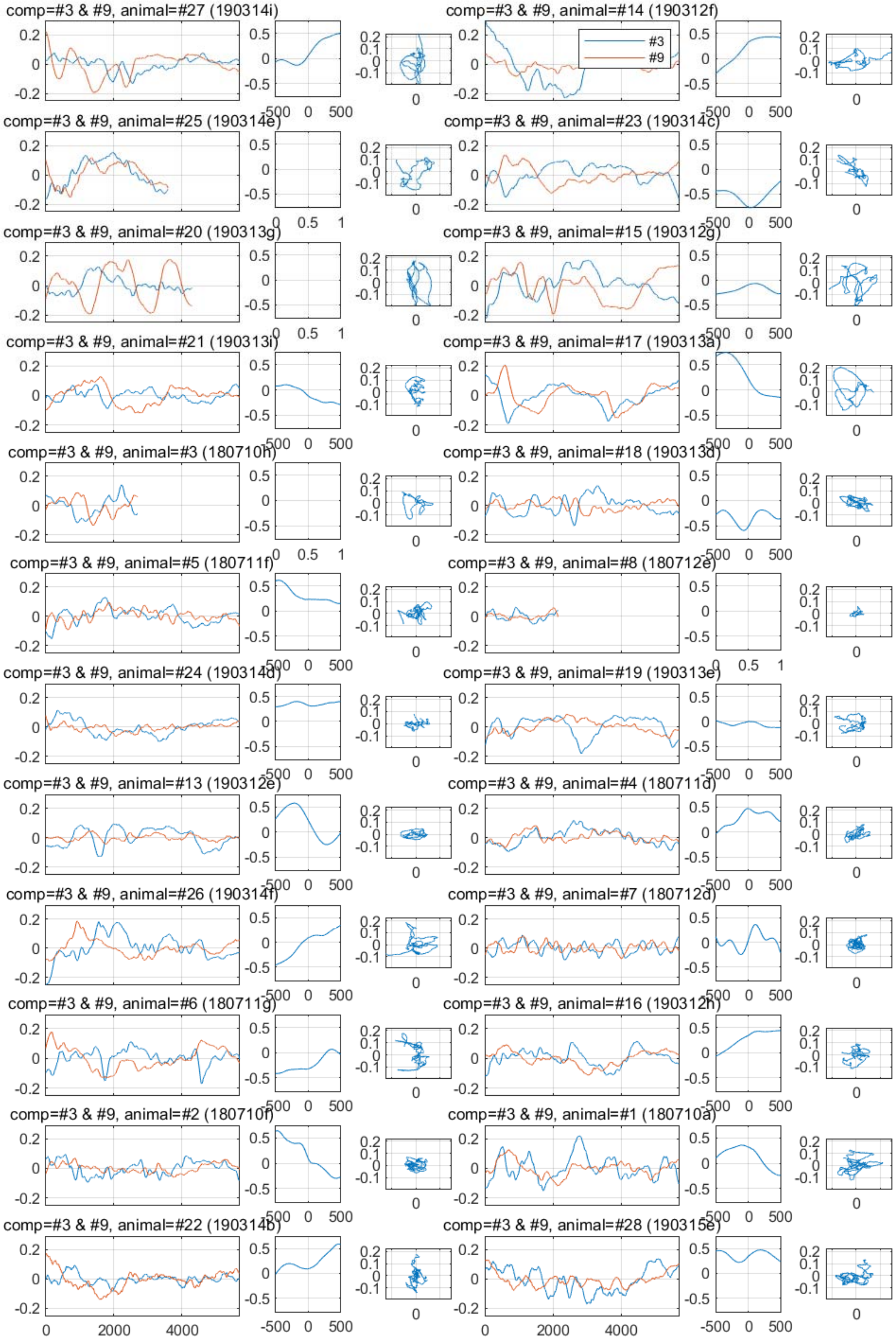
comp=#3 & #9, animal=#27 (190314i)
comp=#3 & #9, animal=#14 (190312f)
#3
#9
comp=#3 & #9, animal=#25 (190314e)
comp=#3 & #9, animal=#23 (190314c)
comp=#3 & #9, animal=#20 (190313g)
comp=#3 & #9, animal=#15 (190312g)
comp=#3 & #9, animal=#21 (190313i)
comp=#3 & #9, animal=#17 (190313a)
comp=#3 & #9, animal=#3 (180710h)
comp=#3 & #9, animal=#18 (190313d)
comp=#3 & #9, animal=#5 (180711f)
comp=#3 & #9, animal=#8 (180712e)
comp=#3 & #9, animal=#24 (190314d)
comp=#3 & #9, animal=#19 (190313e)
comp=#3 & #9, animal=#13 (190312e)
comp=#3 & #9, animal=#4 (180711d)
comp=#3 & #9, animal=#26 (190314f)
comp=#3 & #9, animal=#7 (180712d)
comp=#3 & #9, animal=#6 (180711g)
comp=#3 & #9, animal=#16 (190312h)
comp=#3 & #9, animal=#2 (180710f)
comp=#3 & #9, animal=#1 (180710a)
comp=#3 & #9, animal=#22 (190314b)
comp=#3 & #9, animal=#28 (190315e)

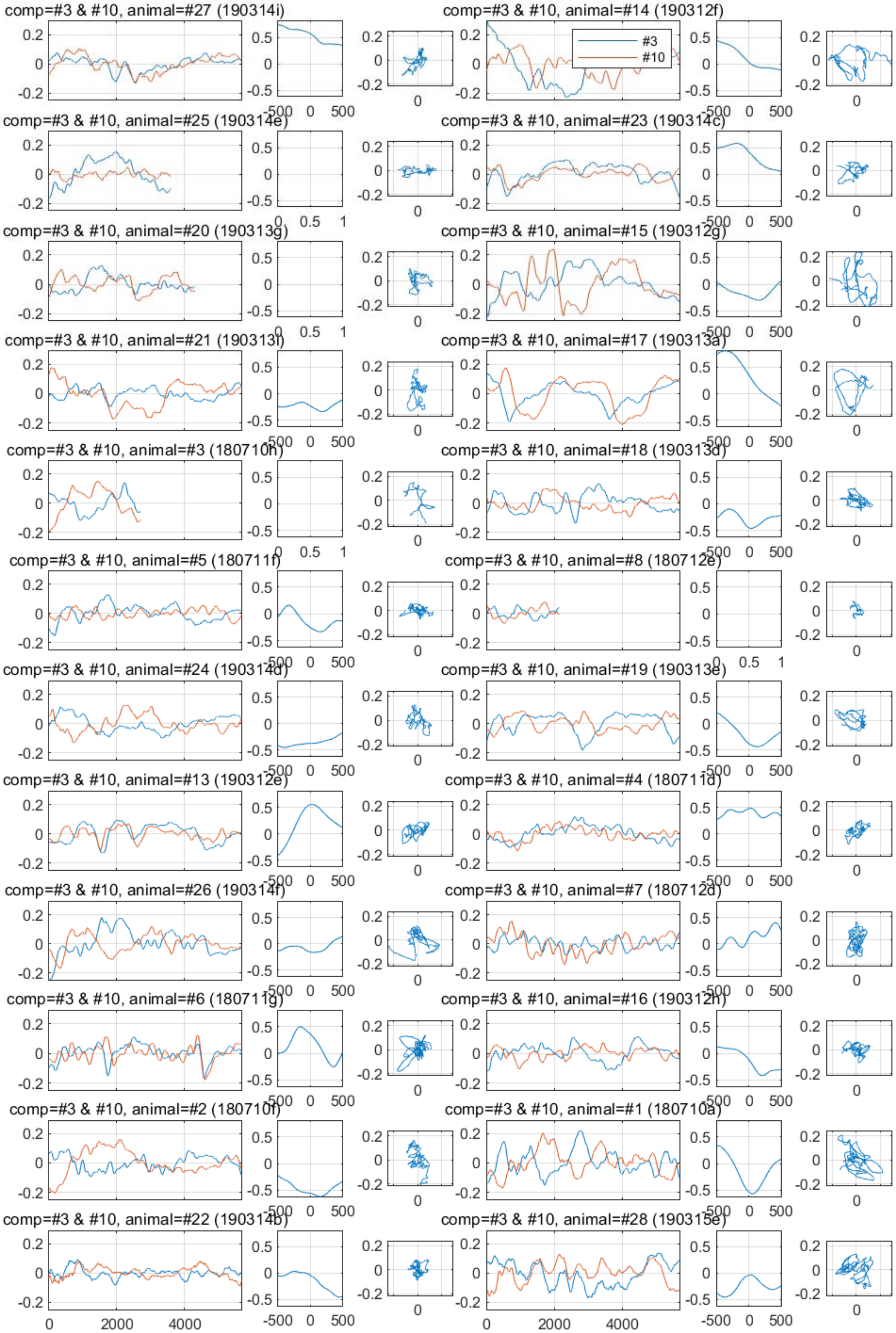

comp=#3 & #10, animal=#27 (190314i)
comp=#3 & #10, animal=#14 (190312f)
#3
#10
comp=#3 & #10, animal=#25 (190314e)
comp=#3 & #10, animal=#23 (190314c)
comp=#3 & #10, animal=#20 (190313g)
comp=#3 & #10, animal=#15 (190312g)
comp=#3 & #10, animal=#21 (190313i)
comp=#3 & #10, animal=#17 (190313a)
comp=#3 & #10, animal=#3 (180710h)
comp=#3 & #10, animal=#18 (190313d)
comp=#3 & #10, animal=#5 (180711f)
comp=#3 & #10, animal=#8 (180712e)
comp=#3 & #10, animal=#24 (190314d)
comp=#3 & #10, animal=#19 (190313e)
comp=#3 & #10, animal=#13 (190312e)
comp=#3 & #10, animal=#4 (180711d)
comp=#3 & #10, animal=#26 (190314f)
comp=#3 & #10, animal=#7 (180712d)
comp=#3 & #10, animal=#6 (180711g)
comp=#3 & #10, animal=#16 (190312h)
comp=#3 & #10, animal=#2 (180710f)
comp=#3 & #10, animal=#1 (180710a)
comp=#3 & #10, animal=#22 (190314b)
comp=#3 & #10, animal=#28 (190315e)

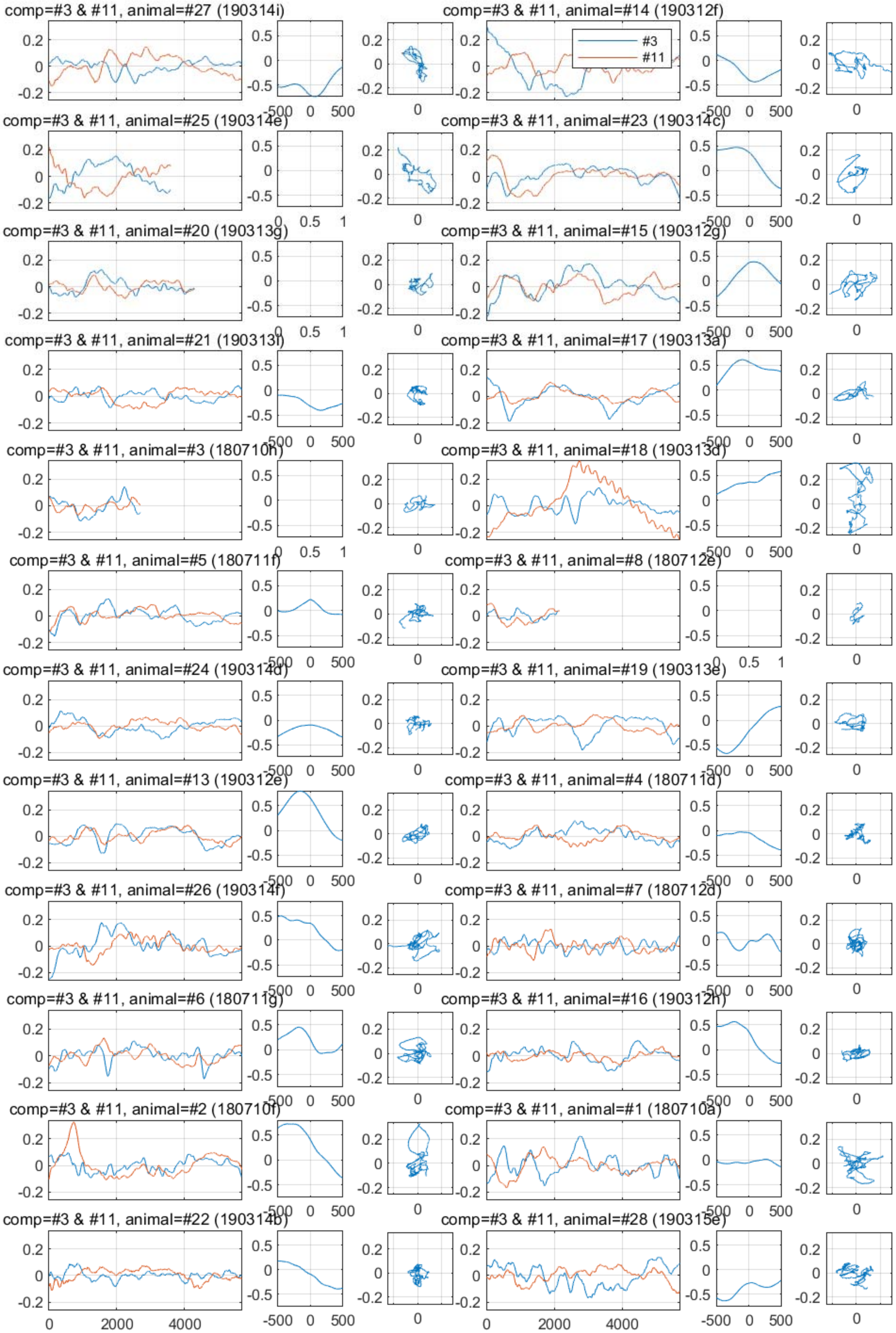

comp=#3 & #11, animal=#27 (190314i)
comp=#3 & #11, animal=#14 (190312f)
#3
#11
comp=#3 & #11, animal=#25 (190314e)
comp=#3 & #11, animal=#23 (190314c)
comp=#3 & #11, animal=#20 (190313g)
comp=#3 & #11, animal=#15 (190312g)
comp=#3 & #11, animal=#21 (190313i)
comp=#3 & #11, animal=#17 (190313a)
comp=#3 & #11, animal=#3 (180710h)
comp=#3 & #11, animal=#18 (190313d)
comp=#3 & #11, animal=#5 (180711f)
comp=#3 & #11, animal=#8 (180712e)
comp=#3 & #11, animal=#24 (190314d)
comp=#3 & #11, animal=#19 (190313e)
comp=#3 & #11, animal=#13 (190312e)
comp=#3 & #11, animal=#4 (180711d)
comp=#3 & #11, animal=#26 (190314f)
comp=#3 & #11, animal=#7 (180712d)
comp=#3 & #11, animal=#6 (180711g)
comp=#3 & #11, animal=#16 (190312h)
comp=#3 & #11, animal=#2 (180710f)
comp=#3 & #11, animal=#1 (180710a)
comp=#3 & #11, animal=#22 (190314b)
comp=#3 & #11, animal=#28 (190315e)

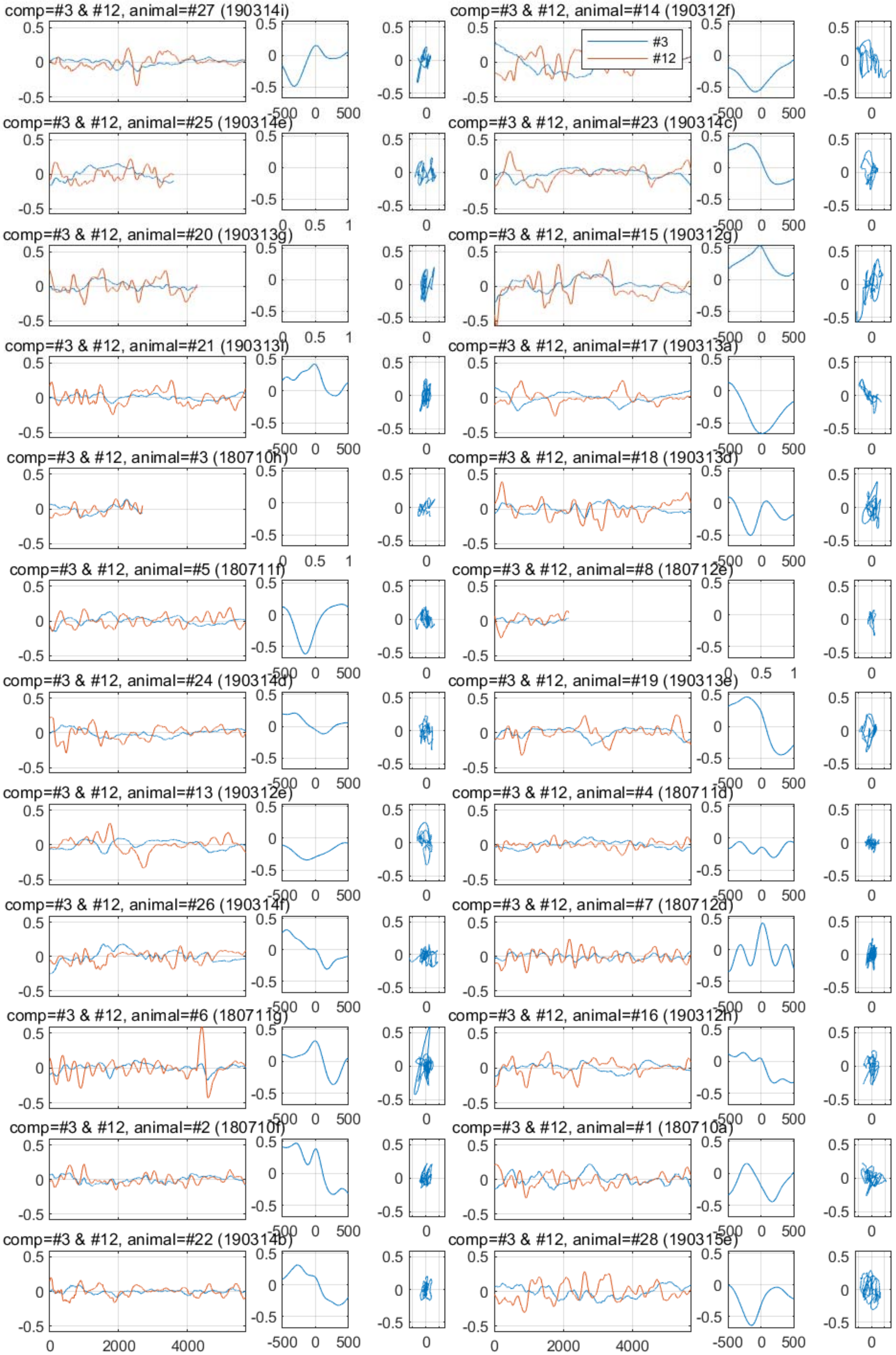
comp=#3 & #12, animal=#27 (190314i)
comp=#3 & #12, animal=#14 (190312f)
#3
#12
comp=#3 & #12, animal=#25 (190314e)
comp=#3 & #12, animal=#23 (190314c)
comp=#3 & #12, animal=#20 (190313g)
comp=#3 & #12, animal=#15 (190312g)
comp=#3 & #12, animal=#21 (190313i)
comp=#3 & #12, animal=#17 (190313a)
comp=#3 & #12, animal=#3 (180710h)
comp=#3 & #12, animal=#18 (190313d)
comp=#3 & #12, animal=#5 (180711f)
comp=#3 & #12, animal=#8 (180712e)
comp=#3 & #12, animal=#24 (190314d)
comp=#3 & #12, animal=#19 (190313e)
comp=#3 & #12, animal=#13 (190312e)
comp=#3 & #12, animal=#4 (180711d)
comp=#3 & #12, animal=#26 (190314f)
comp=#3 & #12, animal=#7 (180712d)
comp=#3 & #12, animal=#6 (180711g)
comp=#3 & #12, animal=#16 (190312h)
comp=#3 & #12, animal=#2 (180710f)
comp=#3 & #12, animal=#1 (180710a)
comp=#3 & #12, animal=#22 (190314b)
comp=#3 & #12, animal=#28 (190315e)

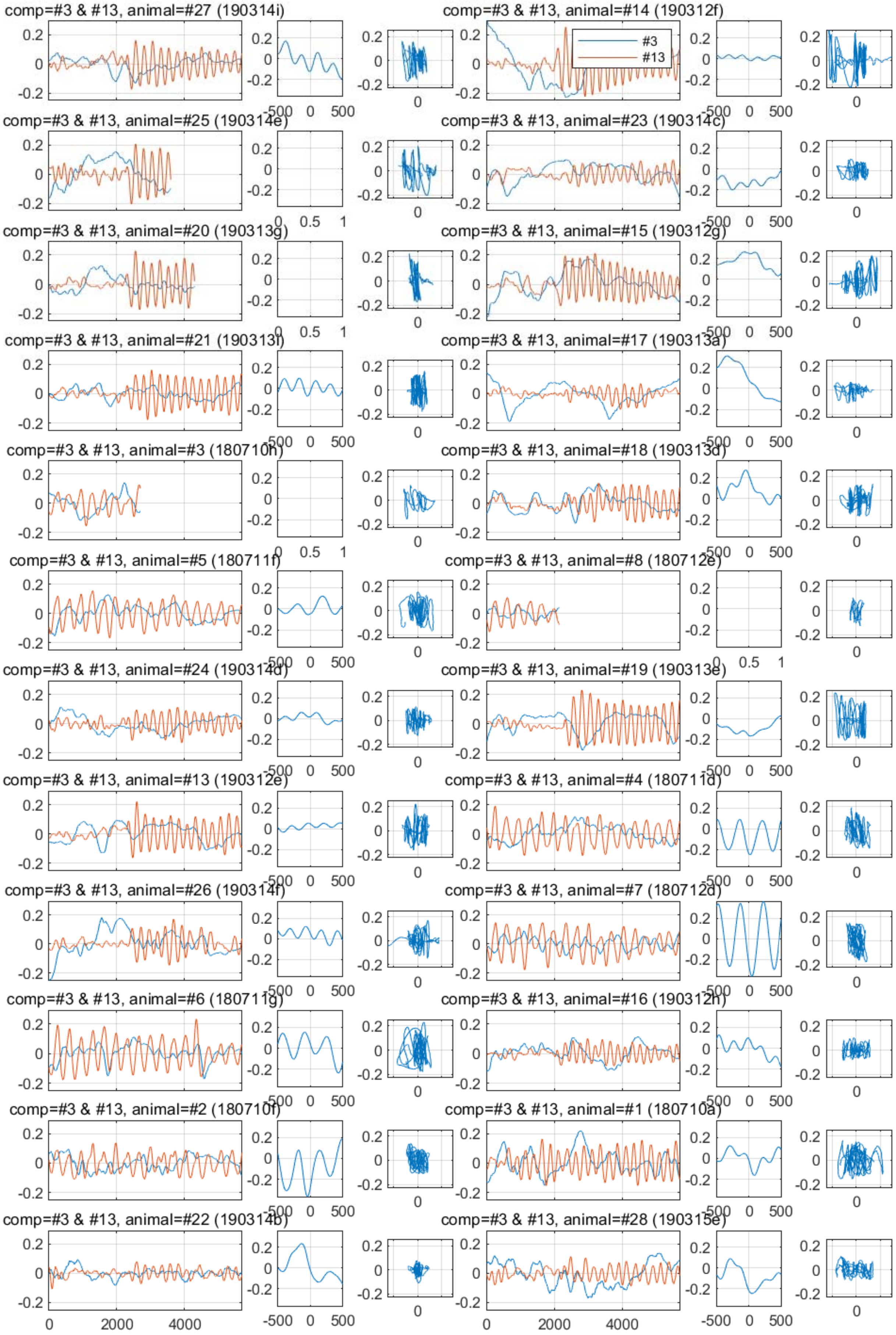
comp=#3 & #13, animal=#27 (190314i)
comp=#3 & #13, animal=#14 (190312f)
#3
#13
comp=#3 & #13, animal=#25 (190314e)
comp=#3 & #13, animal=#23 (190314c)
comp=#3 & #13, animal=#20 (190313g)
comp=#3 & #13, animal=#15 (190312g)
comp=#3 & #13, animal=#21 (190313f)
comp=#3 & #13, animal=#17 (190313a)
comp=#3 & #13, animal=#3 (180710h)
comp=#3 & #13, animal=#18 (190313d)
comp=#3 & #13, animal=#5 (180711f)
comp=#3 & #13, animal=#8 (180712e)
comp=#3 & #13, animal=#24 (190314d)
comp=#3 & #13, animal=#19 (190313e)
comp=#3 & #13, animal=#13 (190312e)
comp=#3 & #13, animal=#4 (180711d)
comp=#3 & #13, animal=#26 (190314f)
comp=#3 & #13, animal=#7 (180712d)
comp=#3 & #13, animal=#6 (180711g)
comp=#3 & #13, animal=#16 (190312h)
comp=#3 & #13, animal=#2 (180710f)
comp=#3 & #13, animal=#1 (180710a)
comp=#3 & #13, animal=#22 (190314b)
comp=#3 & #13, animal=#28 (190315e)

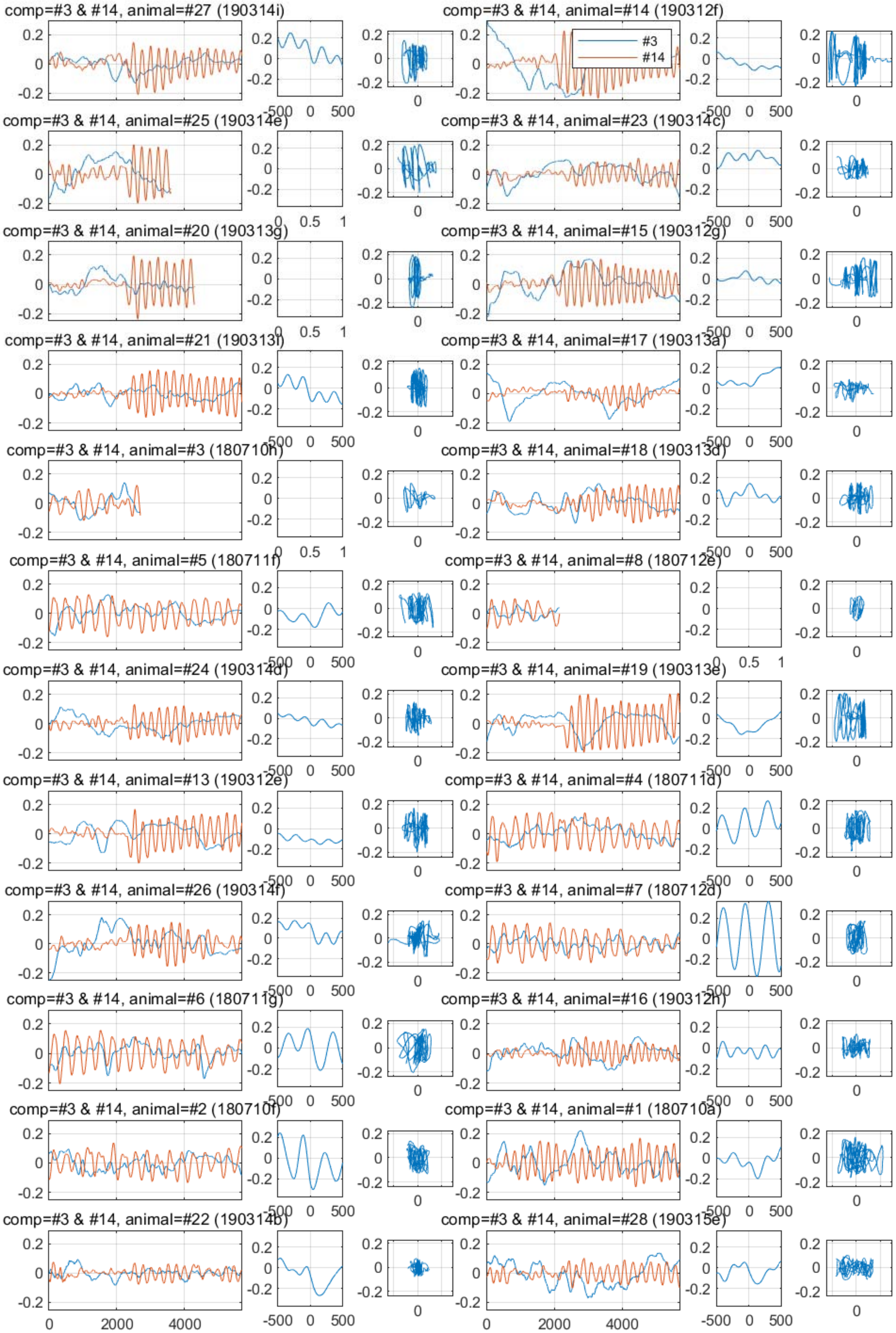
comp=#3 & #14, animal=#27 (190314i)
comp=#3 & #14, animal=#14 (190312f)
#3
#14
comp=#3 & #14, animal=#25 (190314e)
comp=#3 & #14, animal=#23 (190314c)
comp=#3 & #14, animal=#20 (190313g)
comp=#3 & #14, animal=#15 (190312g)
comp=#3 & #14, animal=#21 (190313f)
comp=#3 & #14, animal=#17 (190313a)
comp=#3 & #14, animal=#3 (180710h)
comp=#3 & #14, animal=#18 (190313d)
comp=#3 & #14, animal=#5 (180711f)
comp=#3 & #14, animal=#8 (180712e)
comp=#3 & #14, animal=#24 (190314d)
comp=#3 & #14, animal=#19 (190313e)
comp=#3 & #14, animal=#13 (190312e)
comp=#3 & #14, animal=#4 (180711d)
comp=#3 & #14, animal=#26 (190314f)
comp=#3 & #14, animal=#7 (180712d)
comp=#3 & #14, animal=#6 (180711g)
comp=#3 & #14, animal=#16 (190312h)
comp=#3 & #14, animal=#2 (180710f)
comp=#3 & #14, animal=#1 (180710a)
comp=#3 & #14, animal=#22 (190314b)
comp=#3 & #14, animal=#28 (190315e)
0
2000
4000
-500
0
500

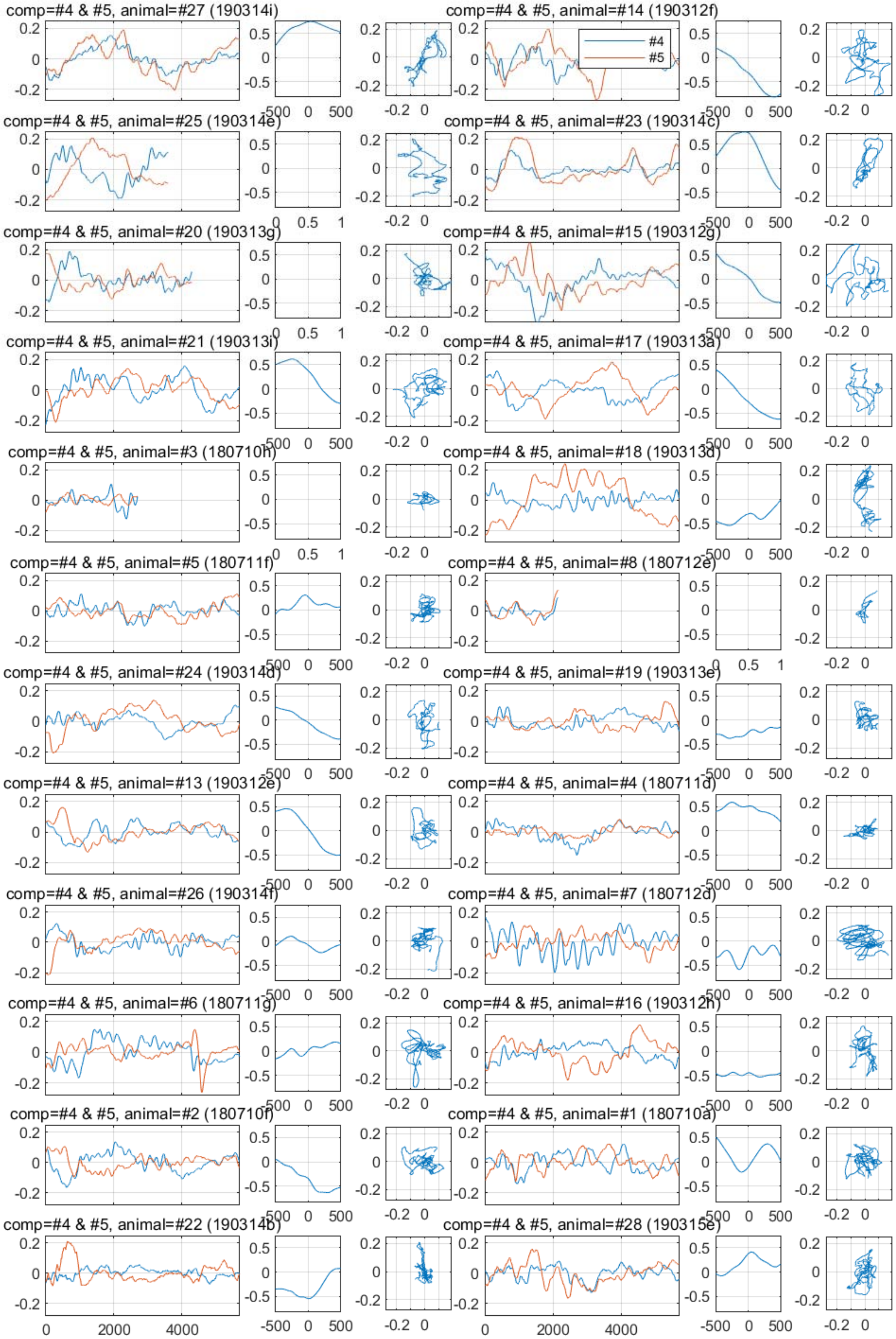

comp=#4 & #5, animal=#27 (190314i)
comp=#4 & #5, animal=#14 (190312f)
#4
#5
comp=#4 & #5, animal=#25 (190314e)
comp=#4 & #5, animal=#23 (190314c)
comp=#4 & #5, animal=#20 (190313g)
comp=#4 & #5, animal=#15 (190312g)
comp=#4 & #5, animal=#21 (190313i)
comp=#4 & #5, animal=#17 (190313a)
comp=#4 & #5, animal=#3 (180710h)
comp=#4 & #5, animal=#18 (190313d)
comp=#4 & #5, animal=#5 (180711f)
comp=#4 & #5, animal=#8 (180712e)
comp=#4 & #5, animal=#24 (190314d)
comp=#4 & #5, animal=#19 (190313e)
comp=#4 & #5, animal=#13 (190312e)
comp=#4 & #5, animal=#4 (180711d)
comp=#4 & #5, animal=#26 (190314f)
comp=#4 & #5, animal=#7 (180712d)
comp=#4 & #5, animal=#6 (180711g)
comp=#4 & #5, animal=#16 (190312h)
comp=#4 & #5, animal=#2 (180710f)
comp=#4 & #5, animal=#1 (180710a)
comp=#4 & #5, animal=#22 (190314b)
comp=#4 & #5, animal=#28 (190315e)

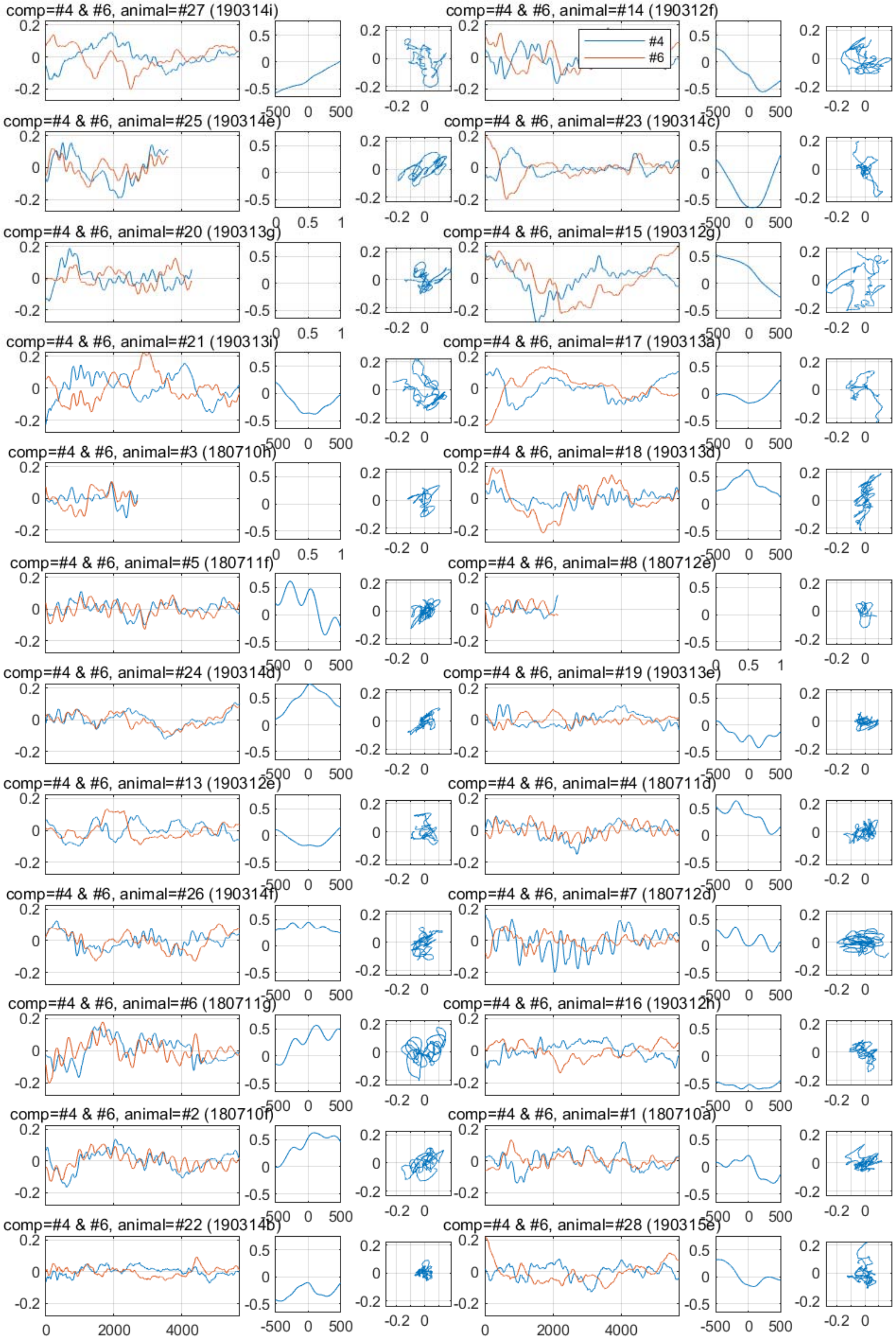
comp=#4 & #6, animal=#27 (190314i)
comp=#4 & #6, animal=#14 (190312f)
#4
#6
comp=#4 & #6, animal=#25 (190314e)
comp=#4 & #6, animal=#23 (190314c)
comp=#4 & #6, animal=#20 (190313g)
comp=#4 & #6, animal=#15 (190312g)
comp=#4 & #6, animal=#21 (190313i)
comp=#4 & #6, animal=#17 (190313a)
comp=#4 & #6, animal=#3 (180710h)
comp=#4 & #6, animal=#18 (190313d)
comp=#4 & #6, animal=#5 (180711f)
comp=#4 & #6, animal=#8 (180712e)
comp=#4 & #6, animal=#24 (190314d)
comp=#4 & #6, animal=#19 (190313e)
comp=#4 & #6, animal=#13 (190312e)
comp=#4 & #6, animal=#4 (180711d)
comp=#4 & #6, animal=#26 (190314f)
comp=#4 & #6, animal=#7 (180712d)
comp=#4 & #6, animal=#6 (180711g)
comp=#4 & #6, animal=#16 (190312h)
comp=#4 & #6, animal=#2 (180710f)
comp=#4 & #6, animal=#1 (180710a)
comp=#4 & #6, animal=#22 (190314b)
comp=#4 & #6, animal=#28 (190315e)

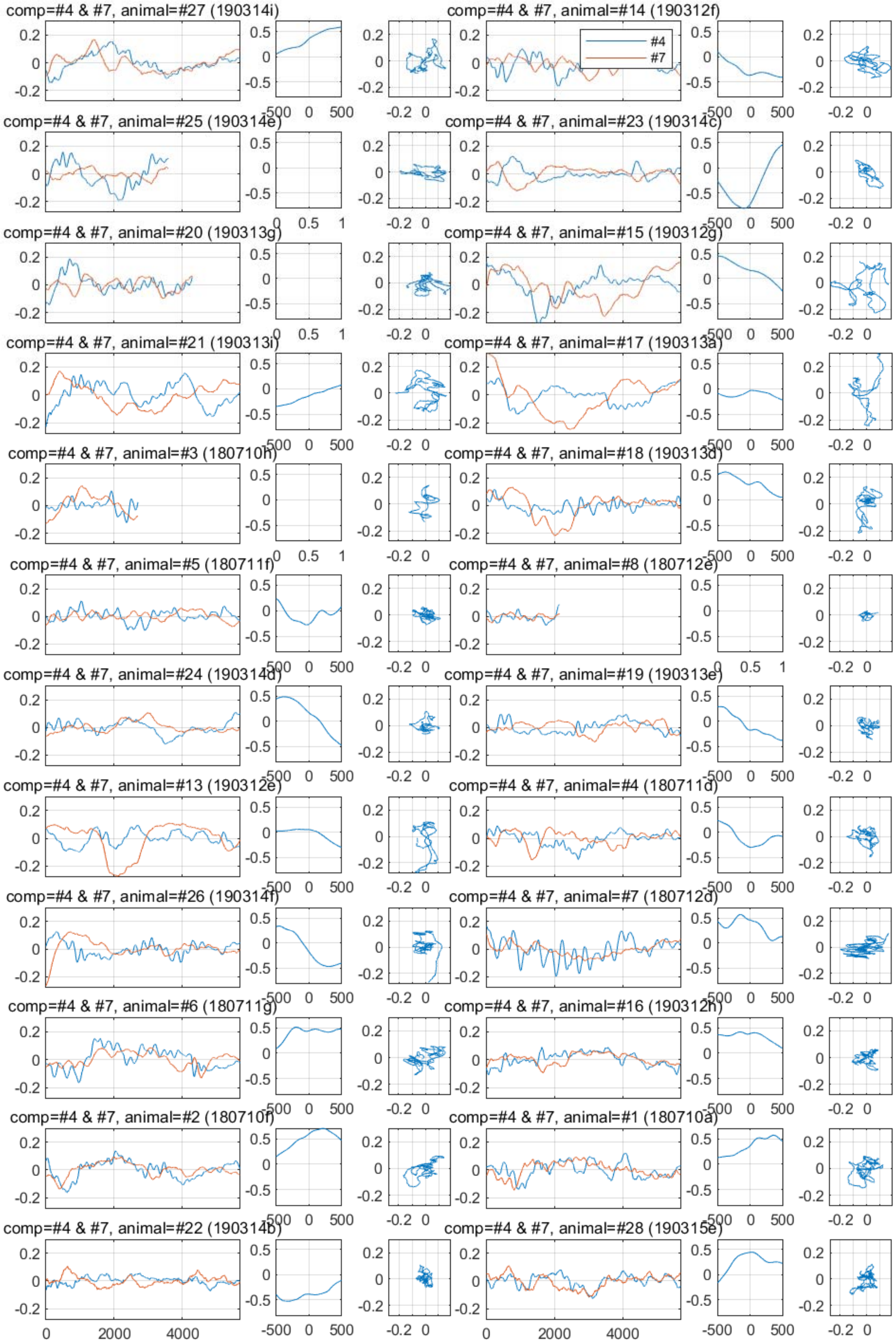
comp=#4 & #7, animal=#27 (190314i)
comp=#4 & #7, animal=#14 (190312f)
#4
#7
comp=#4 & #7, animal=#25 (190314e)
comp=#4 & #7, animal=#23 (190314c)
comp=#4 & #7, animal=#20 (190313g)
comp=#4 & #7, animal=#15 (190312g)
comp=#4 & #7, animal=#21 (190313i)
comp=#4 & #7, animal=#17 (190313a)
comp=#4 & #7, animal=#3 (180710h)
comp=#4 & #7, animal=#18 (190313d)
comp=#4 & #7, animal=#5 (180711f)
comp=#4 & #7, animal=#8 (180712e)
comp=#4 & #7, animal=#24 (190314d)
comp=#4 & #7, animal=#19 (190313e)
comp=#4 & #7, animal=#13 (190312e)
comp=#4 & #7, animal=#4 (180711d)
comp=#4 & #7, animal=#26 (190314f)
comp=#4 & #7, animal=#7 (180712d)
comp=#4 & #7, animal=#6 (180711g)
comp=#4 & #7, animal=#16 (190312h)
comp=#4 & #7, animal=#2 (180710f)
comp=#4 & #7, animal=#1 (180710a)
comp=#4 & #7, animal=#22 (190314b)
comp=#4 & #7, animal=#28 (190315e)

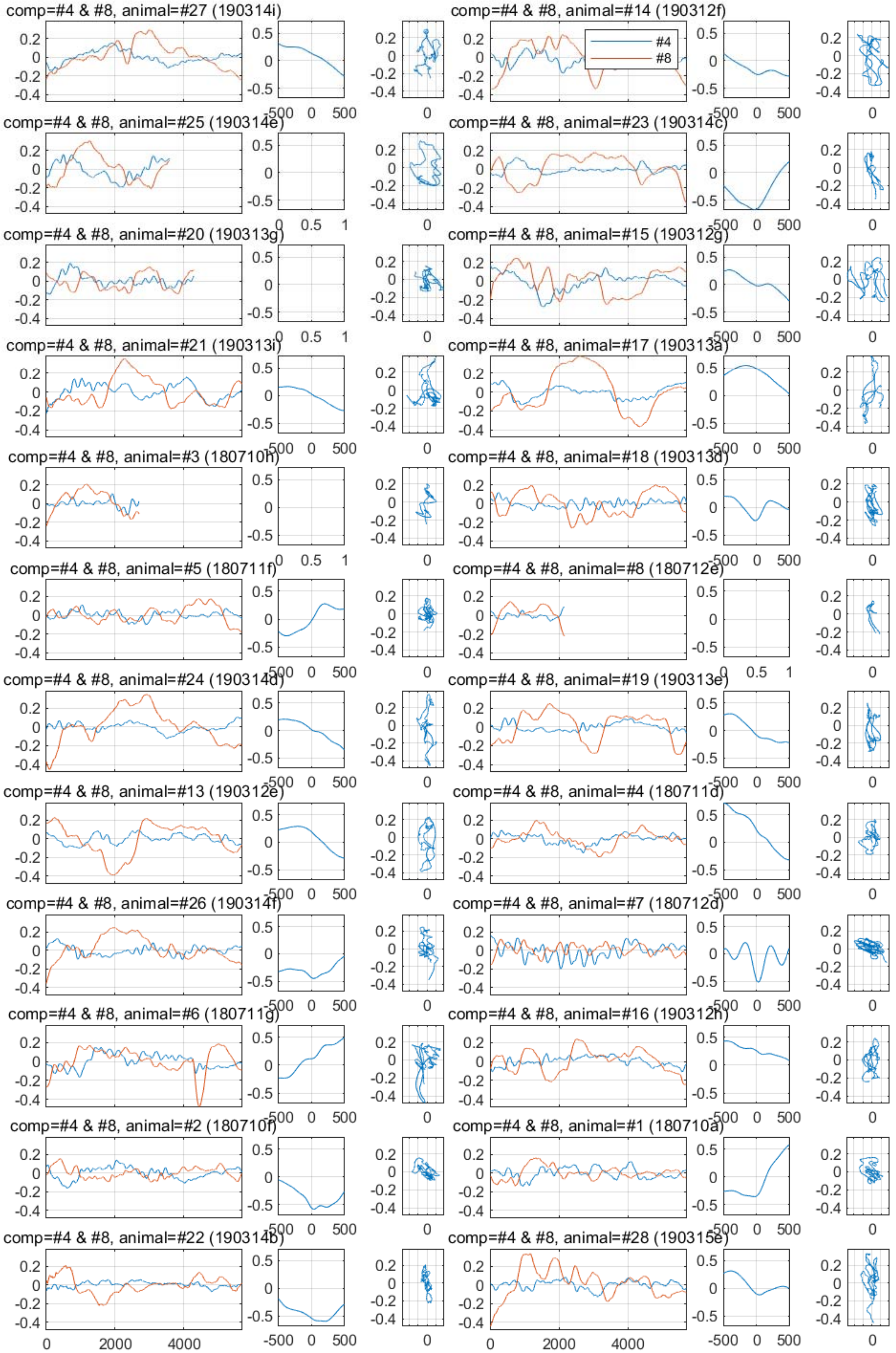
comp=#4 & #8, animal=#27 (190314i)
comp=#4 & #8, animal=#14 (190312f)
#4
#8
comp=#4 & #8, animal=#25 (190314e)
comp=#4 & #8, animal=#23 (190314c)
comp=#4 & #8, animal=#20 (190313g)
comp=#4 & #8, animal=#15 (190312g)
comp=#4 & #8, animal=#21 (190313i)
comp=#4 & #8, animal=#17 (190313a)
comp=#4 & #8, animal=#3 (180710h)
comp=#4 & #8, animal=#18 (190313d)
comp=#4 & #8, animal=#5 (180711f)
comp=#4 & #8, animal=#8 (180712e)
comp=#4 & #8, animal=#24 (190314d)
comp=#4 & #8, animal=#19 (190313e)
comp=#4 & #8, animal=#13 (190312e)
comp=#4 & #8, animal=#4 (180711d)
comp=#4 & #8, animal=#26 (190314f)
comp=#4 & #8, animal=#7 (180712d)
comp=#4 & #8, animal=#6 (180711g)
comp=#4 & #8, animal=#16 (190312h)
comp=#4 & #8, animal=#2 (180710f)
comp=#4 & #8, animal=#1 (180710a)
comp=#4 & #8, animal=#22 (190314b)
comp=#4 & #8, animal=#28 (190315e)

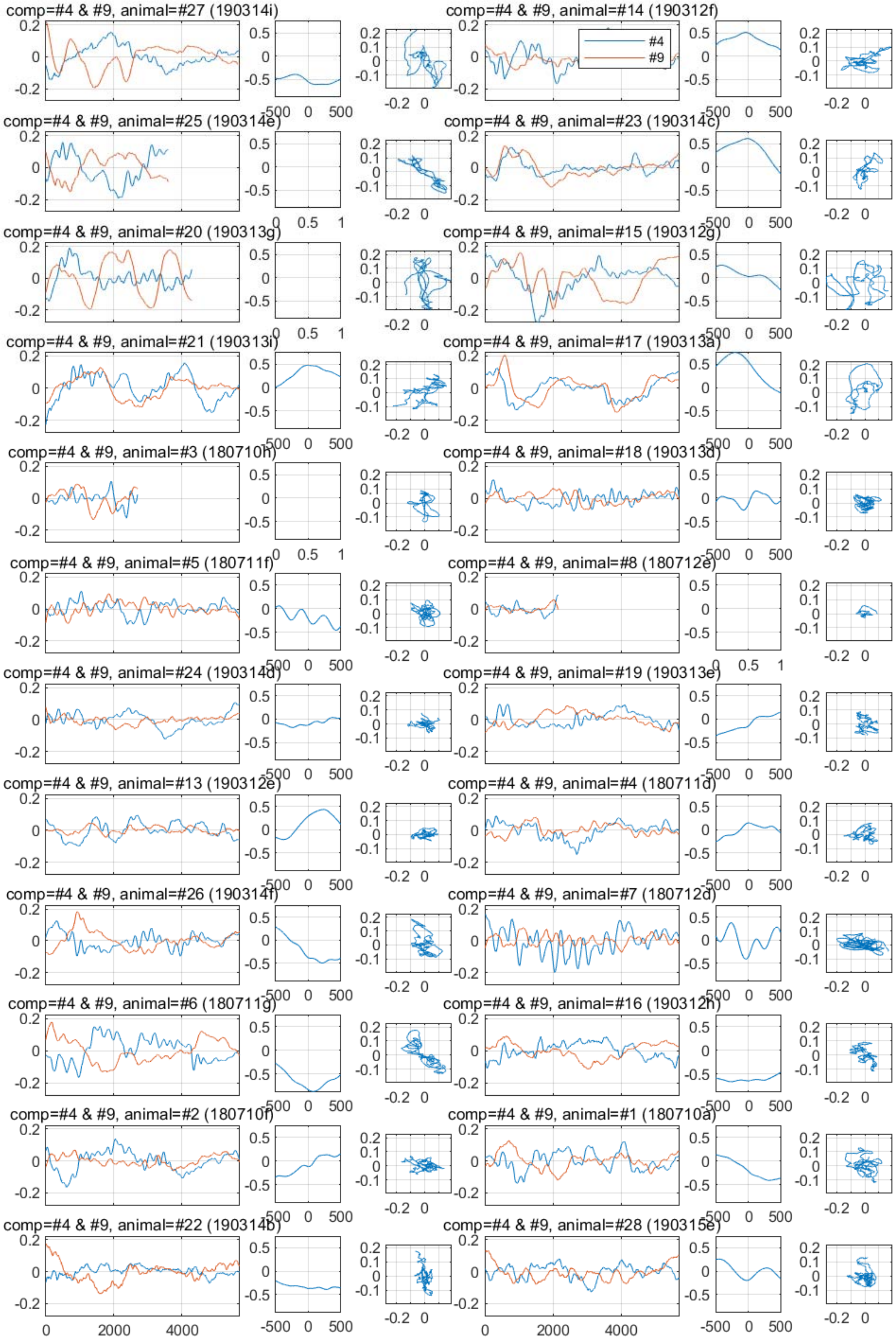
comp=#4 & #9, animal=#27 (190314i)
comp=#4 & #9, animal=#14 (190312f)
#4
#9
comp=#4 & #9, animal=#25 (190314e)
comp=#4 & #9, animal=#23 (190314c)
comp=#4 & #9, animal=#20 (190313g)
comp=#4 & #9, animal=#15 (190312g)
comp=#4 & #9, animal=#21 (190313i)
comp=#4 & #9, animal=#17 (190313a)
comp=#4 & #9, animal=#3 (180710h)
comp=#4 & #9, animal=#18 (190313d)
comp=#4 & #9, animal=#5 (180711f)
comp=#4 & #9, animal=#8 (180712e)
comp=#4 & #9, animal=#24 (190314d)
comp=#4 & #9, animal=#19 (190313e)
comp=#4 & #9, animal=#13 (190312e)
comp=#4 & #9, animal=#4 (180711d)
comp=#4 & #9, animal=#26 (190314f)
comp=#4 & #9, animal=#7 (180712d)
comp=#4 & #9, animal=#6 (180711g)
comp=#4 & #9, animal=#16 (190312h)
comp=#4 & #9, animal=#2 (180710f)
comp=#4 & #9, animal=#1 (180710a)
comp=#4 & #9, animal=#22 (190314b)
comp=#4 & #9, animal=#28 (190315e)

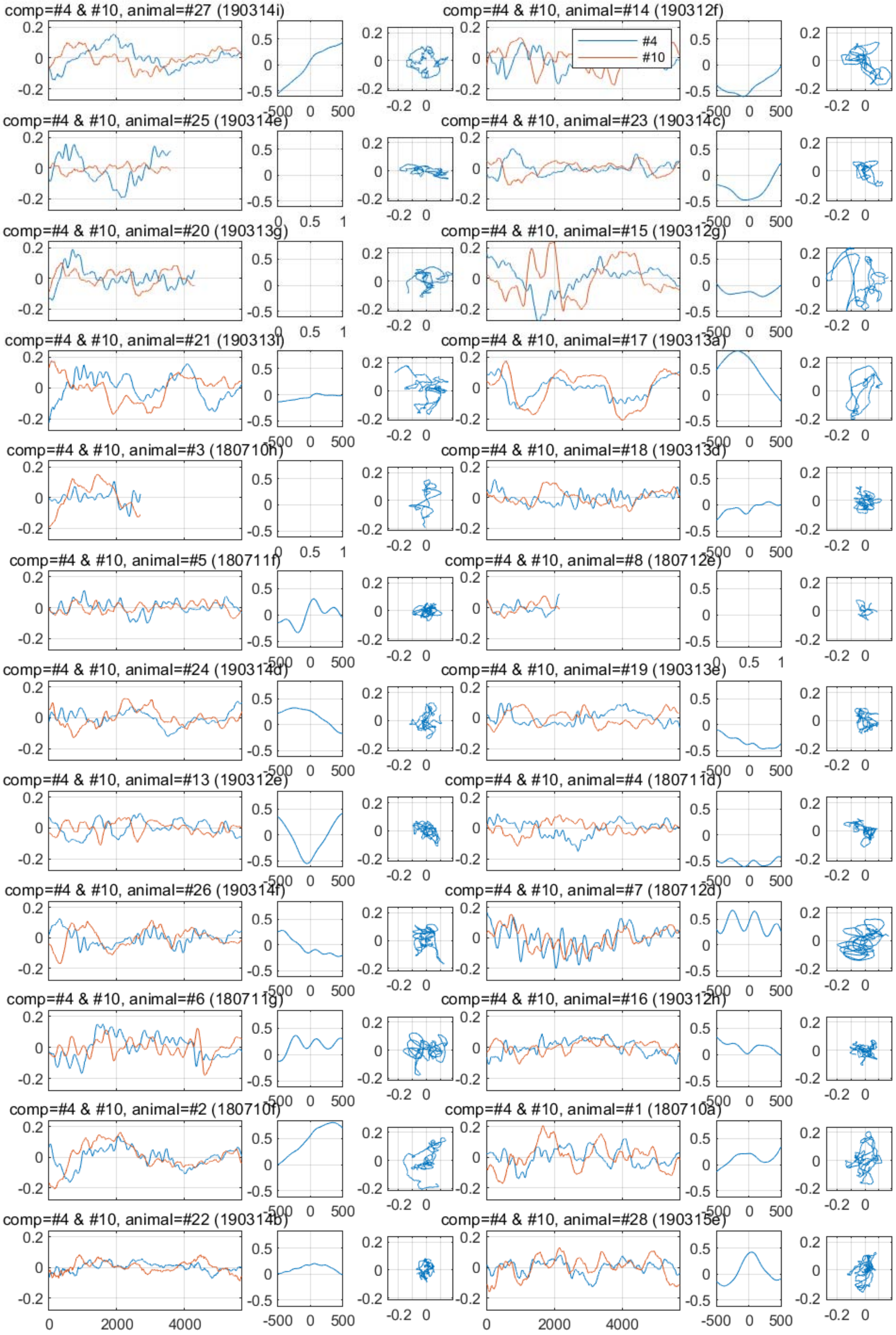

comp=#4 & #10, animal=#27 (190314i)
comp=#4 & #10, animal=#14 (190312f)
#4
#10
comp=#4 & #10, animal=#25 (190314e)
comp=#4 & #10, animal=#23 (190314c)
comp=#4 & #10, animal=#20 (190313g)
comp=#4 & #10, animal=#15 (190312g)
comp=#4 & #10, animal=#21 (190313i)
comp=#4 & #10, animal=#17 (190313a)
comp=#4 & #10, animal=#3 (180710h)
comp=#4 & #10, animal=#18 (190313d)
comp=#4 & #10, animal=#5 (180711f)
comp=#4 & #10, animal=#8 (180712e)
comp=#4 & #10, animal=#24 (190314d)
comp=#4 & #10, animal=#19 (190313e)
comp=#4 & #10, animal=#13 (190312e)
comp=#4 & #10, animal=#4 (180711d)
comp=#4 & #10, animal=#26 (190314f)
comp=#4 & #10, animal=#7 (180712d)
comp=#4 & #10, animal=#6 (180711g)
comp=#4 & #10, animal=#16 (190312h)
comp=#4 & #10, animal=#2 (180710f)
comp=#4 & #10, animal=#1 (180710a)
comp=#4 & #10, animal=#22 (190314b)
comp=#4 & #10, animal=#28 (190315e)
0
2000
4000
-500
0
500
-0.2
0
0.2
0.5
-0.5
1

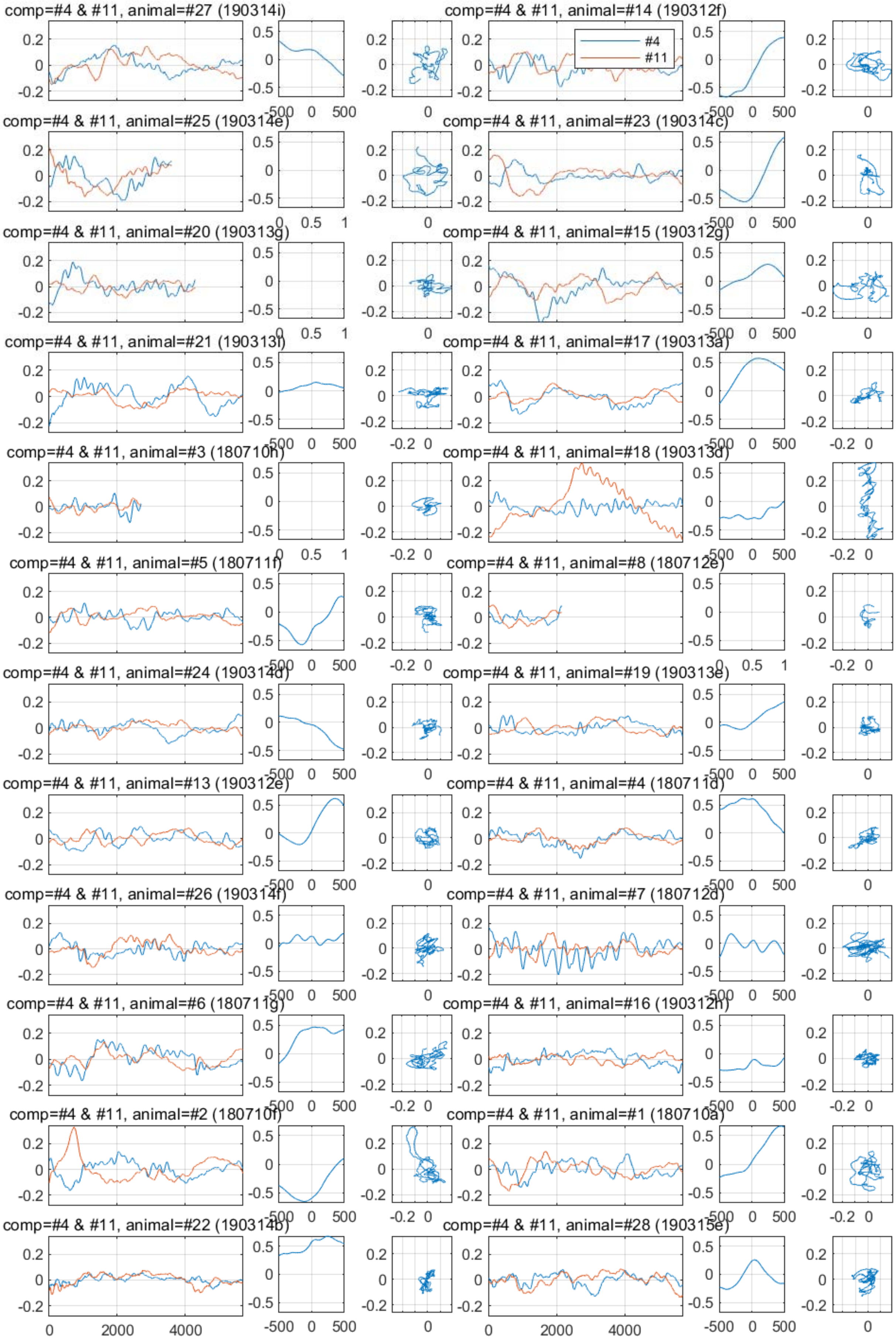
comp=#4 & #11, animal=#27 (190314i)
comp=#4 & #11, animal=#14 (190312f)
#4
#11
comp=#4 & #11, animal=#25 (190314e)
comp=#4 & #11, animal=#23 (190314c)
comp=#4 & #11, animal=#20 (190313g)
comp=#4 & #11, animal=#15 (190312g)
comp=#4 & #11, animal=#21 (190313i)
comp=#4 & #11, animal=#17 (190313a)
comp=#4 & #11, animal=#3 (180710h)
comp=#4 & #11, animal=#18 (190313d)
comp=#4 & #11, animal=#5 (180711f)
comp=#4 & #11, animal=#8 (180712e)
comp=#4 & #11, animal=#24 (190314d)
comp=#4 & #11, animal=#19 (190313e)
comp=#4 & #11, animal=#13 (190312e)
comp=#4 & #11, animal=#4 (180711d)
comp=#4 & #11, animal=#26 (190314f)
comp=#4 & #11, animal=#7 (180712d)
comp=#4 & #11, animal=#6 (180711g)
comp=#4 & #11, animal=#16 (190312h)
comp=#4 & #11, animal=#2 (180710f)
comp=#4 & #11, animal=#1 (180710a)
comp=#4 & #11, animal=#22 (190314b)
comp=#4 & #11, animal=#28 (190315e)

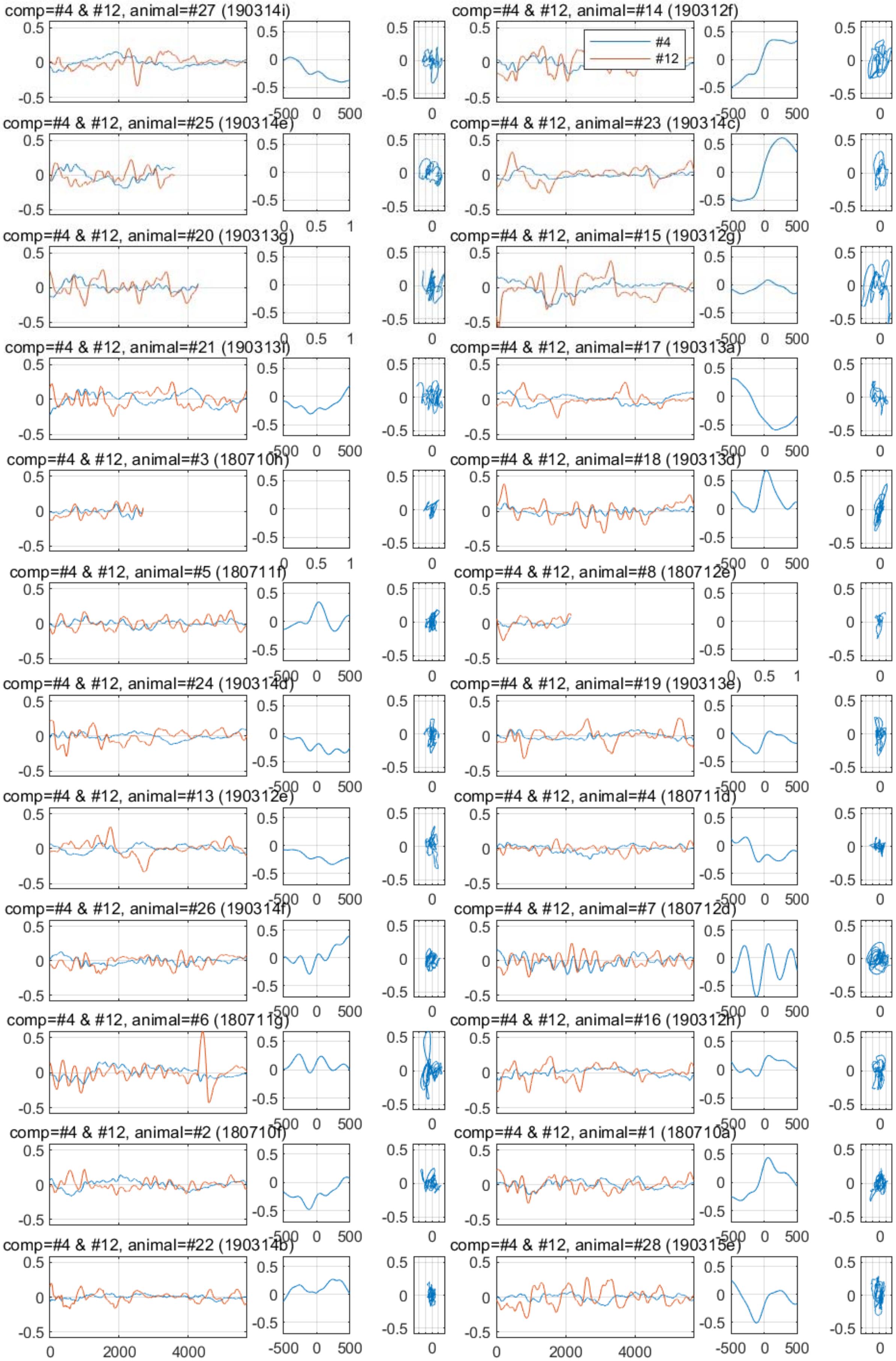

comp=#4 & #12, animal=#27 (190314i)
comp=#4 & #12, animal=#14 (190312f)
#4
#12
comp=#4 & #12, animal=#25 (190314e)
comp=#4 & #12, animal=#23 (190314c)
comp=#4 & #12, animal=#20 (190313g)
comp=#4 & #12, animal=#15 (190312g)
comp=#4 & #12, animal=#21 (190313f)
comp=#4 & #12, animal=#17 (190313a)
comp=#4 & #12, animal=#3 (180710h)
comp=#4 & #12, animal=#18 (190313d)
comp=#4 & #12, animal=#5 (180711f)
comp=#4 & #12, animal=#8 (180712e)
comp=#4 & #12, animal=#24 (190314d)
comp=#4 & #12, animal=#19 (190313e)
comp=#4 & #12, animal=#13 (190312e)
comp=#4 & #12, animal=#4 (180711d)
comp=#4 & #12, animal=#26 (190314f)
comp=#4 & #12, animal=#7 (180712d)
comp=#4 & #12, animal=#6 (180711g)
comp=#4 & #12, animal=#16 (190312h)
comp=#4 & #12, animal=#2 (180710f)
comp=#4 & #12, animal=#1 (180710a)
comp=#4 & #12, animal=#22 (190314b)
comp=#4 & #12, animal=#28 (190315e)

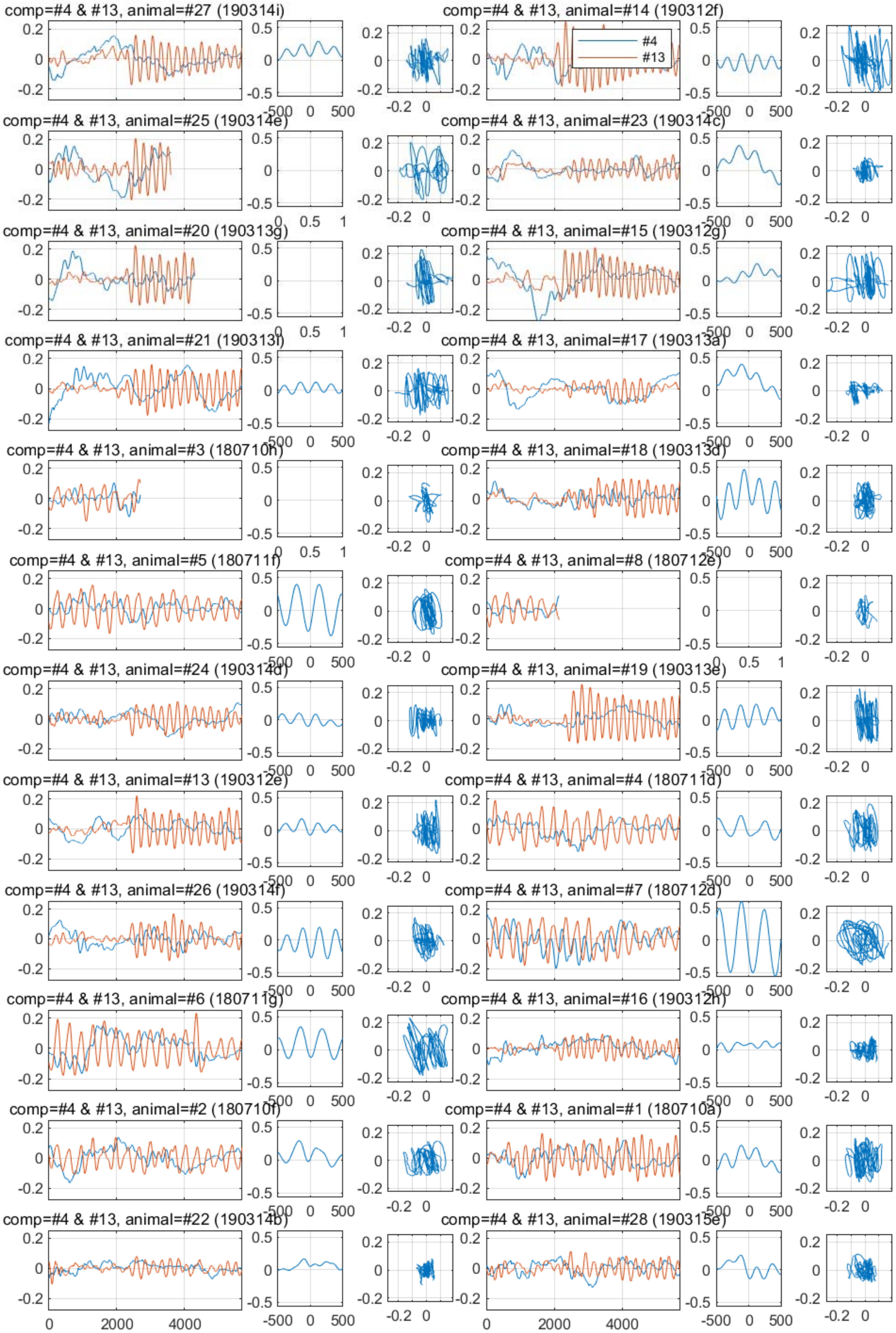

comp=#4 & #13, animal=#27 (190314i)
comp=#4 & #13, animal=#14 (190312f)
#4
#13
comp=#4 & #13, animal=#25 (190314e)
comp=#4 & #13, animal=#23 (190314c)
comp=#4 & #13, animal=#20 (190313g)
comp=#4 & #13, animal=#15 (190312g)
comp=#4 & #13, animal=#21 (190313i)
comp=#4 & #13, animal=#17 (190313a)
comp=#4 & #13, animal=#3 (180710h)
comp=#4 & #13, animal=#18 (190313d)
comp=#4 & #13, animal=#5 (180711f)
comp=#4 & #13, animal=#8 (180712e)
comp=#4 & #13, animal=#24 (190314d)
comp=#4 & #13, animal=#19 (190313e)
comp=#4 & #13, animal=#13 (190312e)
comp=#4 & #13, animal=#4 (180711d)
comp=#4 & #13, animal=#26 (190314f)
comp=#4 & #13, animal=#7 (180712d)
comp=#4 & #13, animal=#6 (180711g)
comp=#4 & #13, animal=#16 (190312h)
comp=#4 & #13, animal=#2 (180710f)
comp=#4 & #13, animal=#1 (180710a)
comp=#4 & #13, animal=#22 (190314b)
comp=#4 & #13, animal=#28 (190315e)

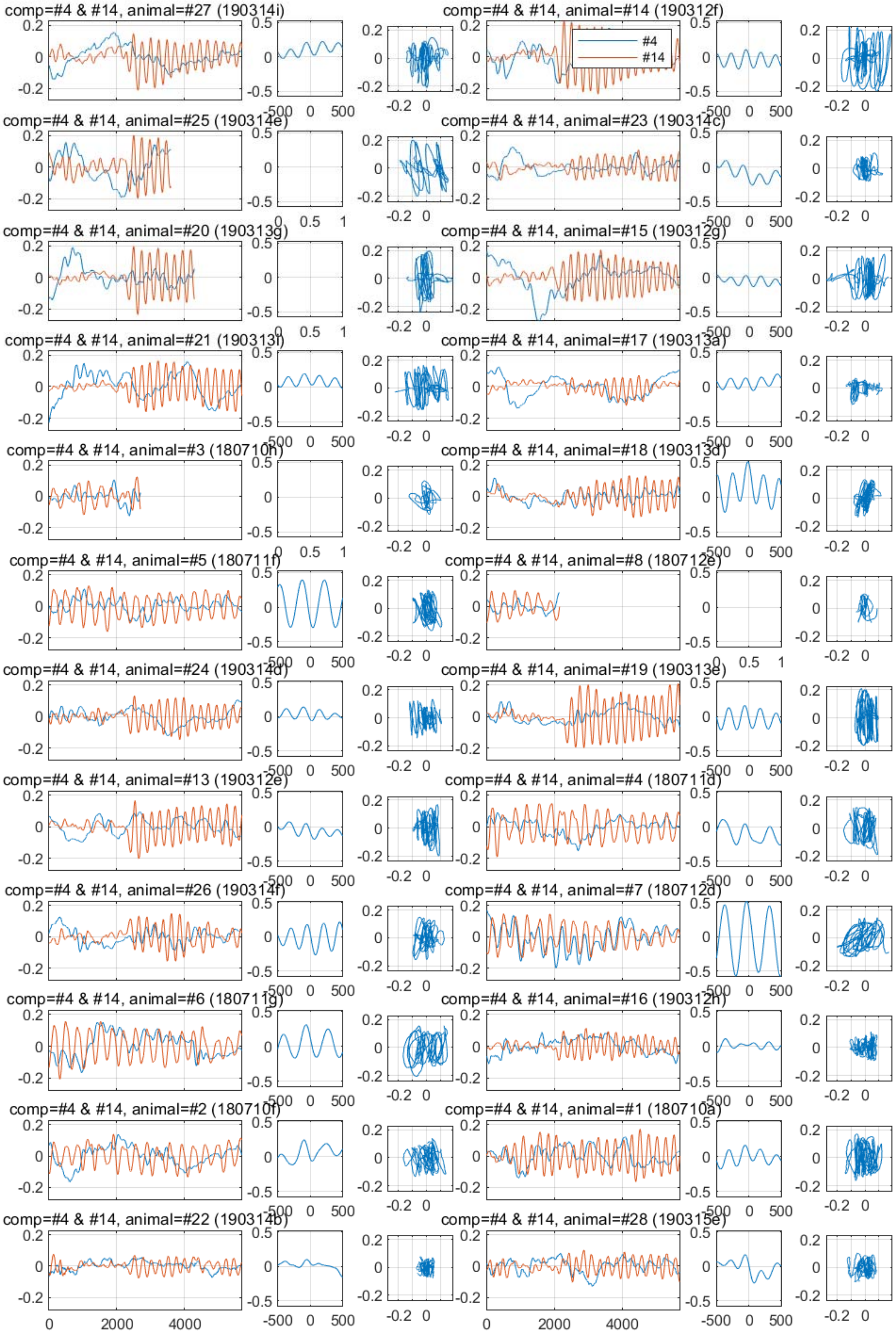
comp=#4 & #14, animal=#27 (190314i)
comp=#4 & #14, animal=#14 (190312f)
#4
#14
comp=#4 & #14, animal=#25 (190314e)
comp=#4 & #14, animal=#23 (190314c)
comp=#4 & #14, animal=#20 (190313g)
comp=#4 & #14, animal=#15 (190312g)
comp=#4 & #14, animal=#21 (190313i)
comp=#4 & #14, animal=#17 (190313a)
comp=#4 & #14, animal=#3 (180710h)
comp=#4 & #14, animal=#18 (190313d)
comp=#4 & #14, animal=#5 (180711f)
comp=#4 & #14, animal=#8 (180712e)
comp=#4 & #14, animal=#24 (190314d)
comp=#4 & #14, animal=#19 (190313e)
comp=#4 & #14, animal=#13 (190312e)
comp=#4 & #14, animal=#4 (180711d)
comp=#4 & #14, animal=#26 (190314f)
comp=#4 & #14, animal=#7 (180712d)
comp=#4 & #14, animal=#6 (180711g)
comp=#4 & #14, animal=#16 (190312h)
comp=#4 & #14, animal=#2 (180710f)
comp=#4 & #14, animal=#1 (180710a)
comp=#4 & #14, animal=#22 (190314b)
comp=#4 & #14, animal=#28 (190315e)

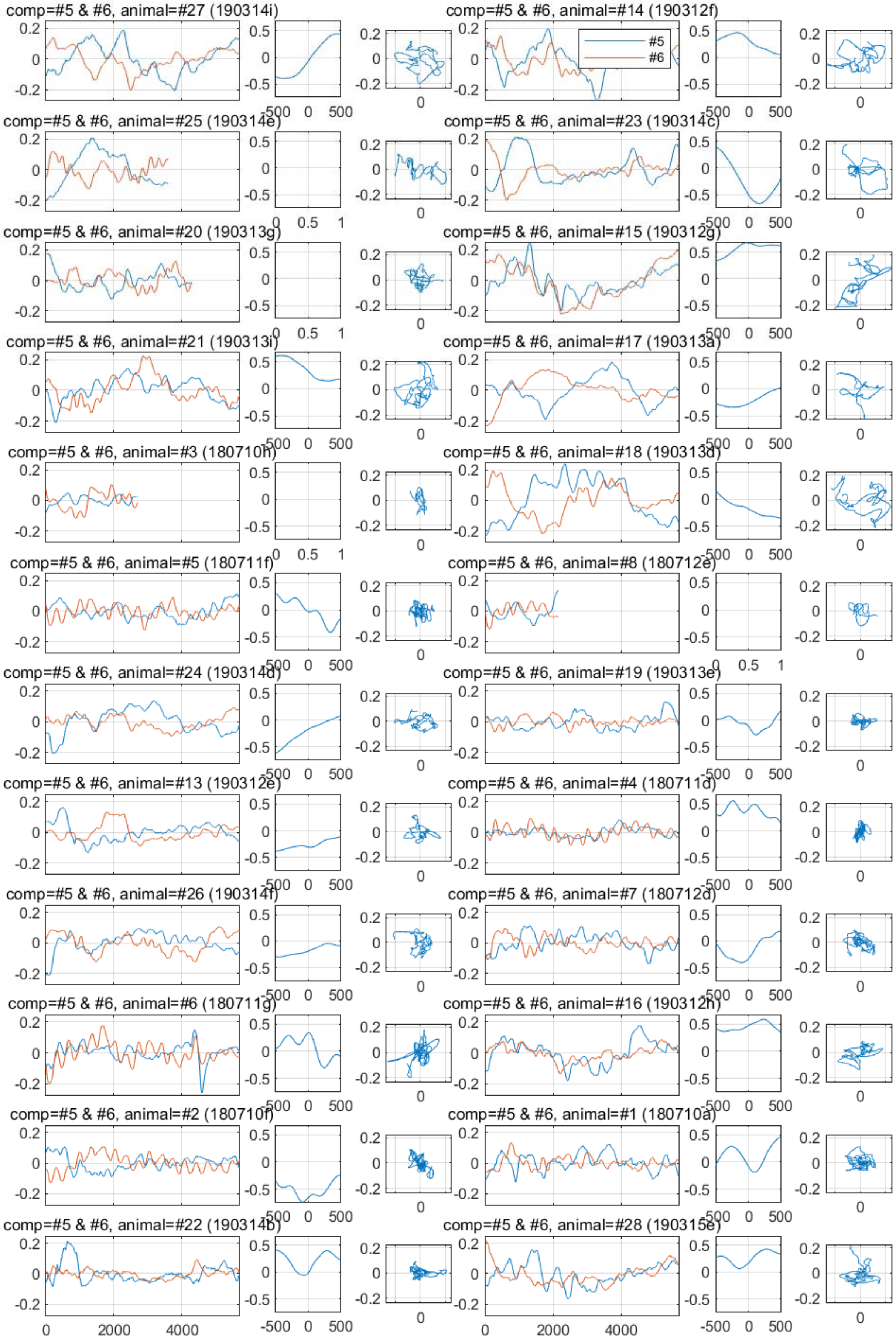
comp=#5 & #6, animal=#27 (190314i)
comp=#5 & #6, animal=#14 (190312f)
#5
#6
comp=#5 & #6, animal=#25 (190314e)
comp=#5 & #6, animal=#23 (190314c)
comp=#5 & #6, animal=#20 (190313g)
comp=#5 & #6, animal=#15 (190312g)
comp=#5 & #6, animal=#21 (190313i)
comp=#5 & #6, animal=#17 (190313a)
comp=#5 & #6, animal=#3 (180710h)
comp=#5 & #6, animal=#18 (190313d)
comp=#5 & #6, animal=#5 (180711f)
comp=#5 & #6, animal=#8 (180712e)
comp=#5 & #6, animal=#24 (190314d)
comp=#5 & #6, animal=#19 (190313e)
comp=#5 & #6, animal=#13 (190312e)
comp=#5 & #6, animal=#4 (180711d)
comp=#5 & #6, animal=#26 (190314f)
comp=#5 & #6, animal=#7 (180712d)
comp=#5 & #6, animal=#6 (180711g)
comp=#5 & #6, animal=#16 (190312h)
comp=#5 & #6, animal=#2 (180710f)
comp=#5 & #6, animal=#1 (180710a)
comp=#5 & #6, animal=#22 (190314b)
comp=#5 & #6, animal=#28 (190315e)
0
2000
4000
-500
0
500

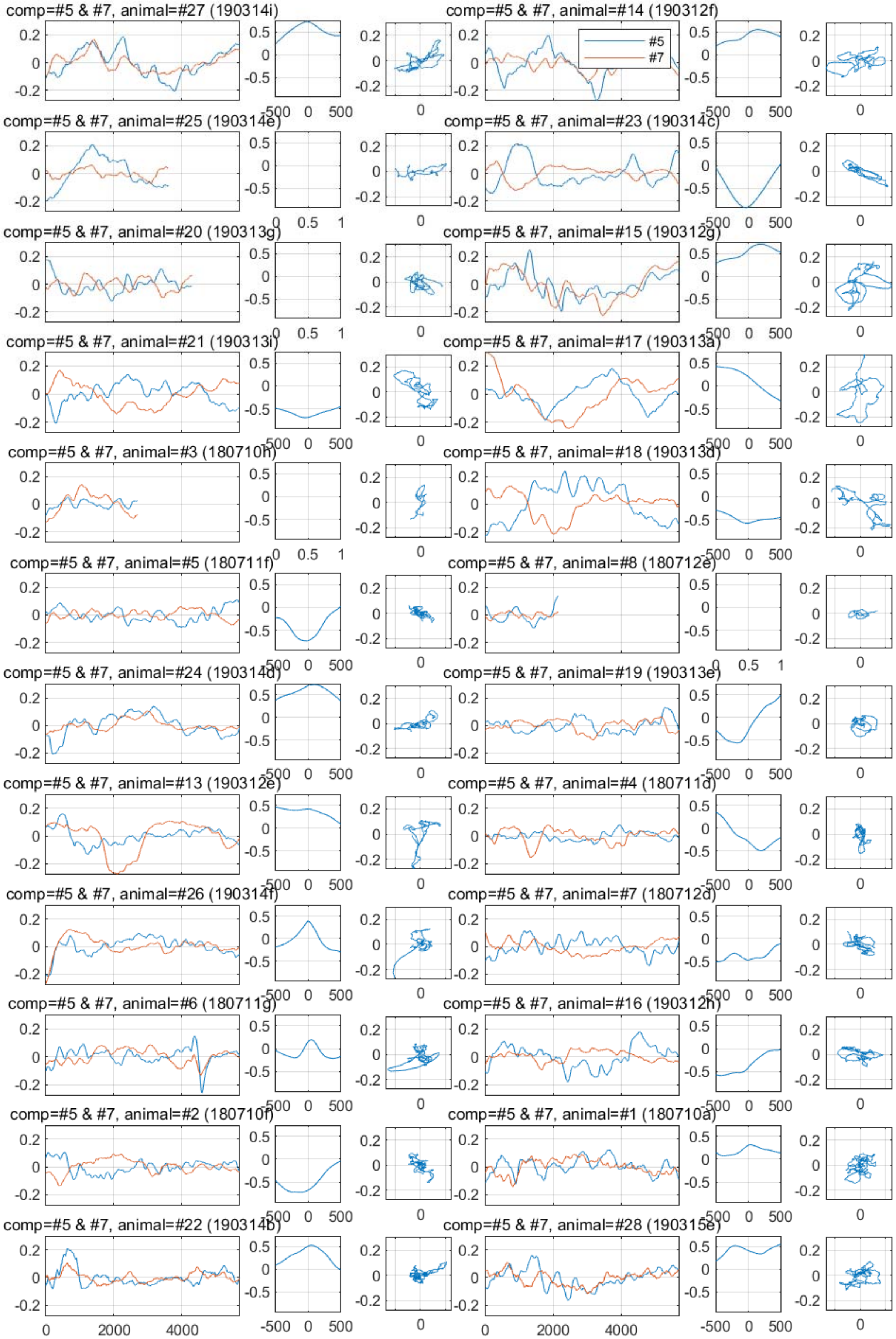

comp=#5 & #7, animal=#27 (190314i)
comp=#5 & #7, animal=#14 (190312f)
#5
#7
comp=#5 & #7, animal=#25 (190314e)
comp=#5 & #7, animal=#23 (190314c)
comp=#5 & #7, animal=#20 (190313g)
comp=#5 & #7, animal=#15 (190312g)
comp=#5 & #7, animal=#21 (190313i)
comp=#5 & #7, animal=#17 (190313a)
comp=#5 & #7, animal=#3 (180710h)
comp=#5 & #7, animal=#18 (190313d)
comp=#5 & #7, animal=#5 (180711f)
comp=#5 & #7, animal=#8 (180712e)
comp=#5 & #7, animal=#24 (190314d)
comp=#5 & #7, animal=#19 (190313e)
comp=#5 & #7, animal=#13 (190312e)
comp=#5 & #7, animal=#4 (180711d)
comp=#5 & #7, animal=#26 (190314f)
comp=#5 & #7, animal=#7 (180712d)
comp=#5 & #7, animal=#6 (180711g)
comp=#5 & #7, animal=#16 (190312h)
comp=#5 & #7, animal=#2 (180710f)
comp=#5 & #7, animal=#1 (180710a)
comp=#5 & #7, animal=#22 (190314b)
comp=#5 & #7, animal=#28 (190315e)
0
2000
4000
-500
0
500

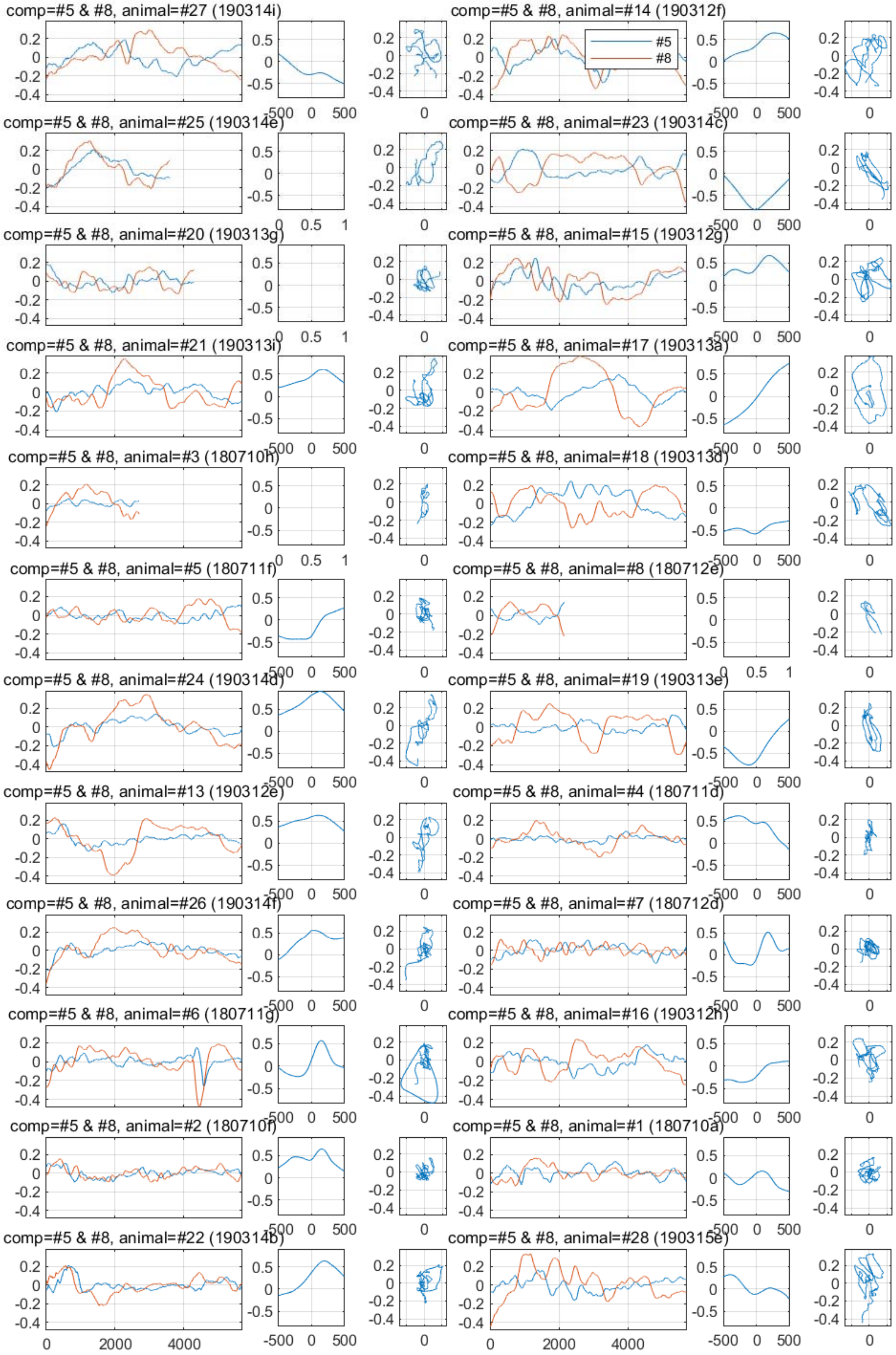

comp=#5 & #8, animal=#27 (190314i)
comp=#5 & #8, animal=#14 (190312f)
#5
#8
comp=#5 & #8, animal=#25 (190314e)
comp=#5 & #8, animal=#23 (190314c)
comp=#5 & #8, animal=#20 (190313g)
comp=#5 & #8, animal=#15 (190312g)
comp=#5 & #8, animal=#21 (190313i)
comp=#5 & #8, animal=#17 (190313a)
comp=#5 & #8, animal=#3 (180710h)
comp=#5 & #8, animal=#18 (190313d)
comp=#5 & #8, animal=#5 (180711f)
comp=#5 & #8, animal=#8 (180712e)
comp=#5 & #8, animal=#24 (190314d)
comp=#5 & #8, animal=#19 (190313e)
comp=#5 & #8, animal=#13 (190312e)
comp=#5 & #8, animal=#4 (180711d)
comp=#5 & #8, animal=#26 (190314f)
comp=#5 & #8, animal=#7 (180712d)
comp=#5 & #8, animal=#6 (180711g)
comp=#5 & #8, animal=#16 (190312h)
comp=#5 & #8, animal=#2 (180710f)
comp=#5 & #8, animal=#1 (180710a)
comp=#5 & #8, animal=#22 (190314b)
comp=#5 & #8, animal=#28 (190315e)

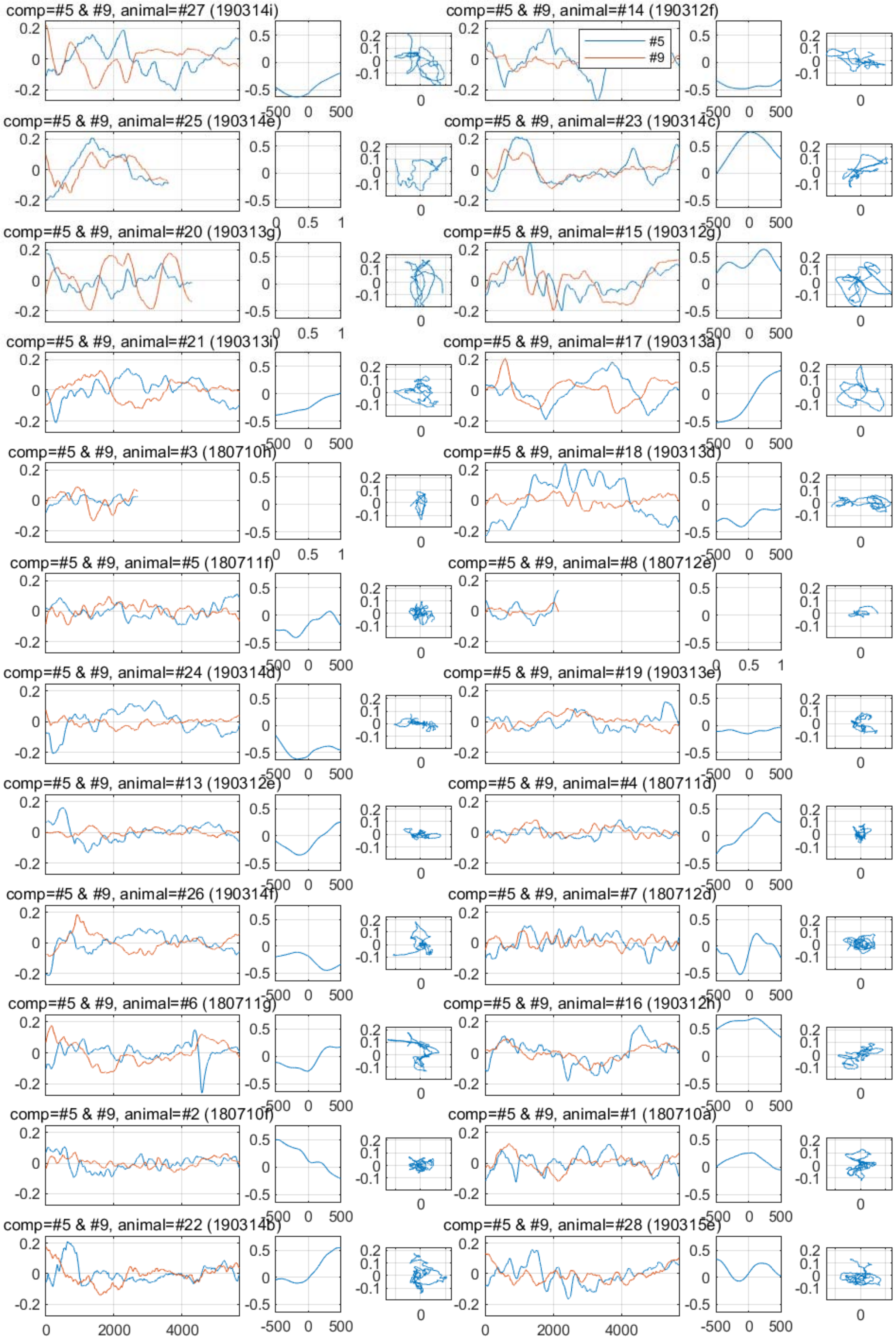
comp=#5 & #9, animal=#27 (190314i)
comp=#5 & #9, animal=#14 (190312f)
#5
#9
comp=#5 & #9, animal=#25 (190314e)
comp=#5 & #9, animal=#23 (190314c)
comp=#5 & #9, animal=#20 (190313g)
comp=#5 & #9, animal=#15 (190312g)
comp=#5 & #9, animal=#21 (190313i)
comp=#5 & #9, animal=#17 (190313a)
comp=#5 & #9, animal=#3 (180710h)
comp=#5 & #9, animal=#18 (190313d)
comp=#5 & #9, animal=#5 (180711f)
comp=#5 & #9, animal=#8 (180712e)
comp=#5 & #9, animal=#24 (190314d)
comp=#5 & #9, animal=#19 (190313e)
comp=#5 & #9, animal=#13 (190312e)
comp=#5 & #9, animal=#4 (180711d)
comp=#5 & #9, animal=#26 (190314f)
comp=#5 & #9, animal=#7 (180712d)
comp=#5 & #9, animal=#6 (180711g)
comp=#5 & #9, animal=#16 (190312h)
comp=#5 & #9, animal=#2 (180710f)
comp=#5 & #9, animal=#1 (180710a)
comp=#5 & #9, animal=#22 (190314b)
comp=#5 & #9, animal=#28 (190315e)
0
2000
4000
-500
0
500

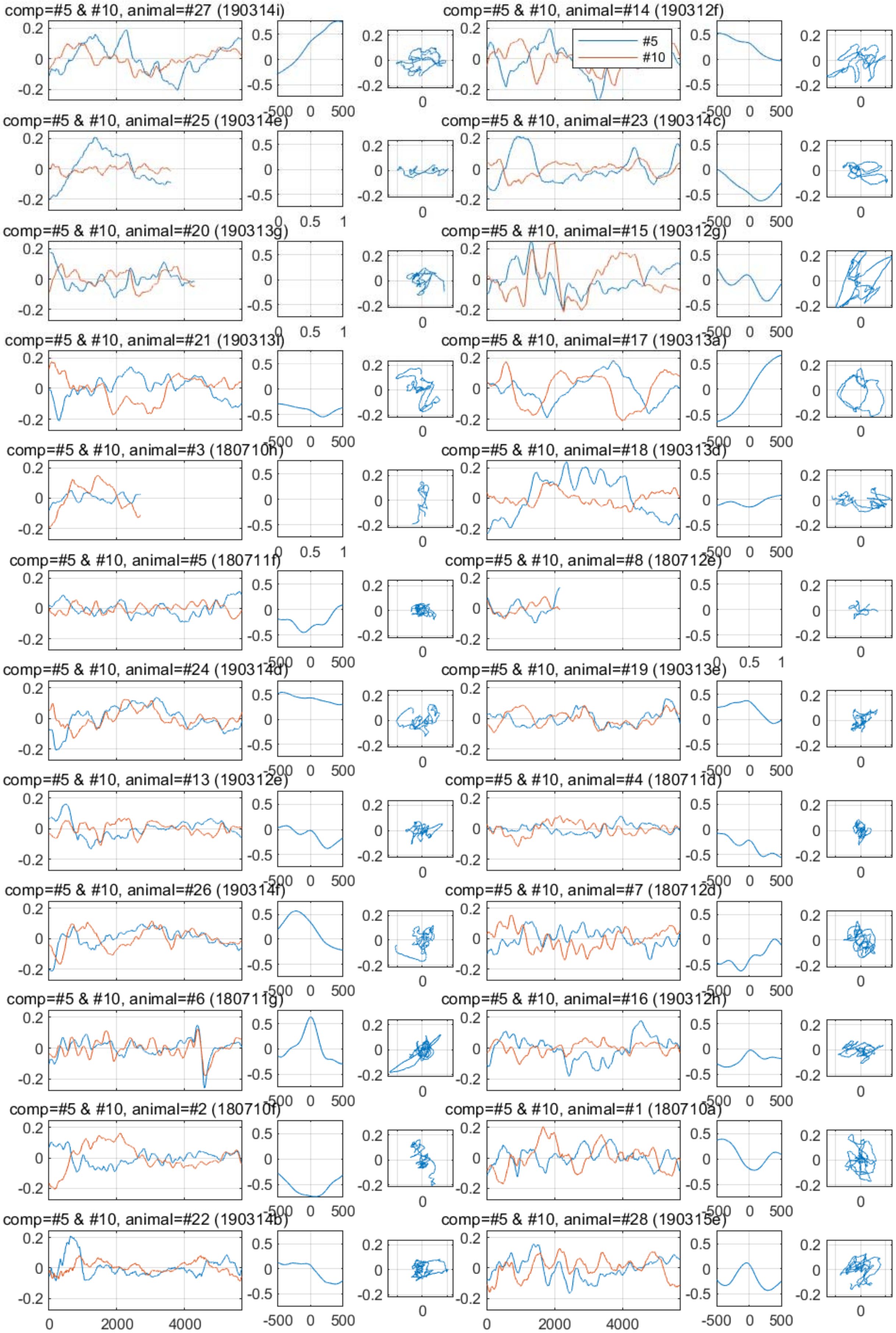
comp=#5 & #10, animal=#27 (190314i)
comp=#5 & #10, animal=#14 (190312f)
#5
#10
comp=#5 & #10, animal=#25 (190314e)
comp=#5 & #10, animal=#23 (190314c)
comp=#5 & #10, animal=#20 (190313g)
comp=#5 & #10, animal=#15 (190312g)
comp=#5 & #10, animal=#21 (190313f)
comp=#5 & #10, animal=#17 (190313a)
comp=#5 & #10, animal=#3 (180710h)
comp=#5 & #10, animal=#18 (190313d)
comp=#5 & #10, animal=#5 (180711f)
comp=#5 & #10, animal=#8 (180712e)
comp=#5 & #10, animal=#24 (190314d)
comp=#5 & #10, animal=#19 (190313e)
comp=#5 & #10, animal=#13 (190312e)
comp=#5 & #10, animal=#4 (180711d)
comp=#5 & #10, animal=#26 (190314f)
comp=#5 & #10, animal=#7 (180712d)
comp=#5 & #10, animal=#6 (180711g)
comp=#5 & #10, animal=#16 (190312h)
comp=#5 & #10, animal=#2 (180710f)
comp=#5 & #10, animal=#1 (180710a)
comp=#5 & #10, animal=#22 (190314b)
comp=#5 & #10, animal=#28 (190315e)
0
2000
4000
-500
0
500

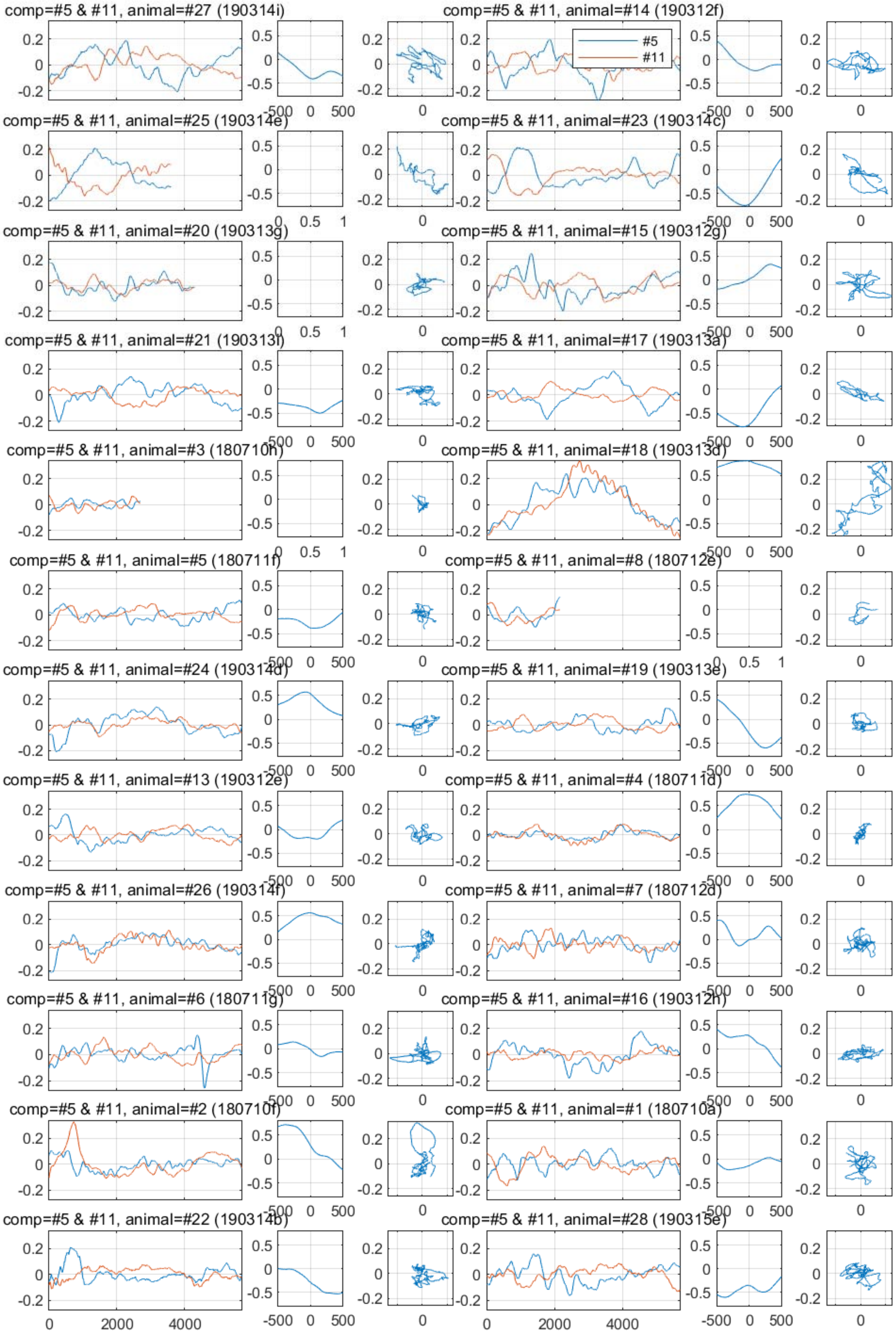
comp=#5 & #11, animal=#27 (190314i)
comp=#5 & #11, animal=#14 (190312f)
#5
#11
comp=#5 & #11, animal=#25 (190314e)
comp=#5 & #11, animal=#23 (190314c)
comp=#5 & #11, animal=#20 (190313g)
comp=#5 & #11, animal=#15 (190312g)
comp=#5 & #11, animal=#21 (190313i)
comp=#5 & #11, animal=#17 (190313a)
comp=#5 & #11, animal=#3 (180710h)
comp=#5 & #11, animal=#18 (190313d)
comp=#5 & #11, animal=#5 (180711f)
comp=#5 & #11, animal=#8 (180712e)
comp=#5 & #11, animal=#24 (190314d)
comp=#5 & #11, animal=#19 (190313e)
comp=#5 & #11, animal=#13 (190312e)
comp=#5 & #11, animal=#4 (180711d)
comp=#5 & #11, animal=#26 (190314f)
comp=#5 & #11, animal=#7 (180712d)
comp=#5 & #11, animal=#6 (180711g)
comp=#5 & #11, animal=#16 (190312h)
comp=#5 & #11, animal=#2 (180710f)
comp=#5 & #11, animal=#1 (180710a)
comp=#5 & #11, animal=#22 (190314b)
comp=#5 & #11, animal=#28 (190315e)

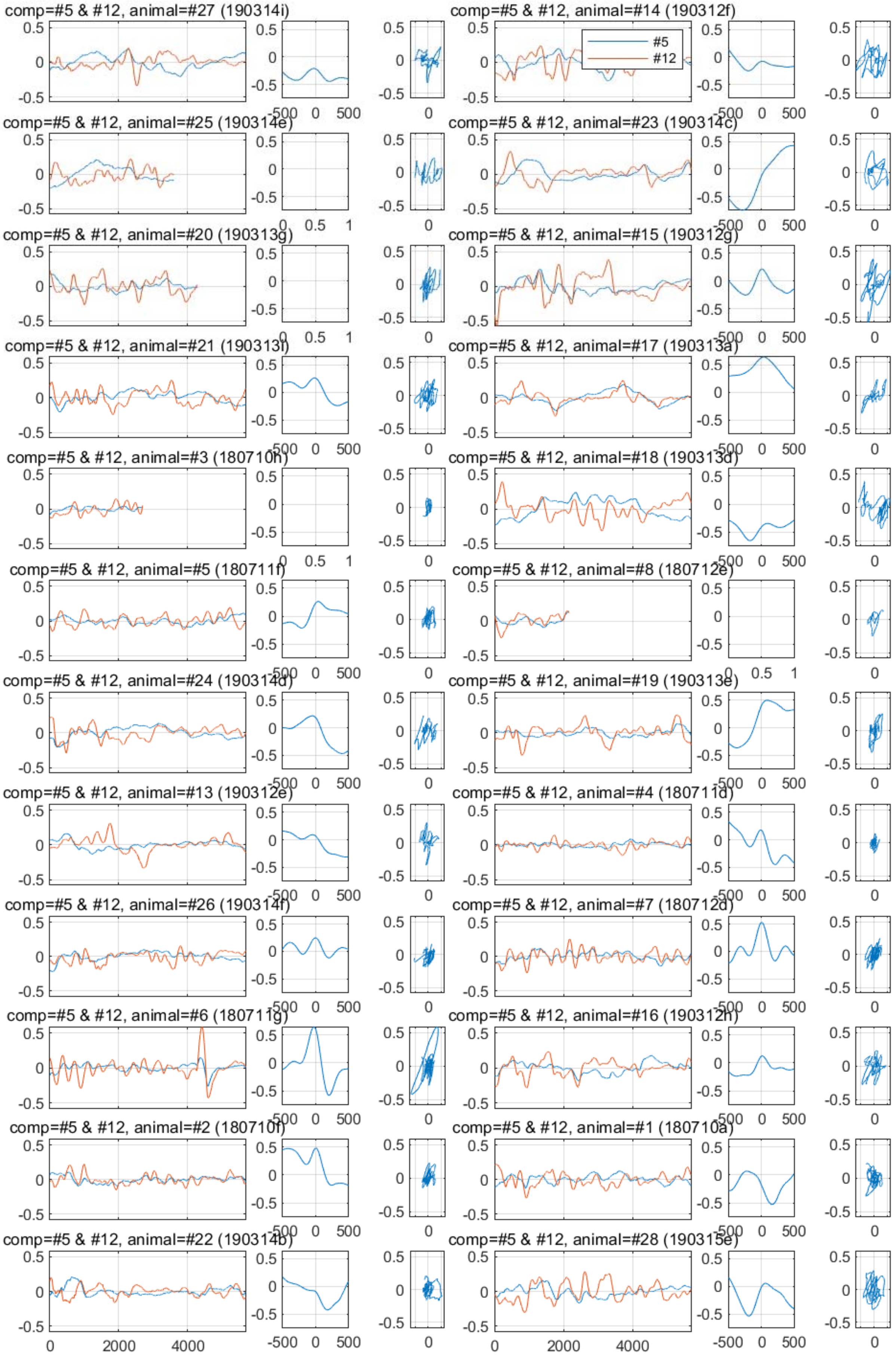
comp=#5 & #12, animal=#27 (190314i)
comp=#5 & #12, animal=#14 (190312f)
#5
#12
comp=#5 & #12, animal=#25 (190314e)
comp=#5 & #12, animal=#23 (190314c)
comp=#5 & #12, animal=#20 (190313g)
comp=#5 & #12, animal=#15 (190312g)
comp=#5 & #12, animal=#21 (190313i)
comp=#5 & #12, animal=#17 (190313a)
comp=#5 & #12, animal=#3 (180710h)
comp=#5 & #12, animal=#18 (190313d)
comp=#5 & #12, animal=#5 (180711f)
comp=#5 & #12, animal=#8 (180712e)
comp=#5 & #12, animal=#24 (190314d)
comp=#5 & #12, animal=#19 (190313e)
comp=#5 & #12, animal=#13 (190312e)
comp=#5 & #12, animal=#4 (180711d)
comp=#5 & #12, animal=#26 (190314f)
comp=#5 & #12, animal=#7 (180712d)
comp=#5 & #12, animal=#6 (180711g)
comp=#5 & #12, animal=#16 (190312h)
comp=#5 & #12, animal=#2 (180710f)
comp=#5 & #12, animal=#1 (180710a)
comp=#5 & #12, animal=#22 (190314b)
comp=#5 & #12, animal=#28 (190315e)
0
2000
4000
-500
0
500

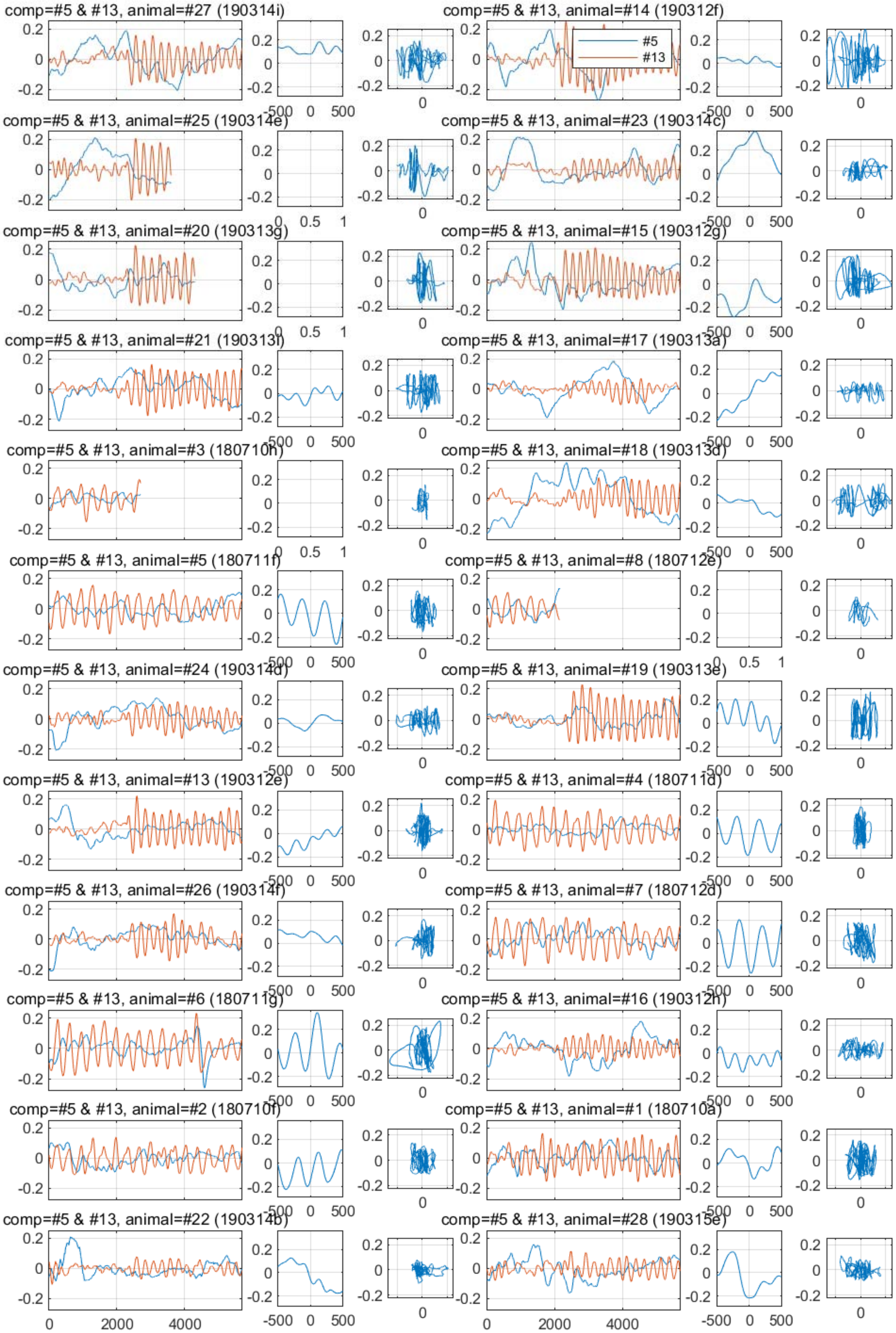
comp=#5 & #13, animal=#27 (190314i)
comp=#5 & #13, animal=#14 (190312f)
#5
#13
comp=#5 & #13, animal=#25 (190314e)
comp=#5 & #13, animal=#23 (190314c)
comp=#5 & #13, animal=#20 (190313g)
comp=#5 & #13, animal=#15 (190312g)
comp=#5 & #13, animal=#21 (190313f)
comp=#5 & #13, animal=#17 (190313a)
comp=#5 & #13, animal=#3 (180710h)
comp=#5 & #13, animal=#18 (190313d)
comp=#5 & #13, animal=#5 (180711f)
comp=#5 & #13, animal=#8 (180712e)
comp=#5 & #13, animal=#24 (190314d)
comp=#5 & #13, animal=#19 (190313e)
comp=#5 & #13, animal=#13 (190312e)
comp=#5 & #13, animal=#4 (180711d)
comp=#5 & #13, animal=#26 (190314f)
comp=#5 & #13, animal=#7 (180712d)
comp=#5 & #13, animal=#6 (180711g)
comp=#5 & #13, animal=#16 (190312h)
comp=#5 & #13, animal=#2 (180710f)
comp=#5 & #13, animal=#1 (180710a)
comp=#5 & #13, animal=#22 (190314b)
comp=#5 & #13, animal=#28 (190315e)

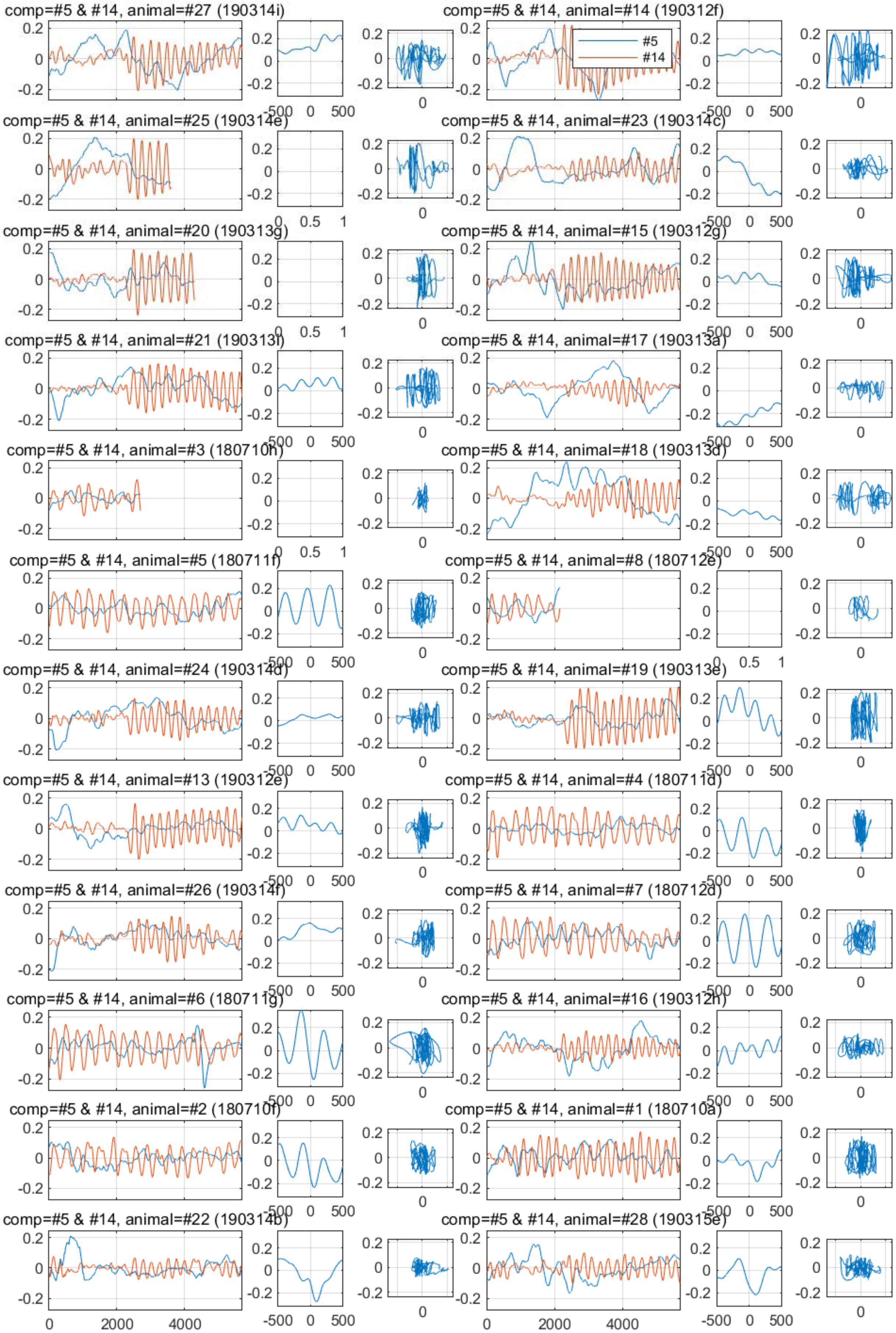
comp=#5 & #14, animal=#27 (190314i)
comp=#5 & #14, animal=#14 (190312f)
#5
#14
comp=#5 & #14, animal=#25 (190314e)
comp=#5 & #14, animal=#23 (190314c)
comp=#5 & #14, animal=#20 (190313g)
comp=#5 & #14, animal=#15 (190312g)
comp=#5 & #14, animal=#21 (190313f)
comp=#5 & #14, animal=#17 (190313a)
comp=#5 & #14, animal=#3 (180710h)
comp=#5 & #14, animal=#18 (190313d)
comp=#5 & #14, animal=#5 (180711f)
comp=#5 & #14, animal=#8 (180712e)
comp=#5 & #14, animal=#24 (190314d)
comp=#5 & #14, animal=#19 (190313e)
comp=#5 & #14, animal=#13 (190312e)
comp=#5 & #14, animal=#4 (180711d)
comp=#5 & #14, animal=#26 (190314f)
comp=#5 & #14, animal=#7 (180712d)
comp=#5 & #14, animal=#6 (180711g)
comp=#5 & #14, animal=#16 (190312h)
comp=#5 & #14, animal=#2 (180710f)
comp=#5 & #14, animal=#1 (180710a)
comp=#5 & #14, animal=#22 (190314b)
comp=#5 & #14, animal=#28 (190315e)

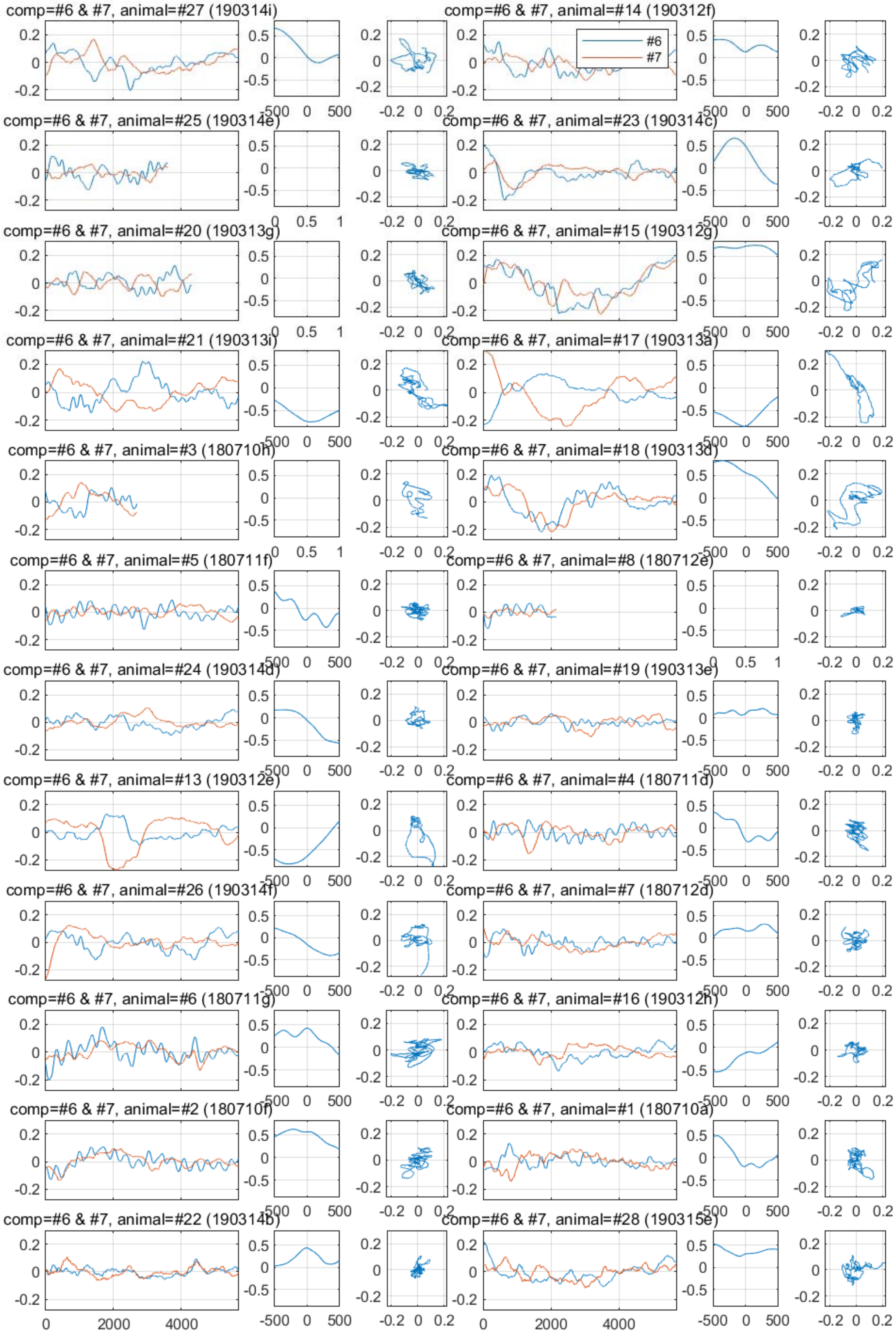

comp=#6 & #7, animal=#27 (190314i)
comp=#6 & #7, animal=#14 (190312f)
#6
#7
comp=#6 & #7, animal=#25 (190314e)
comp=#6 & #7, animal=#23 (190314c)
comp=#6 & #7, animal=#20 (190313g)
comp=#6 & #7, animal=#15 (190312g)
comp=#6 & #7, animal=#21 (190313i)
comp=#6 & #7, animal=#17 (190313a)
comp=#6 & #7, animal=#3 (180710h)
comp=#6 & #7, animal=#18 (190313d)
comp=#6 & #7, animal=#5 (180711f)
comp=#6 & #7, animal=#8 (180712e)
comp=#6 & #7, animal=#24 (190314d)
comp=#6 & #7, animal=#19 (190313e)
comp=#6 & #7, animal=#13 (190312e)
comp=#6 & #7, animal=#4 (180711d)
comp=#6 & #7, animal=#26 (190314f)
comp=#6 & #7, animal=#7 (180712d)
comp=#6 & #7, animal=#6 (180711g)
comp=#6 & #7, animal=#16 (190312h)
comp=#6 & #7, animal=#2 (180710f)
comp=#6 & #7, animal=#1 (180710a)
comp=#6 & #7, animal=#22 (190314b)
comp=#6 & #7, animal=#28 (190315e)
0
2000
4000
-500
0
500
-0.2
0
0.2
0.5
-0.5
1

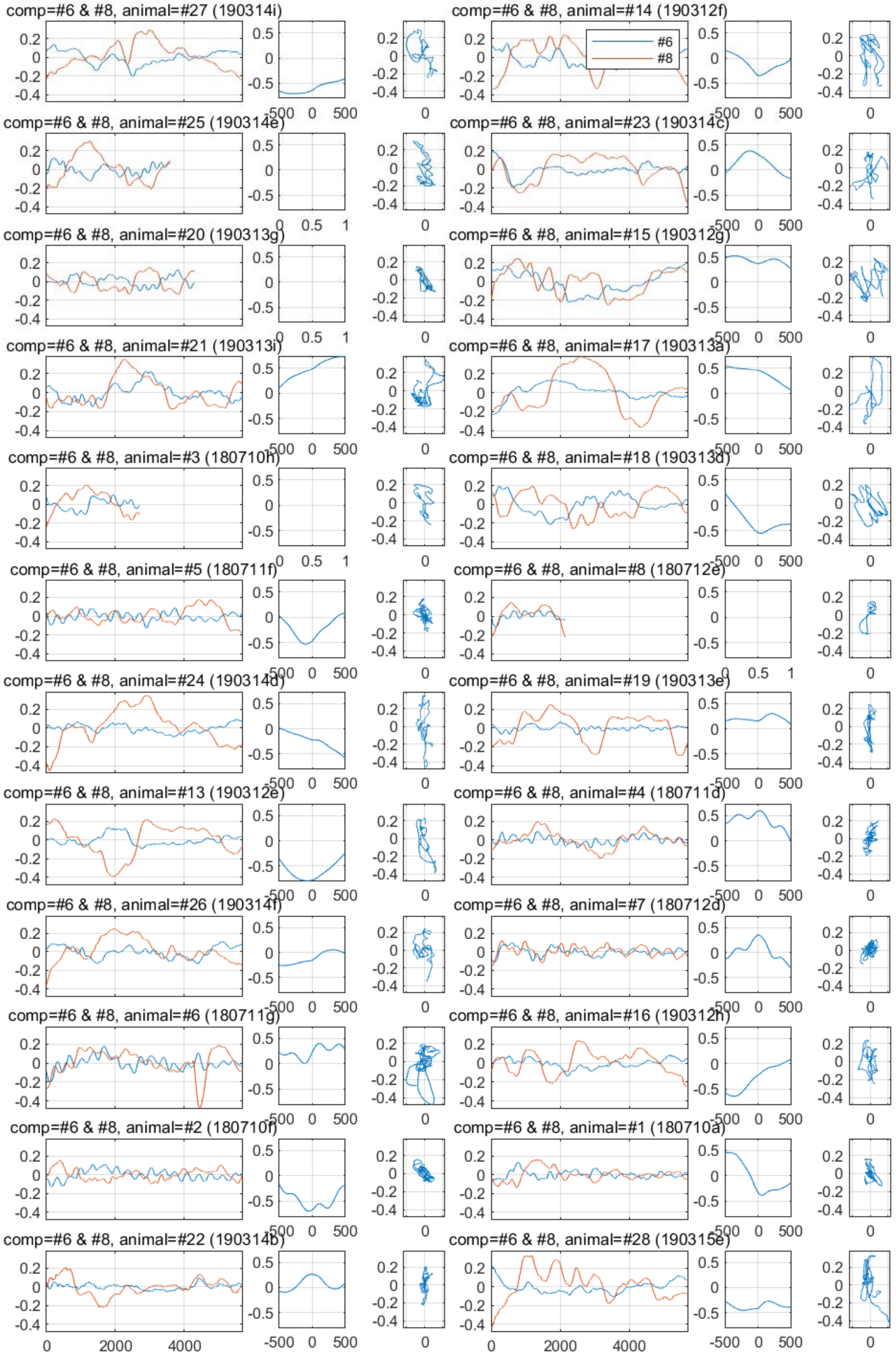
comp=#6 & #8, animal=#27 (190314i)
comp=#6 & #8, animal=#14 (190312f)
#6
#8
comp=#6 & #8, animal=#25 (190314e)
comp=#6 & #8, animal=#23 (190314c)
comp=#6 & #8, animal=#20 (190313g)
comp=#6 & #8, animal=#15 (190312g)
comp=#6 & #8, animal=#21 (190313i)
comp=#6 & #8, animal=#17 (190313a)
comp=#6 & #8, animal=#3 (180710h)
comp=#6 & #8, animal=#18 (190313d)
comp=#6 & #8, animal=#5 (180711f)
comp=#6 & #8, animal=#8 (180712e)
comp=#6 & #8, animal=#24 (190314d)
comp=#6 & #8, animal=#19 (190313e)
comp=#6 & #8, animal=#13 (190312e)
comp=#6 & #8, animal=#4 (180711d)
comp=#6 & #8, animal=#26 (190314f)
comp=#6 & #8, animal=#7 (180712d)
comp=#6 & #8, animal=#6 (180711g)
comp=#6 & #8, animal=#16 (190312h)
comp=#6 & #8, animal=#2 (180710f)
comp=#6 & #8, animal=#1 (180710a)
comp=#6 & #8, animal=#22 (190314b)
comp=#6 & #8, animal=#28 (190315e)
0
2000
4000
-500
0
500

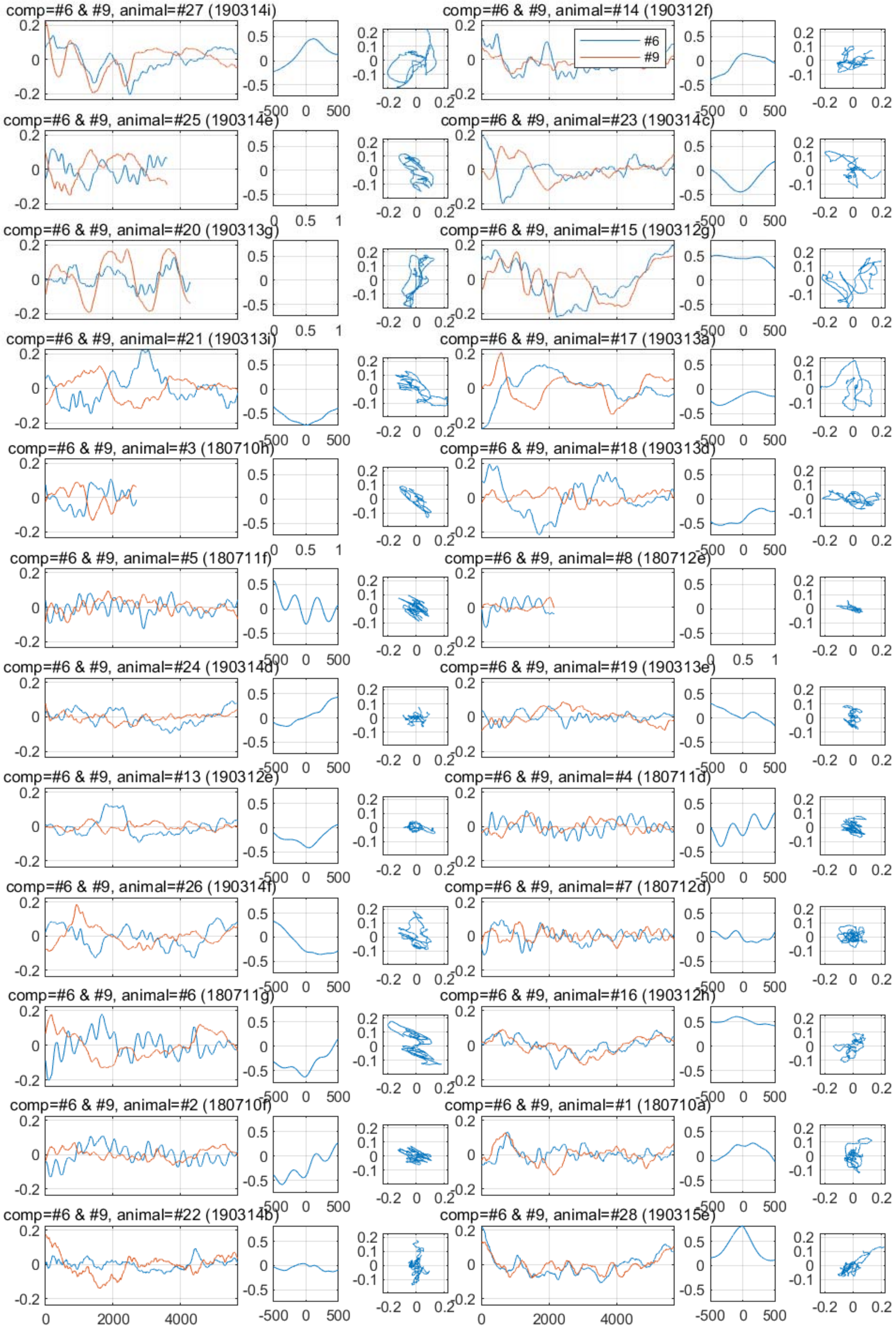
comp=#6 & #9, animal=#27 (190314i)
comp=#6 & #9, animal=#14 (190312f)
#6
#9
comp=#6 & #9, animal=#25 (190314e)
comp=#6 & #9, animal=#23 (190314c)
comp=#6 & #9, animal=#20 (190313g)
comp=#6 & #9, animal=#15 (190312g)
comp=#6 & #9, animal=#21 (190313i)
comp=#6 & #9, animal=#17 (190313a)
comp=#6 & #9, animal=#3 (180710h)
comp=#6 & #9, animal=#18 (190313d)
comp=#6 & #9, animal=#5 (180711f)
comp=#6 & #9, animal=#8 (180712e)
comp=#6 & #9, animal=#24 (190314d)
comp=#6 & #9, animal=#19 (190313e)
comp=#6 & #9, animal=#13 (190312e)
comp=#6 & #9, animal=#4 (180711d)
comp=#6 & #9, animal=#26 (190314f)
comp=#6 & #9, animal=#7 (180712d)
comp=#6 & #9, animal=#6 (180711g)
comp=#6 & #9, animal=#16 (190312h)
comp=#6 & #9, animal=#2 (180710f)
comp=#6 & #9, animal=#1 (180710a)
comp=#6 & #9, animal=#22 (190314b)
comp=#6 & #9, animal=#28 (190315e)

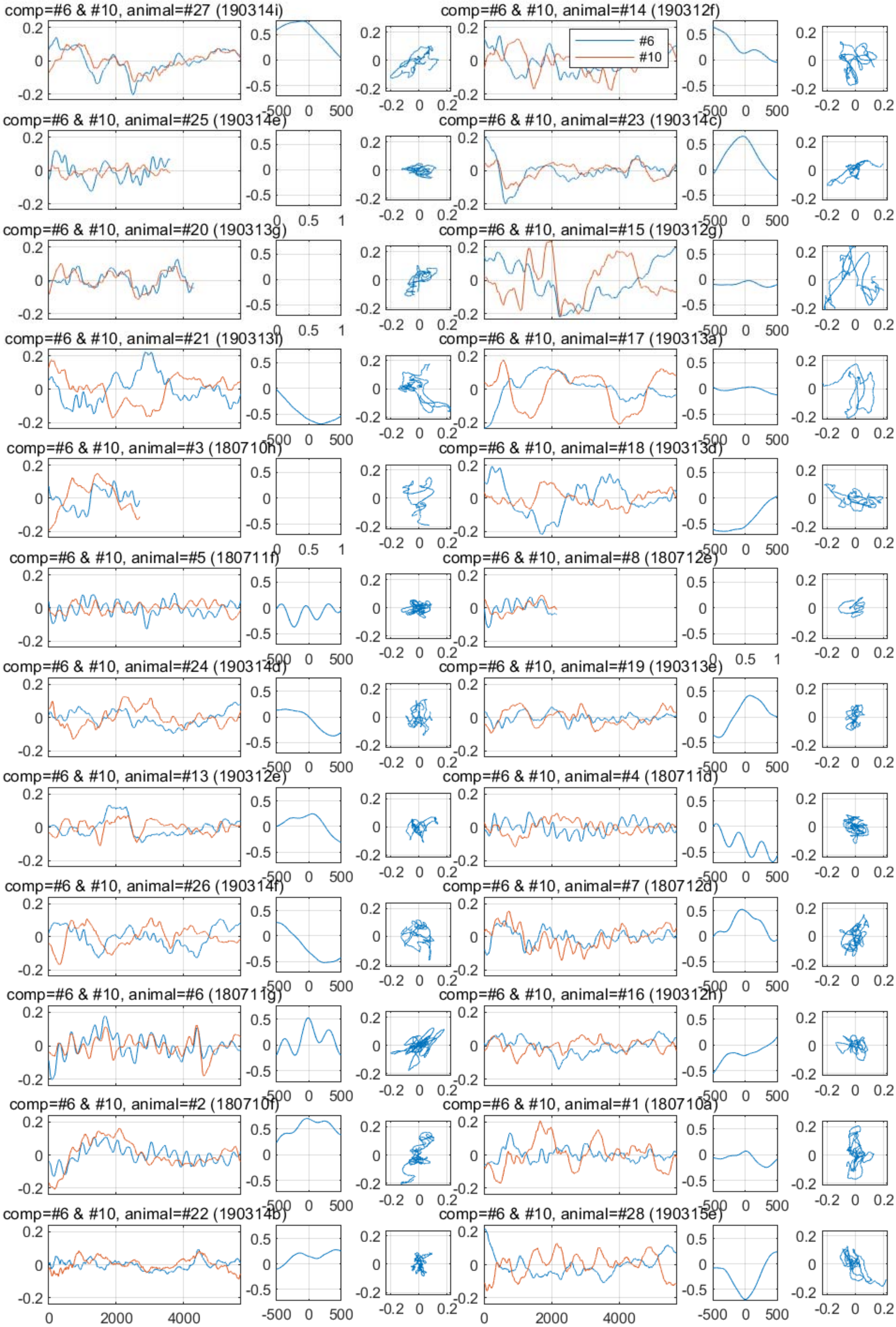

comp=#6 & #10, animal=#27 (190314i)
comp=#6 & #10, animal=#14 (190312f)
#6
#10
comp=#6 & #10, animal=#25 (190314e)
comp=#6 & #10, animal=#23 (190314c)
comp=#6 & #10, animal=#20 (190313g)
comp=#6 & #10, animal=#15 (190312g)
comp=#6 & #10, animal=#21 (190313i)
comp=#6 & #10, animal=#17 (190313a)
comp=#6 & #10, animal=#3 (180710h)
comp=#6 & #10, animal=#18 (190313d)
comp=#6 & #10, animal=#5 (180711f)
comp=#6 & #10, animal=#8 (180712e)
comp=#6 & #10, animal=#24 (190314d)
comp=#6 & #10, animal=#19 (190313e)
comp=#6 & #10, animal=#13 (190312e)
comp=#6 & #10, animal=#4 (180711d)
comp=#6 & #10, animal=#26 (190314f)
comp=#6 & #10, animal=#7 (180712d)
comp=#6 & #10, animal=#6 (180711g)
comp=#6 & #10, animal=#16 (190312h)
comp=#6 & #10, animal=#2 (180710f)
comp=#6 & #10, animal=#1 (180710a)
comp=#6 & #10, animal=#22 (190314b)
comp=#6 & #10, animal=#28 (190315e)

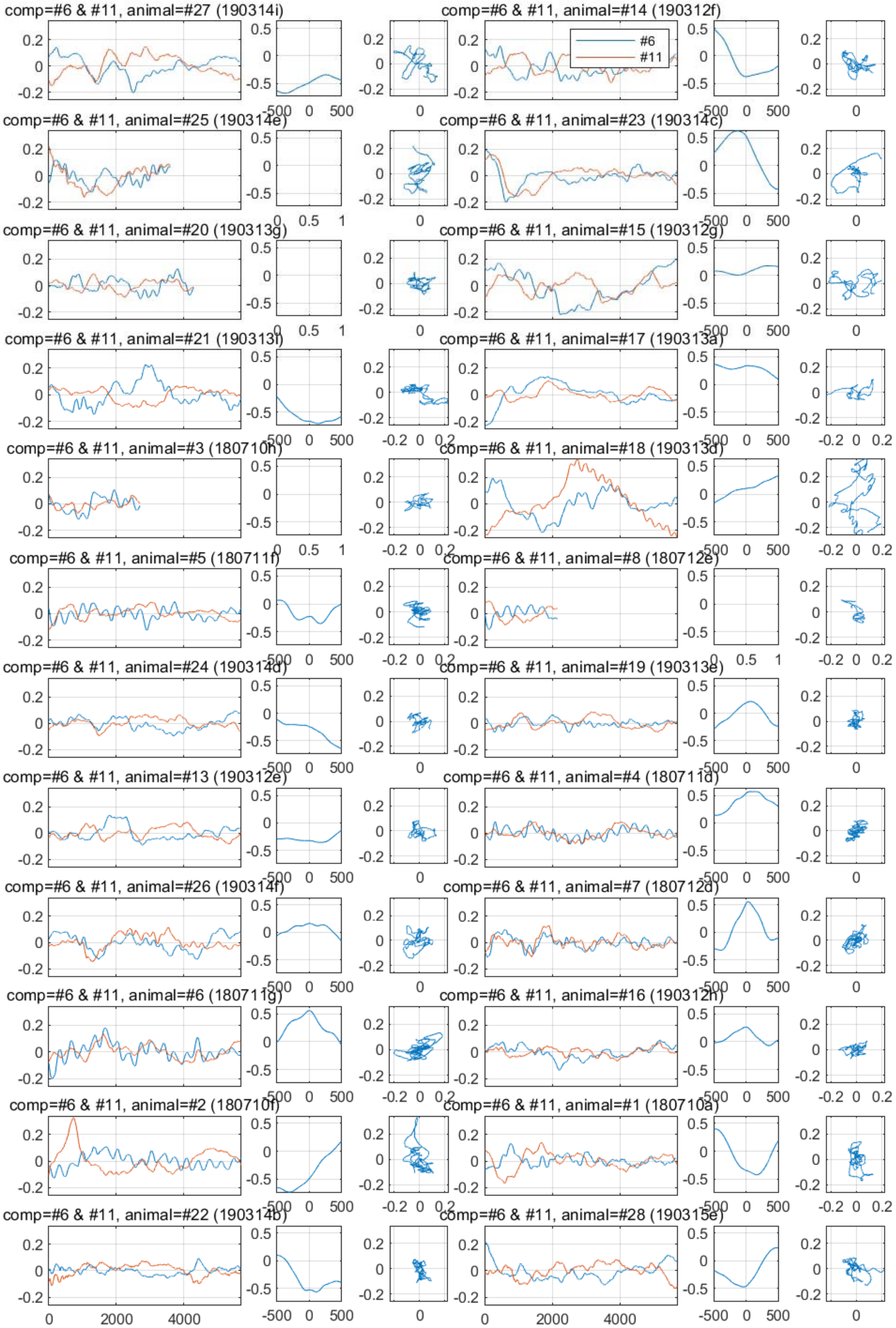

comp=#6 & #11, animal=#27 (190314i)
comp=#6 & #11, animal=#14 (190312f)
#6
#11
comp=#6 & #11, animal=#25 (190314e)
comp=#6 & #11, animal=#23 (190314c)
comp=#6 & #11, animal=#20 (190313g)
comp=#6 & #11, animal=#15 (190312g)
comp=#6 & #11, animal=#21 (190313f)
comp=#6 & #11, animal=#17 (190313a)
comp=#6 & #11, animal=#3 (180710h)
comp=#6 & #11, animal=#18 (190313d)
comp=#6 & #11, animal=#5 (180711f)
comp=#6 & #11, animal=#8 (180712e)
comp=#6 & #11, animal=#24 (190314d)
comp=#6 & #11, animal=#19 (190313e)
comp=#6 & #11, animal=#13 (190312e)
comp=#6 & #11, animal=#4 (180711d)
comp=#6 & #11, animal=#26 (190314f)
comp=#6 & #11, animal=#7 (180712d)
comp=#6 & #11, animal=#6 (180711g)
comp=#6 & #11, animal=#16 (190312h)
comp=#6 & #11, animal=#2 (180710f)
comp=#6 & #11, animal=#1 (180710a)
comp=#6 & #11, animal=#22 (190314b)
comp=#6 & #11, animal=#28 (190315e)

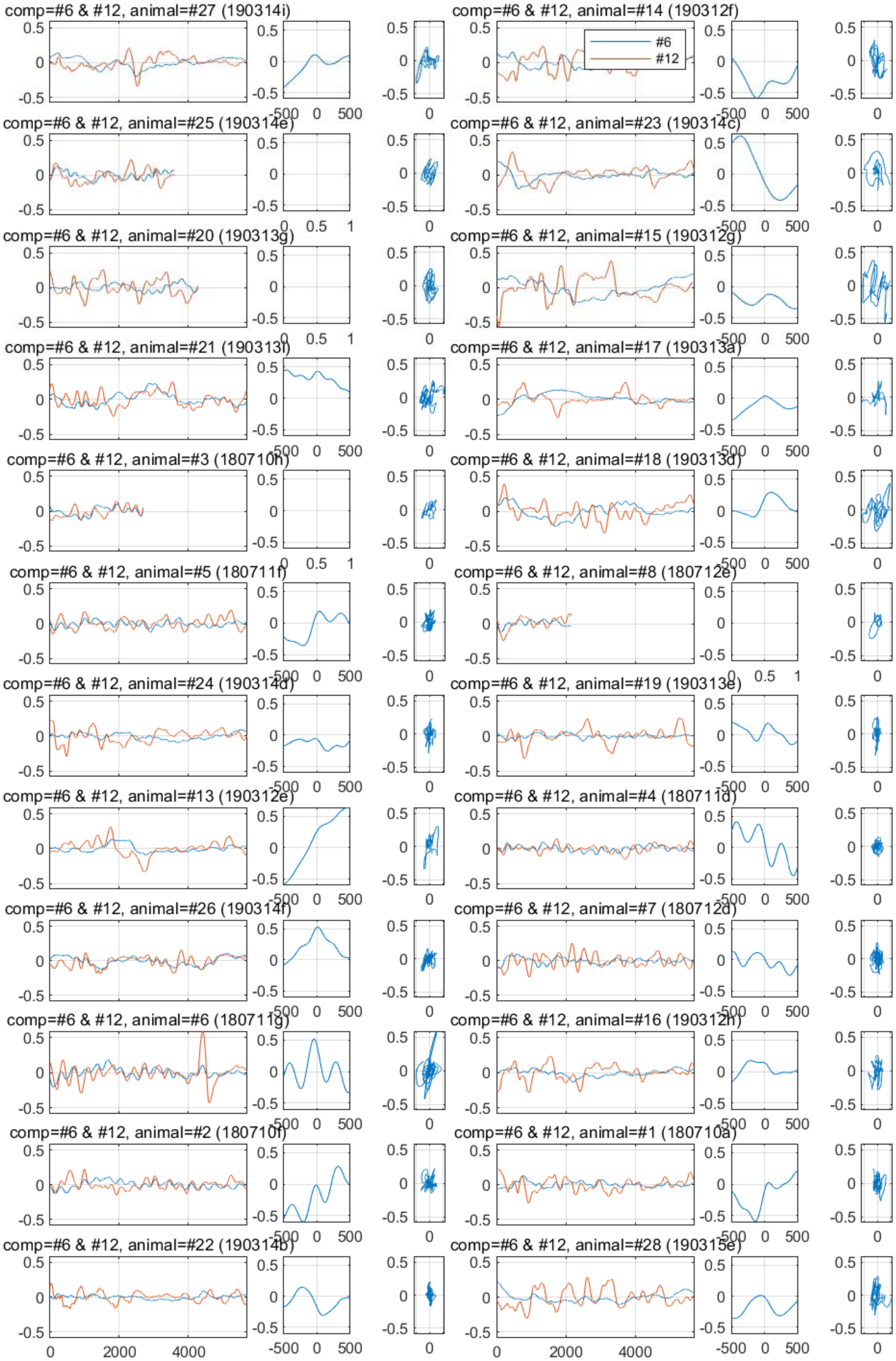

comp=#6 & #12, animal=#27 (190314i)
comp=#6 & #12, animal=#14 (190312f)
#6
#12
comp=#6 & #12, animal=#25 (190314e)
comp=#6 & #12, animal=#23 (190314c)
comp=#6 & #12, animal=#20 (190313g)
comp=#6 & #12, animal=#15 (190312g)
comp=#6 & #12, animal=#21 (190313i)
comp=#6 & #12, animal=#17 (190313a)
comp=#6 & #12, animal=#3 (180710h)
comp=#6 & #12, animal=#18 (190313d)
comp=#6 & #12, animal=#5 (180711f)
comp=#6 & #12, animal=#8 (180712e)
comp=#6 & #12, animal=#24 (190314d)
comp=#6 & #12, animal=#19 (190313e)
comp=#6 & #12, animal=#13 (190312e)
comp=#6 & #12, animal=#4 (180711d)
comp=#6 & #12, animal=#26 (190314f)
comp=#6 & #12, animal=#7 (180712d)
comp=#6 & #12, animal=#6 (180711g)
comp=#6 & #12, animal=#16 (190312h)
comp=#6 & #12, animal=#2 (180710f)
comp=#6 & #12, animal=#1 (180710a)
comp=#6 & #12, animal=#22 (190314b)
comp=#6 & #12, animal=#28 (190315e)

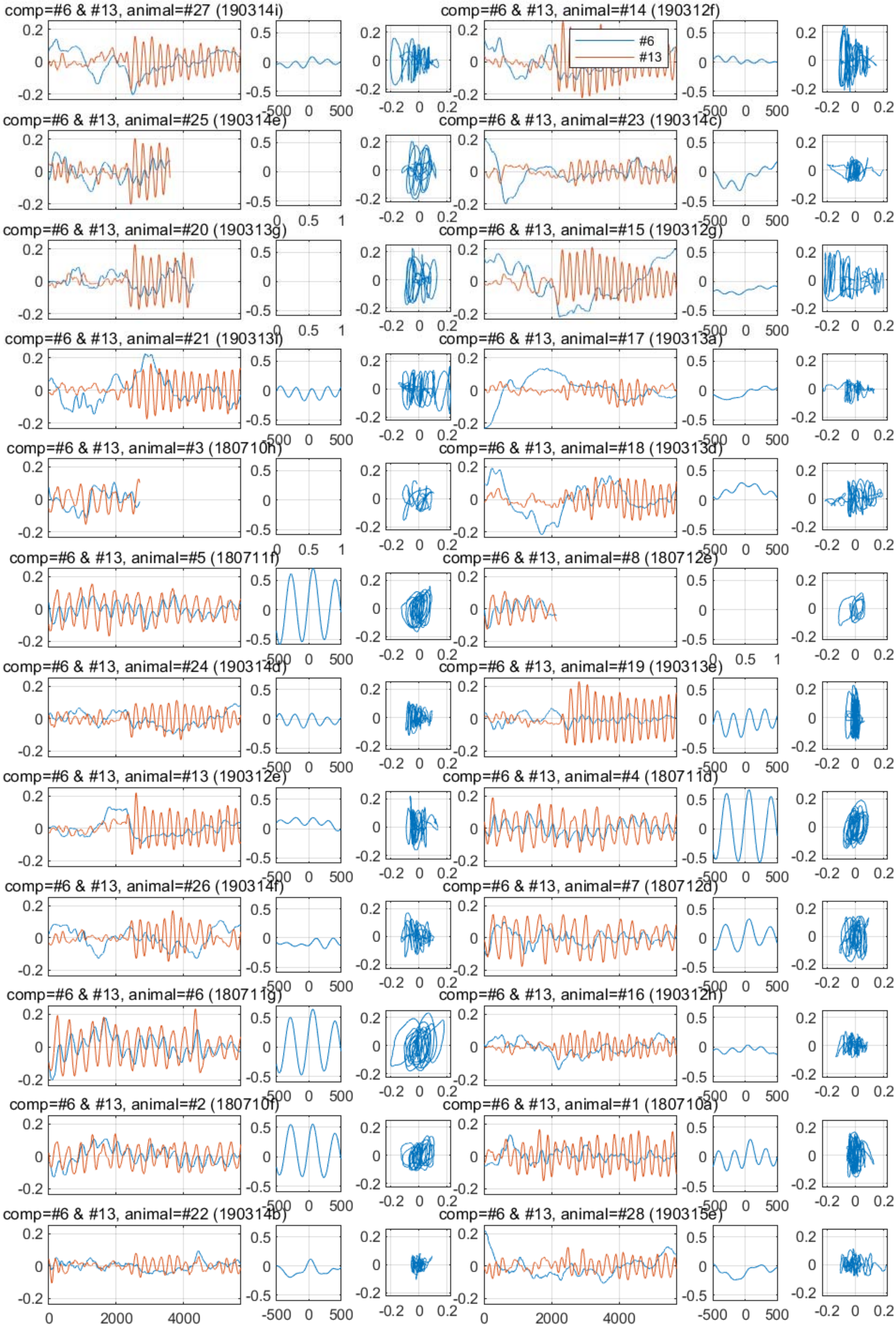
comp=#6 & #13, animal=#27 (190314i)
comp=#6 & #13, animal=#14 (190312f)
#6
#13
comp=#6 & #13, animal=#25 (190314e)
comp=#6 & #13, animal=#23 (190314c)
comp=#6 & #13, animal=#20 (190313g)
comp=#6 & #13, animal=#15 (190312g)
comp=#6 & #13, animal=#21 (190313i)
comp=#6 & #13, animal=#17 (190313a)
comp=#6 & #13, animal=#3 (180710h)
comp=#6 & #13, animal=#18 (190313d)
comp=#6 & #13, animal=#5 (180711f)
comp=#6 & #13, animal=#8 (180712e)
comp=#6 & #13, animal=#24 (190314d)
comp=#6 & #13, animal=#19 (190313e)
comp=#6 & #13, animal=#13 (190312e)
comp=#6 & #13, animal=#4 (180711d)
comp=#6 & #13, animal=#26 (190314f)
comp=#6 & #13, animal=#7 (180712d)
comp=#6 & #13, animal=#6 (180711g)
comp=#6 & #13, animal=#16 (190312h)
comp=#6 & #13, animal=#2 (180710f)
comp=#6 & #13, animal=#1 (180710a)
comp=#6 & #13, animal=#22 (190314b)
comp=#6 & #13, animal=#28 (190315e)
0
2000
4000
-500
0
500
-0.2
0
0.2

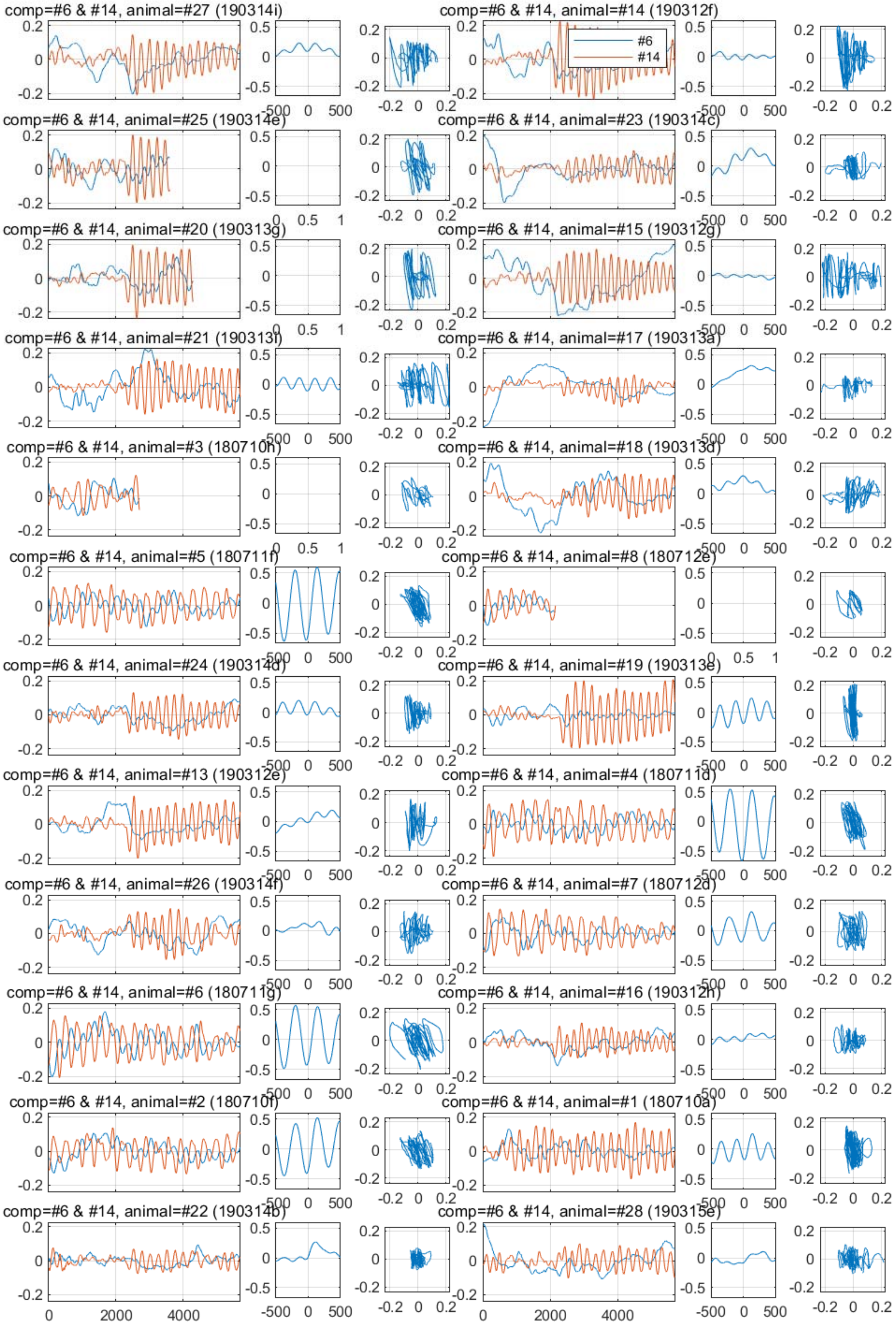
comp=#6 & #14, animal=#27 (190314i)
comp=#6 & #14, animal=#14 (190312f)
#6
#14
comp=#6 & #14, animal=#25 (190314e)
comp=#6 & #14, animal=#23 (190314c)
comp=#6 & #14, animal=#20 (190313g)
comp=#6 & #14, animal=#15 (190312g)
comp=#6 & #14, animal=#21 (190313i)
comp=#6 & #14, animal=#17 (190313a)
comp=#6 & #14, animal=#3 (180710h)
comp=#6 & #14, animal=#18 (190313d)
comp=#6 & #14, animal=#5 (180711f)
comp=#6 & #14, animal=#8 (180712e)
comp=#6 & #14, animal=#24 (190314d)
comp=#6 & #14, animal=#19 (190313e)
comp=#6 & #14, animal=#13 (190312e)
comp=#6 & #14, animal=#4 (180711d)
comp=#6 & #14, animal=#26 (190314f)
comp=#6 & #14, animal=#7 (180712d)
comp=#6 & #14, animal=#6 (180711g)
comp=#6 & #14, animal=#16 (190312h)
comp=#6 & #14, animal=#2 (180710f)
comp=#6 & #14, animal=#1 (180710a)
comp=#6 & #14, animal=#22 (190314b)
comp=#6 & #14, animal=#28 (190315e)

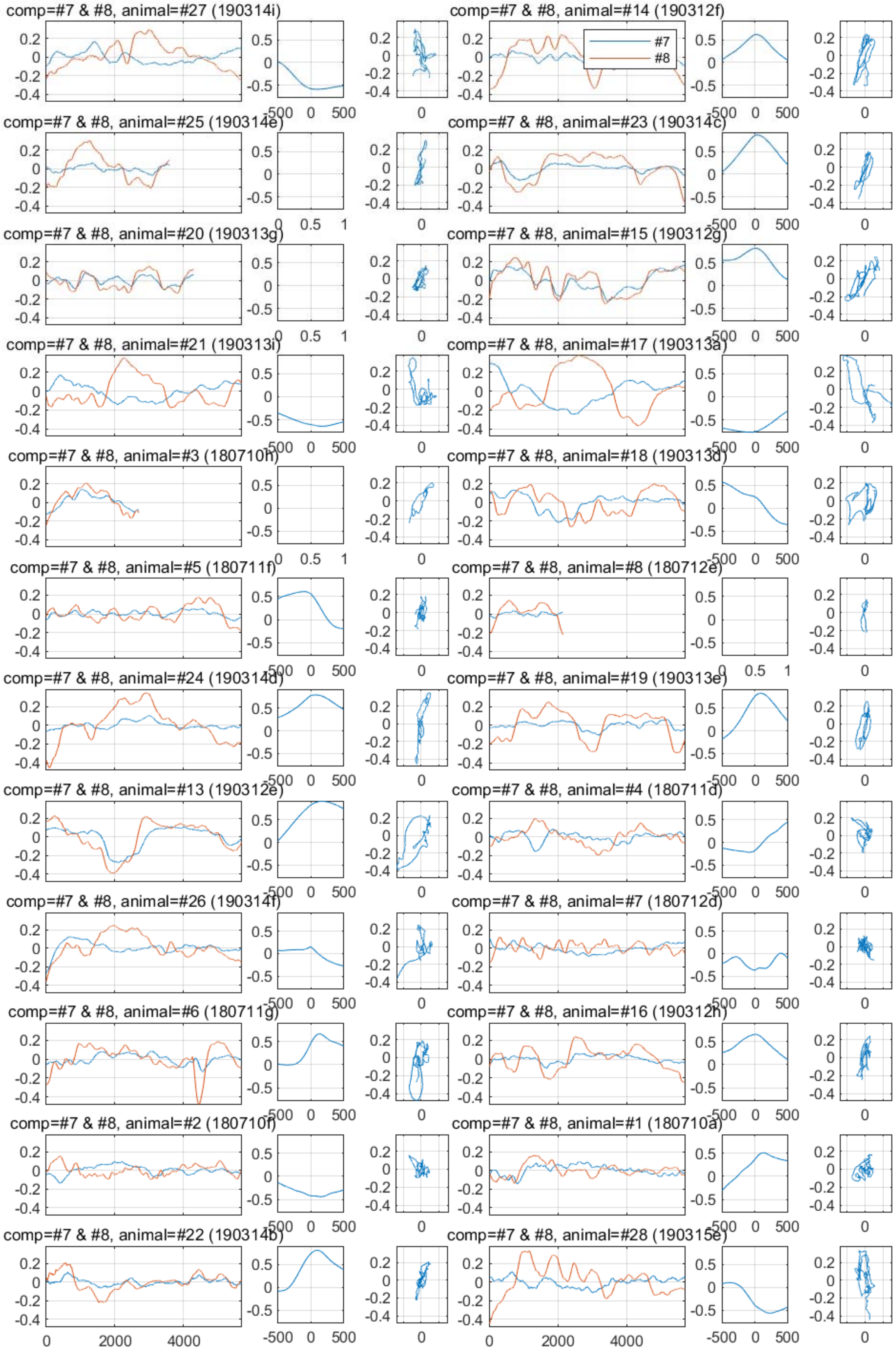
comp=#7 & #8, animal=#27 (190314i)
comp=#7 & #8, animal=#14 (190312f)
#7
#8
comp=#7 & #8, animal=#25 (190314e)
comp=#7 & #8, animal=#23 (190314c)
comp=#7 & #8, animal=#20 (190313g)
comp=#7 & #8, animal=#15 (190312g)
comp=#7 & #8, animal=#21 (190313i)
comp=#7 & #8, animal=#17 (190313a)
comp=#7 & #8, animal=#3 (180710h)
comp=#7 & #8, animal=#18 (190313d)
comp=#7 & #8, animal=#5 (180711f)
comp=#7 & #8, animal=#8 (180712e)
comp=#7 & #8, animal=#24 (190314d)
comp=#7 & #8, animal=#19 (190313e)
comp=#7 & #8, animal=#13 (190312e)
comp=#7 & #8, animal=#4 (180711d)
comp=#7 & #8, animal=#26 (190314f)
comp=#7 & #8, animal=#7 (180712d)
comp=#7 & #8, animal=#6 (180711g)
comp=#7 & #8, animal=#16 (190312h)
comp=#7 & #8, animal=#2 (180710f)
comp=#7 & #8, animal=#1 (180710a)
comp=#7 & #8, animal=#22 (190314b)
comp=#7 & #8, animal=#28 (190315e)
0
2000
4000
-500
0
500

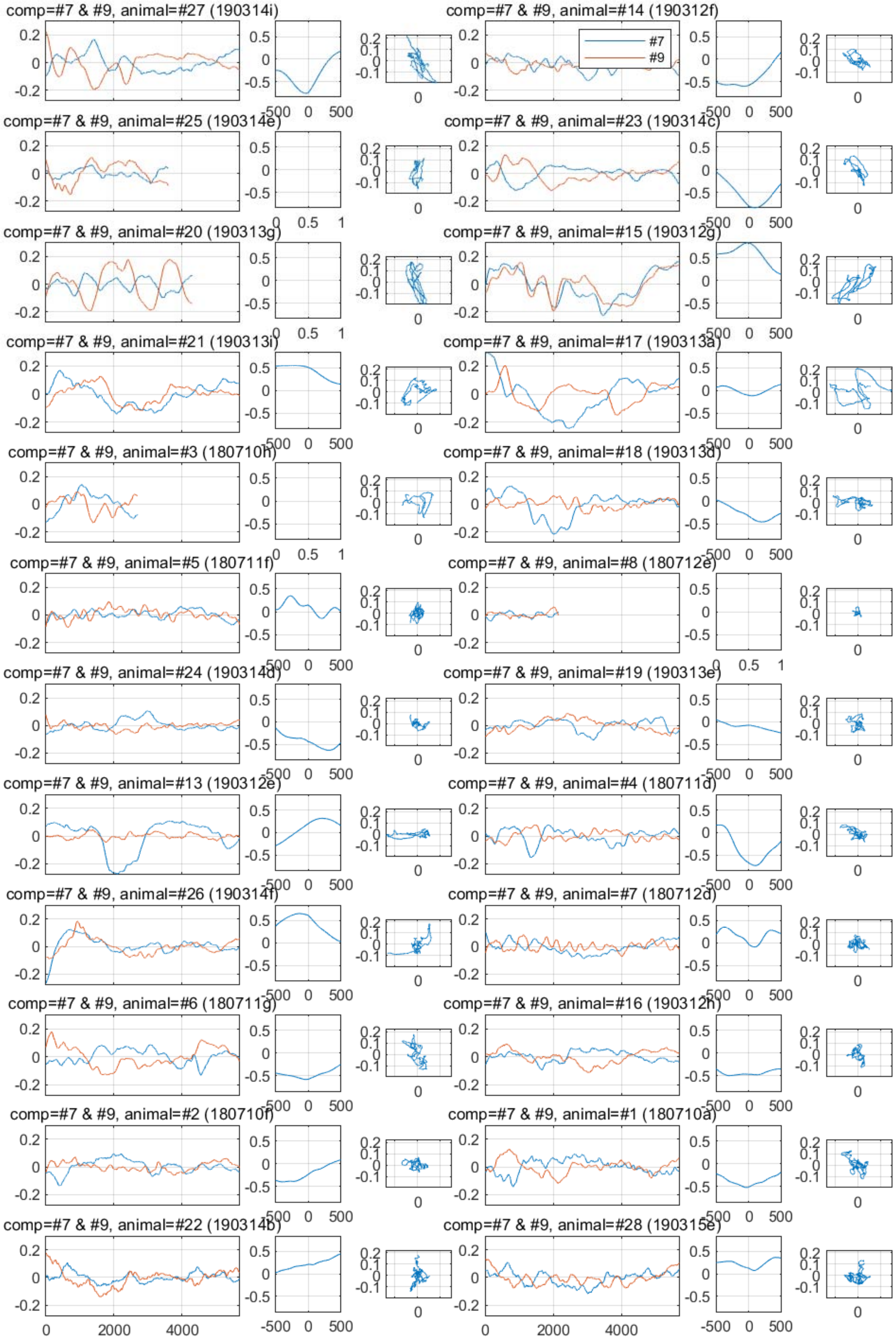
comp=#7 & #9, animal=#27 (190314i)
comp=#7 & #9, animal=#14 (190312f)
#7
#9
comp=#7 & #9, animal=#25 (190314e)
comp=#7 & #9, animal=#23 (190314c)
comp=#7 & #9, animal=#20 (190313g)
comp=#7 & #9, animal=#15 (190312g)
comp=#7 & #9, animal=#21 (190313i)
comp=#7 & #9, animal=#17 (190313a)
comp=#7 & #9, animal=#3 (180710h)
comp=#7 & #9, animal=#18 (190313d)
comp=#7 & #9, animal=#5 (180711f)
comp=#7 & #9, animal=#8 (180712e)
comp=#7 & #9, animal=#24 (190314d)
comp=#7 & #9, animal=#19 (190313e)
comp=#7 & #9, animal=#13 (190312e)
comp=#7 & #9, animal=#4 (180711d)
comp=#7 & #9, animal=#26 (190314f)
comp=#7 & #9, animal=#7 (180712d)
comp=#7 & #9, animal=#6 (180711g)
comp=#7 & #9, animal=#16 (190312h)
comp=#7 & #9, animal=#2 (180710f)
comp=#7 & #9, animal=#1 (180710a)
comp=#7 & #9, animal=#22 (190314b)
comp=#7 & #9, animal=#28 (190315e)

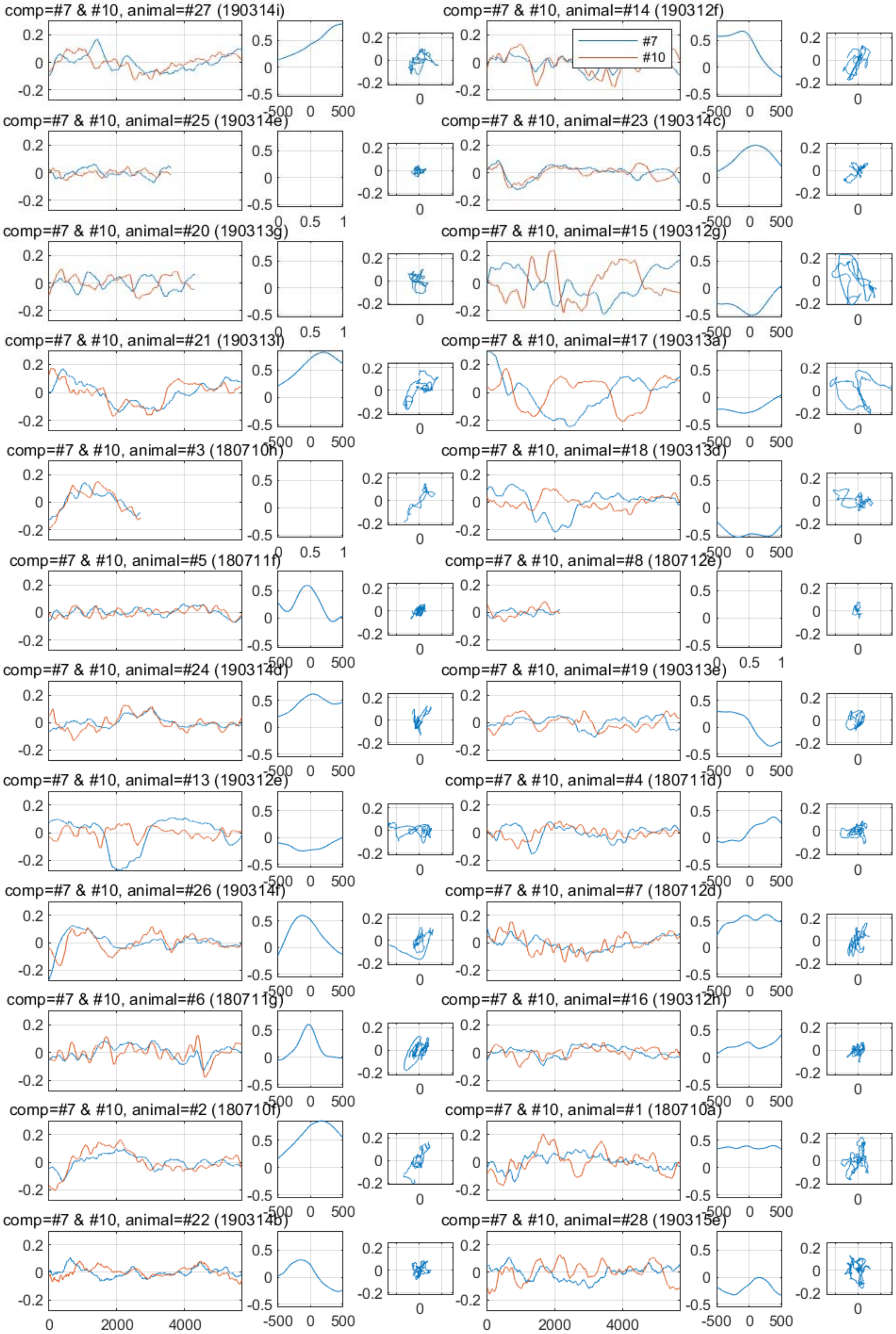
comp=#7 & #10, animal=#27 (190314i)
comp=#7 & #10, animal=#14 (190312f)
#7
#10
comp=#7 & #10, animal=#25 (190314e)
comp=#7 & #10, animal=#23 (190314c)
comp=#7 & #10, animal=#20 (190313g)
comp=#7 & #10, animal=#15 (190312g)
comp=#7 & #10, animal=#21 (190313i)
comp=#7 & #10, animal=#17 (190313a)
comp=#7 & #10, animal=#3 (180710h)
comp=#7 & #10, animal=#18 (190313d)
comp=#7 & #10, animal=#5 (180711f)
comp=#7 & #10, animal=#8 (180712e)
comp=#7 & #10, animal=#24 (190314d)
comp=#7 & #10, animal=#19 (190313e)
comp=#7 & #10, animal=#13 (190312e)
comp=#7 & #10, animal=#4 (180711d)
comp=#7 & #10, animal=#26 (190314f)
comp=#7 & #10, animal=#7 (180712d)
comp=#7 & #10, animal=#6 (180711g)
comp=#7 & #10, animal=#16 (190312h)
comp=#7 & #10, animal=#2 (180710f)
comp=#7 & #10, animal=#1 (180710a)
comp=#7 & #10, animal=#22 (190314b)
comp=#7 & #10, animal=#28 (190315e)

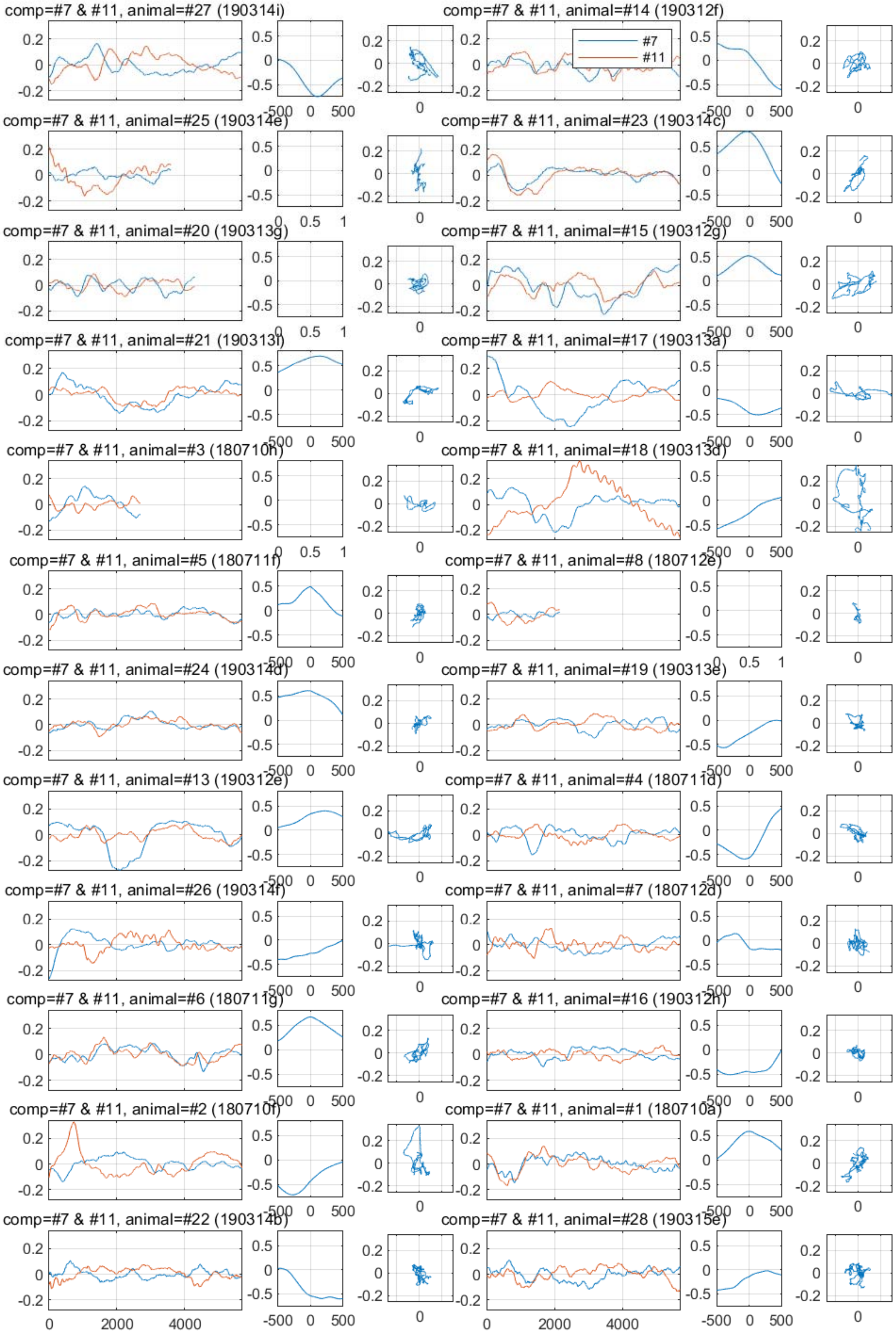
comp=#7 & #11, animal=#27 (190314i)
comp=#7 & #11, animal=#14 (190312f)
#7
#11
comp=#7 & #11, animal=#25 (190314e)
comp=#7 & #11, animal=#23 (190314c)
comp=#7 & #11, animal=#20 (190313g)
comp=#7 & #11, animal=#15 (190312g)
comp=#7 & #11, animal=#21 (190313f)
comp=#7 & #11, animal=#17 (190313a)
comp=#7 & #11, animal=#3 (180710h)
comp=#7 & #11, animal=#18 (190313d)
comp=#7 & #11, animal=#5 (180711f)
comp=#7 & #11, animal=#8 (180712e)
comp=#7 & #11, animal=#24 (190314d)
comp=#7 & #11, animal=#19 (190313e)
comp=#7 & #11, animal=#13 (190312e)
comp=#7 & #11, animal=#4 (180711d)
comp=#7 & #11, animal=#26 (190314f)
comp=#7 & #11, animal=#7 (180712d)
comp=#7 & #11, animal=#6 (180711g)
comp=#7 & #11, animal=#16 (190312h)
comp=#7 & #11, animal=#2 (180710f)
comp=#7 & #11, animal=#1 (180710a)
comp=#7 & #11, animal=#22 (190314b)
comp=#7 & #11, animal=#28 (190315e)
0
2000
4000
-500
0
500

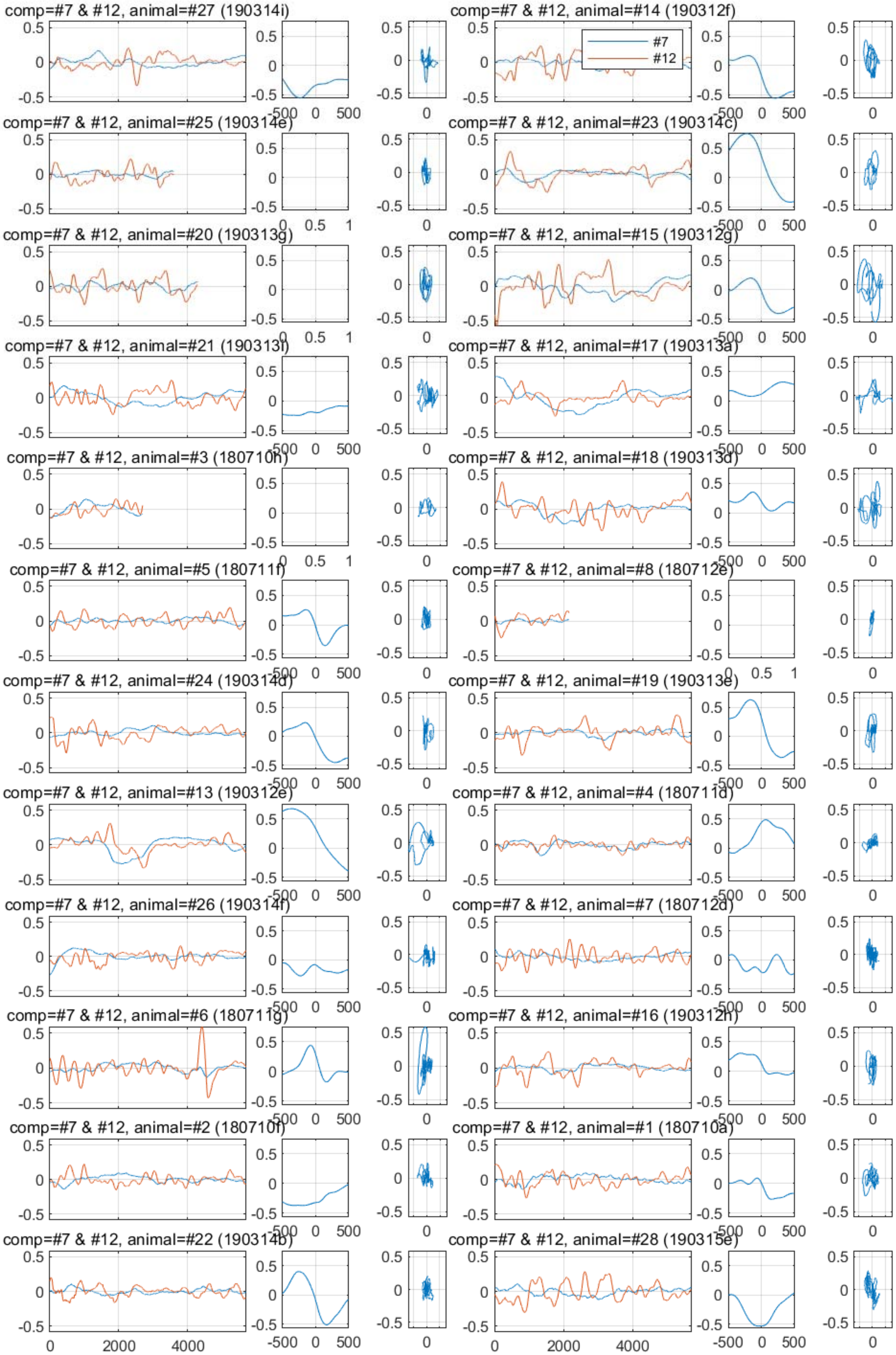
comp=#7 & #12, animal=#27 (190314i)
comp=#7 & #12, animal=#14 (190312f)
#7
#12
comp=#7 & #12, animal=#25 (190314e)
comp=#7 & #12, animal=#23 (190314c)
comp=#7 & #12, animal=#20 (190313g)
comp=#7 & #12, animal=#15 (190312g)
comp=#7 & #12, animal=#21 (190313f)
comp=#7 & #12, animal=#17 (190313a)
comp=#7 & #12, animal=#3 (180710h)
comp=#7 & #12, animal=#18 (190313d)
comp=#7 & #12, animal=#5 (180711f)
comp=#7 & #12, animal=#8 (180712e)
comp=#7 & #12, animal=#24 (190314d)
comp=#7 & #12, animal=#19 (190313e)
comp=#7 & #12, animal=#13 (190312e)
comp=#7 & #12, animal=#4 (180711d)
comp=#7 & #12, animal=#26 (190314f)
comp=#7 & #12, animal=#7 (180712d)
comp=#7 & #12, animal=#6 (180711g)
comp=#7 & #12, animal=#16 (190312h)
comp=#7 & #12, animal=#2 (180710f)
comp=#7 & #12, animal=#1 (180710a)
comp=#7 & #12, animal=#22 (190314b)
comp=#7 & #12, animal=#28 (190315e)

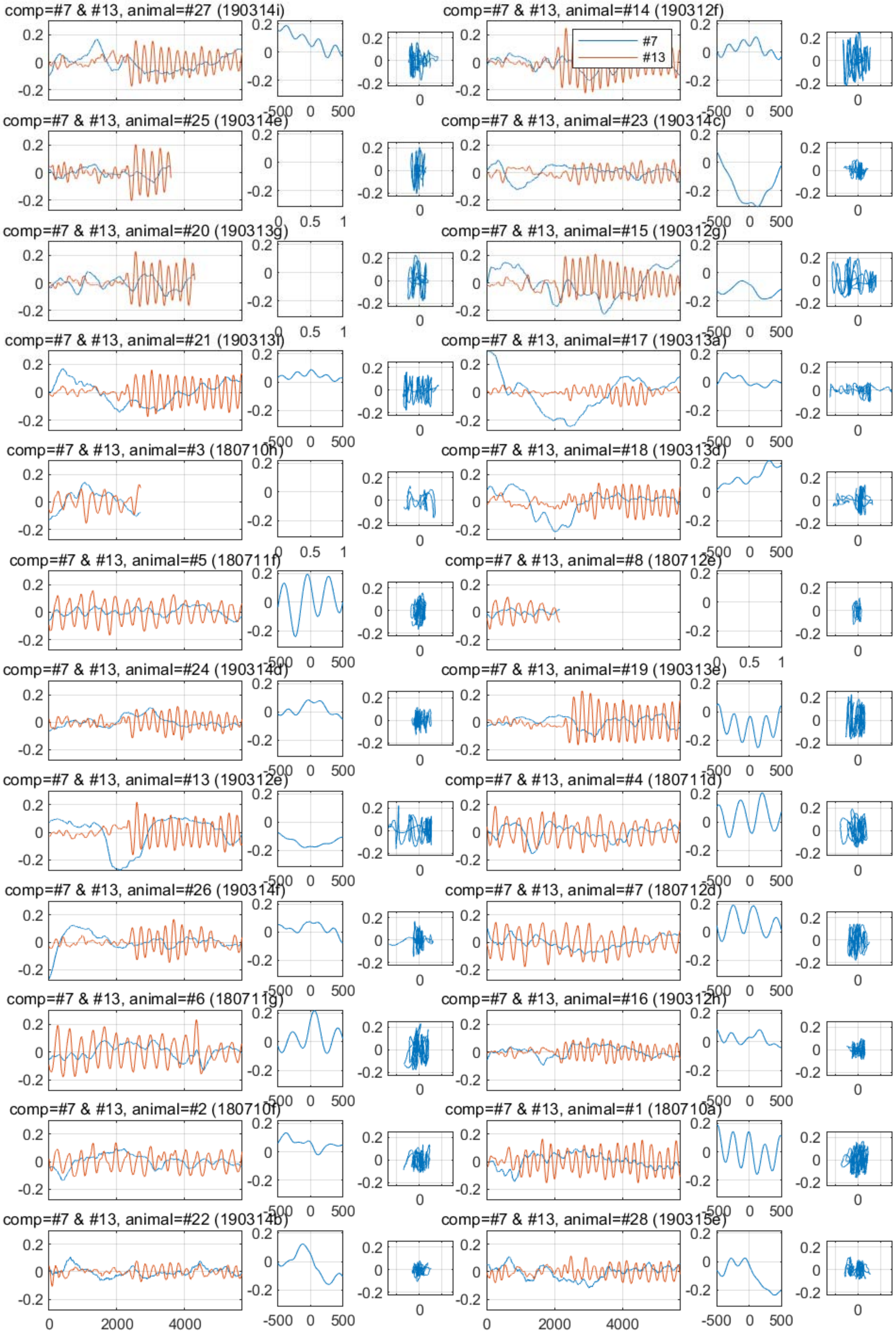

comp=#7 & #13, animal=#27 (190314i)
comp=#7 & #13, animal=#14 (190312f)
#7
#13
comp=#7 & #13, animal=#25 (190314e)
comp=#7 & #13, animal=#23 (190314c)
comp=#7 & #13, animal=#20 (190313g)
comp=#7 & #13, animal=#15 (190312g)
comp=#7 & #13, animal=#21 (190313i)
comp=#7 & #13, animal=#17 (190313a)
comp=#7 & #13, animal=#3 (180710h)
comp=#7 & #13, animal=#18 (190313d)
comp=#7 & #13, animal=#5 (180711f)
comp=#7 & #13, animal=#8 (180712e)
comp=#7 & #13, animal=#24 (190314d)
comp=#7 & #13, animal=#19 (190313e)
comp=#7 & #13, animal=#13 (190312e)
comp=#7 & #13, animal=#4 (180711d)
comp=#7 & #13, animal=#26 (190314f)
comp=#7 & #13, animal=#7 (180712d)
comp=#7 & #13, animal=#6 (180711g)
comp=#7 & #13, animal=#16 (190312h)
comp=#7 & #13, animal=#2 (180710f)
comp=#7 & #13, animal=#1 (180710a)
comp=#7 & #13, animal=#22 (190314b)
comp=#7 & #13, animal=#28 (190315e)
0 2000 4000
-500 0 500
0 0.5 1
0.2 0 -0.2

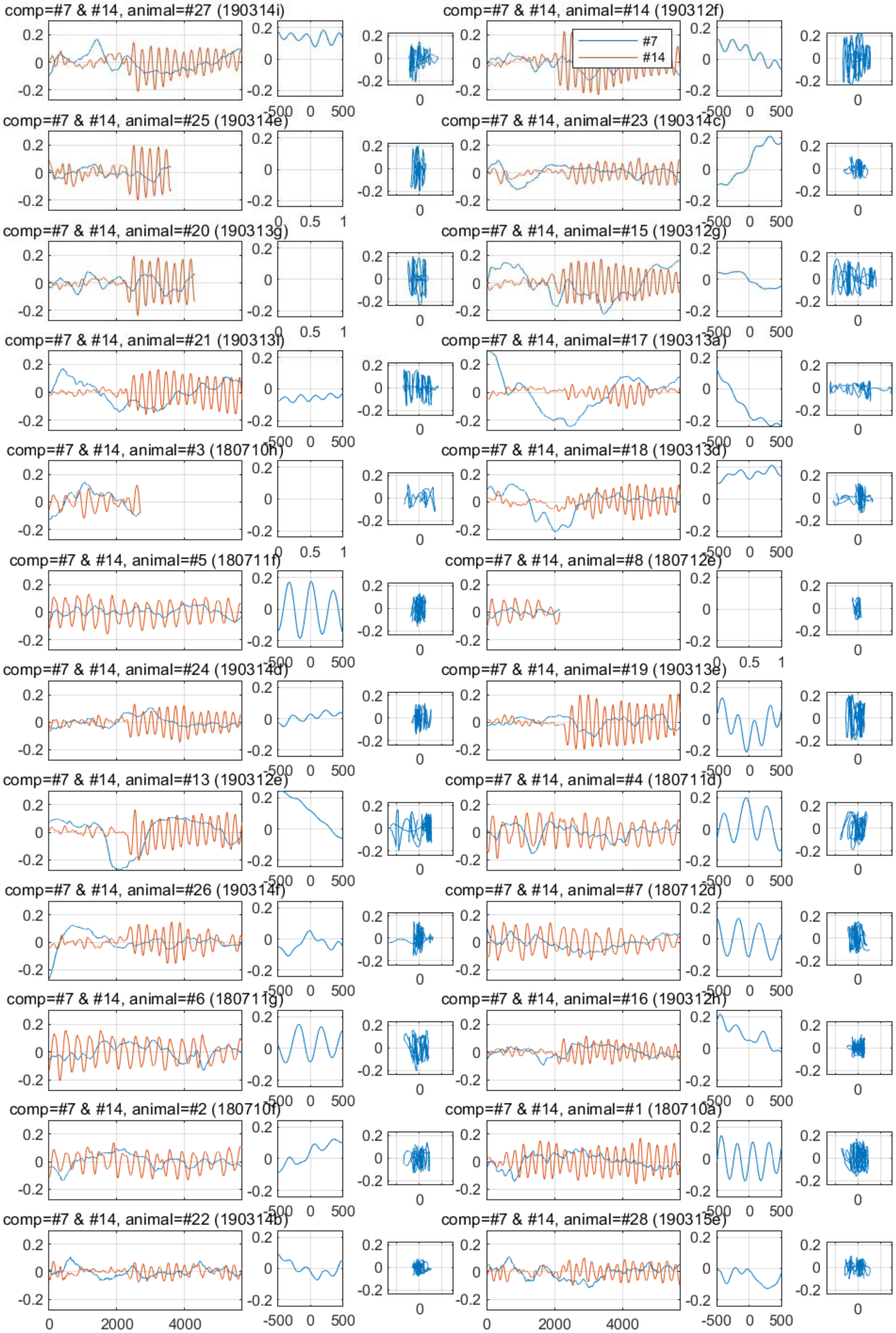
comp=#7 & #14, animal=#27 (190314i)
comp=#7 & #14, animal=#14 (190312f)
#7
#14
comp=#7 & #14, animal=#25 (190314e)
comp=#7 & #14, animal=#23 (190314c)
comp=#7 & #14, animal=#20 (190313g)
comp=#7 & #14, animal=#15 (190312g)
comp=#7 & #14, animal=#21 (190313f)
comp=#7 & #14, animal=#17 (190313a)
comp=#7 & #14, animal=#3 (180710h)
comp=#7 & #14, animal=#18 (190313d)
comp=#7 & #14, animal=#5 (180711f)
comp=#7 & #14, animal=#8 (180712e)
comp=#7 & #14, animal=#24 (190314d)
comp=#7 & #14, animal=#19 (190313e)
comp=#7 & #14, animal=#13 (190312e)
comp=#7 & #14, animal=#4 (180711d)
comp=#7 & #14, animal=#26 (190314f)
comp=#7 & #14, animal=#7 (180712d)
comp=#7 & #14, animal=#6 (180711g)
comp=#7 & #14, animal=#16 (190312h)
comp=#7 & #14, animal=#2 (180710f)
comp=#7 & #14, animal=#1 (180710a)
comp=#7 & #14, animal=#22 (190314b)
comp=#7 & #14, animal=#28 (190315e)

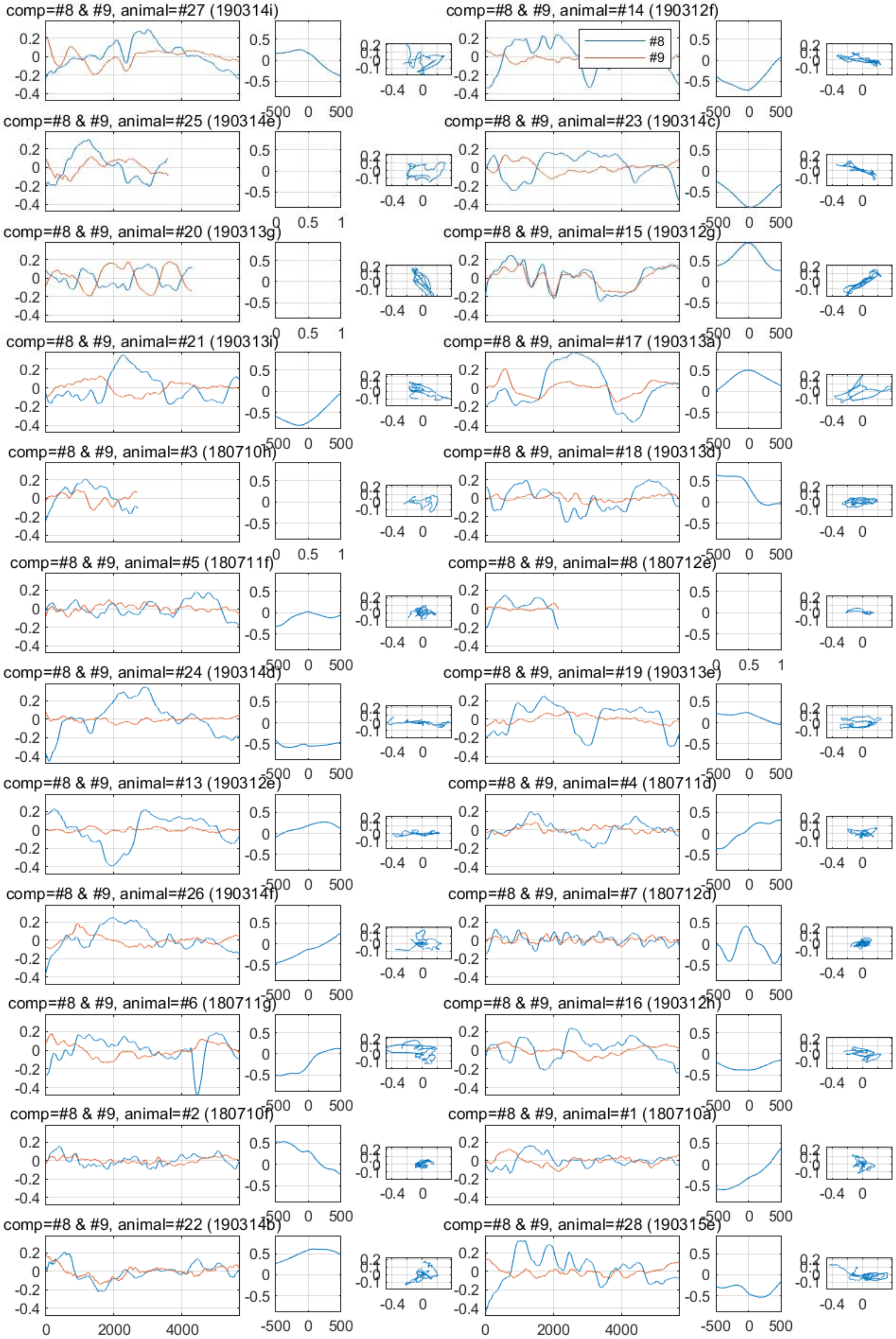
comp=#8 & #9, animal=#27 (190314i)
comp=#8 & #9, animal=#14 (190312f)
#8
#9
comp=#8 & #9, animal=#25 (190314e)
comp=#8 & #9, animal=#23 (190314c)
comp=#8 & #9, animal=#20 (190313g)
comp=#8 & #9, animal=#15 (190312g)
comp=#8 & #9, animal=#21 (190313i)
comp=#8 & #9, animal=#17 (190313a)
comp=#8 & #9, animal=#3 (180710h)
comp=#8 & #9, animal=#18 (190313d)
comp=#8 & #9, animal=#5 (180711f)
comp=#8 & #9, animal=#8 (180712e)
comp=#8 & #9, animal=#24 (190314d)
comp=#8 & #9, animal=#19 (190313e)
comp=#8 & #9, animal=#13 (190312e)
comp=#8 & #9, animal=#4 (180711d)
comp=#8 & #9, animal=#26 (190314f)
comp=#8 & #9, animal=#7 (180712d)
comp=#8 & #9, animal=#6 (180711g)
comp=#8 & #9, animal=#16 (190312h)
comp=#8 & #9, animal=#2 (180710f)
comp=#8 & #9, animal=#1 (180710a)
comp=#8 & #9, animal=#22 (190314b)
comp=#8 & #9, animal=#28 (190315e)

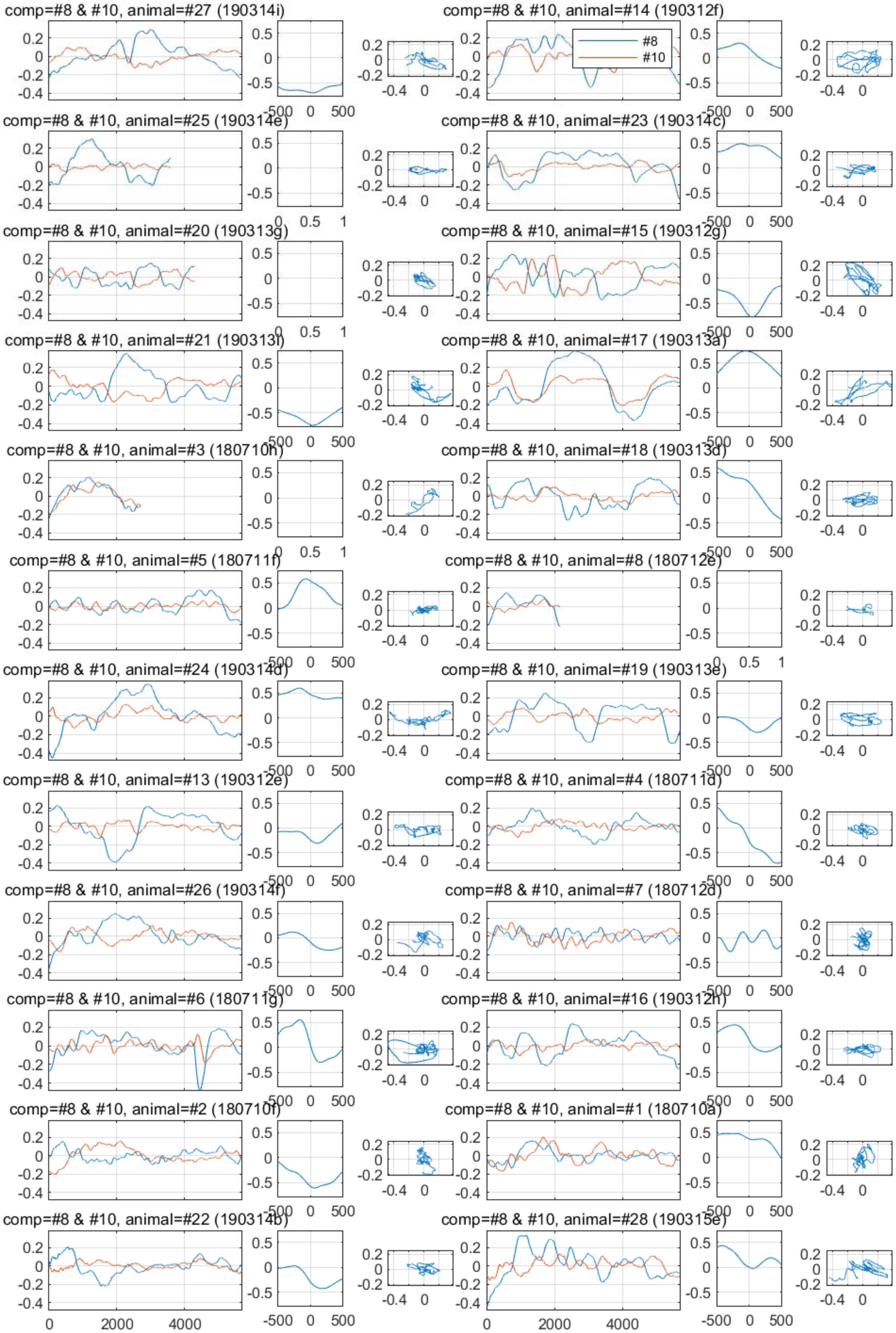

comp=#8 & #10, animal=#27 (190314i)
comp=#8 & #10, animal=#14 (190312f)
#8
#10
comp=#8 & #10, animal=#25 (190314e)
comp=#8 & #10, animal=#23 (190314c)
comp=#8 & #10, animal=#20 (190313g)
comp=#8 & #10, animal=#15 (190312g)
comp=#8 & #10, animal=#21 (190313i)
comp=#8 & #10, animal=#17 (190313a)
comp=#8 & #10, animal=#3 (180710h)
comp=#8 & #10, animal=#18 (190313d)
comp=#8 & #10, animal=#5 (180711f)
comp=#8 & #10, animal=#8 (180712e)
comp=#8 & #10, animal=#24 (190314d)
comp=#8 & #10, animal=#19 (190313e)
comp=#8 & #10, animal=#13 (190312e)
comp=#8 & #10, animal=#4 (180711d)
comp=#8 & #10, animal=#26 (190314f)
comp=#8 & #10, animal=#7 (180712d)
comp=#8 & #10, animal=#6 (180711g)
comp=#8 & #10, animal=#16 (190312h)
comp=#8 & #10, animal=#2 (180710f)
comp=#8 & #10, animal=#1 (180710a)
comp=#8 & #10, animal=#22 (190314b)
comp=#8 & #10, animal=#28 (190315e)
0
2000
4000
-500
0
500
-0.4
0

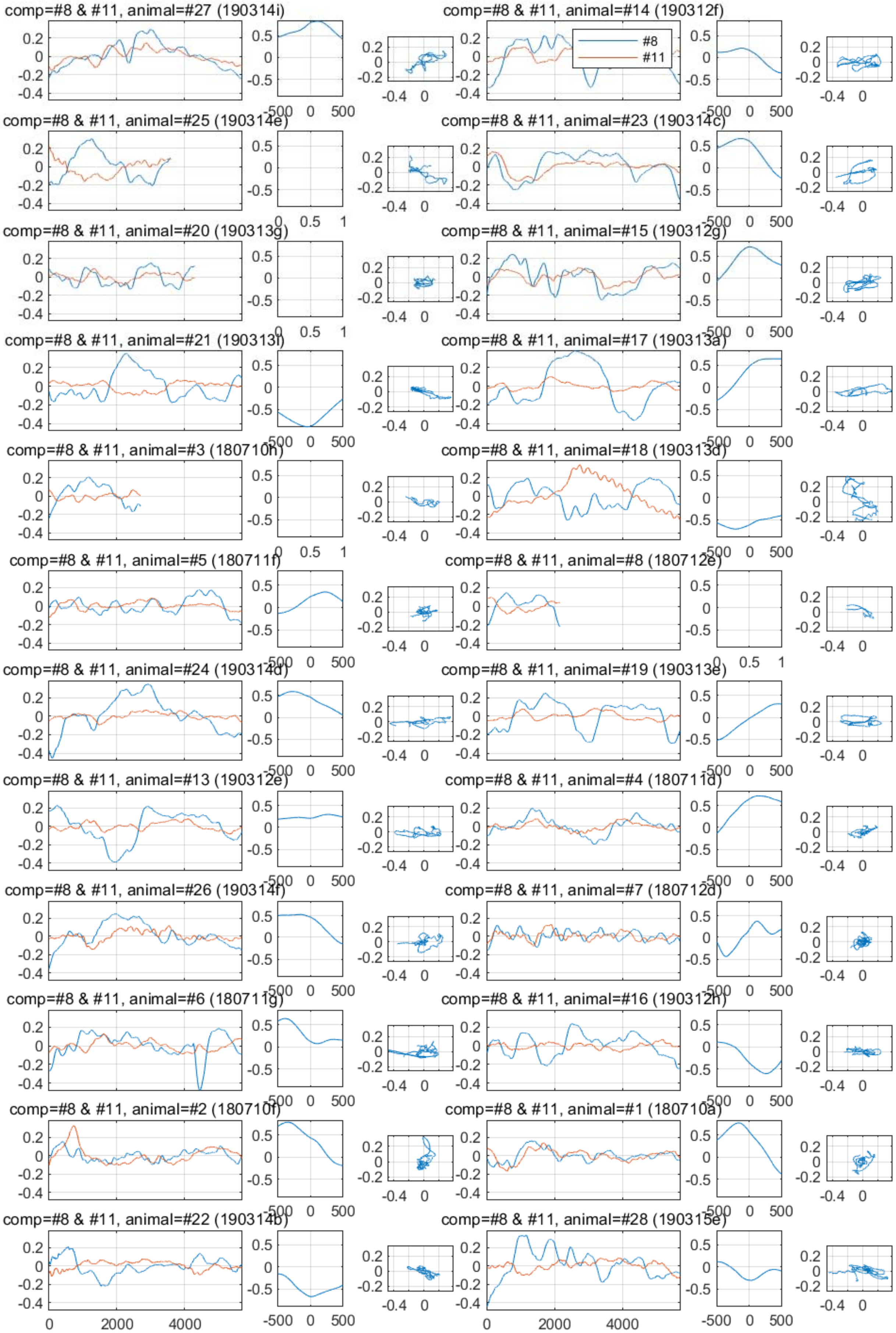
comp=#8 & #11, animal=#27 (190314i)
comp=#8 & #11, animal=#14 (190312f)
#8
#11
comp=#8 & #11, animal=#25 (190314e)
comp=#8 & #11, animal=#23 (190314c)
comp=#8 & #11, animal=#20 (190313g)
comp=#8 & #11, animal=#15 (190312g)
comp=#8 & #11, animal=#21 (190313i)
comp=#8 & #11, animal=#17 (190313a)
comp=#8 & #11, animal=#3 (180710h)
comp=#8 & #11, animal=#18 (190313d)
comp=#8 & #11, animal=#5 (180711f)
comp=#8 & #11, animal=#8 (180712e)
comp=#8 & #11, animal=#24 (190314d)
comp=#8 & #11, animal=#19 (190313e)
comp=#8 & #11, animal=#13 (190312e)
comp=#8 & #11, animal=#4 (180711d)
comp=#8 & #11, animal=#26 (190314f)
comp=#8 & #11, animal=#7 (180712d)
comp=#8 & #11, animal=#6 (180711g)
comp=#8 & #11, animal=#16 (190312h)
comp=#8 & #11, animal=#2 (180710f)
comp=#8 & #11, animal=#1 (180710a)
comp=#8 & #11, animal=#22 (190314b)
comp=#8 & #11, animal=#28 (190315e)

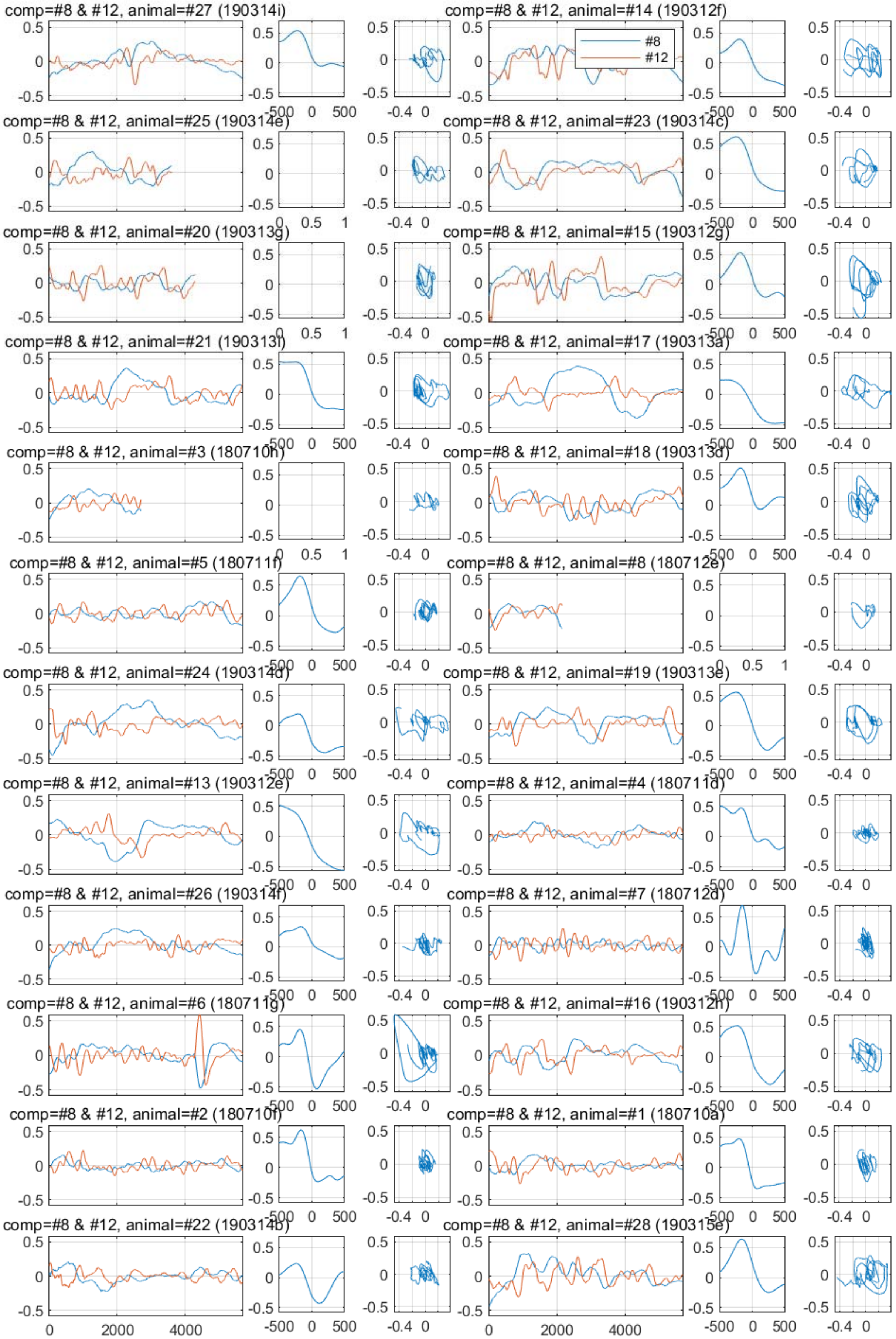
comp=#8 & #12, animal=#27 (190314i)
comp=#8 & #12, animal=#14 (190312f)
#8
#12
comp=#8 & #12, animal=#25 (190314e)
comp=#8 & #12, animal=#23 (190314c)
comp=#8 & #12, animal=#20 (190313g)
comp=#8 & #12, animal=#15 (190312g)
comp=#8 & #12, animal=#21 (190313i)
comp=#8 & #12, animal=#17 (190313a)
comp=#8 & #12, animal=#3 (180710h)
comp=#8 & #12, animal=#18 (190313d)
comp=#8 & #12, animal=#5 (180711f)
comp=#8 & #12, animal=#8 (180712e)
comp=#8 & #12, animal=#24 (190314d)
comp=#8 & #12, animal=#19 (190313e)
comp=#8 & #12, animal=#13 (190312e)
comp=#8 & #12, animal=#4 (180711d)
comp=#8 & #12, animal=#26 (190314f)
comp=#8 & #12, animal=#7 (180712d)
comp=#8 & #12, animal=#6 (180711g)
comp=#8 & #12, animal=#16 (190312h)
comp=#8 & #12, animal=#2 (180710f)
comp=#8 & #12, animal=#1 (180710a)
comp=#8 & #12, animal=#22 (190314b)
comp=#8 & #12, animal=#28 (190315e)

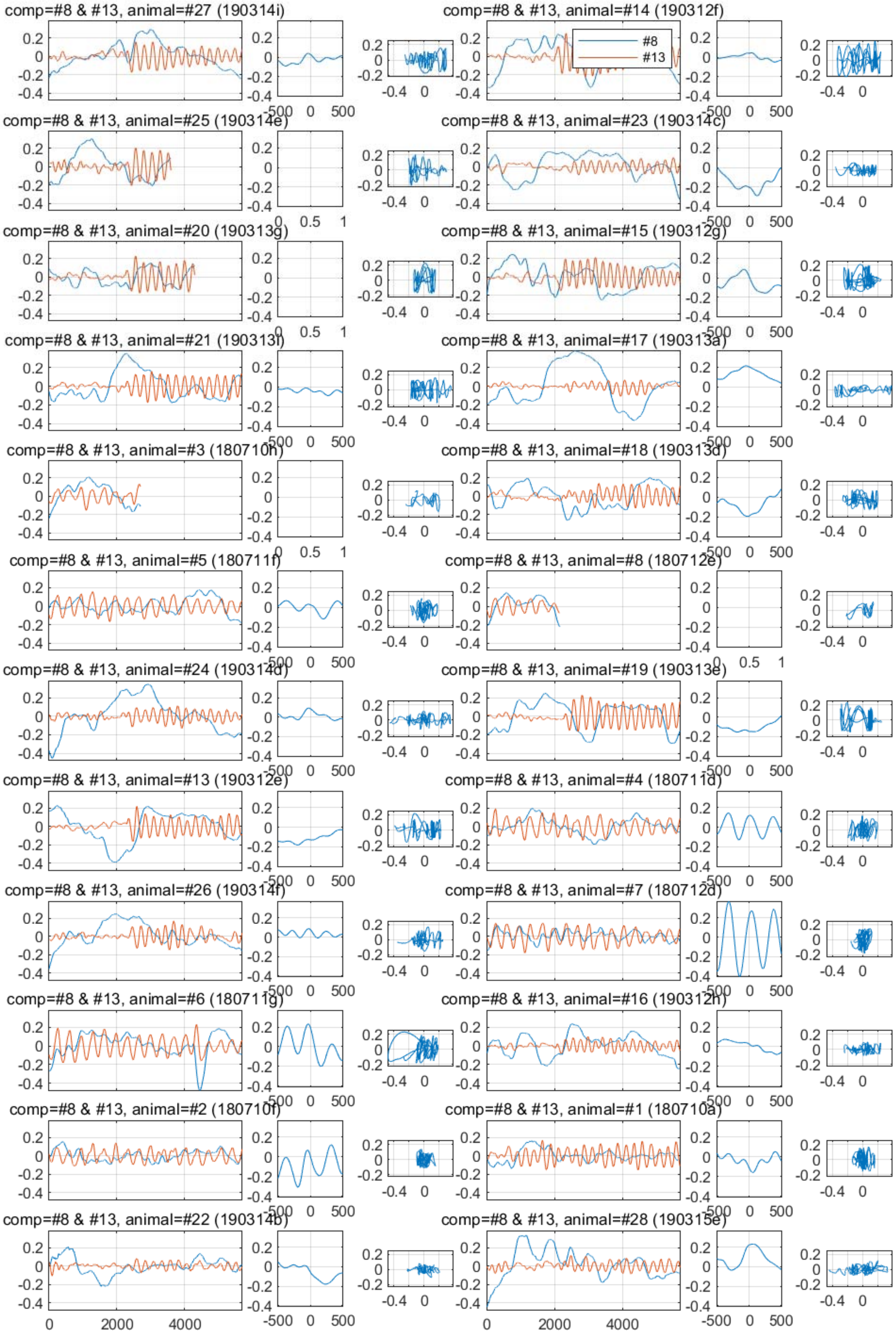

comp=#8 & #13, animal=#27 (190314i)
comp=#8 & #13, animal=#14 (190312f)
#8
#13
comp=#8 & #13, animal=#25 (190314e)
comp=#8 & #13, animal=#23 (190314c)
comp=#8 & #13, animal=#20 (190313g)
comp=#8 & #13, animal=#15 (190312g)
comp=#8 & #13, animal=#21 (190313i)
comp=#8 & #13, animal=#17 (190313a)
comp=#8 & #13, animal=#3 (180710h)
comp=#8 & #13, animal=#18 (190313d)
comp=#8 & #13, animal=#5 (180711f)
comp=#8 & #13, animal=#8 (180712e)
comp=#8 & #13, animal=#24 (190314d)
comp=#8 & #13, animal=#19 (190313e)
comp=#8 & #13, animal=#13 (190312e)
comp=#8 & #13, animal=#4 (180711d)
comp=#8 & #13, animal=#26 (190314f)
comp=#8 & #13, animal=#7 (180712d)
comp=#8 & #13, animal=#6 (180711g)
comp=#8 & #13, animal=#16 (190312h)
comp=#8 & #13, animal=#2 (180710f)
comp=#8 & #13, animal=#1 (180710a)
comp=#8 & #13, animal=#22 (190314b)
comp=#8 & #13, animal=#28 (190315e)

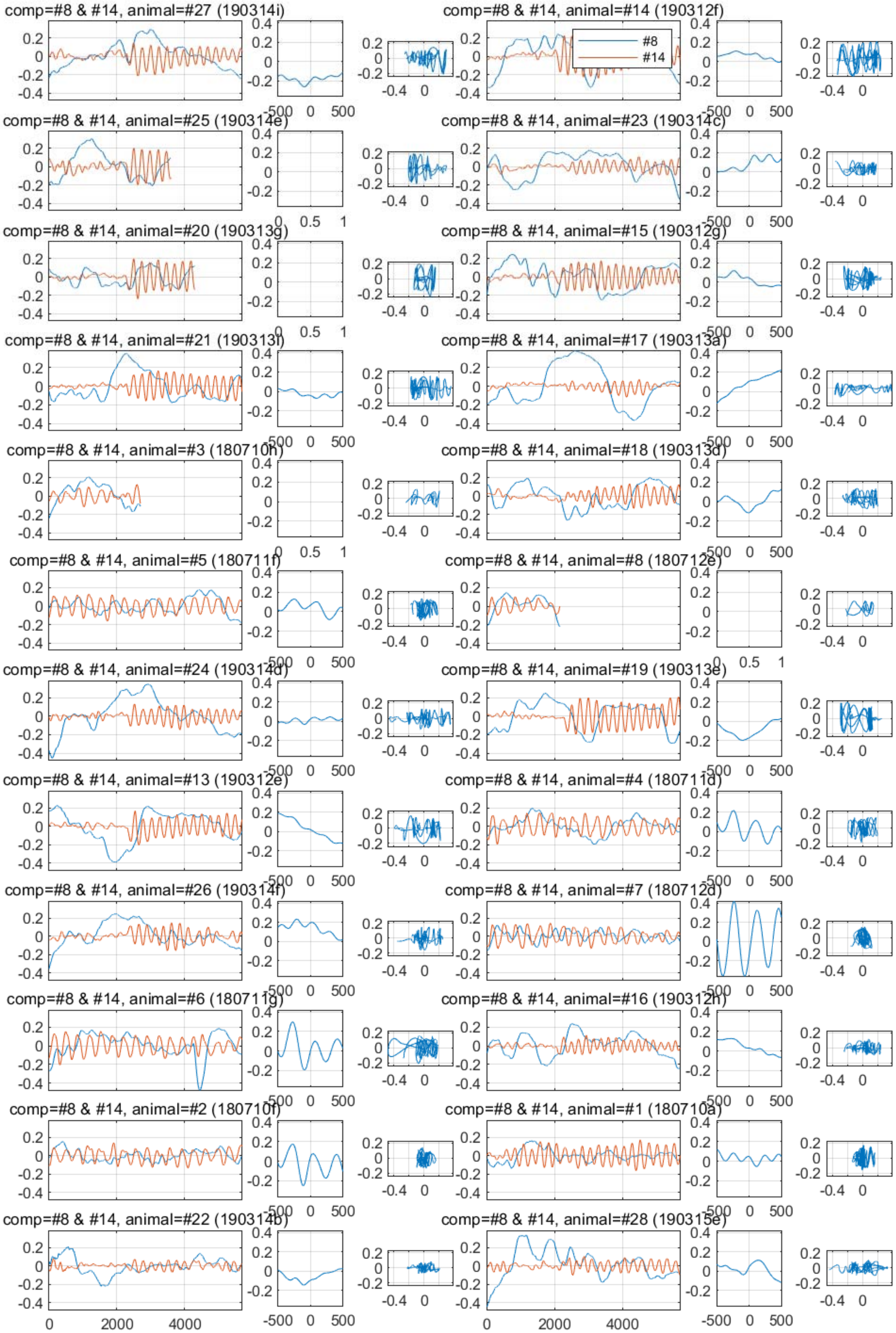

comp=#8 & #14, animal=#27 (190314i)
comp=#8 & #14, animal=#14 (190312f)
#8
#14
comp=#8 & #14, animal=#25 (190314e)
comp=#8 & #14, animal=#23 (190314c)
comp=#8 & #14, animal=#20 (190313g)
comp=#8 & #14, animal=#15 (190312g)
comp=#8 & #14, animal=#21 (190313f)
comp=#8 & #14, animal=#17 (190313a)
comp=#8 & #14, animal=#3 (180710h)
comp=#8 & #14, animal=#18 (190313d)
comp=#8 & #14, animal=#5 (180711f)
comp=#8 & #14, animal=#8 (180712e)
comp=#8 & #14, animal=#24 (190314d)
comp=#8 & #14, animal=#19 (190313e)
comp=#8 & #14, animal=#13 (190312e)
comp=#8 & #14, animal=#4 (180711d)
comp=#8 & #14, animal=#26 (190314f)
comp=#8 & #14, animal=#7 (180712d)
comp=#8 & #14, animal=#6 (180711g)
comp=#8 & #14, animal=#16 (190312h)
comp=#8 & #14, animal=#2 (180710f)
comp=#8 & #14, animal=#1 (180710a)
comp=#8 & #14, animal=#22 (190314b)
comp=#8 & #14, animal=#28 (190315e)

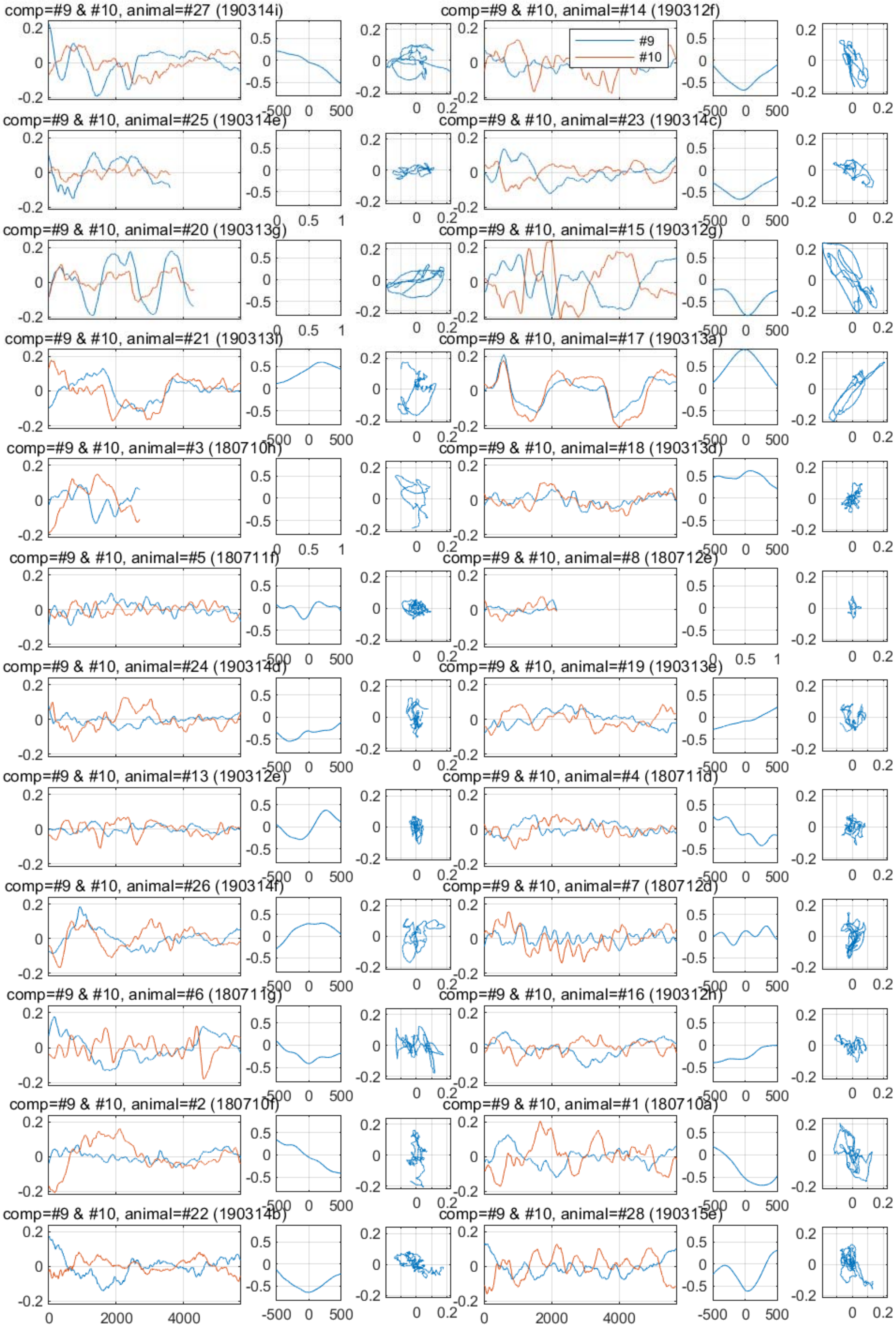
comp=#9 & #10, animal=#27 (190314i)
comp=#9 & #10, animal=#14 (190312f)
#9
#10
comp=#9 & #10, animal=#25 (190314e)
comp=#9 & #10, animal=#23 (190314c)
comp=#9 & #10, animal=#20 (190313g)
comp=#9 & #10, animal=#15 (190312g)
comp=#9 & #10, animal=#21 (190313i)
comp=#9 & #10, animal=#17 (190313a)
comp=#9 & #10, animal=#3 (180710h)
comp=#9 & #10, animal=#18 (190313d)
comp=#9 & #10, animal=#5 (180711f)
comp=#9 & #10, animal=#8 (180712e)
comp=#9 & #10, animal=#24 (190314d)
comp=#9 & #10, animal=#19 (190313e)
comp=#9 & #10, animal=#13 (190312e)
comp=#9 & #10, animal=#4 (180711d)
comp=#9 & #10, animal=#26 (190314f)
comp=#9 & #10, animal=#7 (180712d)
comp=#9 & #10, animal=#6 (180711g)
comp=#9 & #10, animal=#16 (190312h)
comp=#9 & #10, animal=#2 (180710f)
comp=#9 & #10, animal=#1 (180710a)
comp=#9 & #10, animal=#22 (190314b)
comp=#9 & #10, animal=#28 (190315e)
0
2000
4000
-500
0
500
0.2
-0.2

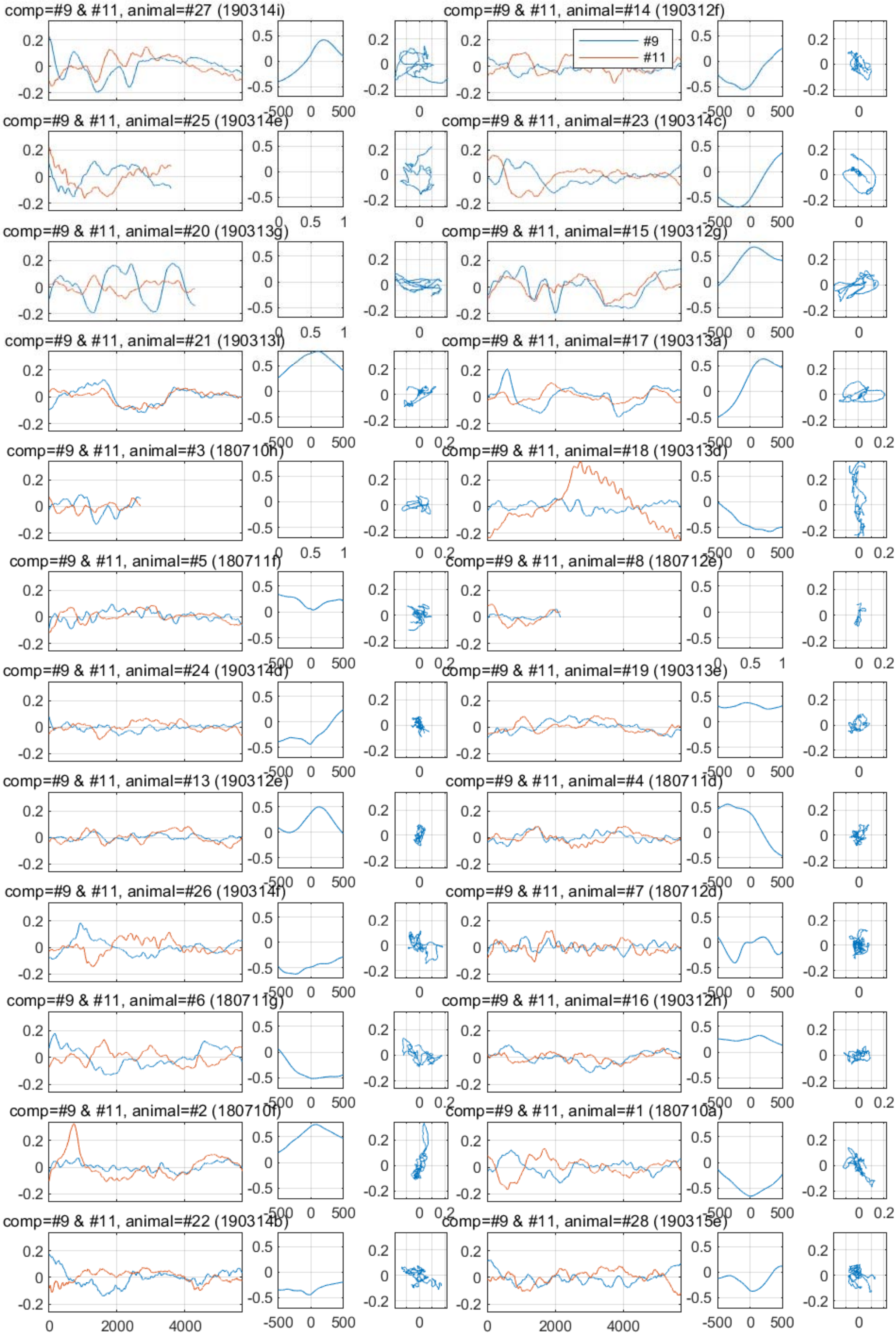

comp=#9 & #11, animal=#27 (190314i)
comp=#9 & #11, animal=#14 (190312f)
#9
#11
comp=#9 & #11, animal=#25 (190314e)
comp=#9 & #11, animal=#23 (190314c)
comp=#9 & #11, animal=#20 (190313g)
comp=#9 & #11, animal=#15 (190312g)
comp=#9 & #11, animal=#21 (190313i)
comp=#9 & #11, animal=#17 (190313a)
comp=#9 & #11, animal=#3 (180710h)
comp=#9 & #11, animal=#18 (190313d)
comp=#9 & #11, animal=#5 (180711f)
comp=#9 & #11, animal=#8 (180712e)
comp=#9 & #11, animal=#24 (190314d)
comp=#9 & #11, animal=#19 (190313e)
comp=#9 & #11, animal=#13 (190312e)
comp=#9 & #11, animal=#4 (180711d)
comp=#9 & #11, animal=#26 (190314f)
comp=#9 & #11, animal=#7 (180712d)
comp=#9 & #11, animal=#6 (180711g)
comp=#9 & #11, animal=#16 (190312h)
comp=#9 & #11, animal=#2 (180710f)
comp=#9 & #11, animal=#1 (180710a)
comp=#9 & #11, animal=#22 (190314b)
comp=#9 & #11, animal=#28 (190315e)
0
2000
4000
-500
0
500

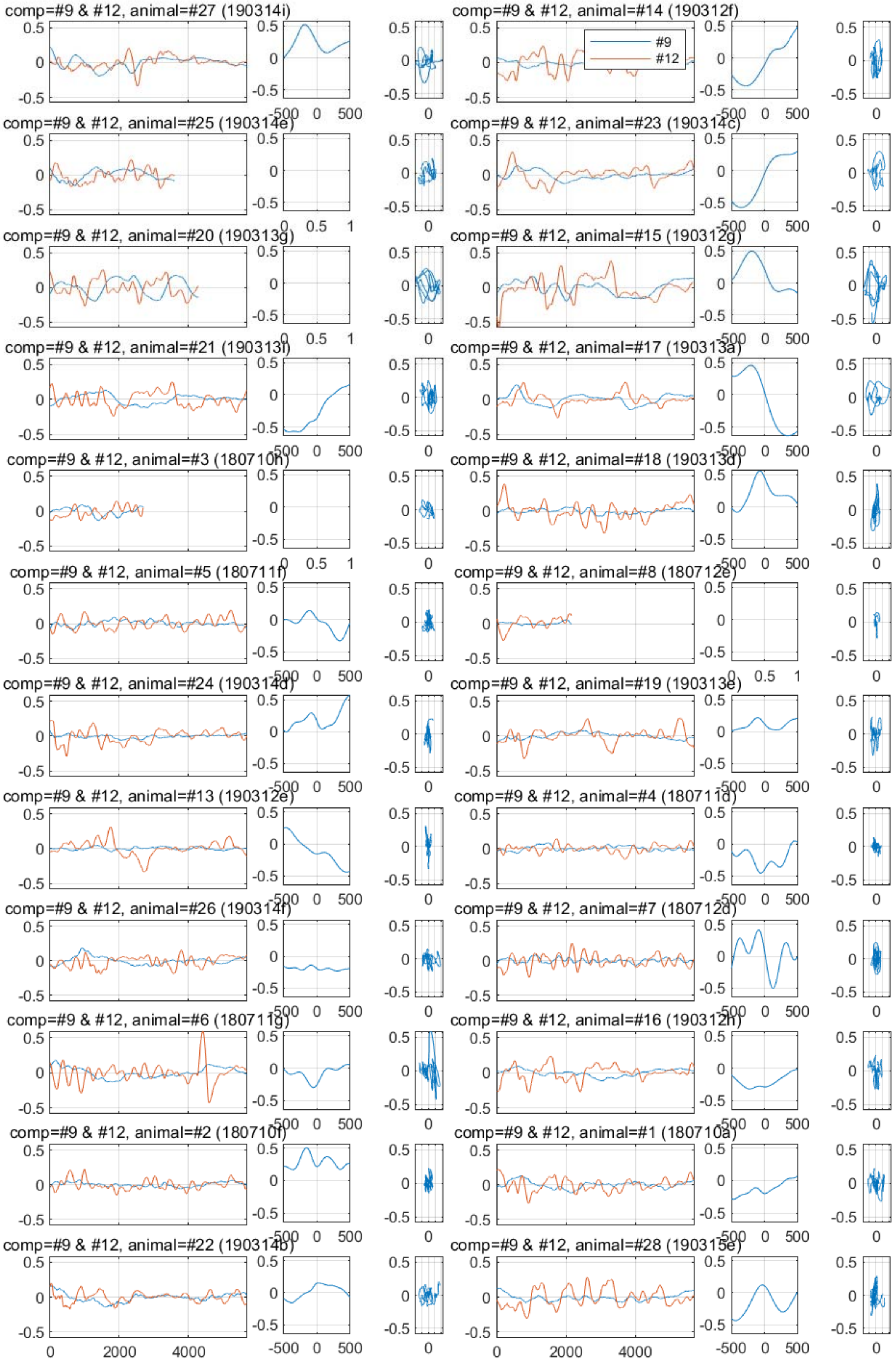
comp=#9 & #12, animal=#27 (190314i)
comp=#9 & #12, animal=#14 (190312f)
#9
#12
comp=#9 & #12, animal=#25 (190314e)
comp=#9 & #12, animal=#23 (190314c)
comp=#9 & #12, animal=#20 (190313g)
comp=#9 & #12, animal=#15 (190312g)
comp=#9 & #12, animal=#21 (190313i)
comp=#9 & #12, animal=#17 (190313a)
comp=#9 & #12, animal=#3 (180710h)
comp=#9 & #12, animal=#18 (190313d)
comp=#9 & #12, animal=#5 (180711f)
comp=#9 & #12, animal=#8 (180712e)
comp=#9 & #12, animal=#24 (190314d)
comp=#9 & #12, animal=#19 (190313e)
comp=#9 & #12, animal=#13 (190312e)
comp=#9 & #12, animal=#4 (180711d)
comp=#9 & #12, animal=#26 (190314f)
comp=#9 & #12, animal=#7 (180712d)
comp=#9 & #12, animal=#6 (180711g)
comp=#9 & #12, animal=#16 (190312h)
comp=#9 & #12, animal=#2 (180710f)
comp=#9 & #12, animal=#1 (180710a)
comp=#9 & #12, animal=#22 (190314b)
comp=#9 & #12, animal=#28 (190315e)

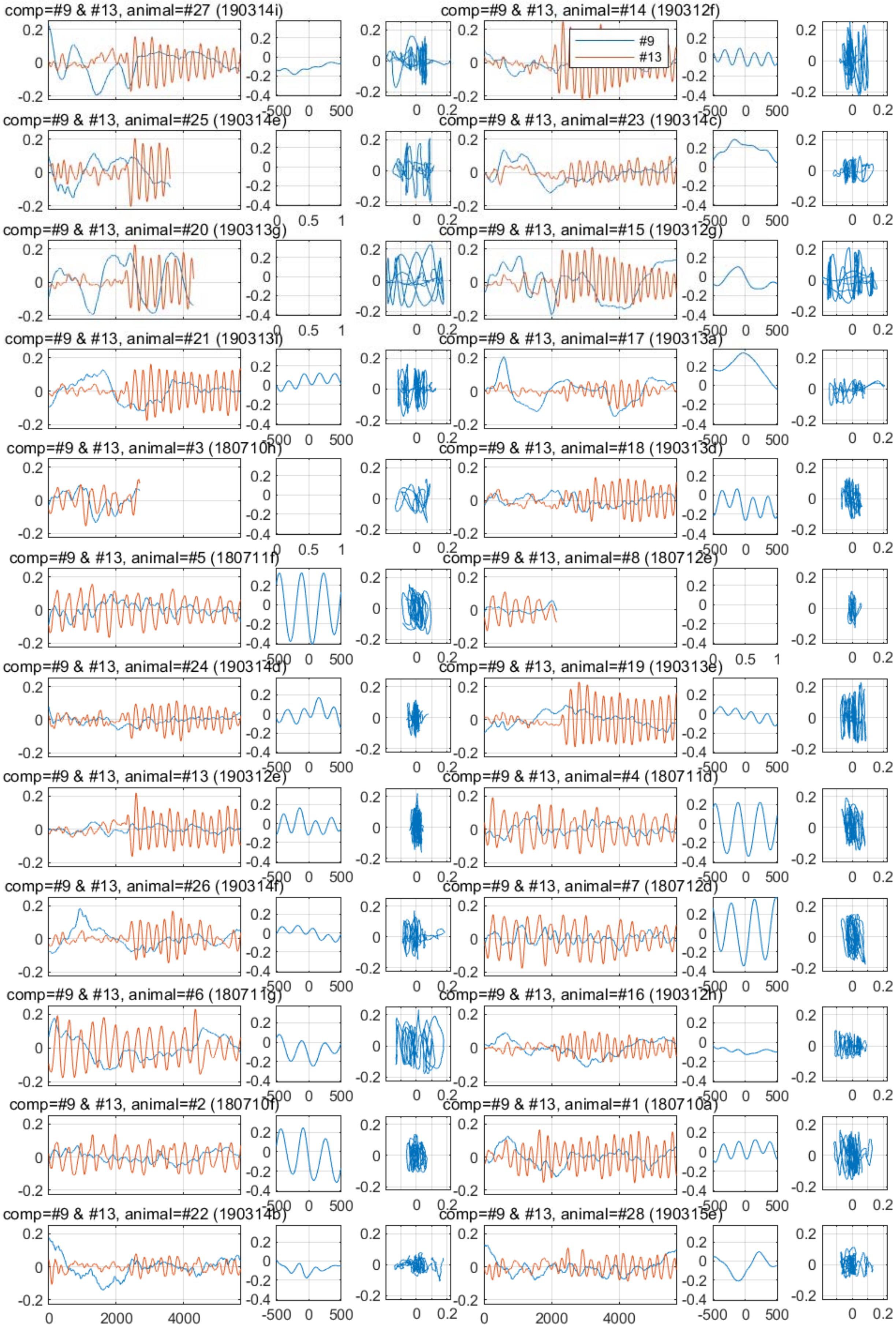
comp=#9 & #13, animal=#27 (190314i)
comp=#9 & #13, animal=#14 (190312f)
#9
#13
comp=#9 & #13, animal=#25 (190314e)
comp=#9 & #13, animal=#23 (190314c)
comp=#9 & #13, animal=#20 (190313g)
comp=#9 & #13, animal=#15 (190312g)
comp=#9 & #13, animal=#21 (190313f)
comp=#9 & #13, animal=#17 (190313a)
comp=#9 & #13, animal=#3 (180710h)
comp=#9 & #13, animal=#18 (190313d)
comp=#9 & #13, animal=#5 (180711f)
comp=#9 & #13, animal=#8 (180712e)
comp=#9 & #13, animal=#24 (190314d)
comp=#9 & #13, animal=#19 (190313e)
comp=#9 & #13, animal=#13 (190312e)
comp=#9 & #13, animal=#4 (180711d)
comp=#9 & #13, animal=#26 (190314f)
comp=#9 & #13, animal=#7 (180712d)
comp=#9 & #13, animal=#6 (180711g)
comp=#9 & #13, animal=#16 (190312h)
comp=#9 & #13, animal=#2 (180710f)
comp=#9 & #13, animal=#1 (180710a)
comp=#9 & #13, animal=#22 (190314b)
comp=#9 & #13, animal=#28 (190315e)
0
2000
4000
-500
0
500
0.2

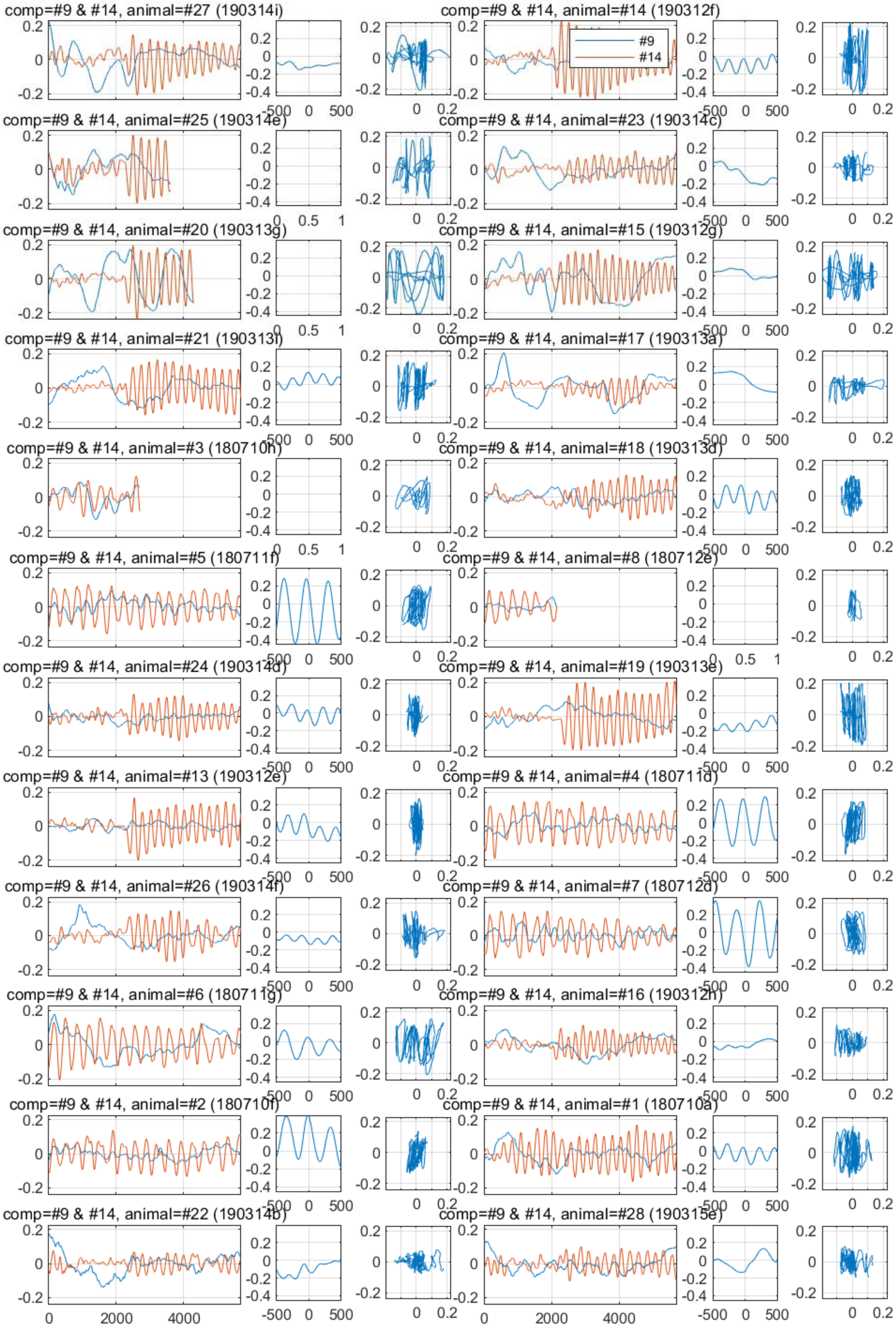
comp=#9 & #14, animal=#27 (190314i)
comp=#9 & #14, animal=#14 (190312f)
#9
#14
comp=#9 & #14, animal=#25 (190314e)
comp=#9 & #14, animal=#23 (190314c)
comp=#9 & #14, animal=#20 (190313g)
comp=#9 & #14, animal=#15 (190312g)
comp=#9 & #14, animal=#21 (190313f)
comp=#9 & #14, animal=#17 (190313a)
comp=#9 & #14, animal=#3 (180710h)
comp=#9 & #14, animal=#18 (190313d)
comp=#9 & #14, animal=#5 (180711f)
comp=#9 & #14, animal=#8 (180712e)
comp=#9 & #14, animal=#24 (190314d)
comp=#9 & #14, animal=#19 (190313e)
comp=#9 & #14, animal=#13 (190312e)
comp=#9 & #14, animal=#4 (180711d)
comp=#9 & #14, animal=#26 (190314f)
comp=#9 & #14, animal=#7 (180712d)
comp=#9 & #14, animal=#6 (180711g)
comp=#9 & #14, animal=#16 (190312h)
comp=#9 & #14, animal=#2 (180710f)
comp=#9 & #14, animal=#1 (180710a)
comp=#9 & #14, animal=#22 (190314b)
comp=#9 & #14, animal=#28 (190315e)

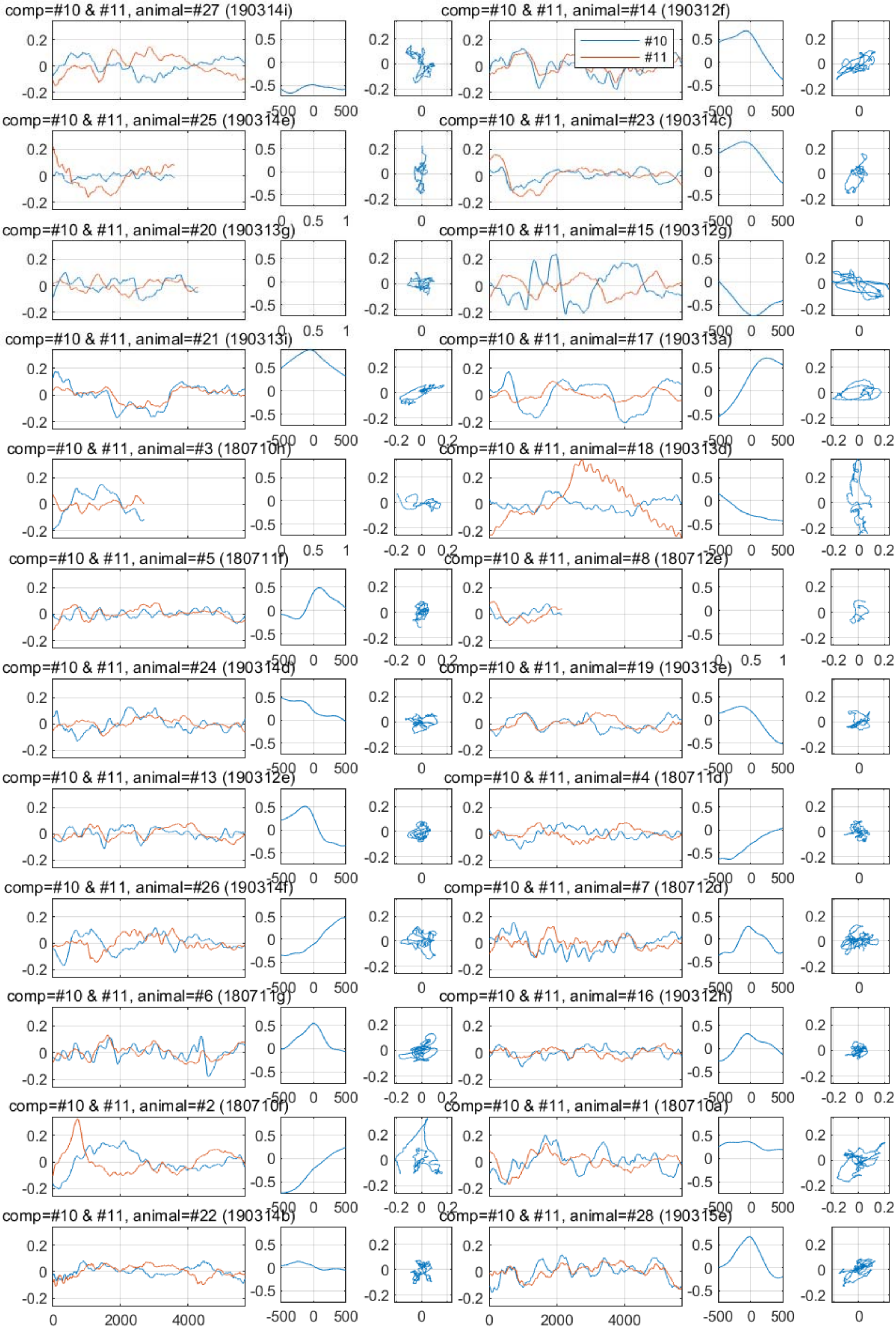
comp=#10 & #11, animal=#27 (190314i)
comp=#10 & #11, animal=#14 (190312f)
#10
#11
comp=#10 & #11, animal=#25 (190314e)
comp=#10 & #11, animal=#23 (190314c)
comp=#10 & #11, animal=#20 (190313g)
comp=#10 & #11, animal=#15 (190312g)
comp=#10 & #11, animal=#21 (190313i)
comp=#10 & #11, animal=#17 (190313a)
comp=#10 & #11, animal=#3 (180710h)
comp=#10 & #11, animal=#18 (190313d)
comp=#10 & #11, animal=#5 (180711f)
comp=#10 & #11, animal=#8 (180712e)
comp=#10 & #11, animal=#24 (190314d)
comp=#10 & #11, animal=#19 (190313e)
comp=#10 & #11, animal=#13 (190312e)
comp=#10 & #11, animal=#4 (180711d)
comp=#10 & #11, animal=#26 (190314f)
comp=#10 & #11, animal=#7 (180712d)
comp=#10 & #11, animal=#6 (180711g)
comp=#10 & #11, animal=#16 (190312h)
comp=#10 & #11, animal=#2 (180710f)
comp=#10 & #11, animal=#1 (180710a)
comp=#10 & #11, animal=#22 (190314b)
comp=#10 & #11, animal=#28 (190315e)

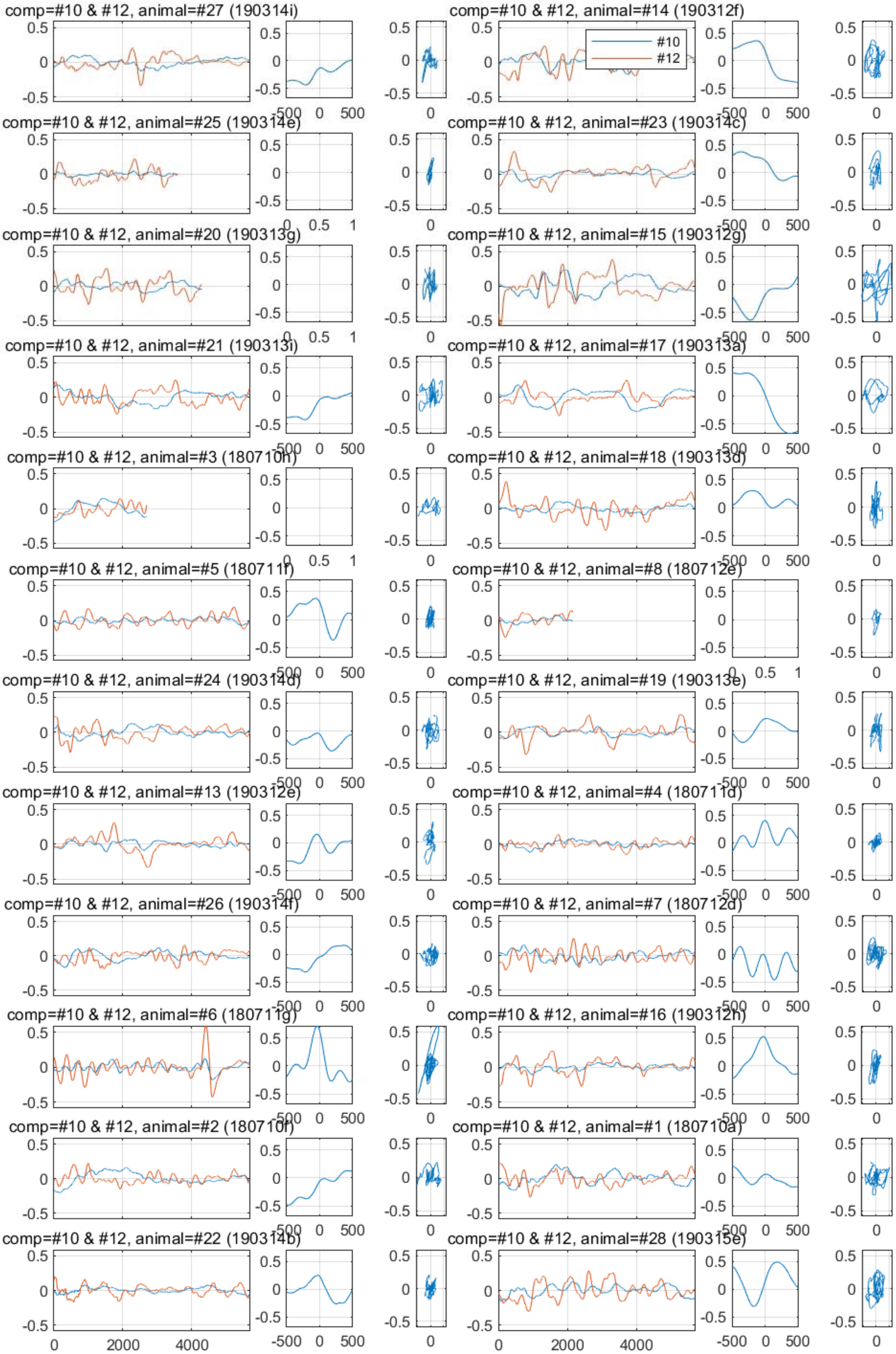
comp=#10 & #12, animal=#27 (190314i)
comp=#10 & #12, animal=#14 (190312f)
#10
#12
comp=#10 & #12, animal=#25 (190314e)
comp=#10 & #12, animal=#23 (190314c)
comp=#10 & #12, animal=#20 (190313g)
comp=#10 & #12, animal=#15 (190312g)
comp=#10 & #12, animal=#21 (190313i)
comp=#10 & #12, animal=#17 (190313a)
comp=#10 & #12, animal=#3 (180710h)
comp=#10 & #12, animal=#18 (190313d)
comp=#10 & #12, animal=#5 (180711f)
comp=#10 & #12, animal=#8 (180712e)
comp=#10 & #12, animal=#24 (190314d)
comp=#10 & #12, animal=#19 (190313e)
comp=#10 & #12, animal=#13 (190312e)
comp=#10 & #12, animal=#4 (180711d)
comp=#10 & #12, animal=#26 (190314f)
comp=#10 & #12, animal=#7 (180712d)
comp=#10 & #12, animal=#6 (180711g)
comp=#10 & #12, animal=#16 (190312h)
comp=#10 & #12, animal=#2 (180710f)
comp=#10 & #12, animal=#1 (180710a)
comp=#10 & #12, animal=#22 (190314b)
comp=#10 & #12, animal=#28 (190315e)
0
2000
4000
-500
0
500

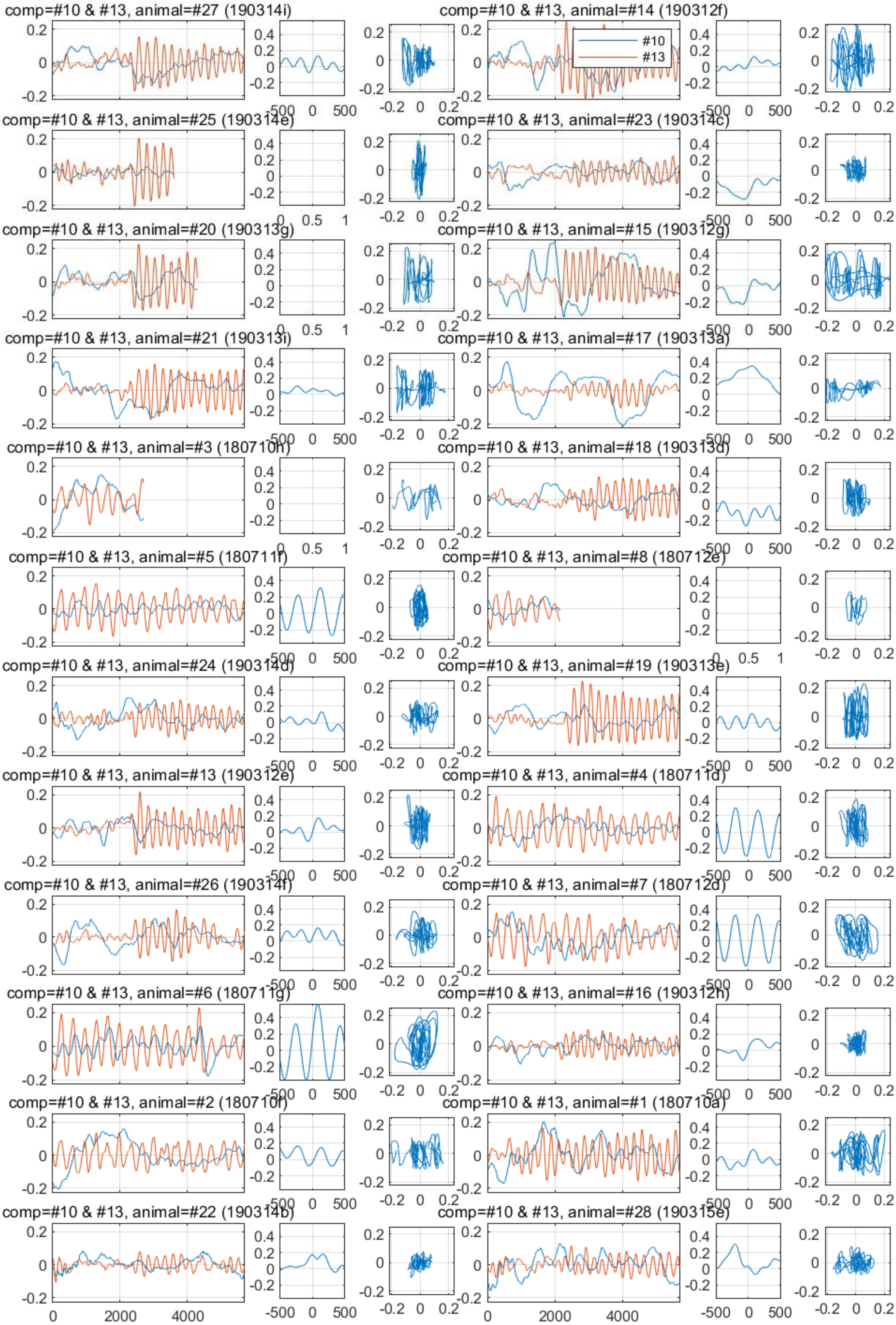
comp=#10 & #13, animal=#27 (190314i)
comp=#10 & #13, animal=#14 (190312f)
#10
#13
comp=#10 & #13, animal=#25 (190314e)
comp=#10 & #13, animal=#23 (190314c)
comp=#10 & #13, animal=#20 (190313g)
comp=#10 & #13, animal=#15 (190312g)
comp=#10 & #13, animal=#21 (190313i)
comp=#10 & #13, animal=#17 (190313a)
comp=#10 & #13, animal=#3 (180710h)
comp=#10 & #13, animal=#18 (190313d)
comp=#10 & #13, animal=#5 (180711f)
comp=#10 & #13, animal=#8 (180712e)
comp=#10 & #13, animal=#24 (190314d)
comp=#10 & #13, animal=#19 (190313e)
comp=#10 & #13, animal=#13 (190312e)
comp=#10 & #13, animal=#4 (180711d)
comp=#10 & #13, animal=#26 (190314f)
comp=#10 & #13, animal=#7 (180712d)
comp=#10 & #13, animal=#6 (180711g)
comp=#10 & #13, animal=#16 (190312h)
comp=#10 & #13, animal=#2 (180710f)
comp=#10 & #13, animal=#1 (180710a)
comp=#10 & #13, animal=#22 (190314b)
comp=#10 & #13, animal=#28 (190315e)

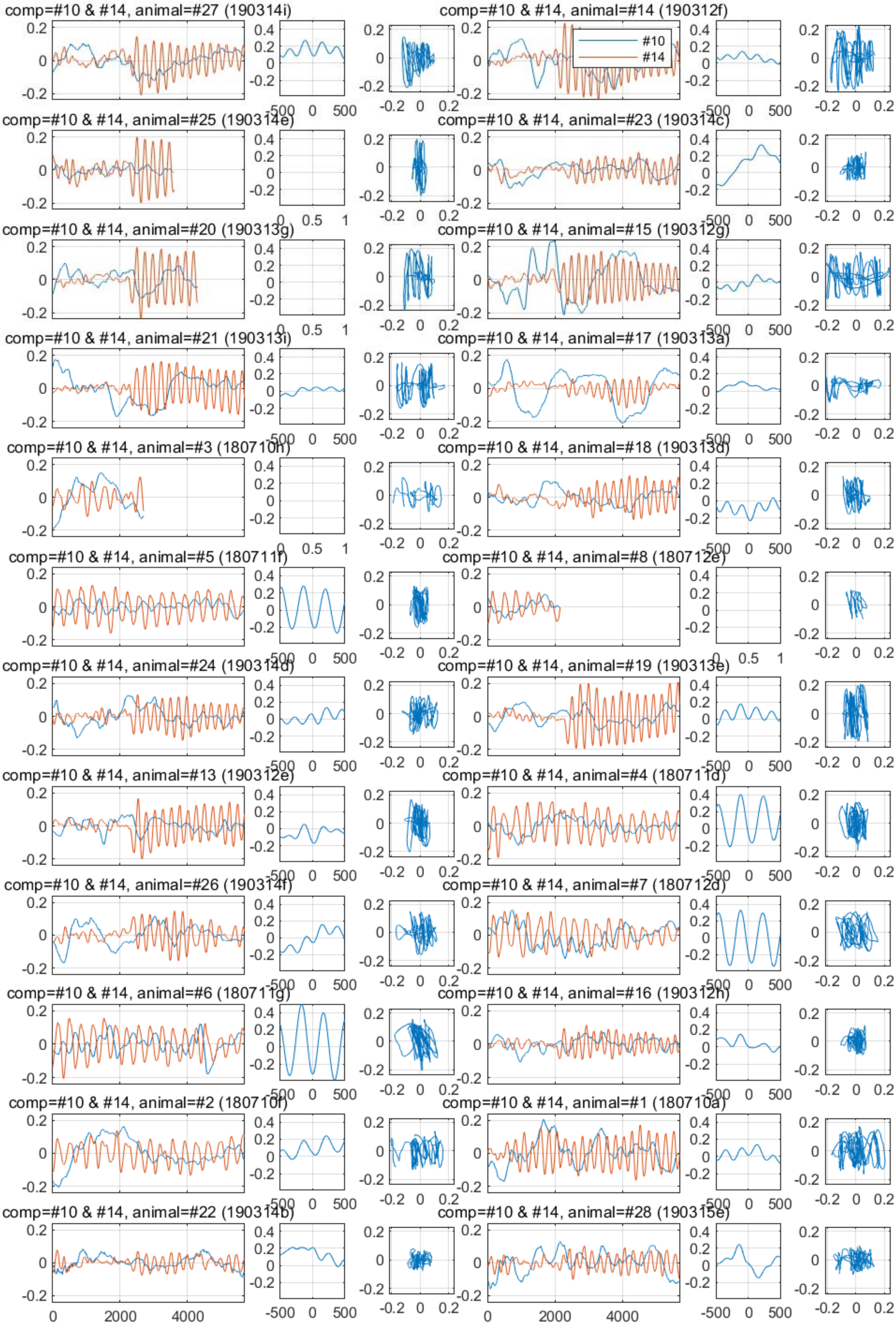
comp=#10 & #14, animal=#27 (190314i)
comp=#10 & #14, animal=#14 (190312f)
#10
#14
comp=#10 & #14, animal=#25 (190314e)
comp=#10 & #14, animal=#23 (190314c)
comp=#10 & #14, animal=#20 (190313g)
comp=#10 & #14, animal=#15 (190312g)
comp=#10 & #14, animal=#21 (190313i)
comp=#10 & #14, animal=#17 (190313a)
comp=#10 & #14, animal=#3 (180710h)
comp=#10 & #14, animal=#18 (190313d)
comp=#10 & #14, animal=#5 (180711f)
comp=#10 & #14, animal=#8 (180712e)
comp=#10 & #14, animal=#24 (190314d)
comp=#10 & #14, animal=#19 (190313e)
comp=#10 & #14, animal=#13 (190312e)
comp=#10 & #14, animal=#4 (180711d)
comp=#10 & #14, animal=#26 (190314f)
comp=#10 & #14, animal=#7 (180712d)
comp=#10 & #14, animal=#6 (180711g)
comp=#10 & #14, animal=#16 (190312h)
comp=#10 & #14, animal=#2 (180710f)
comp=#10 & #14, animal=#1 (180710a)
comp=#10 & #14, animal=#22 (190314b)
comp=#10 & #14, animal=#28 (190315e)

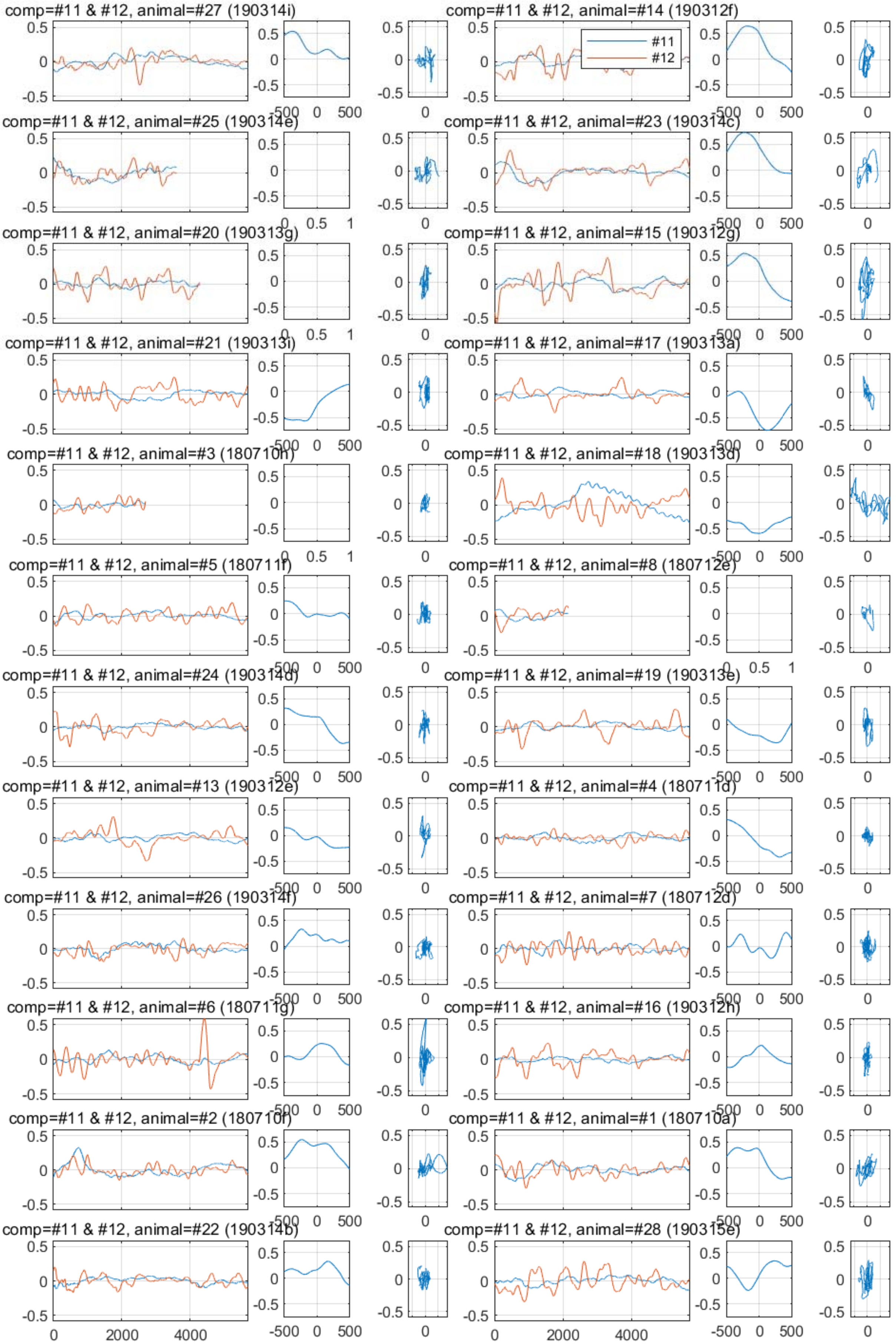

comp=#11 & #12, animal=#27 (190314i)
comp=#11 & #12, animal=#14 (190312f)
#11
#12
comp=#11 & #12, animal=#25 (190314e)
comp=#11 & #12, animal=#23 (190314c)
comp=#11 & #12, animal=#20 (190313g)
comp=#11 & #12, animal=#15 (190312g)
comp=#11 & #12, animal=#21 (190313i)
comp=#11 & #12, animal=#17 (190313a)
comp=#11 & #12, animal=#3 (180710h)
comp=#11 & #12, animal=#18 (190313d)
comp=#11 & #12, animal=#5 (180711f)
comp=#11 & #12, animal=#8 (180712e)
comp=#11 & #12, animal=#24 (190314d)
comp=#11 & #12, animal=#19 (190313e)
comp=#11 & #12, animal=#13 (190312e)
comp=#11 & #12, animal=#4 (180711d)
comp=#11 & #12, animal=#26 (190314f)
comp=#11 & #12, animal=#7 (180712d)
comp=#11 & #12, animal=#6 (180711g)
comp=#11 & #12, animal=#16 (190312h)
comp=#11 & #12, animal=#2 (180710f)
comp=#11 & #12, animal=#1 (180710a)
comp=#11 & #12, animal=#22 (190314b)
comp=#11 & #12, animal=#28 (190315e)

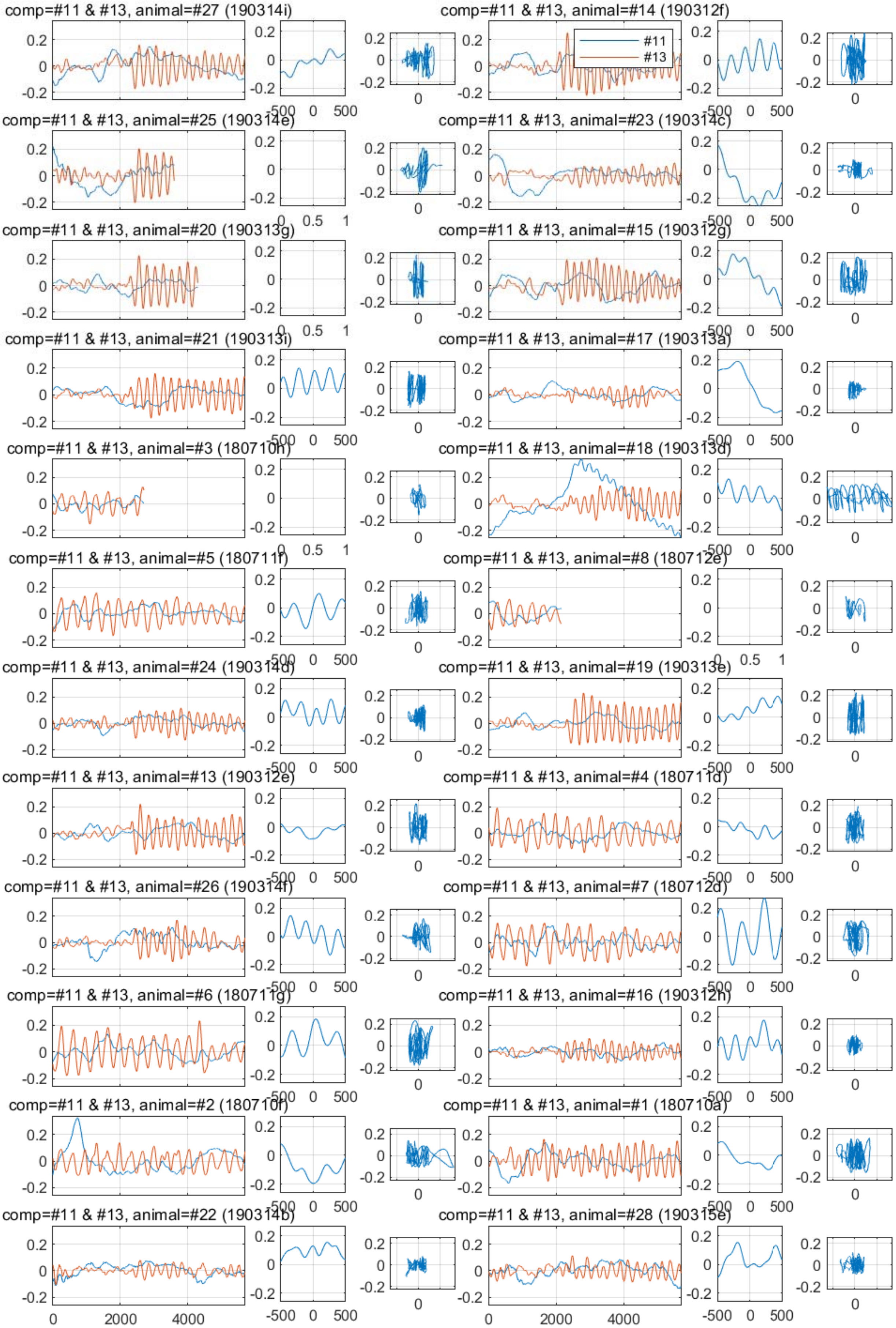

comp=#11 & #13, animal=#27 (190314i)
comp=#11 & #13, animal=#14 (190312f)
#11
#13
comp=#11 & #13, animal=#25 (190314e)
comp=#11 & #13, animal=#23 (190314c)
comp=#11 & #13, animal=#20 (190313g)
comp=#11 & #13, animal=#15 (190312g)
comp=#11 & #13, animal=#21 (190313i)
comp=#11 & #13, animal=#17 (190313a)
comp=#11 & #13, animal=#3 (180710h)
comp=#11 & #13, animal=#18 (190313d)
comp=#11 & #13, animal=#5 (180711f)
comp=#11 & #13, animal=#8 (180712e)
comp=#11 & #13, animal=#24 (190314d)
comp=#11 & #13, animal=#19 (190313e)
comp=#11 & #13, animal=#13 (190312e)
comp=#11 & #13, animal=#4 (180711d)
comp=#11 & #13, animal=#26 (190314f)
comp=#11 & #13, animal=#7 (180712d)
comp=#11 & #13, animal=#6 (180711g)
comp=#11 & #13, animal=#16 (190312h)
comp=#11 & #13, animal=#2 (180710f)
comp=#11 & #13, animal=#1 (180710a)
comp=#11 & #13, animal=#22 (190314b)
comp=#11 & #13, animal=#28 (190315e)

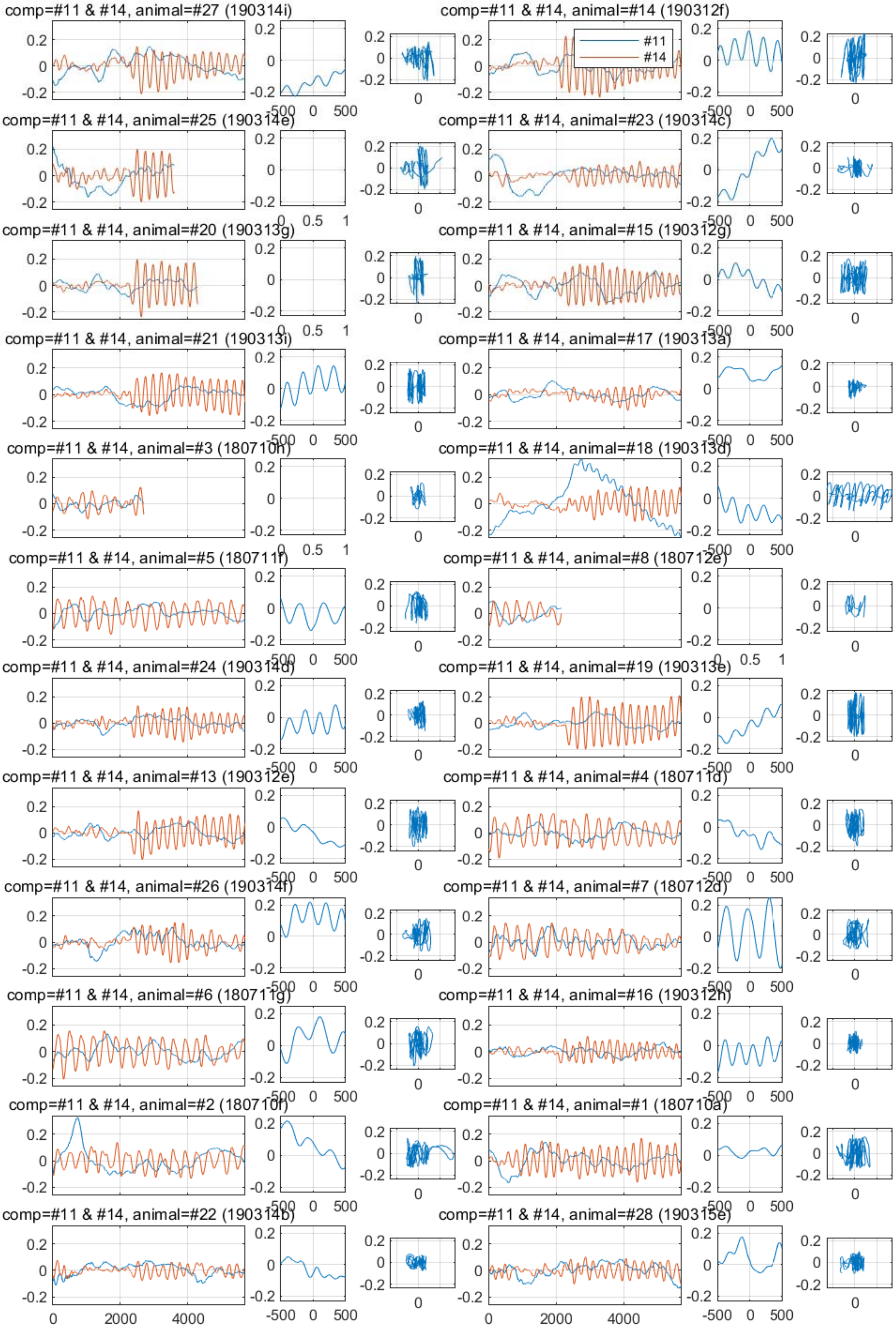
comp=#11 & #14, animal=#27 (190314i)
comp=#11 & #14, animal=#14 (190312f)
#11
#14
comp=#11 & #14, animal=#25 (190314e)
comp=#11 & #14, animal=#23 (190314c)
comp=#11 & #14, animal=#20 (190313g)
comp=#11 & #14, animal=#15 (190312g)
comp=#11 & #14, animal=#21 (190313i)
comp=#11 & #14, animal=#17 (190313a)
comp=#11 & #14, animal=#3 (180710h)
comp=#11 & #14, animal=#18 (190313d)
comp=#11 & #14, animal=#5 (180711f)
comp=#11 & #14, animal=#8 (180712e)
comp=#11 & #14, animal=#24 (190314d)
comp=#11 & #14, animal=#19 (190313e)
comp=#11 & #14, animal=#13 (190312e)
comp=#11 & #14, animal=#4 (180711d)
comp=#11 & #14, animal=#26 (190314f)
comp=#11 & #14, animal=#7 (180712d)
comp=#11 & #14, animal=#6 (180711g)
comp=#11 & #14, animal=#16 (190312h)
comp=#11 & #14, animal=#2 (180710f)
comp=#11 & #14, animal=#1 (180710a)
comp=#11 & #14, animal=#22 (190314b)
comp=#11 & #14, animal=#28 (190315e)

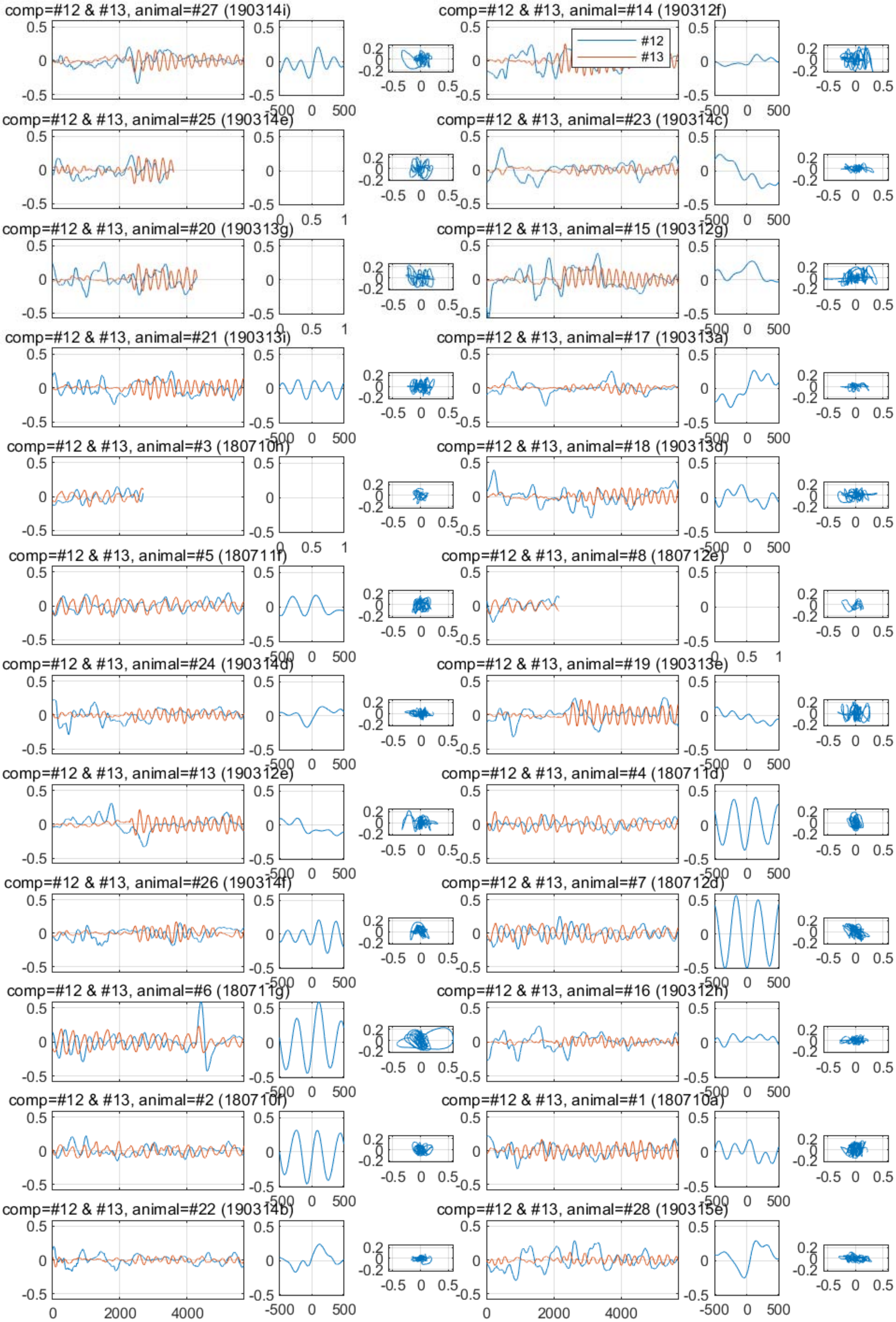
comp=#12 & #13, animal=#27 (190314i)
comp=#12 & #13, animal=#14 (190312f)
#12
#13
comp=#12 & #13, animal=#25 (190314e)
comp=#12 & #13, animal=#23 (190314c)
comp=#12 & #13, animal=#20 (190313g)
comp=#12 & #13, animal=#15 (190312g)
comp=#12 & #13, animal=#21 (190313i)
comp=#12 & #13, animal=#17 (190313a)
comp=#12 & #13, animal=#3 (180710h)
comp=#12 & #13, animal=#18 (190313d)
comp=#12 & #13, animal=#5 (180711f)
comp=#12 & #13, animal=#8 (180712e)
comp=#12 & #13, animal=#24 (190314d)
comp=#12 & #13, animal=#19 (190313e)
comp=#12 & #13, animal=#13 (190312e)
comp=#12 & #13, animal=#4 (180711d)
comp=#12 & #13, animal=#26 (190314f)
comp=#12 & #13, animal=#7 (180712d)
comp=#12 & #13, animal=#6 (180711g)
comp=#12 & #13, animal=#16 (190312h)
comp=#12 & #13, animal=#2 (180710f)
comp=#12 & #13, animal=#1 (180710a)
comp=#12 & #13, animal=#22 (190314b)
comp=#12 & #13, animal=#28 (190315e)
0
2000
4000
-500
0
500

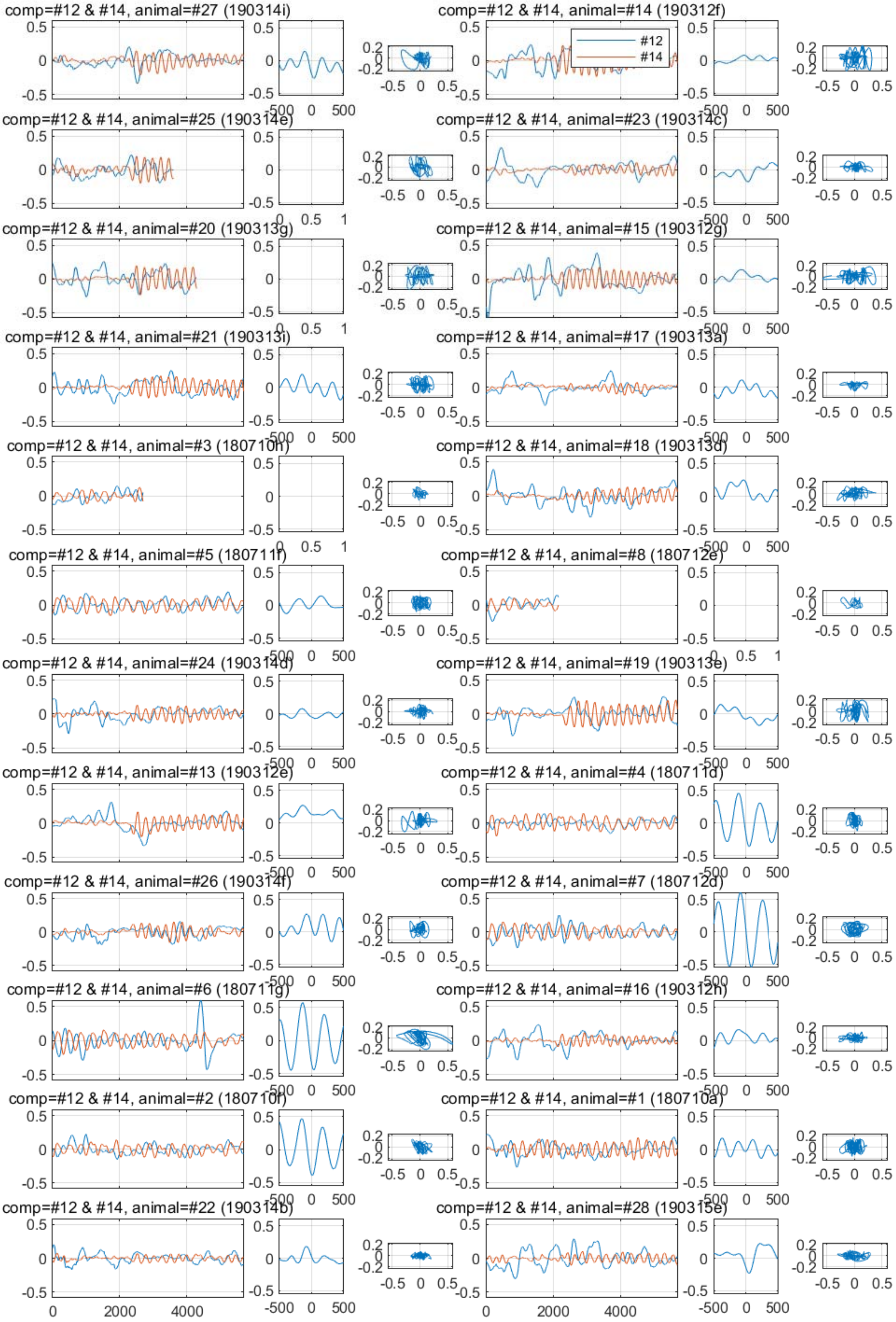
comp=#12 & #14, animal=#27 (190314i)
comp=#12 & #14, animal=#14 (190312f)
#12
#14
comp=#12 & #14, animal=#25 (190314e)
comp=#12 & #14, animal=#23 (190314c)
comp=#12 & #14, animal=#20 (190313g)
comp=#12 & #14, animal=#15 (190312g)
comp=#12 & #14, animal=#21 (190313i)
comp=#12 & #14, animal=#17 (190313a)
comp=#12 & #14, animal=#3 (180710h)
comp=#12 & #14, animal=#18 (190313d)
comp=#12 & #14, animal=#5 (180711f)
comp=#12 & #14, animal=#8 (180712e)
comp=#12 & #14, animal=#24 (190314d)
comp=#12 & #14, animal=#19 (190313e)
comp=#12 & #14, animal=#13 (190312e)
comp=#12 & #14, animal=#4 (180711d)
comp=#12 & #14, animal=#26 (190314f)
comp=#12 & #14, animal=#7 (180712d)
comp=#12 & #14, animal=#6 (180711g)
comp=#12 & #14, animal=#16 (190312h)
comp=#12 & #14, animal=#2 (180710f)
comp=#12 & #14, animal=#1 (180710a)
comp=#12 & #14, animal=#22 (190314b)
comp=#12 & #14, animal=#28 (190315e)

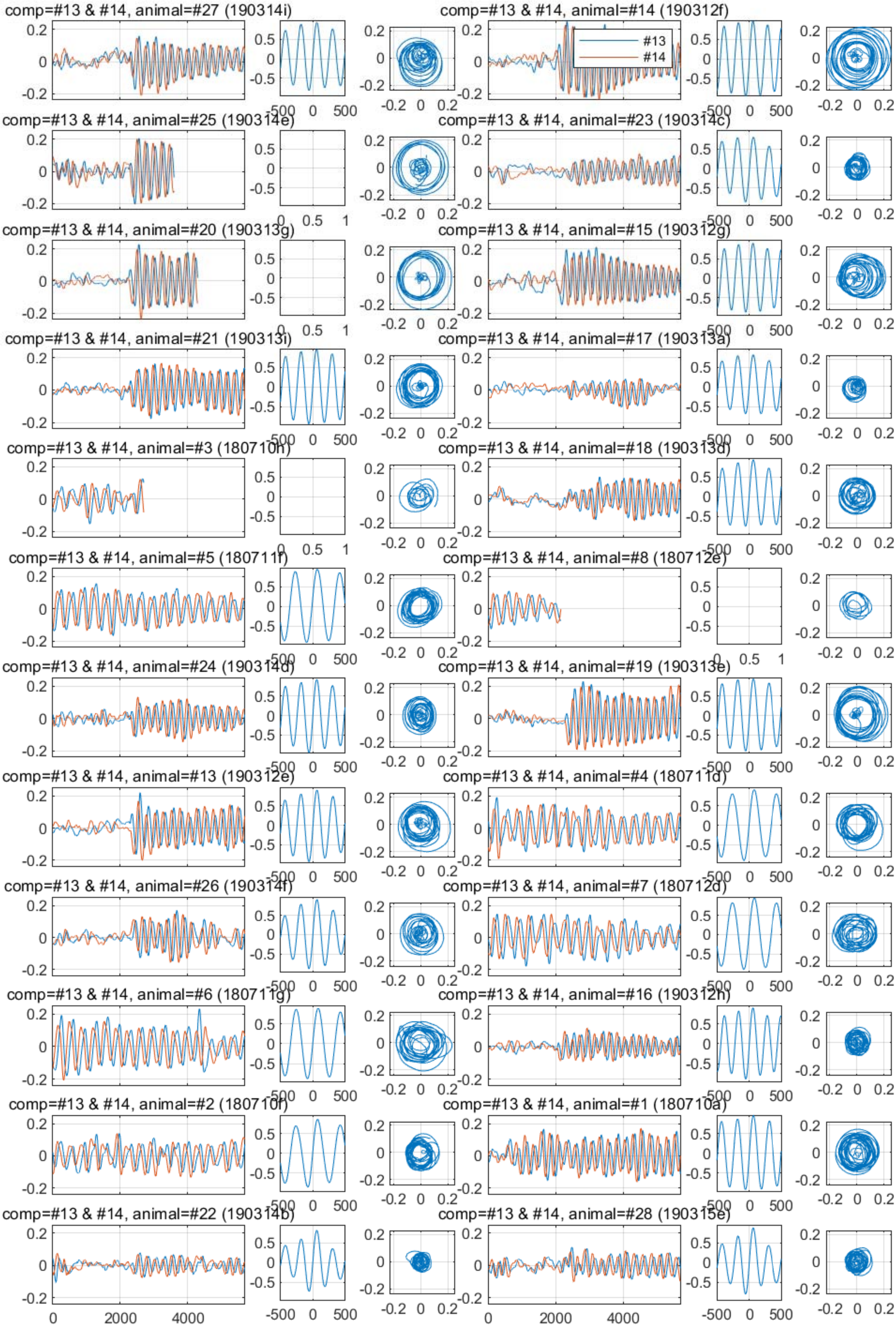
comp=#13 & #14, animal=#27 (190314i)
comp=#13 & #14, animal=#14 (190312f)
#13
#14
comp=#13 & #14, animal=#25 (190314e)
comp=#13 & #14, animal=#23 (190314c)
comp=#13 & #14, animal=#20 (190313g)
comp=#13 & #14, animal=#15 (190312g)
comp=#13 & #14, animal=#21 (190313i)
comp=#13 & #14, animal=#17 (190313a)
comp=#13 & #14, animal=#3 (180710h)
comp=#13 & #14, animal=#18 (190313d)
comp=#13 & #14, animal=#5 (180711f)
comp=#13 & #14, animal=#8 (180712e)
comp=#13 & #14, animal=#24 (190314d)
comp=#13 & #14, animal=#19 (190313e)
comp=#13 & #14, animal=#13 (190312e)
comp=#13 & #14, animal=#4 (180711d)
comp=#13 & #14, animal=#26 (190314f)
comp=#13 & #14, animal=#7 (180712d)
comp=#13 & #14, animal=#6 (180711g)
comp=#13 & #14, animal=#16 (190312h)
comp=#13 & #14, animal=#2 (180710f)
comp=#13 & #14, animal=#1 (180710a)
comp=#13 & #14, animal=#22 (190314b)
comp=#13 & #14, animal=#28 (190315e)
