## Supplementary File 4A for "Deducing ensemble dynamics and information flow from the whole-brain imaging data"

### Sample1

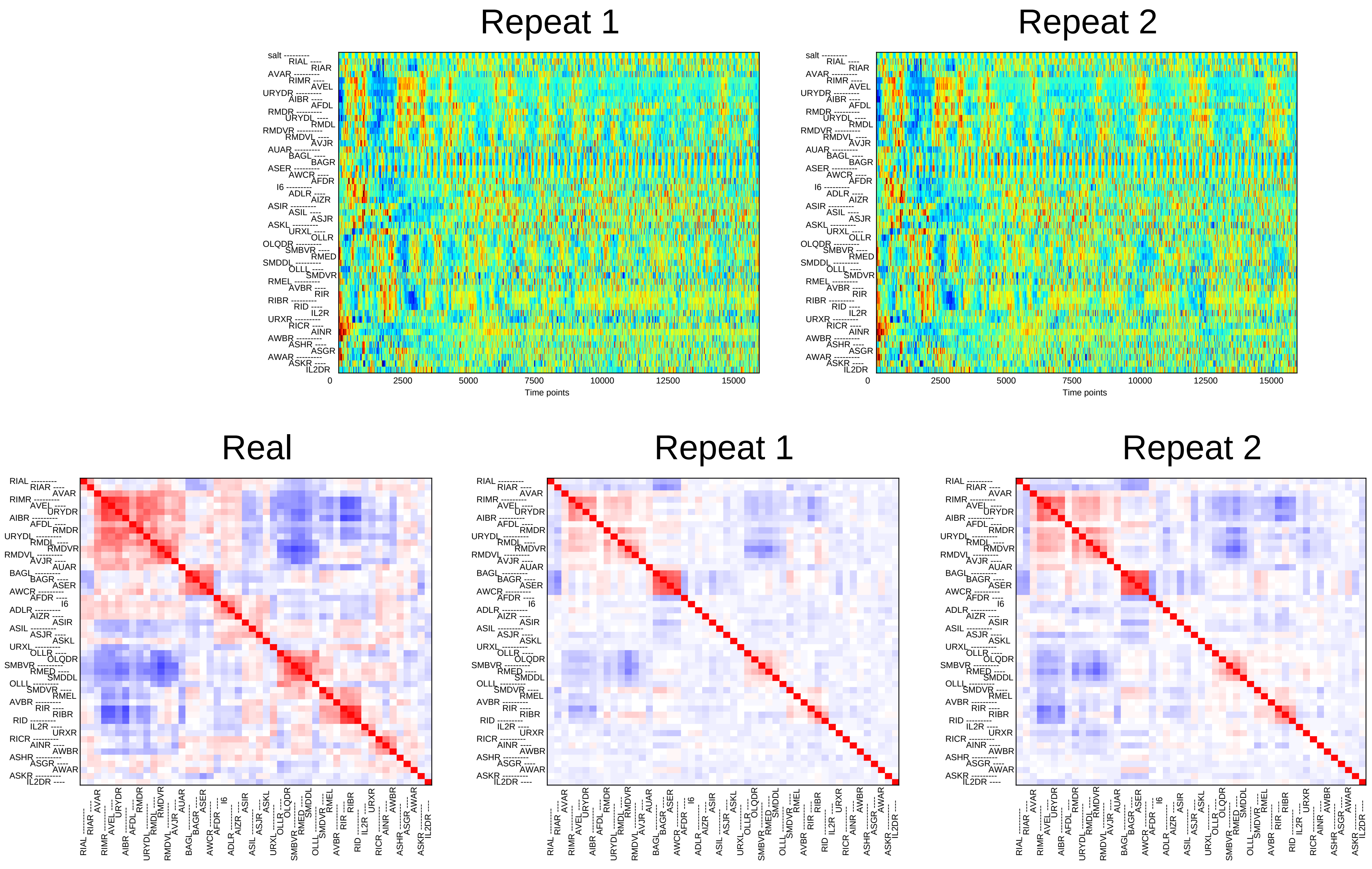
Repeat 1
Repeat 2
salt
RIAL
RIAR
AVAR
RIMR
AVEL
URYDR
AIBR
AFDL
RMDR
URYDL
RMDL
RMDVR
RMDVL
AVJR
AUAR
BAGL
BAGR
ASER
AWCR
AFDR
I6
ADLR
AIZR
ASIR
ASIL
ASJR
ASKL
URXL
OLLR
OLQDR
SMBVR
RMED
SMDDL
OLLL
SMDVR
RMEL
AVBR
RIR
RIBR
RID
IL2R
URXR
RICR
AINR
AWBR
ASHR
ASGR
AWAR
ASKR
IL2DR
0
2500
5000
7500
10000
12500
15000
Time points
Real
Repeat 1
Repeat 2

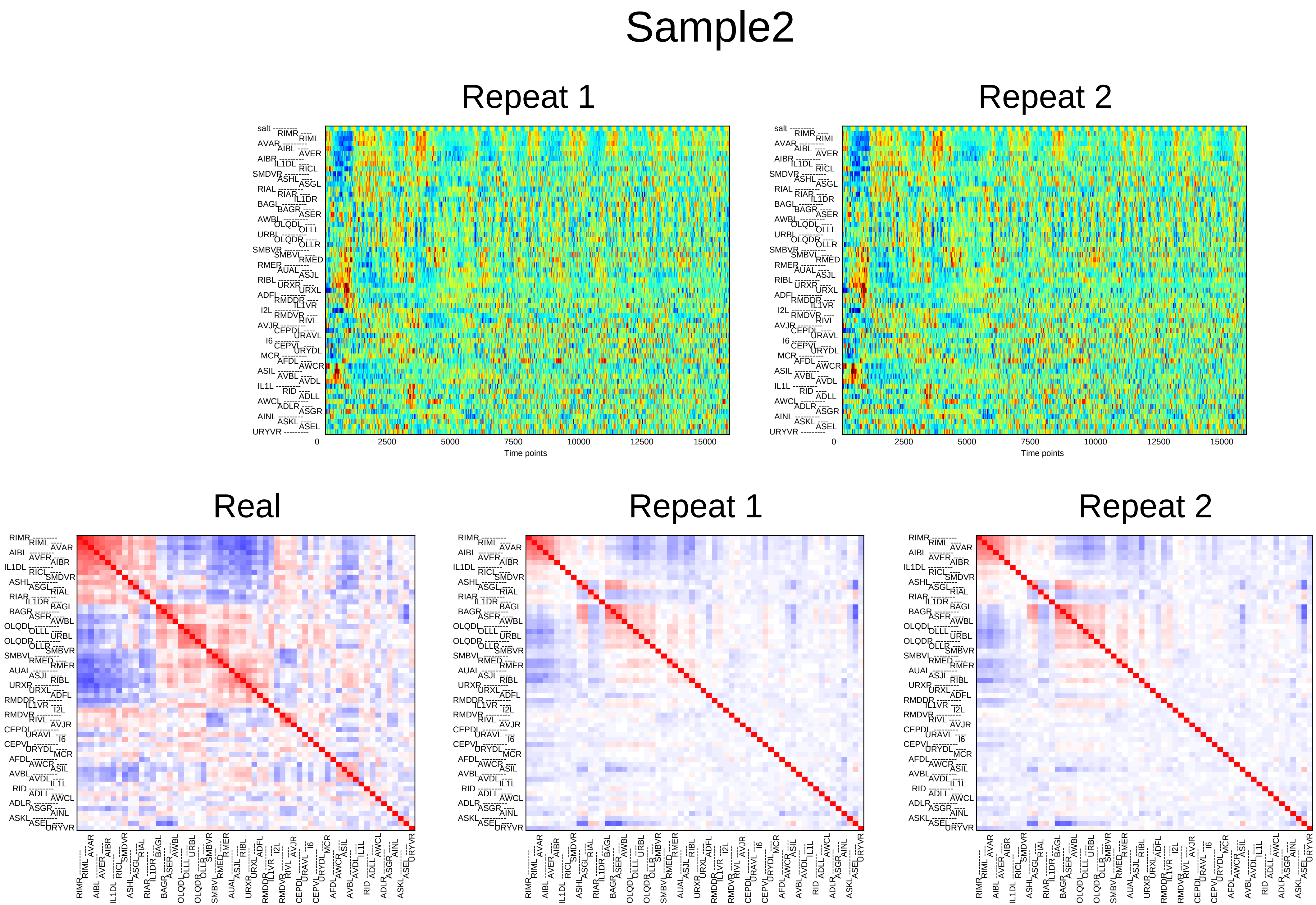

Sample2
Repeat 1
Repeat 2
Time points
0
2500
5000
7500
10000
12500
15000
salt
RIMR
RIML
AVAR
AIBL
AVER
AIBR
IL1DL
RICL
SMDVR
ASHL
ASGL
RIAL
RIAR
IL1DR
BAGL
BAGR
ASER
AWBL
OLQDL
OLLL
URBL
OLQDR
OLLR
SMBVR
SMBVL
RMED
RMER
AUAL
ASJL
RIBL
URXR
URXL
ADFL
RMDDR
IL1VR
I2L
RMDVR
RIVL
AVJR
CEPDL
URAVL
I6
CEPVL
URYDL
MCR
AFDL
AWCR
ASIL
AVBL
AVDL
IL1L
RID
ADLL
AWCL
ADLR
ASGR
AINL
ASKL
ASEL
URYVR
Real
Repeat 1
Repeat 2

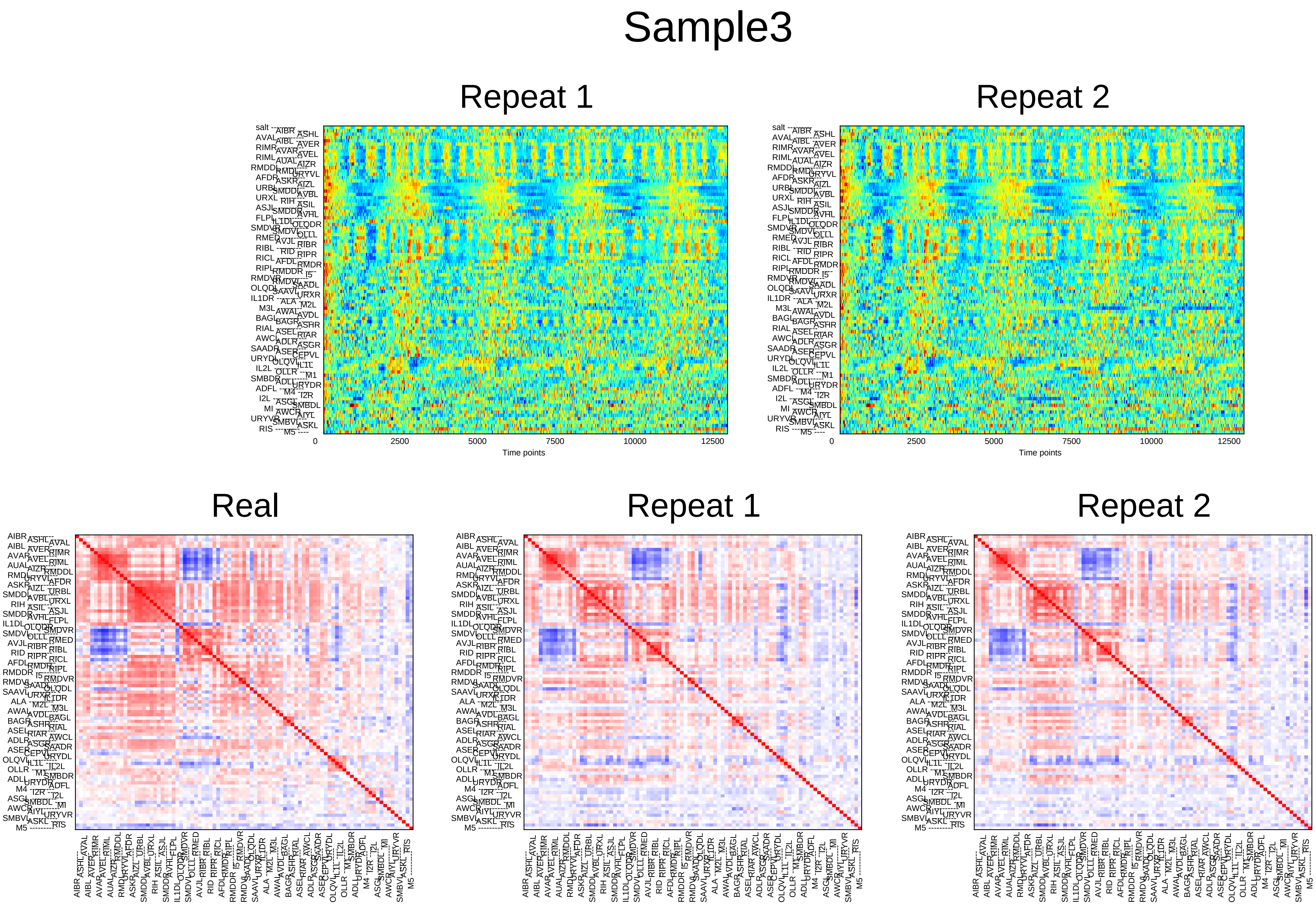
Sample3
Repeat 1
Repeat 2
0
2500
5000
7500
10000
12500
Time points
Real
Repeat 1
Repeat 2

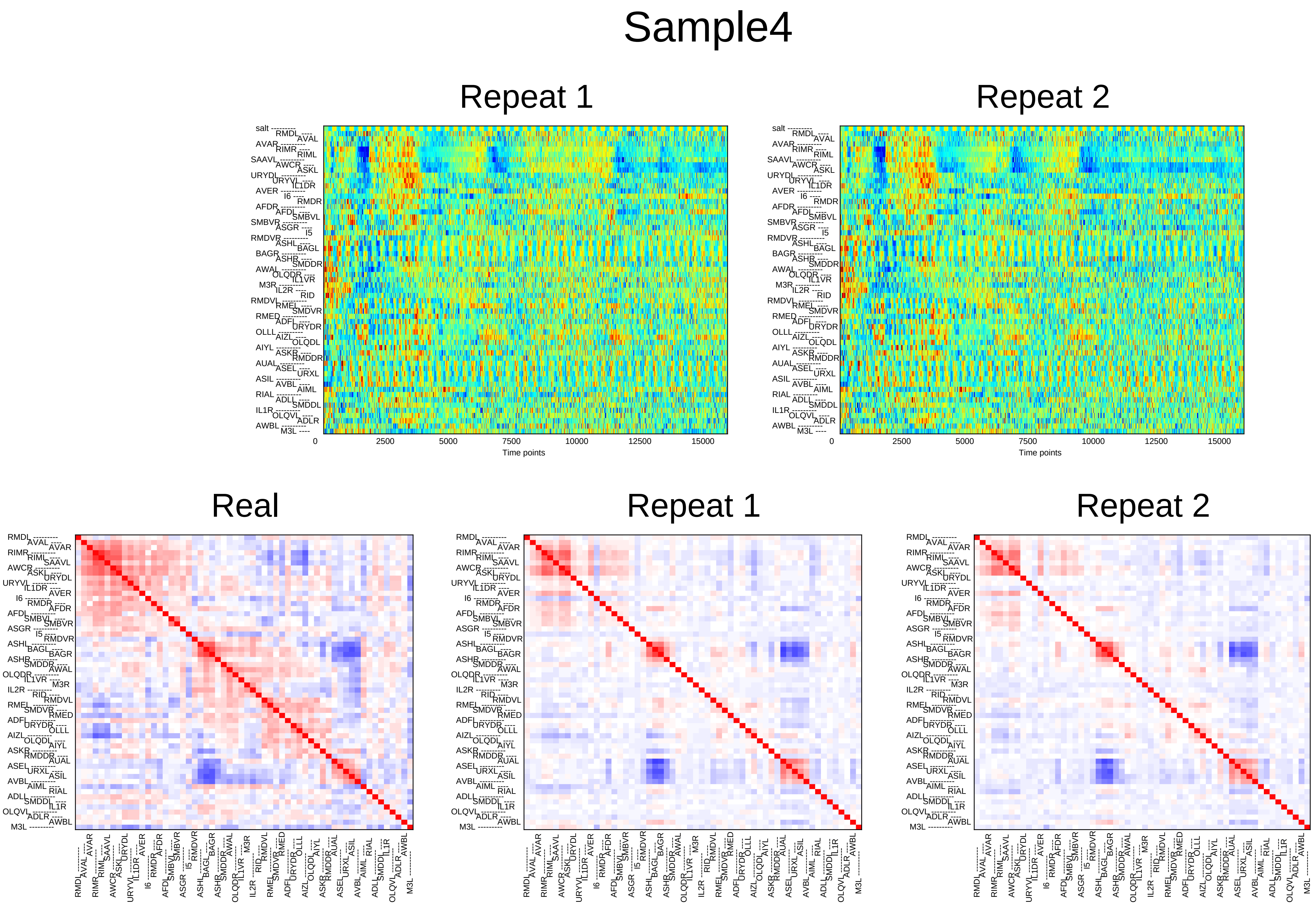
Sample4
Repeat 1
Repeat 2
salt
RMDL
AVAL
AVAR
RIMR
RIML
SAAVL
AWCR
ASKL
URYDL
URYVL
IL1DR
AVER
I6
RMDR
AFDR
AFDL
SMBVL
SMBVR
ASGR
I5
RMDVR
ASHL
BAGL
BAGR
ASHR
SMDDR
AWAL
OLQDR
IL1VR
M3R
IL2R
RID
RMDVL
RMEL
SMDVR
RMED
ADFL
URYDR
OLLL
AIZL
OLQDL
AIYL
ASKR
RMDDR
AUAL
ASEL
URXL
ASIL
AVBL
AIML
RIAL
ADLL
SMDDL
IL1R
OLQVL
ADLR
AWBL
M3L
0
2500
5000
7500
10000
12500
15000
Time points
Real
Repeat 1
Repeat 2

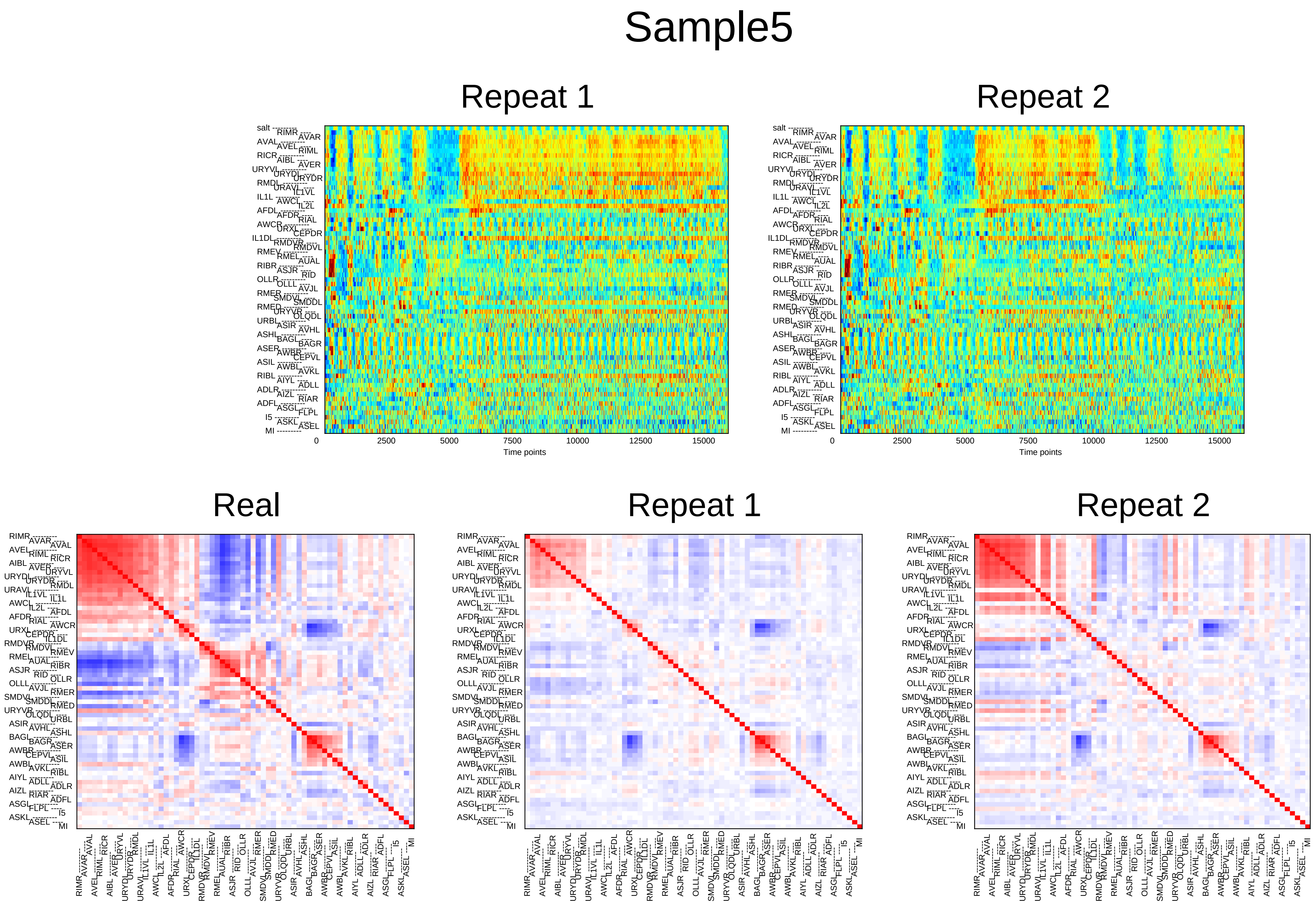

Sample5
Repeat 1
Repeat 2
salt
RIMR
AVAR
AVAL
AVEL
RIML
RICR
AIBL
AVER
URYVL
URYDL
URYDR
RMDL
URAVL
IL1VL
IL1L
AWCL
IL2L
AFDL
AFDR
RIAL
AWCR
URXL
CEPDR
IL1DL
RMDVR
RMDVL
RMEV
RMEL
AUAL
RIBR
ASJR
RID
OLLR
OLLL
AVJL
RMER
SMDVL
SMDDL
RMED
URYVR
OLQDL
URBL
ASIR
AVHL
ASHL
BAGL
BAGR
ASER
AWBR
CEPVL
ASIL
AWBL
AVKL
RIBL
AIYL
ADLL
ADLR
AIZL
RIAR
ADFL
ASGL
FLPL
I5
ASKL
ASEL
MI
0
2500
5000
7500
10000
12500
15000
Time points
Real
Repeat 1
Repeat 2

### Sample6

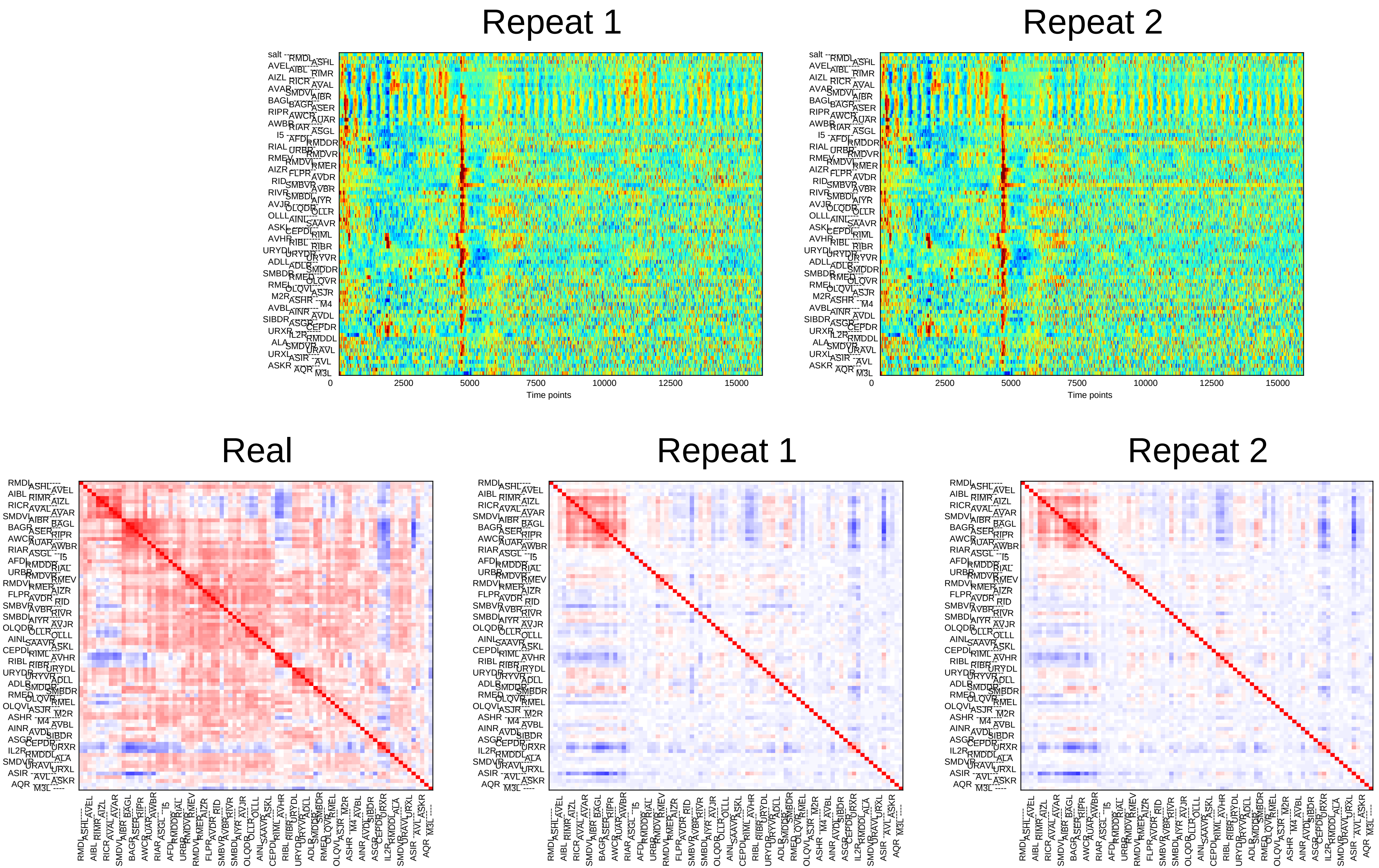
Repeat 1
Repeat 2
0
2500
5000
7500
10000
12500
15000
Time points
Real
Repeat 1
Repeat 2

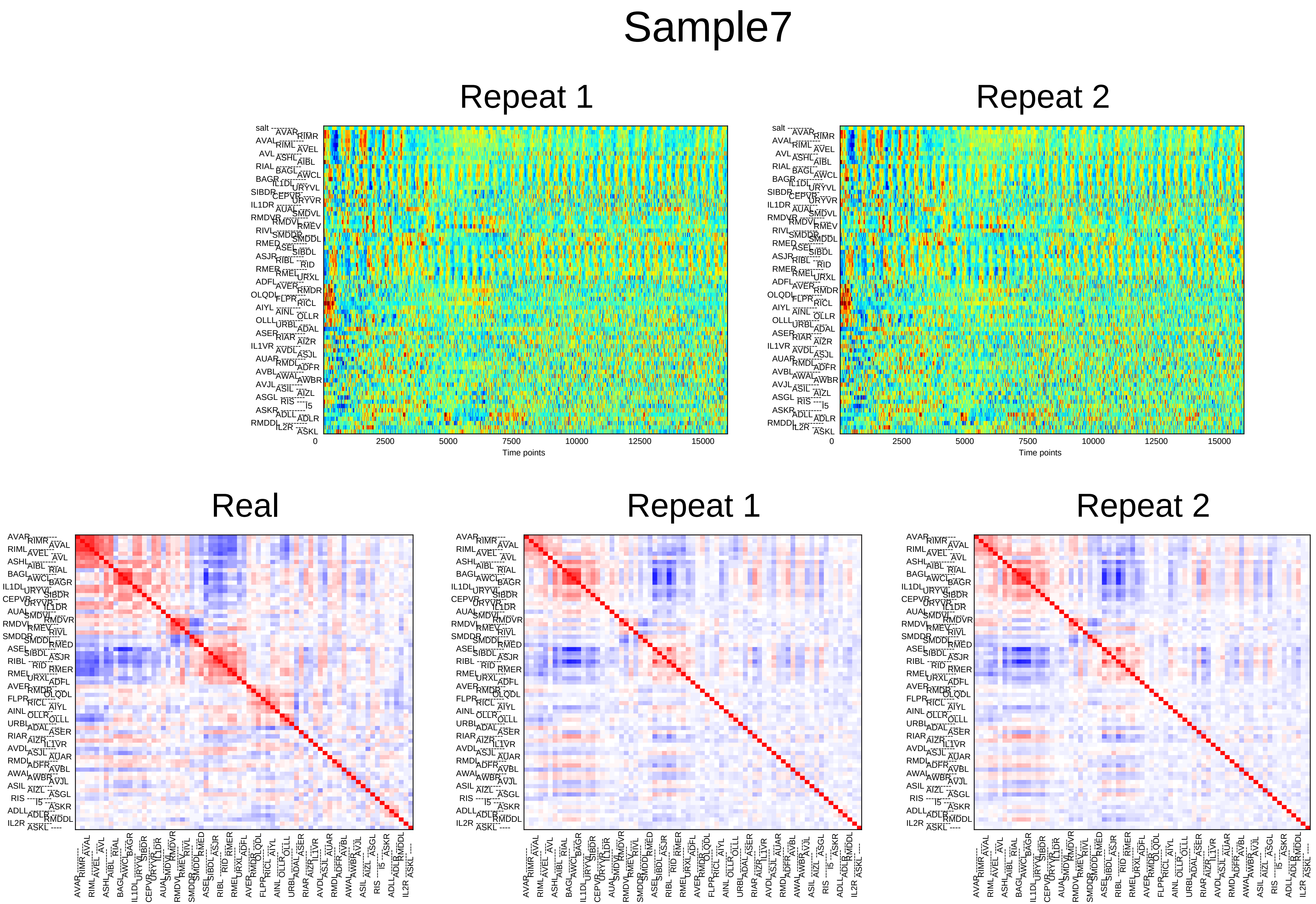
Sample7
Repeat 1
Repeat 2
0
2500
5000
7500
10000
12500
15000
Time points
Real
Repeat 1
Repeat 2

### Sample8

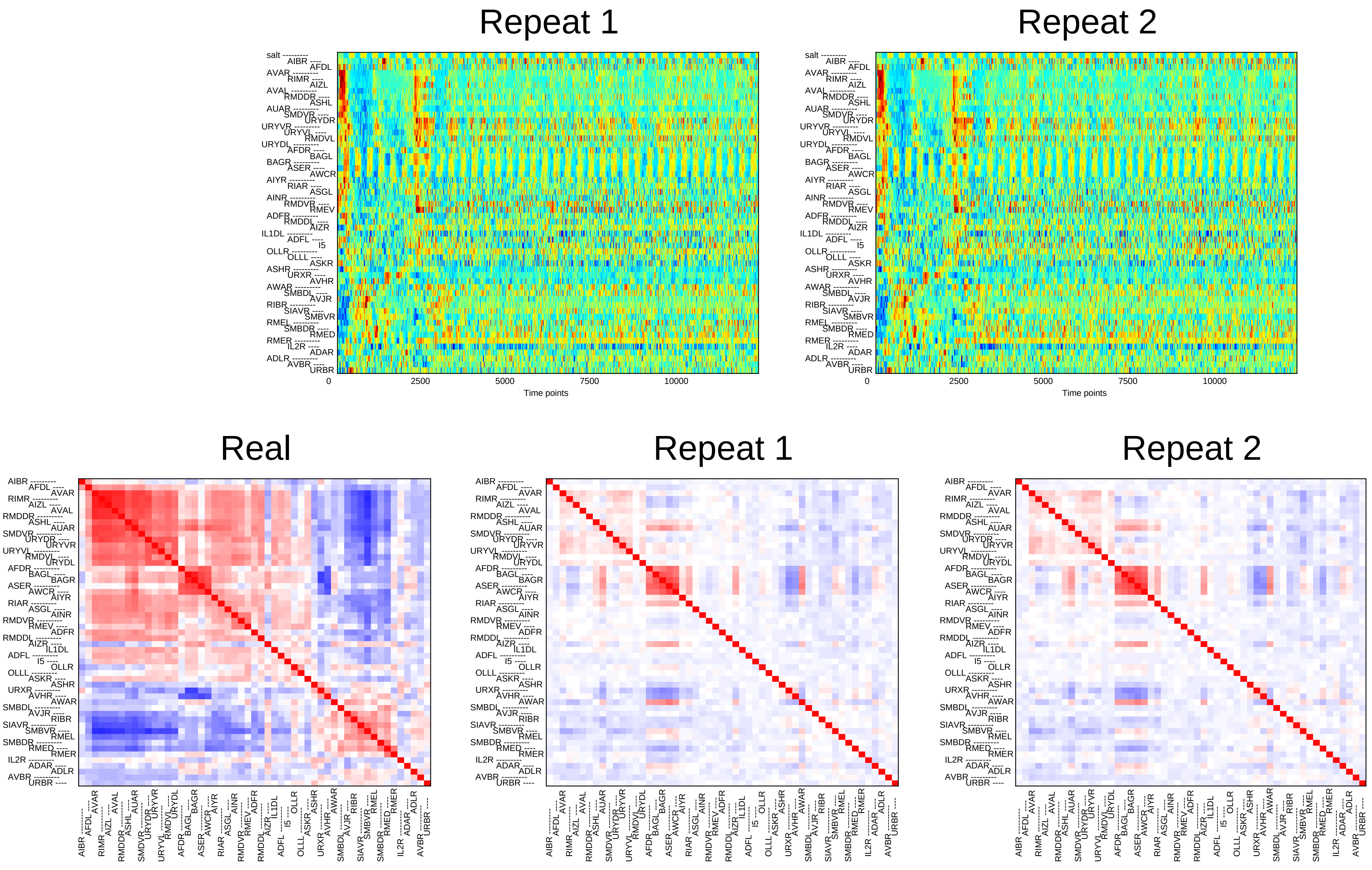
Repeat 1
Repeat 2
salt
AIBR AFDL AVAR RIMR AIZL AVAL RMDDR ASHL AUAR SMDVR URYDR URYVR URYVL RMDVL URYDL AFDR BAGL BAGR ASER AWCR AIYR RIAR ASGL AINR RMDVR RMEV ADFR RMDDL AIZR IL1DL ADFL I5 OLLR OLLL ASKR ASHR URXR AVHR AWAR SMBDL AVJR RIBR SIAVR SMBVR RMEL SMBDR RMED RMER IL2R ADAR ADLR AVBR URBR
0
2500
5000
7500
10000
Time points
Real
Repeat 1
Repeat 2

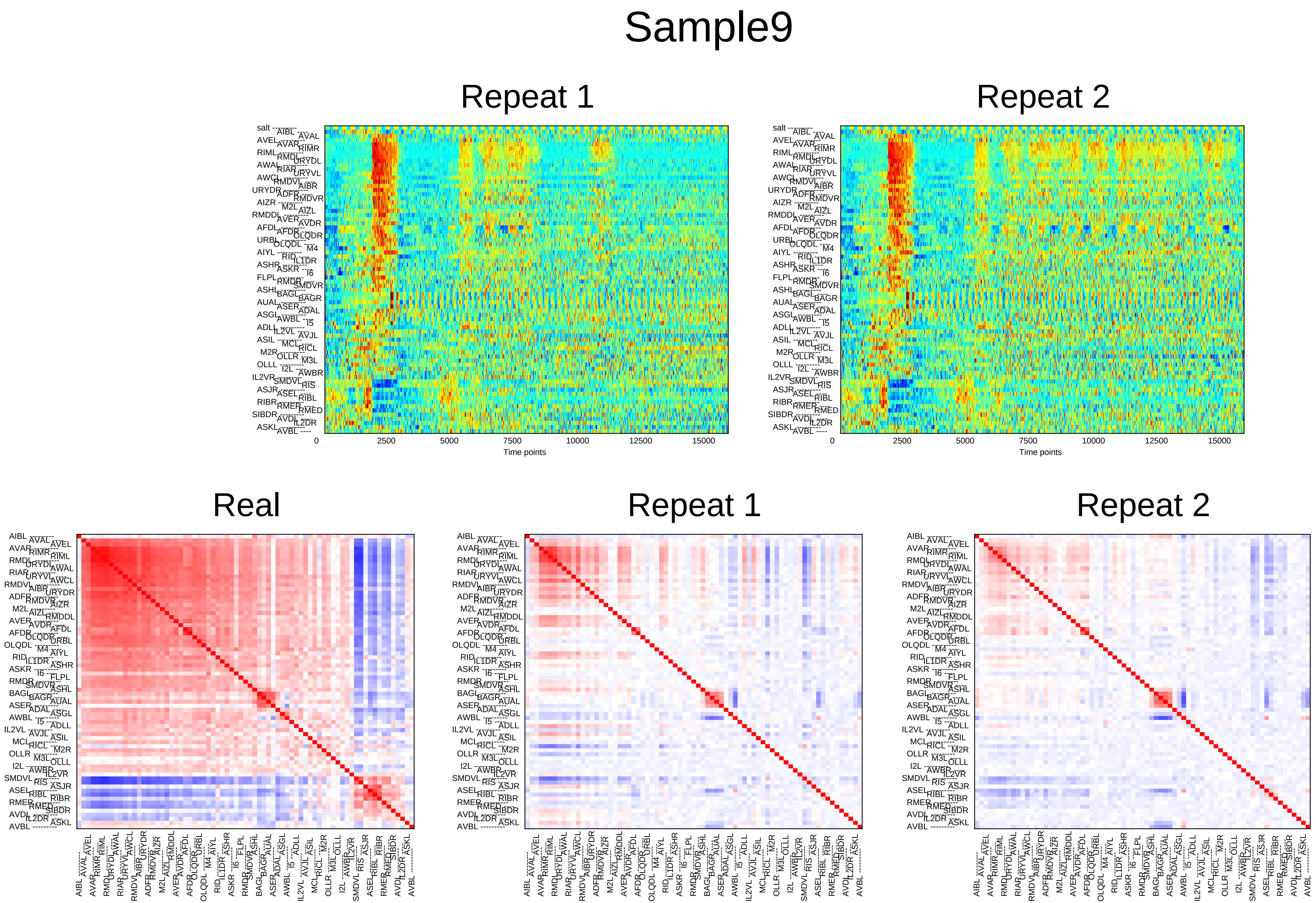
Sample9
Repeat 1
Repeat 2
0
2500
5000
7500
10000
12500
15000
Time points
Real
Repeat 1
Repeat 2

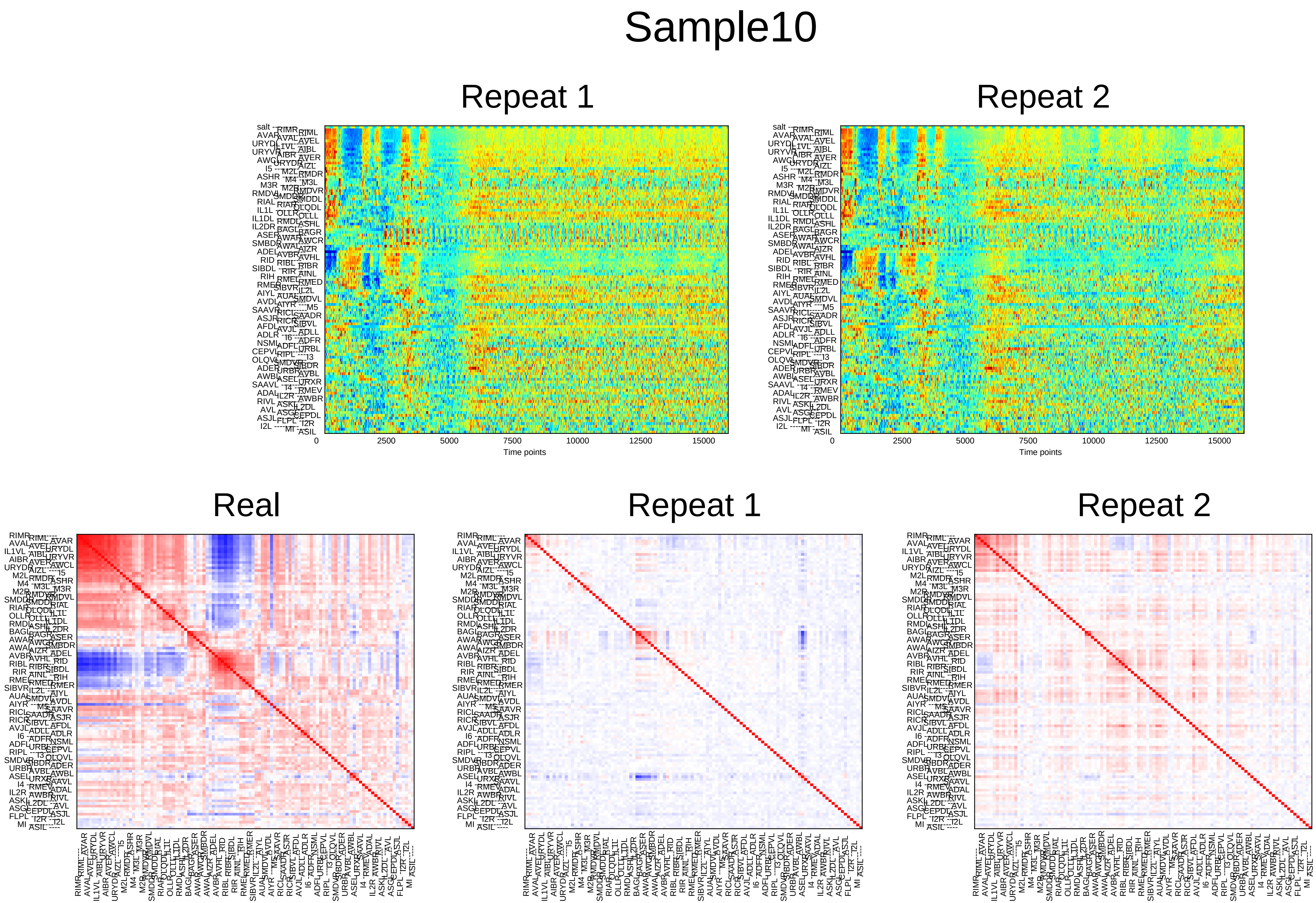
Sample10
Repeat 1
Repeat 2
0
2500
5000
7500
10000
12500
15000
Time points
Real
Repeat 1
Repeat 2

### Sample11

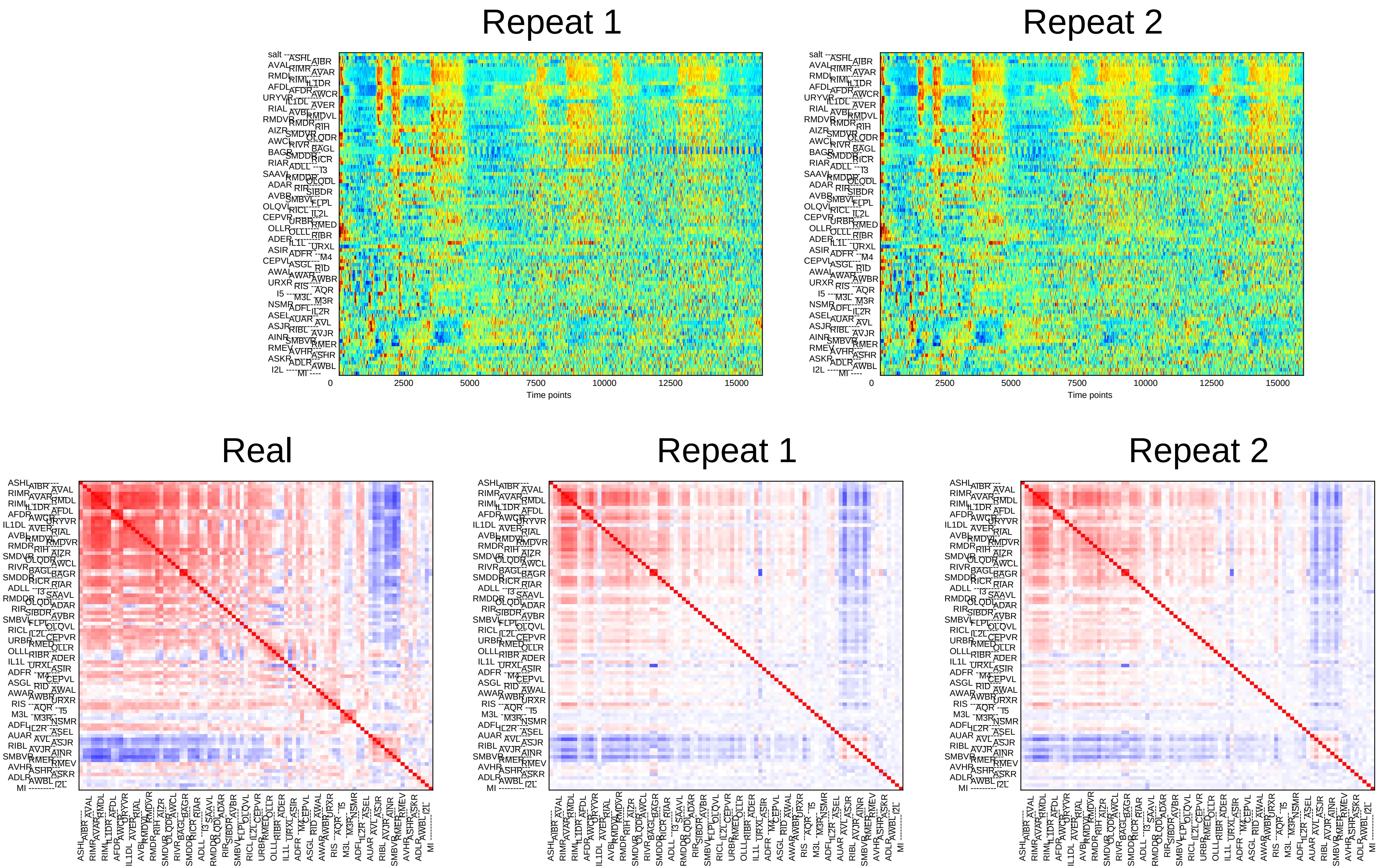
Repeat 1
Repeat 2
0
2500
5000
7500
10000
12500
15000
Time points
Real
Repeat 1
Repeat 2

### Sample12

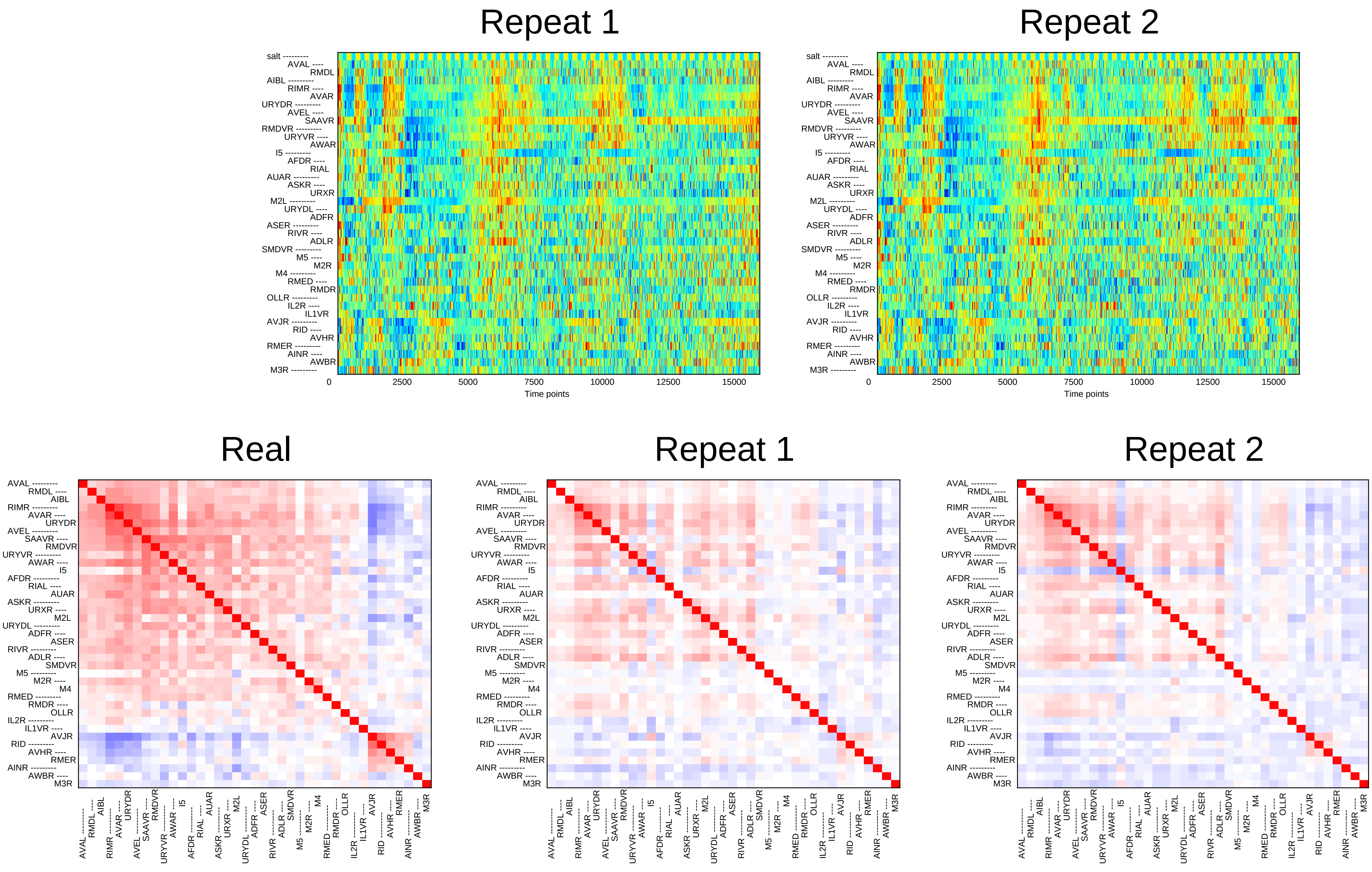

Repeat 1
Repeat 2
salt
AVAL
RMDL
AIBL
RIMR
AVAR
URYDR
AVEL
SAAVR
RMDVR
URYVR
AWAR
I5
AFDR
RIAL
AUAR
ASKR
URXR
M2L
URYDL
ADFR
ASER
RIVR
ADLR
SMDVR
M5
M2R
M4
RMED
RMDR
OLLR
IL2R
IL1VR
AVJR
RID
AVHR
RMER
AINR
AWBR
M3R
0
2500
5000
7500
10000
12500
15000
Time points
Real

### Sample13

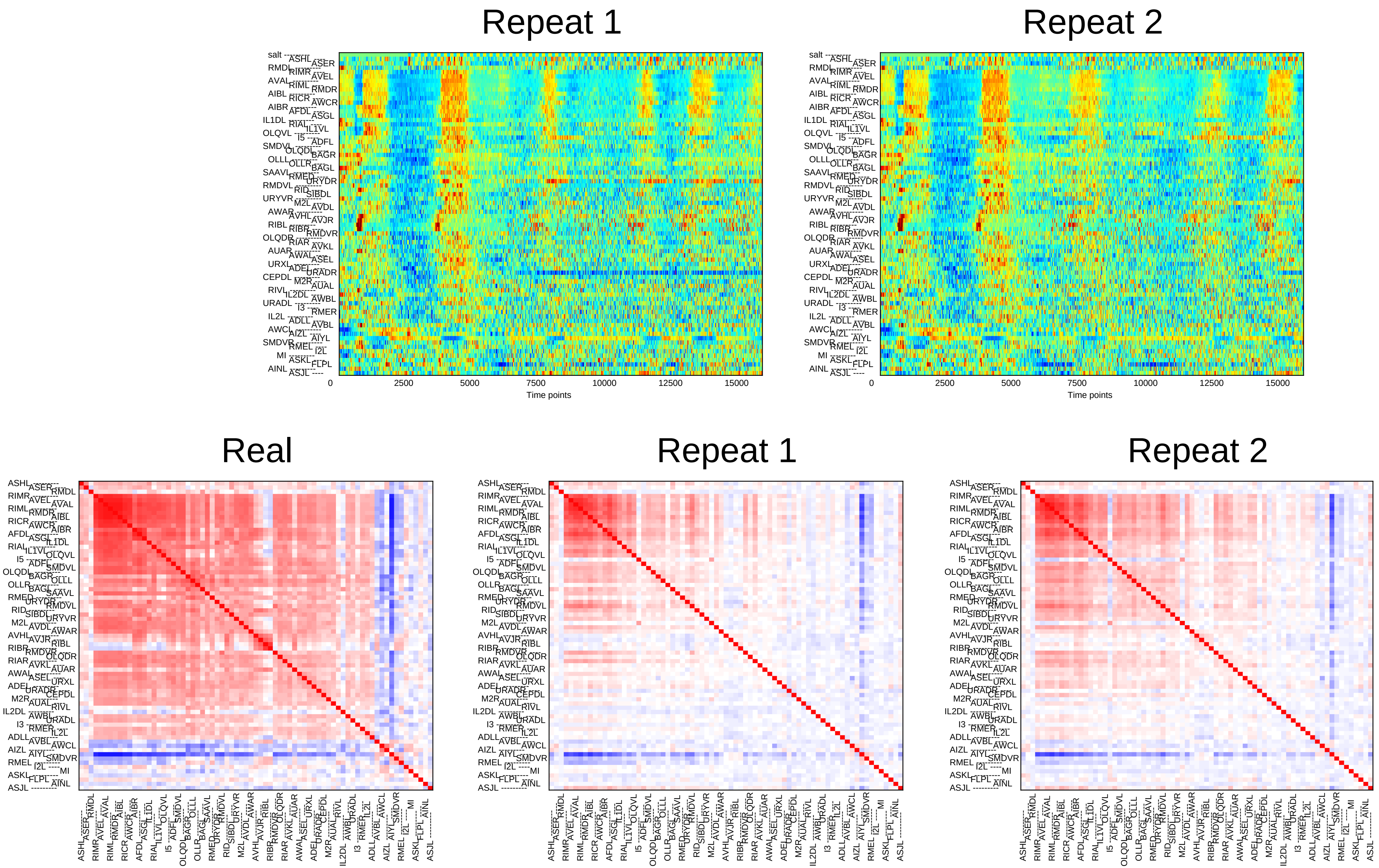

Repeat 1
Repeat 2
0
2500
5000
7500
10000
12500
15000
Time points
Real
Repeat 1
Repeat 2

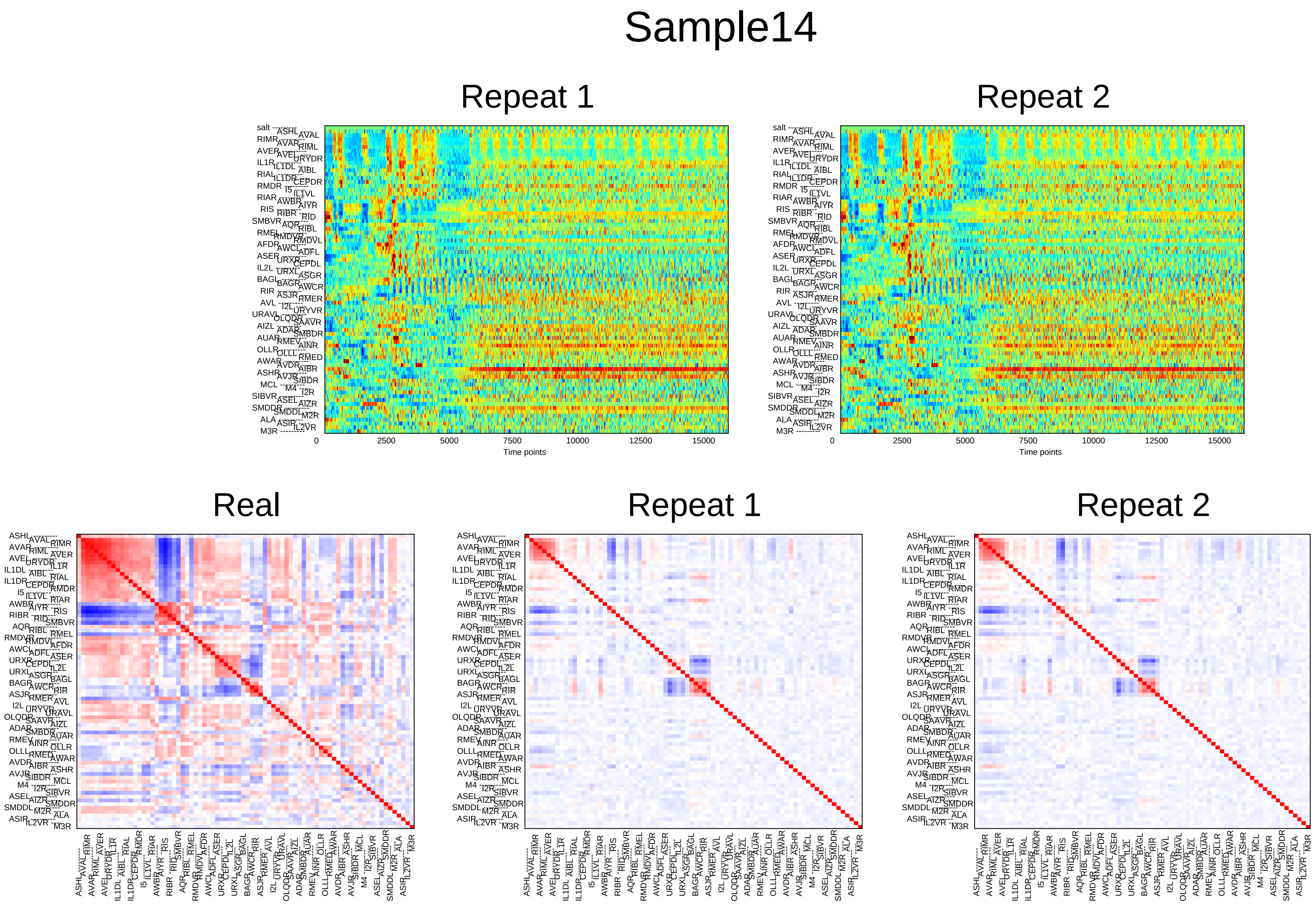
Sample14
Repeat 1
Repeat 2
0
2500
5000
7500
10000
12500
15000
Time points
Real
Repeat 1
Repeat 2

### Sample15

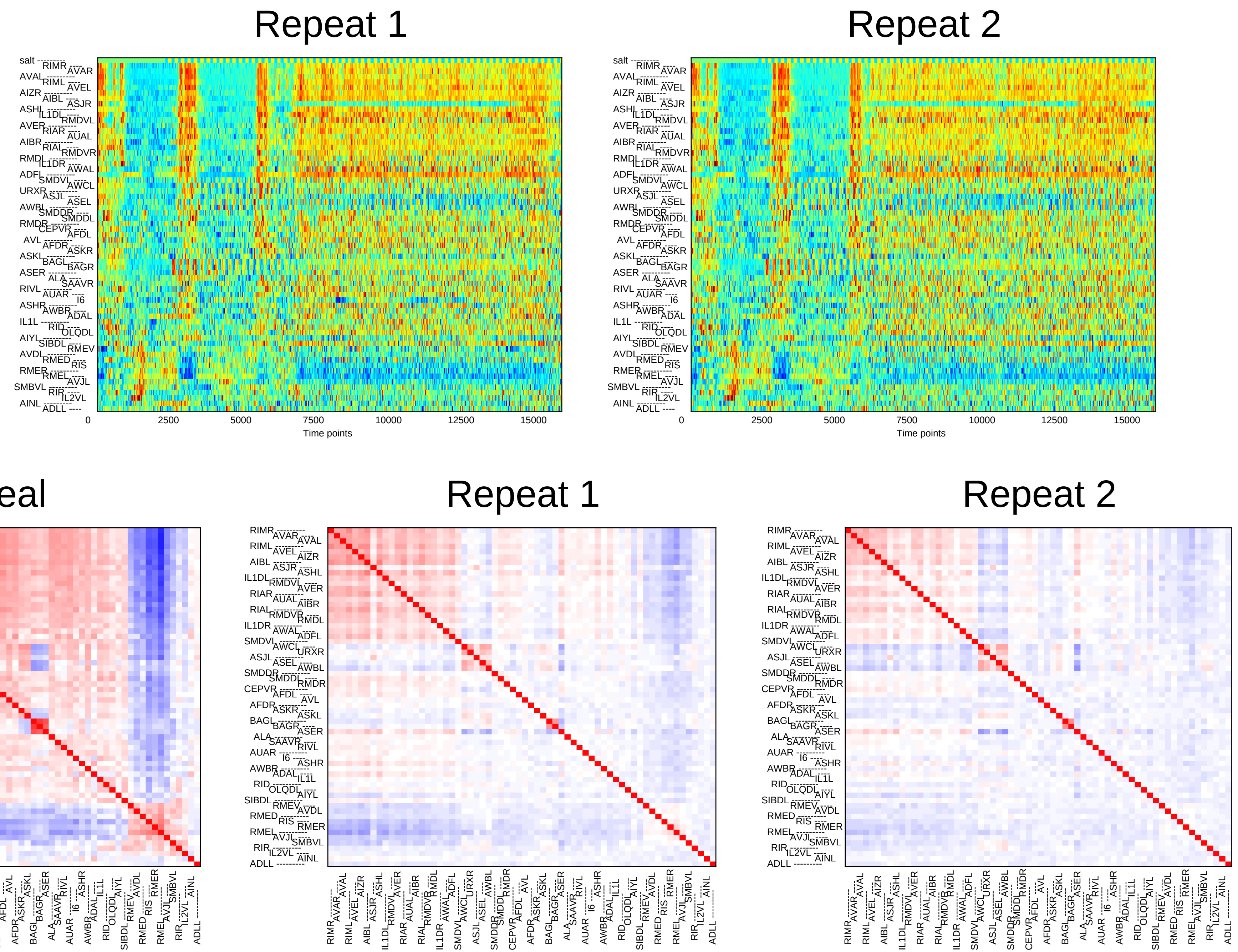
Repeat 1
Repeat 2
Time points
Time points
Real
Repeat 1
Repeat 2

### Sample16

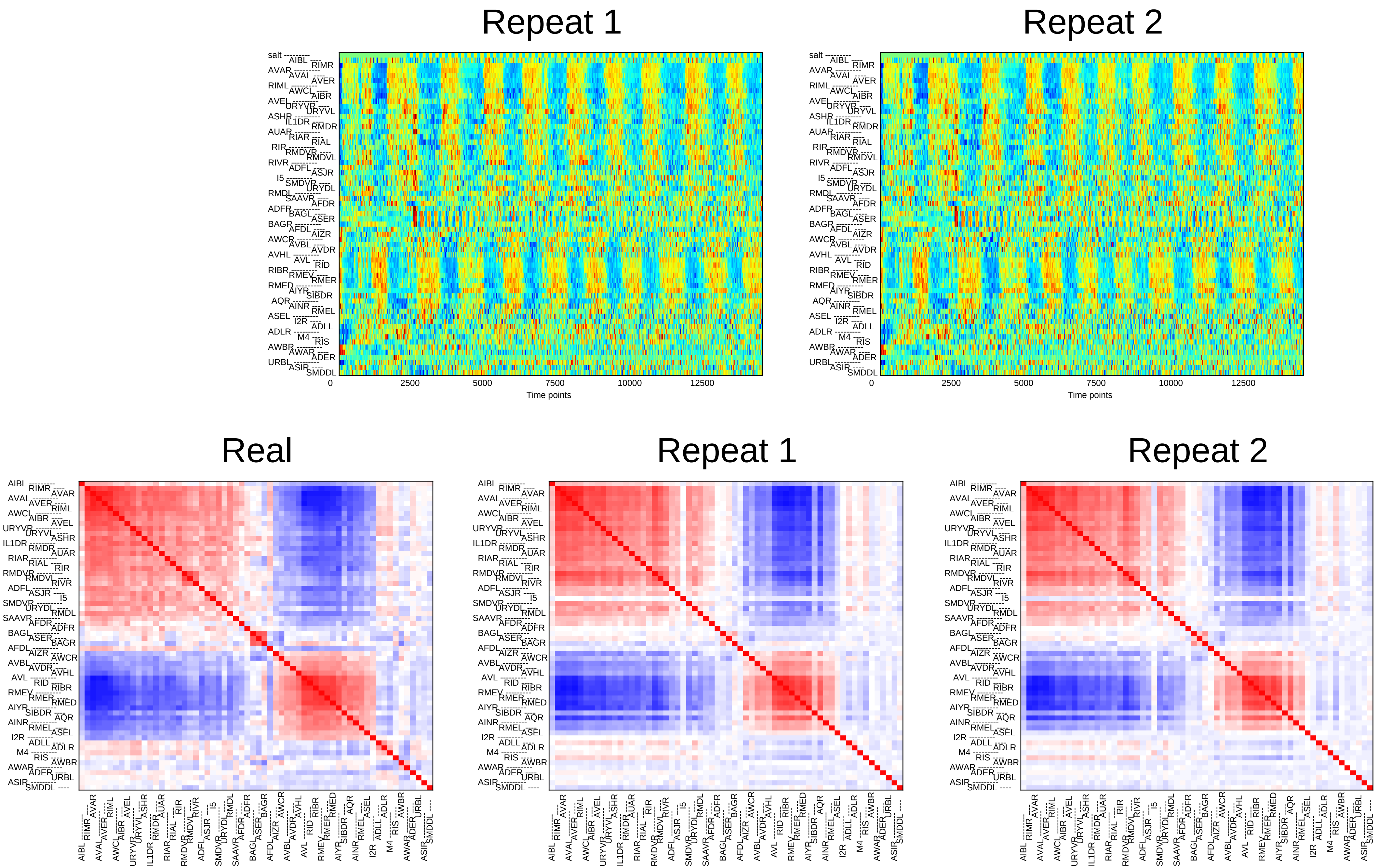
Repeat 1
Repeat 2
0
2500
5000
7500
10000
12500
Time points
Real
Repeat 1
Repeat 2

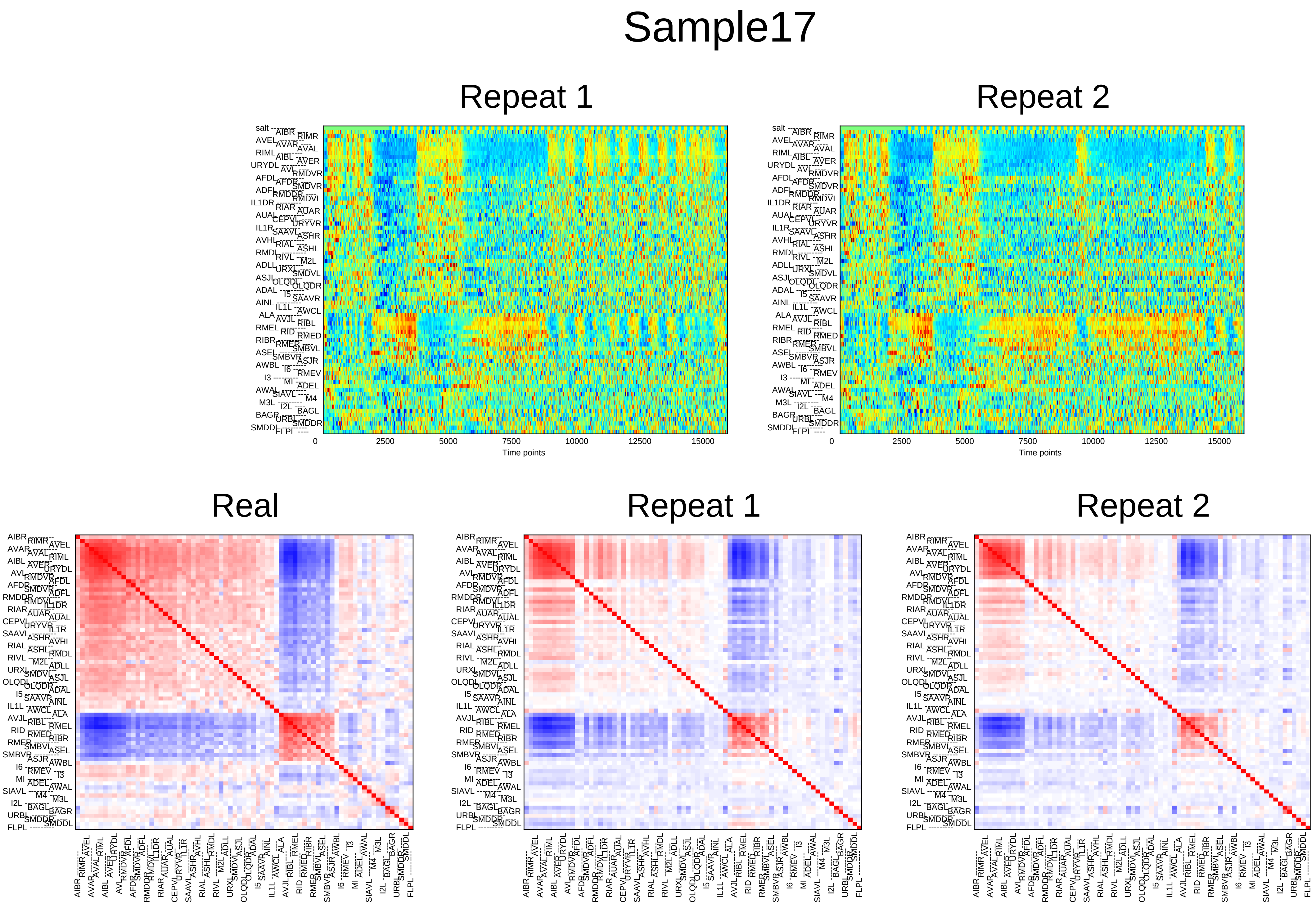
Sample17
Repeat 1
Repeat 2
Time points
Real
Repeat 1
Repeat 2

### Sample18

#### Repeat 1

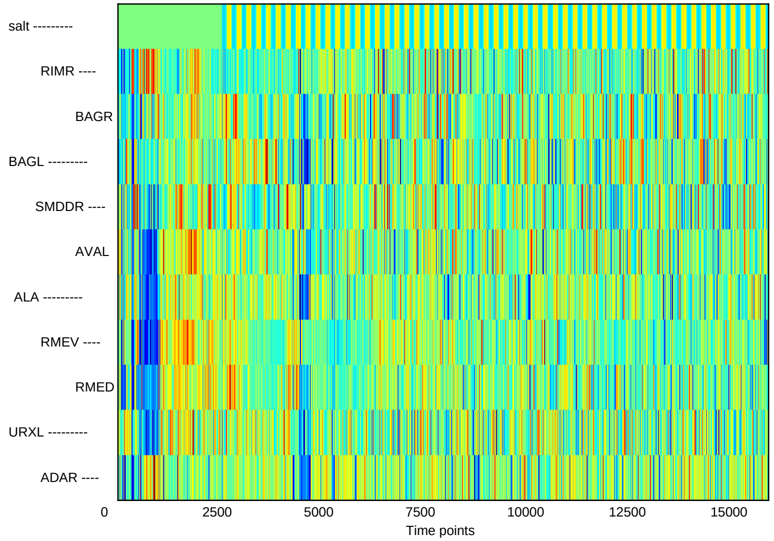


#### Repeat 2

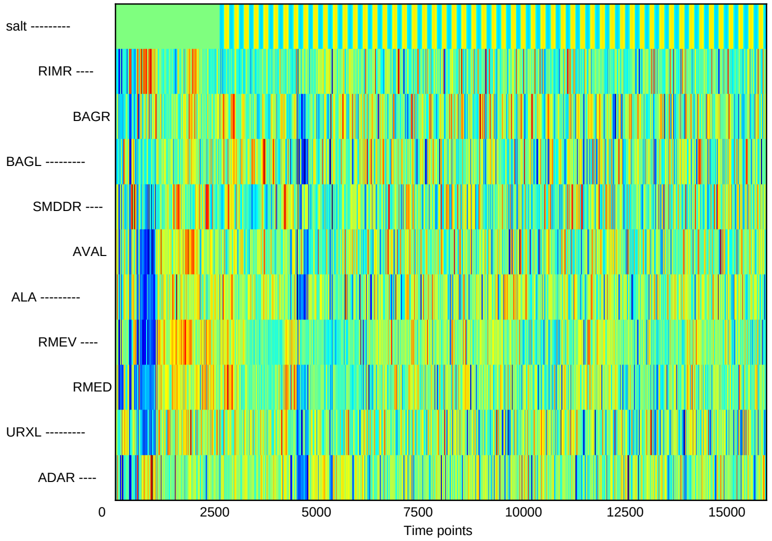


#### Real

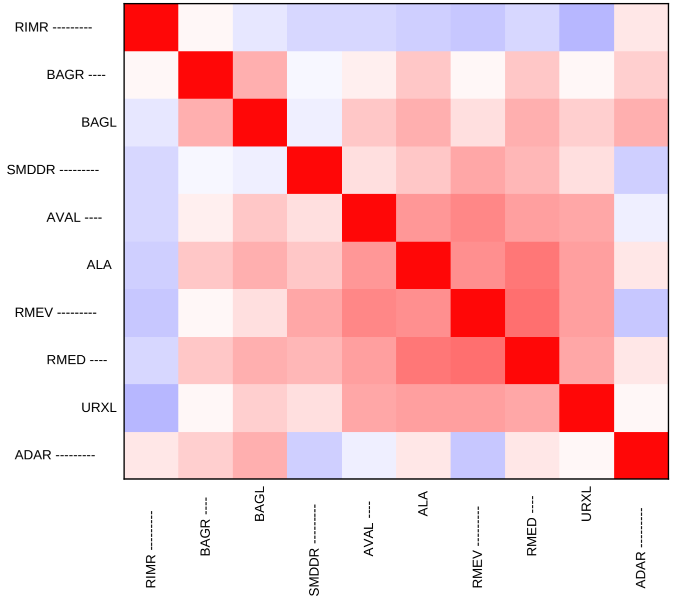


#### Repeat 1

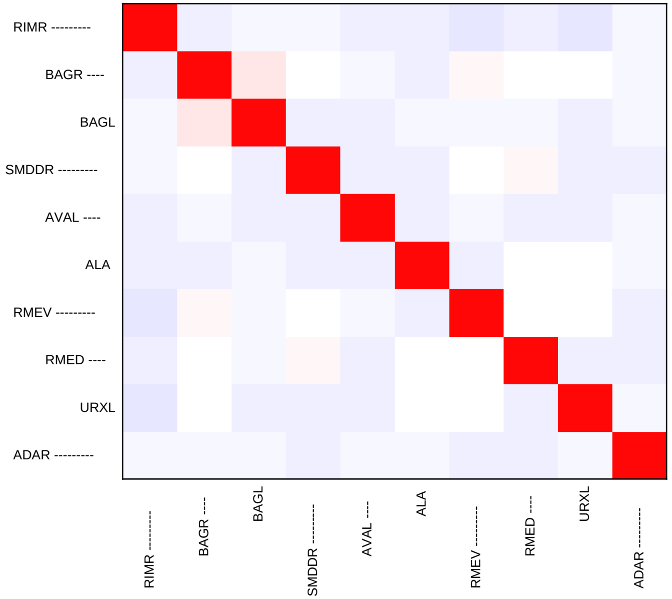


#### Repeat 2

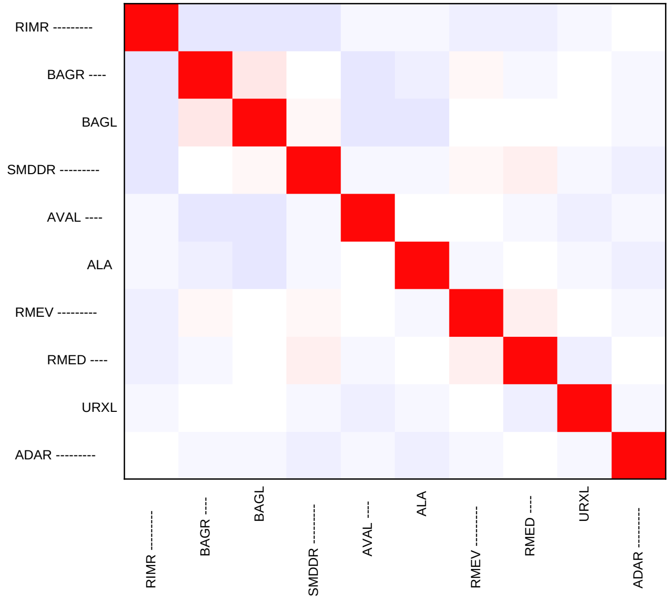

### Sample19

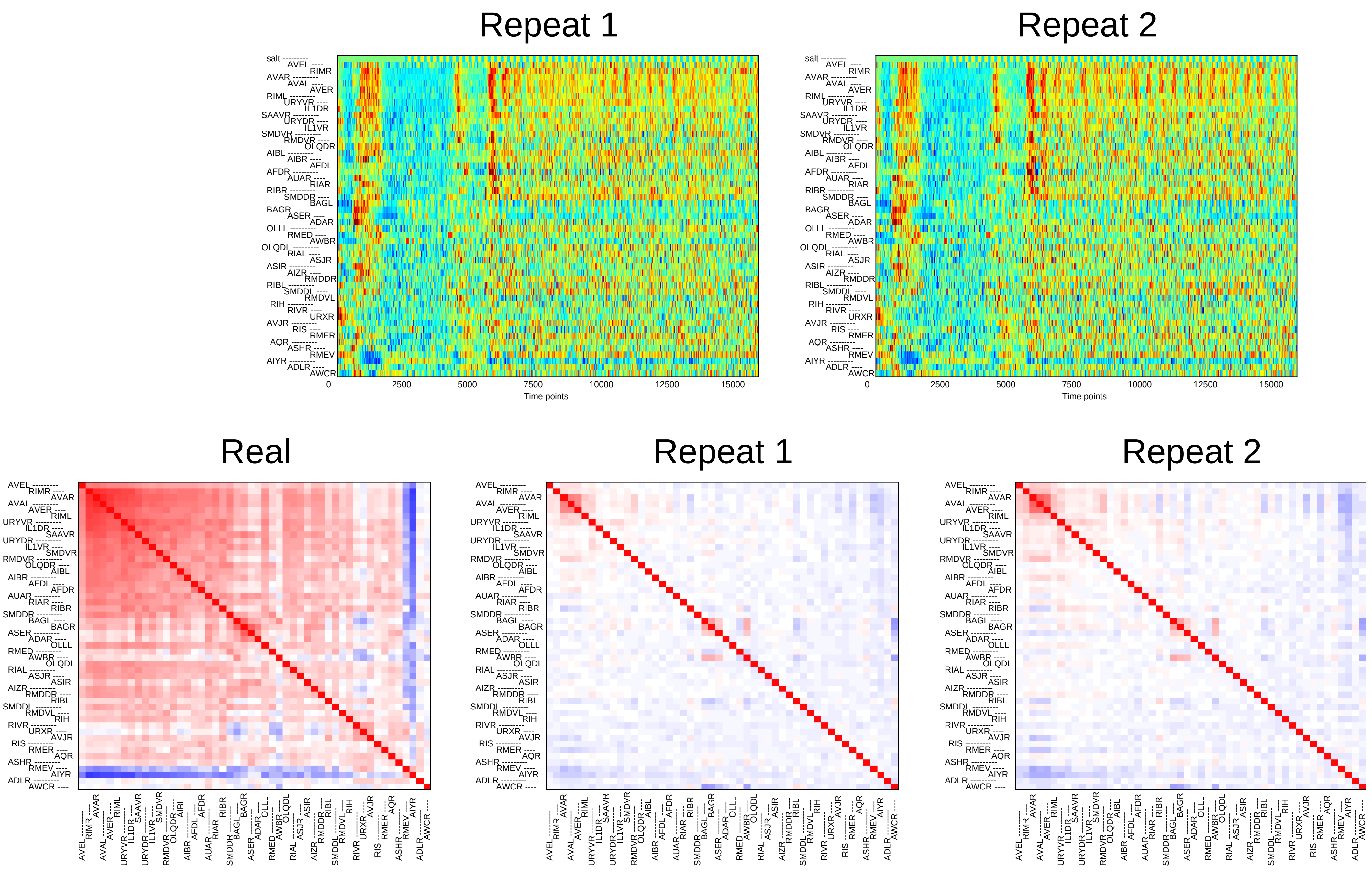

Repeat 1
Repeat 2
salt
AVEL
RIMR
AVAR
AVAL
AVER
RIML
URYVR
IL1DR
SAAVR
URYDR
IL1VR
SMDVR
RMDVR
OLQDR
AIBL
AIBR
AFDL
AFDR
AUAR
RIAR
RIBR
SMDDR
BAGL
BAGR
ASER
ADAR
OLLL
RMED
AWBR
OLQDL
RIAL
ASJR
ASIR
AIZR
RMDDR
RIBL
SMDDL
RMDVL
RIH
RIVR
URXR
AVJR
RIS
RMER
AQR
ASHR
RMEV
AIYR
ADLR
AWCR
0
2500
5000
7500
10000
12500
15000
Time points
Real
Repeat 1
Repeat 2

### Sample20

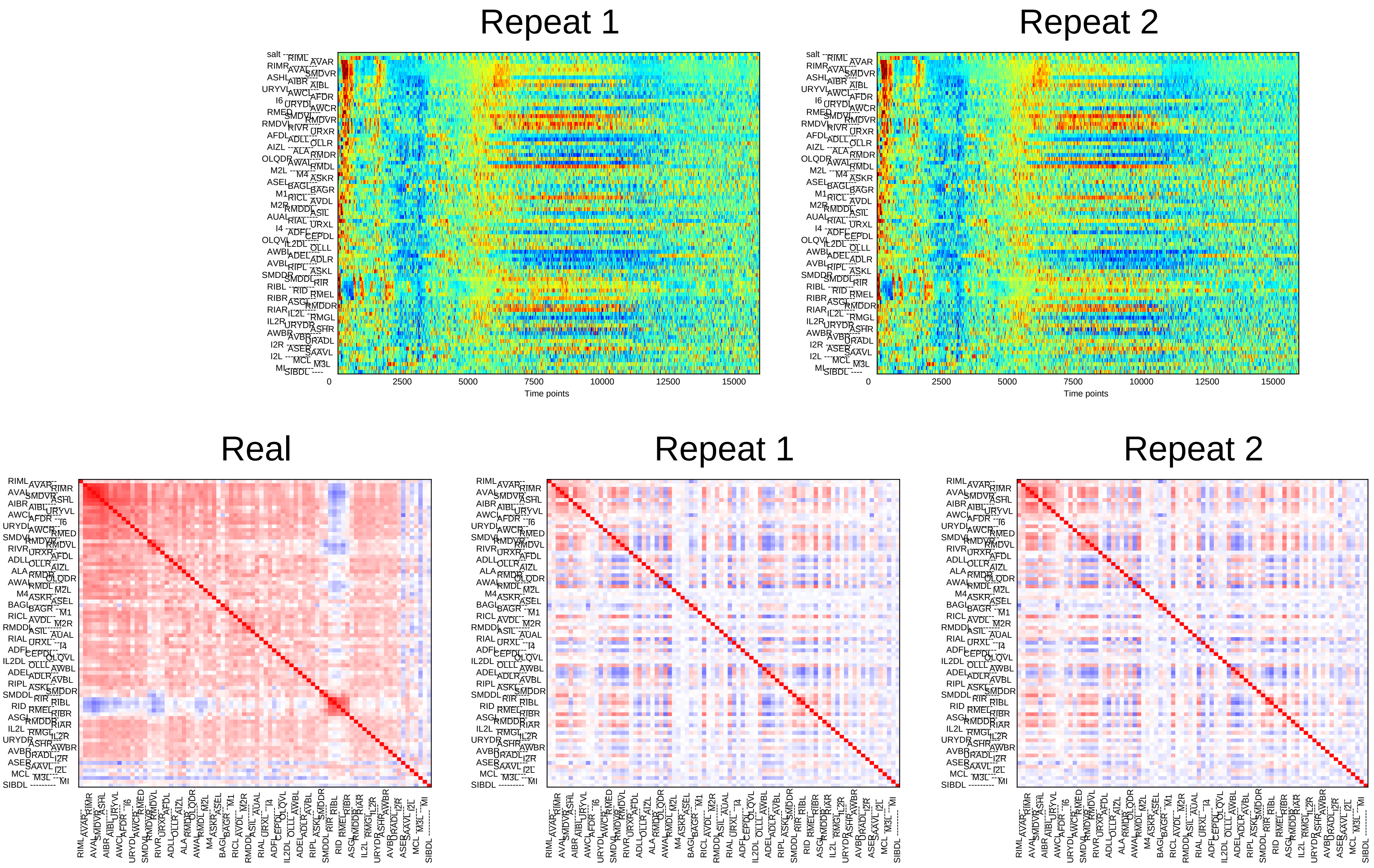
Repeat 1
Repeat 2
0
2500
5000
7500
10000
12500
15000
Time points
Real
Repeat 1
Repeat 2

### Sample21

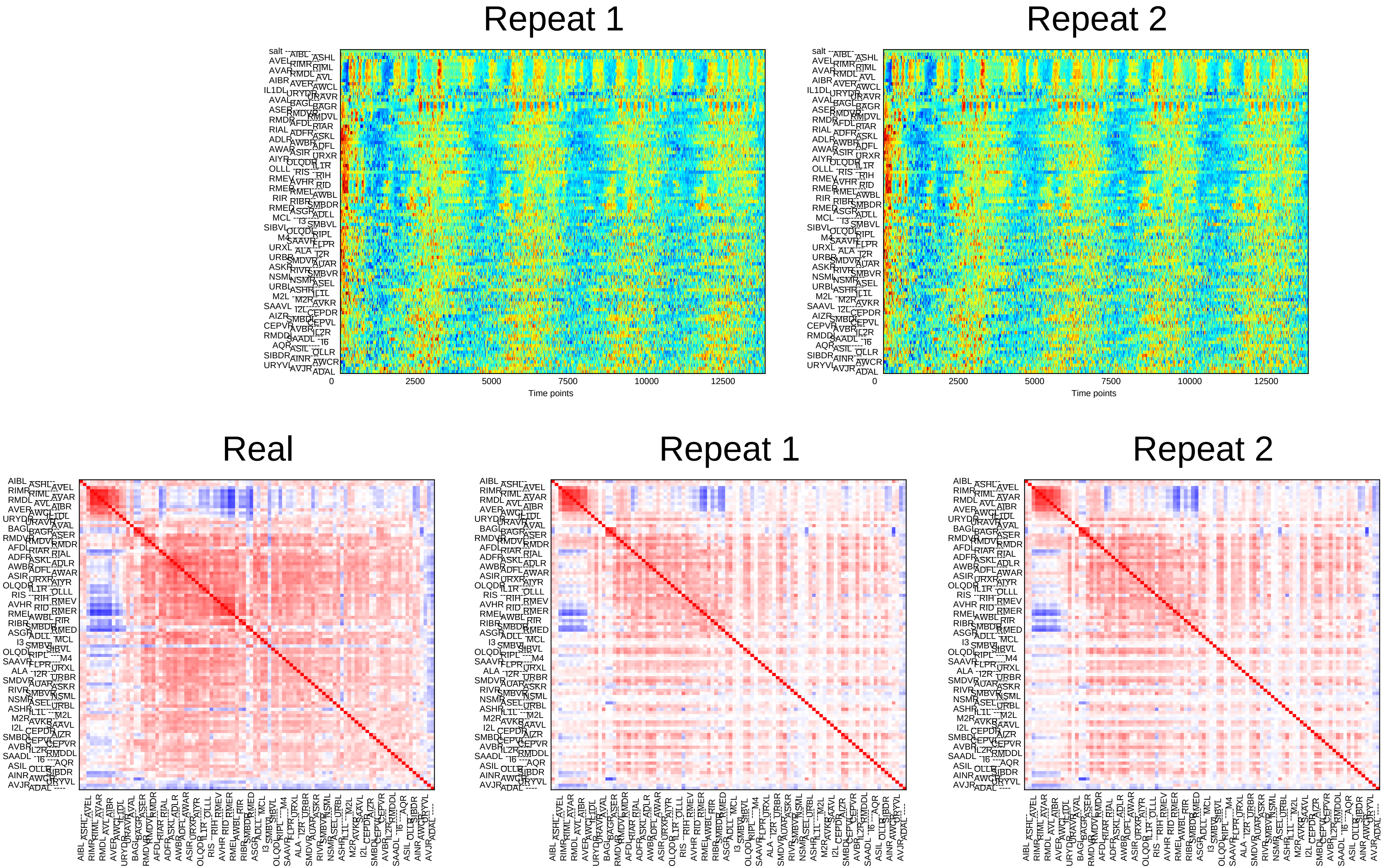
Repeat 1
Repeat 2
0
2500
5000
7500
10000
12500
Time points
Real
Repeat 1
Repeat 2

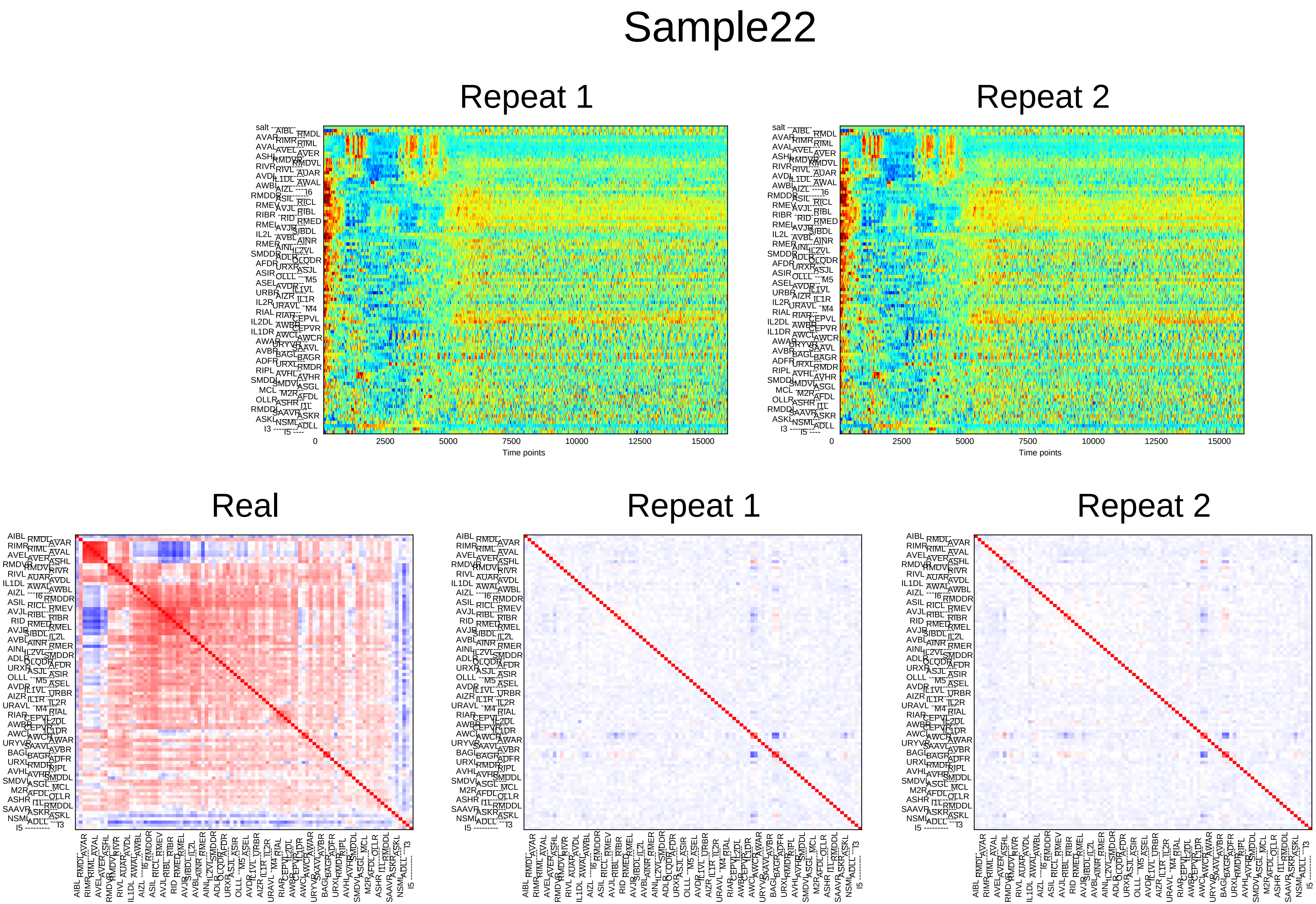
Sample22
Repeat 1
Repeat 2
0
2500
5000
7500
10000
12500
15000
Time points
Real
Repeat 1
Repeat 2

### Sample23

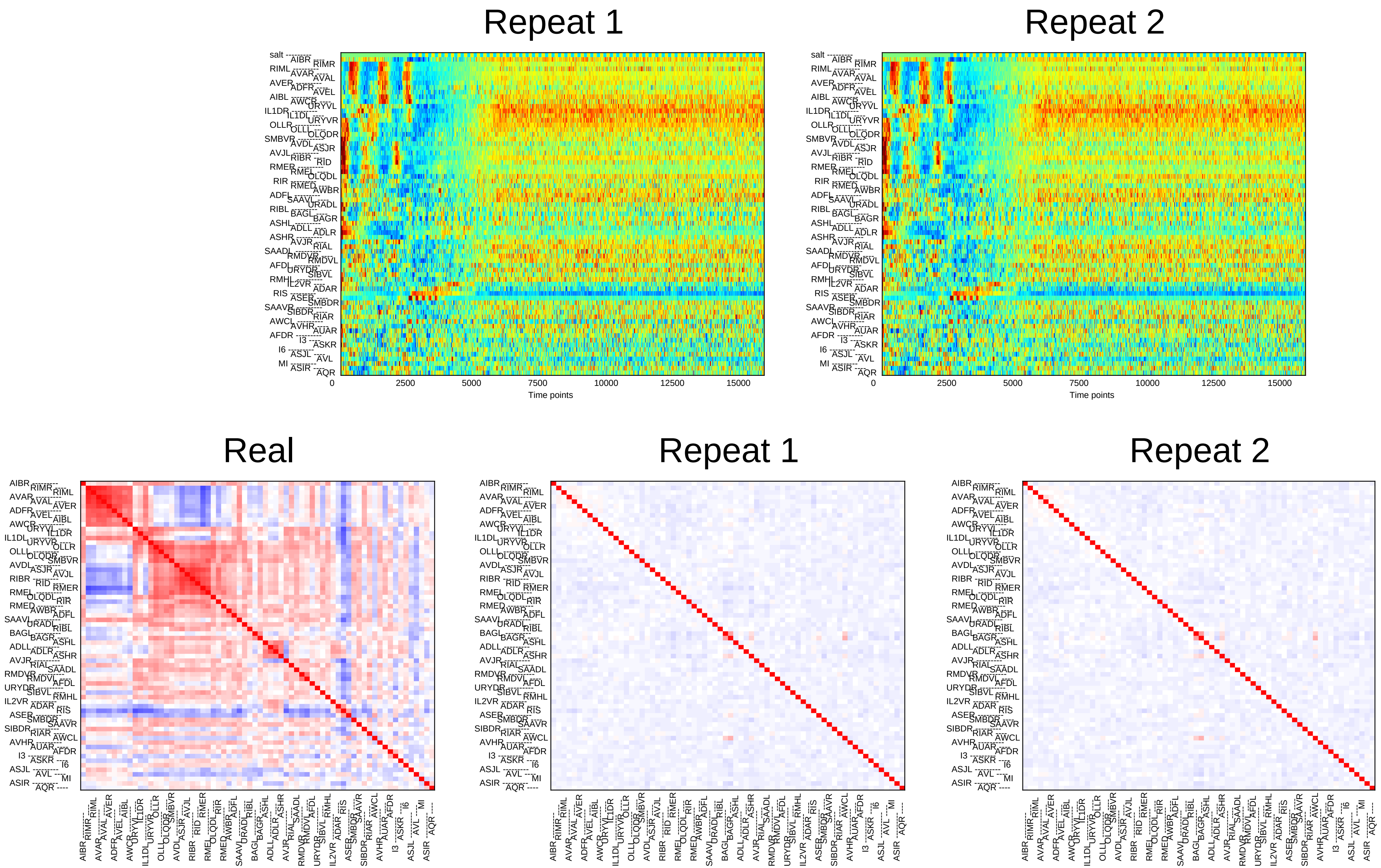

Repeat 1
Repeat 2
0
2500
5000
7500
10000
12500
15000
Time points
Real
Repeat 1
Repeat 2

### Sample24

#### Repeat 1

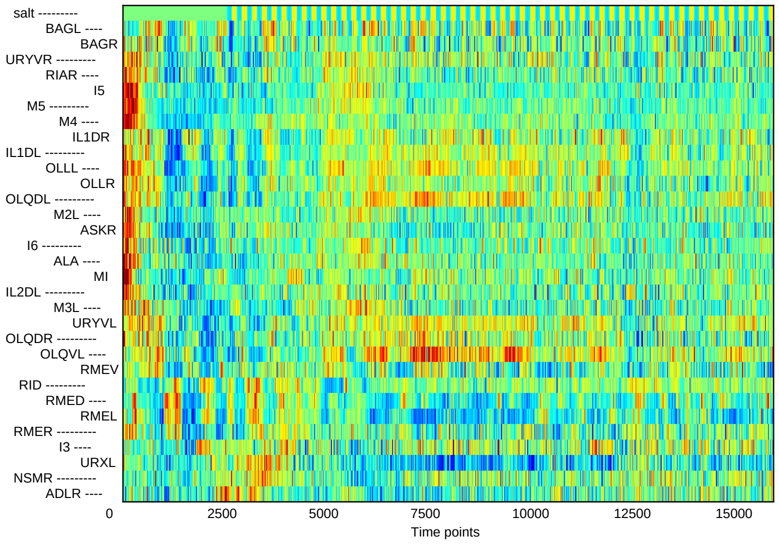


#### Repeat 2

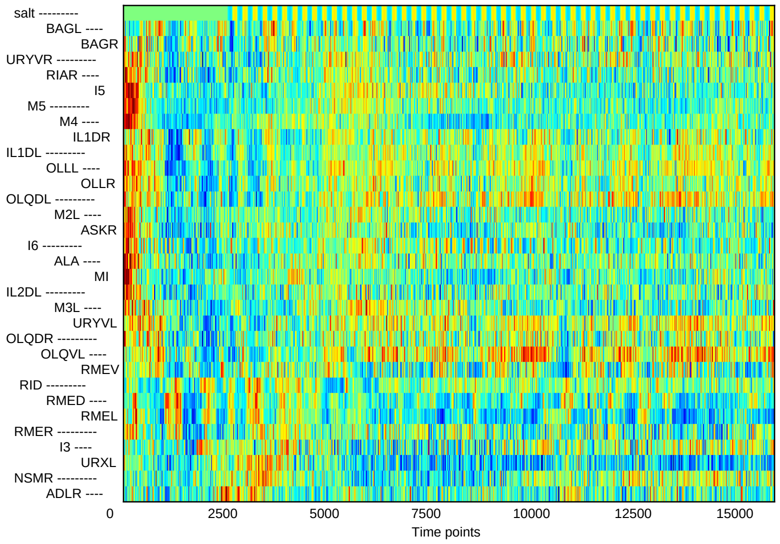


#### Real

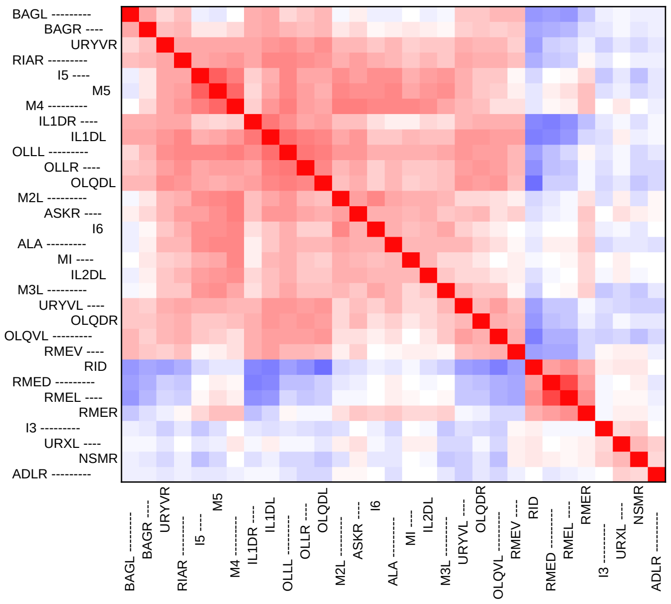


#### Repeat 1

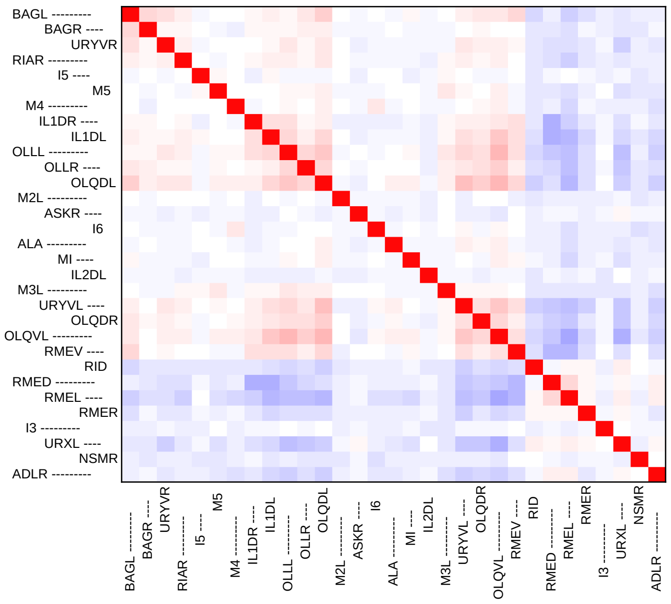


#### Repeat 2

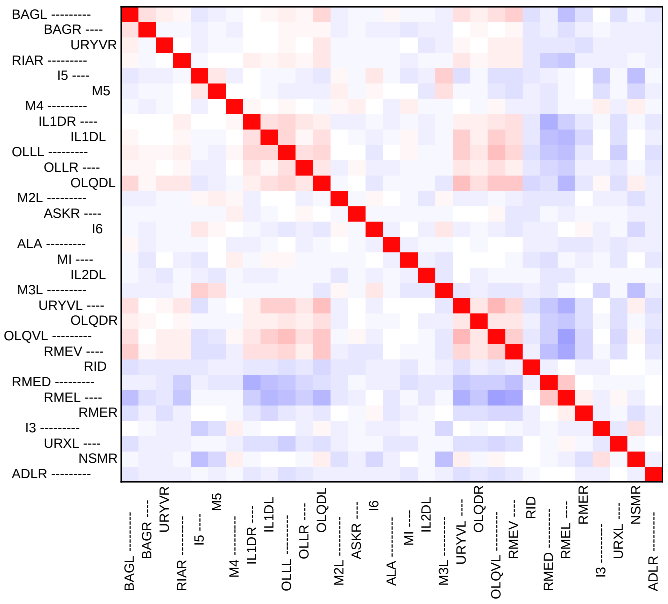
