## Supplementary File 4B for "Deducing ensemble dynamics and information flow from the whole-brain imaging data"

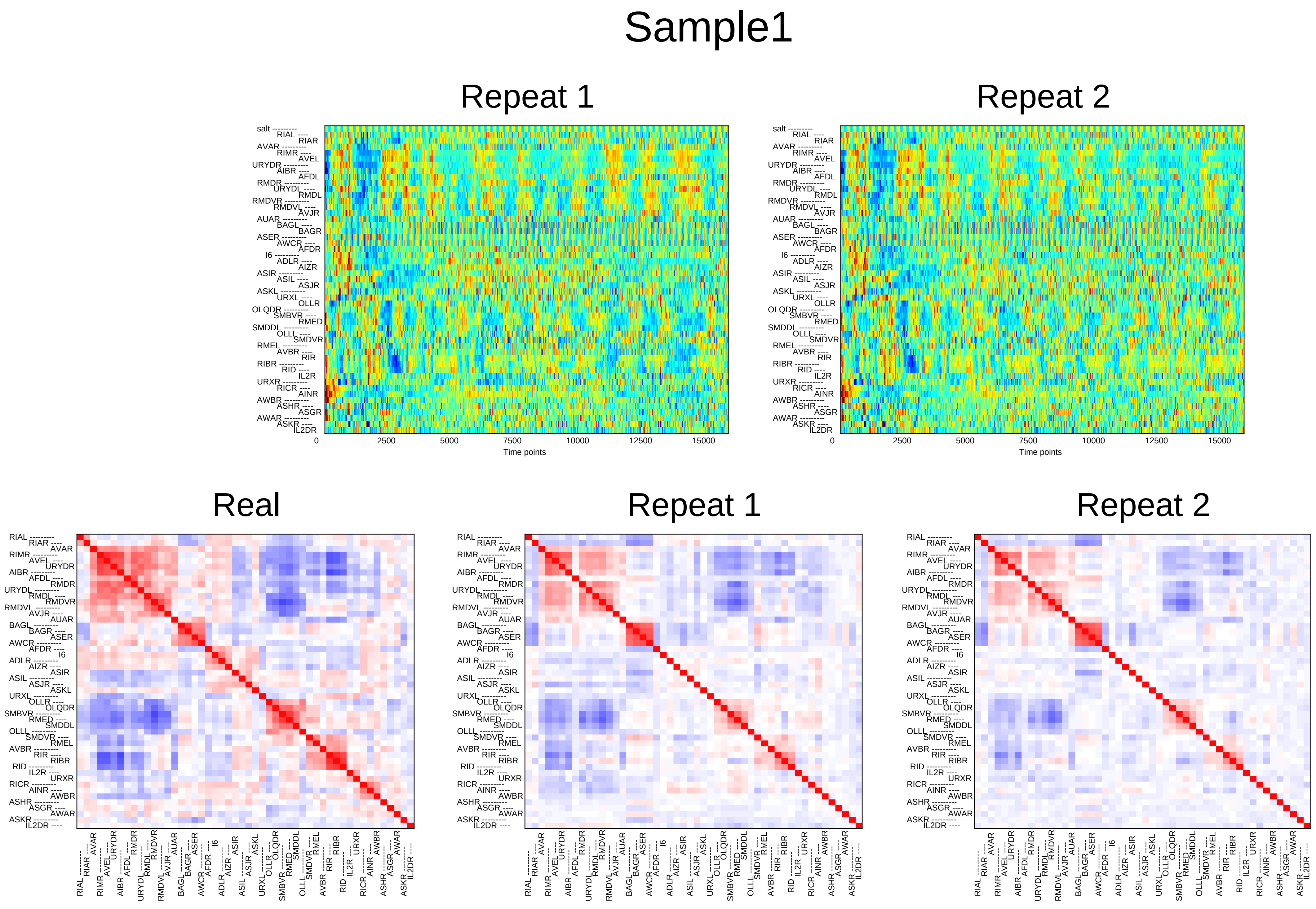
Sample1
Repeat 1
Repeat 2
Time points
0
2500
5000
7500
10000
12500
15000
Real
Repeat 1
Repeat 2

### Sample2

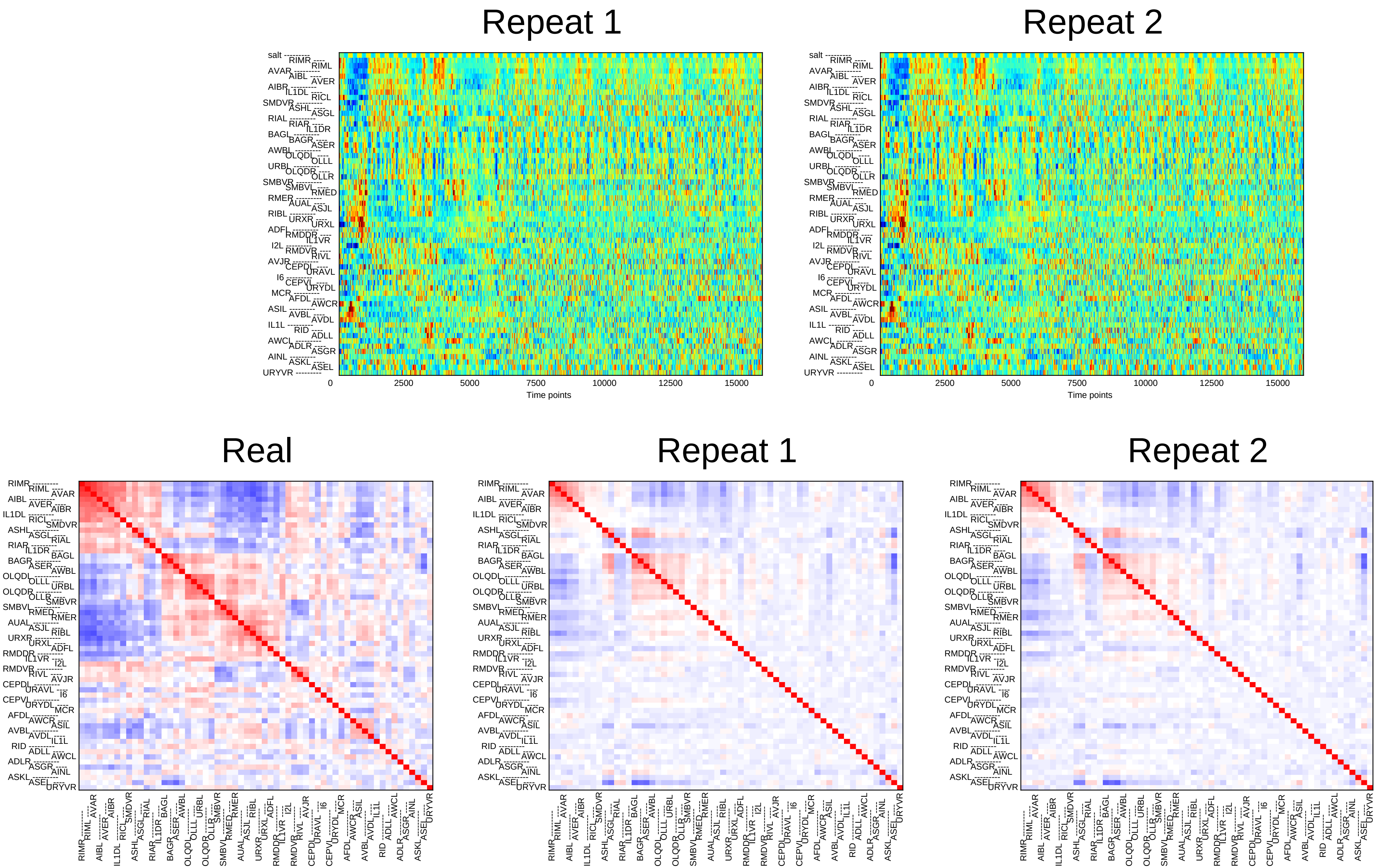
Repeat 1
Repeat 2
salt
RIMR RIML AVAR AIBL AVER AIBR IL1DL RICL SMDVR ASHL ASGL RIAL RIAR IL1DR BAGL BAGR ASER AWBL OLQDL OLLL URBL OLQDR OLLR SMBVR SMBVL RMED RMER AUAL ASJL RIBL URXR URXL ADFL RMDDR IL1VR I2L RMDVR RIVL AVJR CEPDL URAVL I6 CEPVL URYDL MCR AFDL AWCR ASIL AVBL AVDL IL1L RID ADLL AWCL ADLR ASGR AINL ASKL ASEL URYVR
0
2500
5000
7500
10000
12500
15000
Time points
Real
Repeat 1
Repeat 2

### Sample3

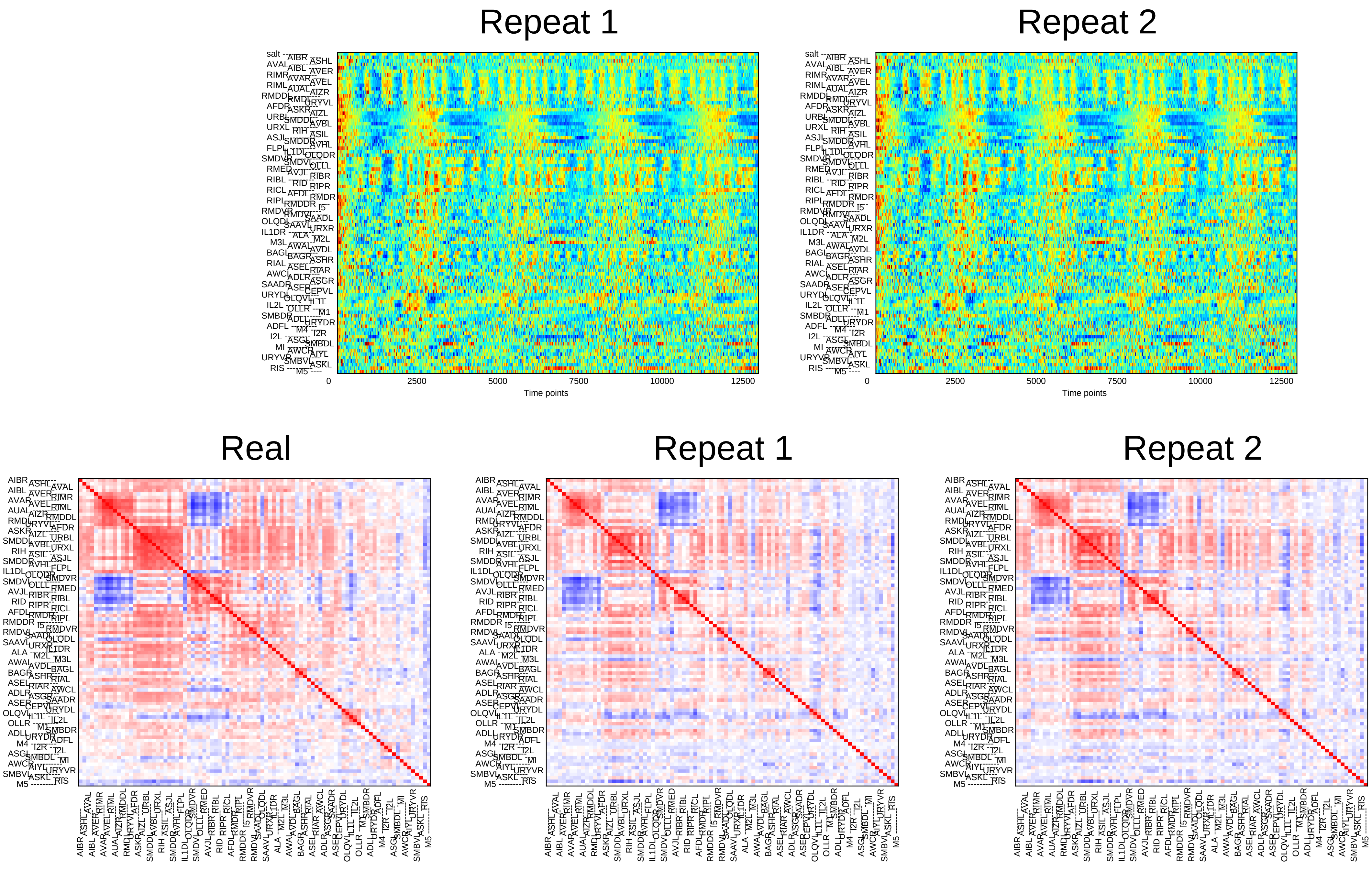

Repeat 1
Repeat 2
0
2500
5000
7500
10000
12500
Time points
Real
Repeat 1
Repeat 2

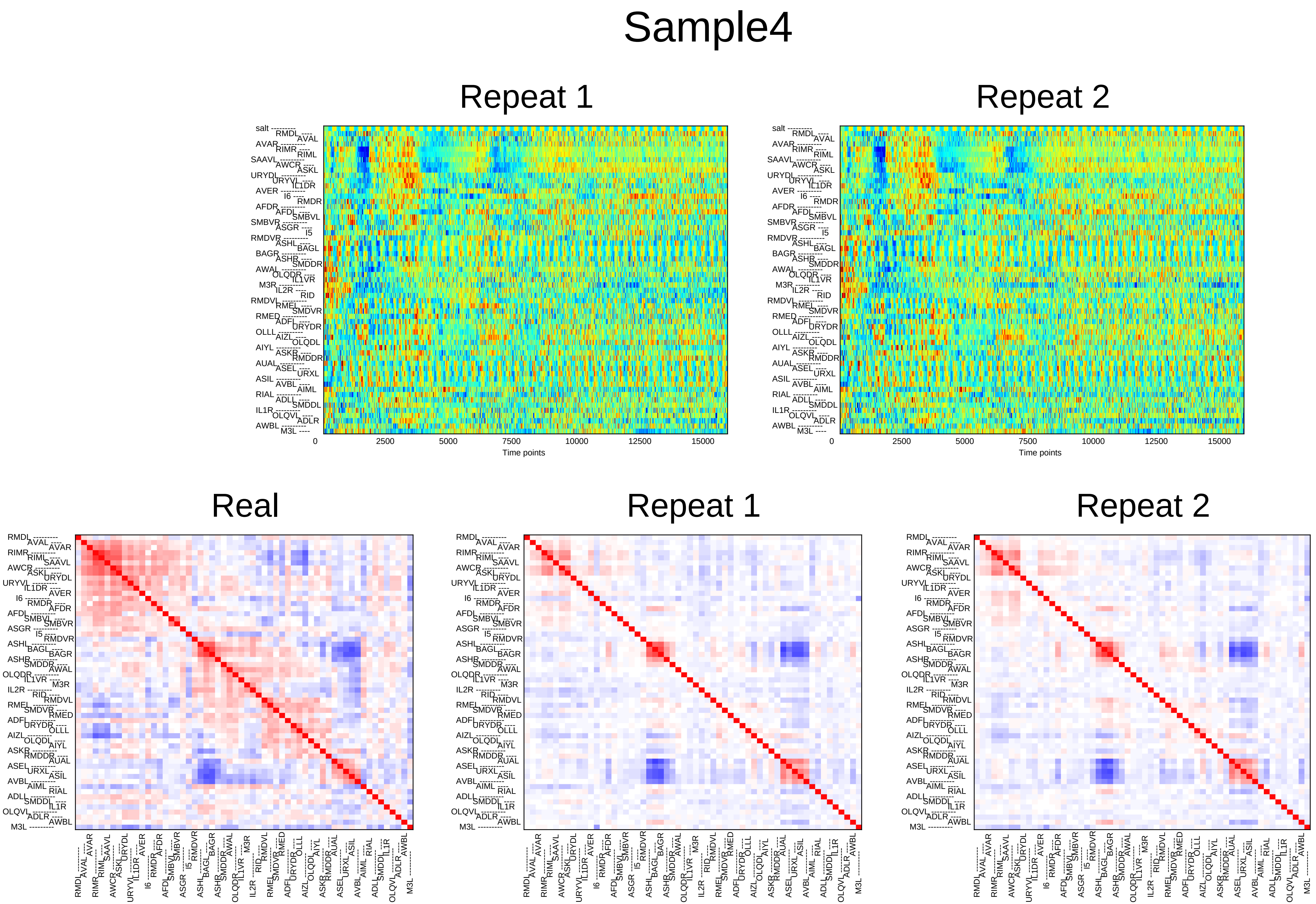
Sample4
Repeat 1
Repeat 2
0
2500
5000
7500
10000
12500
15000
Time points
Real
Repeat 1
Repeat 2

### Sample5

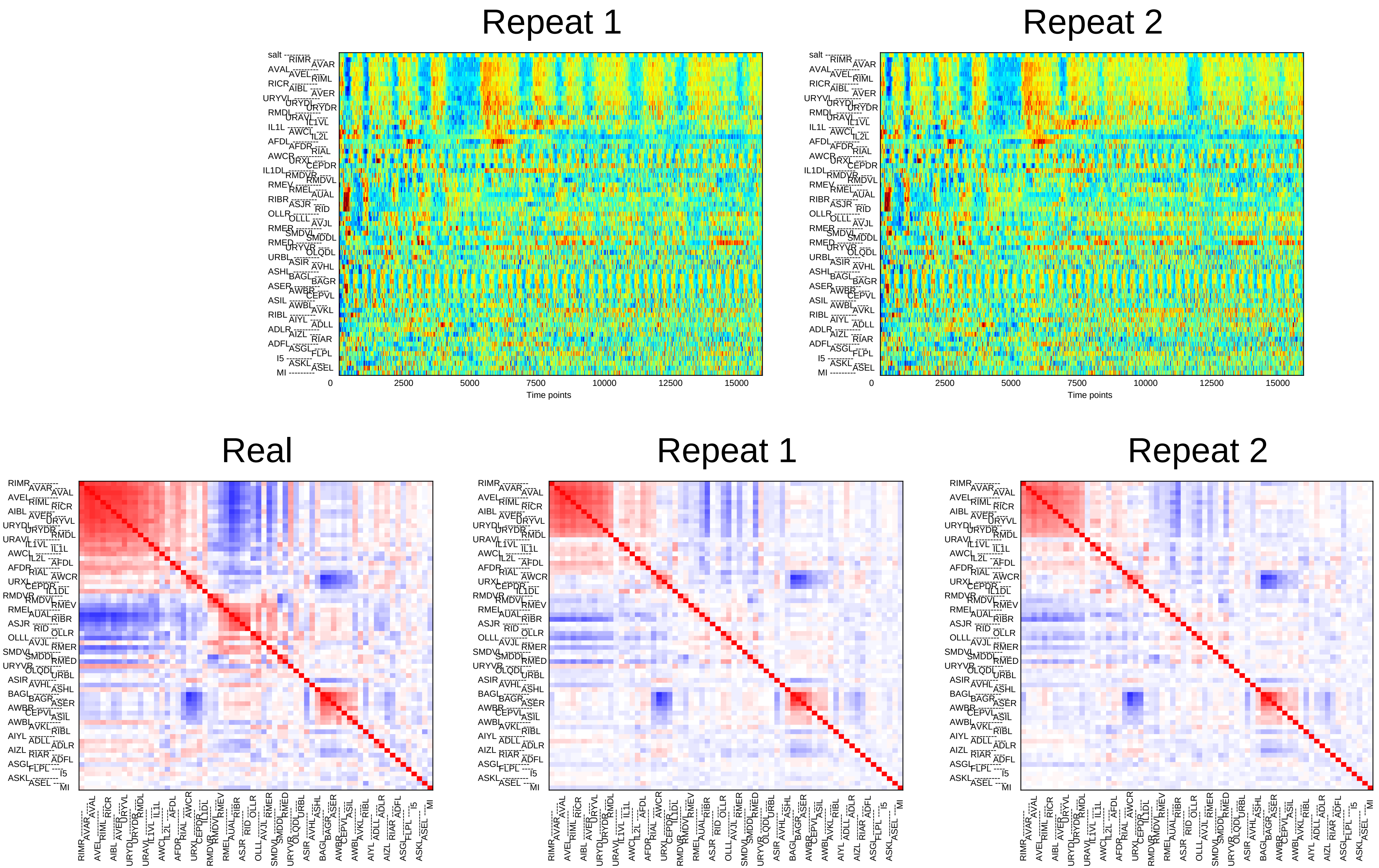
Repeat 1
Repeat 2
0
2500
5000
7500
10000
12500
15000
Time points
Real
Repeat 1
Repeat 2

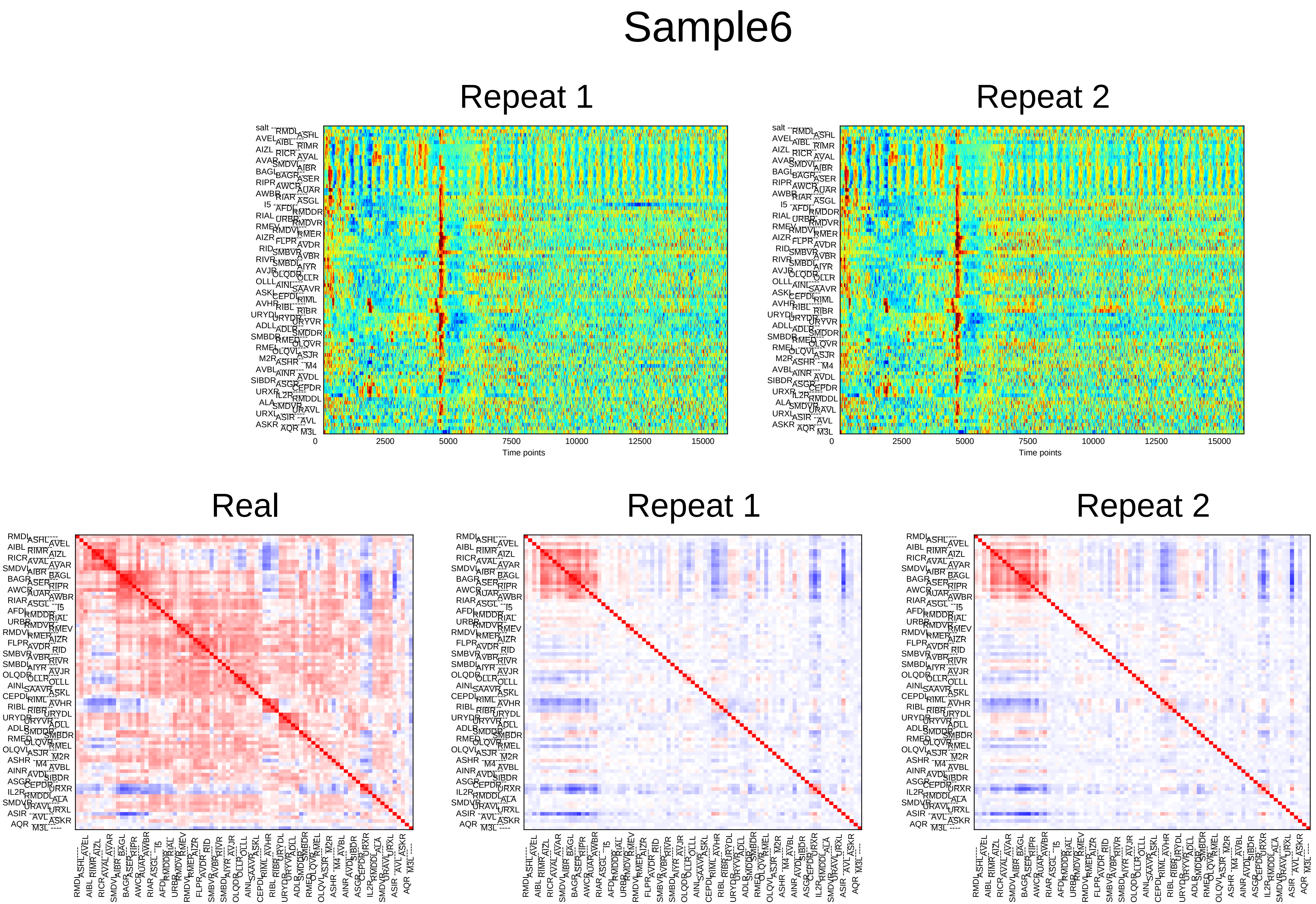
Sample6
Repeat 1
Repeat 2
0
2500
5000
7500
10000
12500
15000
Time points
Real
Repeat 1
Repeat 2

### Sample7

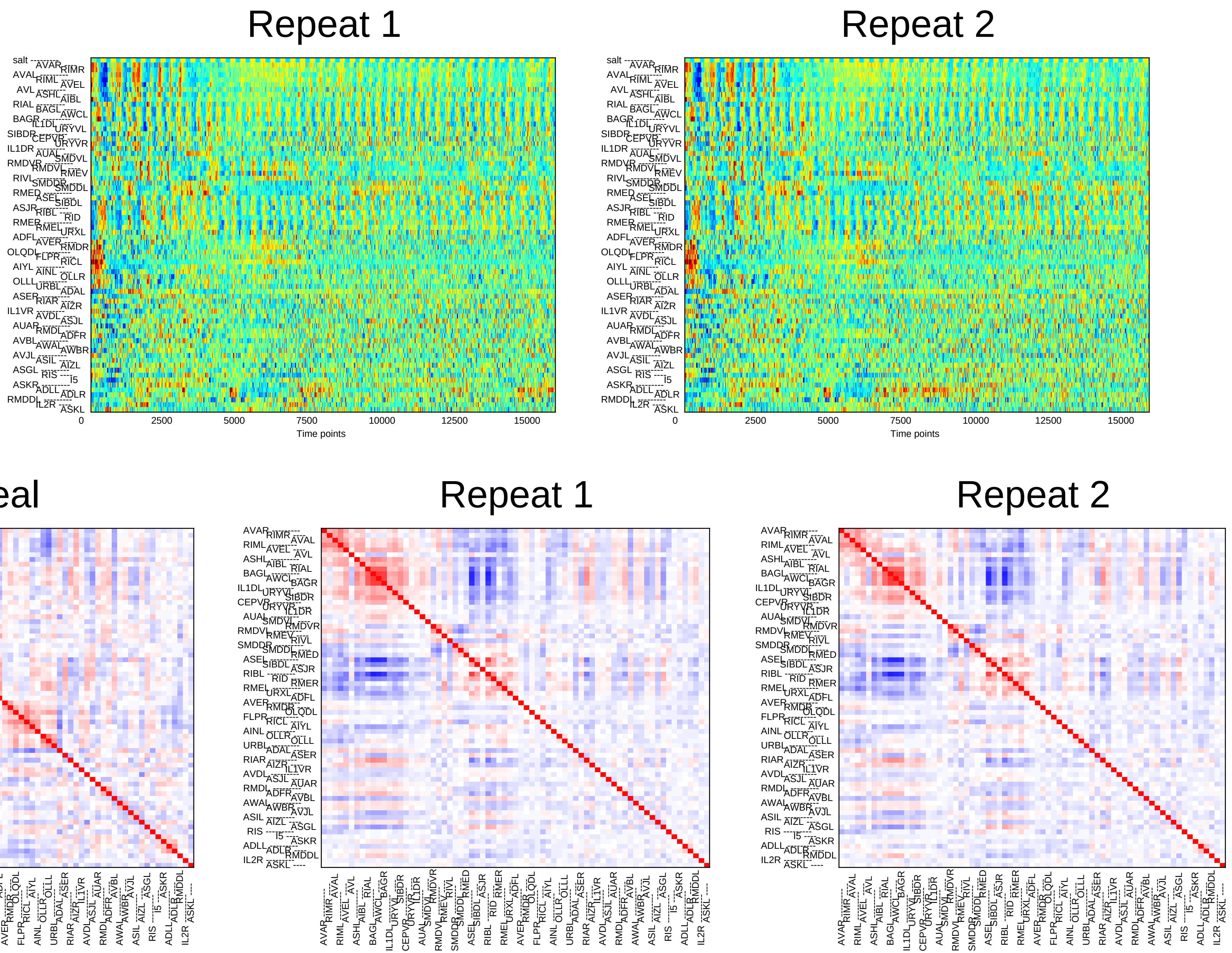
Repeat 1
Repeat 2
0
2500
5000
7500
10000
12500
15000
Time points
Real
Repeat 1
Repeat 2

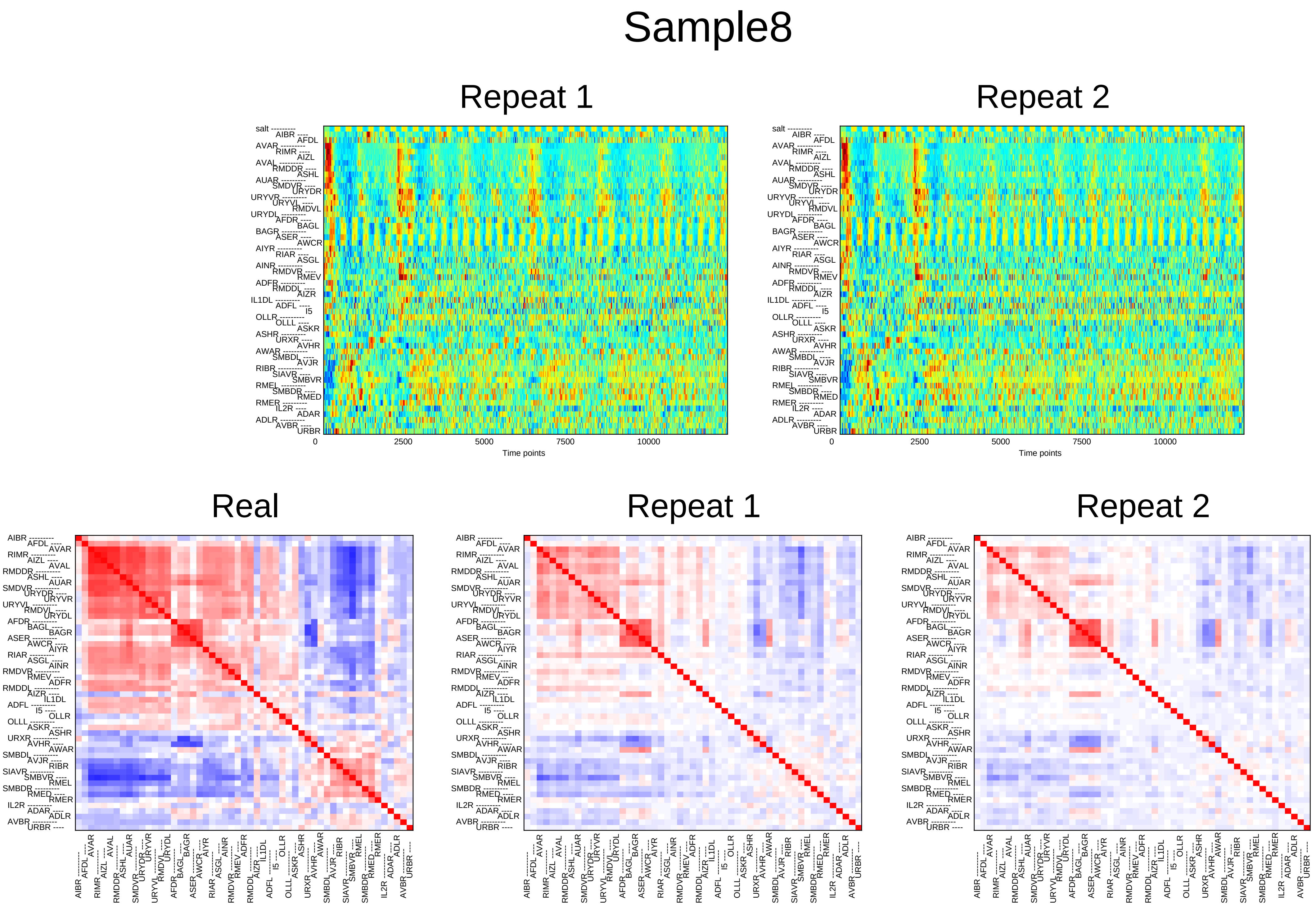

Sample8
Repeat 1
Repeat 2
salt
AIBR AFDL AVAR RIMR AIZL AVAL RMDDR ASHL AUAR SMDVR URYDR URYVR URYVL RMDVL URYDL AFDR BAGL BAGR ASER AWCR AIYR RIAR ASGL AINR RMDVR RMEV ADFR RMDDL AIZR IL1DL ADFL I5 OLLR OLLL ASKR ASHR URXR AVHR AWAR SMBDL AVJR RIBR SIAVR SMBVR RMEL SMBDR RMED RMER IL2R ADAR ADLR AVBR URBR
0
2500
5000
7500
10000
Time points
Real
Repeat 1
Repeat 2

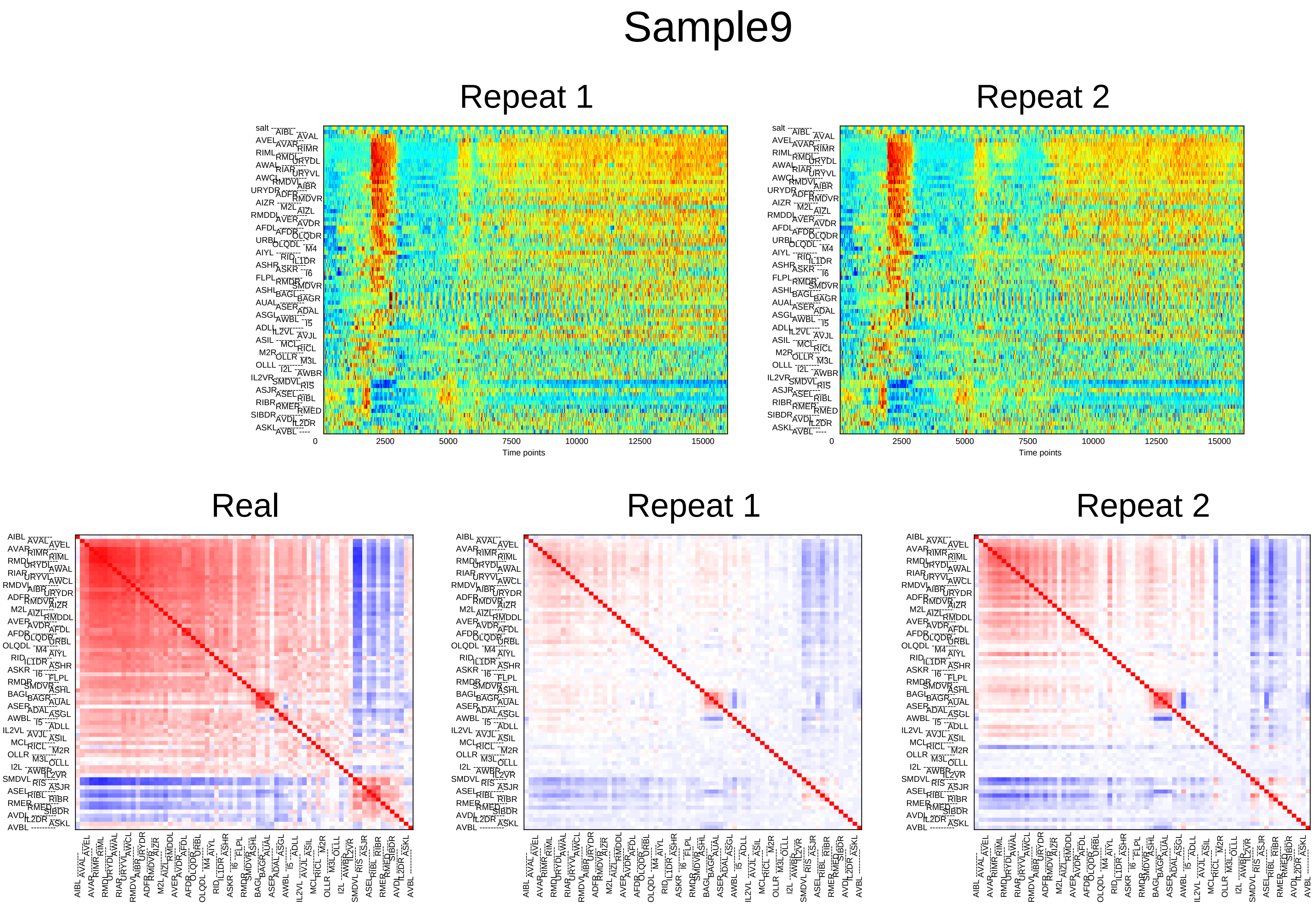
Sample9
Repeat 1
Repeat 2
0
2500
5000
7500
10000
12500
15000
Time points
Real
Repeat 1
Repeat 2

### Sample10

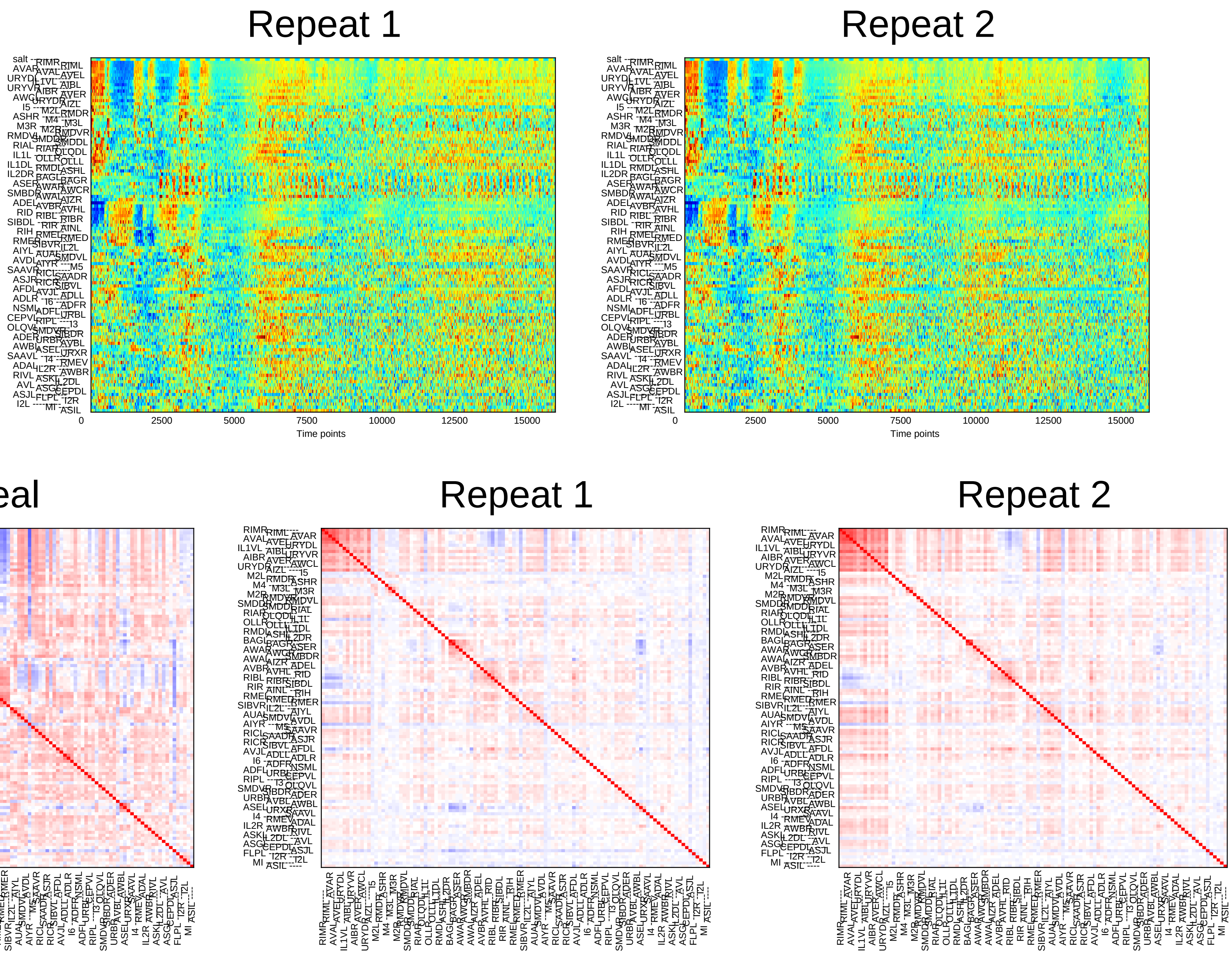
Repeat 1
Repeat 2
0
2500
5000
7500
10000
12500
15000
Time points
Real
Repeat 1
Repeat 2

### Sample11

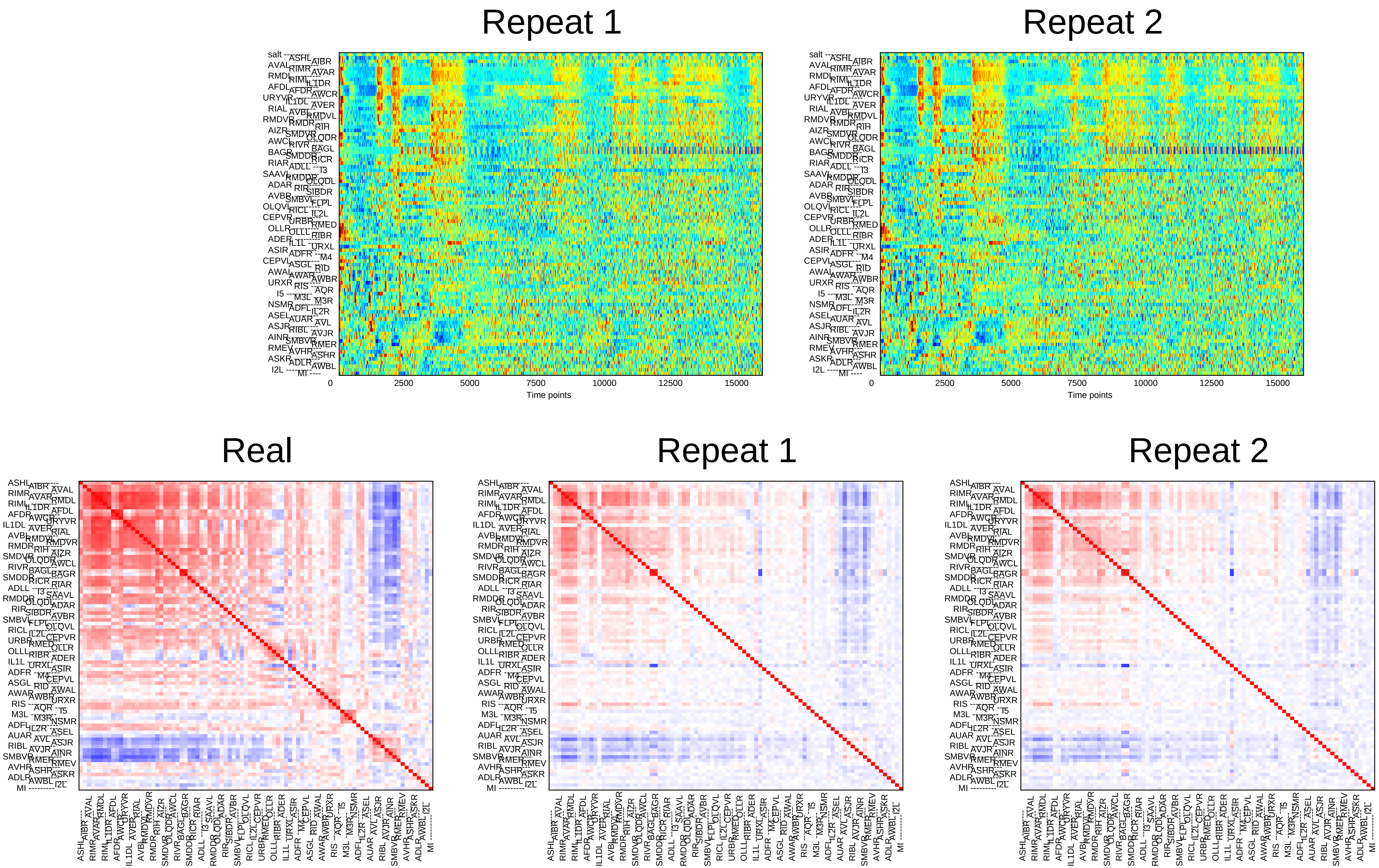

Repeat 1
Repeat 2
0
2500
5000
7500
10000
12500
15000
Time points
Real
Repeat 1
Repeat 2

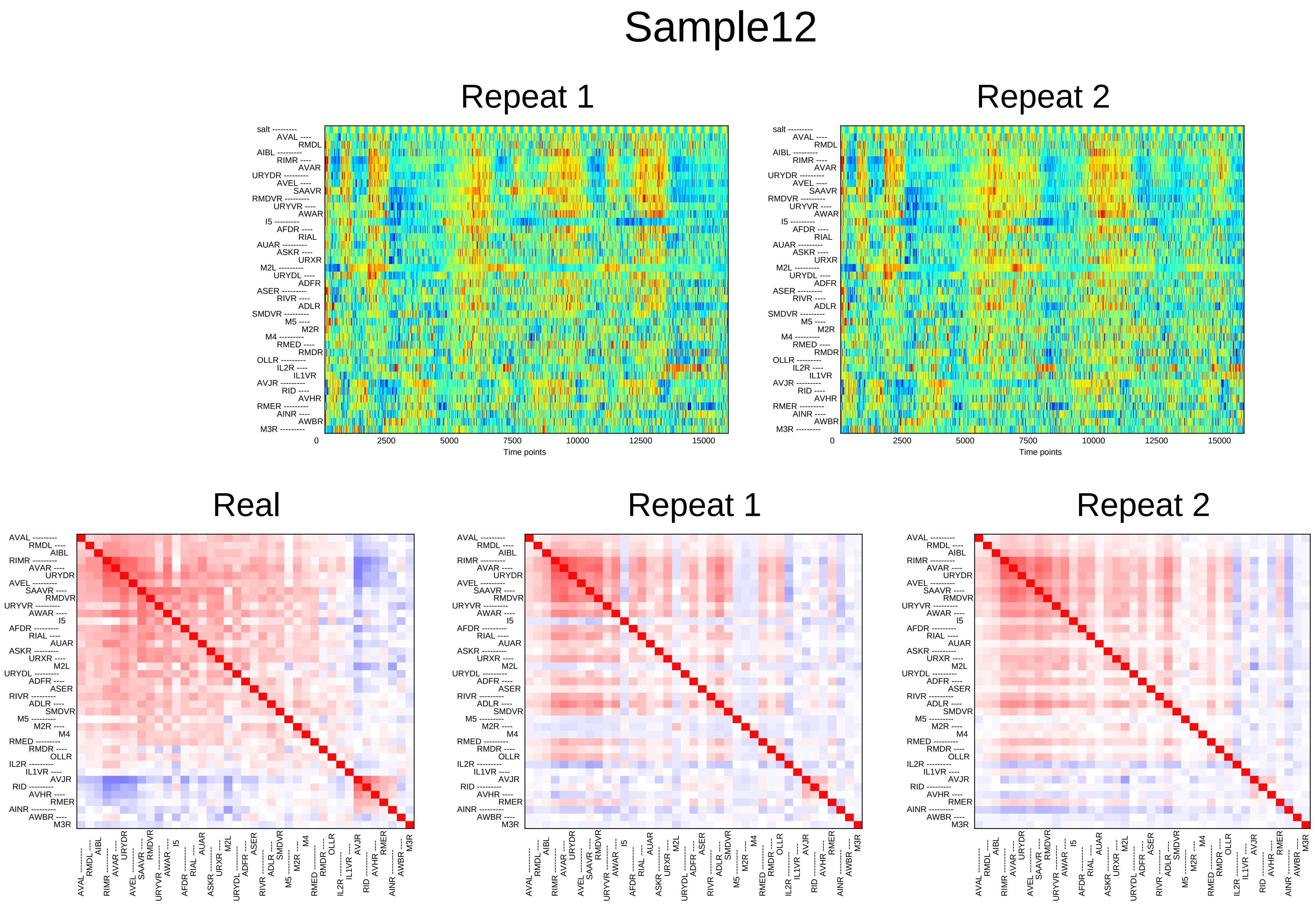
Sample12
Repeat 1
Repeat 2
salt
AVAL
RMDL
AIBL
RIMR
AVAR
URYDR
AVEL
SAAVR
RMDVR
URYVR
AWAR
I5
AFDR
RIAL
AUAR
ASKR
URXR
M2L
URYDL
ADFR
ASER
RIVR
ADLR
SMDVR
M5
M2R
M4
RMED
RMDR
OLLR
IL2R
IL1VR
AVJR
RID
AVHR
RMER
AINR
AWBR
M3R
0
2500
5000
7500
10000
12500
15000
Time points
Real
Repeat 1
Repeat 2

### Sample13

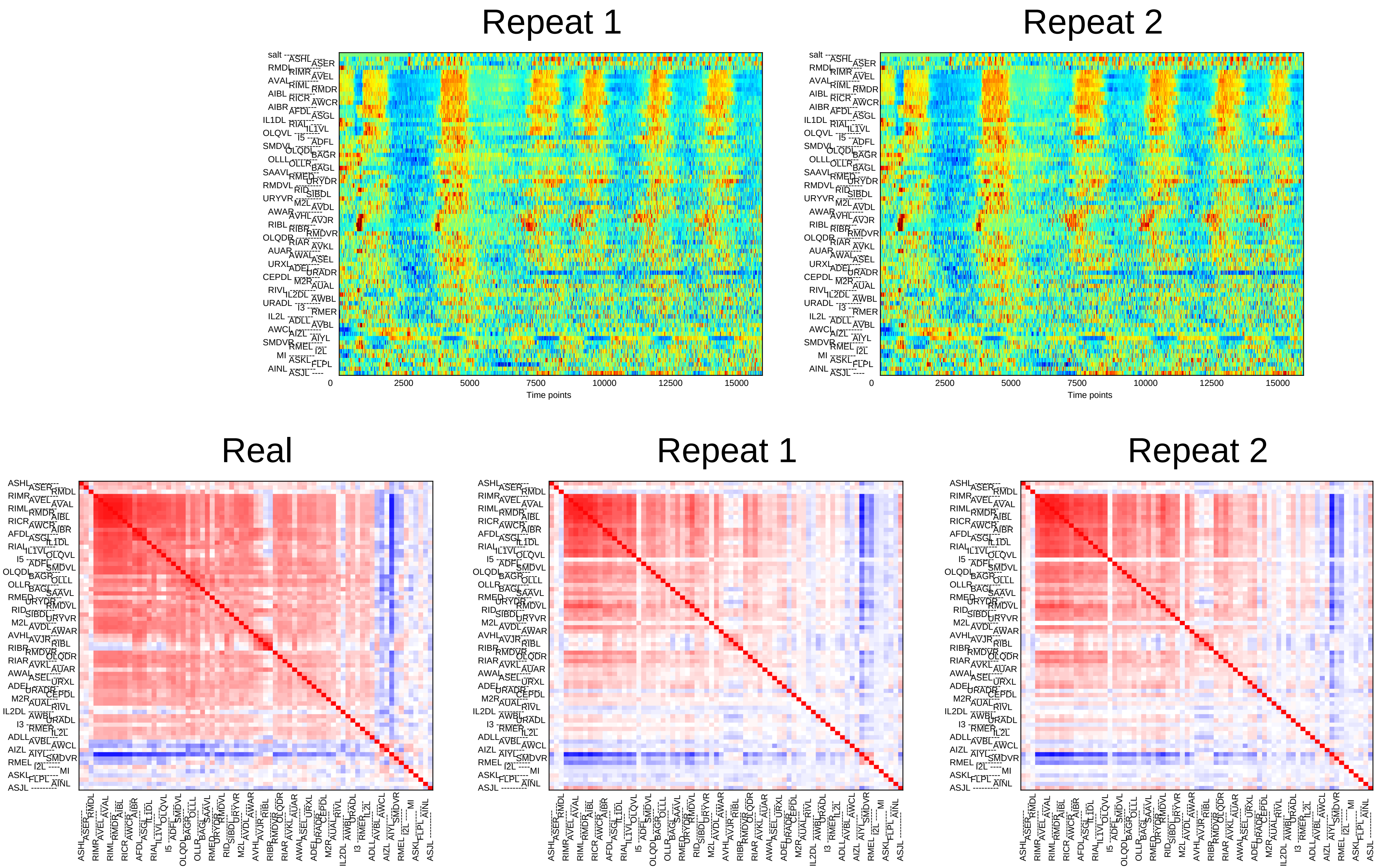
Repeat 1
Repeat 2
0
2500
5000
7500
10000
12500
15000
Time points
Real
Repeat 1
Repeat 2

### Sample14

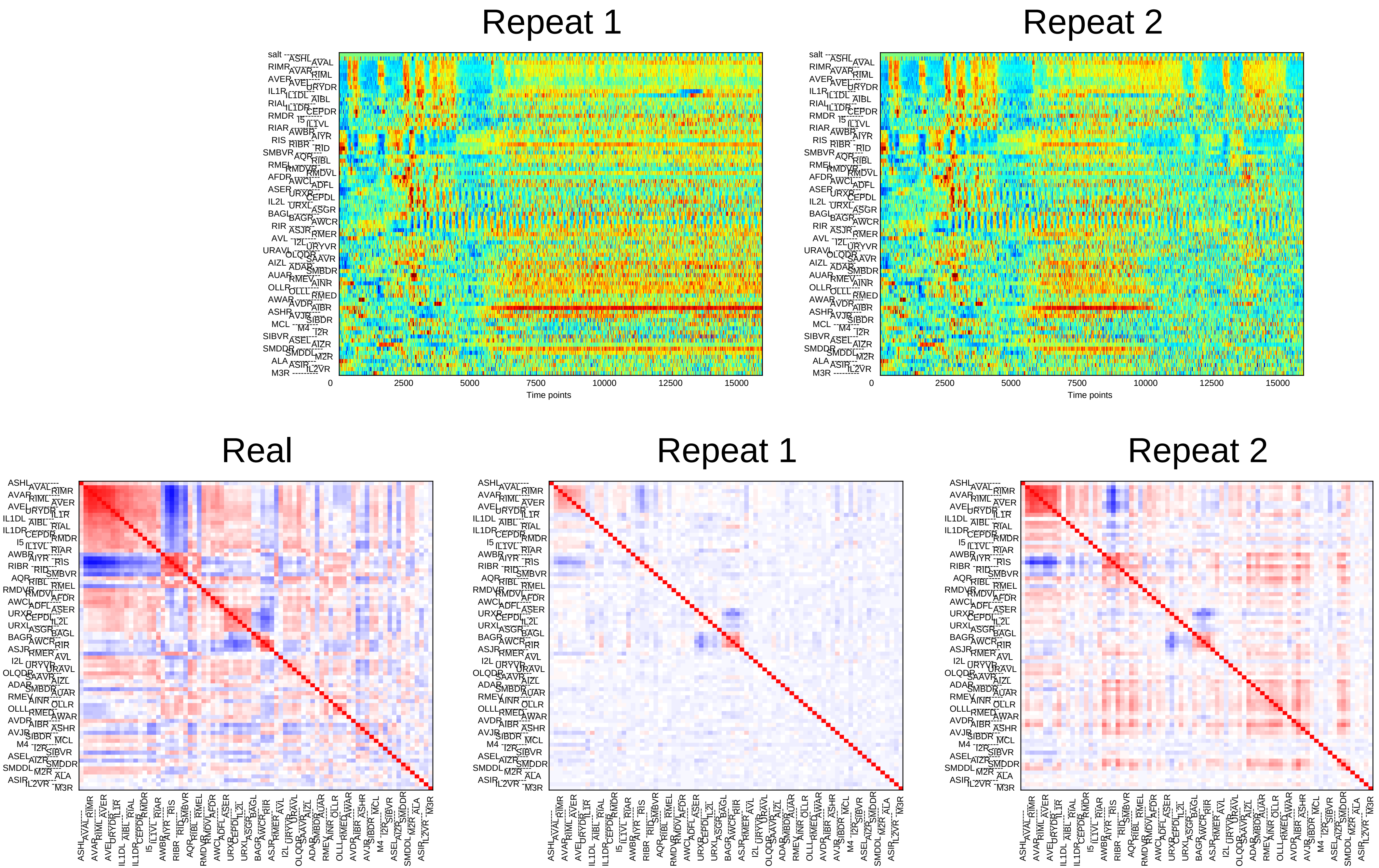
Repeat 1
Repeat 2
Time points
Time points
Real
Repeat 1
Repeat 2

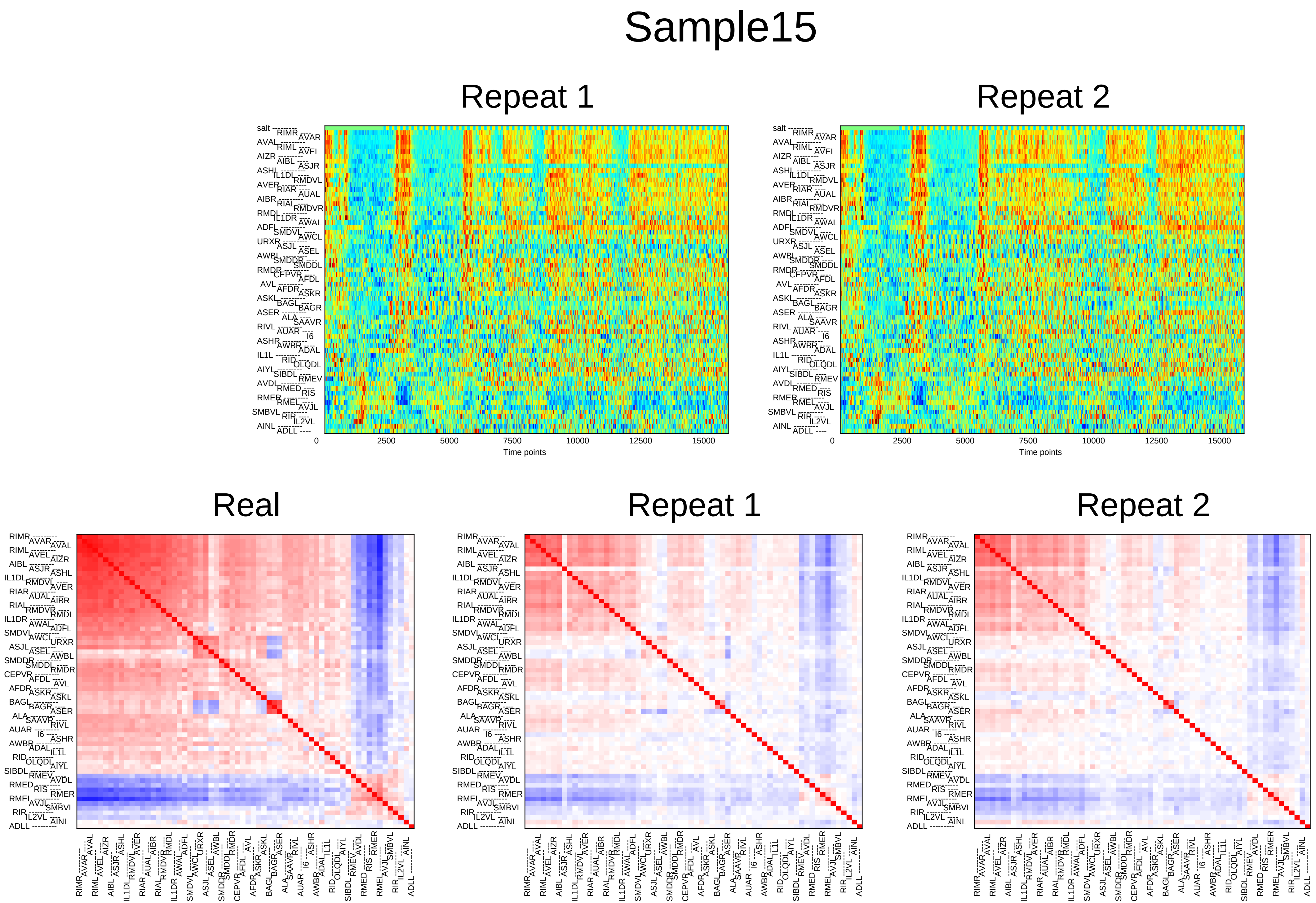

Sample15
Repeat 1
Repeat 2
0
2500
5000
7500
10000
12500
15000
Time points
Real
Repeat 1
Repeat 2

### Sample16

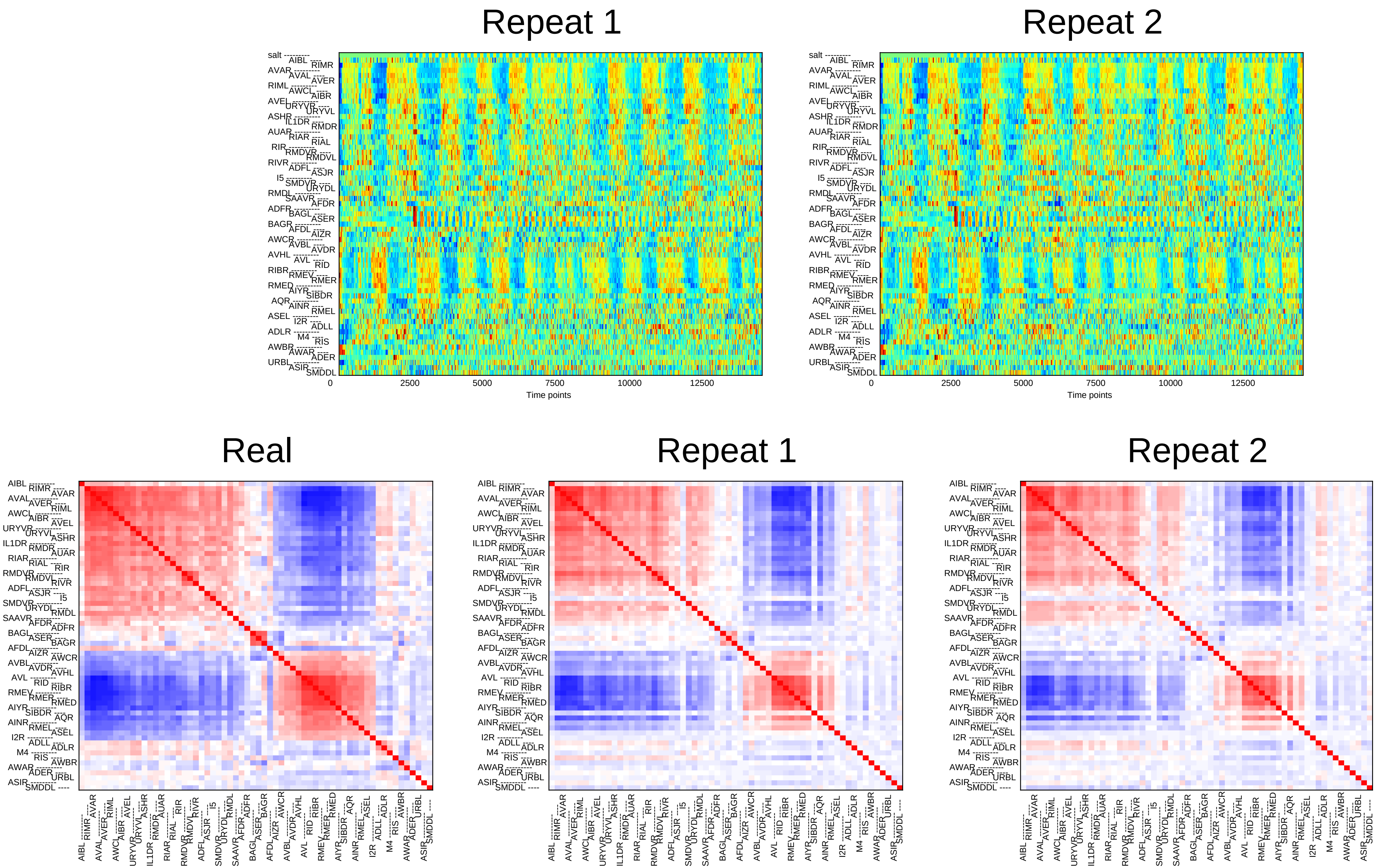

Repeat 1
Repeat 2
0
2500
5000
7500
10000
12500
Time points
Real
Repeat 1
Repeat 2

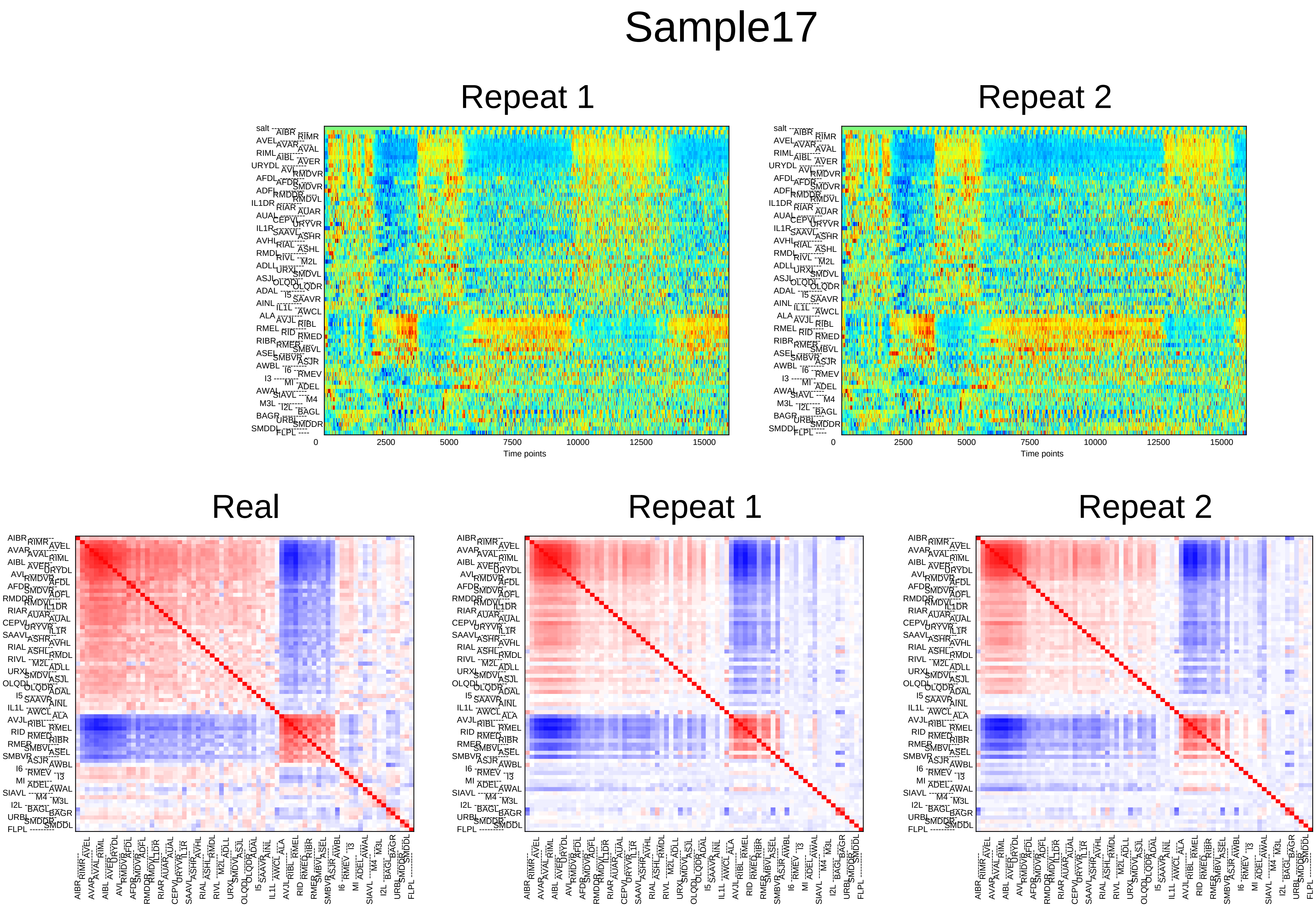

Sample17
Repeat 1
Repeat 2
0
2500
5000
7500
10000
12500
15000
Time points
Real
Repeat 1
Repeat 2

### Sample18

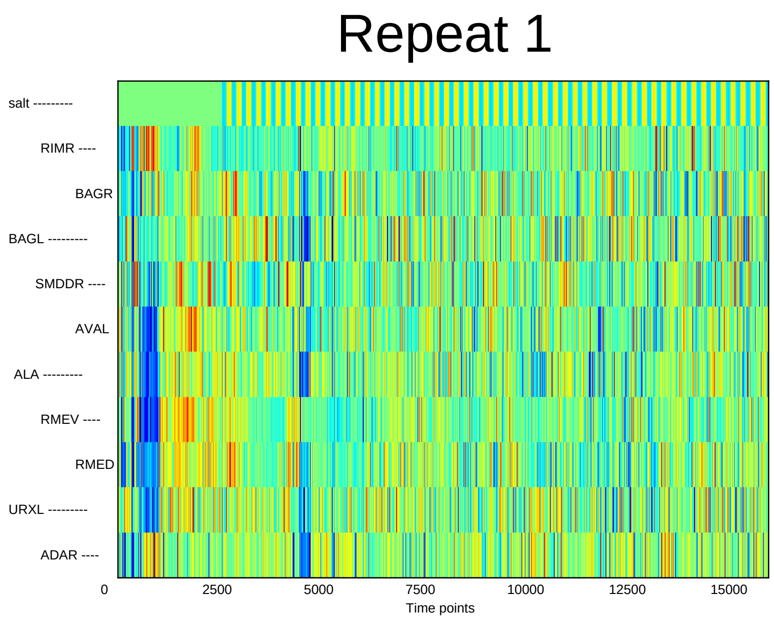
Repeat 1
salt ---------
RIMR ----
BAGR
BAGL ---------
SMDDR ----
AVAL
ALA ---------
RMEV ----
RMED
URXL ---------
ADAR ----
0
2500
5000
7500
10000
12500
15000
Time points

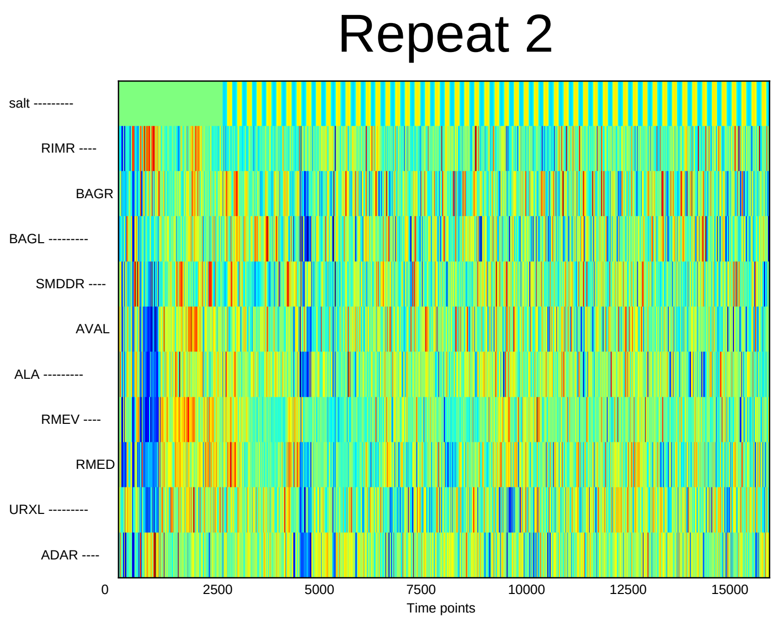
Repeat 2
salt ---------
RIMR ----
BAGR
BAGL ---------
SMDDR ----
AVAL
ALA ---------
RMEV ----
RMED
URXL ---------
ADAR ----
0
2500
5000
7500
10000
12500
15000
Time points

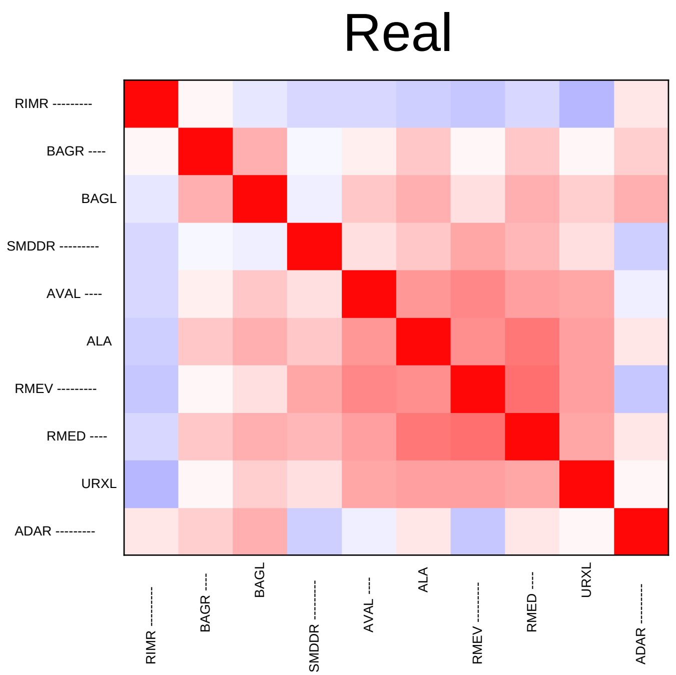
Real
RIMR ---------
BAGR ----
BAGL
SMDDR ---------
AVAL ----
ALA
RMEV ---------
RMED ----
URXL
ADAR ---------

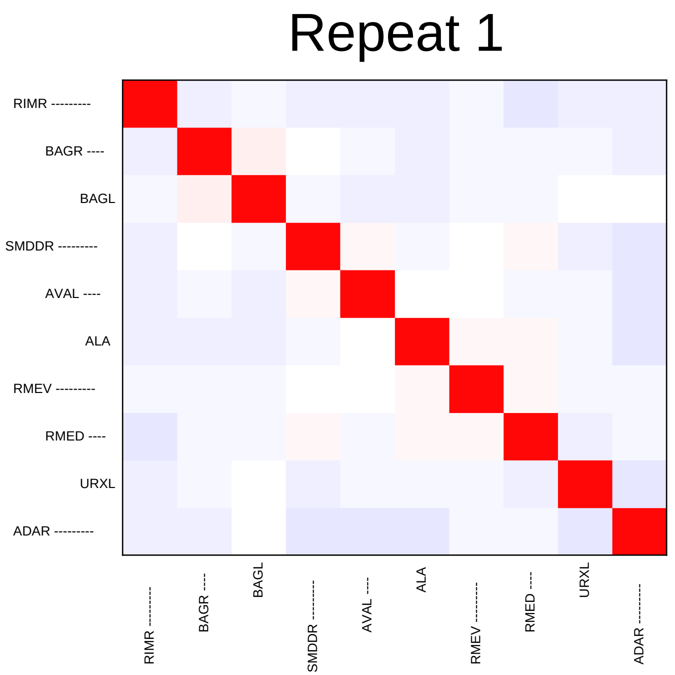
Repeat 1
RIMR ---------
BAGR ----
BAGL
SMDDR ---------
AVAL ----
ALA
RMEV ---------
RMED ----
URXL
ADAR ---------

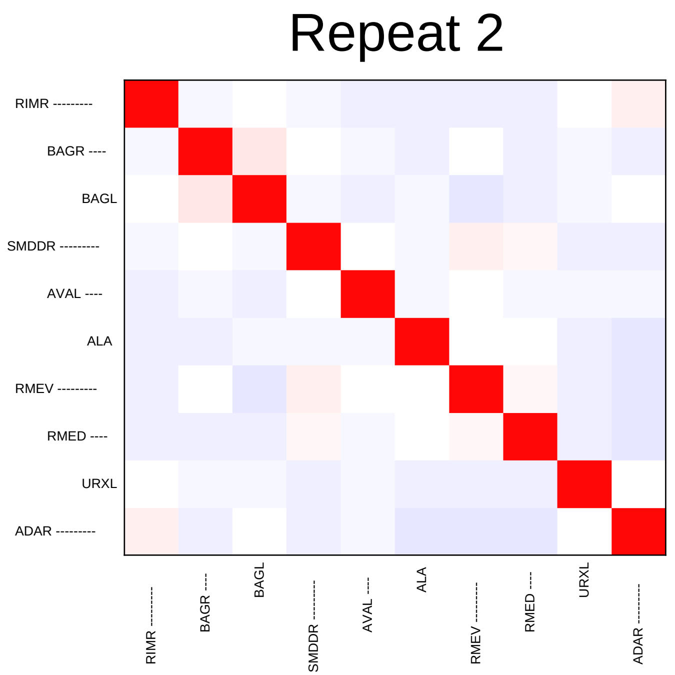
Repeat 2
RIMR ---------
BAGR ----
BAGL
SMDDR ---------
AVAL ----
ALA
RMEV ---------
RMED ----
URXL
ADAR ---------

### Sample19

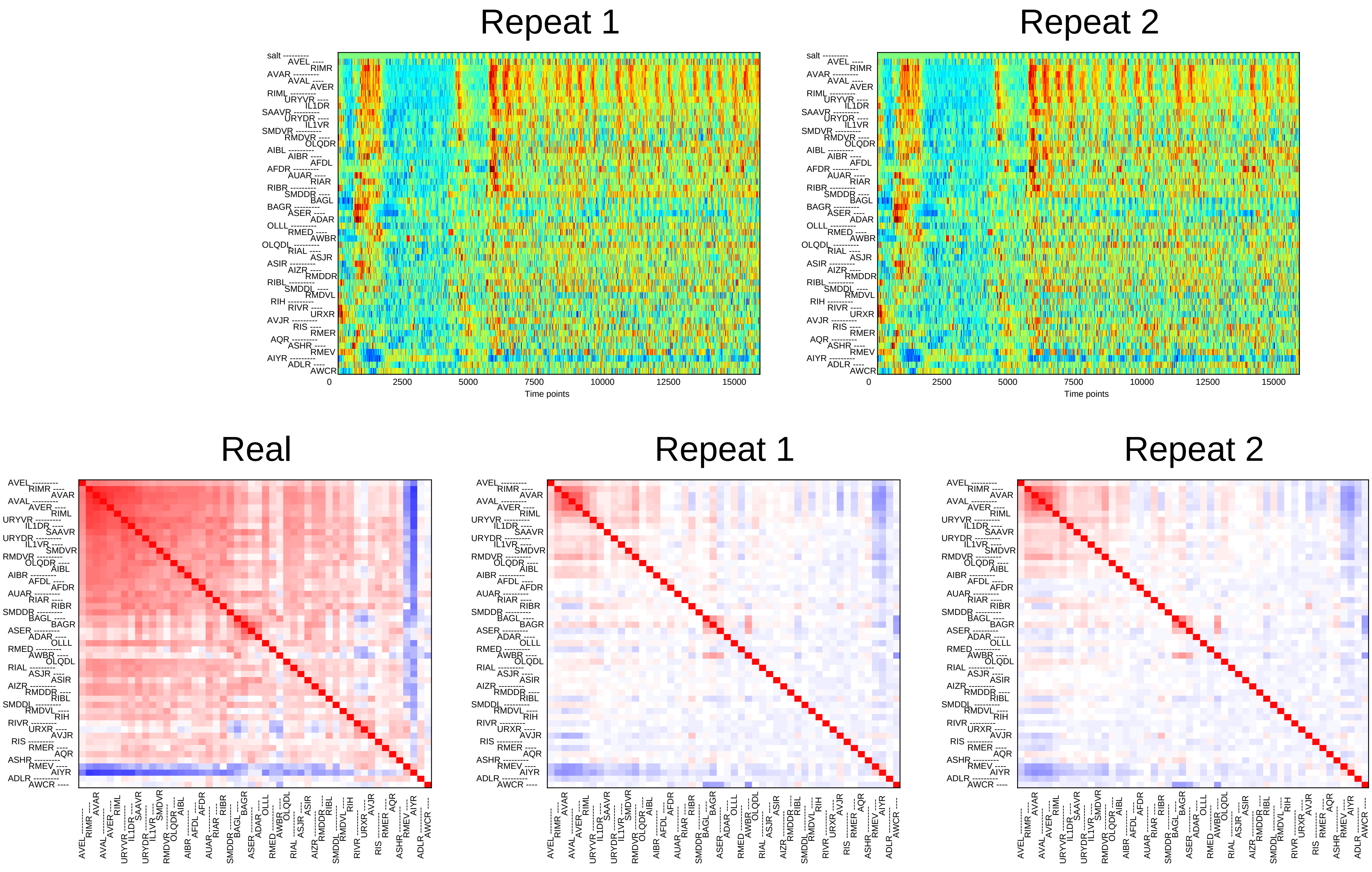
Repeat 1
Repeat 2
salt
AVEL
RIMR
AVAR
AVAL
AVER
RIML
URYVR
IL1DR
SAAVR
URYDR
IL1VR
SMDVR
RMDVR
OLQDR
AIBL
AIBR
AFDL
AFDR
AUAR
RIAR
RIBR
SMDDR
BAGL
BAGR
ASER
ADAR
OLLL
RMED
AWBR
OLQDL
RIAL
ASJR
ASIR
AIZR
RMDDR
RIBL
SMDDL
RMDVL
RIH
RIVR
URXR
AVJR
RIS
RMER
AQR
ASHR
RMEV
AIYR
ADLR
AWCR
0
2500
5000
7500
10000
12500
15000
Time points
Real
Repeat 1
Repeat 2

### Sample20

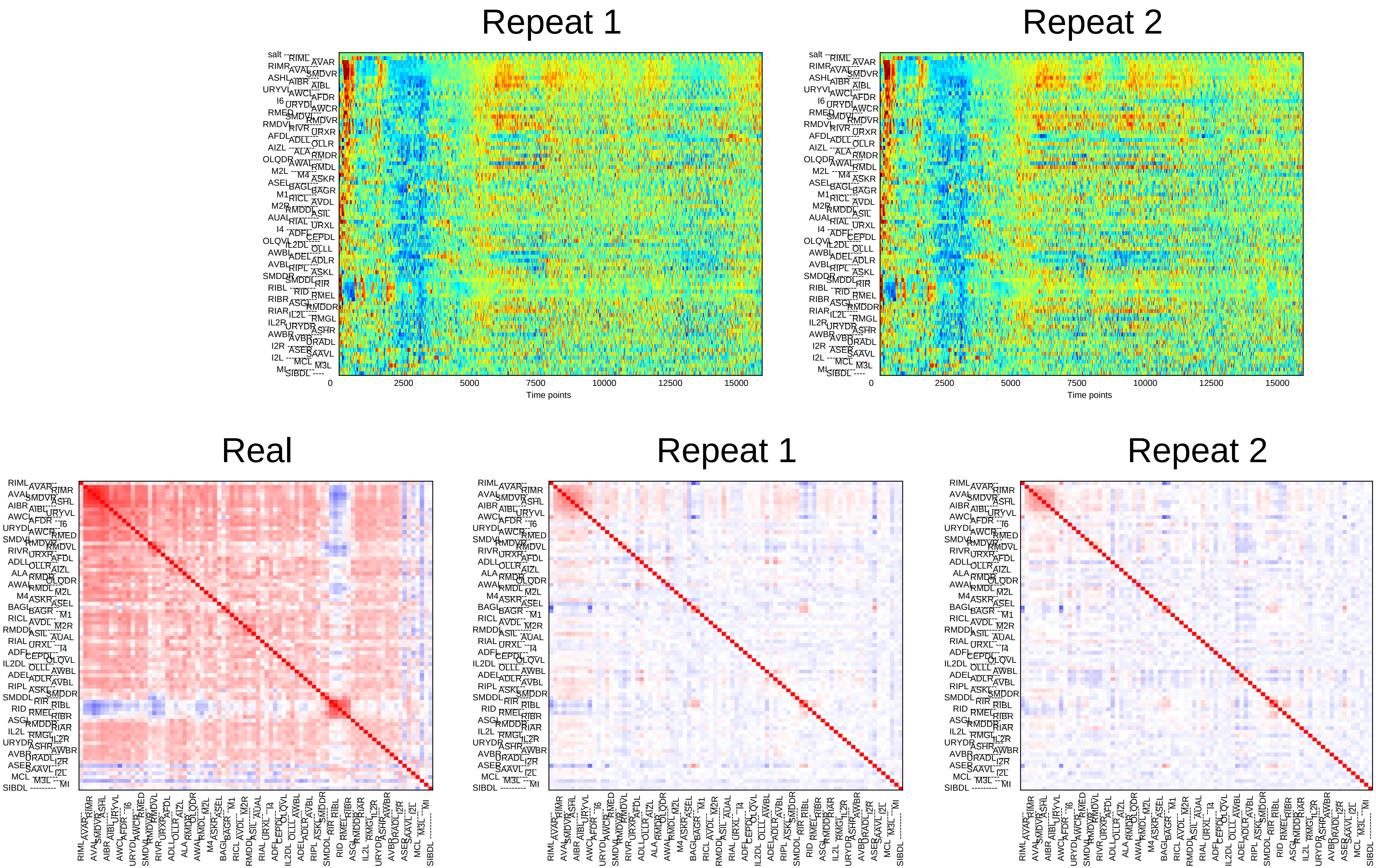
Repeat 1
Repeat 2
0
2500
5000
7500
10000
12500
15000
Time points
Real
Repeat 1
Repeat 2

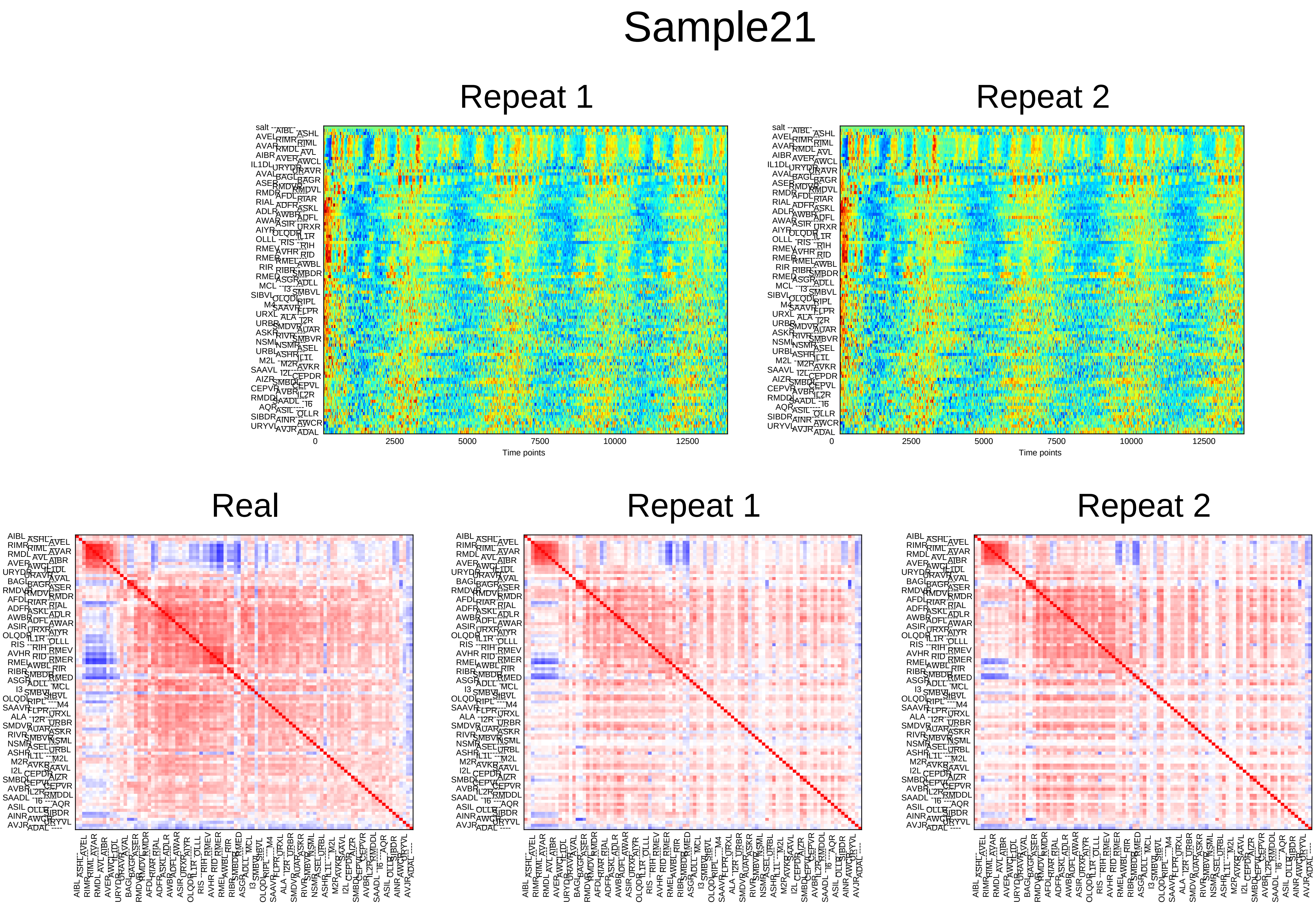
Sample21
Repeat 1
Repeat 2
0
2500
5000
7500
10000
12500
Time points
Real
Repeat 1
Repeat 2

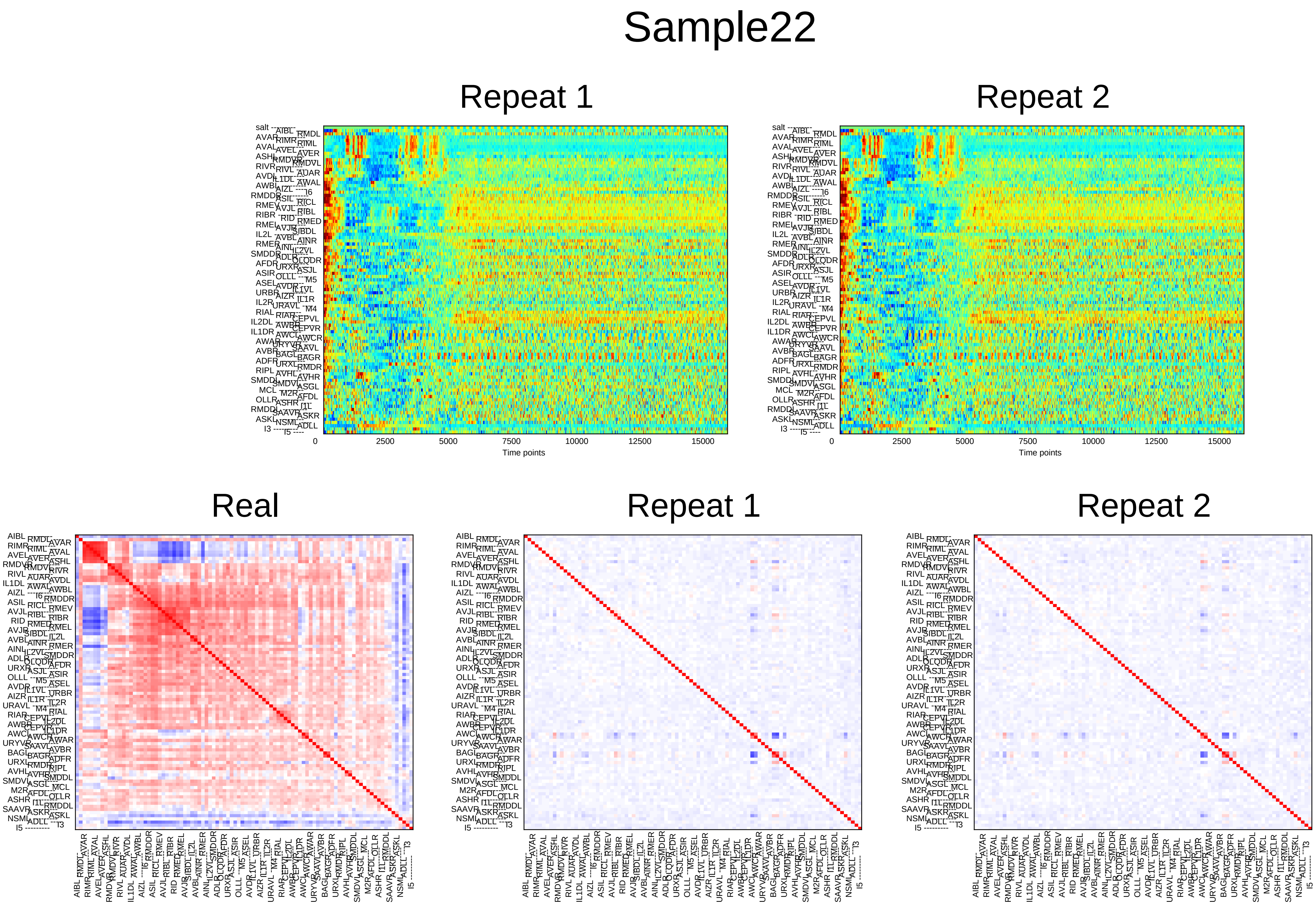
Sample22
Repeat 1
Repeat 2
0
2500
5000
7500
10000
12500
15000
Time points
Real
Repeat 1
Repeat 2

### Sample23

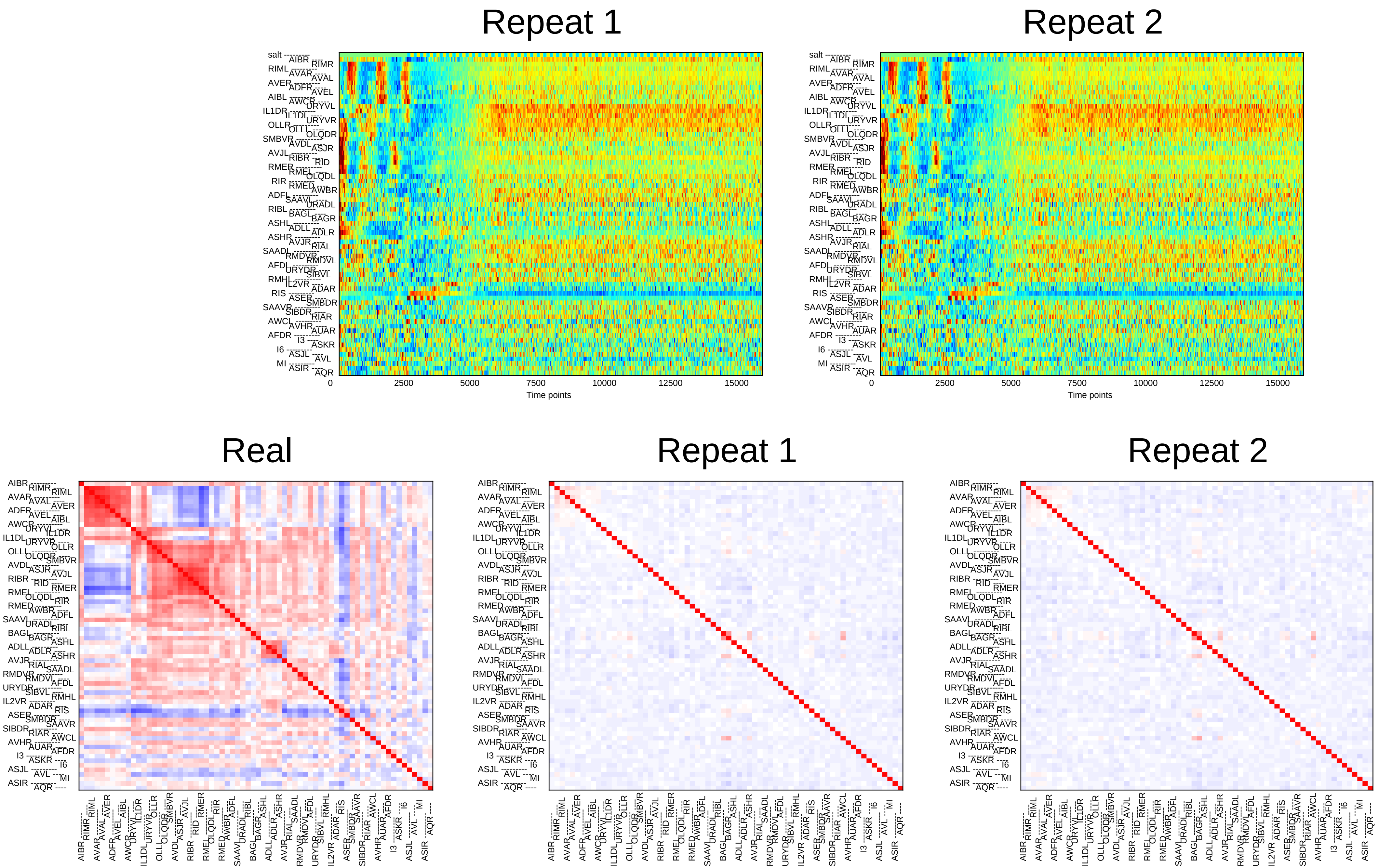
Repeat 1
Repeat 2
0
2500
5000
7500
10000
12500
15000
Time points
Real
Repeat 1
Repeat 2

### Sample24

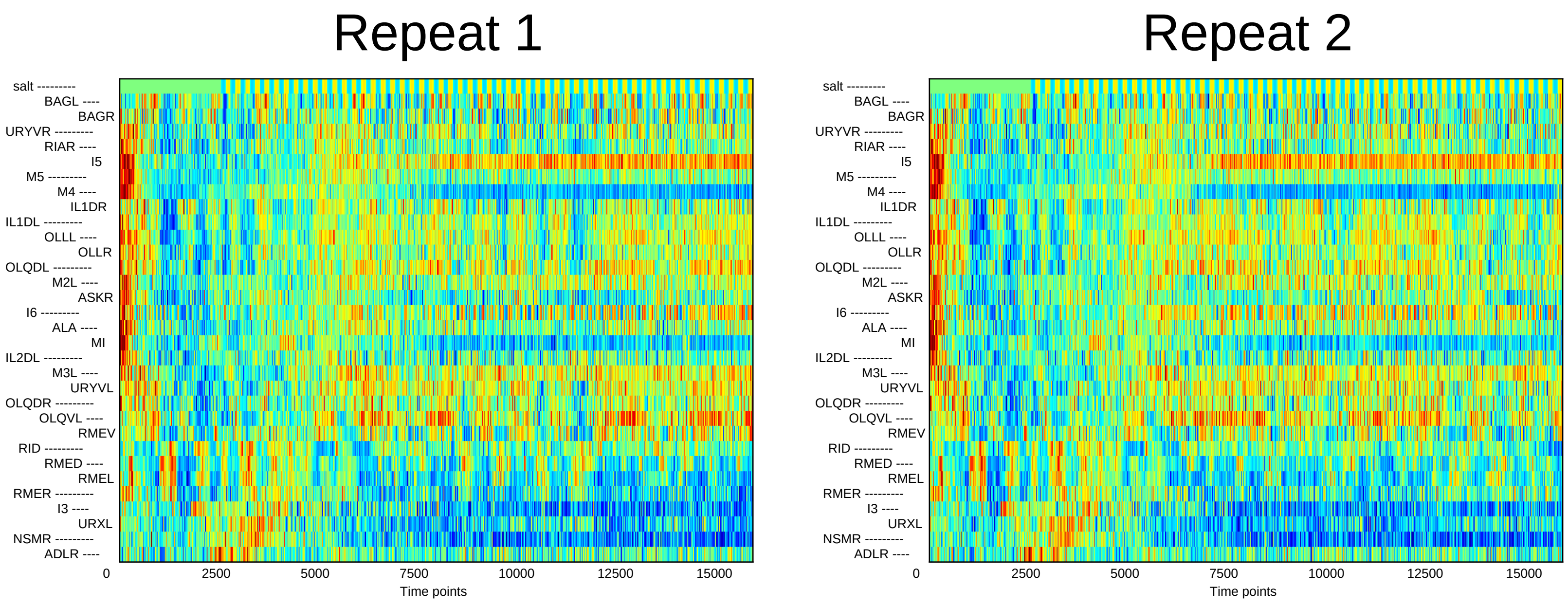
Repeat 1
Repeat 2
salt
BAGL
BAGR
URYVR
RIAR
I5
M5
M4
IL1DR
IL1DL
OLLL
OLLR
OLQDL
M2L
ASKR
I6
ALA
MI
IL2DL
M3L
URYVL
OLQDR
OLQVL
RMEV
RID
RMED
RMEL
RMER
I3
URXL
NSMR
ADLR
0
2500
5000
7500
10000
12500
15000
Time points

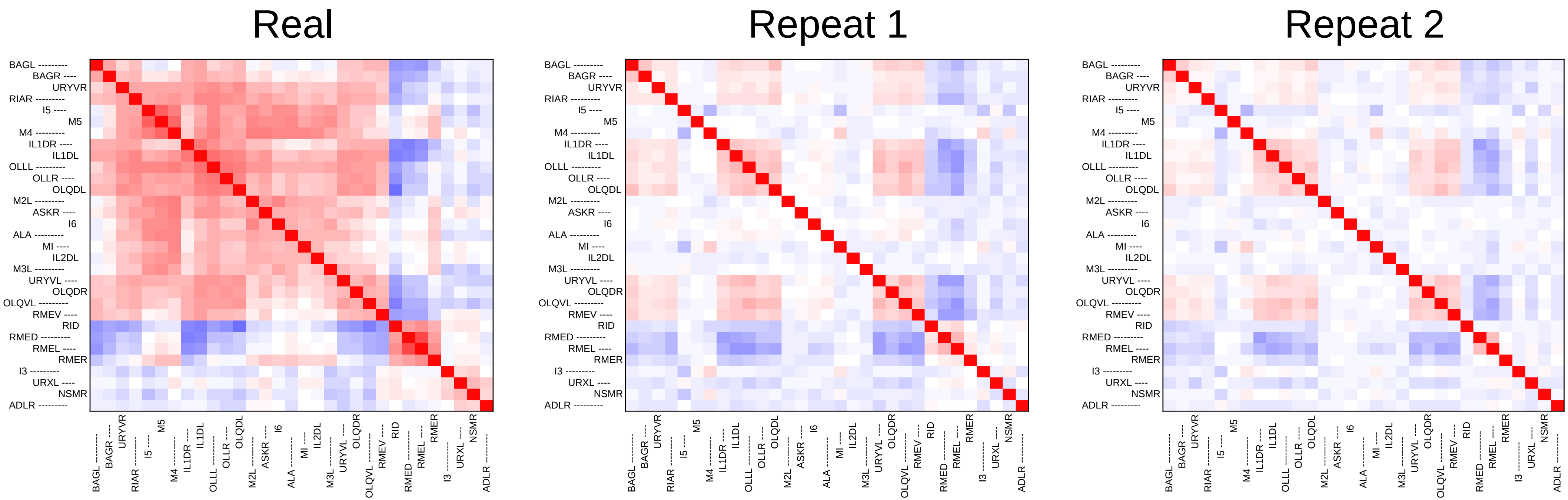
Real
Repeat 1
Repeat 2
BAGL
BAGR
URYVR
RIAR
I5
M5
M4
IL1DR
IL1DL
OLLL
OLLR
OLQDL
M2L
ASKR
I6
ALA
MI
IL2DL
M3L
URYVL
OLQDR
OLQVL
RMEV
RID
RMED
RMEL
RMER
I3
URXL
NSMR
ADLR
