## Supplementary File 4C for "Deducing ensemble dynamics and information flow from the whole-brain imaging data"

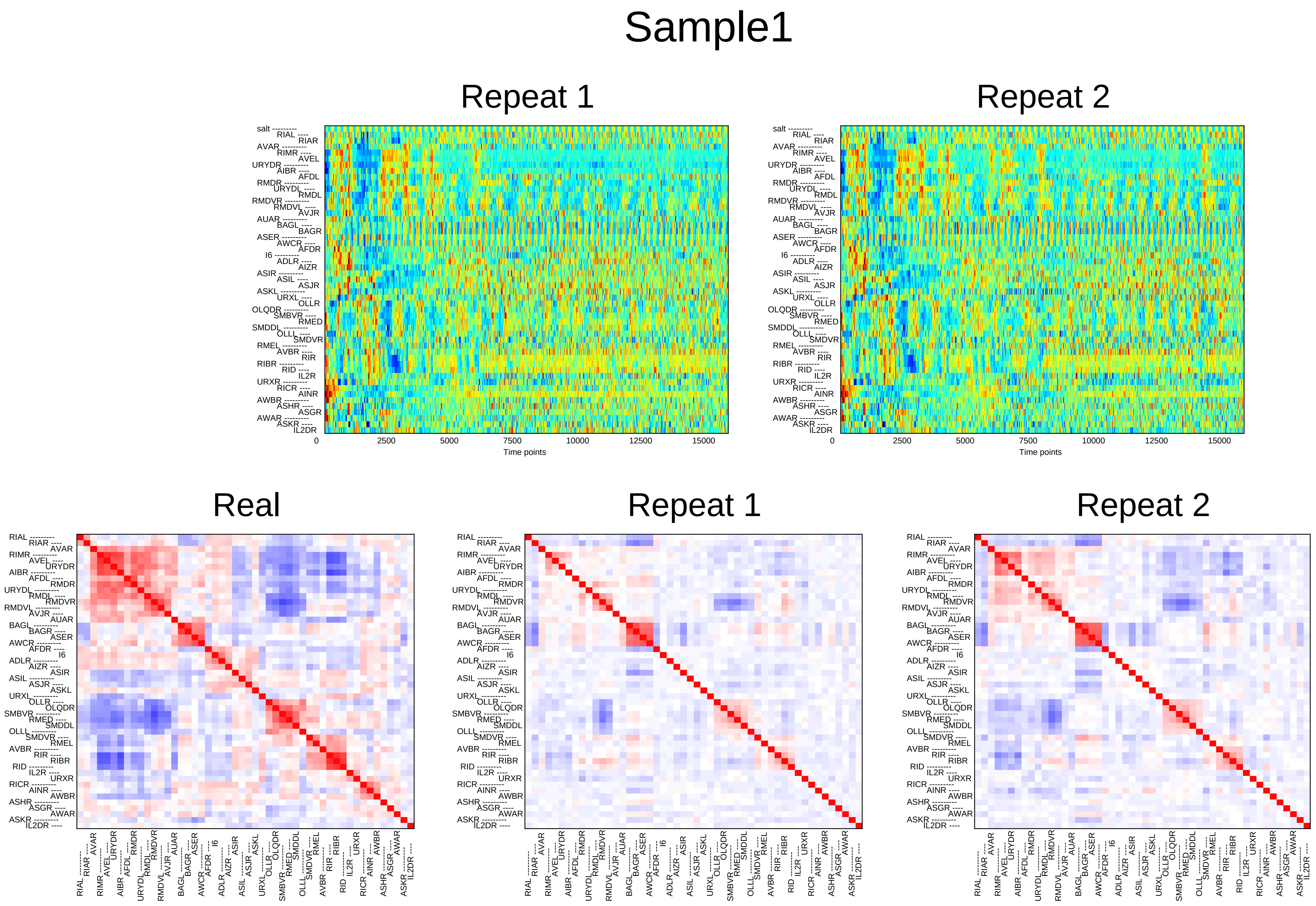
Sample1
Repeat 1
Repeat 2
salt RIAL RIAR AVAR RIMR AVEL URYDR AIBR AFDL RMDR URYDL RMDL RMDVR RMDVL AVJR AUAR BAGL BAGR ASER AWCR AFDR I6 ADLR AIZR ASIR ASIL ASJR ASKL URXL OLLR OLQDR SMBVR RMED SMDDL OLLL SMDVR RMEL AVBR RIR RIBR RID IL2R URXR RICR AINR AWBR ASHR ASGR AWAR ASKR IL2DR
0
2500
5000
7500
10000
12500
15000
Time points
Real
Repeat 1
Repeat 2
RIAL RIAR AVAR RIMR AVEL URYDR AIBR AFDL RMDR URYDL RMDL RMDVR RMDVL AVJR AUAR BAGL BAGR ASER AWCR AFDR I6 ADLR AIZR ASIR ASIL ASJR ASKL URXL OLLR OLQDR SMBVR RMED SMDDL OLLL SMDVR RMEL AVBR RIR RIBR RID IL2R URXR RICR AINR AWBR ASHR ASGR AWAR ASKR IL2DR

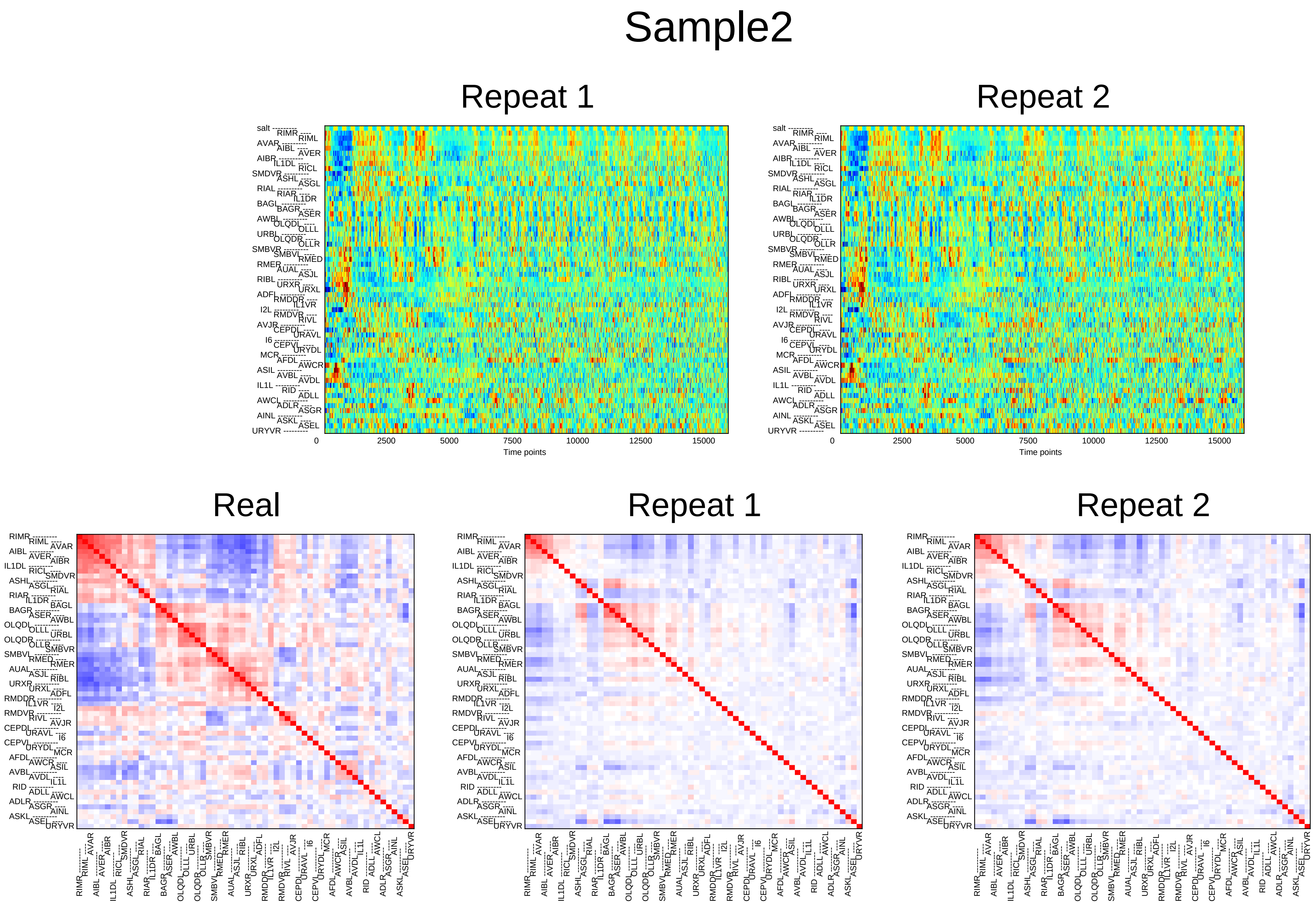

Sample2
Repeat 1
Repeat 2
Time points
0
2500
5000
7500
10000
12500
15000
Real
Repeat 1
Repeat 2

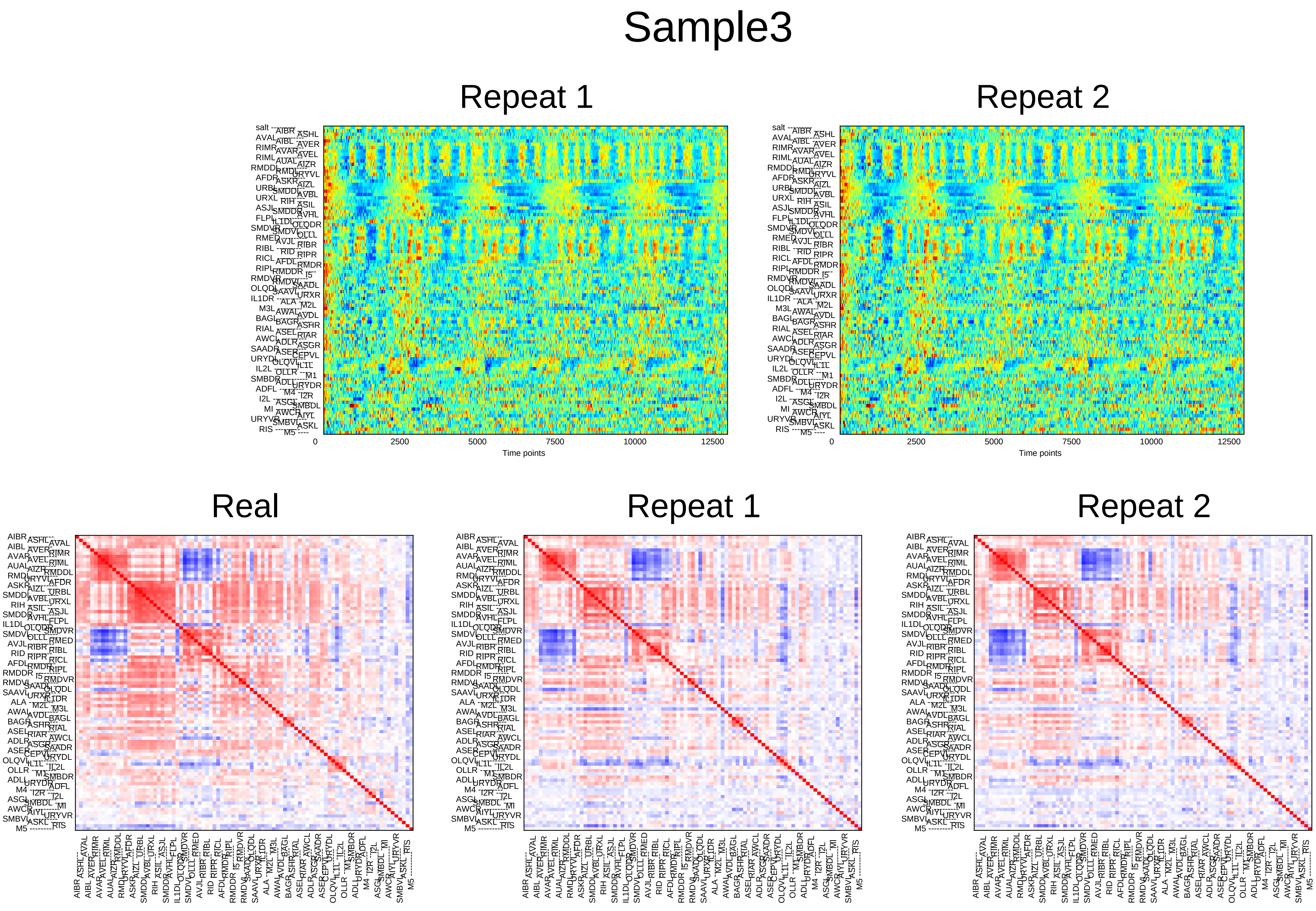

Sample3
Repeat 1
Repeat 2
0
2500
5000
7500
10000
12500
Time points
Real
Repeat 1
Repeat 2

### Sample4

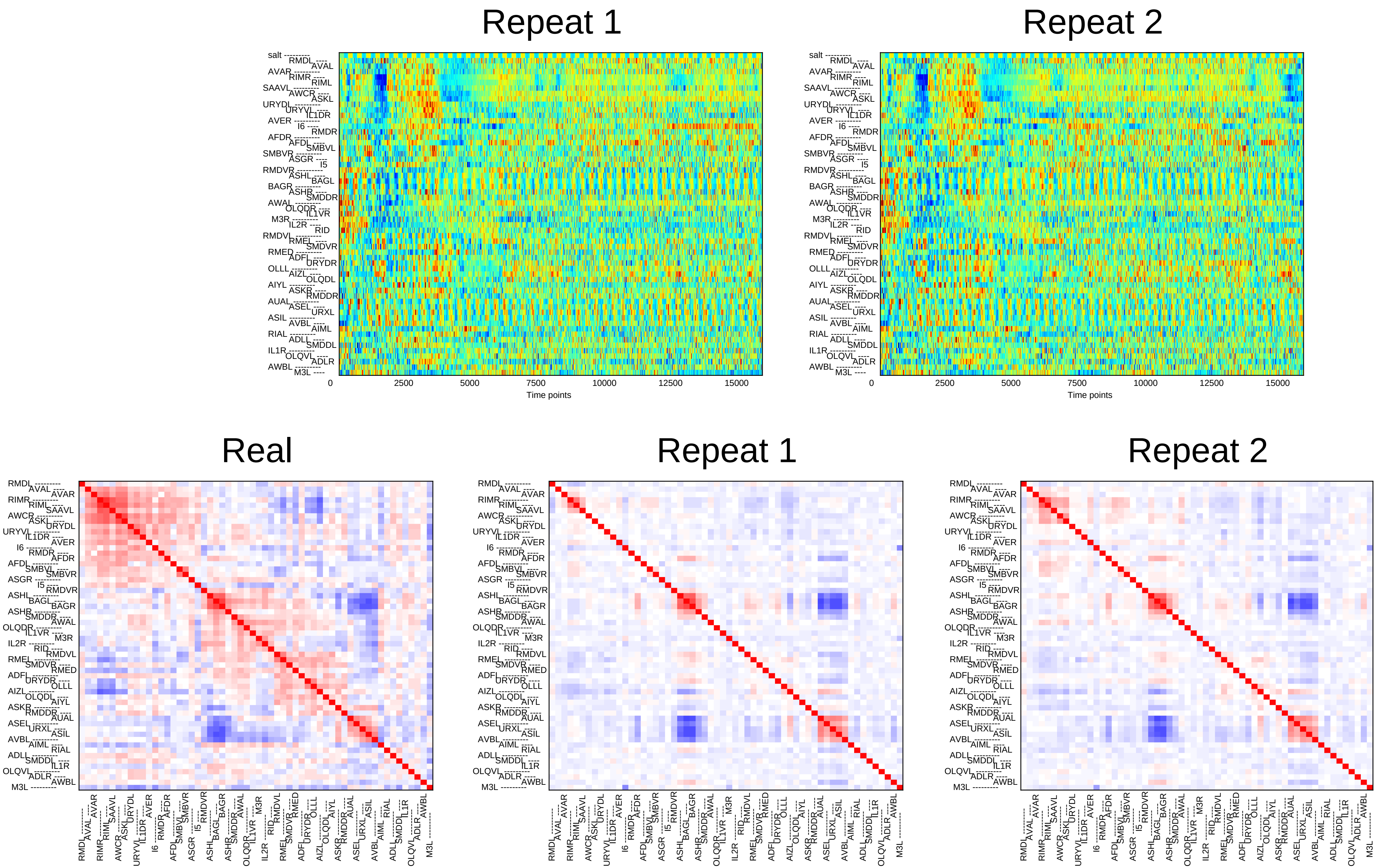
Repeat 1
Repeat 2
Time points
0
2500
5000
7500
10000
12500
15000
Real
Repeat 1
Repeat 2

### Sample5

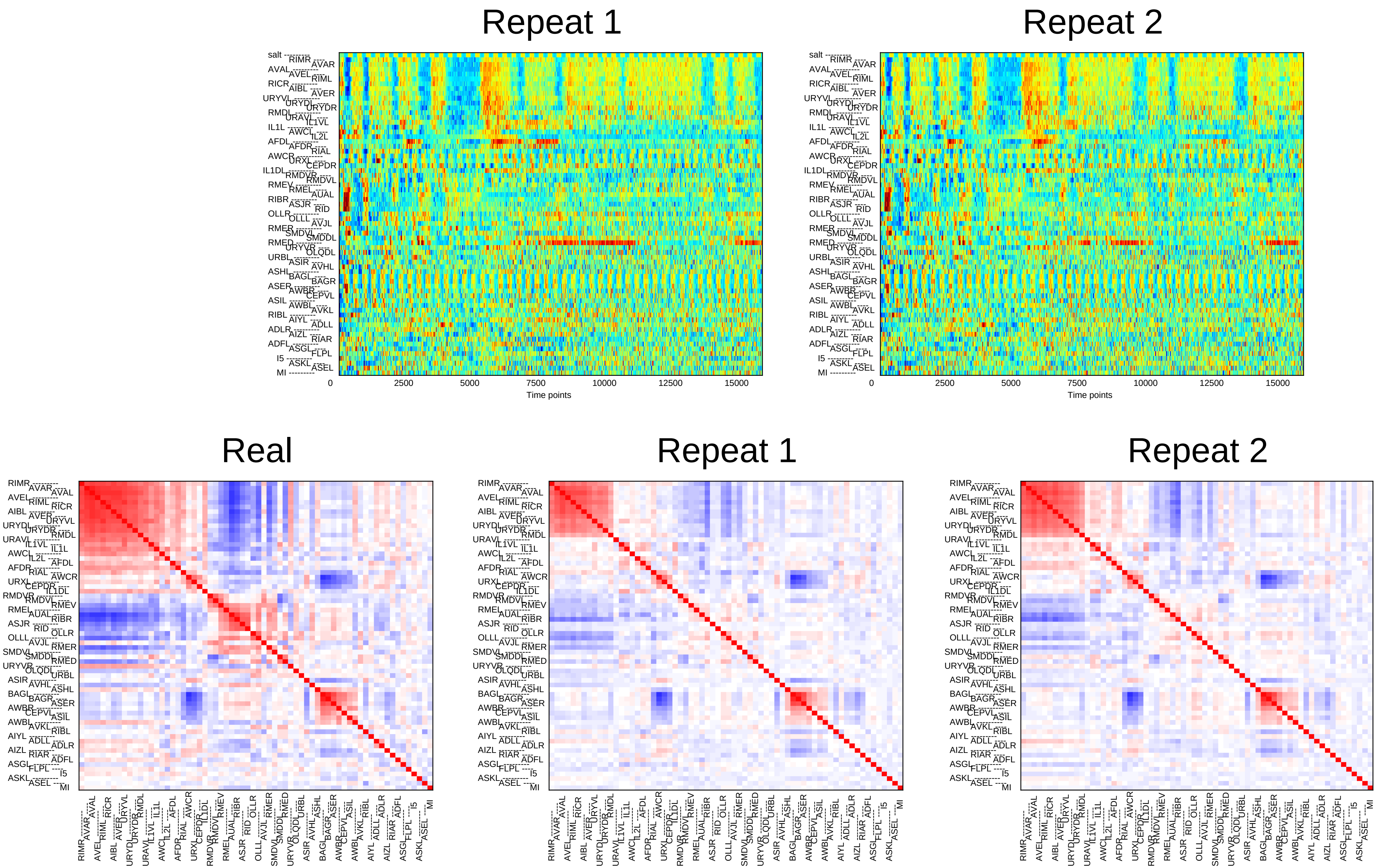

Repeat 1
Repeat 2
0
2500
5000
7500
10000
12500
15000
Time points
Real
Repeat 1
Repeat 2

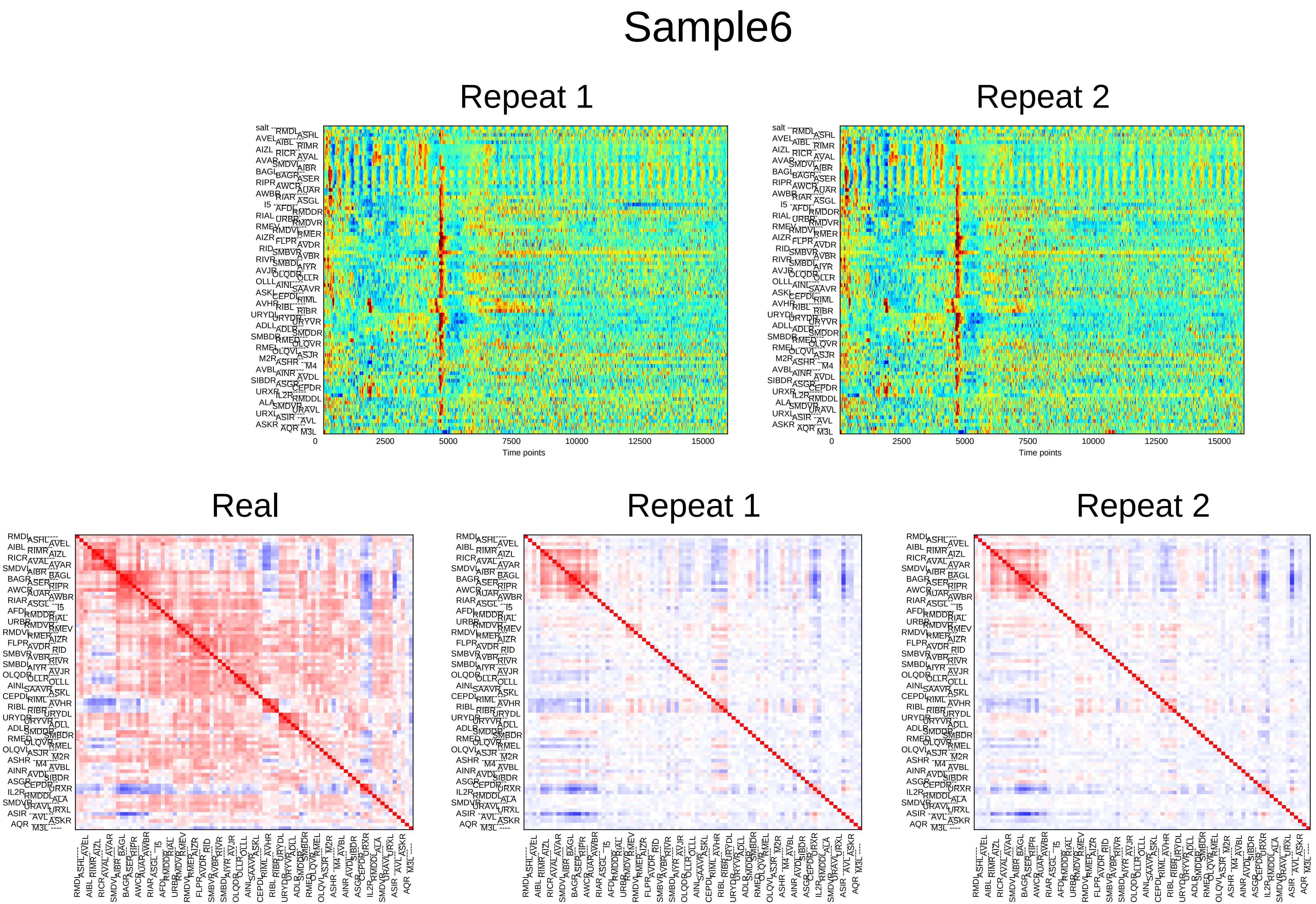
Sample6
Repeat 1
Repeat 2
0
2500
5000
7500
10000
12500
15000
Time points
Real
Repeat 1
Repeat 2

### Sample7

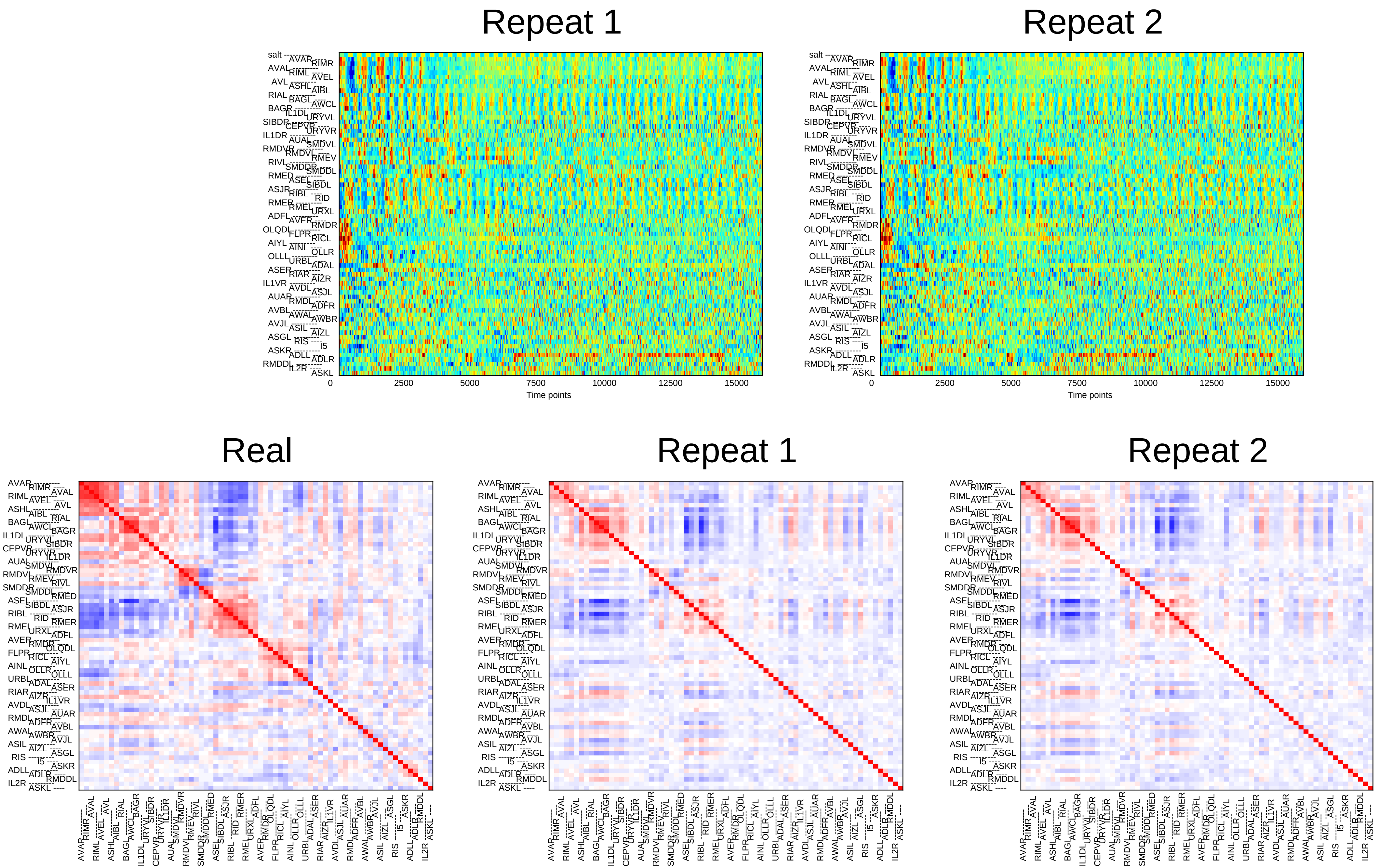
Repeat 1
Repeat 2
0
2500
5000
7500
10000
12500
15000
Time points
Real
Repeat 1
Repeat 2

### Sample8

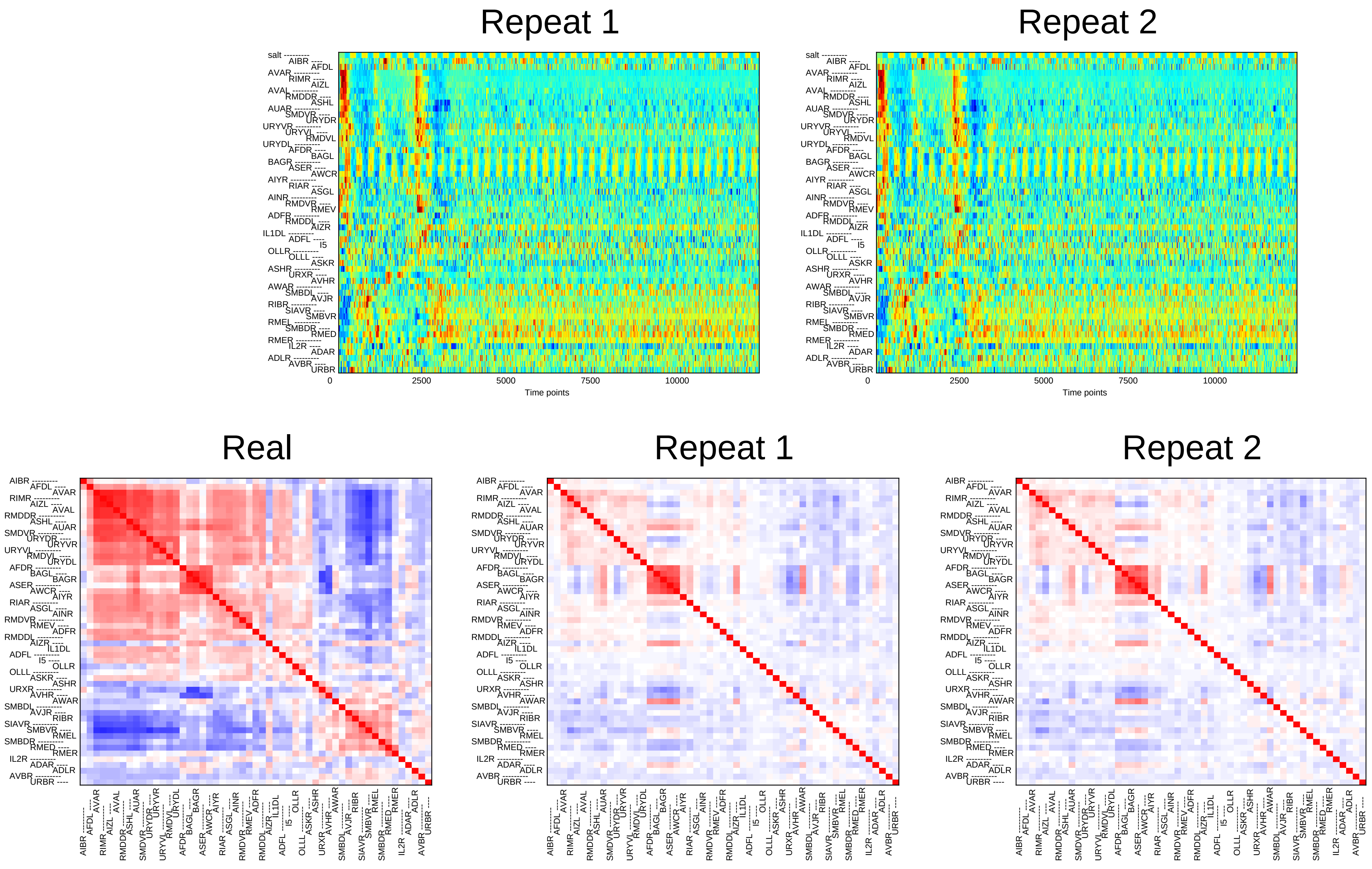
Repeat 1
Repeat 2
0
2500
5000
7500
10000
Time points
Real
Repeat 1
Repeat 2

### Sample9

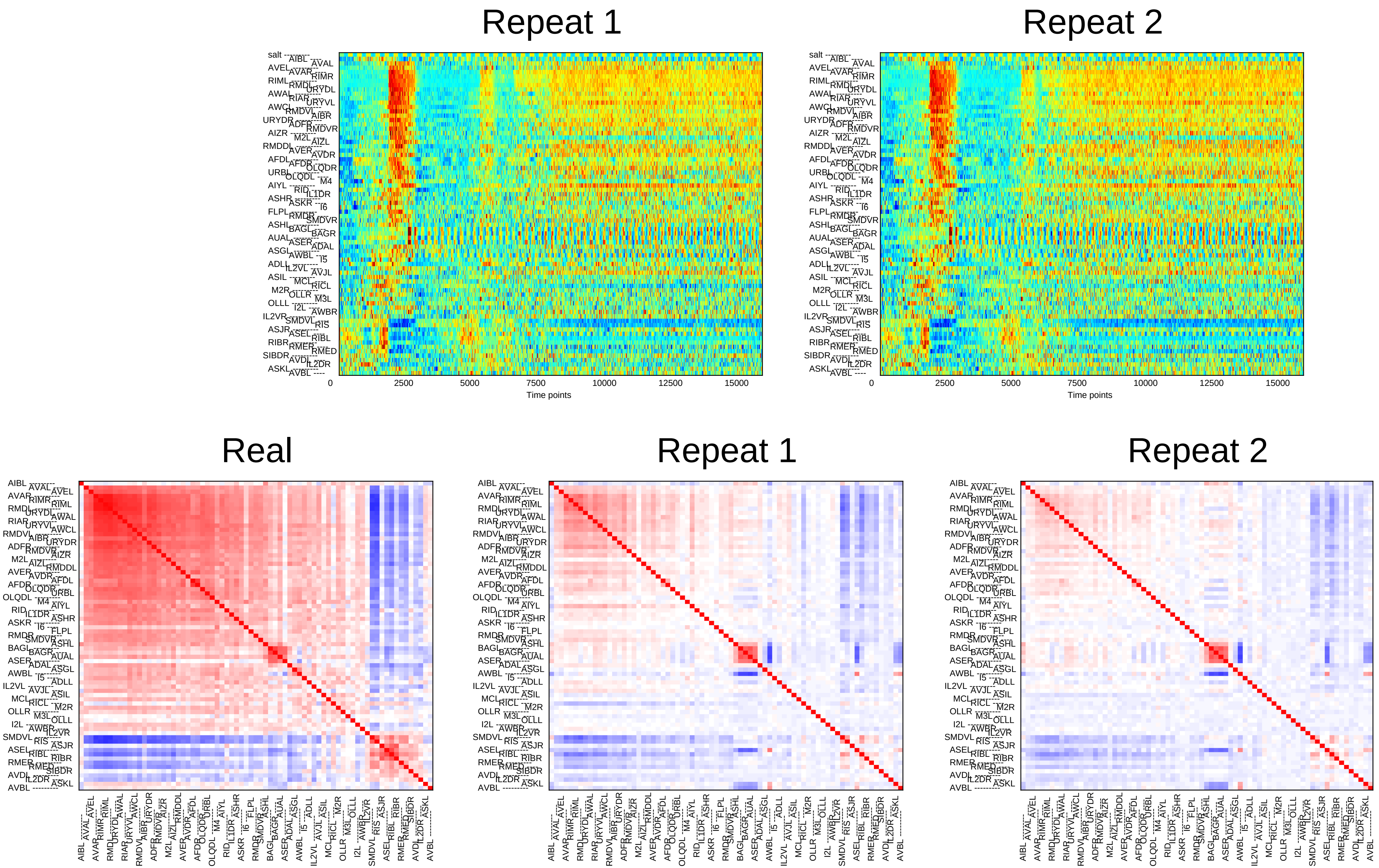
Repeat 1
Repeat 2
Time points
0
2500
5000
7500
10000
12500
15000
Real
Repeat 1
Repeat 2

### Sample10

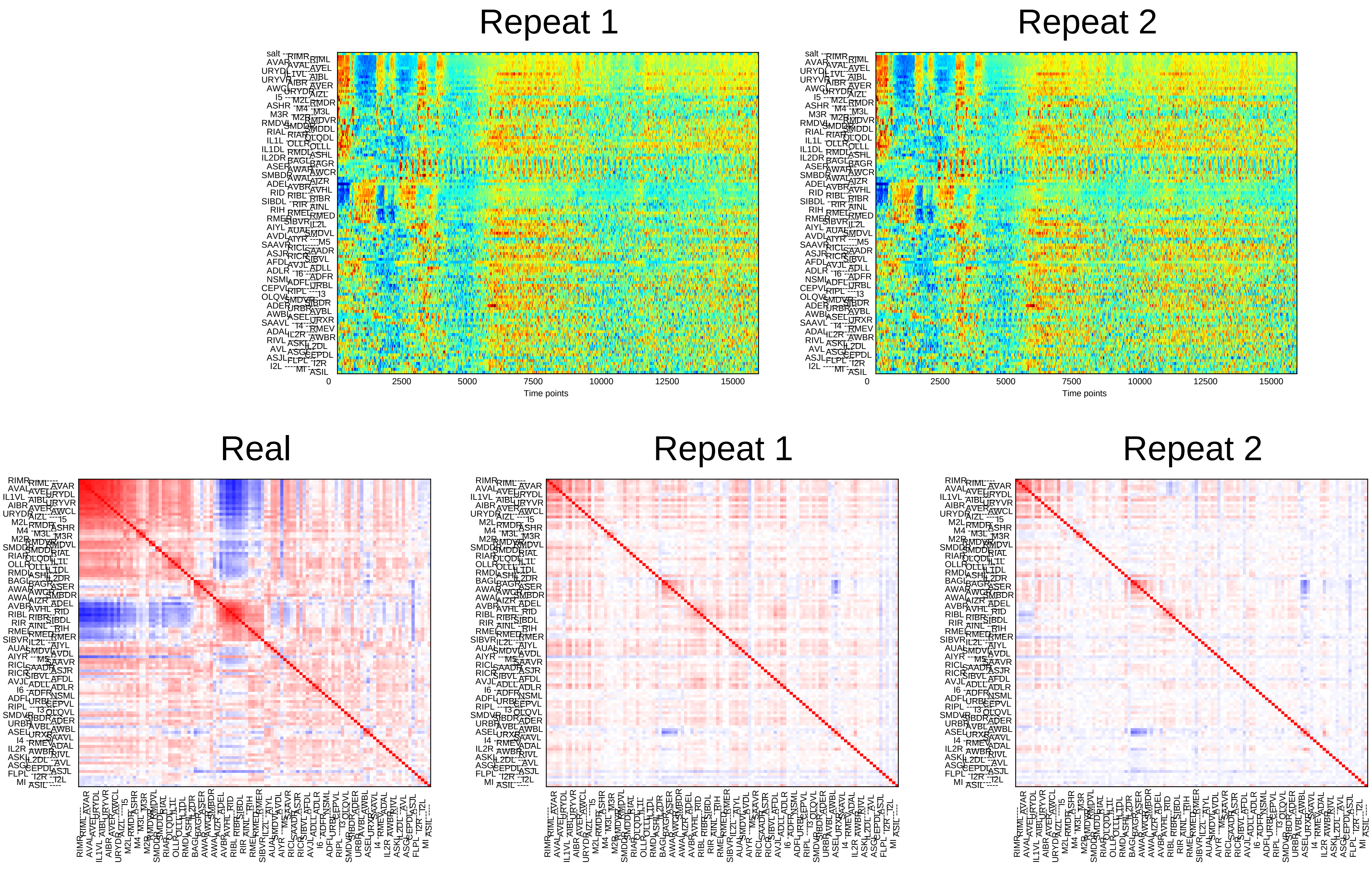
Repeat 1
Repeat 2
0
2500
5000
7500
10000
12500
15000
Time points
Real
Repeat 1
Repeat 2

### Sample11

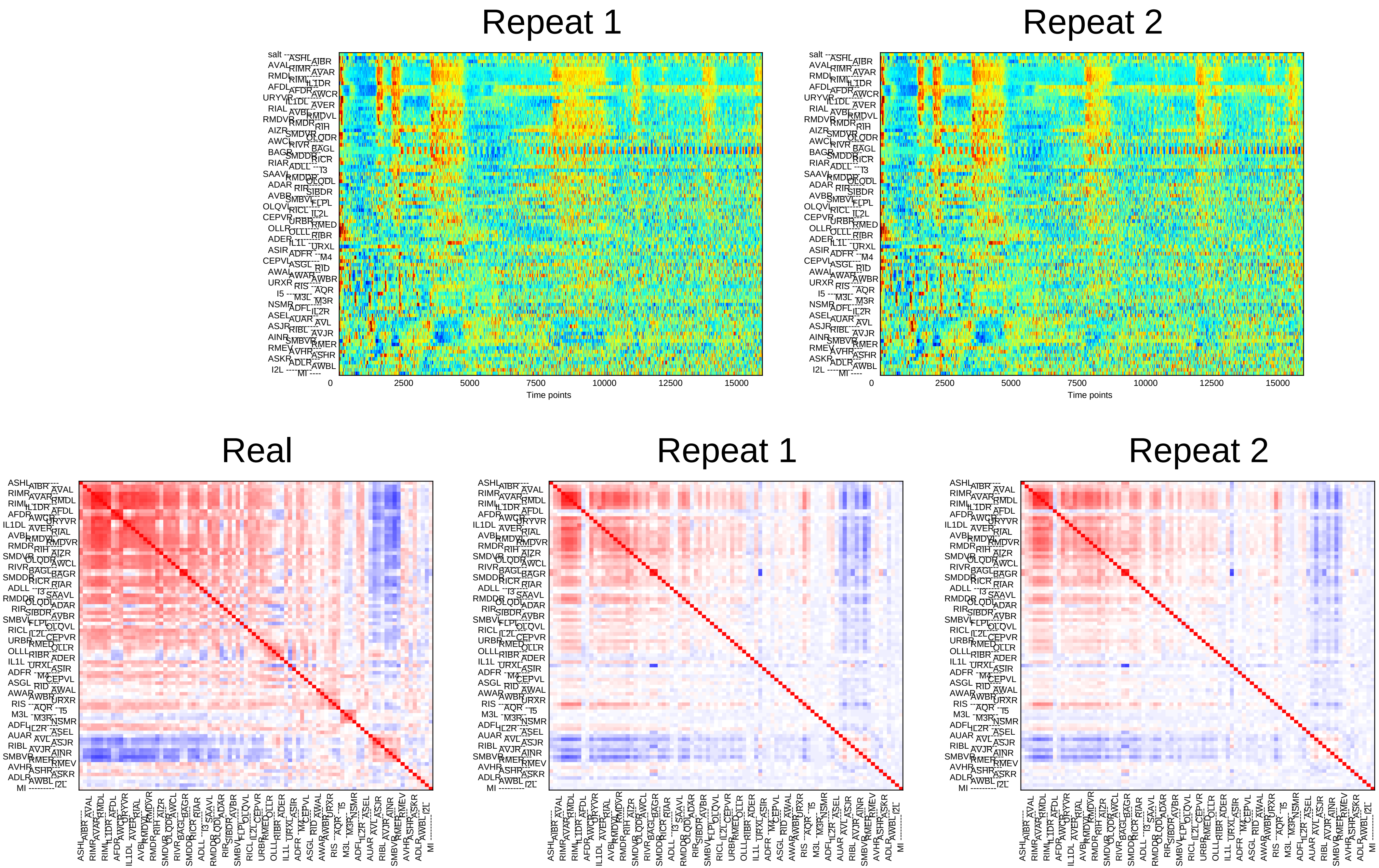
Repeat 1
Repeat 2
0
2500
5000
7500
10000
12500
15000
Time points
Real
Repeat 1
Repeat 2

### Sample12

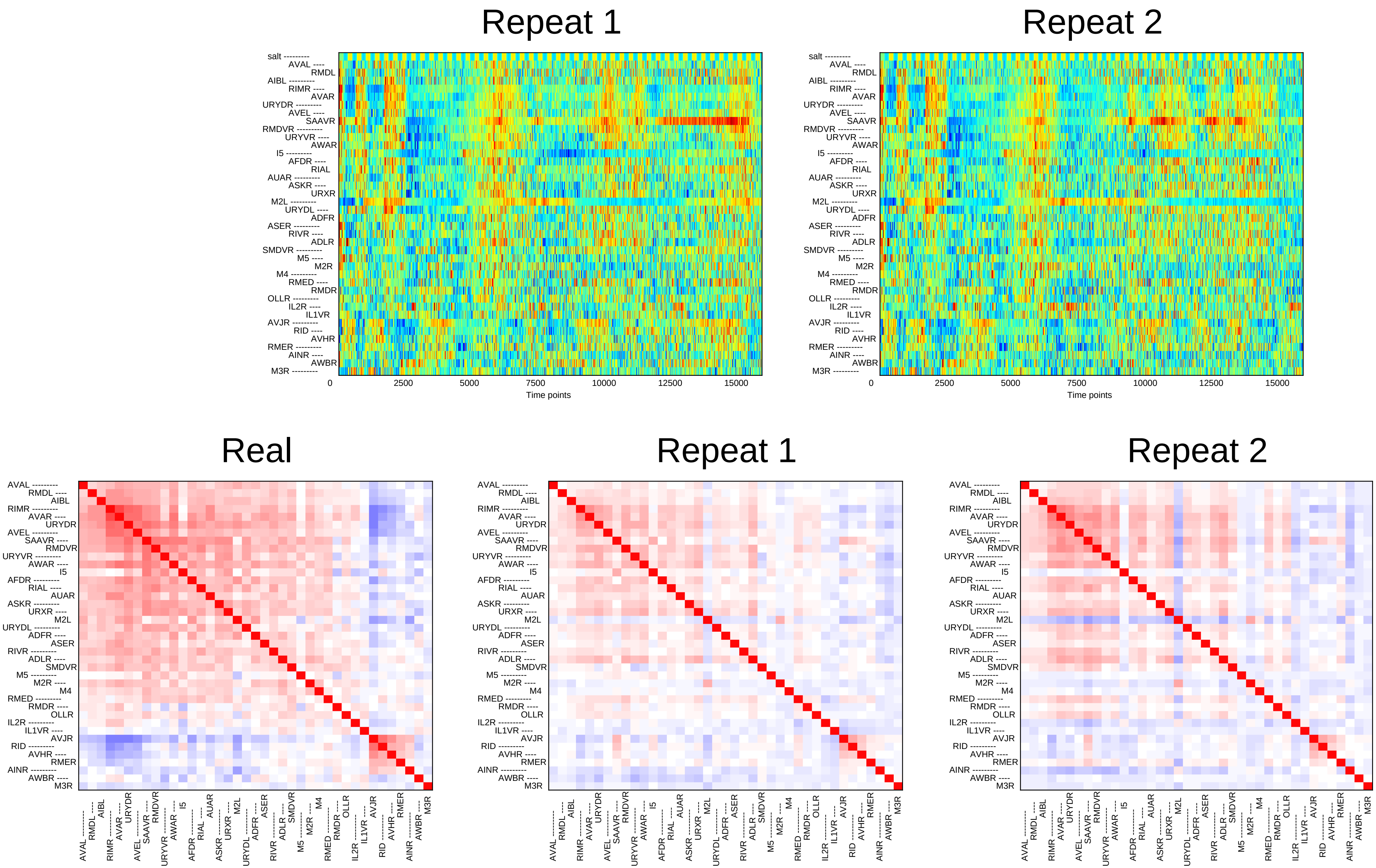
Repeat 1
Repeat 2
salt
AVAL RMDL AIBL RIMR AVAR URYDR AVEL SAAVR RMDVR URYVR AWAR I5 AFDR RIAL AUAR ASKR URXR M2L URYDL ADFR ASER RIVR ADLR SMDVR M5 M2R M4 RMED RMDR OLLR IL2R IL1VR AVJR RID AVHR RMER AINR AWBR M3R
0
2500
5000
7500
10000
12500
15000
Time points
Real
Repeat 1
Repeat 2

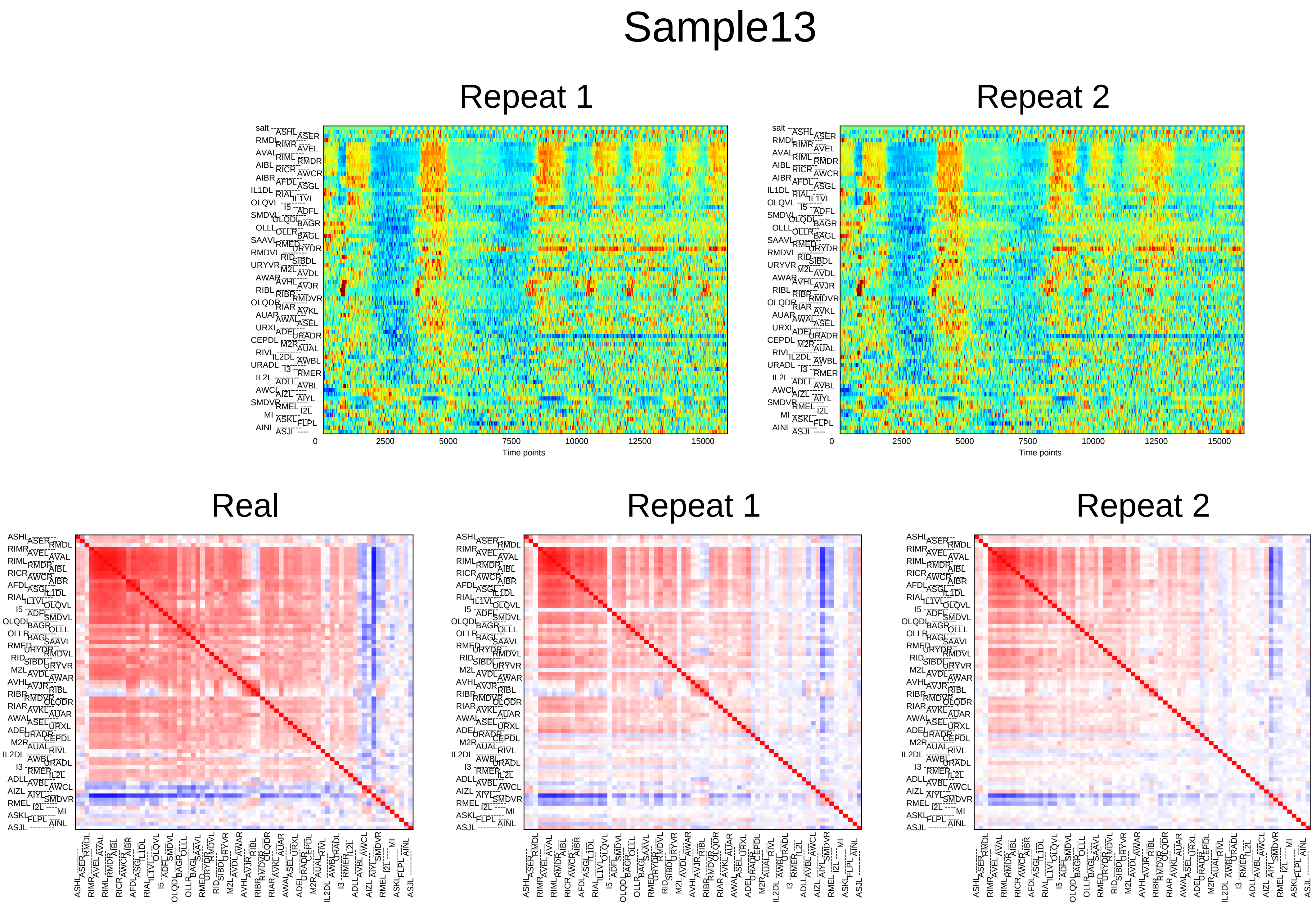
Sample13
Repeat 1
Repeat 2
0
2500
5000
7500
10000
12500
15000
Time points
Real
Repeat 1
Repeat 2

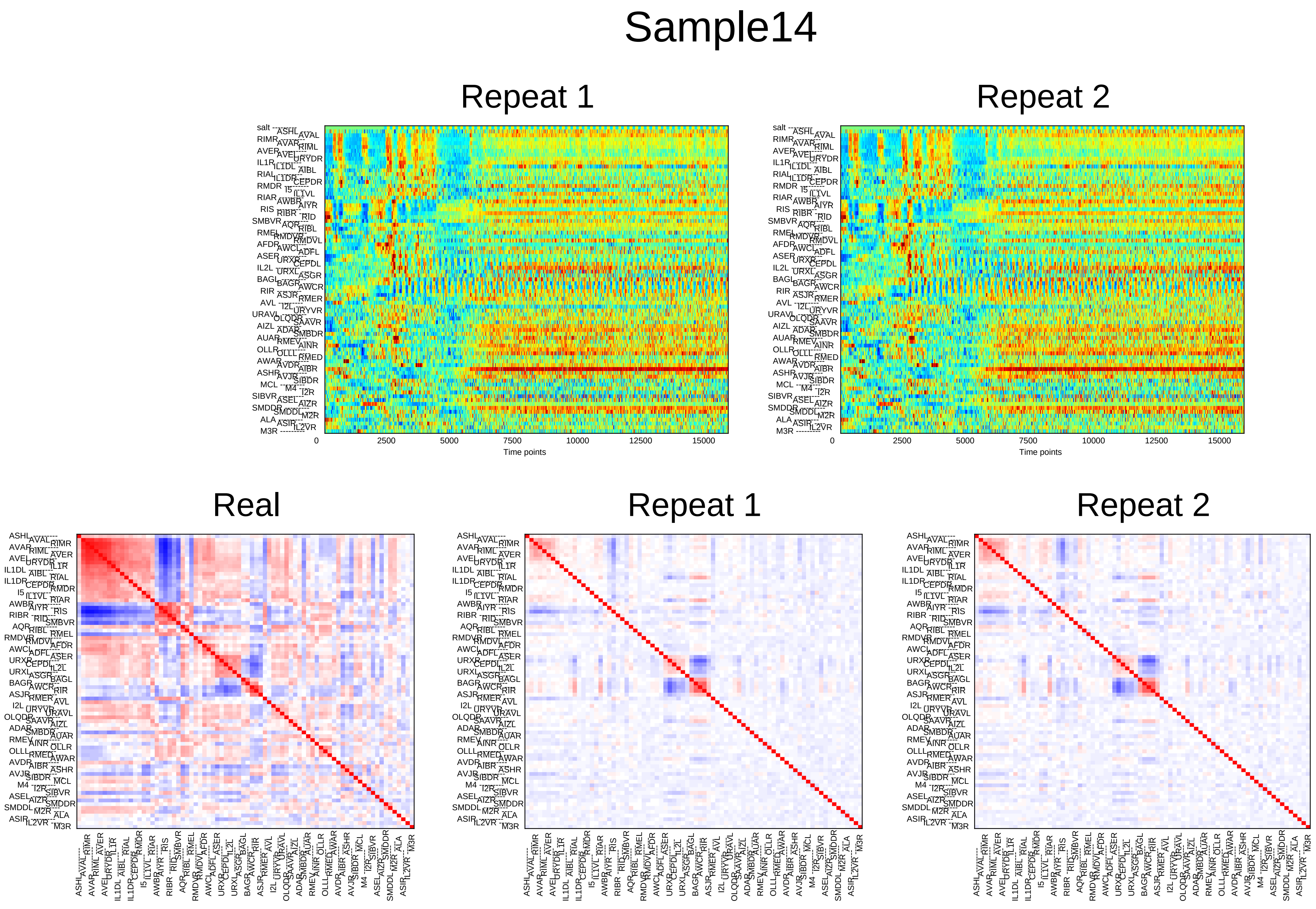
Sample14
Repeat 1
Repeat 2
0
2500
5000
7500
10000
12500
15000
Time points
Real
Repeat 1
Repeat 2

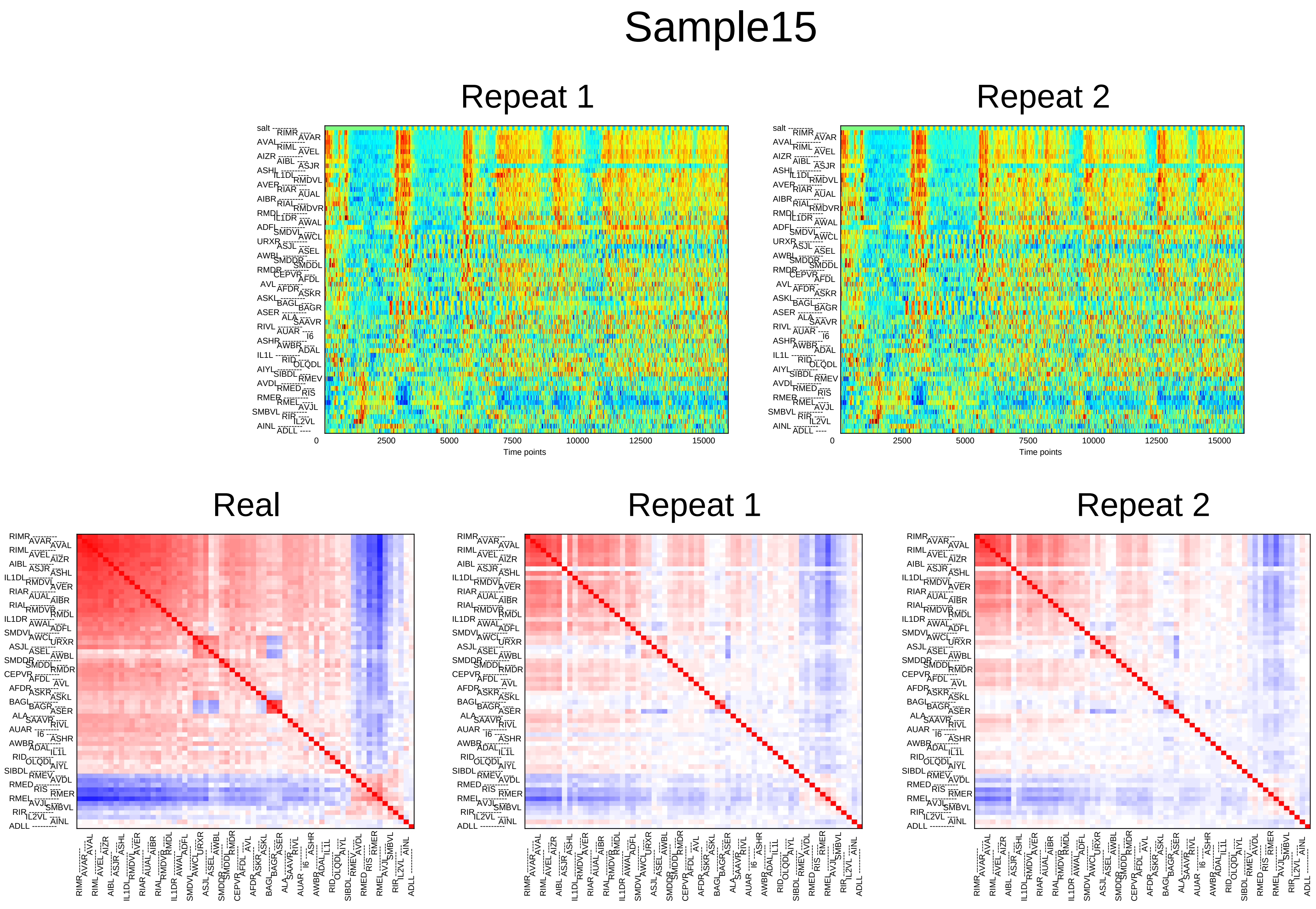
Sample15
Repeat 1
Repeat 2
0
2500
5000
7500
10000
12500
15000
Time points
Real
Repeat 1
Repeat 2

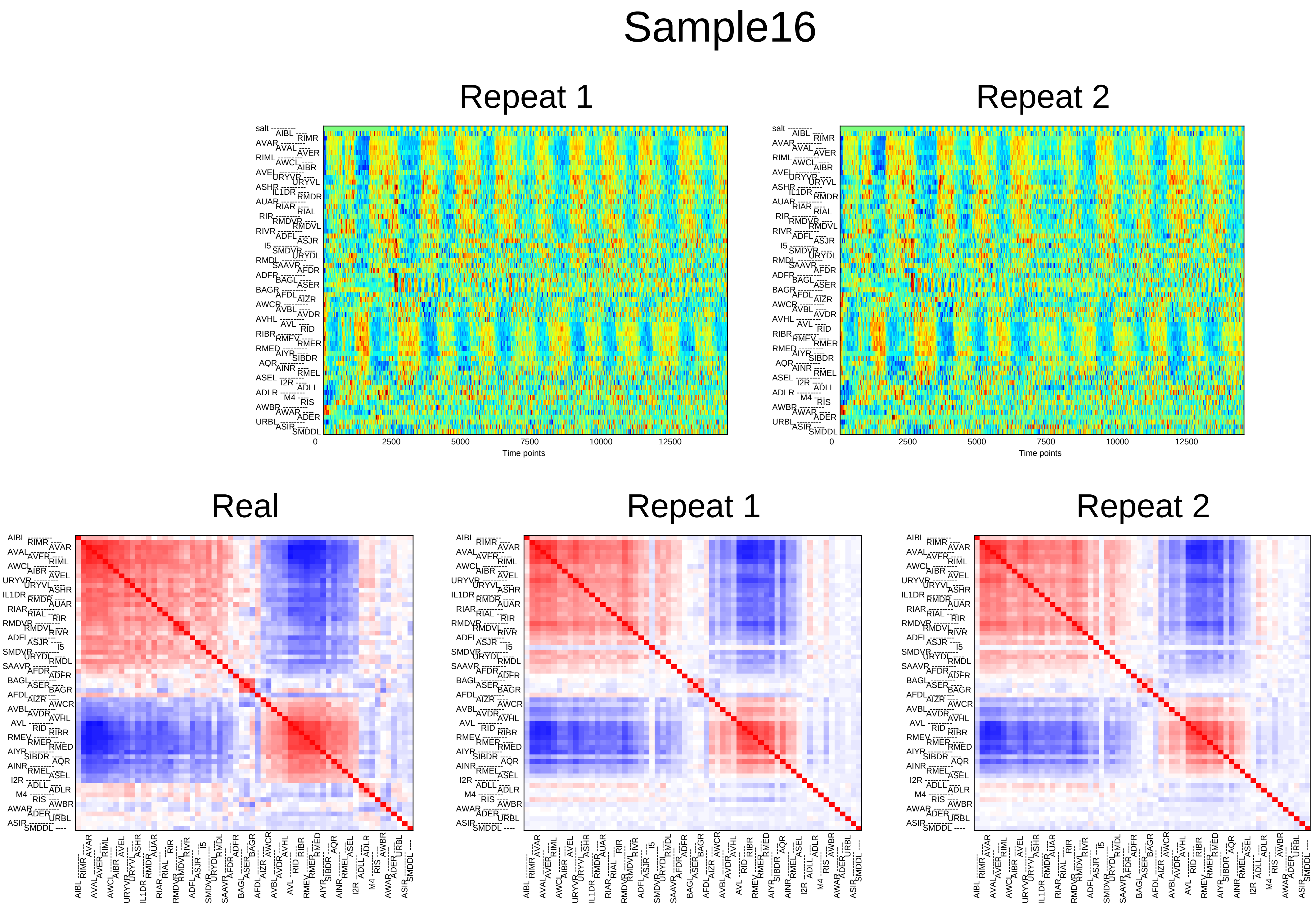
Sample16
Repeat 1
Repeat 2
salt
AIBL
RIMR
AVAR
AVAL
AVER
RIML
AWCL
AIBR
AVEL
URYVR
URYVL
ASHR
IL1DR
RMDR
AUAR
RIAR
RIAL
RIR
RMDVR
RMDVL
RIVR
ADFL
ASJR
I5
SMDVR
URYDL
RMDL
SAAVR
AFDR
ADFR
BAGL
ASER
BAGR
AFDL
AIZR
AWCR
AVBL
AVDR
AVHL
AVL
RID
RIBR
RMEV
RMER
RMED
AIYR
SIBDR
AQR
AINR
RMEL
ASEL
I2R
ADLL
ADLR
M4
RIS
AWBR
AWAR
ADER
URBL
ASIR
SMDDL
0
2500
5000
7500
10000
12500
Time points
Real
Repeat 1
Repeat 2

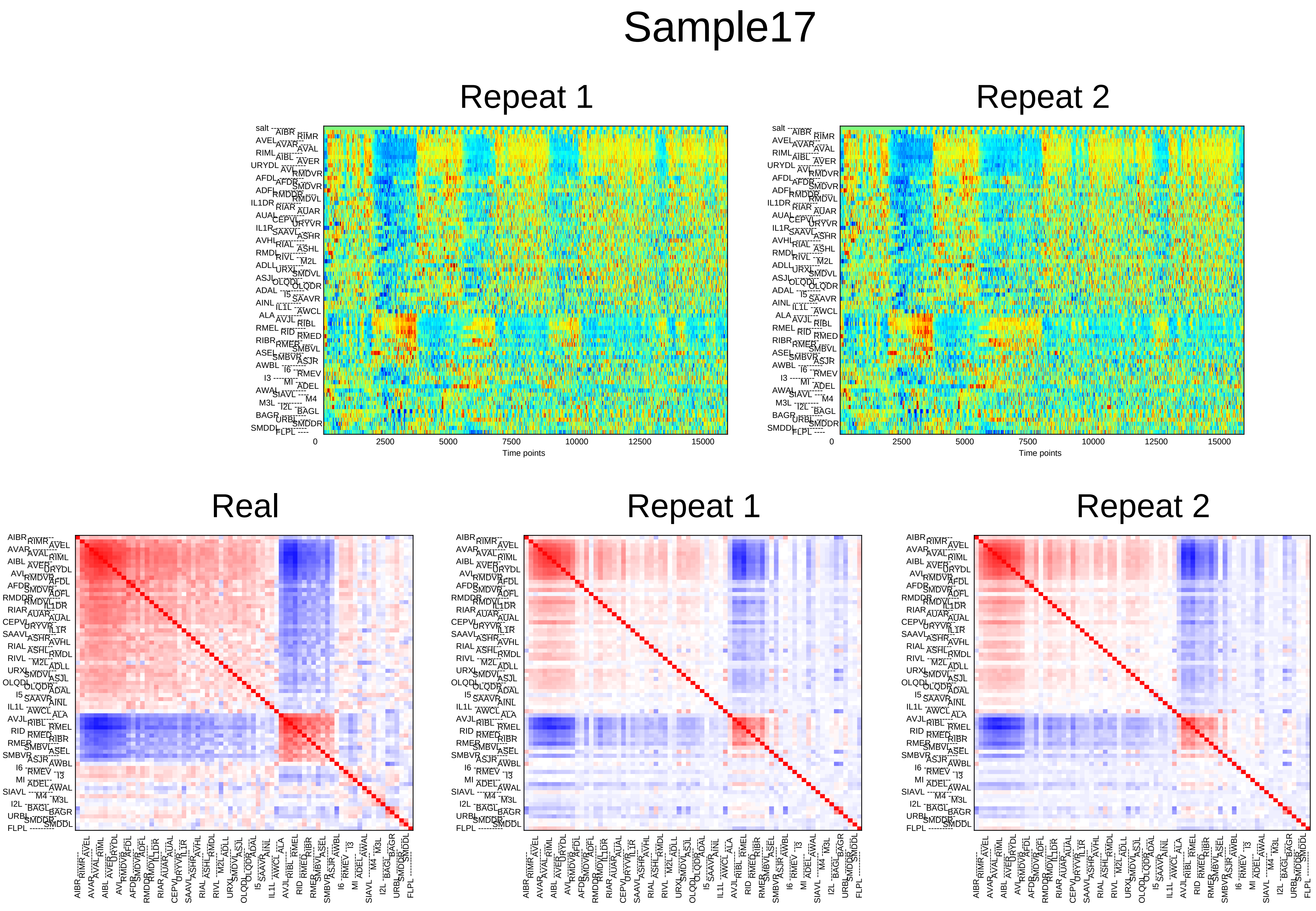

Sample17
Repeat 1
Repeat 2
0
2500
5000
7500
10000
12500
15000
Time points
Real
Repeat 1
Repeat 2

### Sample18

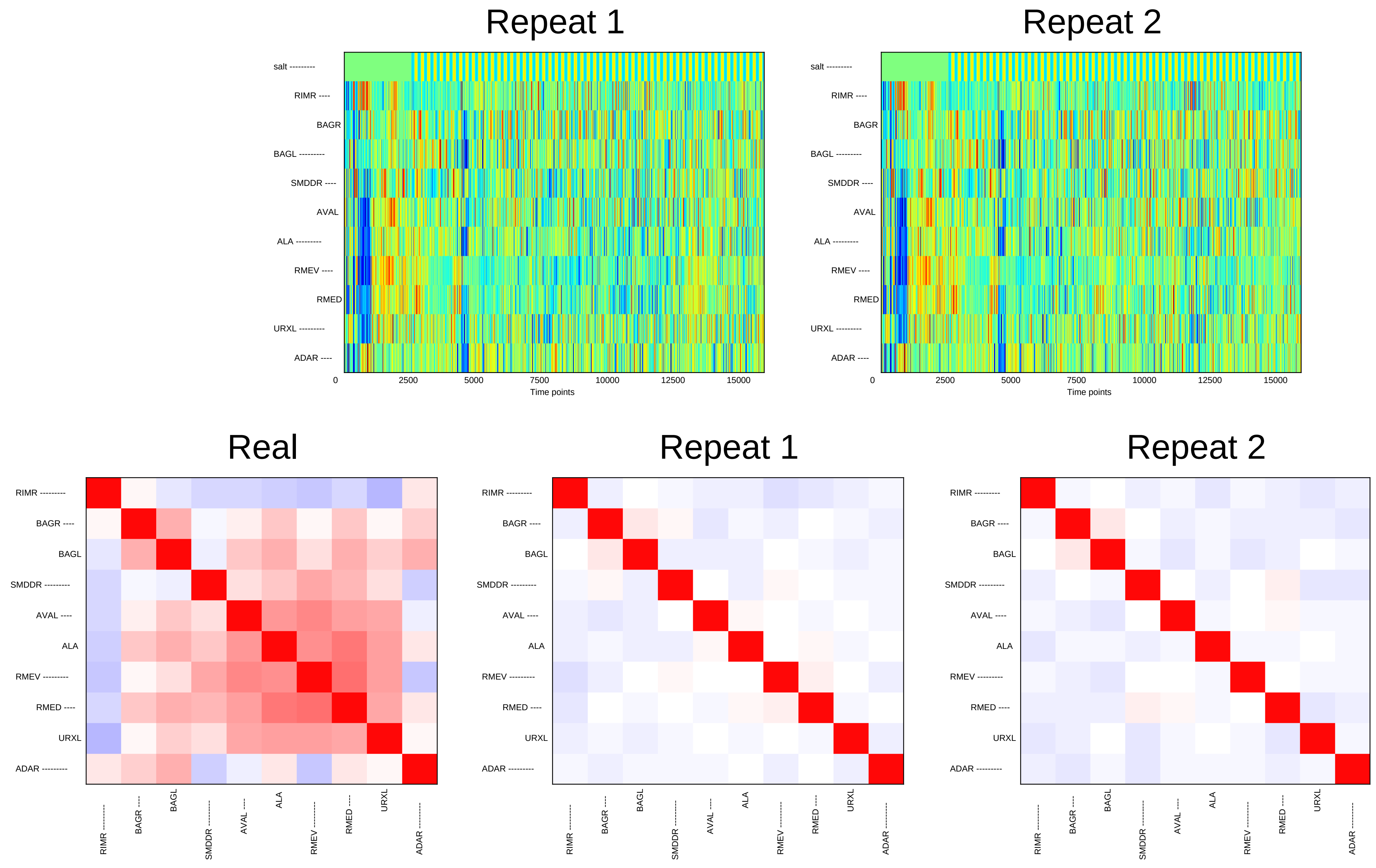

Repeat 1
Repeat 2
salt
RIMR
BAGR
BAGL
SMDDR
AVAL
ALA
RMEV
RMED
URXL
ADAR
0
2500
5000
7500
10000
12500
15000
Time points
Real
Repeat 1
Repeat 2

### Sample19

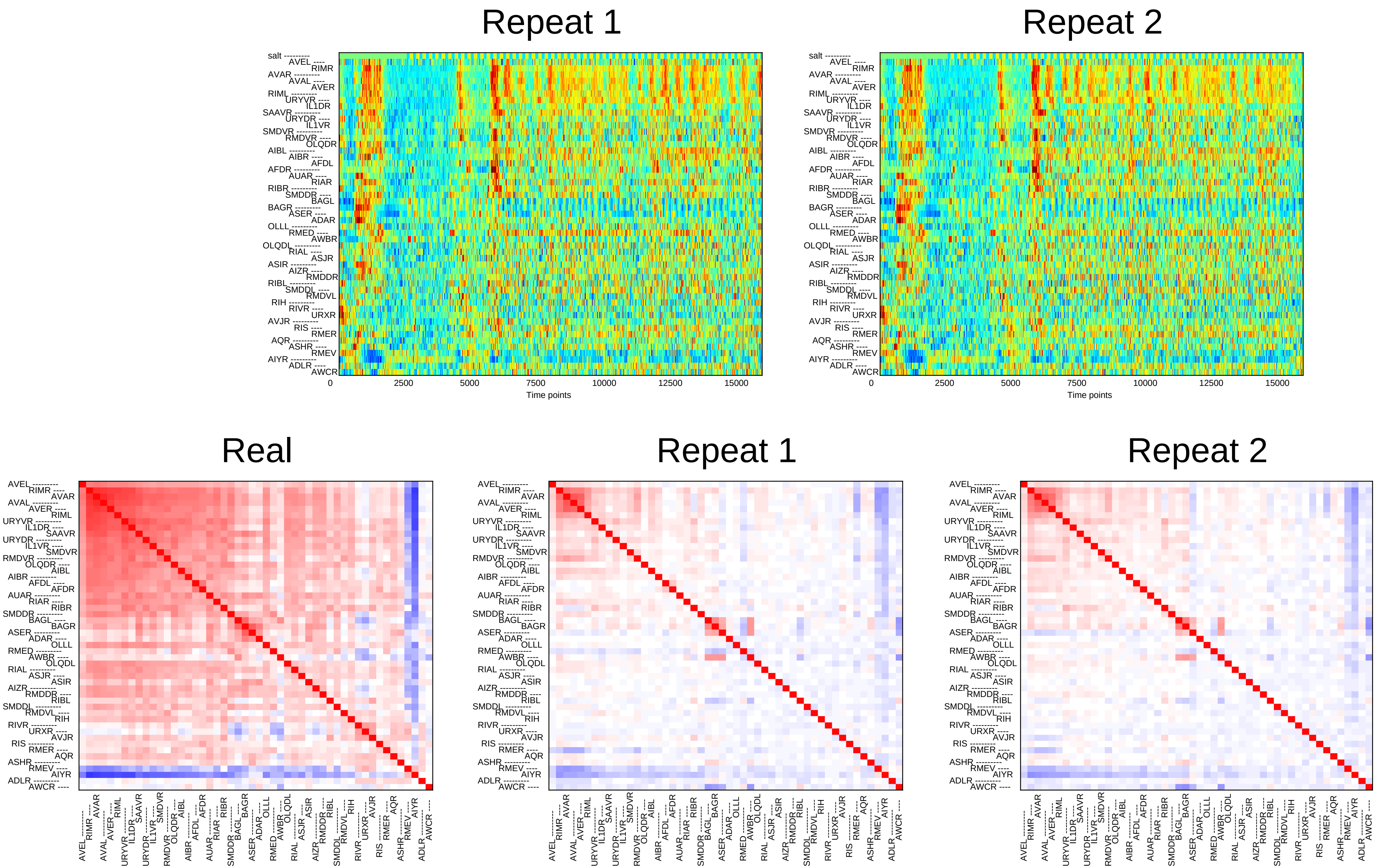

Repeat 1
Repeat 2
Time points
0
2500
5000
7500
10000
12500
15000
salt
Real
Repeat 1
Repeat 2

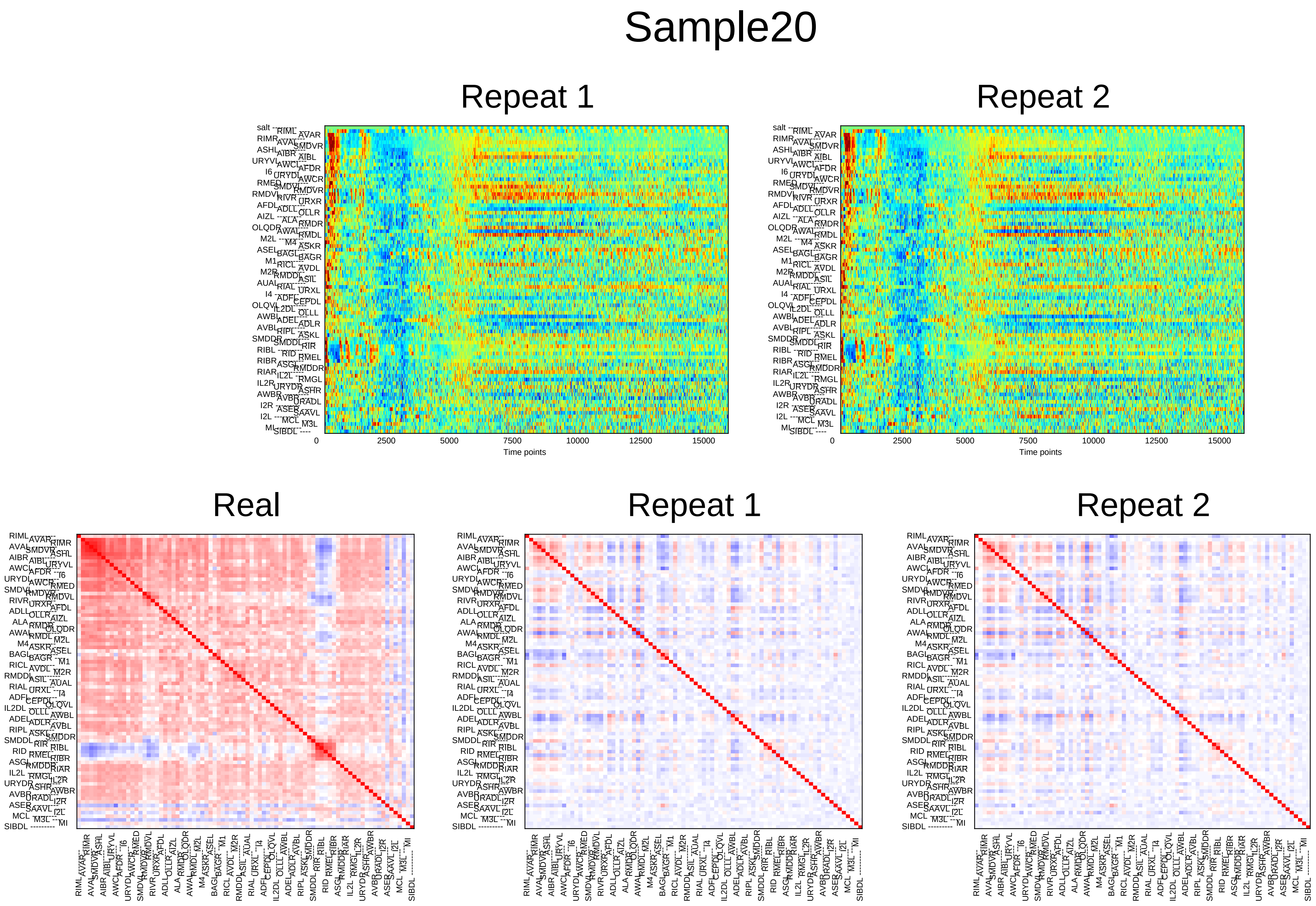

Sample20
Repeat 1
Repeat 2
0
2500
5000
7500
10000
12500
15000
Time points
Real
Repeat 1
Repeat 2

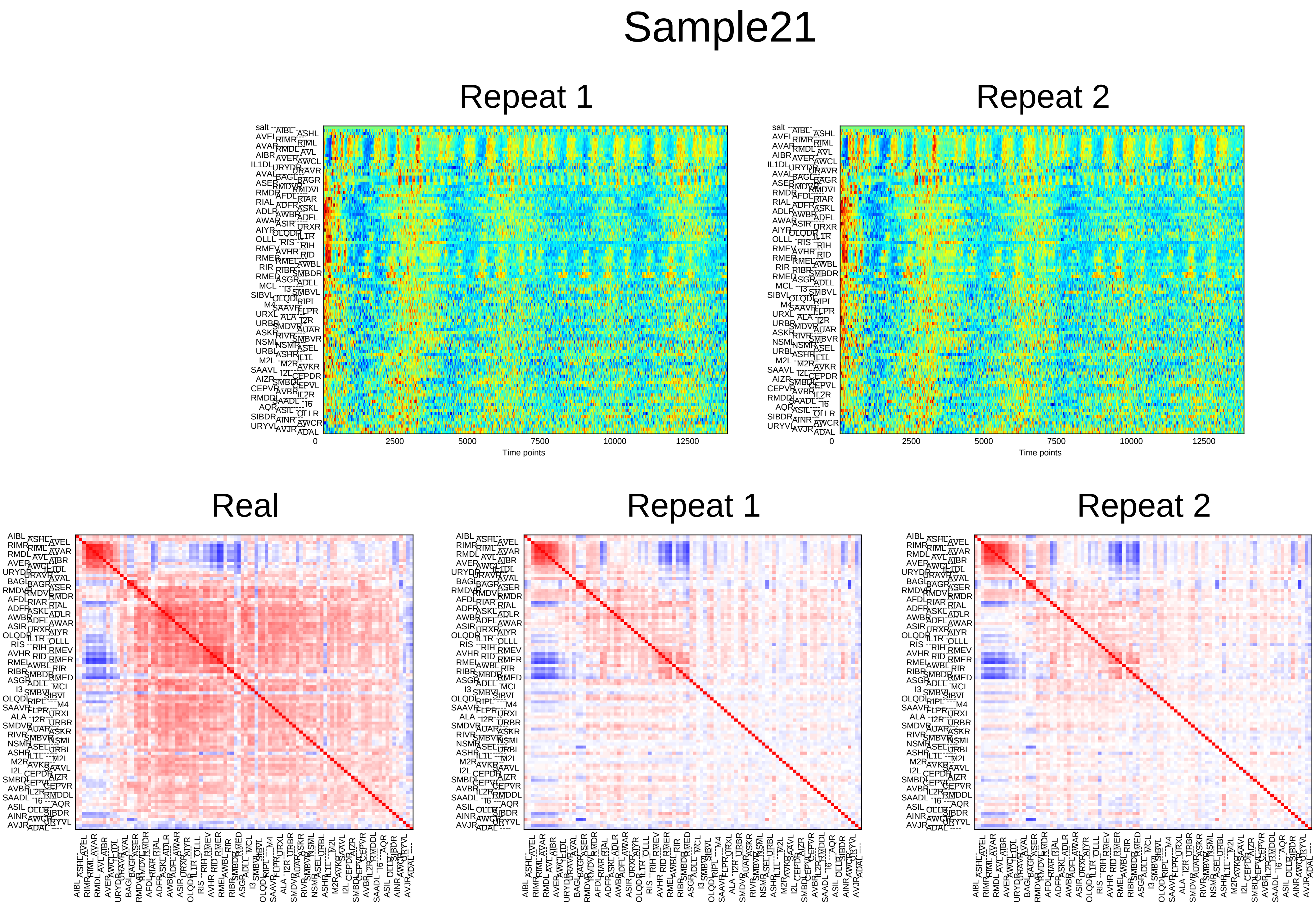
Sample21
Repeat 1
Repeat 2
0
2500
5000
7500
10000
12500
Time points
Real
Repeat 1
Repeat 2

### Sample22

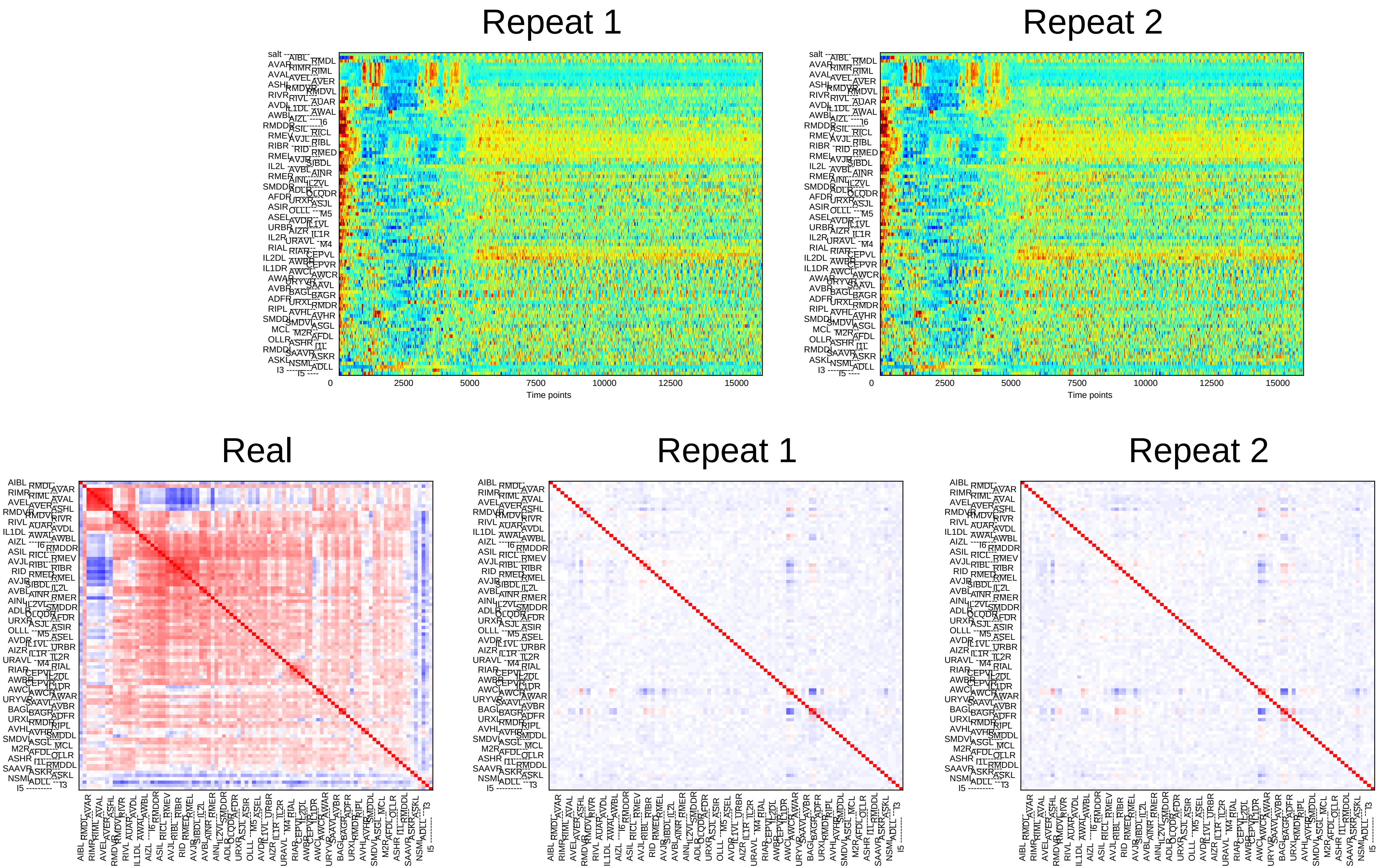

Repeat 1
Repeat 2
Time points
0
2500
5000
7500
10000
12500
15000
Real
Repeat 1
Repeat 2

### Sample23

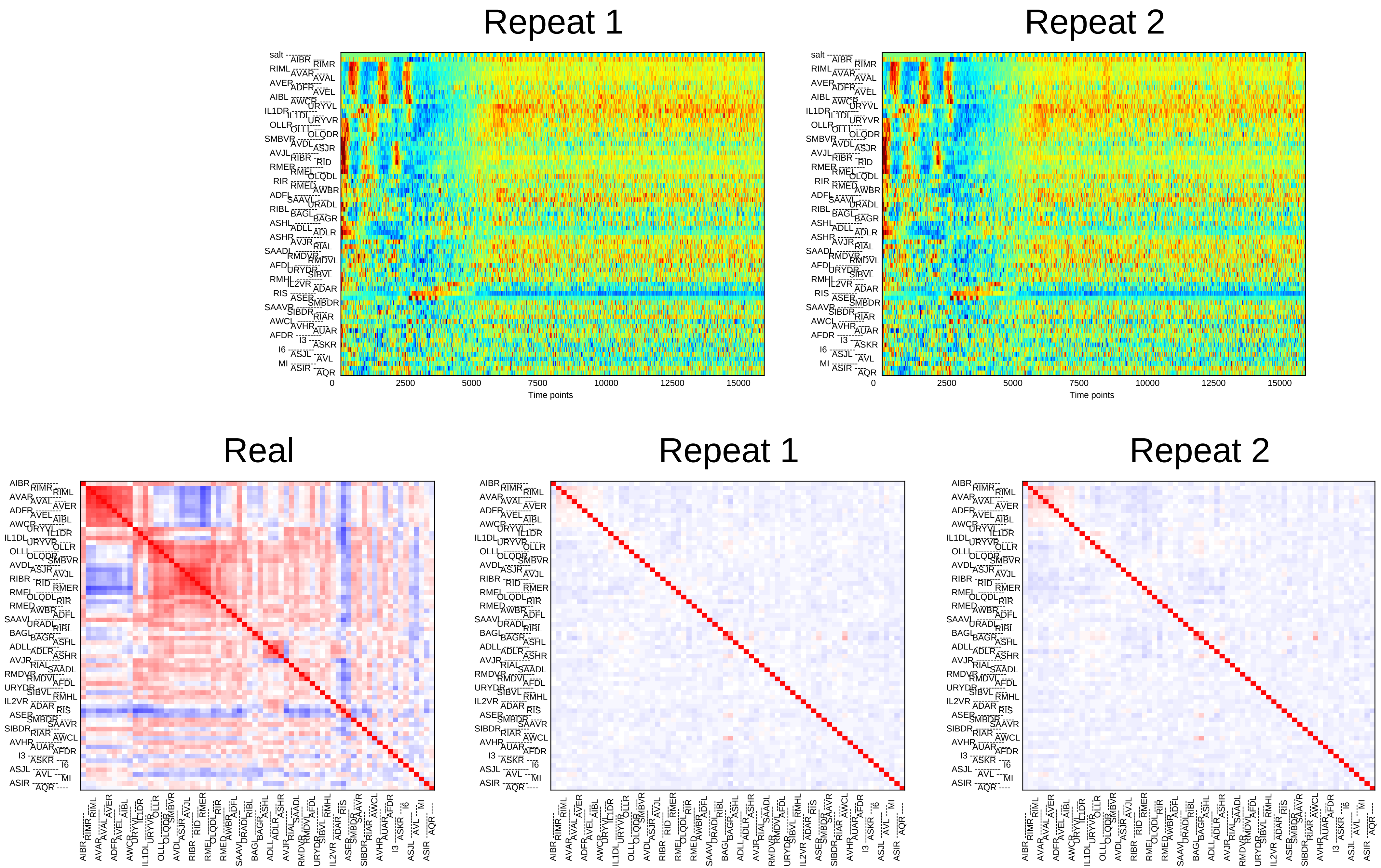
Repeat 1
Repeat 2
0
2500
5000
7500
10000
12500
15000
Time points
Real
Repeat 1
Repeat 2

### Sample24

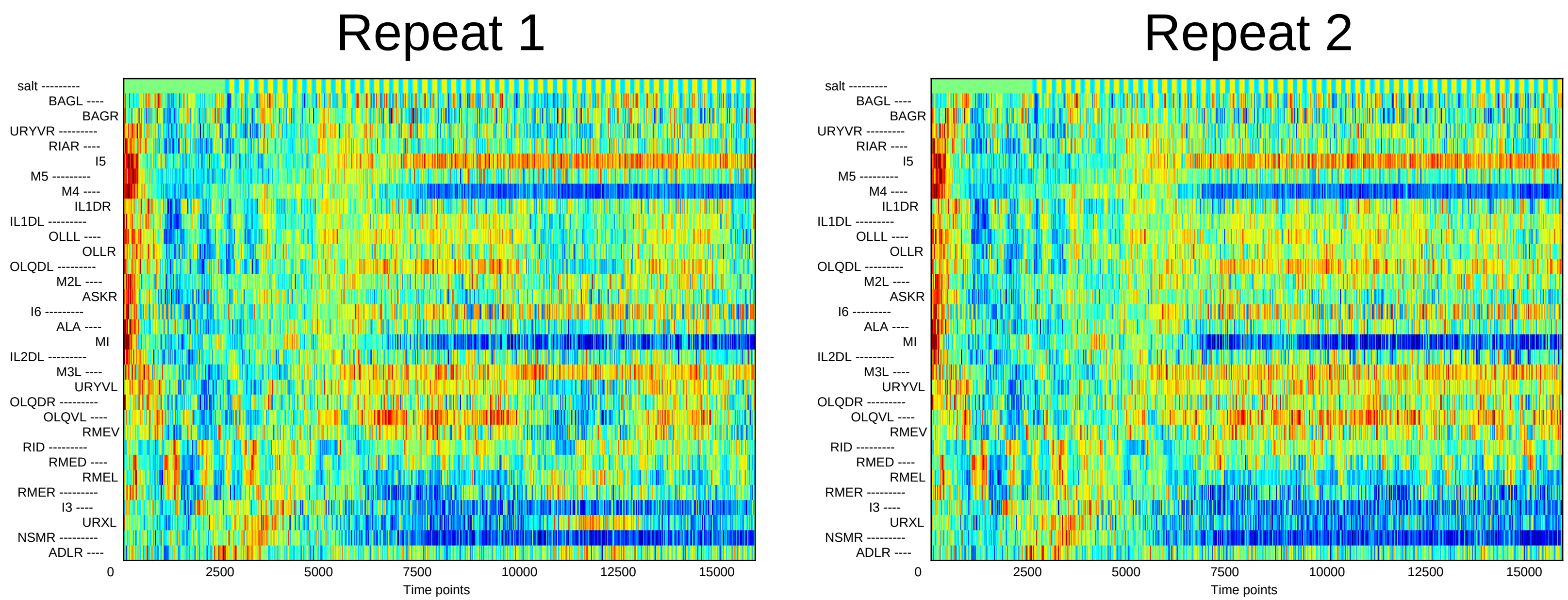
Repeat 1
Repeat 2
salt
BAGL
BAGR
URYVR
RIAR
I5
M5
M4
IL1DR
IL1DL
OLLL
OLLR
OLQDL
M2L
ASKR
I6
ALA
MI
IL2DL
M3L
URYVL
OLQDR
OLQVL
RMEV
RID
RMED
RMEL
RMER
I3
URXL
NSMR
ADLR
0
2500
5000
7500
10000
12500
15000
Time points

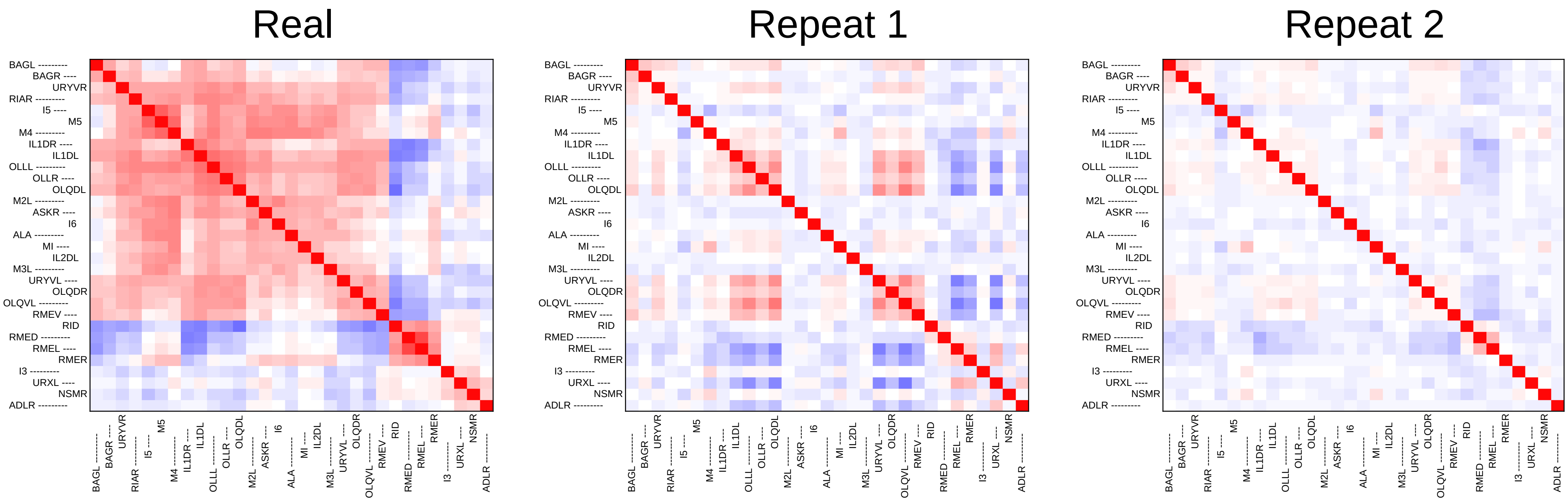
Real
Repeat 1
Repeat 2
BAGL
BAGR
URYVR
RIAR
I5
M5
M4
IL1DR
IL1DL
OLLL
OLLR
OLQDL
M2L
ASKR
I6
ALA
MI
IL2DL
M3L
URYVL
OLQDR
OLQVL
RMEV
RID
RMED
RMEL
RMER
I3
URXL
NSMR
ADLR
