## Supplementary File 5A for "Deducing ensemble dynamics and information flow from the whole-brain imaging data"

### Sample1

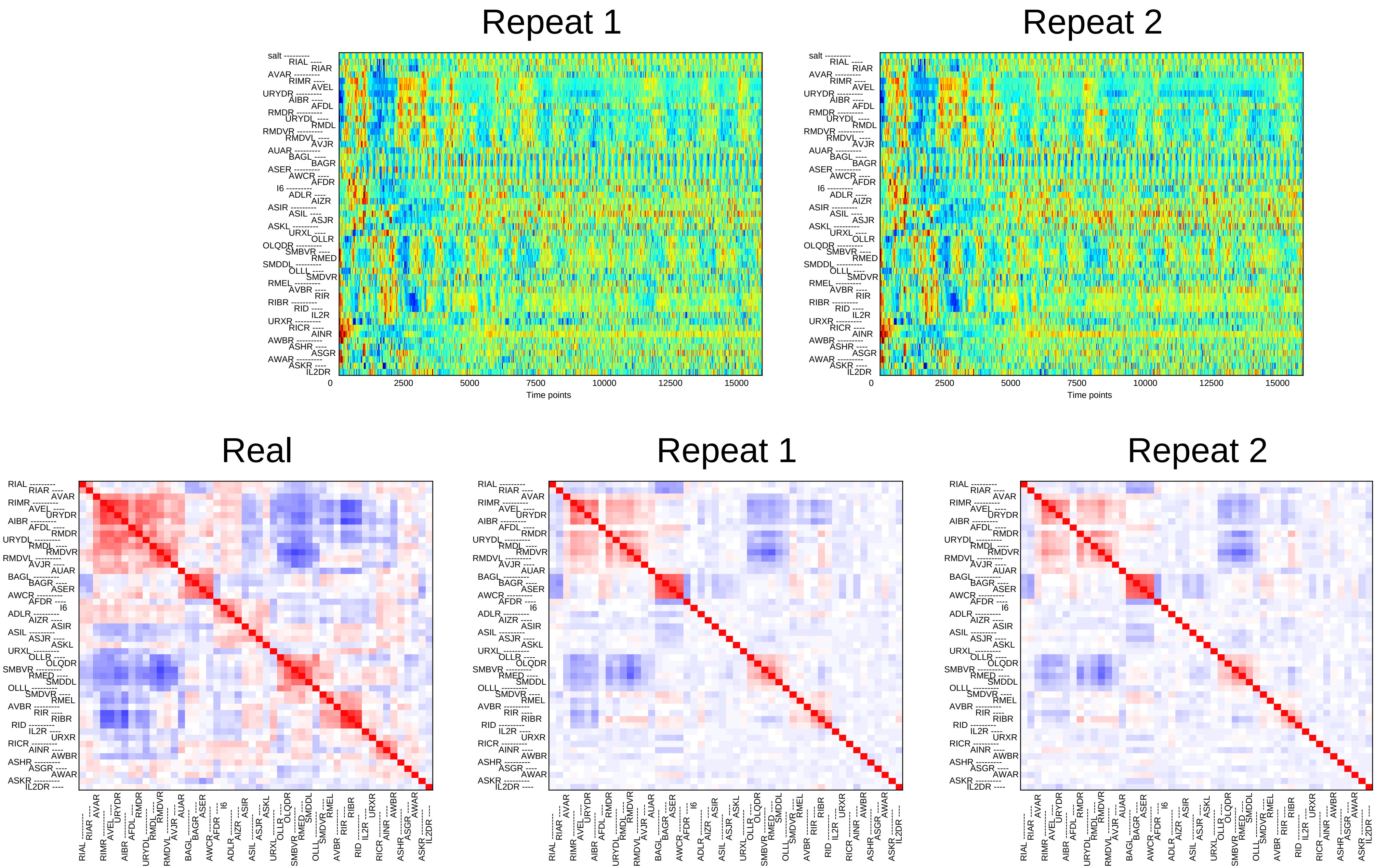

Repeat 1
Repeat 2
salt
RIAL RIAR AVAR RIMR AVEL URYDR AIBR AFDL RMDR URYDL RMDL RMDVR RMDVL AVJR AUAR BAGL BAGR ASER AWCR AFDR I6 ADLR AIZR ASIR ASIL ASJR ASKL URXL OLLR OLQDR SMBVR RMED SMDDL OLLL SMDVR RMEL AVBR RIR RIBR RID IL2R URXR RICR AINR AWBR ASHR ASGR AWAR ASKR IL2DR
0
2500
5000
7500
10000
12500
15000
Time points
Real
Repeat 1
Repeat 2

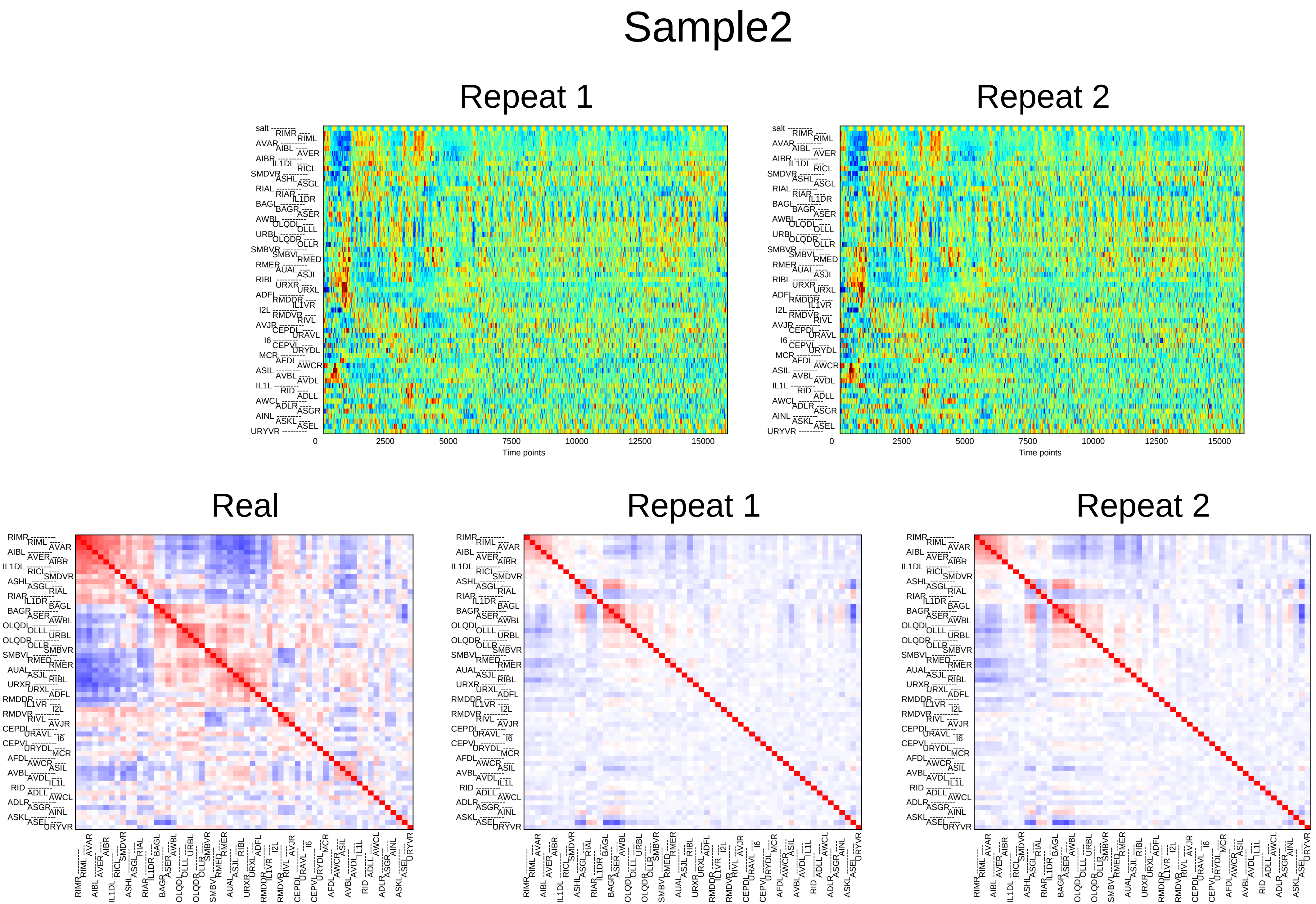
Sample2
Repeat 1
Repeat 2
salt
RIMR RIML AVAR AIBL AVER AIBR IL1DL RICL SMDVR ASHL ASGL RIAL RIAR IL1DR BAGL BAGR ASER AWBL OLQDL OLLL URBL OLQDR OLLR SMBVR SMBVL RMED RMER AUAL ASJL RIBL URXR URXL ADFL RMDDR IL1VR I2L RMDVR RIVL AVJR CEPDL URAVL I6 CEPVL URYDL MCR AFDL AWCR ASIL AVBL AVDL IL1L RID ADLL AWCL ADLR ASGR AINL ASKL ASEL URYVR
0
2500
5000
7500
10000
12500
15000
Time points
Real

### Sample3

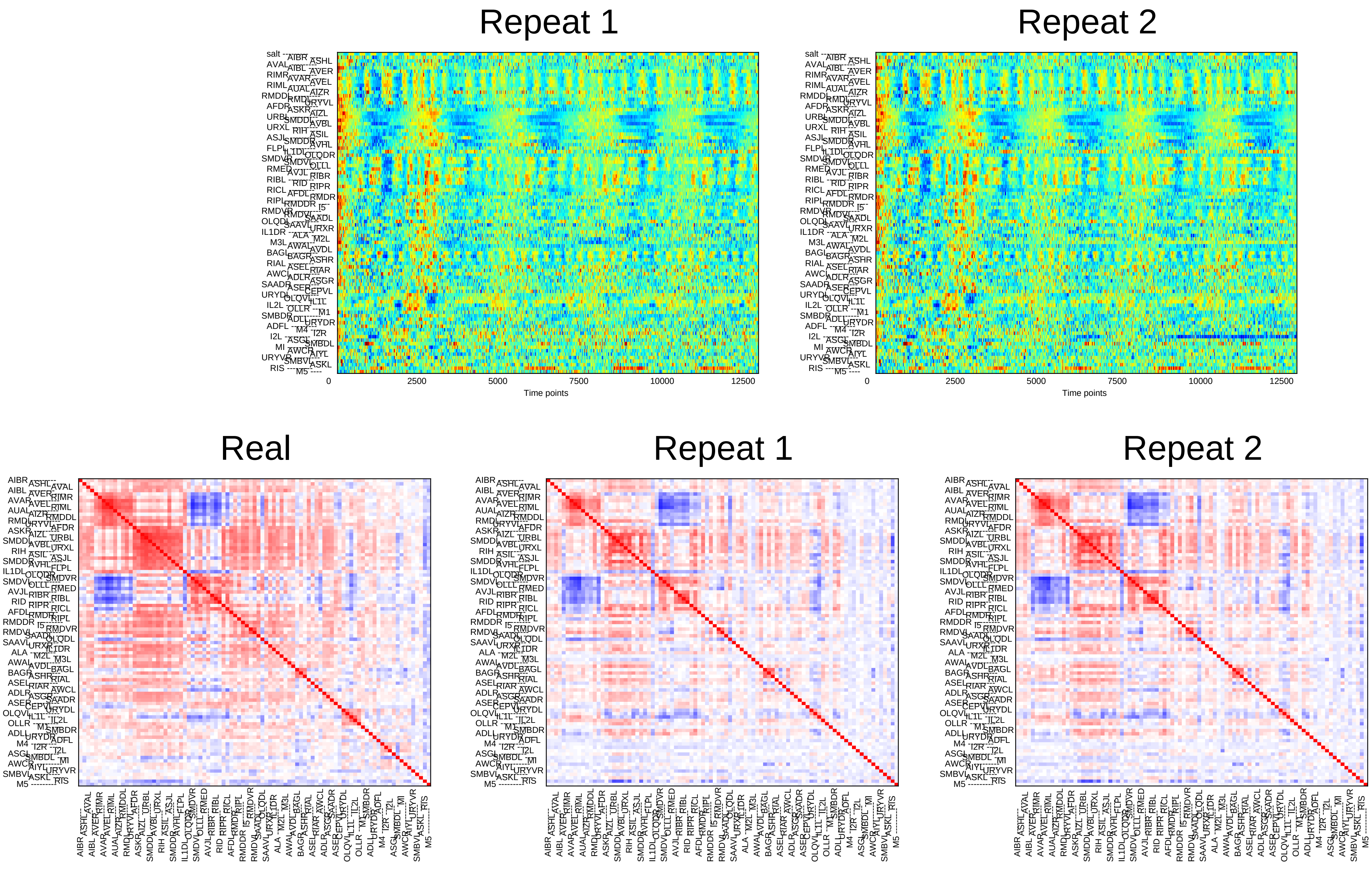
Repeat 1
Repeat 2
0
2500
5000
7500
10000
12500
Time points
Real
Repeat 1
Repeat 2

### Sample4

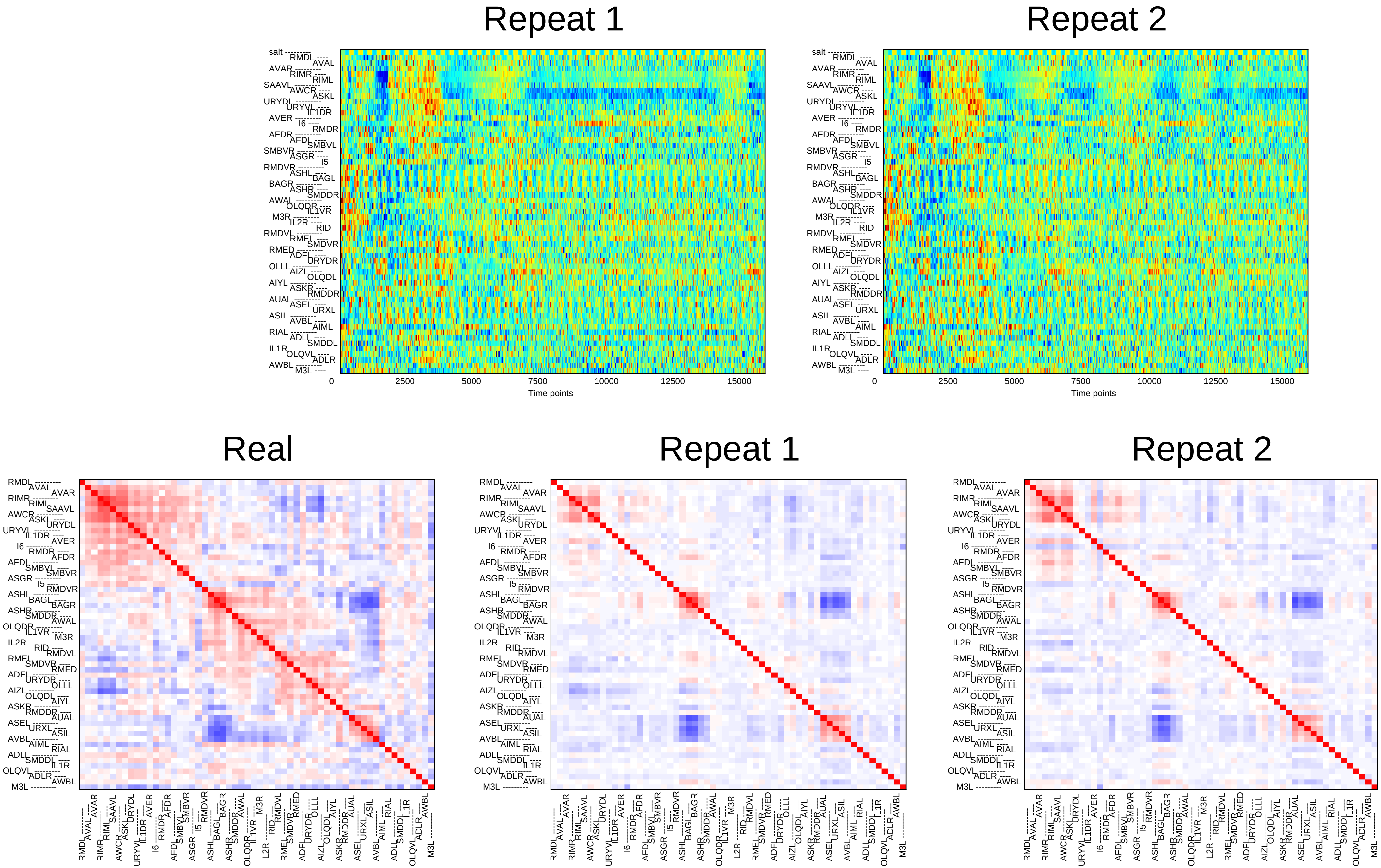

Repeat 1
Repeat 2
Time points
0
2500
5000
7500
10000
12500
15000
Real
Repeat 1
Repeat 2

### Sample5

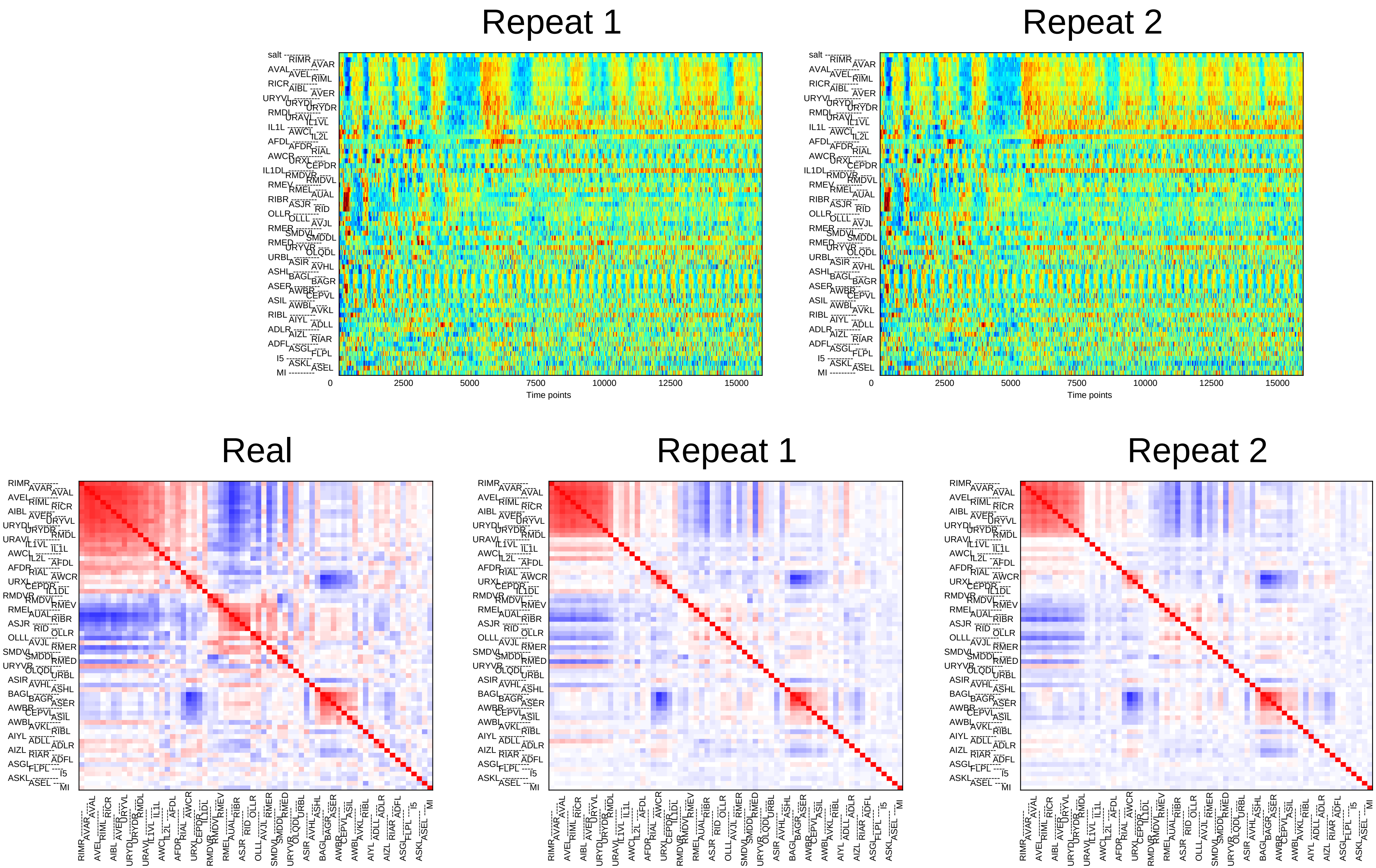

Repeat 1
Repeat 2
0
2500
5000
7500
10000
12500
15000
Time points
Real
Repeat 1
Repeat 2

### Sample6

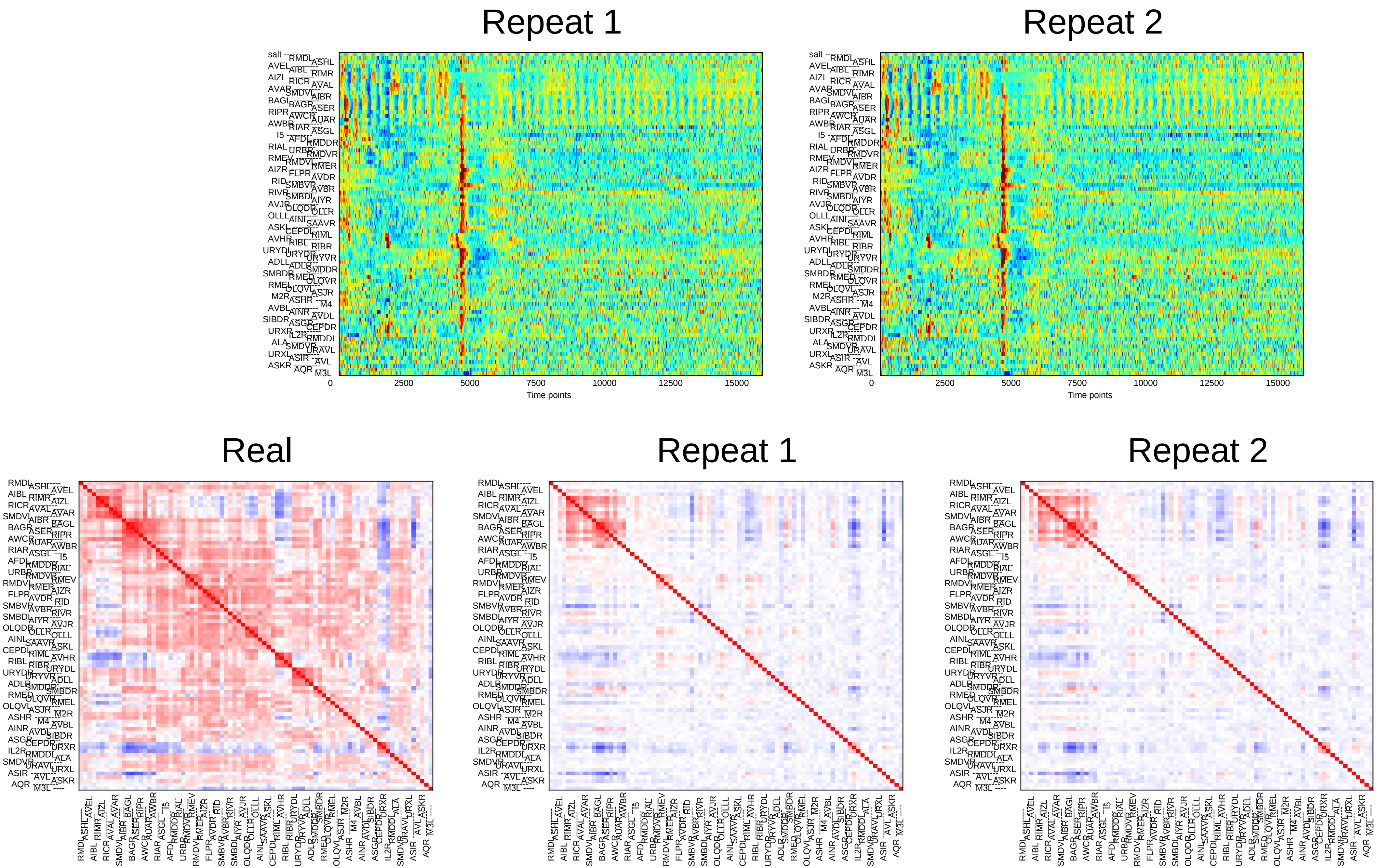
Repeat 1
Repeat 2
0
2500
5000
7500
10000
12500
15000
Time points
Real
Repeat 1
Repeat 2

### Sample7

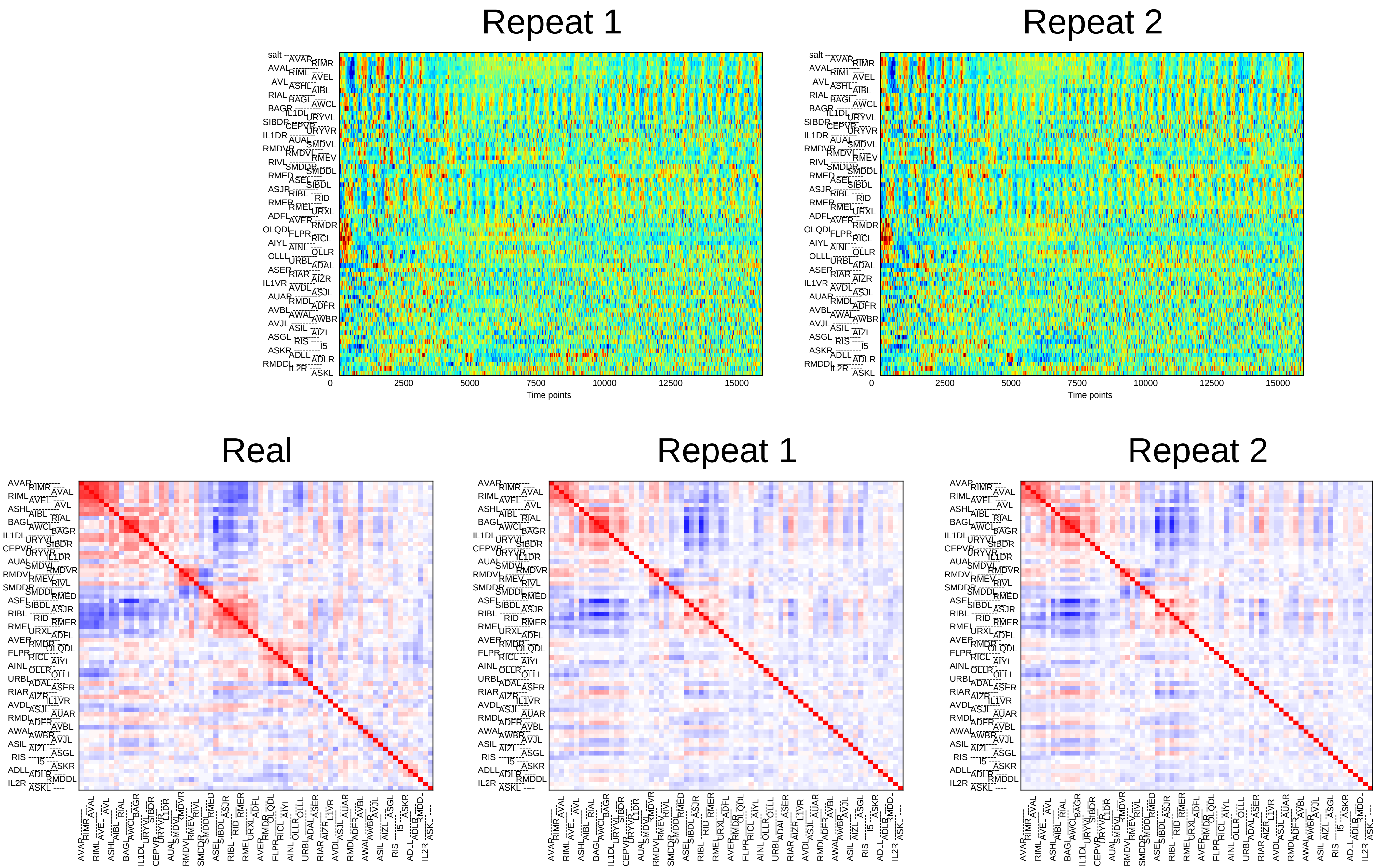
Repeat 1
Repeat 2
0
2500
5000
7500
10000
12500
15000
Time points
Real
Repeat 1
Repeat 2

### Sample8

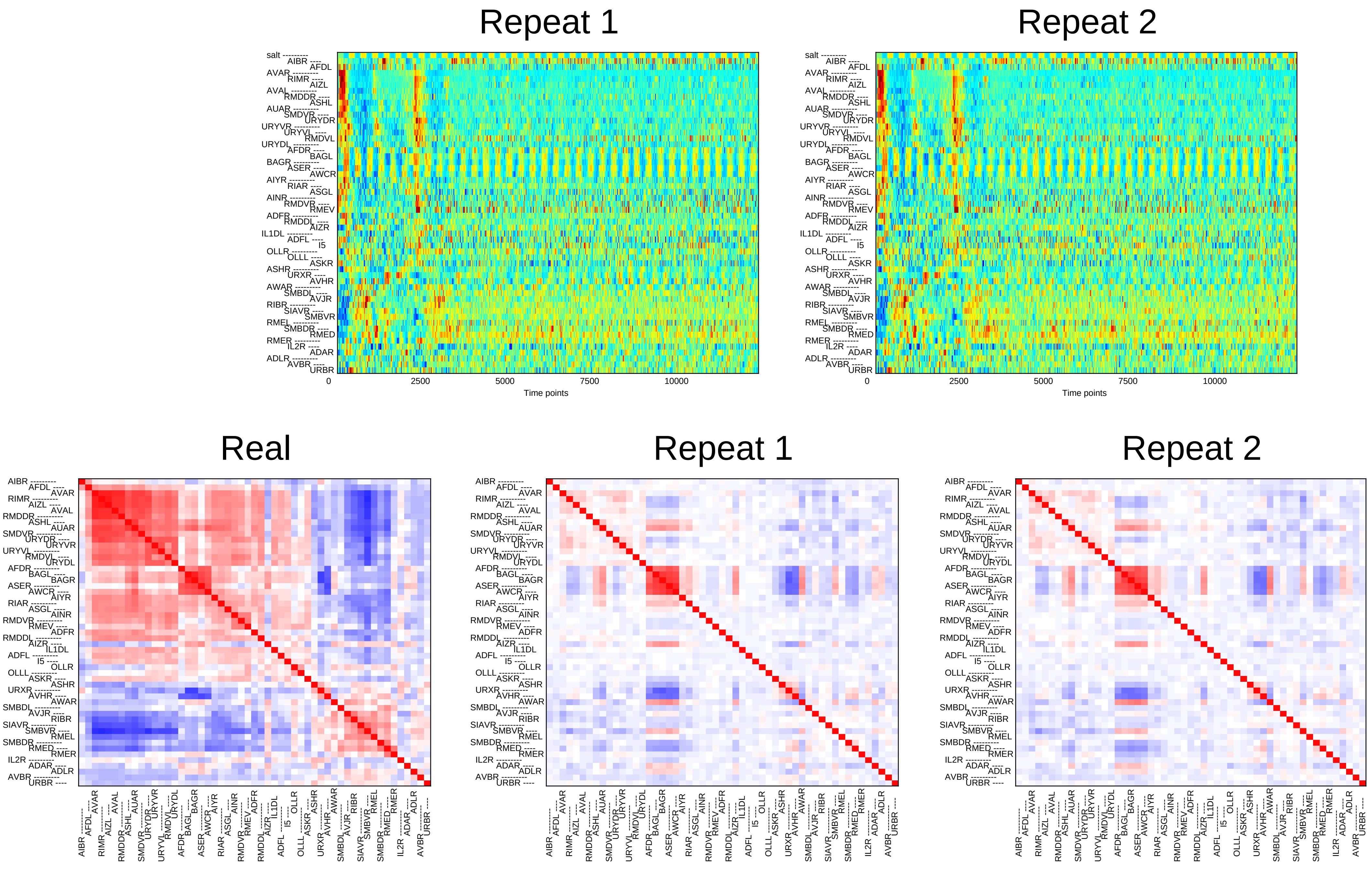

Repeat 1
Repeat 2
0
2500
5000
7500
10000
Time points
Real
Repeat 1
Repeat 2

### Sample9

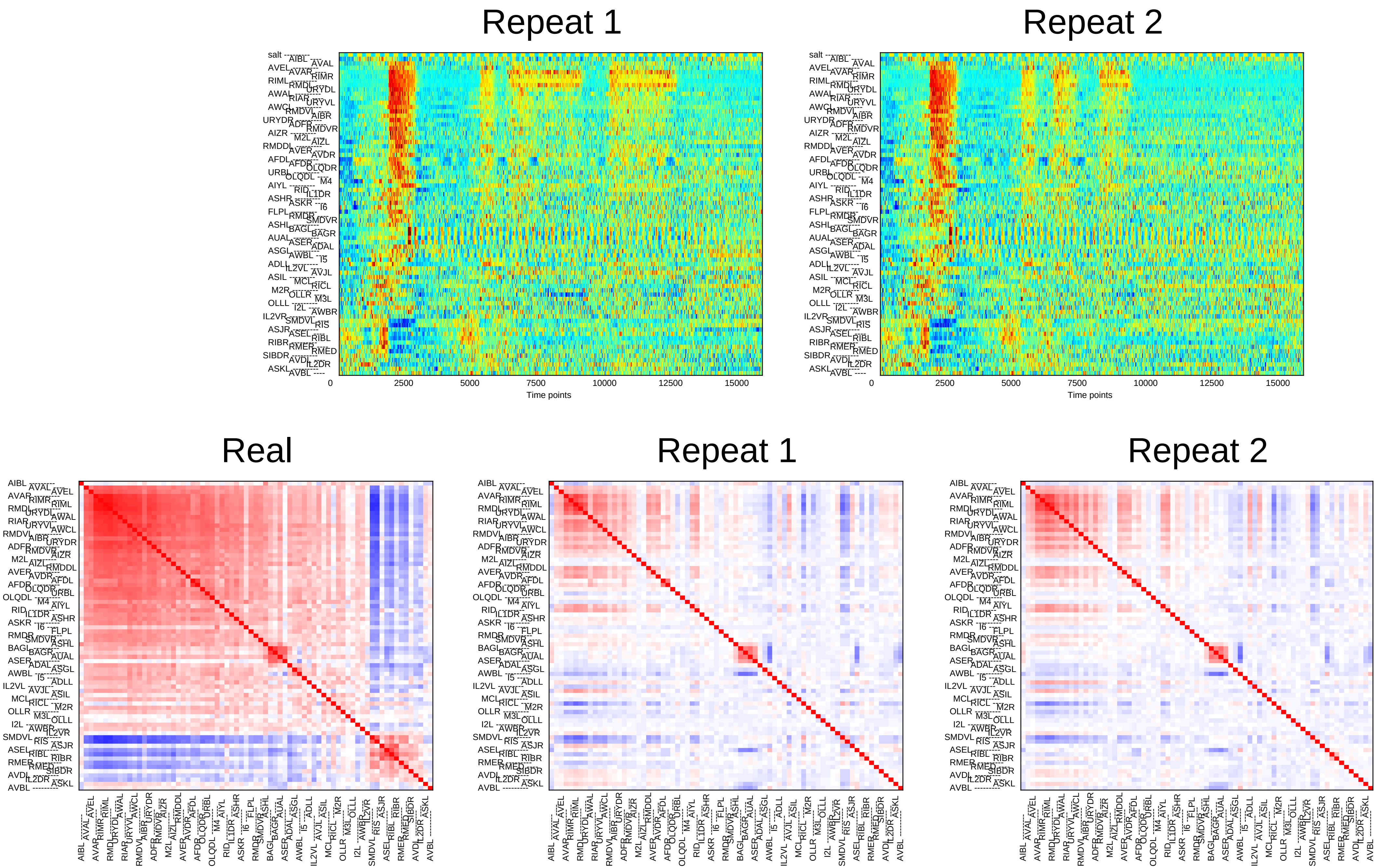

Repeat 1
Repeat 2
0
2500
5000
7500
10000
12500
15000
Time points
Real
Repeat 1
Repeat 2

### Sample10

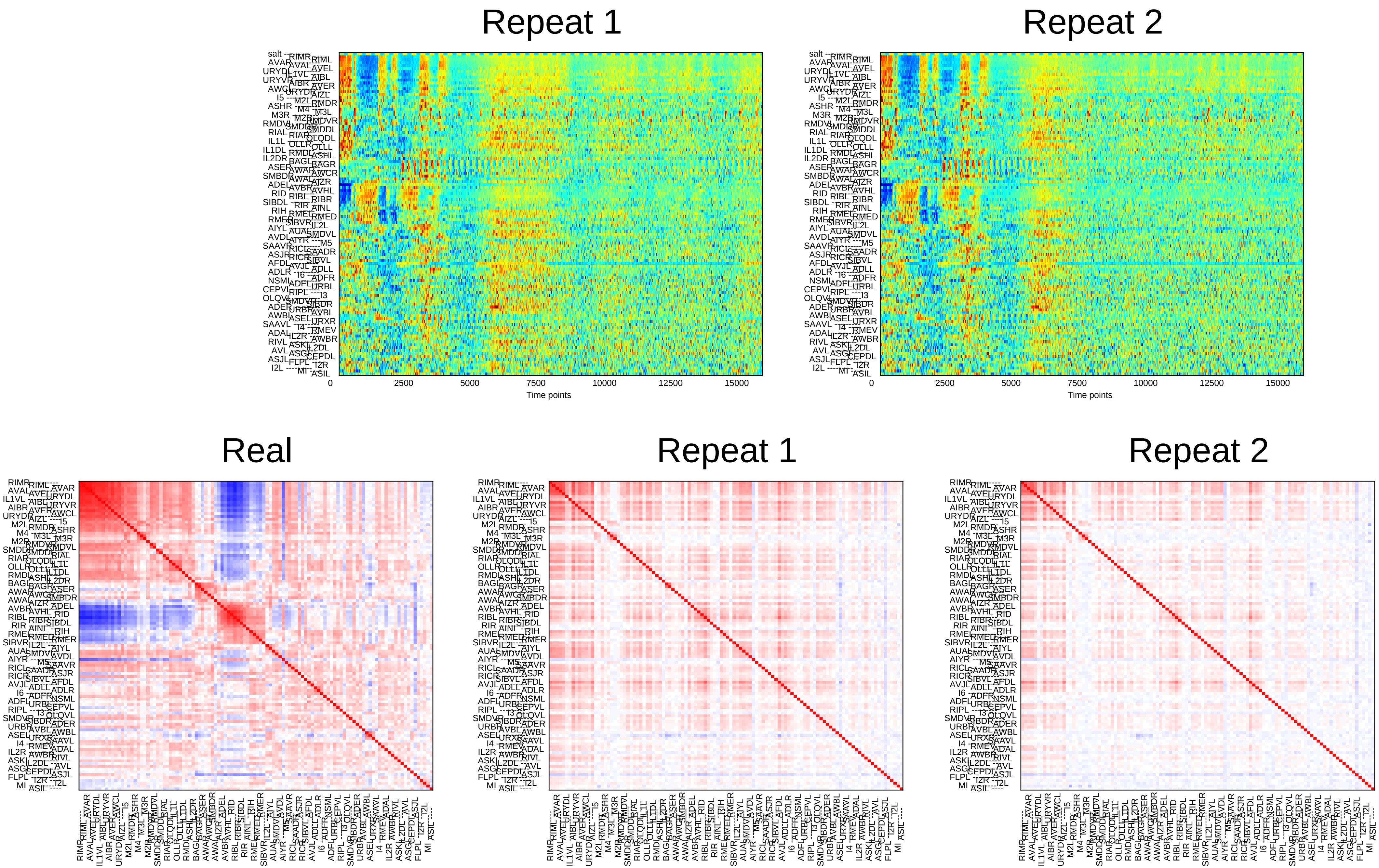
Repeat 1
Repeat 2
0
2500
5000
7500
10000
12500
15000
Time points
Real
Repeat 1
Repeat 2

### Sample11

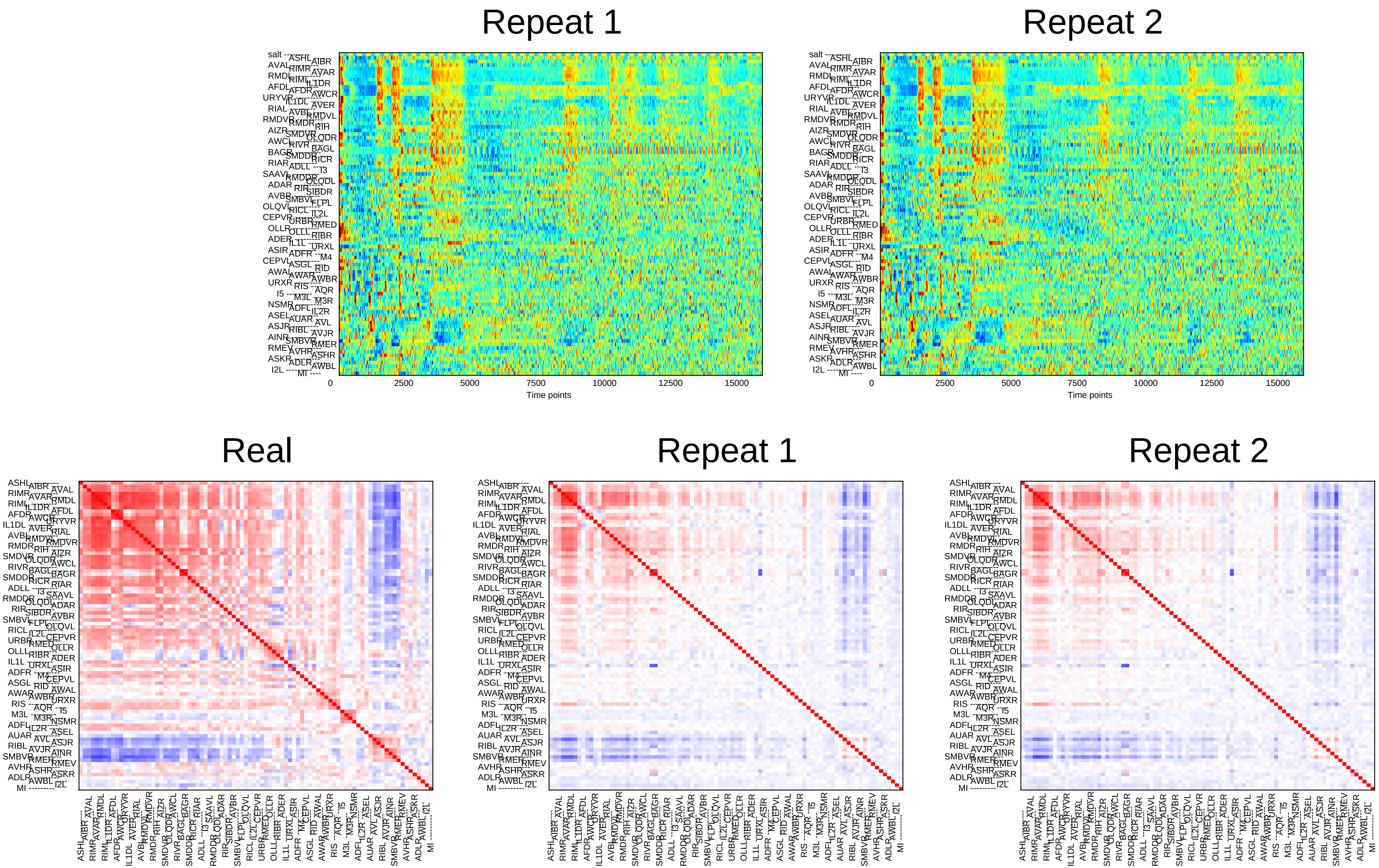
Repeat 1
Repeat 2
0
2500
5000
7500
10000
12500
15000
Time points
Real
Repeat 1
Repeat 2

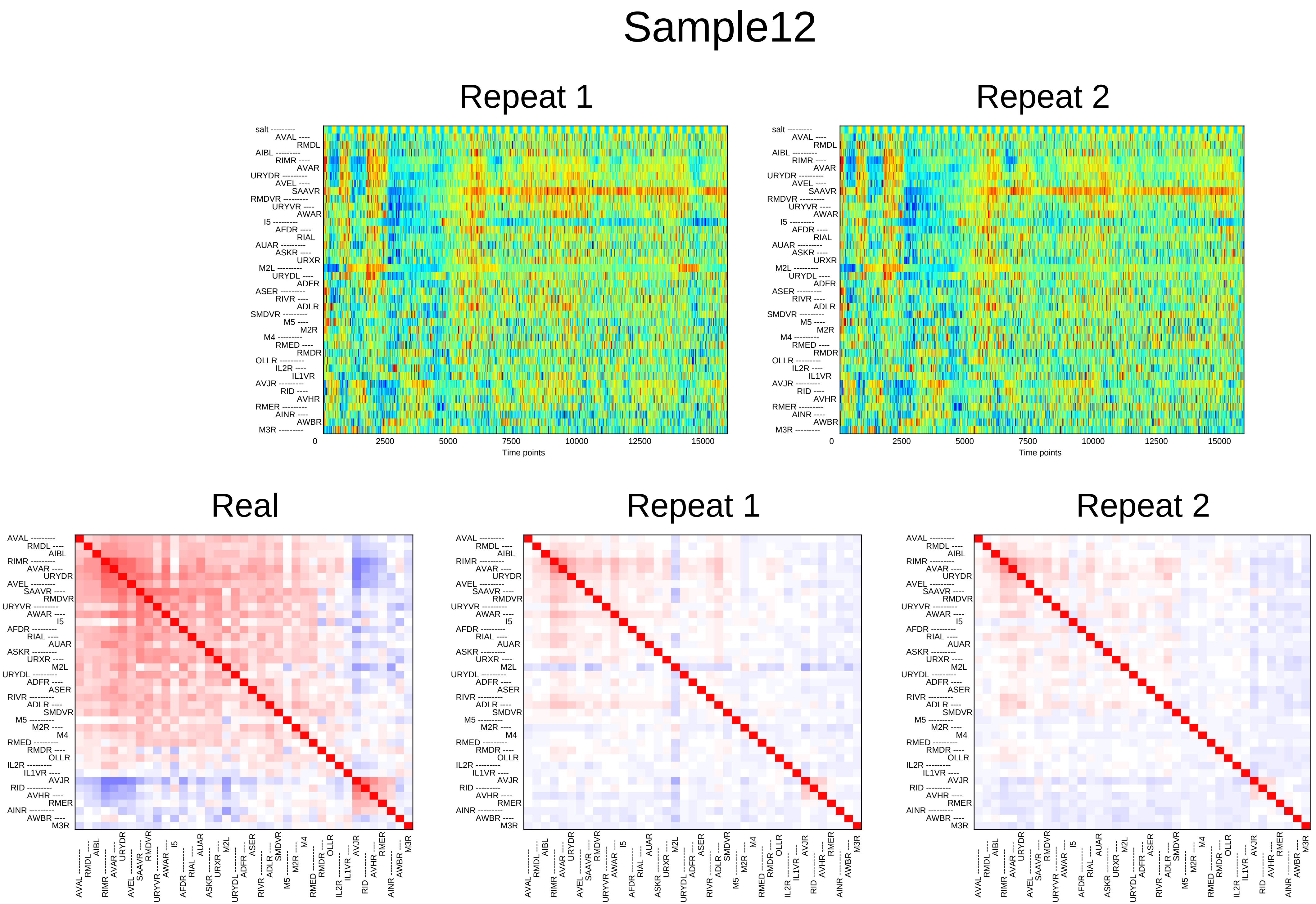

Sample12
Repeat 1
Repeat 2
salt
AVAL
RMDL
AIBL
RIMR
AVAR
URYDR
AVEL
SAAVR
RMDVR
URYVR
AWAR
I5
AFDR
RIAL
AUAR
ASKR
URXR
M2L
URYDL
ADFR
ASER
RIVR
ADLR
SMDVR
M5
M2R
M4
RMED
RMDR
OLLR
IL2R
IL1VR
AVJR
RID
AVHR
RMER
AINR
AWBR
M3R
0
2500
5000
7500
10000
12500
15000
Time points
Real
Repeat 1
Repeat 2

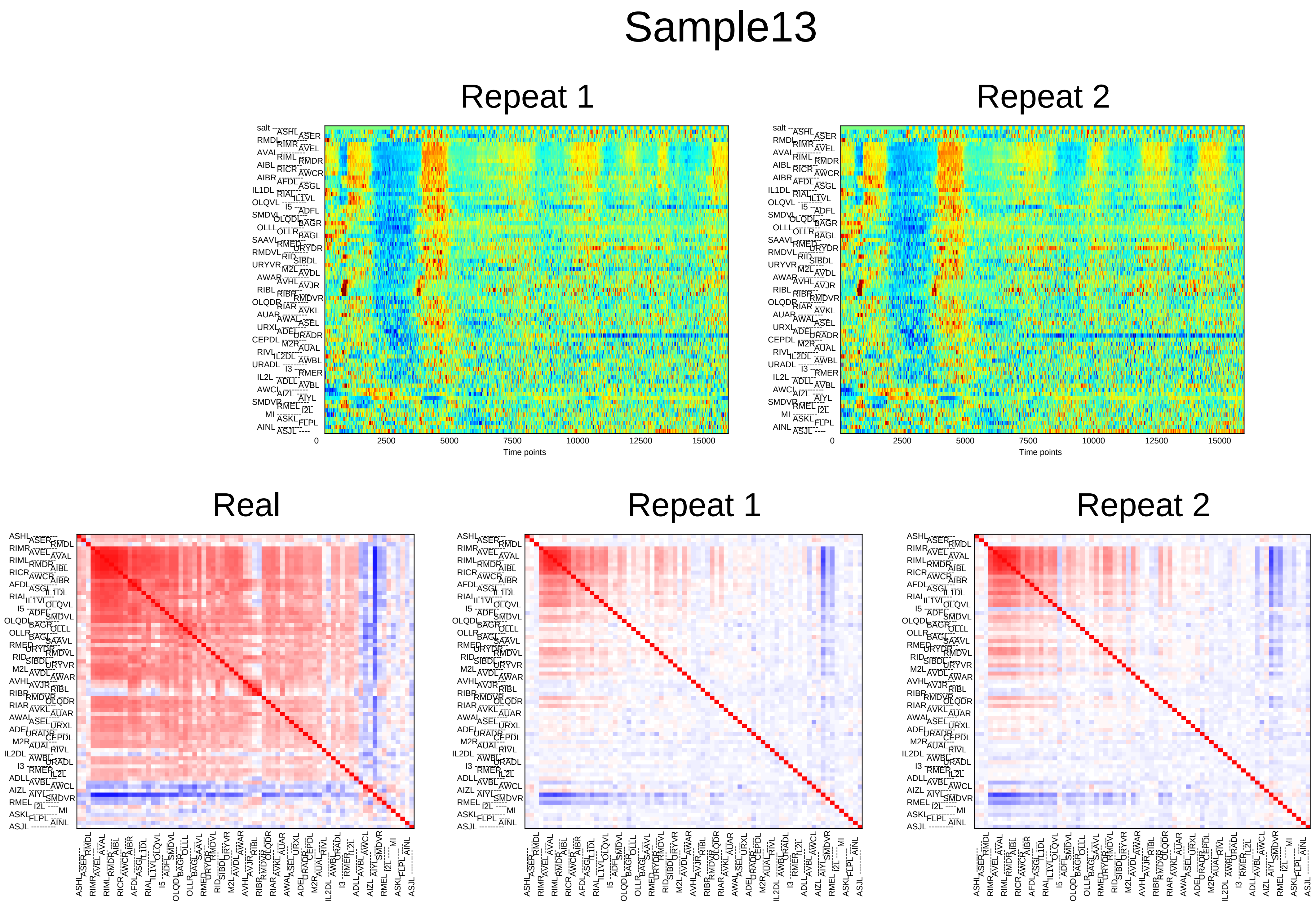

Sample13
Repeat 1
Repeat 2
0
2500
5000
7500
10000
12500
15000
Time points
Real
Repeat 1
Repeat 2

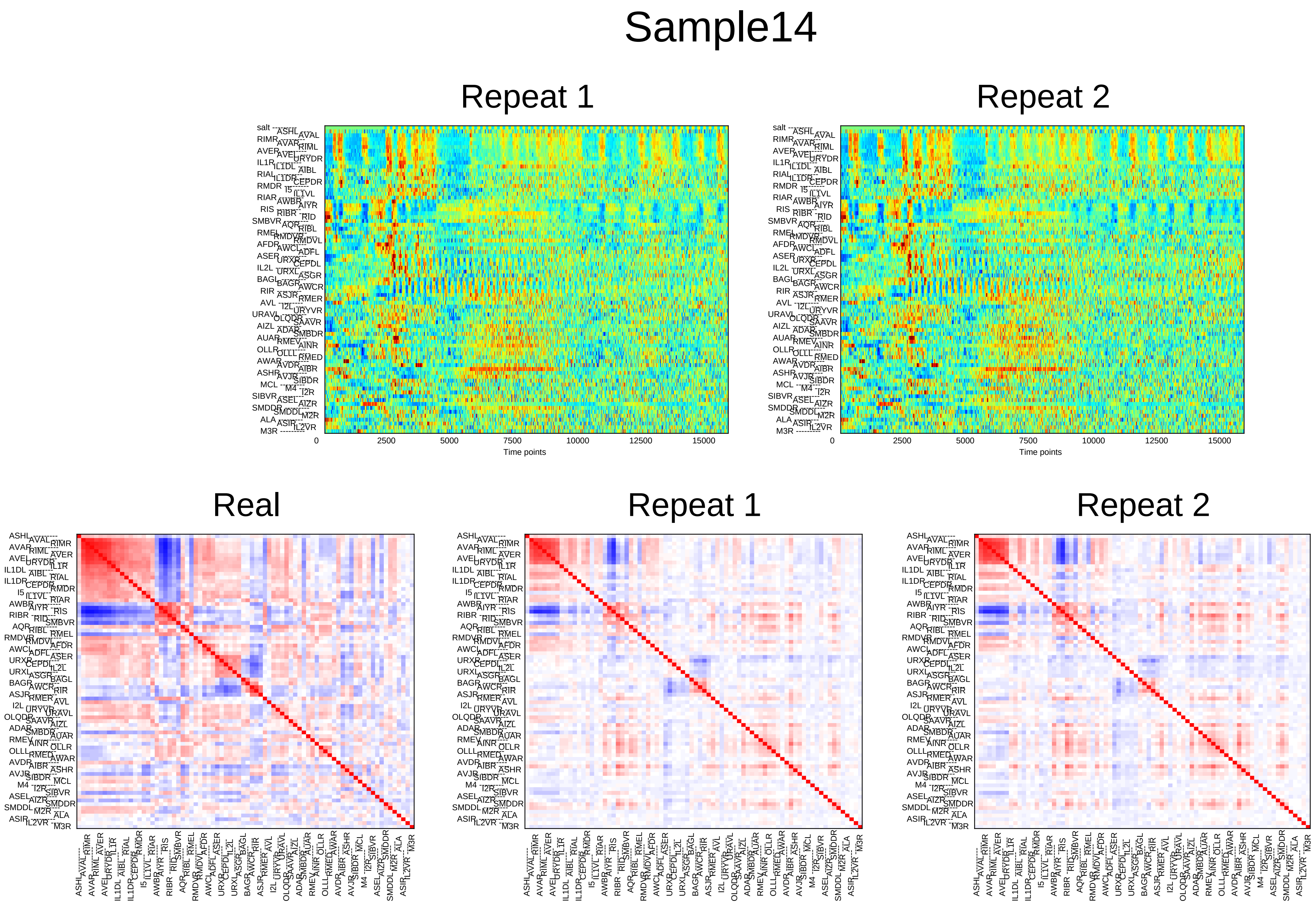
Sample14
Repeat 1
Repeat 2
0
2500
5000
7500
10000
12500
15000
Time points
Real
Repeat 1
Repeat 2

### Sample15

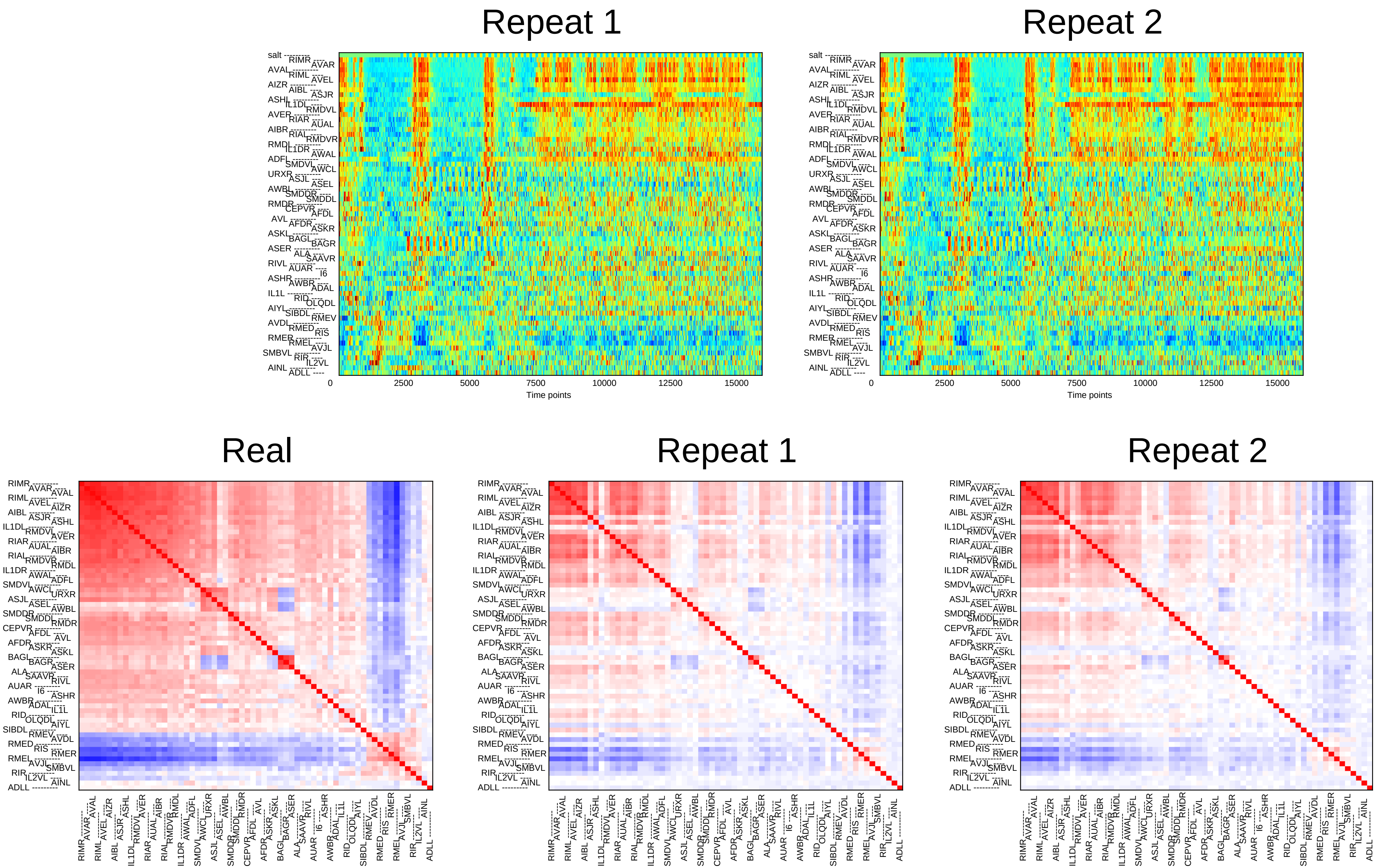

Repeat 1
Repeat 2
0
2500
5000
7500
10000
12500
15000
Time points
Real
Repeat 1
Repeat 2

### Sample16

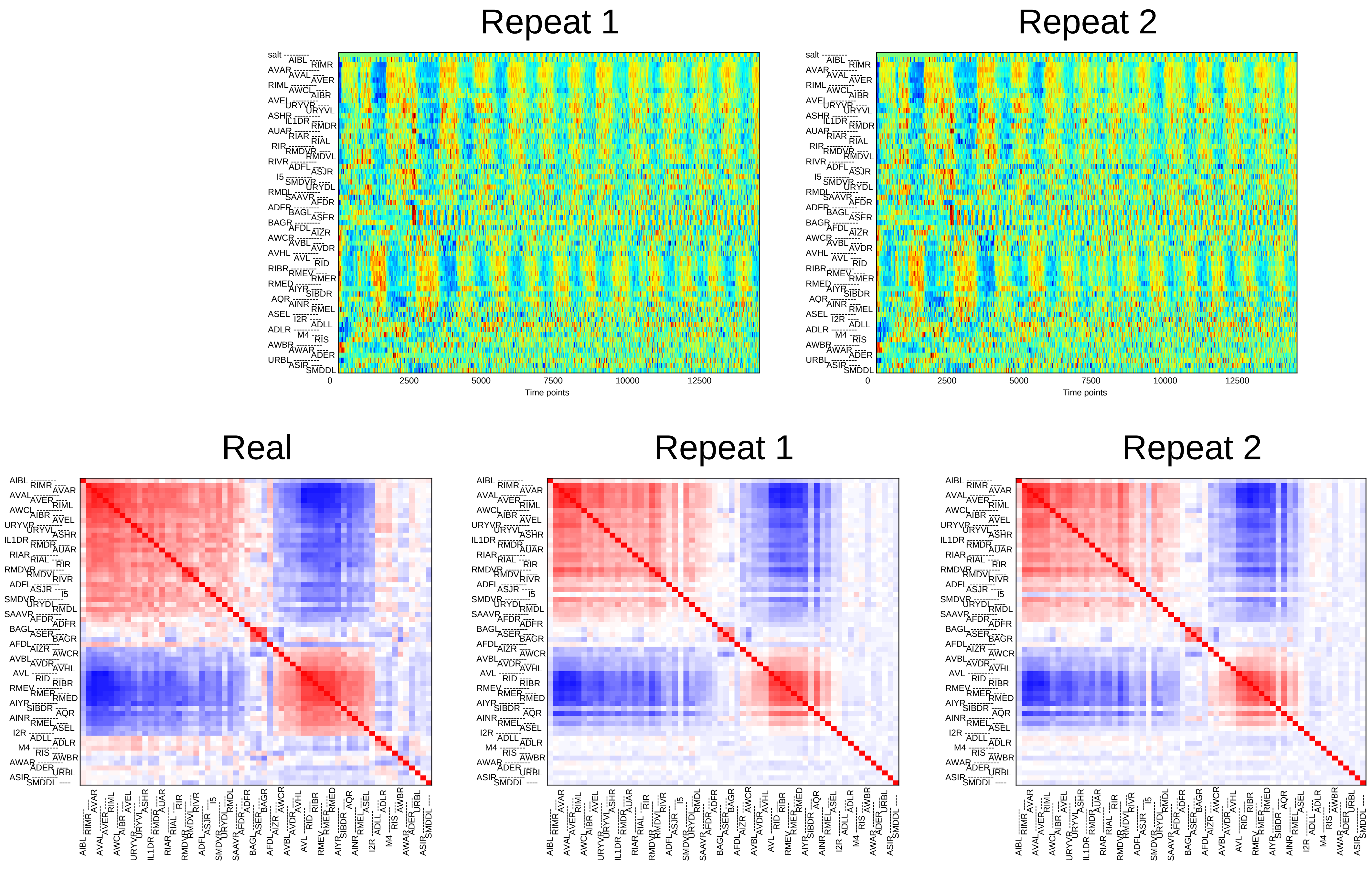

Repeat 1
Repeat 2
0
2500
5000
7500
10000
12500
Time points
Real
Repeat 1
Repeat 2

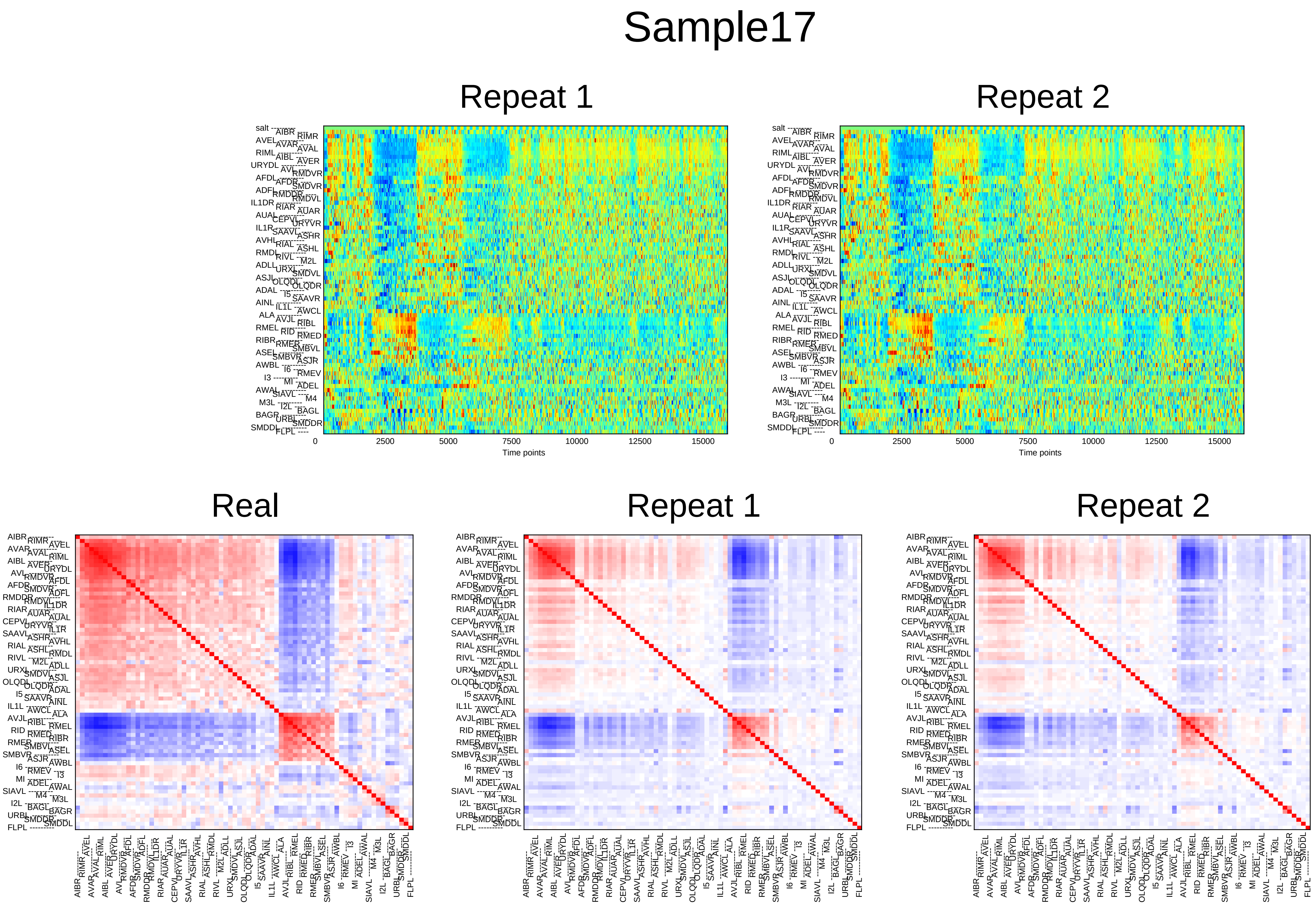
Sample17
Repeat 1
Repeat 2
0
2500
5000
7500
10000
12500
15000
Time points
Real
Repeat 1
Repeat 2

### Sample18

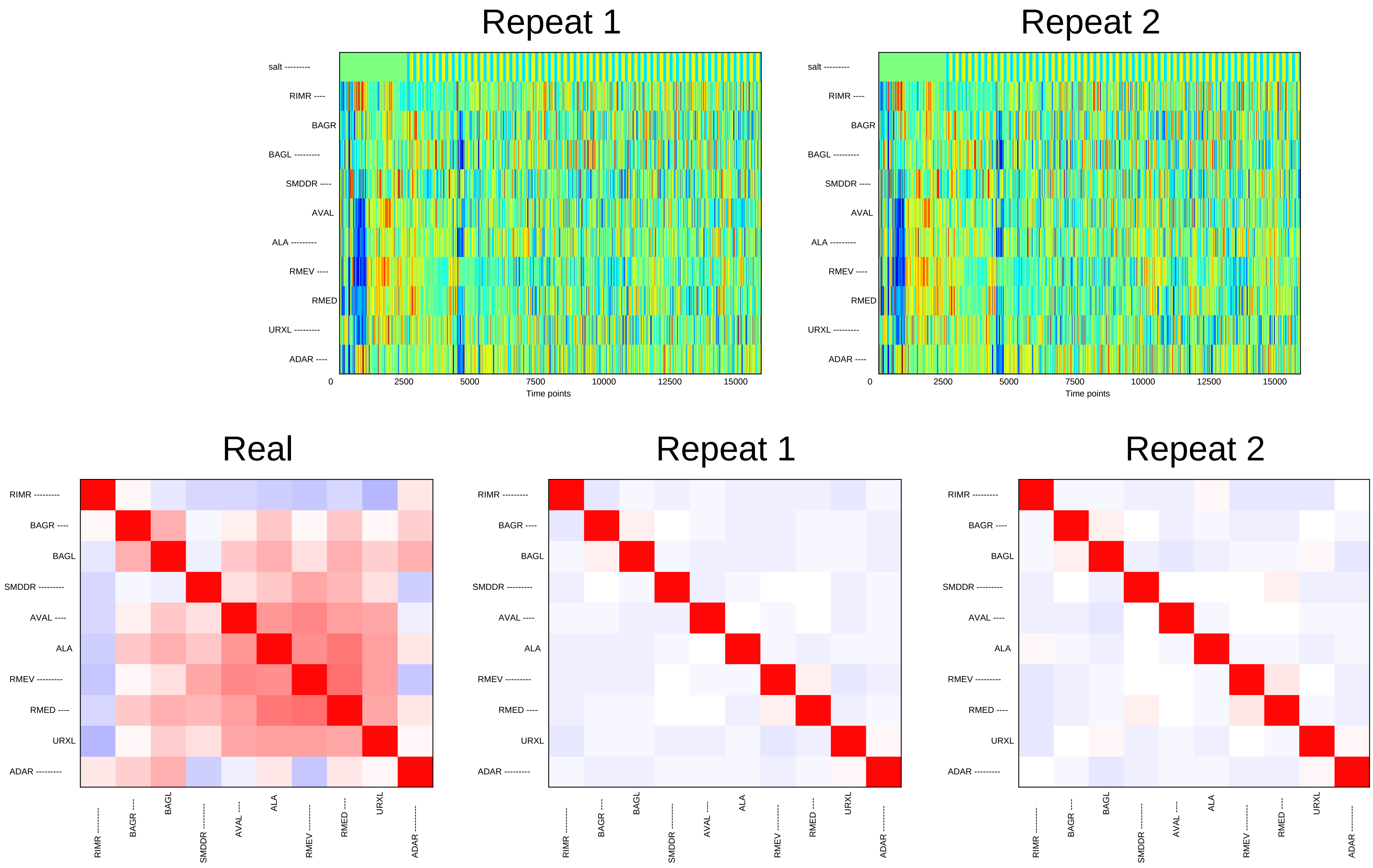

Repeat 1
Repeat 2
salt ---------
RIMR ----
BAGR
BAGL ---------
SMDDR ----
AVAL
ALA ---------
RMEV ----
RMED
URXL ---------
ADAR ----
0
2500
5000
7500
10000
12500
15000
Time points
Real
Repeat 1
Repeat 2
RIMR ---------
BAGR ----
BAGL
SMDDR ---------
AVAL ----
ALA
RMEV ---------
RMED ----
URXL
ADAR ---------

### Sample19

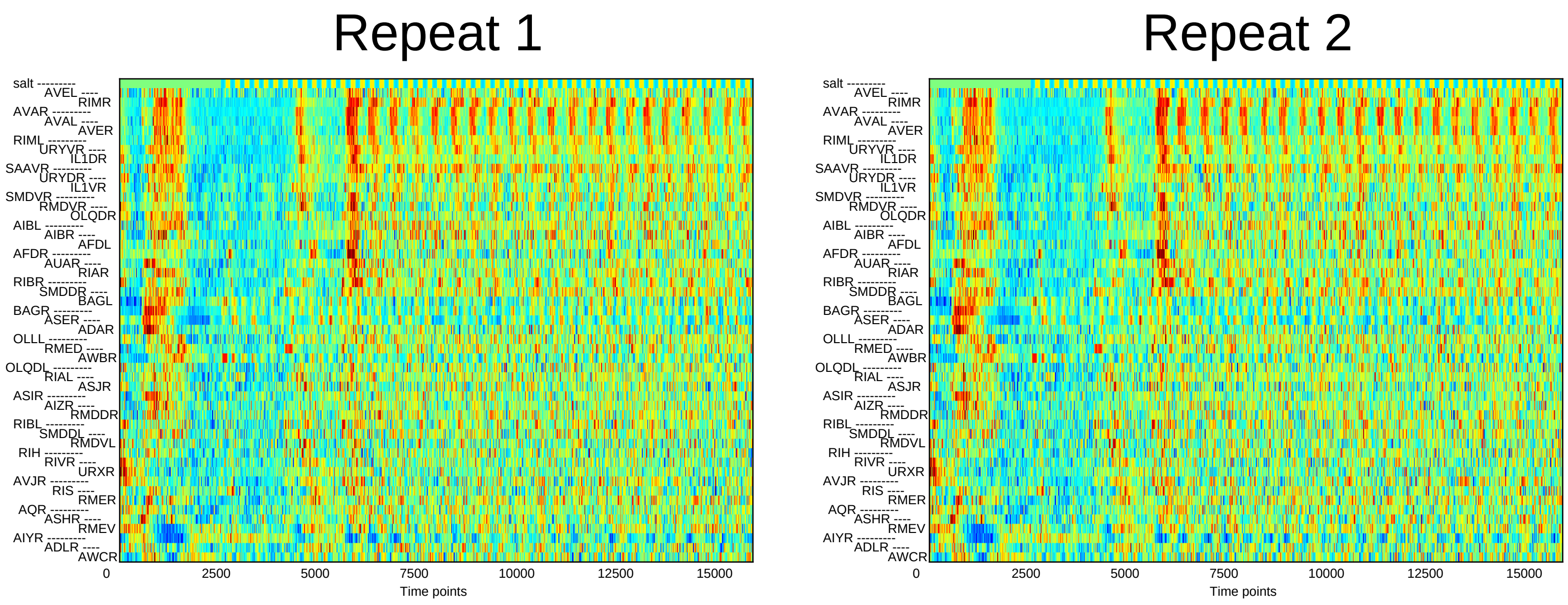
Repeat 1
Repeat 2
salt
AVEL
RIMR
AVAR
AVAL
AVER
RIML
URYVR
IL1DR
SAAVR
URYDR
IL1VR
SMDVR
RMDVR
OLQDR
AIBL
AIBR
AFDL
AFDR
AUAR
RIAR
RIBR
SMDDR
BAGL
BAGR
ASER
ADAR
OLLL
RMED
AWBR
OLQDL
RIAL
ASJR
ASIR
AIZR
RMDDR
RIBL
SMDDL
RMDVL
RIH
RIVR
URXR
AVJR
RIS
RMER
AQR
ASHR
RMEV
AIYR
ADLR
AWCR
0
2500
5000
7500
10000
12500
15000
Time points

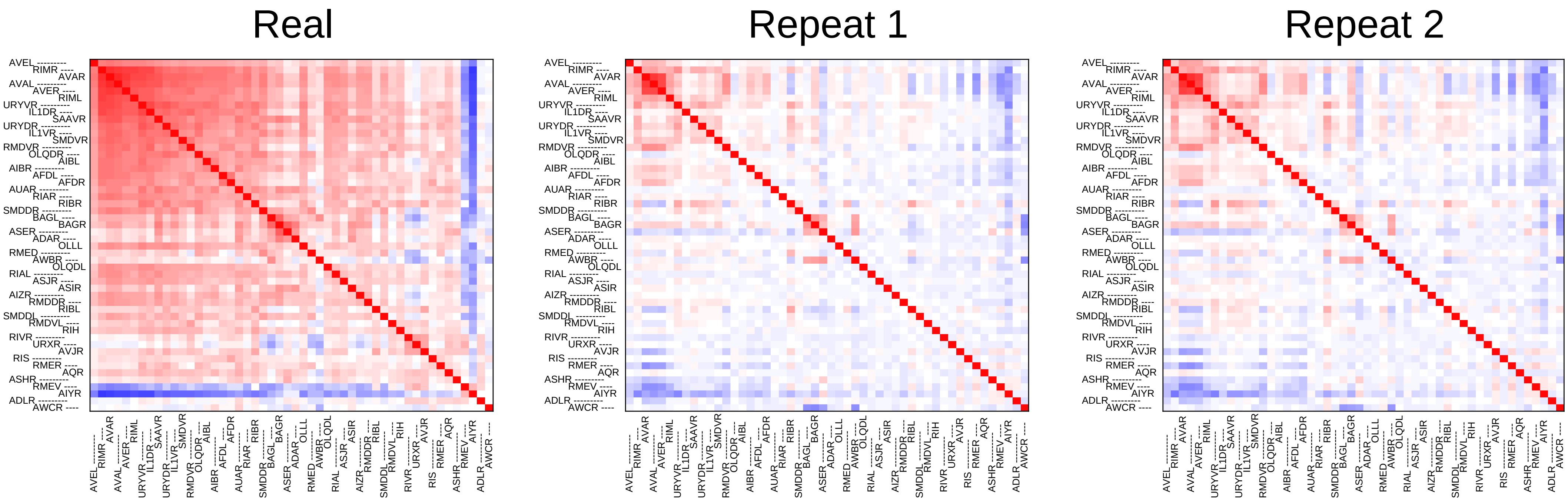
Real
Repeat 1
Repeat 2
AVEL
RIMR
AVAR
AVAL
AVER
RIML
URYVR
IL1DR
SAAVR
URYDR
IL1VR
SMDVR
RMDVR
OLQDR
AIBL
AIBR
AFDL
AFDR
AUAR
RIAR
RIBR
SMDDR
BAGL
BAGR
ASER
ADAR
OLLL
RMED
AWBR
OLQDL
RIAL
ASJR
ASIR
AIZR
RMDDR
RIBL
SMDDL
RMDVL
RIH
RIVR
URXR
AVJR
RIS
RMER
AQR
ASHR
RMEV
AIYR
ADLR
AWCR

### Sample20

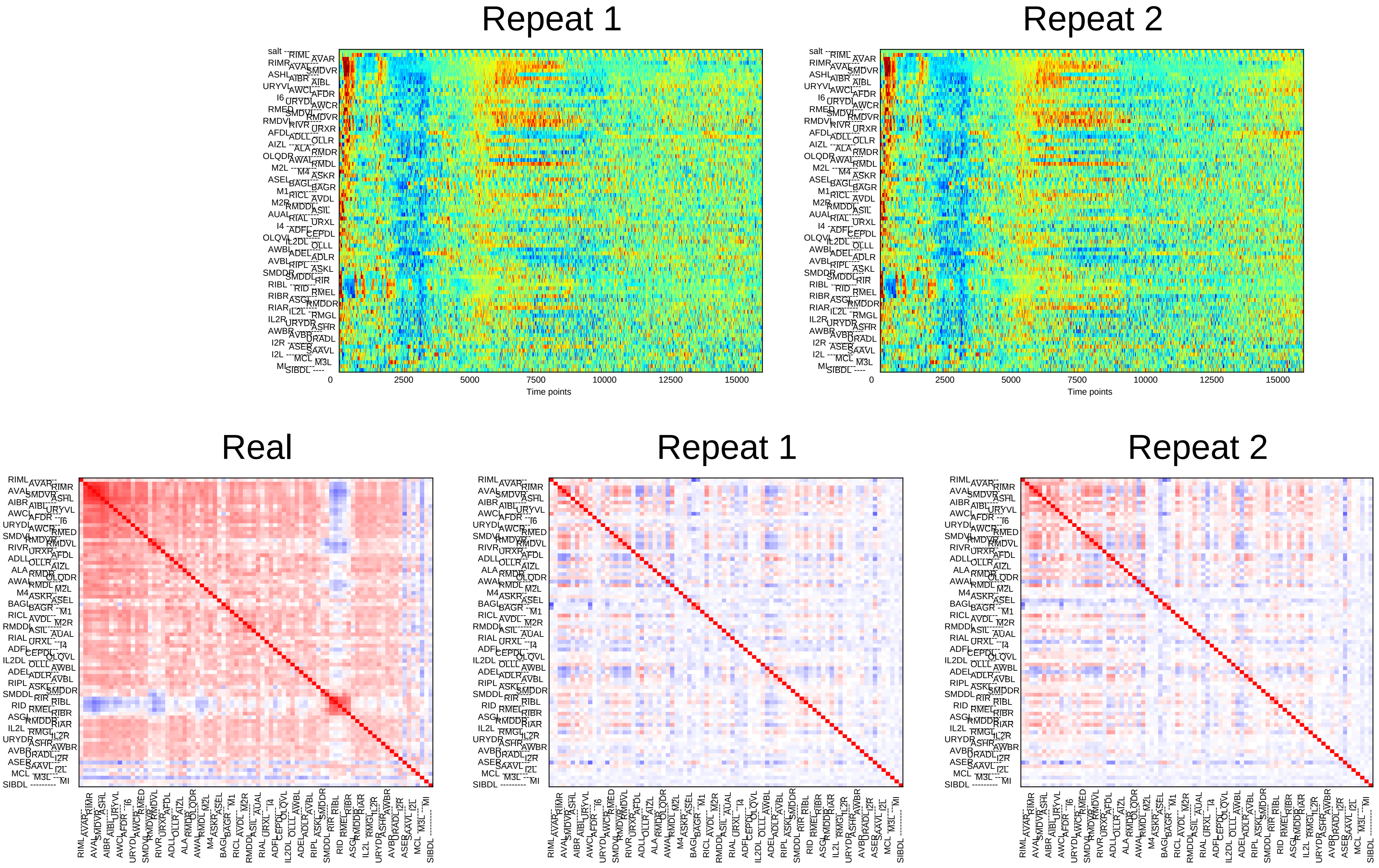

Repeat 1
Repeat 2
0
2500
5000
7500
10000
12500
15000
Time points
Real
Repeat 1
Repeat 2

### Sample21

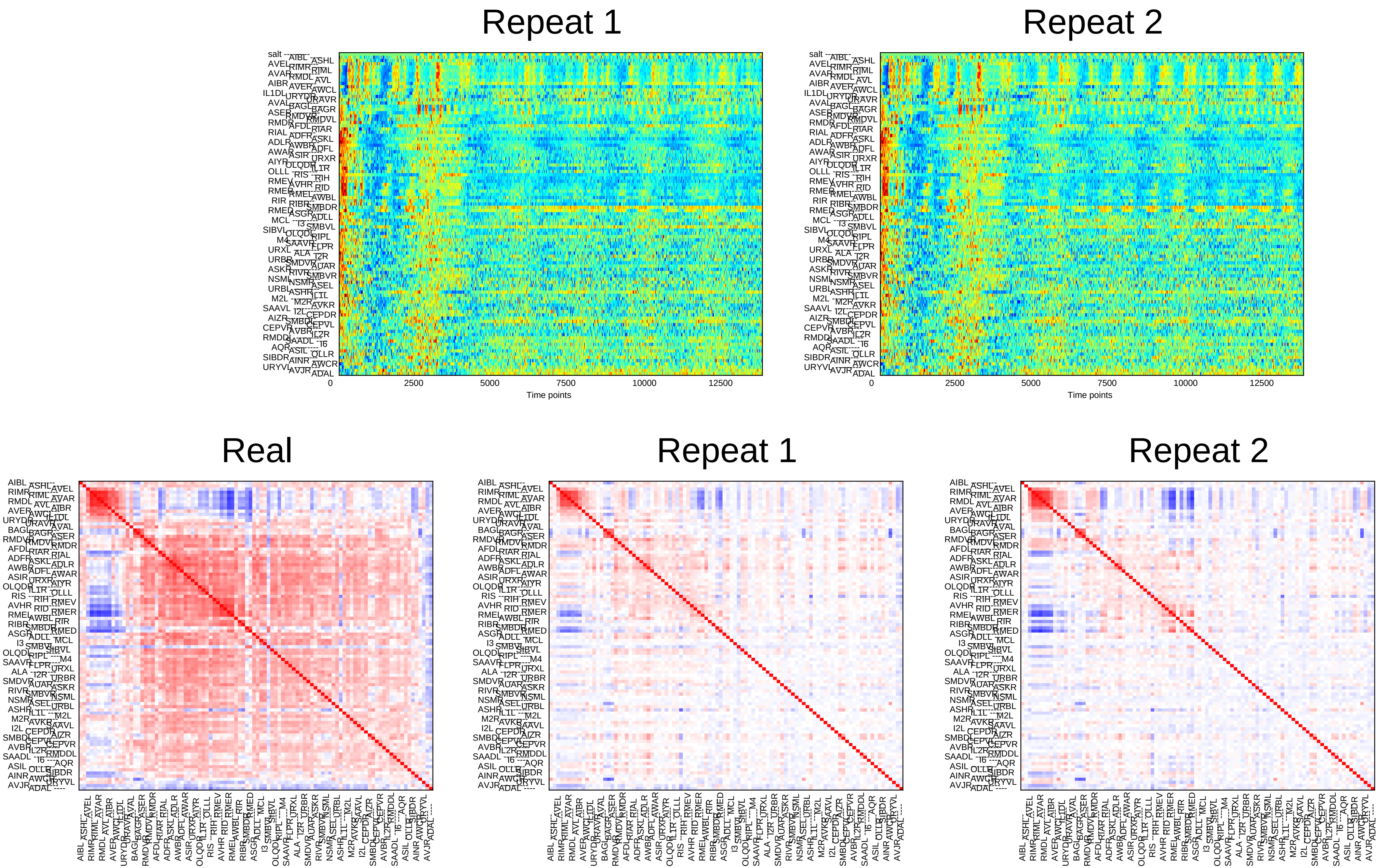

Repeat 1
Repeat 2
0
2500
5000
7500
10000
12500
Time points
Real
Repeat 1
Repeat 2

### Sample22

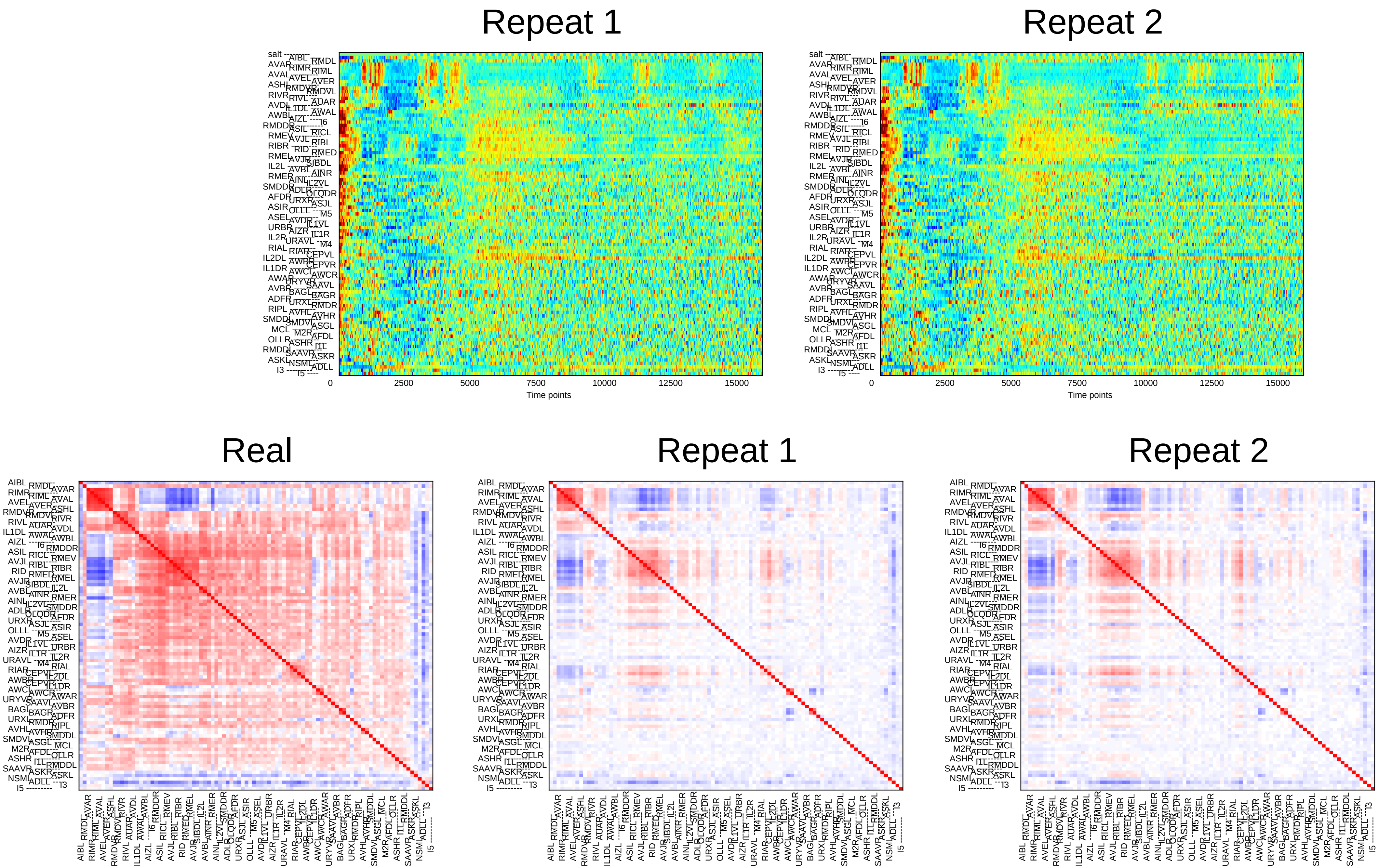
Repeat 1
Repeat 2
Time points
0
2500
5000
7500
10000
12500
15000
Real
Repeat 1
Repeat 2

### Sample23

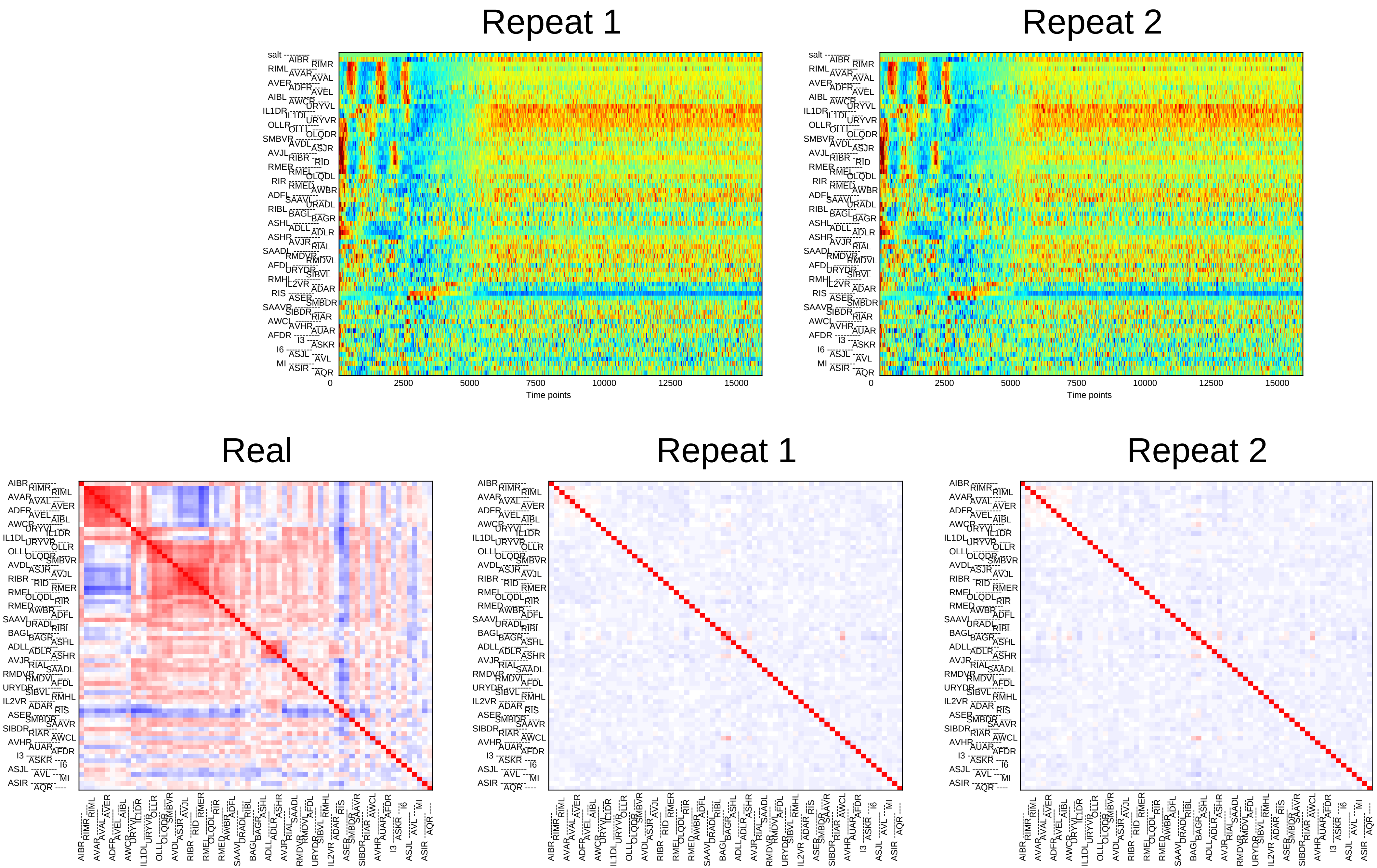
Repeat 1
Repeat 2
0
2500
5000
7500
10000
12500
15000
Time points
Real
Repeat 1
Repeat 2

### Sample24

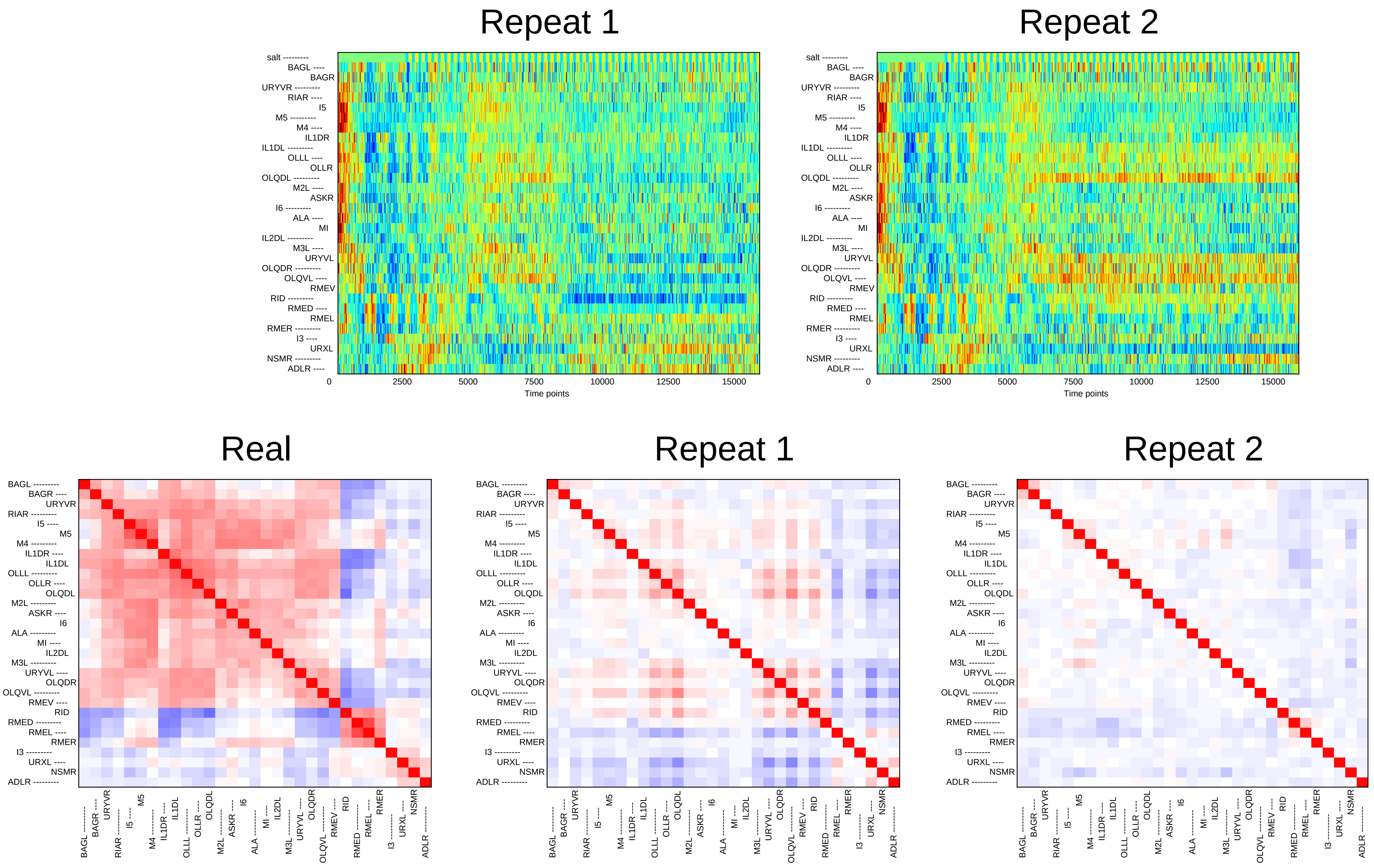

Repeat 1
Repeat 2
salt
BAGL
BAGR
URYVR
RIAR
I5
M5
M4
IL1DR
IL1DL
OLLL
OLLR
OLQDL
M2L
ASKR
I6
ALA
MI
IL2DL
M3L
URYVL
OLQDR
OLQVL
RMEV
RID
RMED
RMEL
RMER
I3
URXL
NSMR
ADLR
0
2500
5000
7500
10000
12500
15000
Time points
Real
Repeat 1
Repeat 2
