## Supplementary File 5B for "Deducing ensemble dynamics and information flow from the whole-brain imaging data"

### Sample1

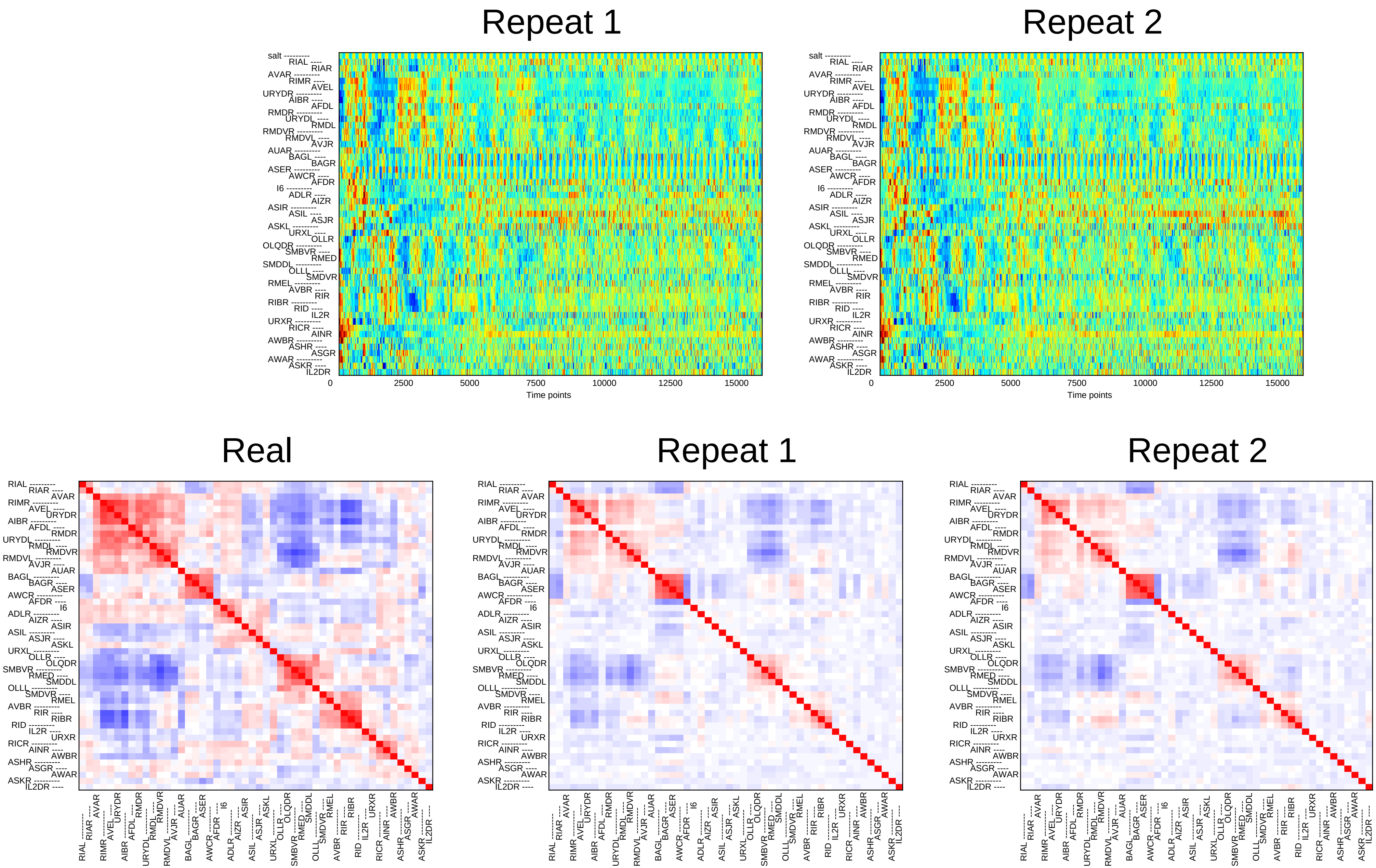
Repeat 1
Repeat 2
salt
0
2500
5000
7500
10000
12500
15000
Time points
Real
Repeat 1
Repeat 2

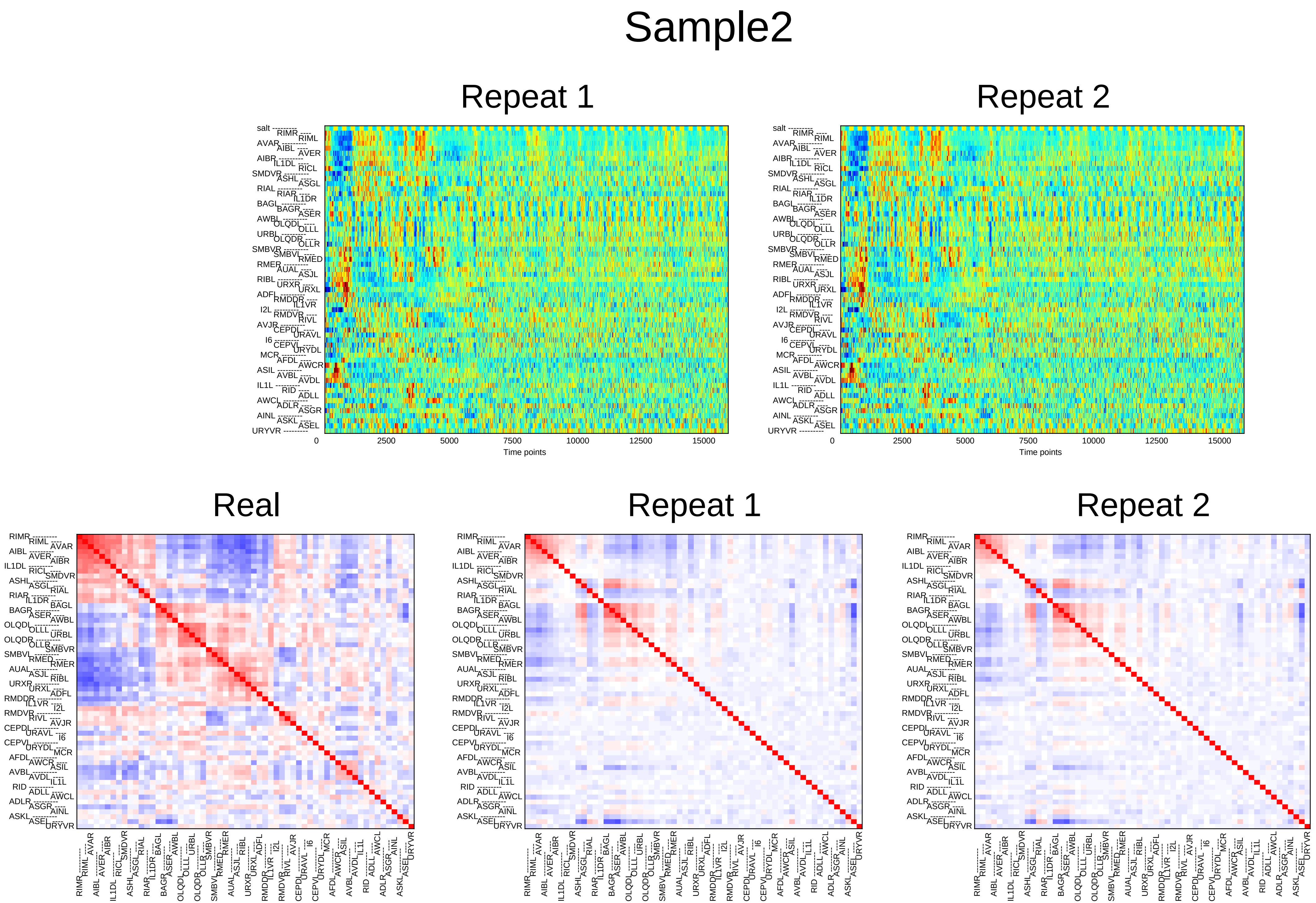
Sample2
Repeat 1
Repeat 2
0
2500
5000
7500
10000
12500
15000
Time points
Real
Repeat 1
Repeat 2

### Sample3

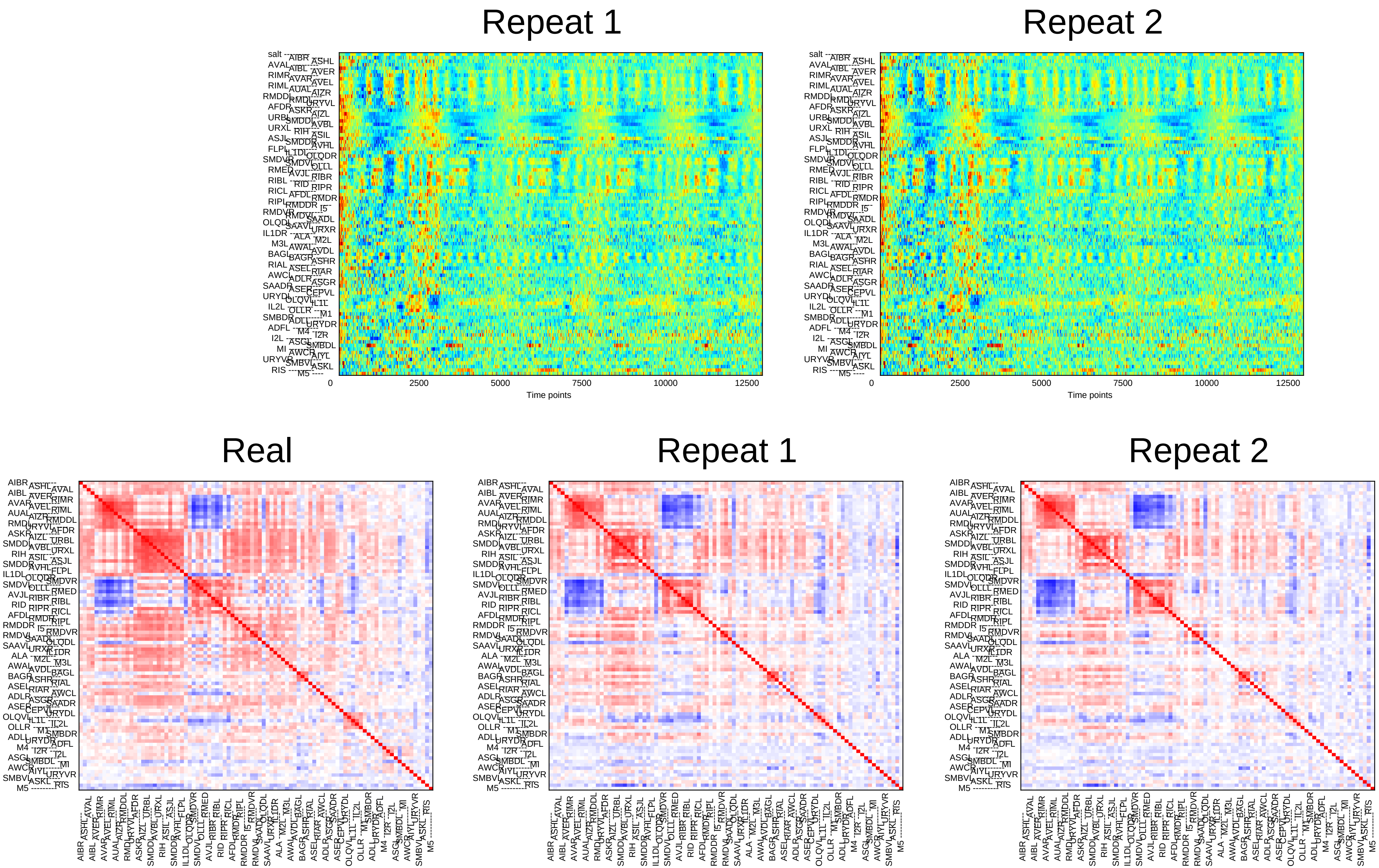

Repeat 1
Repeat 2
0
2500
5000
7500
10000
12500
Time points
Real
Repeat 1
Repeat 2

### Sample4

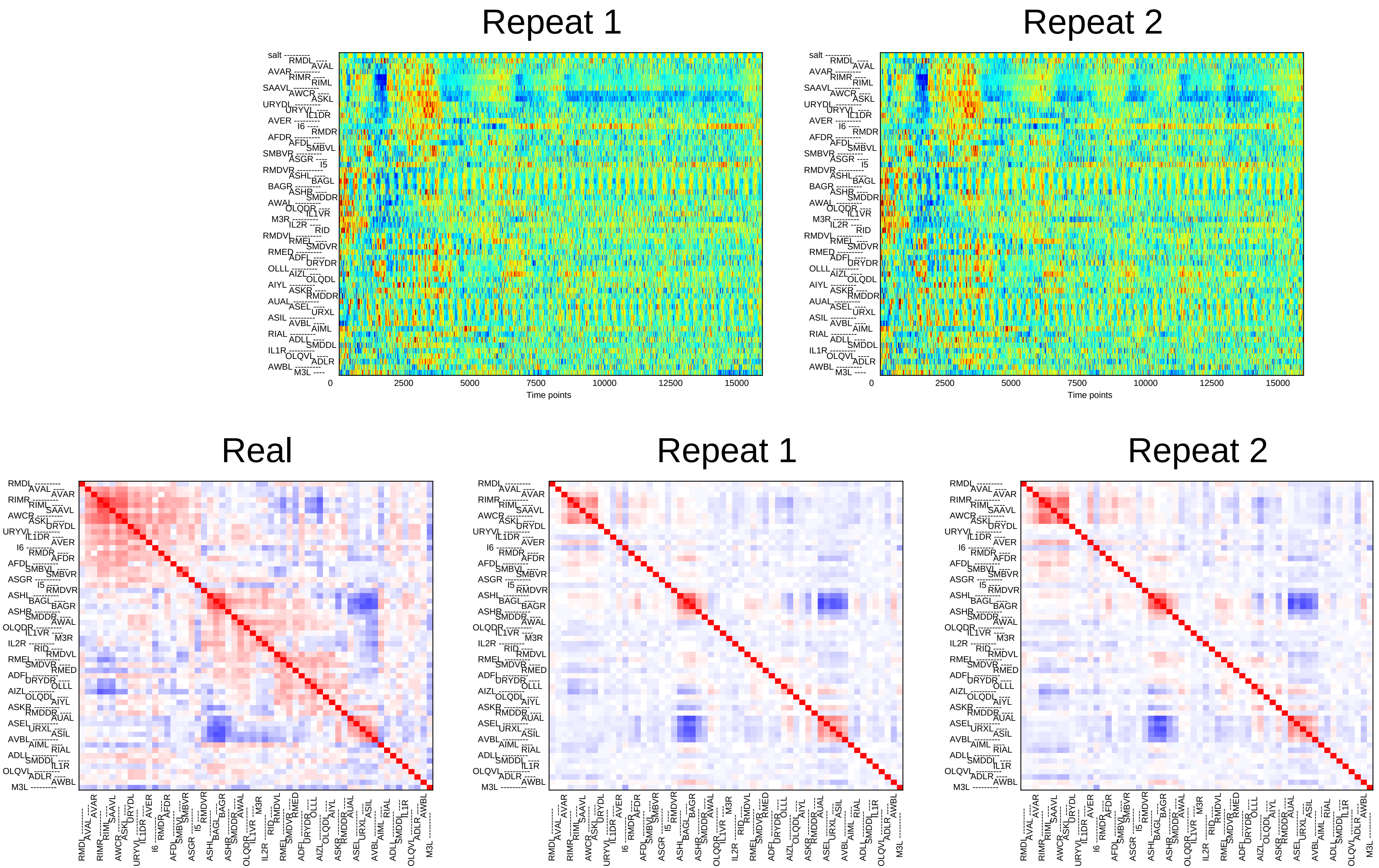

Repeat 1
Repeat 2
Time points
0
2500
5000
7500
10000
12500
15000
Real
Repeat 1
Repeat 2

### Sample5

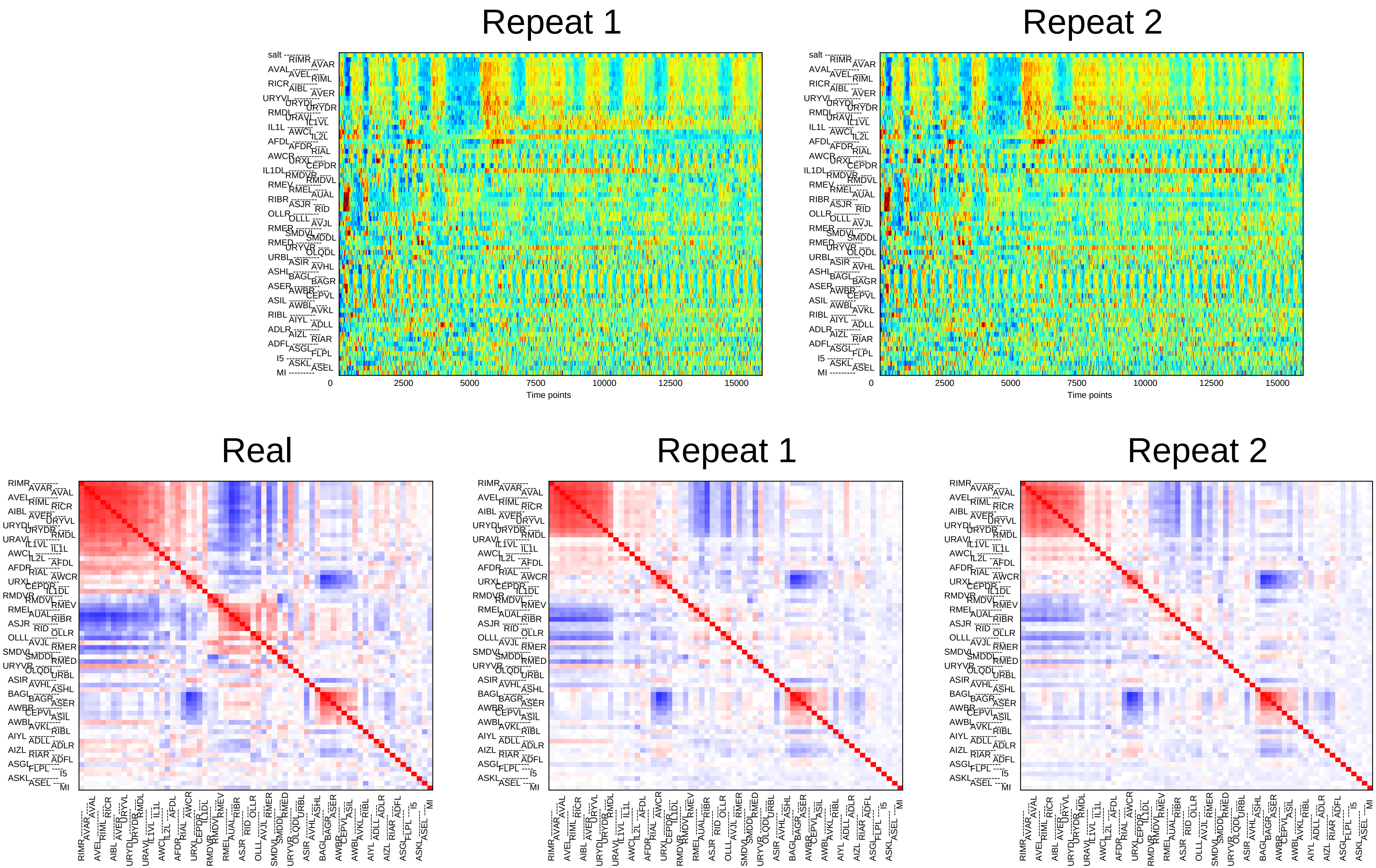

Repeat 1
Repeat 2
0
2500
5000
7500
10000
12500
15000
Time points
Real
Repeat 1
Repeat 2

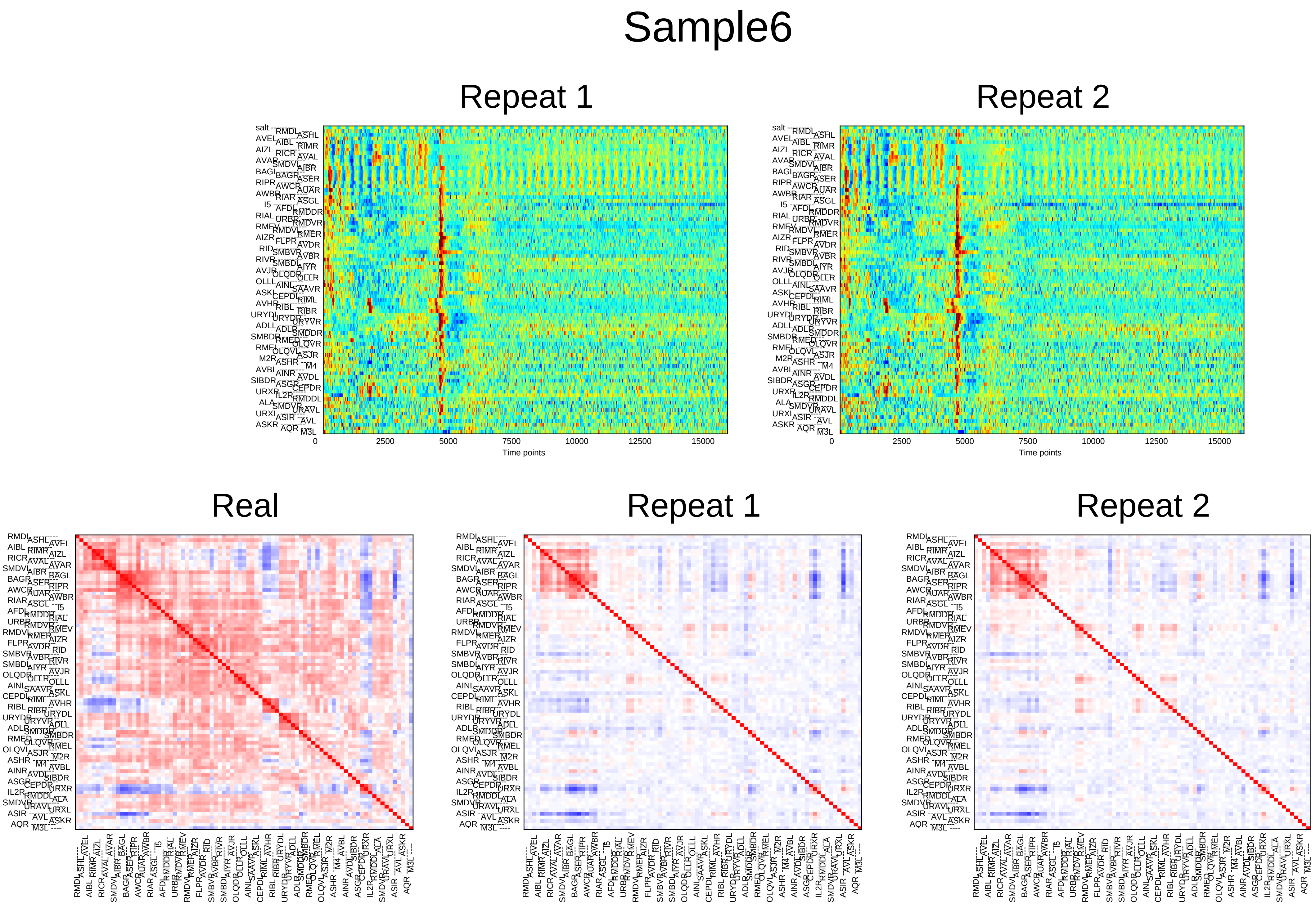
Sample6
Repeat 1
Repeat 2
0
2500
5000
7500
10000
12500
15000
Time points
Real
Repeat 1
Repeat 2

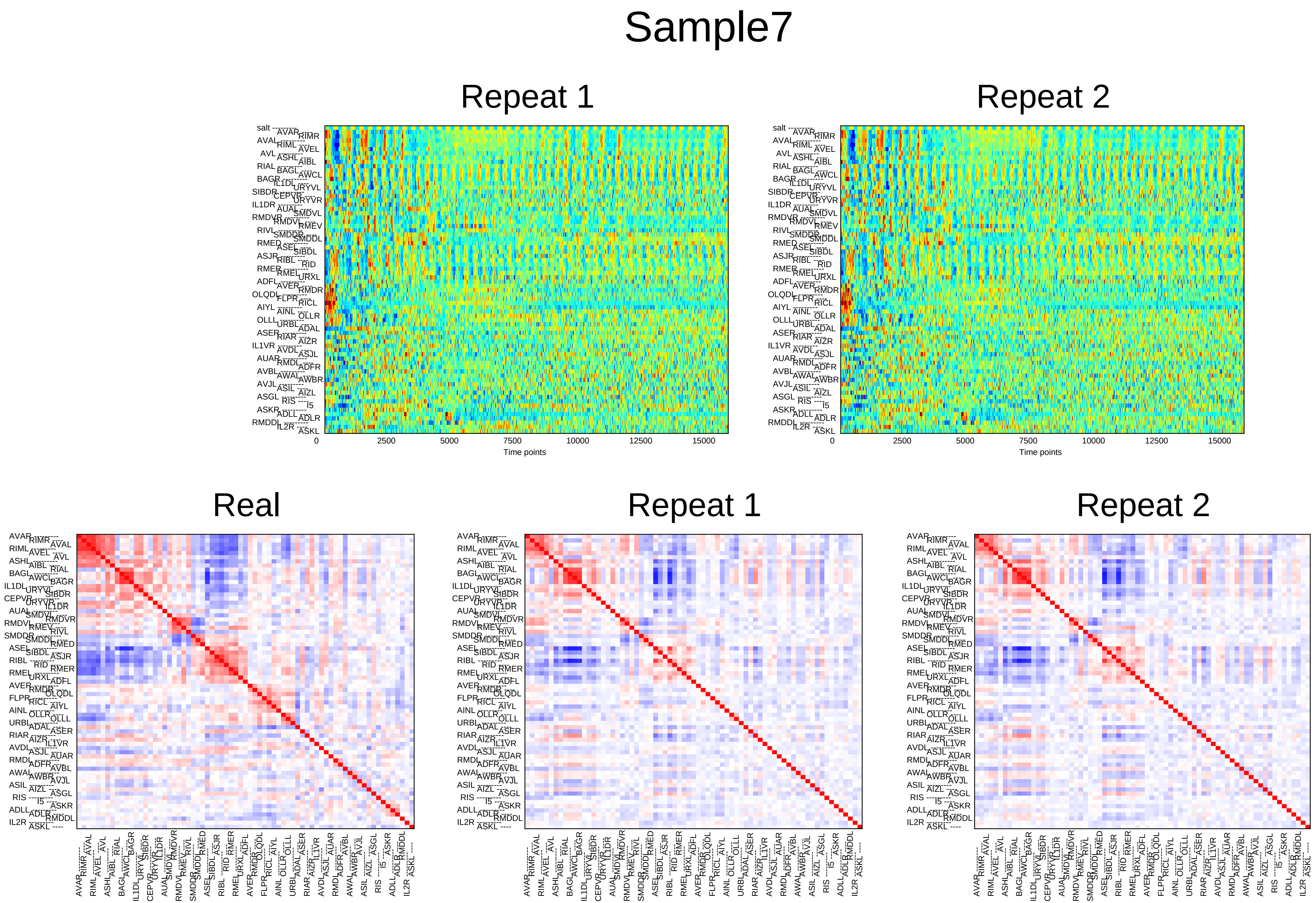
Sample7
Repeat 1
Repeat 2
Time points
0
2500
5000
7500
10000
12500
15000
Real
Repeat 1
Repeat 2

### Sample8

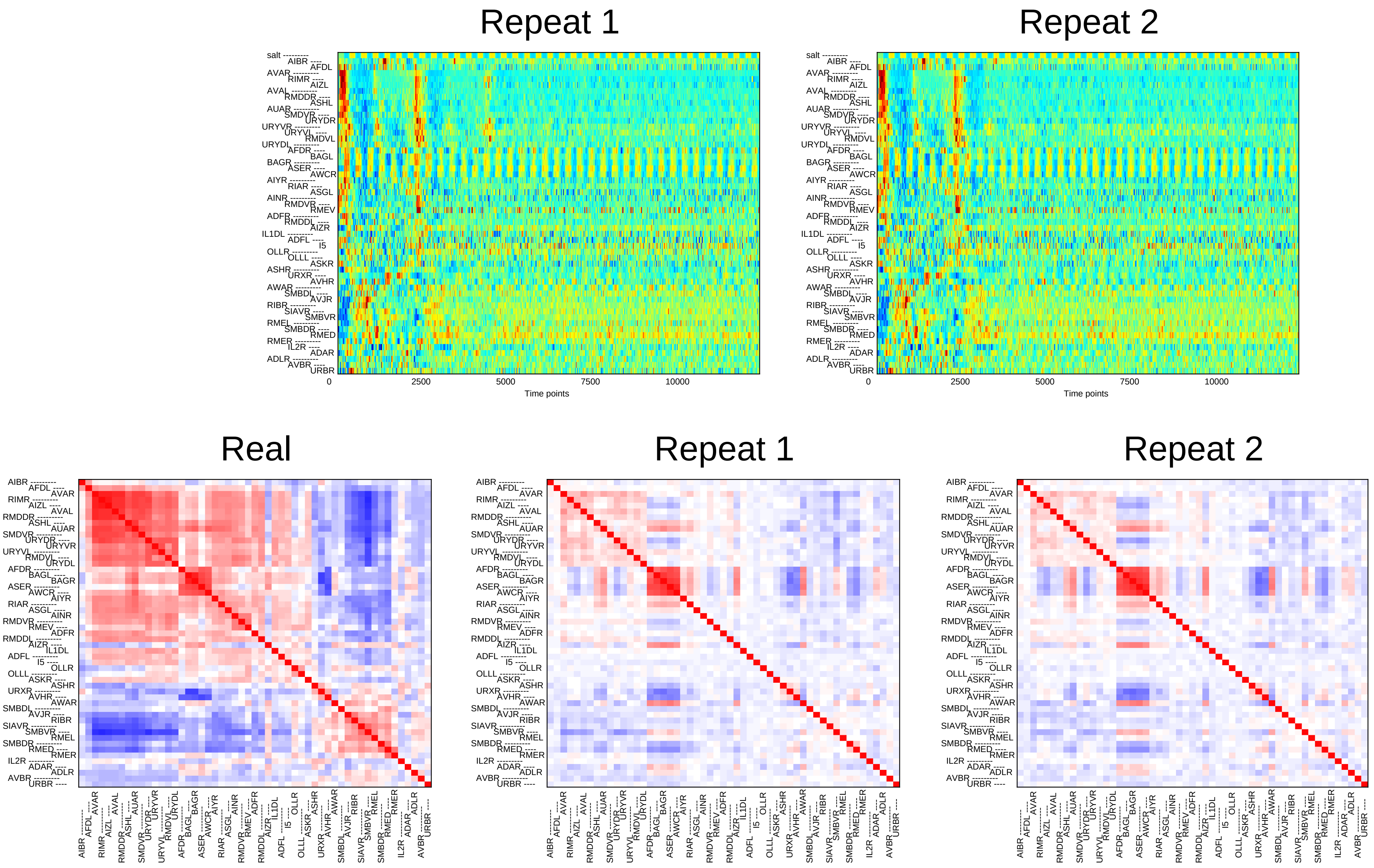

Repeat 1
Repeat 2
salt
AIBR
AFDL
AVAR
RIMR
AIZL
AVAL
RMDDR
ASHL
AUAR
SMDVR
URYDR
URYVR
URYVL
RMDVL
URYDL
AFDR
BAGL
BAGR
ASER
AWCR
AIYR
RIAR
ASGL
AINR
RMDVR
RMEV
ADFR
RMDDL
AIZR
IL1DL
ADFL
I5
OLLR
OLLL
ASKR
ASHR
URXR
AVHR
AWAR
SMBDL
AVJR
RIBR
SIAVR
SMBVR
RMEL
SMBDR
RMED
RMER
IL2R
ADAR
ADLR
AVBR
URBR
0
2500
5000
7500
10000
Time points
Real
Repeat 1
Repeat 2

### Sample9

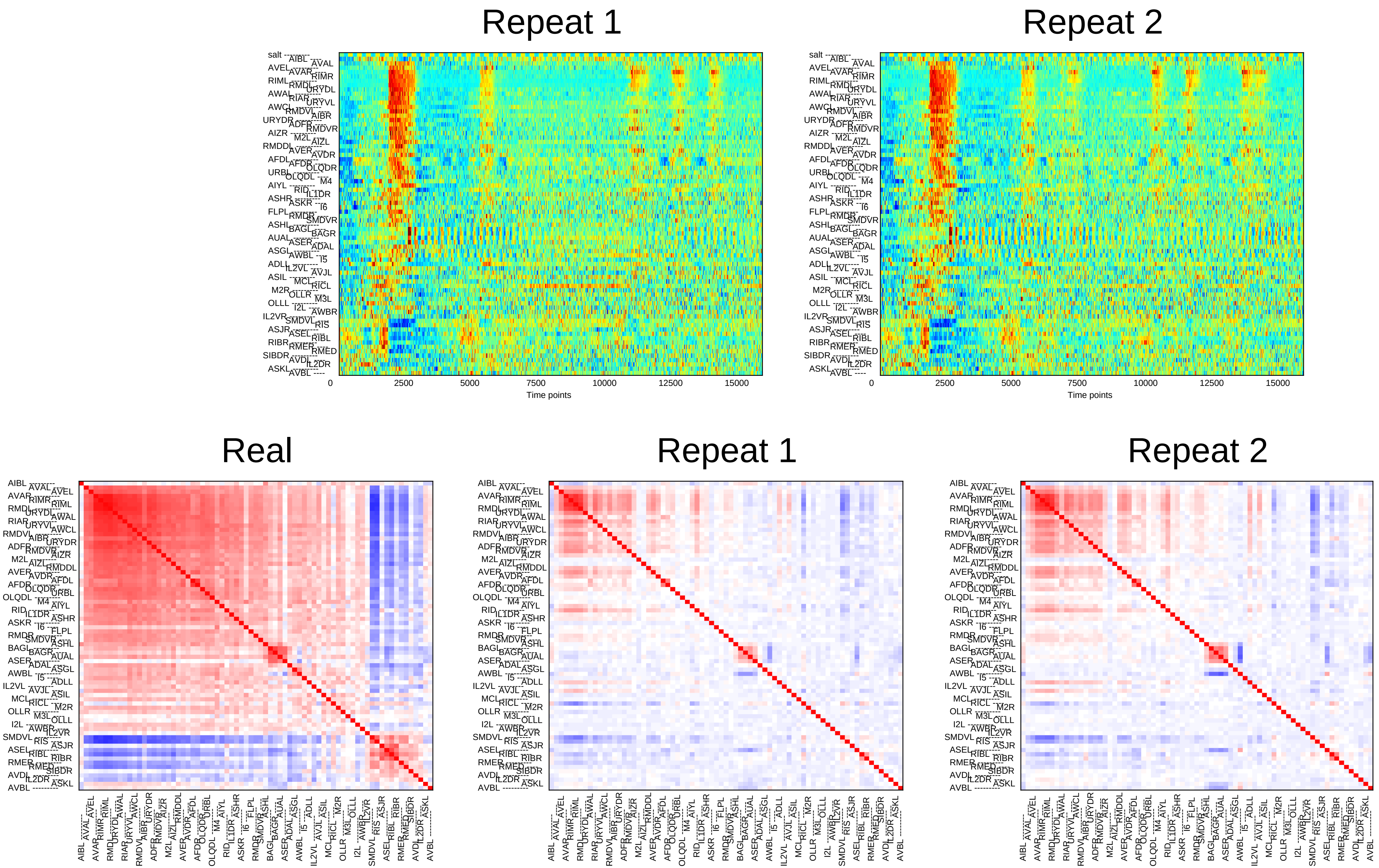
Repeat 1
Repeat 2
0
2500
5000
7500
10000
12500
15000
Time points
Real
Repeat 1
Repeat 2

### Sample10

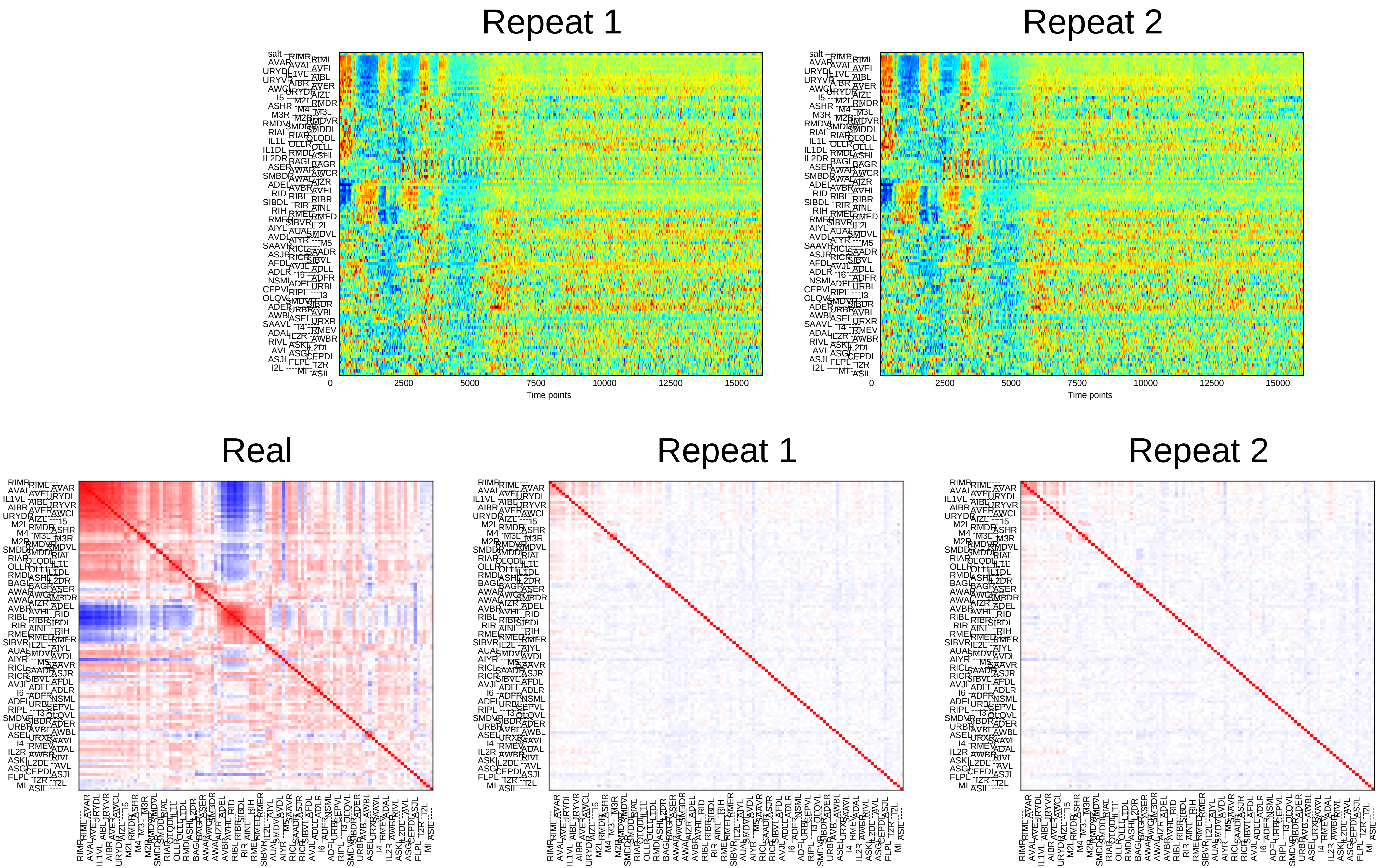

Repeat 1
Repeat 2
0
2500
5000
7500
10000
12500
15000
Time points
Real
Repeat 1
Repeat 2

### Sample11

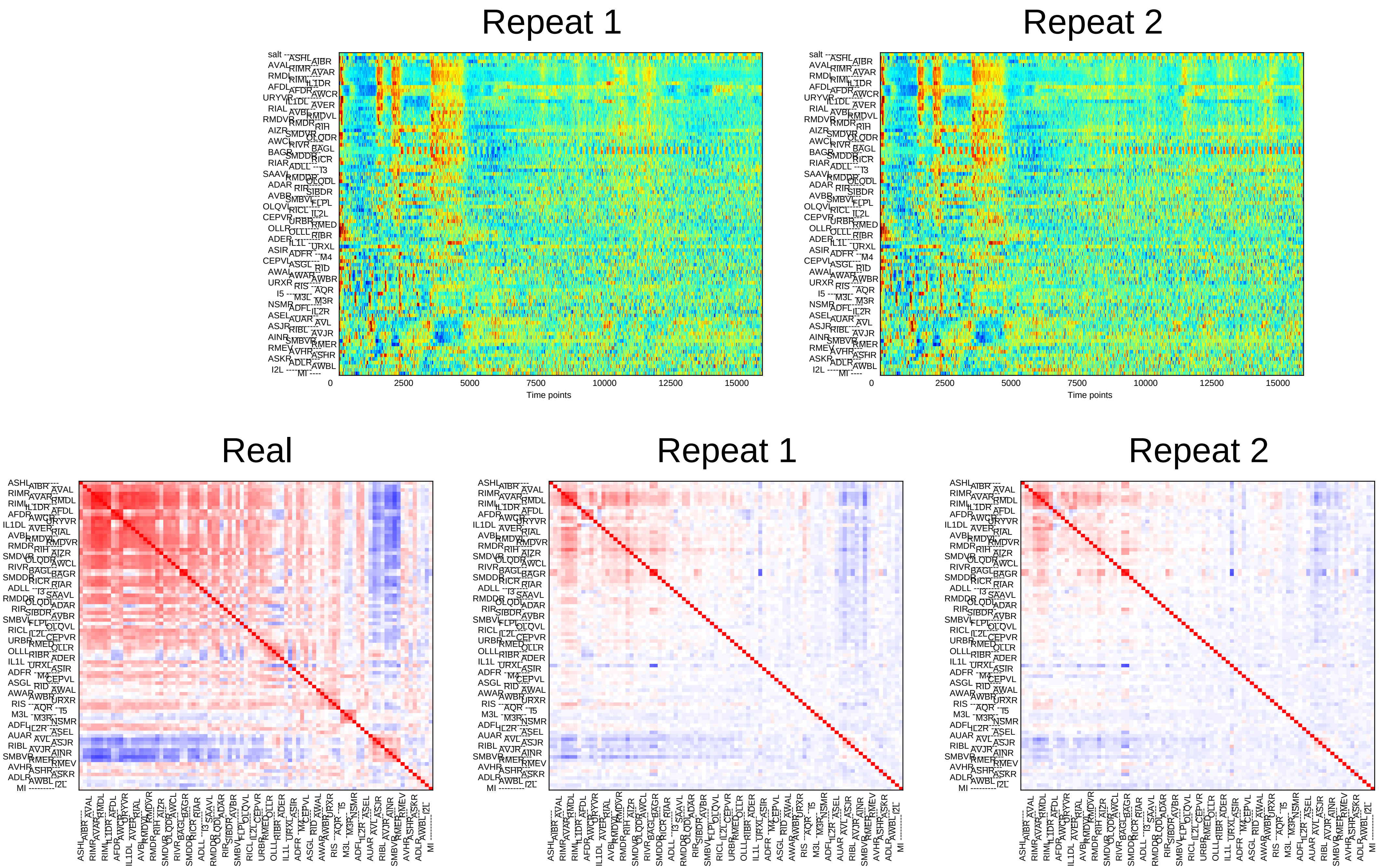

Repeat 1
Repeat 2
0
2500
5000
7500
10000
12500
15000
Time points
Real
Repeat 1
Repeat 2

### Sample12

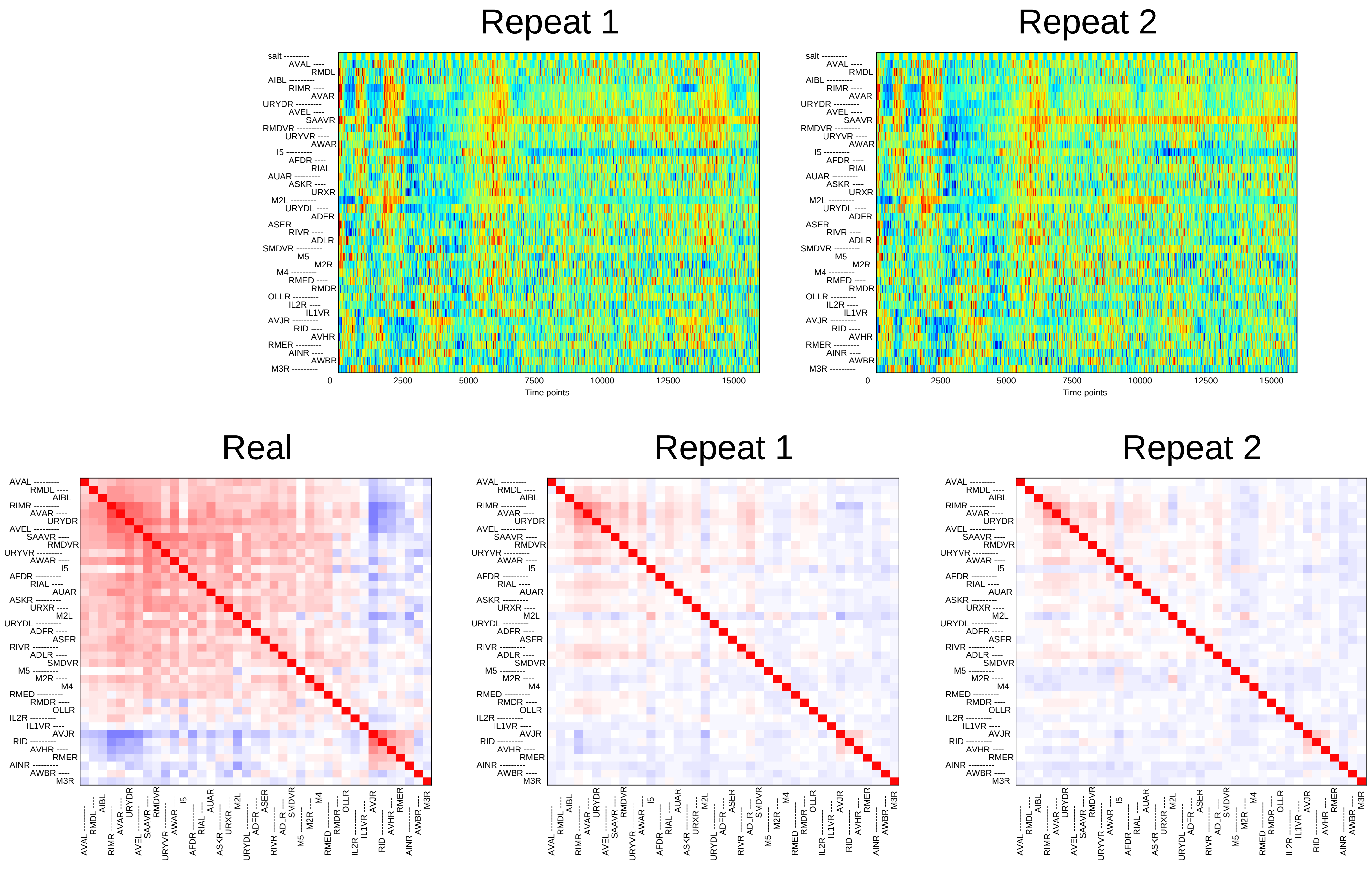

Repeat 1
Repeat 2
salt
AVAL
RMDL
AIBL
RIMR
AVAR
URYDR
AVEL
SAAVR
RMDVR
URYVR
AWAR
I5
AFDR
RIAL
AUAR
ASKR
URXR
M2L
URYDL
ADFR
ASER
RIVR
ADLR
SMDVR
M5
M2R
M4
RMED
RMDR
OLLR
IL2R
IL1VR
AVJR
RID
AVHR
RMER
AINR
AWBR
M3R
0
2500
5000
7500
10000
12500
15000
Time points
Real
Repeat 1
Repeat 2

### Sample13

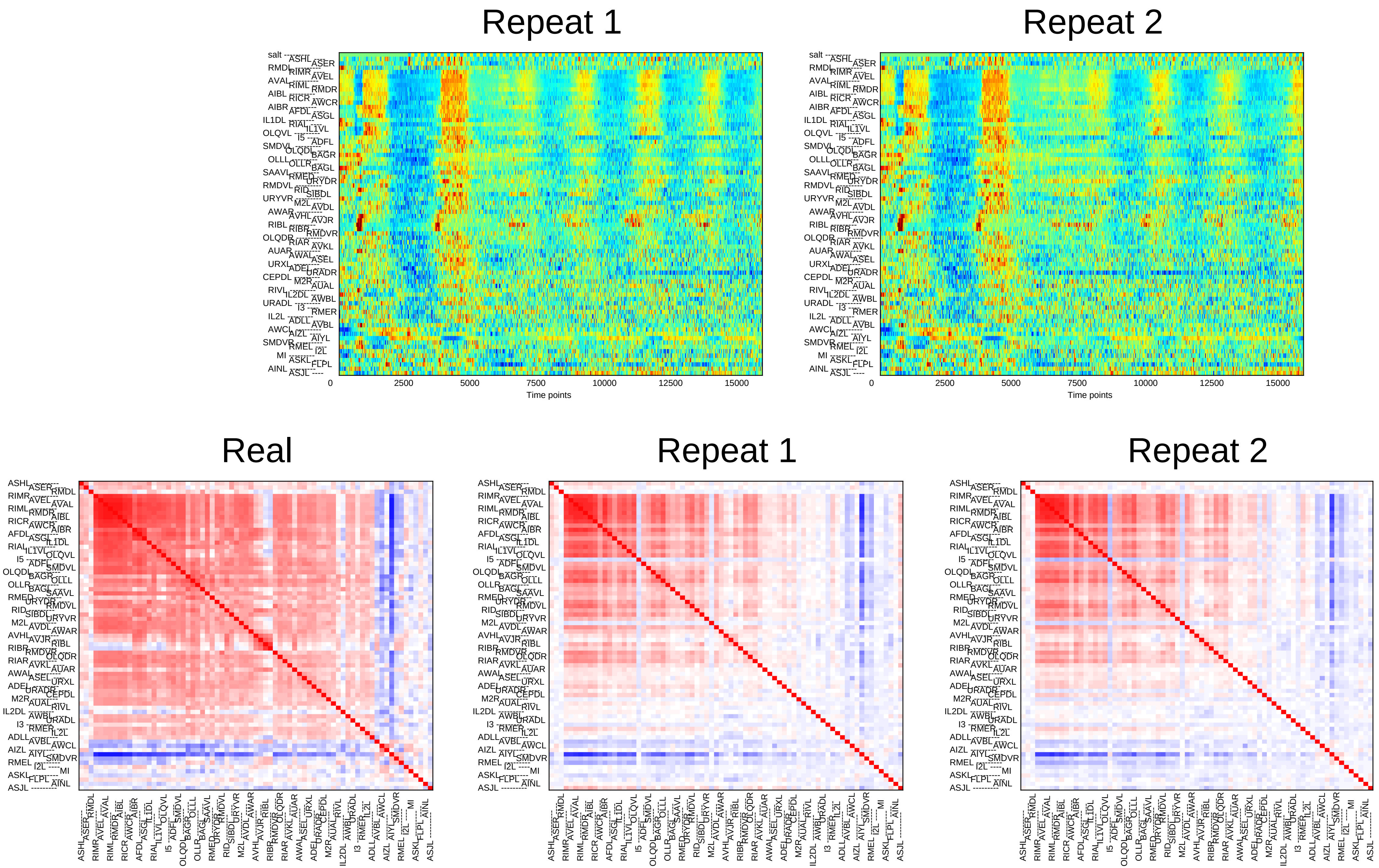
Repeat 1
Repeat 2
0
2500
5000
7500
10000
12500
15000
Time points
Real
Repeat 1
Repeat 2

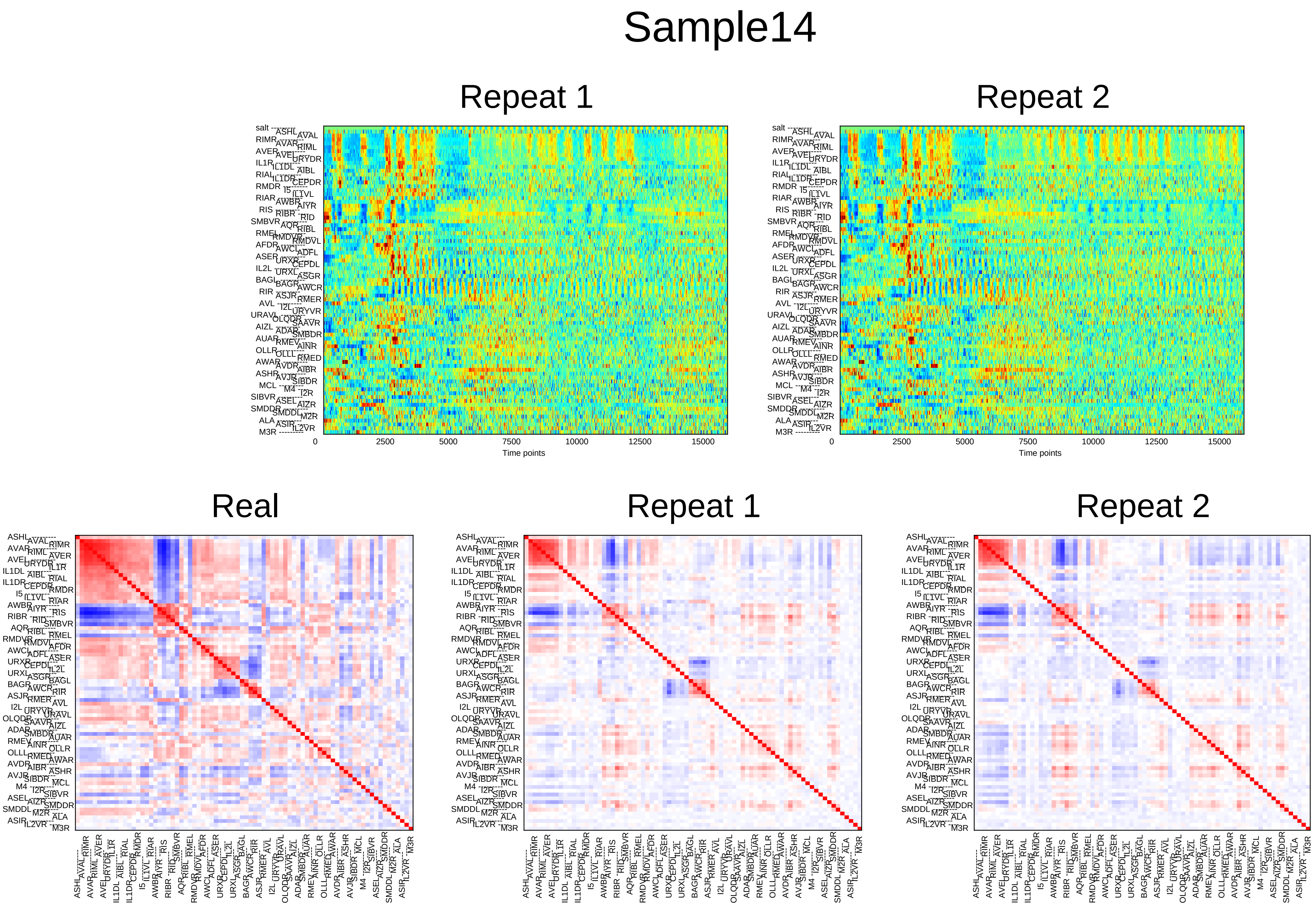
Sample14
Repeat 1
Repeat 2
0
2500
5000
7500
10000
12500
15000
Time points
Real
Repeat 1
Repeat 2

### Sample15

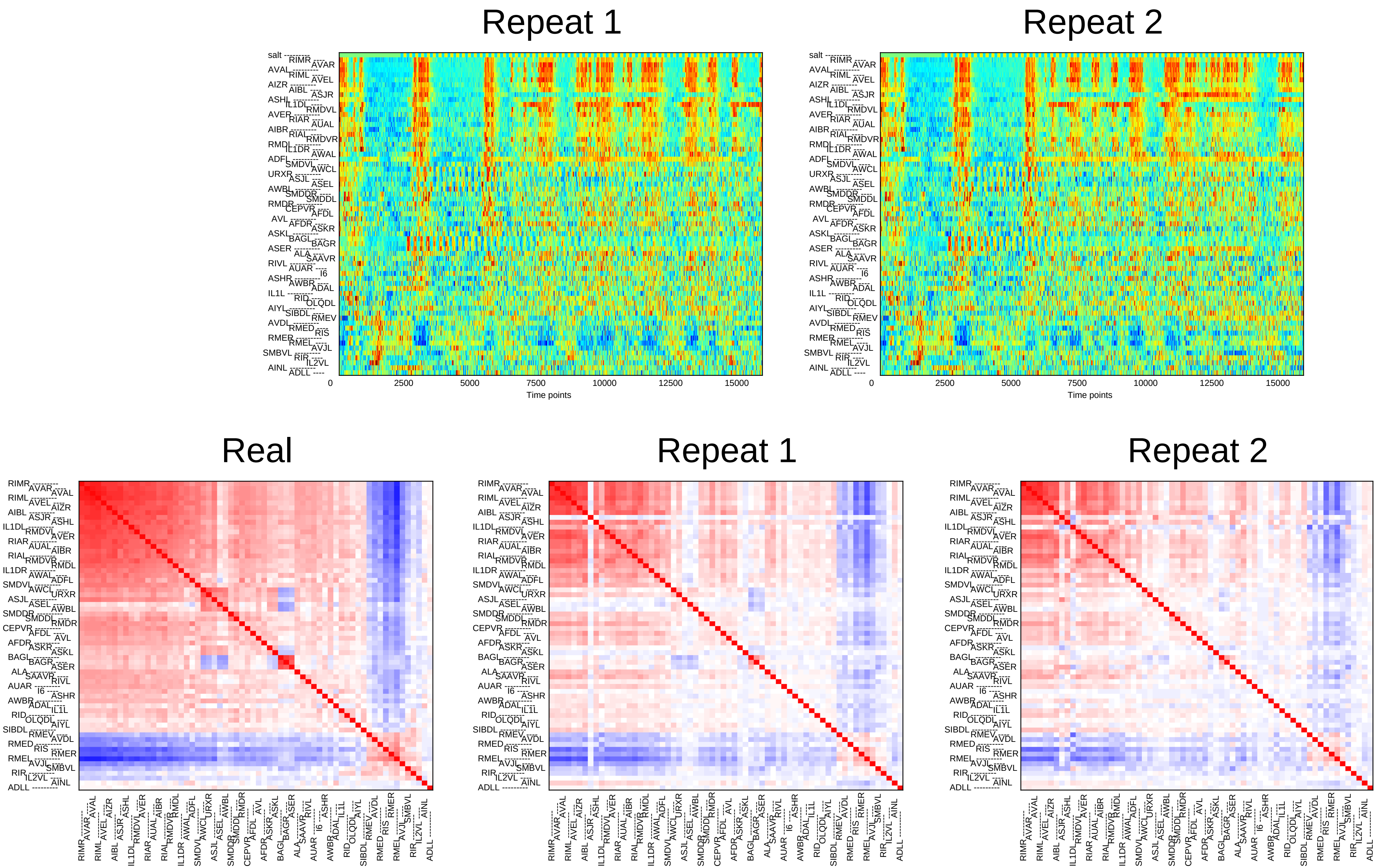
Repeat 1
Repeat 2
Real
Repeat 1
Repeat 2
Time points
0
2500
5000
7500
10000
12500
15000
salt
RIMR AVAR AVAL RIML AVEL AIZR AIBL ASJR ASHL IL1DL RMDVL AVER RIAR AUAL AIBR RIAL RMDVR RMDL IL1DR AWAL ADFL SMDVL AWCL URXR ASJL ASEL AWBL SMDDR SMDDL RMDR CEPVR AFDL AVL AFDR ASKR ASKL BAGL BAGR ASER ALA SAAVR RIVL AUAR I6 ASHR AWBR ADAL IL1L RID OLQDL AIYL SIBDL RMEV AVDL RMED RIS RMER RMEL AVJL SMBVL RIR IL2VL AINL ADLL

### Sample16

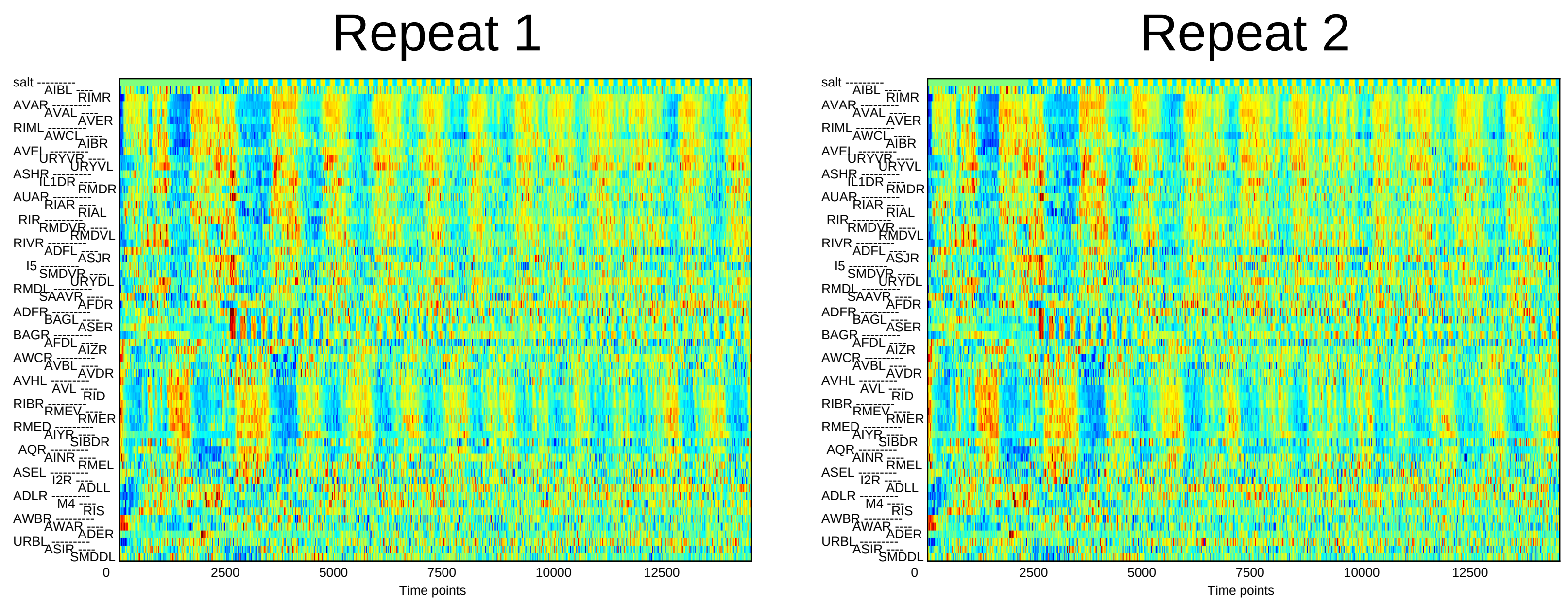

Repeat 1
Repeat 2
Time points
0
2500
5000
7500
10000
12500


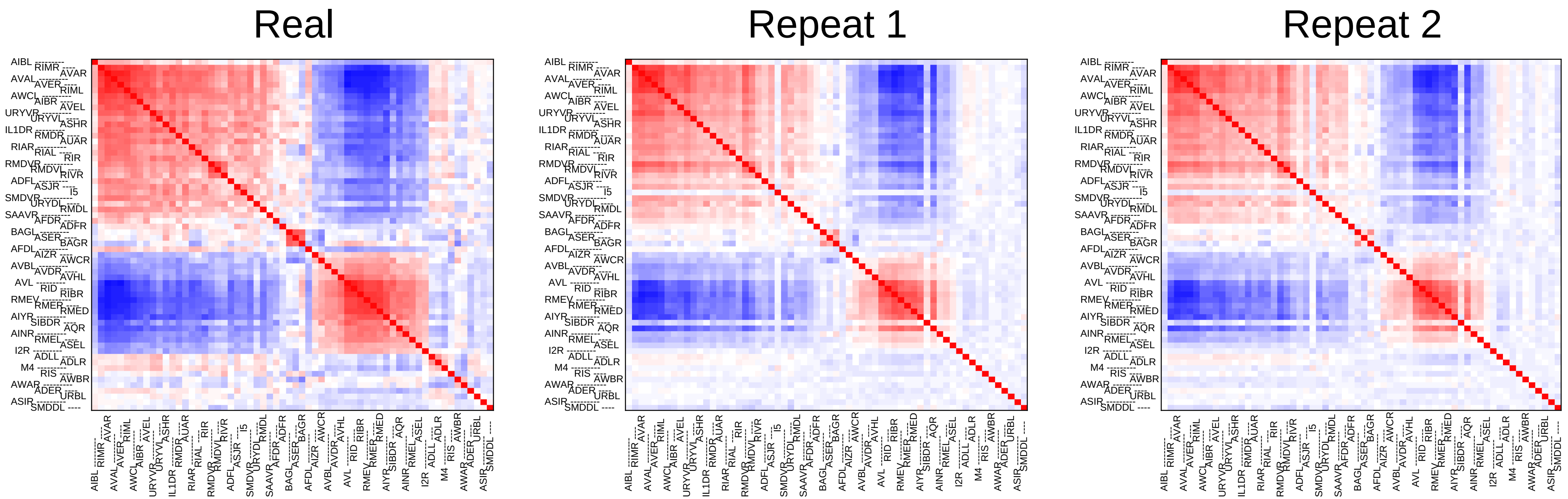

Real
Repeat 1
Repeat 2

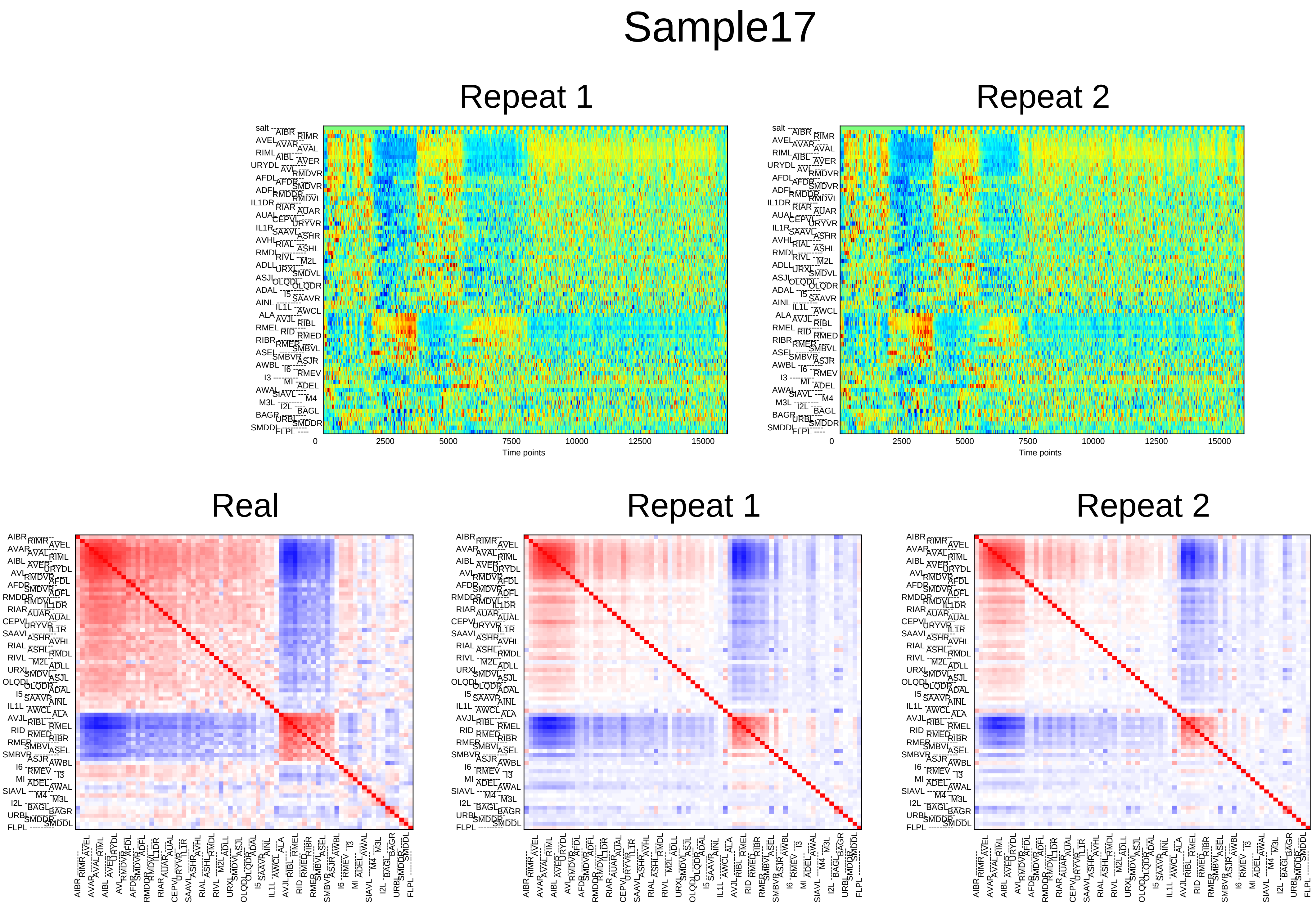

Sample17
Repeat 1
Repeat 2
0
2500
5000
7500
10000
12500
15000
Time points
Real
Repeat 1
Repeat 2

### Sample18

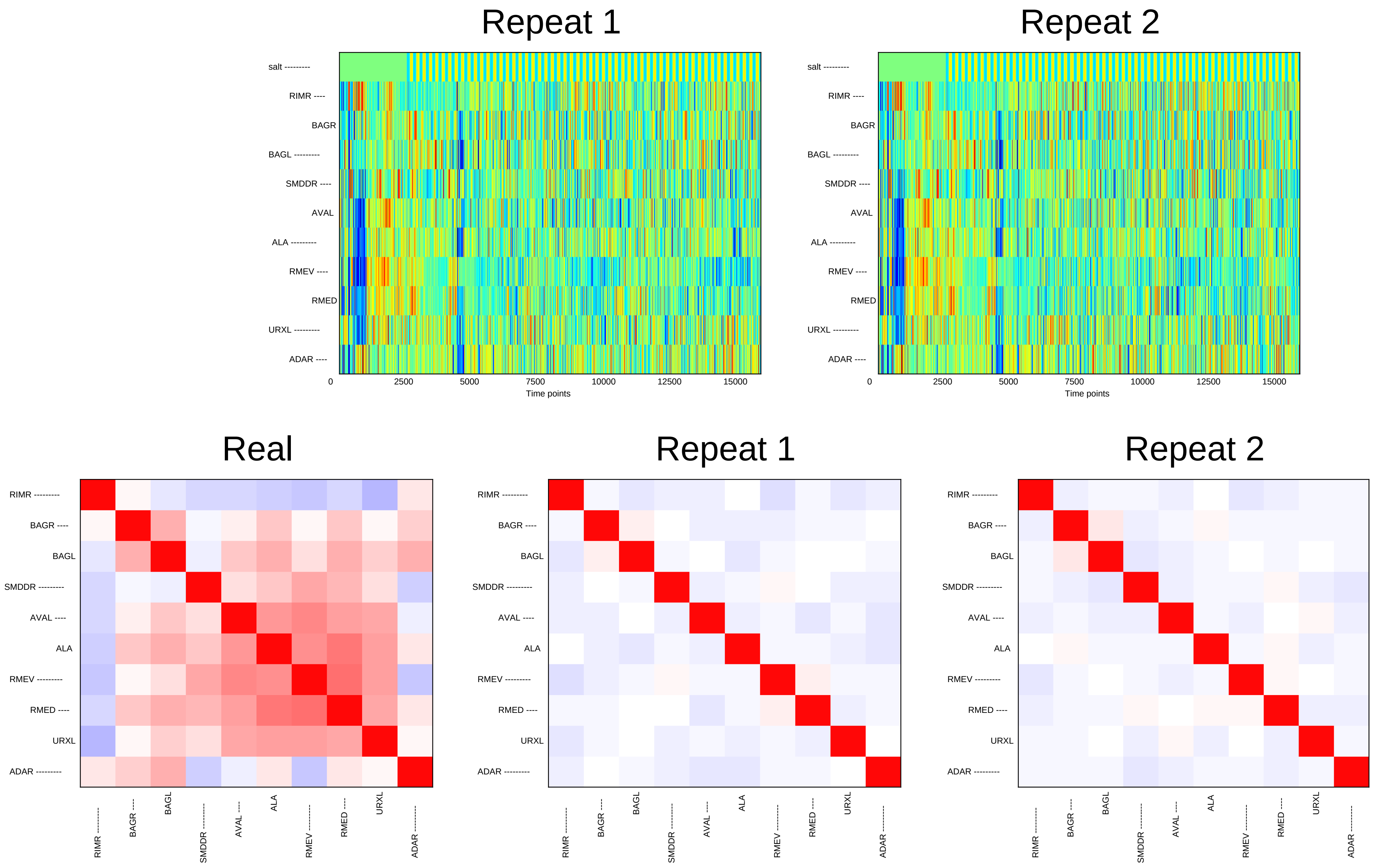
Repeat 1
Repeat 2
salt ---------
RIMR ----
BAGR
BAGL ---------
SMDDR ----
AVAL
ALA ---------
RMEV ----
RMED
URXL ---------
ADAR ----
0
2500
5000
7500
10000
12500
15000
Time points
Real
Repeat 1
Repeat 2
RIMR ---------
BAGR ----
BAGL
SMDDR ---------
AVAL ----
ALA
RMEV ---------
RMED ----
URXL
ADAR ---------

### Sample19

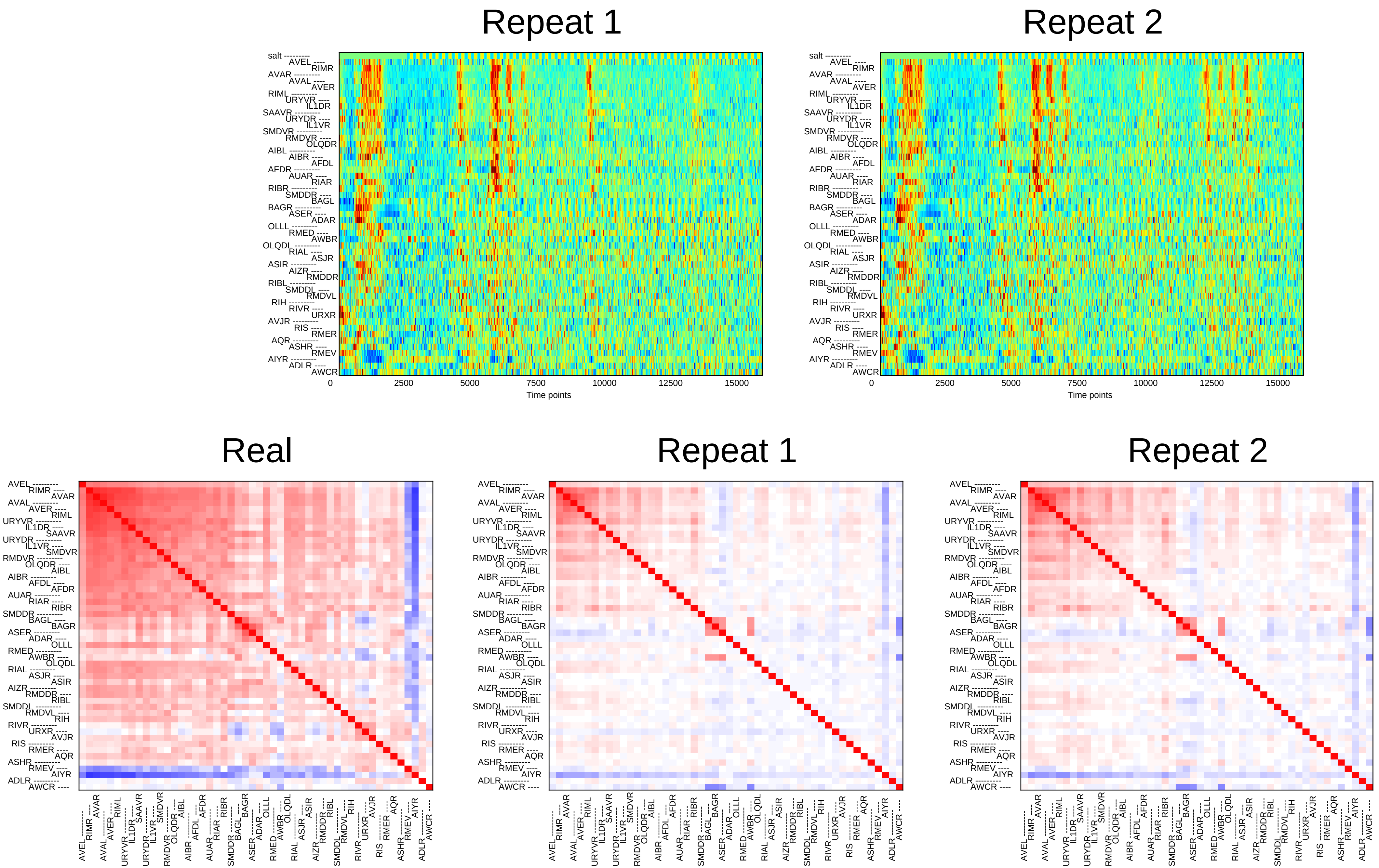

Repeat 1
Repeat 2
0
2500
5000
7500
10000
12500
15000
Time points
Real
Repeat 1
Repeat 2

### Sample20

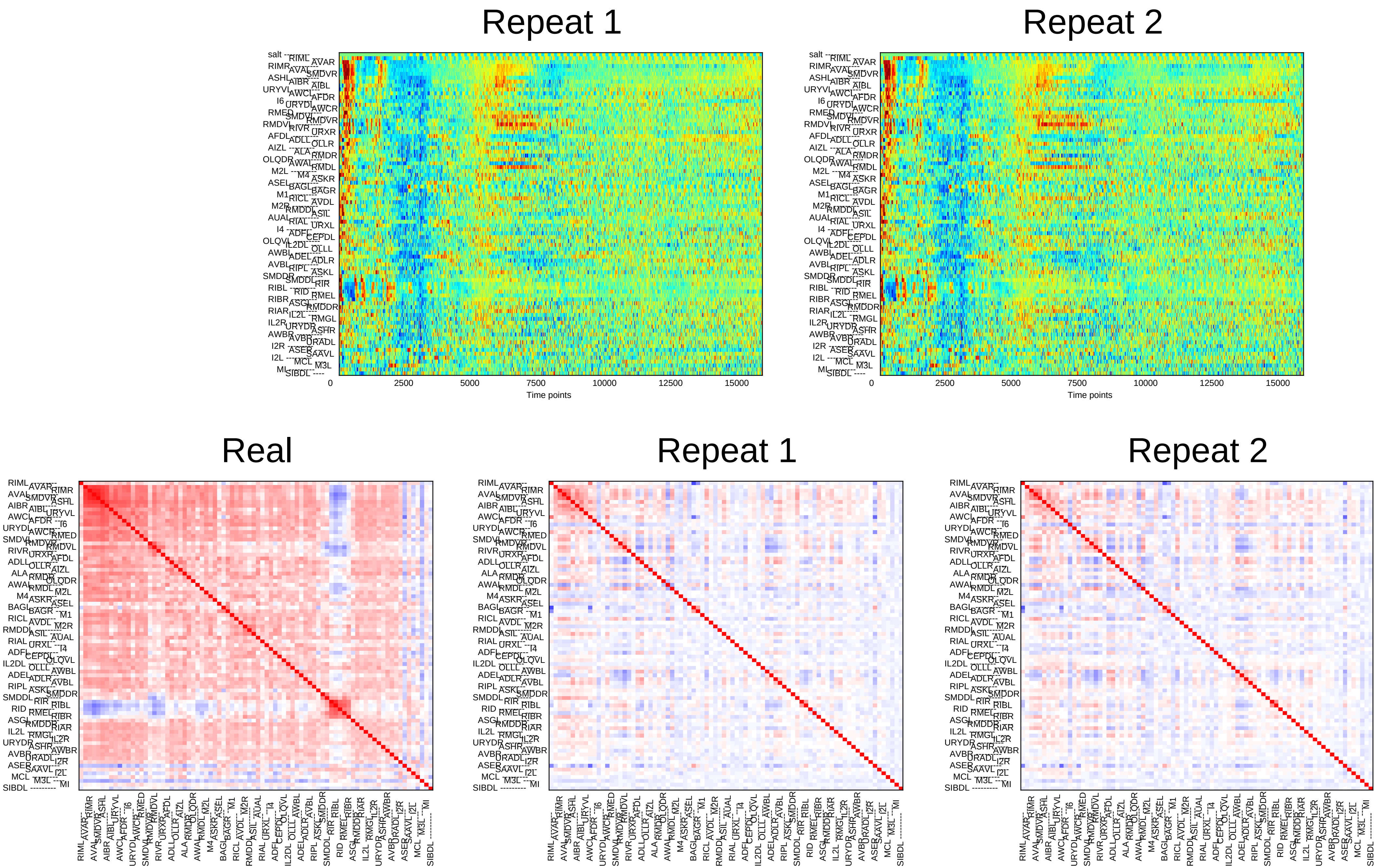
Repeat 1
Repeat 2
0
2500
5000
7500
10000
12500
15000
Time points
Real
Repeat 1
Repeat 2

### Sample21

#### Repeat 1

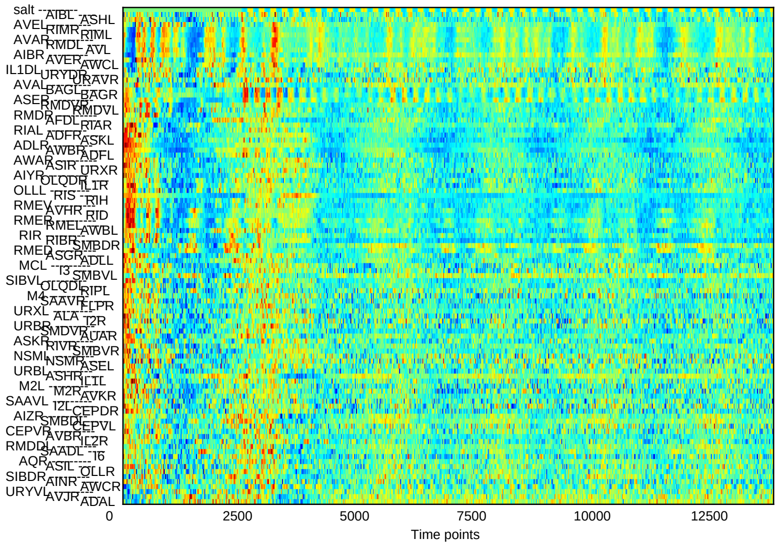


#### Repeat 2

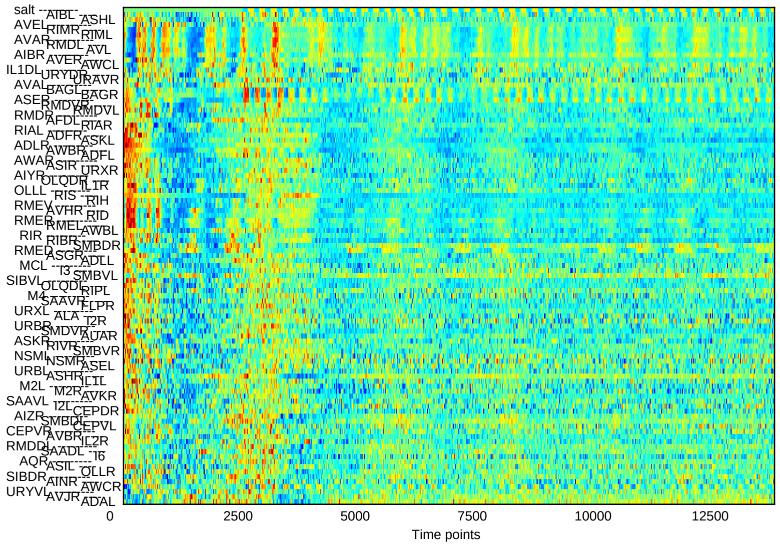


#### Real

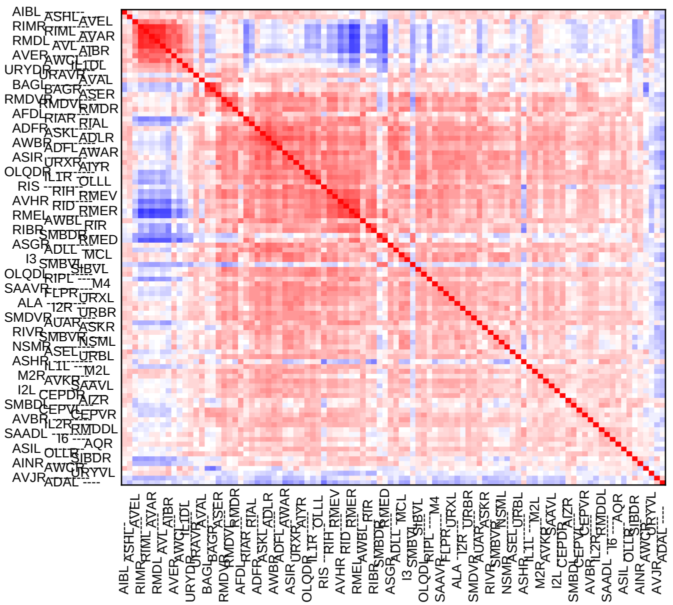

#### Repeat 1

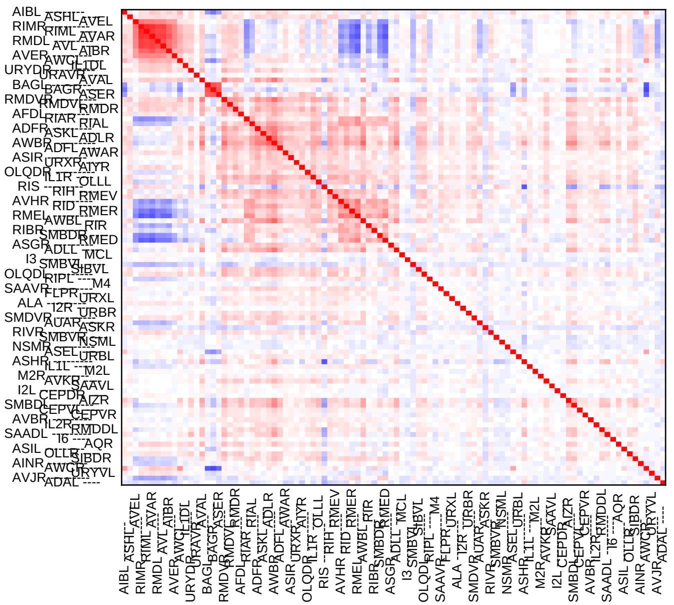

#### Repeat 2

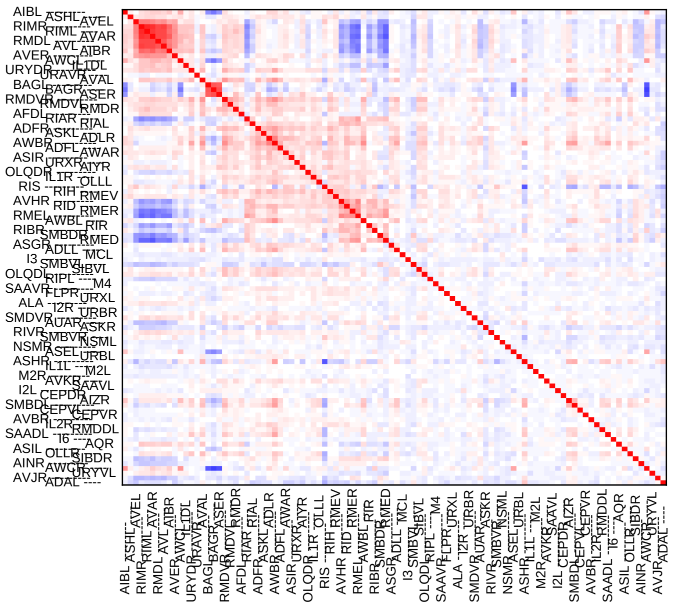

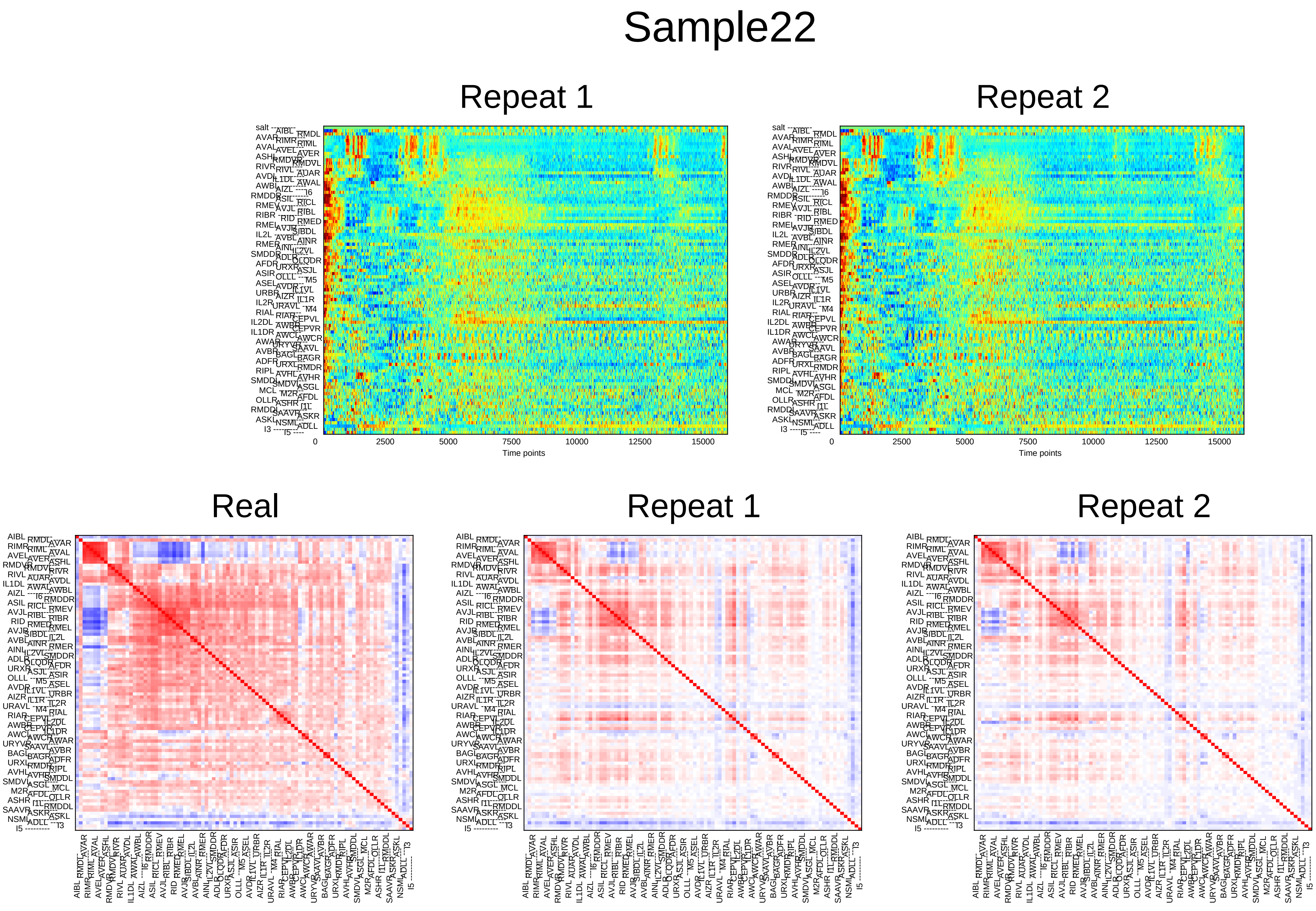
Sample22
Repeat 1
Repeat 2
0
2500
5000
7500
10000
12500
15000
Time points
Real
Repeat 1
Repeat 2

### Sample23

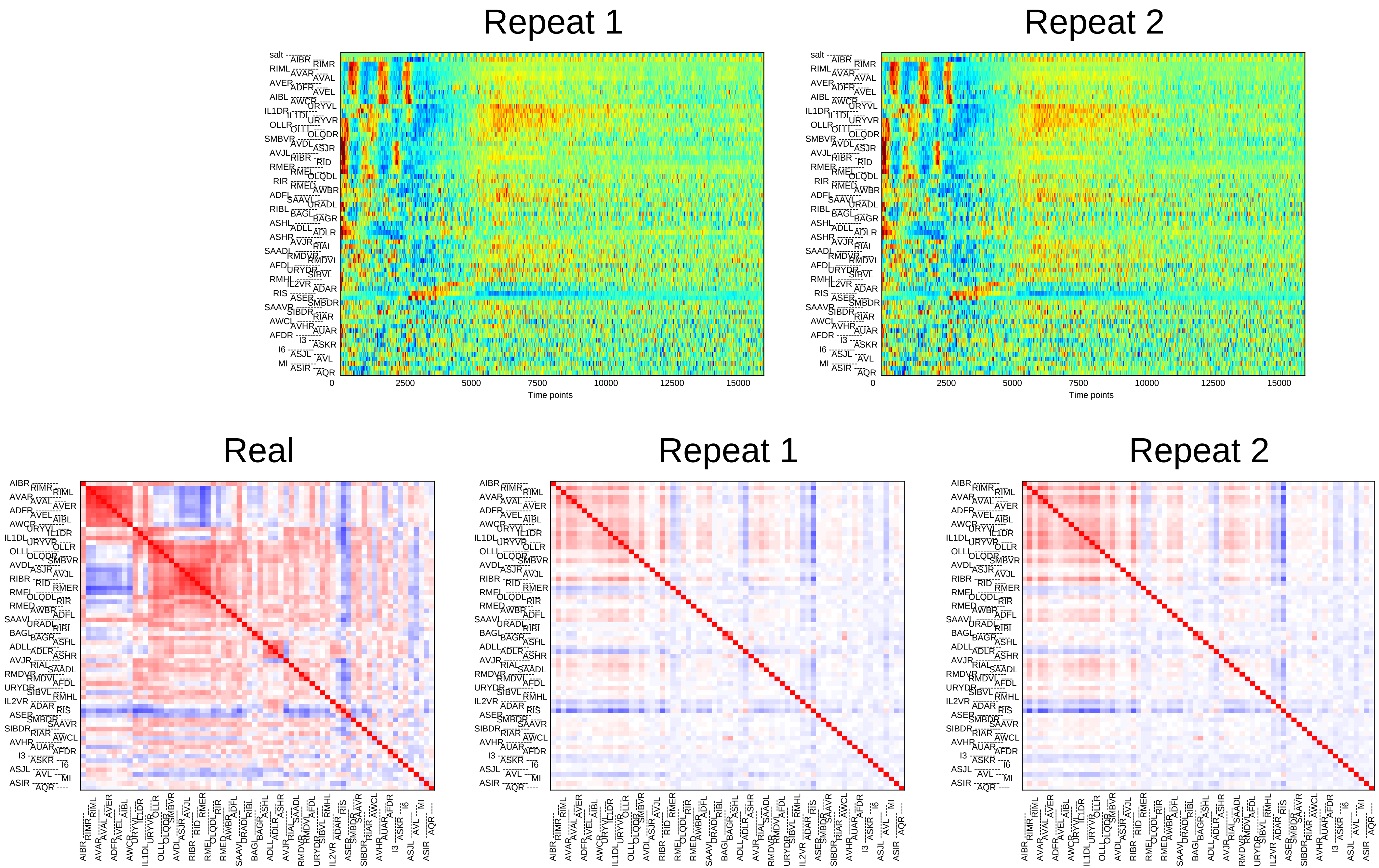
Repeat 1
Repeat 2
0
2500
5000
7500
10000
12500
15000
Time points
Real
Repeat 1
Repeat 2

### Sample24

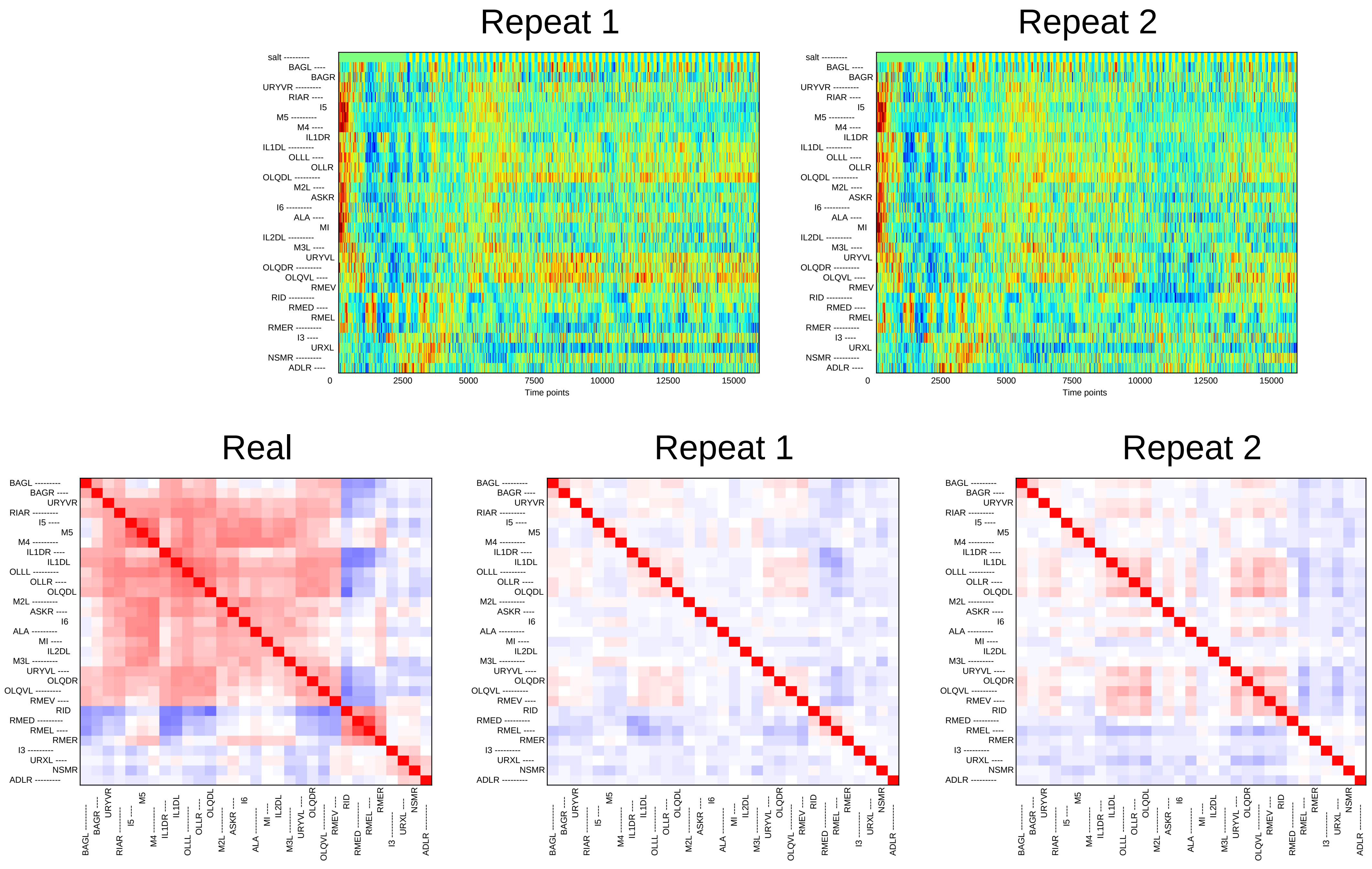
Repeat 1
Repeat 2
salt
BAGL
BAGR
URYVR
RIAR
I5
M5
M4
IL1DR
IL1DL
OLLL
OLLR
OLQDL
M2L
ASKR
I6
ALA
MI
IL2DL
M3L
URYVL
OLQDR
OLQVL
RMEV
RID
RMED
RMEL
RMER
I3
URXL
NSMR
ADLR
0
2500
5000
7500
10000
12500
15000
Time points
Real
Repeat 1
Repeat 2
