## Supplementary File 5C for "Deducing ensemble dynamics and information flow from the whole-brain imaging data"

### Sample1

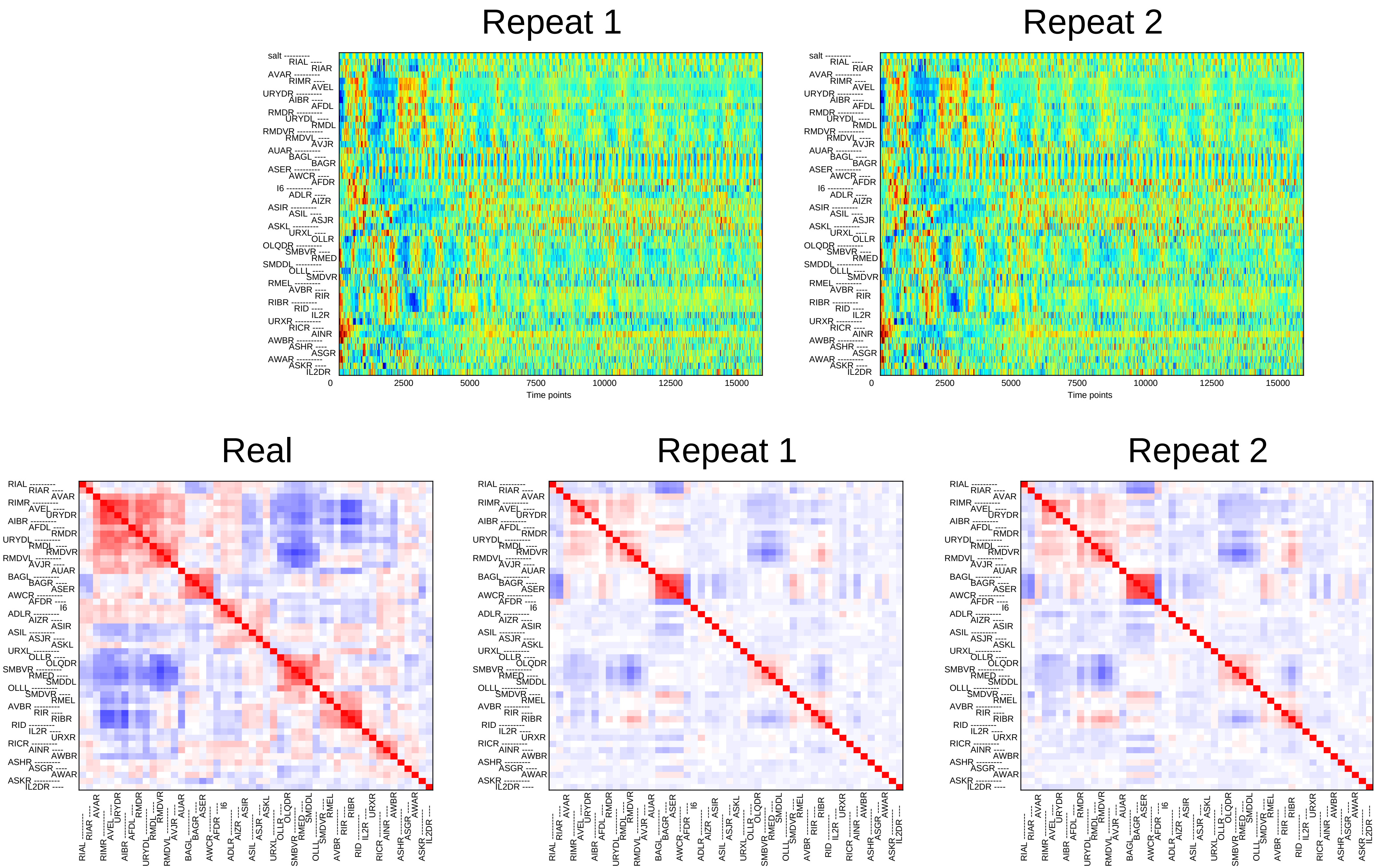
Repeat 1
Repeat 2
salt
RIAL RIAR AVAR RIMR AVEL URYDR AIBR AFDL RMDR URYDL RMDL RMDVR RMDVL AVJR AUAR BAGL BAGR ASER AWCR AFDR I6 ADLR AIZR ASIR ASIL ASJR ASKL URXL OLLR OLQDR SMBVR RMED SMDDL OLLL SMDVR RMEL AVBR RIR RIBR RID IL2R URXR RICR AINR AWBR ASHR ASGR AWAR ASKR IL2DR
0
2500
5000
7500
10000
12500
15000
Time points
Real
Repeat 1
Repeat 2

### Sample2

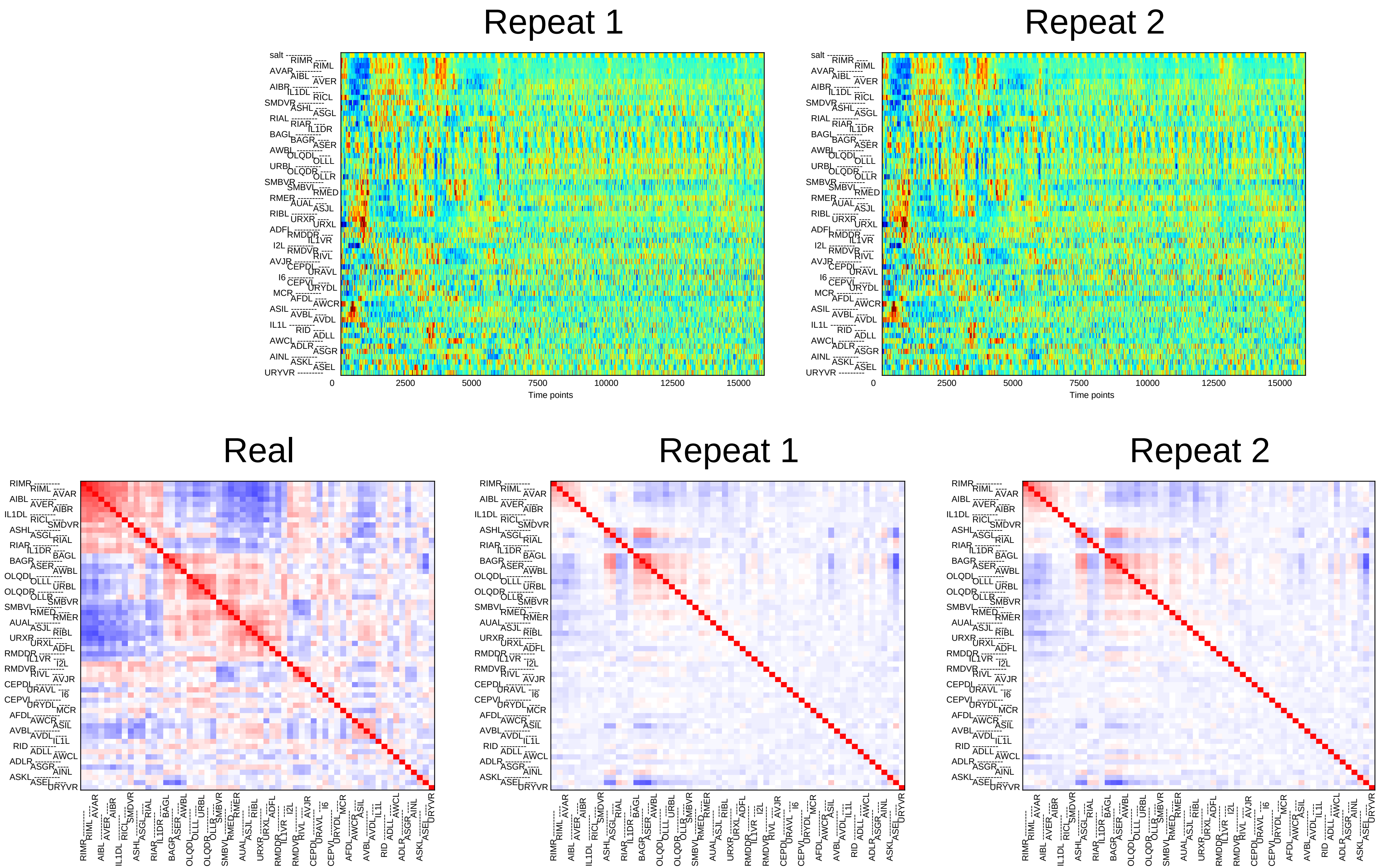

Repeat 1
Repeat 2
salt
0
2500
5000
7500
10000
12500
15000
Time points
Real
Repeat 1
Repeat 2

### Sample3

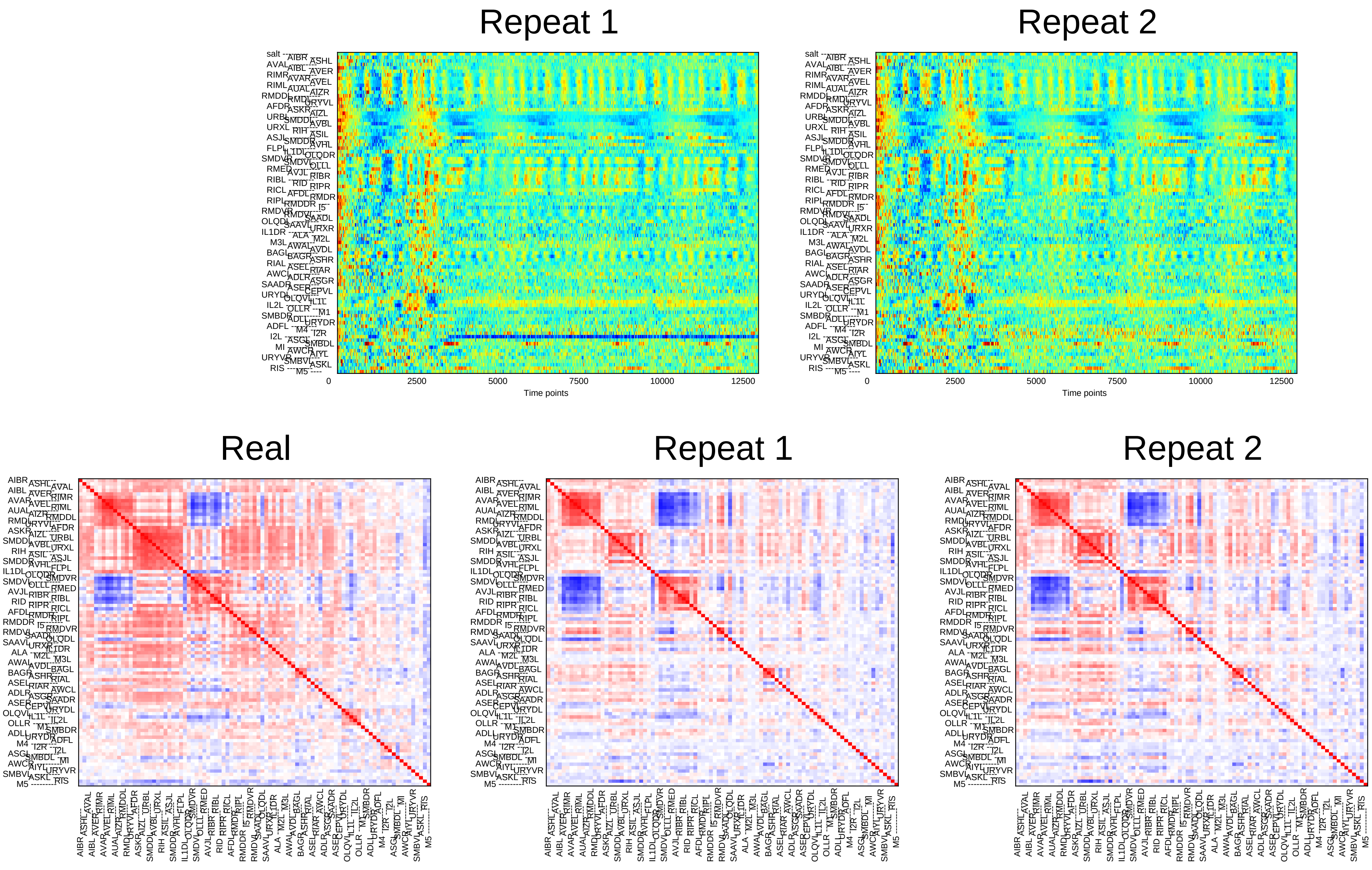
Repeat 1
Repeat 2
0
2500
5000
7500
10000
12500
Time points
Real
Repeat 1
Repeat 2

### Sample4

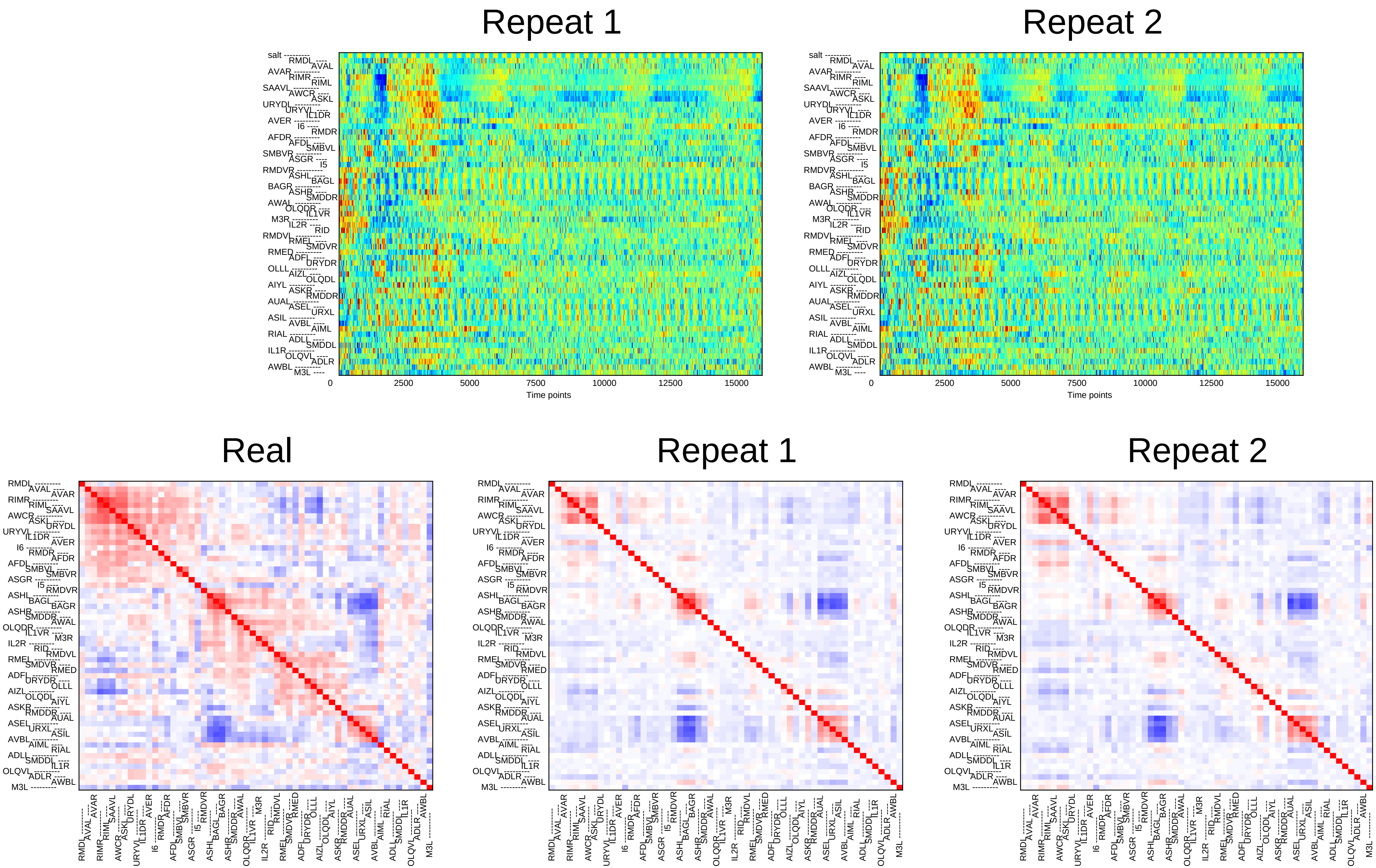

Repeat 1
Repeat 2
Time points
0
2500
5000
7500
10000
12500
15000
Real
Repeat 1
Repeat 2

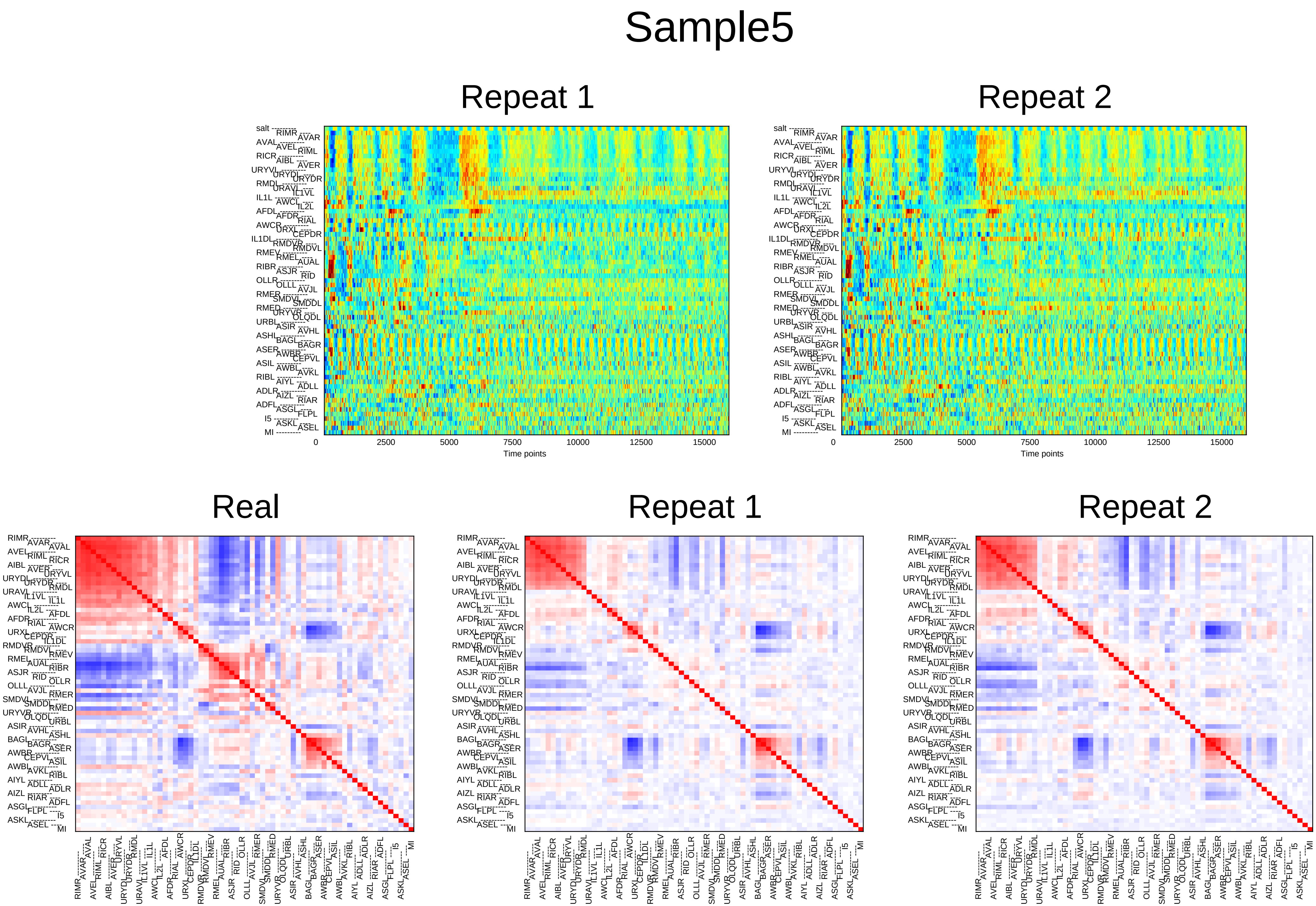
Sample5
Repeat 1
Repeat 2
Time points
0
2500
5000
7500
10000
12500
15000
Real
Repeat 1
Repeat 2

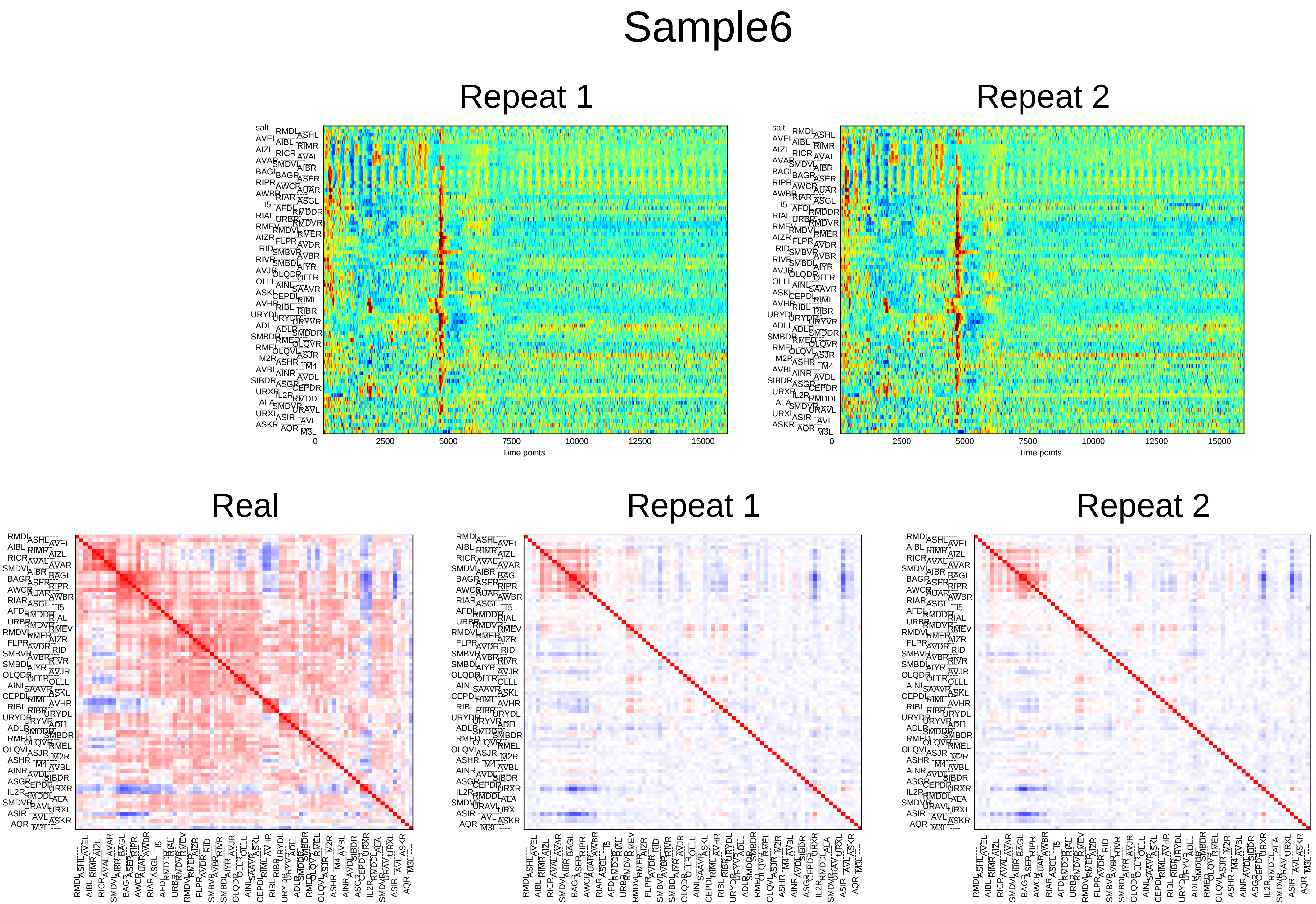

Sample6
Repeat 1
Repeat 2
0
2500
5000
7500
10000
12500
15000
Time points
Real
Repeat 1
Repeat 2

### Sample7

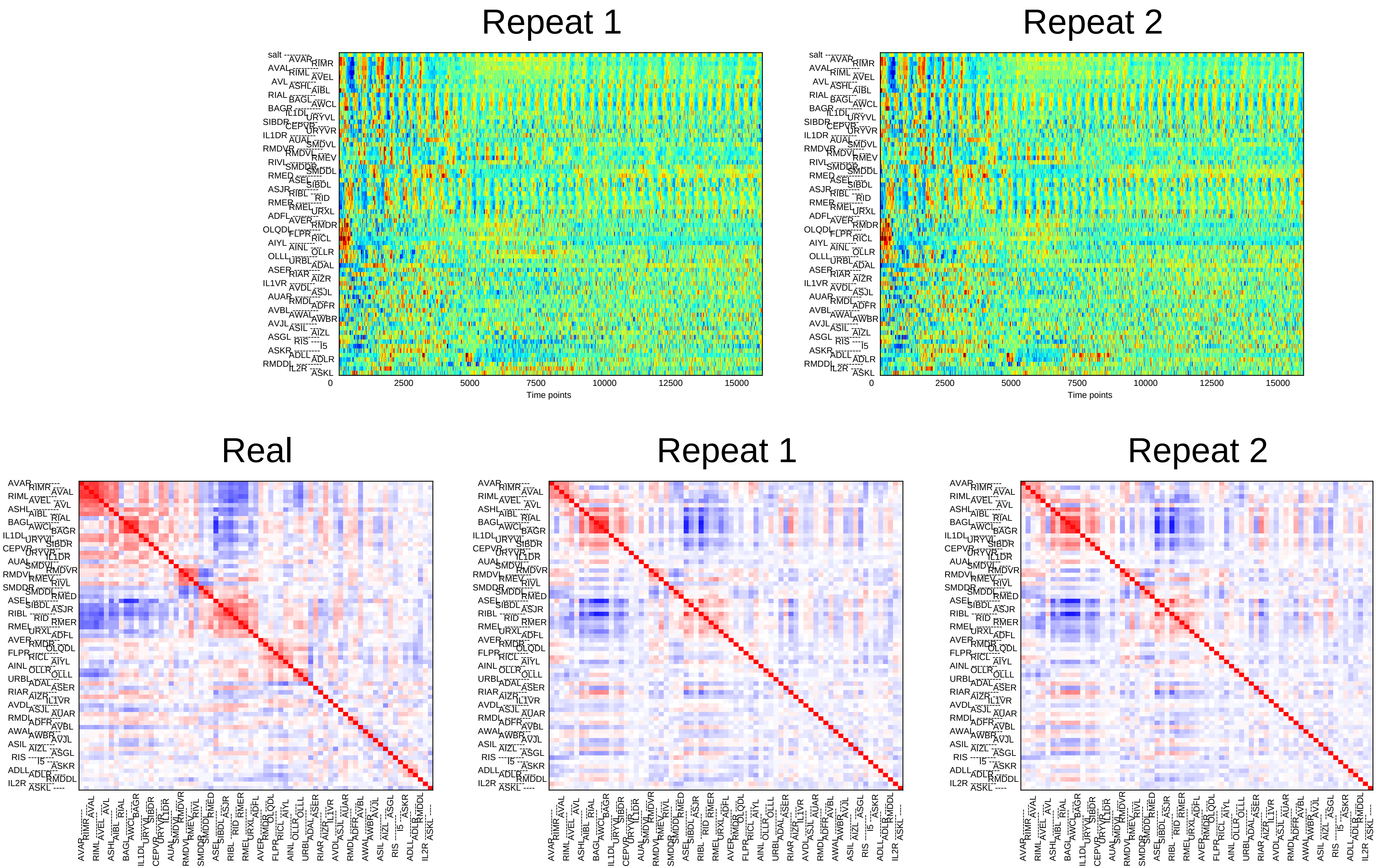

Repeat 1
Repeat 2
0
2500
5000
7500
10000
12500
15000
Time points
Real
Repeat 1
Repeat 2

### Sample8

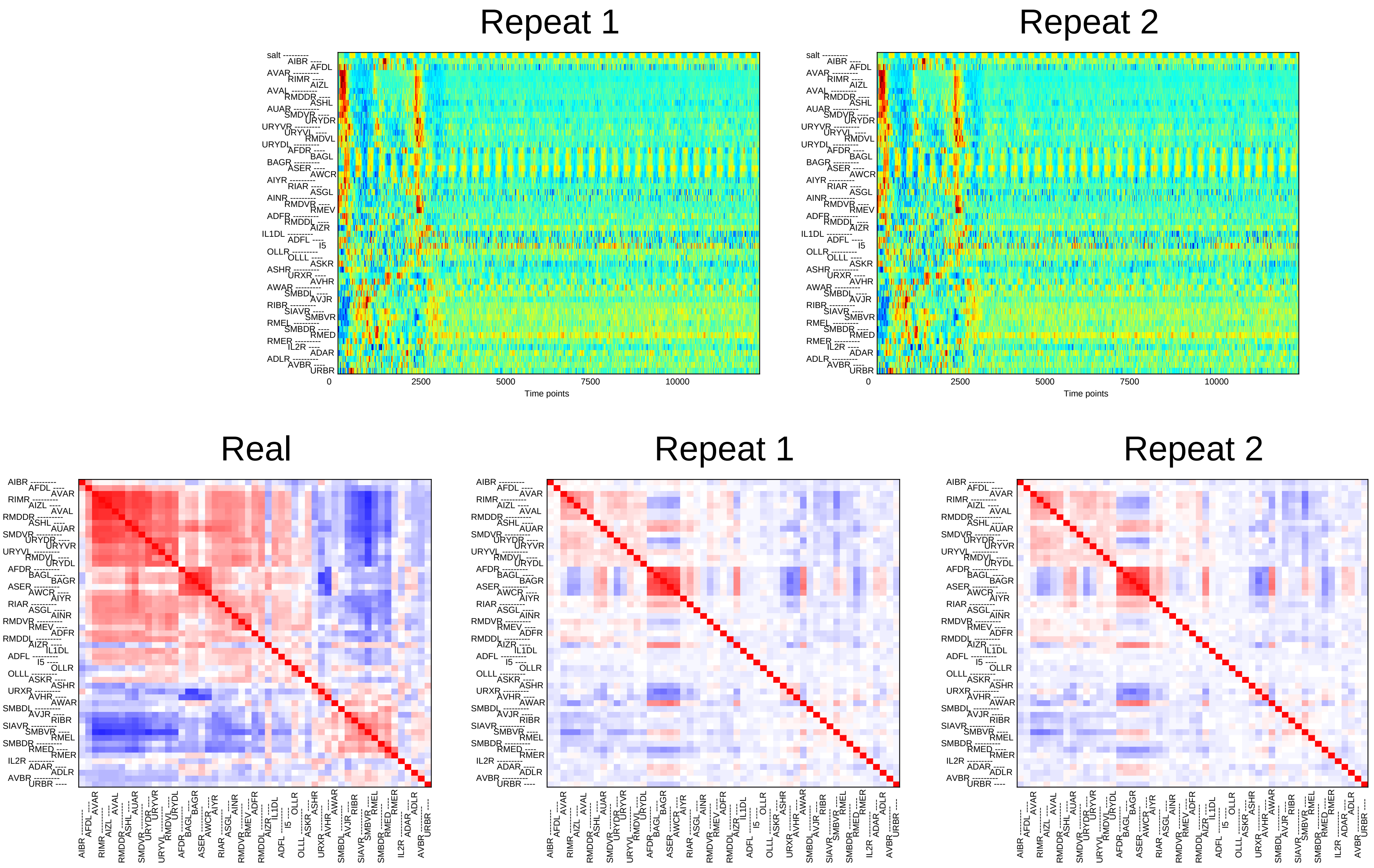
Repeat 1
Repeat 2
0
2500
5000
7500
10000
Time points
Real
Repeat 1
Repeat 2

### Sample9

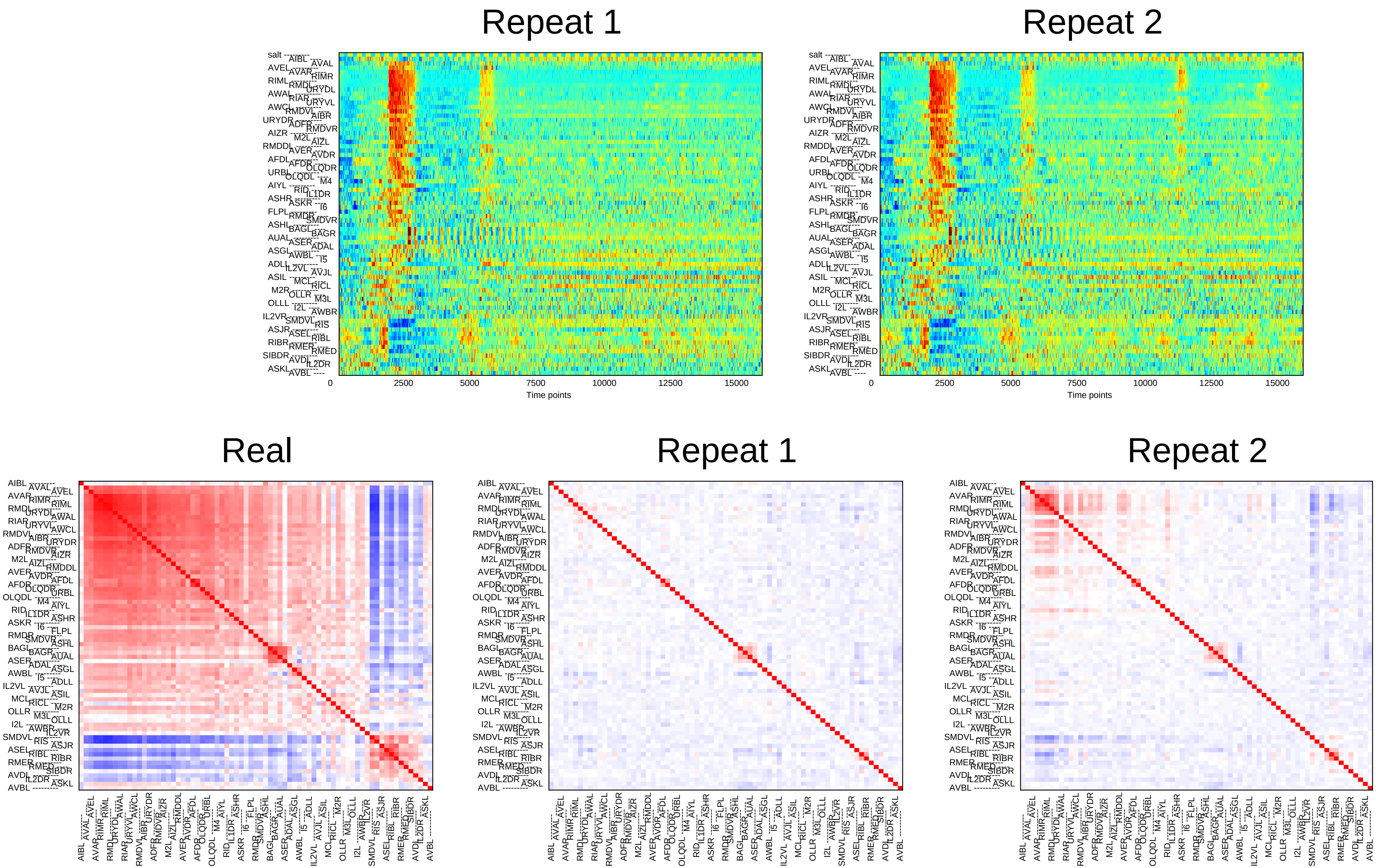
Repeat 1
Repeat 2
0
2500
5000
7500
10000
12500
15000
Time points
Real
Repeat 1
Repeat 2

### Sample10

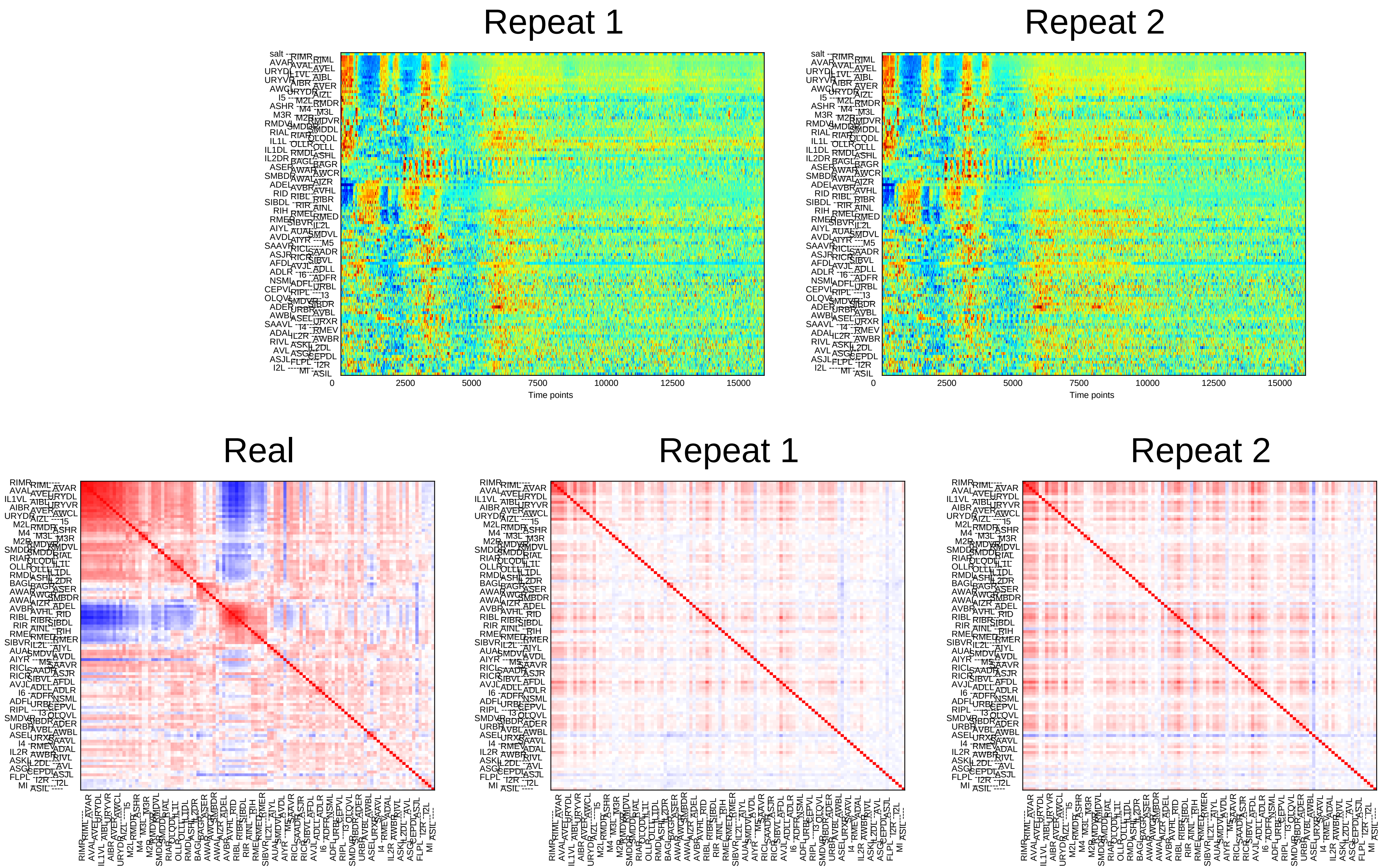

Repeat 1
Repeat 2
0
2500
5000
7500
10000
12500
15000
Time points
Real
Repeat 1
Repeat 2

### Sample11

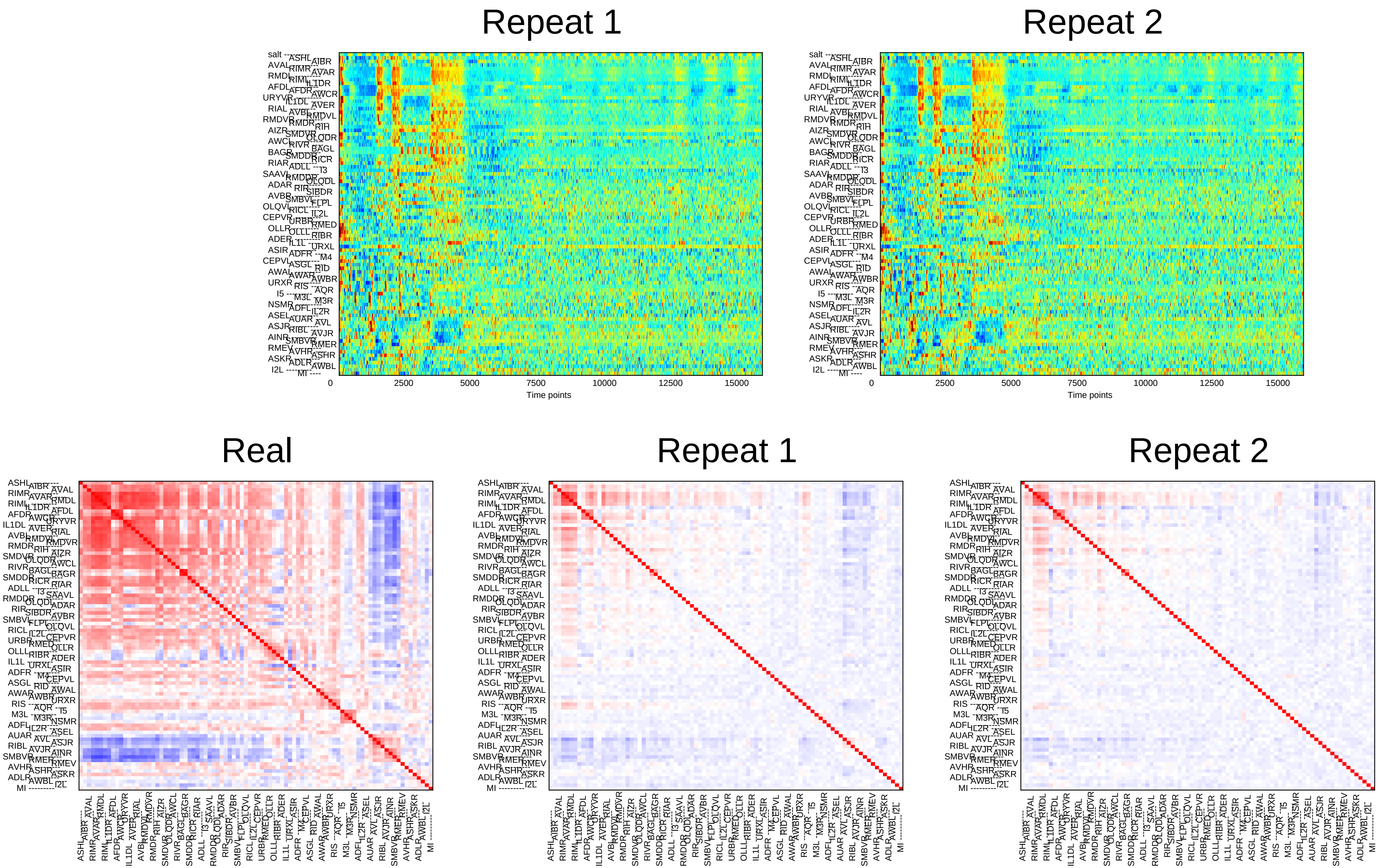

Repeat 1
Repeat 2
0
2500
5000
7500
10000
12500
15000
Time points
Real
Repeat 1
Repeat 2

### Sample12

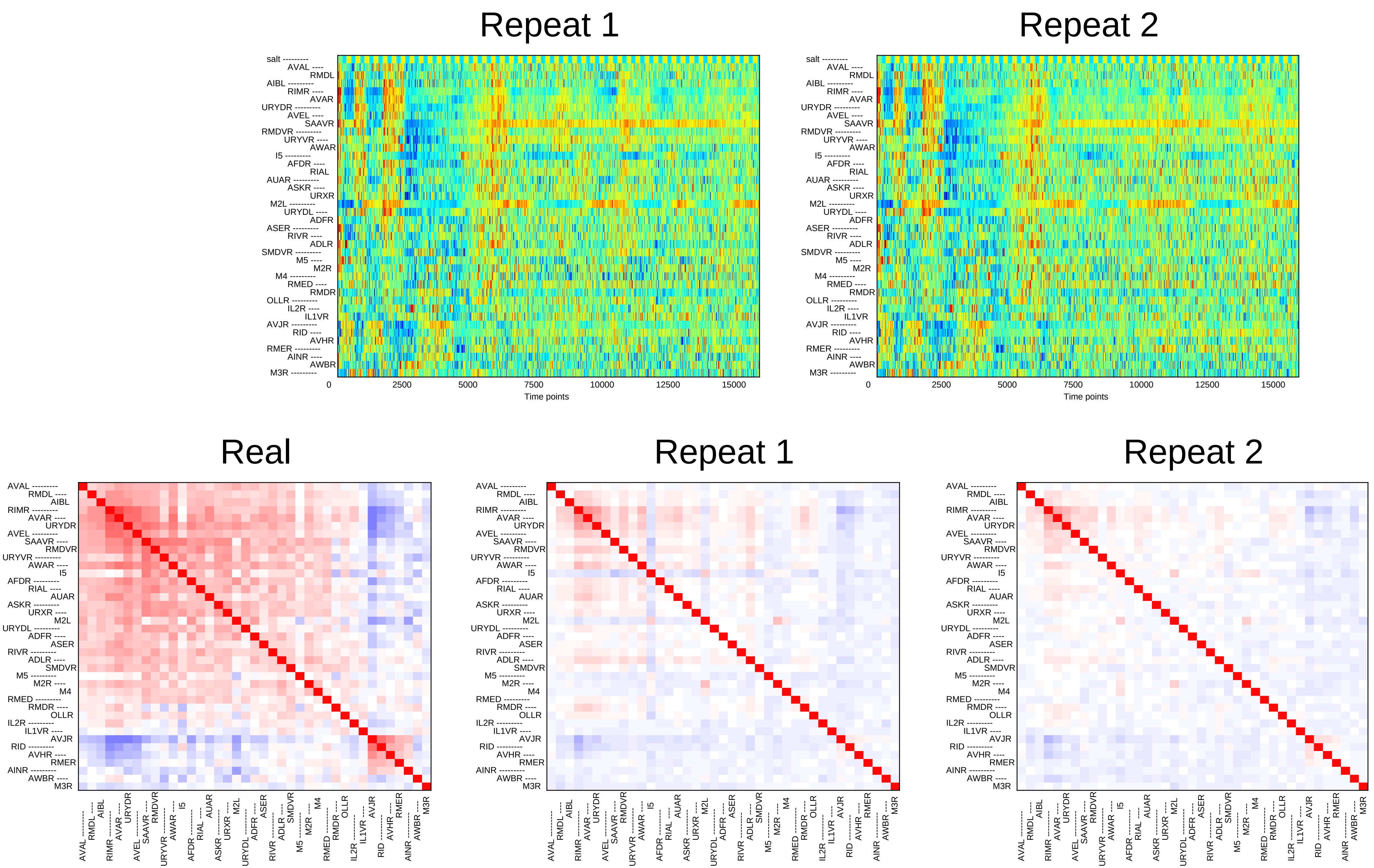

Repeat 1
Repeat 2
salt
AVAL
RMDL
AIBL
RIMR
AVAR
URYDR
AVEL
SAAVR
RMDVR
URYVR
AWAR
I5
AFDR
RIAL
AUAR
ASKR
URXR
M2L
URYDL
ADFR
ASER
RIVR
ADLR
SMDVR
M5
M2R
M4
RMED
RMDR
OLLR
IL2R
IL1VR
AVJR
RID
AVHR
RMER
AINR
AWBR
M3R
0
2500
5000
7500
10000
12500
15000
Time points
Real
Repeat 1
Repeat 2

### Sample13

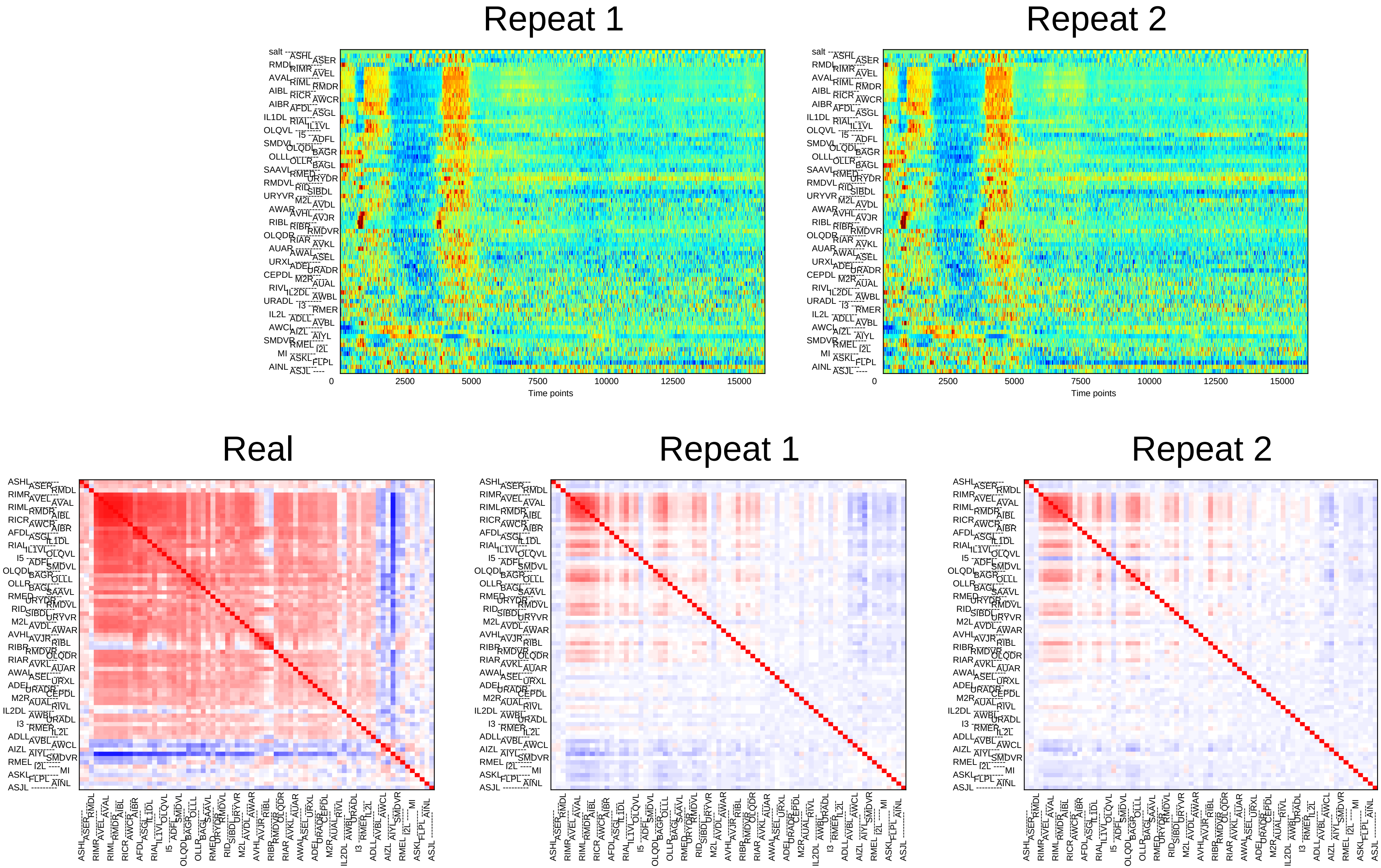

Repeat 1
Repeat 2
0
2500
5000
7500
10000
12500
15000
Time points
Real
Repeat 1
Repeat 2

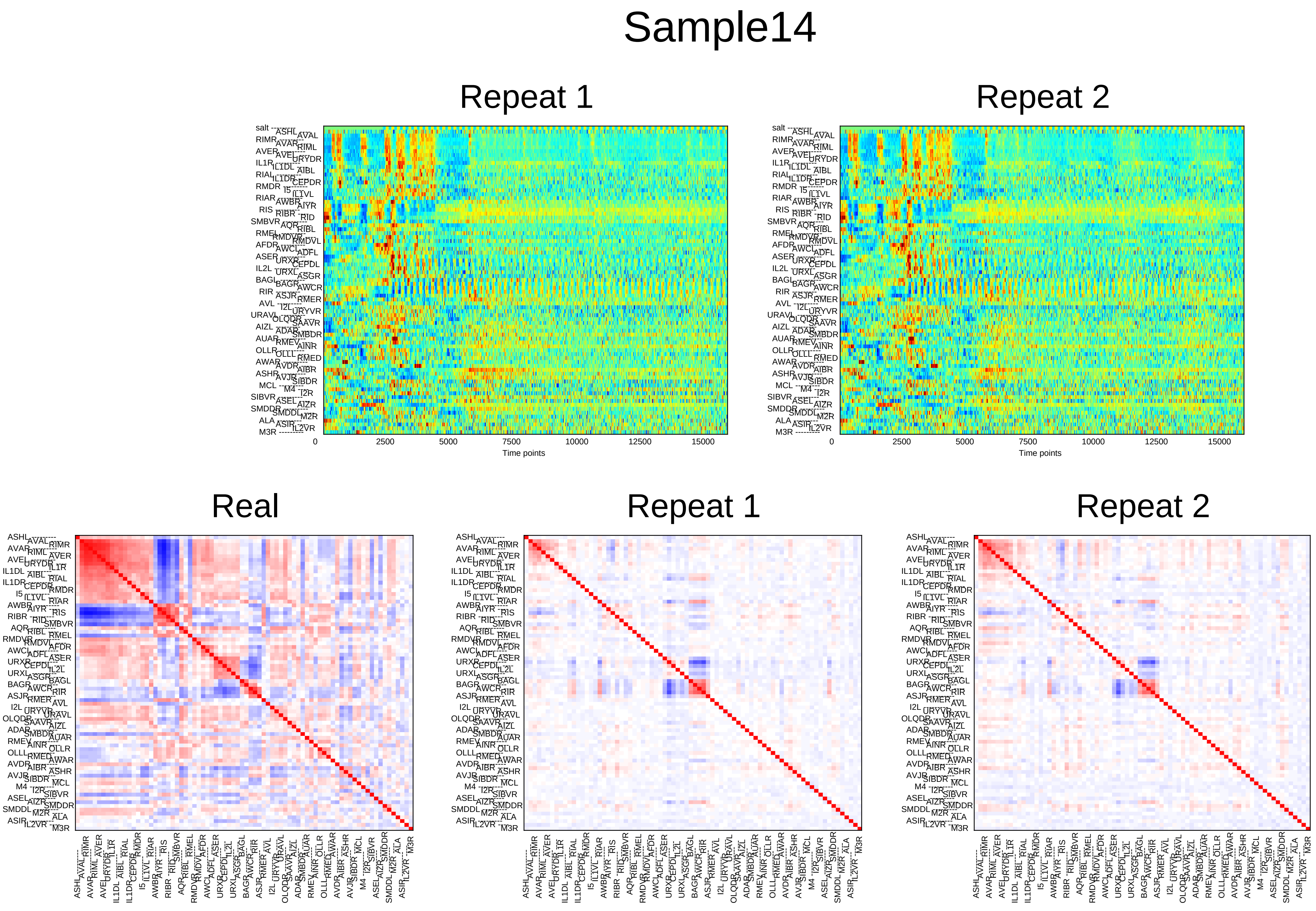
Sample14
Repeat 1
Repeat 2
0
2500
5000
7500
10000
12500
15000
Time points
Real
Repeat 1
Repeat 2

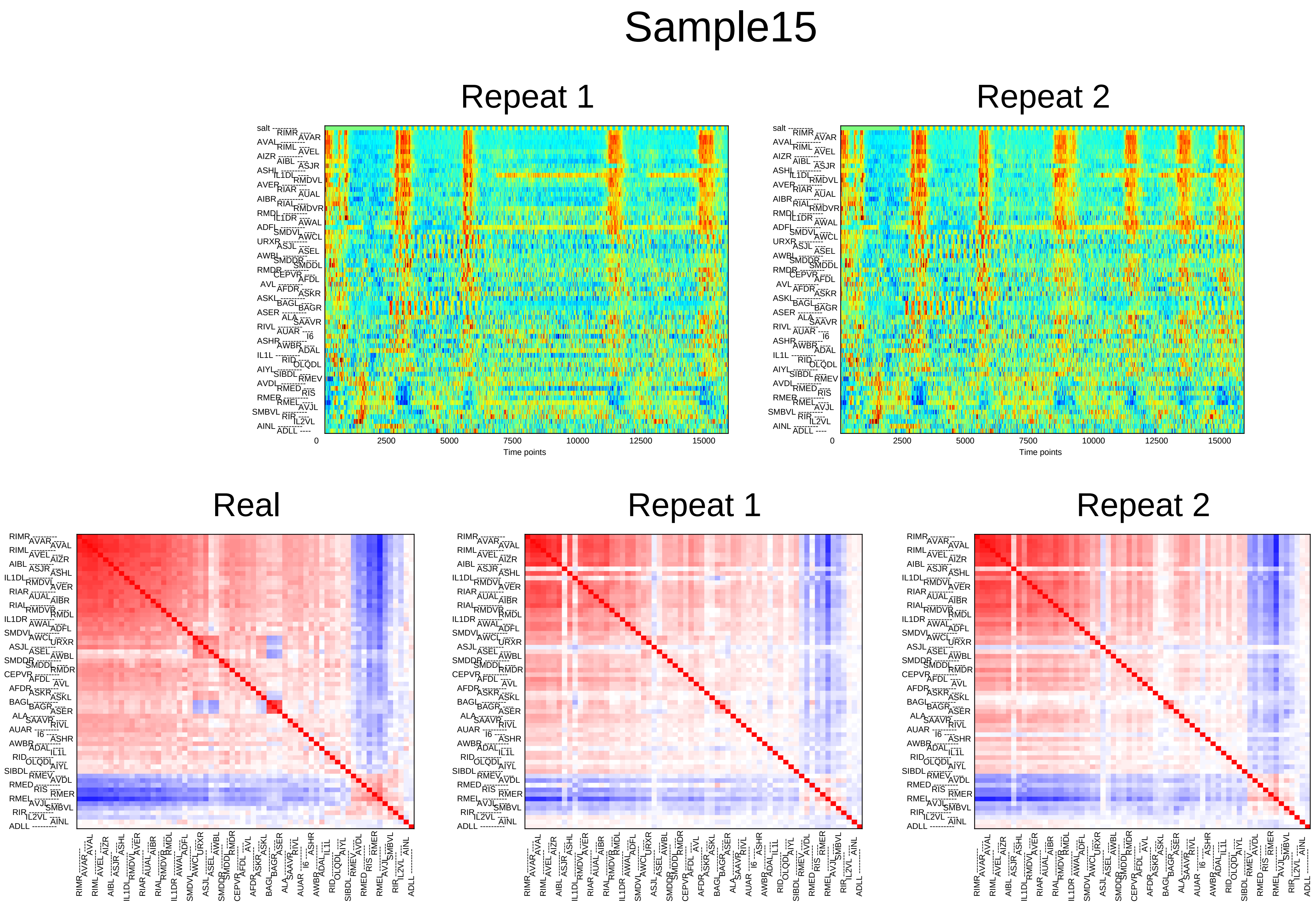
Sample15
Repeat 1
Repeat 2
0
2500
5000
7500
10000
12500
15000
Time points
Real
Repeat 1
Repeat 2

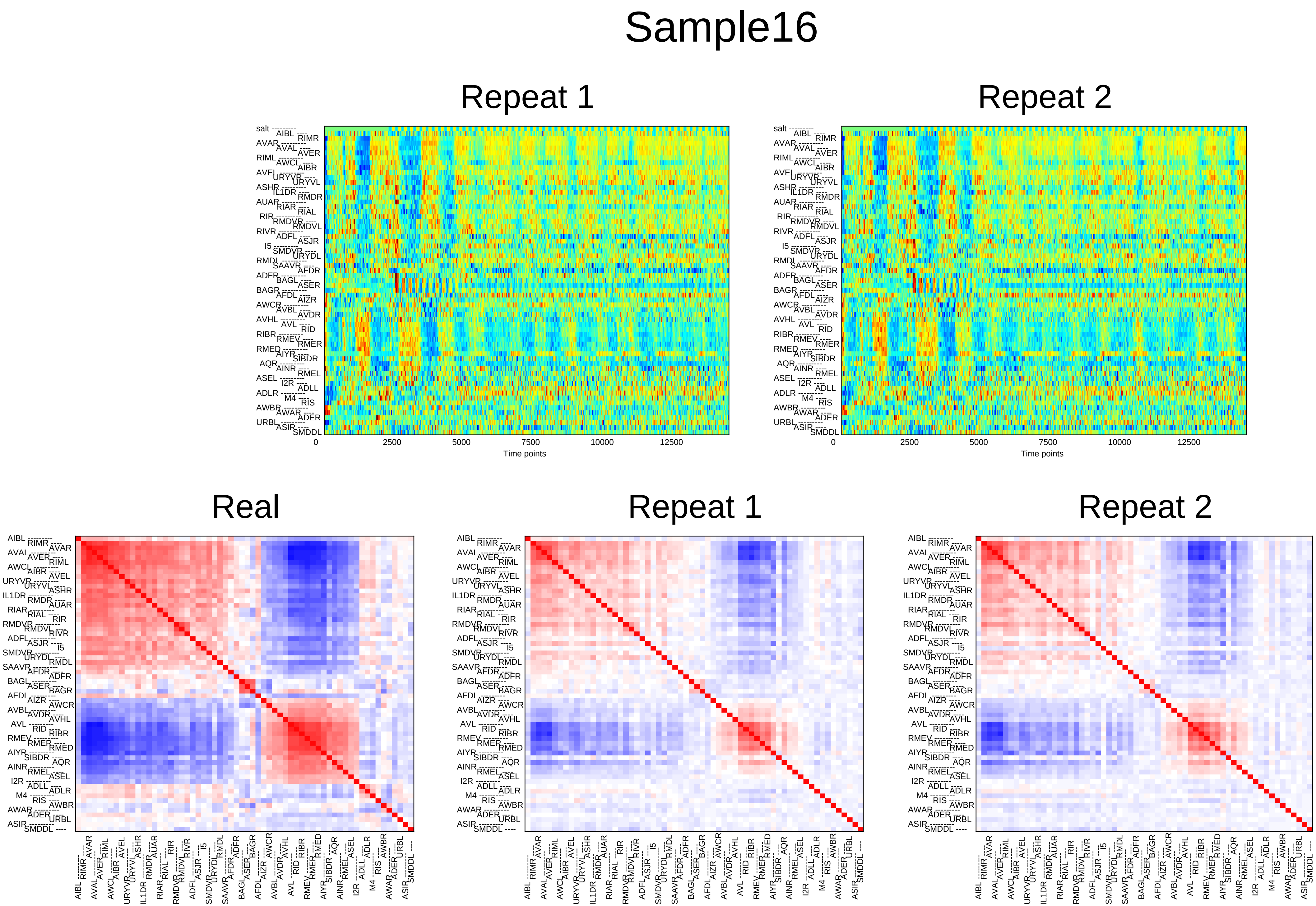

Sample16
Repeat 1
Repeat 2
0
2500
5000
7500
10000
12500
Time points
Real
Repeat 1
Repeat 2

### Sample17

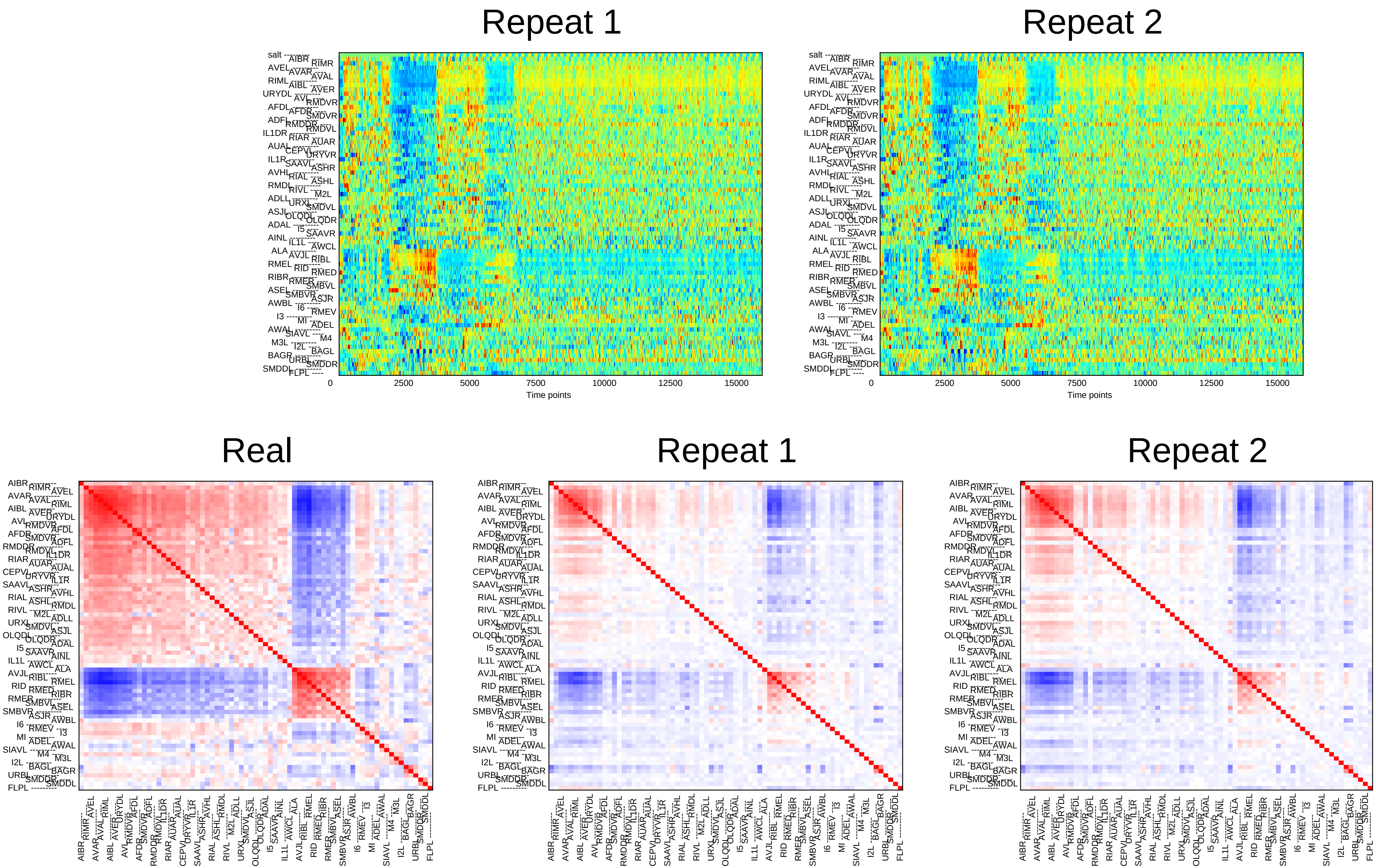

Repeat 1
Repeat 2
0
2500
5000
7500
10000
12500
15000
Time points
Real
Repeat 1
Repeat 2

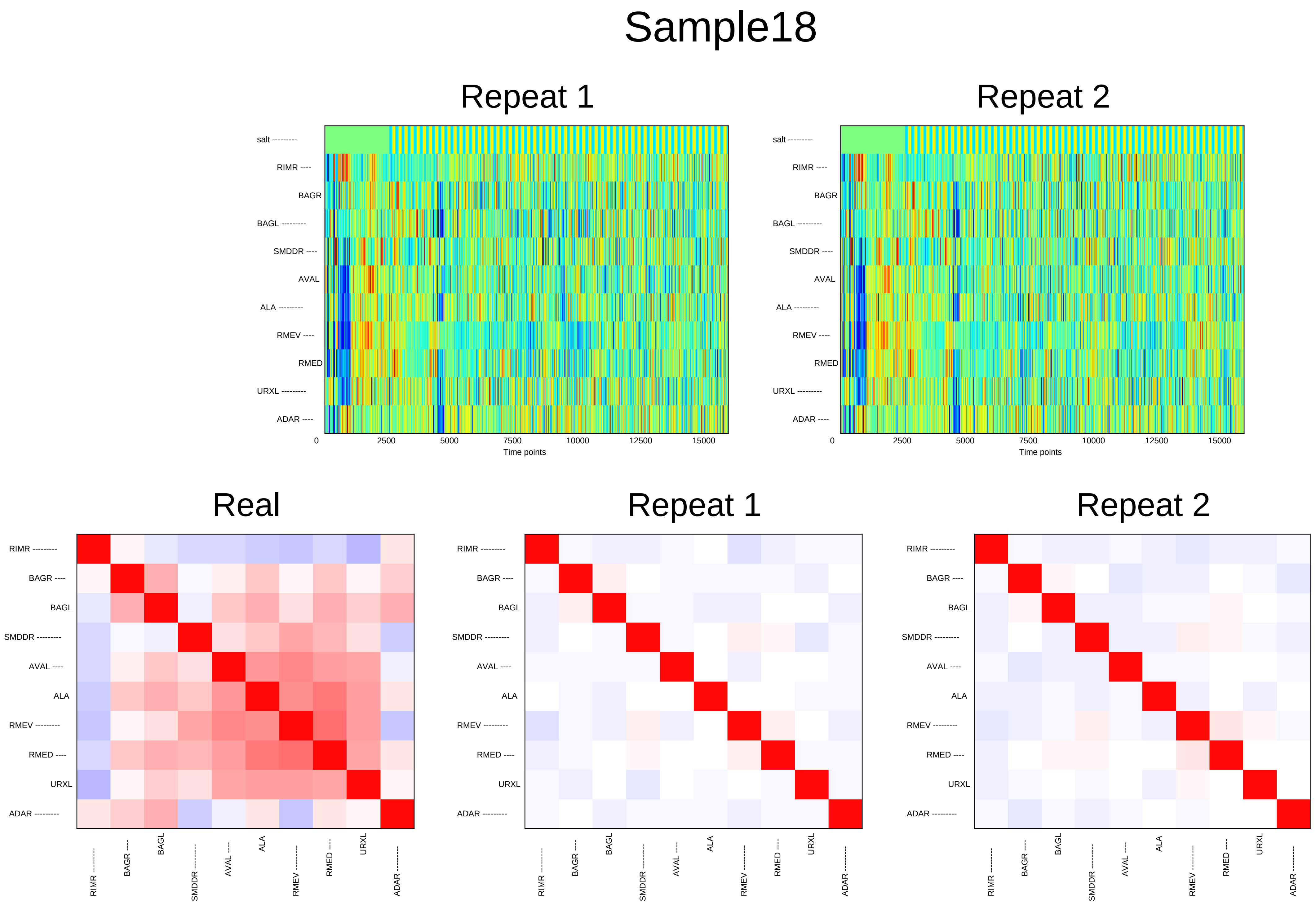
Sample18
Repeat 1
Repeat 2
salt
RIMR
BAGR
BAGL
SMDDR
AVAL
ALA
RMEV
RMED
URXL
ADAR
0
2500
5000
7500
10000
12500
15000
Time points
Real
Repeat 1
Repeat 2

### Sample19

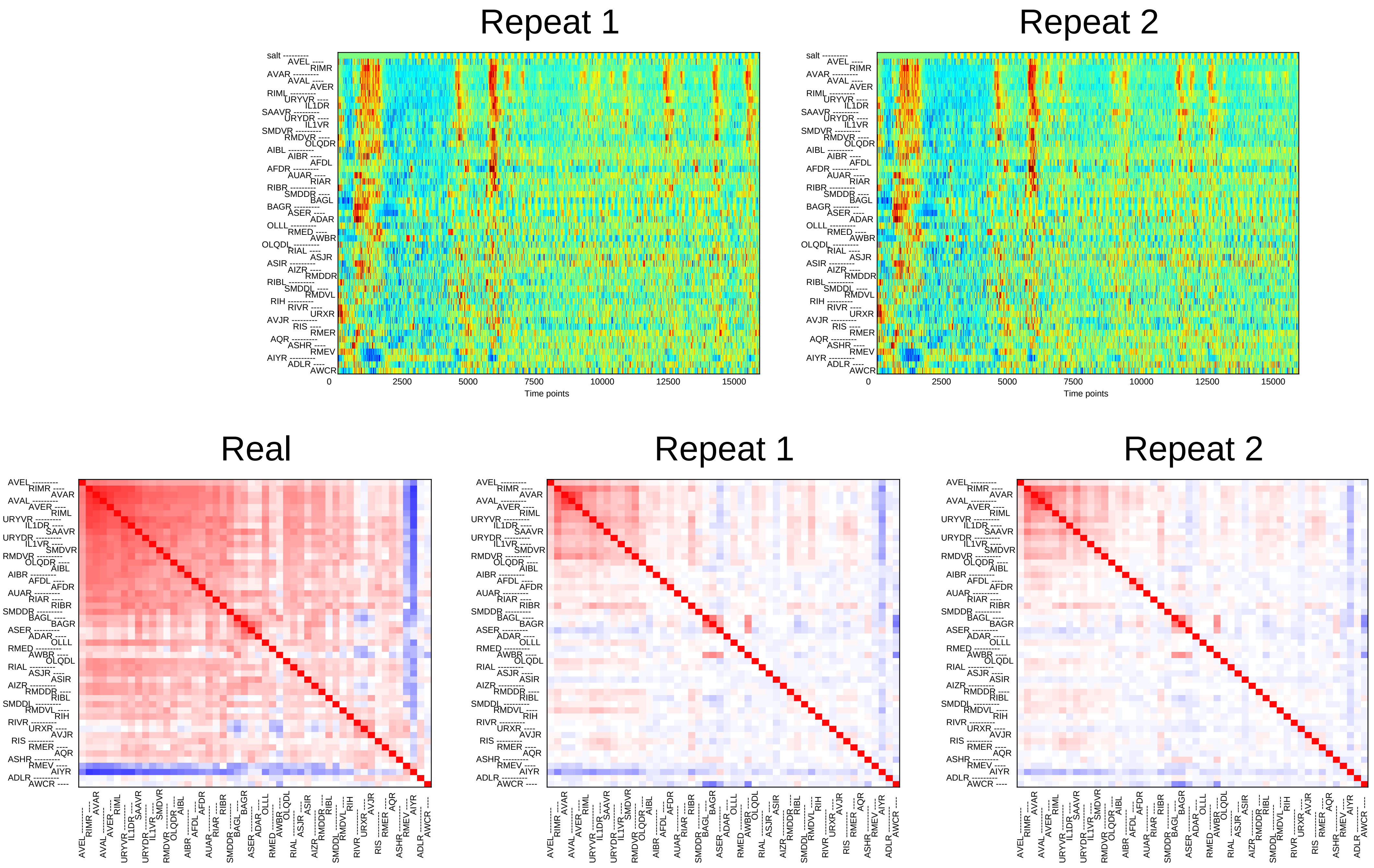
Repeat 1
Repeat 2
salt
AVEL RIMR AVAR AVAL AVER RIML URYVR IL1DR SAAVR URYDR IL1VR SMDVR RMDVR OLQDR AIBL AIBR AFDL AFDR AUAR RIAR RIBR SMDDR BAGL BAGR ASER ADAR OLLL RMED AWBR OLQDL RIAL ASJR ASIR AIZR RMDDR RIBL SMDDL RMDVL RIH RIVR URXR AVJR RIS RMER AQR ASHR RMEV AIYR ADLR AWCR
0
2500
5000
7500
10000
12500
15000
Time points
Real
Repeat 1
Repeat 2

### Sample20

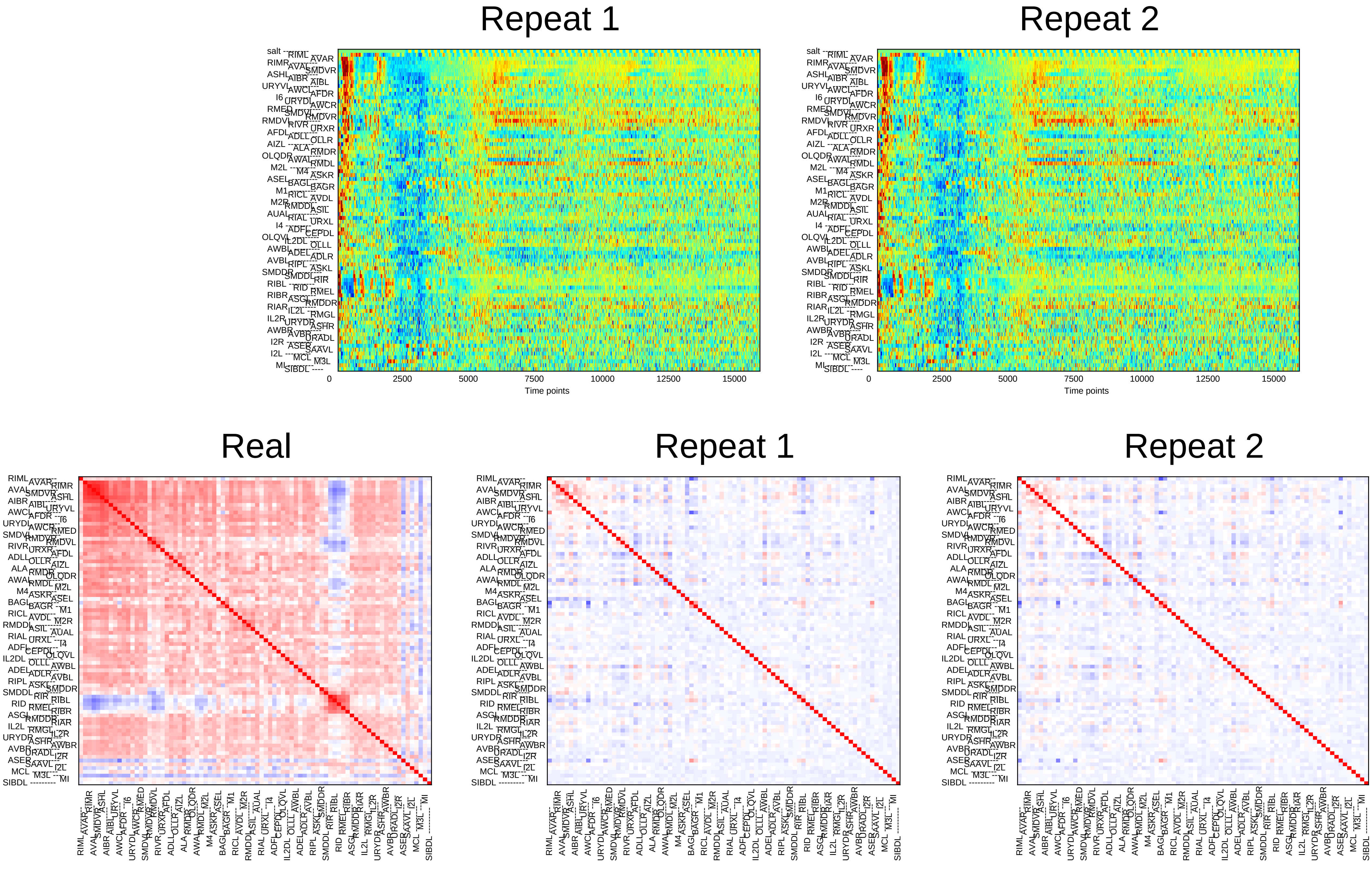

Repeat 1
Repeat 2
salt
0
2500
5000
7500
10000
12500
15000
Time points
Real
Repeat 1
Repeat 2

### Sample21

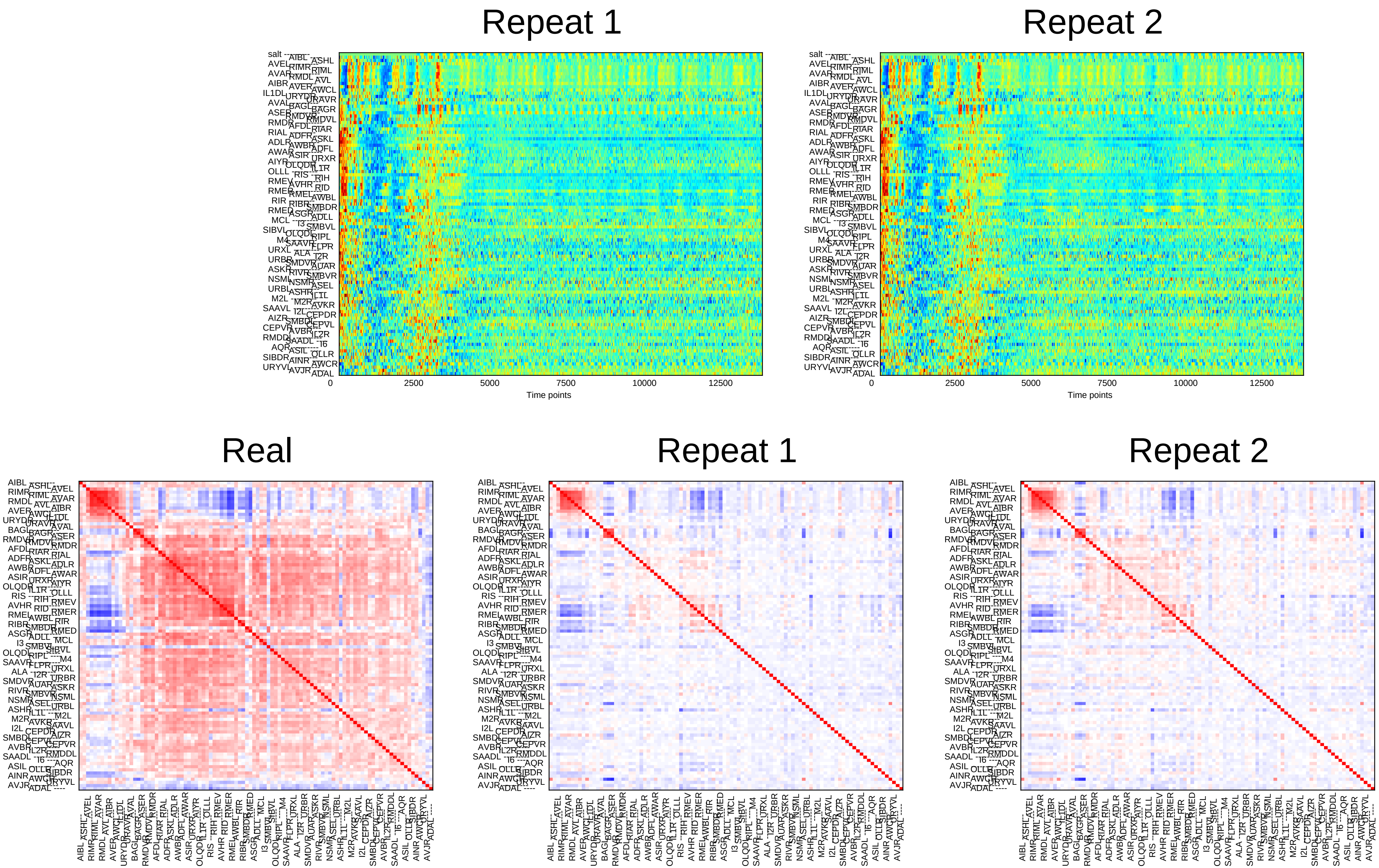
Repeat 1
Repeat 2
0
2500
5000
7500
10000
12500
Time points
Real
Repeat 1
Repeat 2

### Sample22

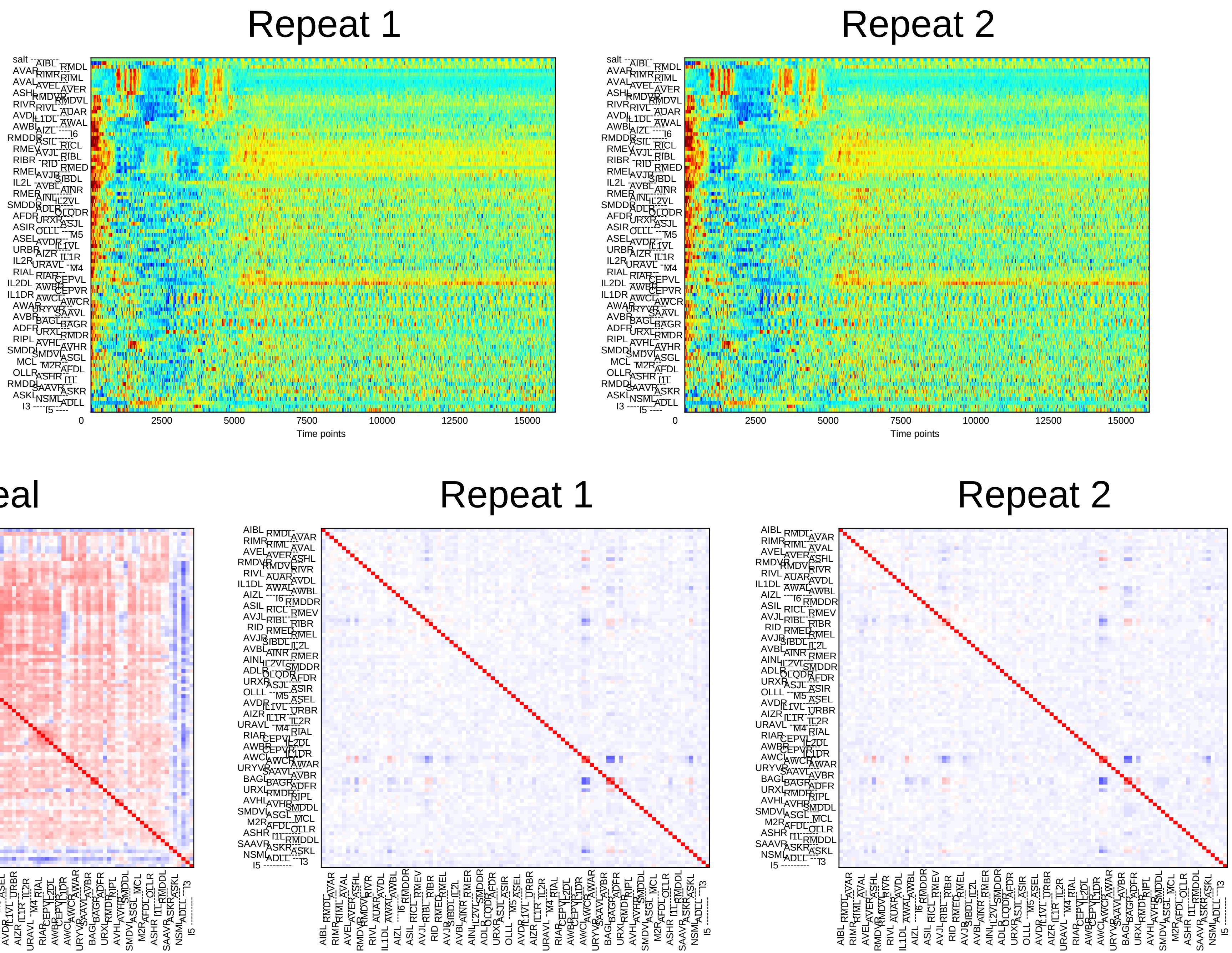

Repeat 1
Repeat 2
0
2500
5000
7500
10000
12500
15000
Time points
Real
Repeat 1
Repeat 2

### Sample23

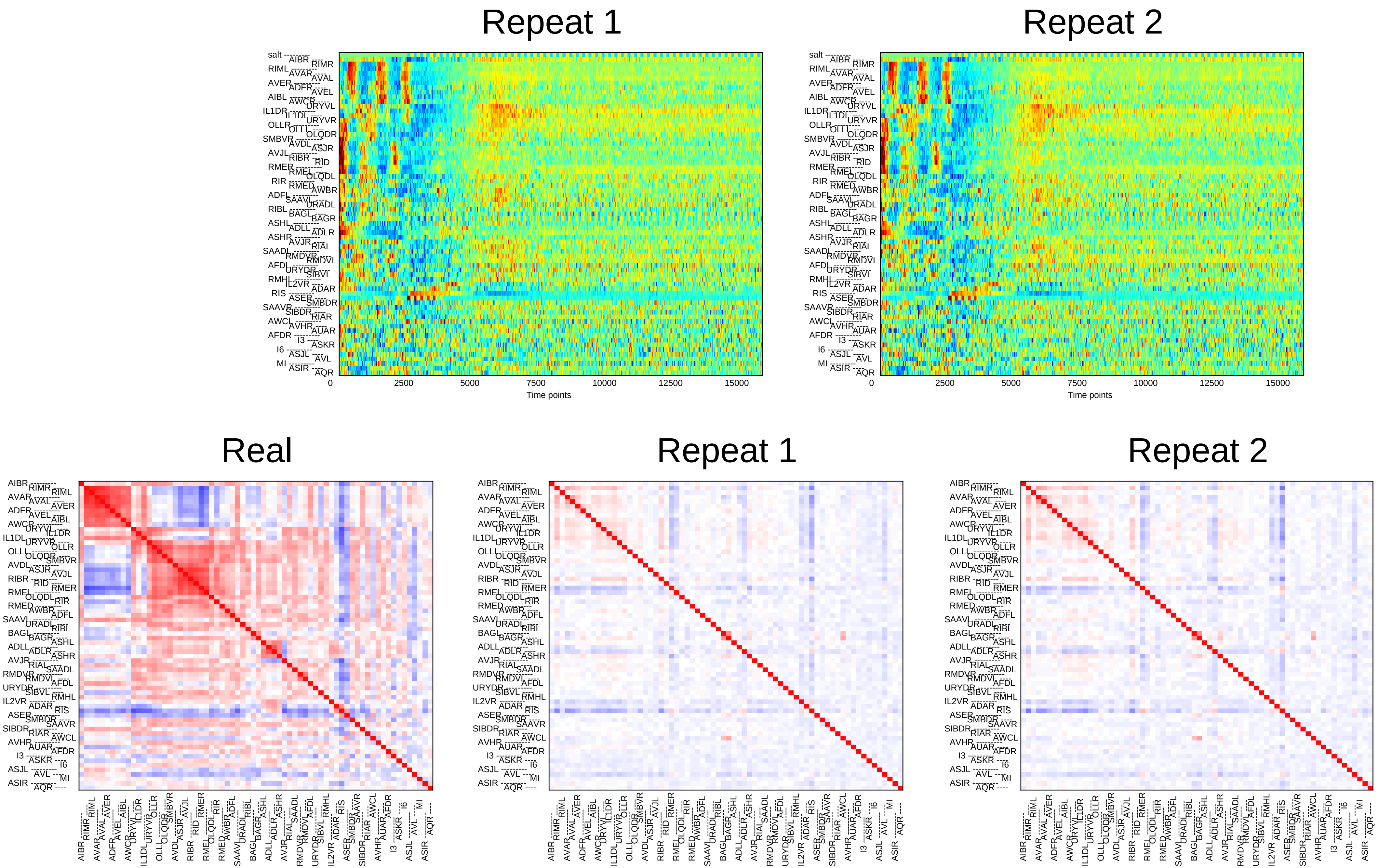

Repeat 1
Repeat 2
0
2500
5000
7500
10000
12500
15000
Time points
Real
Repeat 1
Repeat 2

### Sample24

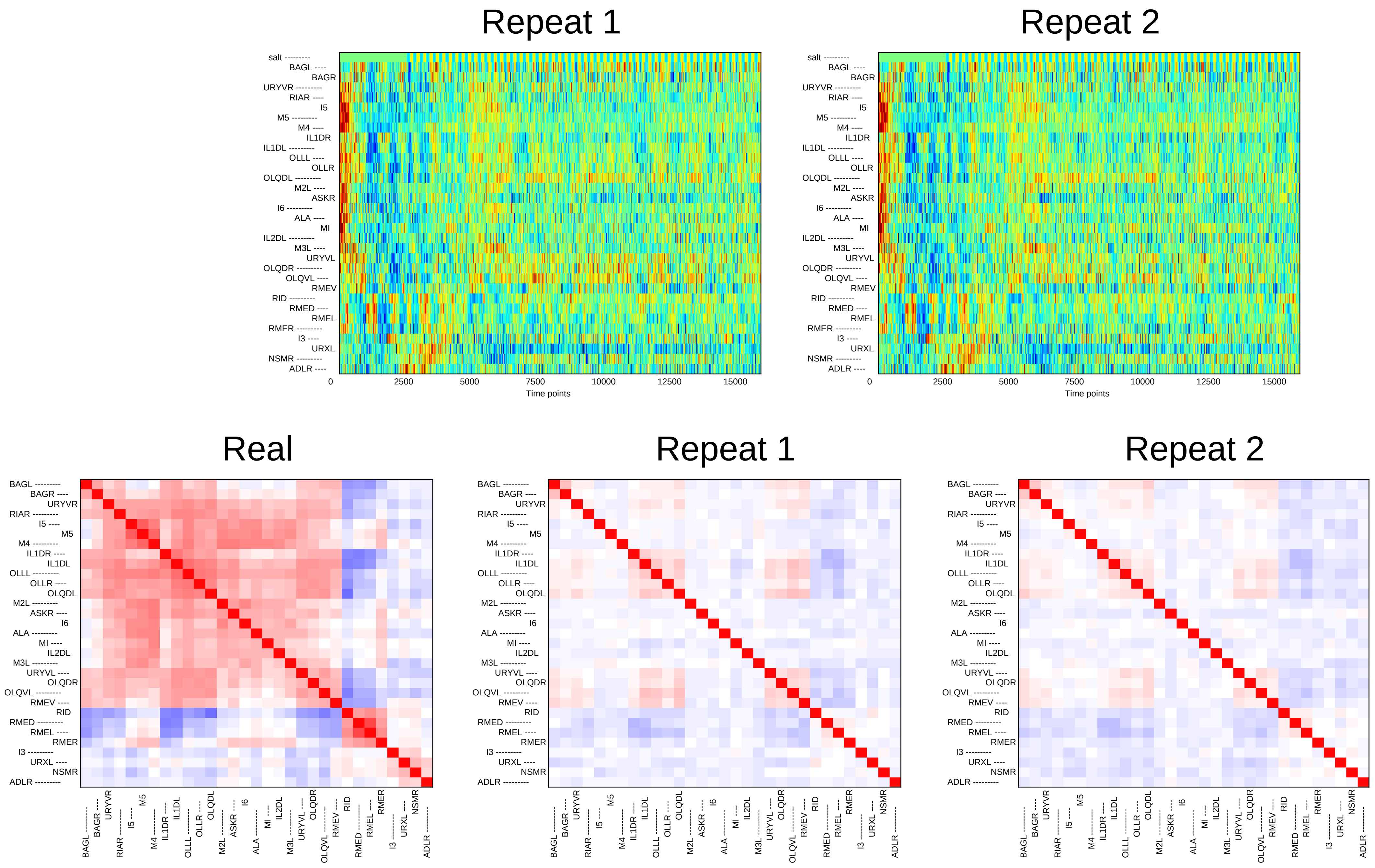
Repeat 1
Repeat 2
salt
BAGL
BAGR
URYVR
RIAR
I5
M5
M4
IL1DR
IL1DL
OLLL
OLLR
OLQDL
M2L
ASKR
I6
ALA
MI
IL2DL
M3L
URYVL
OLQDR
OLQVL
RMEV
RID
RMED
RMEL
RMER
I3
URXL
NSMR
ADLR
0
2500
5000
7500
10000
12500
15000
Time points
Real
Repeat 1
Repeat 2
