## Supplementary File 6B for "Deducing ensemble dynamics and information flow from the whole-brain imaging data"

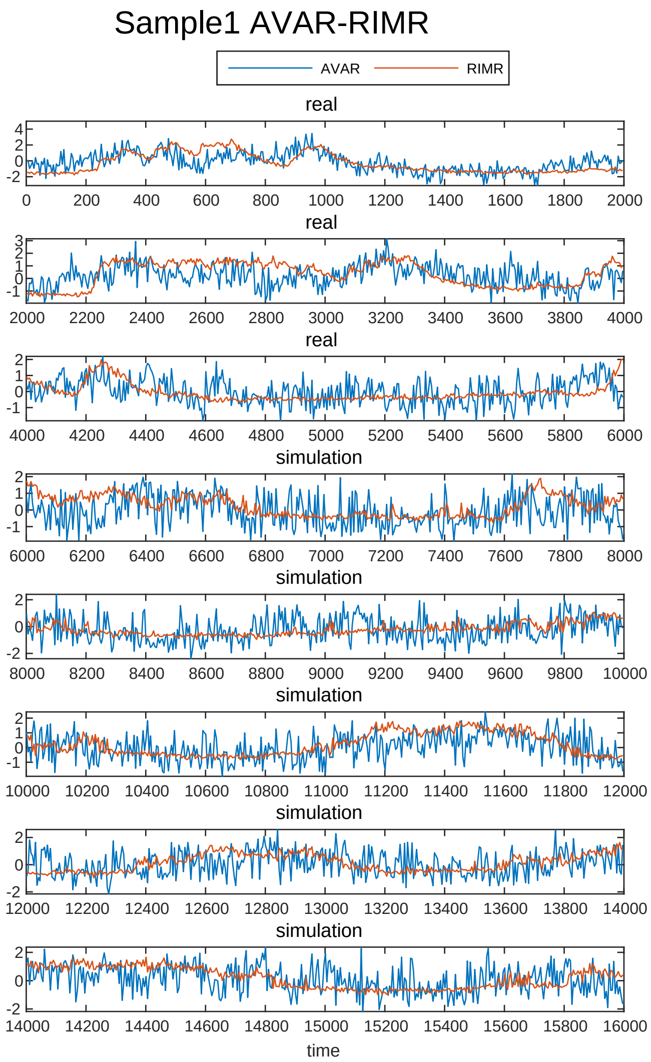
Sample1 AVAR-RIMR
AVAR
RIMR
real
real
real
simulation
simulation
simulation
simulation
simulation
time

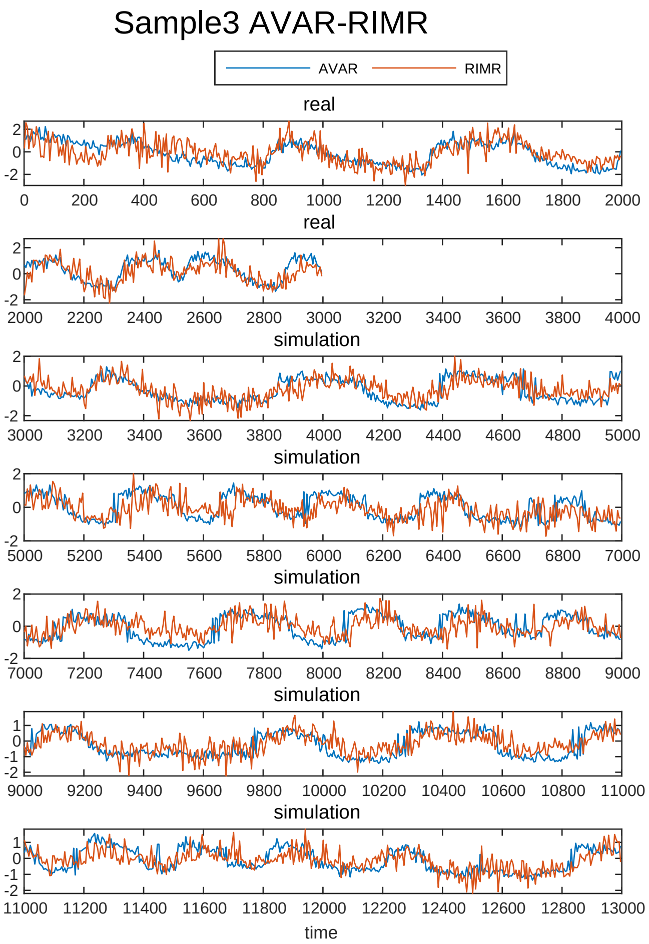
Sample3 AVAR-RIMR
AVAR
RIMR
real
real
simulation
simulation
simulation
simulation
simulation
time

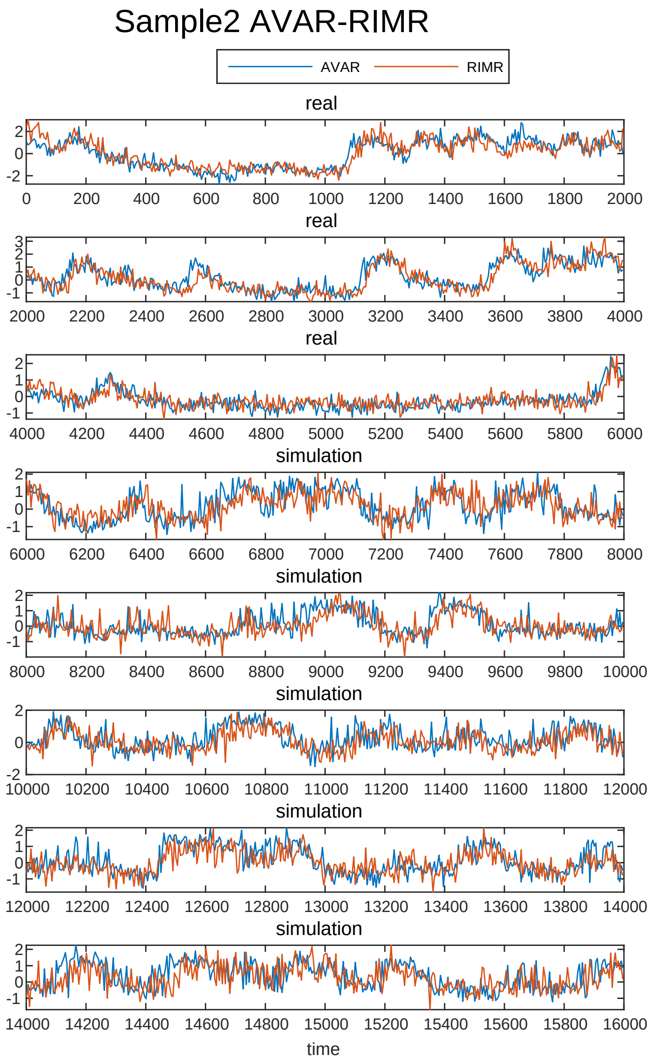
Sample2 AVAR-RIMR
AVAR
RIMR
real
real
real
simulation
simulation
simulation
simulation
simulation
time

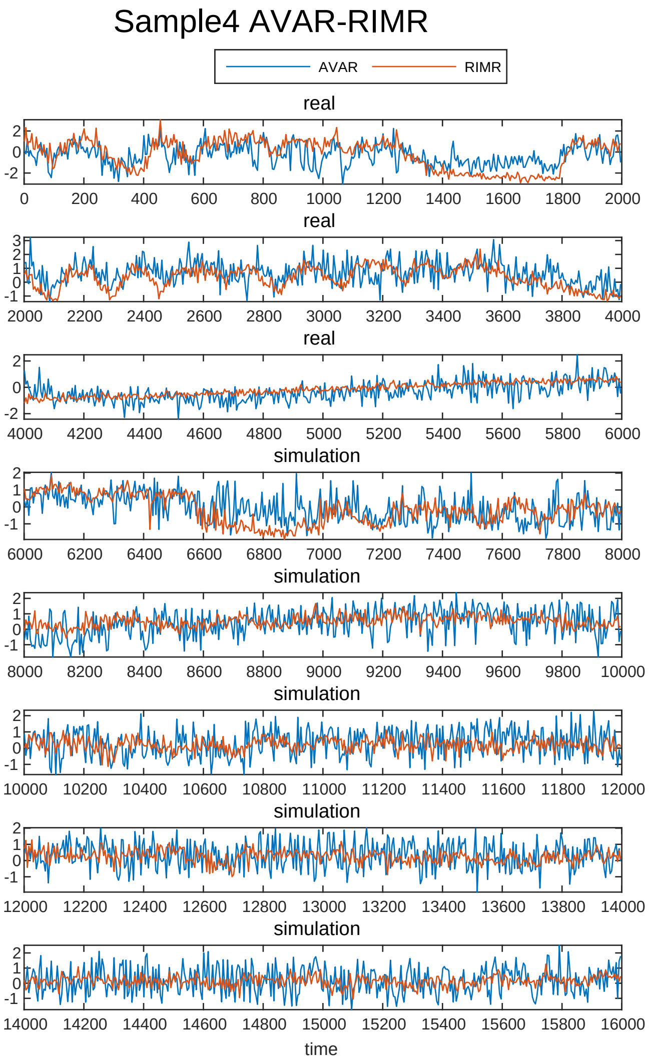
Sample4 AVAR-RIMR
AVAR
RIMR
real
real
real
simulation
simulation
simulation
simulation
simulation
time

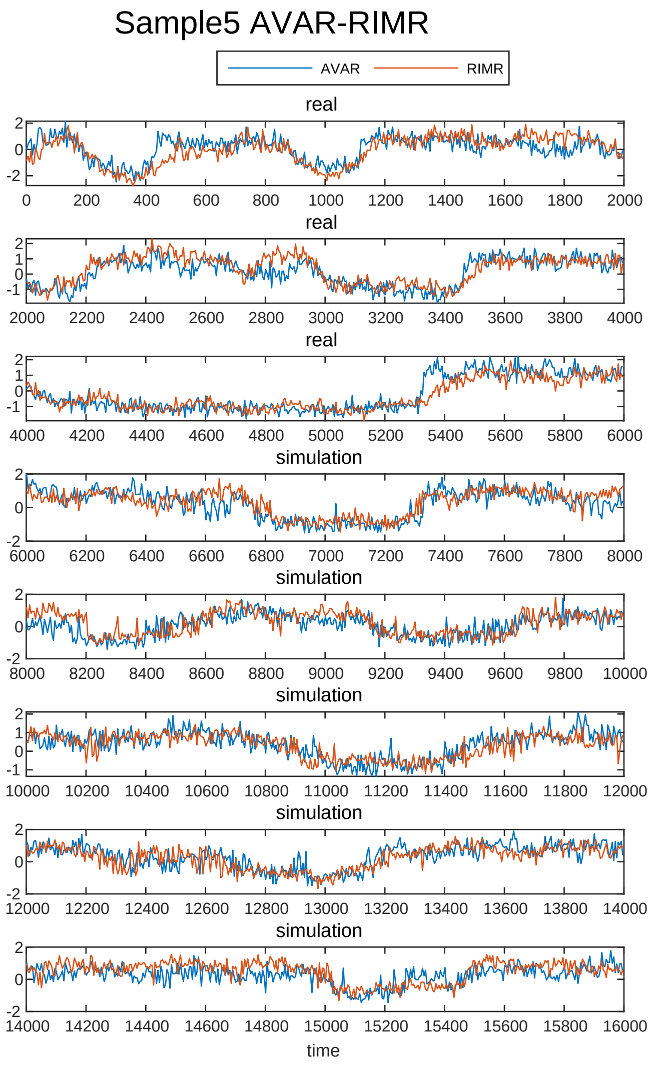

Sample5 AVAR-RIMR
AVAR
RIMR
real
real
real
simulation
simulation
simulation
simulation
simulation
time


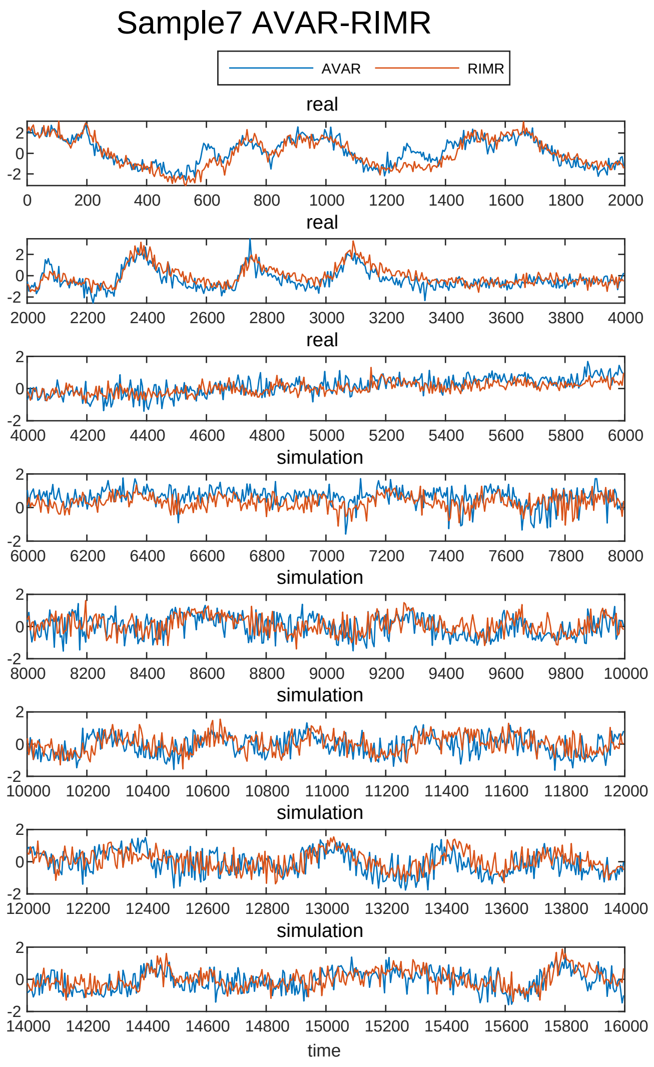

Sample7 AVAR-RIMR
AVAR
RIMR
real
real
real
simulation
simulation
simulation
simulation
simulation
time


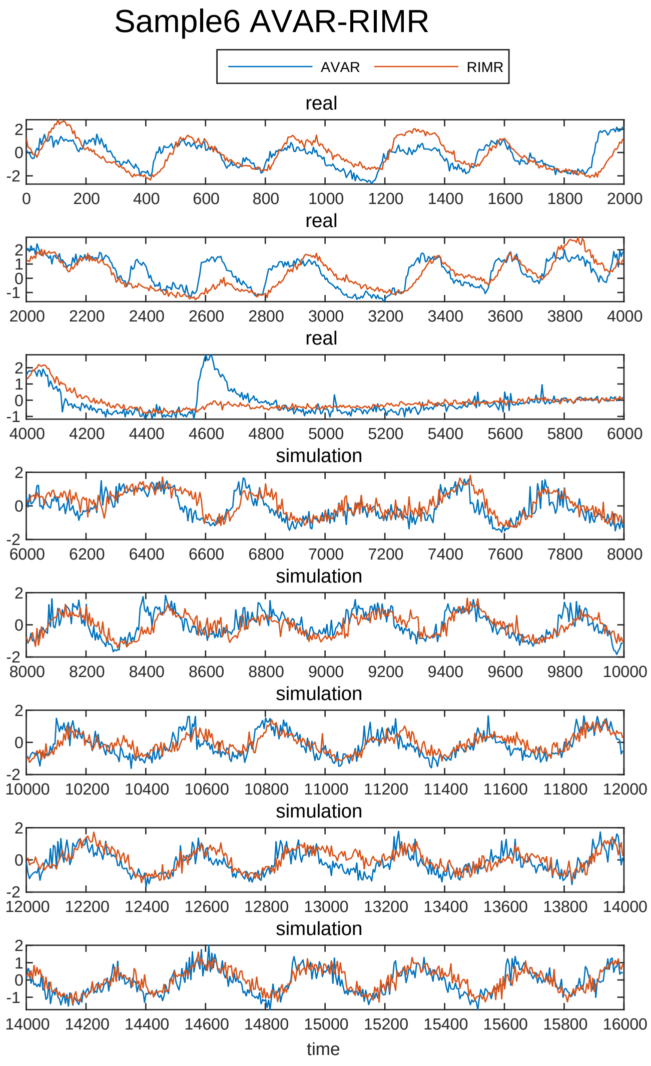

Sample6 AVAR-RIMR
AVAR
RIMR
real
real
real
simulation
simulation
simulation
simulation
simulation
time


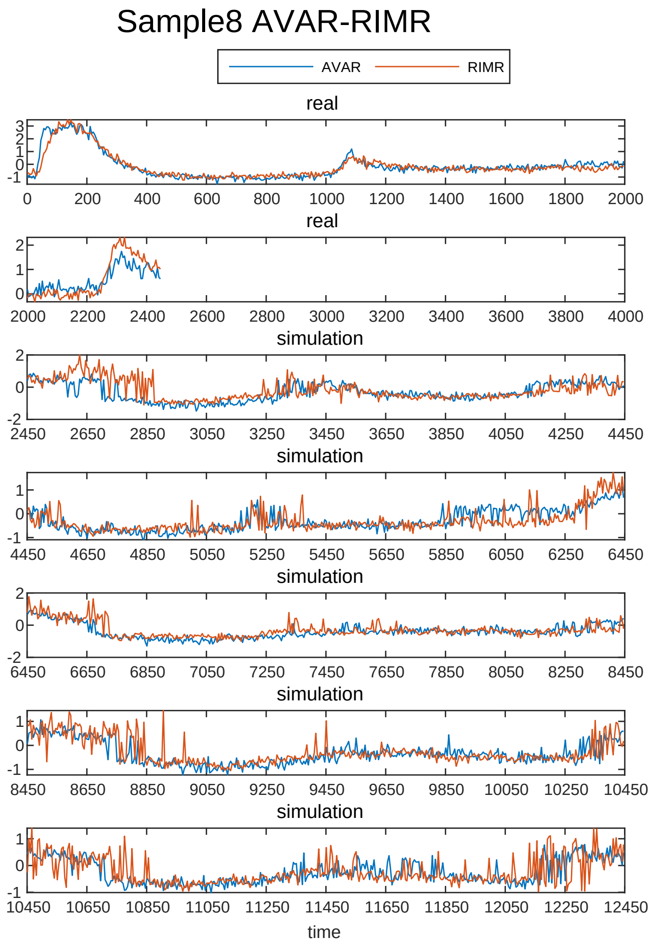

Sample8 AVAR-RIMR
AVAR
RIMR
real
real
simulation
simulation
simulation
simulation
simulation
time

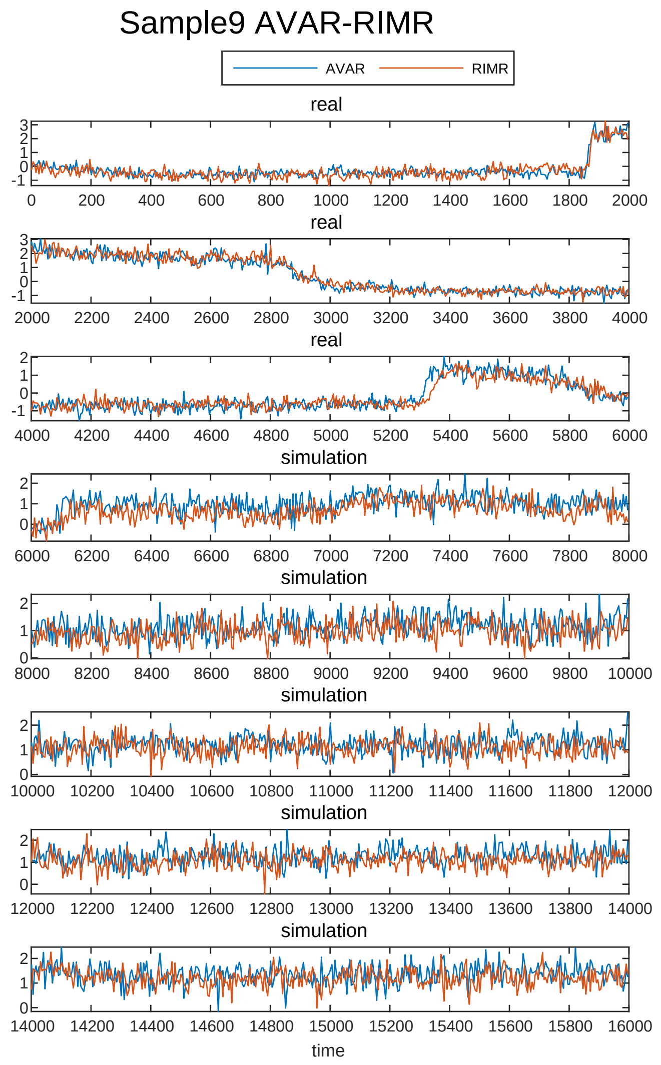
Sample9 AVAR-RIMR
AVAR
RIMR
real
real
real
simulation
simulation
simulation
simulation
simulation
time

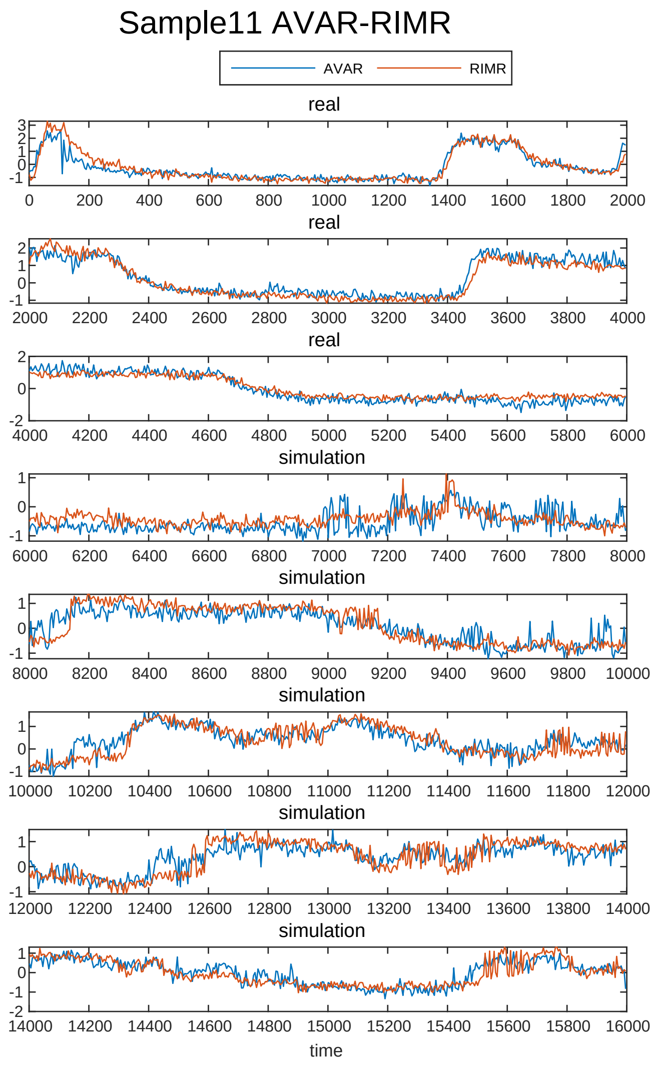
Sample11 AVAR-RIMR
AVAR
RIMR
real
real
real
simulation
simulation
simulation
simulation
simulation
time

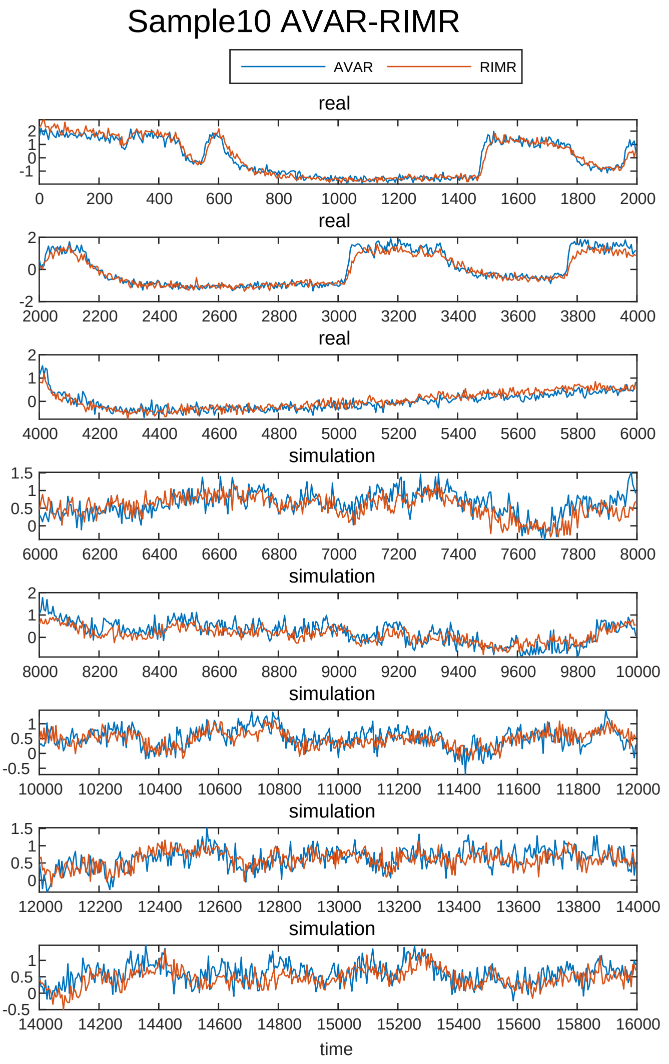
Sample10 AVAR-RIMR
AVAR
RIMR
real
real
real
simulation
simulation
simulation
simulation
simulation
time

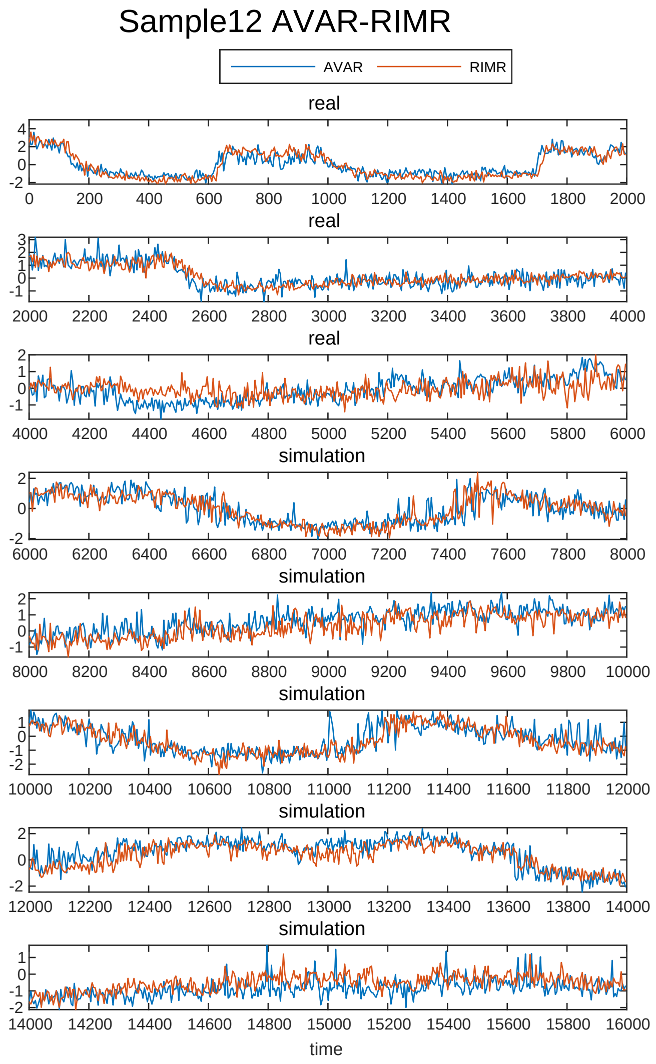
Sample12 AVAR-RIMR
AVAR
RIMR
real
real
real
simulation
simulation
simulation
simulation
simulation
time

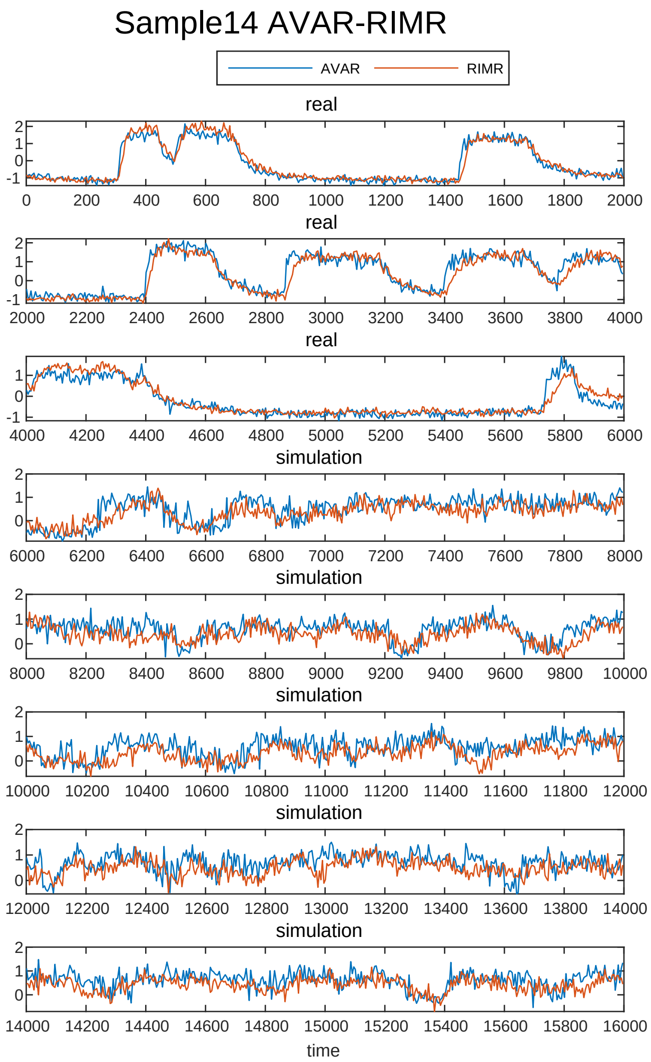
Sample14 AVAR-RIMR
AVAR
RIMR
real
real
real
simulation
simulation
simulation
simulation
simulation
time

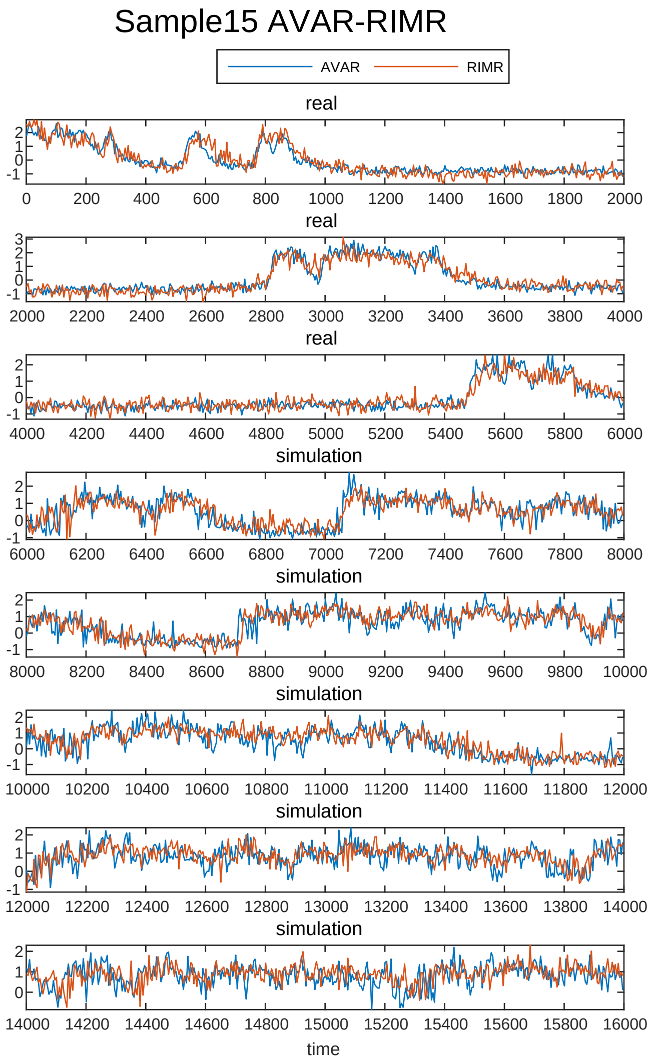
Sample15 AVAR-RIMR
AVAR
RIMR
real
real
real
simulation
simulation
simulation
simulation
simulation
time

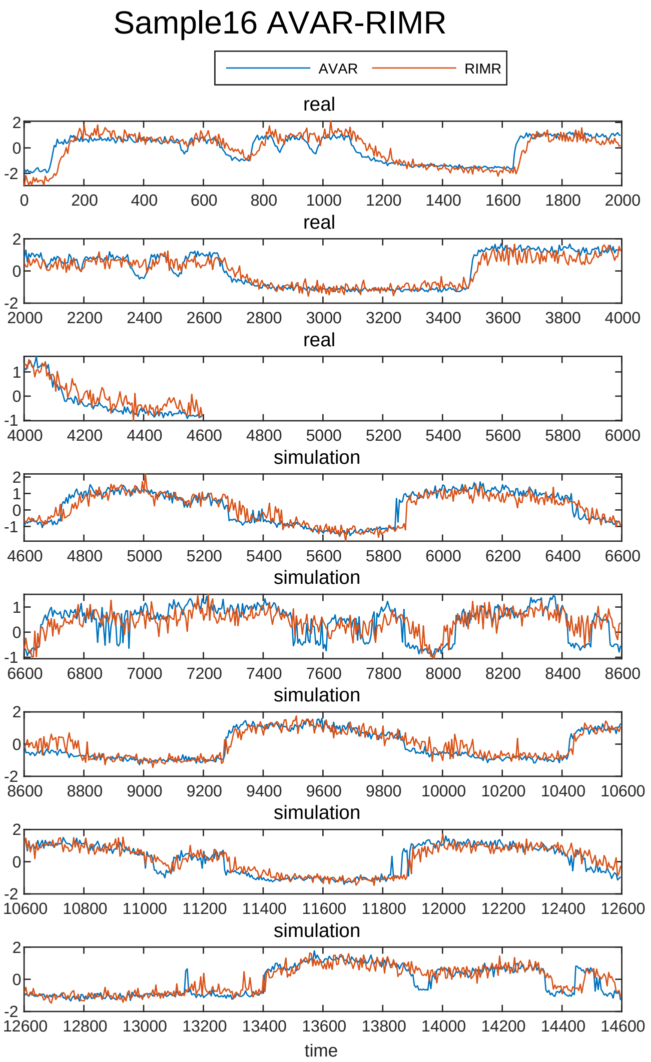
Sample16 AVAR-RIMR
AVAR
RIMR
real
real
real
simulation
simulation
simulation
simulation
simulation
time

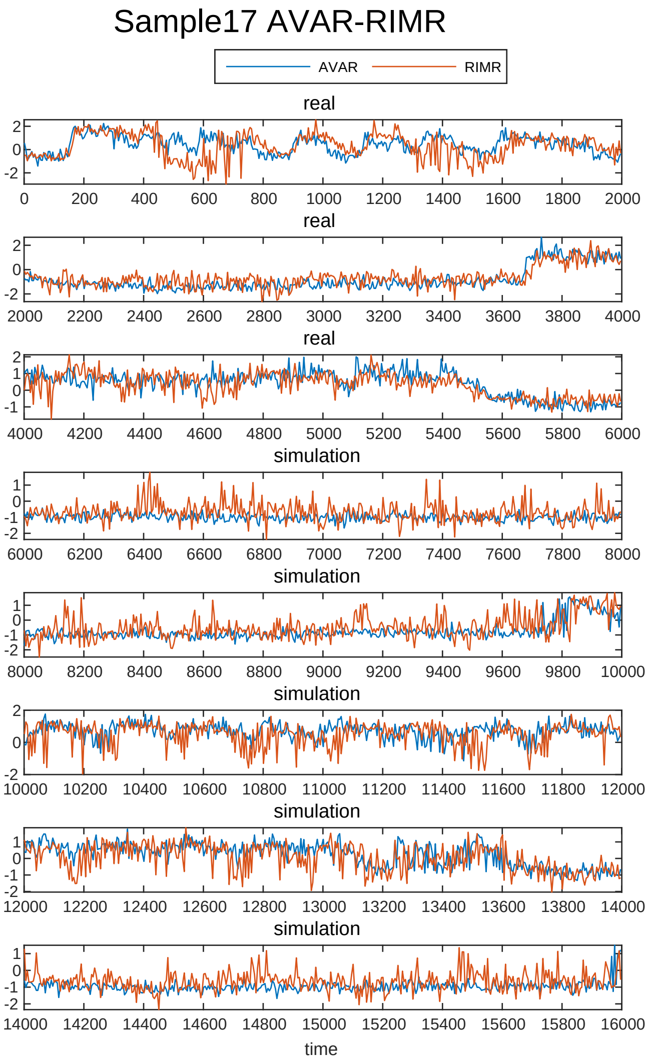
Sample17 AVAR-RIMR
AVAR
RIMR
real
real
real
simulation
simulation
simulation
simulation
simulation
time

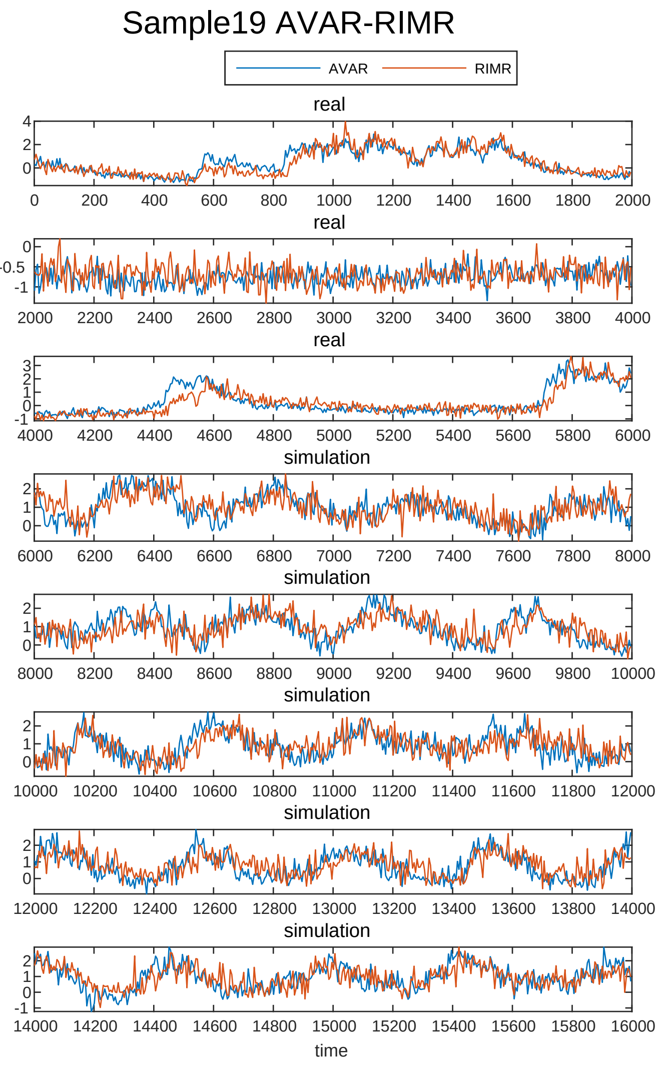
Sample19 AVAR-RIMR
AVAR
RIMR
real
real
real
simulation
simulation
simulation
simulation
simulation
time

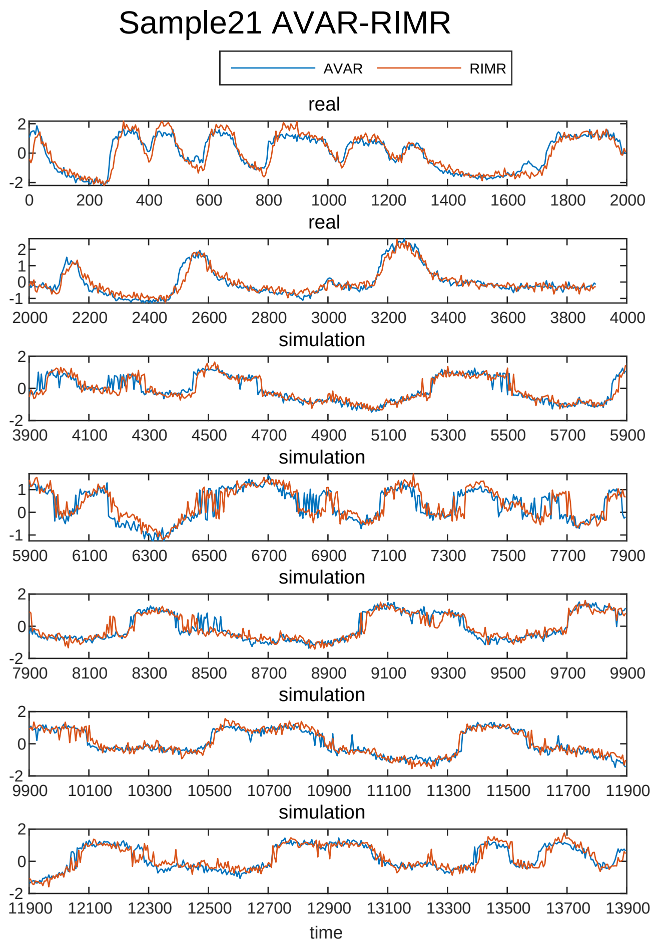
Sample21 AVAR-RIMR
AVAR
RIMR
real
real
simulation
simulation
simulation
simulation
simulation
time

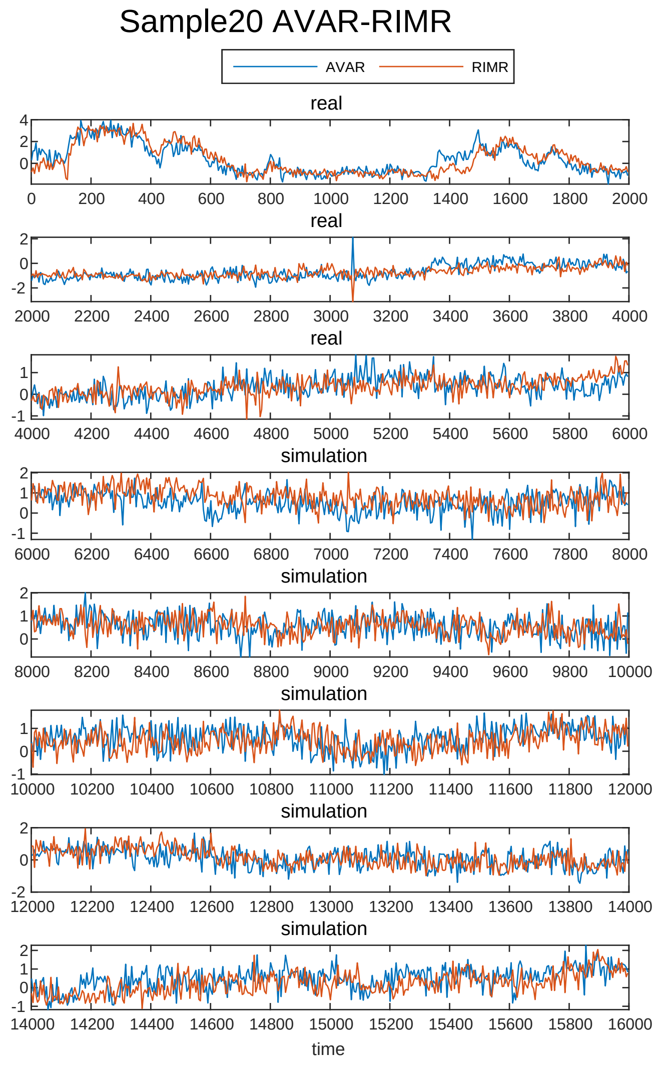
Sample20 AVAR-RIMR
AVAR
RIMR
real
real
real
simulation
simulation
simulation
simulation
simulation
time

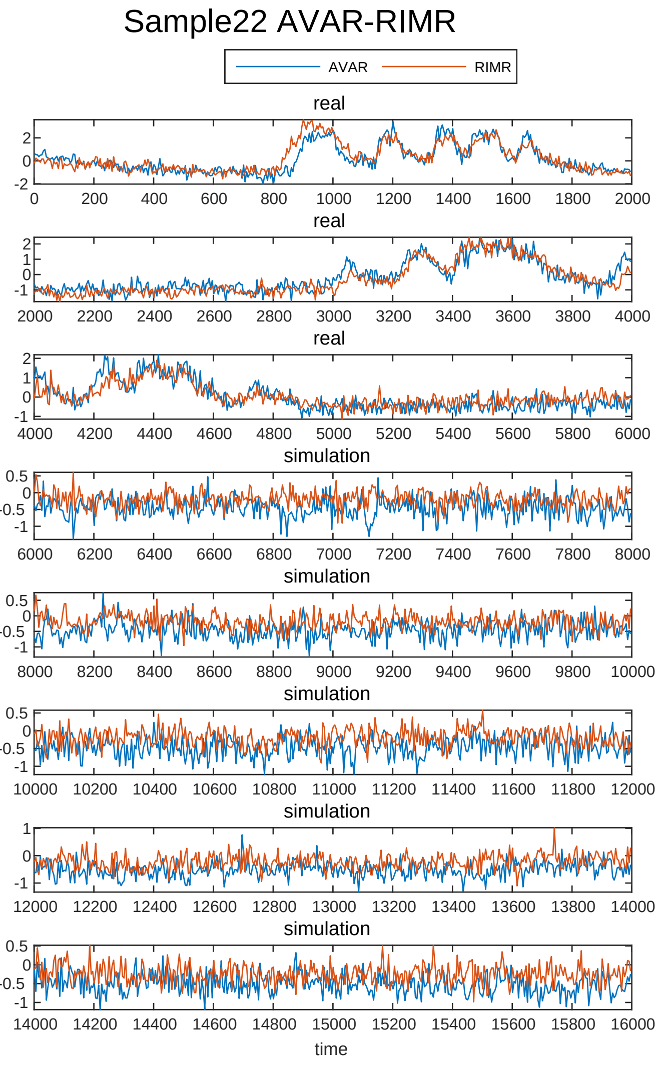
Sample22 AVAR-RIMR
AVAR
RIMR
real
real
real
simulation
simulation
simulation
simulation
simulation
time

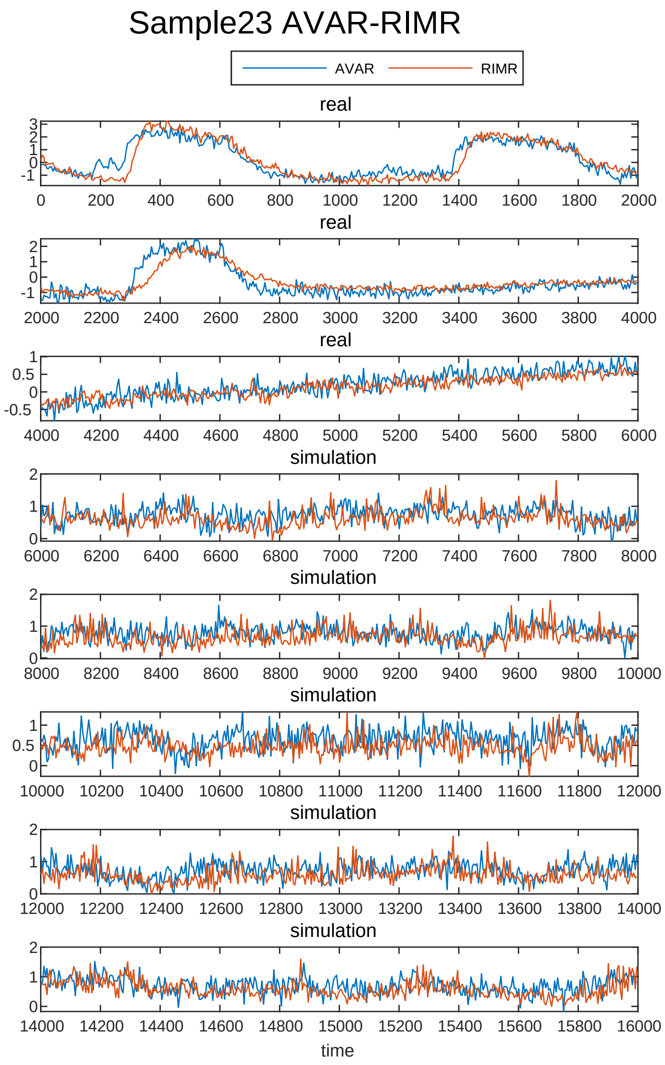
Sample23 AVAR-RIMR
AVAR
RIMR
real
real
real
simulation
simulation
simulation
simulation
simulation
time

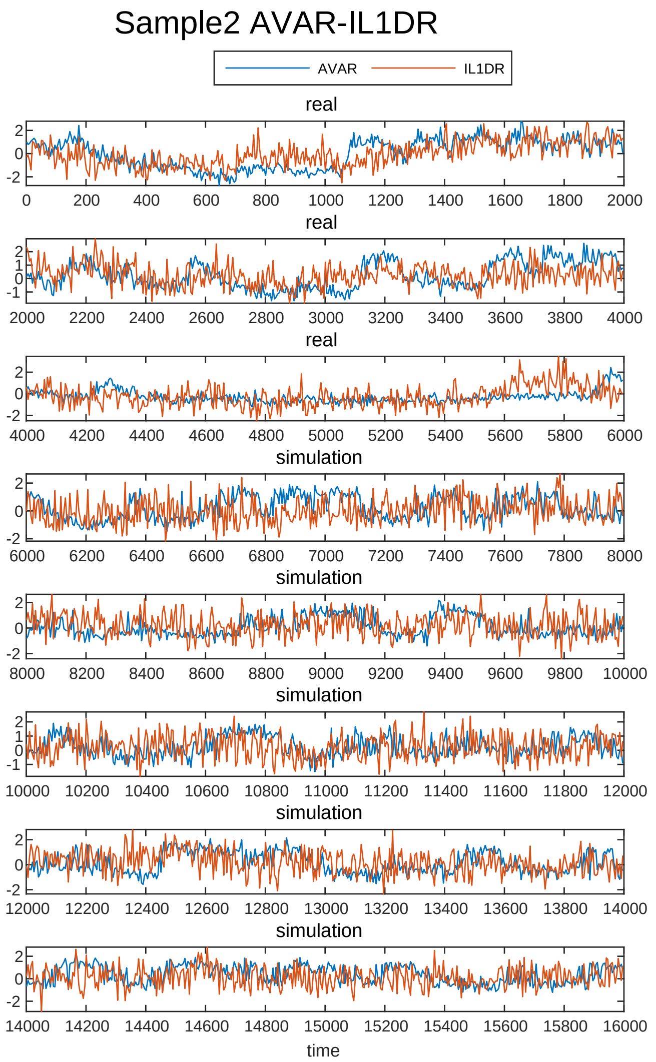
Sample2 AVAR-IL1DR
AVAR
IL1DR
real
real
real
simulation
simulation
simulation
simulation
simulation
time

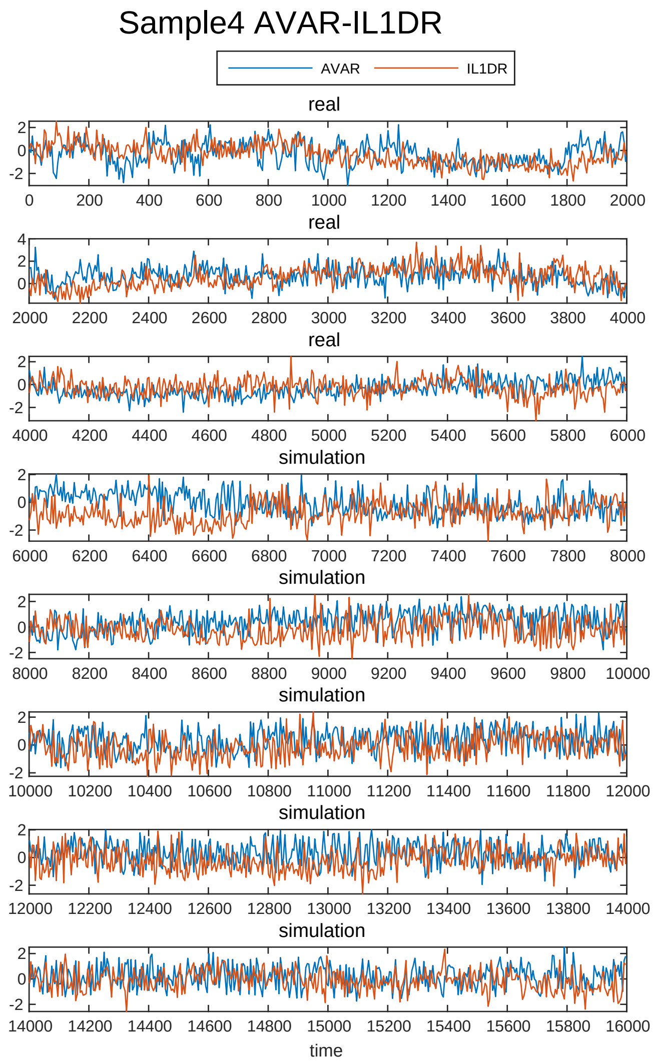
Sample4 AVAR-IL1DR
AVAR
IL1DR
real
real
real
simulation
simulation
simulation
simulation
simulation
time

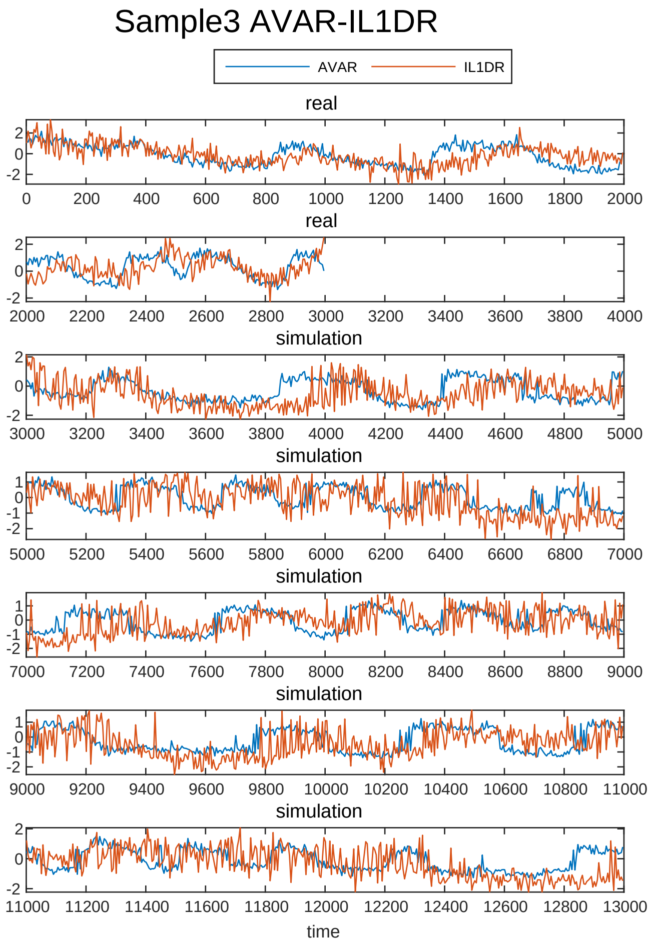
Sample3 AVAR-IL1DR
AVAR
IL1DR
real
real
simulation
simulation
simulation
simulation
simulation
time

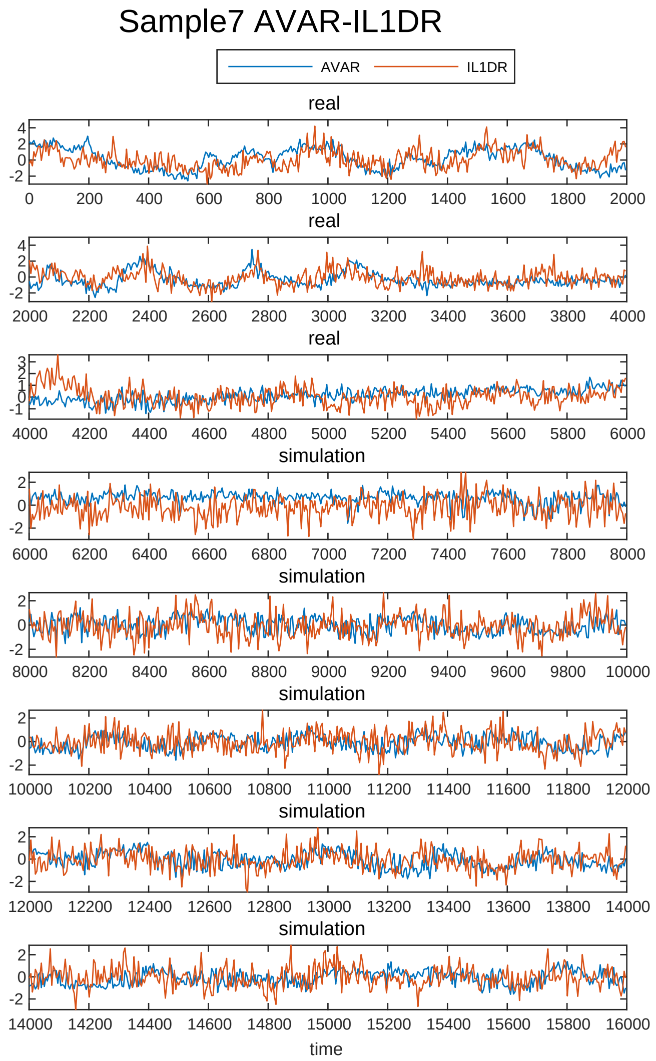
Sample7 AVAR-IL1DR
AVAR
IL1DR
real
real
real
simulation
simulation
simulation
simulation
simulation
time

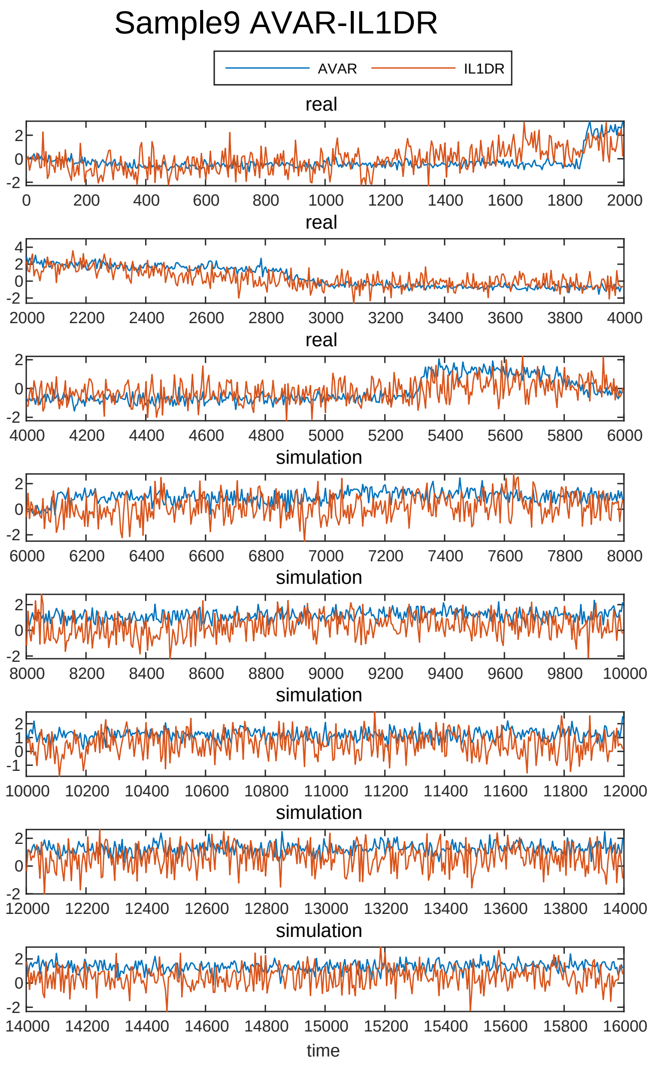
Sample9 AVAR-IL1DR
AVAR
IL1DR
real
real
real
simulation
simulation
simulation
simulation
simulation
time

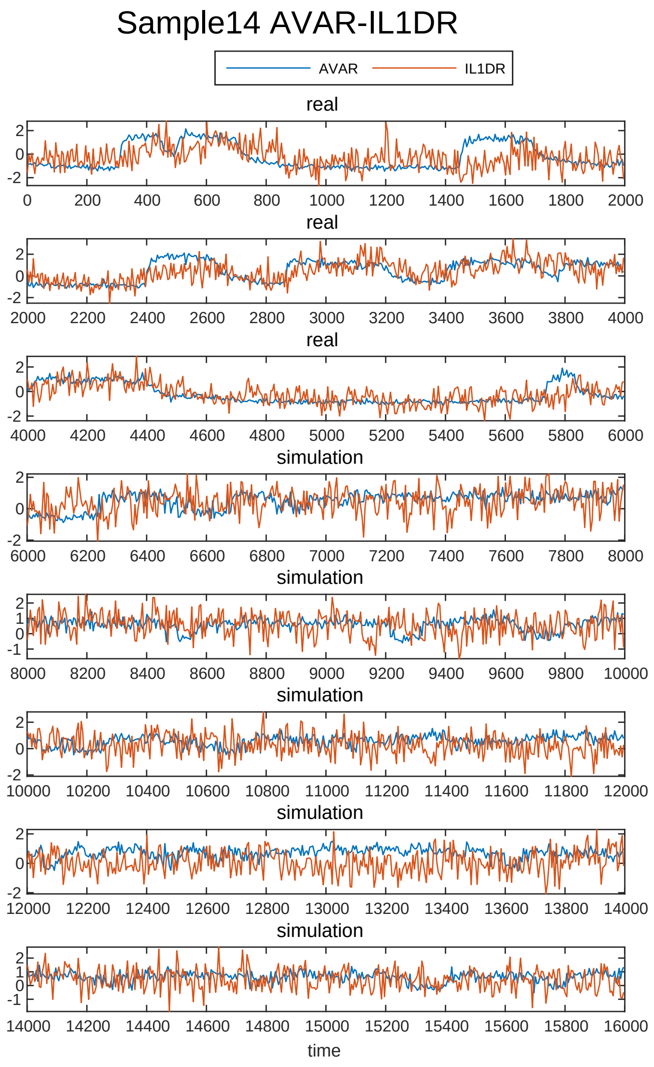
Sample14 AVAR-IL1DR
AVAR
IL1DR
real
real
real
simulation
simulation
simulation
simulation
simulation
time

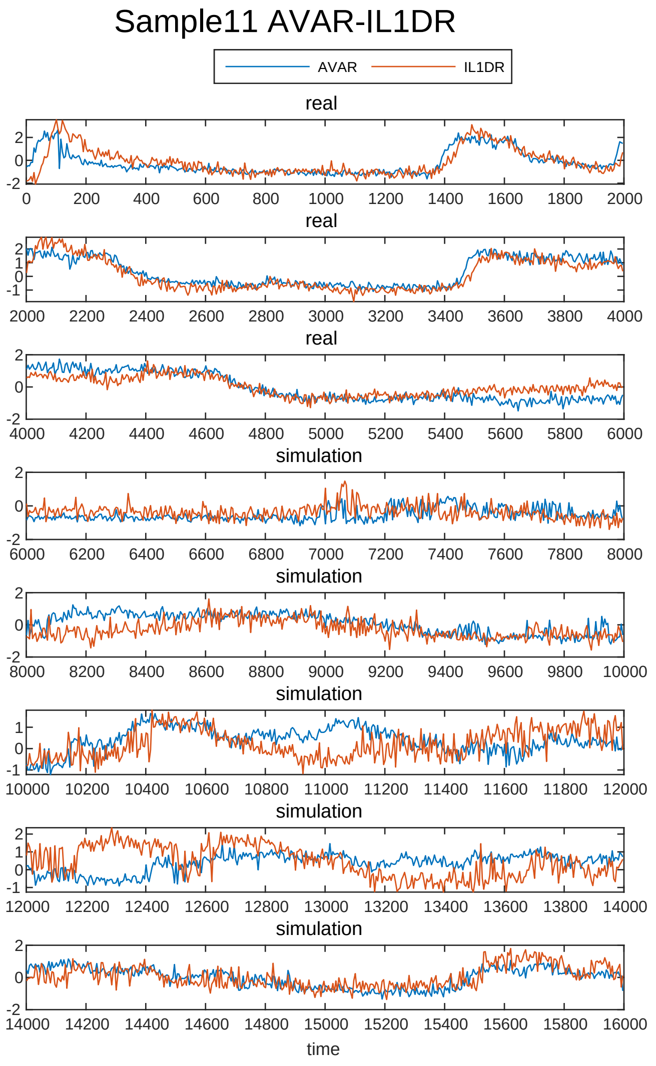
Sample11 AVAR-IL1DR
AVAR
IL1DR
real
real
real
simulation
simulation
simulation
simulation
simulation
time

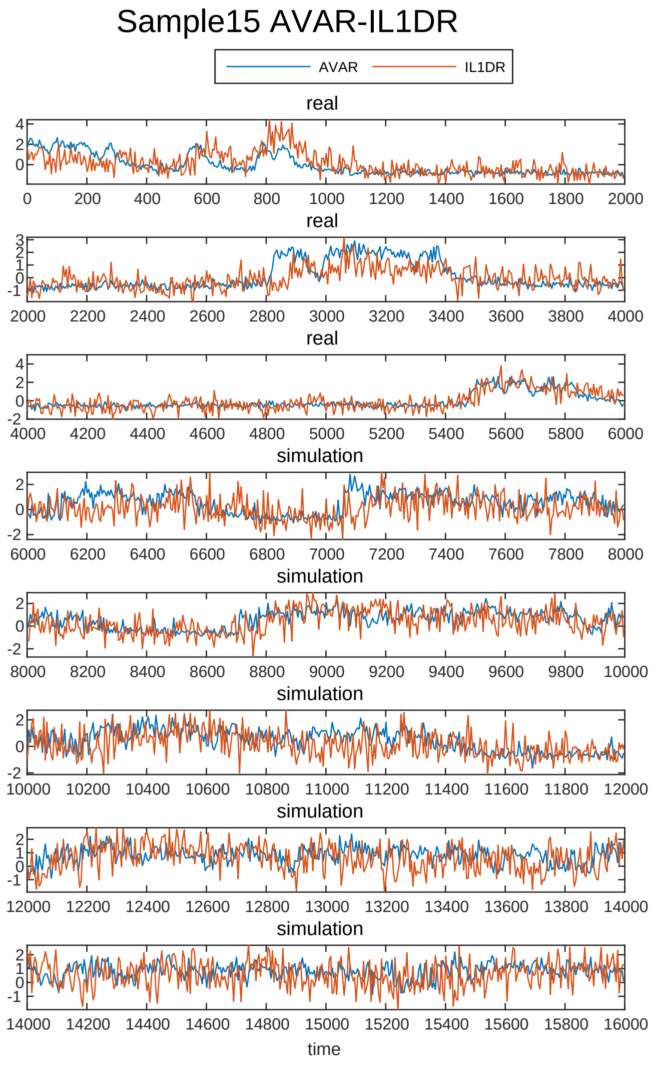
Sample15 AVAR-IL1DR
AVAR
IL1DR
real
real
real
simulation
simulation
simulation
simulation
simulation
time

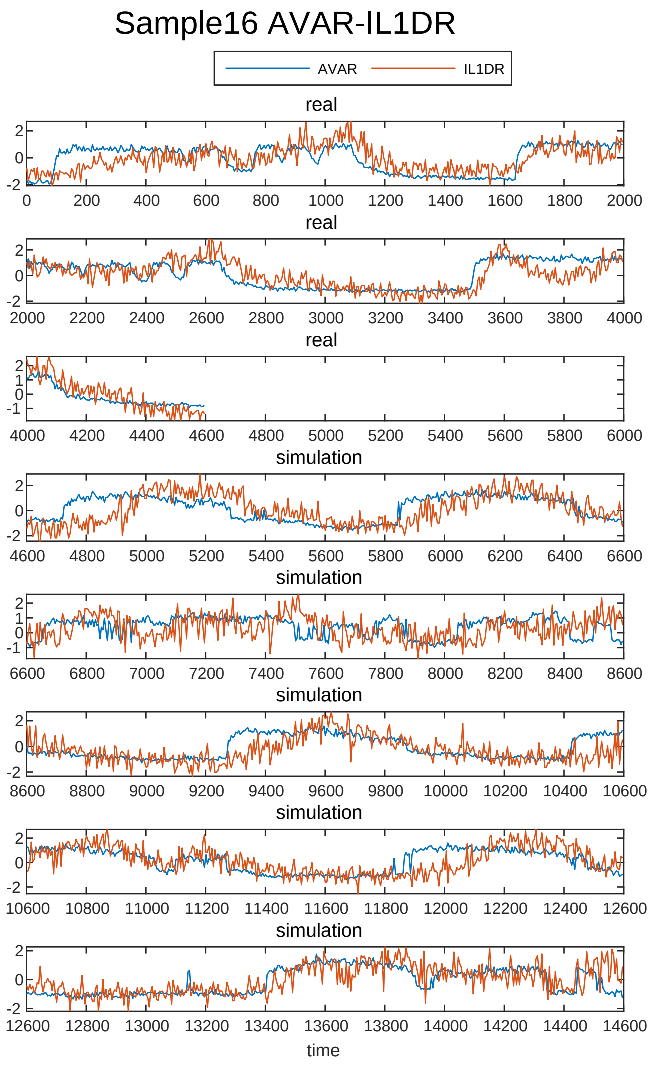
Sample16 AVAR-IL1DR
AVAR
IL1DR
real
real
real
simulation
simulation
simulation
simulation
simulation
time

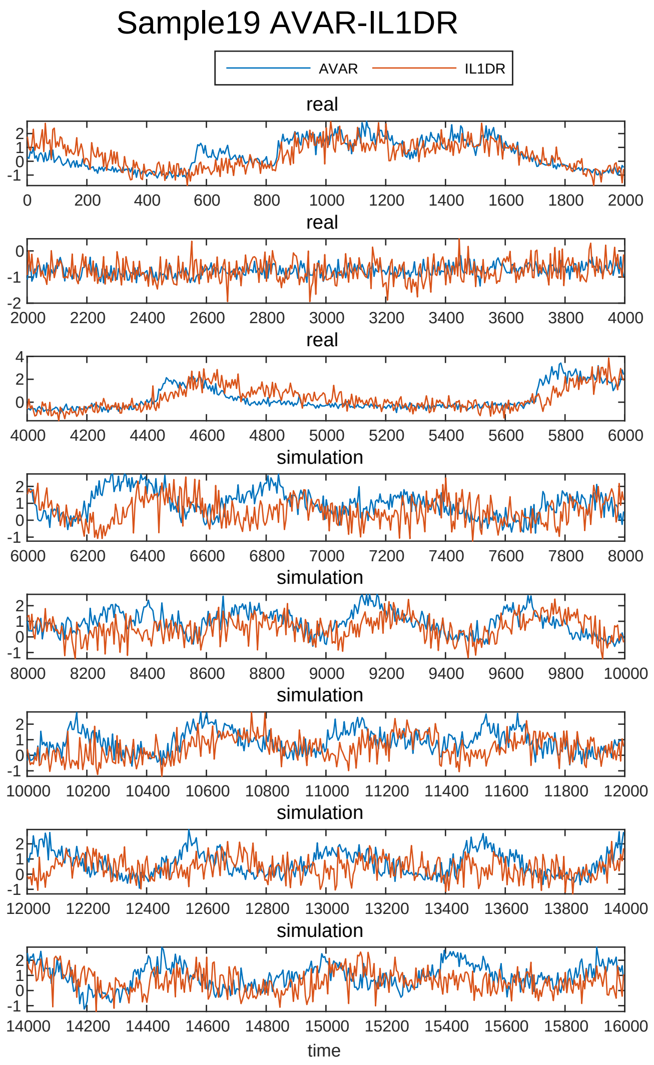
Sample19 AVAR-IL1DR
AVAR
IL1DR
real
real
real
simulation
simulation
simulation
simulation
simulation
time

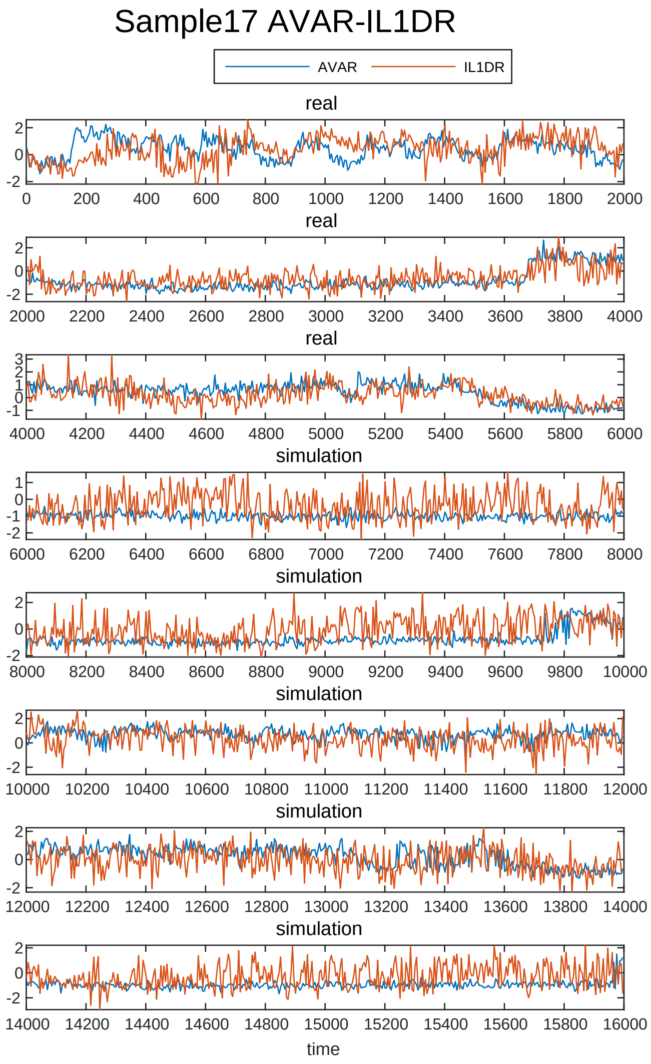
Sample17 AVAR-IL1DR
AVAR
IL1DR
real
real
real
simulation
simulation
simulation
simulation
simulation
time

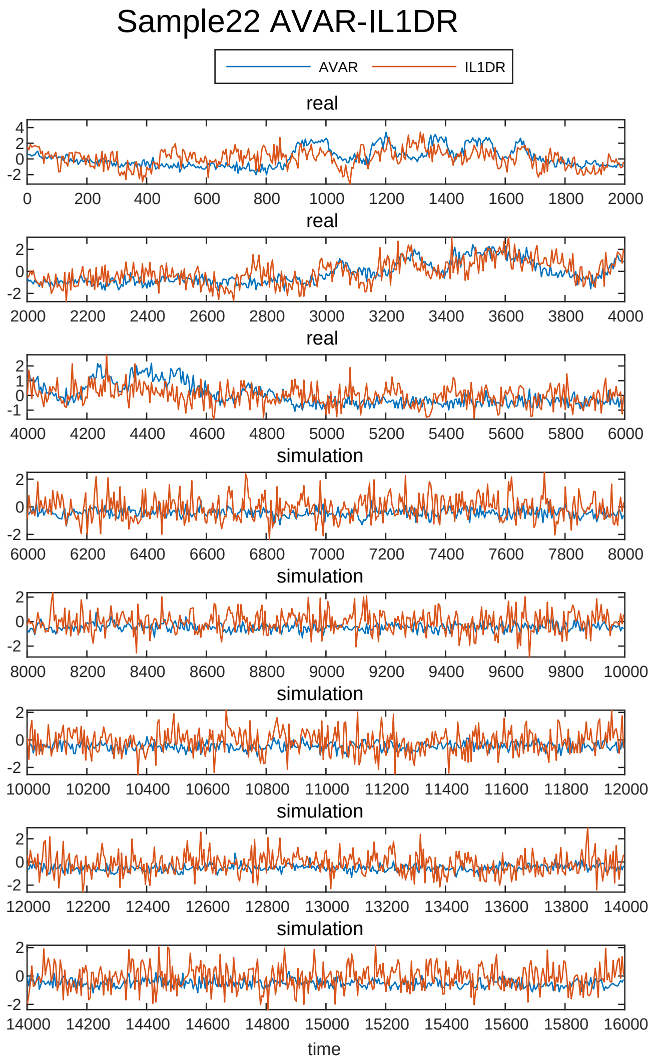
Sample22 AVAR-IL1DR
AVAR
IL1DR
real
real
real
simulation
simulation
simulation
simulation
simulation
time

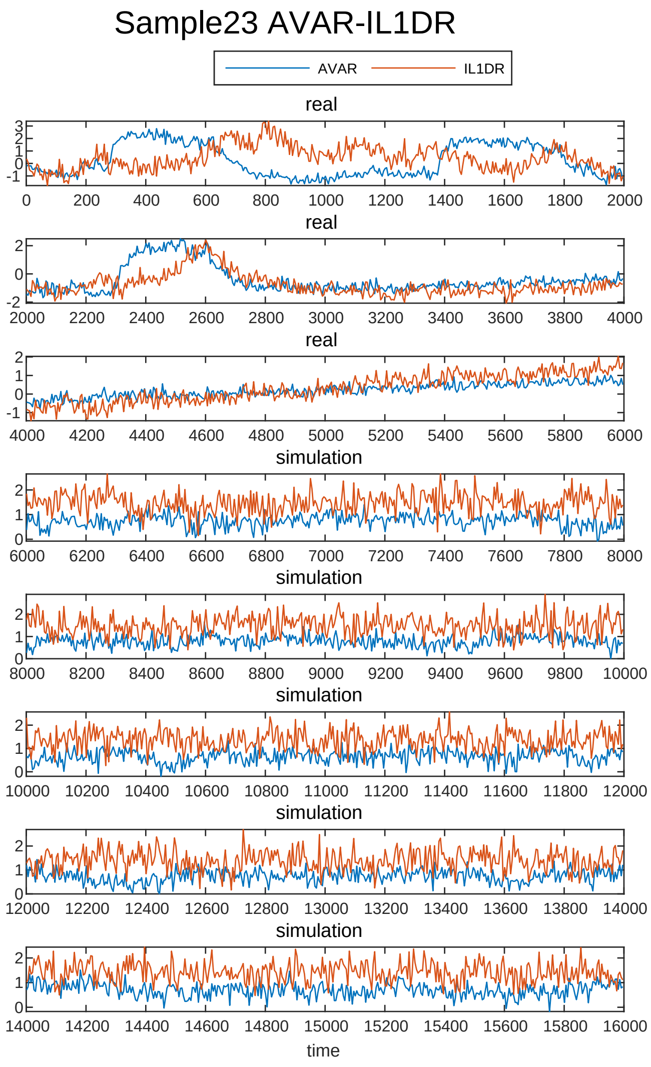

Sample23 AVAR-IL1DR
AVAR
IL1DR
real
real
real
simulation
simulation
simulation
simulation
simulation
time

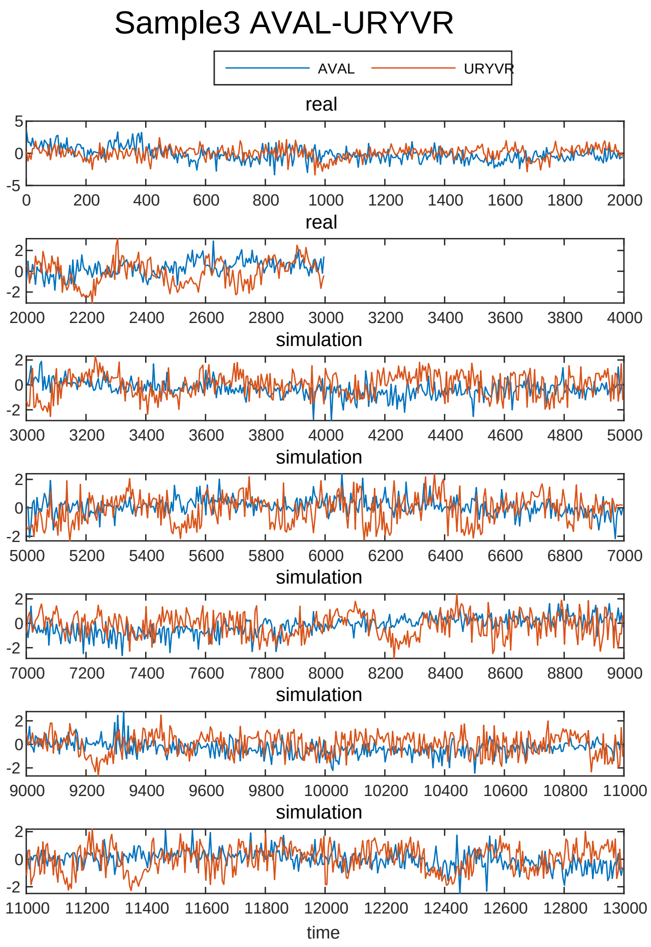
Sample3 AVAL-URYVR
AVAL
URYVR
real
real
simulation
simulation
simulation
simulation
simulation
time

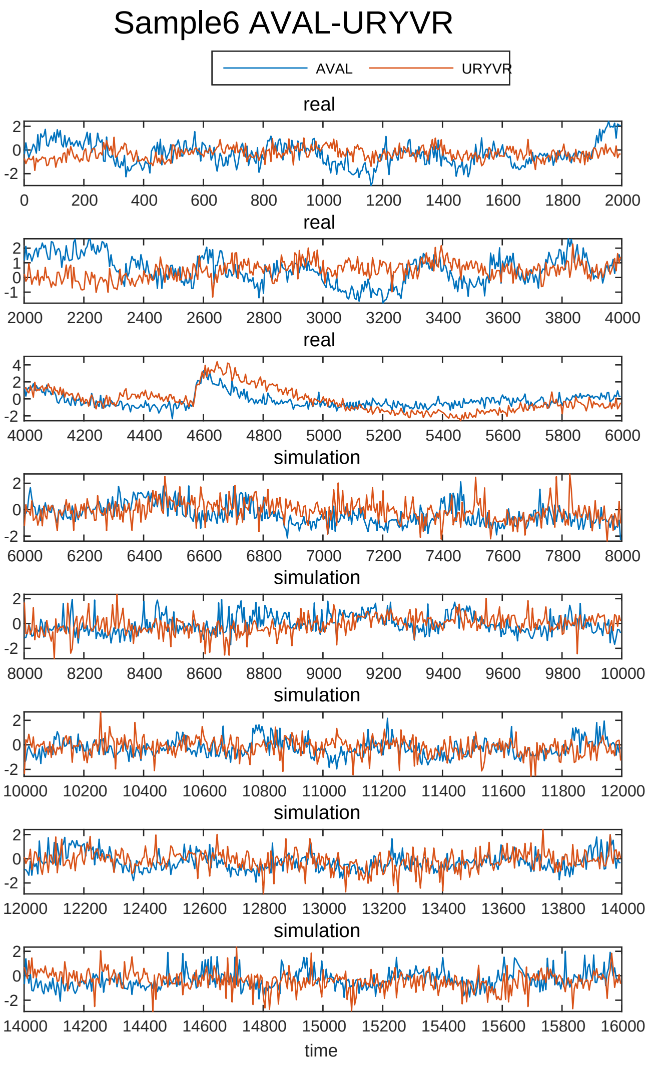
Sample6 AVAL-URYVR
AVAL
URYVR
real
real
real
simulation
simulation
simulation
simulation
simulation
time

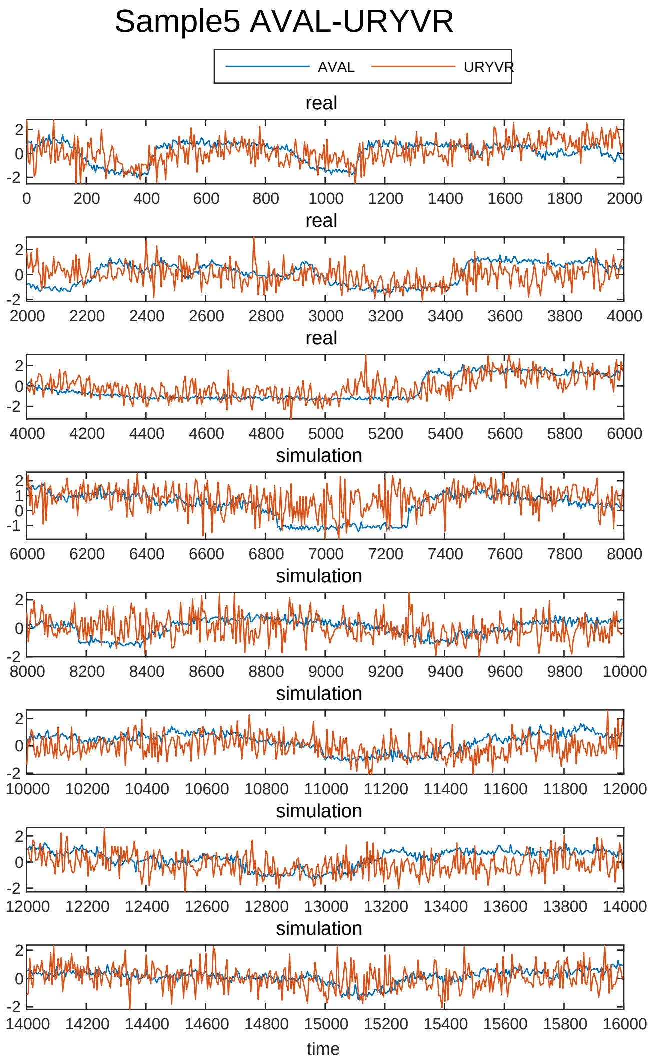
Sample5 AVAL-URYVR
AVAL
URYVR
real
real
real
simulation
simulation
simulation
simulation
simulation
time

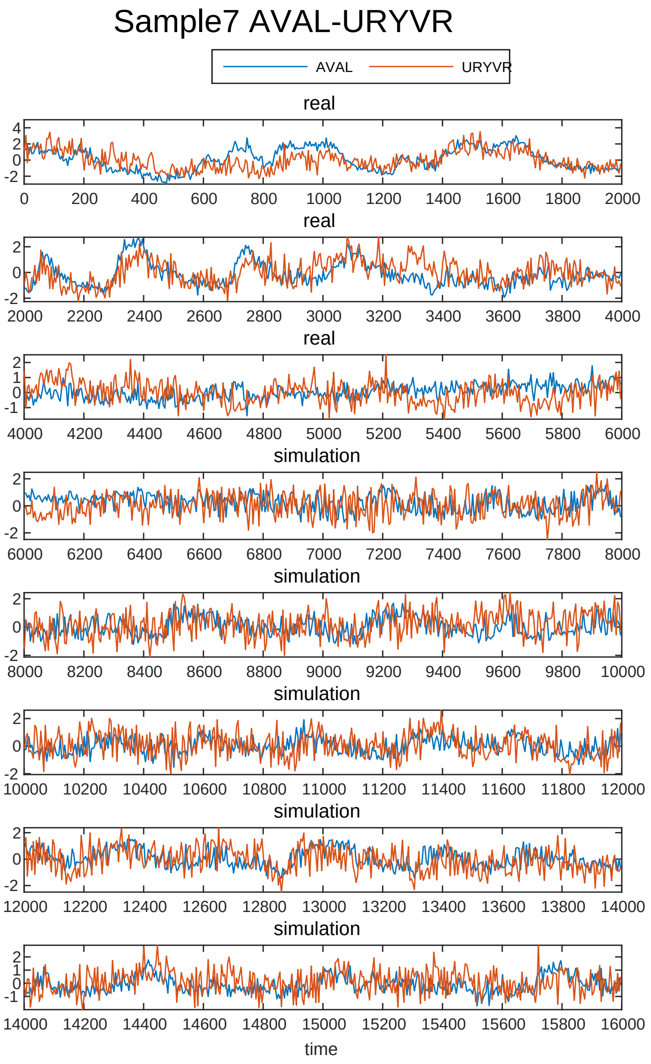
Sample7 AVAL-URYVR
AVAL
URYVR
real
real
real
simulation
simulation
simulation
simulation
simulation
time

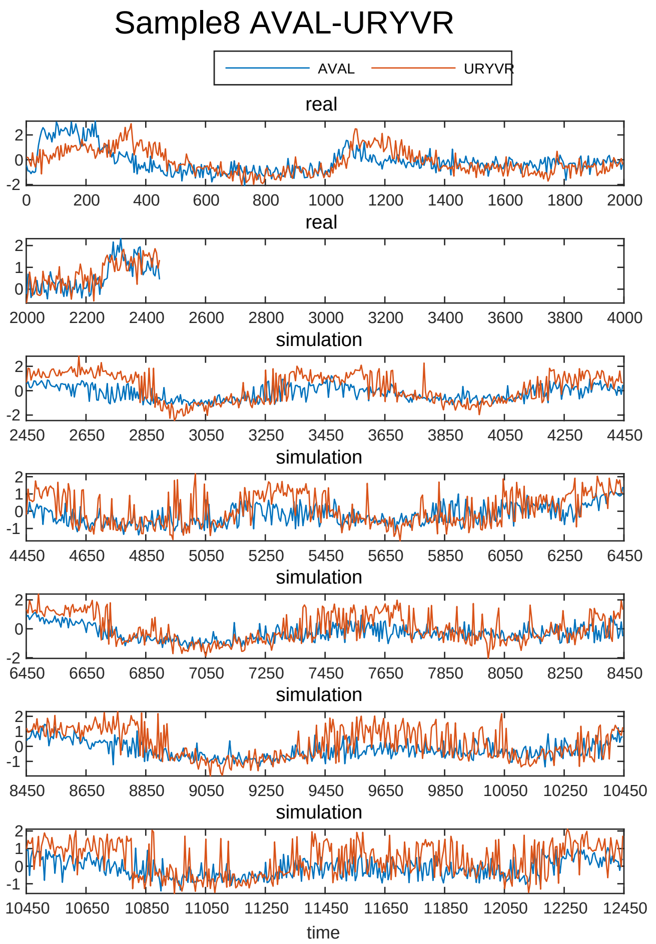
Sample8 AVAL-URYVR
AVAL
URYVR
real
real
simulation
simulation
simulation
simulation
simulation
time

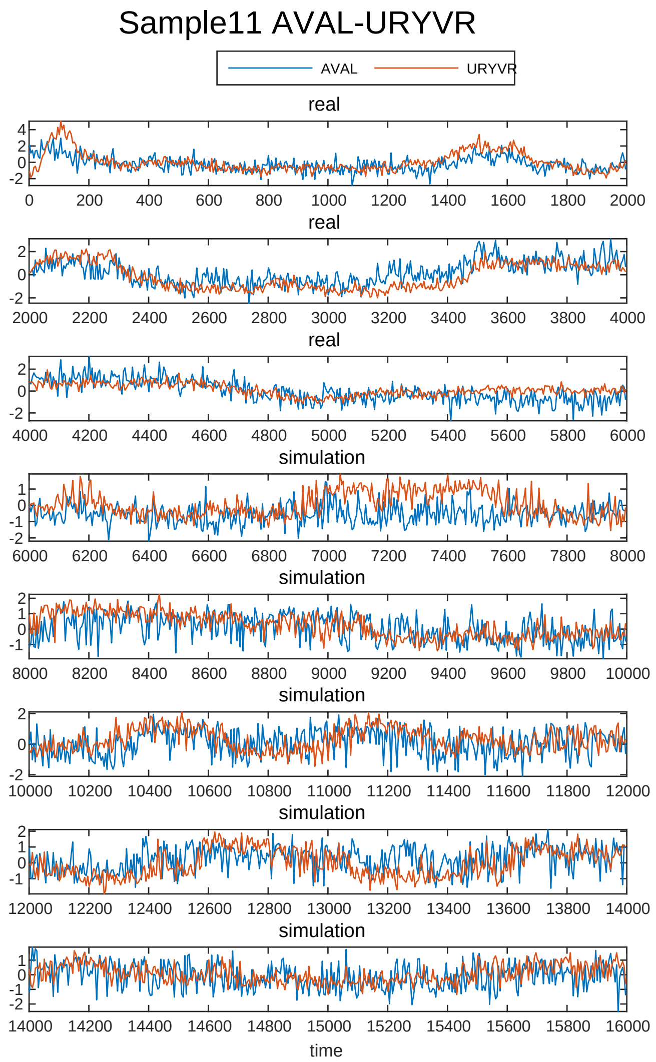
Sample11 AVAL-URYVR
AVAL
URYVR
real
real
real
simulation
simulation
simulation
simulation
simulation
time

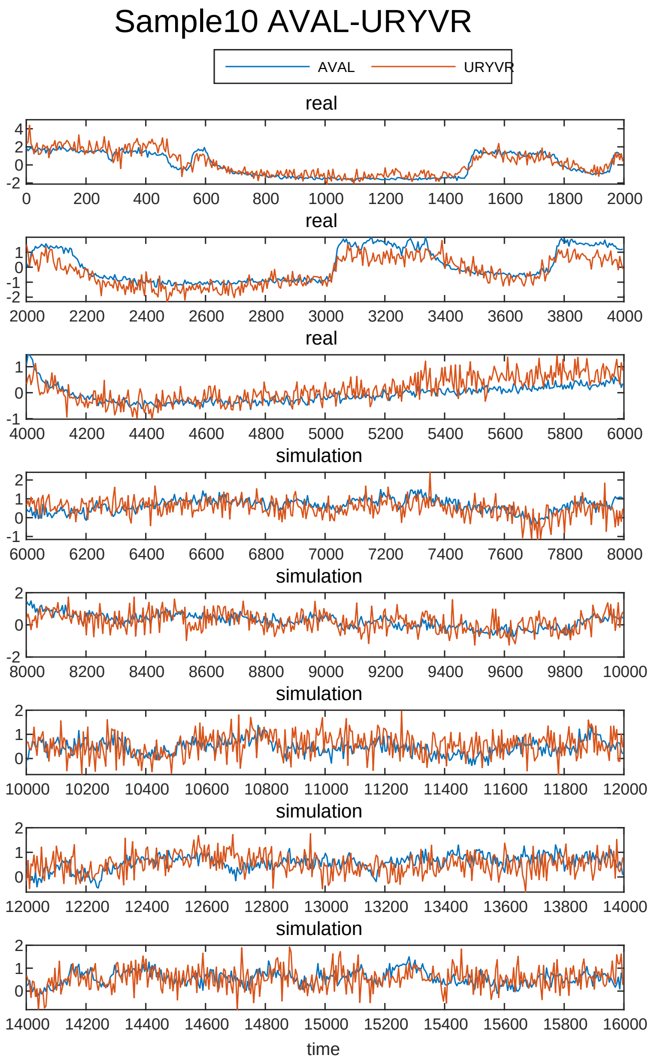
Sample10 AVAL-URYVR
AVAL
URYVR
real
real
real
simulation
simulation
simulation
simulation
simulation
time

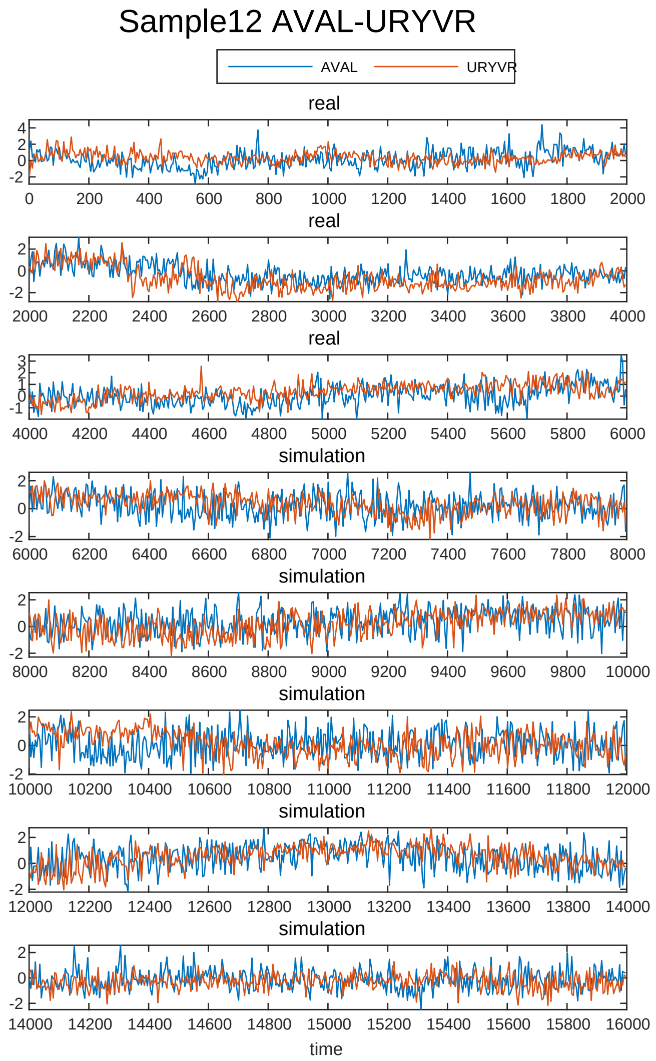
Sample12 AVAL-URYVR
AVAL
URYVR
real
real
real
simulation
simulation
simulation
simulation
simulation
time

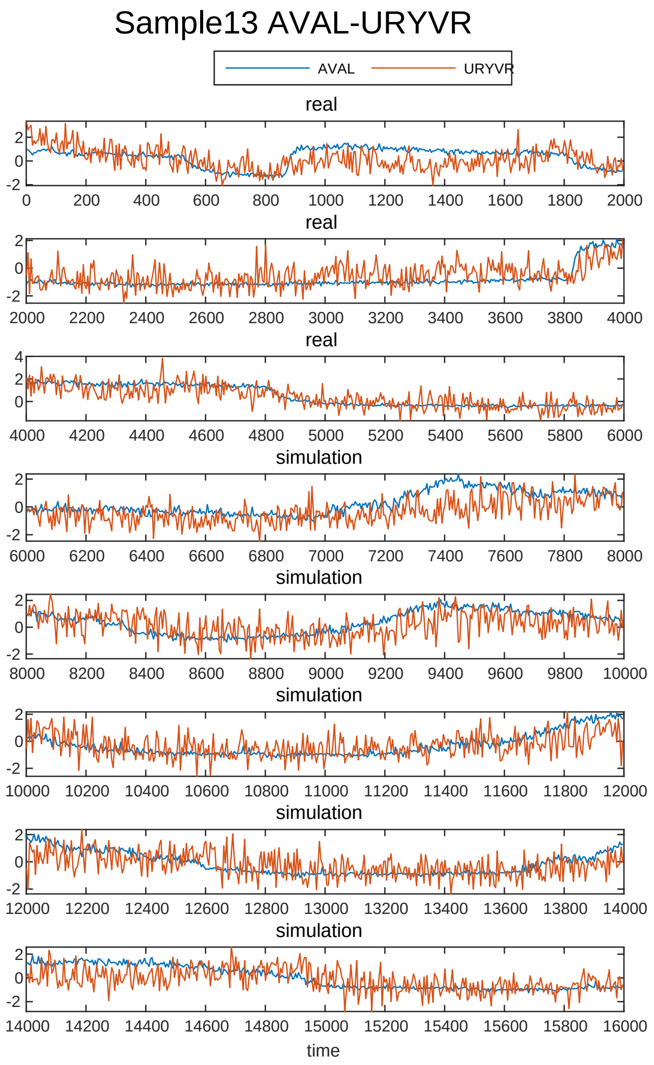
Sample13 AVAL-URYVR
AVAL
URYVR
real
simulation
time

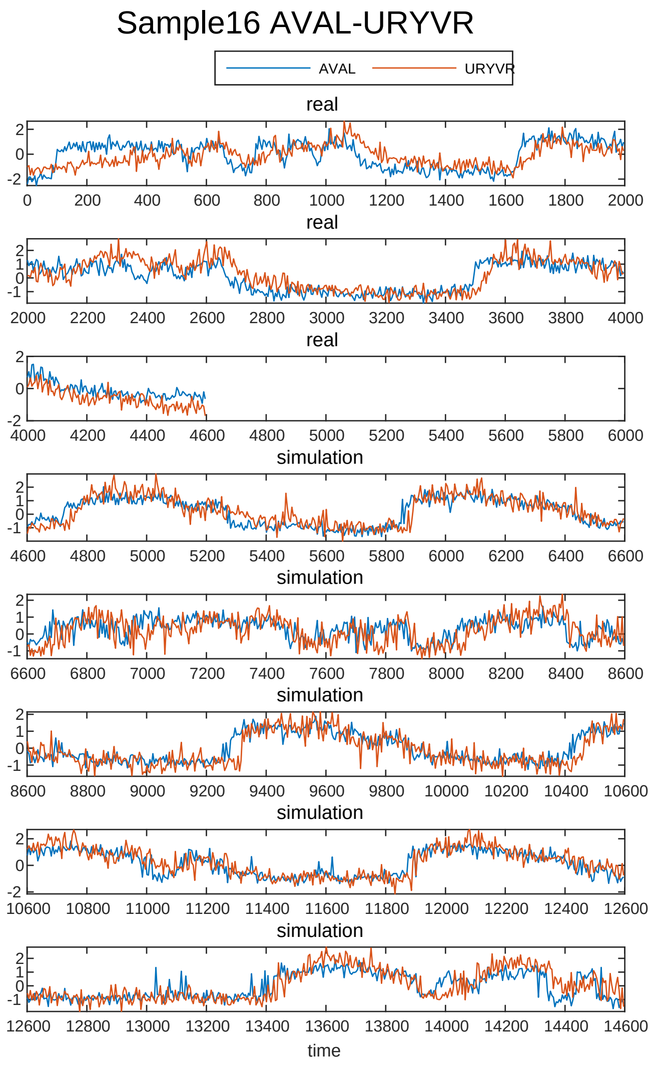
Sample16 AVAL-URYVR
AVAL
URYVR
real
simulation
time

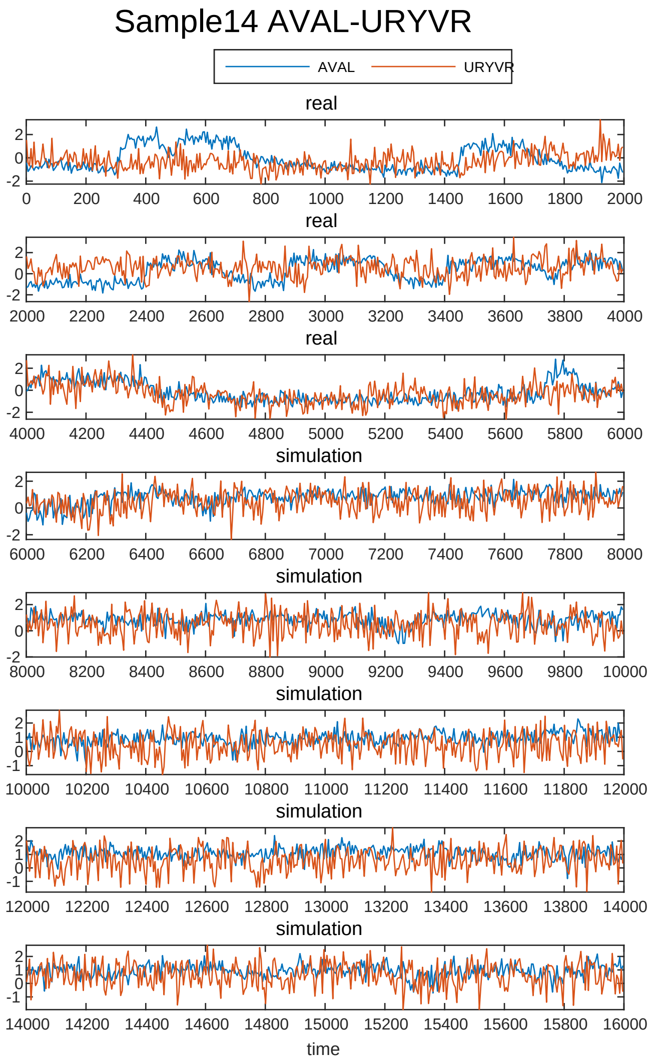
Sample14 AVAL-URYVR
AVAL
URYVR
real
simulation
time

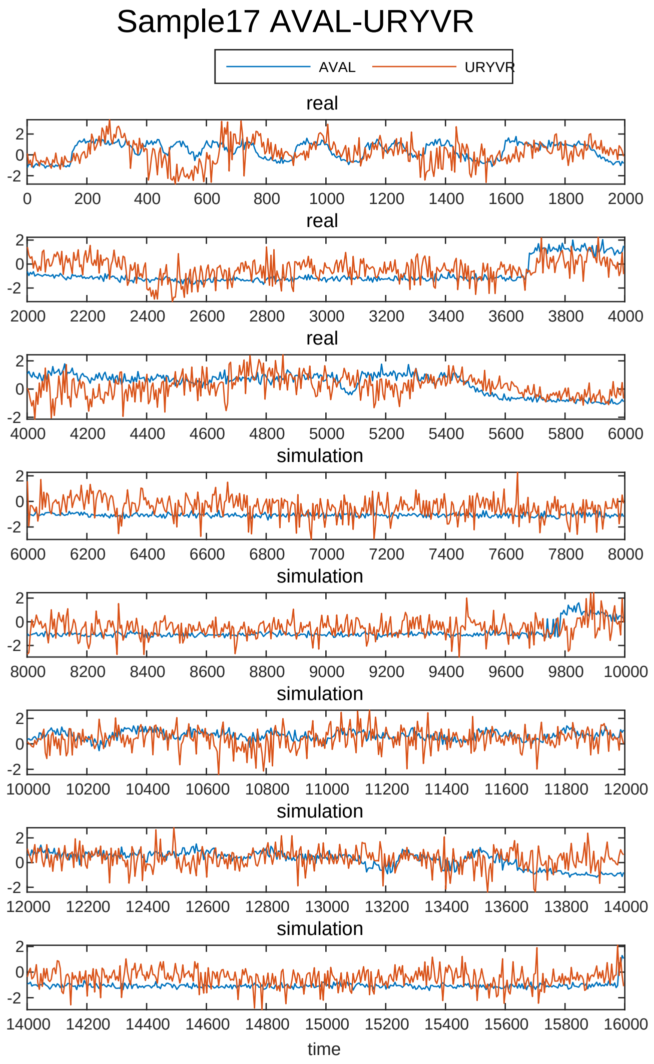
Sample17 AVAL-URYVR
AVAL
URYVR
real
simulation
time

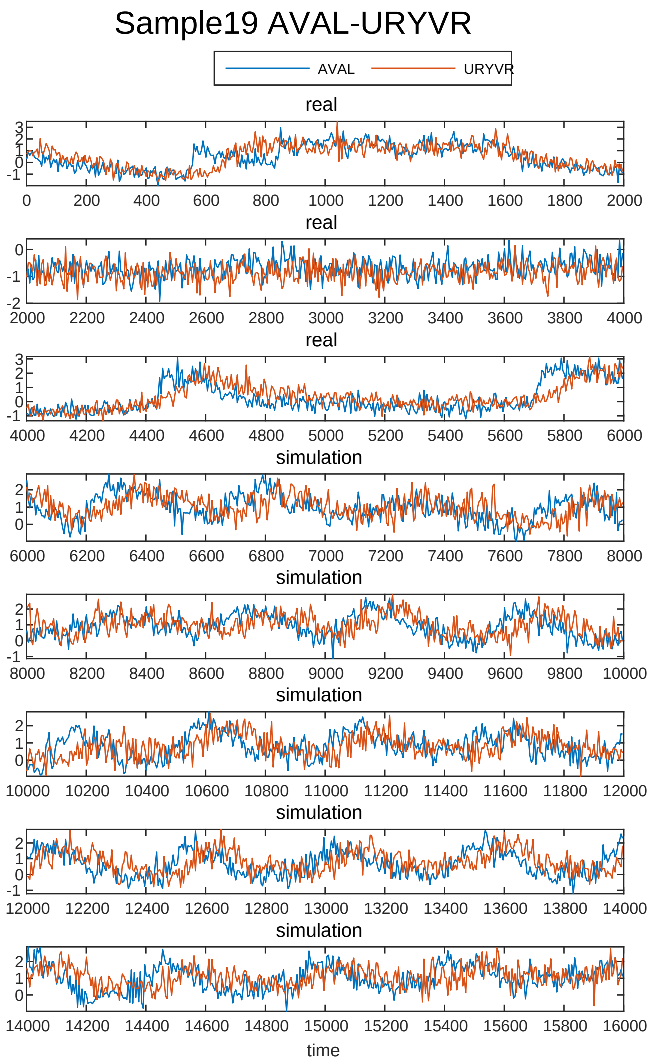

Sample19 AVAL-URYVR
AVAL
URYVR
real
real
real
simulation
simulation
simulation
simulation
simulation
time


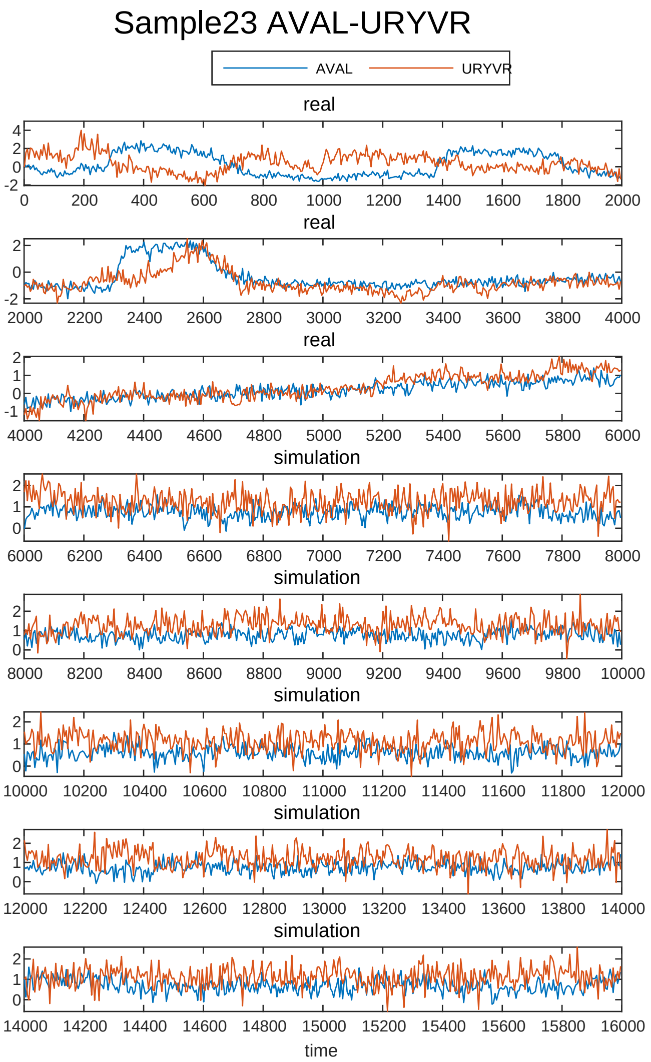

Sample23 AVAL-URYVR
AVAL
URYVR
real
real
real
simulation
simulation
simulation
simulation
simulation
time


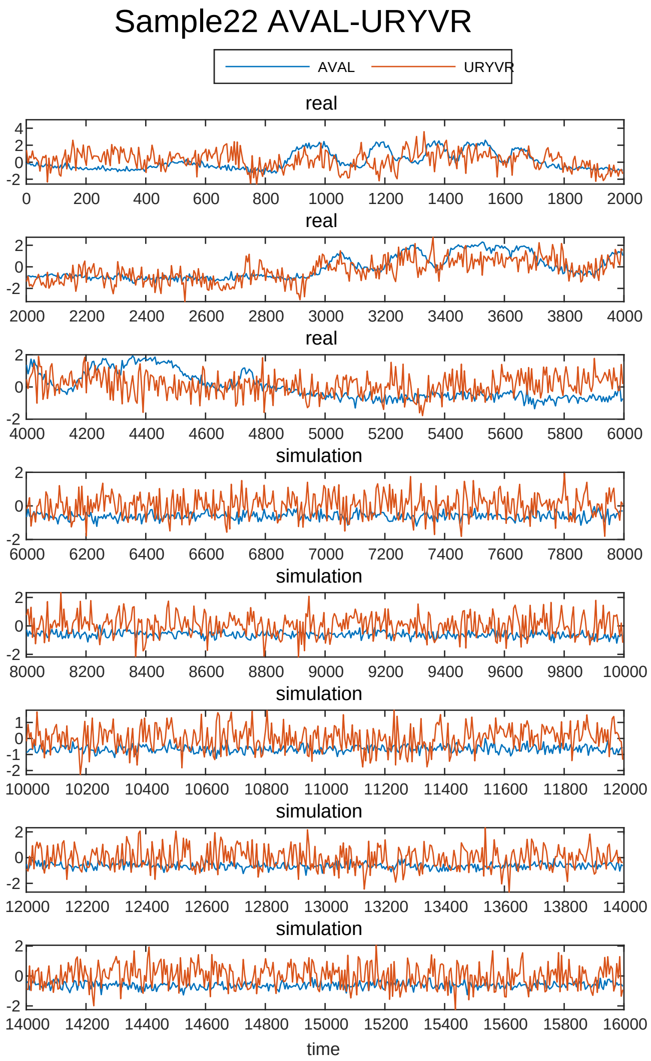

Sample22 AVAL-URYVR
AVAL
URYVR
real
real
real
simulation
simulation
simulation
simulation
simulation
time

Sample2 AVAR-URYVR

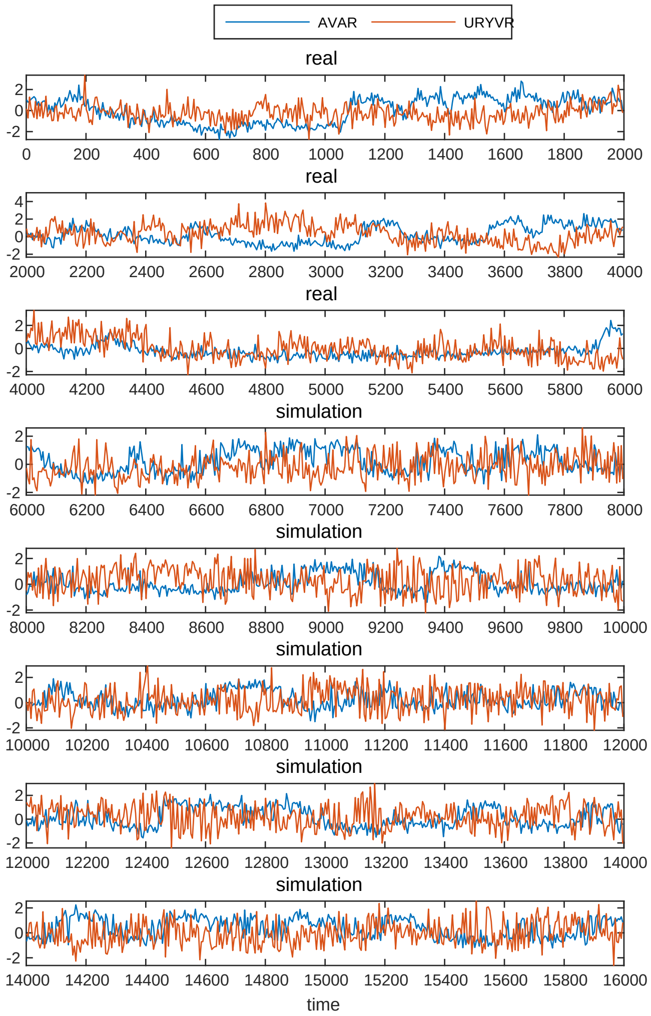


Sample5 AVAR-URYVR

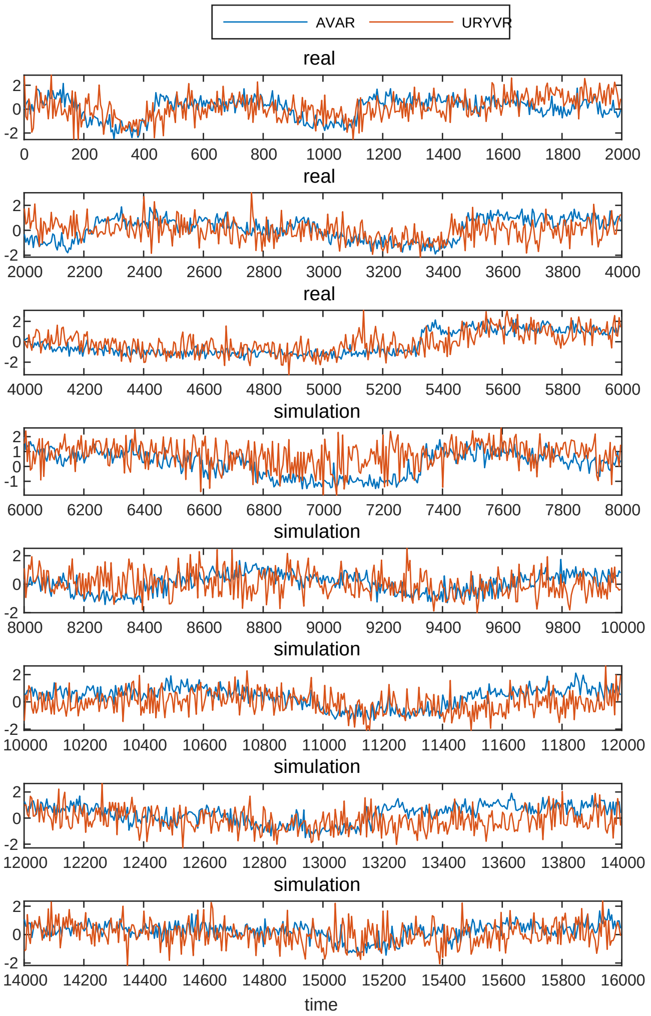


Sample3 AVAR-URYVR

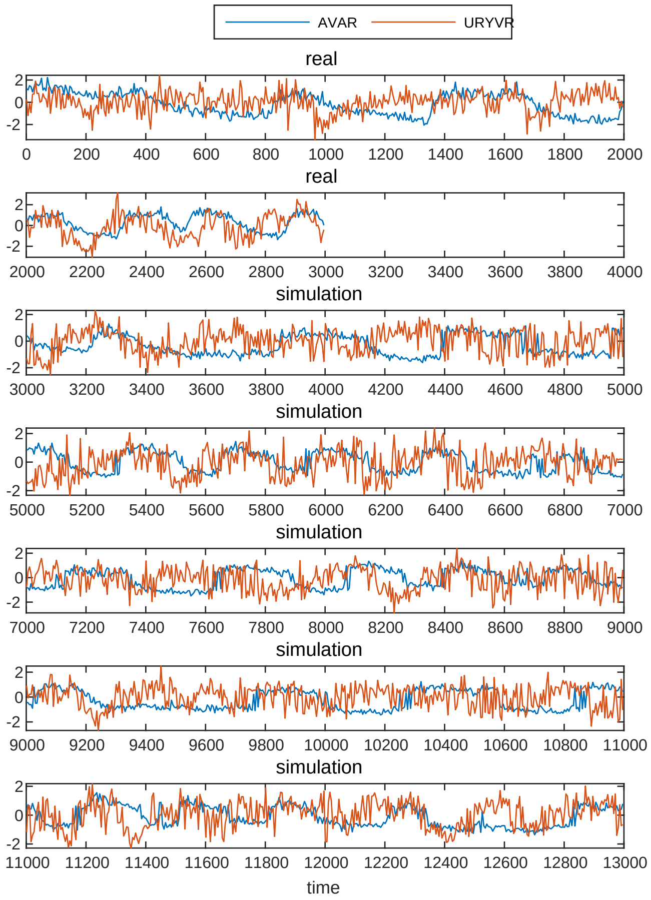


Sample6 AVAR-URYVR

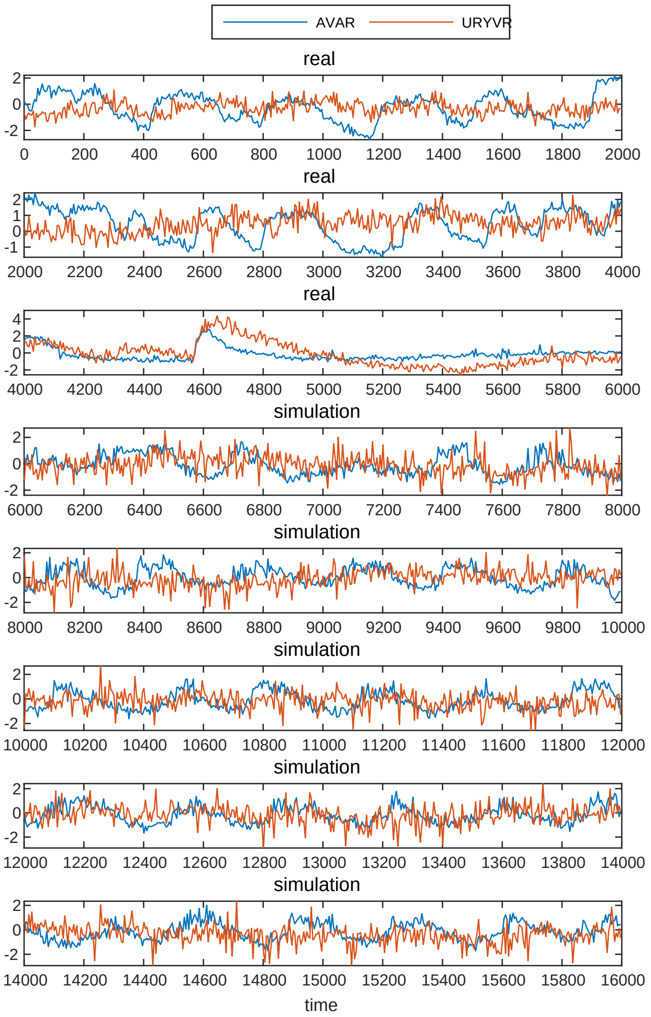

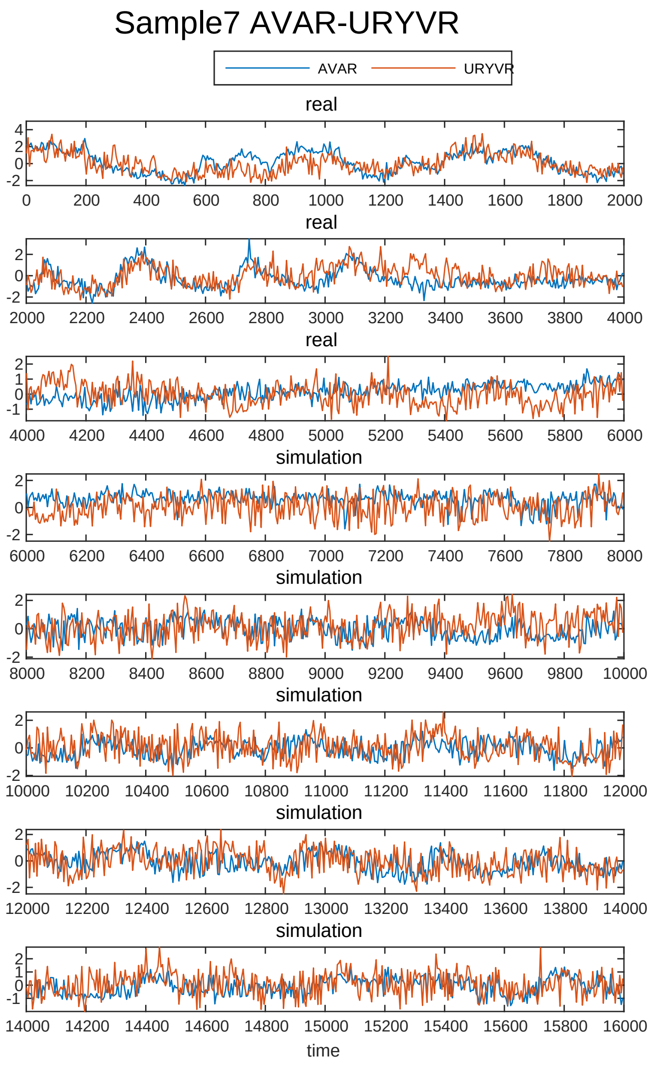
Sample7 AVAR-URYVR
AVAR
URYVR
real
real
real
simulation
simulation
simulation
simulation
simulation
time

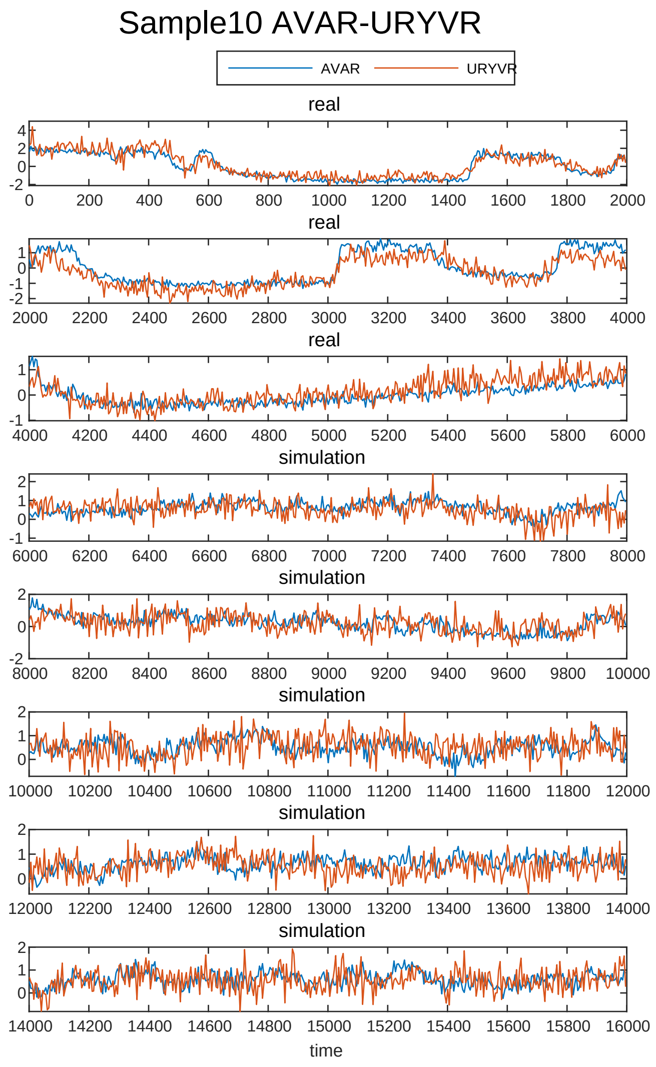
Sample10 AVAR-URYVR
AVAR
URYVR
real
real
real
simulation
simulation
simulation
simulation
simulation
time

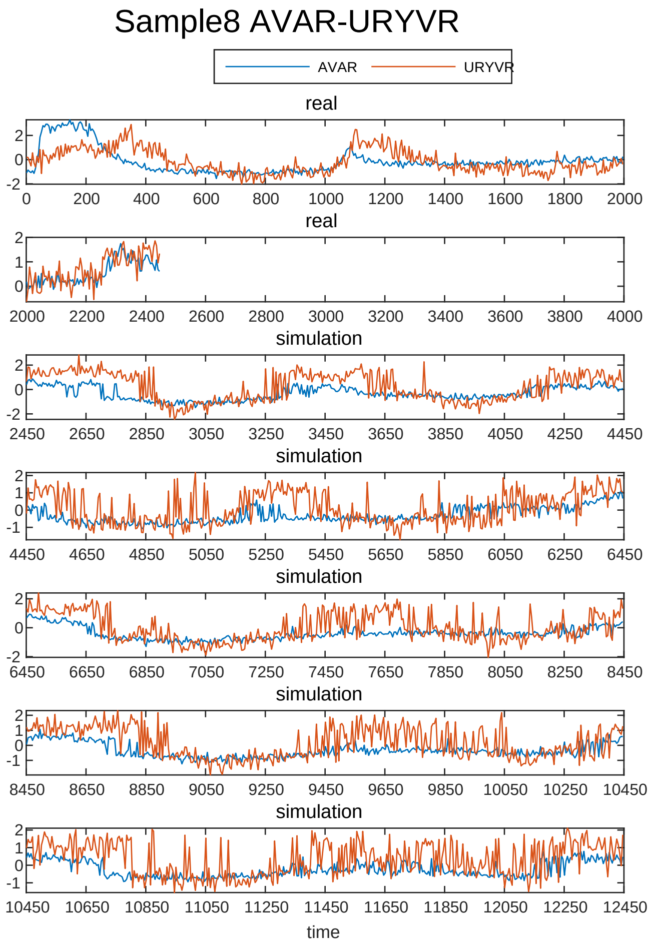
Sample8 AVAR-URYVR
AVAR
URYVR
real
real
simulation
simulation
simulation
simulation
simulation
time

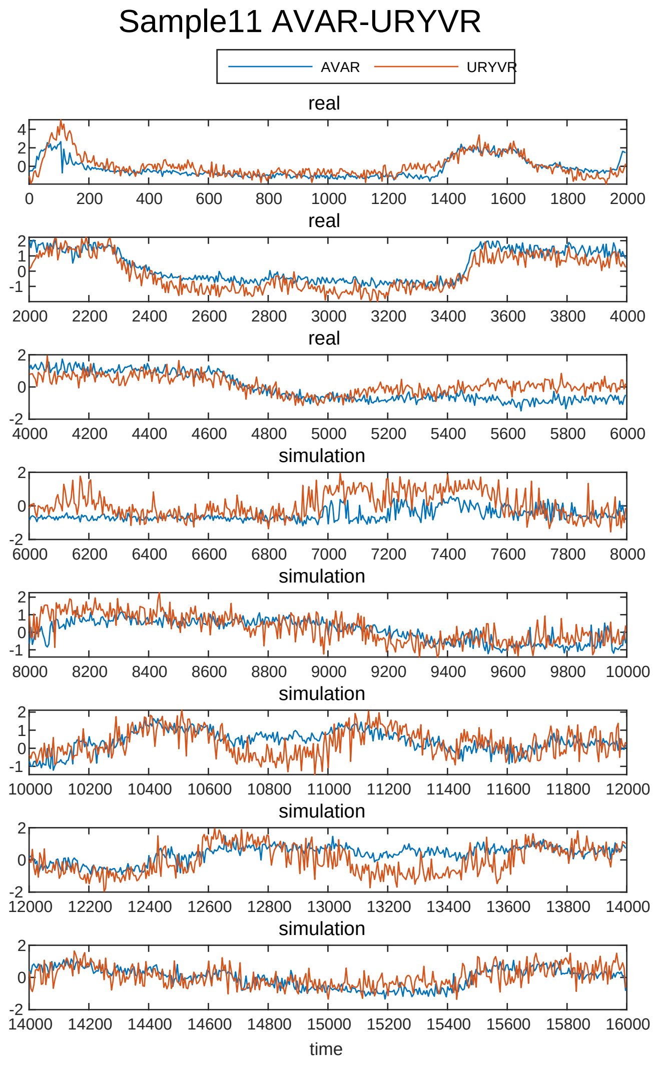
Sample11 AVAR-URYVR
AVAR
URYVR
real
real
real
simulation
simulation
simulation
simulation
simulation
time

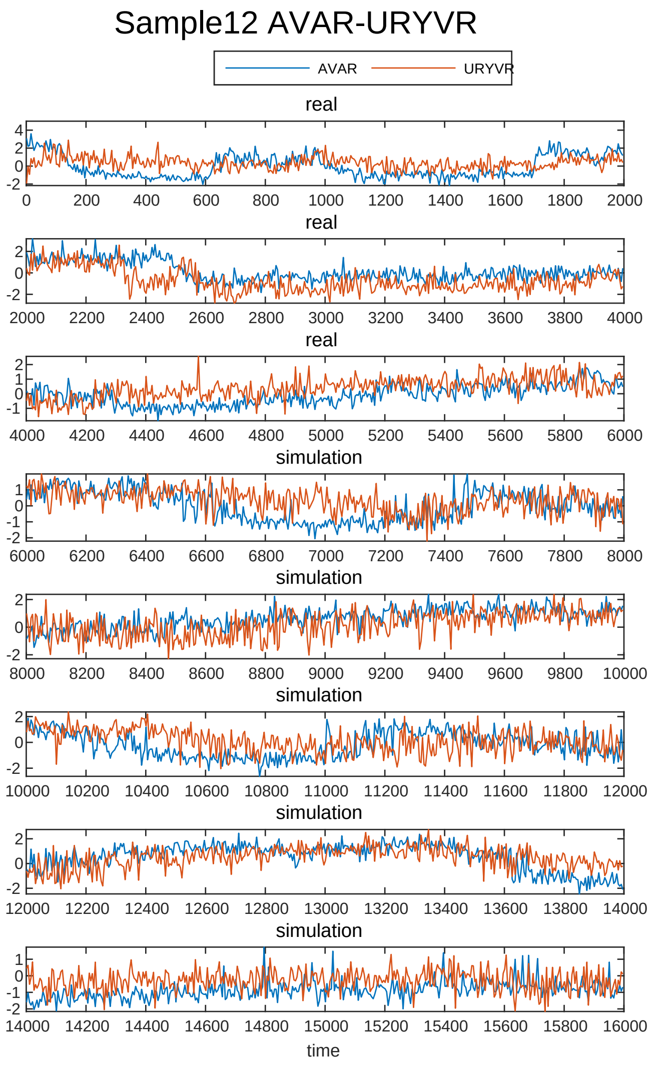
Sample12 AVAR-URYVR
AVAR
URYVR
real
real
real
simulation
simulation
simulation
simulation
simulation
time

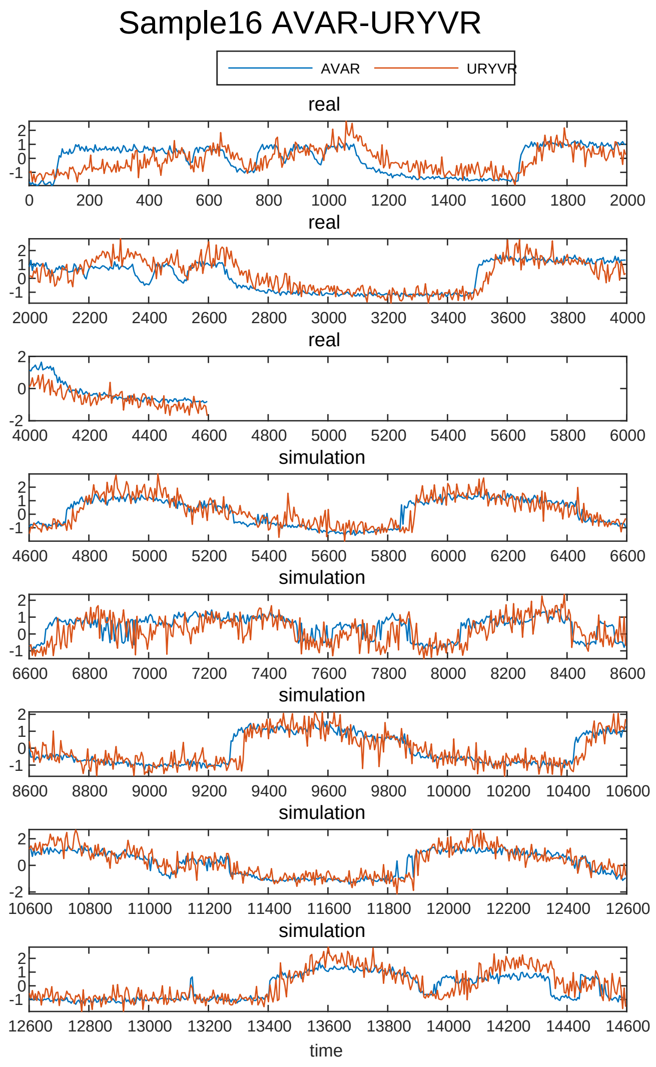
Sample16 AVAR-URYVR
AVAR
URYVR
real
real
real
simulation
simulation
simulation
simulation
simulation
time

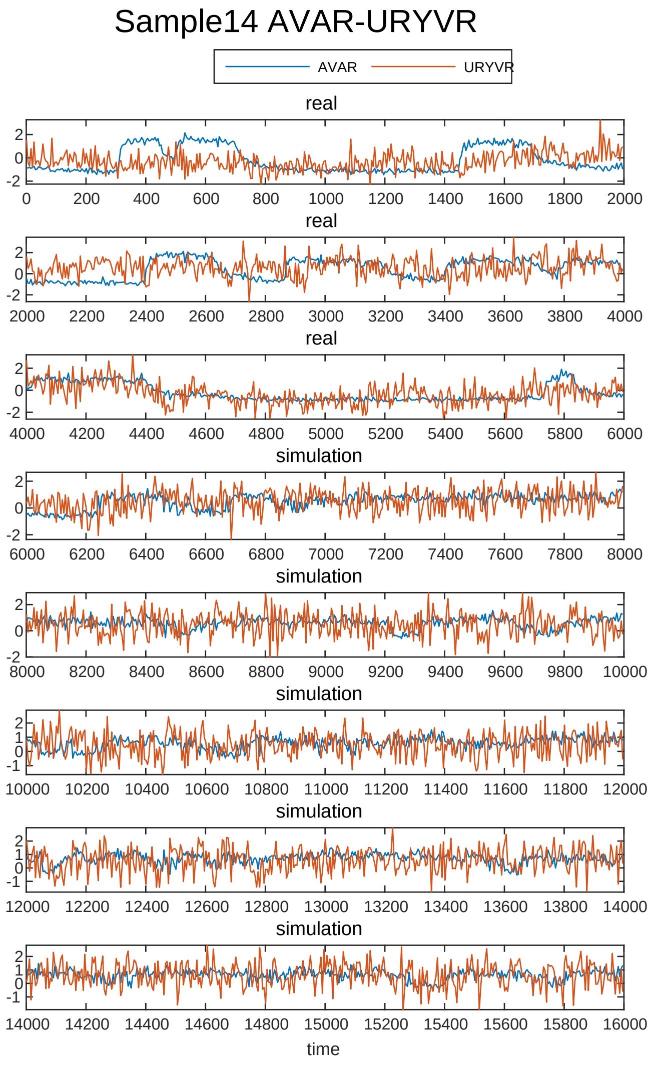
Sample14 AVAR-URYVR
AVAR
URYVR
real
real
real
simulation
simulation
simulation
simulation
simulation
time

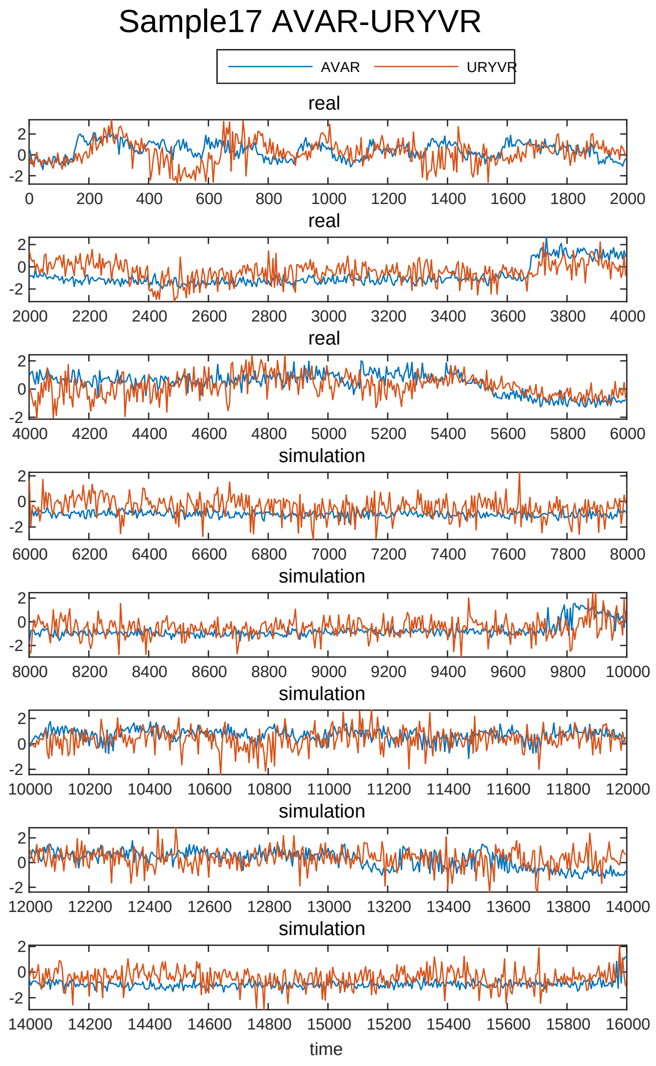
Sample17 AVAR-URYVR
AVAR
URYVR
real
real
real
simulation
simulation
simulation
simulation
simulation
time

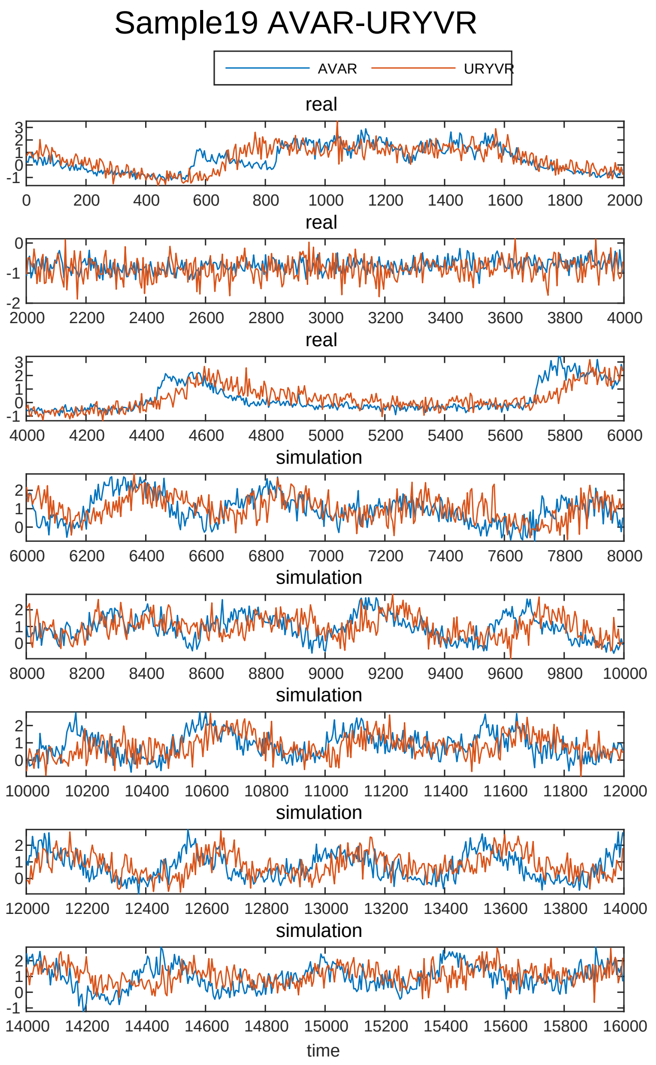
Sample19 AVAR-URYVR
AVAR
URYVR
real
real
real
simulation
simulation
simulation
simulation
simulation
time

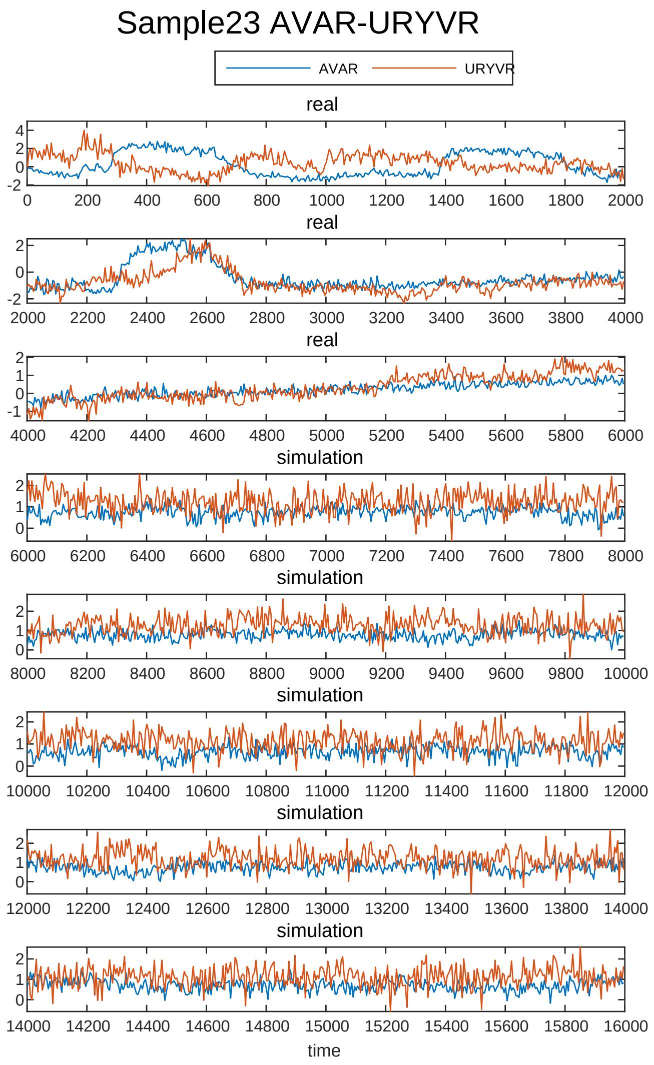
Sample23 AVAR-URYVR
AVAR
URYVR
real
real
real
simulation
simulation
simulation
simulation
simulation
time

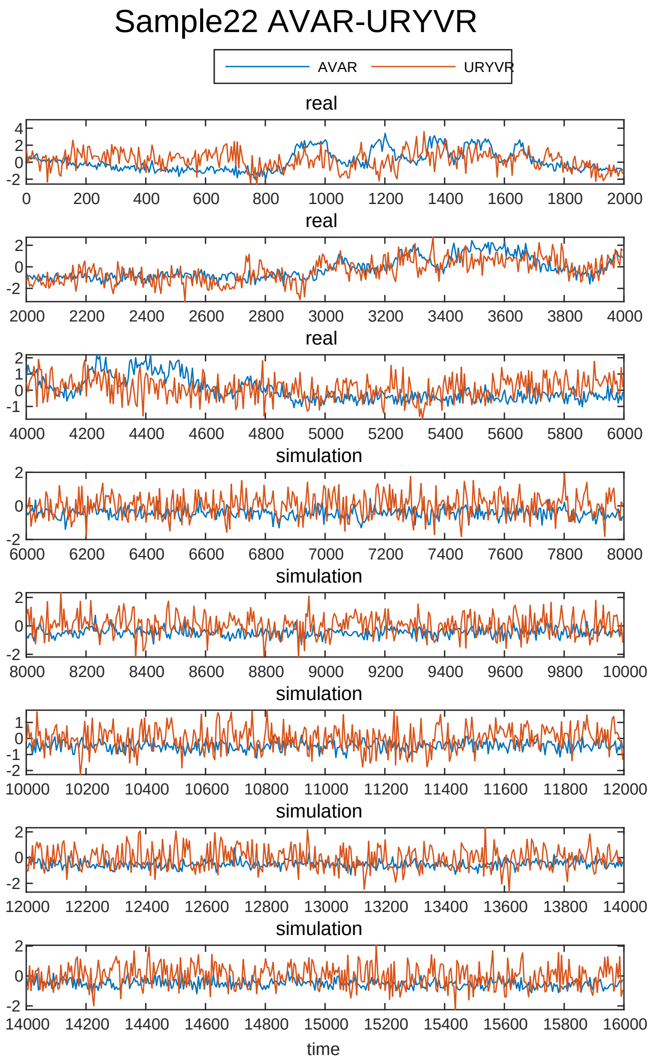
Sample22 AVAR-URYVR
AVAR
URYVR
real
real
real
simulation
simulation
simulation
simulation
simulation
time

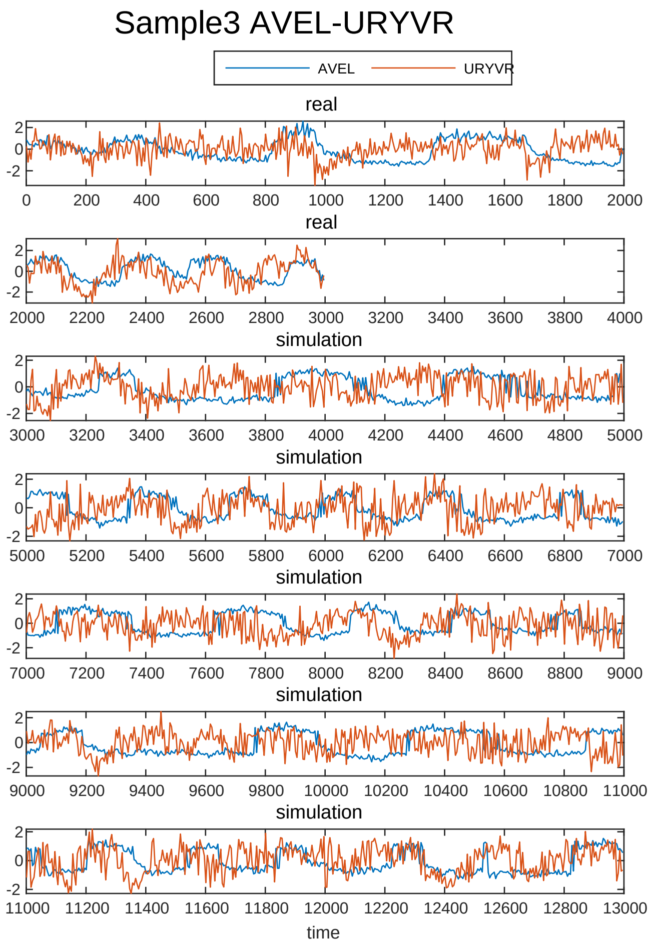
Sample3 AVEL-URYVR
AVEL
URYVR
real
real
simulation
simulation
simulation
simulation
simulation
time

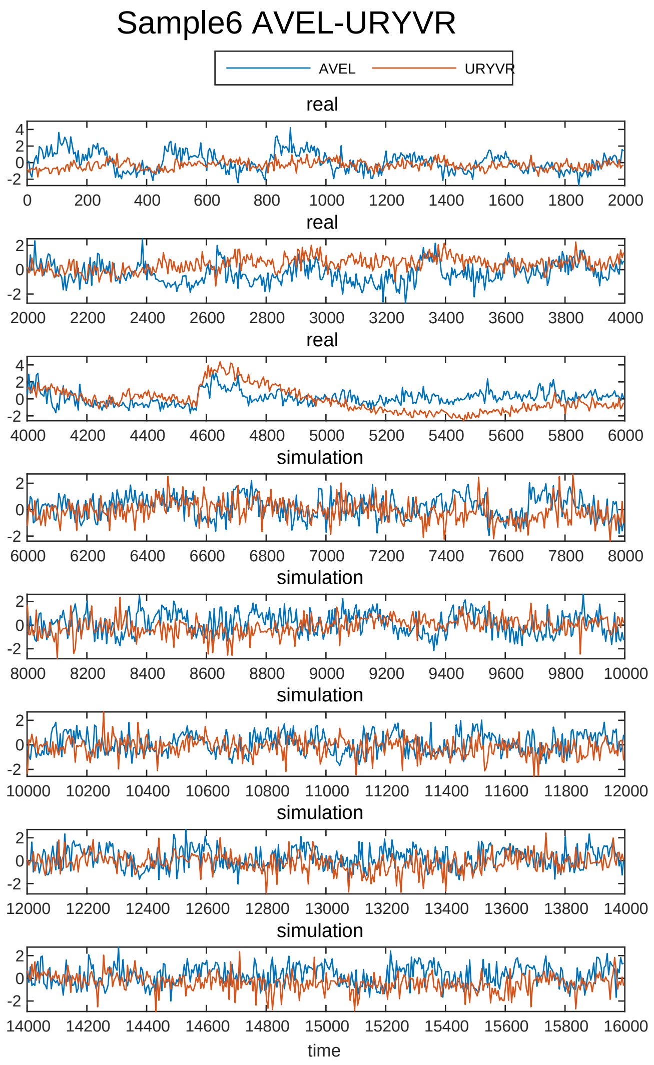
Sample6 AVEL-URYVR
AVEL
URYVR
real
real
real
simulation
simulation
simulation
simulation
simulation
time

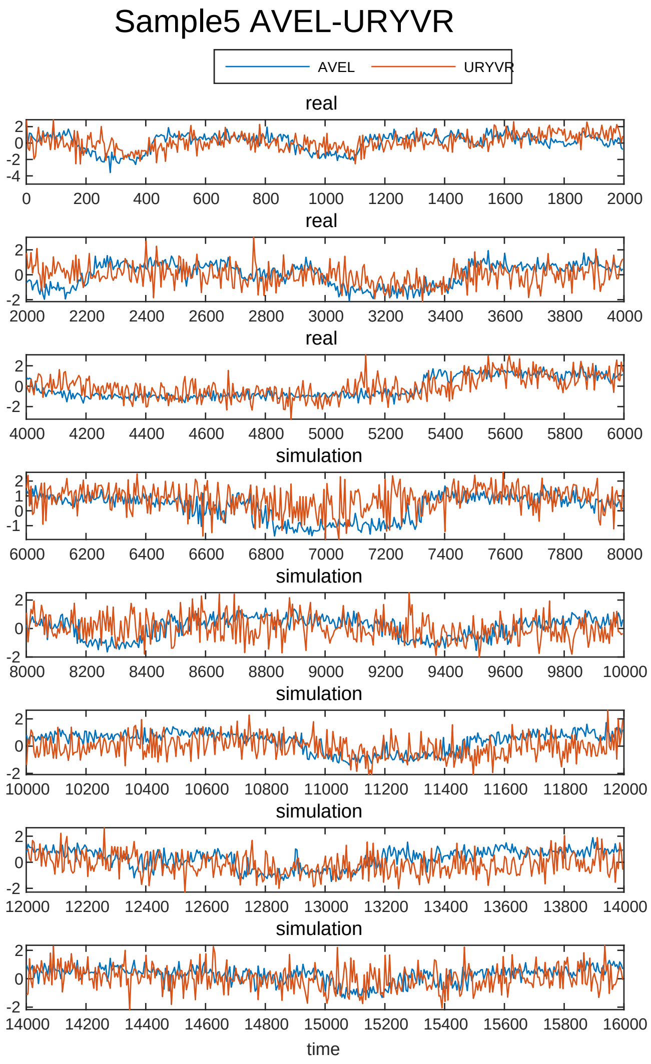
Sample5 AVEL-URYVR
AVEL
URYVR
real
real
real
simulation
simulation
simulation
simulation
simulation
time

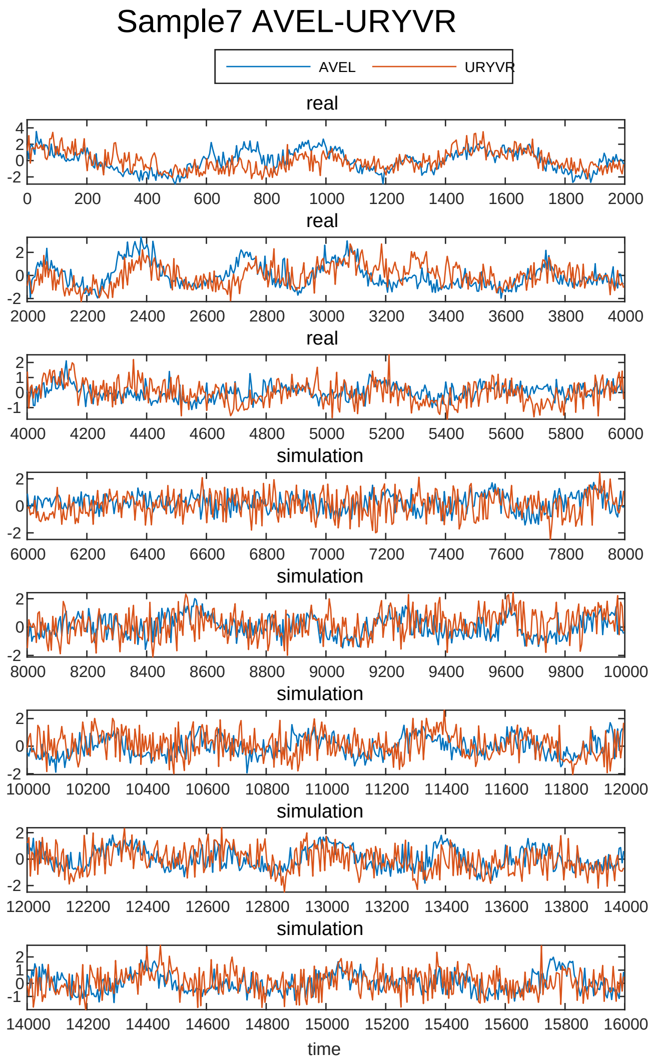
Sample7 AVEL-URYVR
AVEL
URYVR
real
real
real
simulation
simulation
simulation
simulation
simulation
time

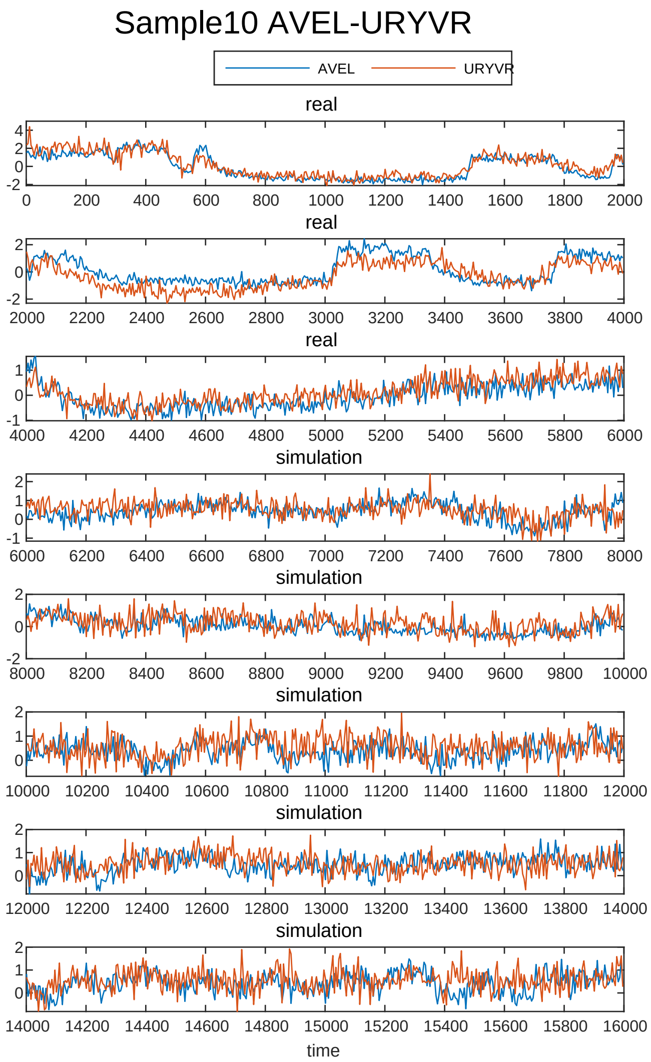
Sample10 AVEL-URYVR
AVEL
URYVR
real
real
real
simulation
simulation
simulation
simulation
simulation
time

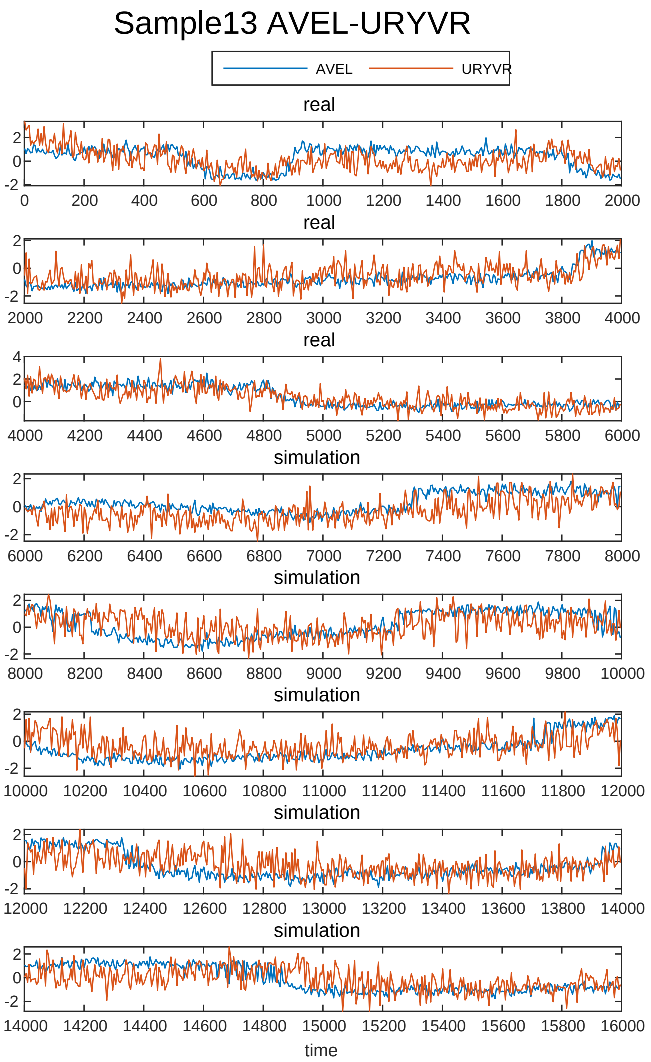
Sample13 AVEL-URYVR
AVEL
URYVR
real
real
real
simulation
simulation
simulation
simulation
simulation
time

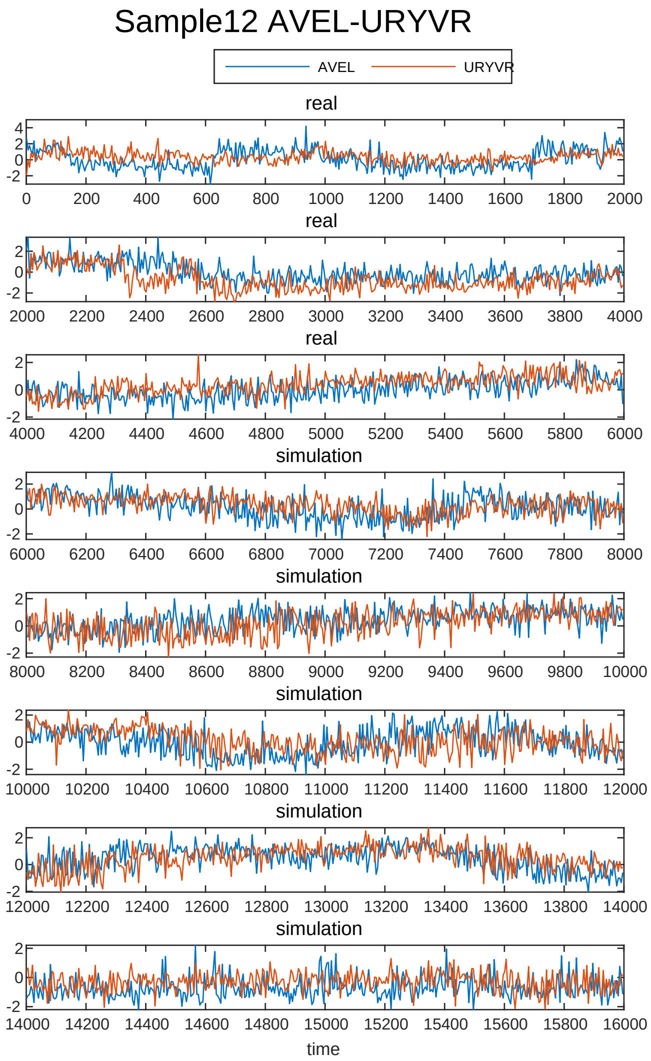
Sample12 AVEL-URYVR
AVEL
URYVR
real
real
real
simulation
simulation
simulation
simulation
simulation
time

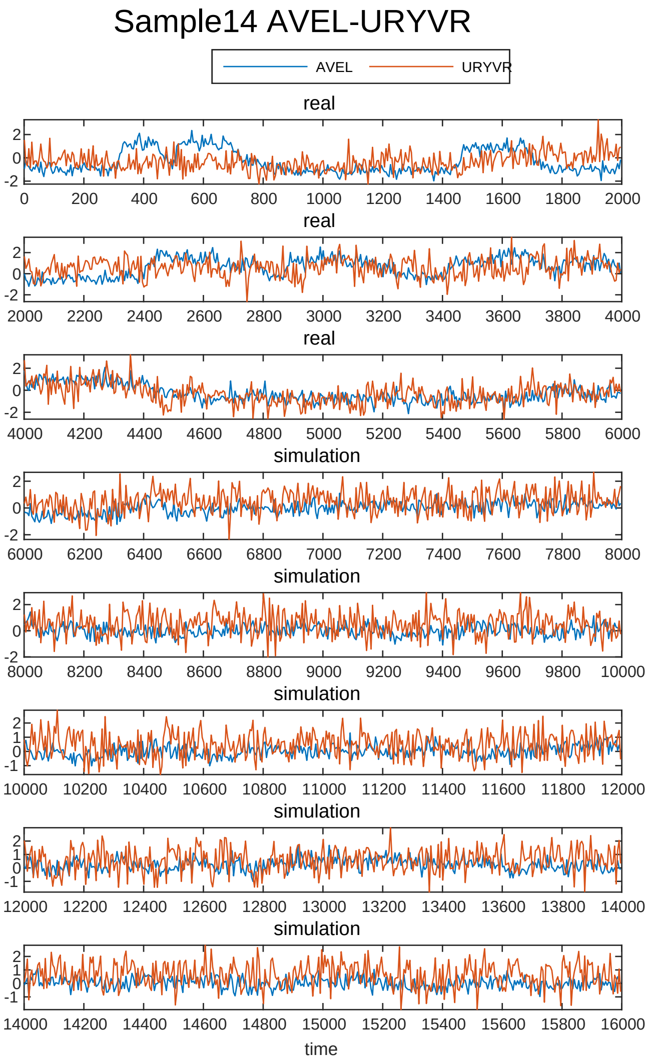
Sample14 AVEL-URYVR
AVEL
URYVR
real
real
real
simulation
simulation
simulation
simulation
simulation
time

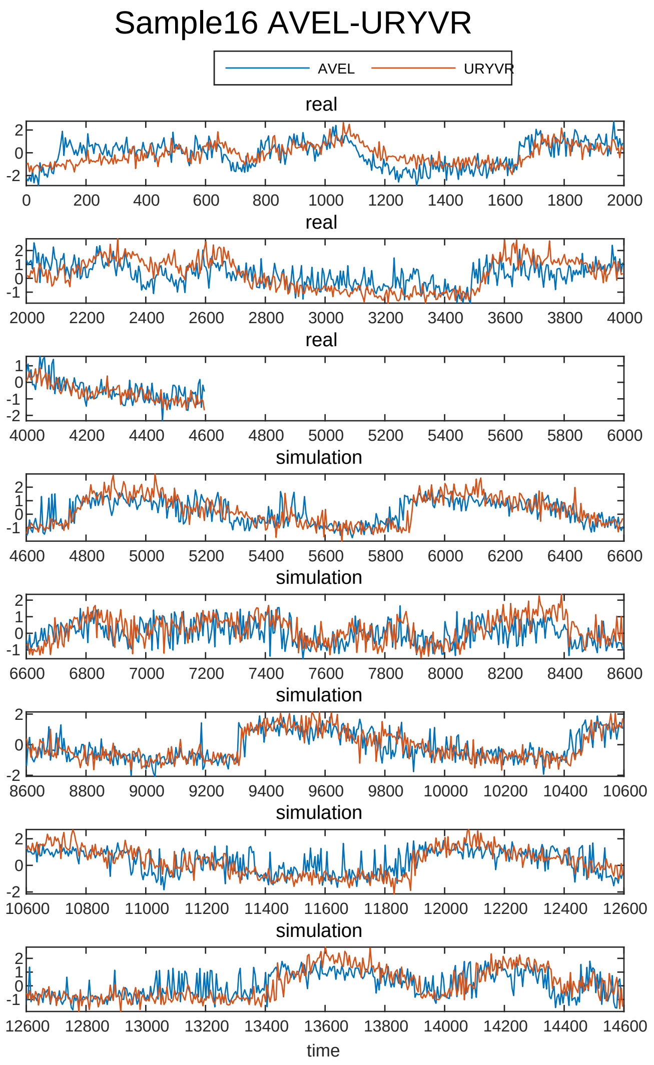
Sample16 AVEL-URYVR
AVEL
URYVR
real
simulation
time

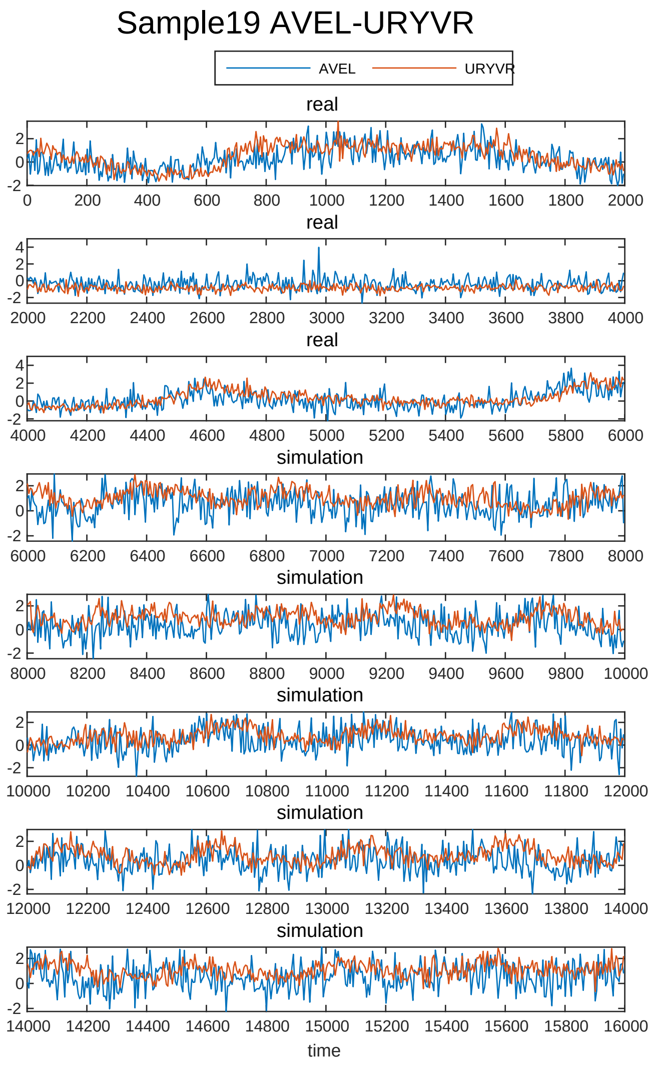
Sample19 AVEL-URYVR
AVEL
URYVR
real
simulation
time

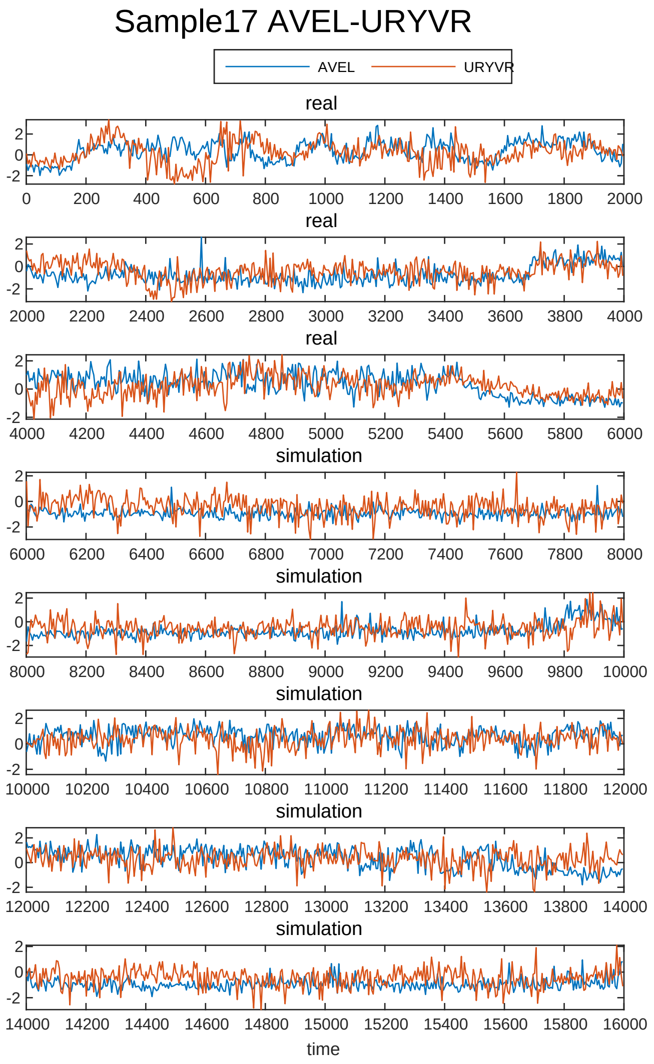
Sample17 AVEL-URYVR
AVEL
URYVR
real
simulation
time

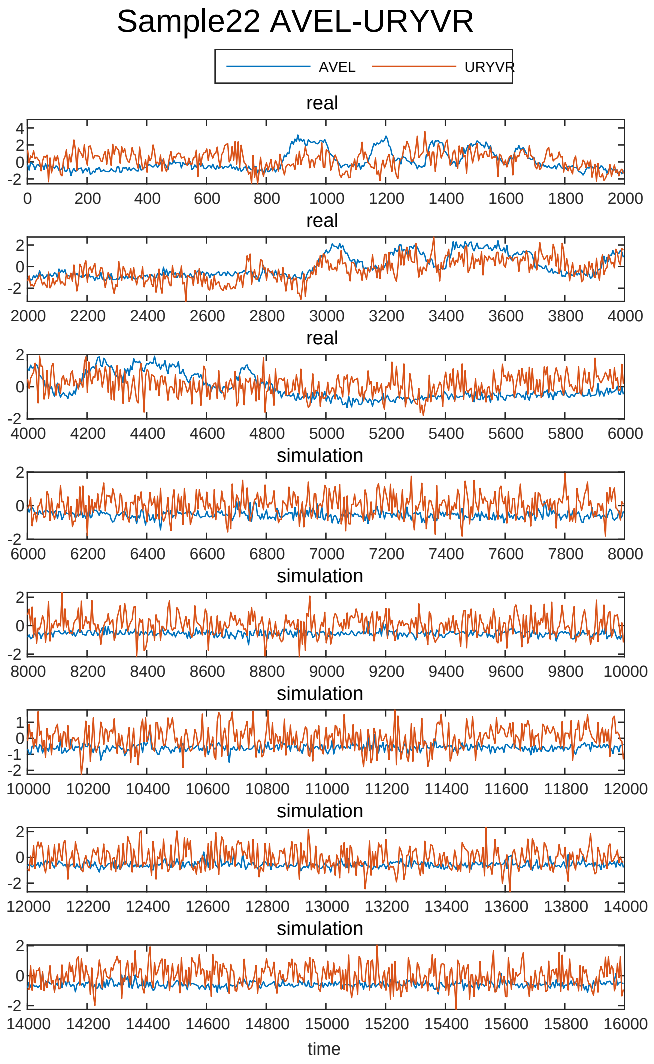
Sample22 AVEL-URYVR
AVEL
URYVR
real
simulation
time

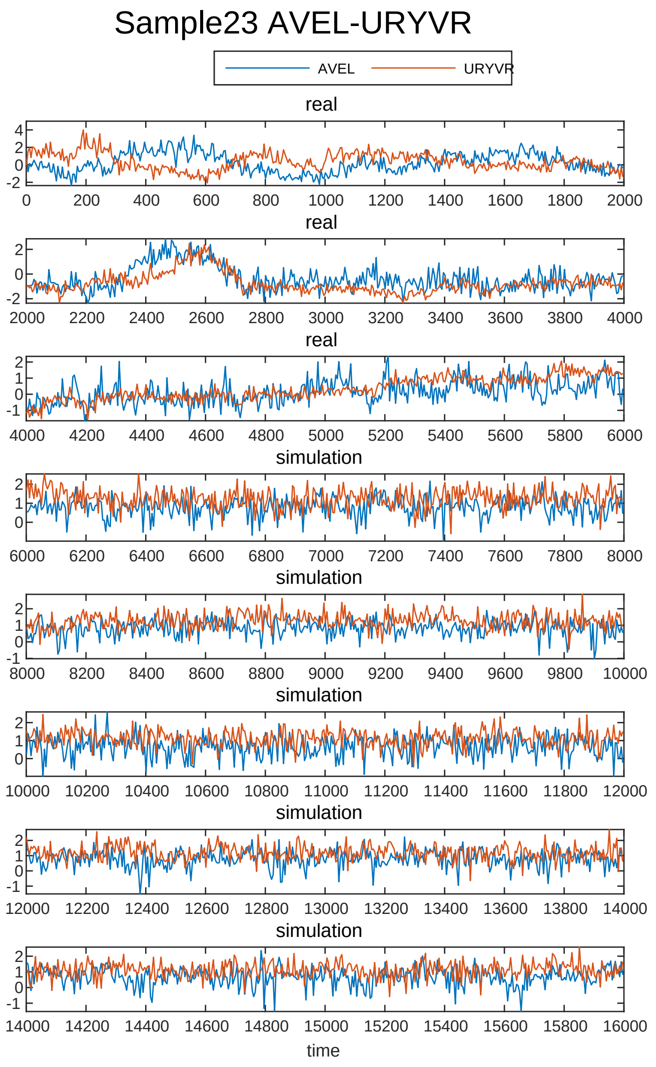
Sample23 AVEL-URYVR
AVEL
URYVR
real
real
real
simulation
simulation
simulation
simulation
simulation
time

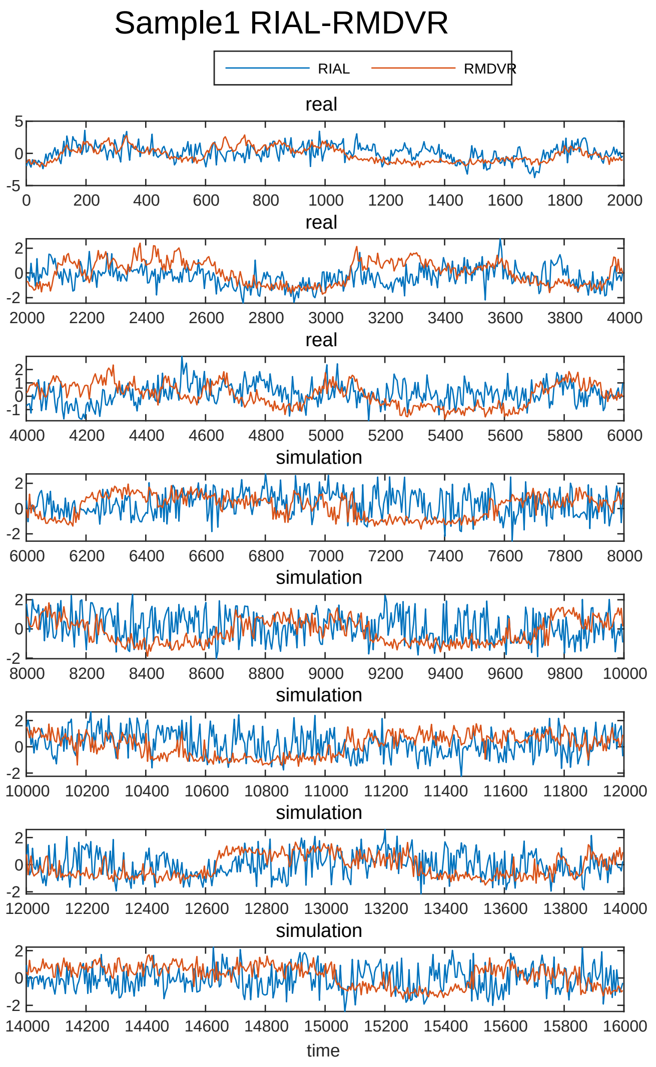
Sample1 RIAL-RMDVR
RIAL
RMDVR
real
real
real
simulation
simulation
simulation
simulation
simulation
time

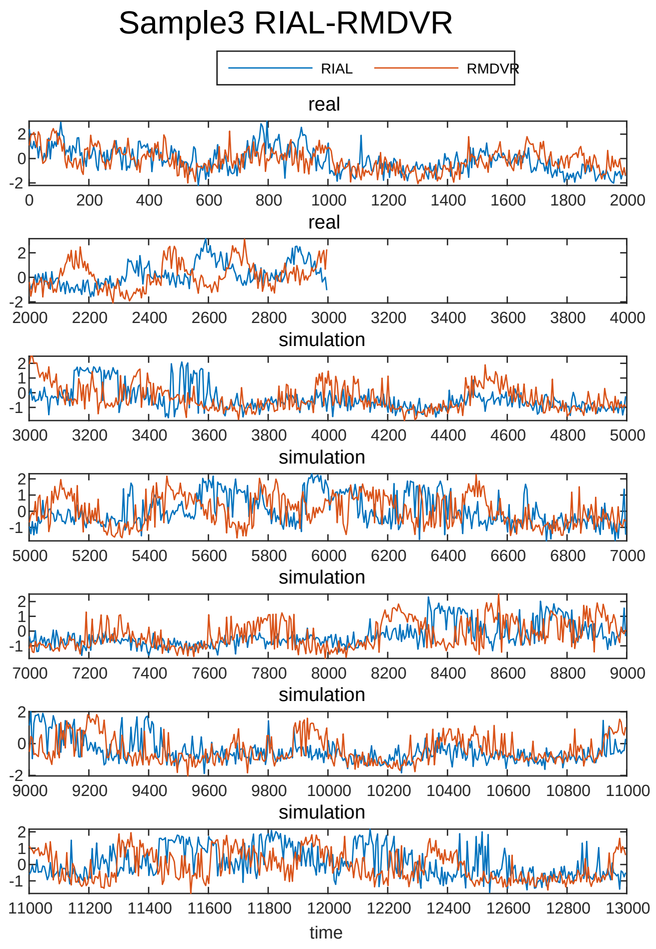
Sample3 RIAL-RMDVR
RIAL
RMDVR
real
real
simulation
simulation
simulation
simulation
simulation
time

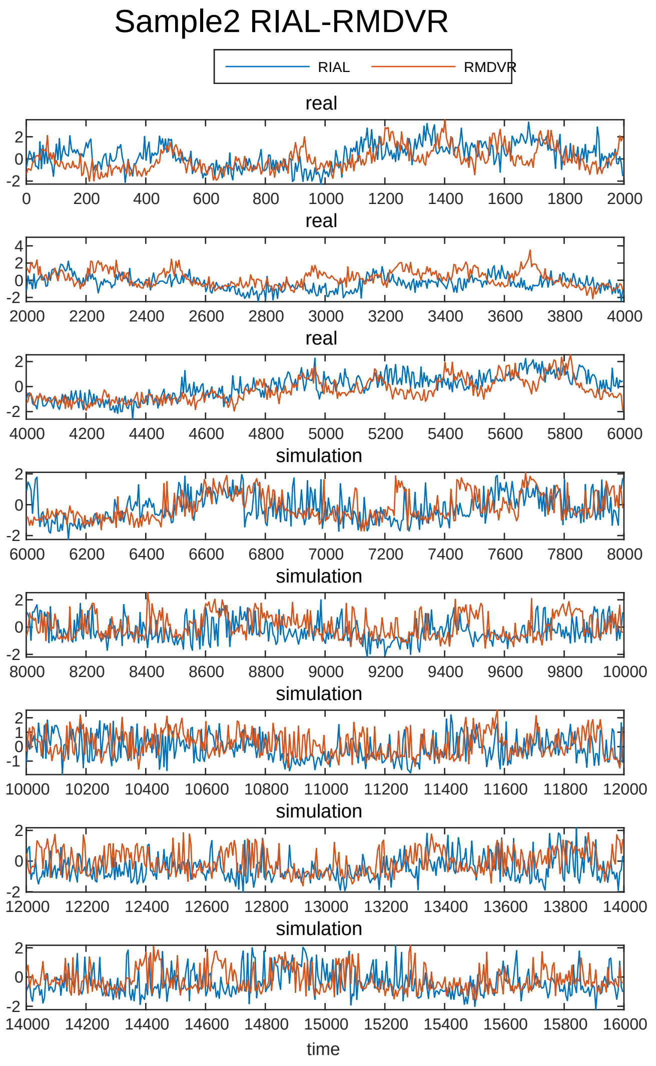
Sample2 RIAL-RMDVR
RIAL
RMDVR
real
real
real
simulation
simulation
simulation
simulation
simulation
time

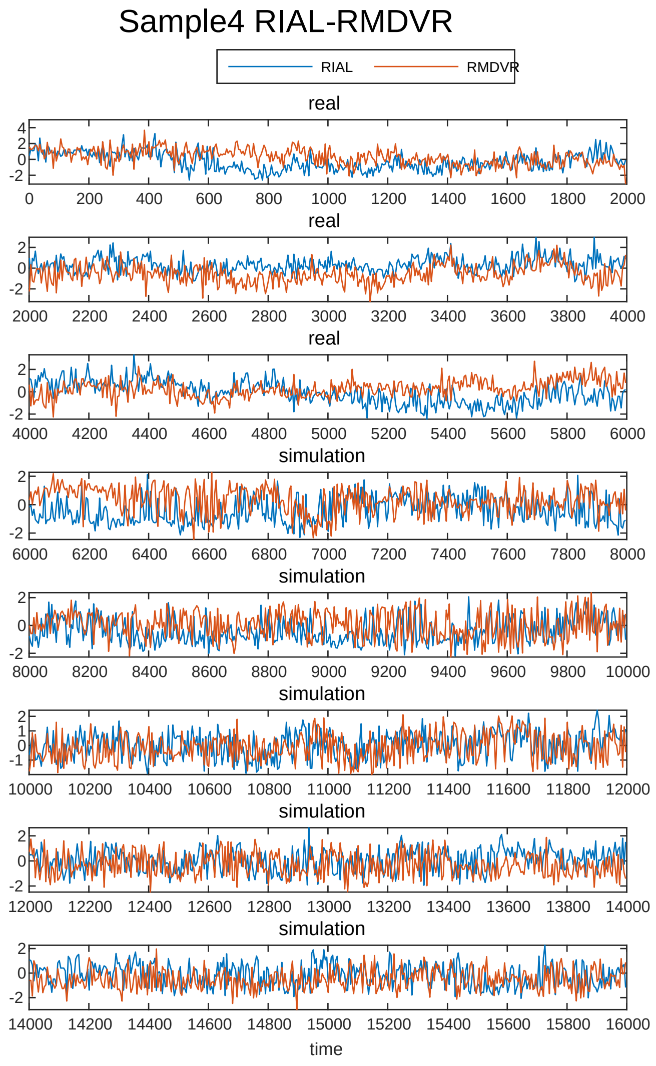
Sample4 RIAL-RMDVR
RIAL
RMDVR
real
real
real
simulation
simulation
simulation
simulation
simulation
time

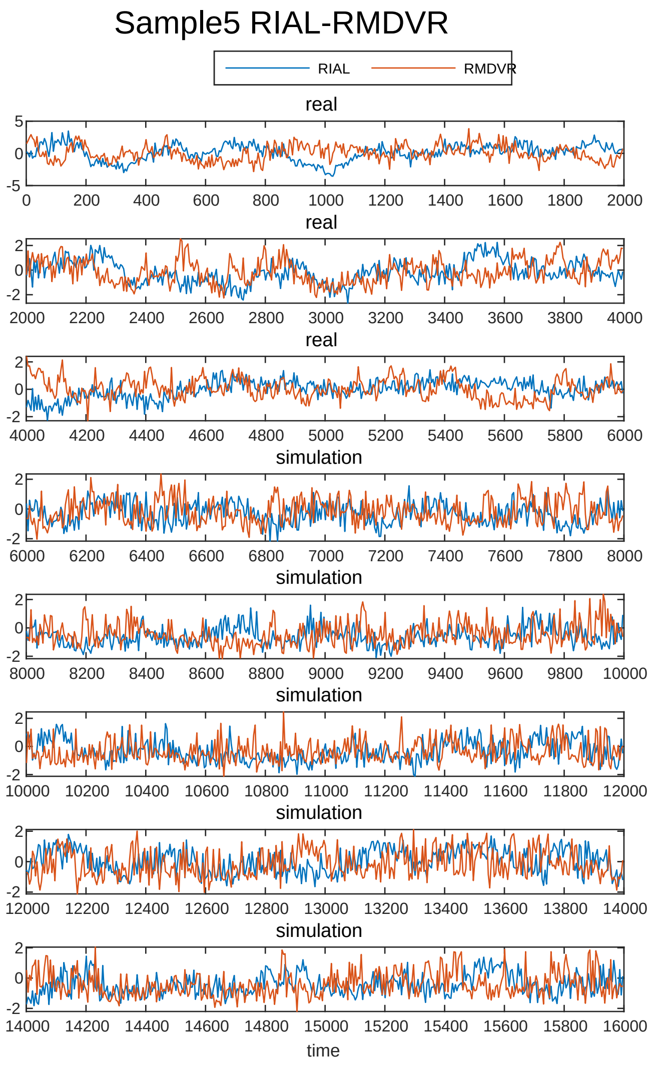
Sample5 RIAL-RMDVR
RIAL
RMDVR
real
real
real
simulation
simulation
simulation
simulation
simulation
time

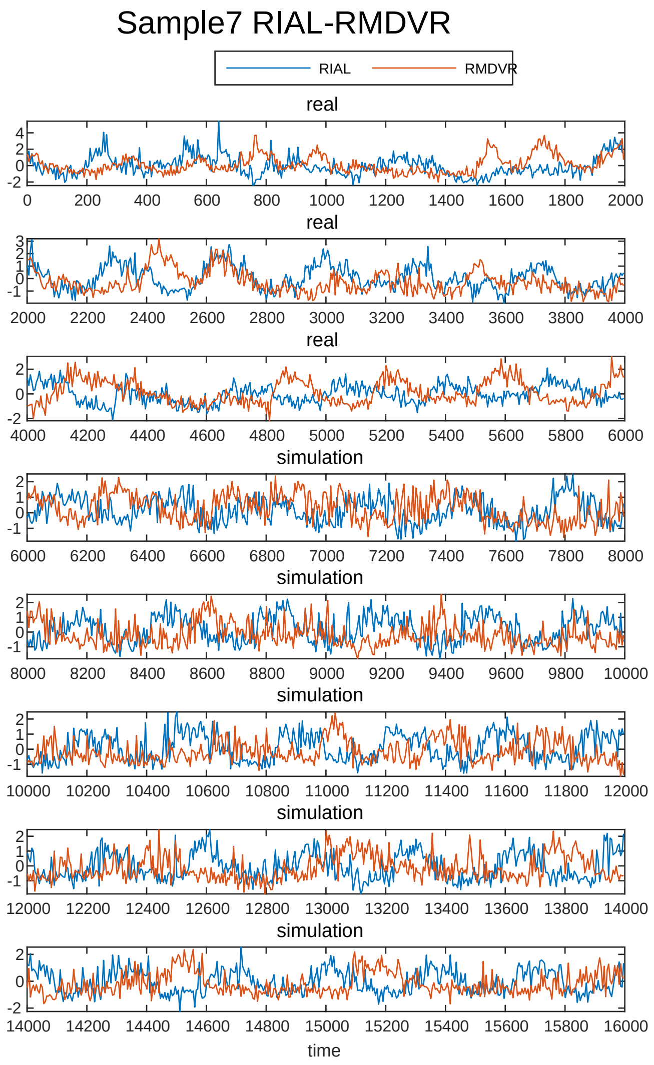
Sample7 RIAL-RMDVR
RIAL
RMDVR
real
real
real
simulation
simulation
simulation
simulation
simulation
time

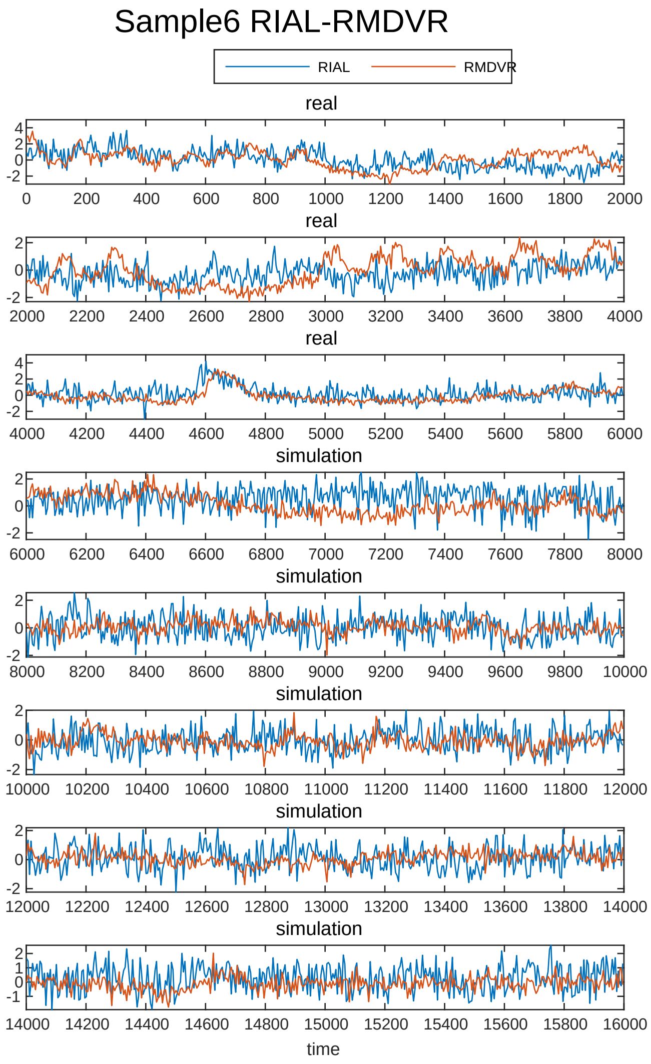
Sample6 RIAL-RMDVR
RIAL
RMDVR
real
real
real
simulation
simulation
simulation
simulation
simulation
time

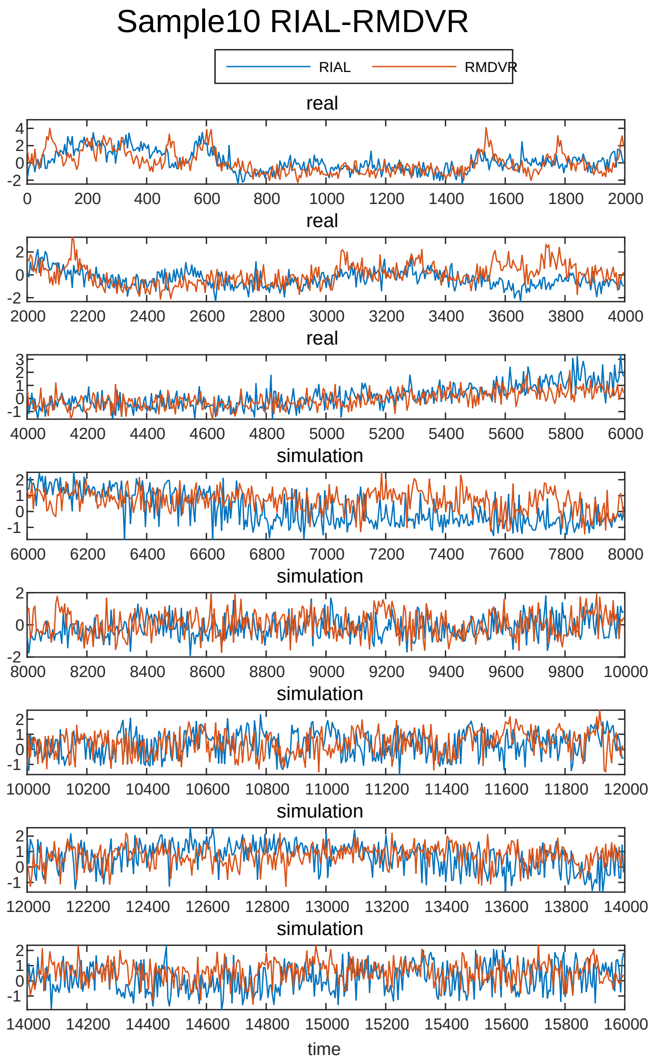
Sample10 RIAL-RMDVR
RIAL
RMDVR
real
real
real
simulation
simulation
simulation
simulation
simulation
time

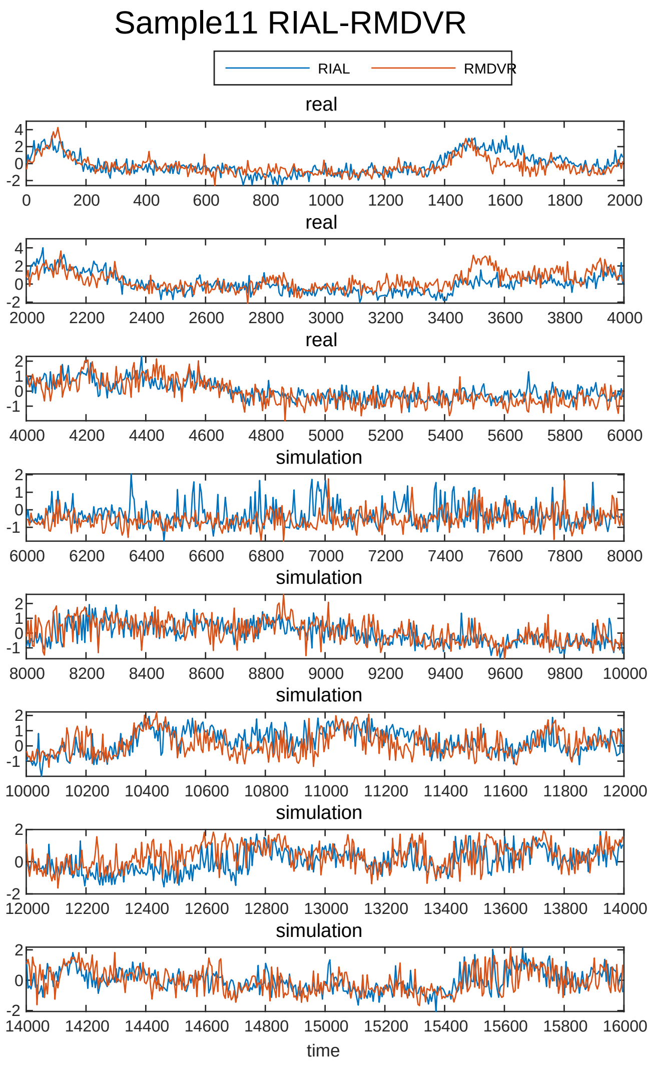
Sample11 RIAL-RMDVR
RIAL
RMDVR
real
real
real
simulation
simulation
simulation
simulation
simulation
time

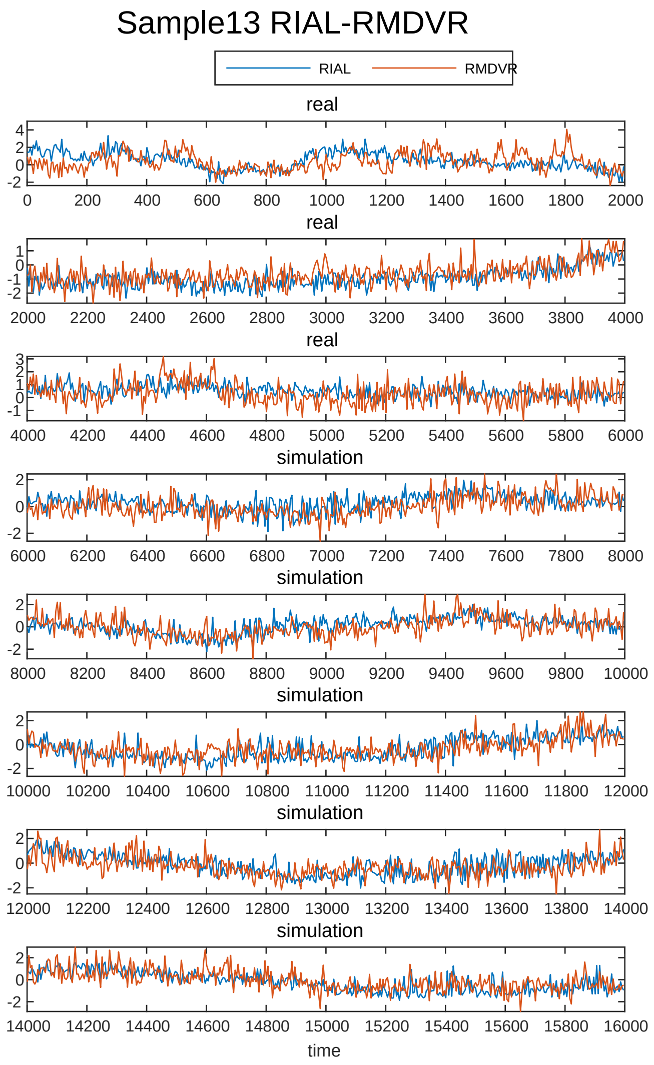
Sample13 RIAL-RMDVR
RIAL
RMDVR
real
real
real
simulation
simulation
simulation
simulation
simulation
time

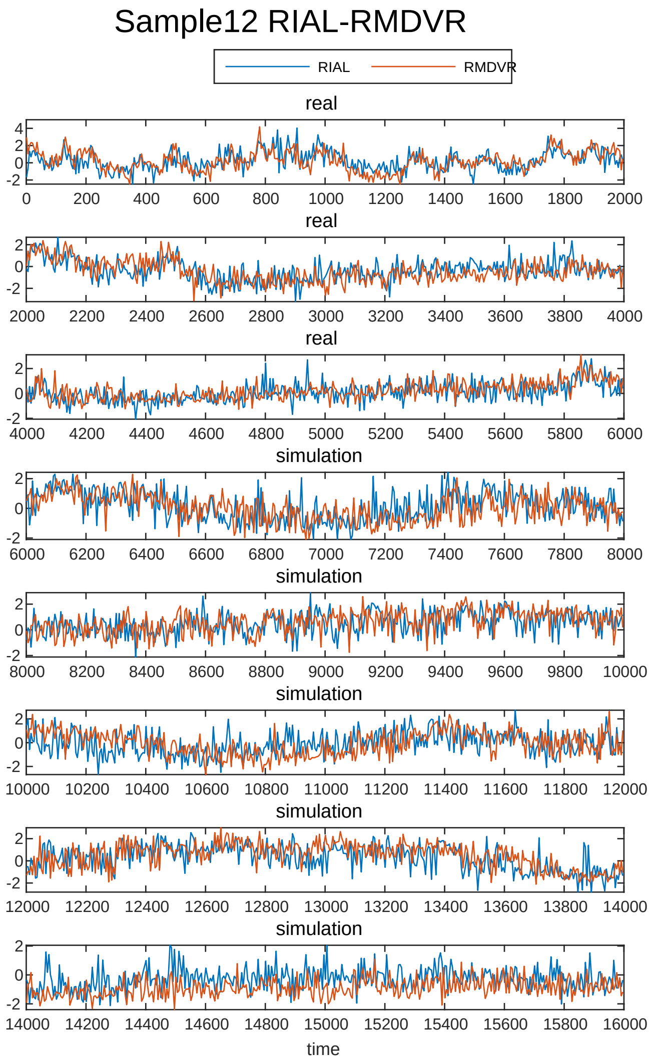
Sample12 RIAL-RMDVR
RIAL
RMDVR
real
real
real
simulation
simulation
simulation
simulation
simulation
time

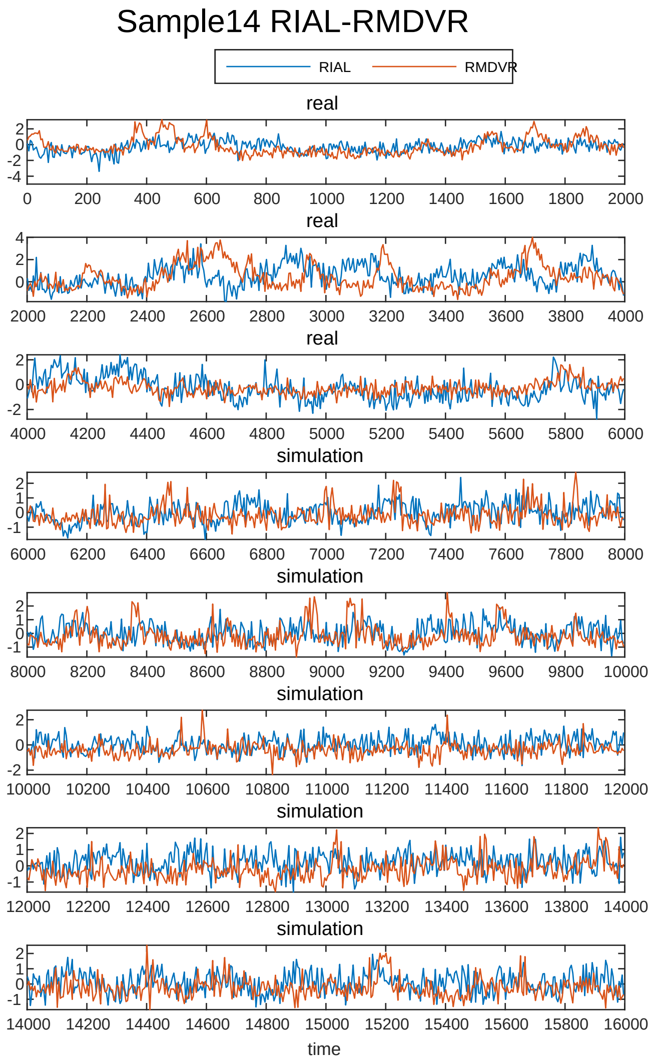
Sample14 RIAL-RMDVR
RIAL
RMDVR
real
real
real
simulation
simulation
simulation
simulation
simulation
time

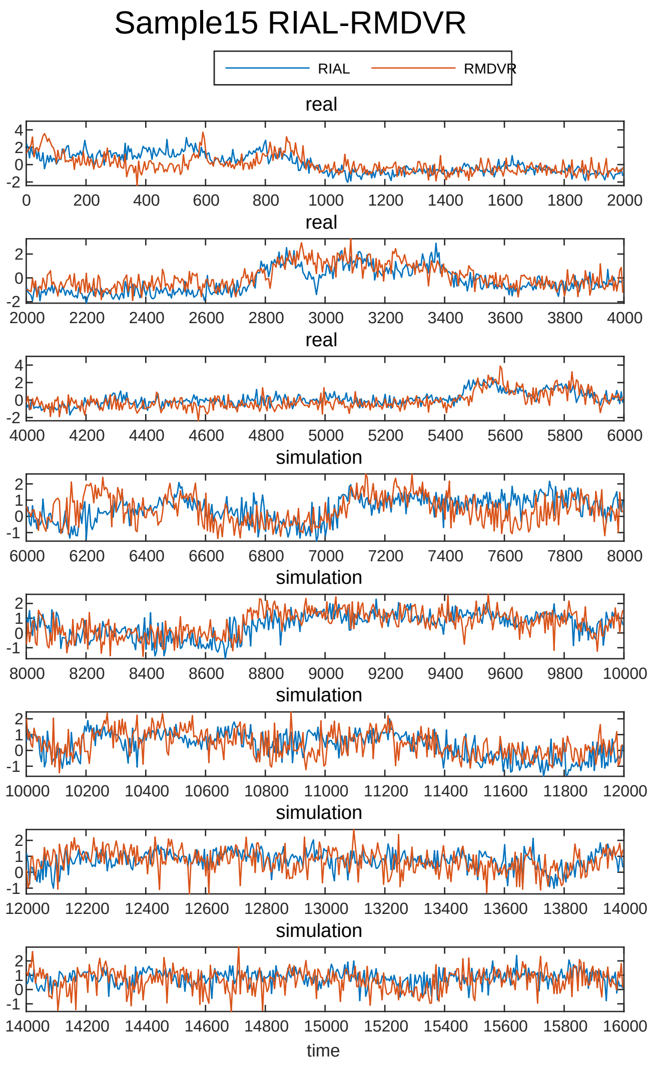
Sample15 RIAL-RMDVR
RIAL
RMDVR
real
simulation
time

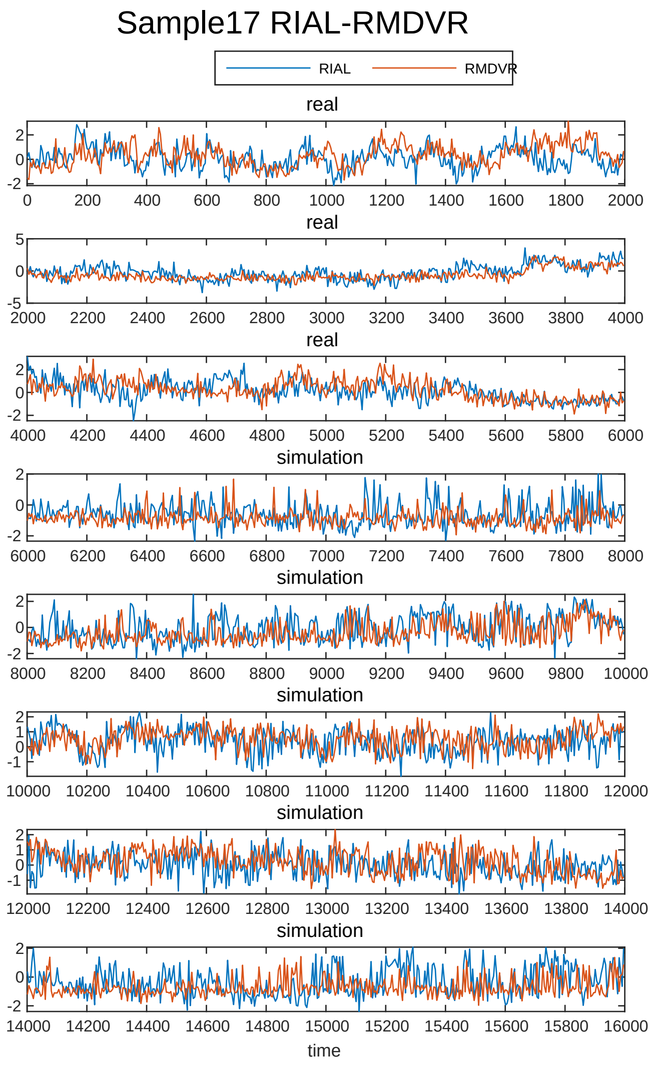
Sample17 RIAL-RMDVR
RIAL
RMDVR
real
simulation
time

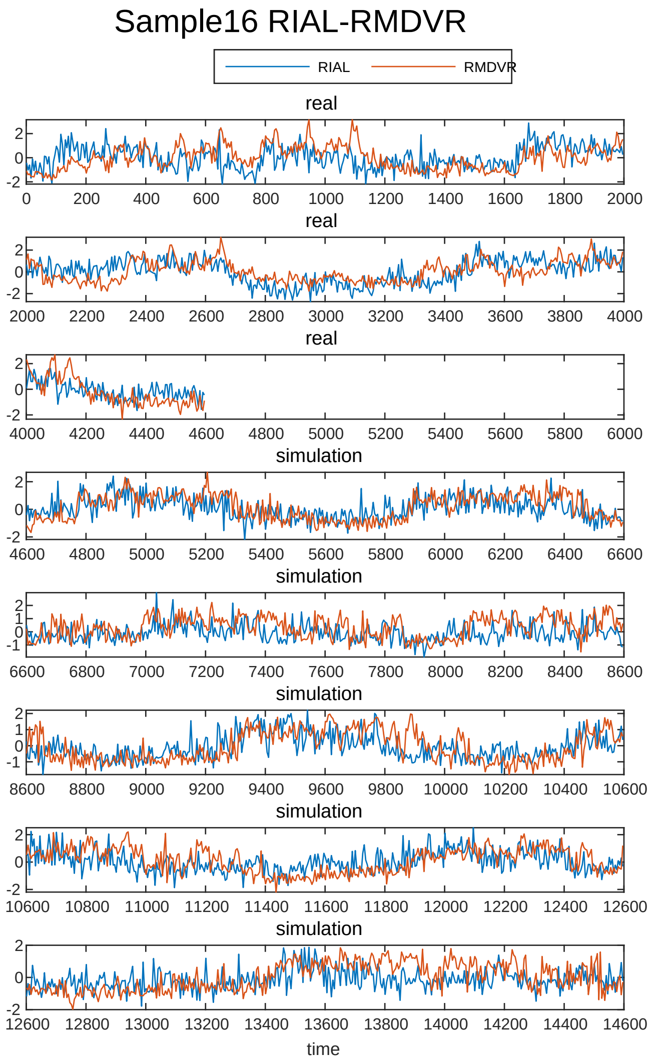
Sample16 RIAL-RMDVR
RIAL
RMDVR
real
simulation
time

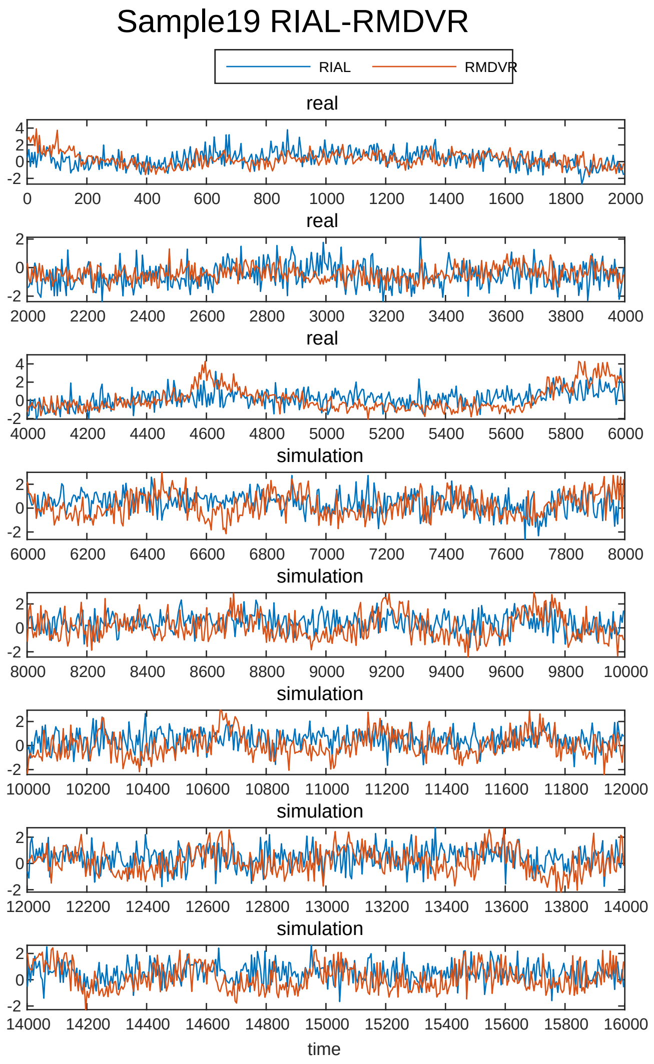
Sample19 RIAL-RMDVR
RIAL
RMDVR
real
simulation
time

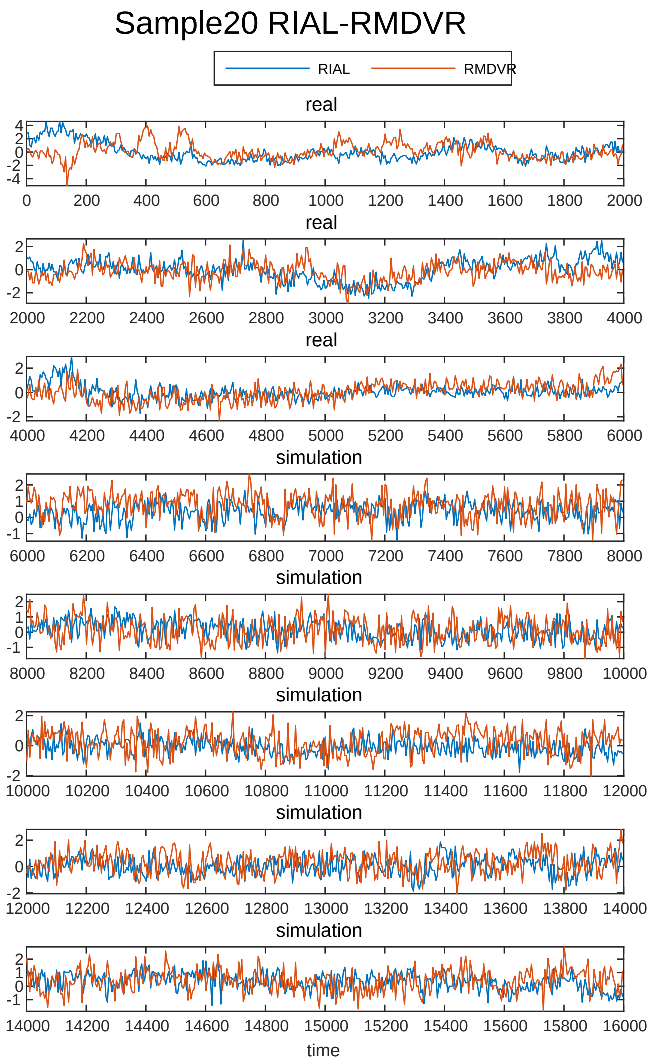
Sample20 RIAL-RMDVR
RIAL
RMDVR
real
real
real
simulation
simulation
simulation
simulation
simulation
time

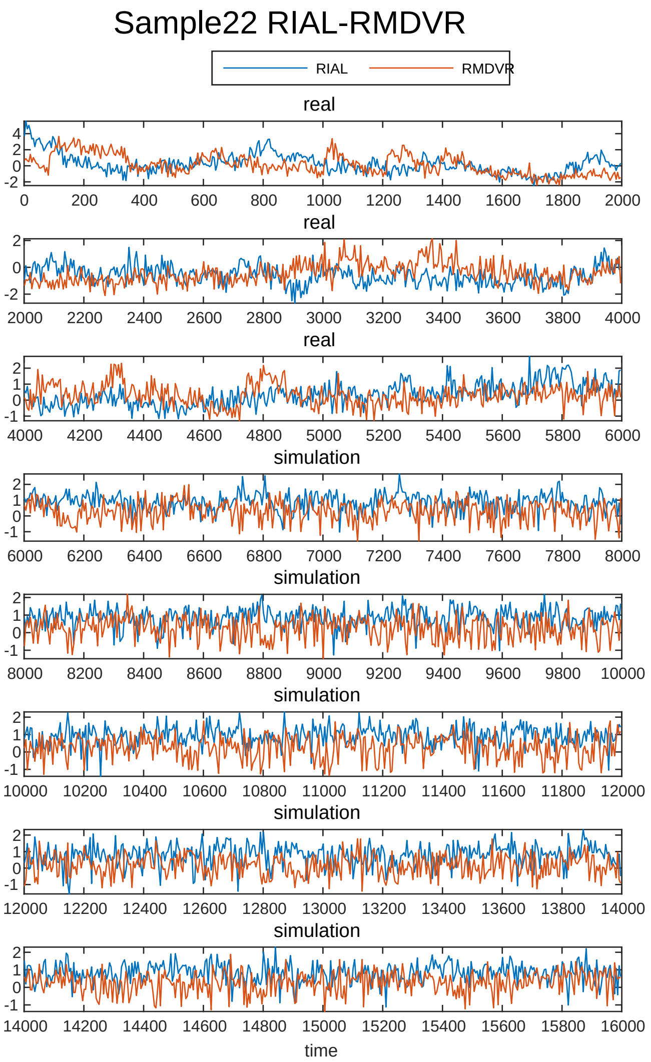
Sample22 RIAL-RMDVR
RIAL
RMDVR
real
real
real
simulation
simulation
simulation
simulation
simulation
time

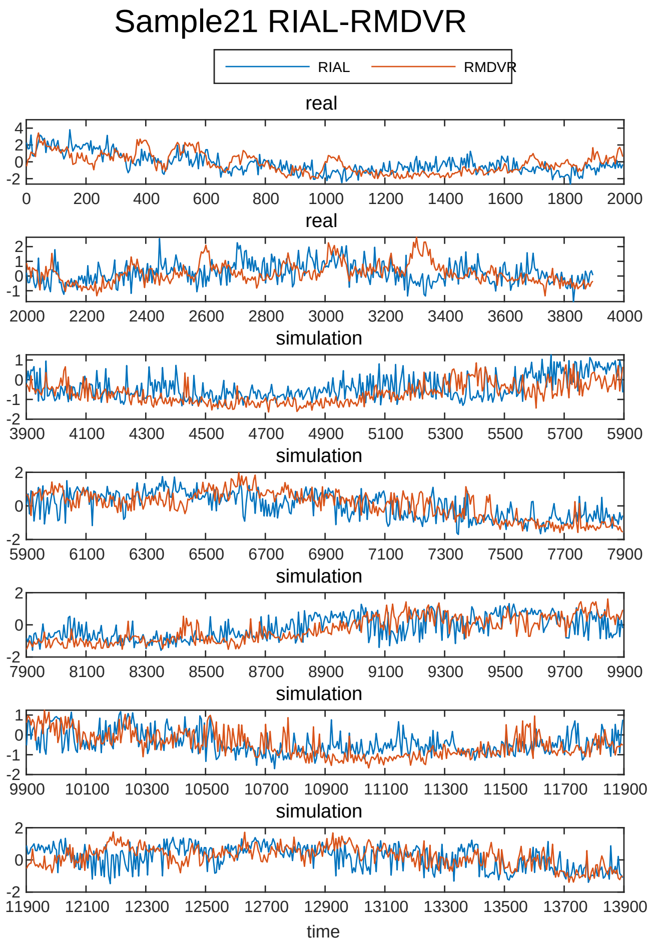
Sample21 RIAL-RMDVR
RIAL
RMDVR
real
real
simulation
simulation
simulation
simulation
simulation
time

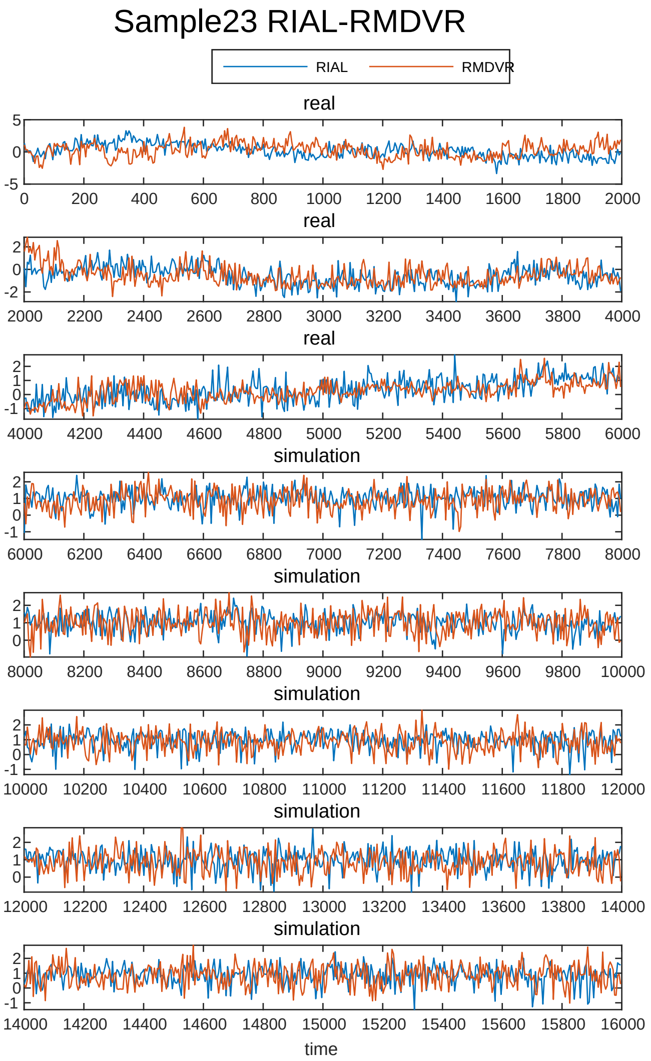
Sample23 RIAL-RMDVR
RIAL
RMDVR
real
real
real
simulation
simulation
simulation
simulation
simulation
time
