## Supplementary File 7 for "Deducing ensemble dynamics and information flow from the whole-brain imaging data"

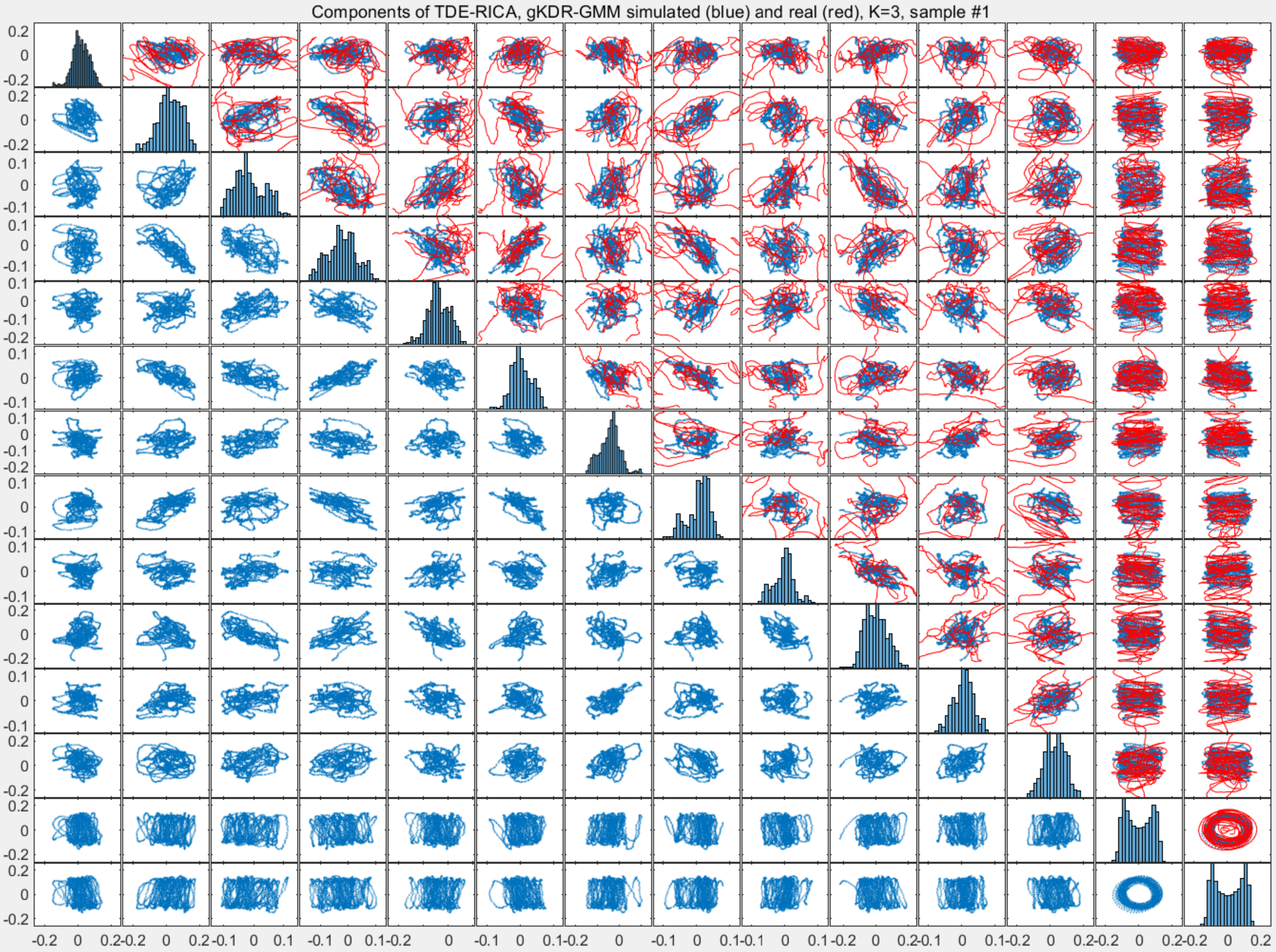
Components of TDE-RICA, gKDR-GMM simulated (blue) and real (red), K=3, sample #1

Components of TDE-RICA, gKDR-GMM simulated (blue) and real (red), K=4, sample #1

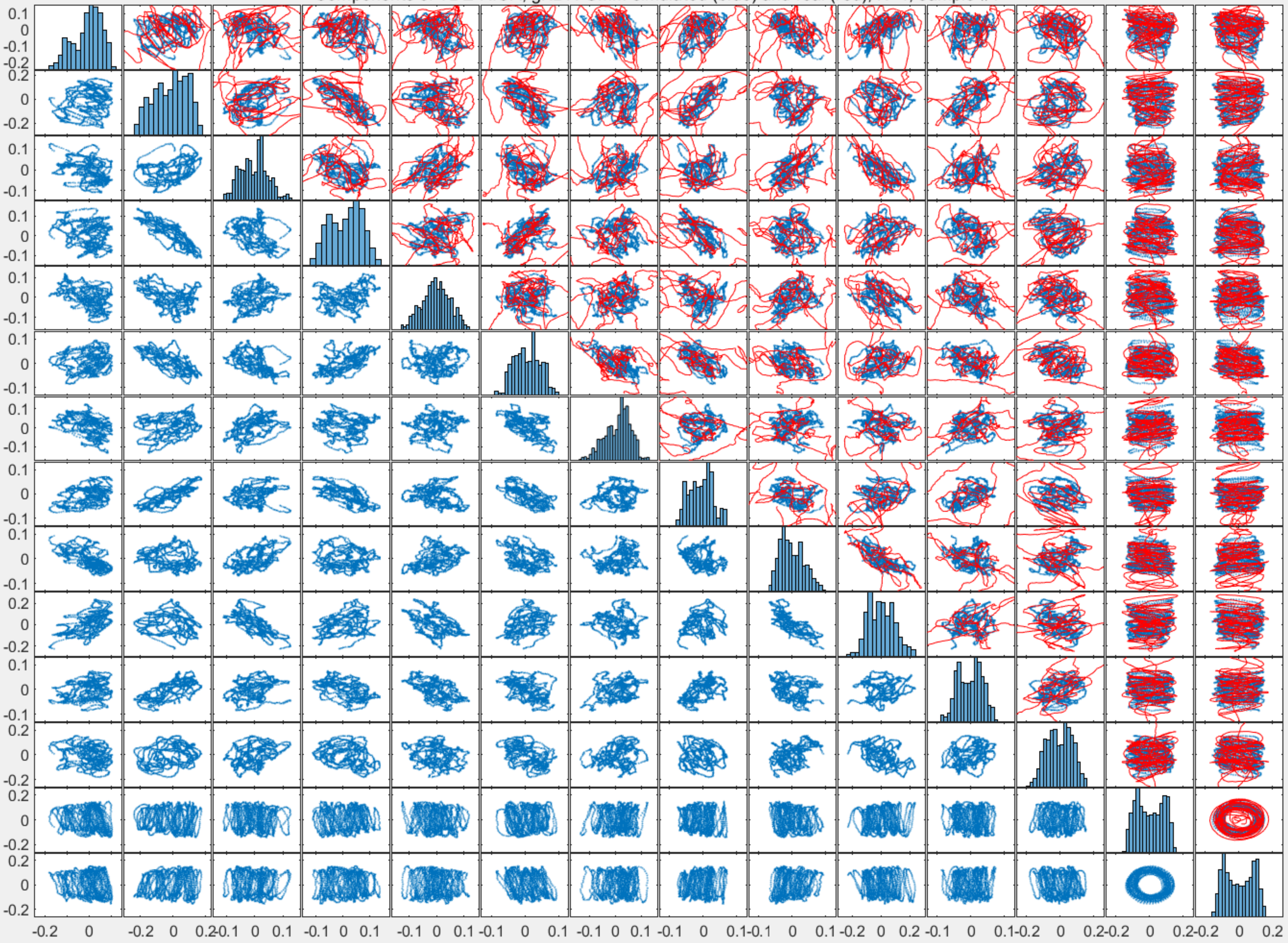

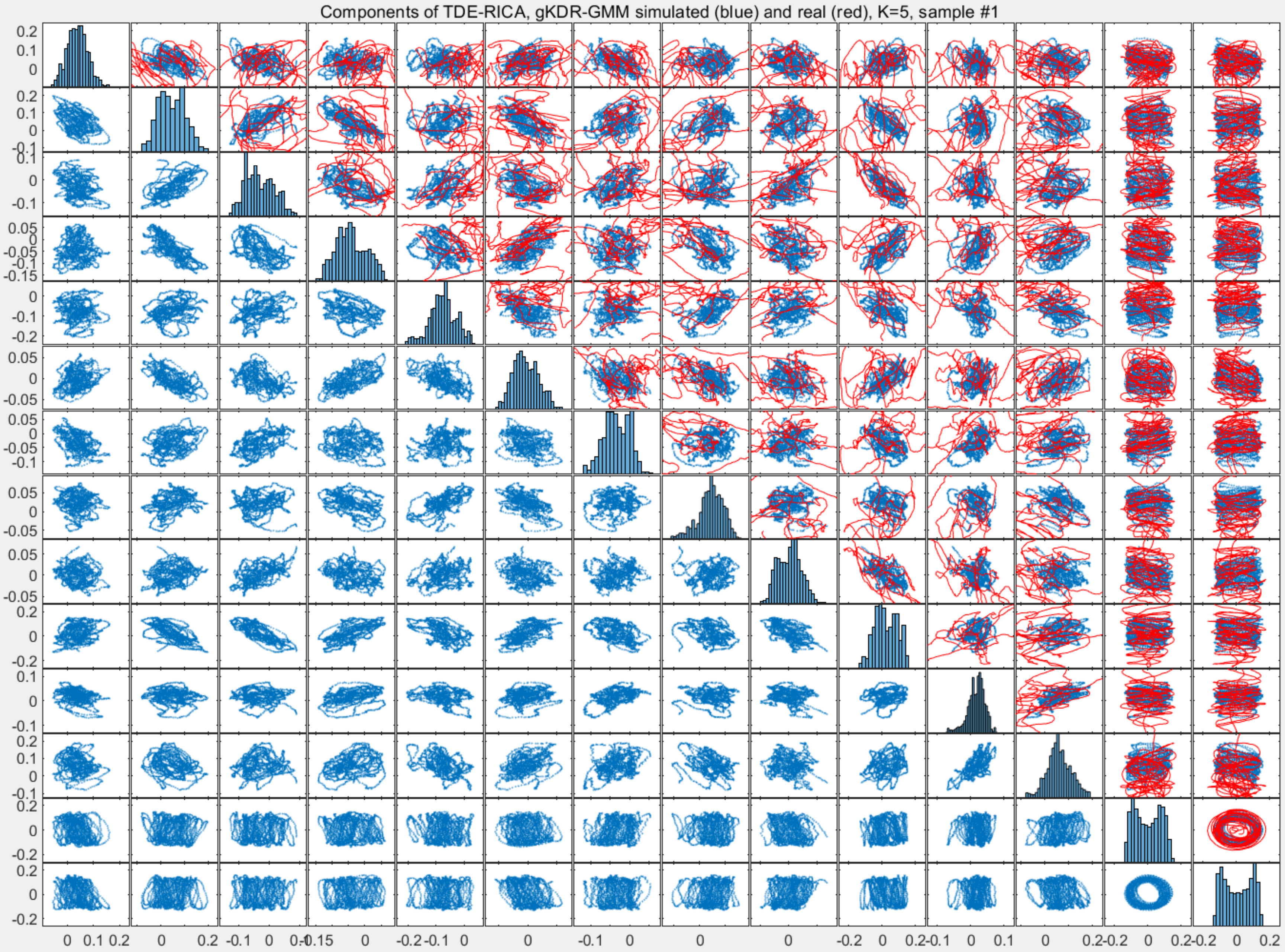
Components of TDE-RICA, gKDR-GMM simulated (blue) and real (red), K=5, sample #1

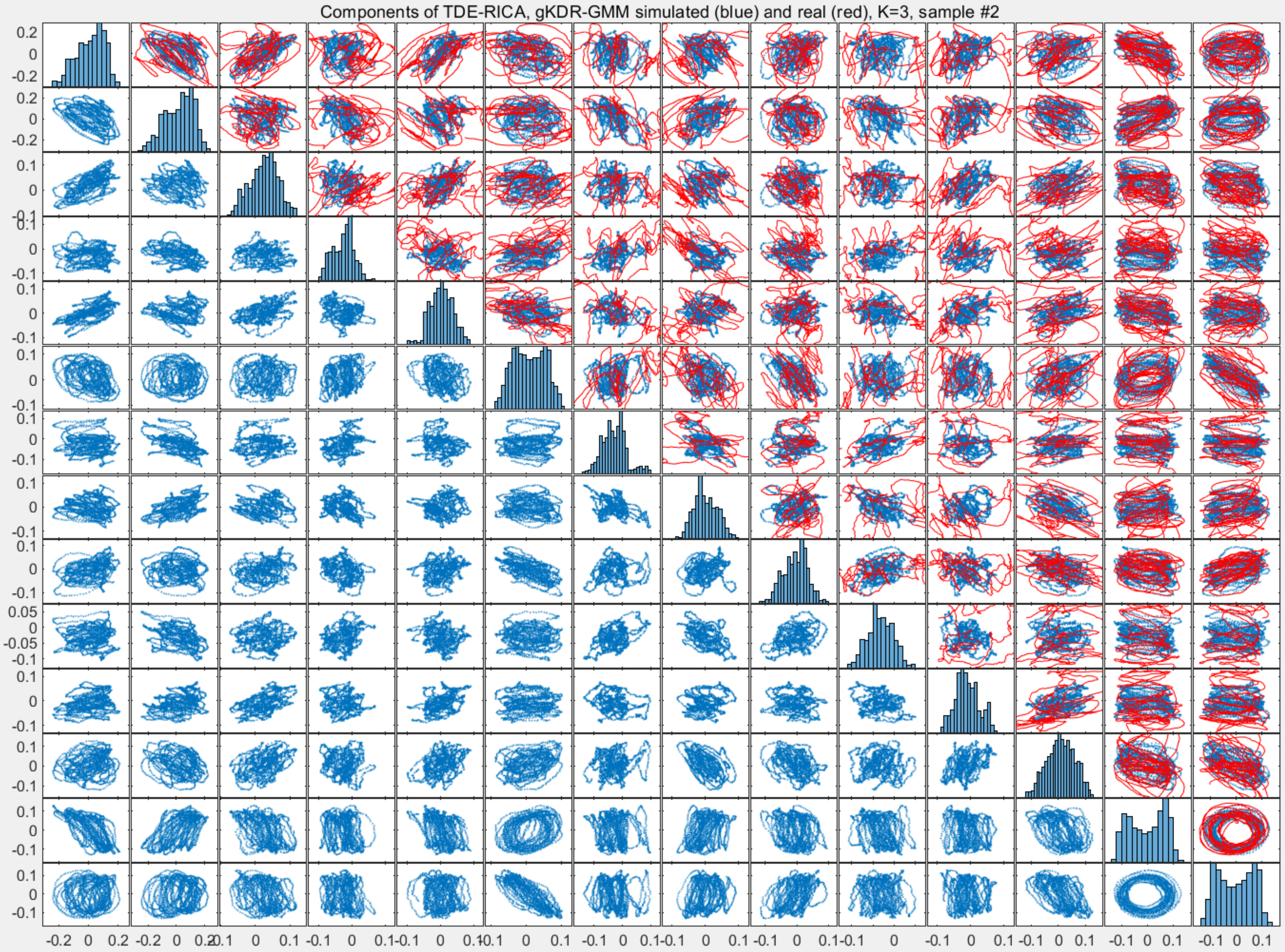
Components of TDE-RICA, gKDR-GMM simulated (blue) and real (red), K=3, sample #2

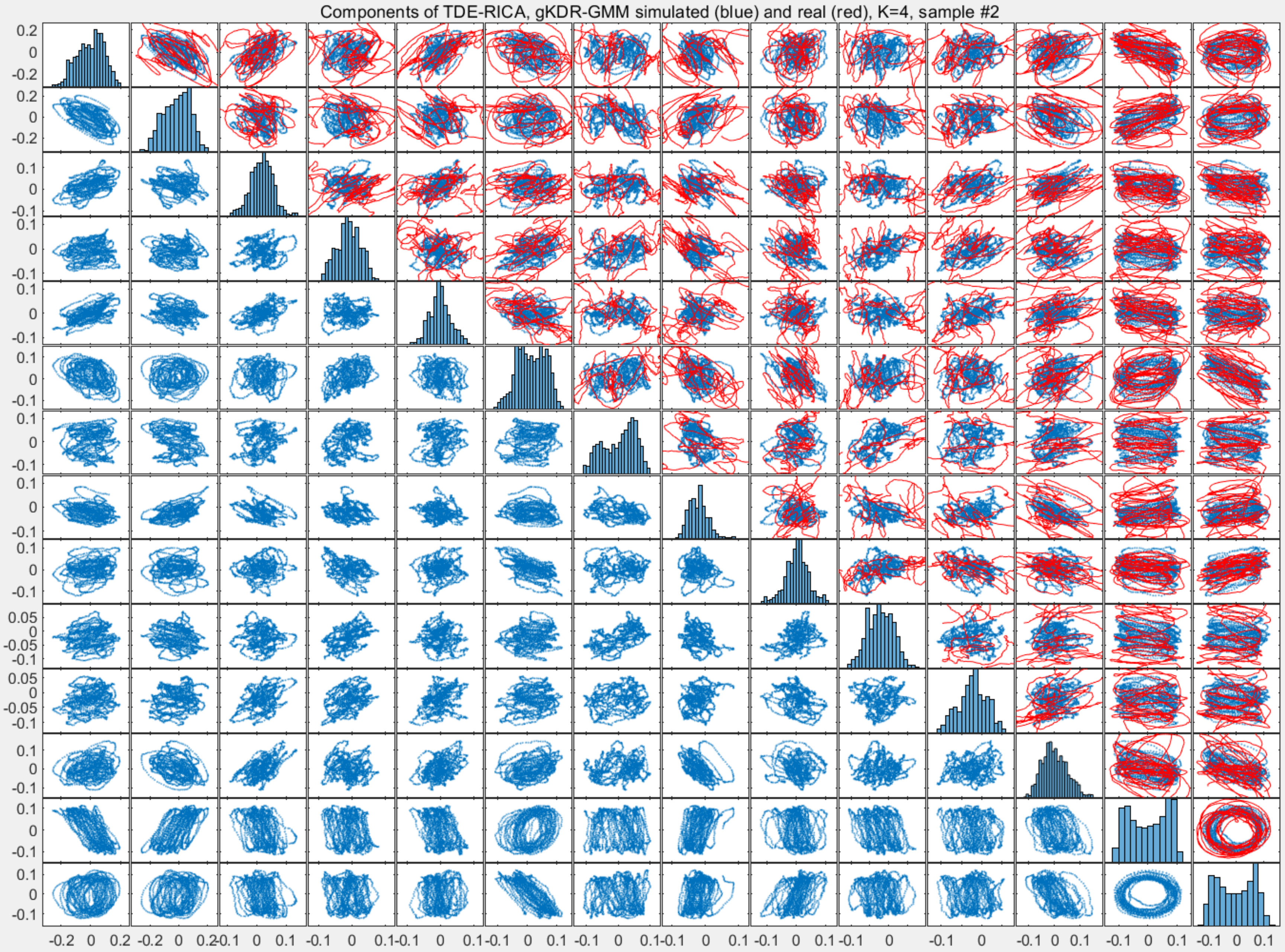
Components of TDE-RICA, gKDR-GMM simulated (blue) and real (red), K=4, sample #2

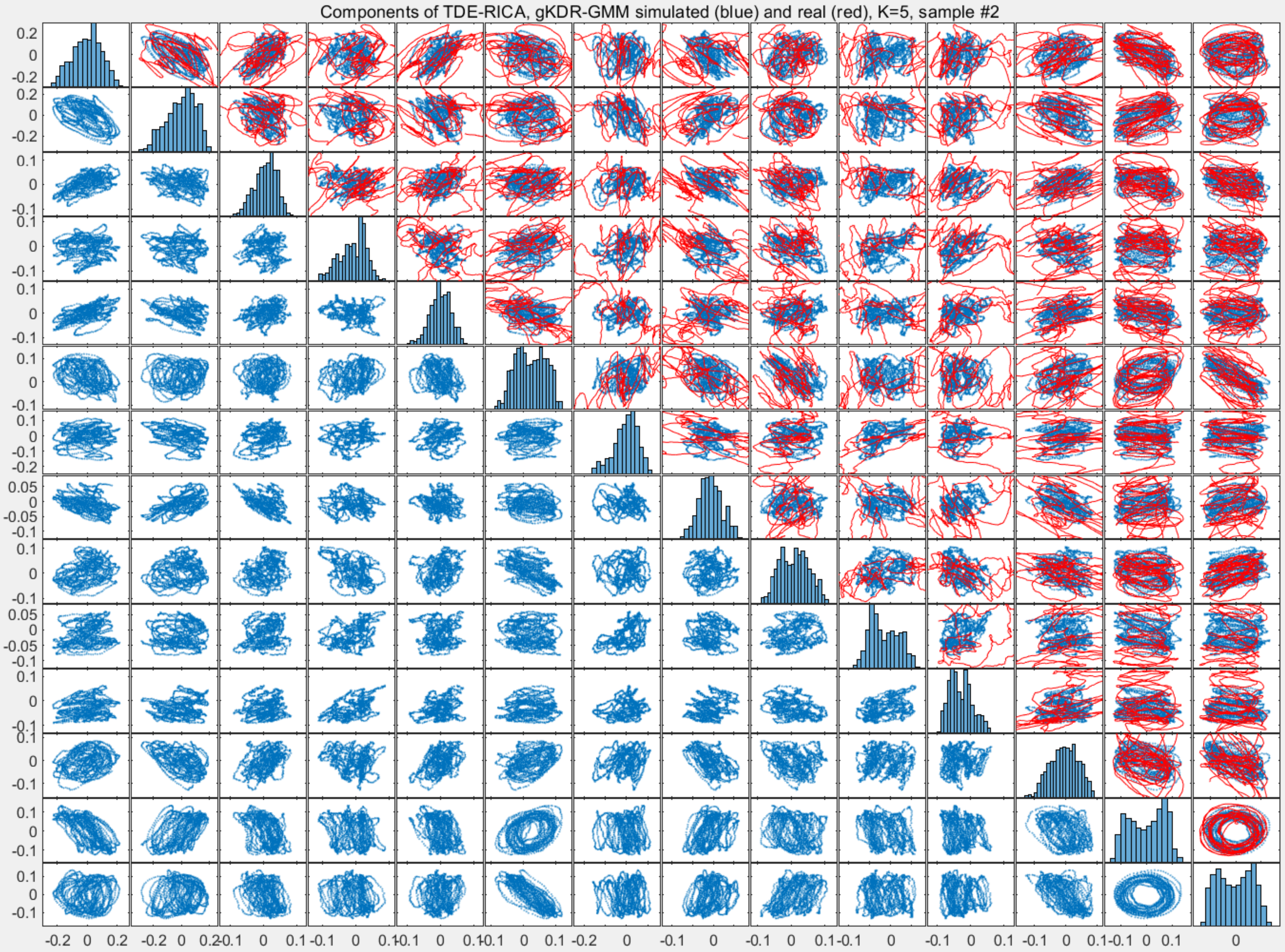

Components of TDE-RICA, gKDR-GMM simulated (blue) and real (red), K=5, sample #2

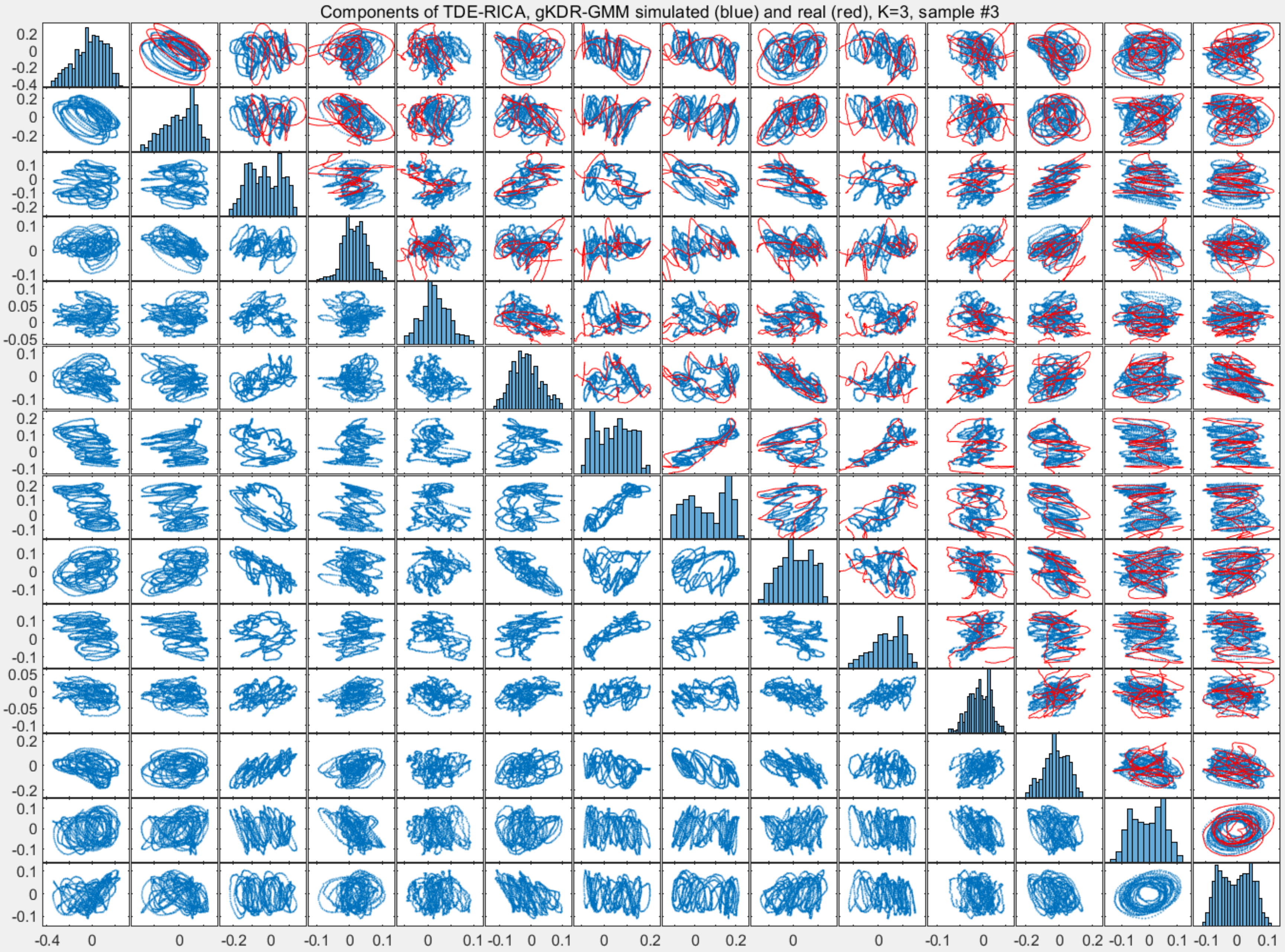
Components of TDE-RICA, gKDR-GMM simulated (blue) and real (red), K=3, sample #3

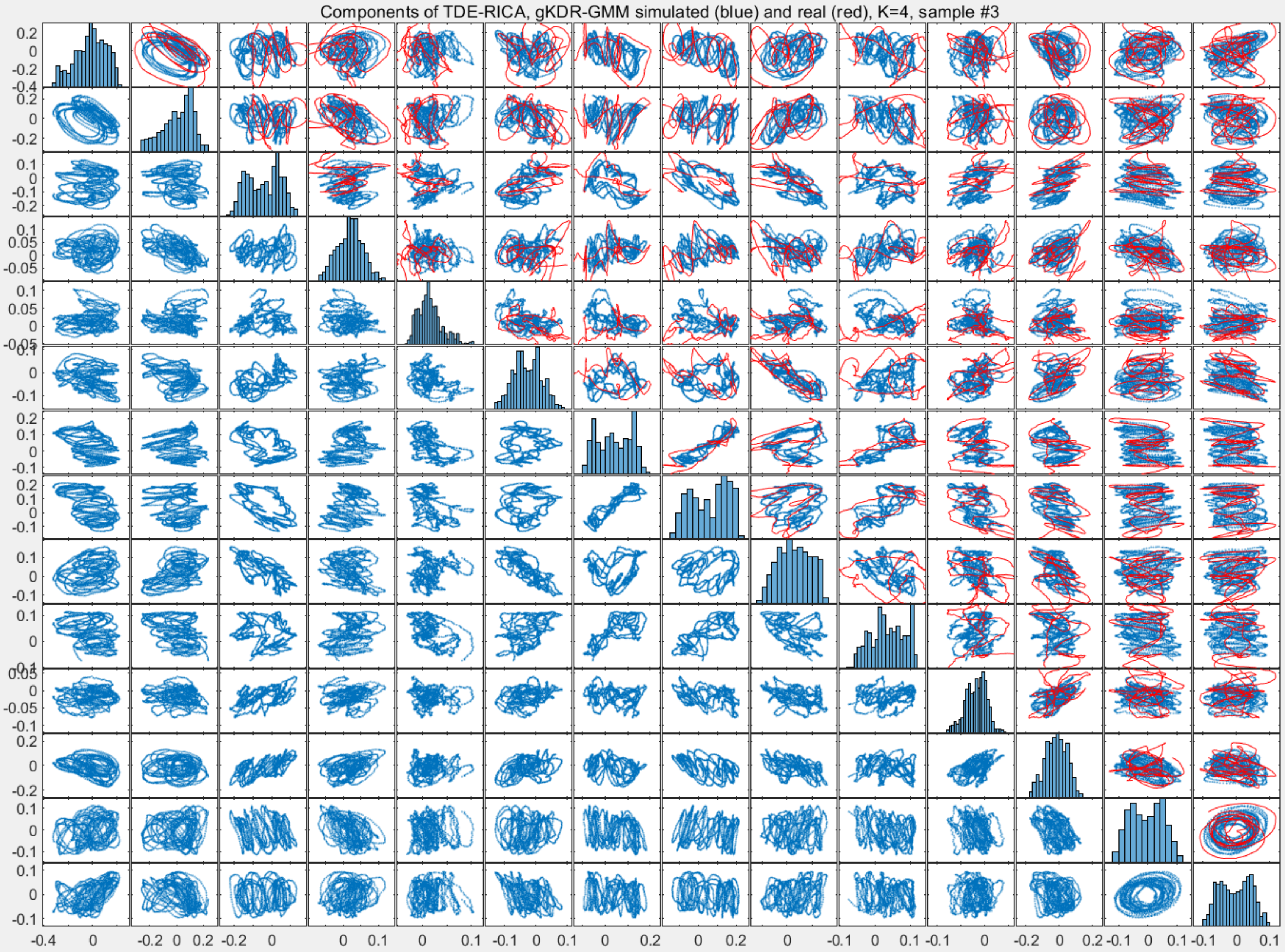
Components of TDE-RICA, gKDR-GMM simulated (blue) and real (red), K=4, sample #3

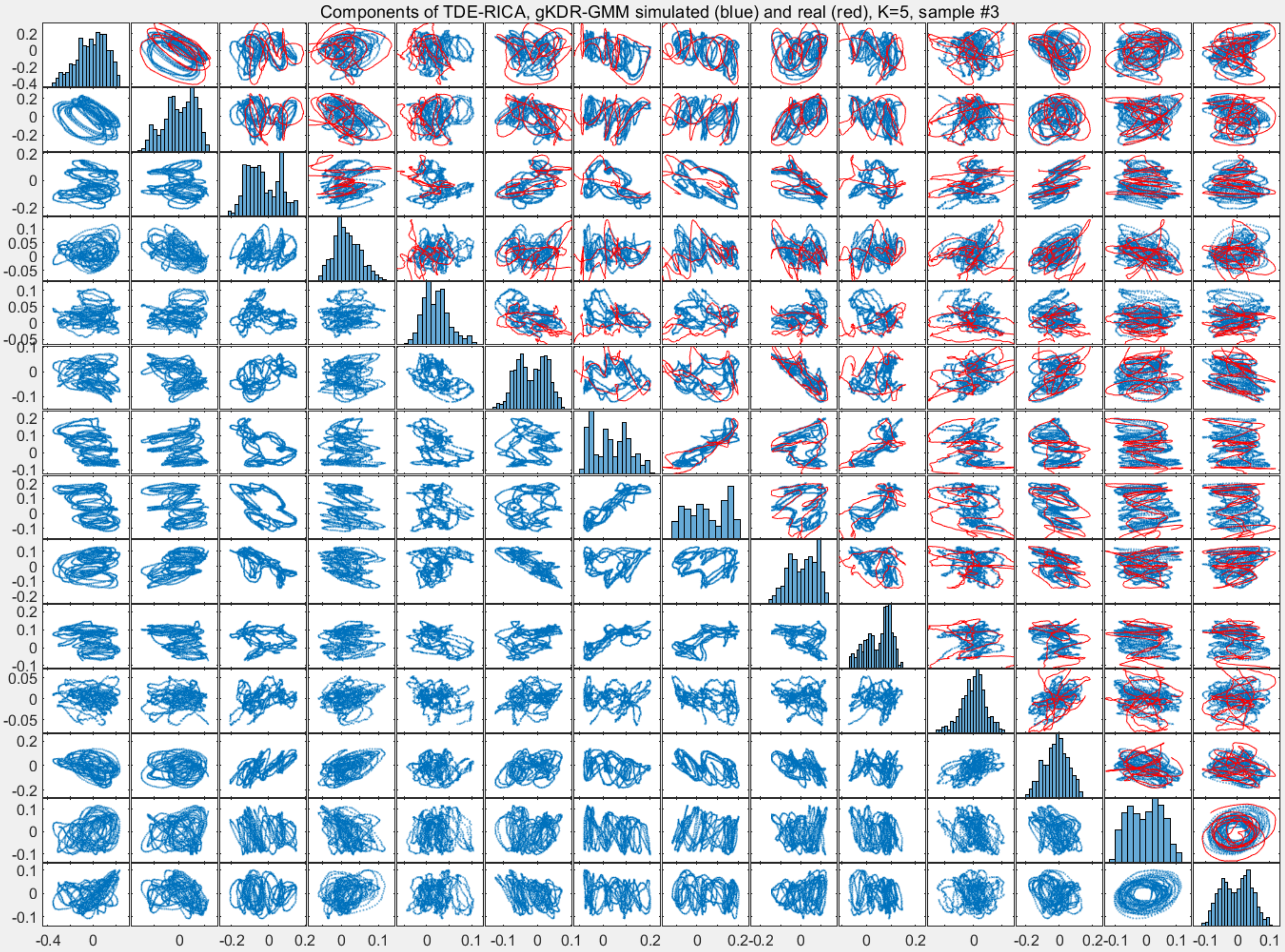

Components of TDE-RICA, gKDR-GMM simulated (blue) and real (red), K=5, sample #3

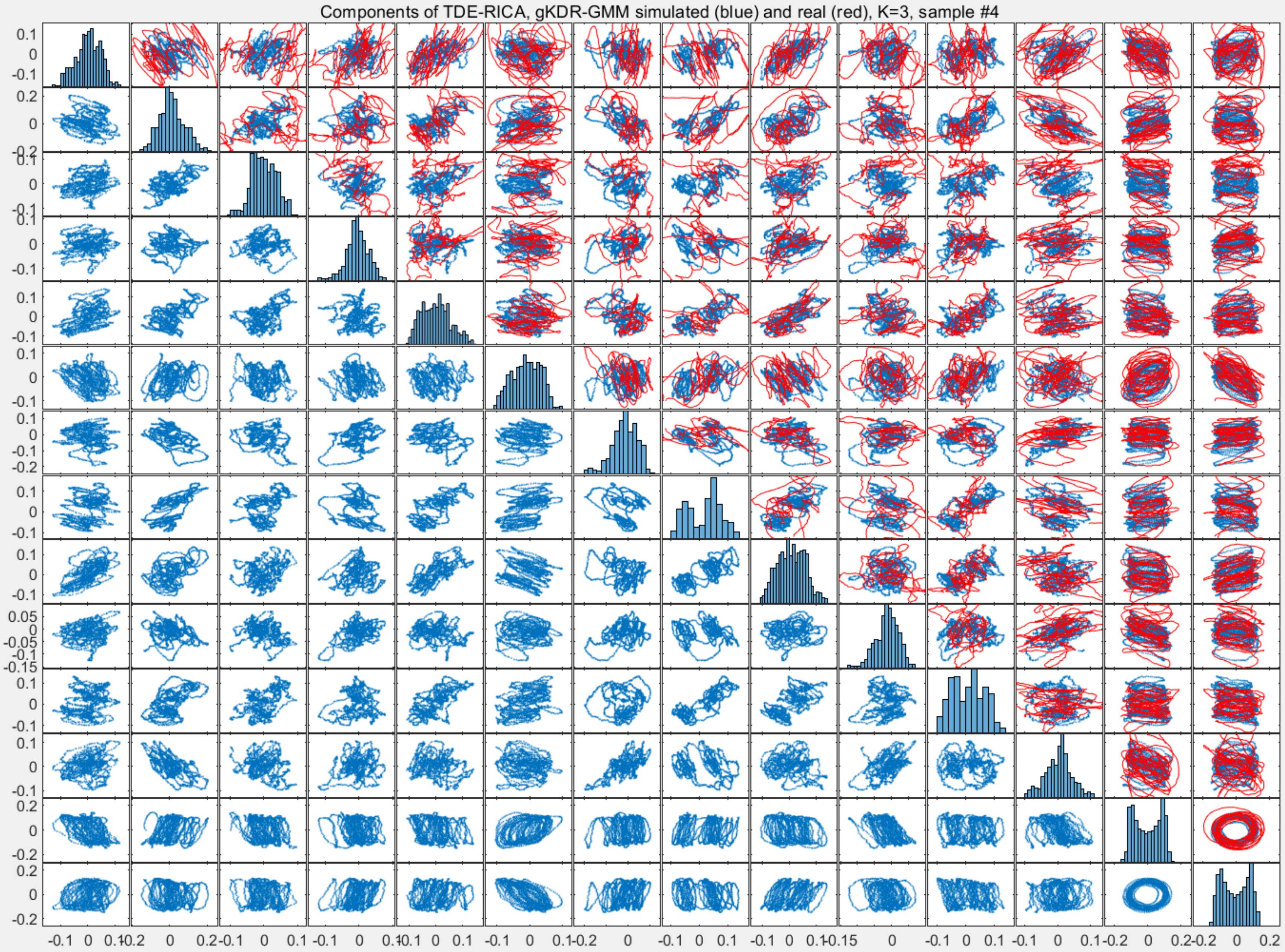

Components of TDE-RICA, gKDR-GMM simulated (blue) and real (red), K=3, sample #4

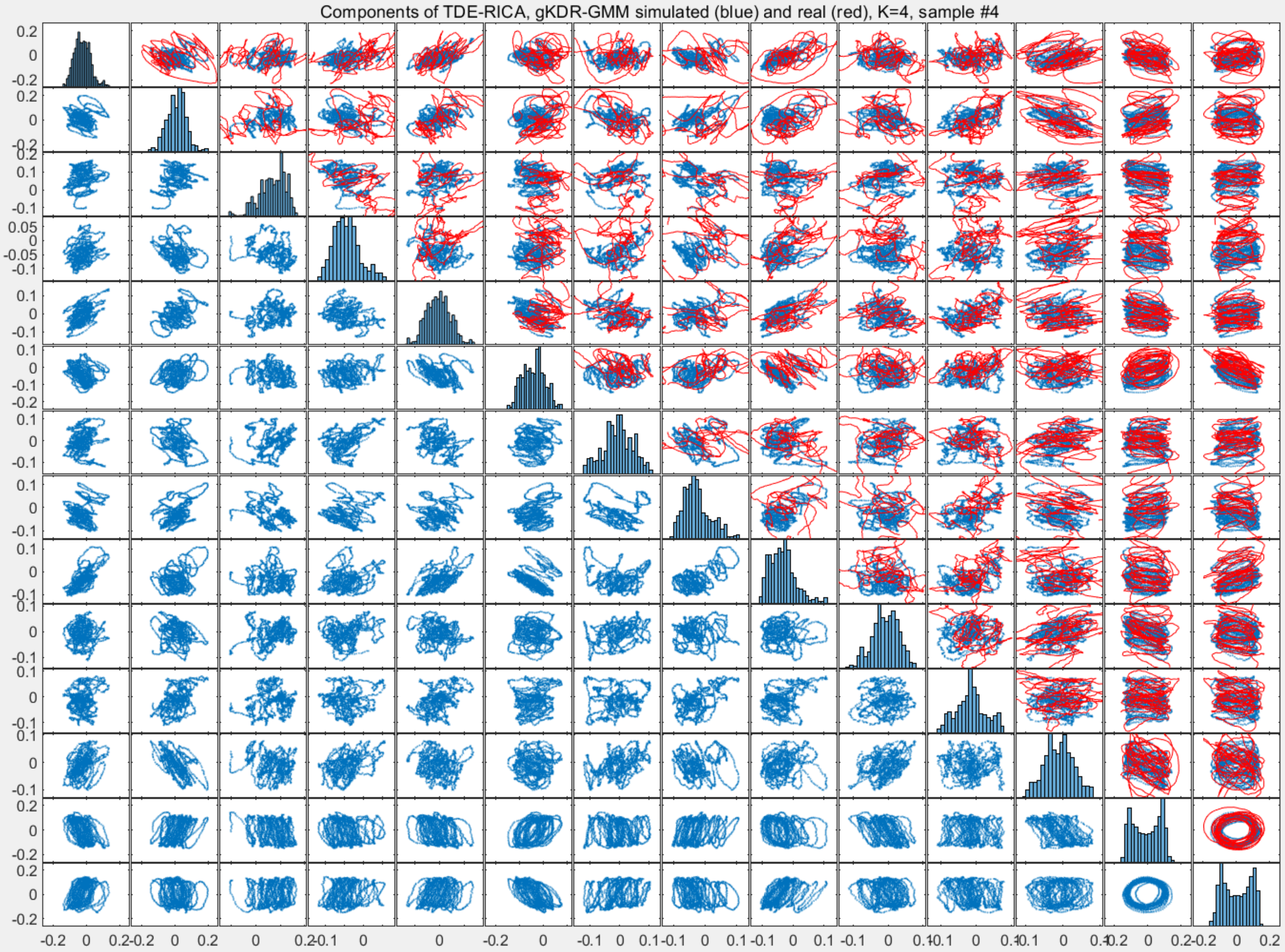
Components of TDE-RICA, gKDR-GMM simulated (blue) and real (red), K=4, sample #4

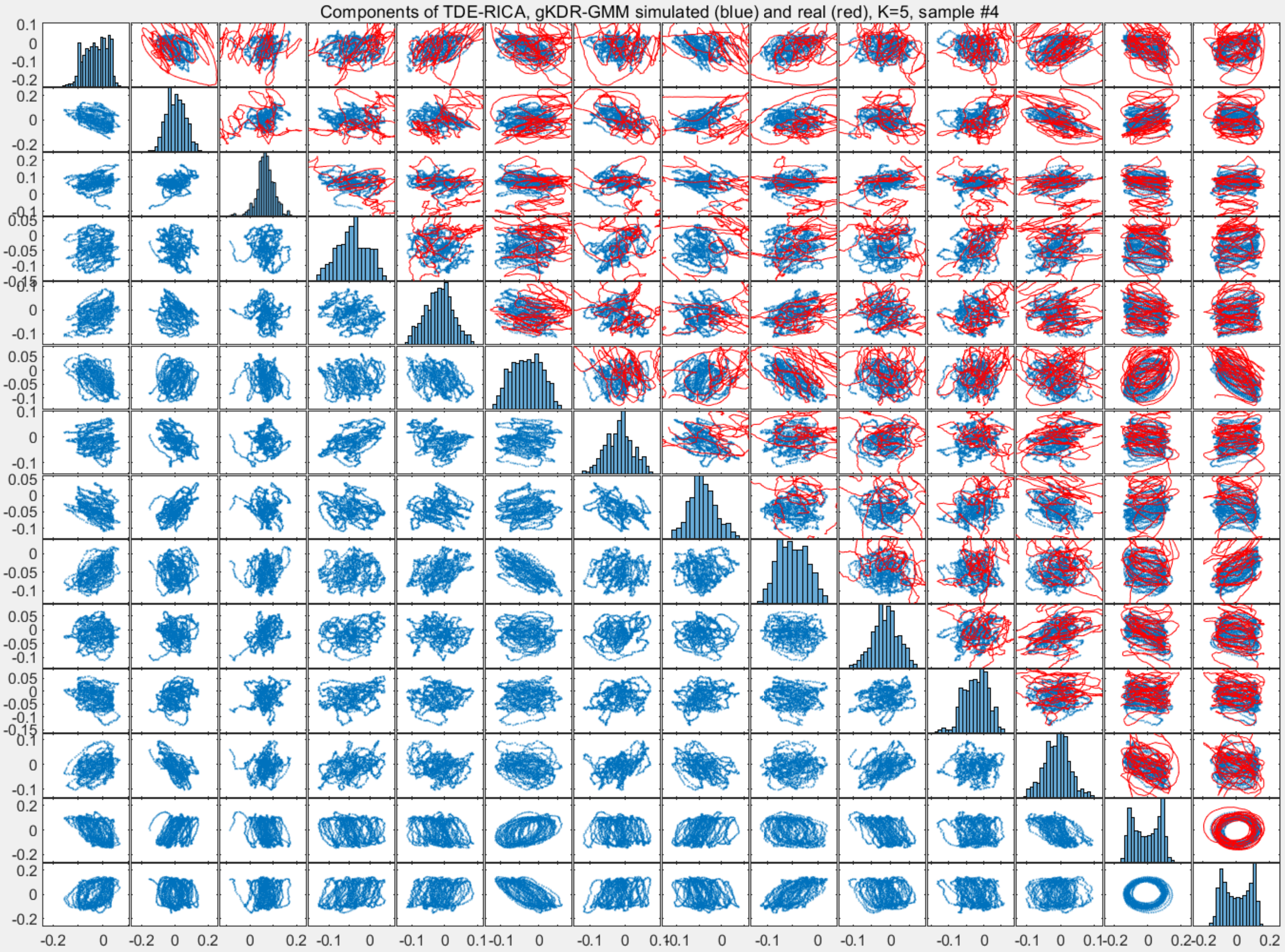

Components of TDE-RICA, gKDR-GMM simulated (blue) and real (red), K=5, sample #4

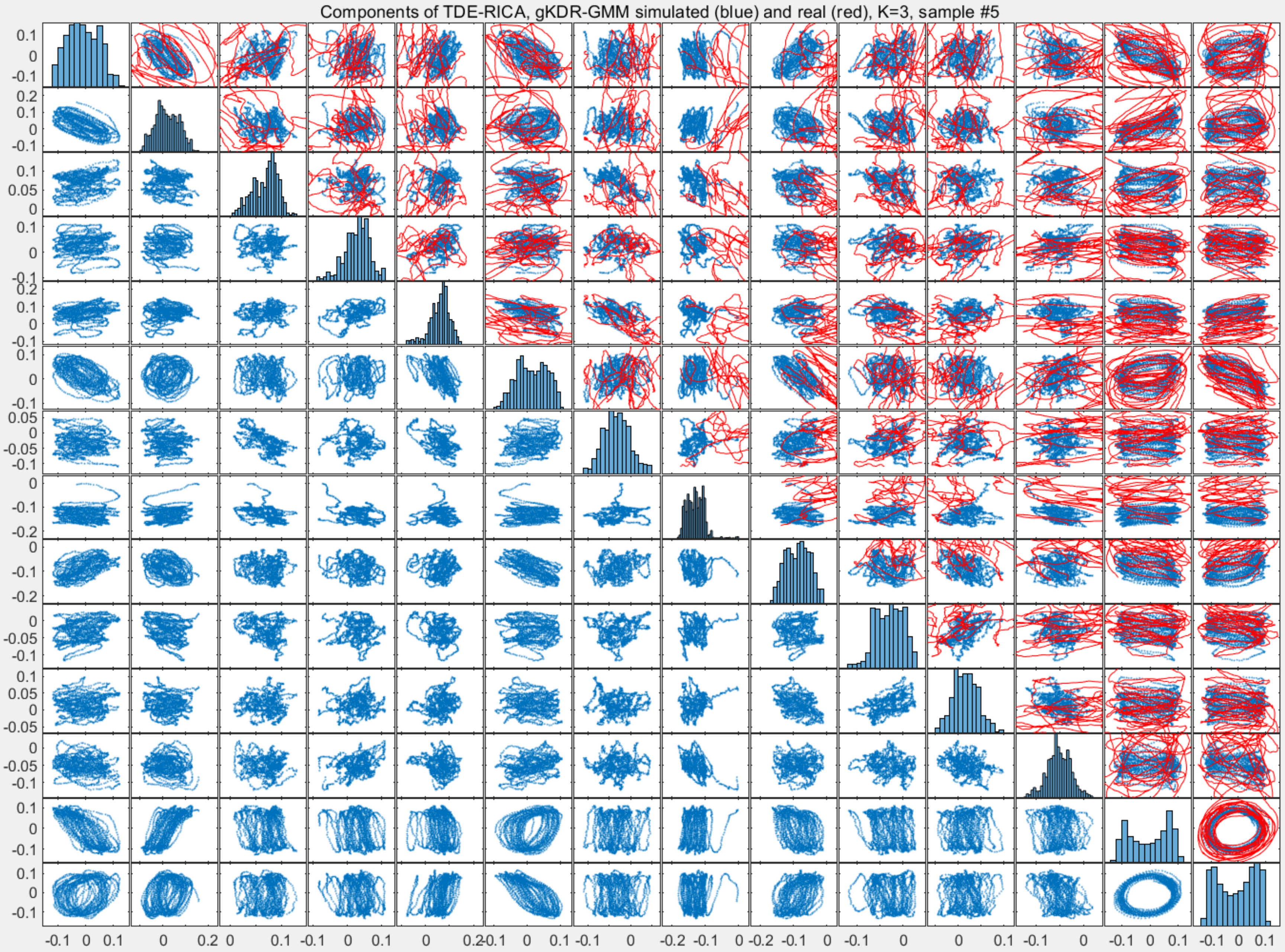
Components of TDE-RICA, gKDR-GMM simulated (blue) and real (red), K=3, sample #5
0.1
0
-0.1
0.2
0.05
-0.05
-0.2
0.2-0.1

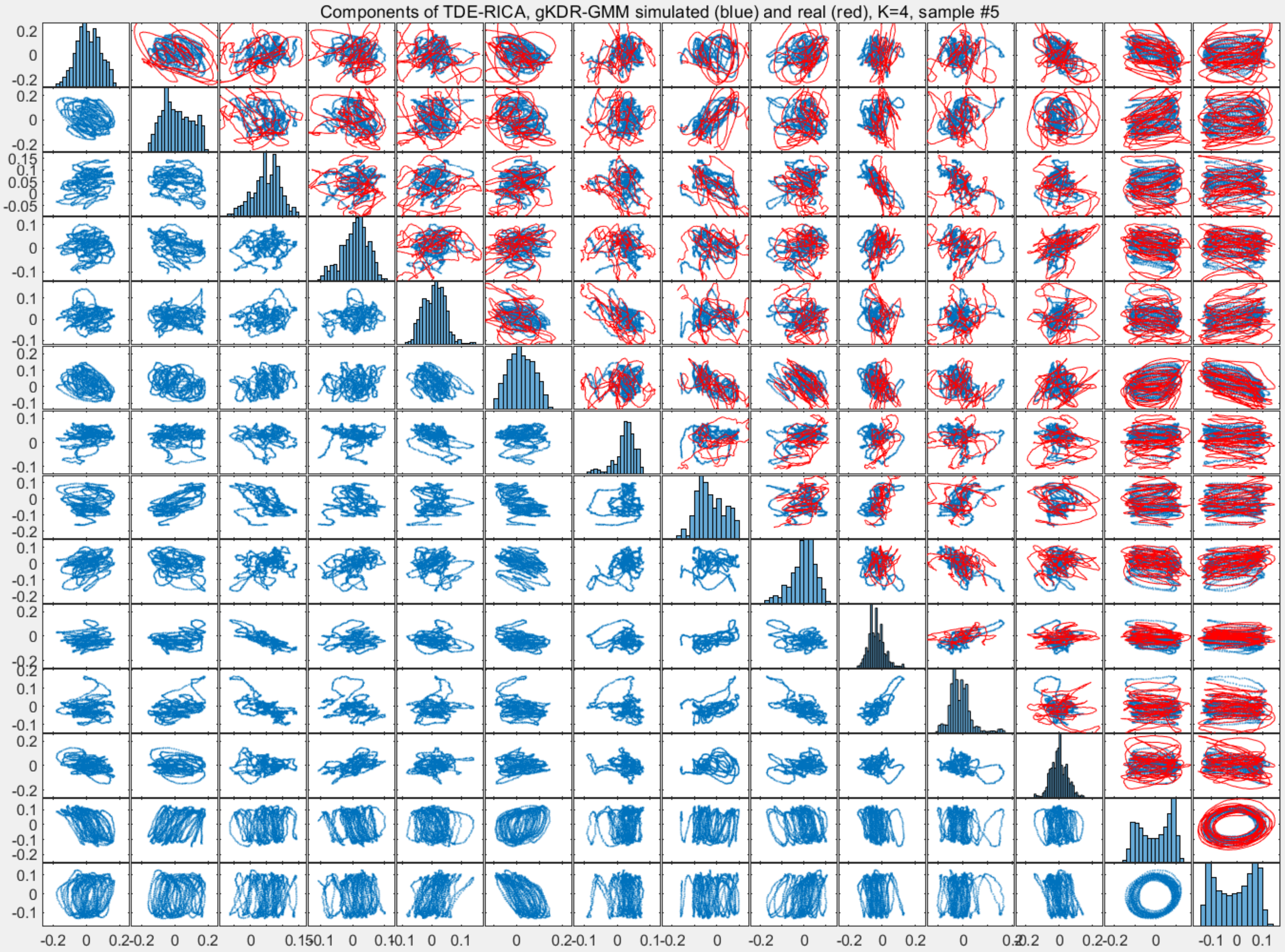
Components of TDE-RICA, gKDR-GMM simulated (blue) and real (red), K=4, sample #5

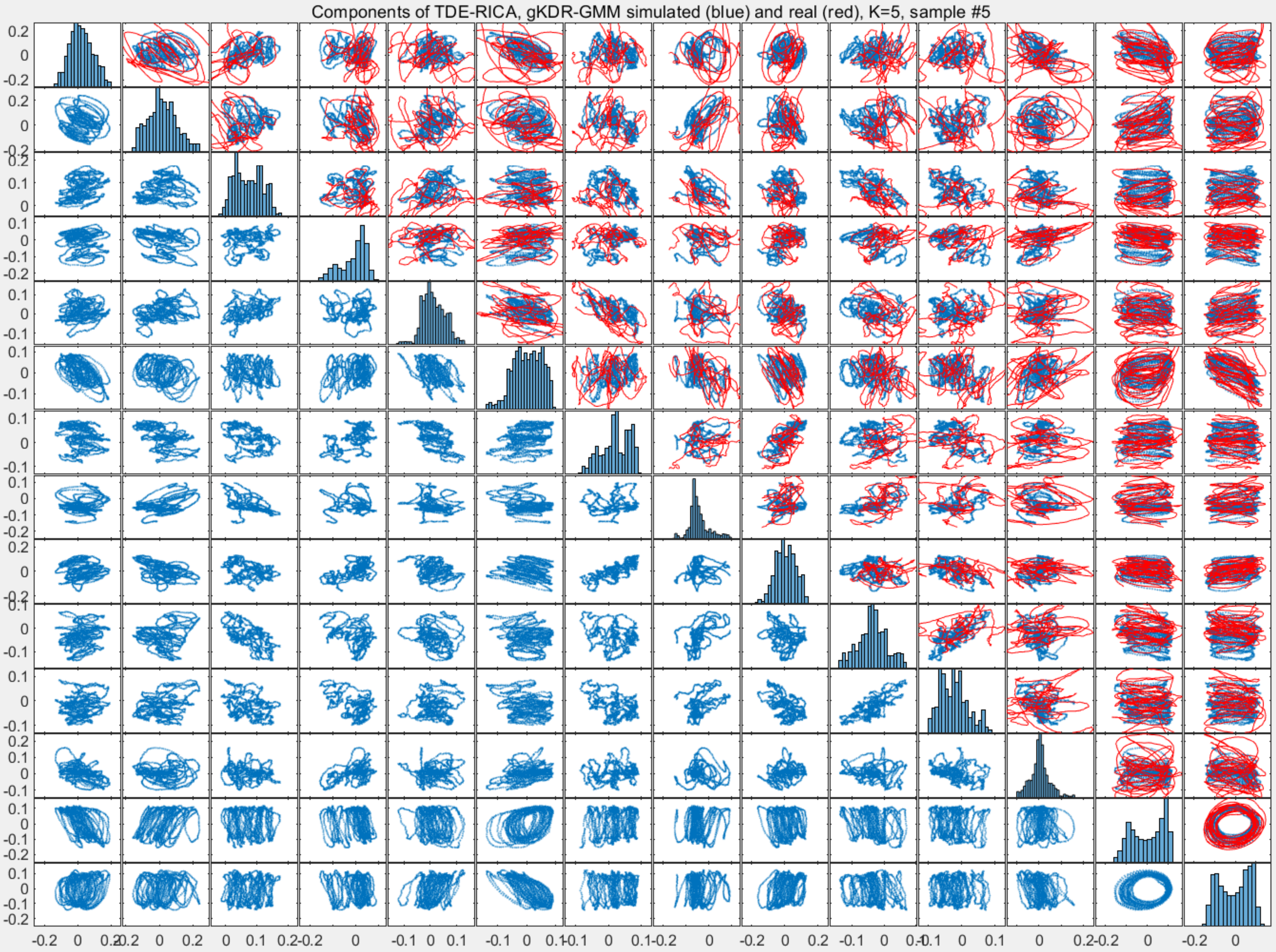
Components of TDE-RICA, gKDR-GMM simulated (blue) and real (red), K=5, sample #5

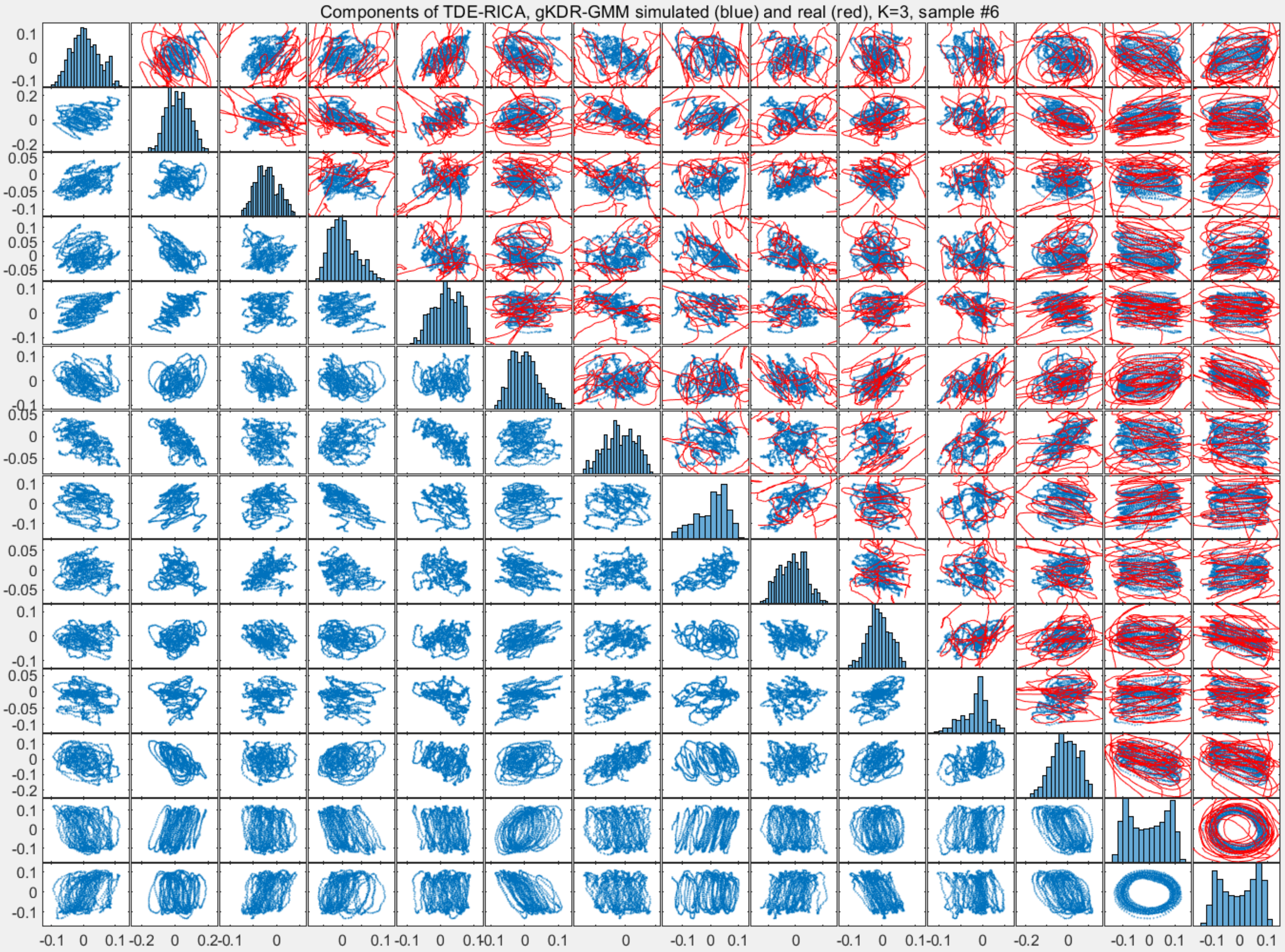

Components of TDE-RICA, gKDR-GMM simulated (blue) and real (red), K=3, sample #6

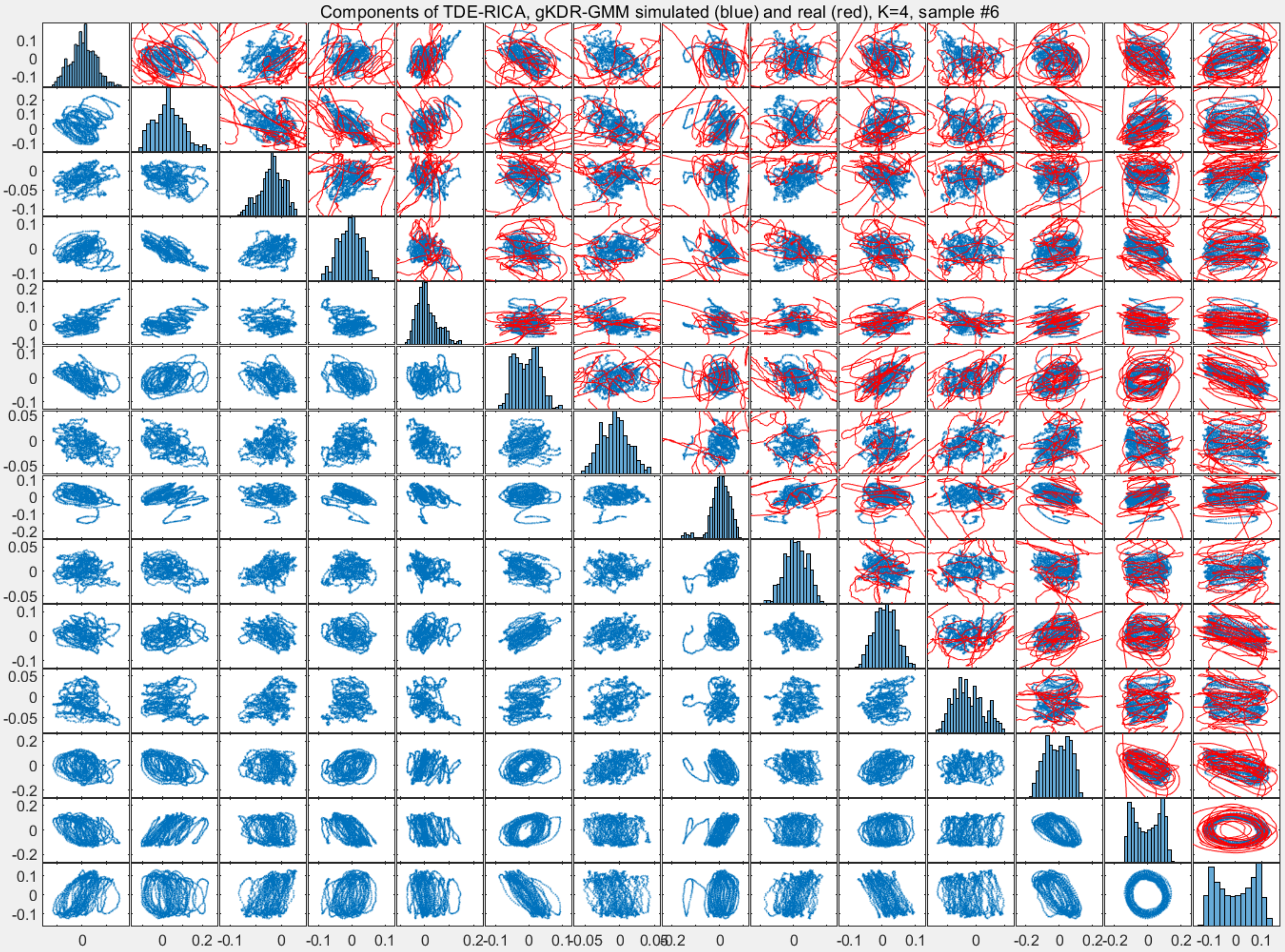

Components of TDE-RICA, gKDR-GMM simulated (blue) and real (red), K=4, sample #6

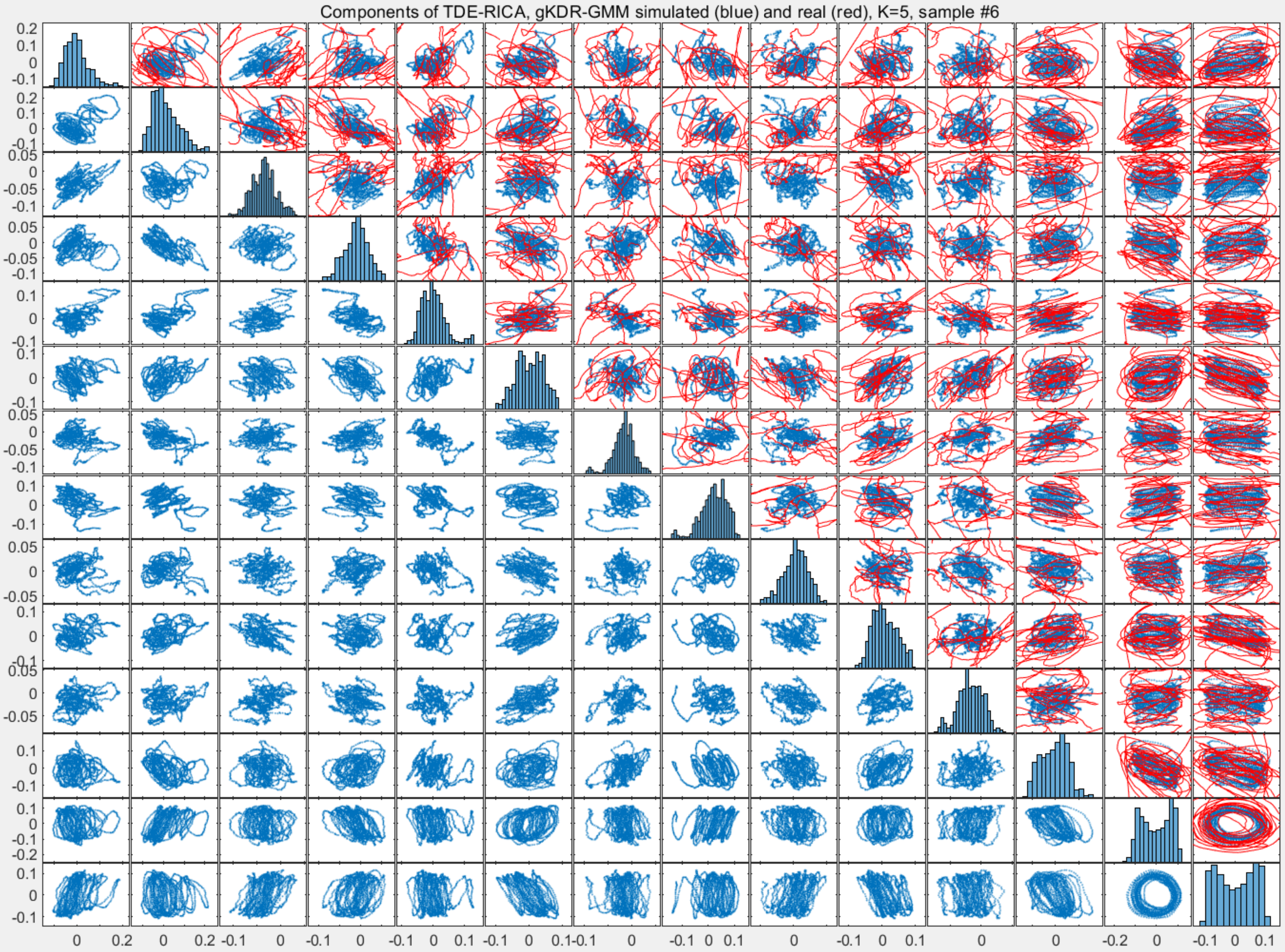
Components of TDE-RICA, gKDR-GMM simulated (blue) and real (red), K=5, sample #6

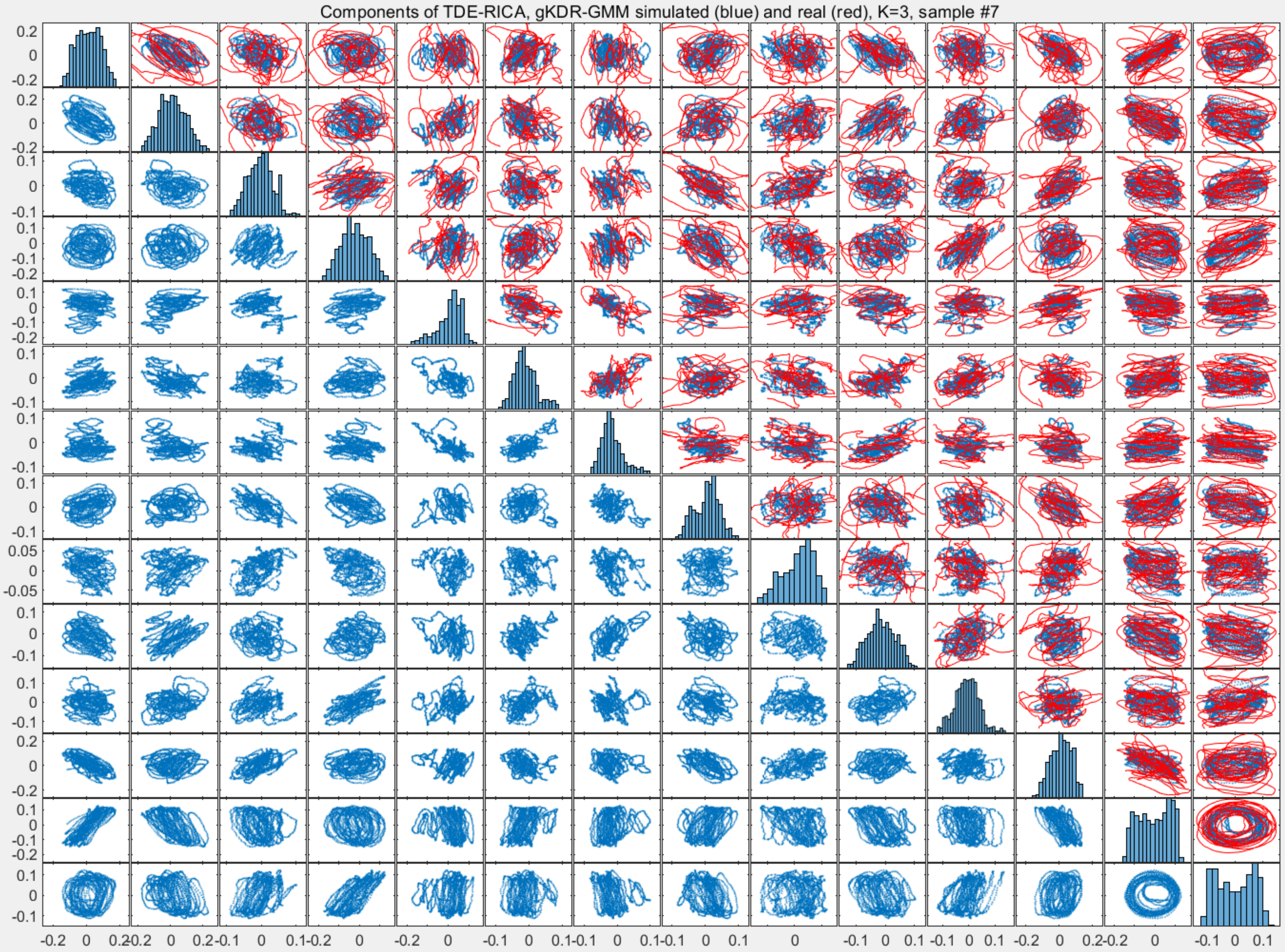
Components of TDE-RICA, gKDR-GMM simulated (blue) and real (red), K=3, sample #7

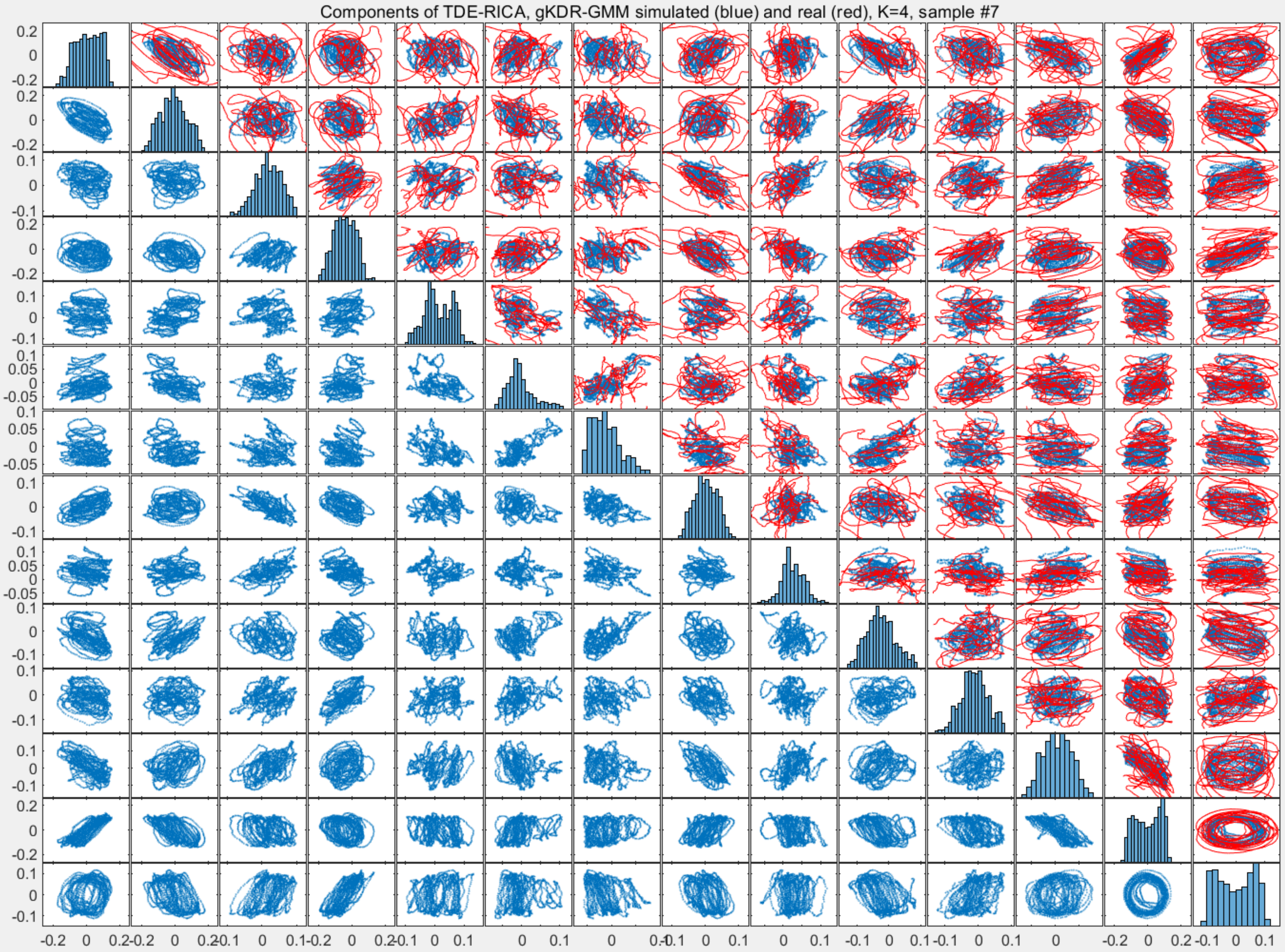

Components of TDE-RICA, gKDR-GMM simulated (blue) and real (red), K=4, sample #7

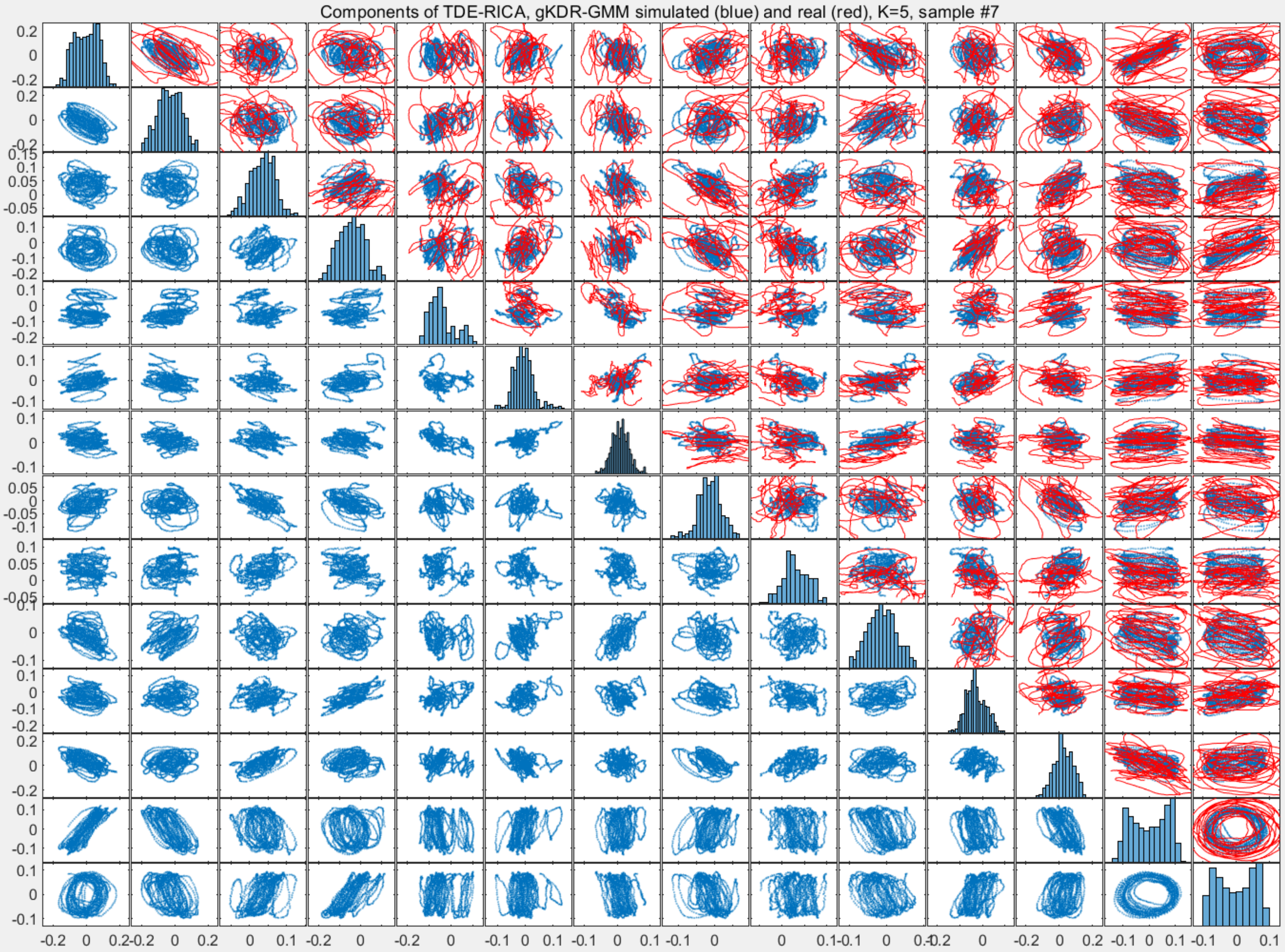
Components of TDE-RICA, gKDR-GMM simulated (blue) and real (red), K=5, sample #7

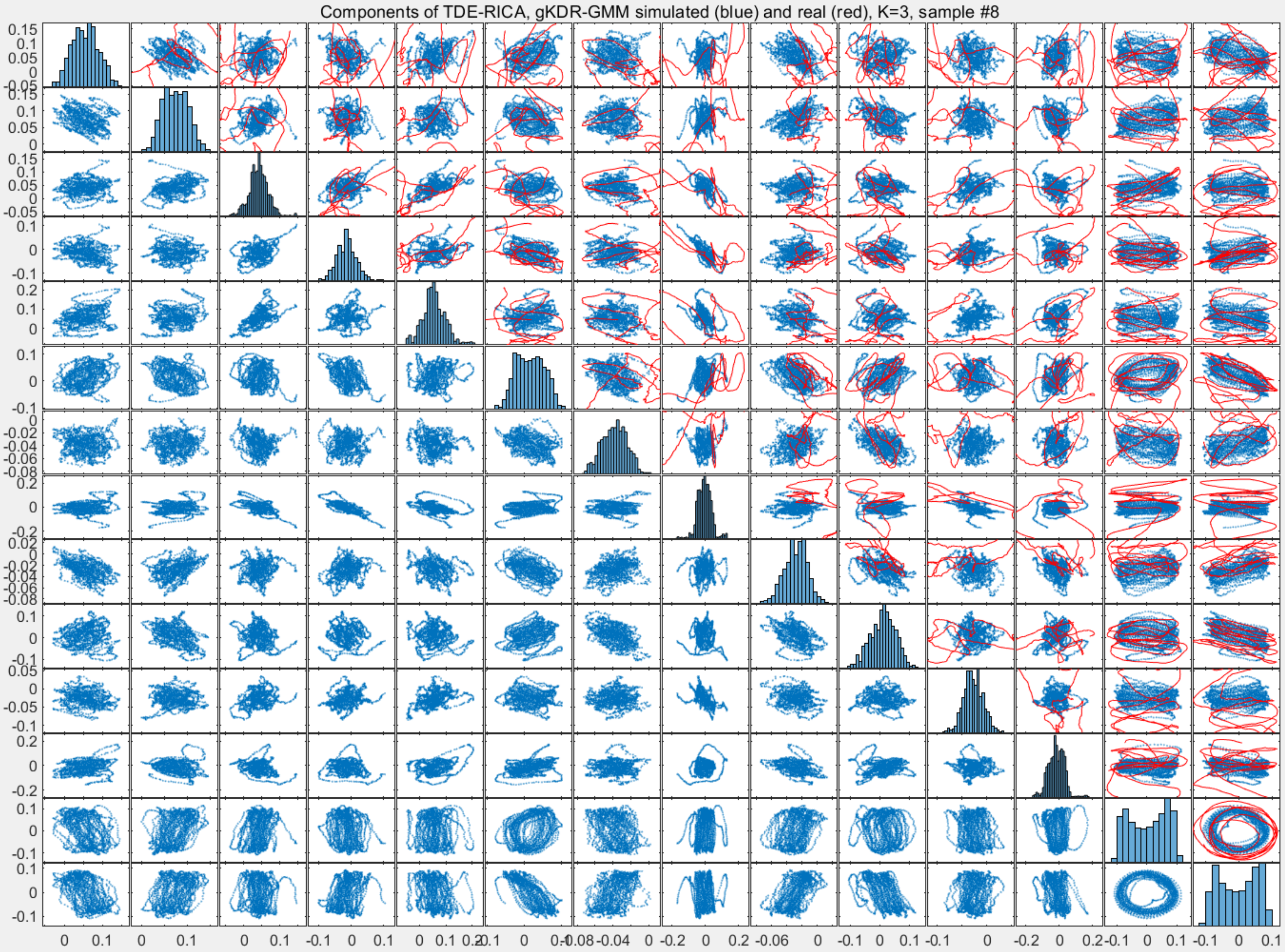

Components of TDE-RICA, gKDR-GMM simulated (blue) and real (red), K=3, sample #8

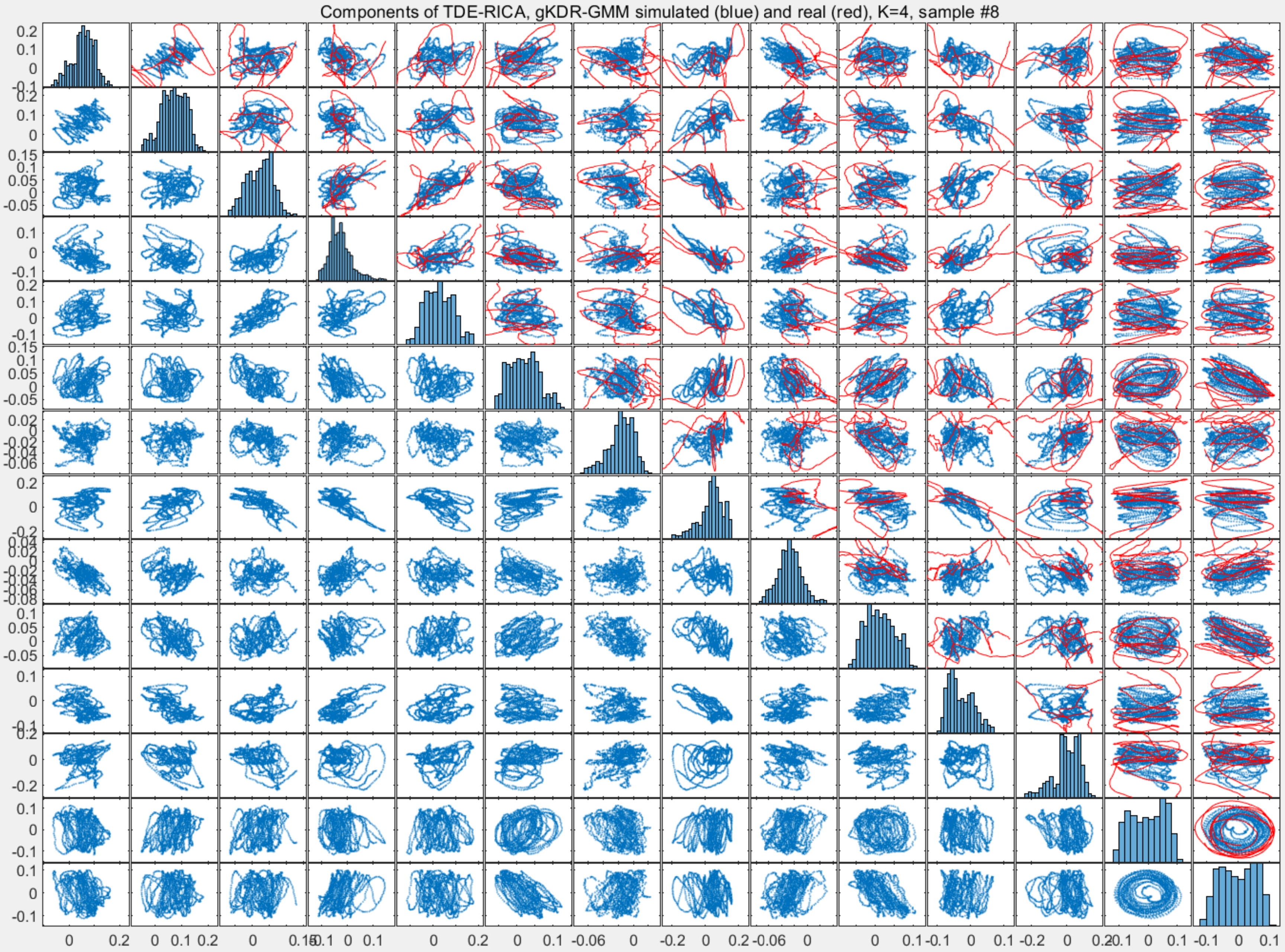

Components of TDE-RICA, gKDR-GMM simulated (blue) and real (red), K=4, sample #8

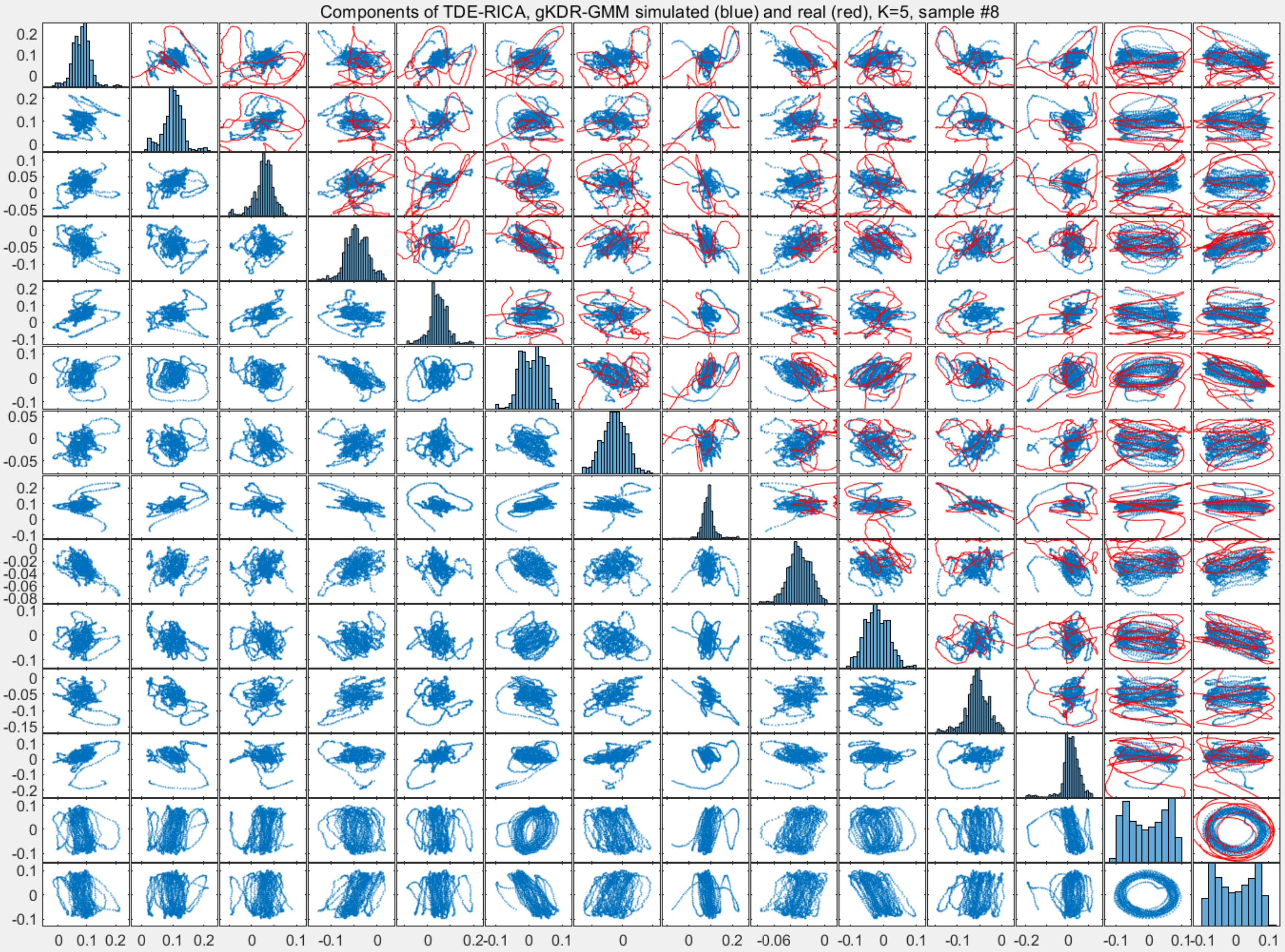
Components of TDE-RICA, gKDR-GMM simulated (blue) and real (red), K=5, sample #8

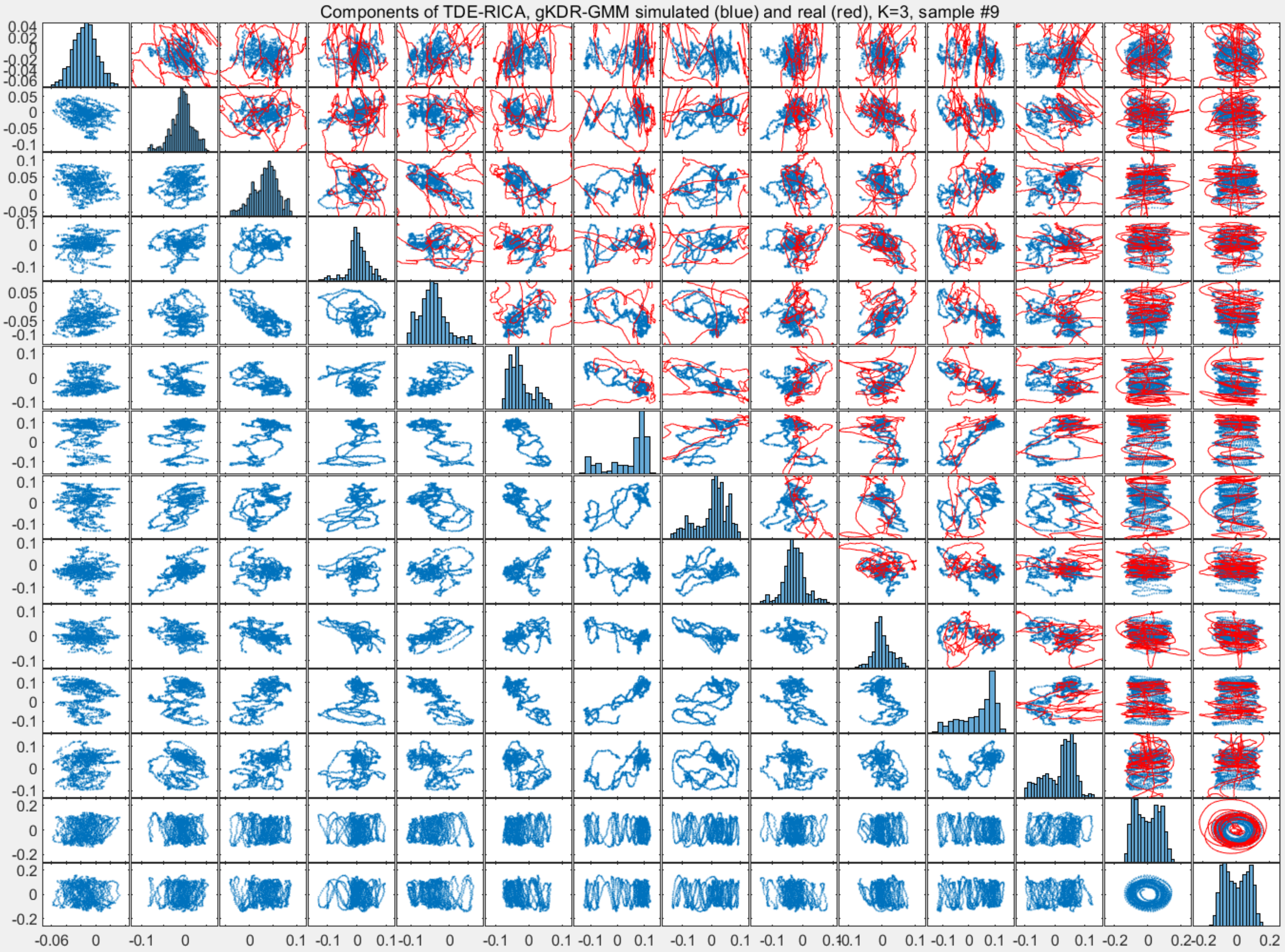
Components of TDE-RICA, gKDR-GMM simulated (blue) and real (red), K=3, sample #9

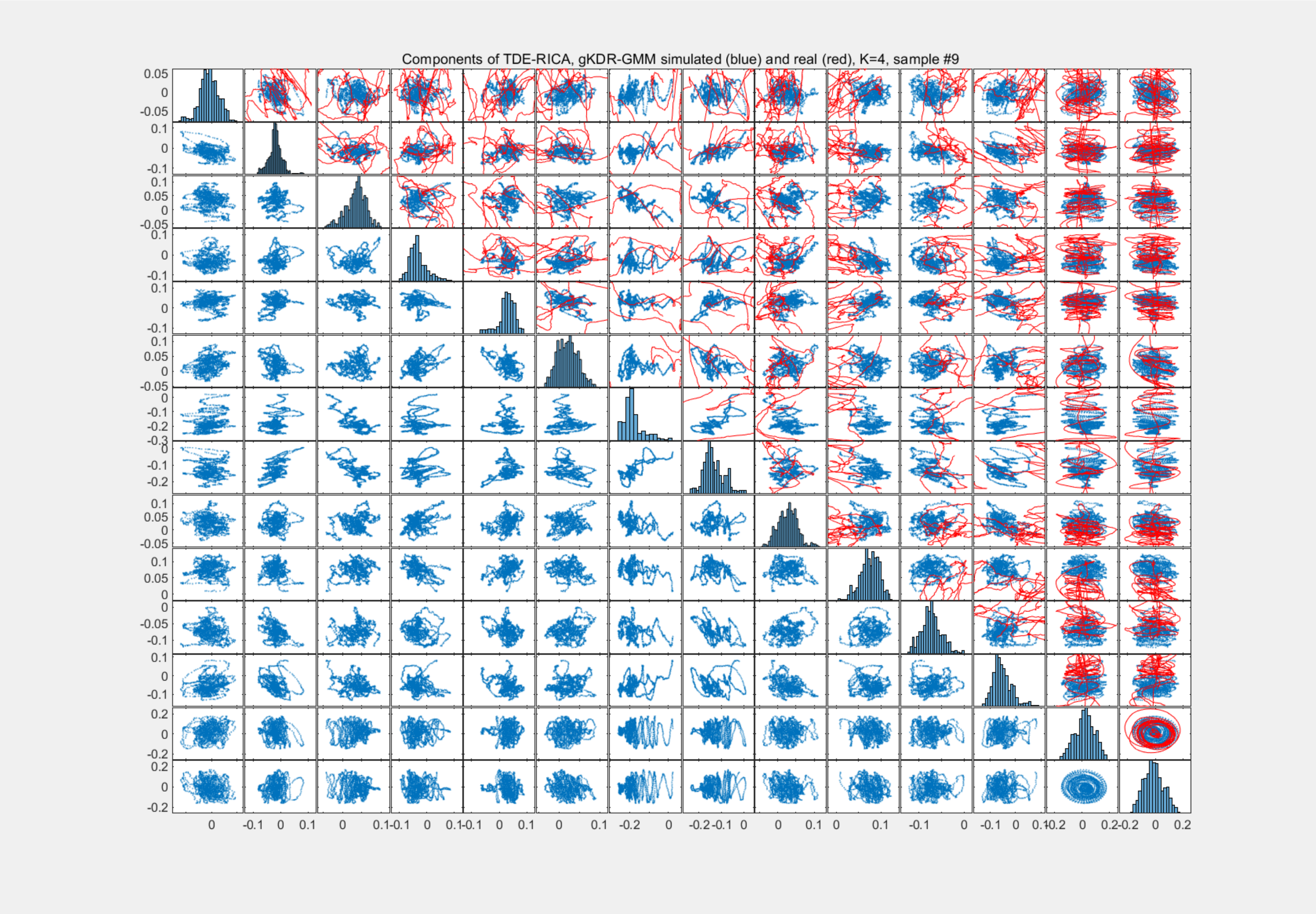
Components of TDE-RICA, gKDR-GMM simulated (blue) and real (red), K=4, sample #9

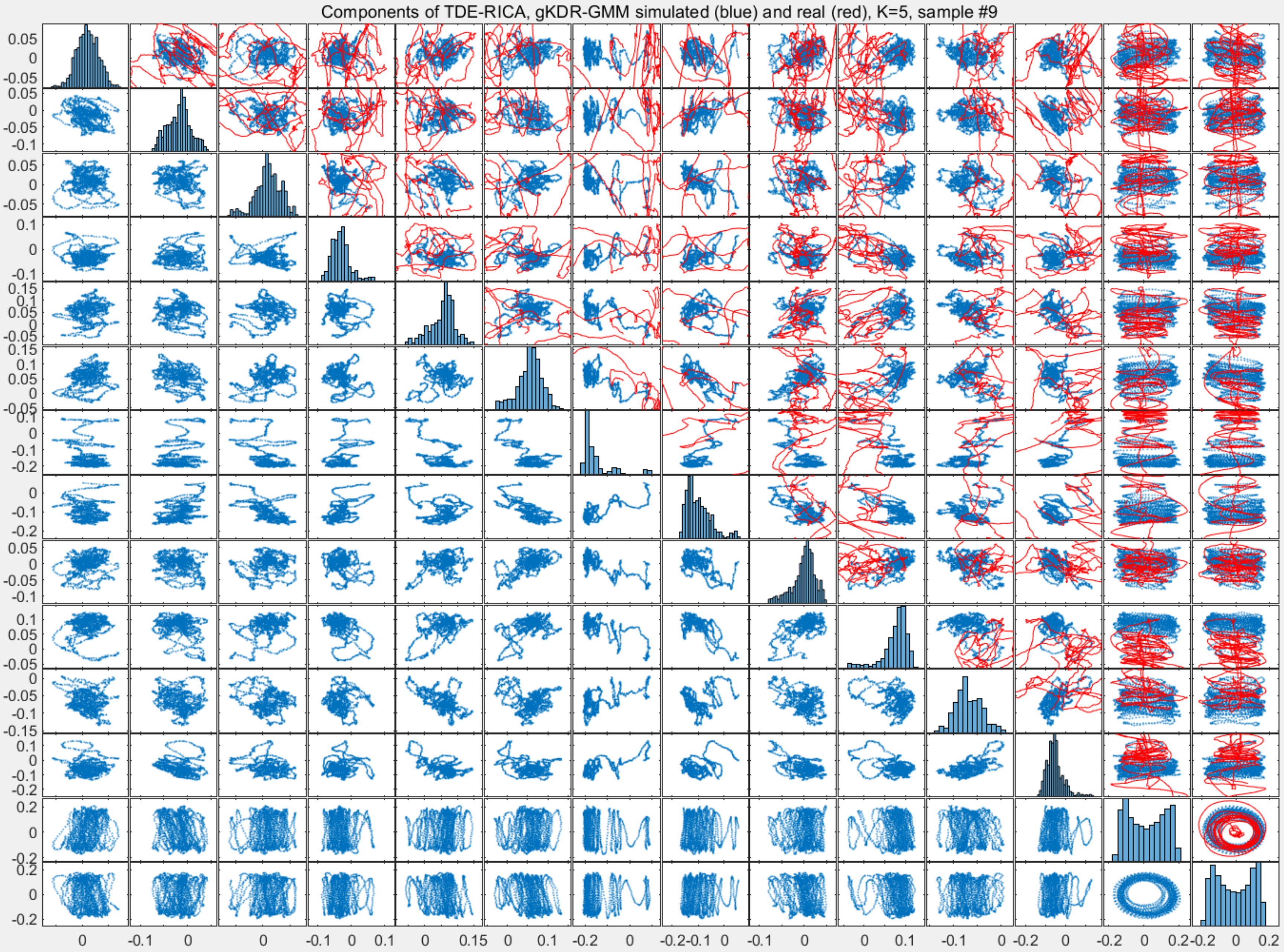

Components of TDE-RICA, gKDR-GMM simulated (blue) and real (red), K=5, sample #9

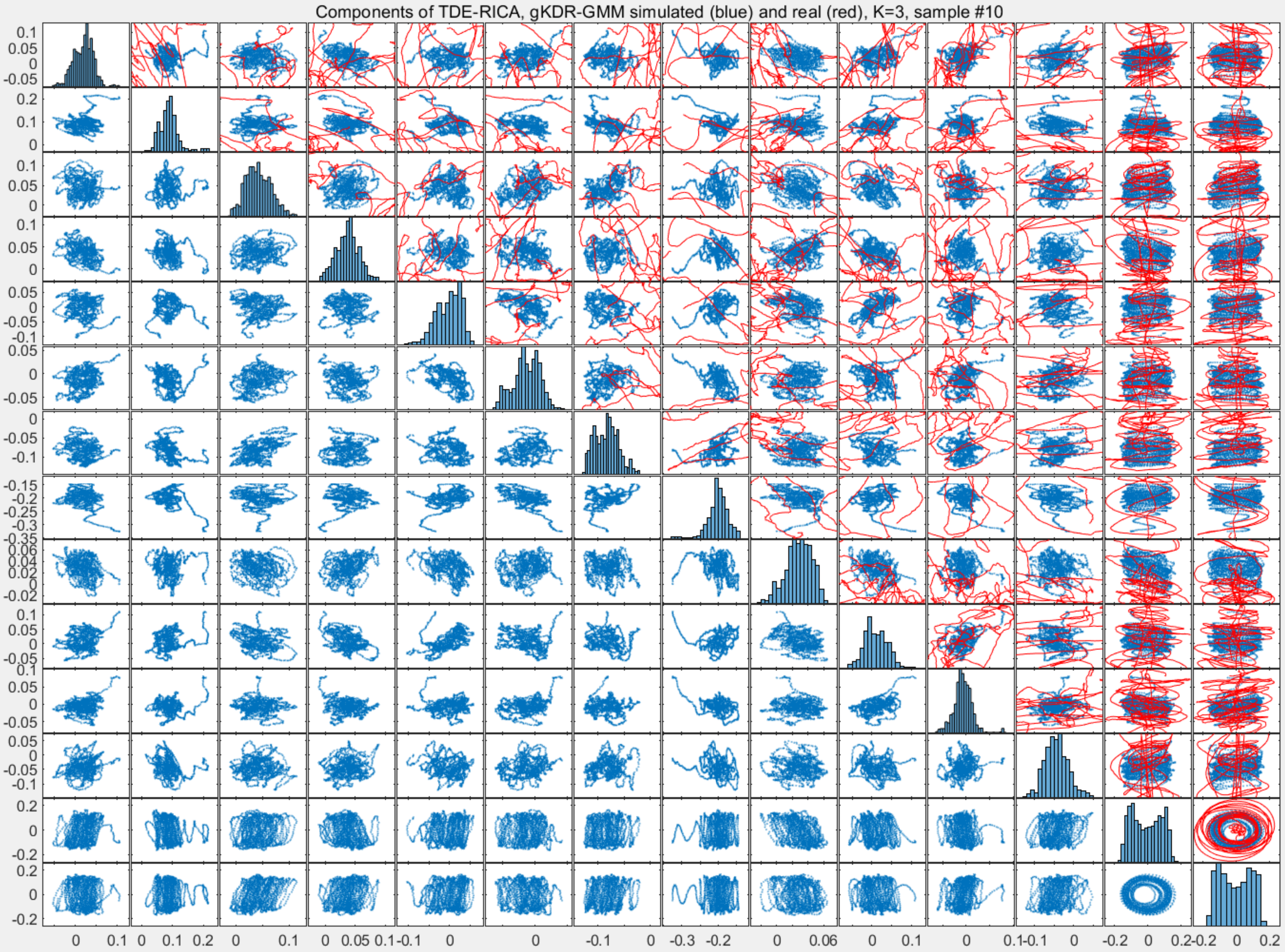

Components of TDE-RICA, gKDR-GMM simulated (blue) and real (red), K=3, sample #10

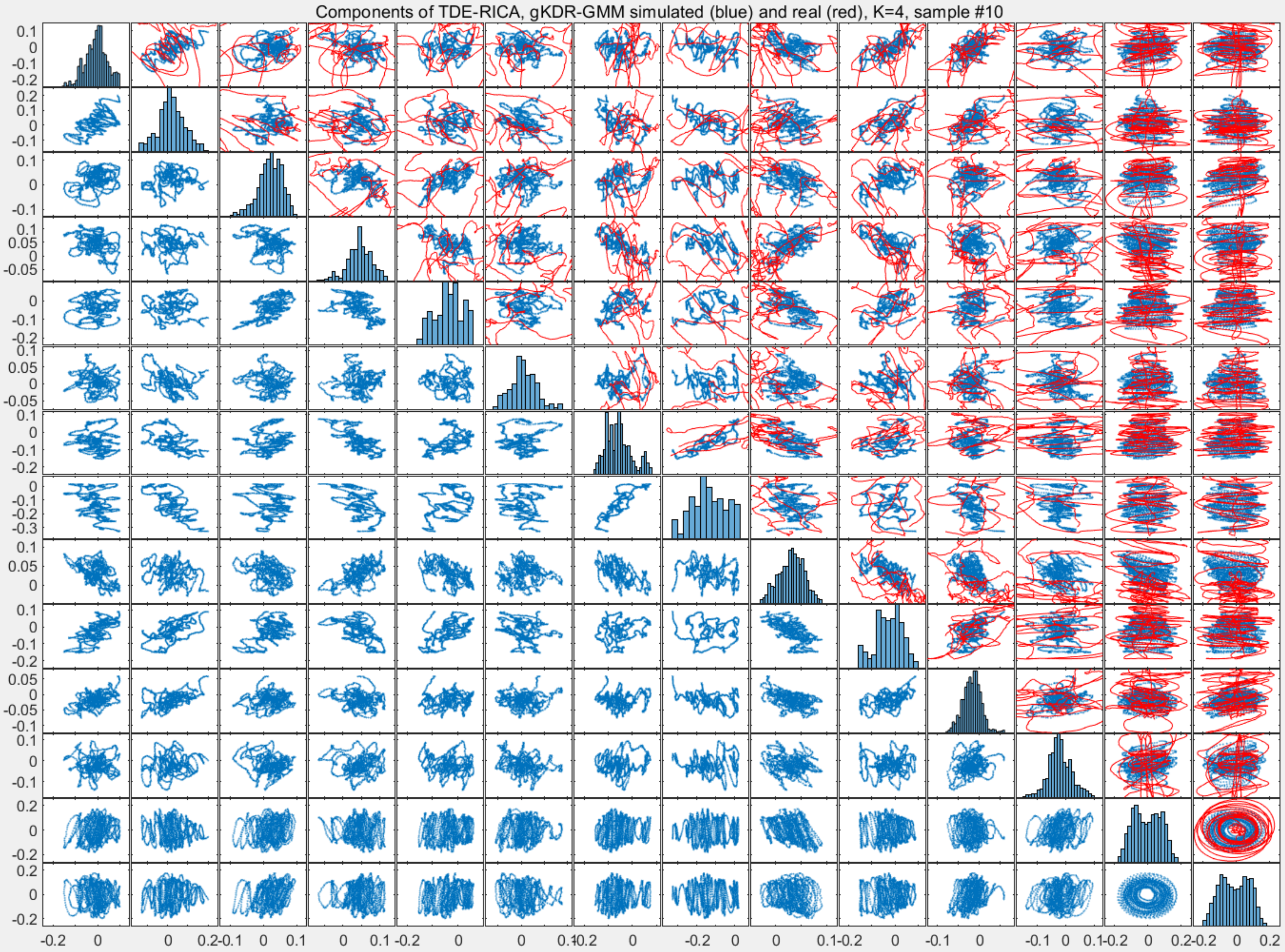
Components of TDE-RICA, gKDR-GMM simulated (blue) and real (red), K=4, sample #10

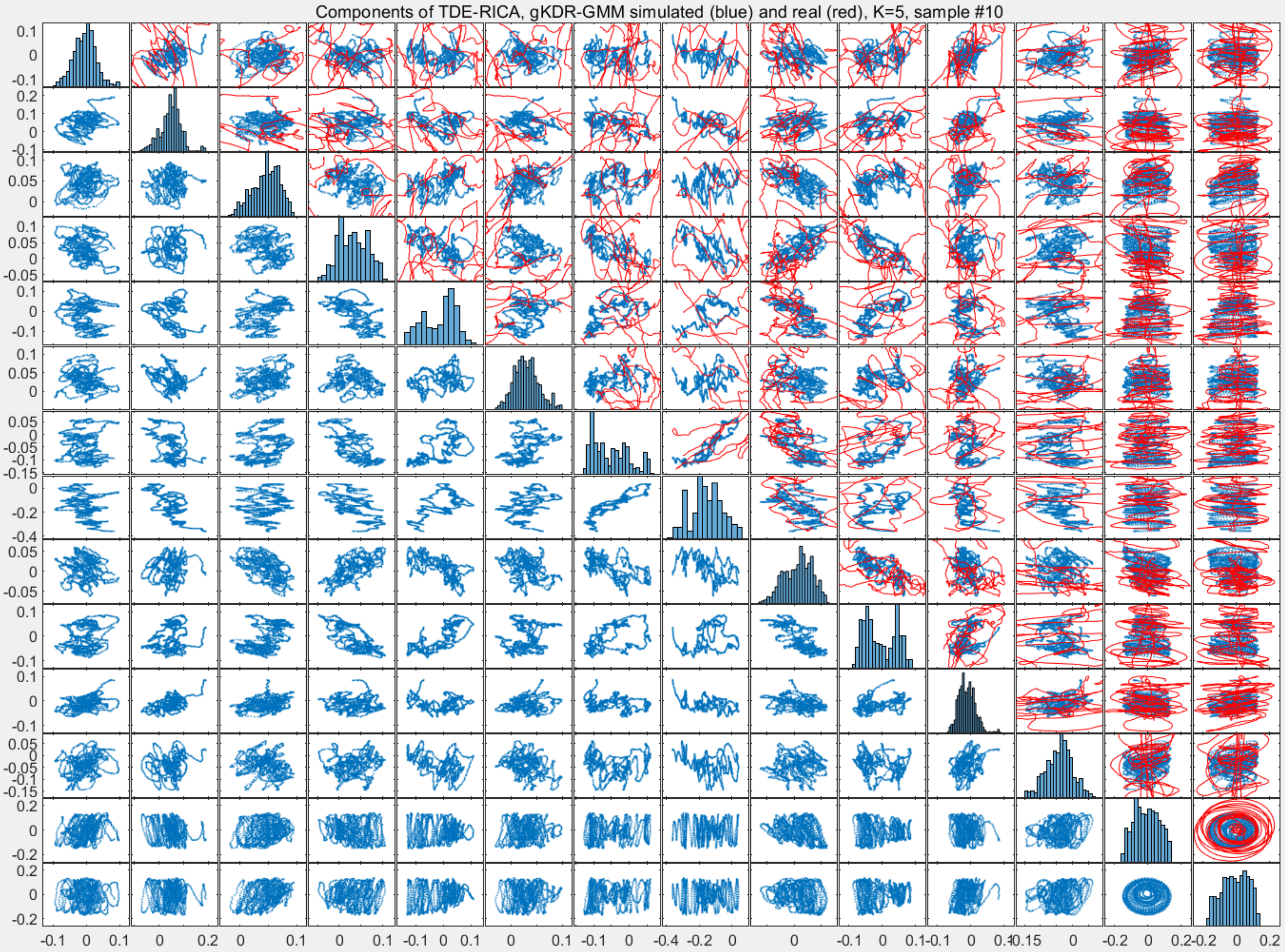

Components of TDE-RICA, gKDR-GMM simulated (blue) and real (red), K=5, sample #10

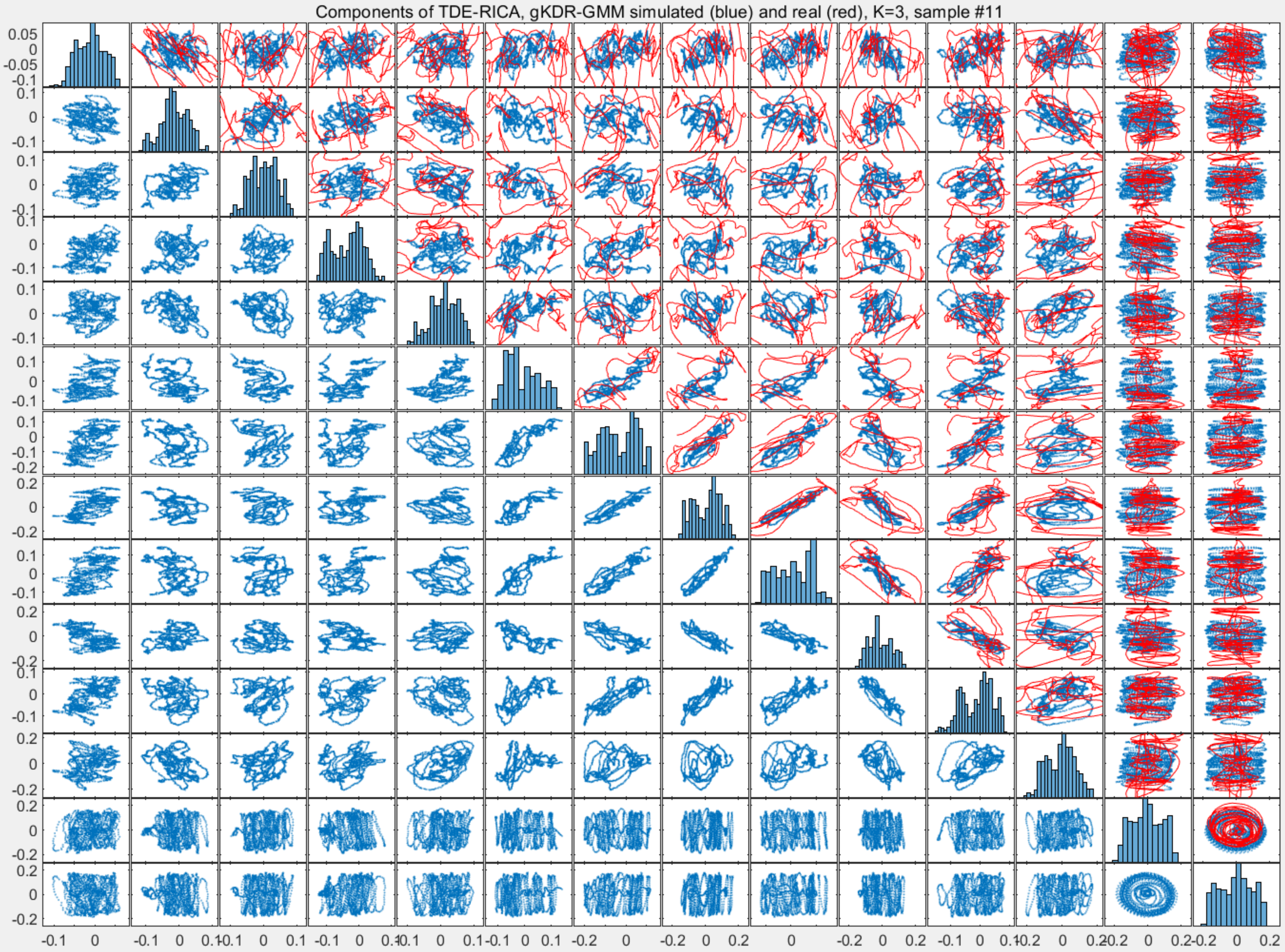
Components of TDE-RICA, gKDR-GMM simulated (blue) and real (red), K=3, sample #11

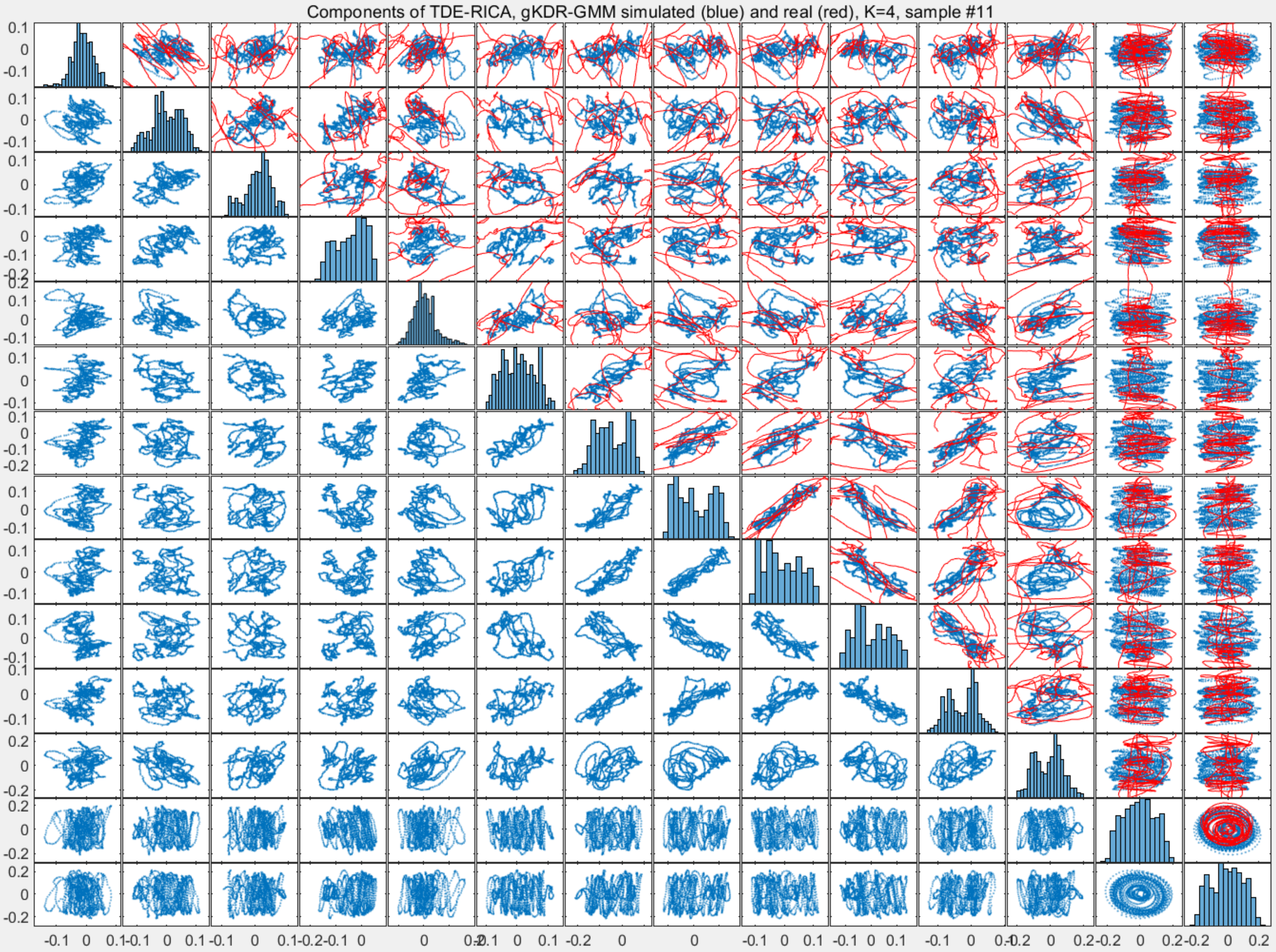

Components of TDE-RICA, gKDR-GMM simulated (blue) and real (red), K=4, sample #11

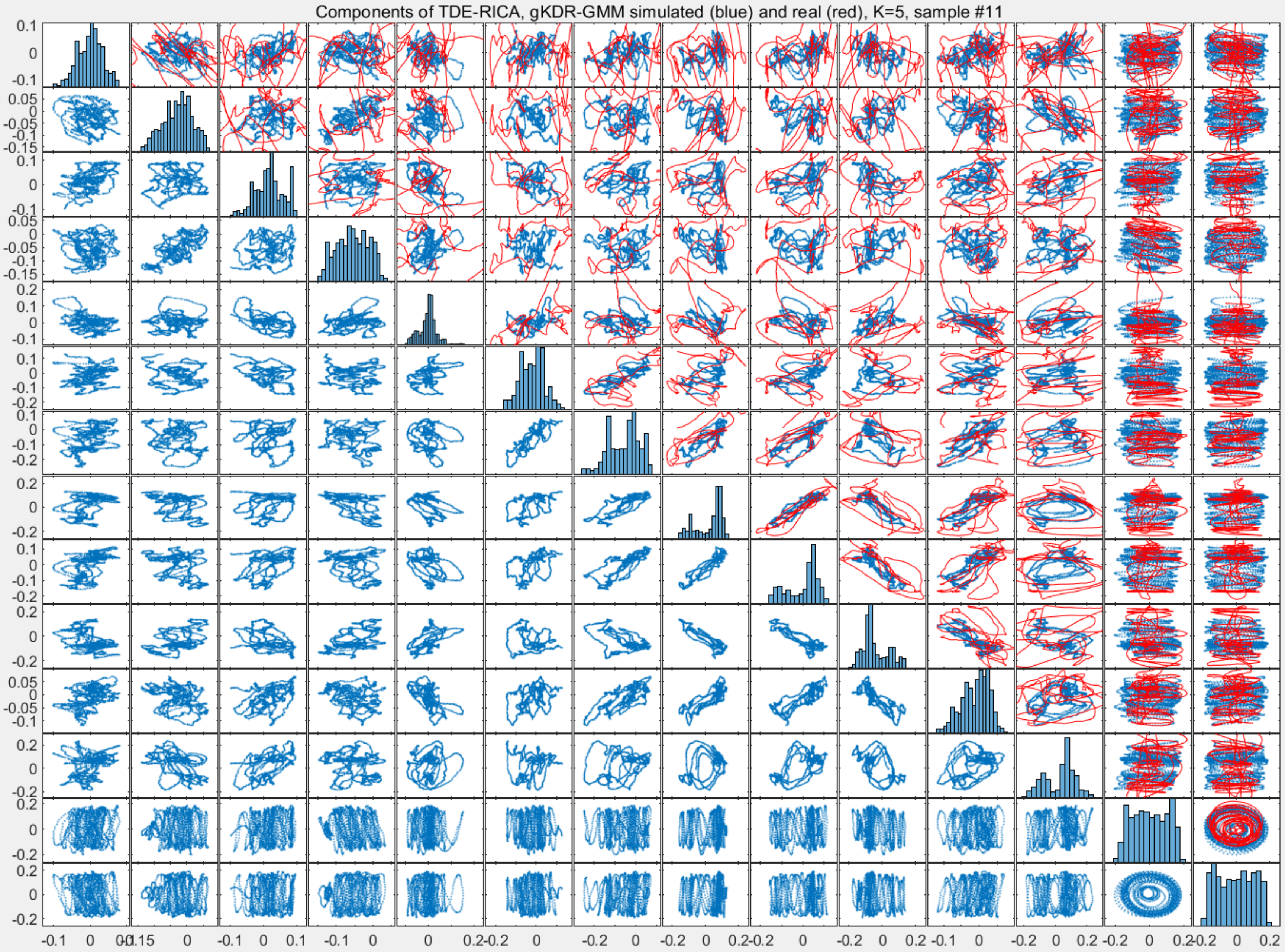
Components of TDE-RICA, gKDR-GMM simulated (blue) and real (red), K=5, sample #11

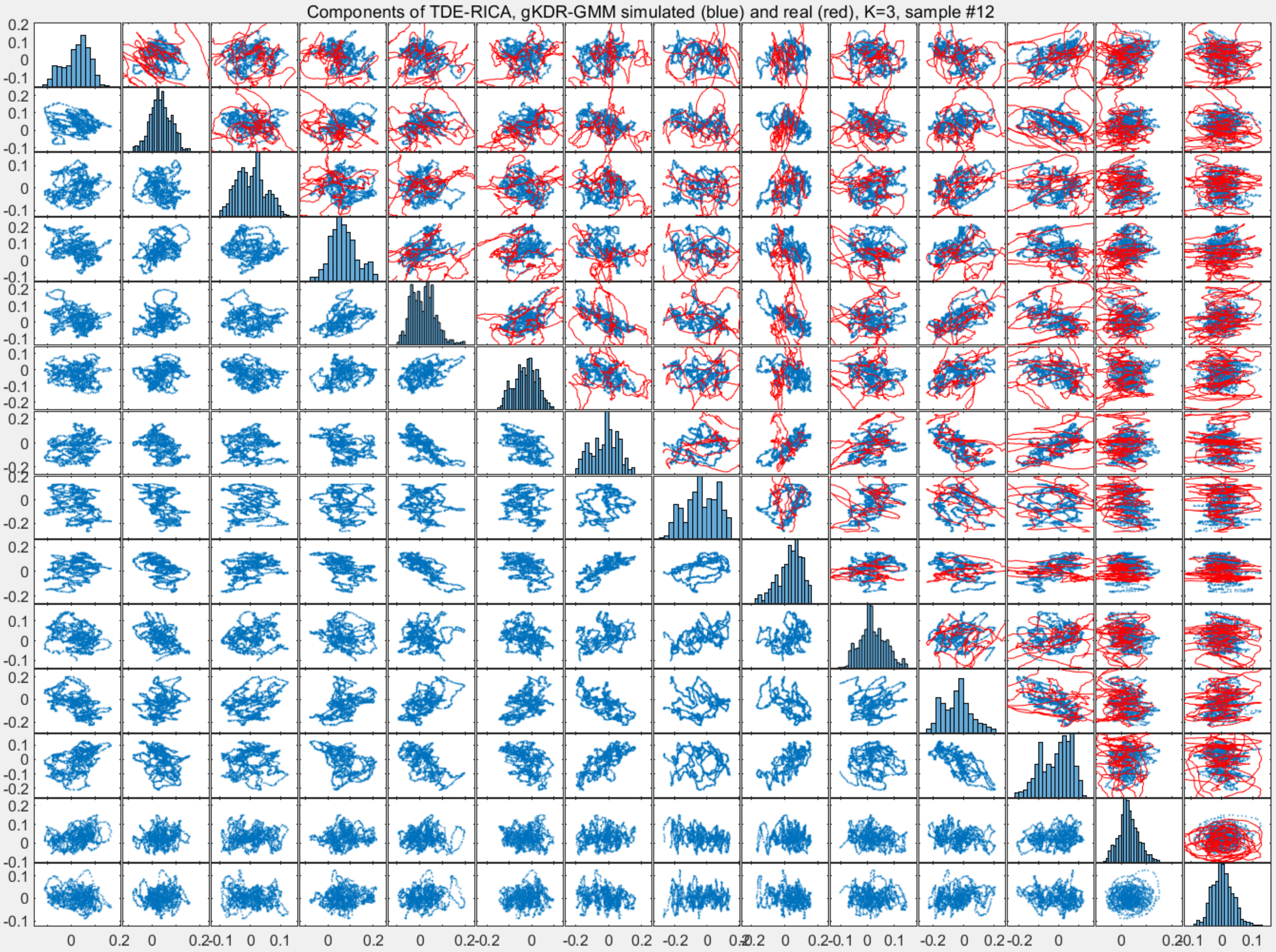

Components of TDE-RICA, gKDR-GMM simulated (blue) and real (red), K=3, sample #12

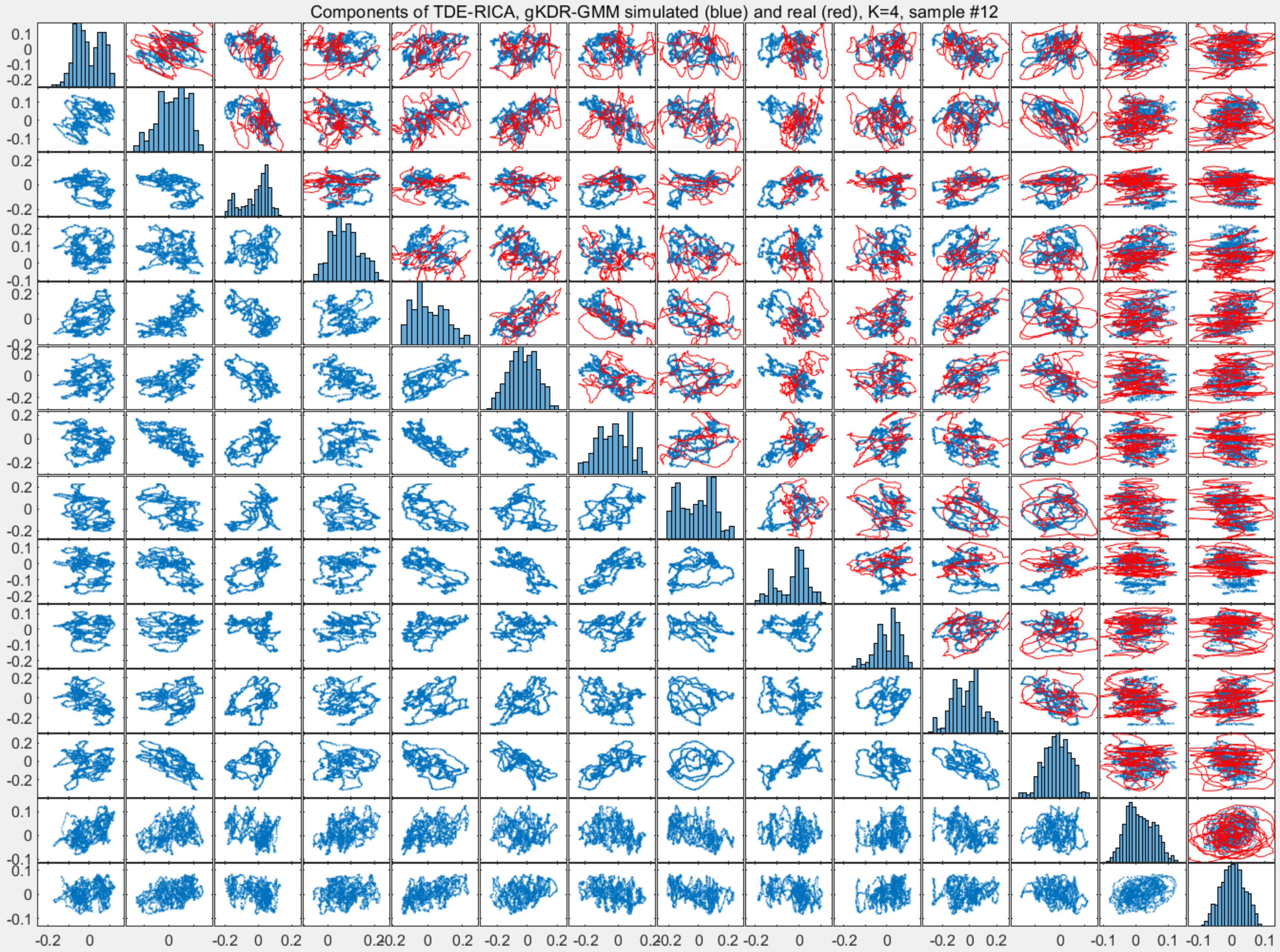
Components of TDE-RICA, gKDR-GMM simulated (blue) and real (red), K=4, sample #12

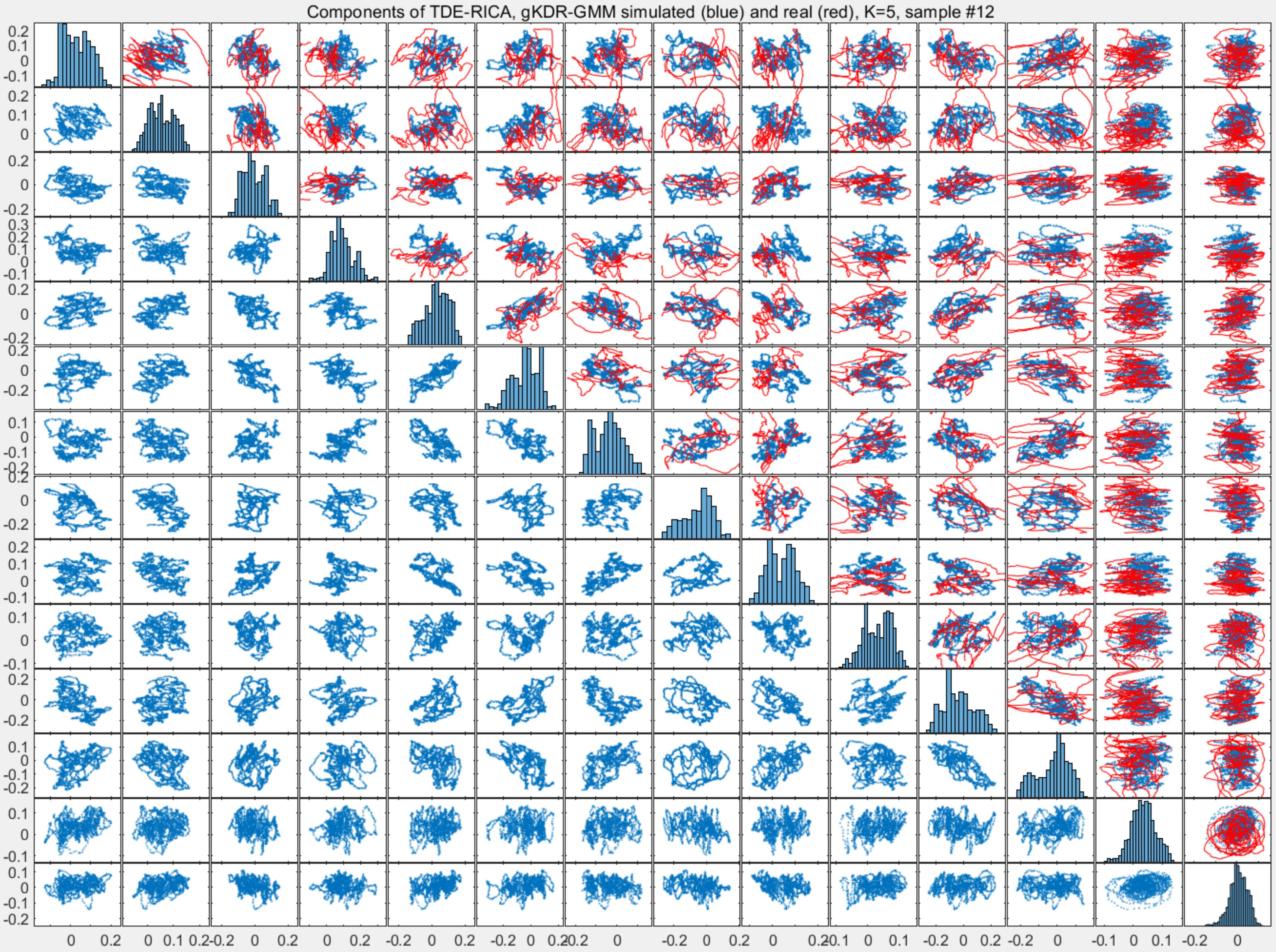

Components of TDE-RICA, gKDR-GMM simulated (blue) and real (red), K=5, sample #12

Components of TDE-RICA, gKDR-GMM simulated (blue) and real (red), K=3, sample #13

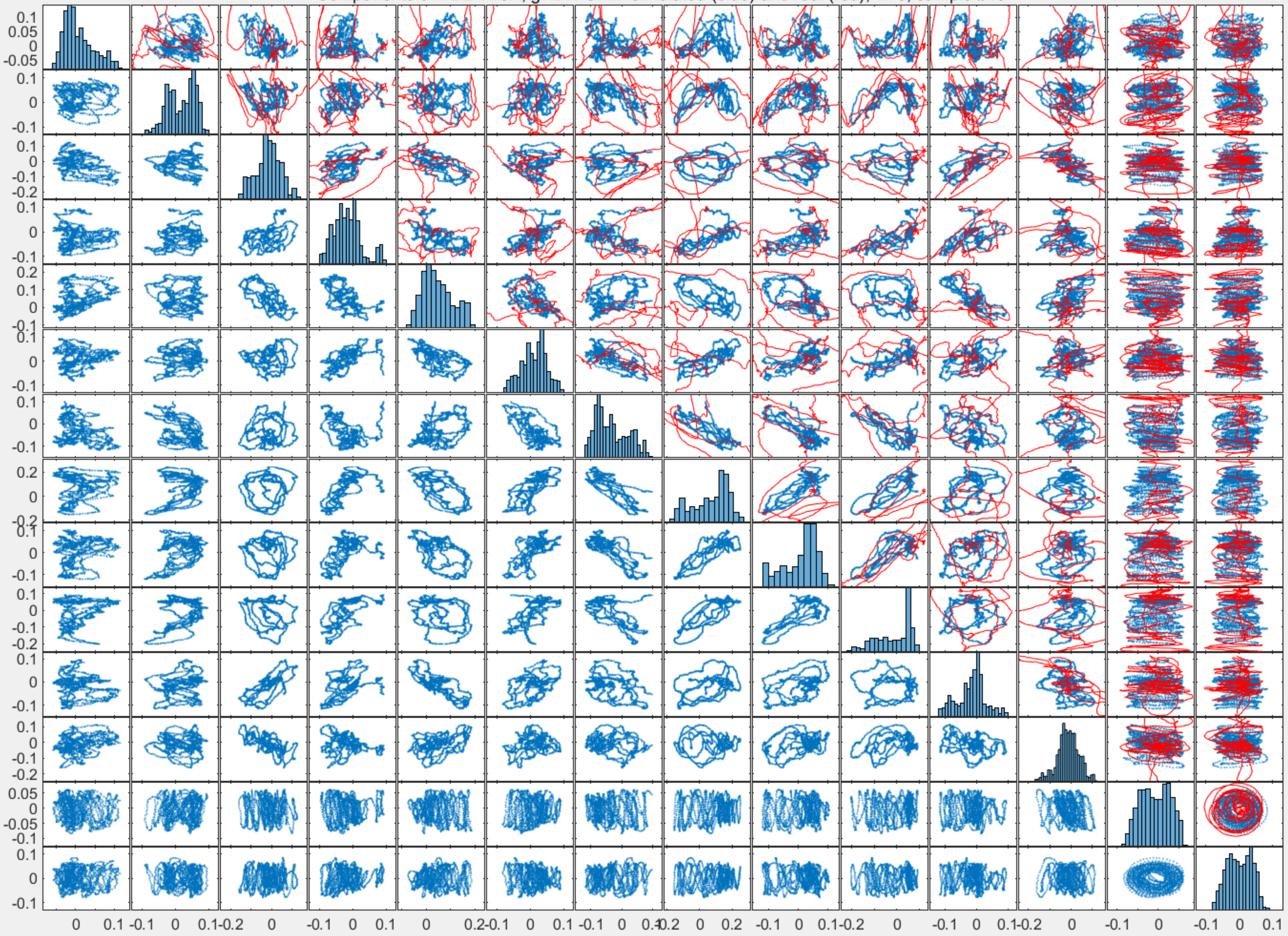

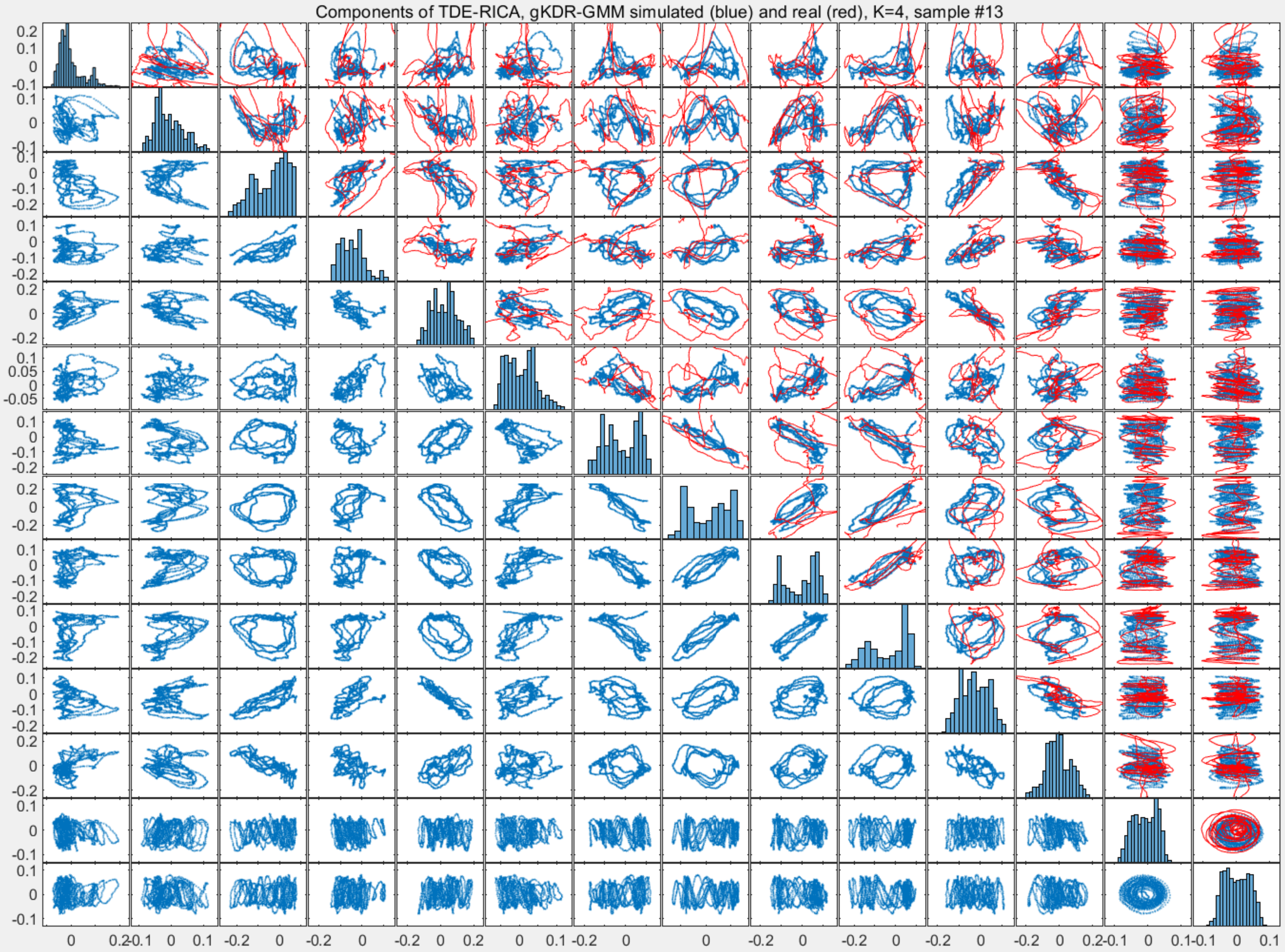
Components of TDE-RICA, gKDR-GMM simulated (blue) and real (red), K=4, sample #13

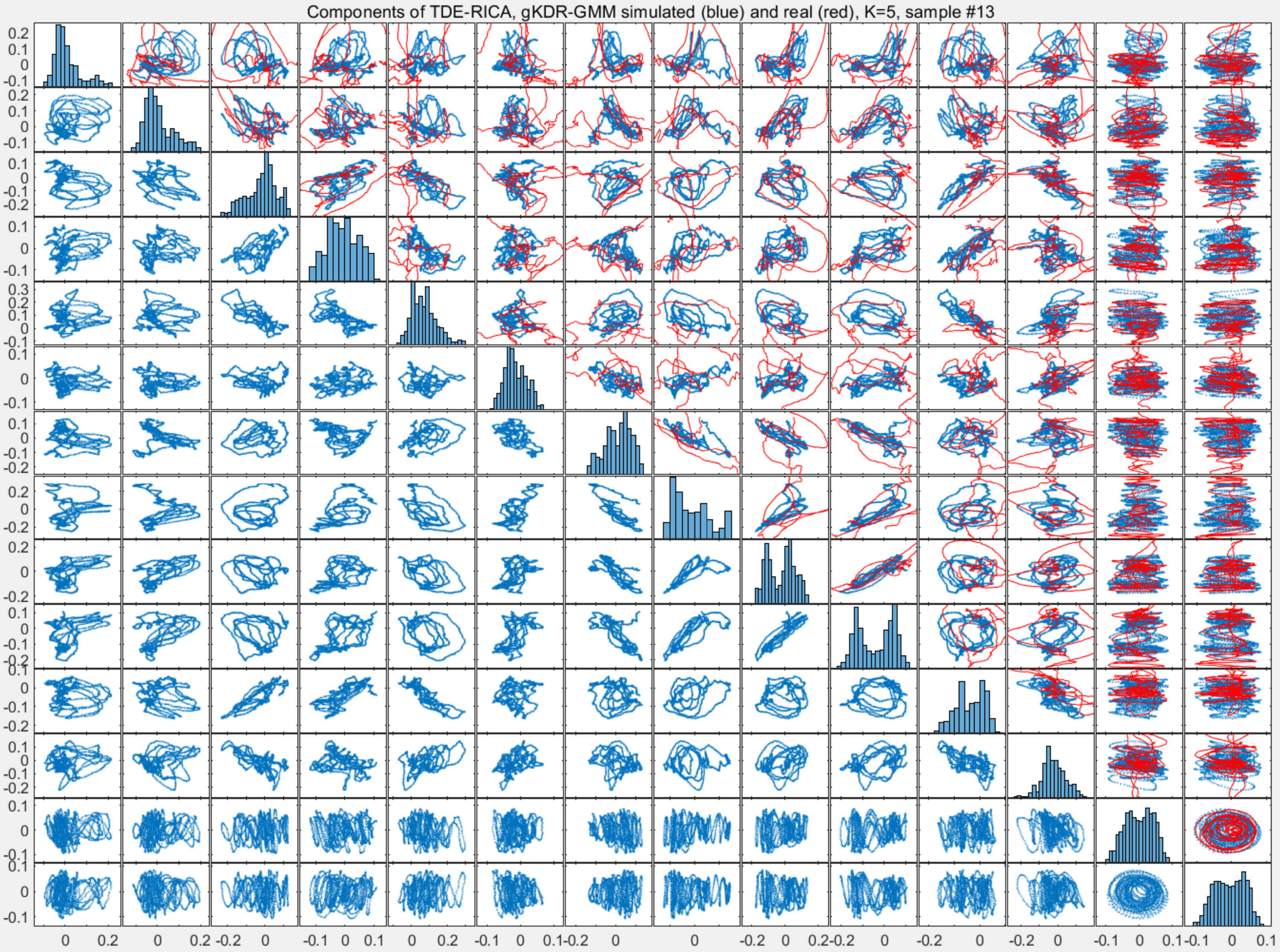
Components of TDE-RICA, gKDR-GMM simulated (blue) and real (red), K=5, sample #13

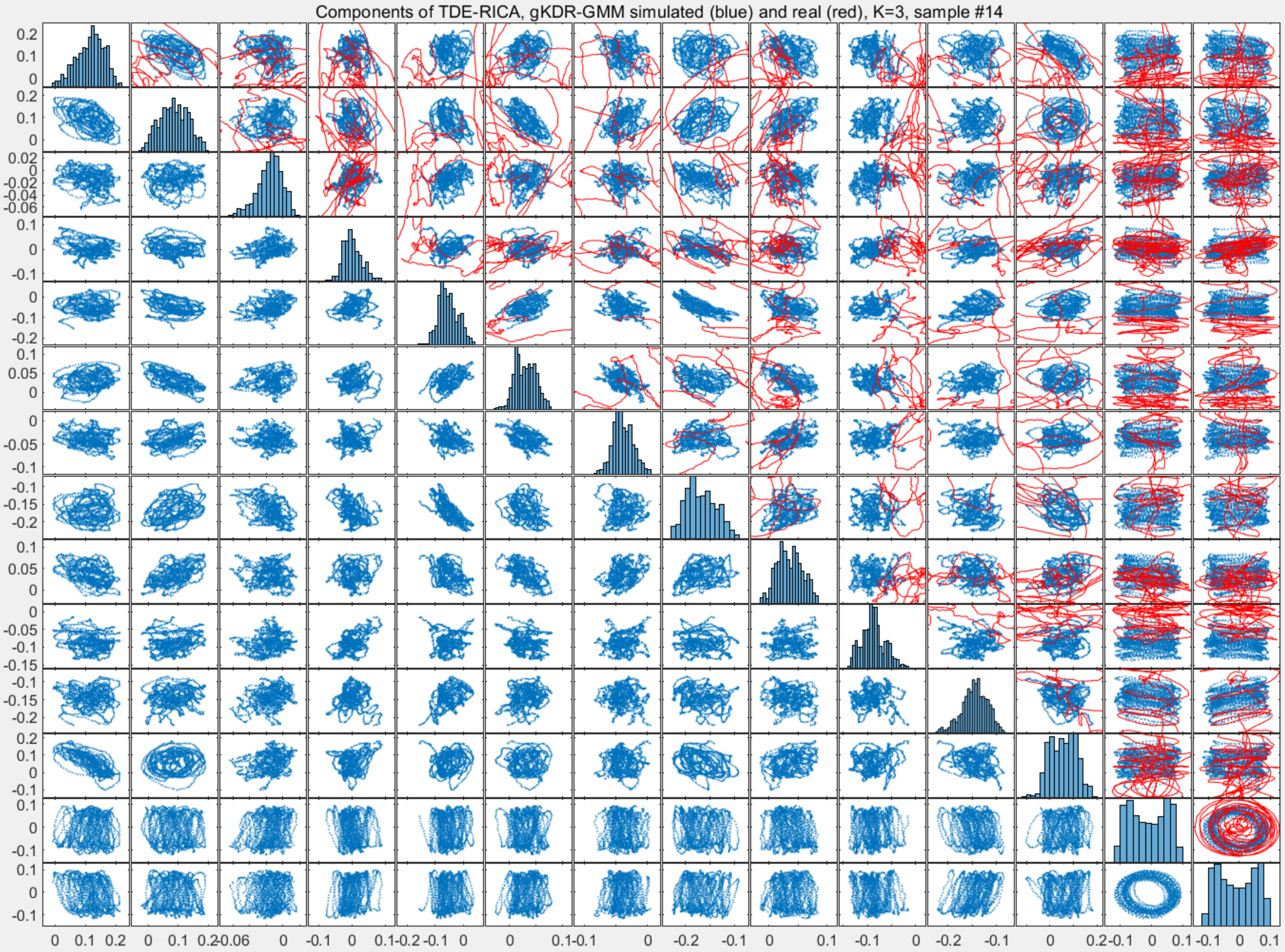
Components of TDE-RICA, gKDR-GMM simulated (blue) and real (red), K=3, sample #14

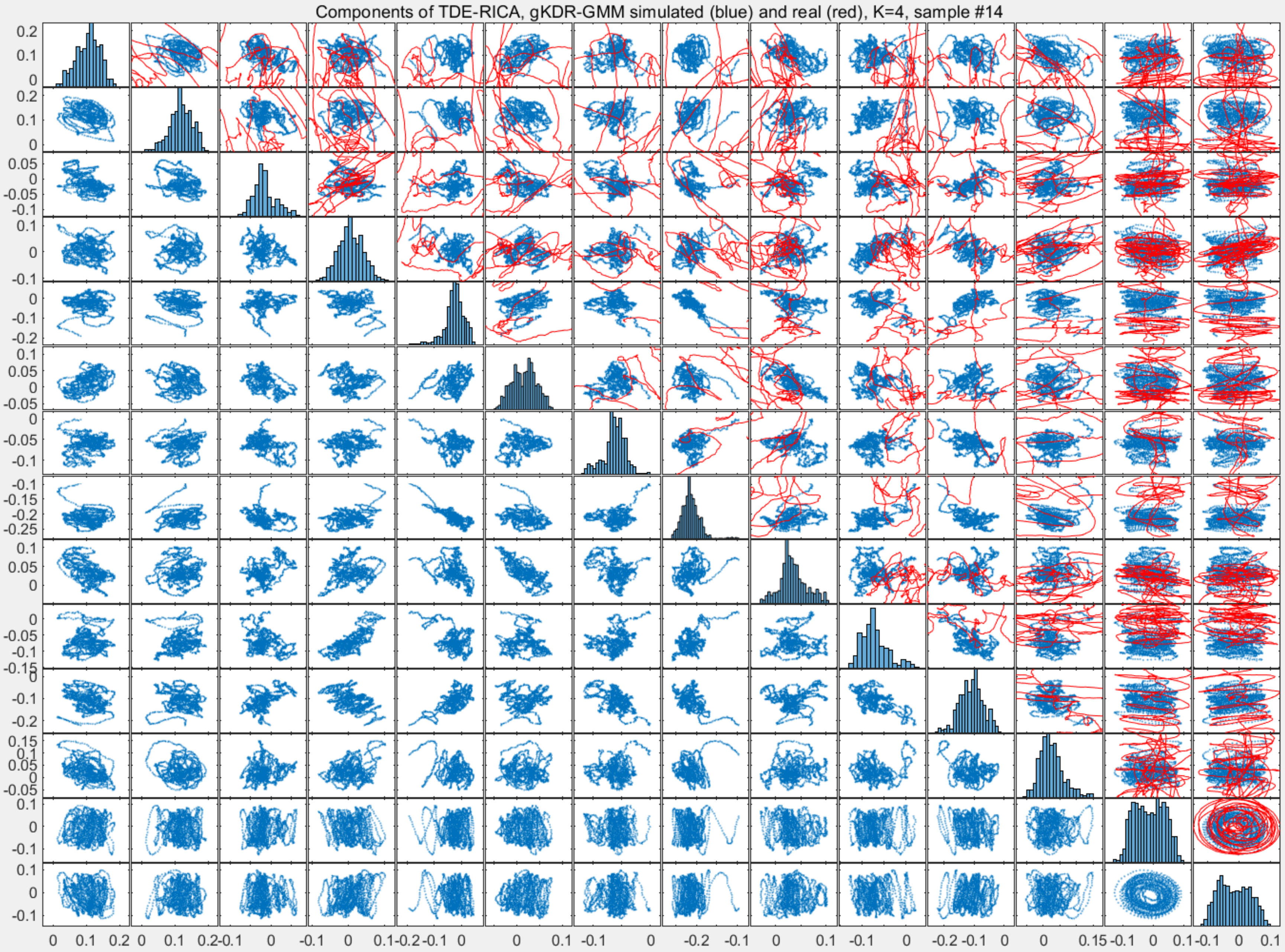
Components of TDE-RICA, gKDR-GMM simulated (blue) and real (red), K=4, sample #14

Components of TDE-RICA, gKDR-GMM simulated (blue) and real (red), K=5, sample #14

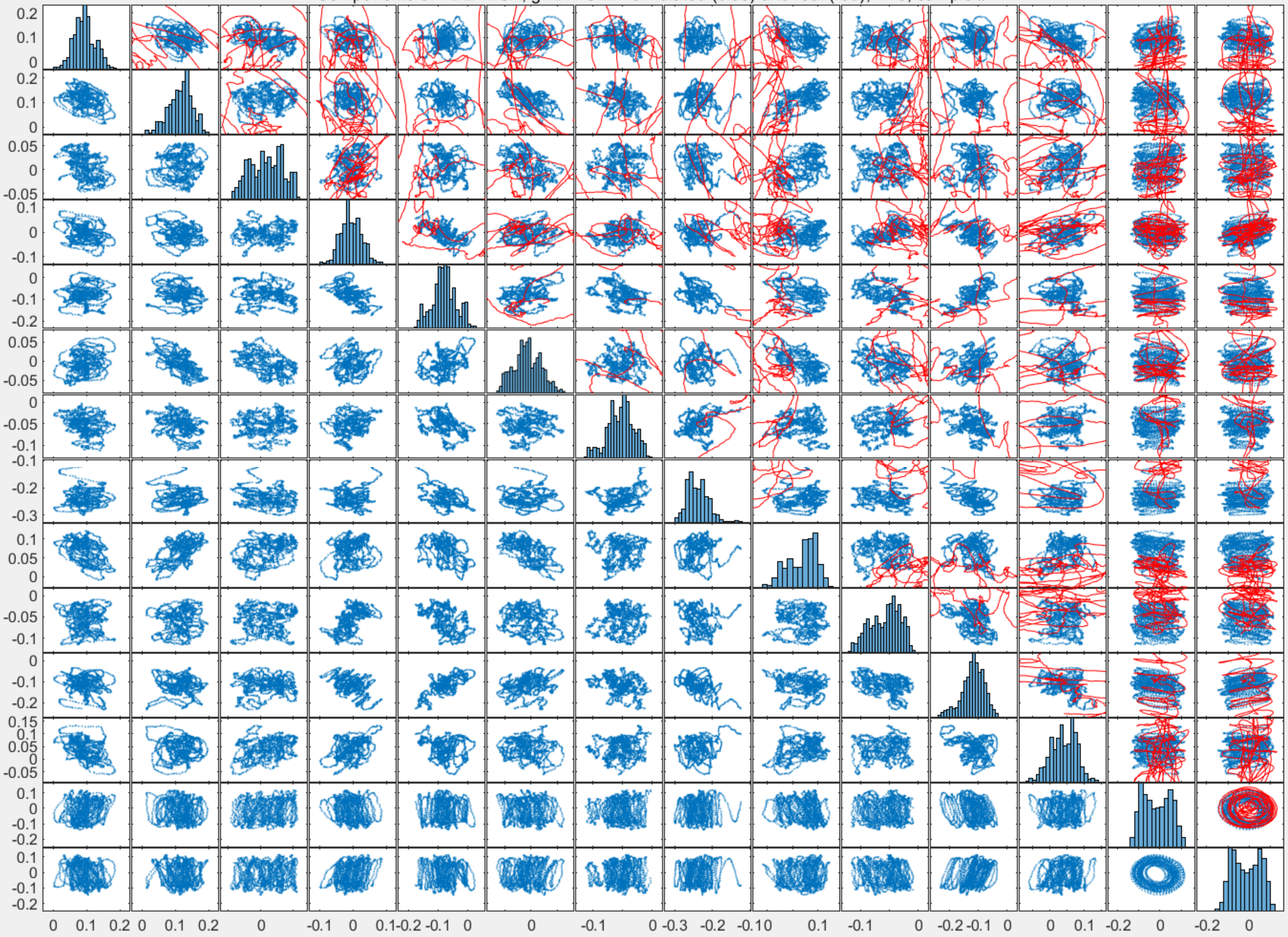

Components of TDE-RICA, gKDR-GMM simulated (blue) and real (red), K=3, sample #15

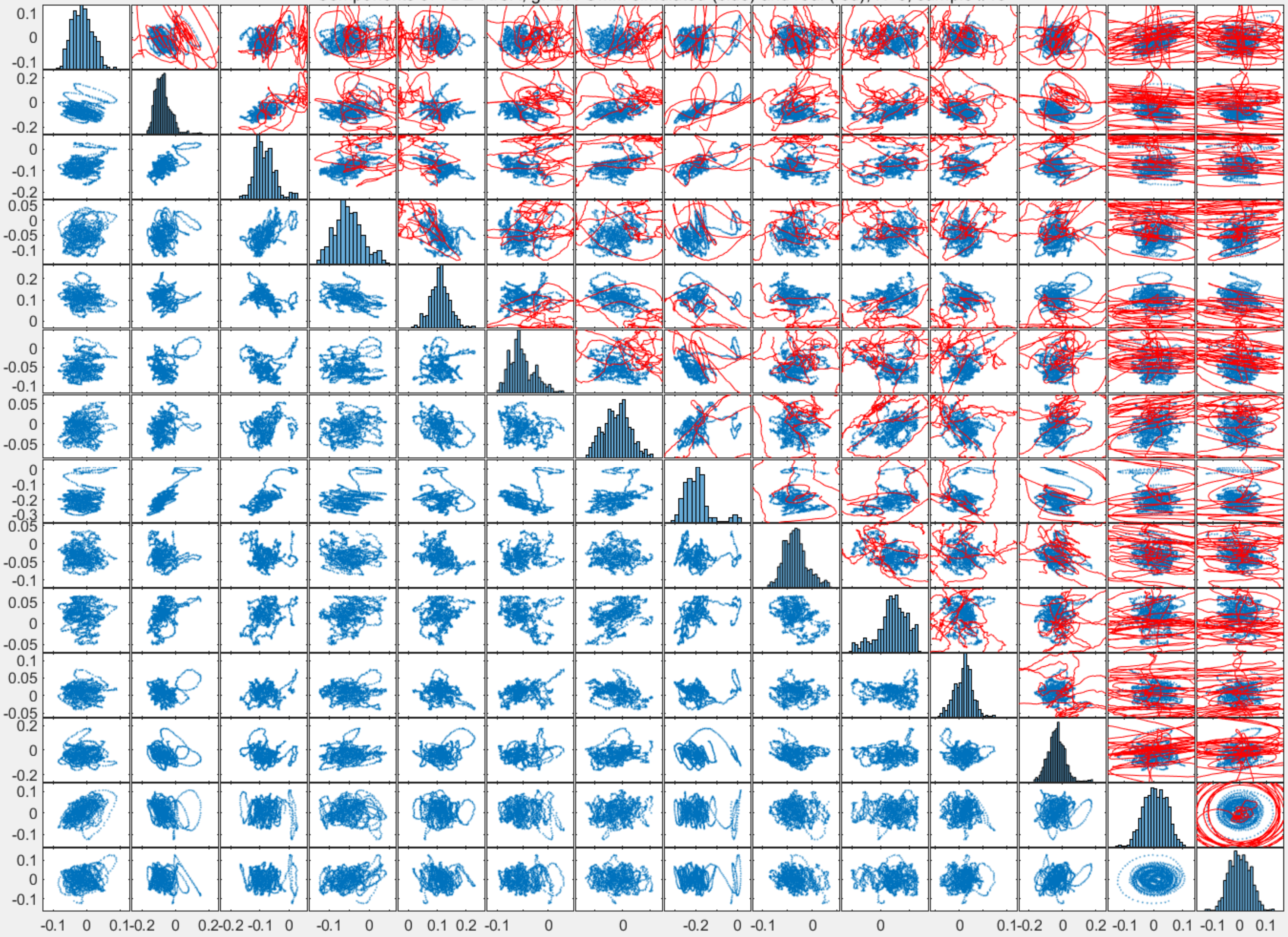

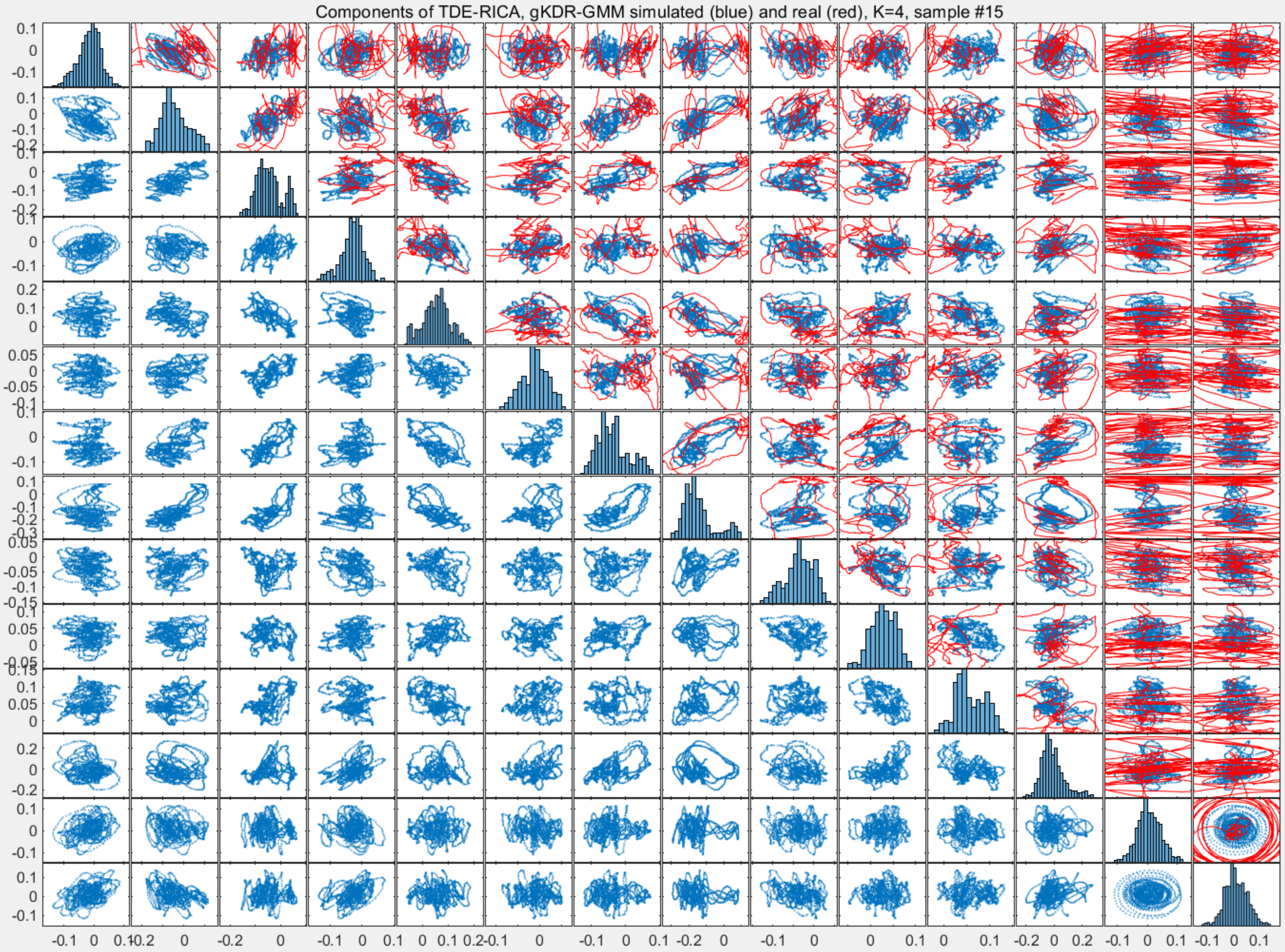
Components of TDE-RICA, gKDR-GMM simulated (blue) and real (red), K=4, sample #15

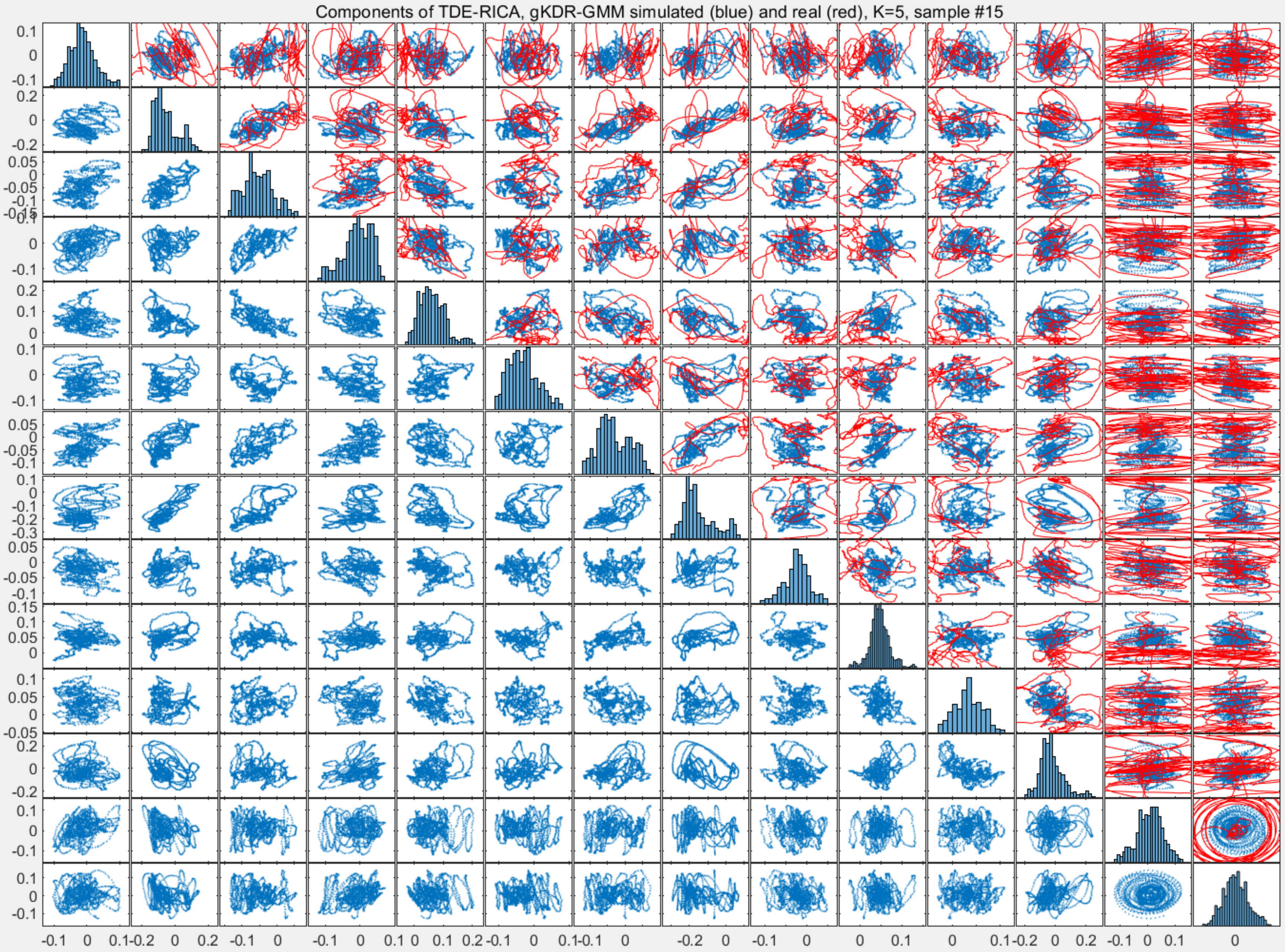
Components of TDE-RICA, gKDR-GMM simulated (blue) and real (red), K=5, sample #15

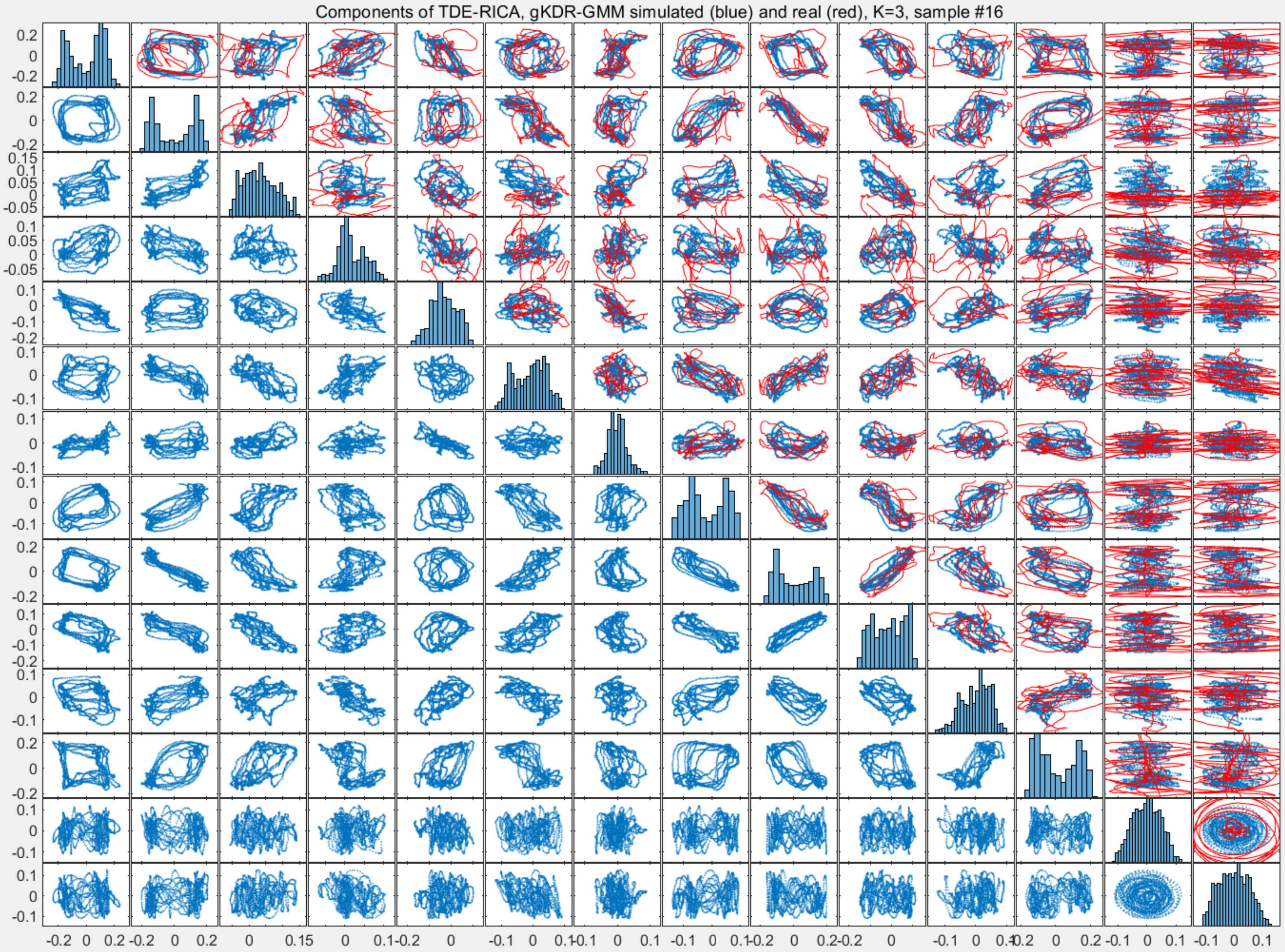
Components of TDE-RICA, gKDR-GMM simulated (blue) and real (red), K=3, sample #16

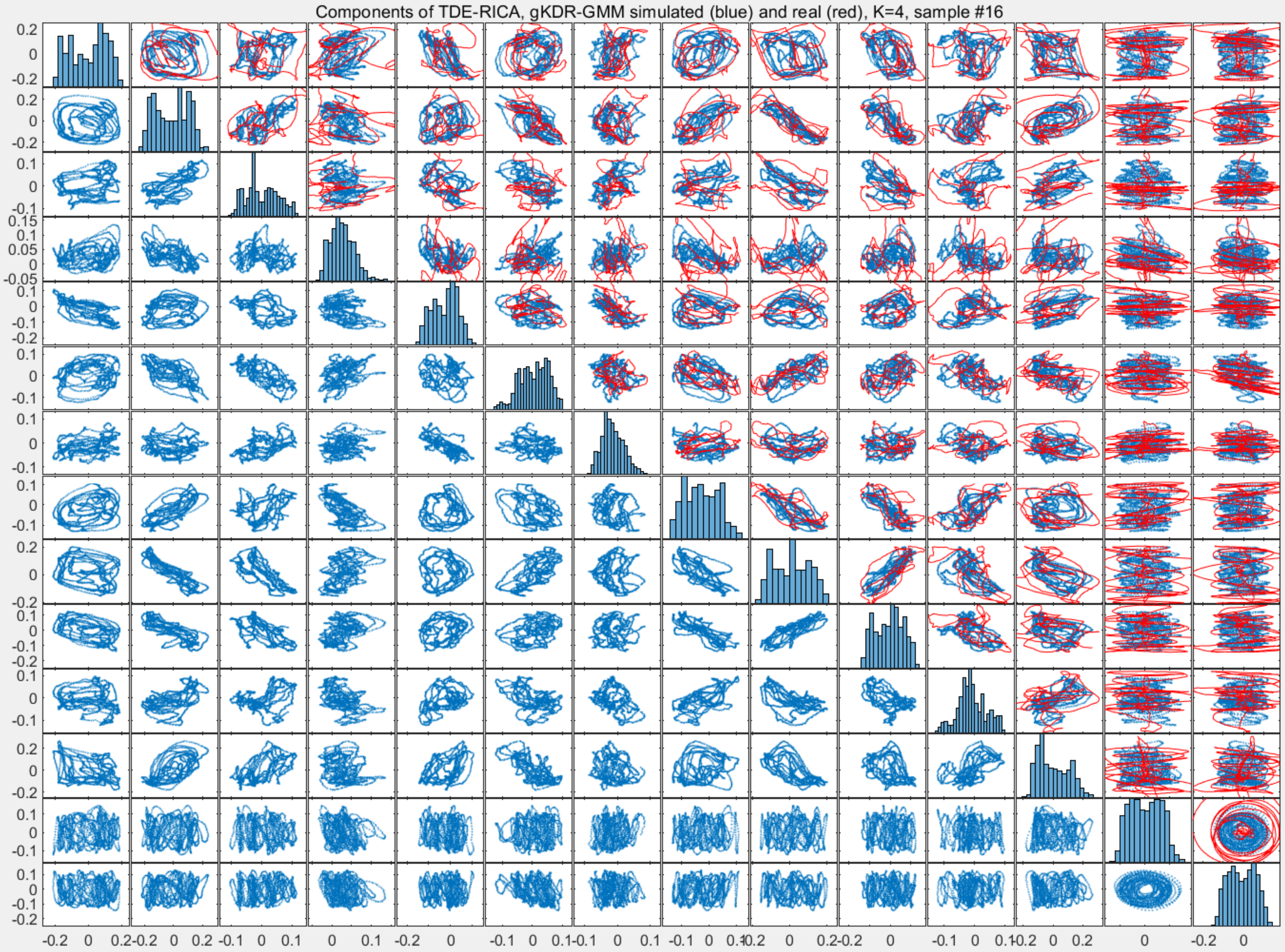
Components of TDE-RICA, gKDR-GMM simulated (blue) and real (red), K=4, sample #16

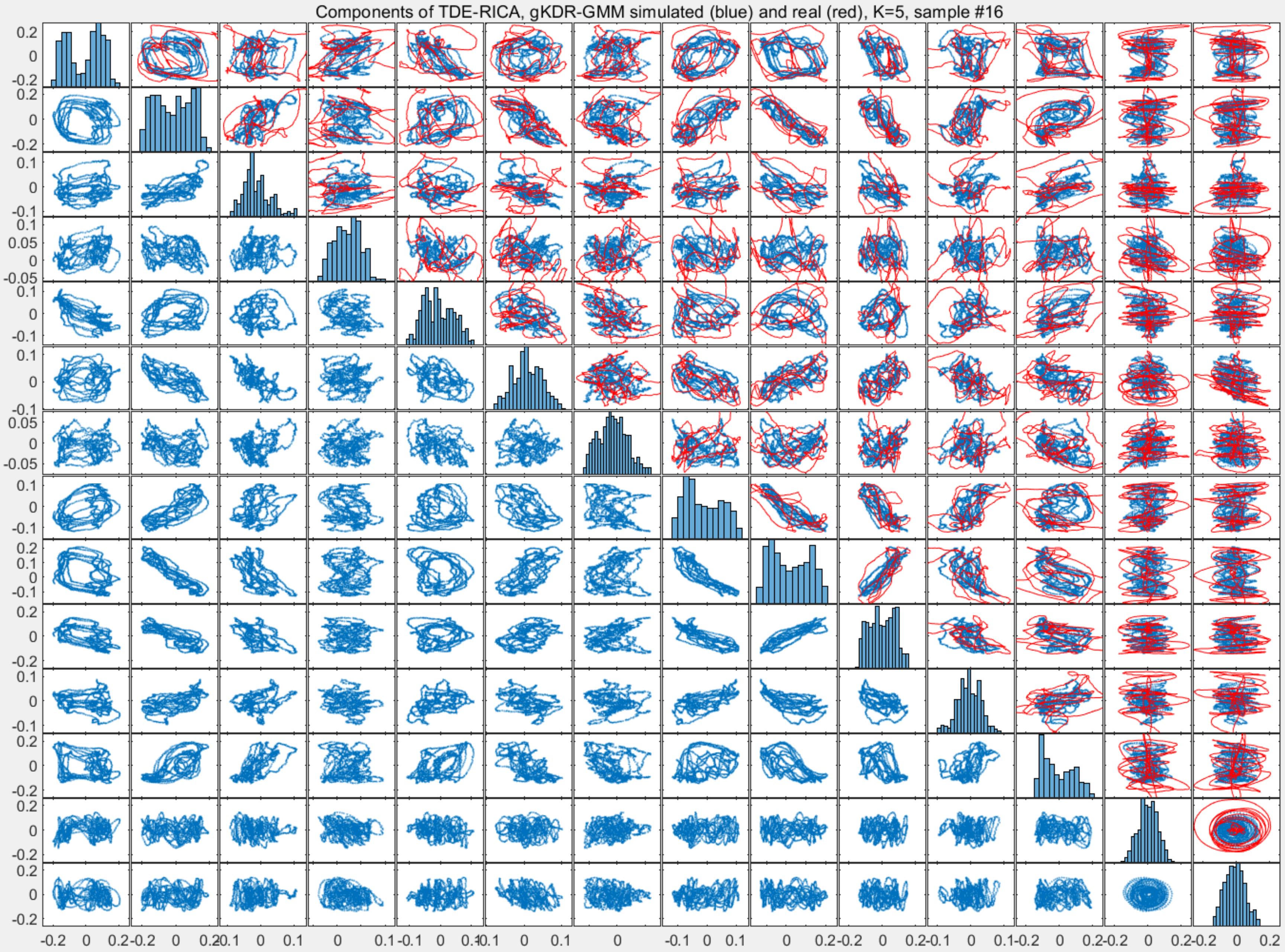
Components of TDE-RICA, gKDR-GMM simulated (blue) and real (red), K=5, sample #16

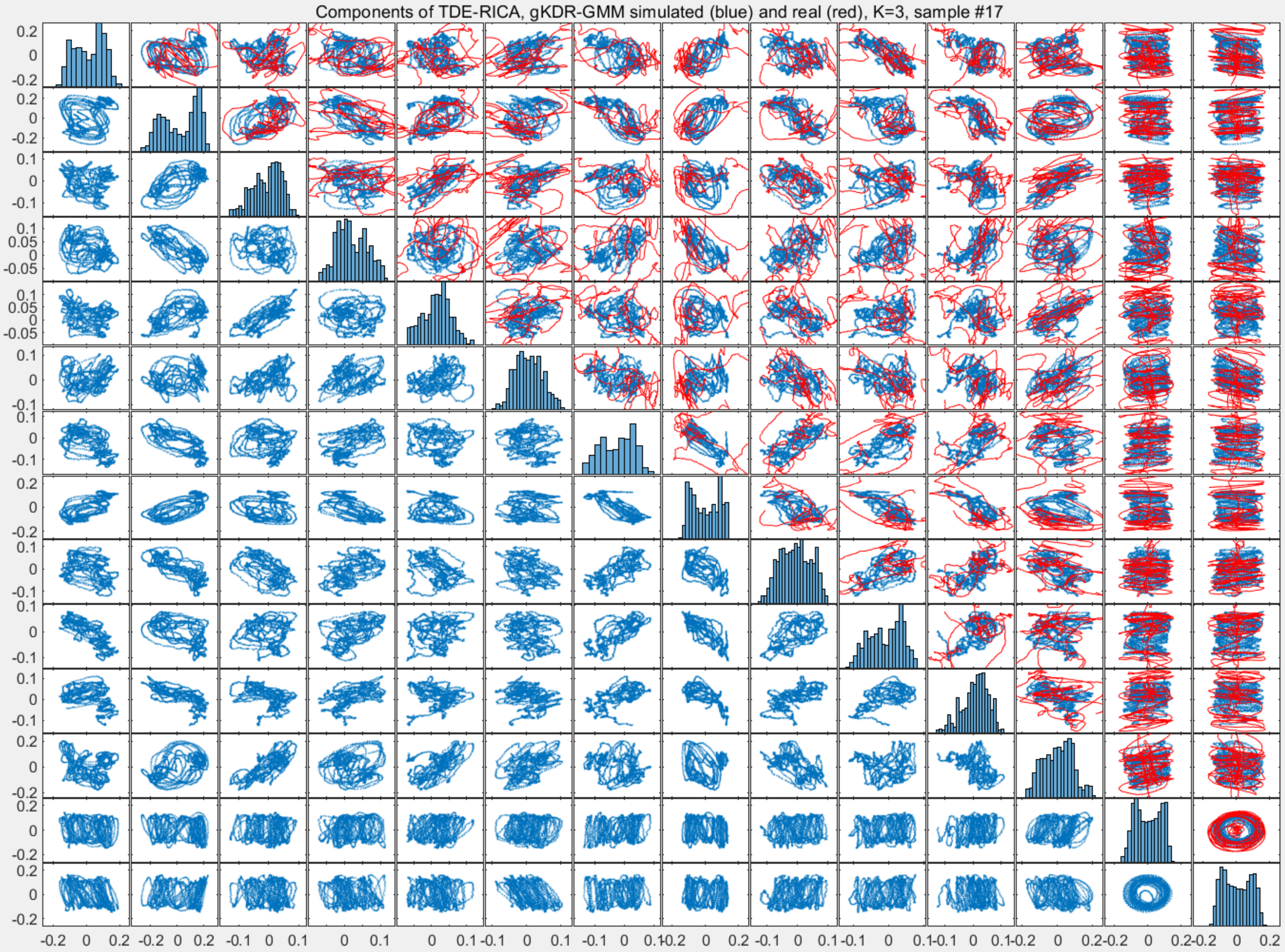
Components of TDE-RICA, gKDR-GMM simulated (blue) and real (red), K=3, sample #17

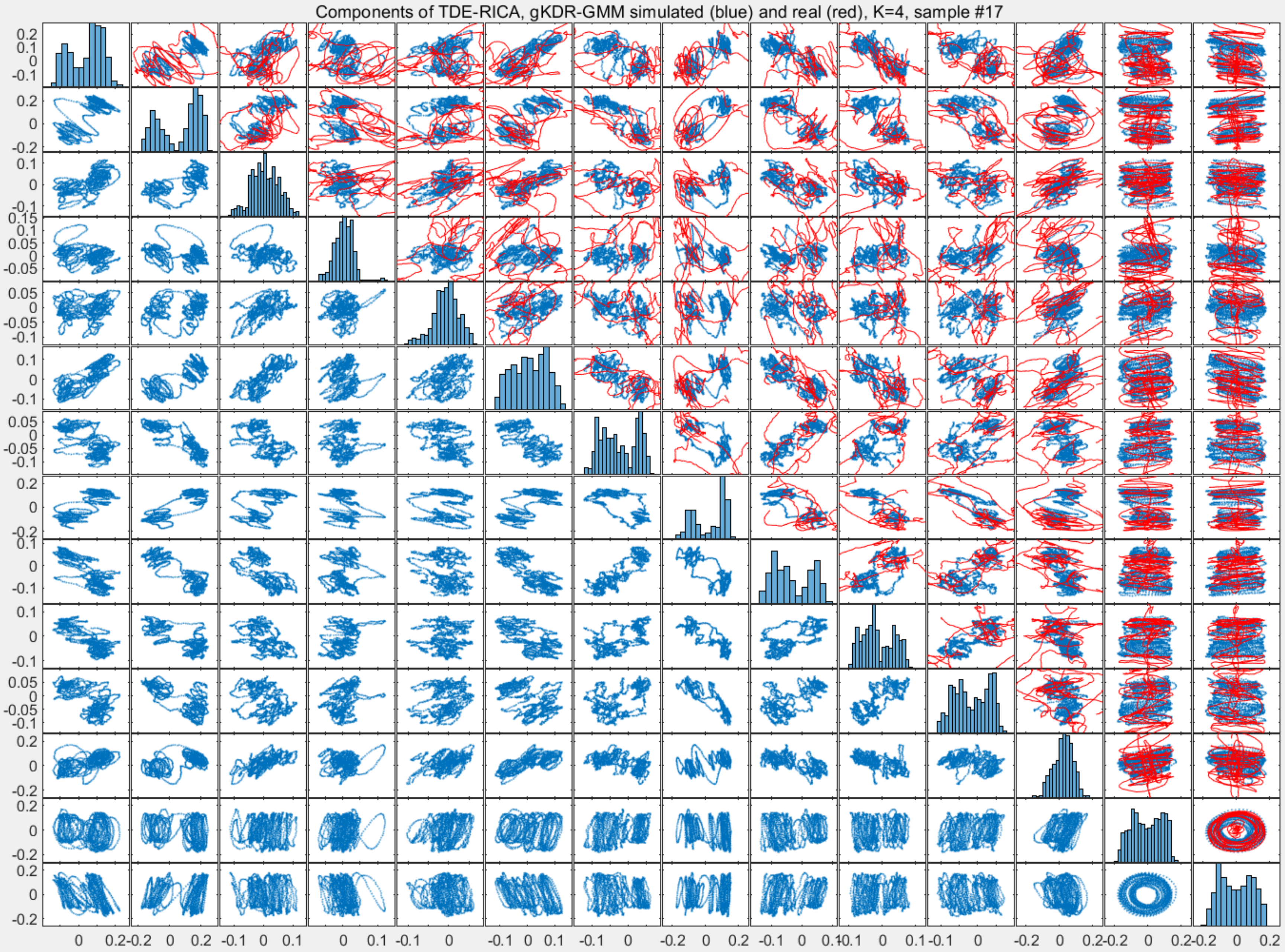

Components of TDE-RICA, gKDR-GMM simulated (blue) and real (red), K=4, sample #17

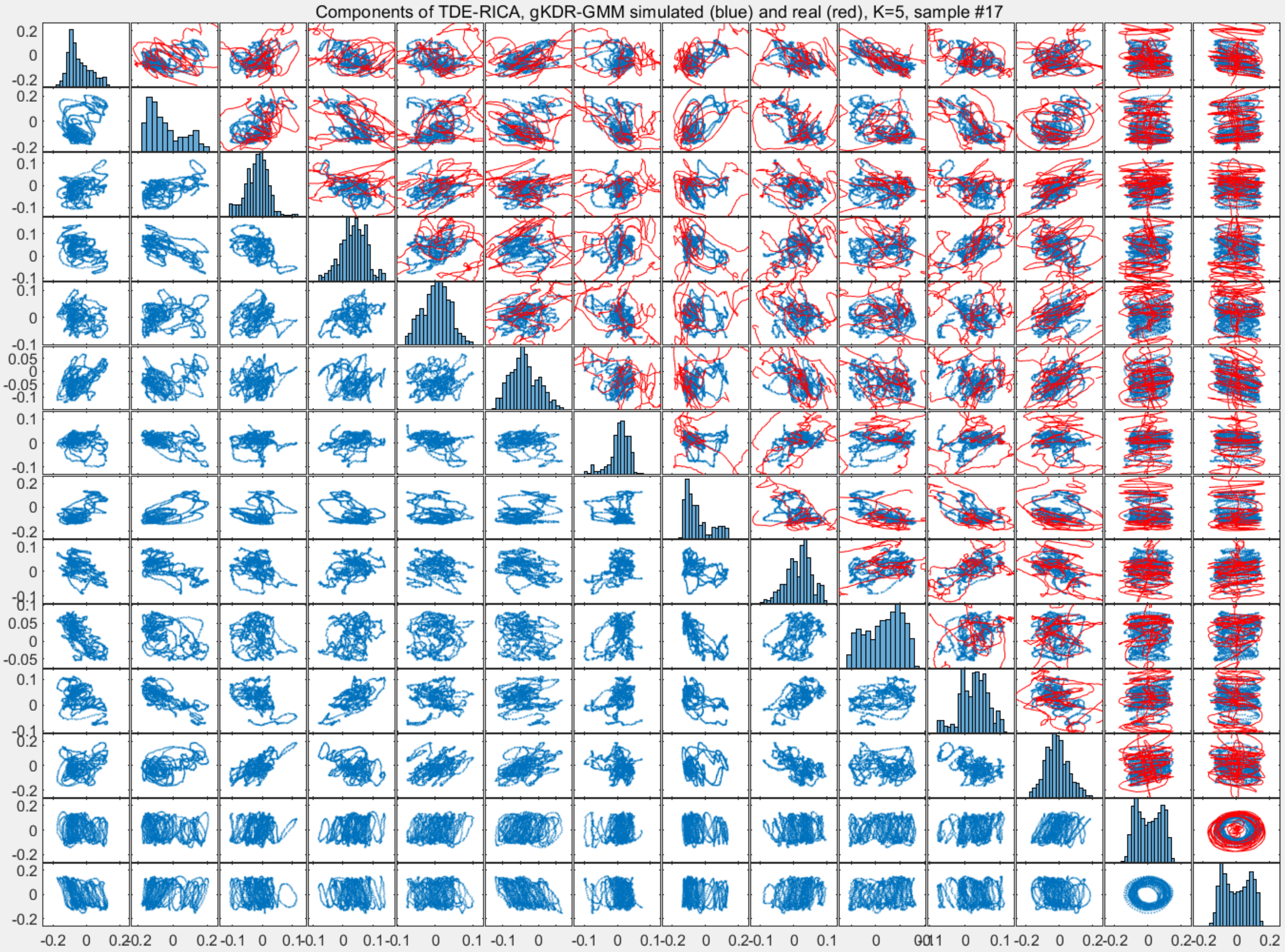
Components of TDE-RICA, gKDR-GMM simulated (blue) and real (red), K=5, sample #17

Components of TDE-RICA, gKDR-GMM simulated (blue) and real (red), K=3, sample #18

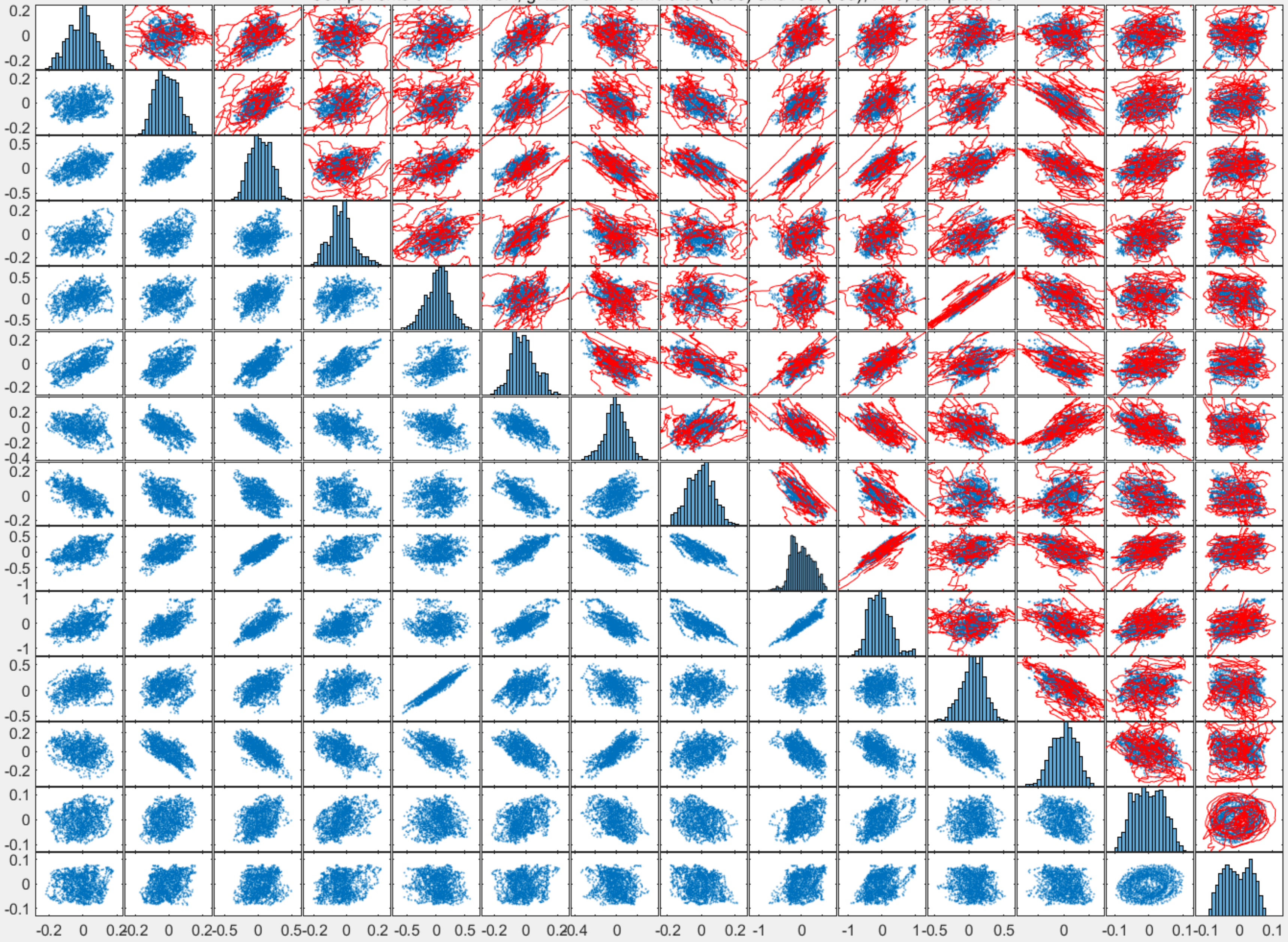

Components of TDE-RICA, gKDR-GMM simulated (blue) and real (red), K=4, sample #18

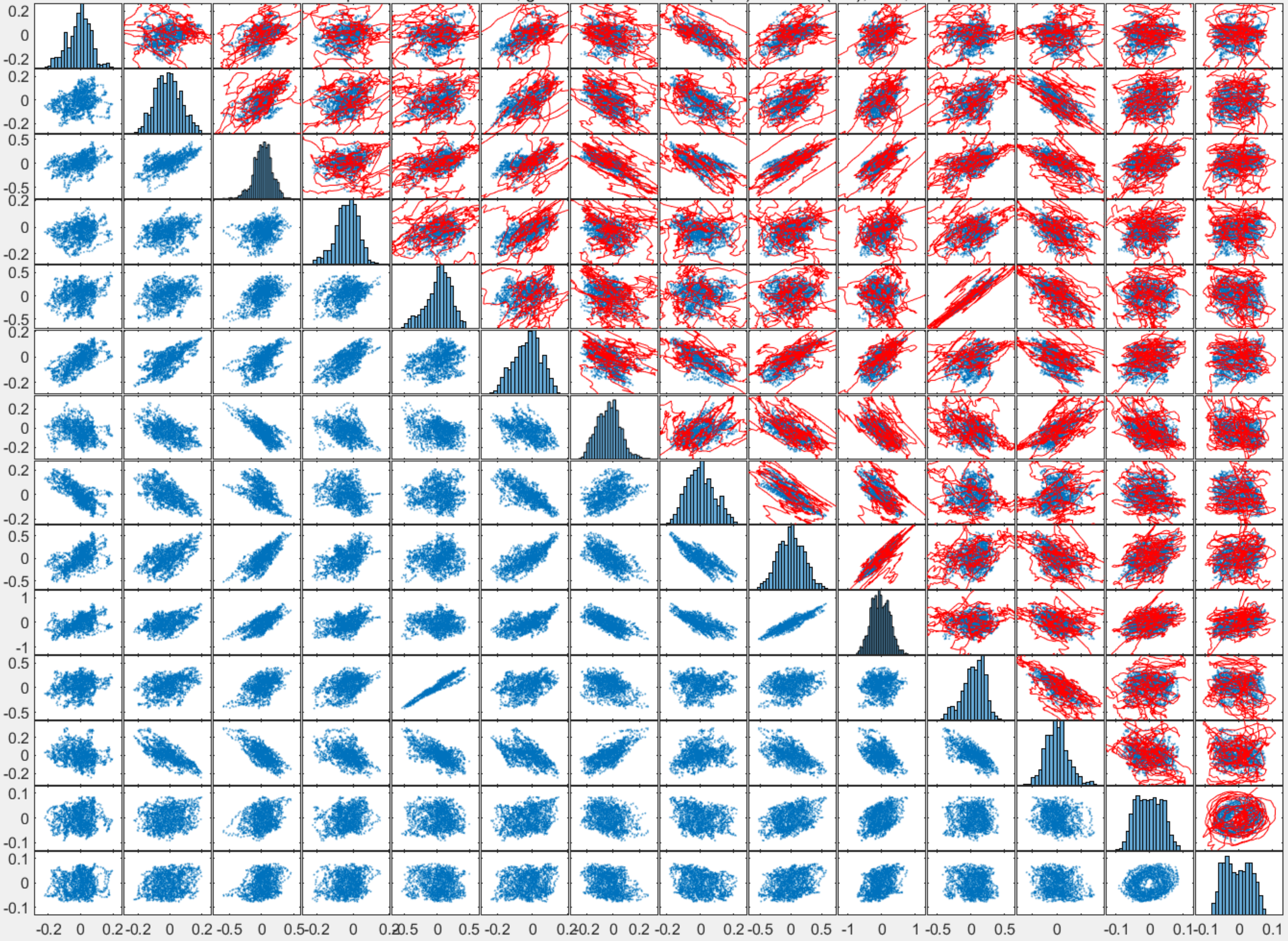

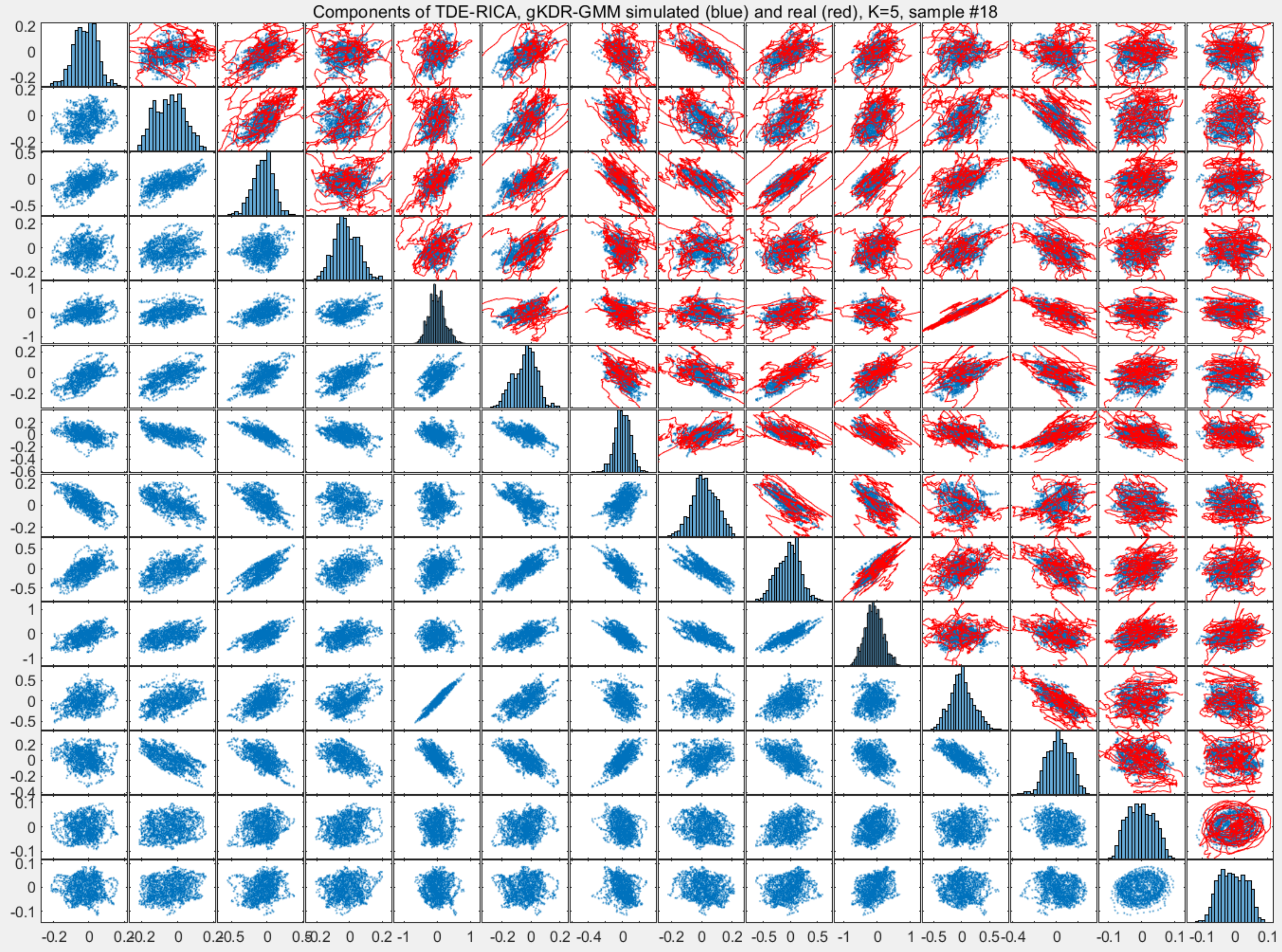
Components of TDE-RICA, gKDR-GMM simulated (blue) and real (red), K=5, sample #18

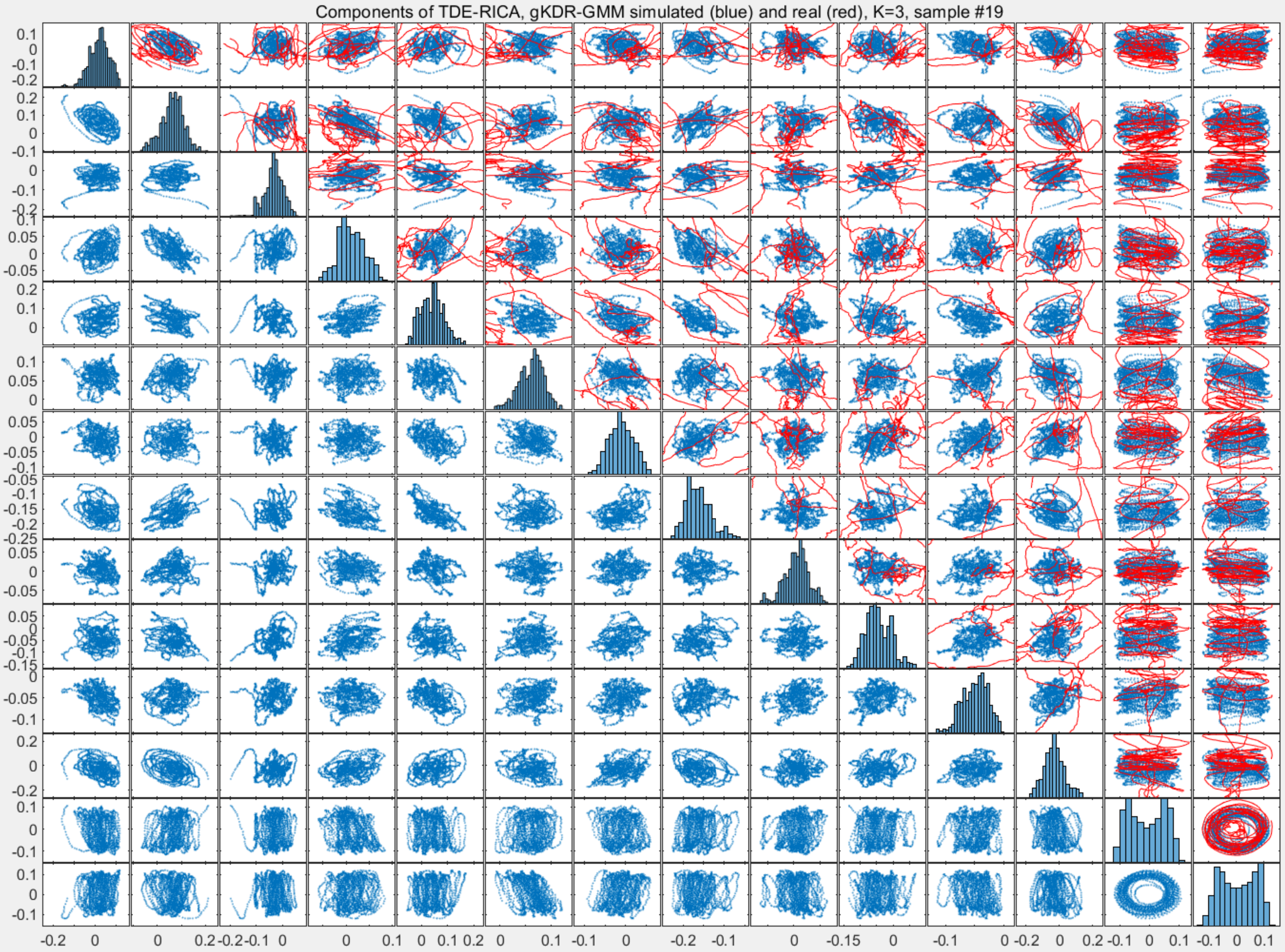

Components of TDE-RICA, gKDR-GMM simulated (blue) and real (red), K=3, sample #19

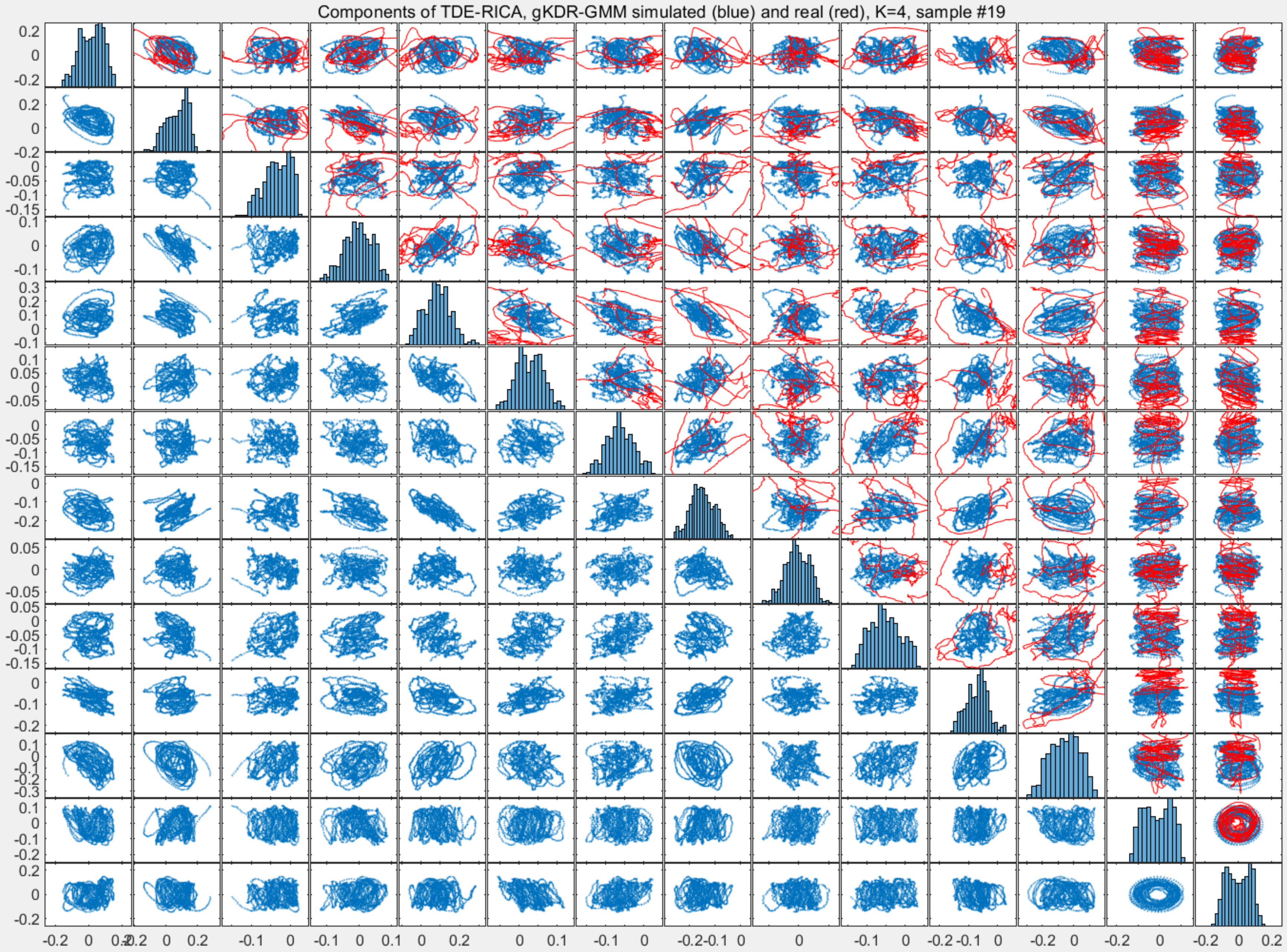
Components of TDE-RICA, gKDR-GMM simulated (blue) and real (red), K=4, sample #19

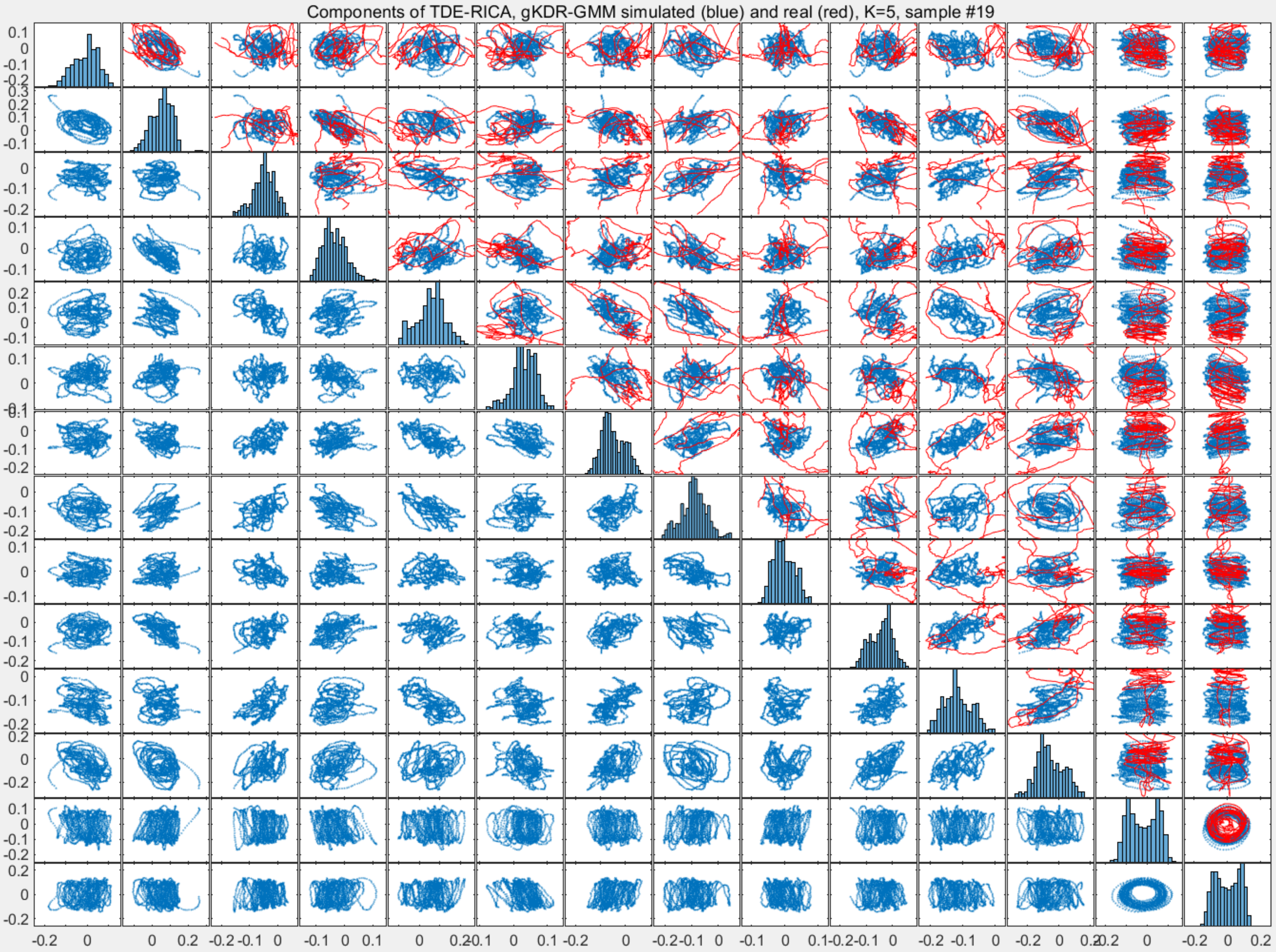
Components of TDE-RICA, gKDR-GMM simulated (blue) and real (red), K=5, sample #19

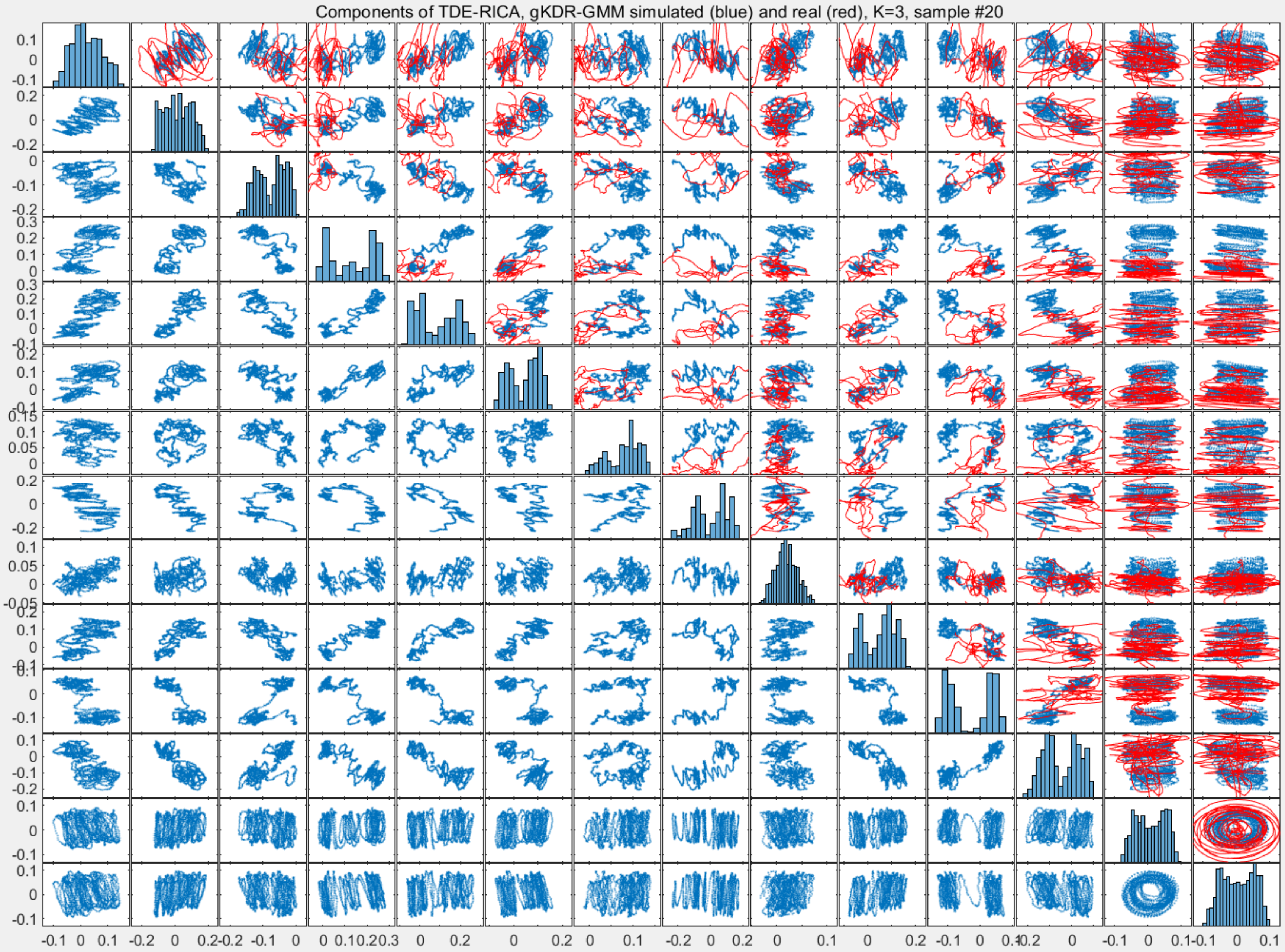

Components of TDE-RICA, gKDR-GMM simulated (blue) and real (red), K=3, sample #20

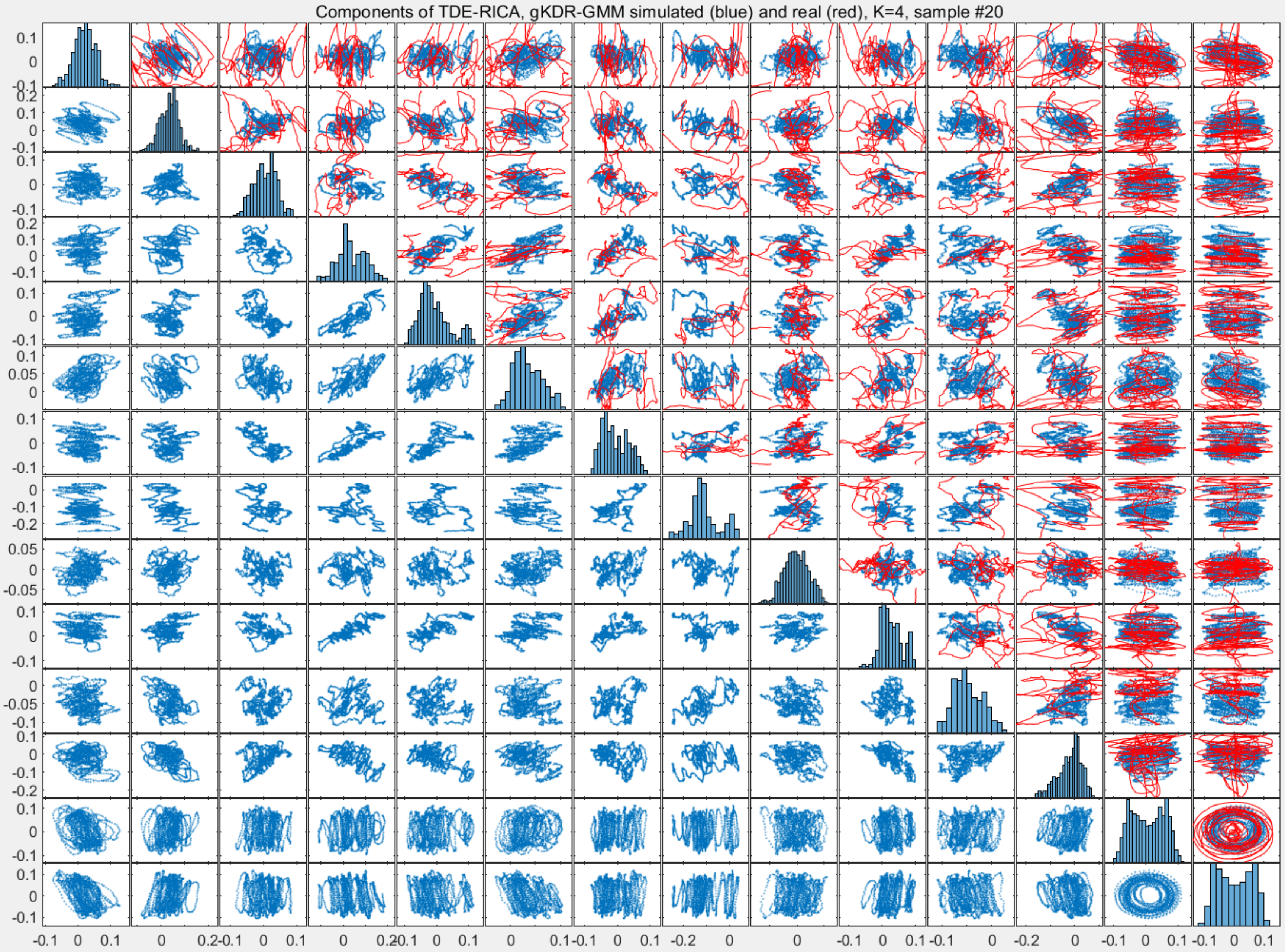
Components of TDE-RICA, gKDR-GMM simulated (blue) and real (red), K=4, sample #20

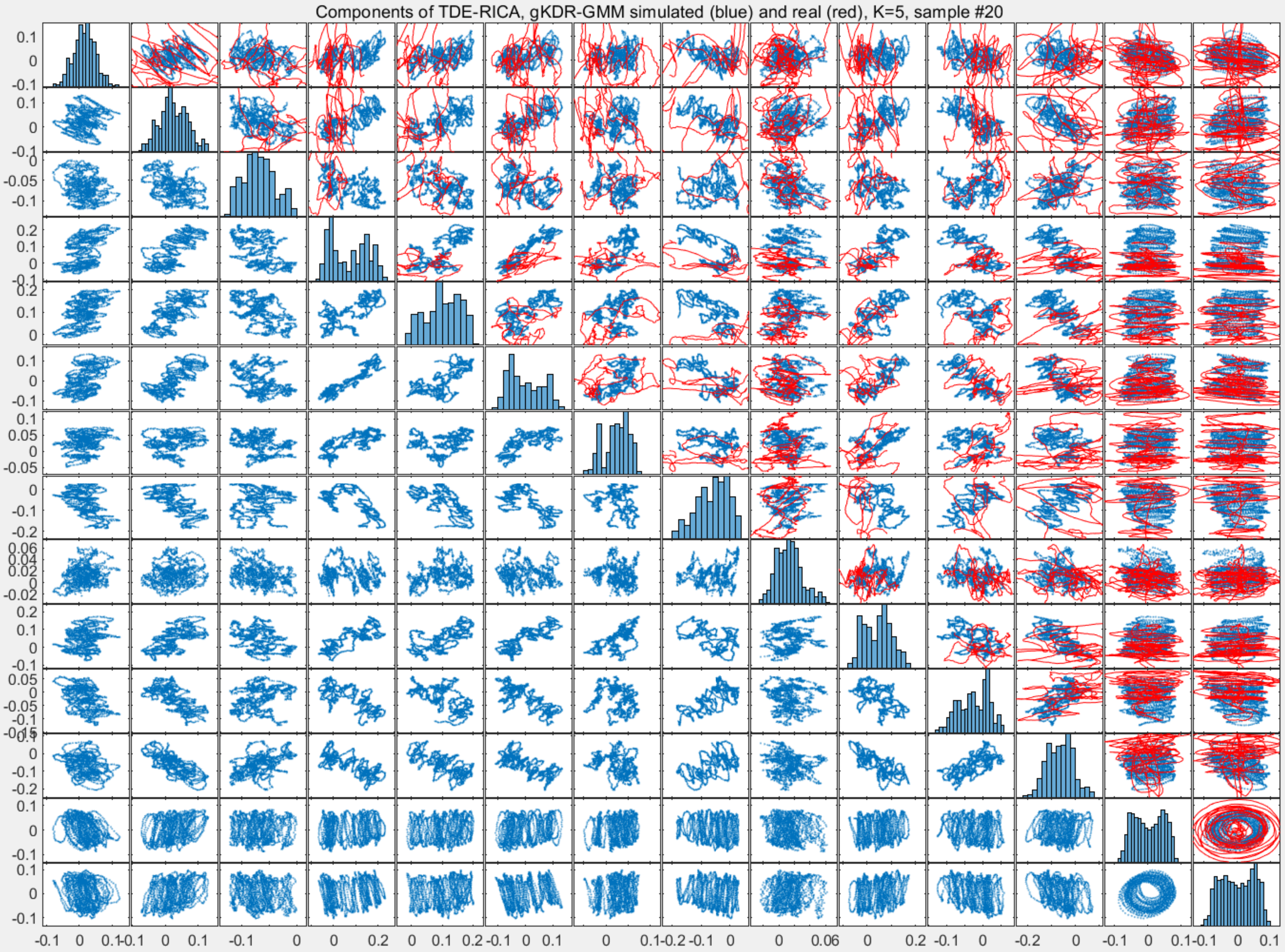

Components of TDE-RICA, gKDR-GMM simulated (blue) and real (red), K=5, sample #20

Components of TDE-RICA, gKDR-GMM simulated (blue) and real (red), K=3, sample #21

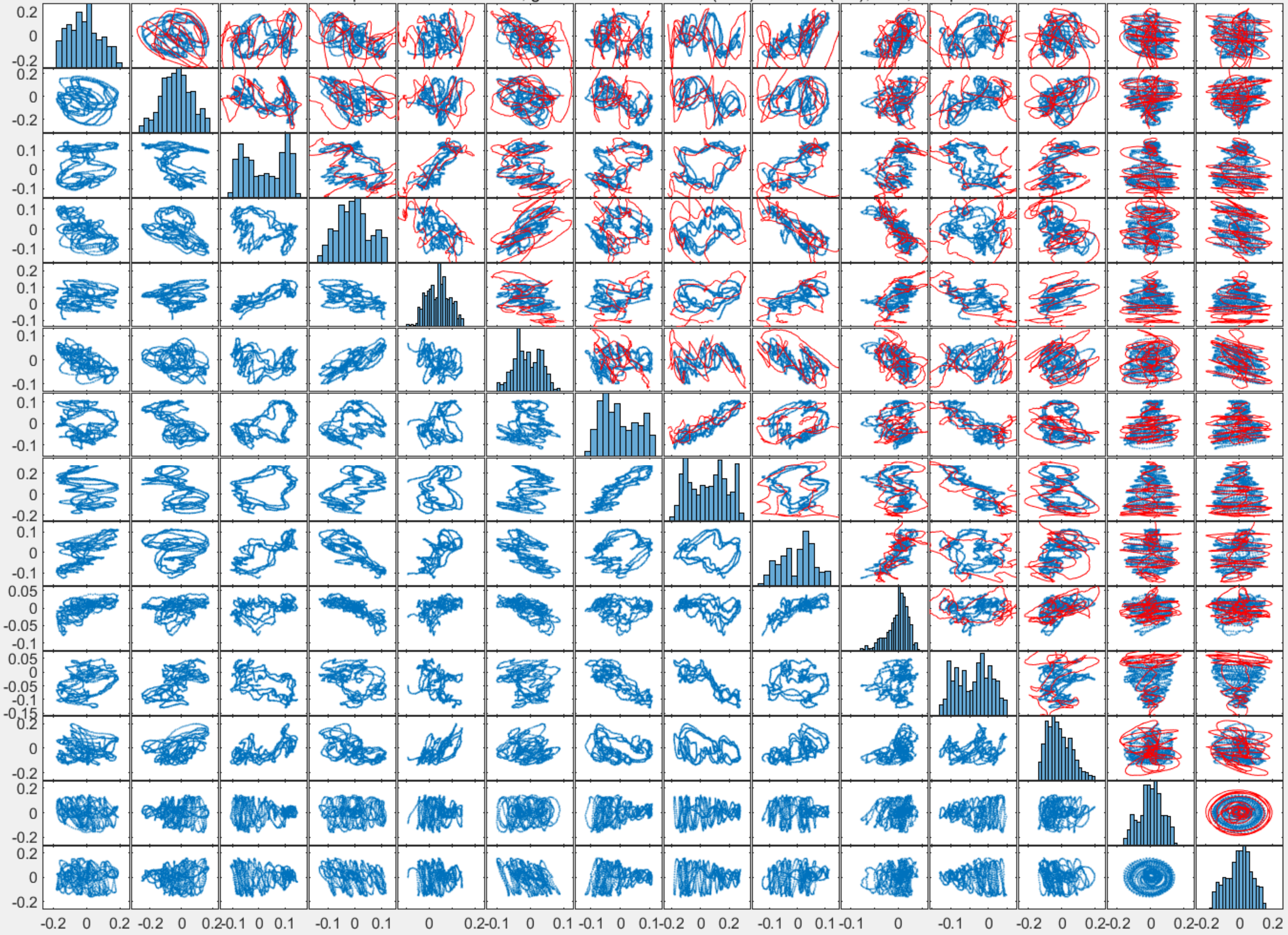

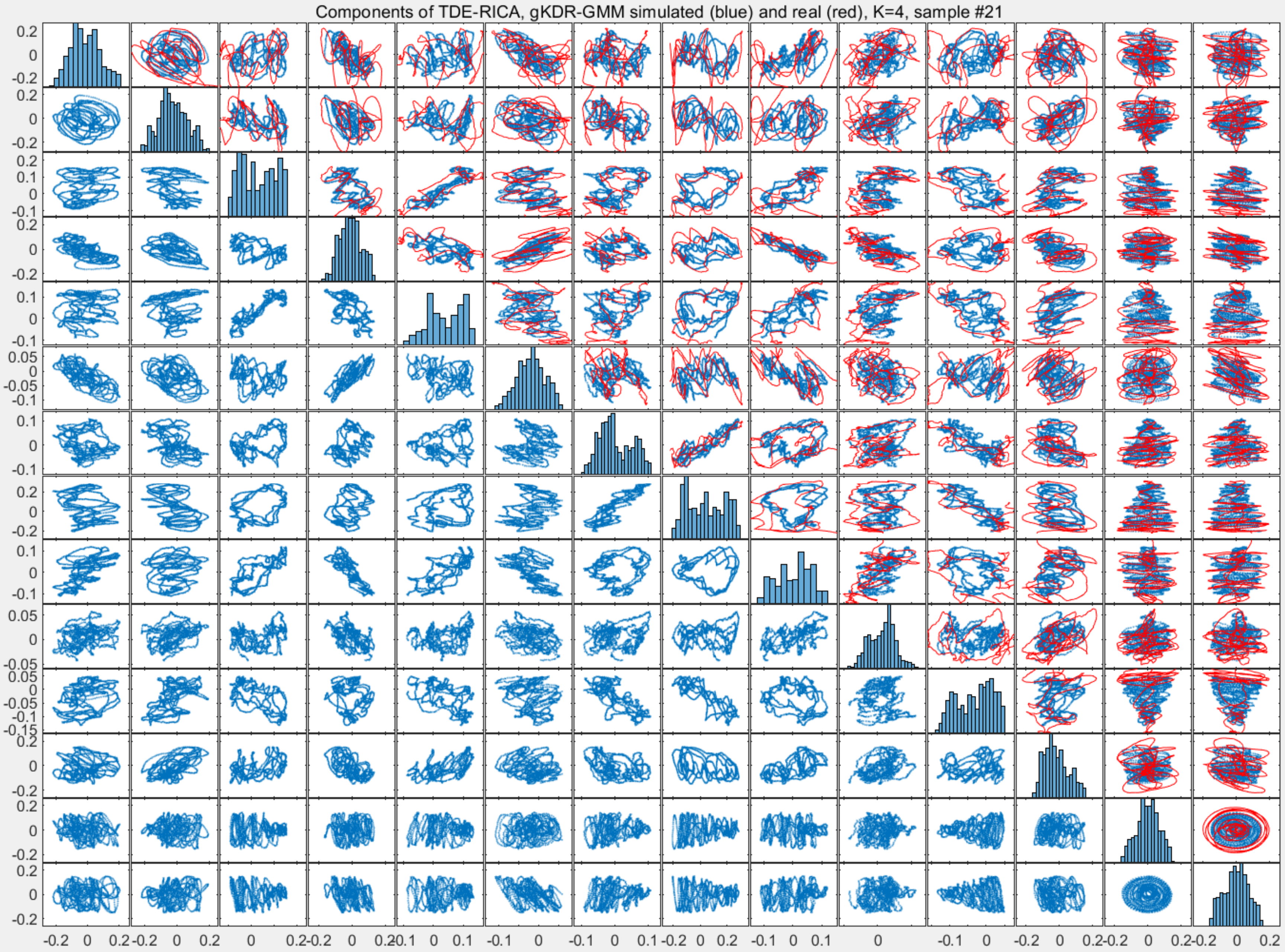

Components of TDE-RICA, gKDR-GMM simulated (blue) and real (red), K=4, sample #21

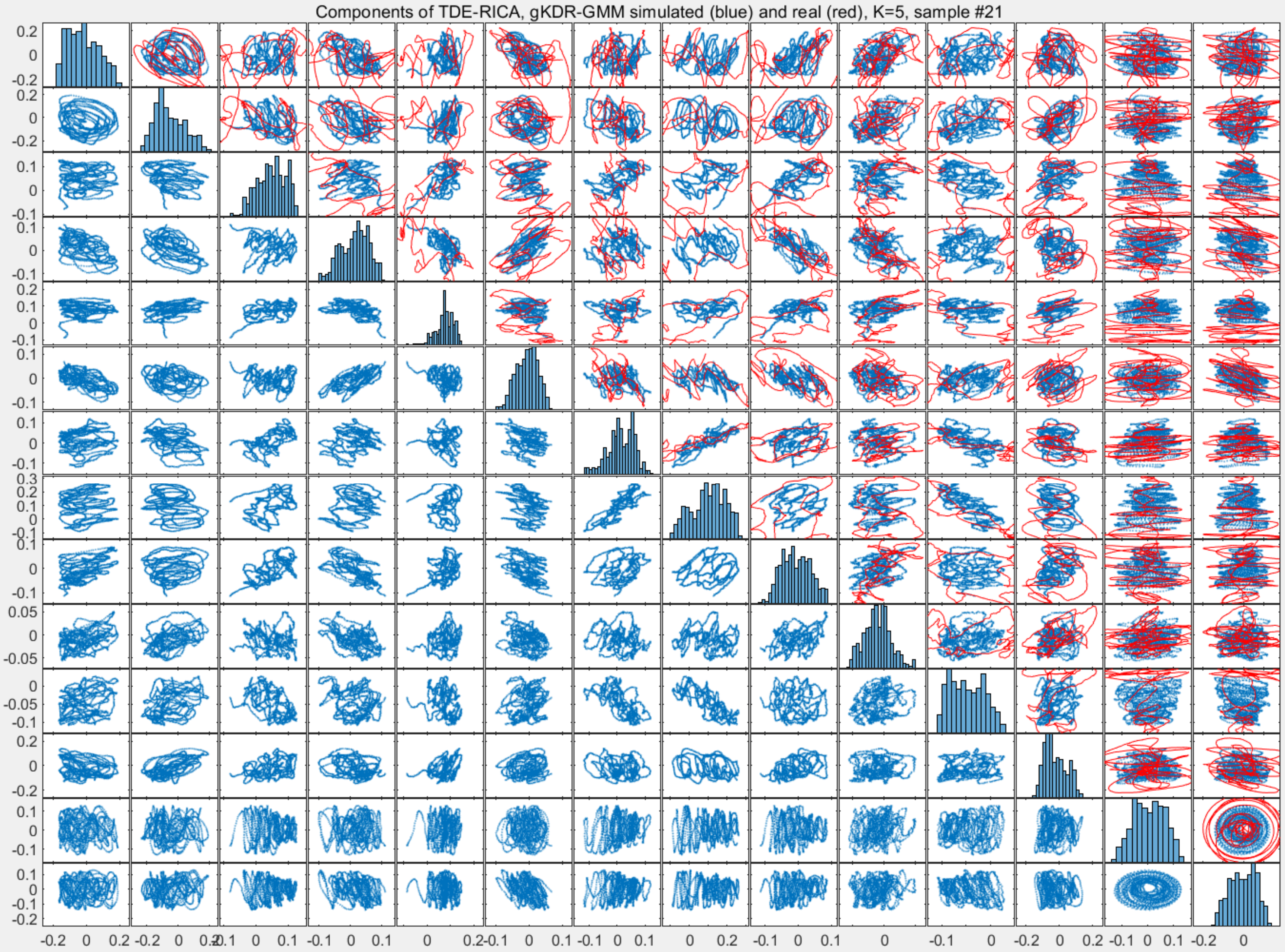
Components of TDE-RICA, gKDR-GMM simulated (blue) and real (red), K=5, sample #21

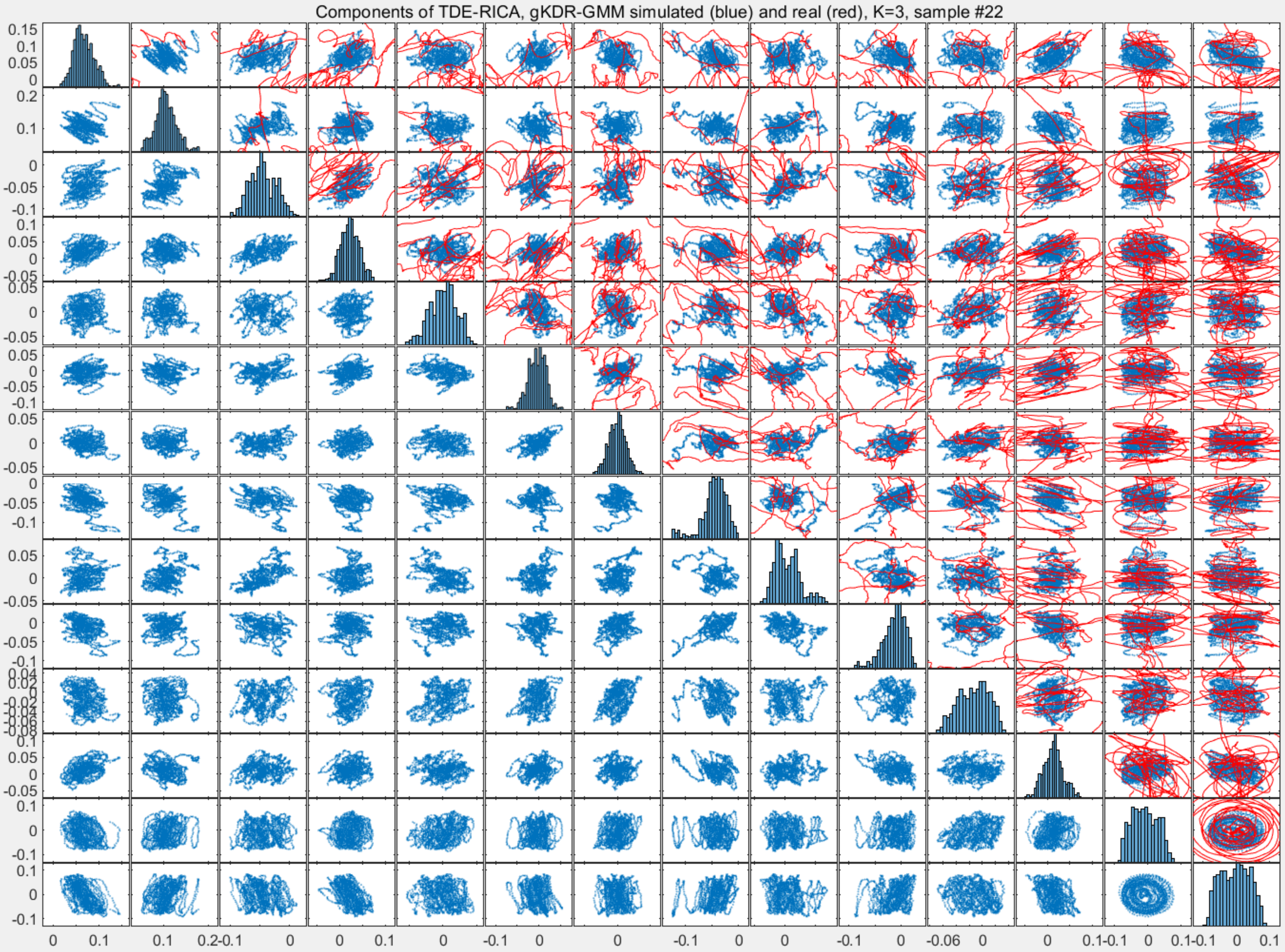
Components of TDE-RICA, gKDR-GMM simulated (blue) and real (red), K=3, sample #22

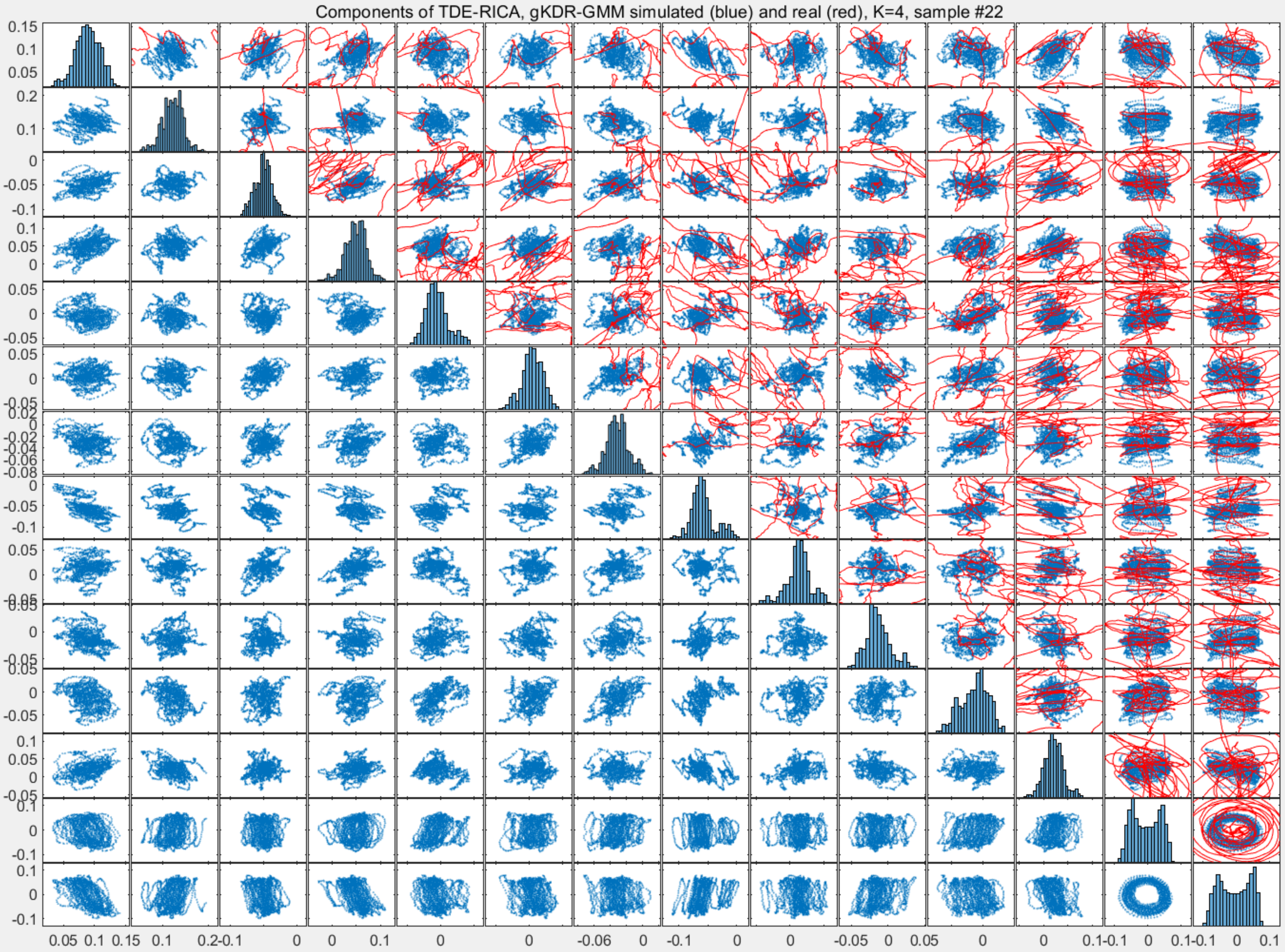
Components of TDE-RICA, gKDR-GMM simulated (blue) and real (red), K=4, sample #22

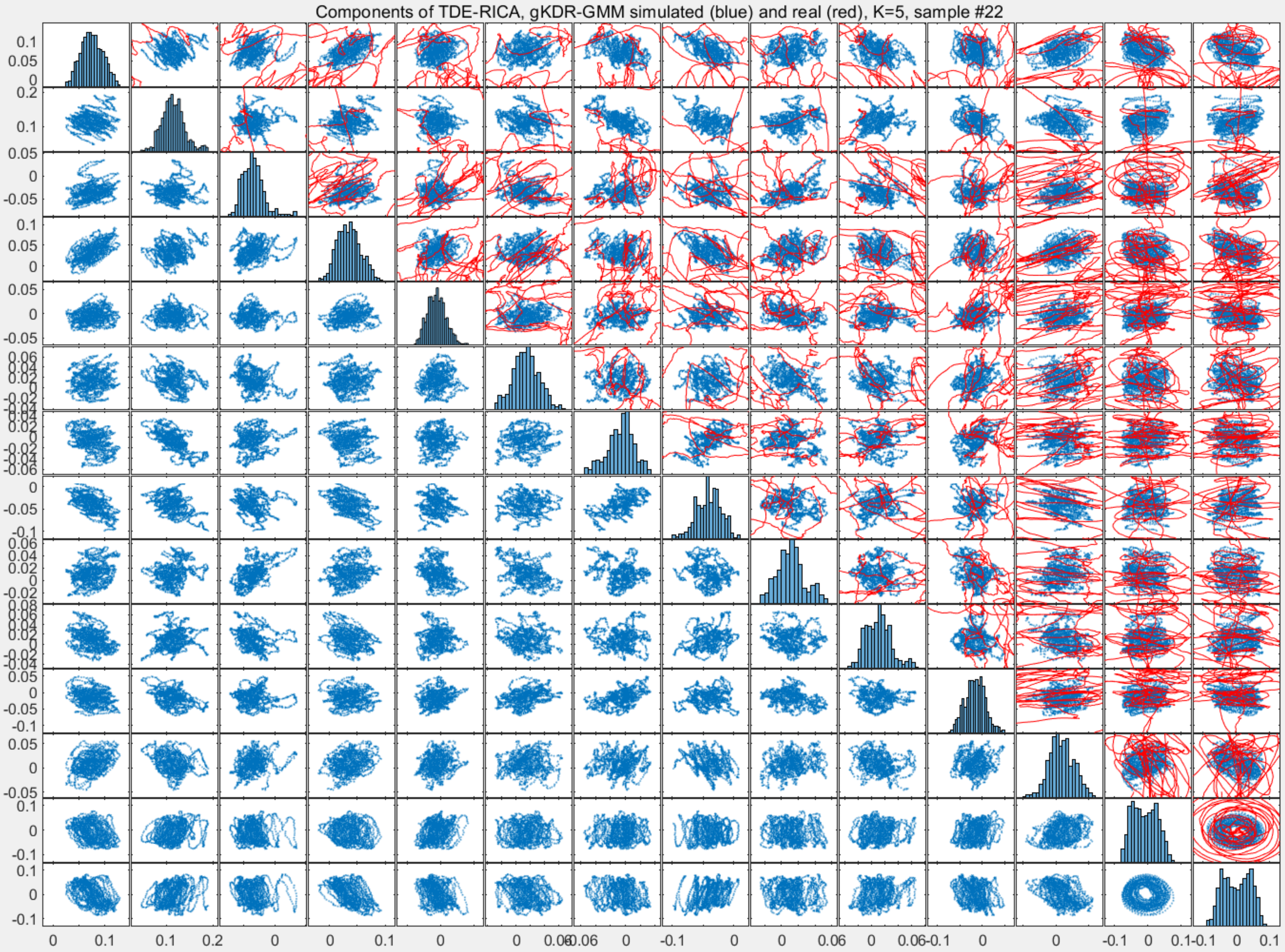
Components of TDE-RICA, gKDR-GMM simulated (blue) and real (red), K=5, sample #22

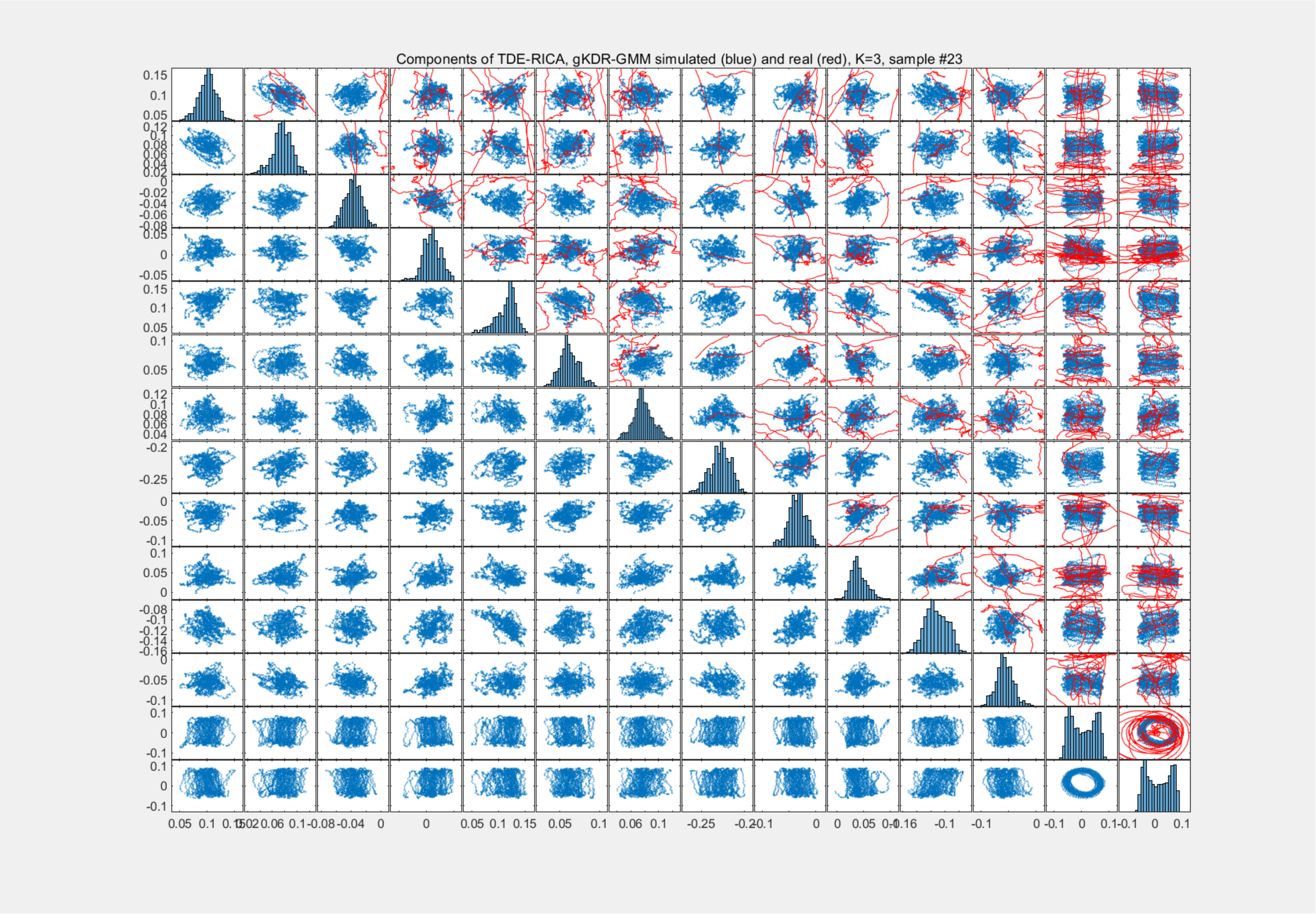
Components of TDE-RICA, gKDR-GMM simulated (blue) and real (red), K=3, sample #23

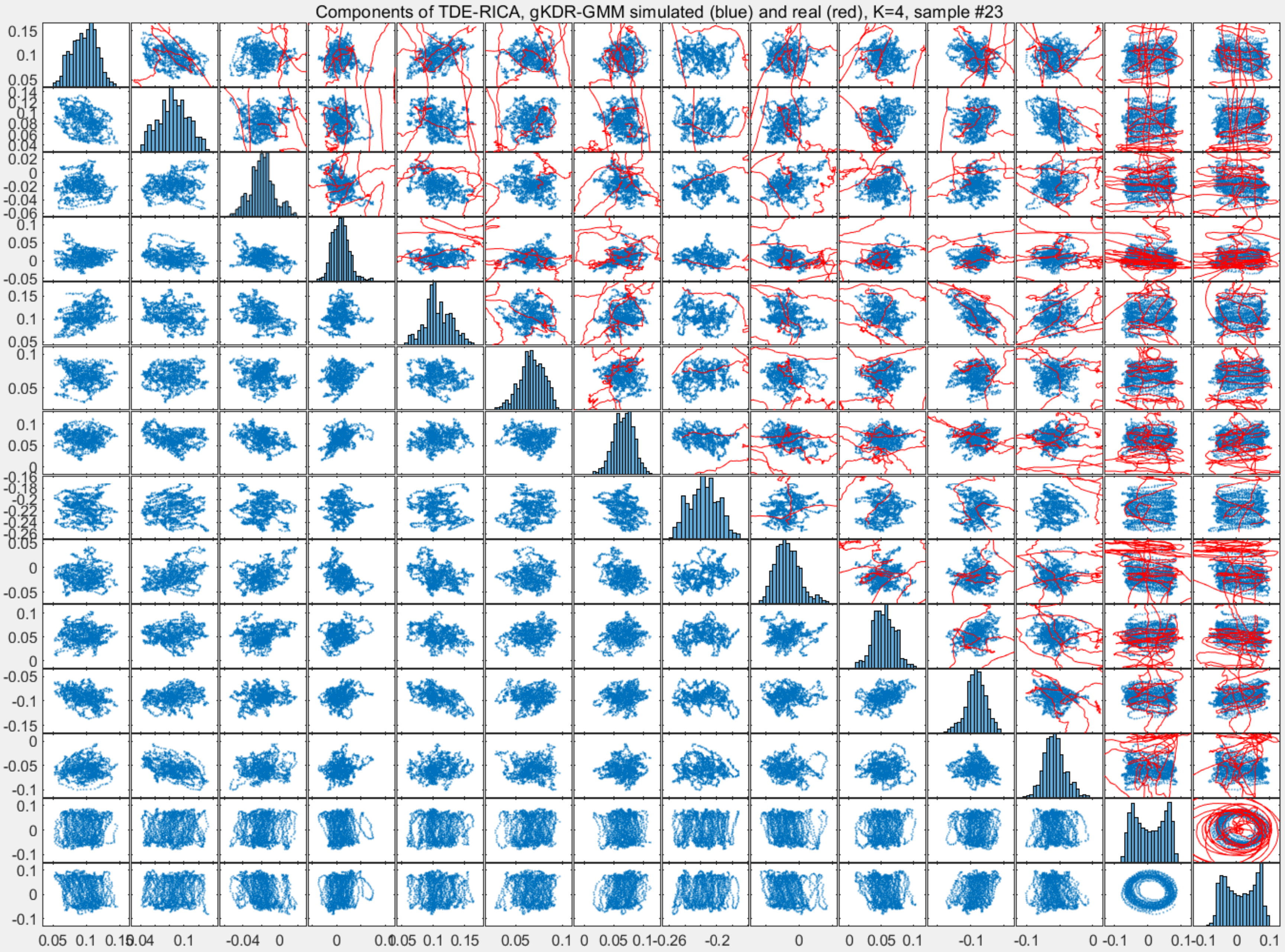
Components of TDE-RICA, gKDR-GMM simulated (blue) and real (red), K=4, sample #23

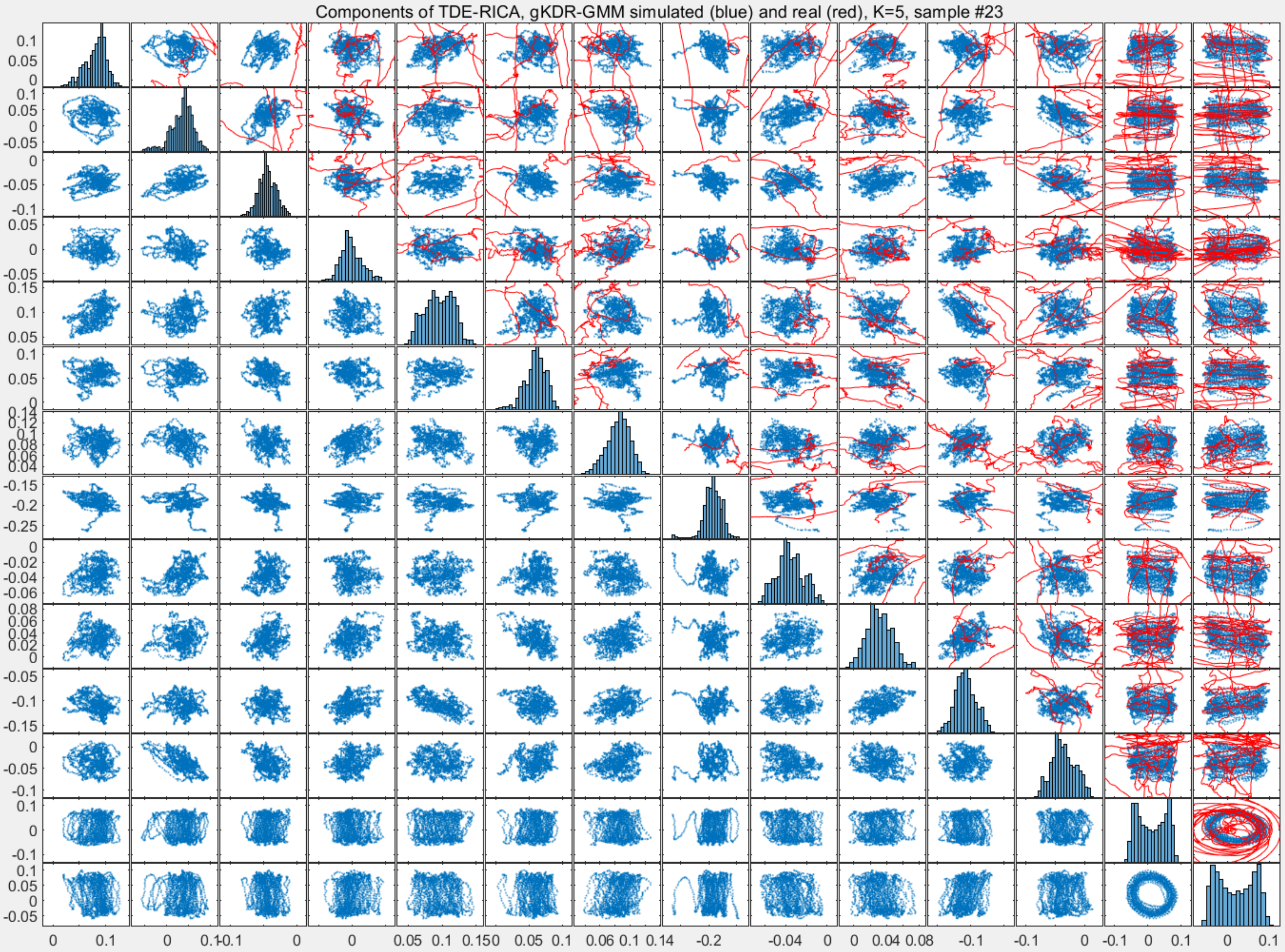

Components of TDE-RICA, gKDR-GMM simulated (blue) and real (red), K=5, sample #23

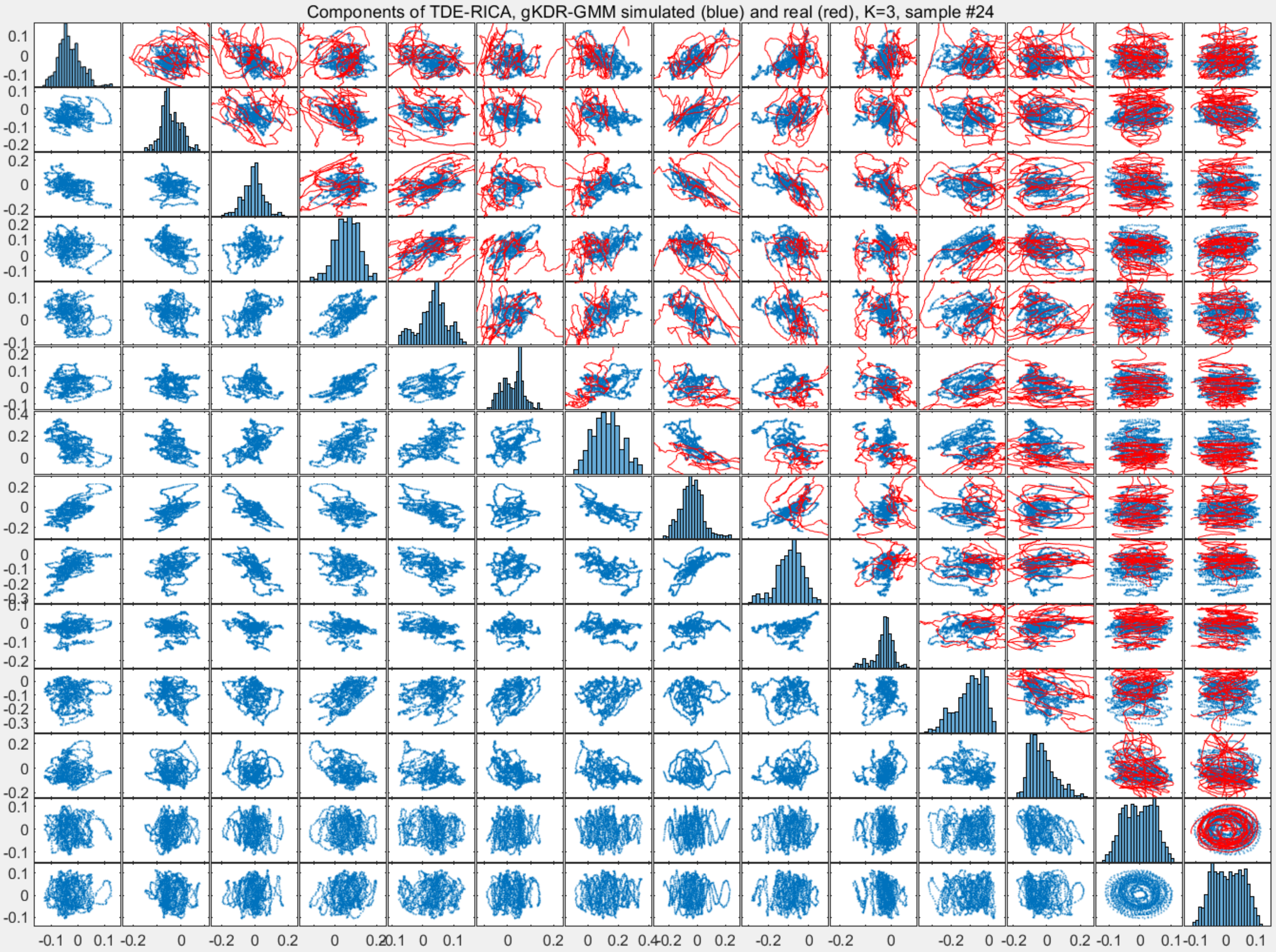

Components of TDE-RICA, gKDR-GMM simulated (blue) and real (red), K=3, sample #24

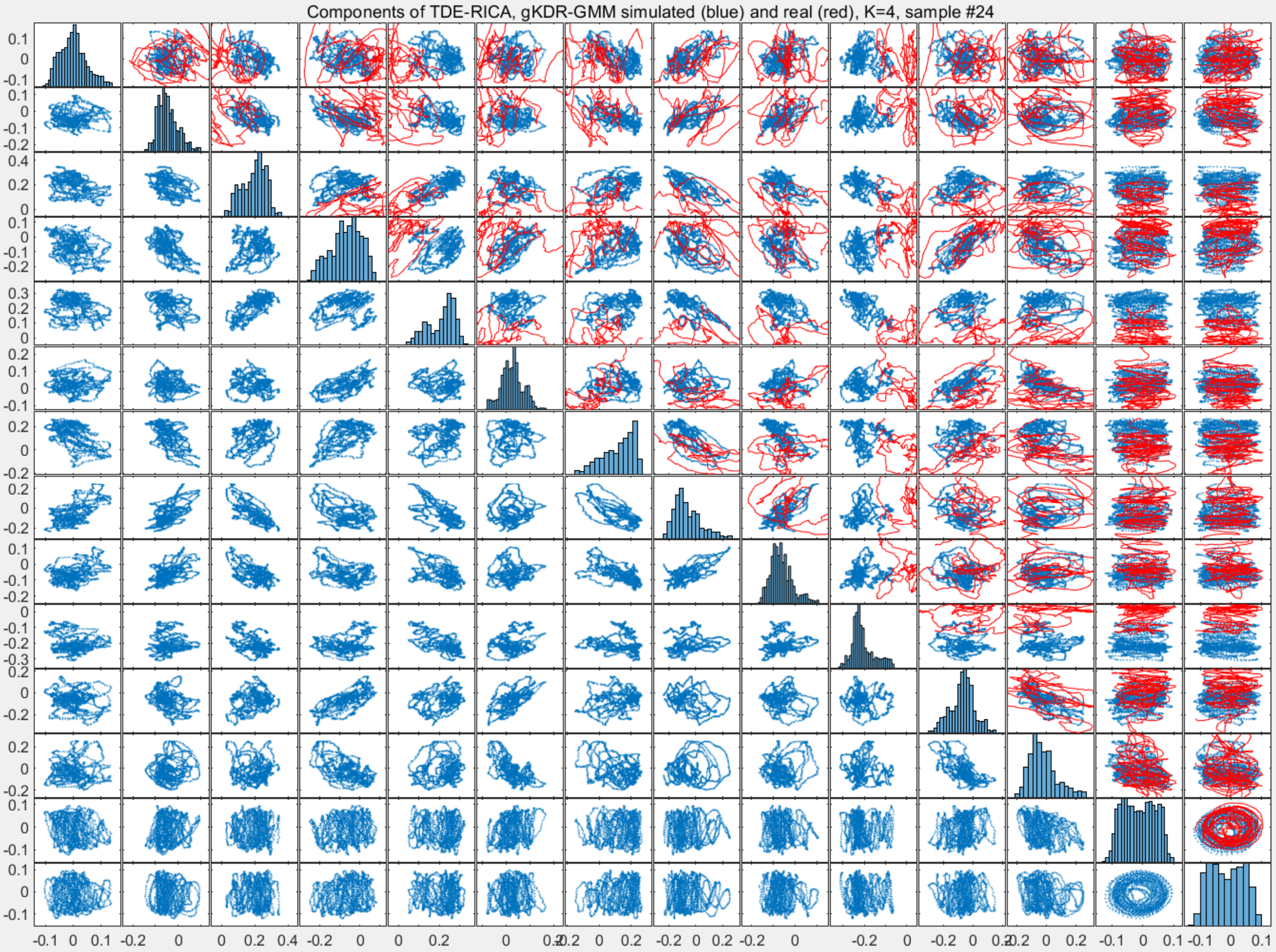
Components of TDE-RICA, gKDR-GMM simulated (blue) and real (red), K=4, sample #24

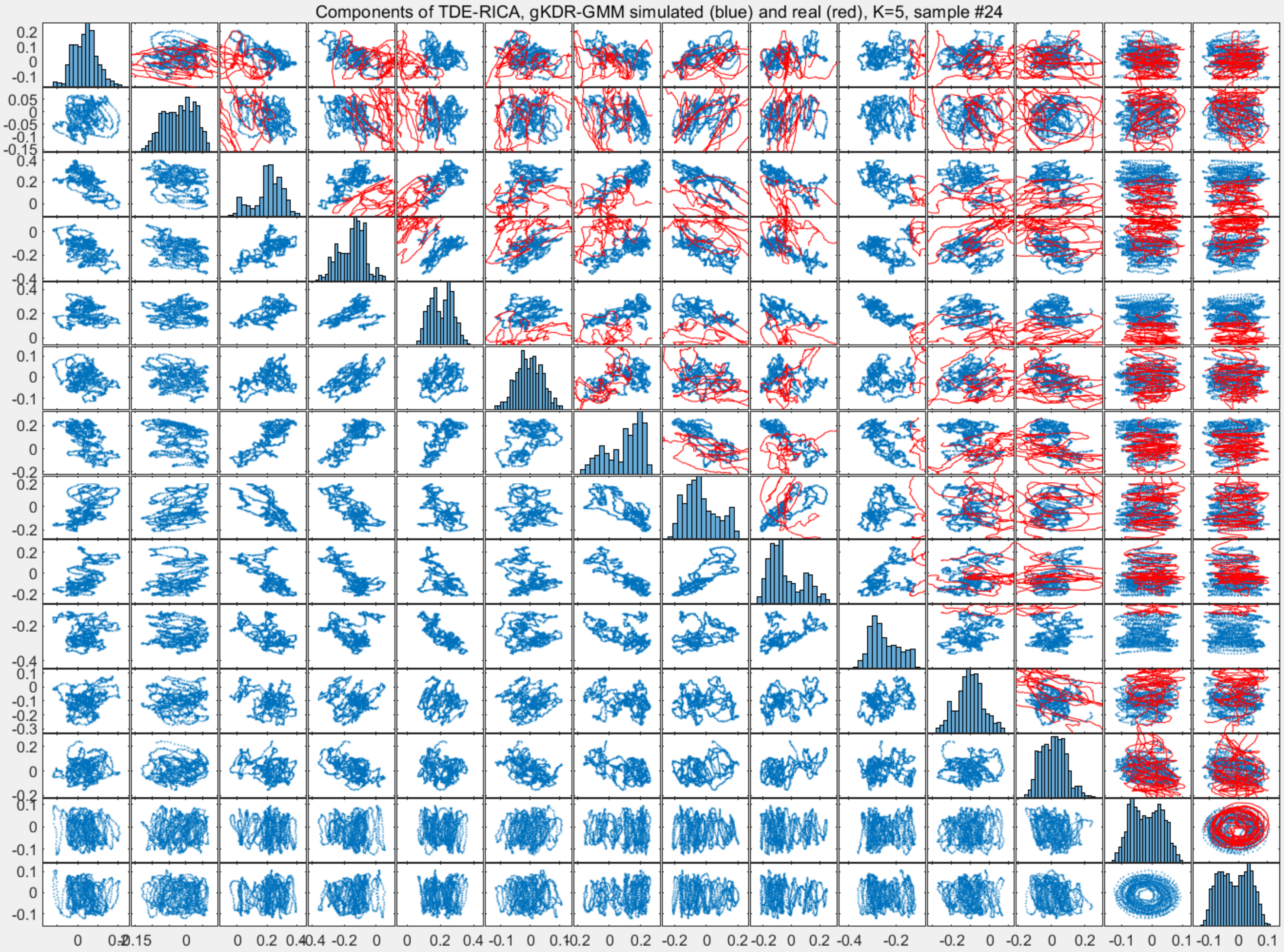
Components of TDE-RICA, gKDR-GMM simulated (blue) and real (red), K=5, sample #24
