## Supplementary File 9A for "Deducing ensemble dynamics and information flow from the whole-brain imaging data"

### Sample1

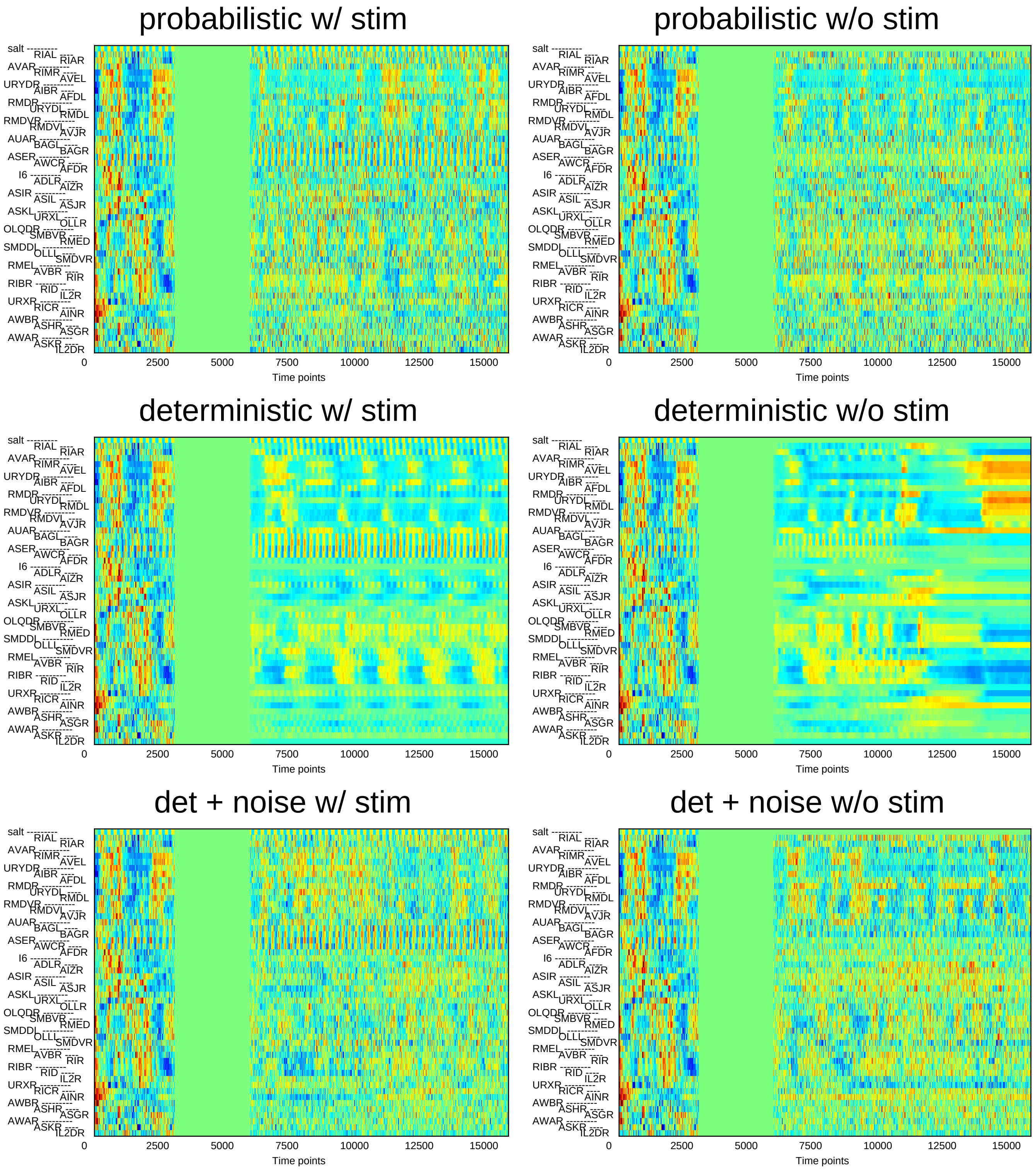
probabilistic w/ stim
probabilistic w/o stim
deterministic w/ stim
deterministic w/o stim
det + noise w/ stim
det + noise w/o stim
salt
RIAL
RIAR
AVAR
RIMR
AVEL
URYDR
AIBR
AFDL
RMDR
URYDL
RMDL
RMDVR
RMDVL
AVJR
AUAR
BAGL
BAGR
ASER
AWCR
AFDR
I6
ADLR
AIZR
ASIR
ASIL
ASJR
ASKL
URXL
OLLR
OLQDR
SMBVR
RMED
SMDDL
OLLL
SMDVR
RMEL
AVBR
RIR
RIBR
RID
IL2R
URXR
RICR
AINR
AWBR
ASHR
ASGR
AWAR
ASKR
IL2DR
0
2500
5000
7500
10000
12500
15000
Time points

### Sample2

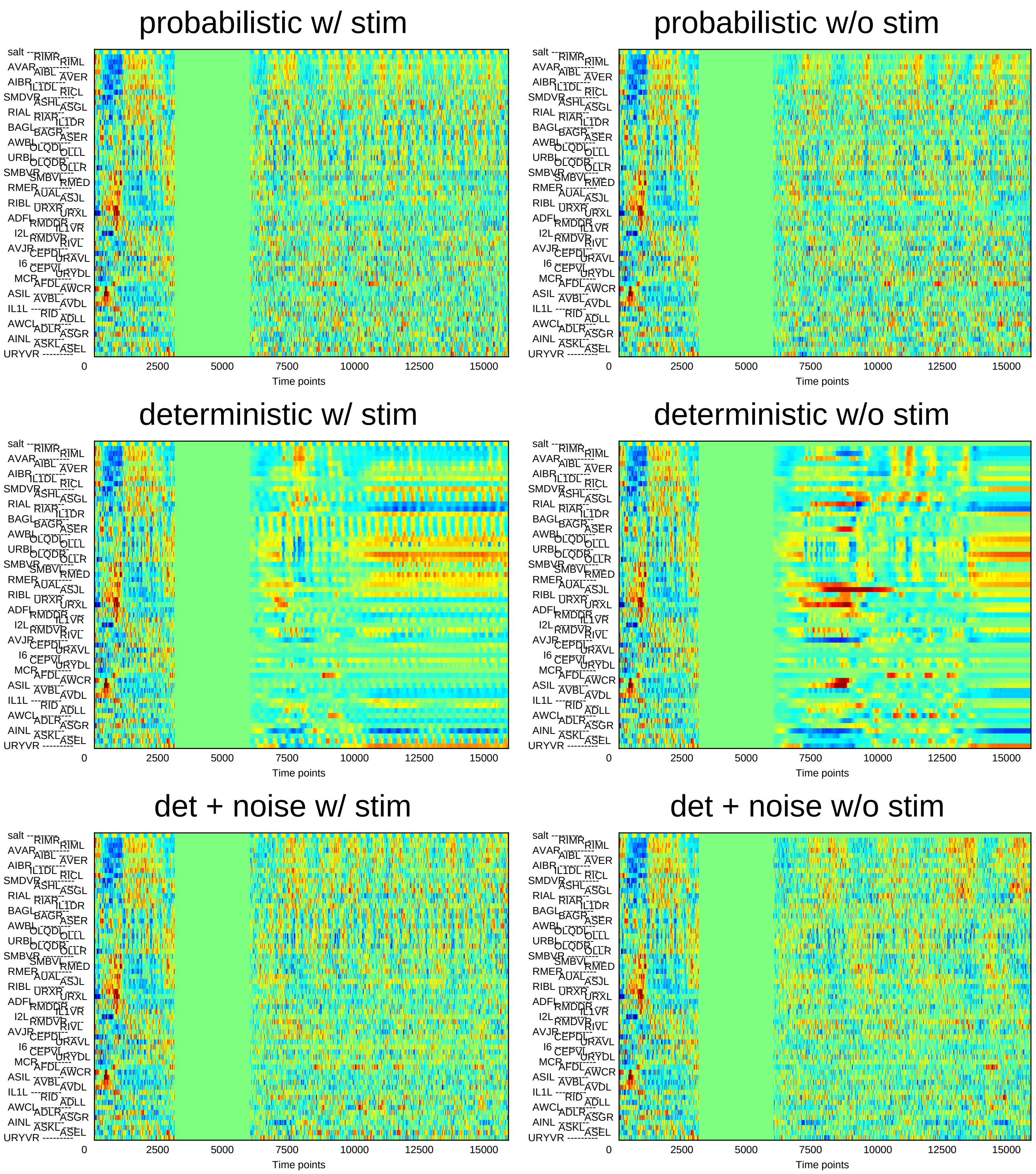
probabilistic w/ stim
probabilistic w/o stim
deterministic w/ stim
deterministic w/o stim
det + noise w/ stim
det + noise w/o stim
Time points
0
2500
5000
7500
10000
12500
15000
salt
RIMR
RIML
AVAR
AIBL
AVER
AIBR
IL1DL
RICL
SMDVR
ASHL
ASGL
RIAL
RIAR
IL1DR
BAGL
BAGR
ASER
AWBL
OLQDL
OLLL
URBL
OLQDR
OLLR
SMBVR
SMBVL
RMED
RMER
AUAL
ASJL
RIBL
URXR
URXL
ADFL
RMDDR
IL1VR
I2L
RMDVR
RIVL
AVJR
CEPDL
URAVL
I6
CEPVL
URYDL
MCR
AFDL
AWCR
ASIL
AVBL
AVDL
IL1L
RID
ADLL
AWCL
ADLR
ASGR
AINL
ASKL
ASEL
URYVR

### Sample3

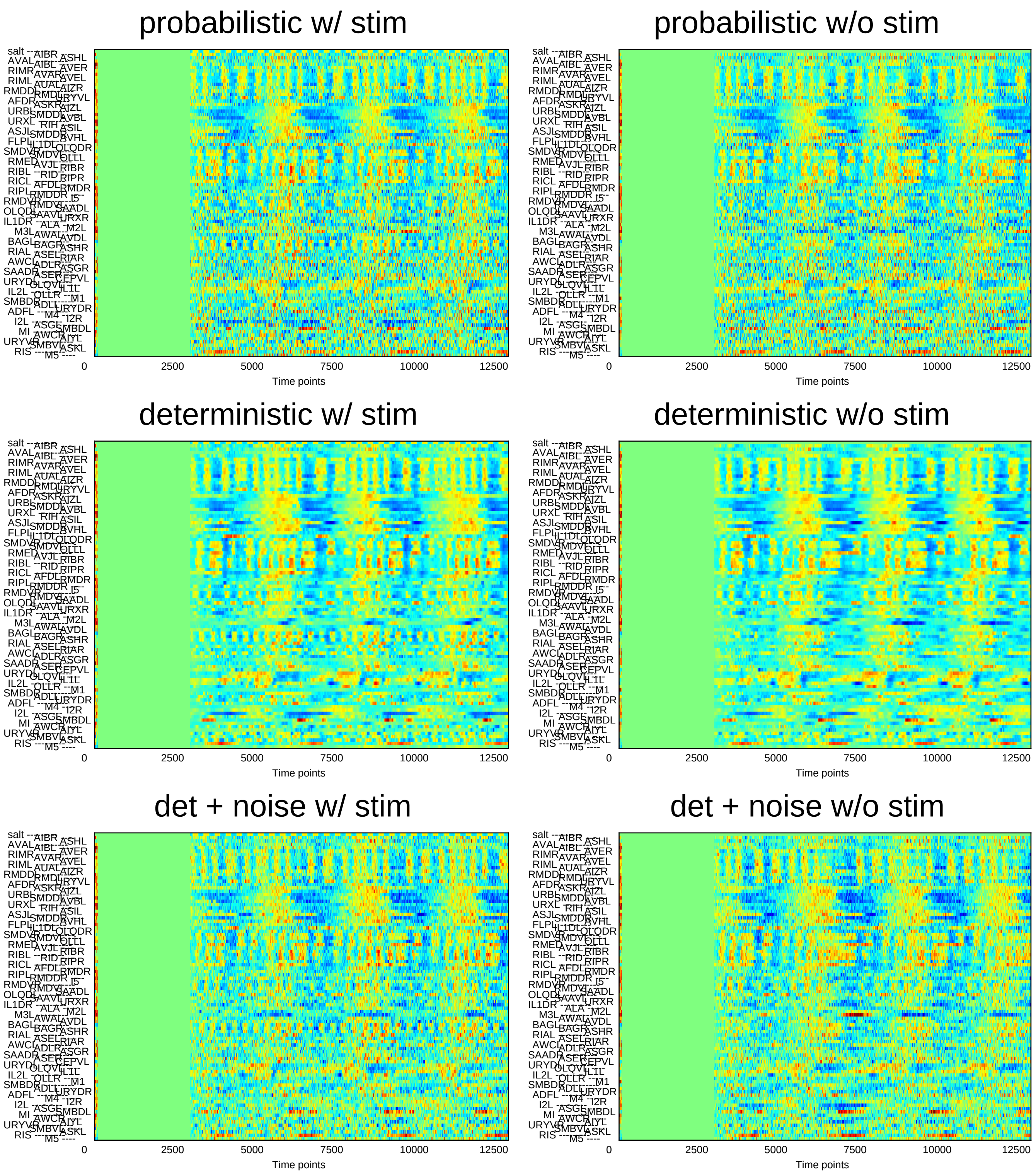
probabilistic w/ stim
probabilistic w/o stim
deterministic w/ stim
deterministic w/o stim
det + noise w/ stim
det + noise w/o stim
0
2500
5000
7500
10000
12500
Time points

### Sample4

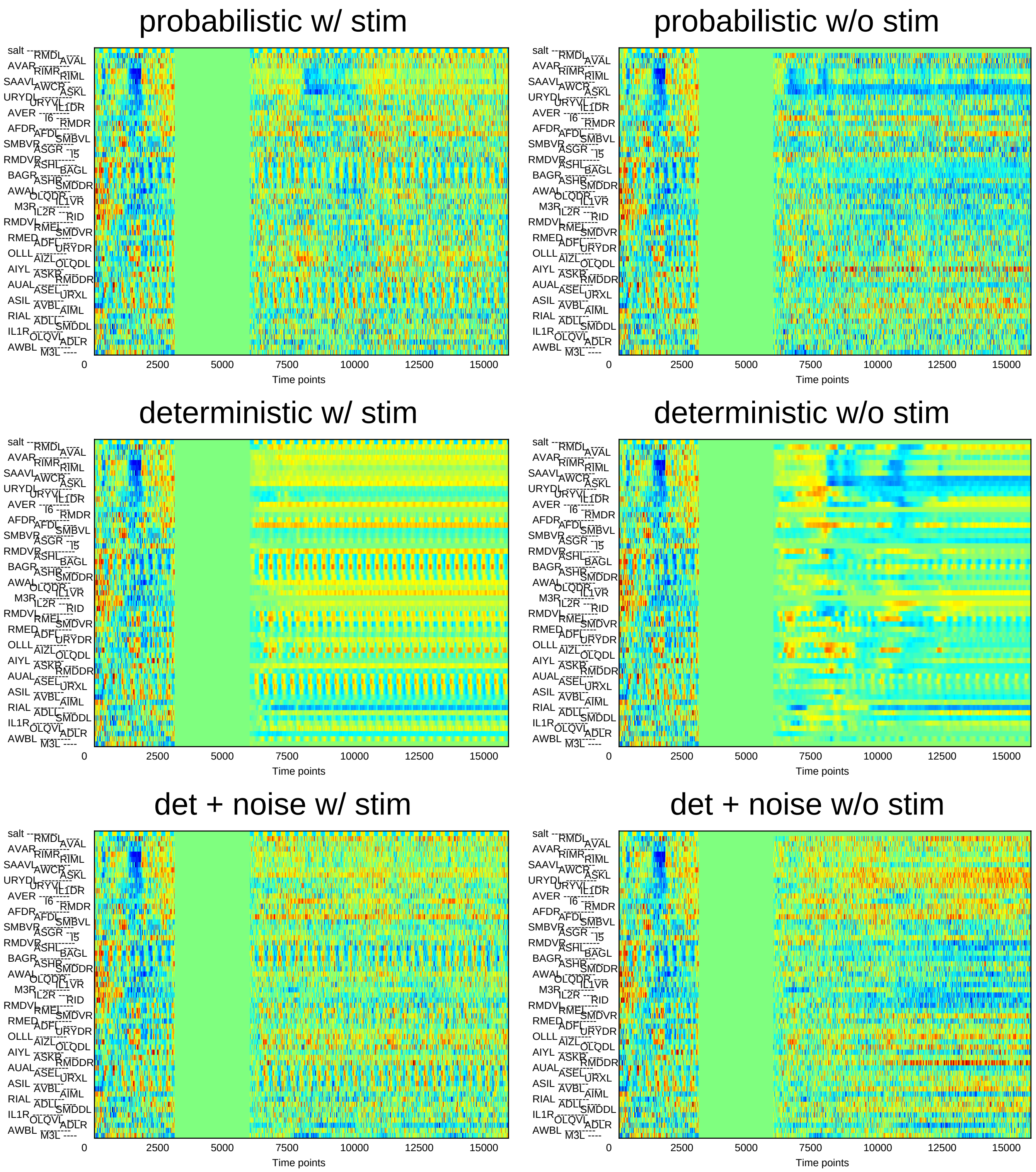
probabilistic w/ stim
probabilistic w/o stim
deterministic w/ stim
deterministic w/o stim
det + noise w/ stim
det + noise w/o stim
salt RMDL AVAL AVAR RIMR RIML SAAVL AWCR ASKL URYDL URYVL IL1DR AVER I6 RMDR AFDR AFDL SMBVL SMBVR ASGR I5 RMDVR ASHL BAGL BAGR ASHR SMDDR AWAL OLQDR IL1VR M3R IL2R RID RMDVL RMEL SMDVR RMED ADFL URYDR OLLL AIZL OLQDL AIYL ASKR RMDDR AUAL ASEL URXL ASIL AVBL AIML RIAL ADLL SMDDL IL1R OLQVL ADLR AWBL M3L
0 2500 5000 7500 10000 12500 15000
Time points

### Sample5

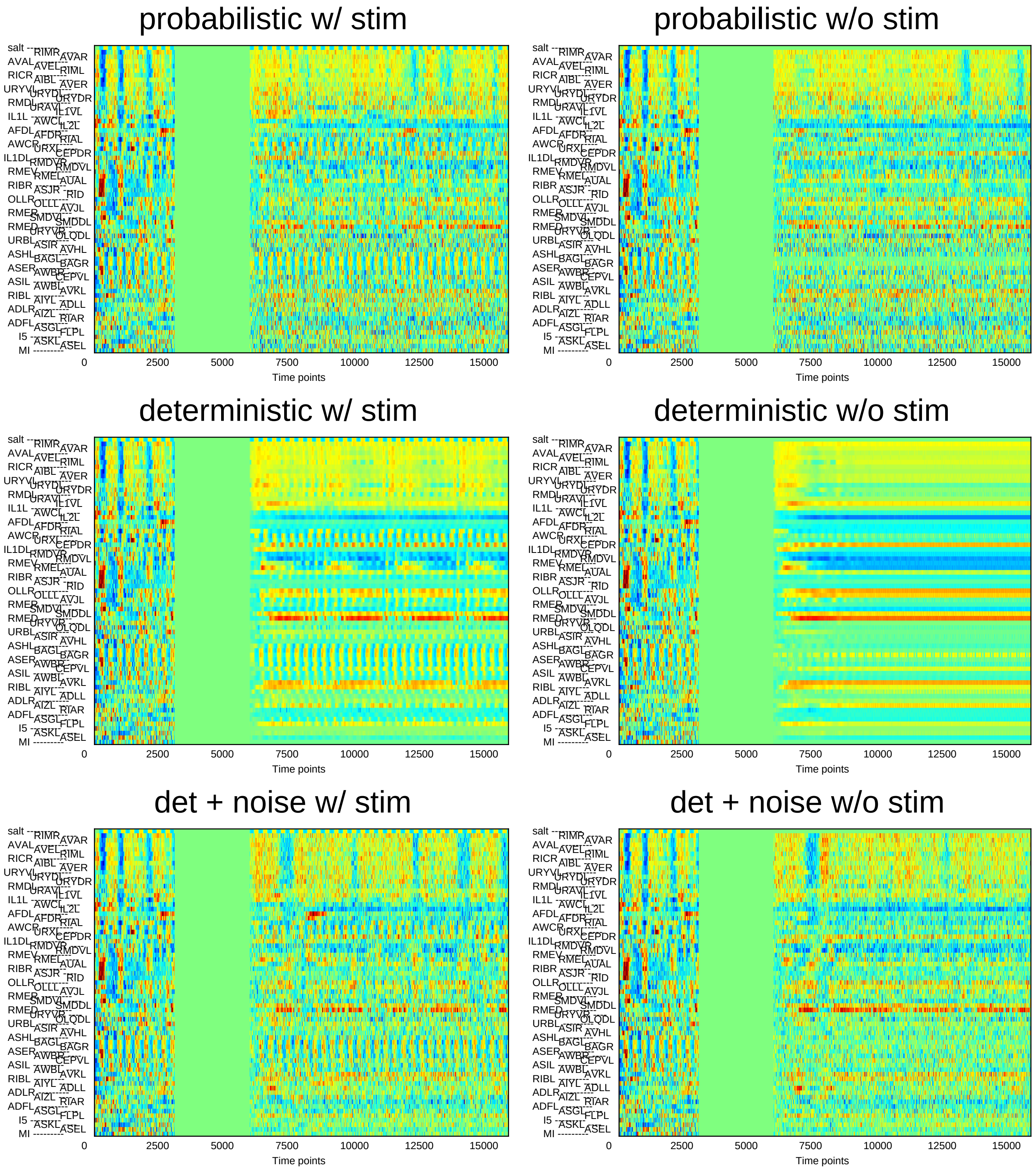
probabilistic w/ stim
probabilistic w/o stim
deterministic w/ stim
deterministic w/o stim
det + noise w/ stim
det + noise w/o stim
Time points
0
2500
5000
7500
10000
12500
15000
salt
AVAL
RICR
URYVL
RMDL
IL1L
AFDL
AWCR
IL1DL
RMEV
RIBR
OLLR
RMER
RMED
URBL
ASHL
ASER
ASIL
RIBL
ADLR
ADFL
I5
MI
RIMR
AVEL
AIBL
URYDL
URAVL
AWCL
AFDR
URXL
RMDVR
RMEL
ASJR
OLLL
SMDVL
URYVR
ASIR
BAGL
AWBR
AWBL
AIYL
AIZL
ASGL
ASKL
AVAR
RIML
AVER
URYDR
IL1VL
IL2L
RIAL
CEPDR
RMDVL
AUAL
RID
AVJL
SMDDL
OLQDL
AVHL
BAGR
CEPVL
AVKL
ADLL
RIAR
FLPL
ASEL

### Sample6

#### probabilistic w/ stim

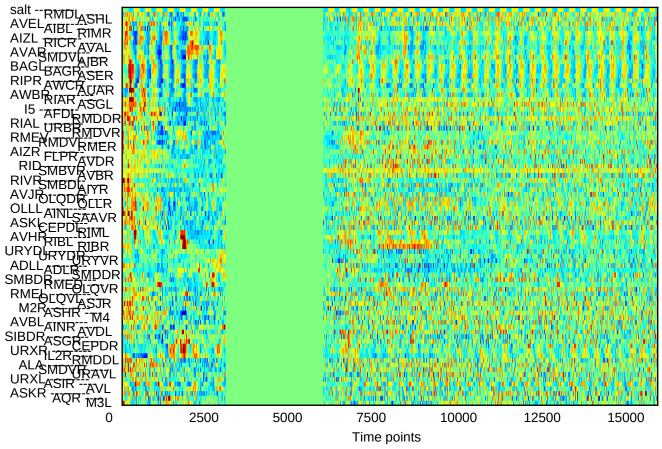


#### probabilistic w/o stim

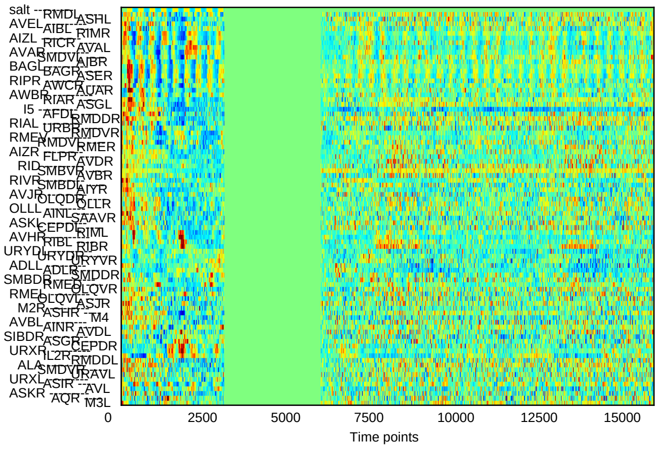


#### deterministic w/ stim

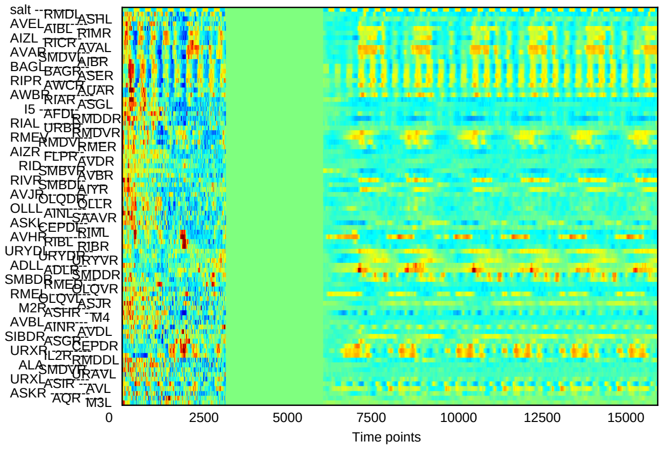


#### deterministic w/o stim

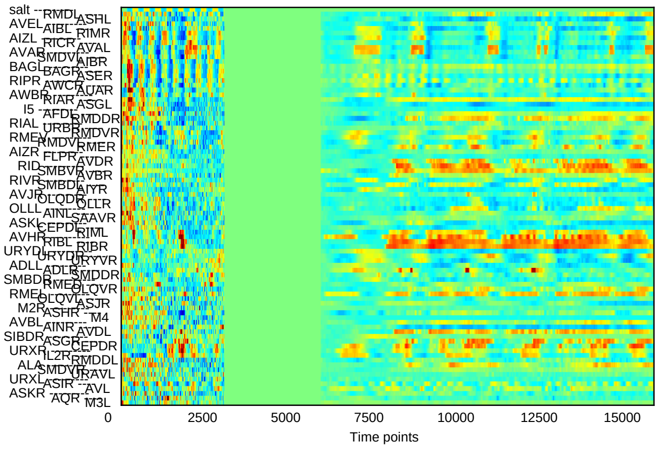


#### det + noise w/ stim

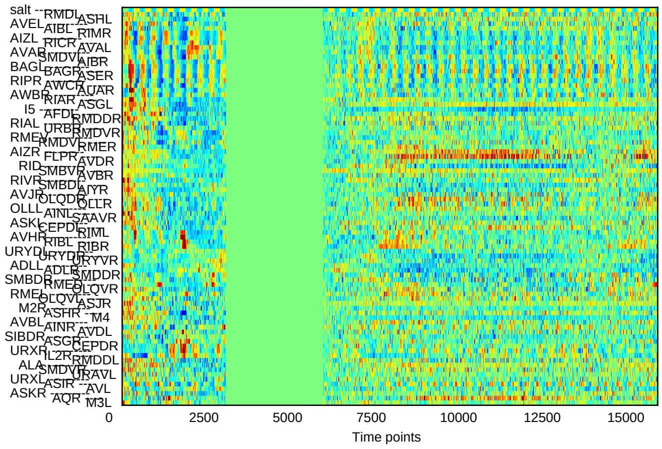


#### det + noise w/o stim

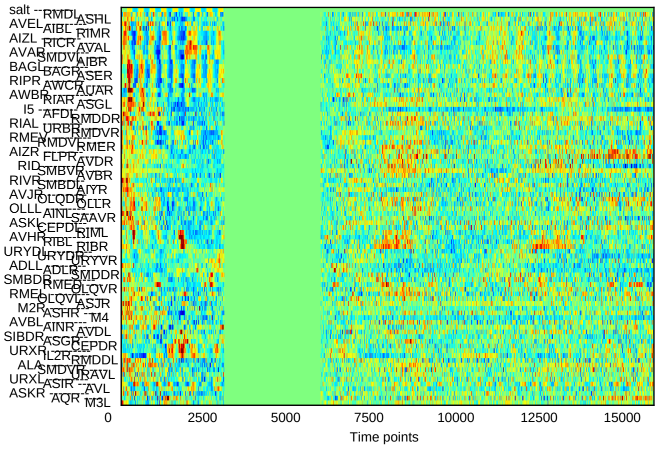

### Sample7

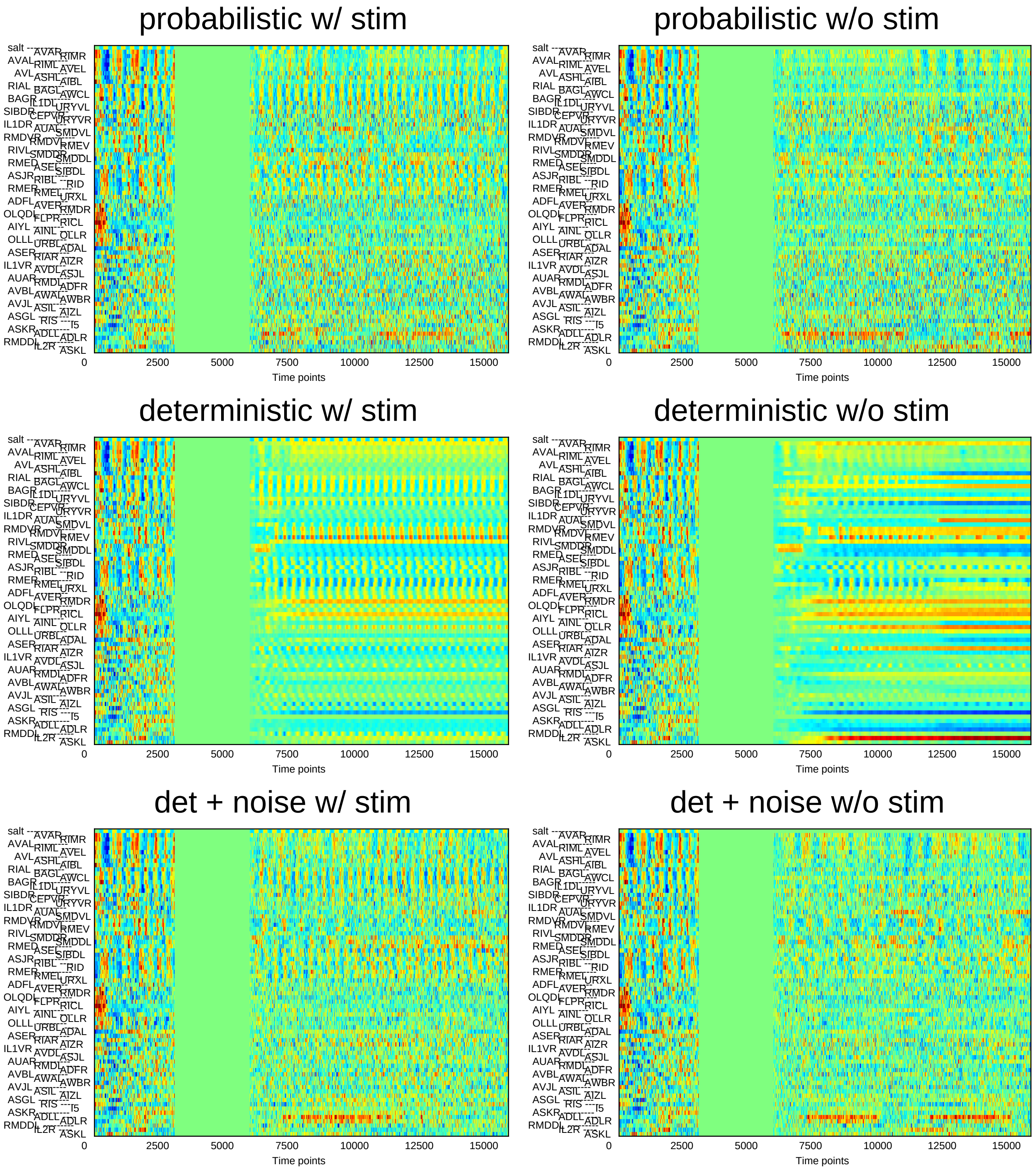

probabilistic w/ stim
probabilistic w/o stim
deterministic w/ stim
deterministic w/o stim
det + noise w/ stim
det + noise w/o stim
Time points
0
2500
5000
7500
10000
12500
15000

### Sample8

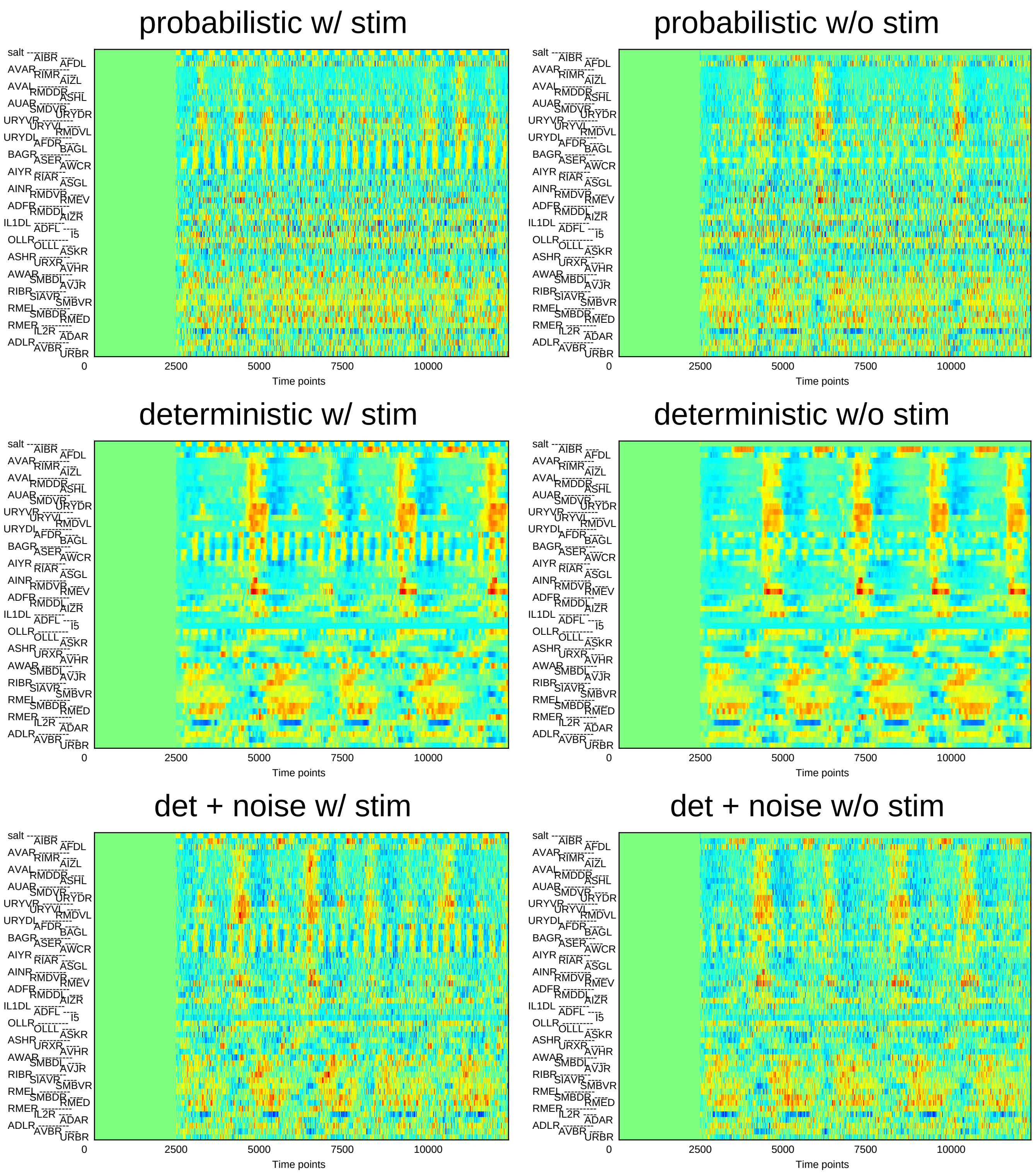
probabilistic w/ stim
probabilistic w/o stim
deterministic w/ stim
deterministic w/o stim
det + noise w/ stim
det + noise w/o stim
salt
AIBR
AFDL
AVAR
RIMR
AIZL
AVAL
RMDDR
ASHL
AUAR
SMDVR
URYDR
URYVR
URYVL
RMDVL
URYDL
AFDR
BAGL
BAGR
ASER
AWCR
AIYR
RIAR
ASGL
AINR
RMDVR
RMEV
ADFR
RMDDL
AIZR
IL1DL
ADFL
I5
OLLR
OLLL
ASKR
ASHR
URXR
AVHR
AWAR
SMBDL
AVJR
RIBR
SIAVR
SMBVR
RMEL
SMBDR
RMED
RMER
IL2R
ADAR
ADLR
AVBR
URBR
0
2500
5000
7500
10000
Time points

### Sample9

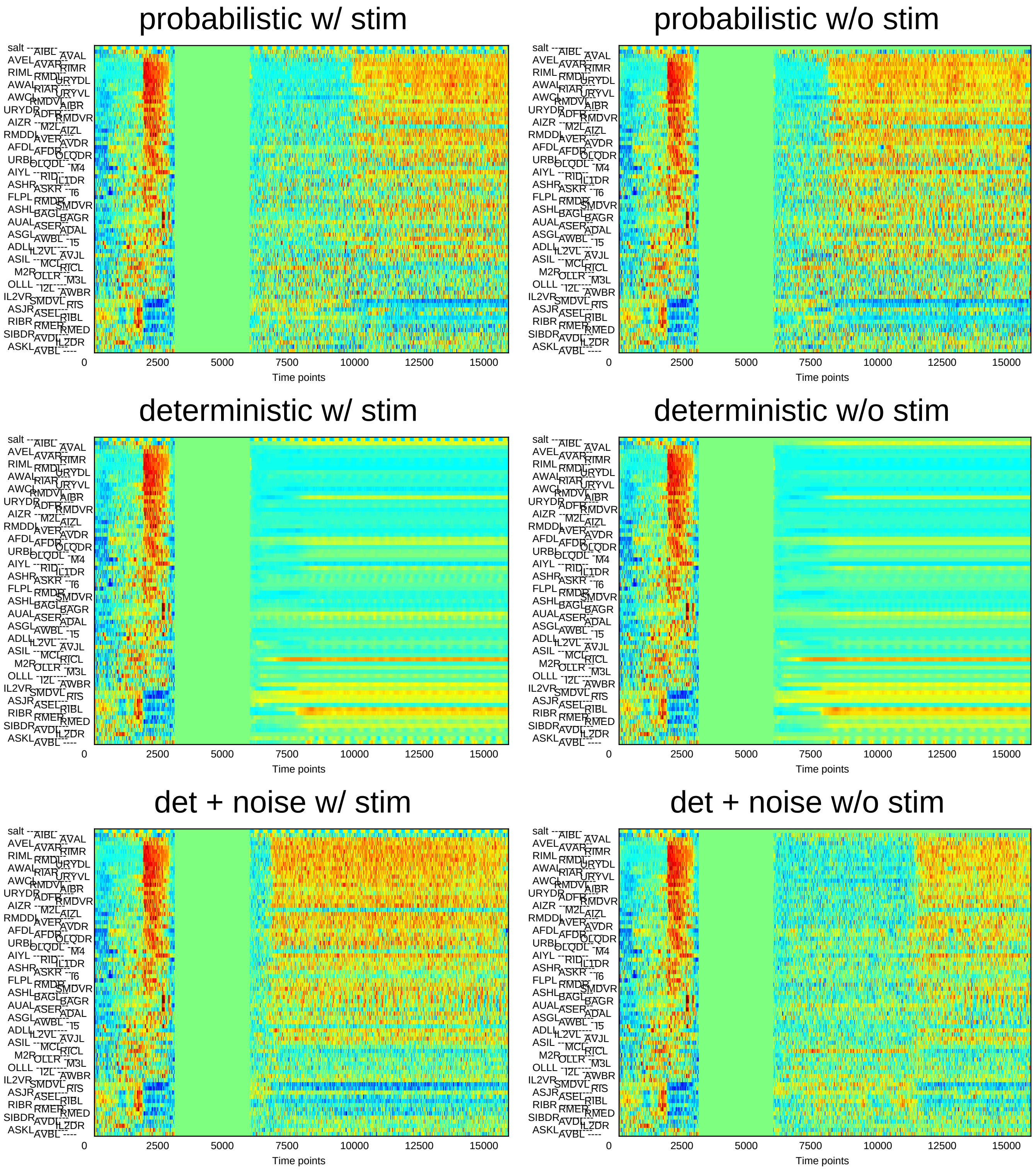
probabilistic w/ stim
probabilistic w/o stim
deterministic w/ stim
deterministic w/o stim
det + noise w/ stim
det + noise w/o stim
0 2500 5000 7500 10000 12500 15000
Time points

### Sample10

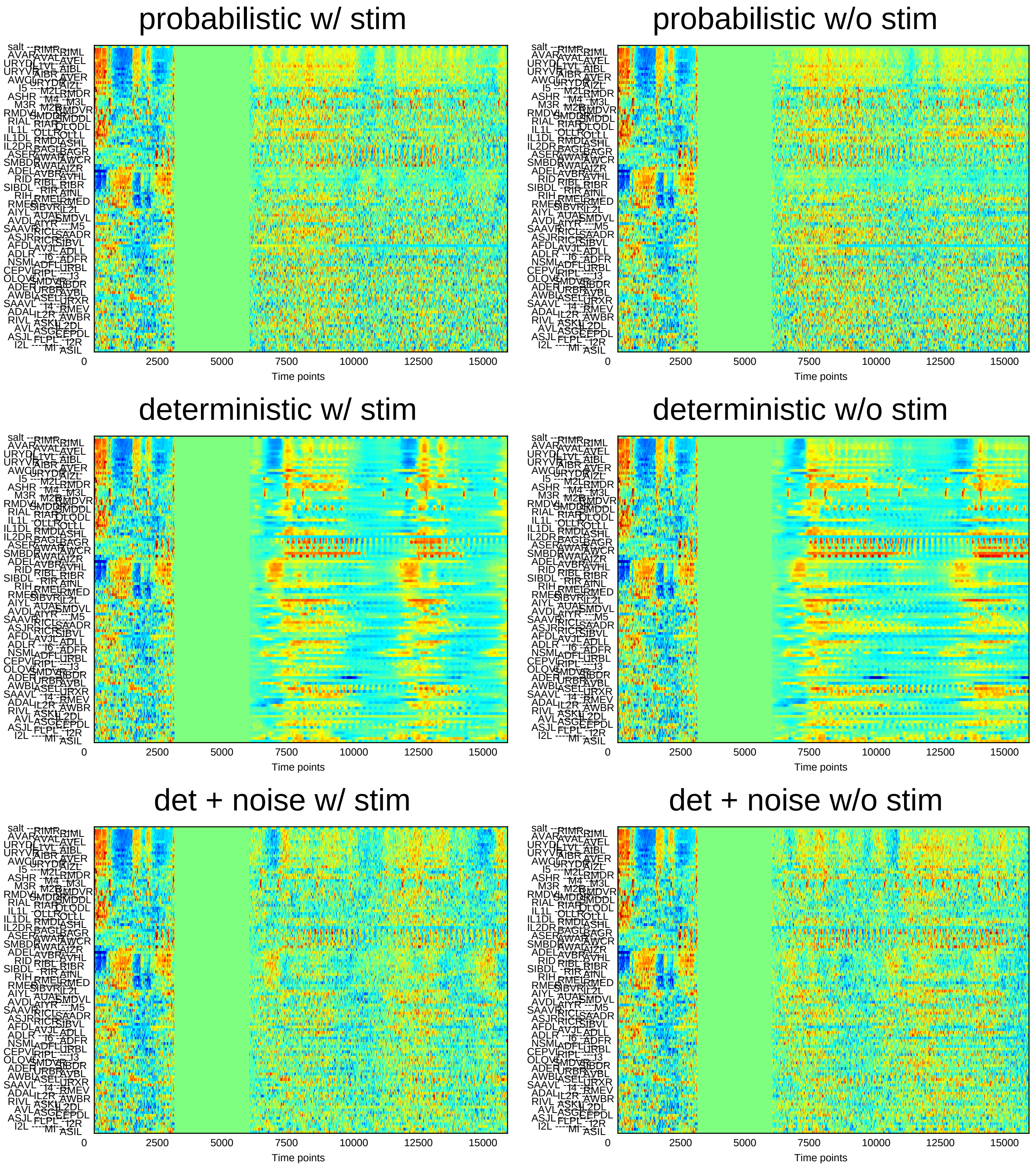
probabilistic w/ stim
probabilistic w/o stim
deterministic w/ stim
deterministic w/o stim
det + noise w/ stim
det + noise w/o stim
0
2500
5000
7500
10000
12500
15000
Time points

### Sample11

#### probabilistic w/ stim

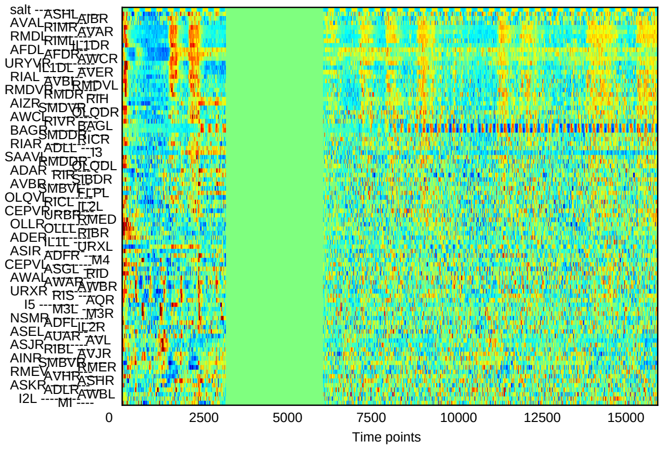


#### probabilistic w/o stim

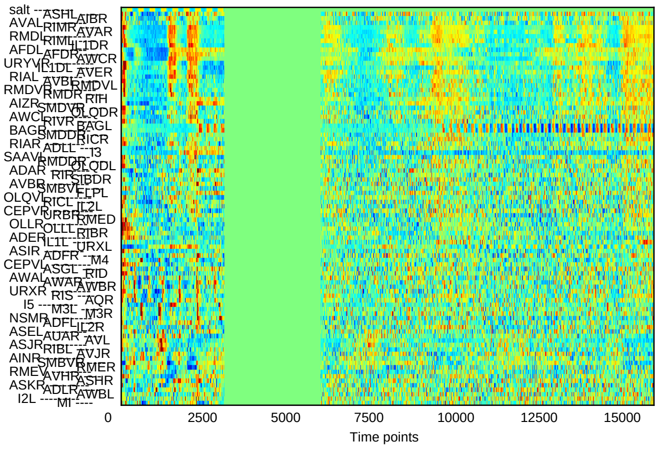


#### deterministic w/ stim

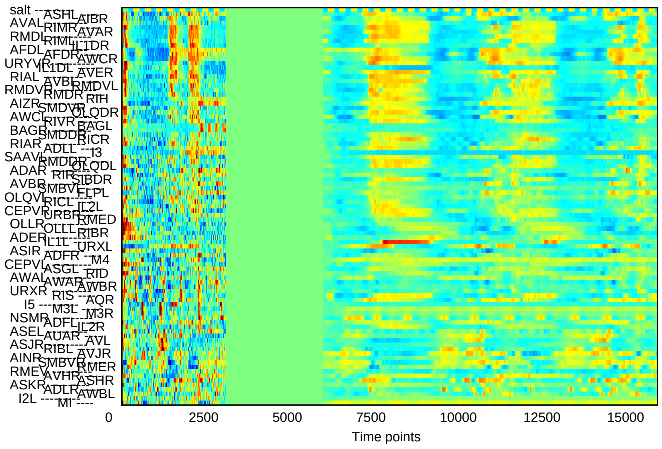


#### deterministic w/o stim

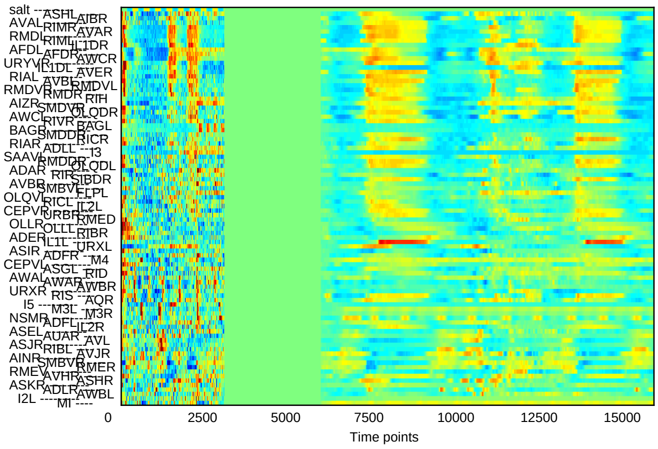


#### det + noise w/ stim

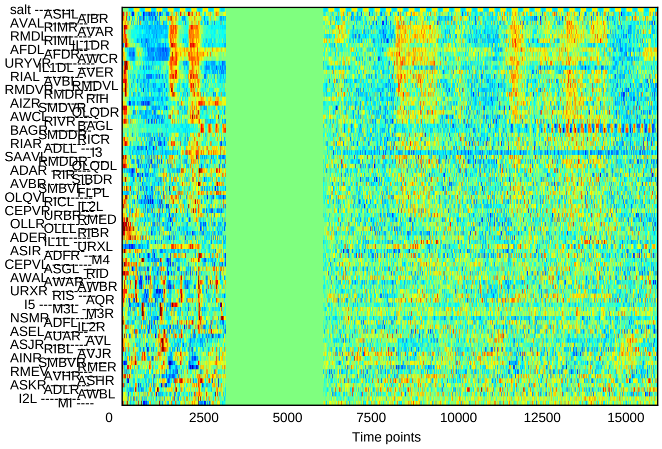


#### det + noise w/o stim

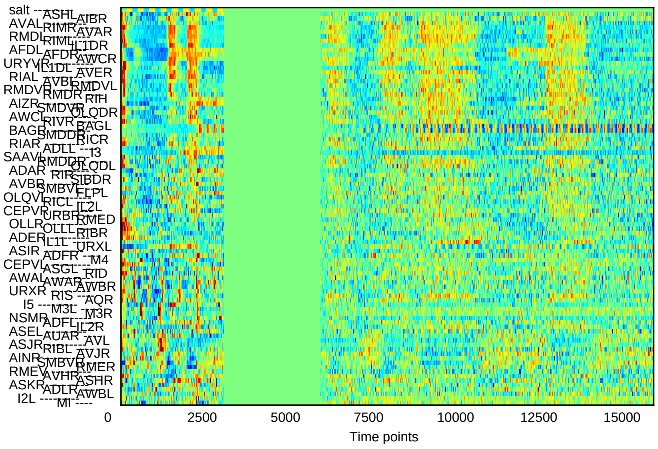

### Sample12

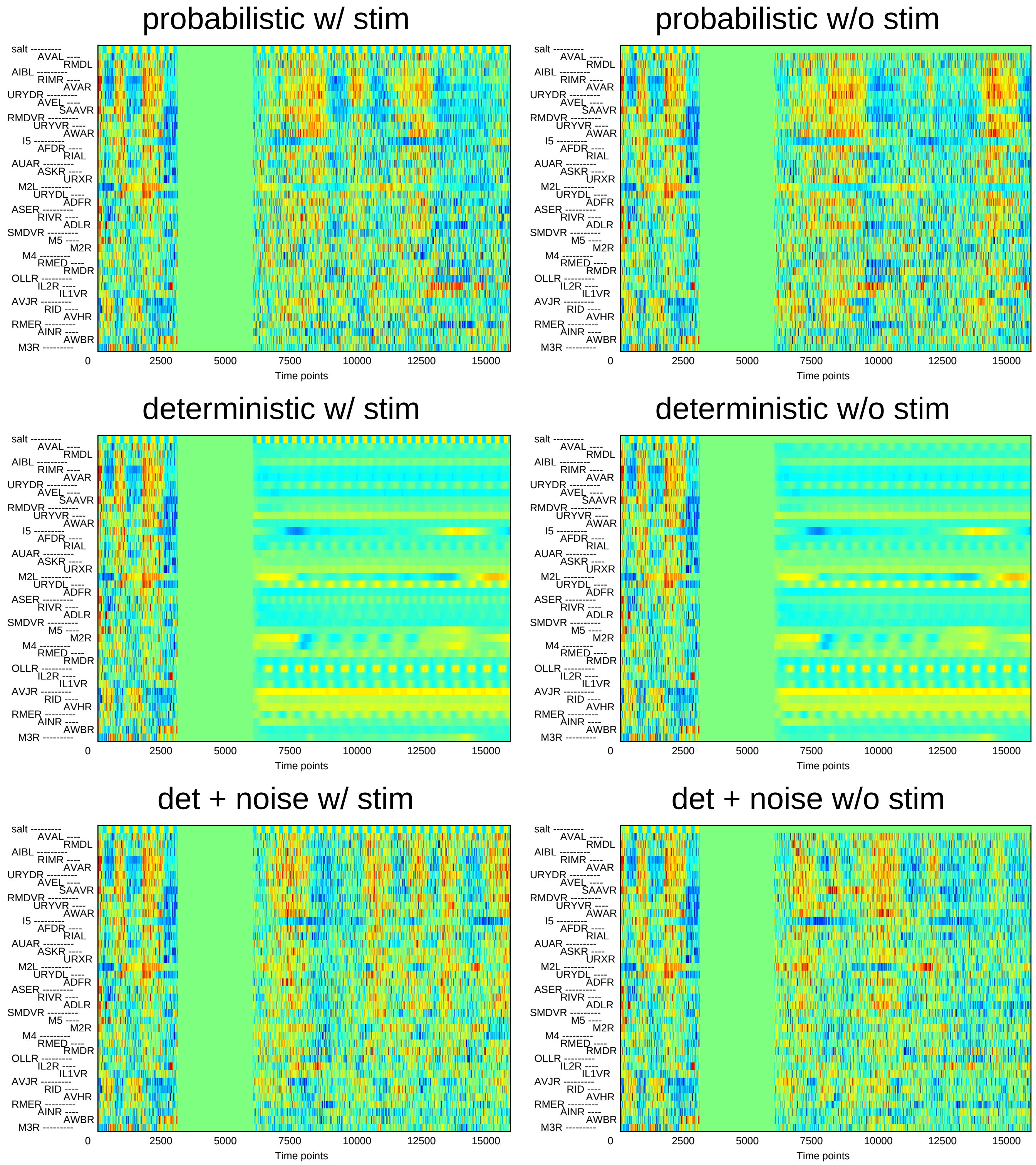
probabilistic w/ stim
probabilistic w/o stim
deterministic w/ stim
deterministic w/o stim
det + noise w/ stim
det + noise w/o stim
salt
AVAL
RMDL
AIBL
RIMR
AVAR
URYDR
AVEL
SAAVR
RMDVR
URYVR
AWAR
I5
AFDR
RIAL
AUAR
ASKR
URXR
M2L
URYDL
ADFR
ASER
RIVR
ADLR
SMDVR
M5
M2R
M4
RMED
RMDR
OLLR
IL2R
IL1VR
AVJR
RID
AVHR
RMER
AINR
AWBR
M3R
0
2500
5000
7500
10000
12500
15000
Time points

### Sample13

#### probabilistic w/ stim

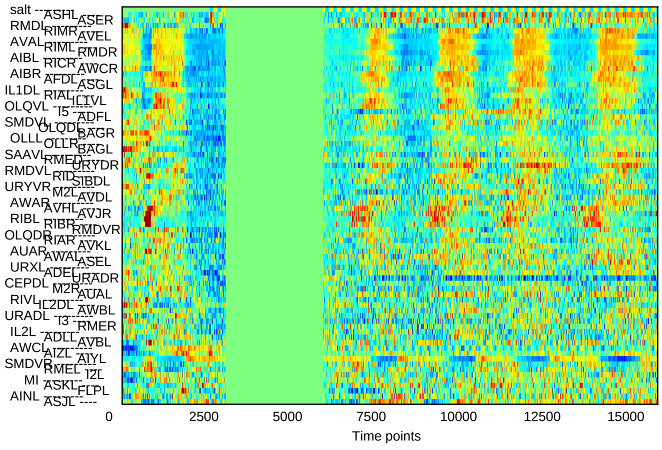


#### probabilistic w/o stim

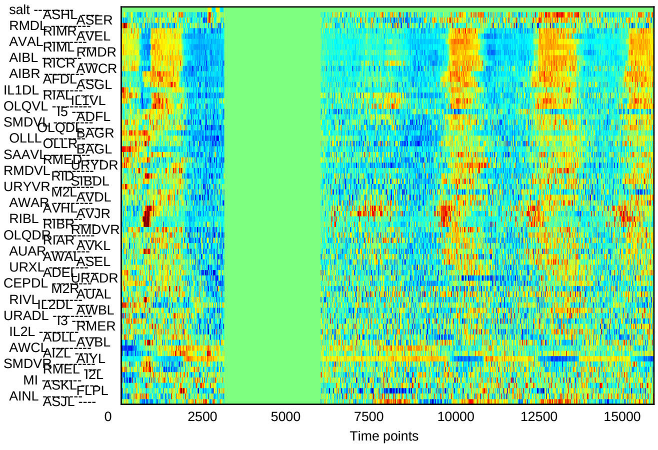


#### deterministic w/ stim

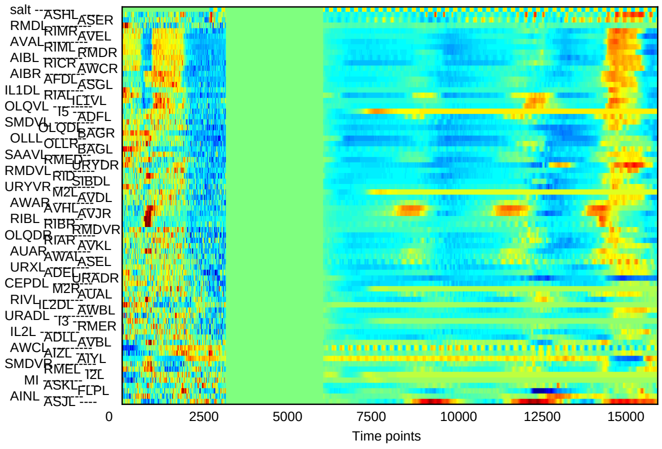


#### deterministic w/o stim

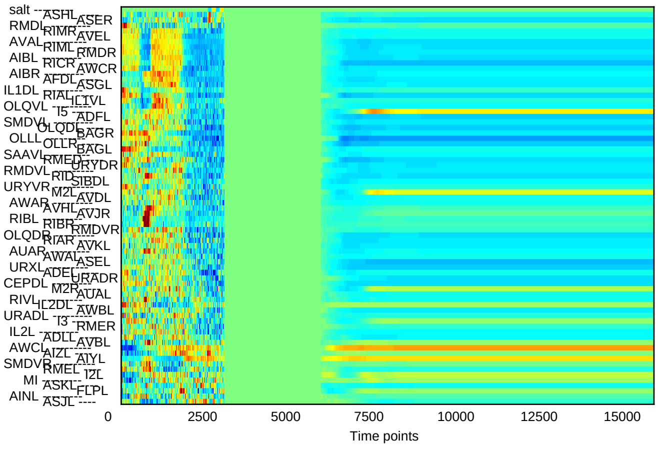


#### det + noise w/ stim

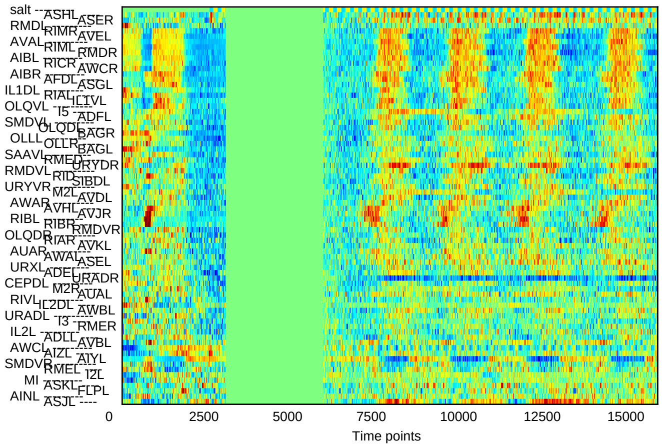


#### det + noise w/o stim

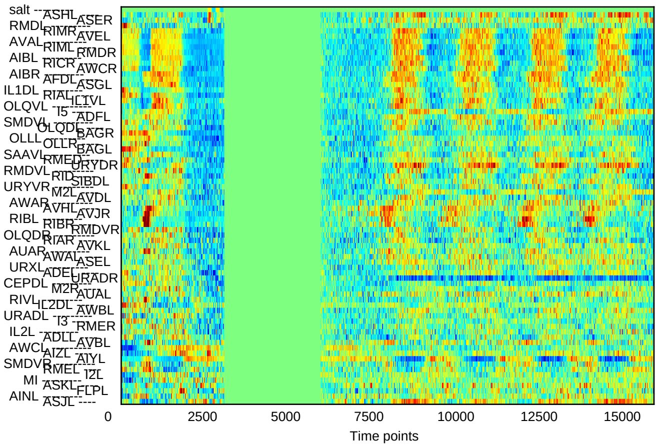

### Sample14

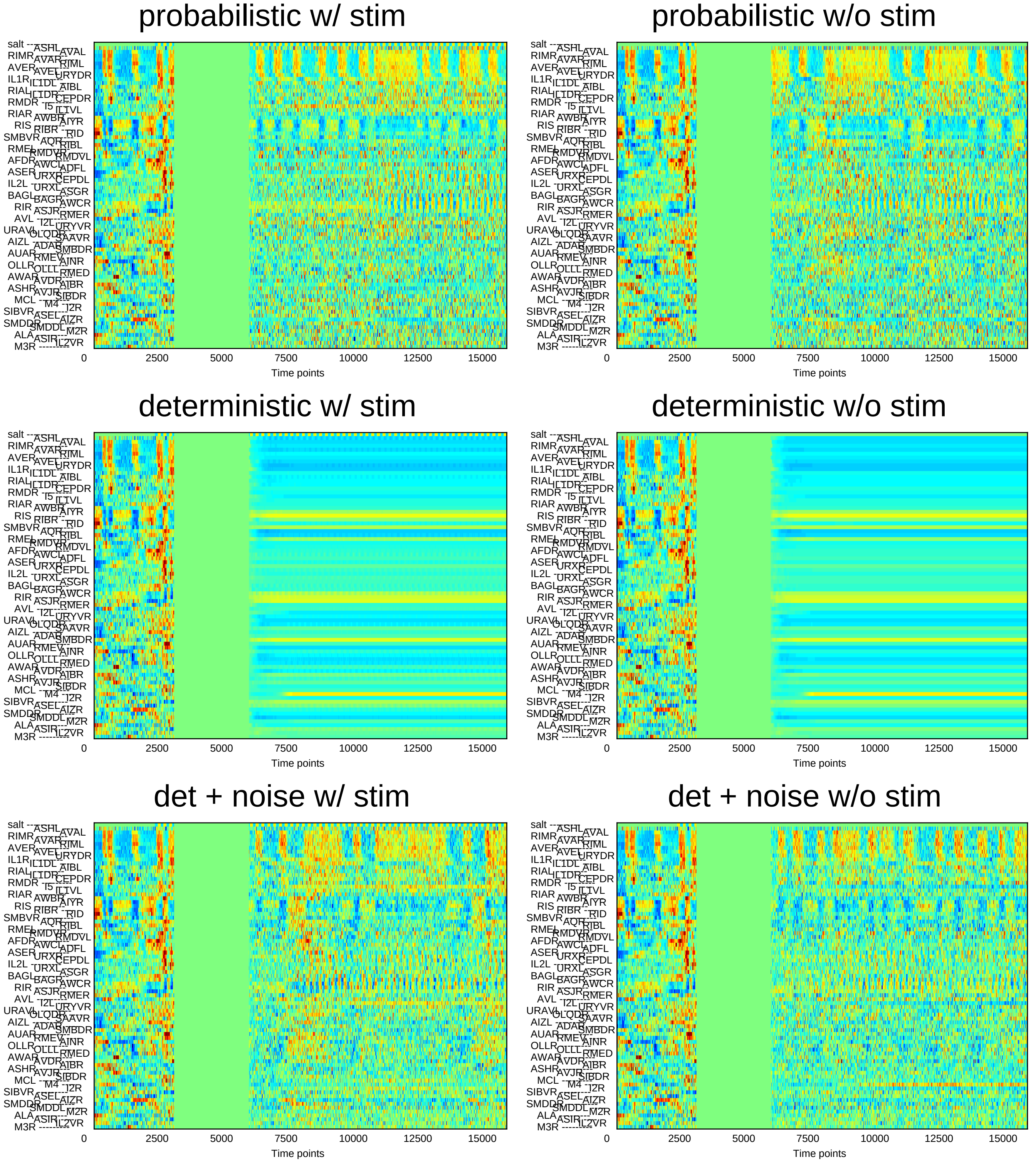
probabilistic w/ stim
probabilistic w/o stim
deterministic w/ stim
deterministic w/o stim
det + noise w/ stim
det + noise w/o stim
Time points
0
2500
5000
7500
10000
12500
15000

### Sample15

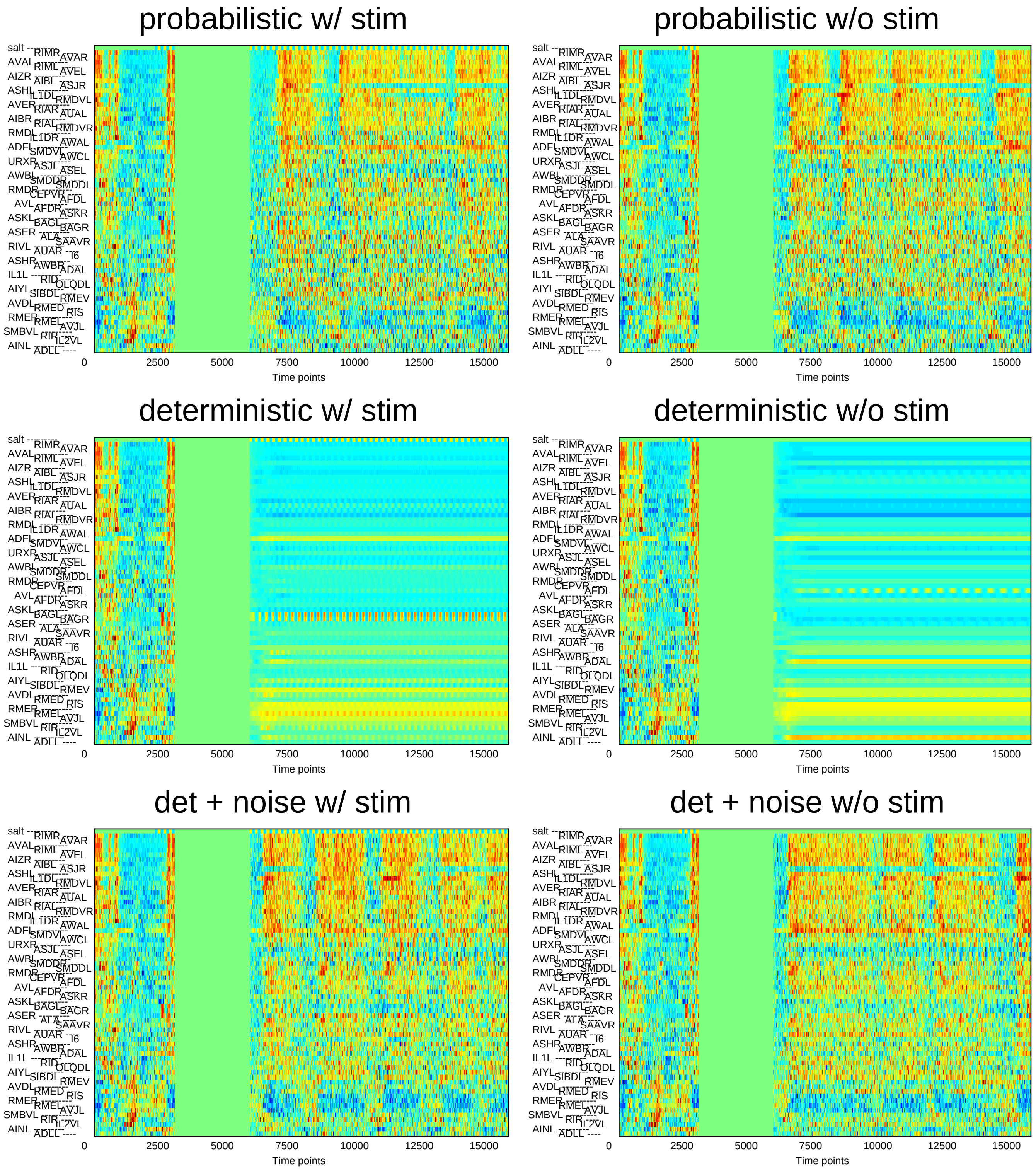
probabilistic w/ stim
probabilistic w/o stim
deterministic w/ stim
deterministic w/o stim
det + noise w/ stim
det + noise w/o stim
salt
RIMR
AVAR
AVAL
RIML
AVEL
AIZR
AIBL
ASJR
ASHL
IL1DL
RMDVL
AVER
RIAR
AUAL
AIBR
RIAL
RMDVR
RMDL
IL1DR
AWAL
ADFL
SMDVL
AWCL
URXR
ASJL
ASEL
AWBL
SMDDR
SMDDL
RMDR
CEPVR
AFDL
AVL
AFDR
ASKR
ASKL
BAGL
BAGR
ASER
ALA
SAAVR
RIVL
AUAR
I6
ASHR
AWBR
ADAL
IL1L
RID
OLQDL
AIYL
SIBDL
RMEV
AVDL
RMED
RIS
RMER
RMEL
AVJL
SMBVL
RIR
IL2VL
AINL
ADLL
0
2500
5000
7500
10000
12500
15000
Time points

### Sample16

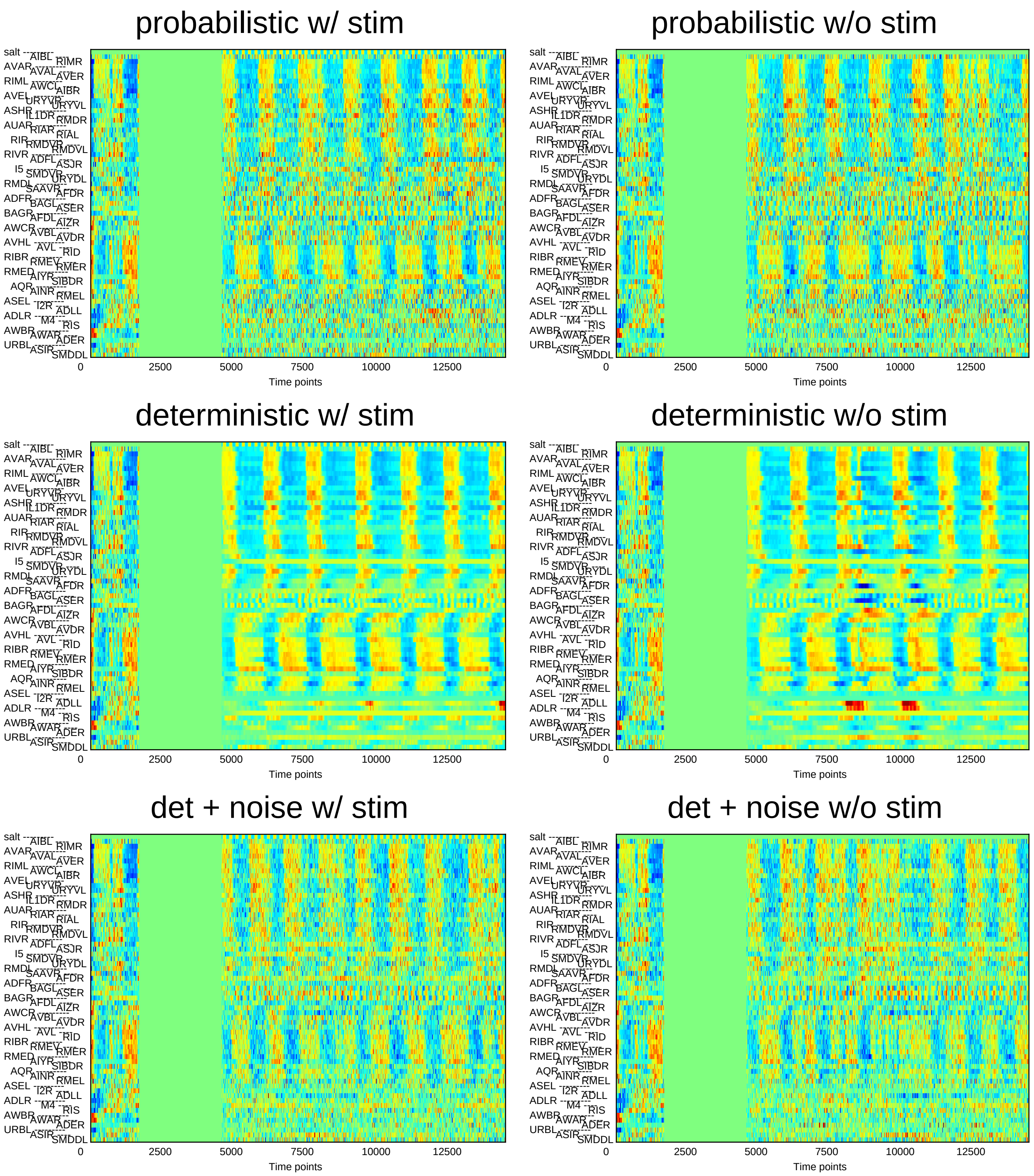
probabilistic w/ stim
probabilistic w/o stim
deterministic w/ stim
deterministic w/o stim
det + noise w/ stim
det + noise w/o stim
0 2500 5000 7500 10000 12500
Time points
salt AIBL RIMR AVAR AVAL AVER RIML AWCL AIBR AVEL URYVR URYVL ASHR IL1DR RMDR AUAR RIAR RIAL RIR RMDVR RMDVL RIVR ADFL ASJR I5 SMDVR URYDL RMDL SAAVR AFDR ADFR BAGL ASER BAGR AFDL AIZR AWCR AVBL AVDR AVHL AVL RID RIBR RMEV RMER RMED AIYR SIBDR AQR AINR RMEL ASEL I2R ADLL ADLR M4 RIS AWBR AWAR ADER URBL ASIR SMDDL

### Sample17

#### probabilistic w/ stim

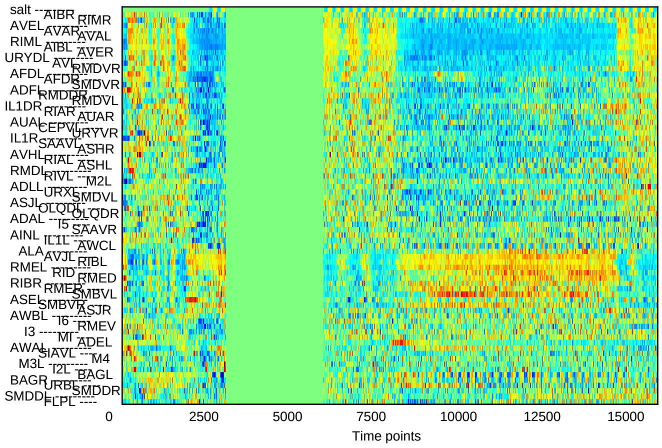


#### probabilistic w/o stim

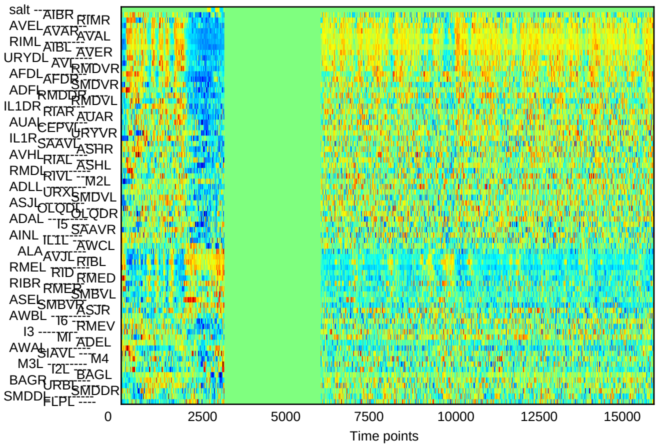


#### deterministic w/ stim

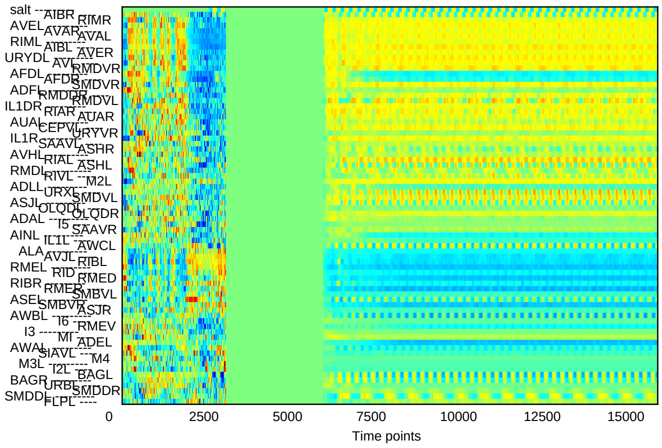


#### deterministic w/o stim

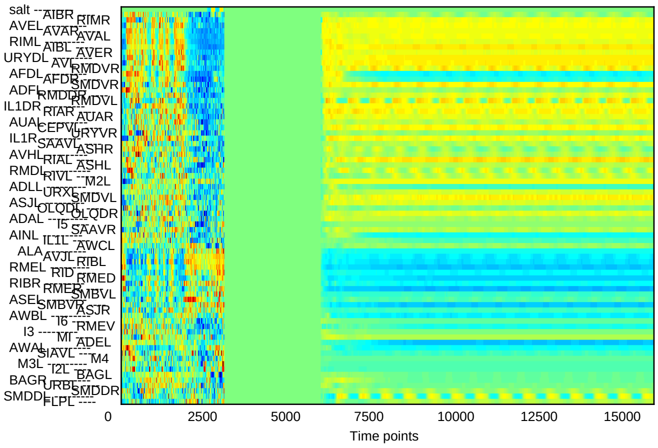


#### det + noise w/ stim

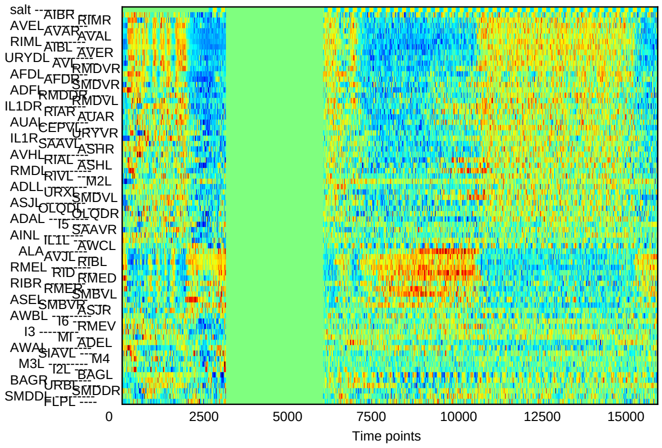


#### det + noise w/o stim

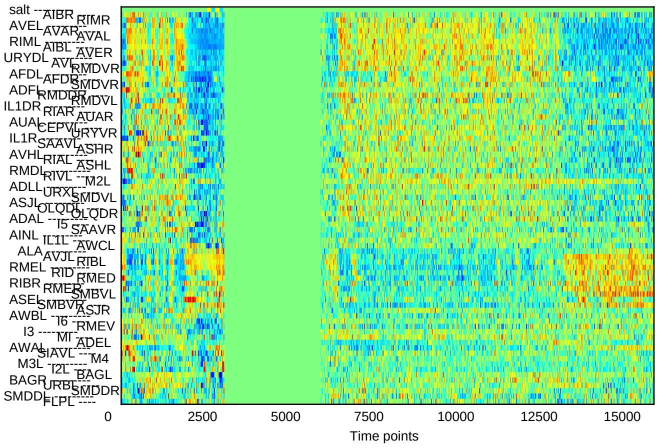

### Sample18

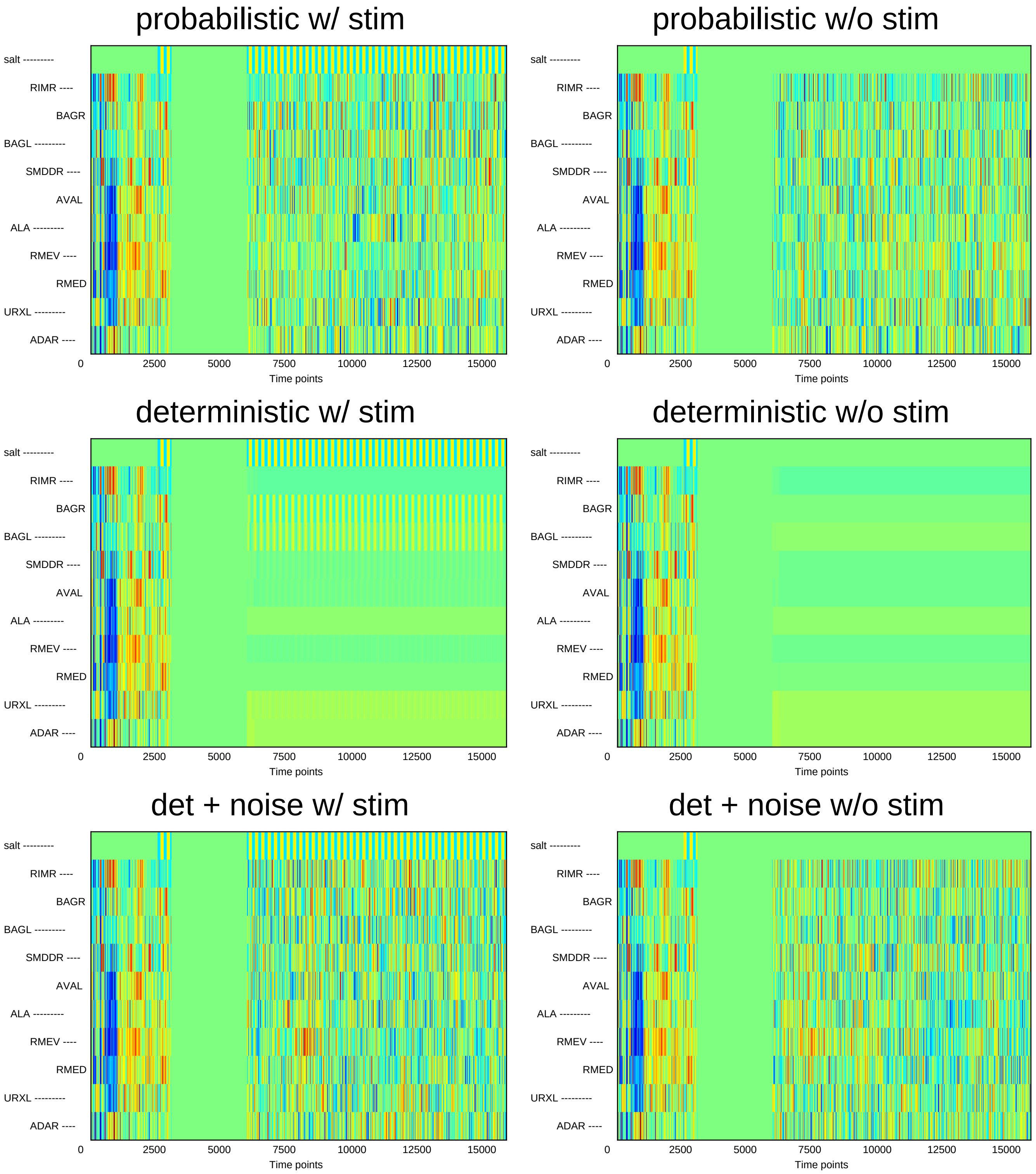
probabilistic w/ stim
probabilistic w/o stim
deterministic w/ stim
deterministic w/o stim
det + noise w/ stim
det + noise w/o stim
salt ---------
RIMR ----
BAGR
BAGL ---------
SMDDR ----
AVAL
ALA ---------
RMEV ----
RMED
URXL ---------
ADAR ----
0
2500
5000
7500
10000
12500
15000
Time points

### Sample19

#### probabilistic w/ stim

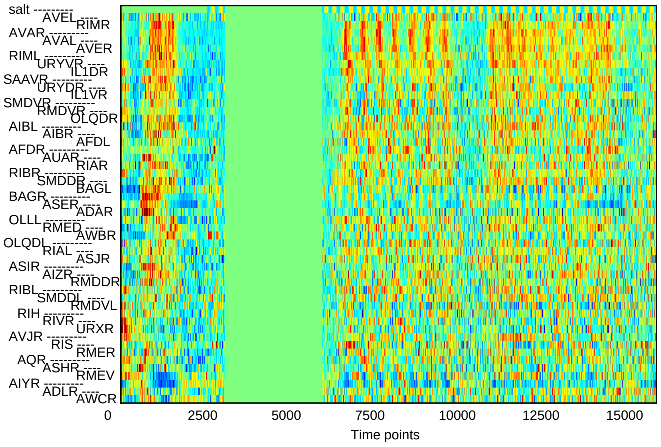


#### probabilistic w/o stim

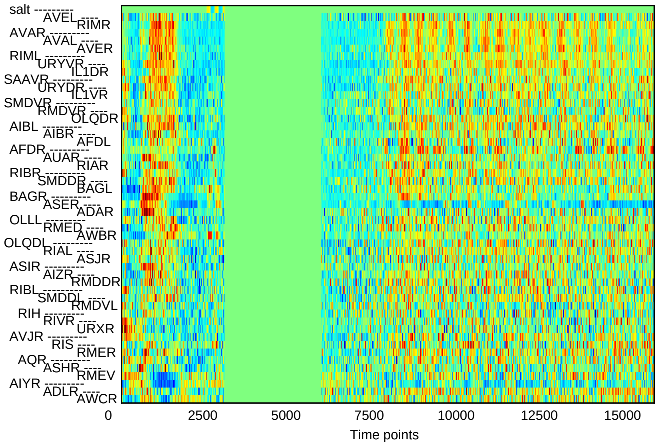


#### deterministic w/ stim

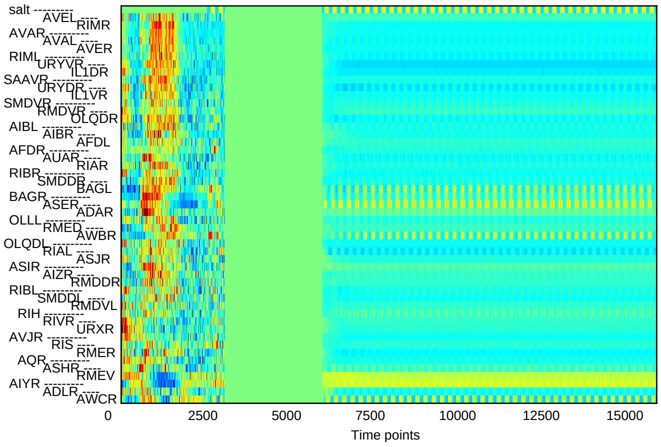


#### deterministic w/o stim

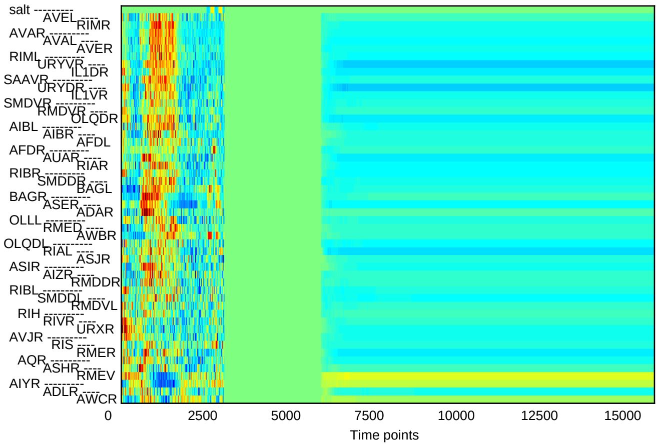


#### det + noise w/ stim

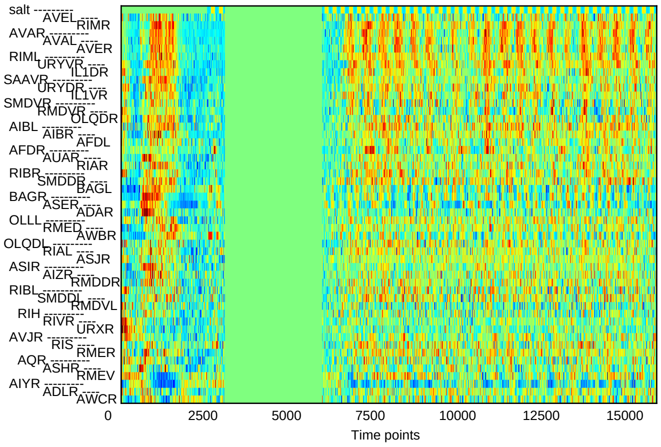


#### det + noise w/o stim

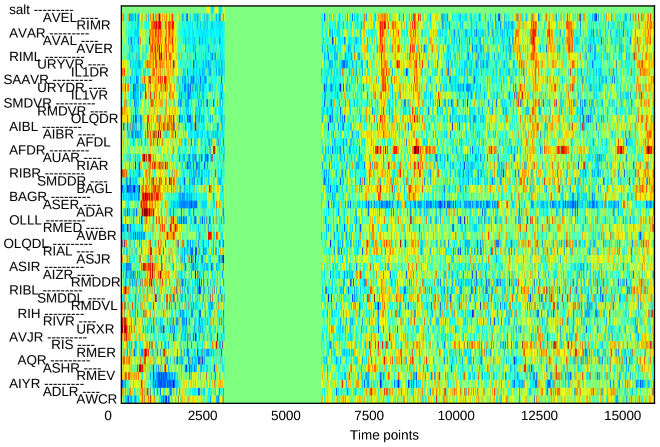

### Sample20

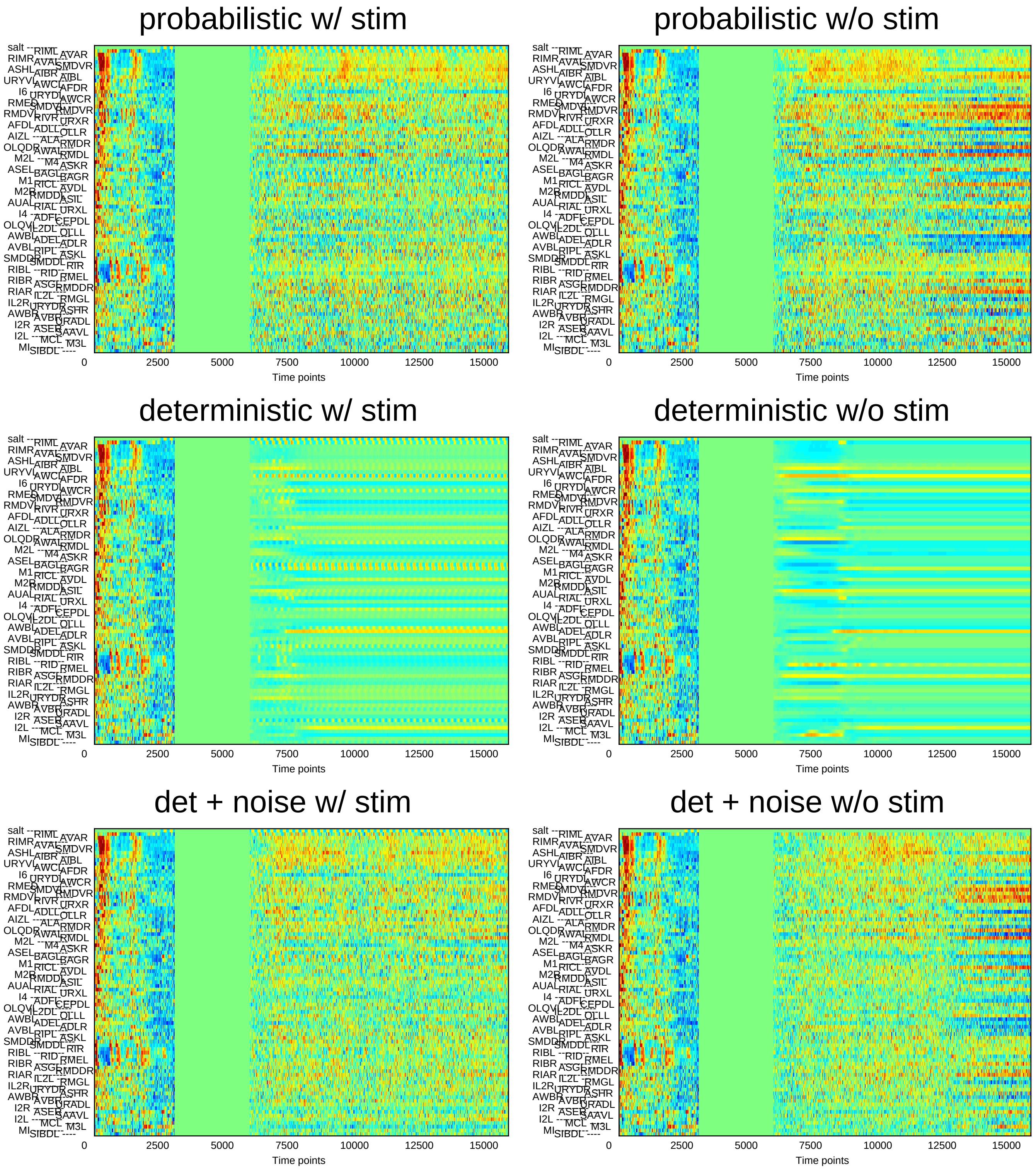
probabilistic w/ stim
probabilistic w/o stim
deterministic w/ stim
deterministic w/o stim
det + noise w/ stim
det + noise w/o stim
Time points
0
2500
5000
7500
10000
12500
15000

### Sample21

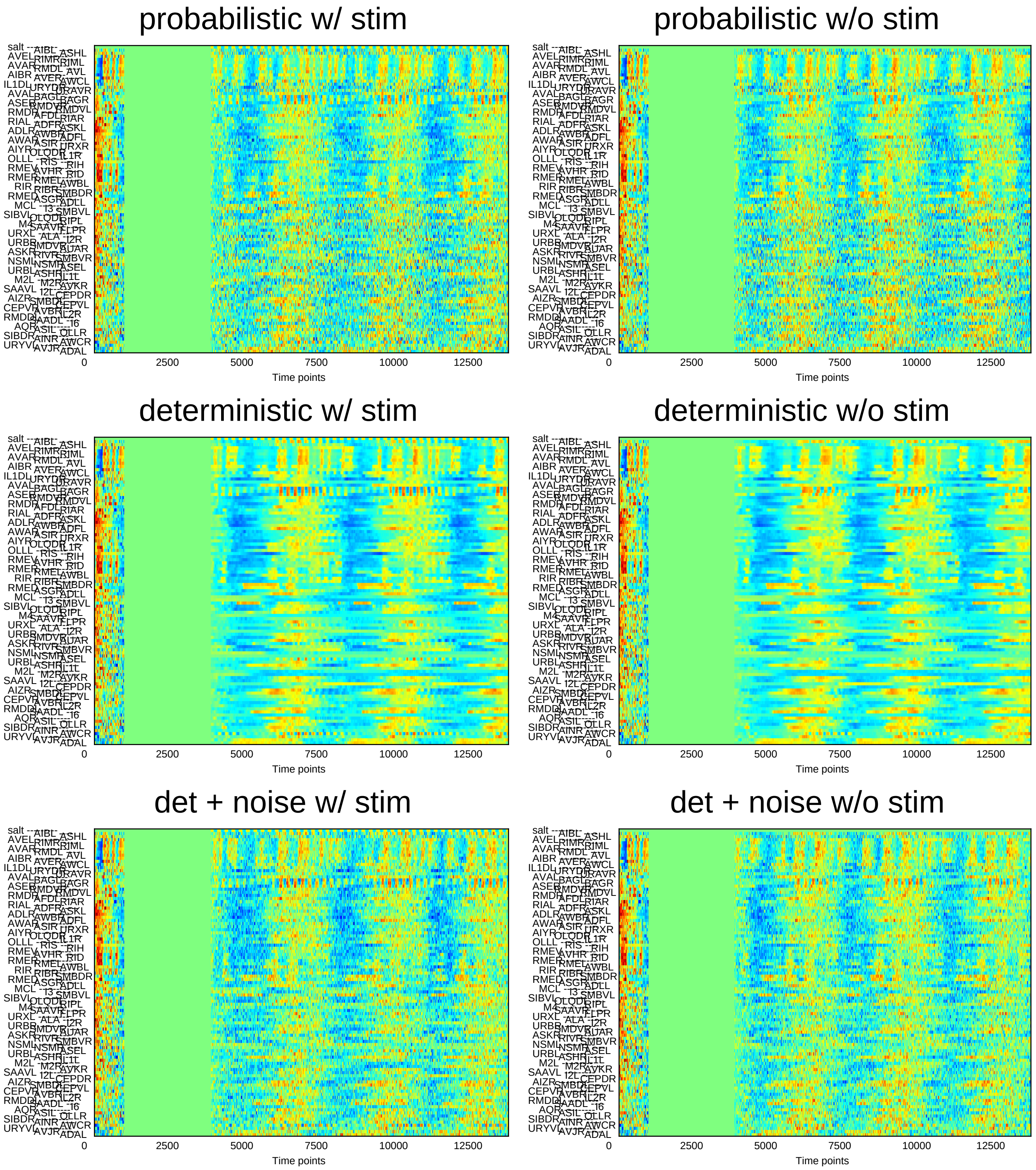
probabilistic w/ stim
probabilistic w/o stim
deterministic w/ stim
deterministic w/o stim
det + noise w/ stim
det + noise w/o stim
0
2500
5000
7500
10000
12500
Time points

### Sample22

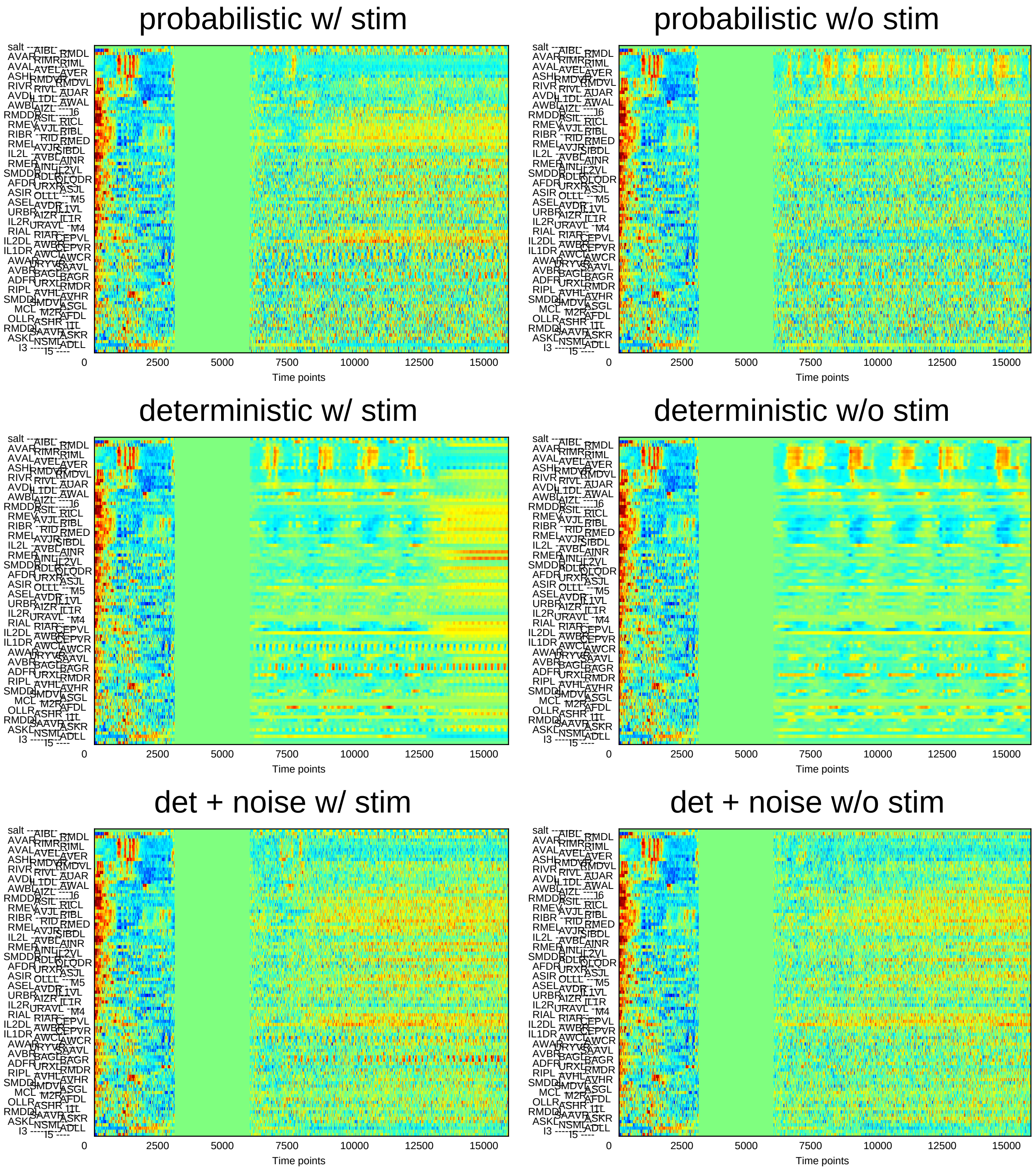
probabilistic w/ stim
probabilistic w/o stim
deterministic w/ stim
deterministic w/o stim
det + noise w/ stim
det + noise w/o stim
0
2500
5000
7500
10000
12500
15000
Time points

### Sample23

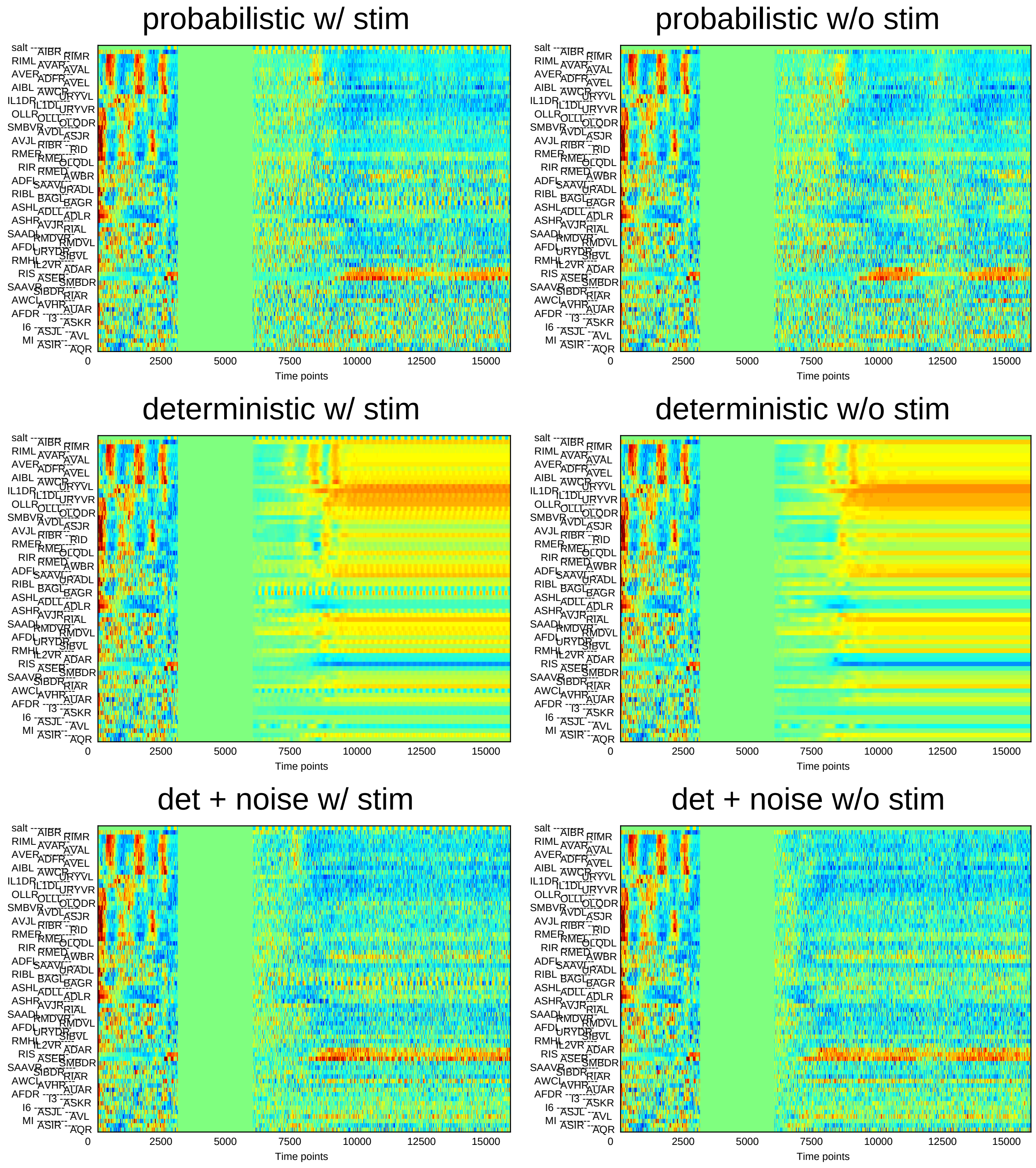
probabilistic w/ stim
probabilistic w/o stim
deterministic w/ stim
deterministic w/o stim
det + noise w/ stim
det + noise w/o stim
salt AIBR RIMR RIML AVAR AVAL AVER ADFR AVEL AIBL AWCR URYVL IL1DR IL1DL URYVR OLLR OLLL OLQDR SMBVR AVDL ASJR AVJL RIBR RID RMER RMEL OLQDL RIR RMED AWBR ADFL SAAVL URADL RIBL BAGL BAGR ASHL ADLL ADLR ASHR AVJR RIAL SAADL RMDVR RMDVL AFDL URYDR SIBVL RMHL IL2VR ADAR RIS ASER SMBDR SAAVR SIBDR RIAR AWCL AVHR AUAR AFDR I3 ASKR I6 ASJL AVL MI ASIR AQR
0
2500
5000
7500
10000
12500
15000
Time points

### Sample24

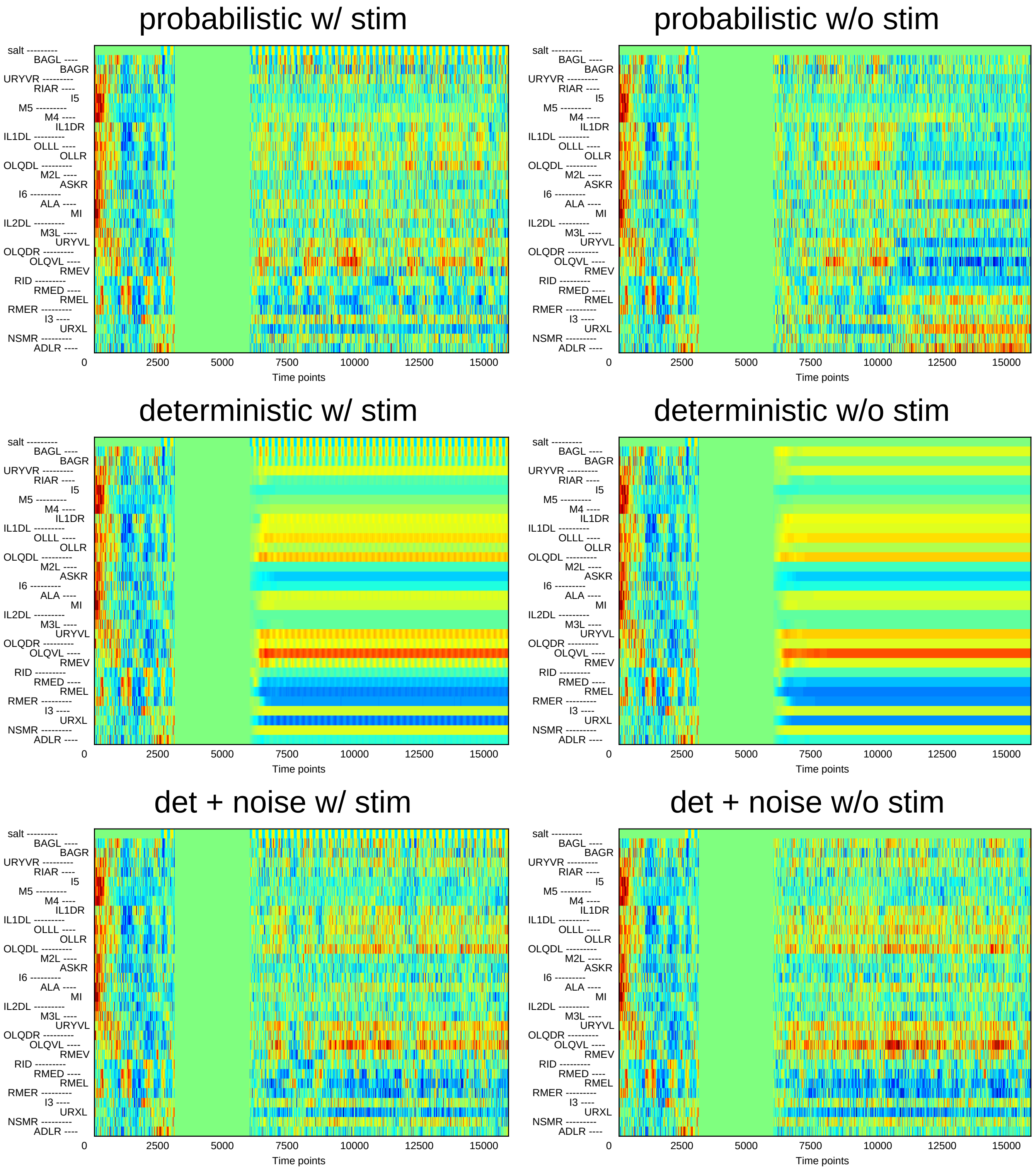
probabilistic w/ stim
probabilistic w/o stim
deterministic w/ stim
deterministic w/o stim
det + noise w/ stim
det + noise w/o stim
salt ---------
BAGL ----
BAGR
URYVR ---------
RIAR ----
I5
M5 ---------
M4 ----
IL1DR
IL1DL ---------
OLLL ----
OLLR
OLQDL ---------
M2L ----
ASKR
I6 ---------
ALA ----
MI
IL2DL ---------
M3L ----
URYVL
OLQDR ---------
OLQVL ----
RMEV
RID ---------
RMED ----
RMEL
RMER ---------
I3 ----
URXL
NSMR ---------
ADLR ----
0
2500
5000
7500
10000
12500
15000
Time points
