## Supplementary File 9B for "Deducing ensemble dynamics and information flow from the whole-brain imaging data"

### Sample1

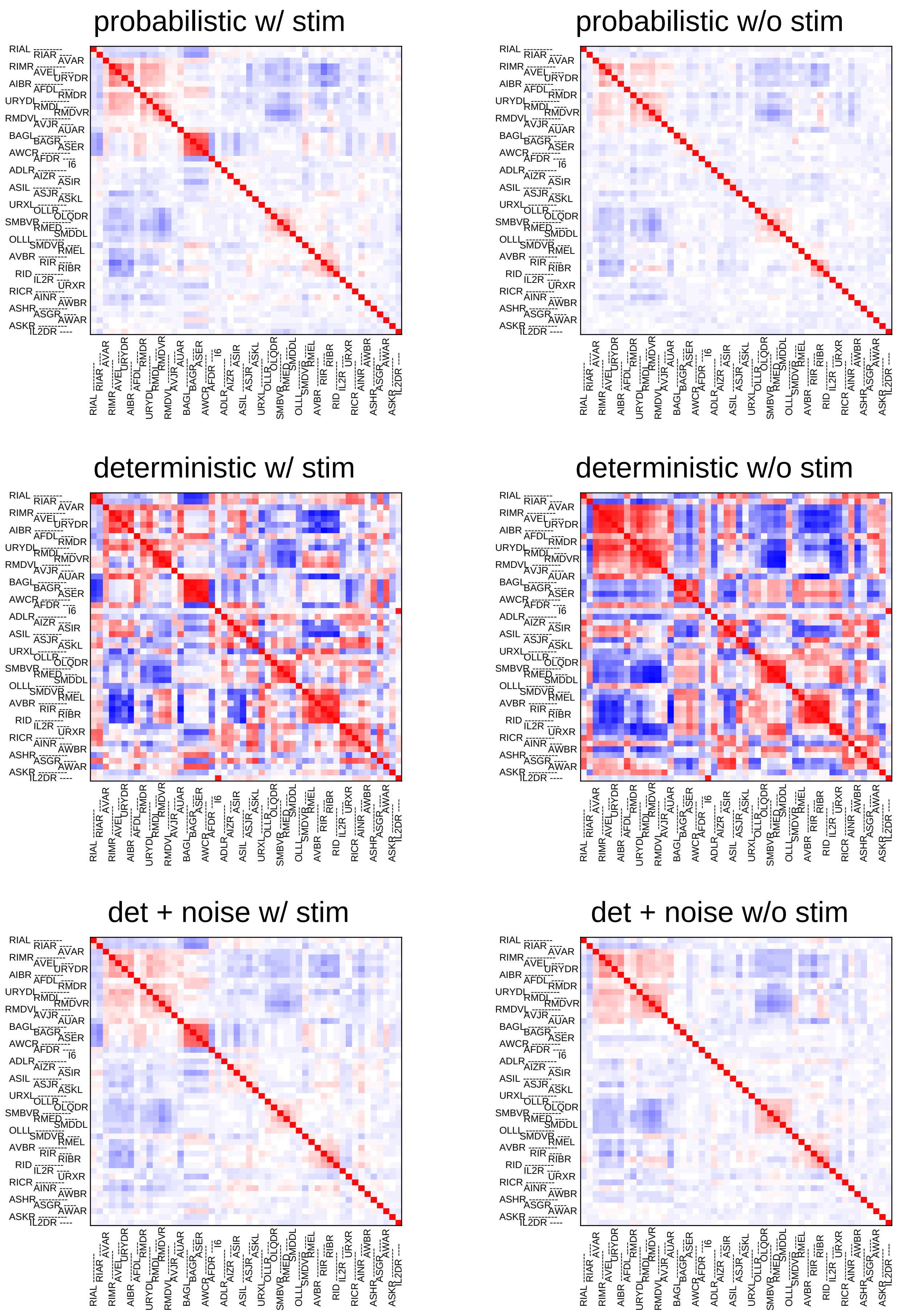
probabilistic w/ stim
probabilistic w/o stim
deterministic w/ stim
deterministic w/o stim
det + noise w/ stim
det + noise w/o stim
RIAL
RIAR
AVAR
RIMR
AVEL
URYDR
AIBR
AFDL
RMDR
URYDL
RMDL
RMDVR
RMDVL
AVJR
AUAR
BAGL
BAGR
ASER
AWCR
AFDR
I6
ADLR
AIZR
ASIR
ASIL
ASJR
ASKL
URXL
OLLR
OLQDR
SMBVR
RMED
SMDDL
OLLL
SMDVR
RMEL
AVBR
RIR
RIBR
RID
IL2R
URXR
RICR
AINR
AWBR
ASHR
ASGR
AWAR
ASKR
IL2DR

### Sample2

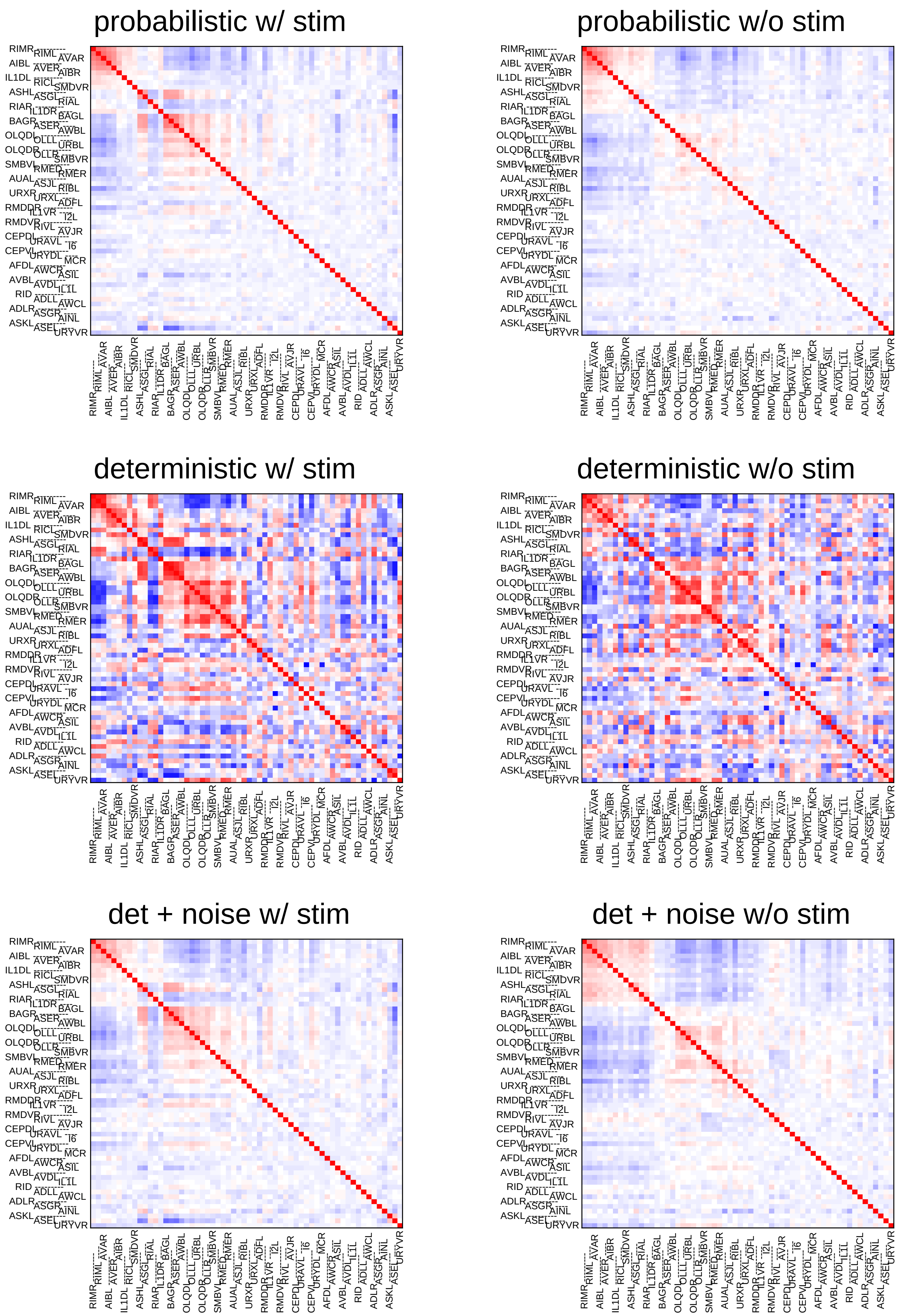
probabilistic w/ stim
probabilistic w/o stim
deterministic w/ stim
deterministic w/o stim
det + noise w/ stim
det + noise w/o stim
RIMR RIML AVAR AIBL AVER AIBR IL1DL RICL SMDVR ASHL ASGL RIAL RIAR IL1DR BAGL BAGR ASER AWBL OLQDL OLLL URBL OLQDR OLLR SMBVR SMBVL RMED RMER AUAL ASJL RIBL URXR URXL ADFL RMDDR IL1VR I2L RMDVR RIVL AVJR CEPDL URAVL I6 CEPVL URYDL MCR AFDL AWCR ASIL AVBL AVDL IL1L RID ADLL AWCL ADLR ASGR AINL ASKL ASEL URYVR

### Sample3

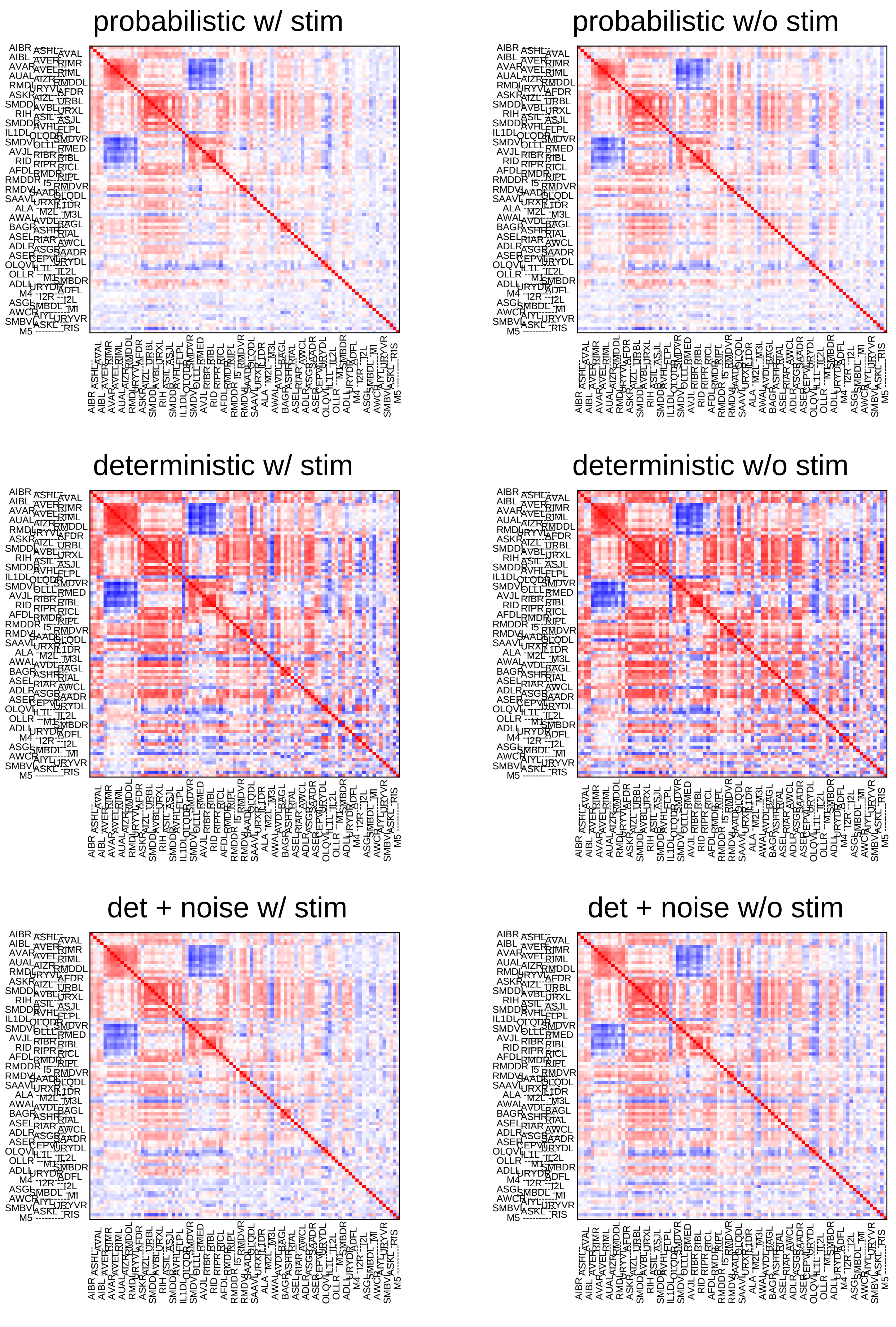
probabilistic w/ stim
probabilistic w/o stim
deterministic w/ stim
deterministic w/o stim
det + noise w/ stim
det + noise w/o stim

### Sample4

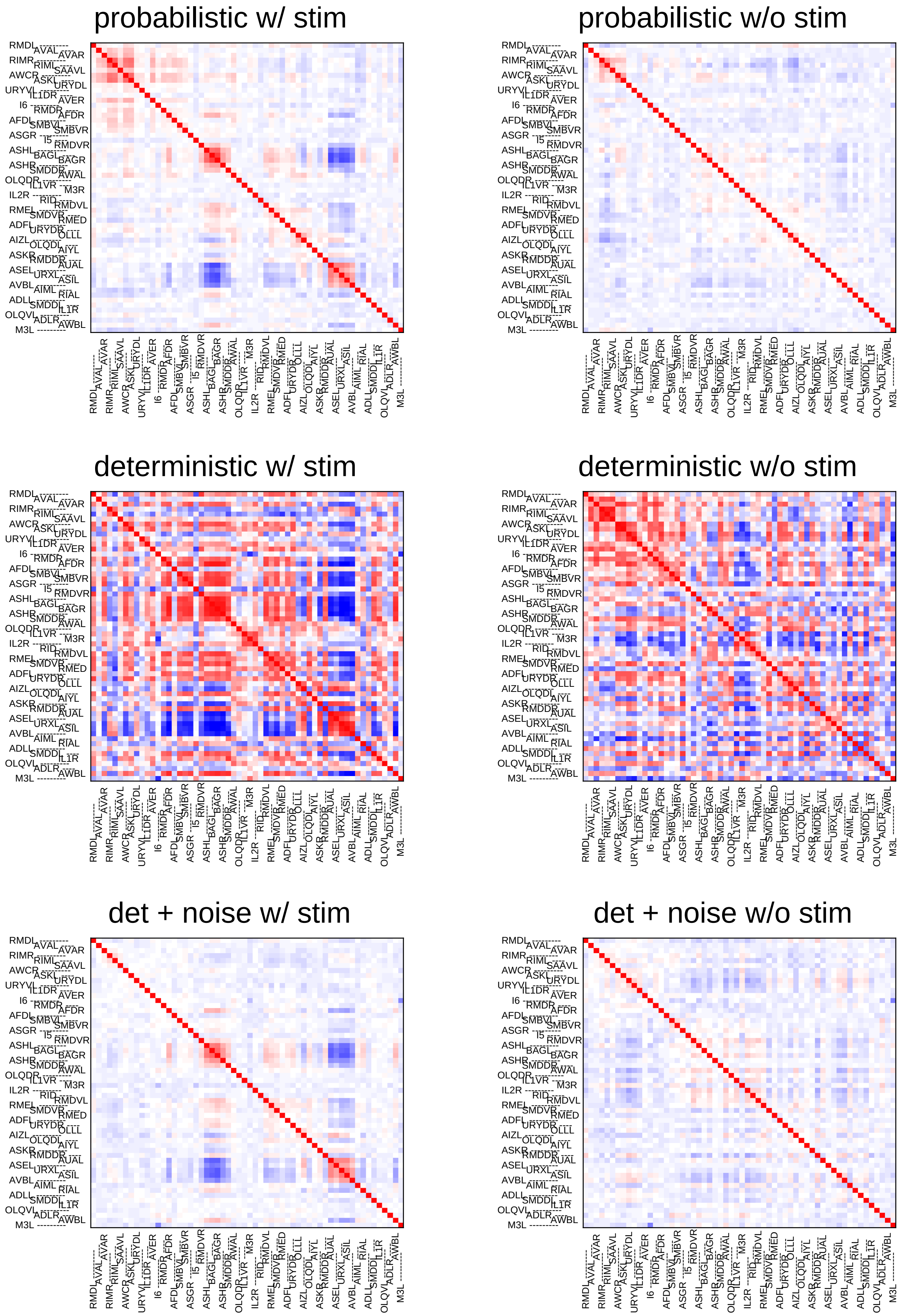
probabilistic w/ stim
probabilistic w/o stim
deterministic w/ stim
deterministic w/o stim
det + noise w/ stim
det + noise w/o stim

### Sample5

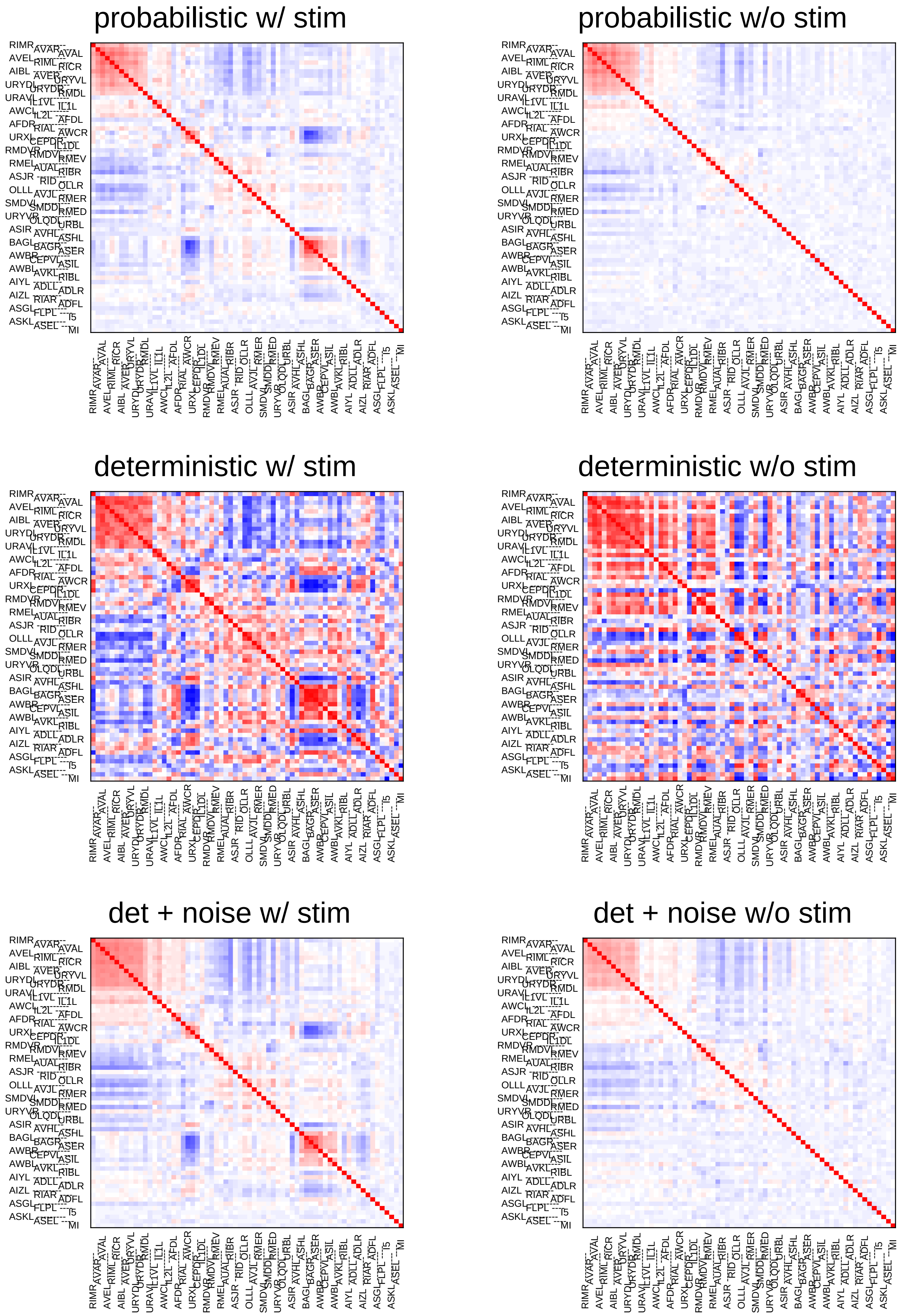
probabilistic w/ stim
probabilistic w/o stim
deterministic w/ stim
deterministic w/o stim
det + noise w/ stim
det + noise w/o stim

### Sample6

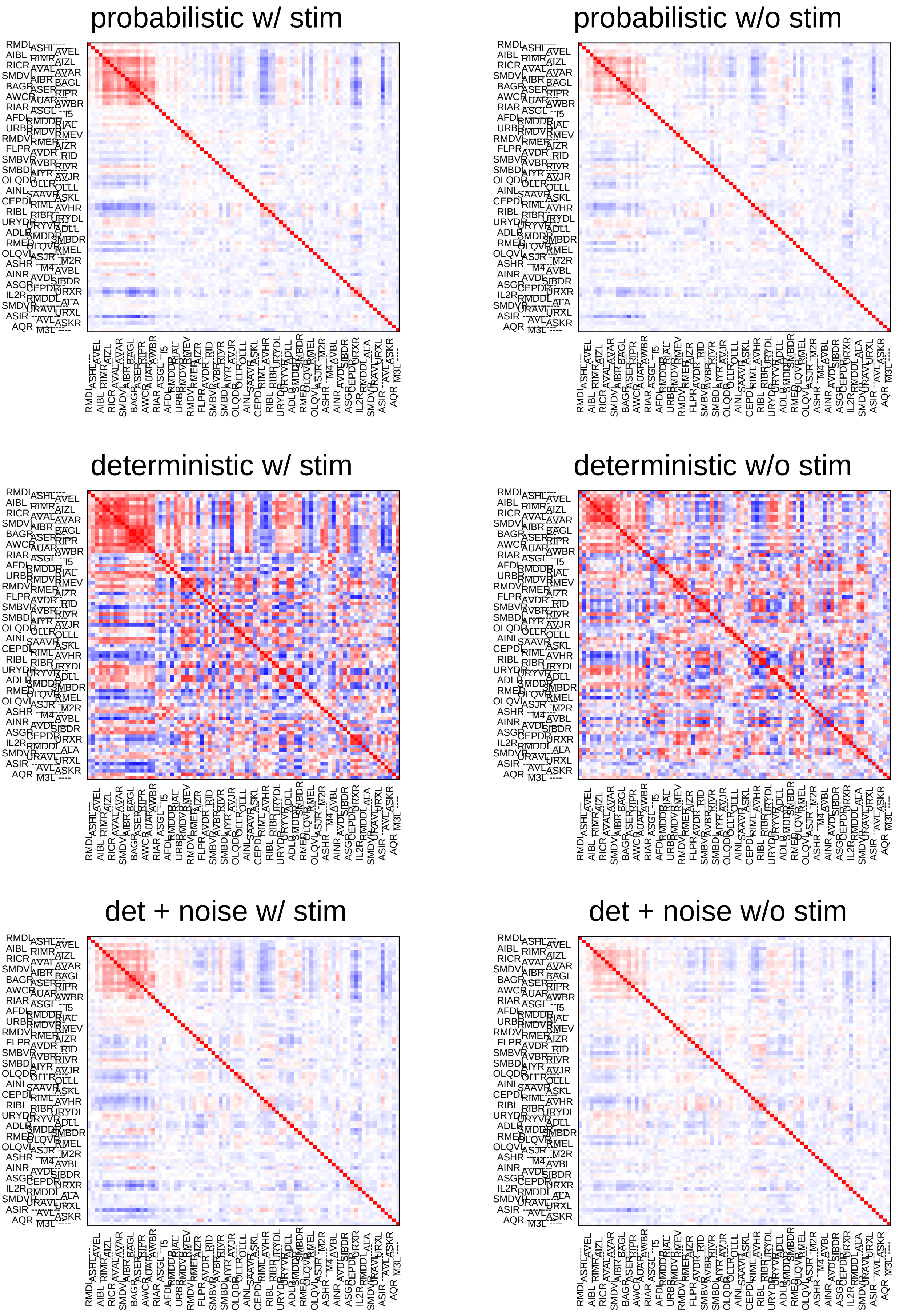
probabilistic w/ stim
probabilistic w/o stim
deterministic w/ stim
deterministic w/o stim
det + noise w/ stim
det + noise w/o stim

### Sample7

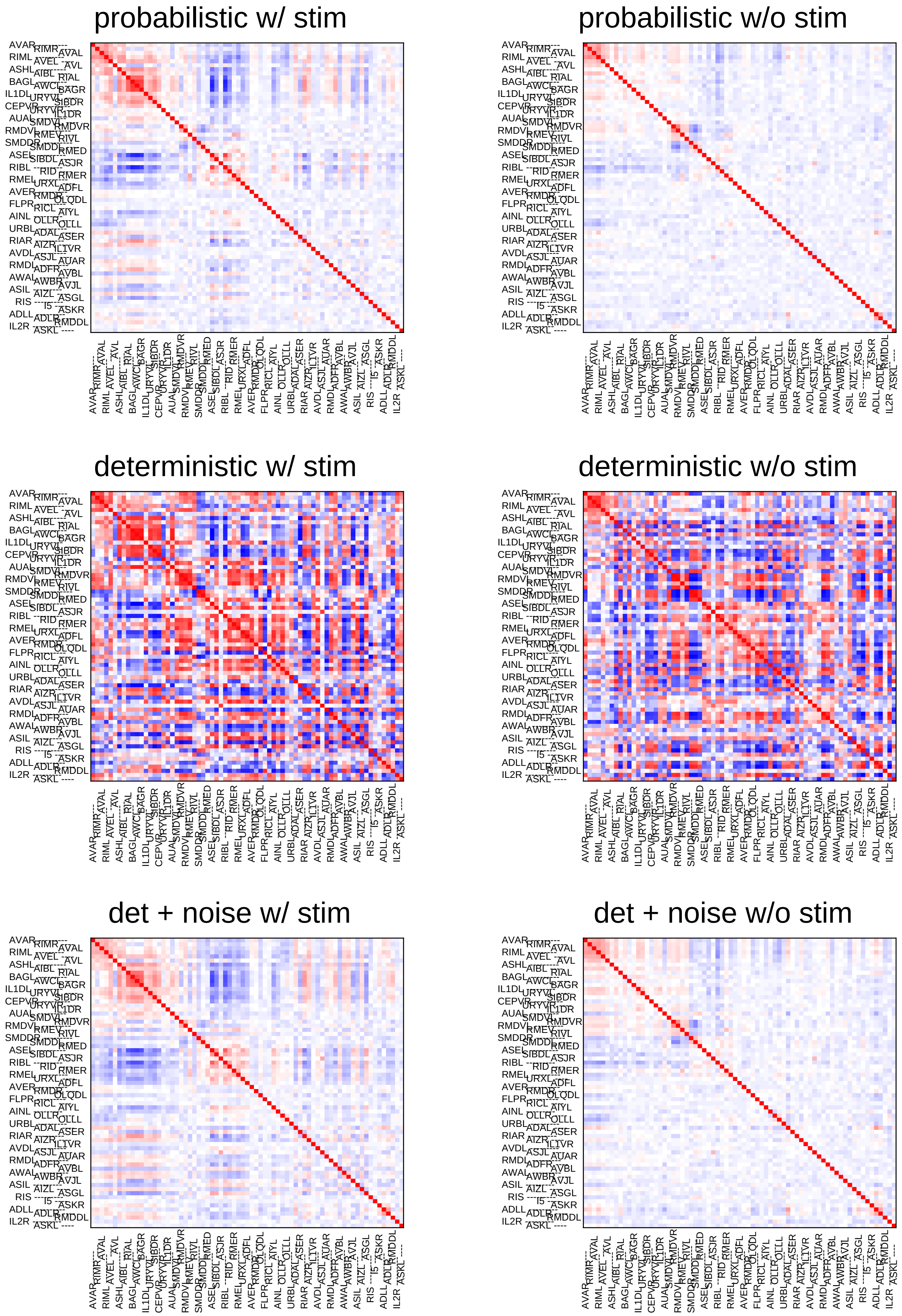
probabilistic w/ stim
probabilistic w/o stim
deterministic w/ stim
deterministic w/o stim
det + noise w/ stim
det + noise w/o stim

### Sample8

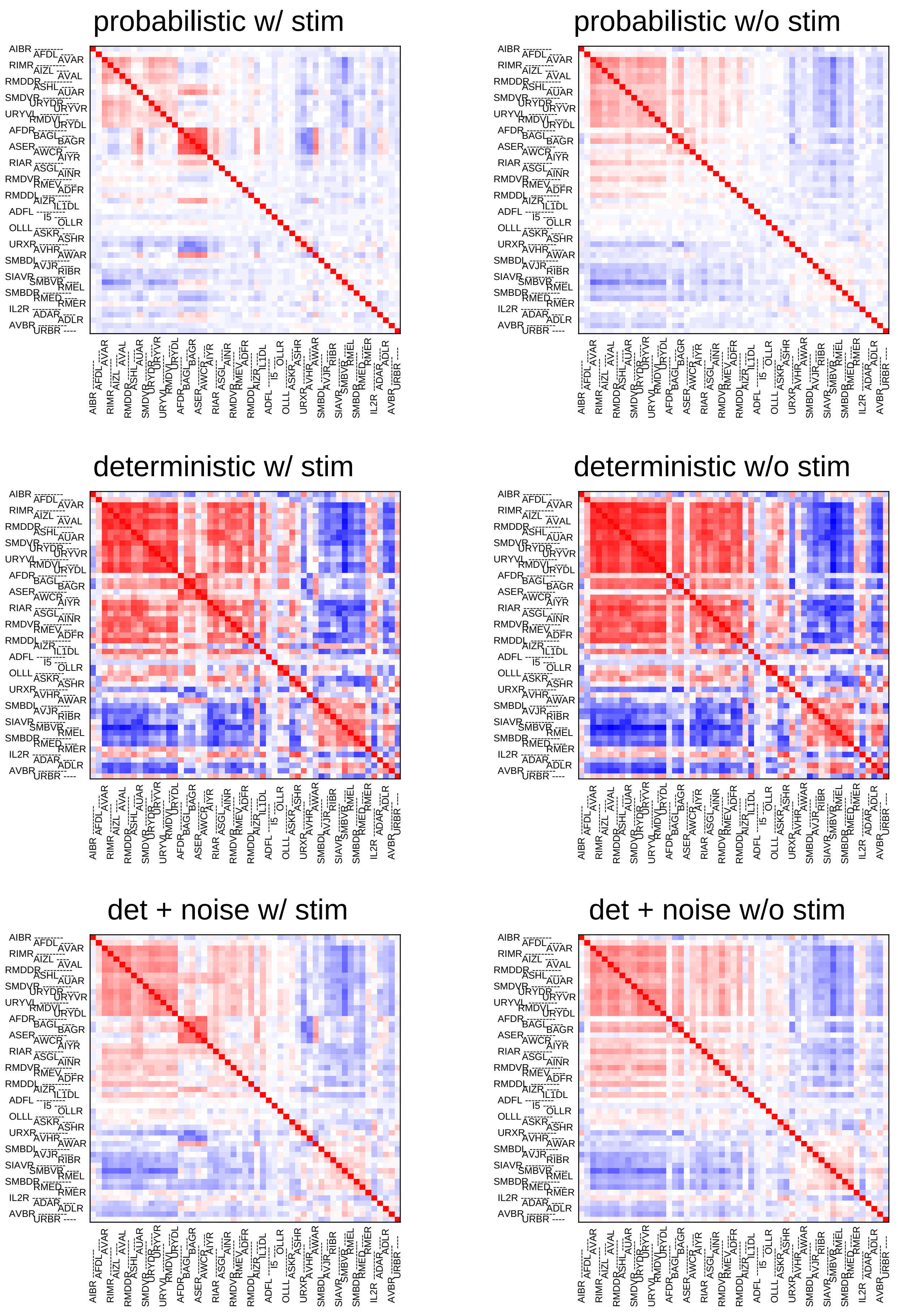
probabilistic w/ stim
probabilistic w/o stim
deterministic w/ stim
deterministic w/o stim
det + noise w/ stim
det + noise w/o stim

### Sample9

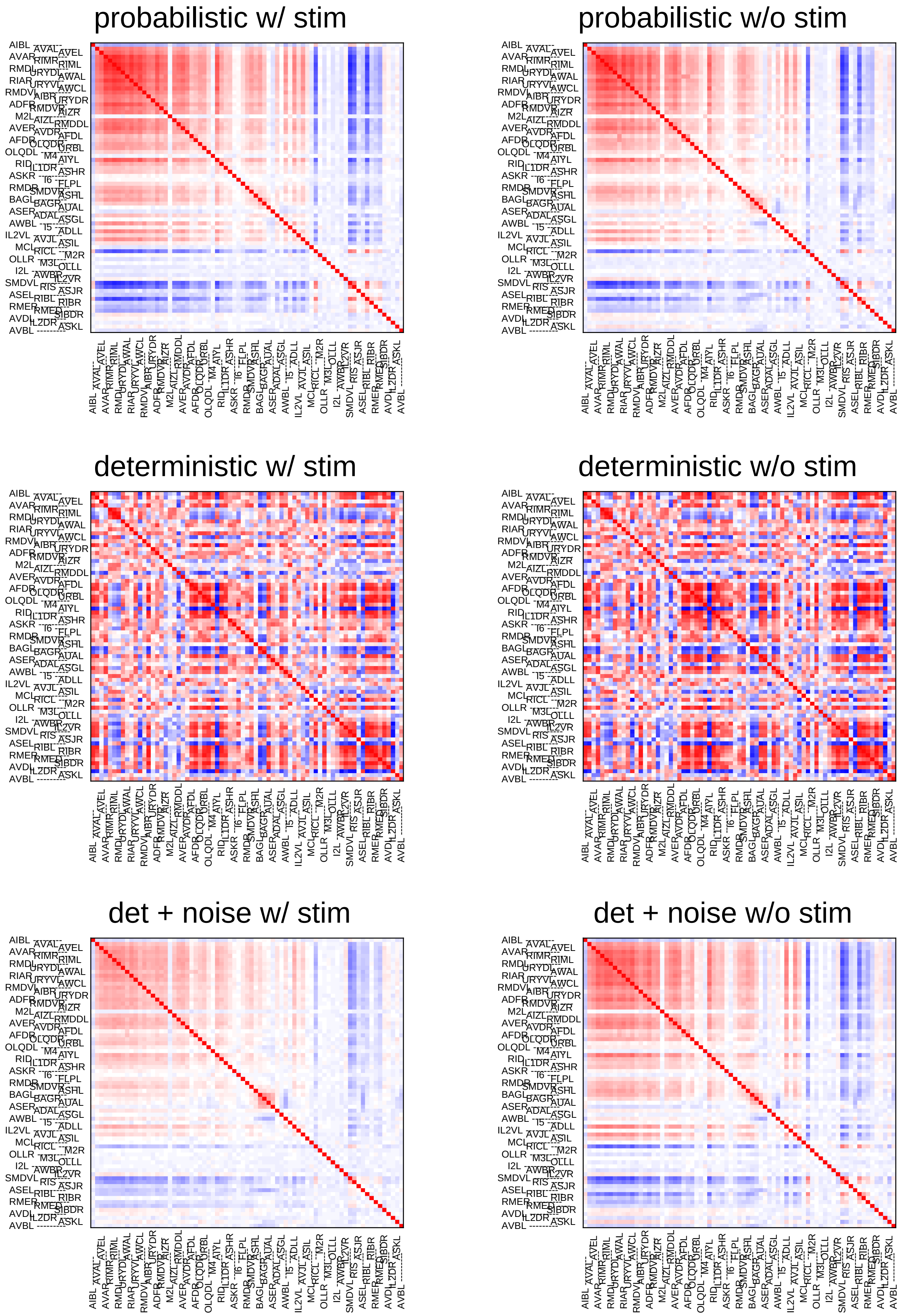
probabilistic w/ stim
probabilistic w/o stim
deterministic w/ stim
deterministic w/o stim
det + noise w/ stim
det + noise w/o stim

### Sample10

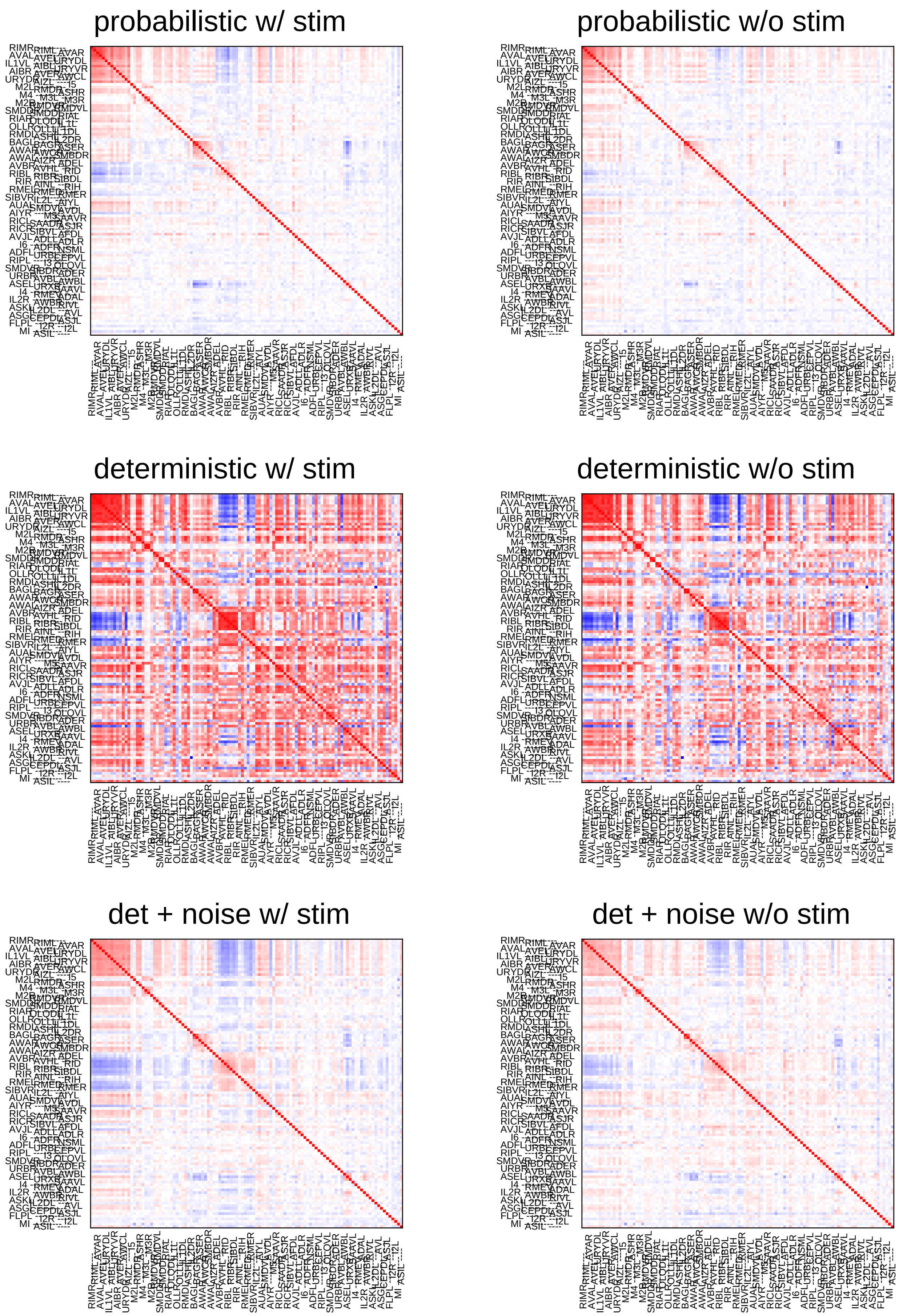

probabilistic w/ stim
probabilistic w/o stim
deterministic w/ stim
deterministic w/o stim
det + noise w/ stim
det + noise w/o stim

### Sample11

#### probabilistic w/ stim

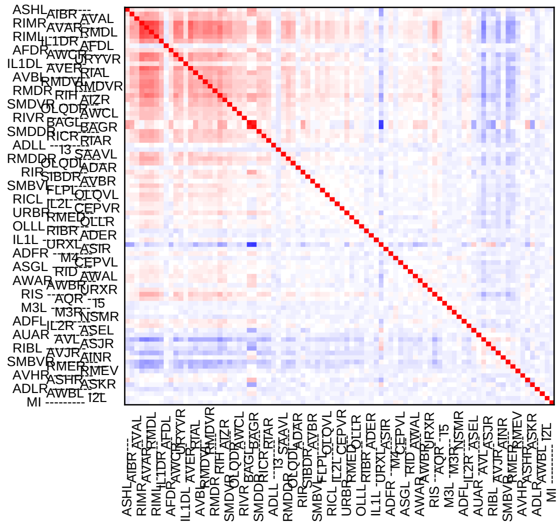

#### probabilistic w/o stim

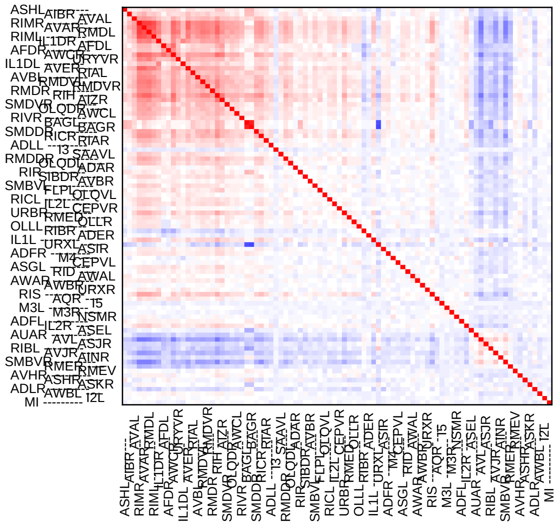

#### deterministic w/ stim

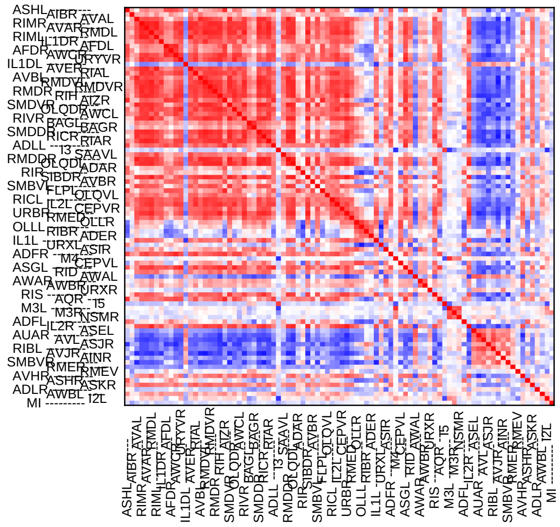

#### deterministic w/o stim

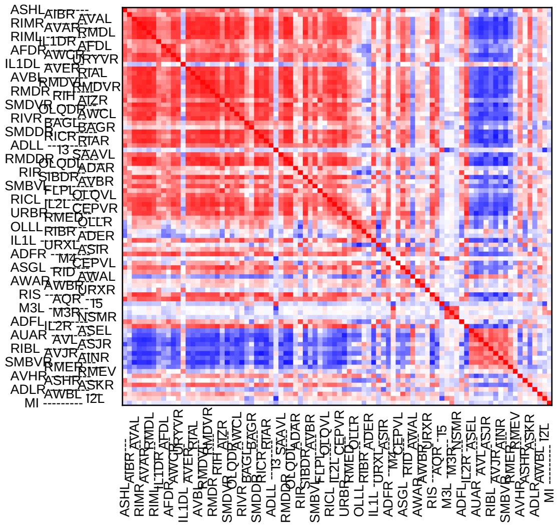

#### det + noise w/ stim

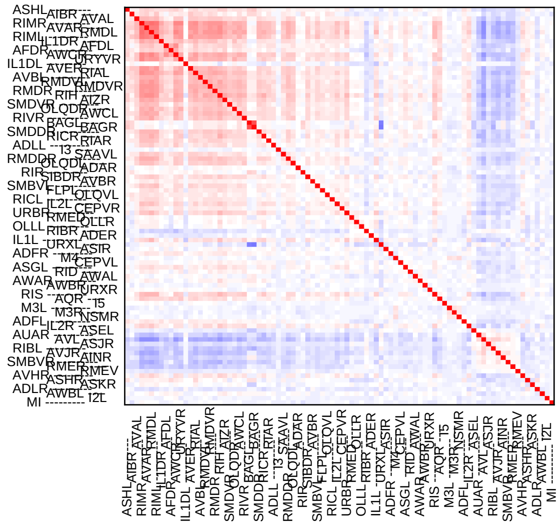

#### det + noise w/o stim

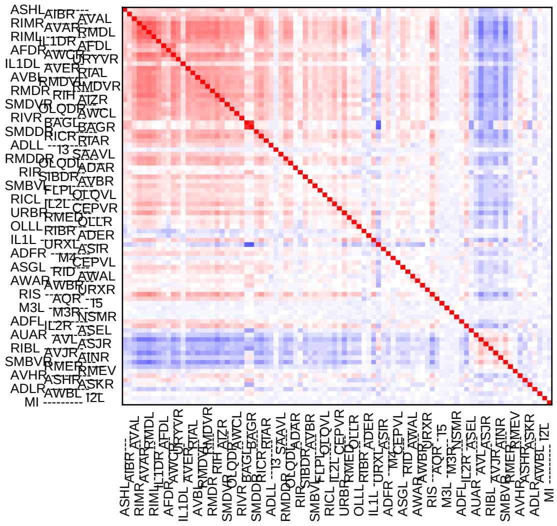

### Sample12

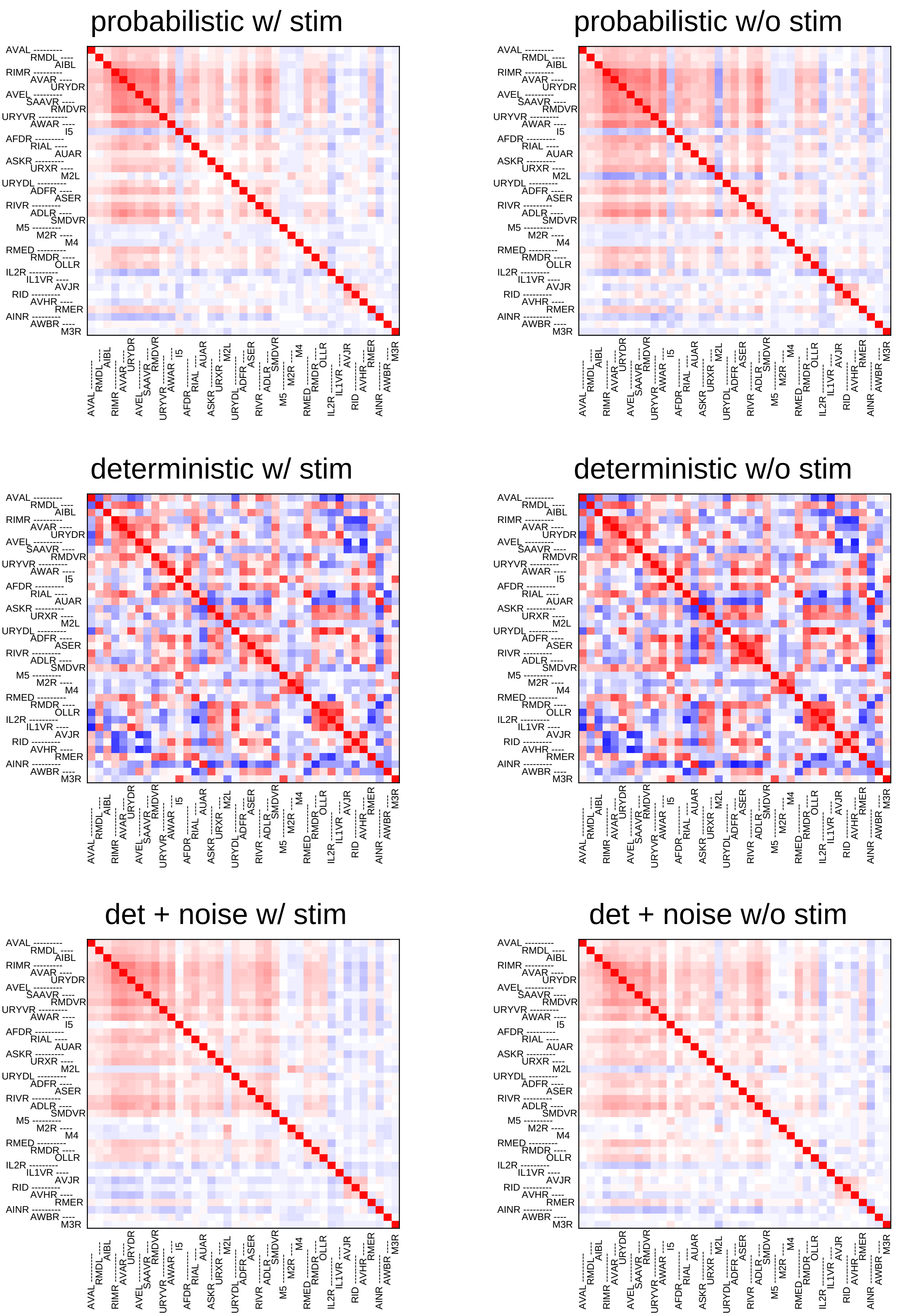
probabilistic w/ stim
probabilistic w/o stim
deterministic w/ stim
deterministic w/o stim
det + noise w/ stim
det + noise w/o stim
AVAL
RMDL
AIBL
RIMR
AVAR
URYDR
AVEL
SAAVR
RMDVR
URYVR
AWAR
I5
AFDR
RIAL
AUAR
ASKR
URXR
M2L
URYDL
ADFR
ASER
RIVR
ADLR
SMDVR
M5
M2R
M4
RMED
RMDR
OLLR
IL2R
IL1VR
AVJR
RID
AVHR
RMER
AINR
AWBR
M3R

### Sample13

#### probabilistic w/ stim

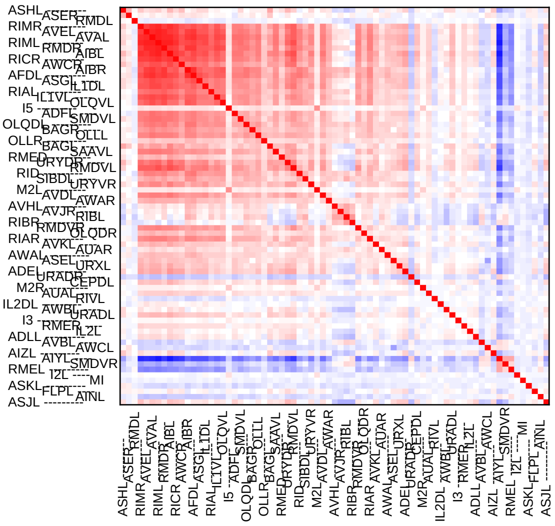

#### probabilistic w/o stim

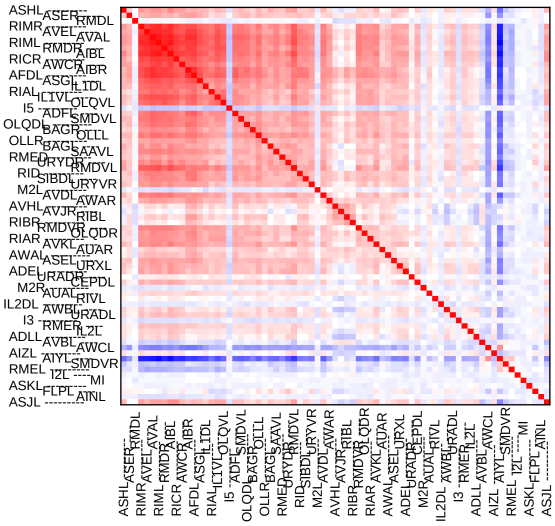

#### deterministic w/ stim

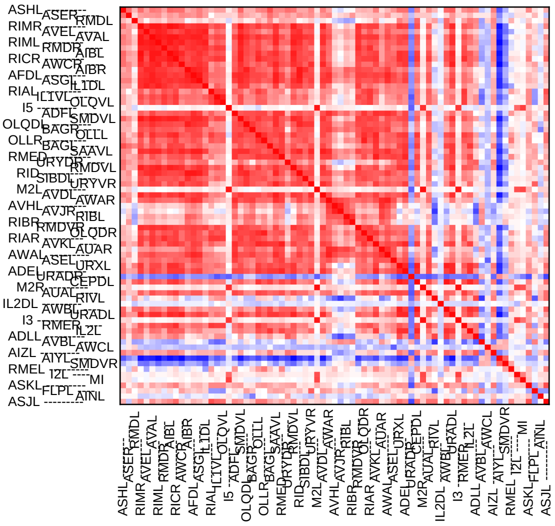

#### deterministic w/o stim

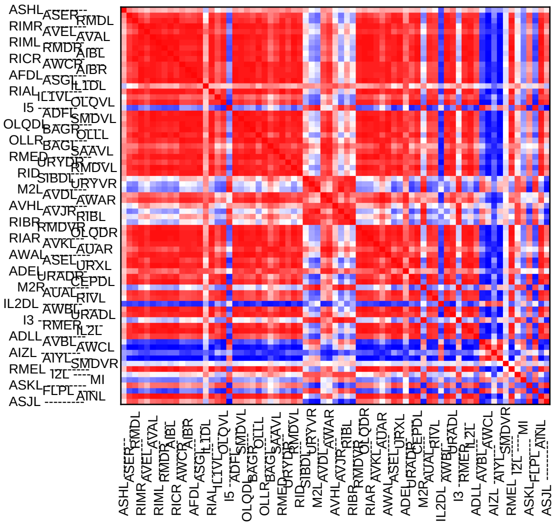

#### det + noise w/ stim

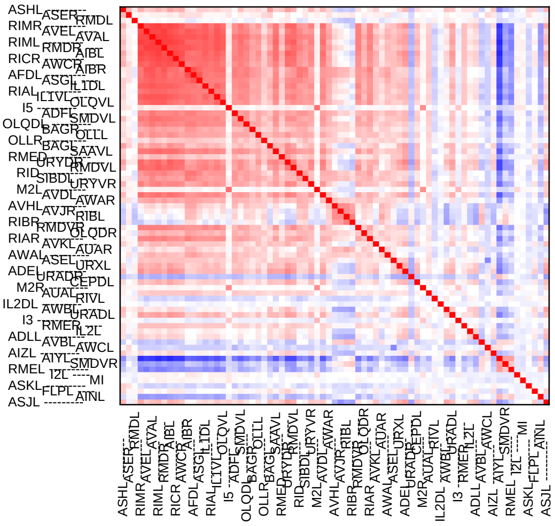

#### det + noise w/o stim

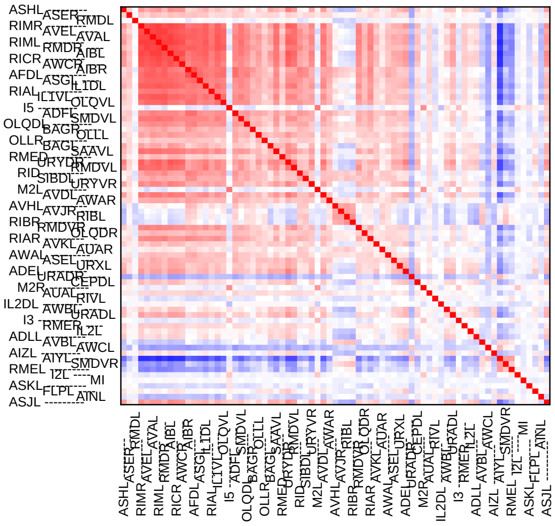

### Sample14

#### probabilistic w/ stim

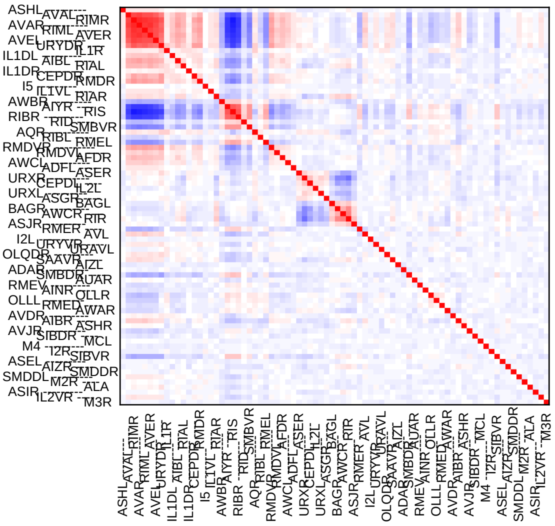

#### probabilistic w/o stim

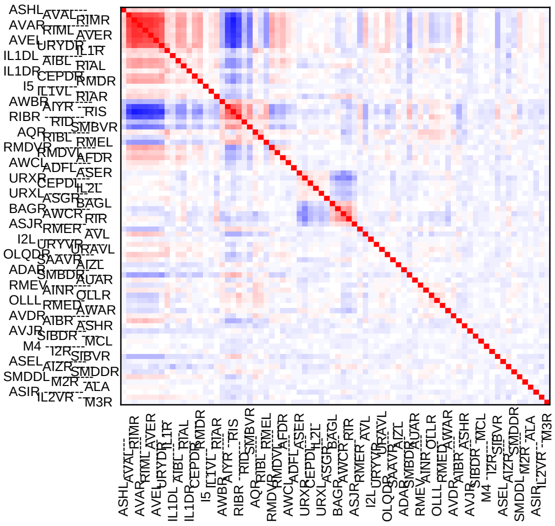

#### deterministic w/ stim

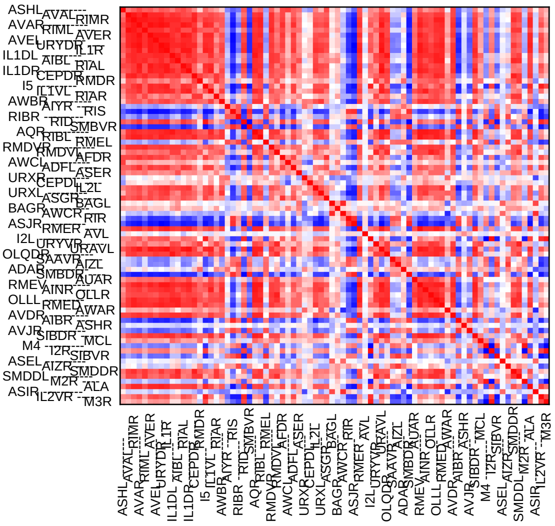

#### deterministic w/o stim

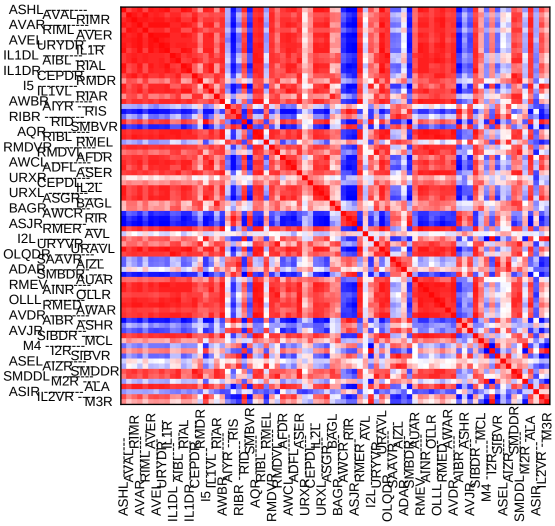

#### det + noise w/ stim

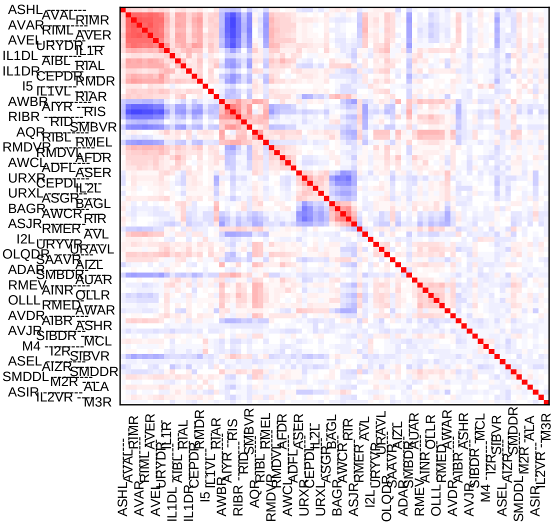

#### det + noise w/o stim

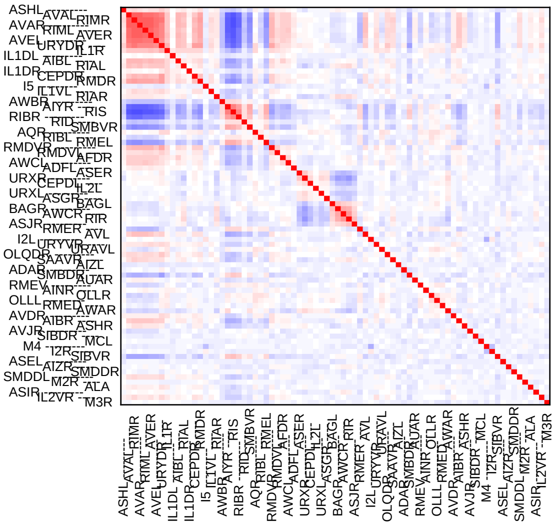

### Sample15

#### probabilistic w/ stim

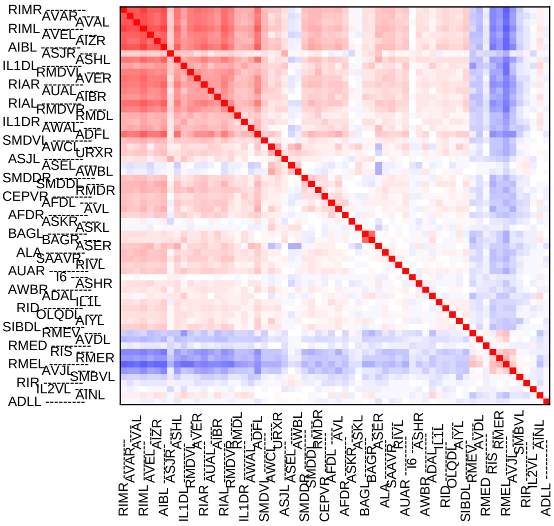


#### probabilistic w/o stim

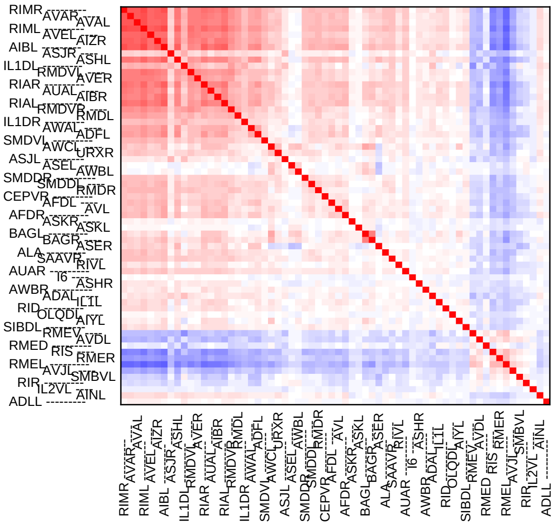


#### deterministic w/ stim

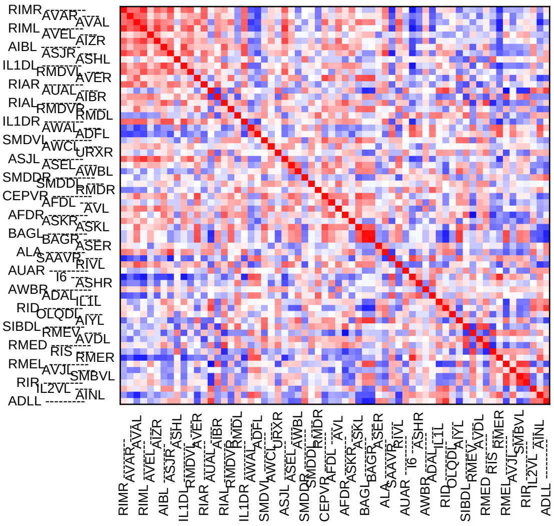


#### deterministic w/o stim

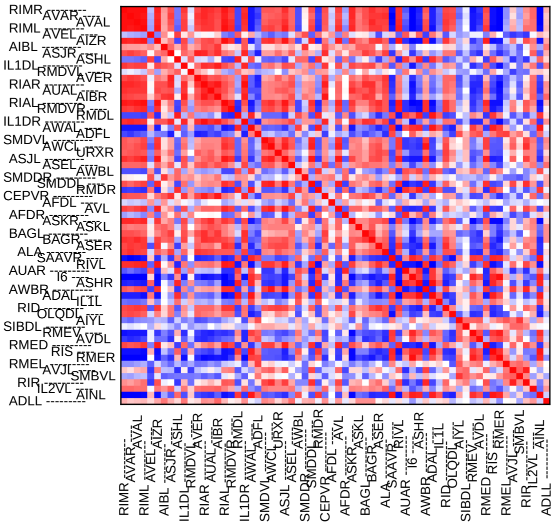


#### det + noise w/ stim

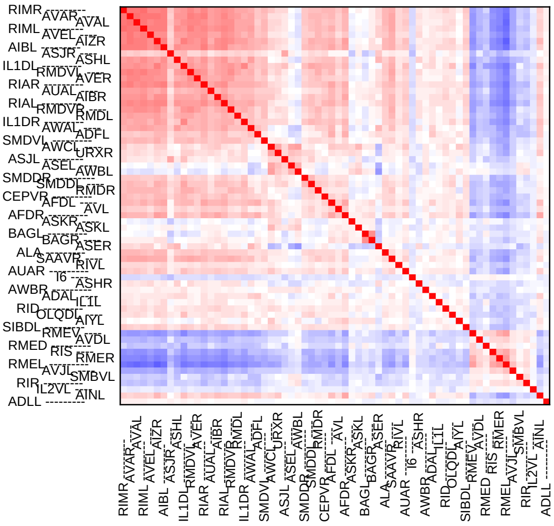


#### det + noise w/o stim

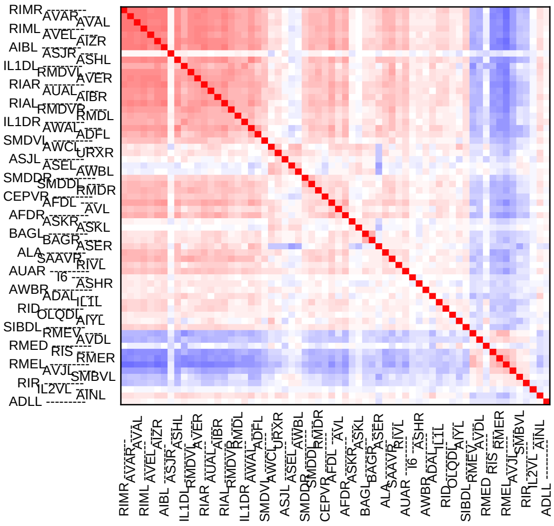

### Sample16

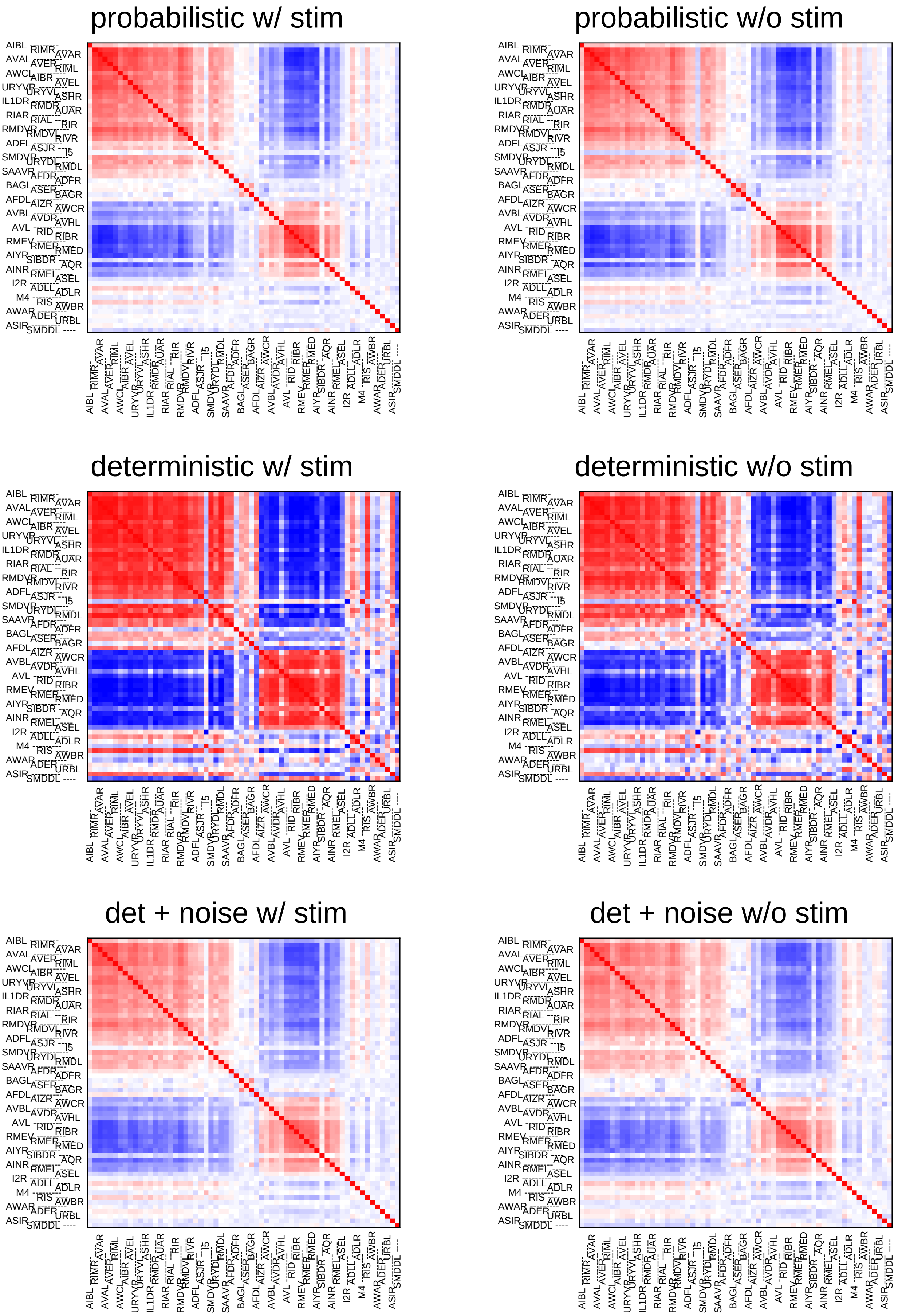
probabilistic w/ stim
probabilistic w/o stim
deterministic w/ stim
deterministic w/o stim
det + noise w/ stim
det + noise w/o stim

### Sample17

#### probabilistic w/ stim

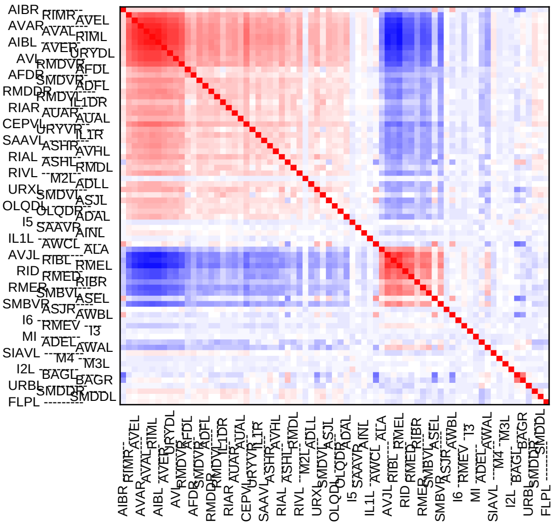

#### probabilistic w/o stim

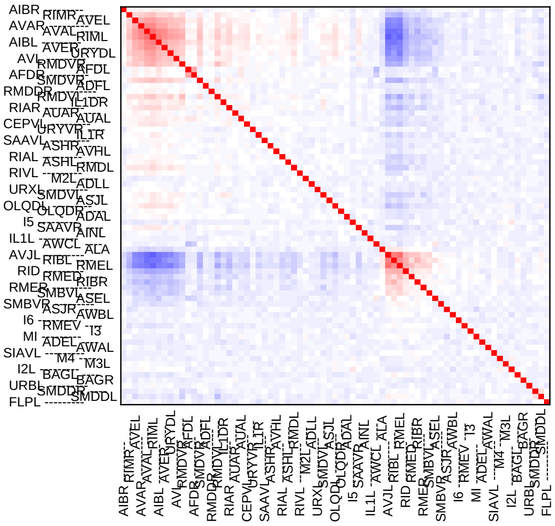

#### deterministic w/ stim

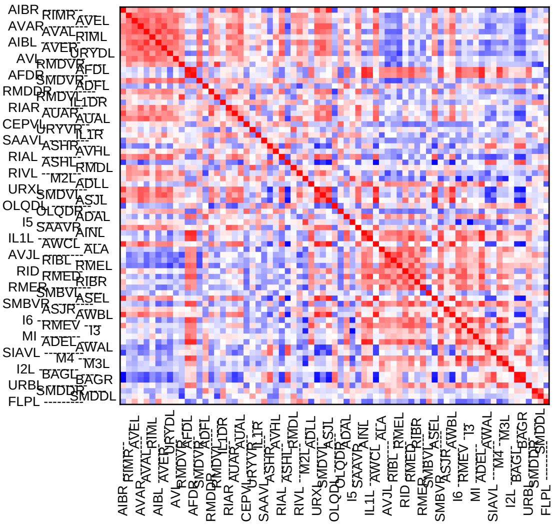

#### deterministic w/o stim

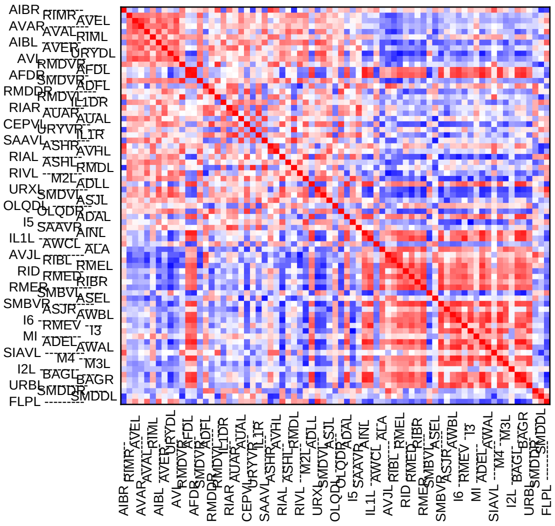

#### det + noise w/ stim

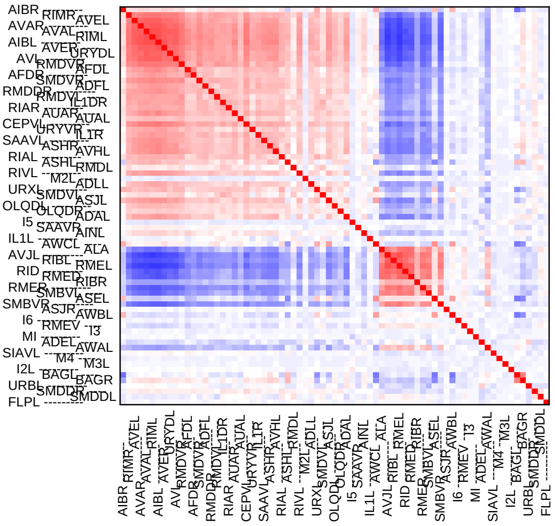

#### det + noise w/o stim

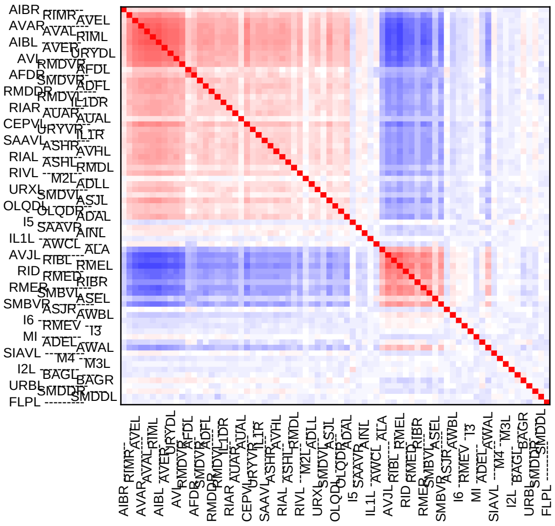

### Sample18

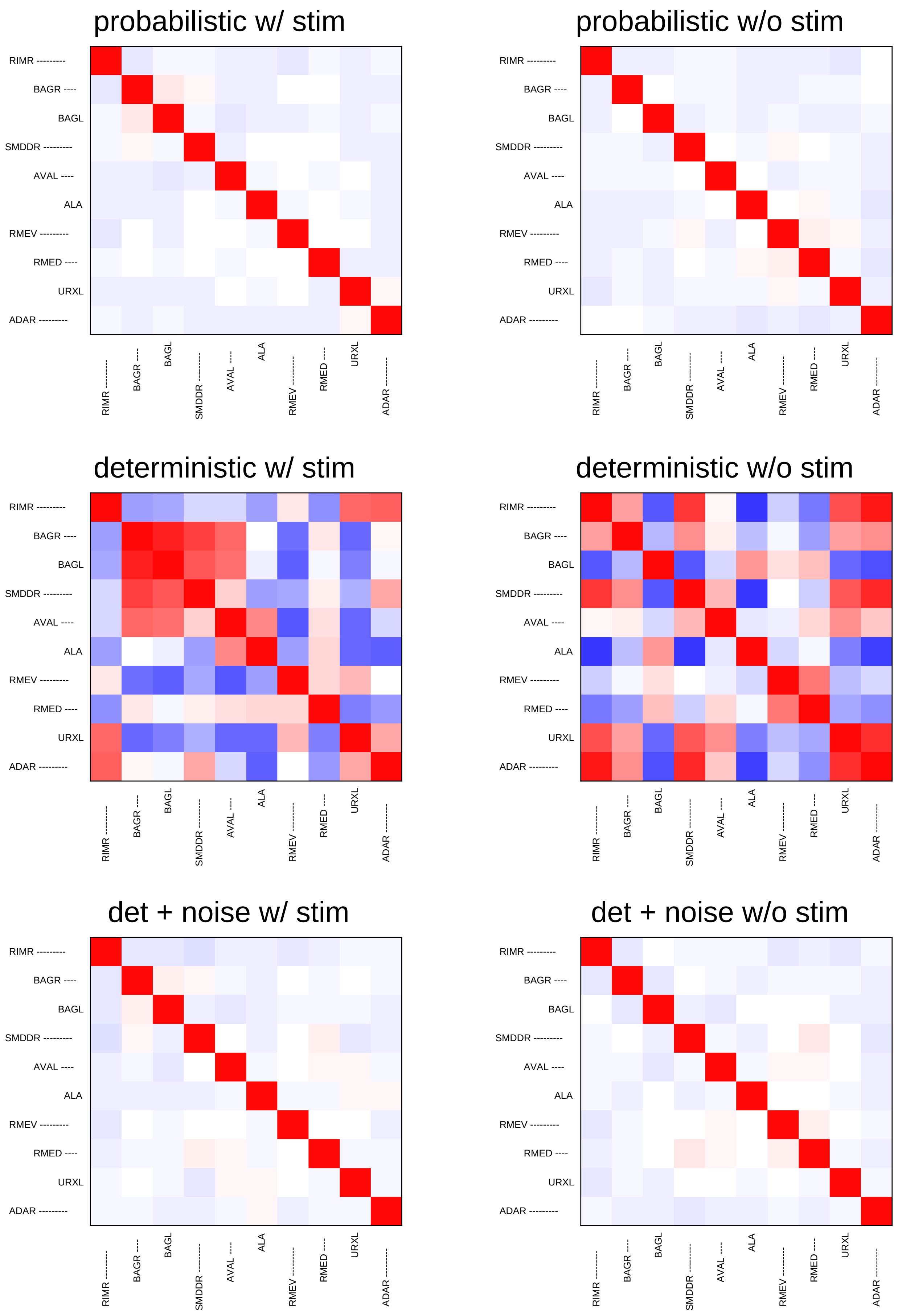
probabilistic w/ stim
probabilistic w/o stim
deterministic w/ stim
deterministic w/o stim
det + noise w/ stim
det + noise w/o stim
RIMR ---------
BAGR ----
BAGL
SMDDR ---------
AVAL ----
ALA
RMEV ---------
RMED ----
URXL
ADAR ---------

### Sample19

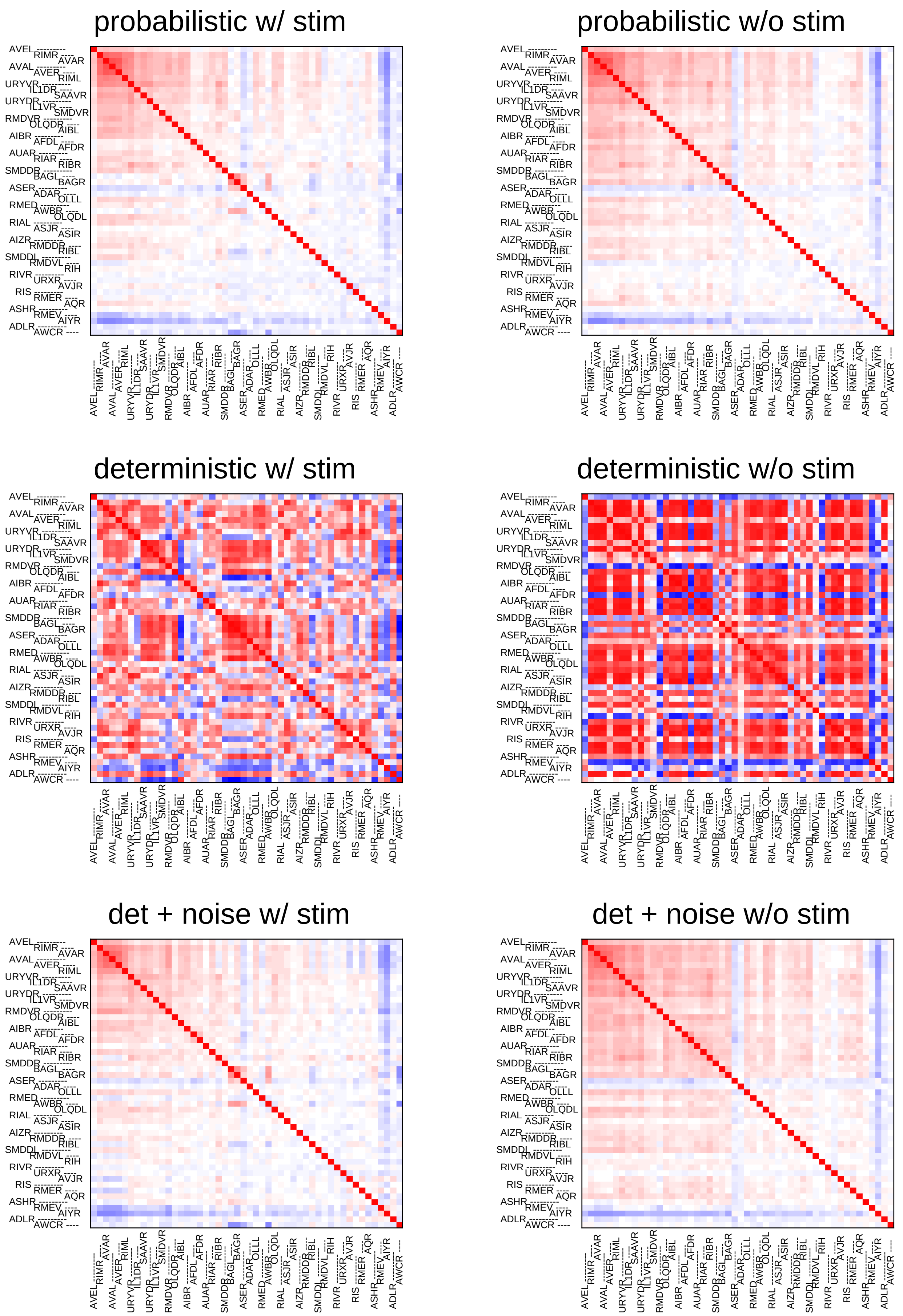
probabilistic w/ stim
probabilistic w/o stim
deterministic w/ stim
deterministic w/o stim
det + noise w/ stim
det + noise w/o stim
AVEL RIMR AVAR AVAL AVER RIML URYVR IL1DR SAAVR URYDR IL1VR SMDVR RMDVR OLQDR AIBL AIBR AFDL AFDR AUAR RIAR RIBR SMDDR BAGL BAGR ASER ADAR OLLL RMED AWBR OLQDL RIAL ASJR ASIR AIZR RMDDR RIBL SMDDL RMDVL RIH RIVR URXR AVJR RIS RMER AQR ASHR RMEV AIYR ADLR AWCR

### Sample20

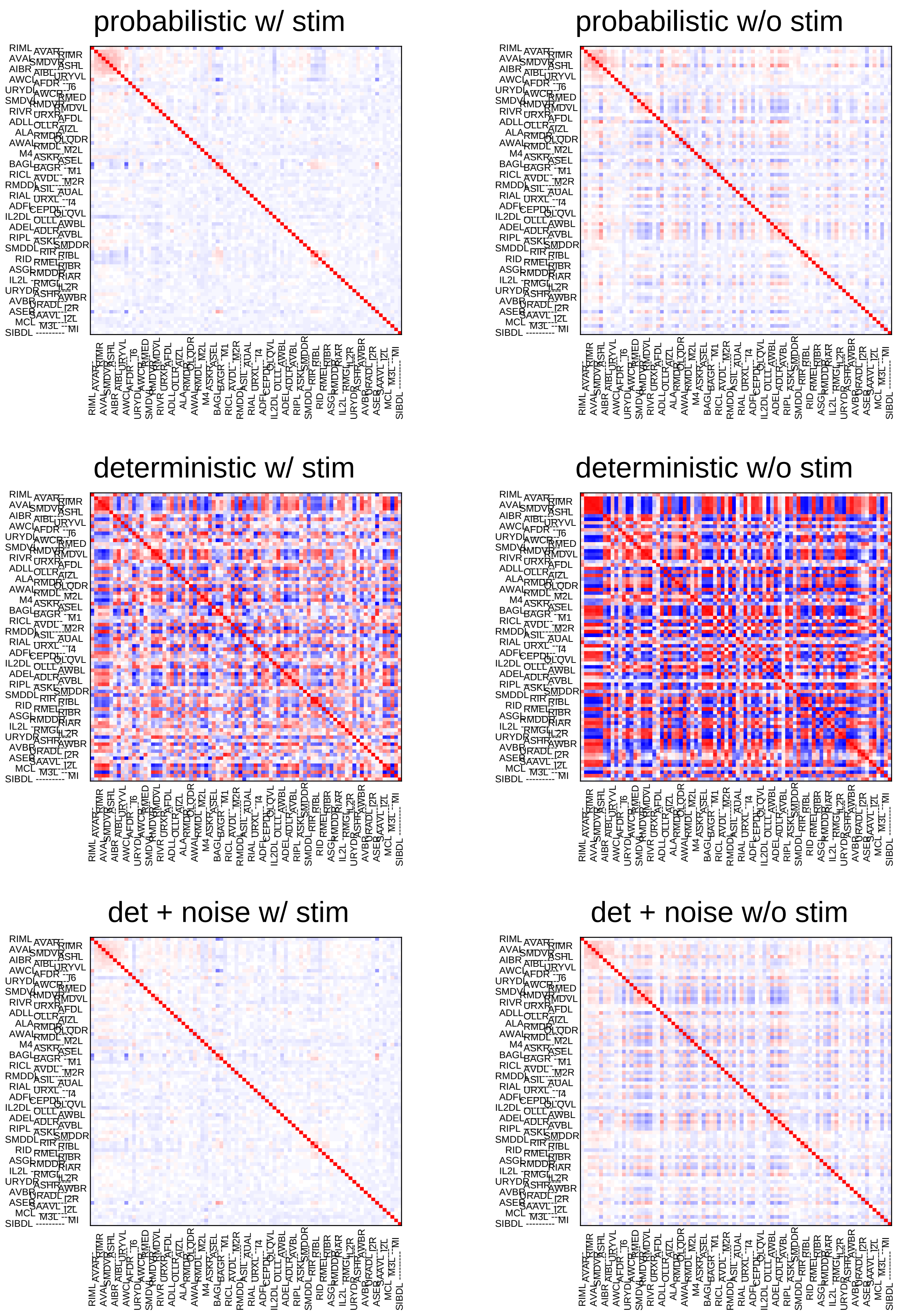
probabilistic w/ stim
probabilistic w/o stim
deterministic w/ stim
deterministic w/o stim
det + noise w/ stim
det + noise w/o stim

### Sample21

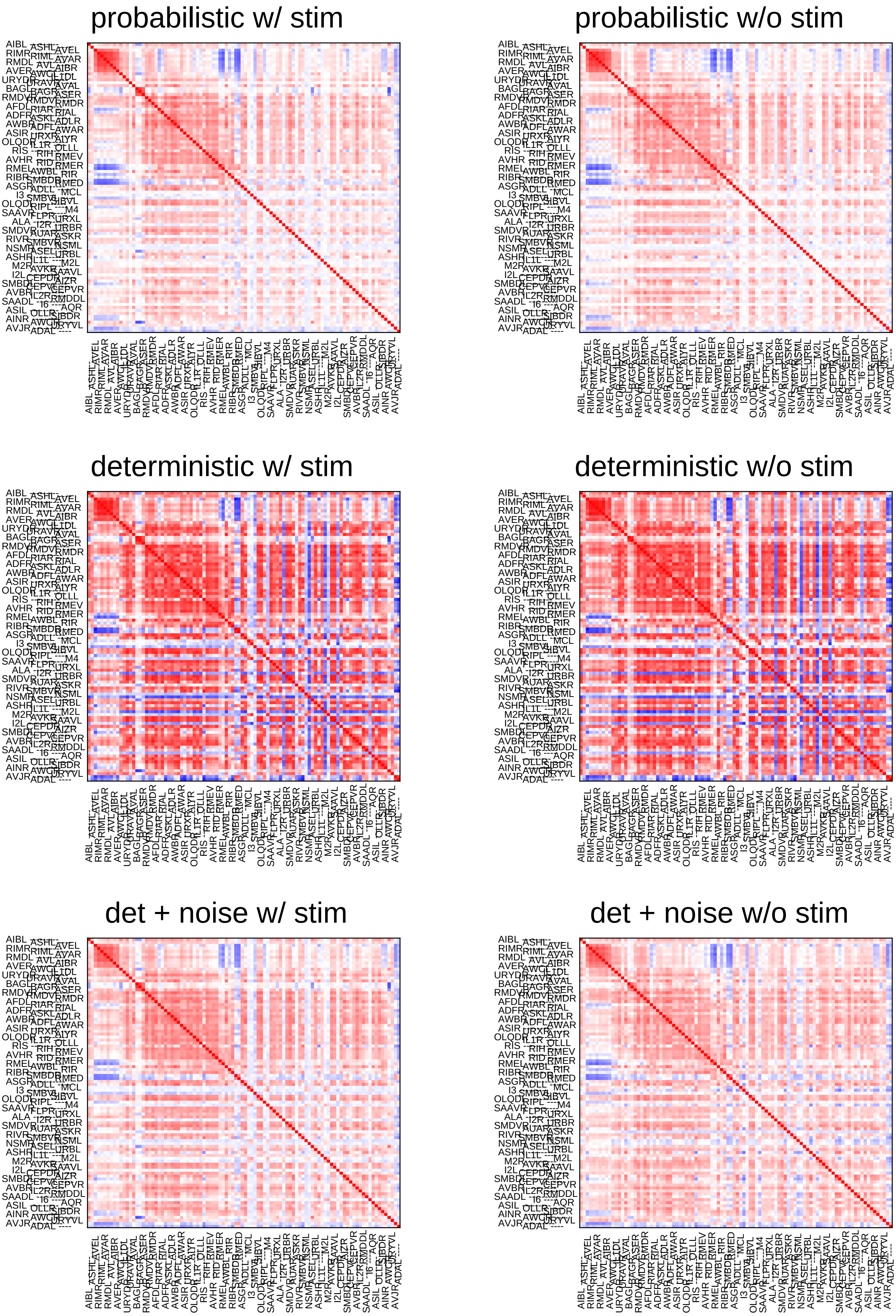

probabilistic w/ stim
probabilistic w/o stim
deterministic w/ stim
deterministic w/o stim
det + noise w/ stim
det + noise w/o stim

### Sample22

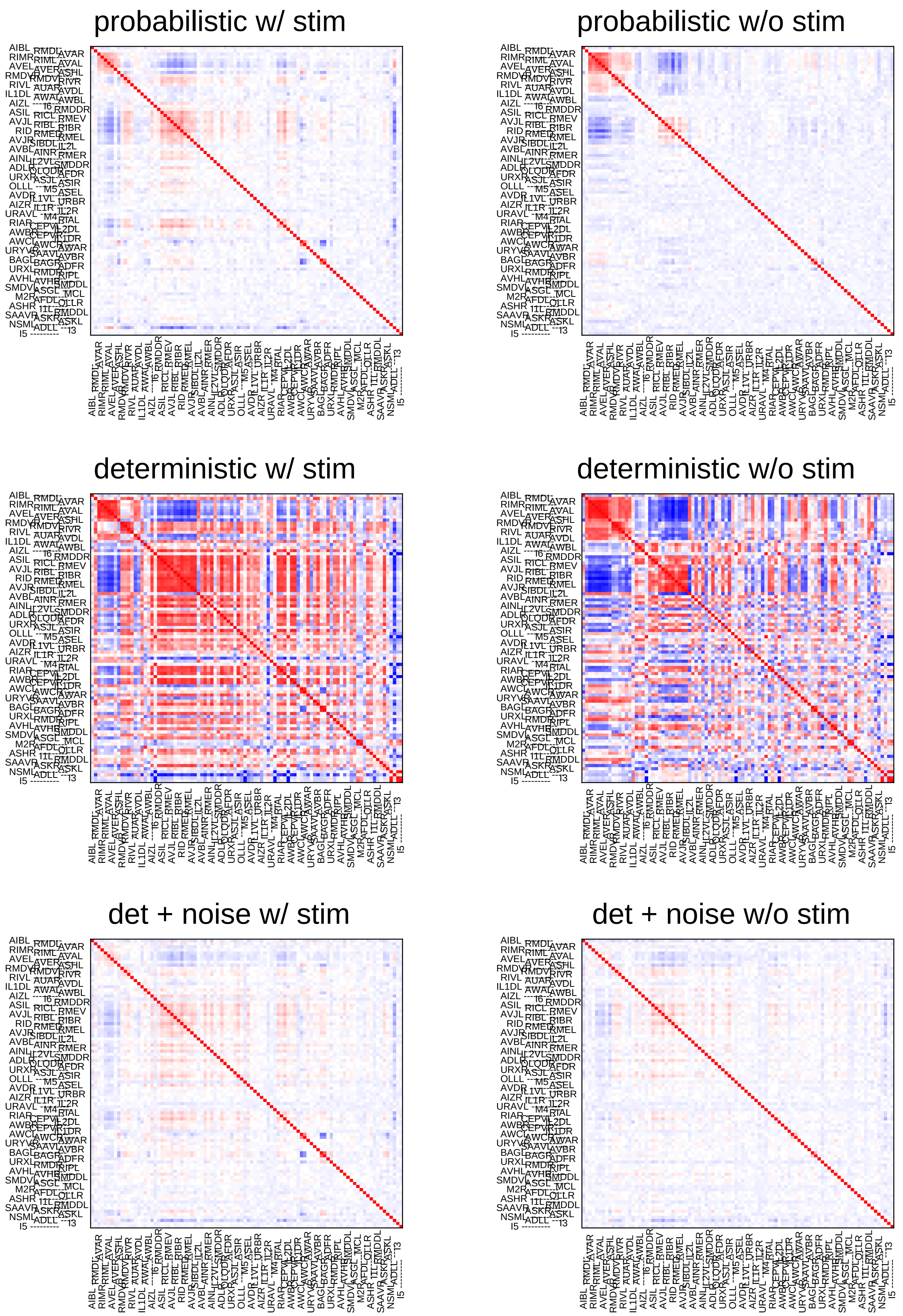

probabilistic w/ stim
probabilistic w/o stim
deterministic w/ stim
deterministic w/o stim
det + noise w/ stim
det + noise w/o stim

### Sample23

#### probabilistic w/ stim

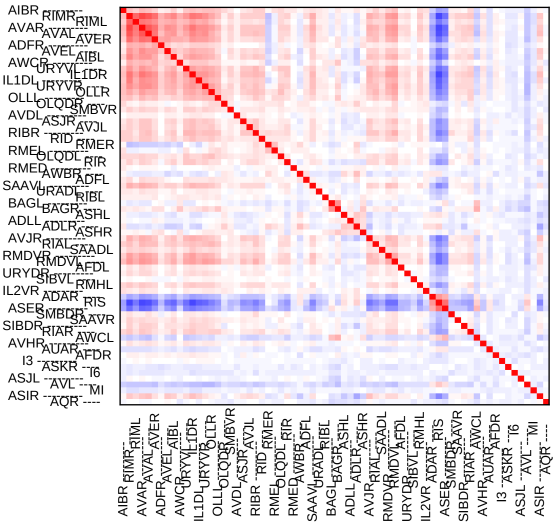

#### probabilistic w/o stim

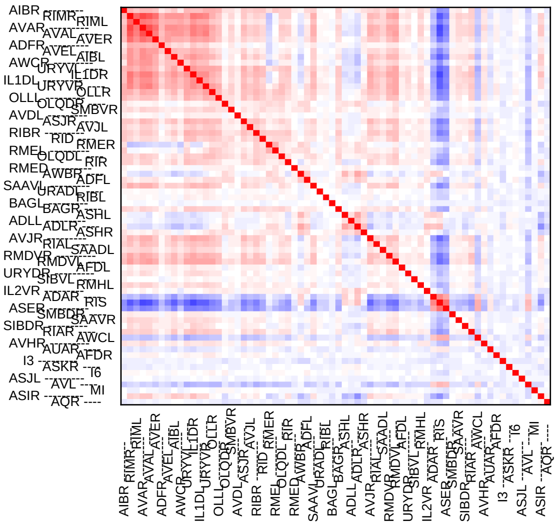

#### deterministic w/ stim

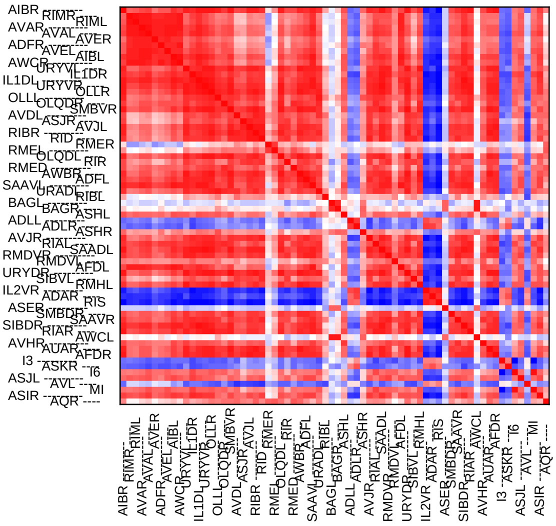

#### deterministic w/o stim

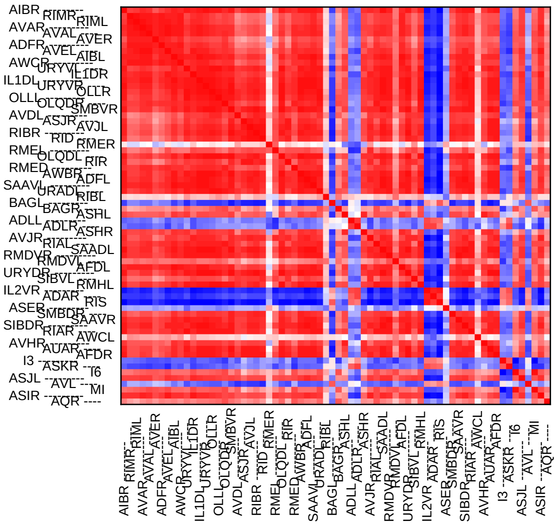

#### det + noise w/ stim

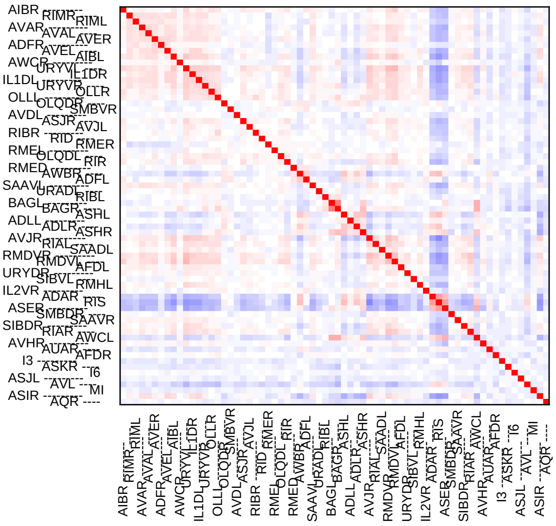

#### det + noise w/o stim

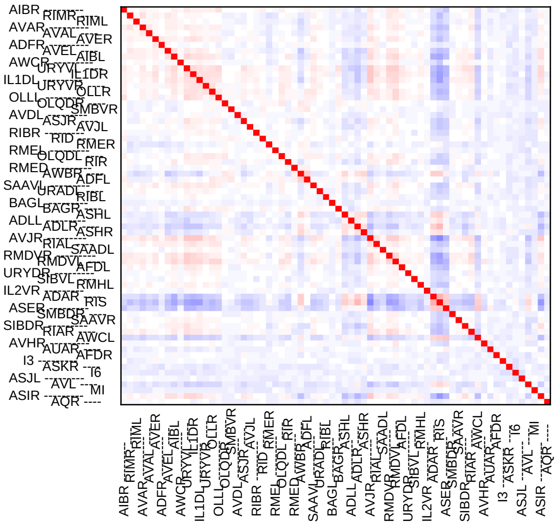

### Sample24

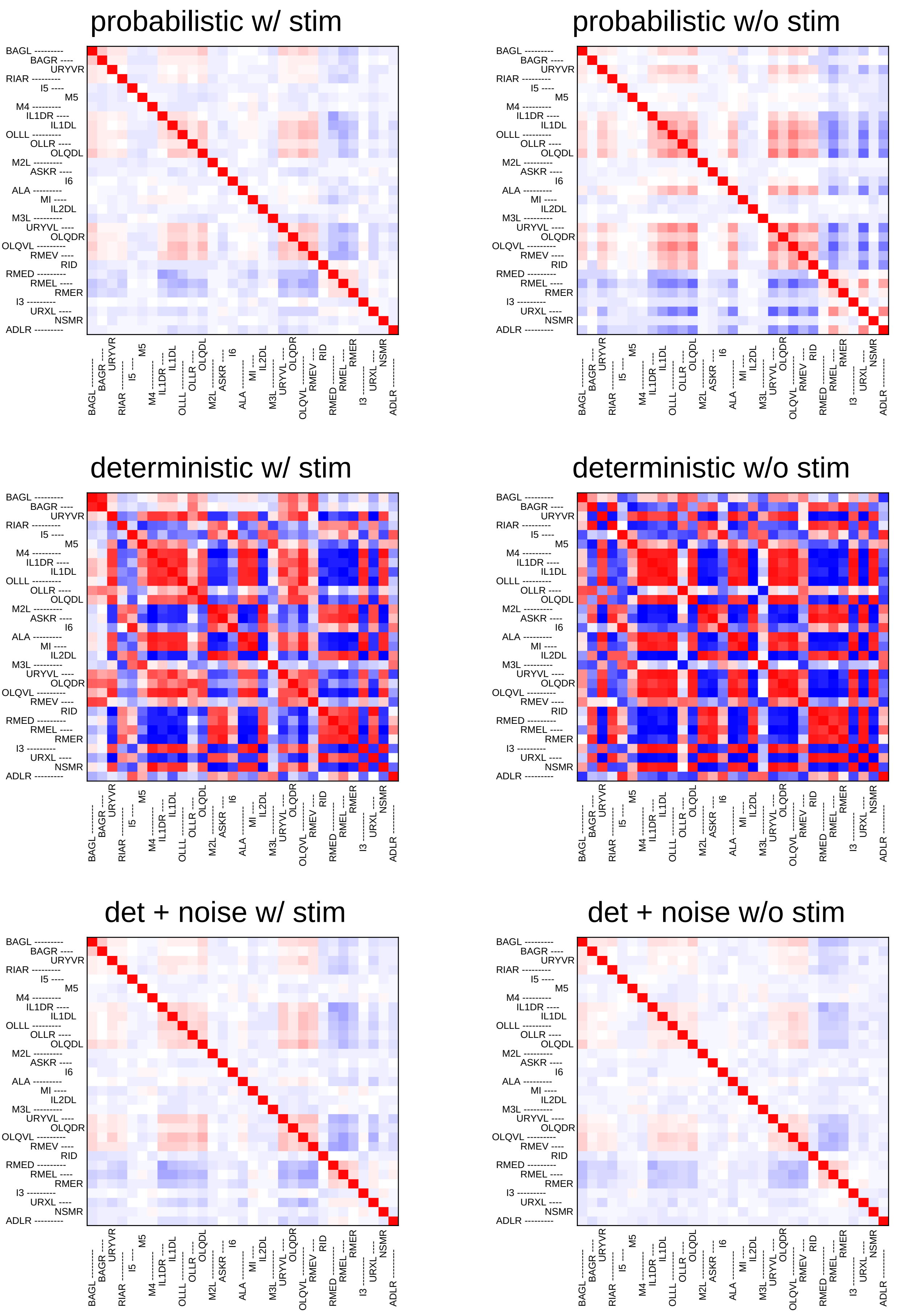
probabilistic w/ stim
probabilistic w/o stim
deterministic w/ stim
deterministic w/o stim
det + noise w/ stim
det + noise w/o stim
