## Supplementary File 10 for "Deducing ensemble dynamics and information flow from the whole-brain imaging data"

### Sample1

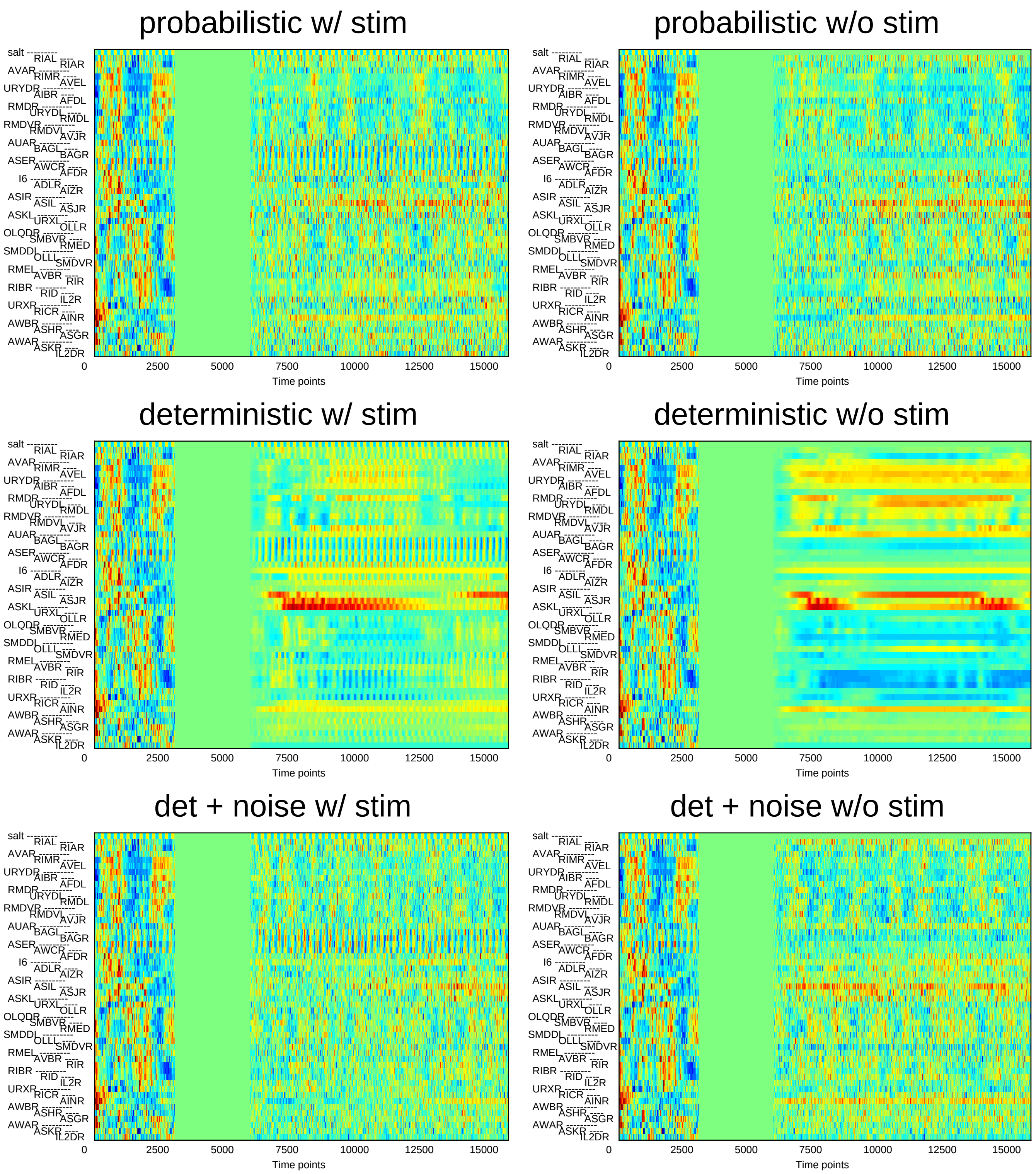
probabilistic w/ stim
probabilistic w/o stim
deterministic w/ stim
deterministic w/o stim
det + noise w/ stim
det + noise w/o stim
salt
RIAL
RIAR
AVAR
RIMR
AVEL
URYDR
AIBR
AFDL
RMDR
URYDL
RMDL
RMDVR
RMDVL
AVJR
AUAR
BAGL
BAGR
ASER
AWCR
AFDR
I6
ADLR
AIZR
ASIR
ASIL
ASJR
ASKL
URXL
OLLR
OLQDR
SMBVR
RMED
SMDDL
OLLL
SMDVR
RMEL
AVBR
RIR
RIBR
RID
IL2R
URXR
RICR
AINR
AWBR
ASHR
ASGR
AWAR
ASKR
IL2DR
0
2500
5000
7500
10000
12500
15000
Time points

### Sample2

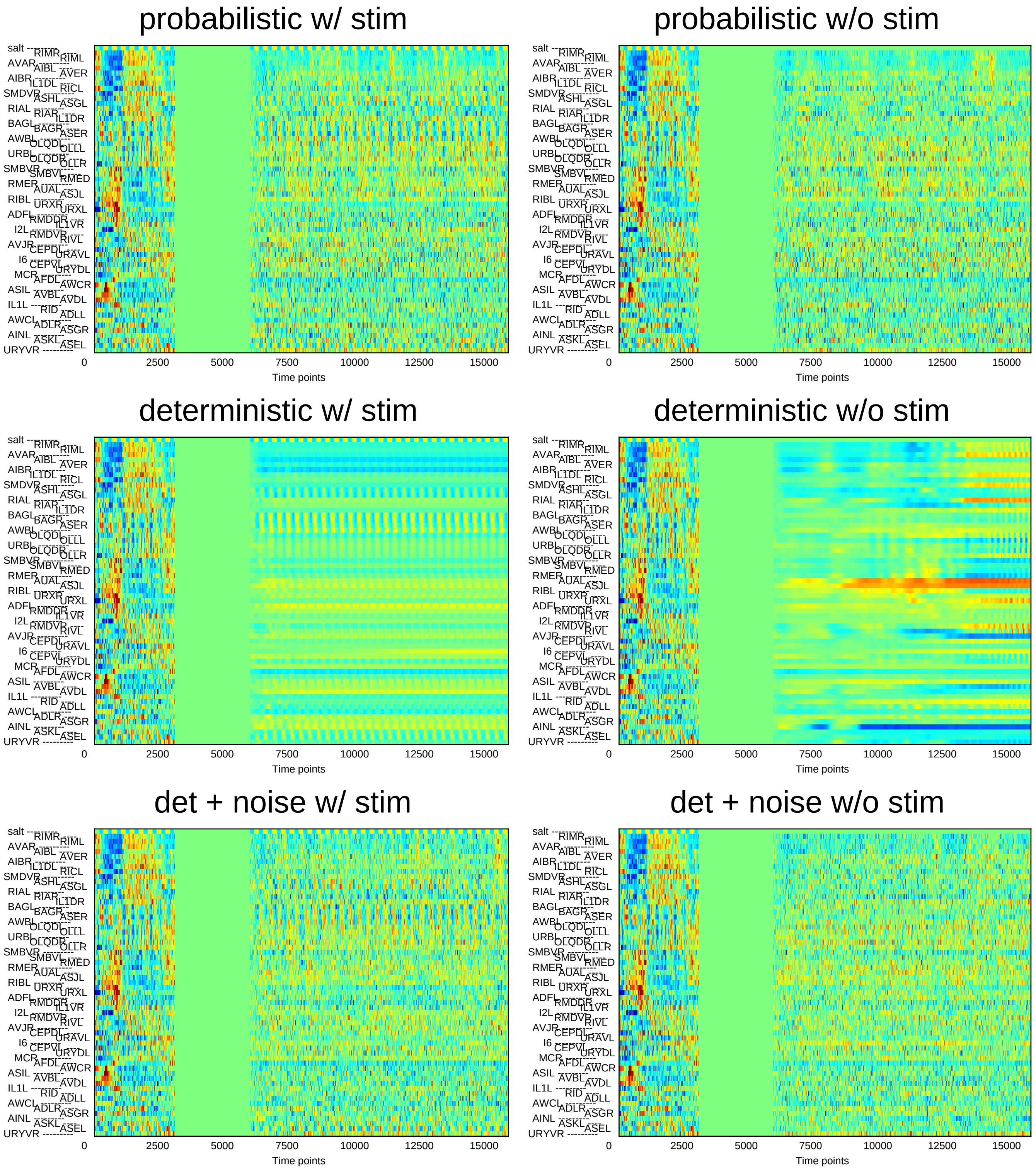
probabilistic w/ stim
probabilistic w/o stim
deterministic w/ stim
deterministic w/o stim
det + noise w/ stim
det + noise w/o stim
salt
RIMR
RIML
AVAR
AIBL
AVER
AIBR
IL1DL
RICL
SMDVR
ASHL
ASGL
RIAL
RIAR
IL1DR
BAGL
BAGR
ASER
AWBL
OLQDL
OLLL
URBL
OLQDR
OLLR
SMBVR
SMBVL
RMED
RMER
AUAL
ASJL
RIBL
URXR
URXL
ADFL
RMDDR
IL1VR
I2L
RMDVR
RIVL
AVJR
CEPDL
URAVL
I6
CEPVL
URYDL
MCR
AFDL
AWCR
ASIL
AVBL
AVDL
IL1L
RID
ADLL
AWCL
ADLR
ASGR
AINL
ASKL
ASEL
URYVR
0
2500
5000
7500
10000
12500
15000
Time points

### Sample3

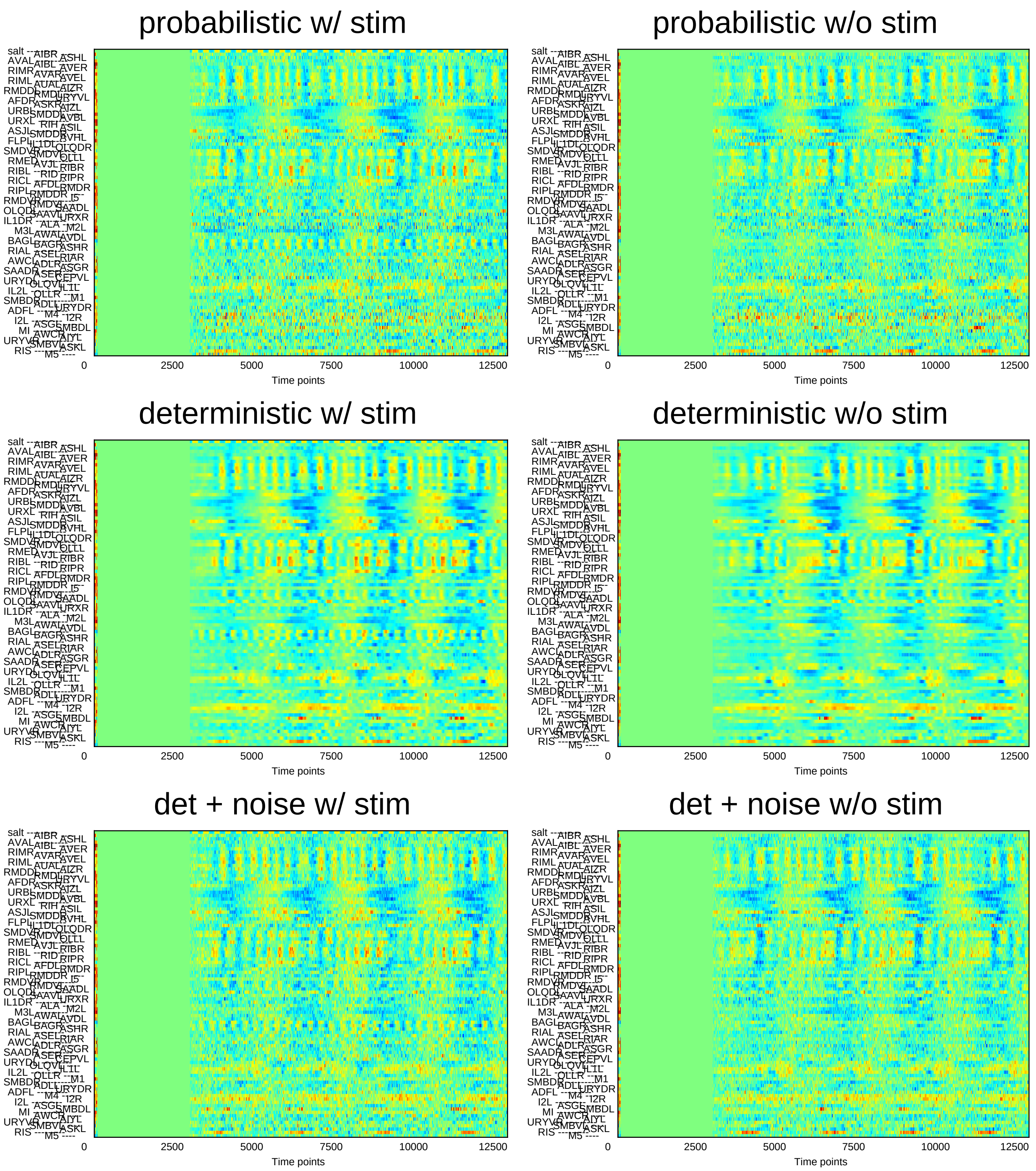
probabilistic w/ stim
probabilistic w/o stim
deterministic w/ stim
deterministic w/o stim
det + noise w/ stim
det + noise w/o stim
0
2500
5000
7500
10000
12500
Time points

### Sample4

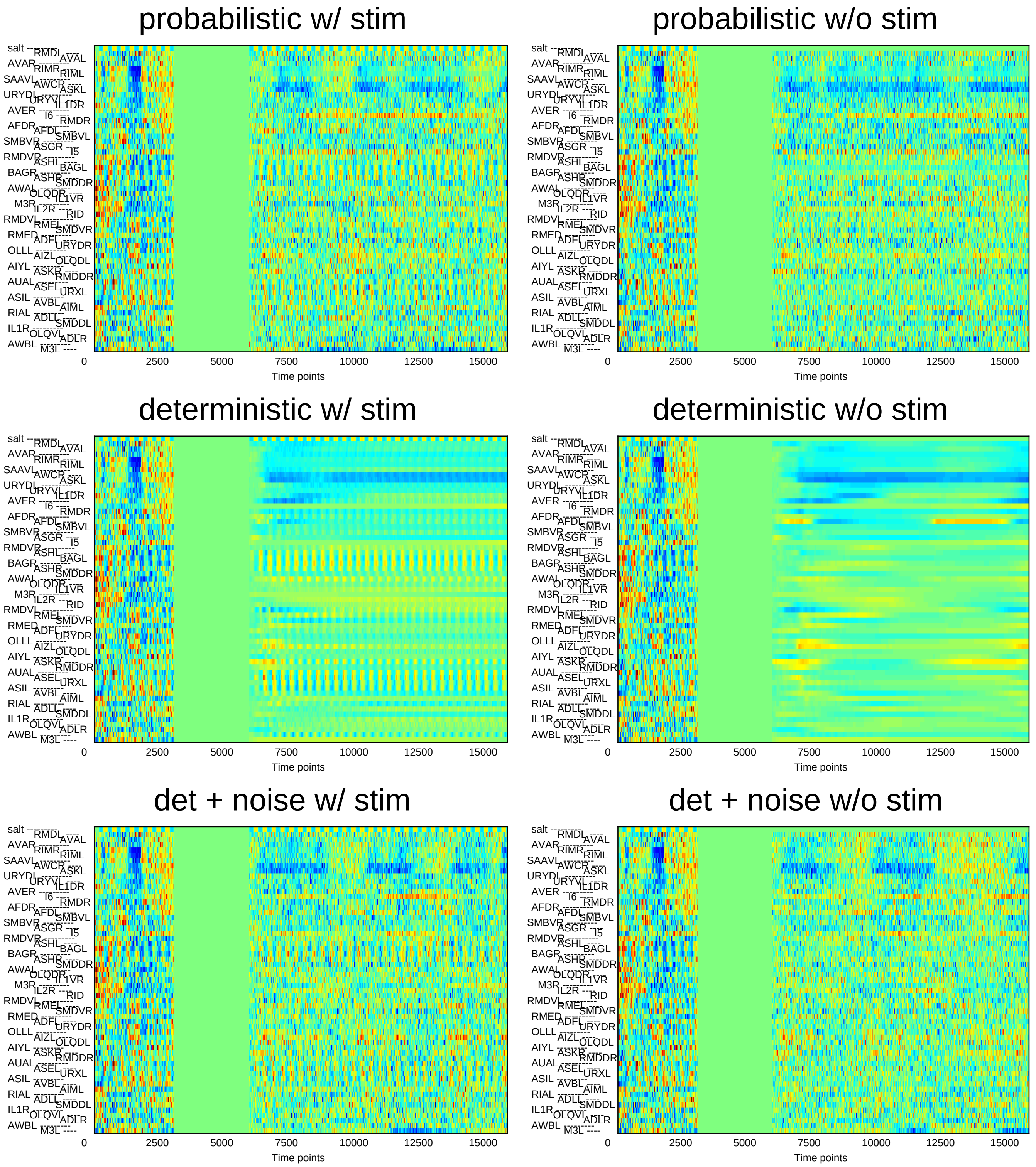
probabilistic w/ stim
probabilistic w/o stim
deterministic w/ stim
deterministic w/o stim
det + noise w/ stim
det + noise w/o stim
salt
AVAR
SAAVL
URYDL
AVER
AFDR
SMBVR
RMDVR
BAGR
AWAL
M3R
RMDVL
RMED
OLLL
AIYL
AUAL
ASIL
RIAL
IL1R
AWBL
RMDL
RIMR
AWCR
URYVL
I6
AFDL
ASGR
ASHL
ASHR
OLQDR
IL2R
RMEL
ADFL
AIZL
ASKR
ASEL
AVBL
ADLL
OLQVL
M3L
AVAL
RIML
ASKL
IL1DR
RMDR
SMBVL
I5
BAGL
SMDDR
IL1VR
RID
SMDVR
URYDR
OLQDL
RMDDR
URXL
AIML
SMDDL
ADLR
0
2500
5000
7500
10000
12500
15000
Time points

### Sample5

#### probabilistic w/ stim

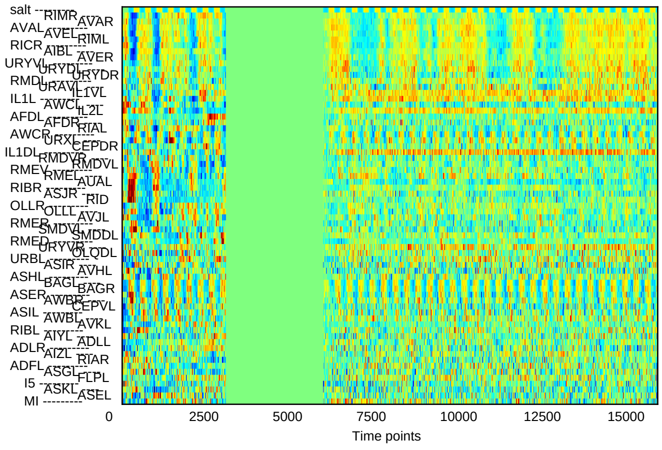


#### probabilistic w/o stim

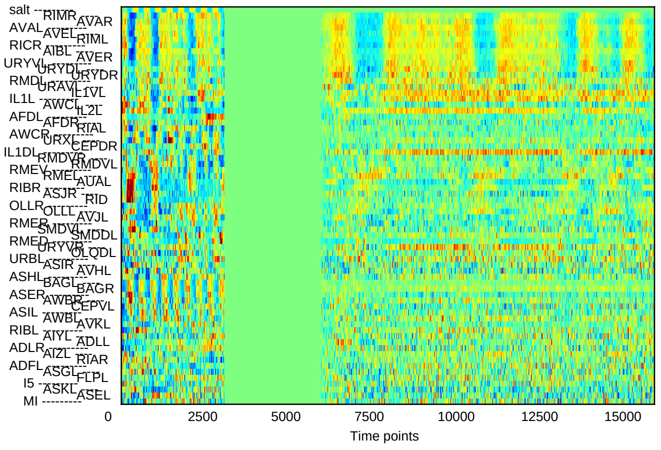


#### deterministic w/ stim

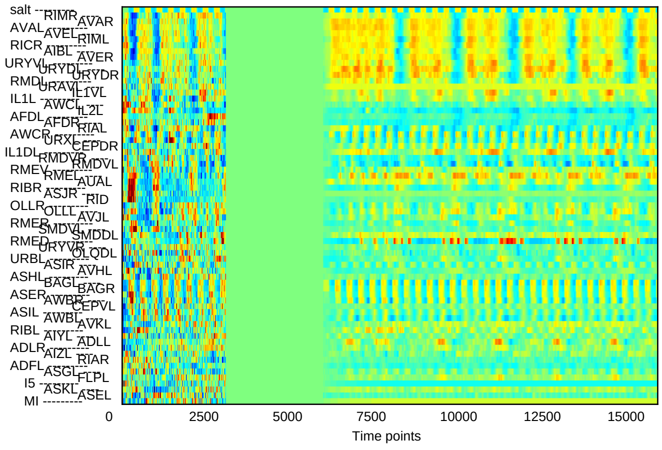


#### deterministic w/o stim

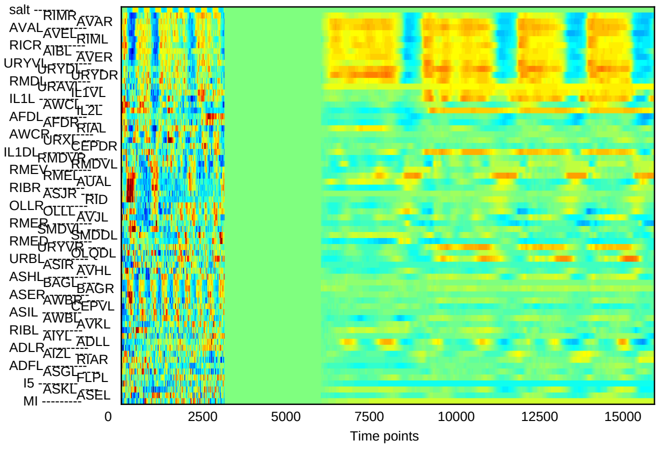


#### det + noise w/ stim

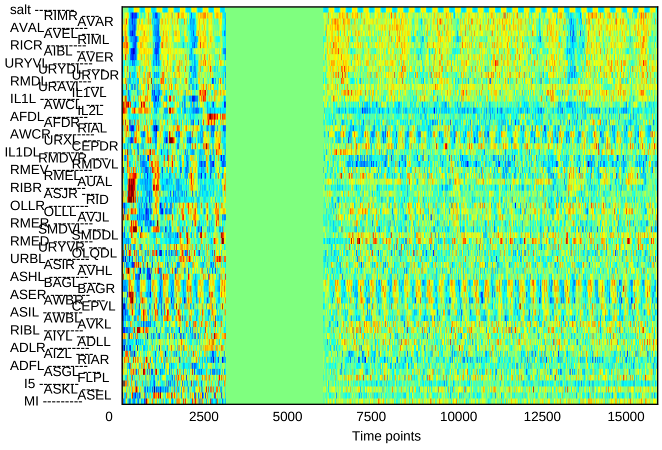


#### det + noise w/o stim

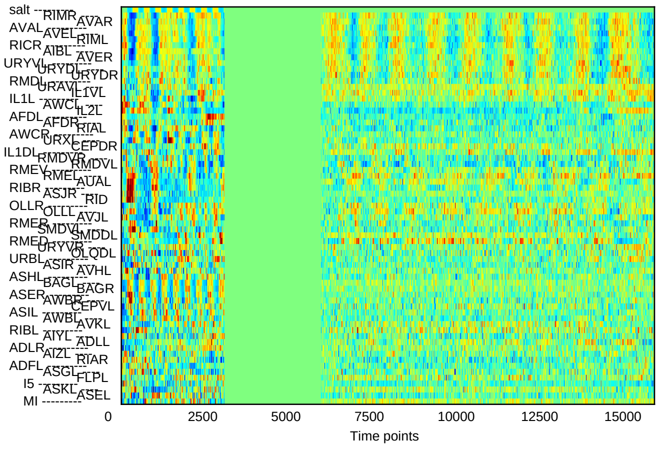

### Sample6

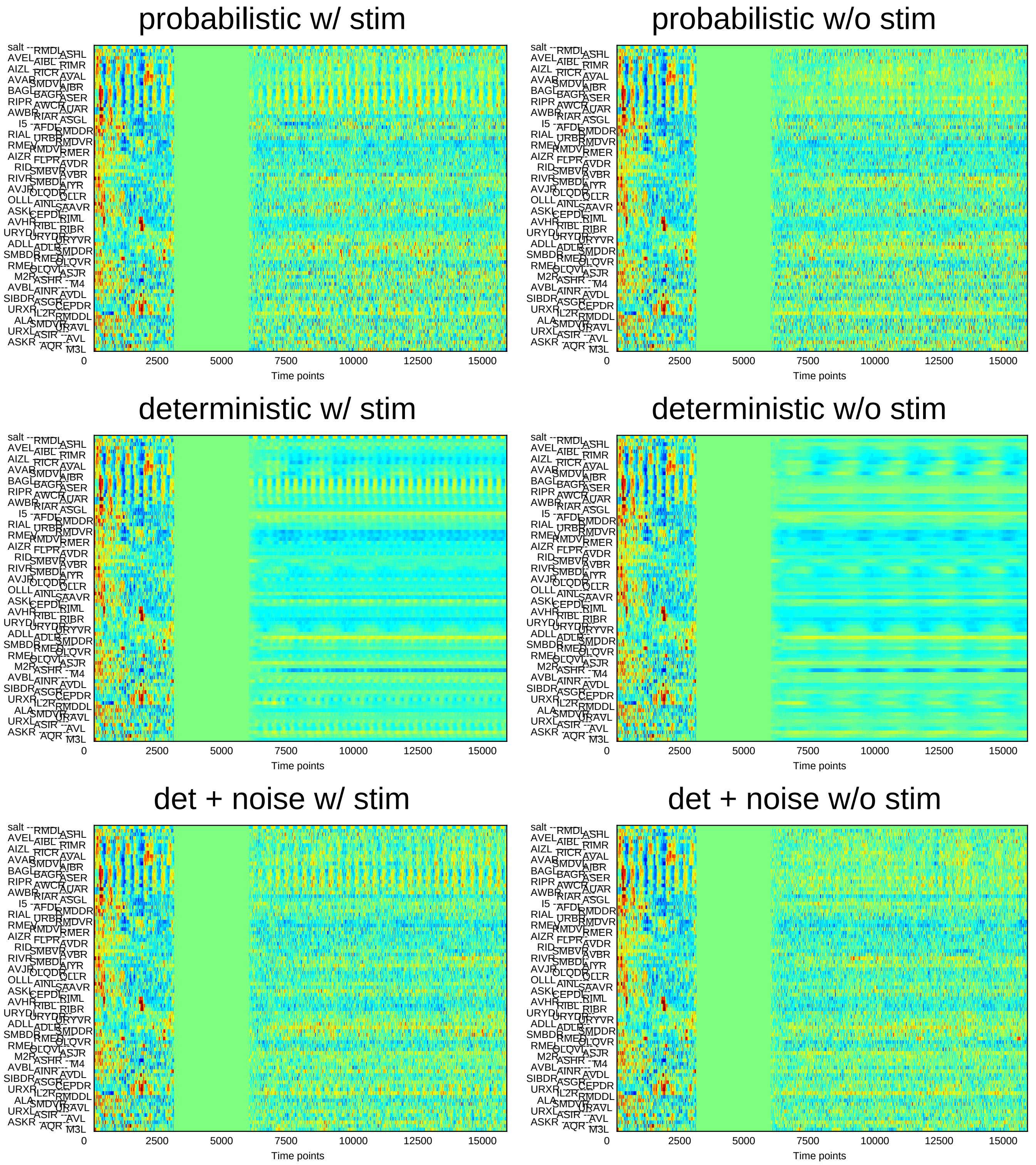
probabilistic w/ stim
probabilistic w/o stim
deterministic w/ stim
deterministic w/o stim
det + noise w/ stim
det + noise w/o stim
0
2500
5000
7500
10000
12500
15000
Time points

### Sample7

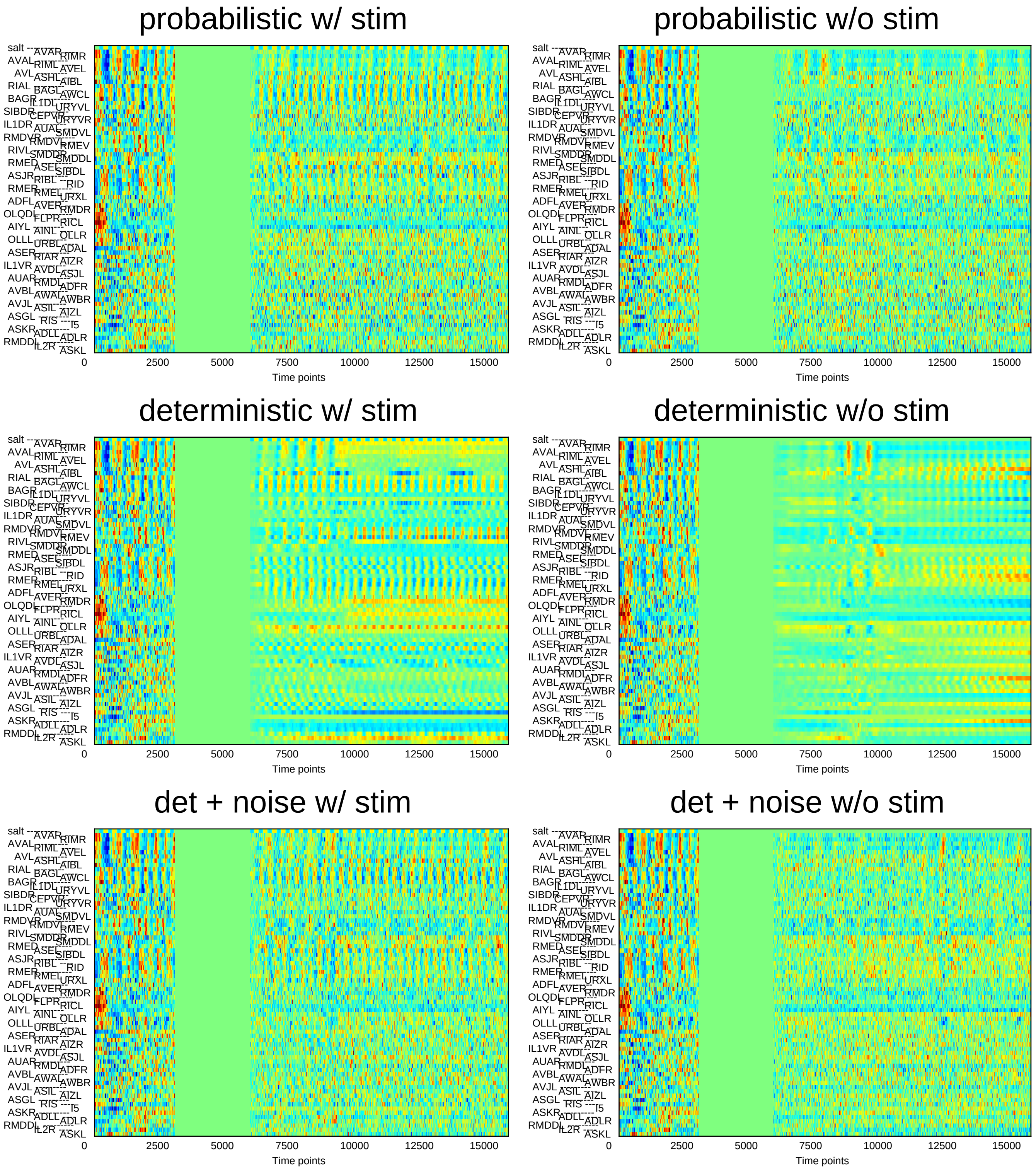
probabilistic w/ stim
probabilistic w/o stim
deterministic w/ stim
deterministic w/o stim
det + noise w/ stim
det + noise w/o stim
0 2500 5000 7500 10000 12500 15000
Time points

### Sample8

#### probabilistic w/ stim

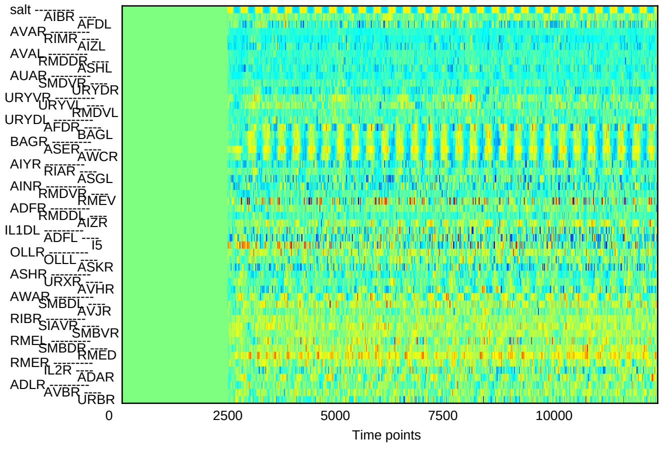


#### probabilistic w/o stim

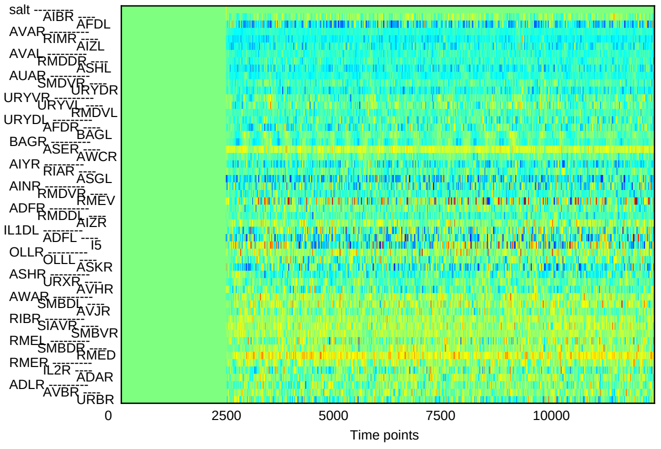


#### deterministic w/ stim

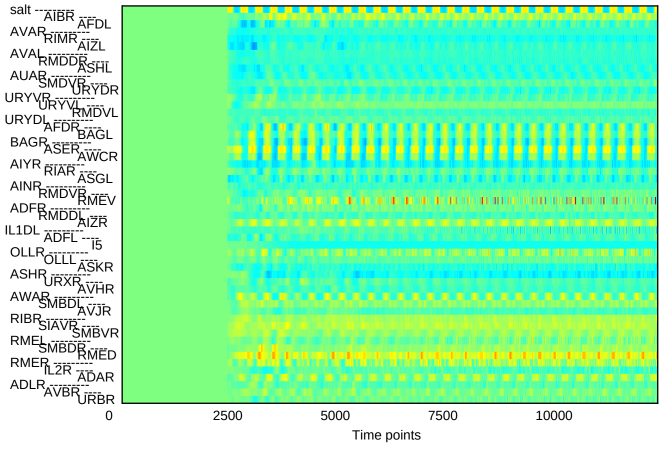


#### deterministic w/o stim

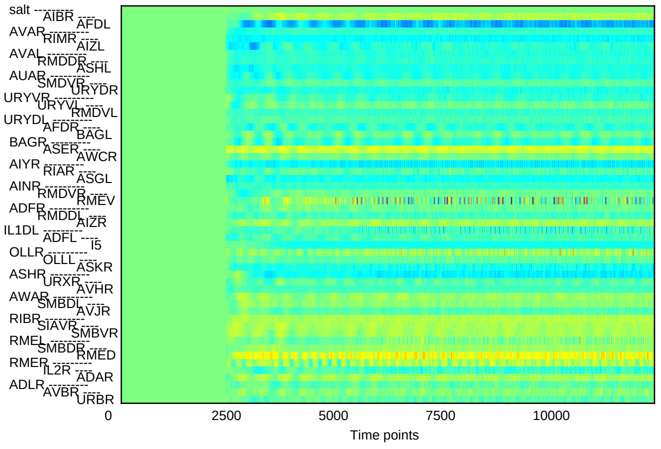


#### det + noise w/ stim

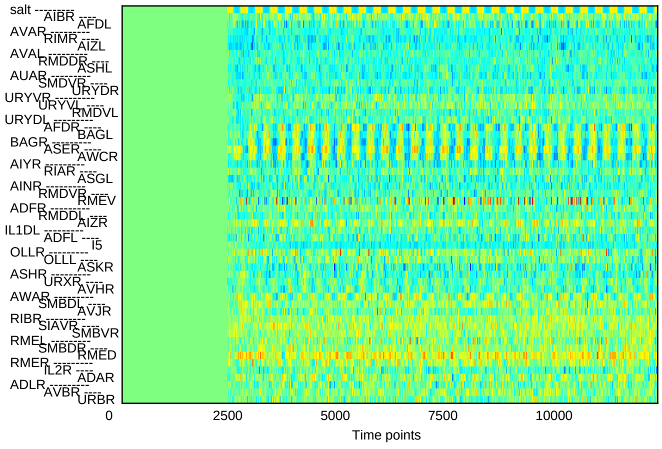


#### det + noise w/o stim

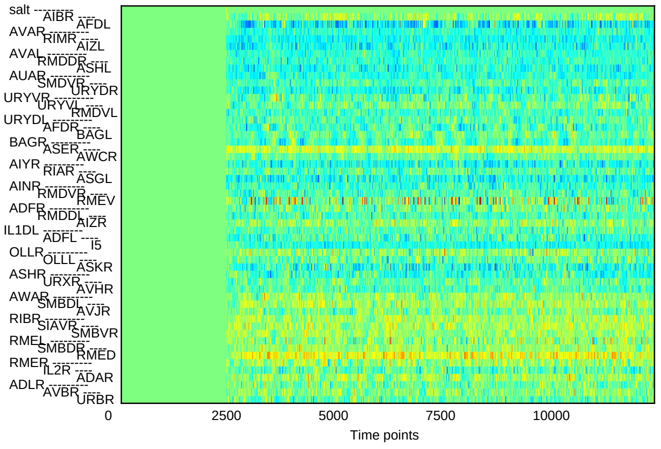

### Sample9

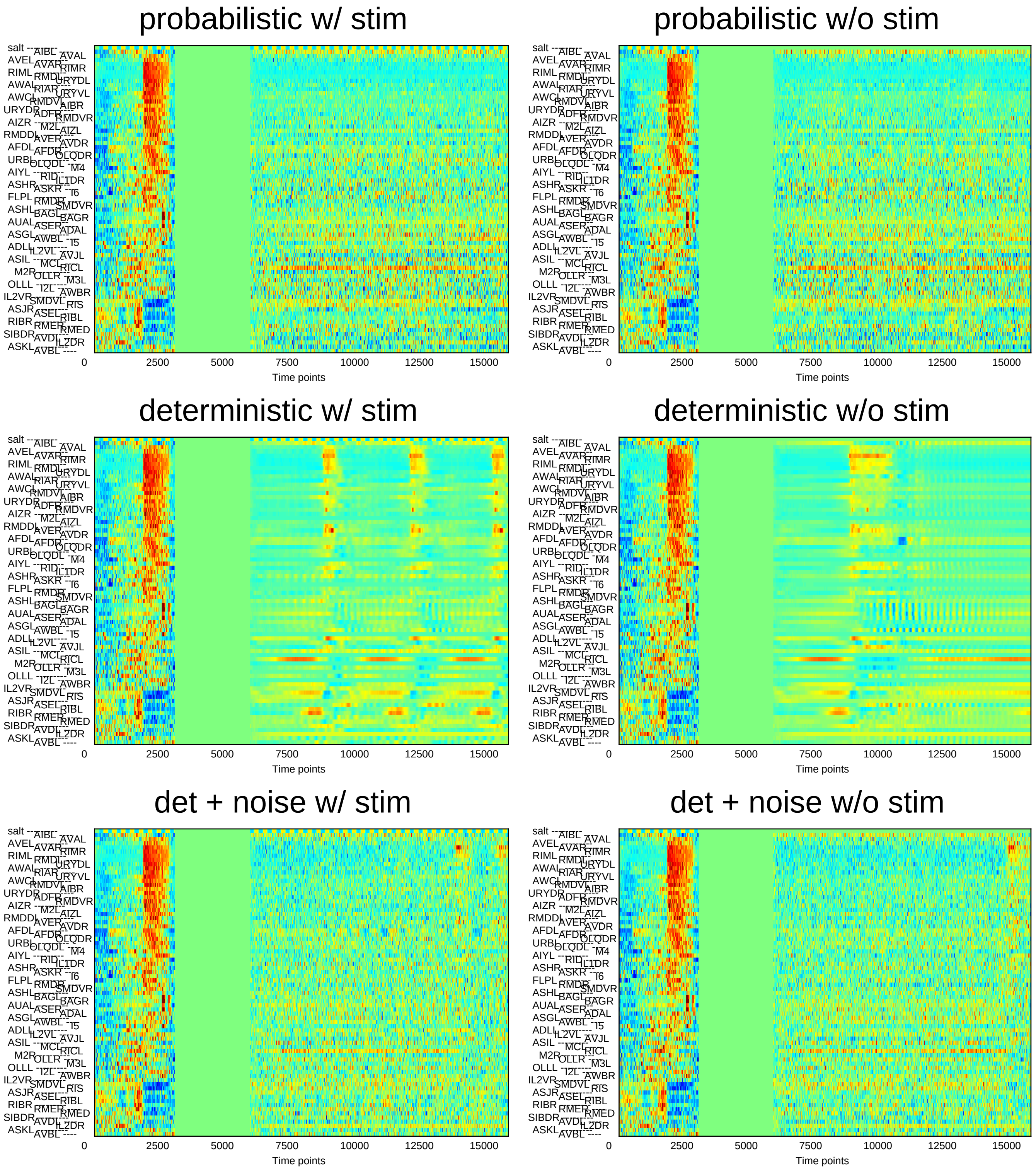
probabilistic w/ stim
probabilistic w/o stim
deterministic w/ stim
deterministic w/o stim
det + noise w/ stim
det + noise w/o stim
0
2500
5000
7500
10000
12500
15000
Time points

### Sample10

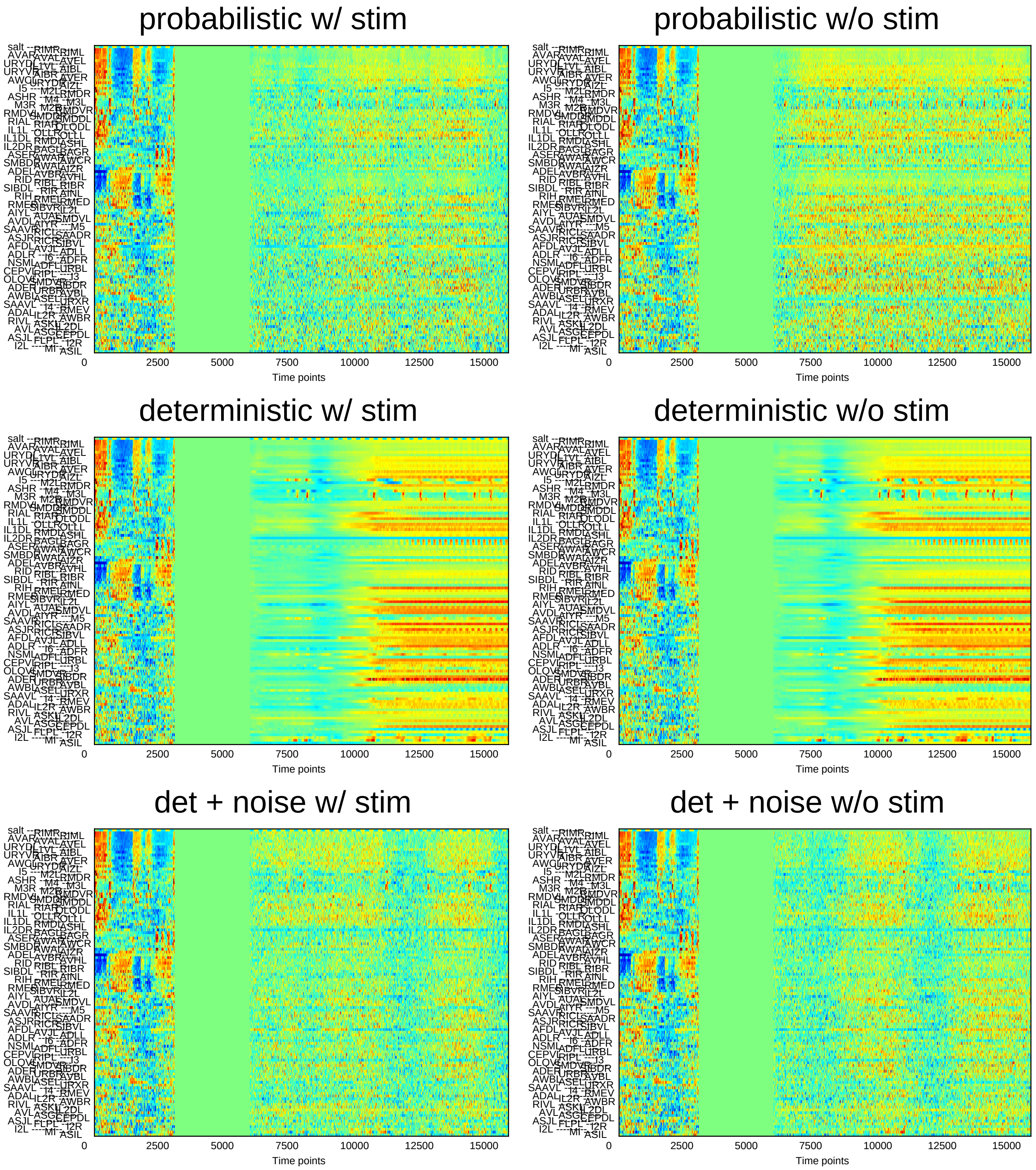
probabilistic w/ stim
probabilistic w/o stim
deterministic w/ stim
deterministic w/o stim
det + noise w/ stim
det + noise w/o stim
0 2500 5000 7500 10000 12500 15000
Time points

### Sample11

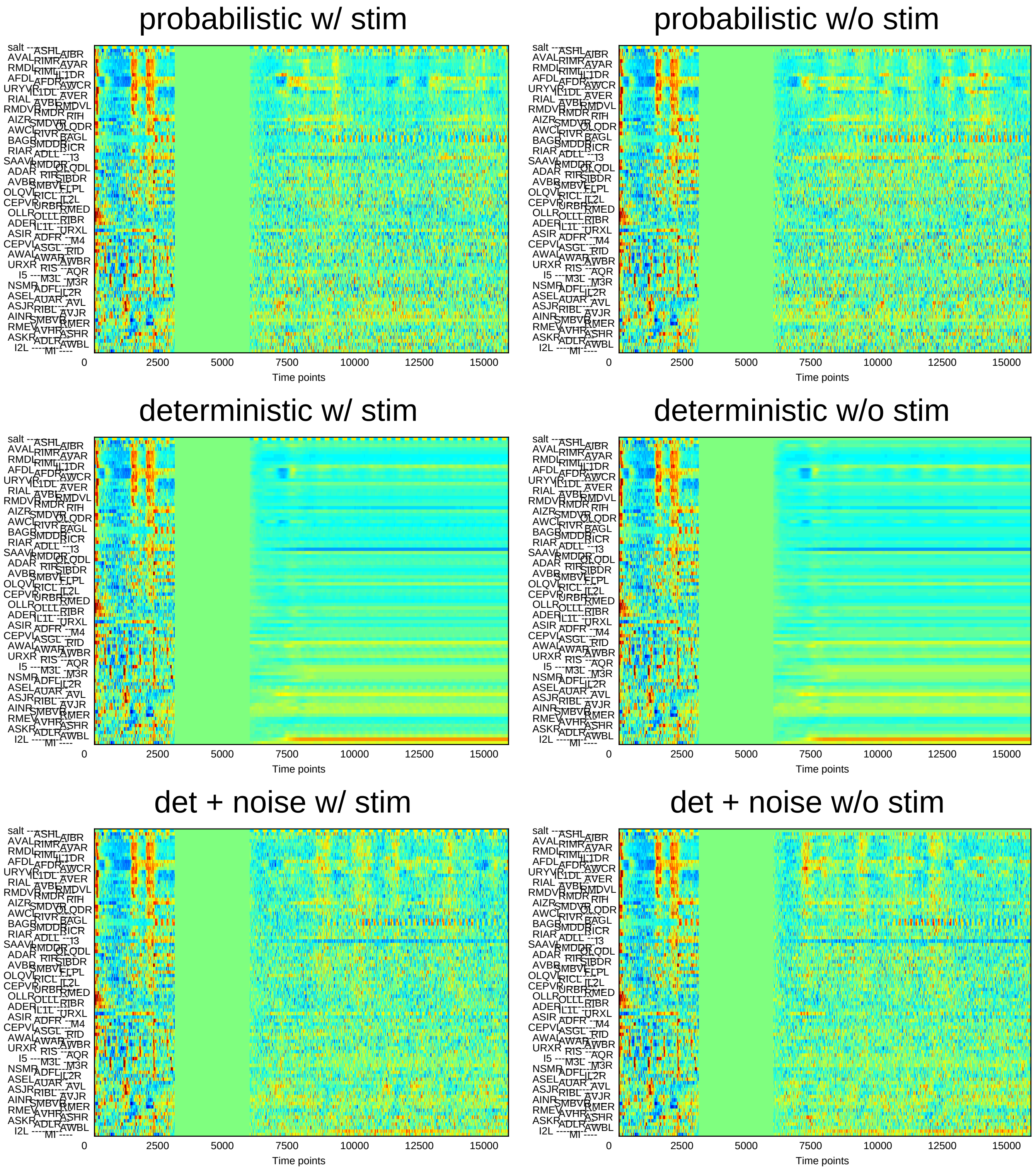
probabilistic w/ stim
probabilistic w/o stim
deterministic w/ stim
deterministic w/o stim
det + noise w/ stim
det + noise w/o stim
0
2500
5000
7500
10000
12500
15000
Time points

### Sample12

#### probabilistic w/ stim

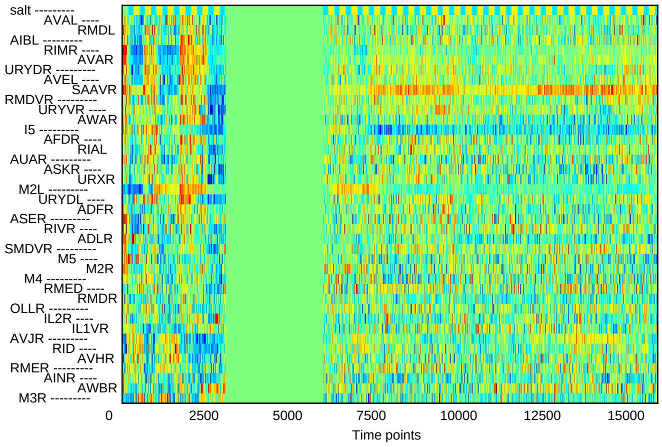


#### probabilistic w/o stim

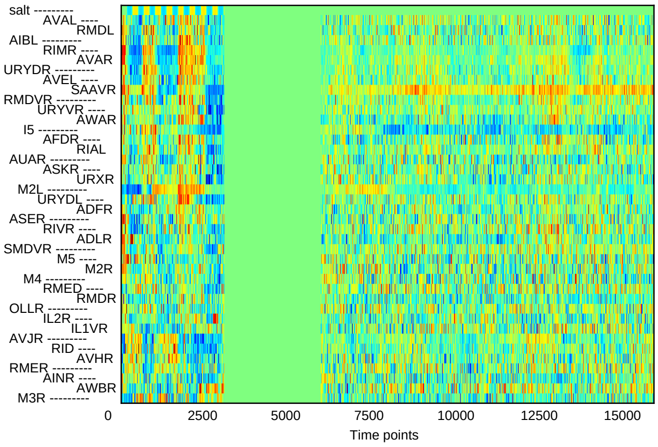


#### deterministic w/ stim

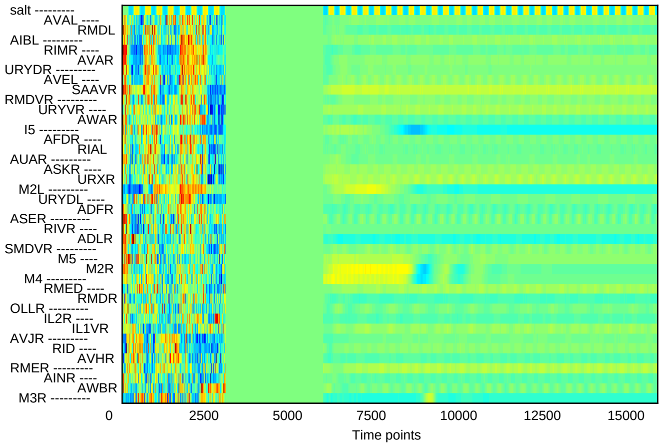


#### deterministic w/o stim

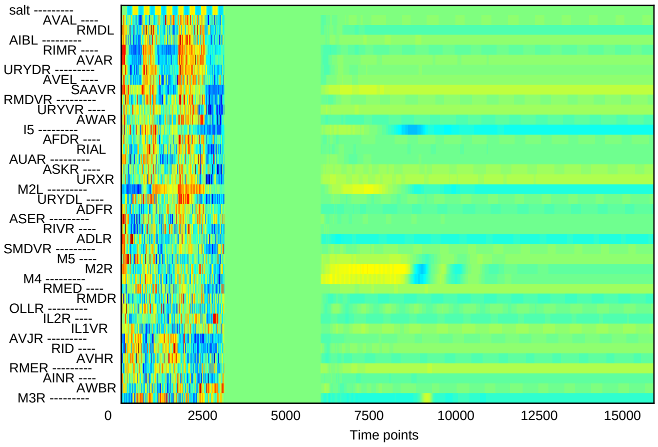


#### det + noise w/ stim

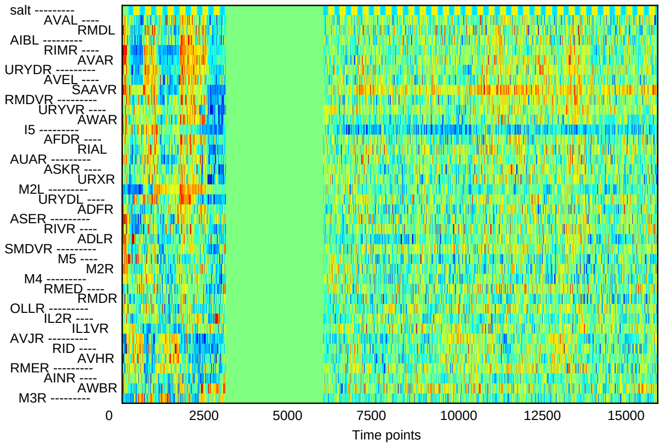


#### det + noise w/o stim

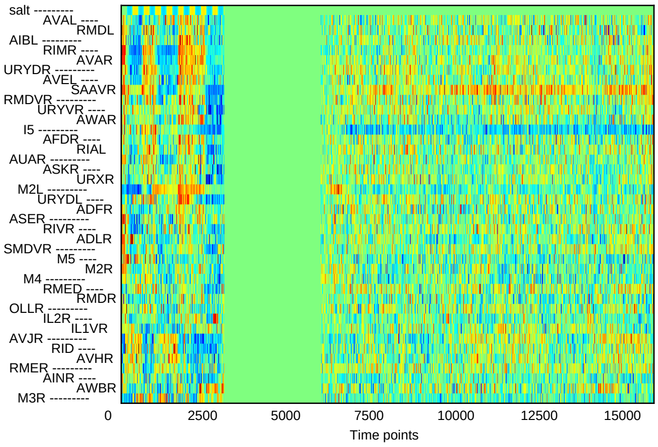

### Sample13

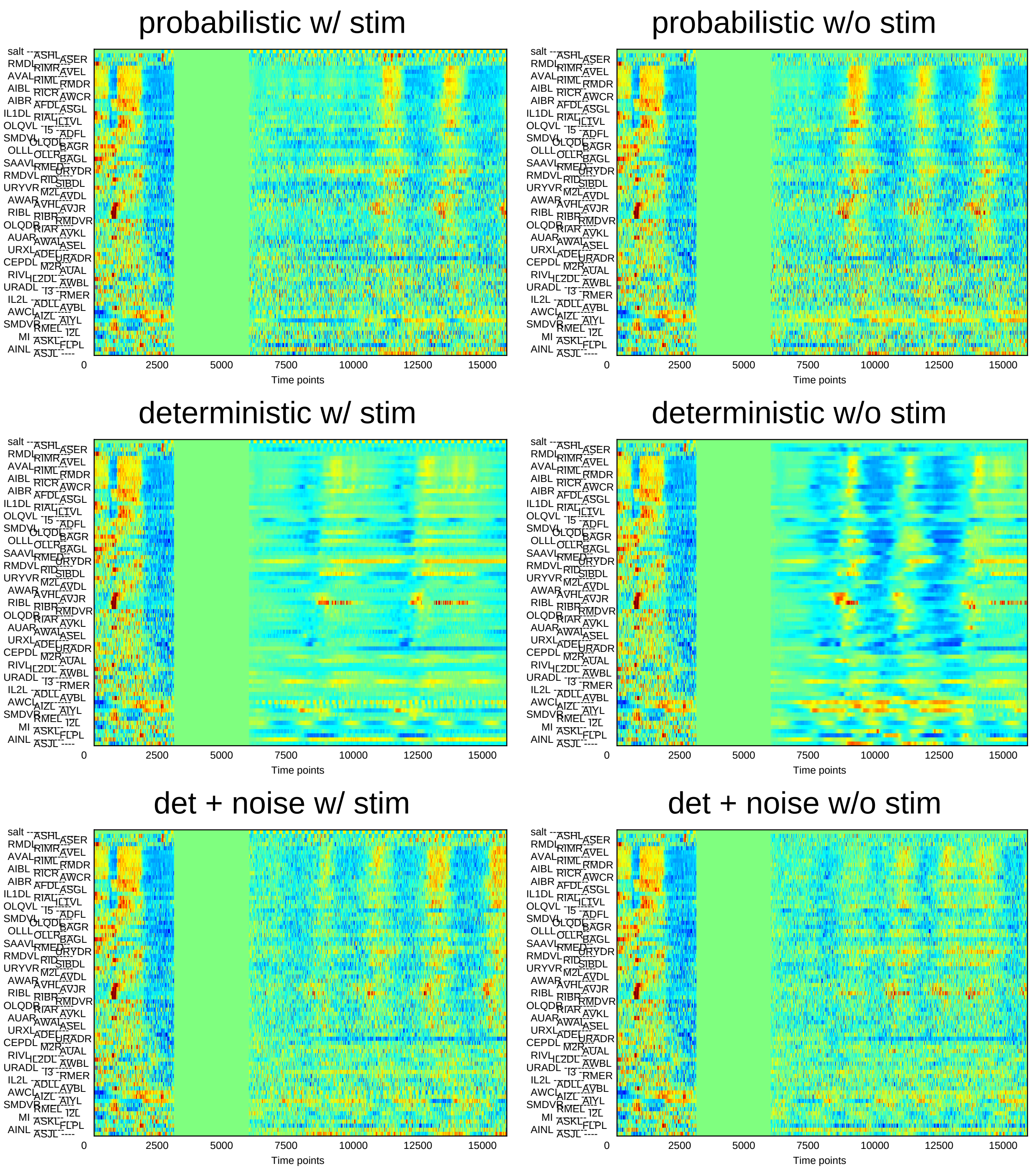
probabilistic w/ stim
probabilistic w/o stim
deterministic w/ stim
deterministic w/o stim
det + noise w/ stim
det + noise w/o stim
0
2500
5000
7500
10000
12500
15000
Time points

### Sample14

#### probabilistic w/ stim

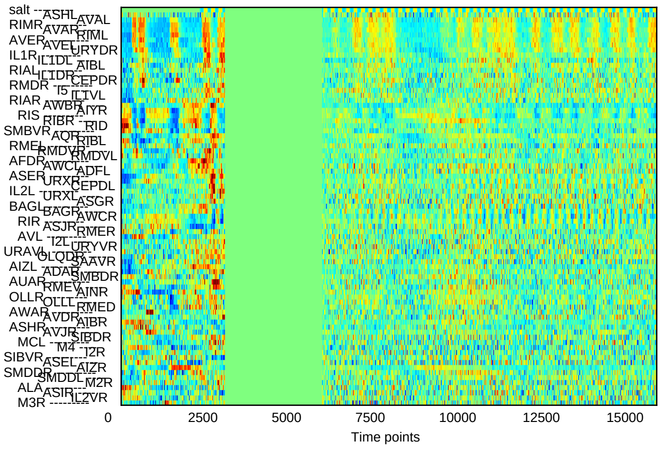


#### probabilistic w/o stim

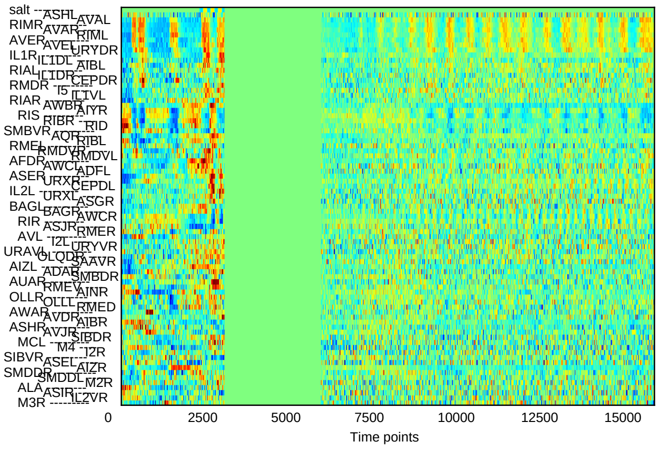


#### deterministic w/ stim

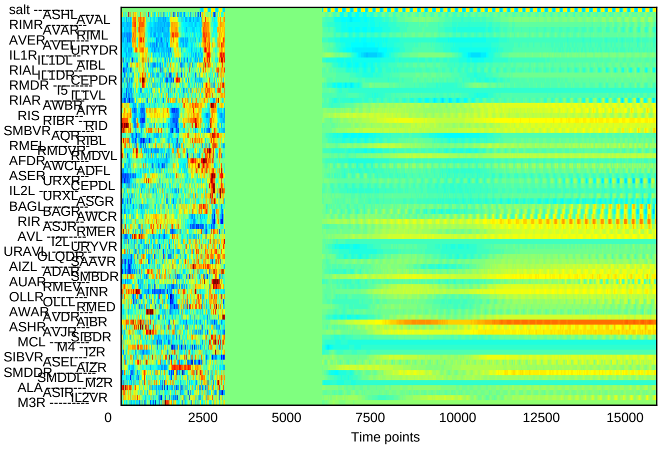


#### deterministic w/o stim

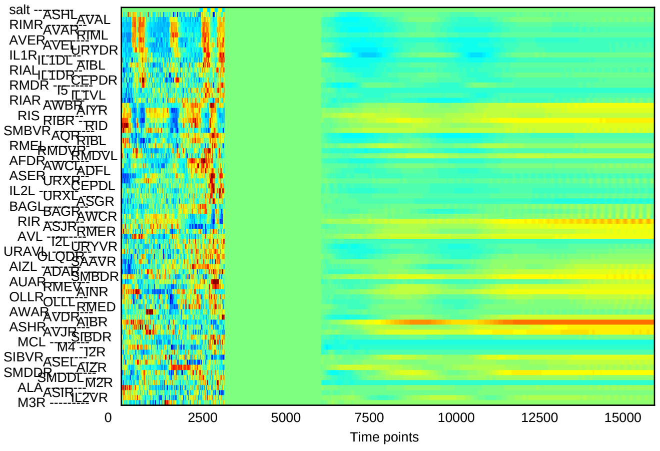


#### det + noise w/ stim

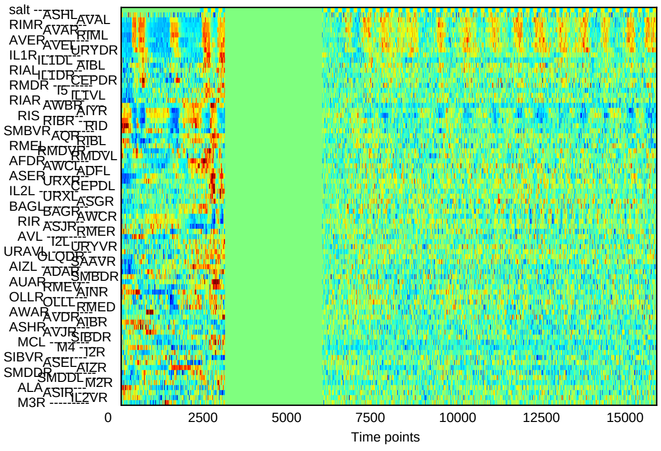


#### det + noise w/o stim

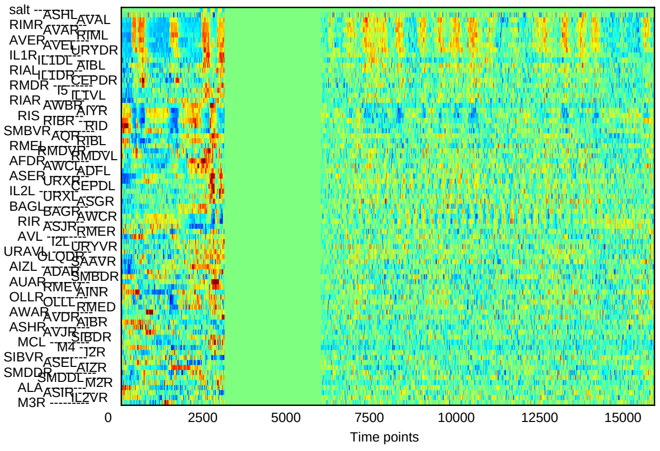

### Sample15

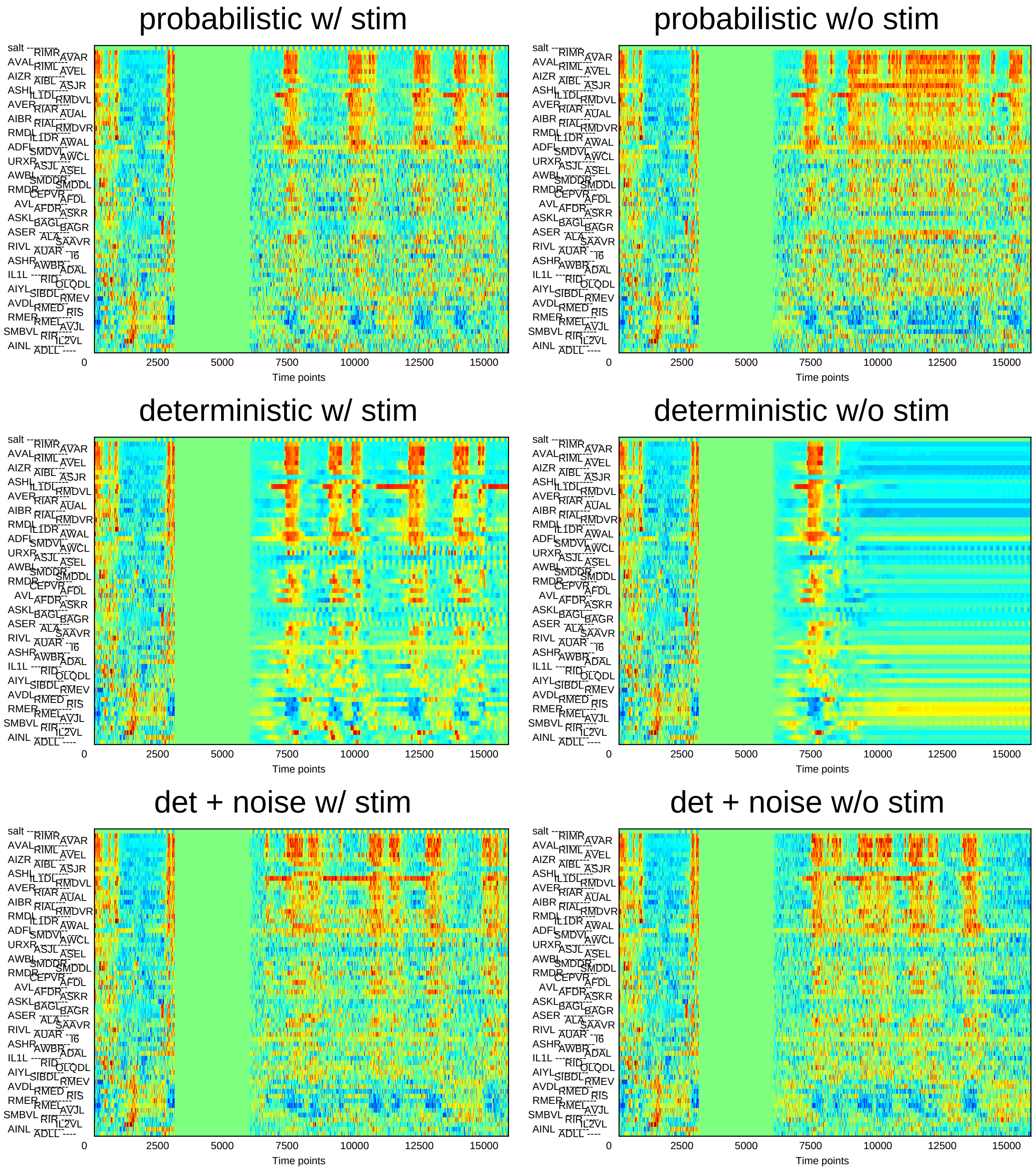
probabilistic w/ stim
probabilistic w/o stim
deterministic w/ stim
deterministic w/o stim
det + noise w/ stim
det + noise w/o stim
salt
AVAL
AIZR
ASHL
AVER
AIBR
RMDL
ADFL
URXR
AWBL
RMDR
AVL
ASKL
ASER
RIVL
ASHR
IL1L
AIYL
AVDL
RMER
SMBVL
AINL
RIMR
RIML
AIBL
IL1DL
RIAR
RIAL
IL1DR
SMDVL
ASJL
SMDDR
CEPVR
AFDR
BAGL
ALA
AUAR
AWBR
RID
SIBDL
RMED
RMEL
RIR
ADLL
AVAR
AVEL
ASJR
RMDVL
AUAL
RMDVR
AWAL
AWCL
ASEL
SMDDL
AFDL
ASKR
BAGR
SAAVR
I6
ADAL
OLQDL
RMEV
RIS
AVJL
IL2VL
0
2500
5000
7500
10000
12500
15000
Time points

### Sample16

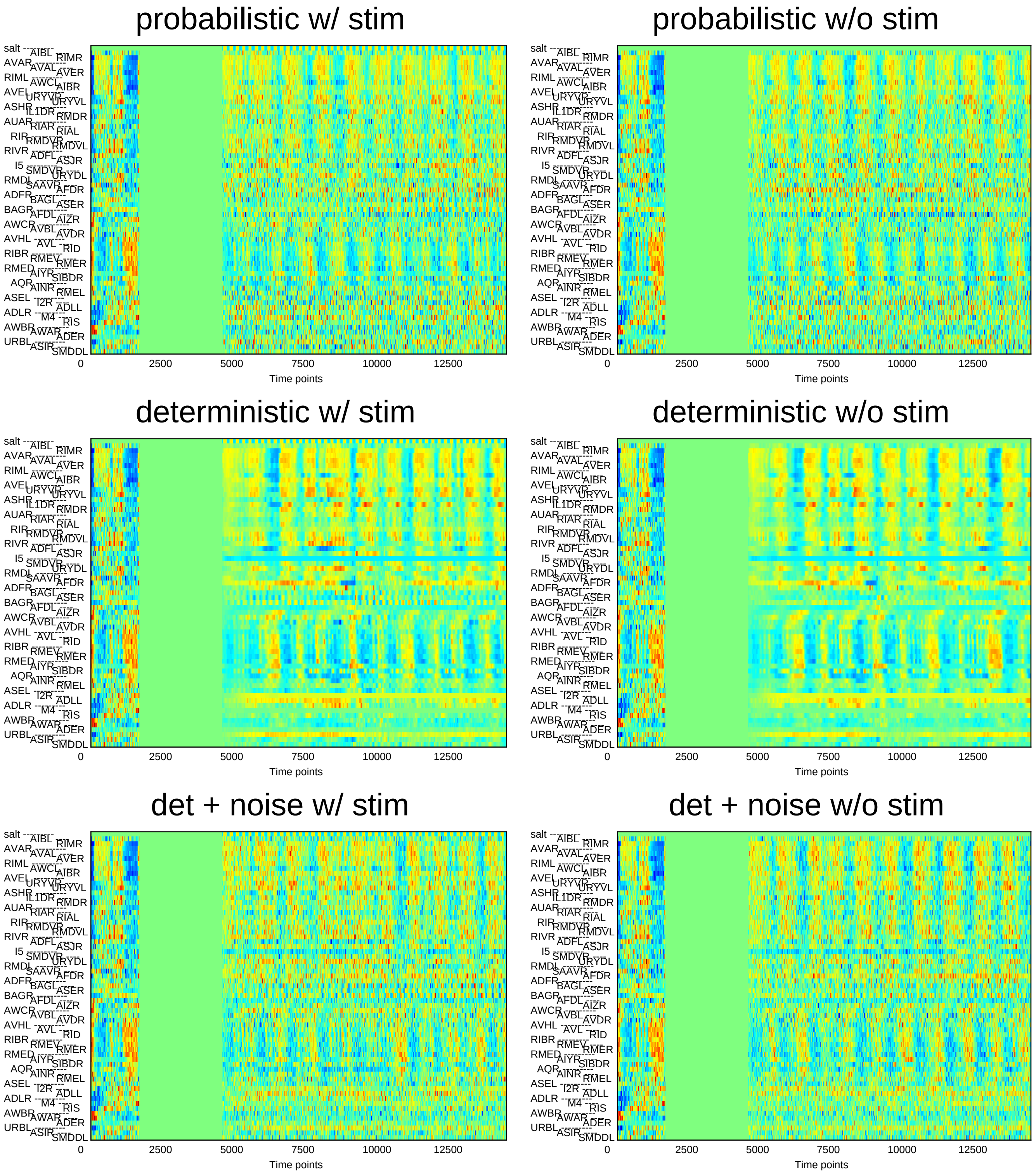
probabilistic w/ stim
probabilistic w/o stim
deterministic w/ stim
deterministic w/o stim
det + noise w/ stim
det + noise w/o stim
salt
AIBL
RIMR
AVAR
AVAL
AVER
RIML
AWCL
AIBR
AVEL
URYVR
URYVL
ASHR
IL1DR
RMDR
AUAR
RIAR
RIAL
RIR
RMDVR
RMDVL
RIVR
ADFL
ASJR
I5
SMDVR
URYDL
RMDL
SAAVR
AFDR
ADFR
BAGL
ASER
BAGR
AFDL
AIZR
AWCR
AVBL
AVDR
AVHL
AVL
RID
RIBR
RMEV
RMER
RMED
AIYR
SIBDR
AQR
AINR
RMEL
ASEL
I2R
ADLL
ADLR
M4
RIS
AWBR
AWAR
ADER
URBL
ASIR
SMDDL
0
2500
5000
7500
10000
12500
Time points

### Sample17

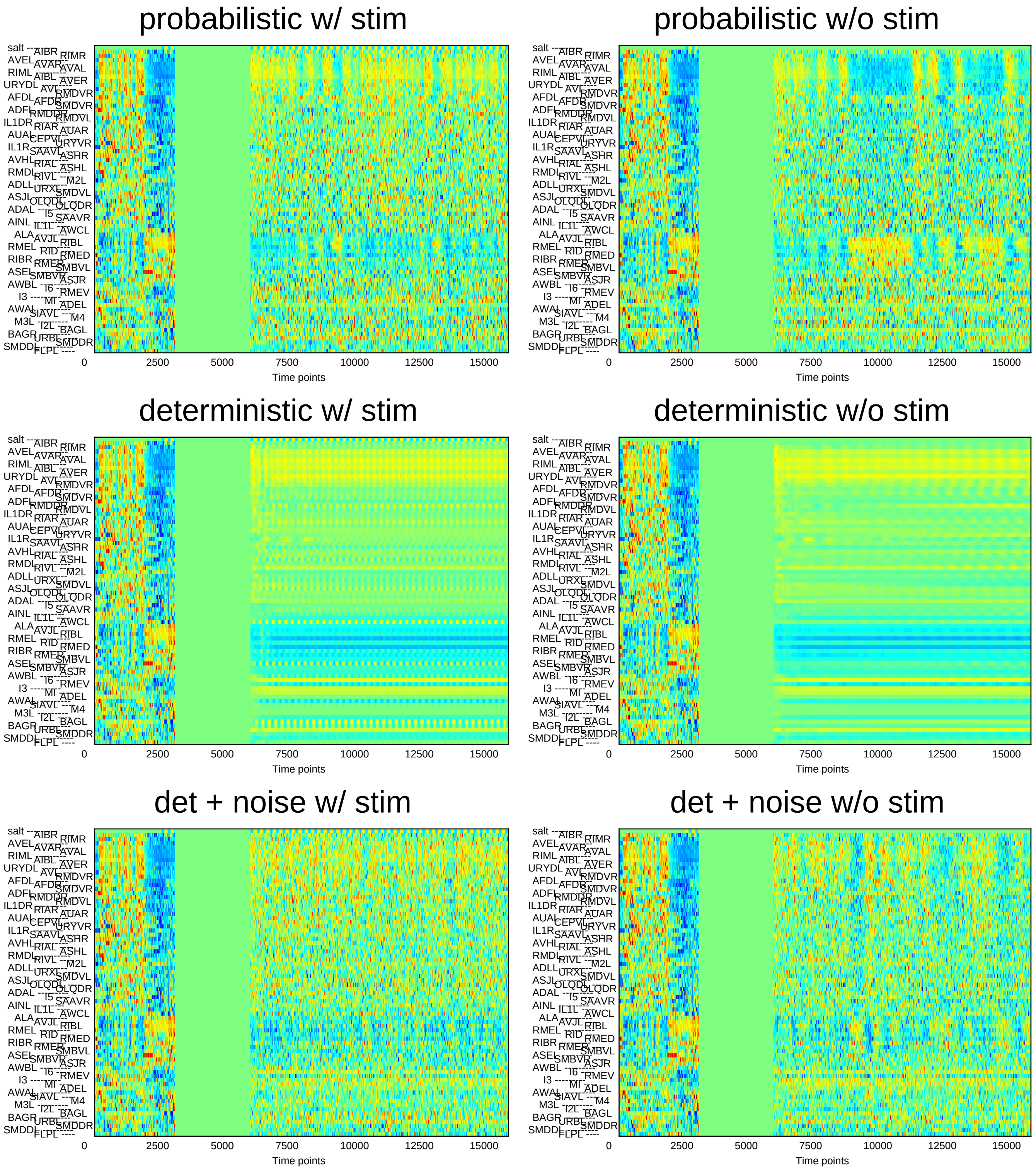
probabilistic w/ stim
probabilistic w/o stim
deterministic w/ stim
deterministic w/o stim
det + noise w/ stim
det + noise w/o stim
0
2500
5000
7500
10000
12500
15000
Time points

### Sample18

#### probabilistic w/ stim

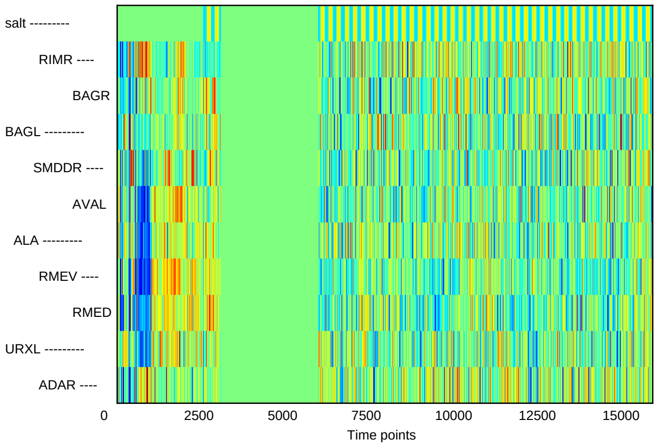


#### probabilistic w/o stim

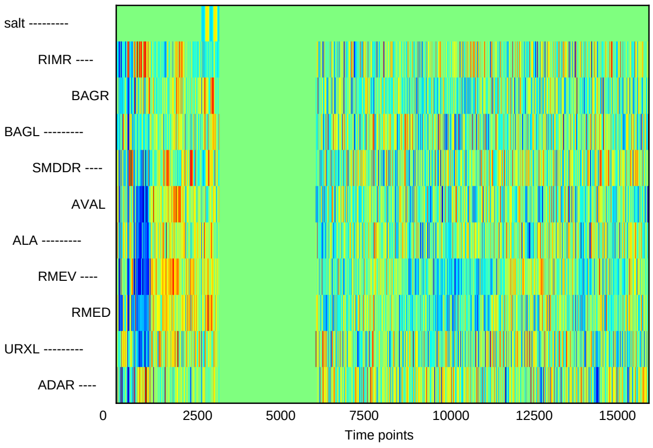


#### deterministic w/ stim

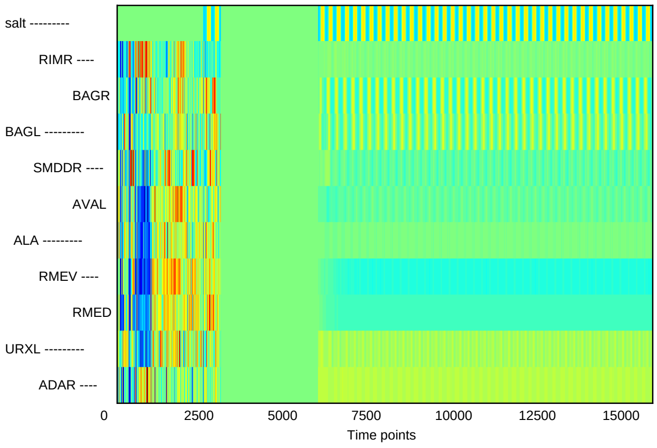


#### deterministic w/o stim

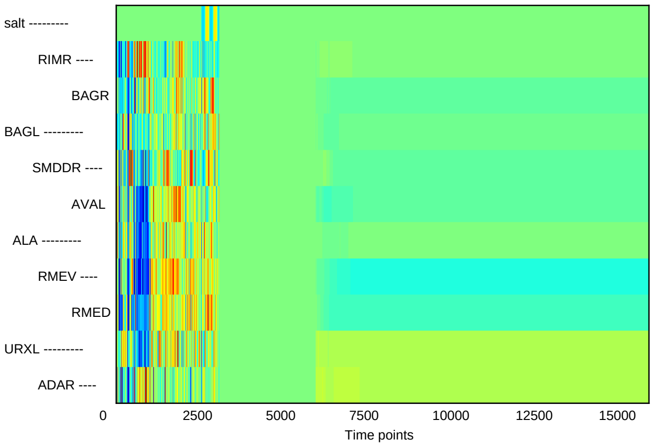


#### det + noise w/ stim

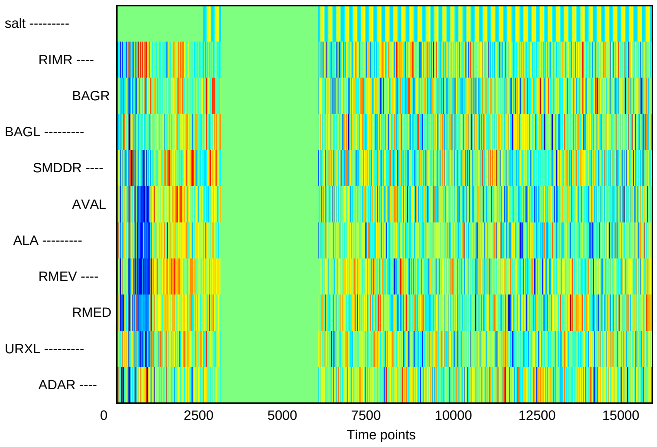


#### det + noise w/o stim

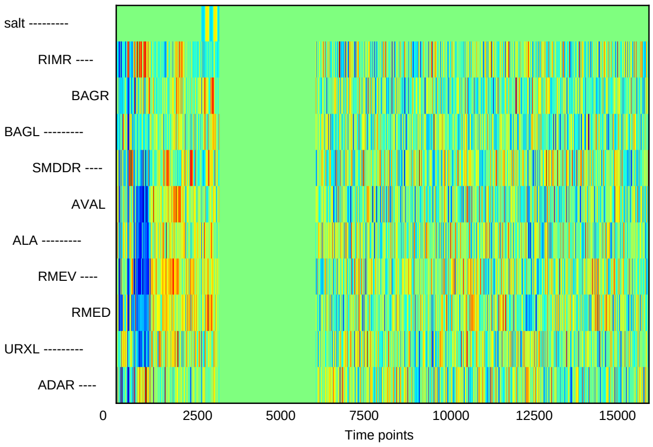

### Sample19

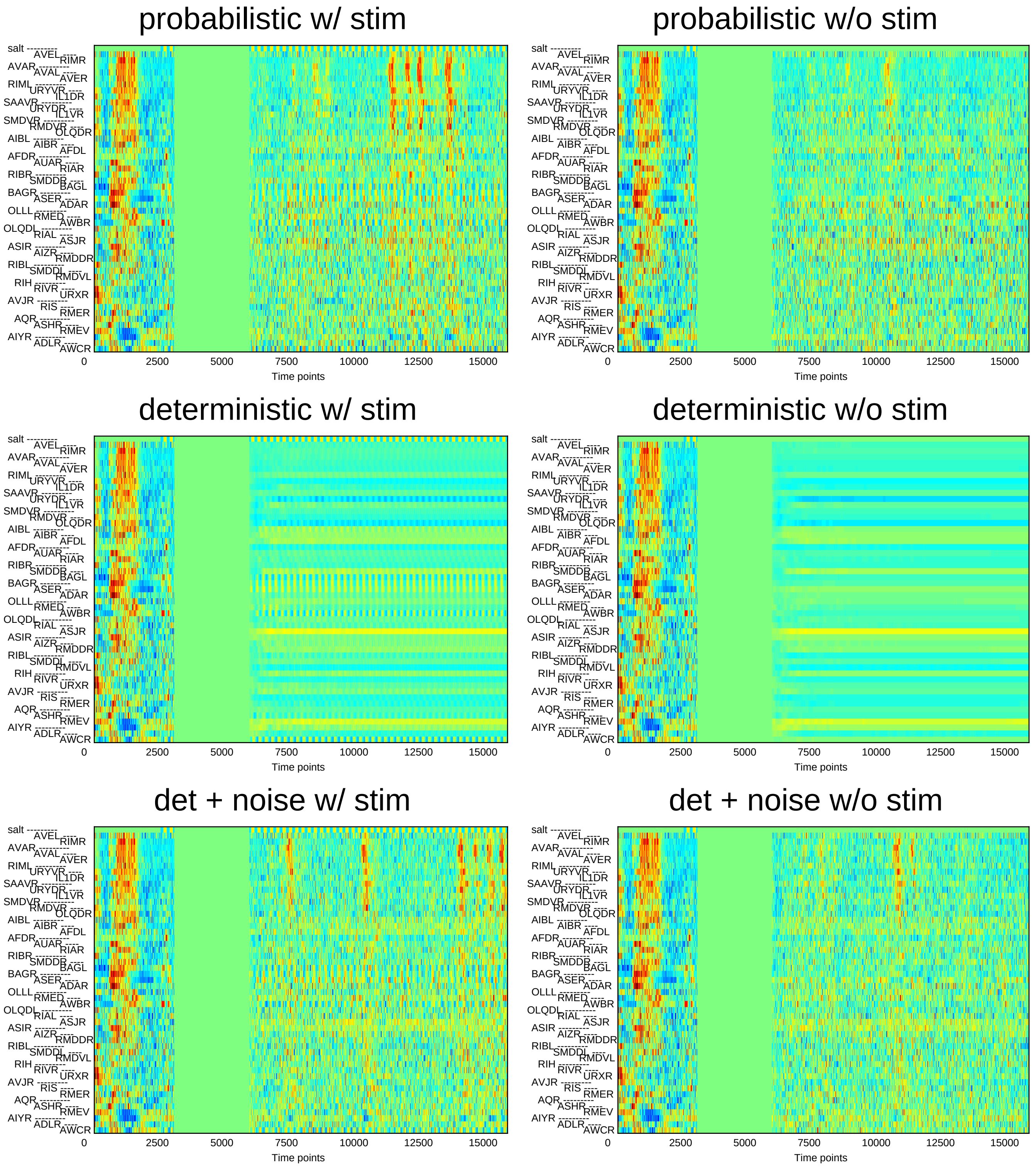
probabilistic w/ stim
probabilistic w/o stim
deterministic w/ stim
deterministic w/o stim
det + noise w/ stim
det + noise w/o stim
salt
AVEL
RIMR
AVAR
AVAL
AVER
RIML
URYVR
IL1DR
SAAVR
URYDR
IL1VR
SMDVR
RMDVR
OLQDR
AIBL
AIBR
AFDL
AFDR
AUAR
RIAR
RIBR
SMDDR
BAGL
BAGR
ASER
ADAR
OLLL
RMED
AWBR
OLQDL
RIAL
ASJR
ASIR
AIZR
RMDDR
RIBL
SMDDL
RMDVL
RIH
RIVR
URXR
AVJR
RIS
RMER
AQR
ASHR
RMEV
AIYR
ADLR
AWCR
0
2500
5000
7500
10000
12500
15000
Time points

### Sample20

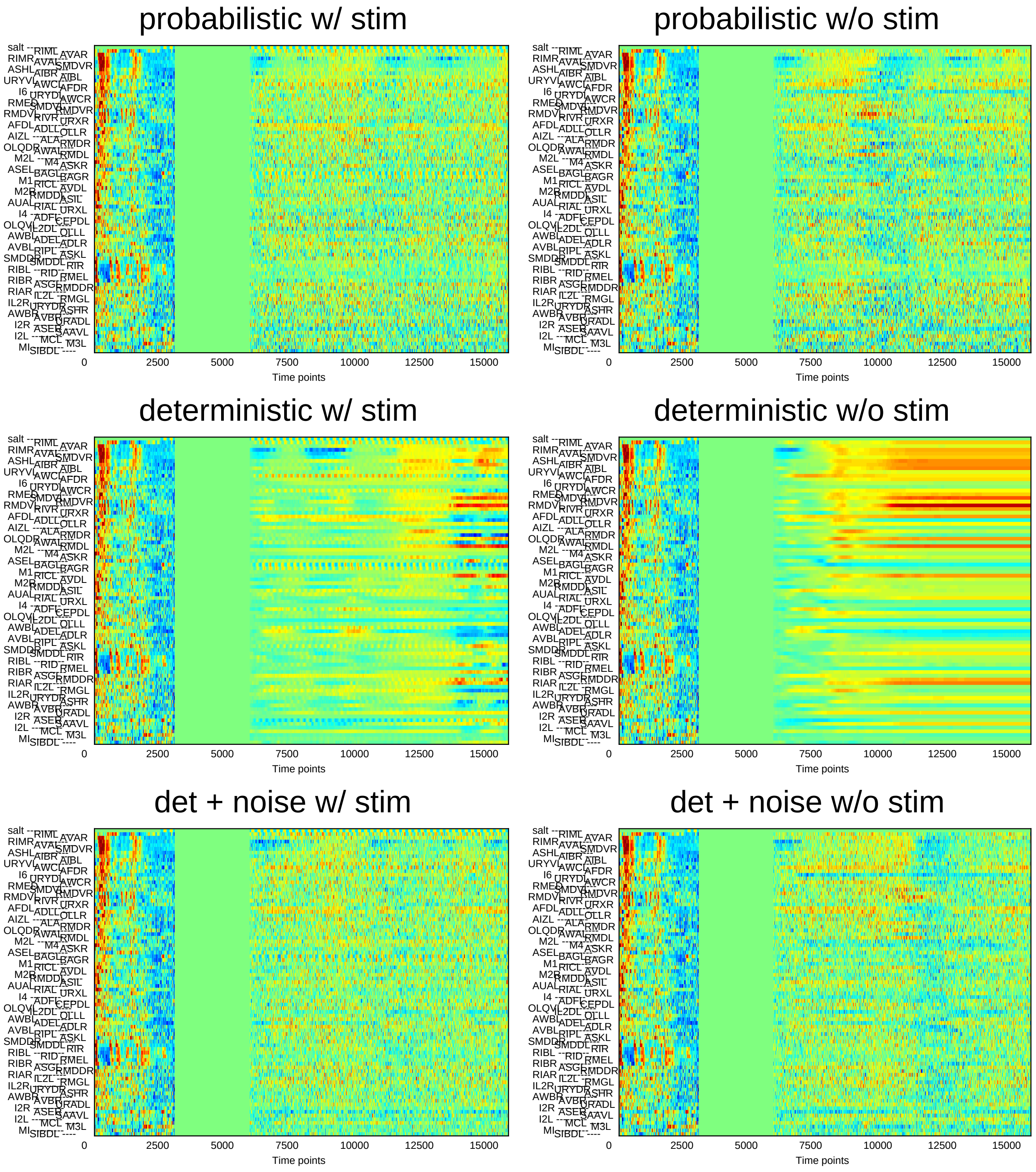

probabilistic w/ stim
probabilistic w/o stim
deterministic w/ stim
deterministic w/o stim
det + noise w/ stim
det + noise w/o stim
Time points
0
2500
5000
7500
10000
12500
15000

### Sample21

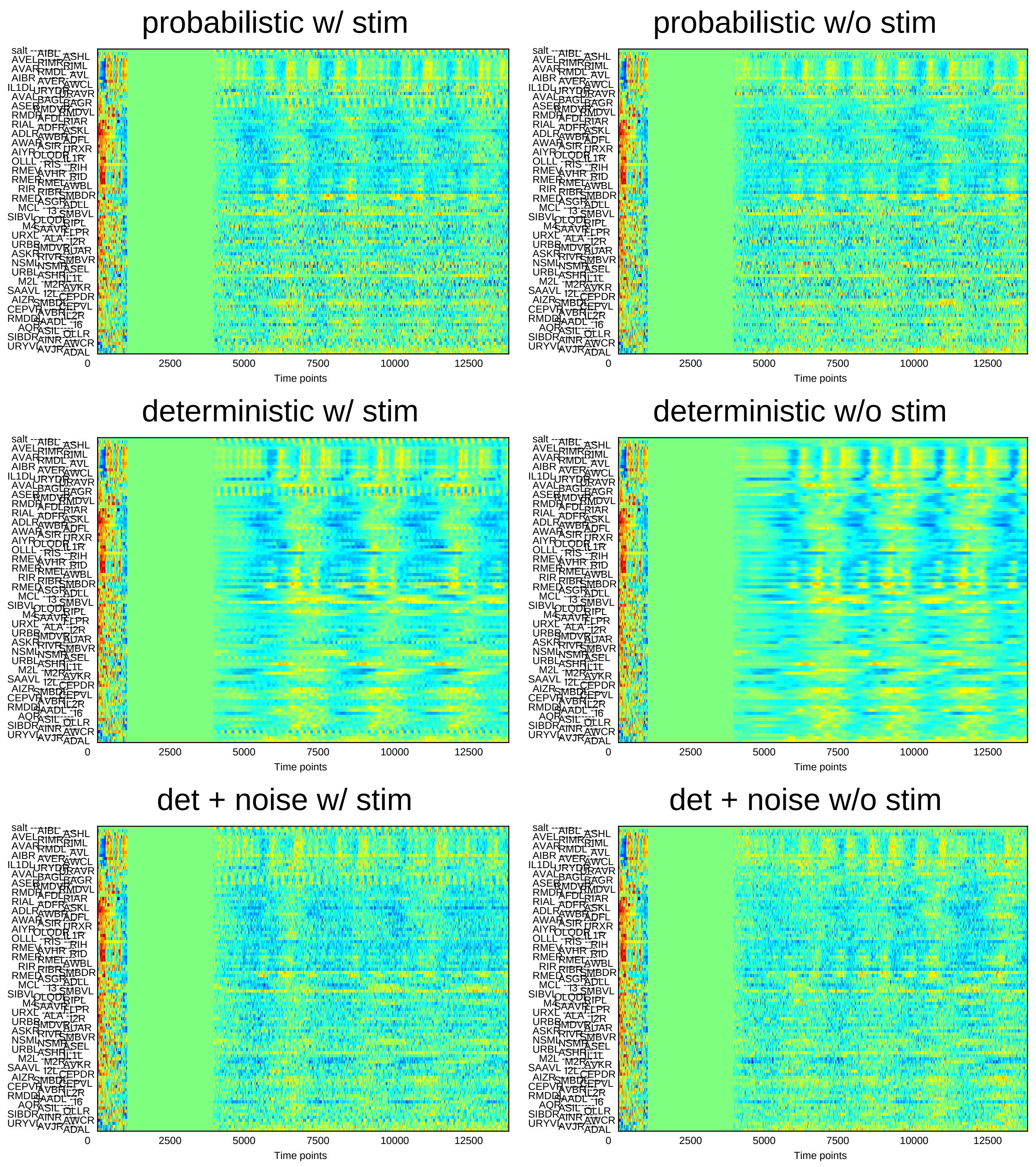
probabilistic w/ stim
probabilistic w/o stim
deterministic w/ stim
deterministic w/o stim
det + noise w/ stim
det + noise w/o stim
0
2500
5000
7500
10000
12500
Time points

### Sample22

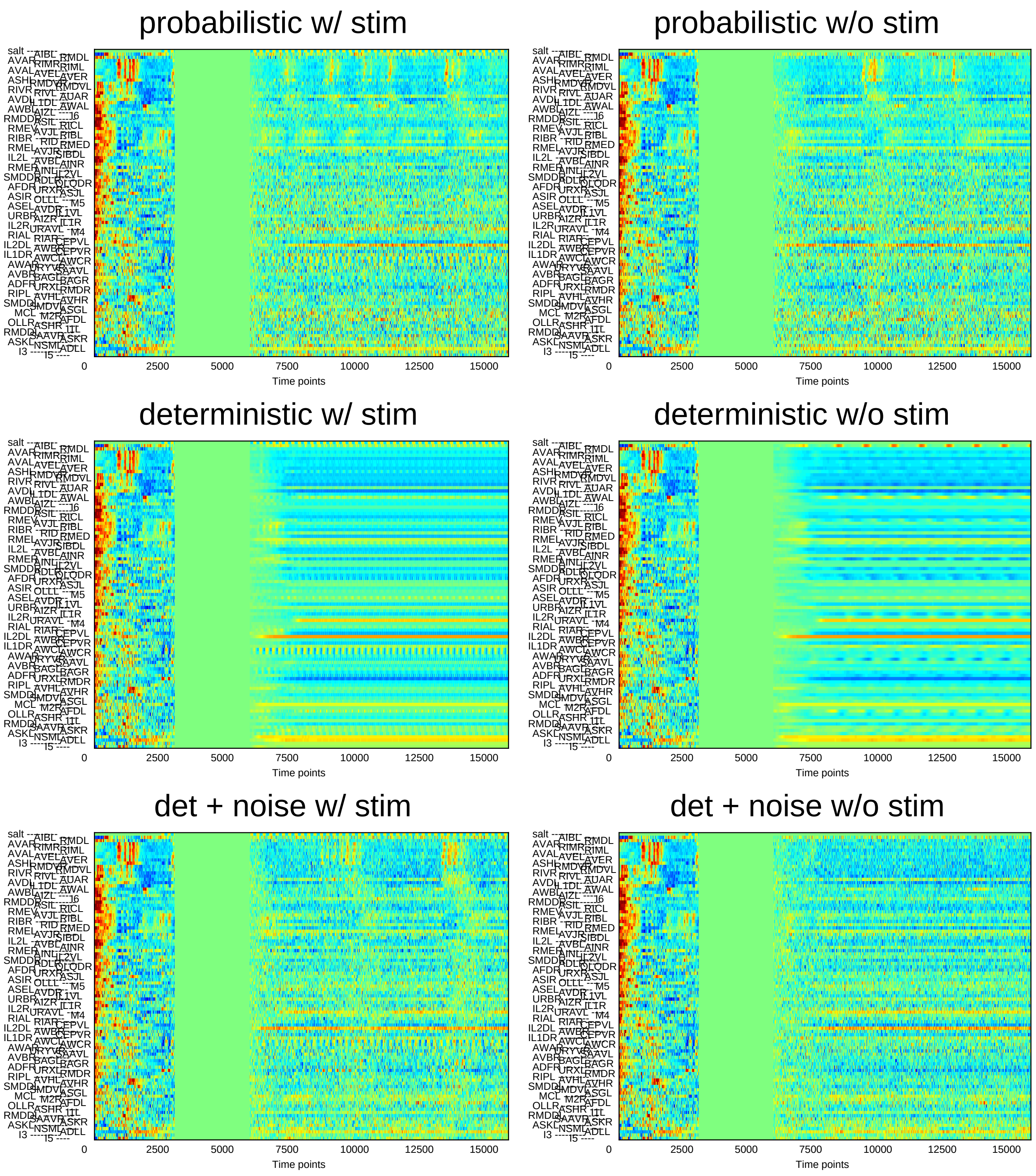
probabilistic w/ stim
probabilistic w/o stim
deterministic w/ stim
deterministic w/o stim
det + noise w/ stim
det + noise w/o stim
0
2500
5000
7500
10000
12500
15000
Time points

### Sample23

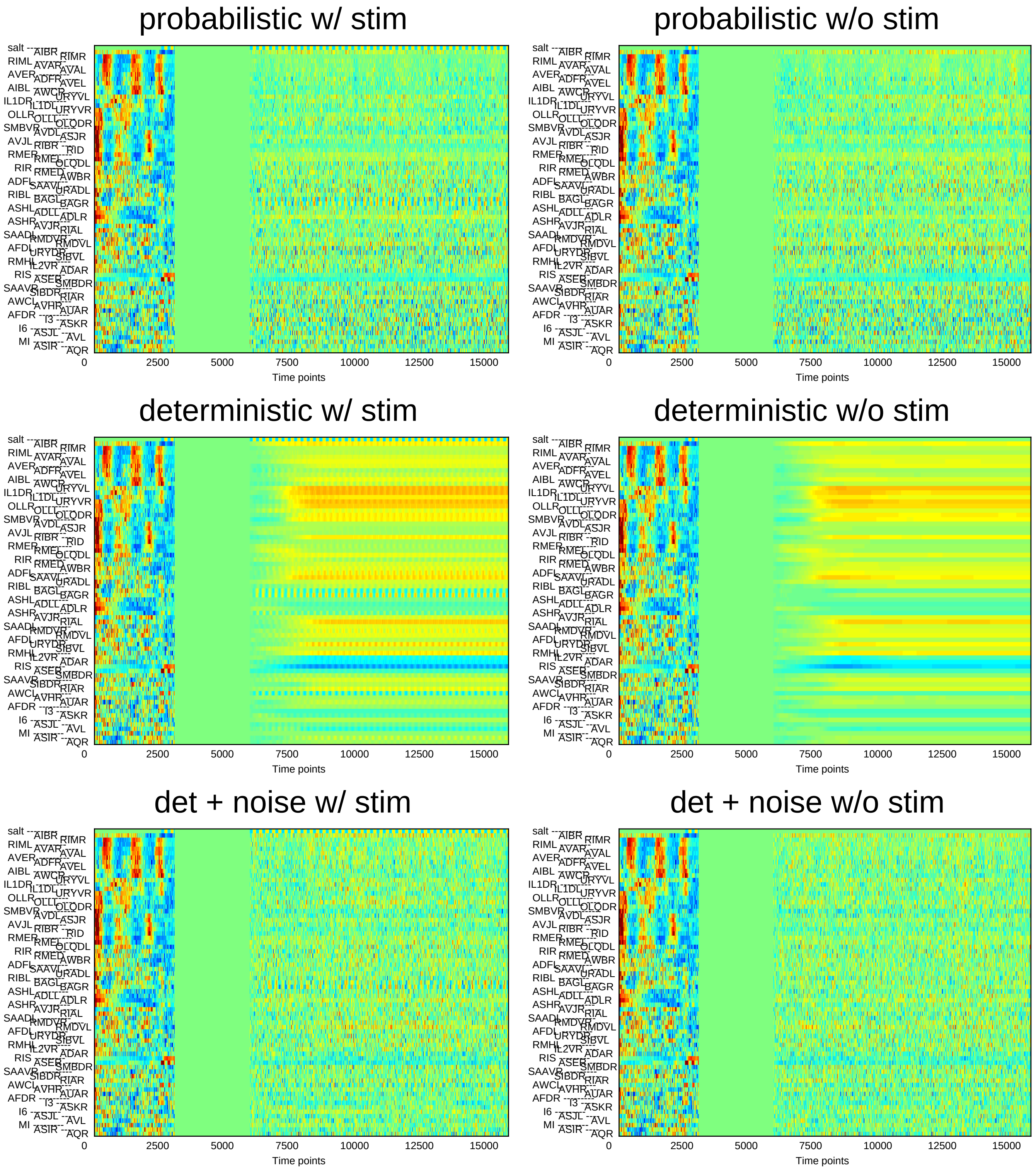
probabilistic w/ stim
probabilistic w/o stim
deterministic w/ stim
deterministic w/o stim
det + noise w/ stim
det + noise w/o stim
0 2500 5000 7500 10000 12500 15000
Time points

### Sample24

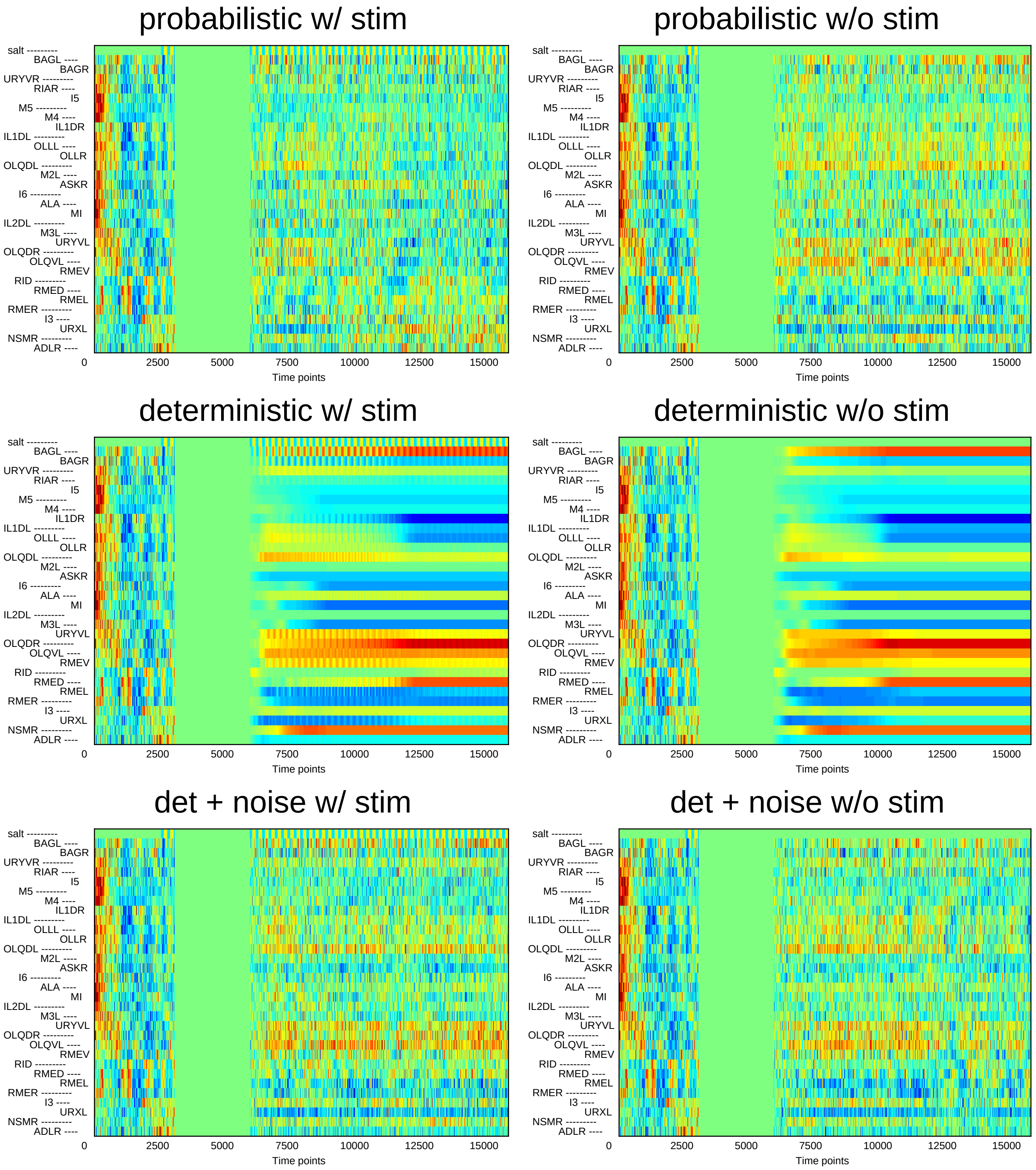
probabilistic w/ stim
probabilistic w/o stim
deterministic w/ stim
deterministic w/o stim
det + noise w/ stim
det + noise w/o stim
salt
BAGL
BAGR
URYVR
RIAR
I5
M5
M4
IL1DR
IL1DL
OLLL
OLLR
OLQDL
M2L
ASKR
I6
ALA
MI
IL2DL
M3L
URYVL
OLQDR
OLQVL
RMEV
RID
RMED
RMEL
RMER
I3
URXL
NSMR
ADLR
0
2500
5000
7500
10000
12500
15000
Time points
