## Supplementary File 11A for "Deducing ensemble dynamics and information flow from the whole-brain imaging data"

### Sample1 connection weight matrix

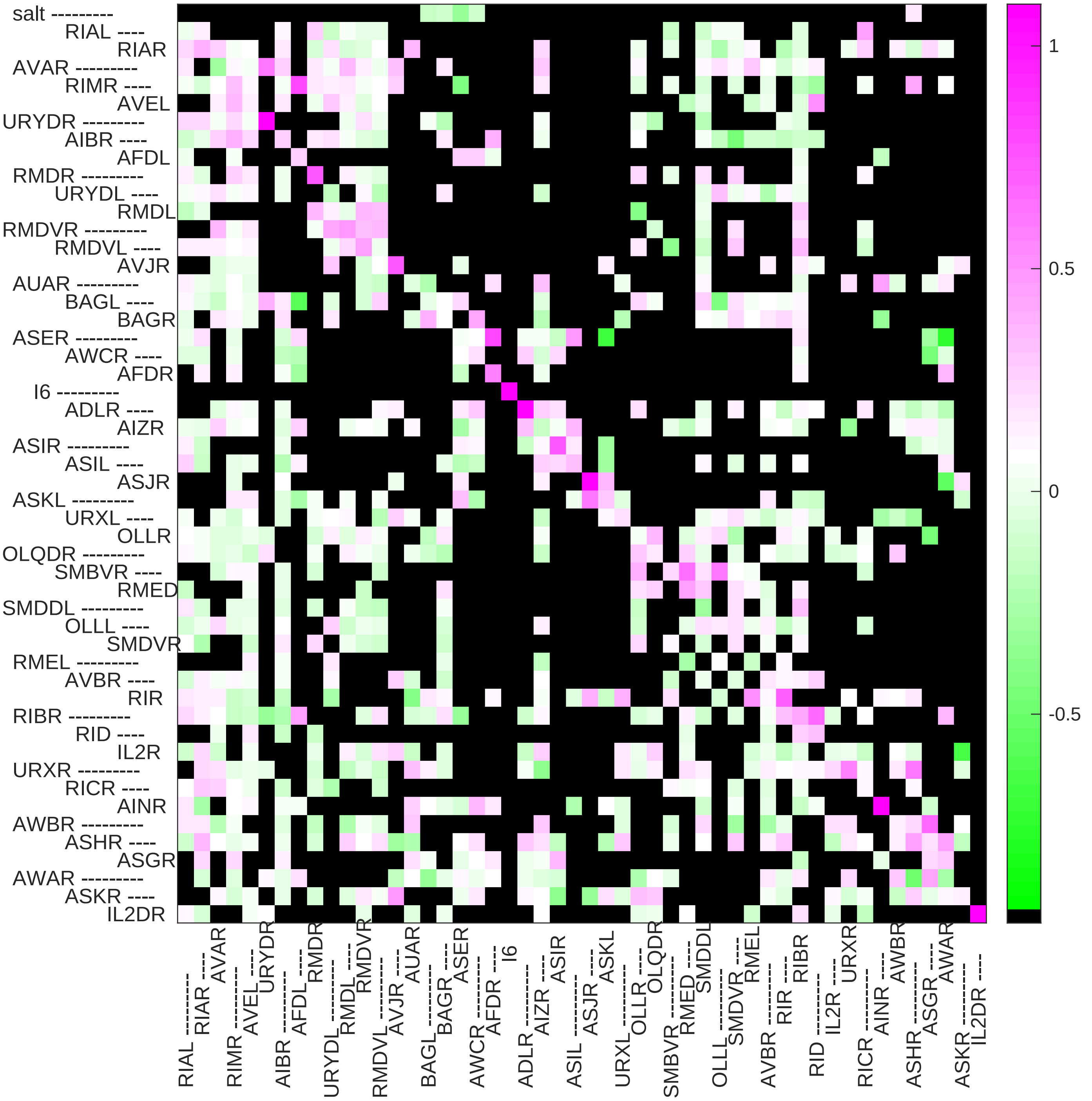

### Sample2 connection weight matrix

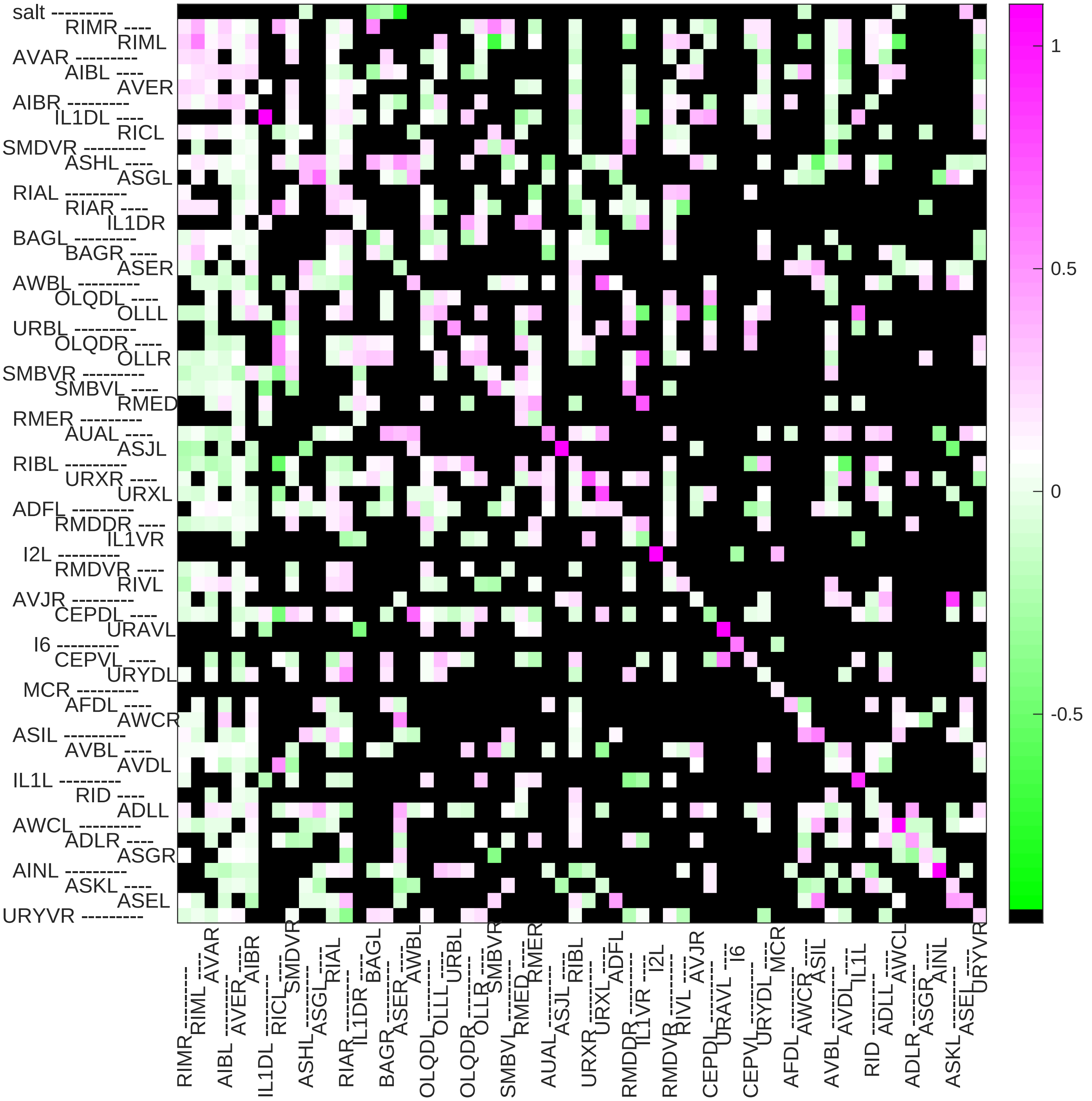

### Sample3 connection weight matrix

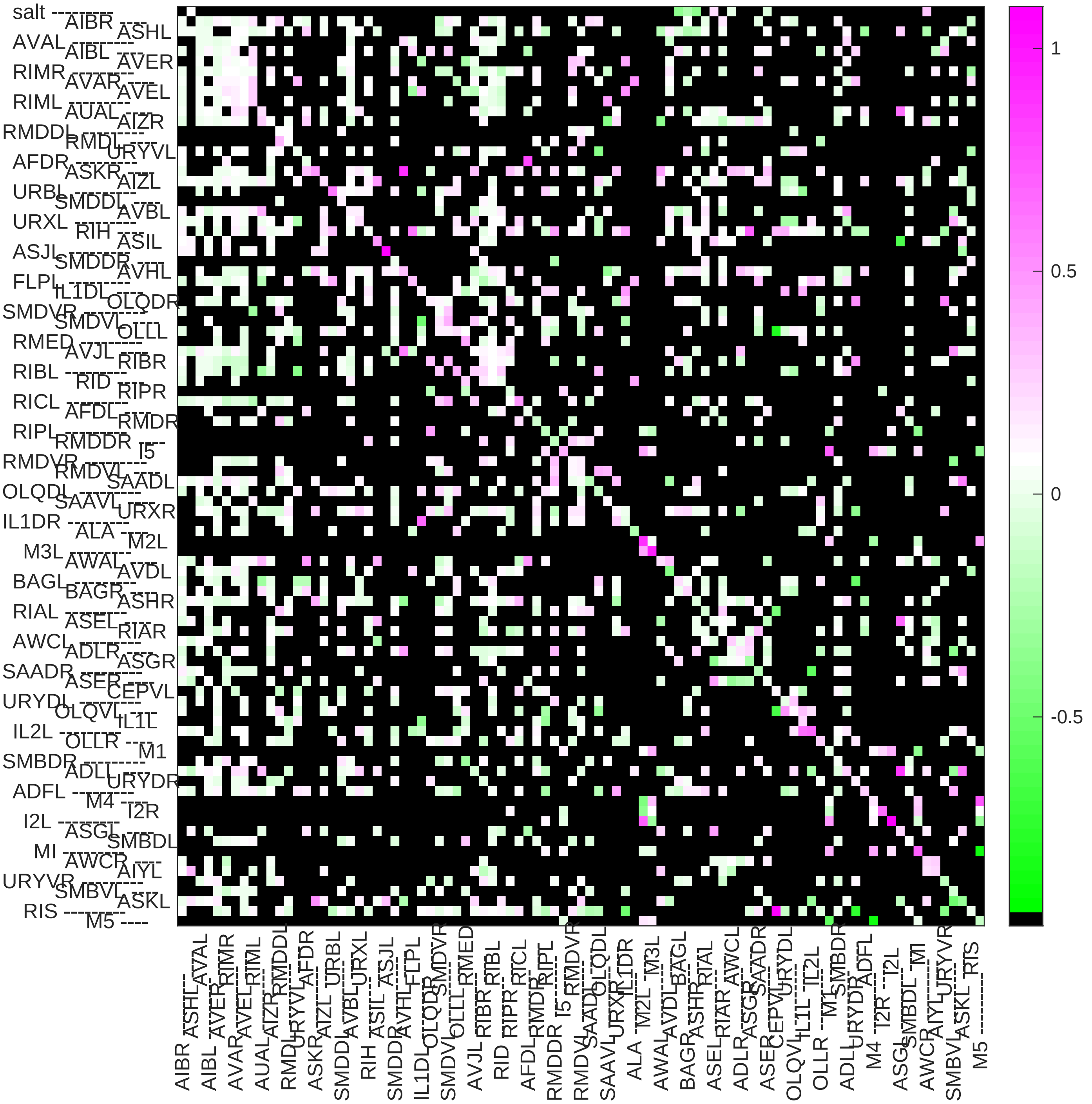

### Sample4 connection weight matrix

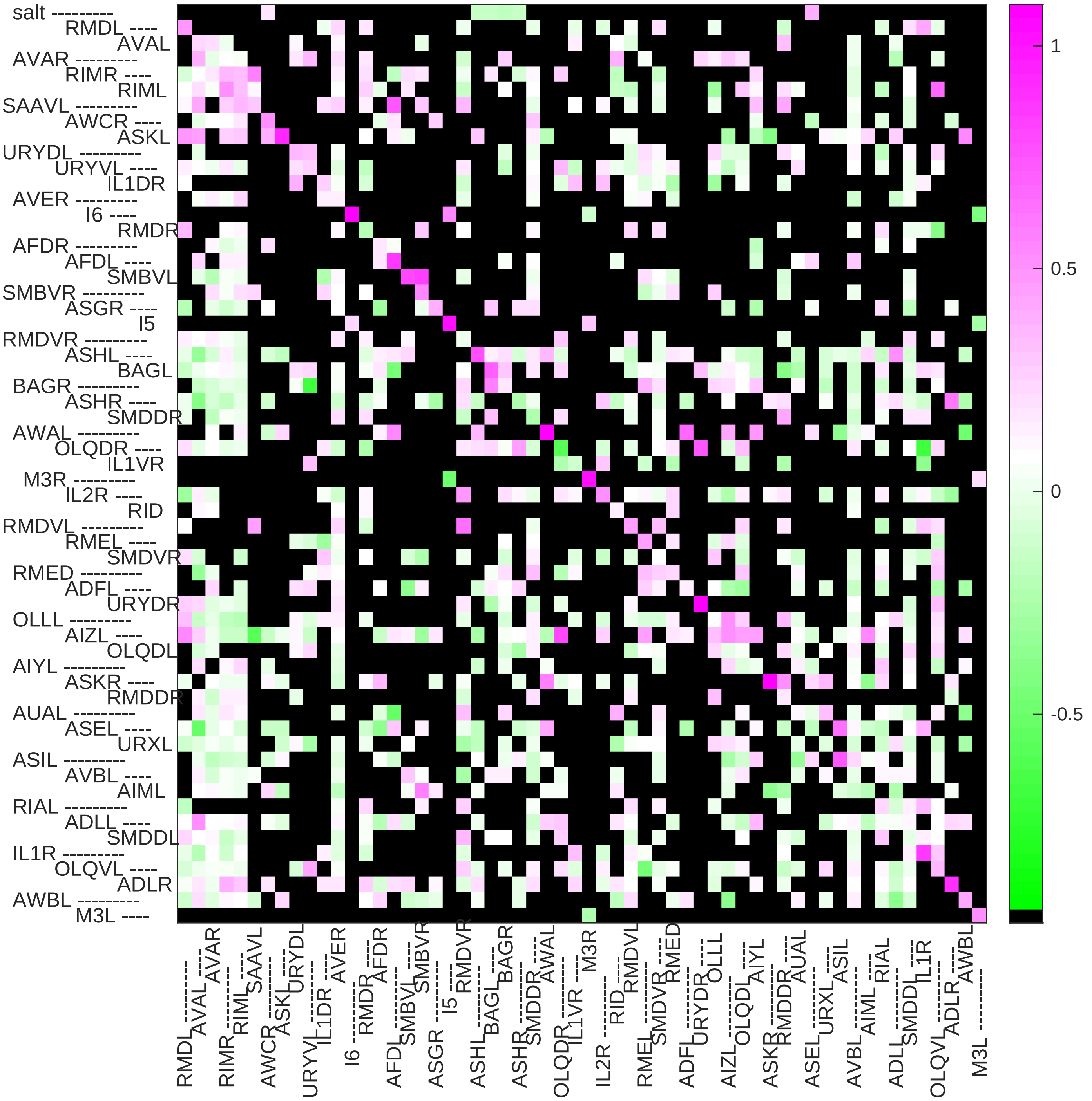

Sample5 connection weight matrix

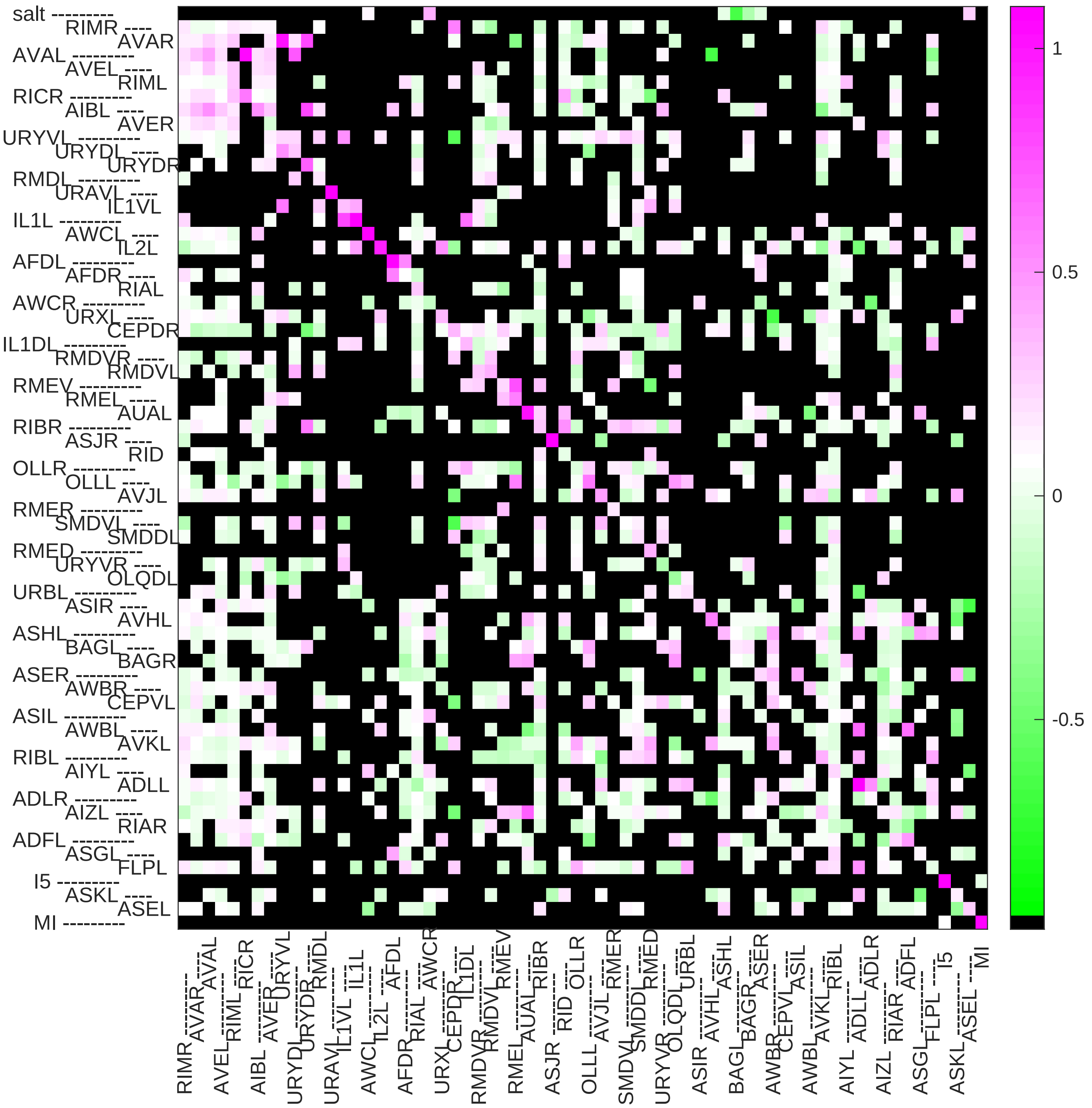

### Sample6 connection weight matrix

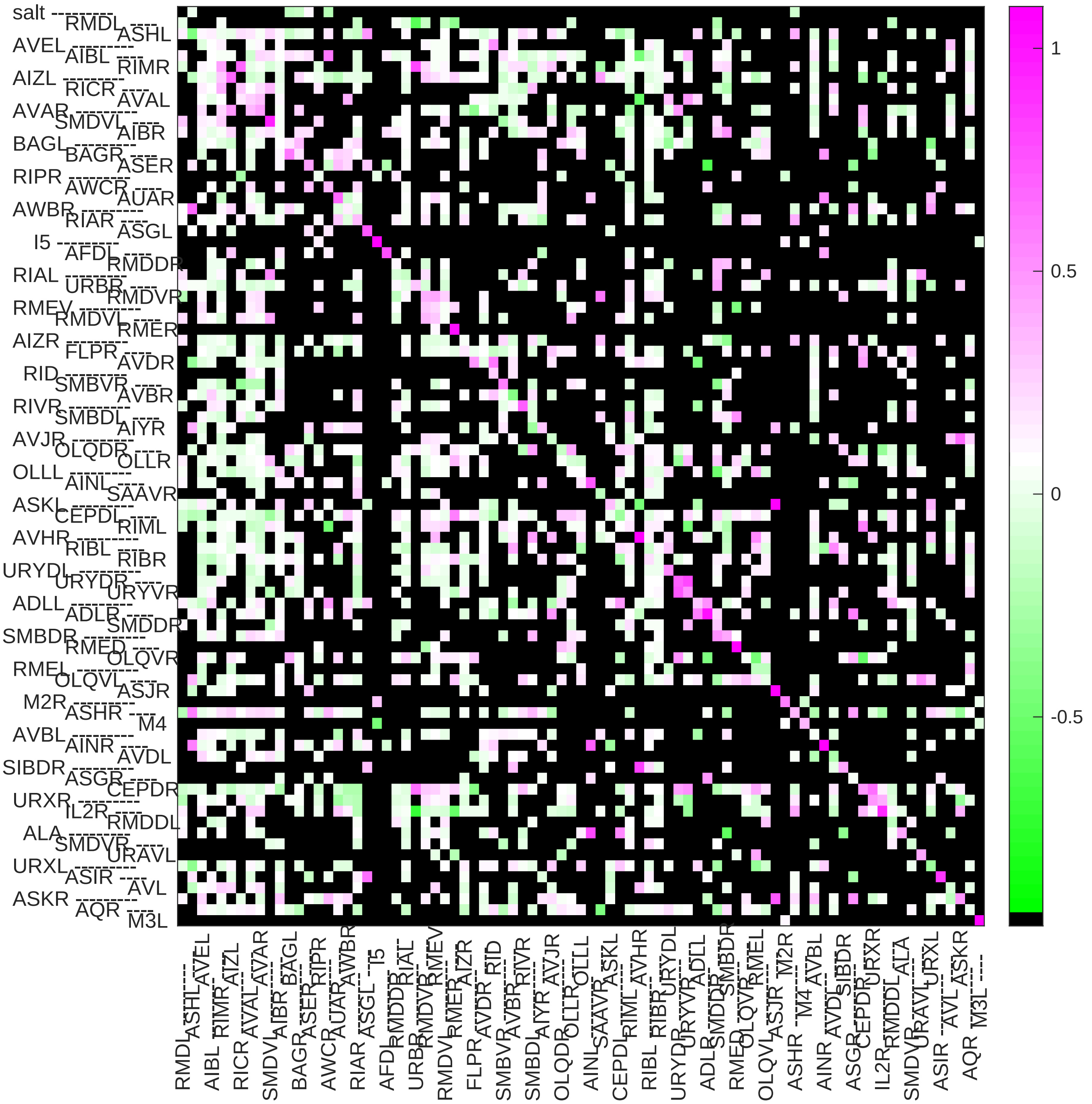

### Sample7 connection weight matrix

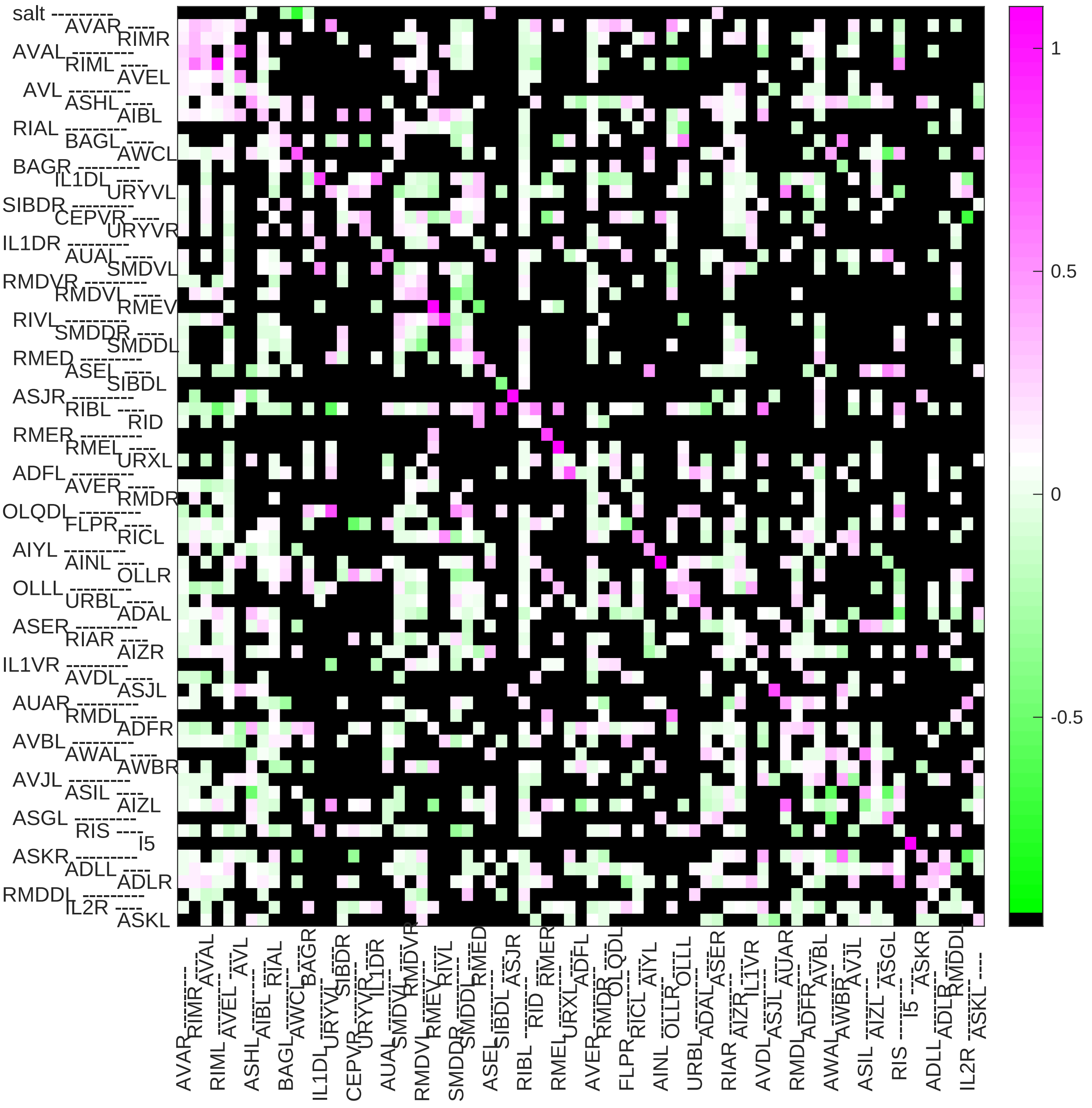

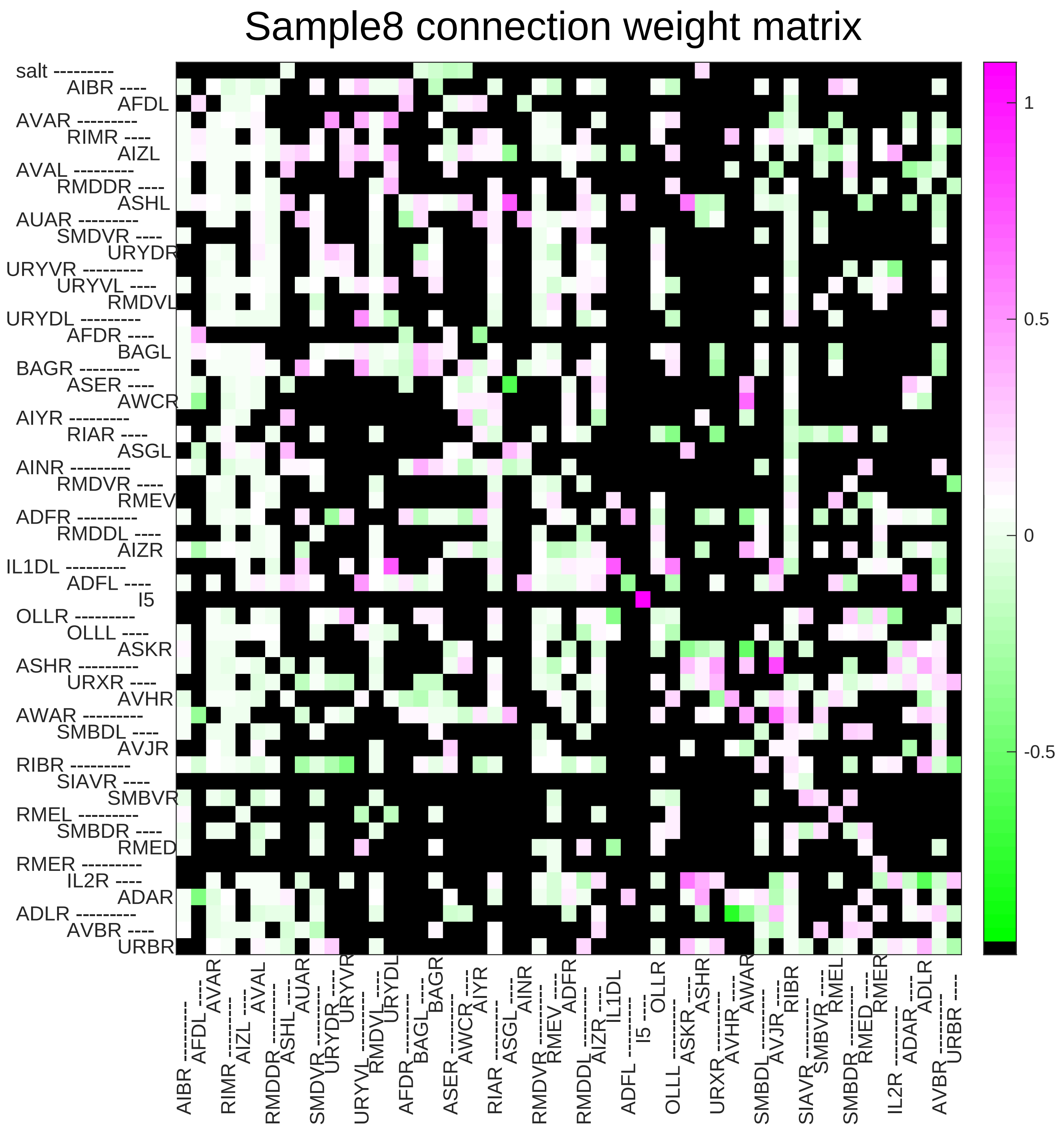

Sample8 connection weight matrix
salt ---------
AIBR ----
AFDL
AVAR ---------
RIMR ----
AIZL
AVAL ---------
RMDDR ----
ASHL
AUAR ---------
SMDVR ----
URYDR
URYVR ---------
URYVL ----
RMDVL
URYDL ---------
AFDR ----
BAGL
BAGR ---------
ASER ----
AWCR
AIYR ---------
RIAR ----
ASGL
AINR ---------
RMDVR ----
RMEV
ADFR ---------
RMDDL ----
AIZR
IL1DL ---------
ADFL ----
I5
OLLR ---------
OLLL ----
ASKR
ASHR ---------
URXR ----
AVHR
AWAR ---------
SMBDL ----
AVJR
RIBR ---------
SIAVR ----
SMBVR
RMEL ---------
SMBDR ----
RMED
RMER ---------
IL2R ----
ADAR
ADLR ---------
AVBR ----
URBR
1
0.5
0
-0.5

### Sample9 connection weight matrix

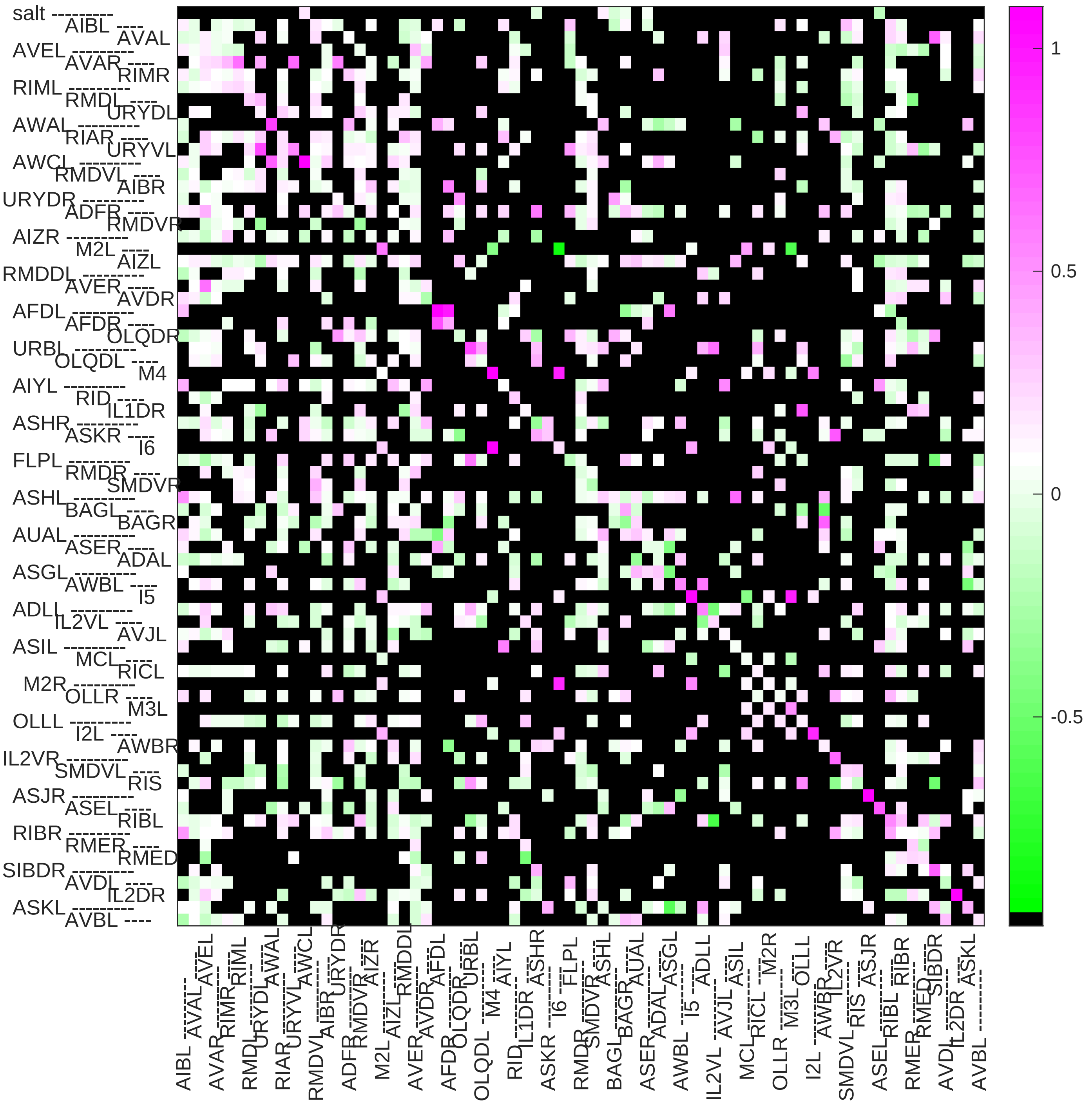

Sample10 connection weight matrix

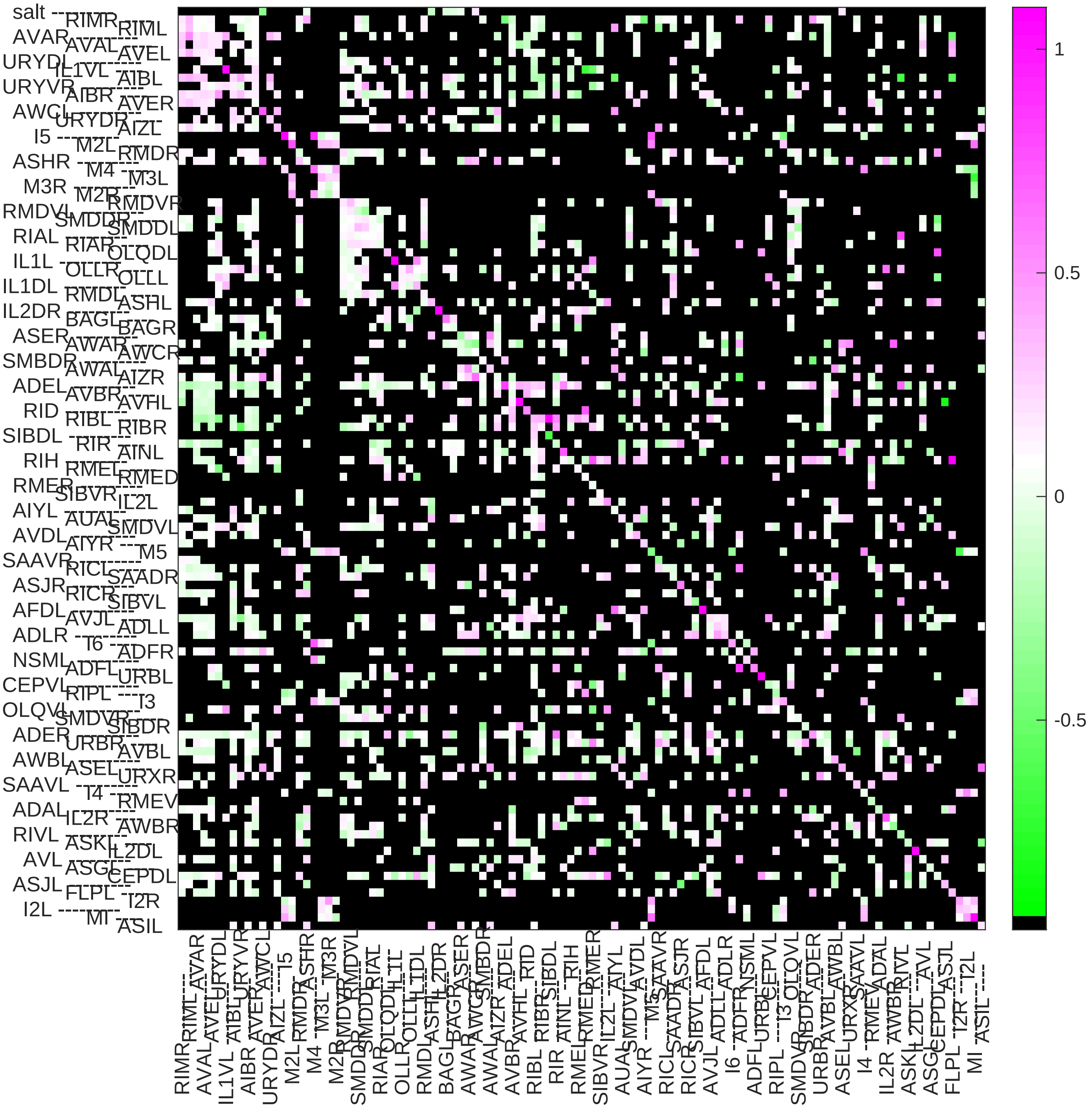

### Sample11 connection weight matrix

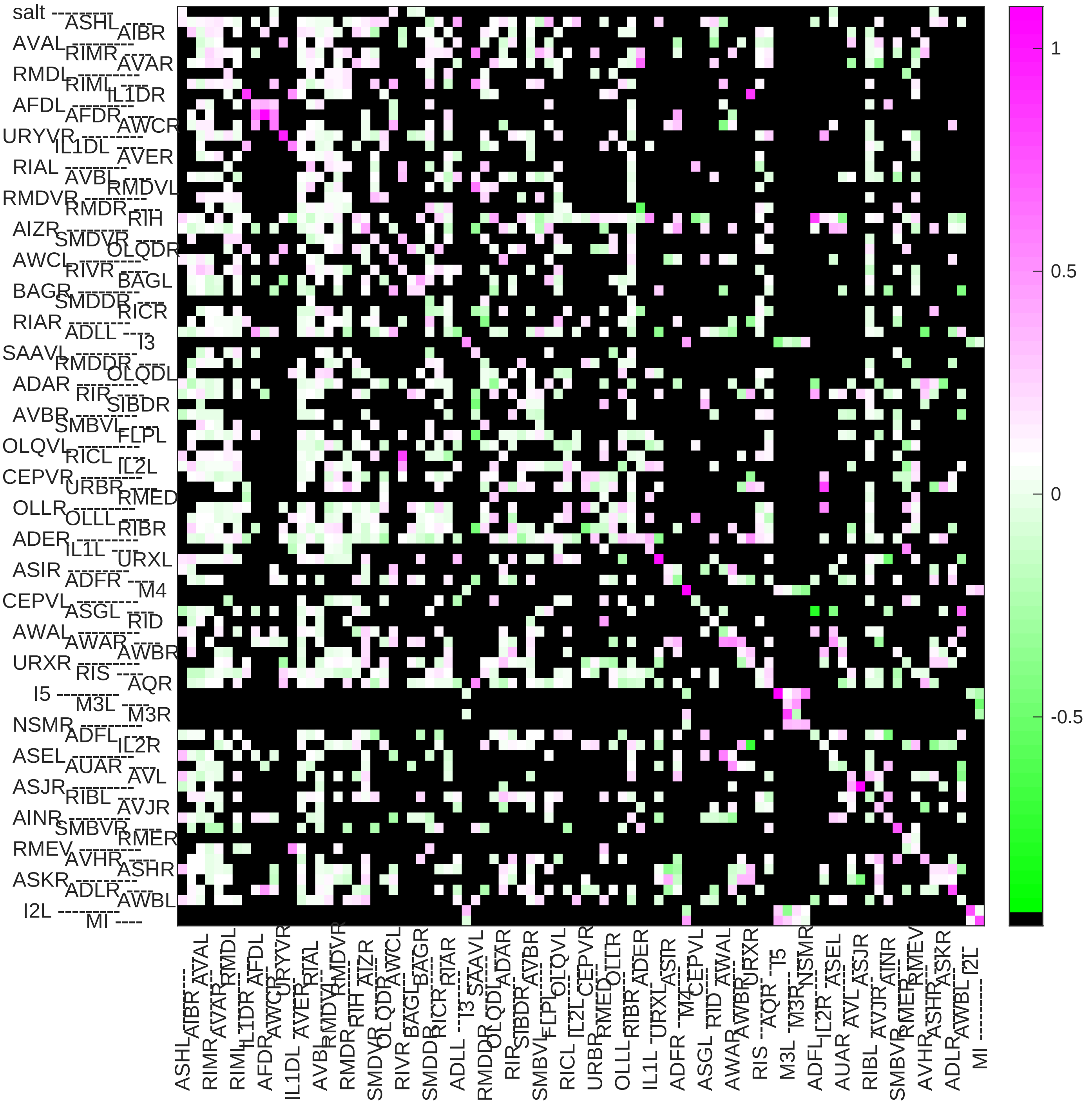

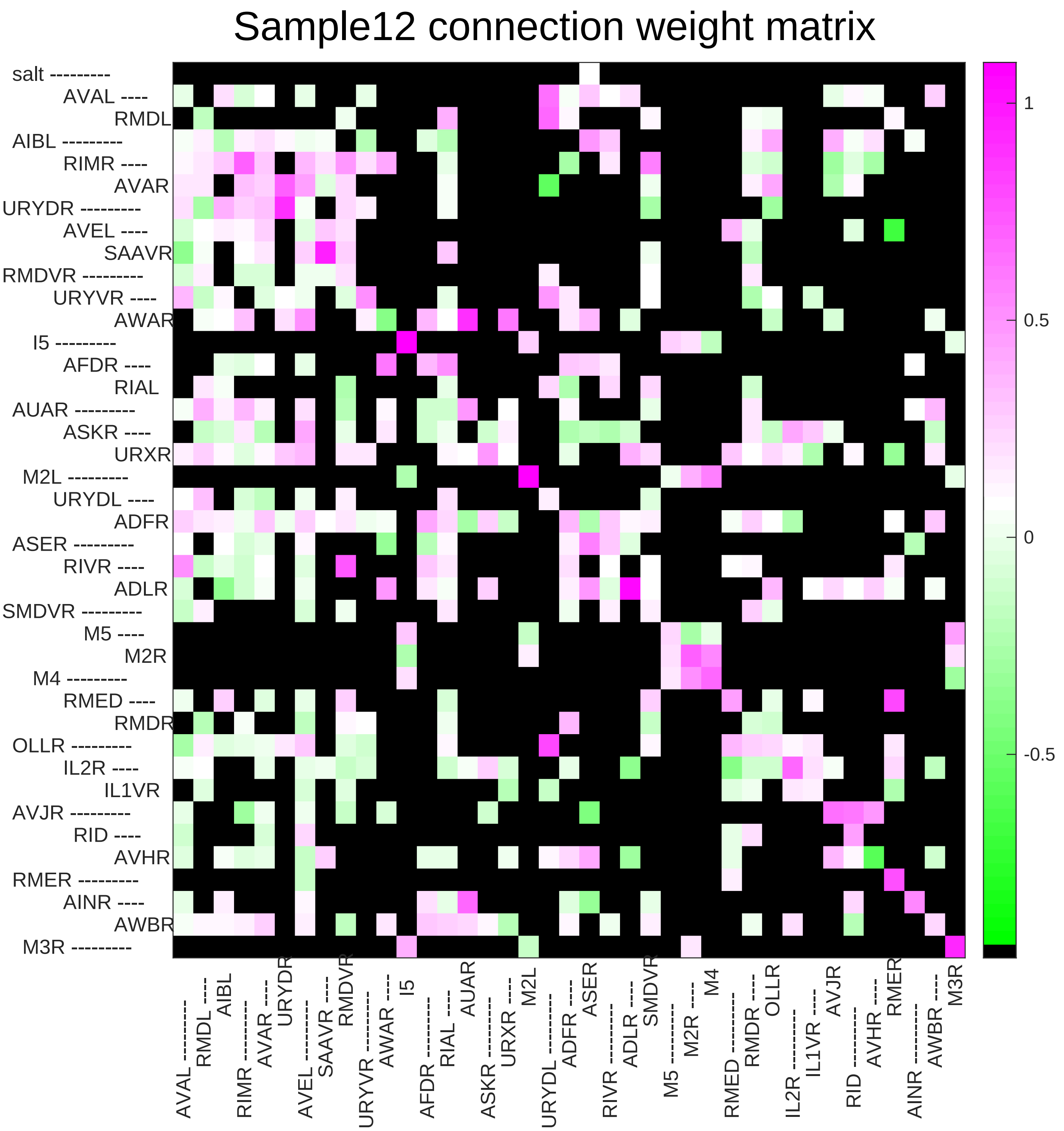

Sample12 connection weight matrix
salt
AVAL
RMDL
AIBL
RIMR
AVAR
URYDR
AVEL
SAAVR
RMDVR
URYVR
AWAR
I5
AFDR
RIAL
AUAR
ASKR
URXR
M2L
URYDL
ADFR
ASER
RIVR
ADLR
SMDVR
M5
M2R
M4
RMED
RMDR
OLLR
IL2R
IL1VR
AVJR
RID
AVHR
RMER
AINR
AWBR
M3R
1
0.5
0
-0.5

### Sample13 connection weight matrix

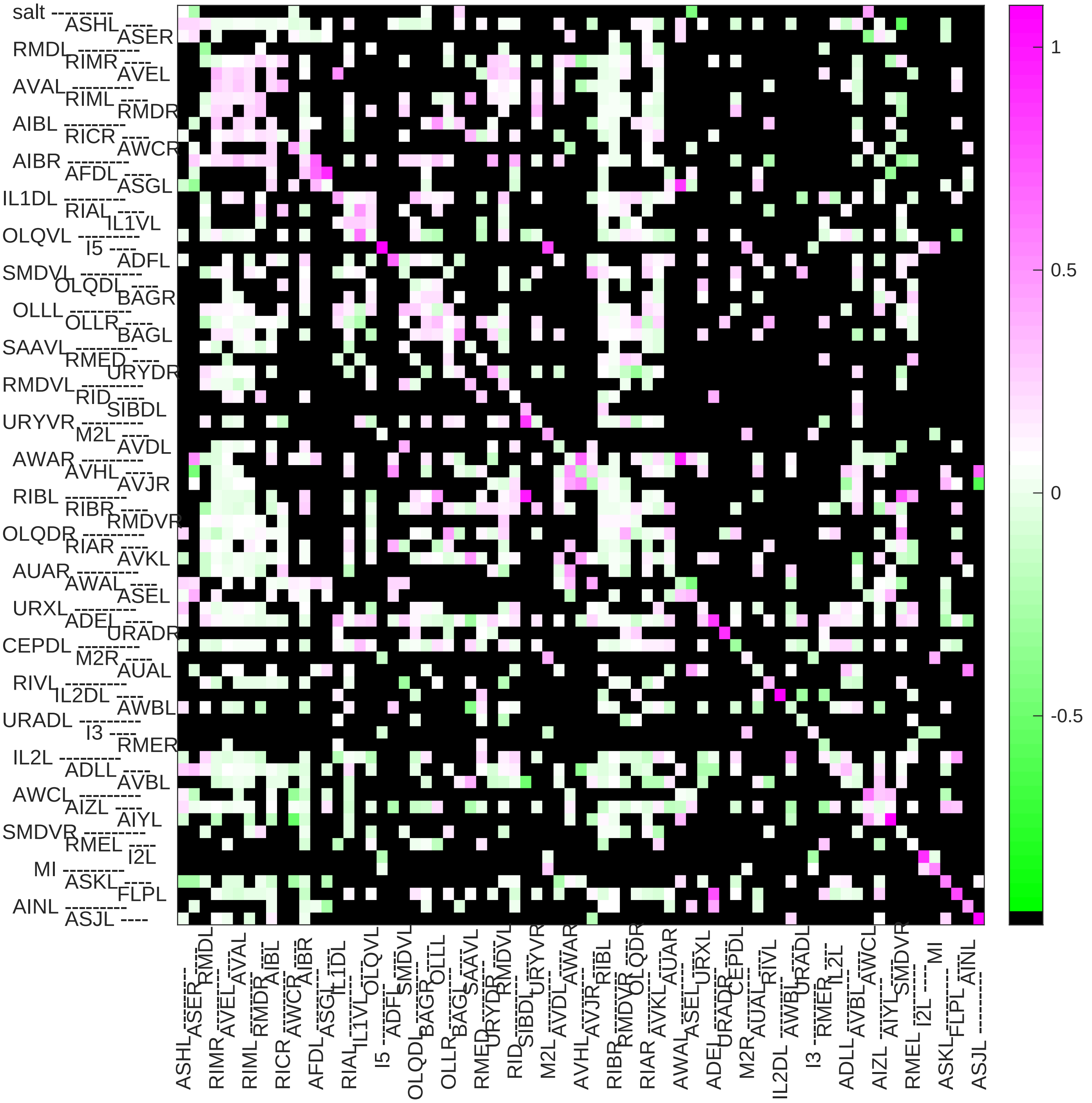

### Sample14 connection weight matrix

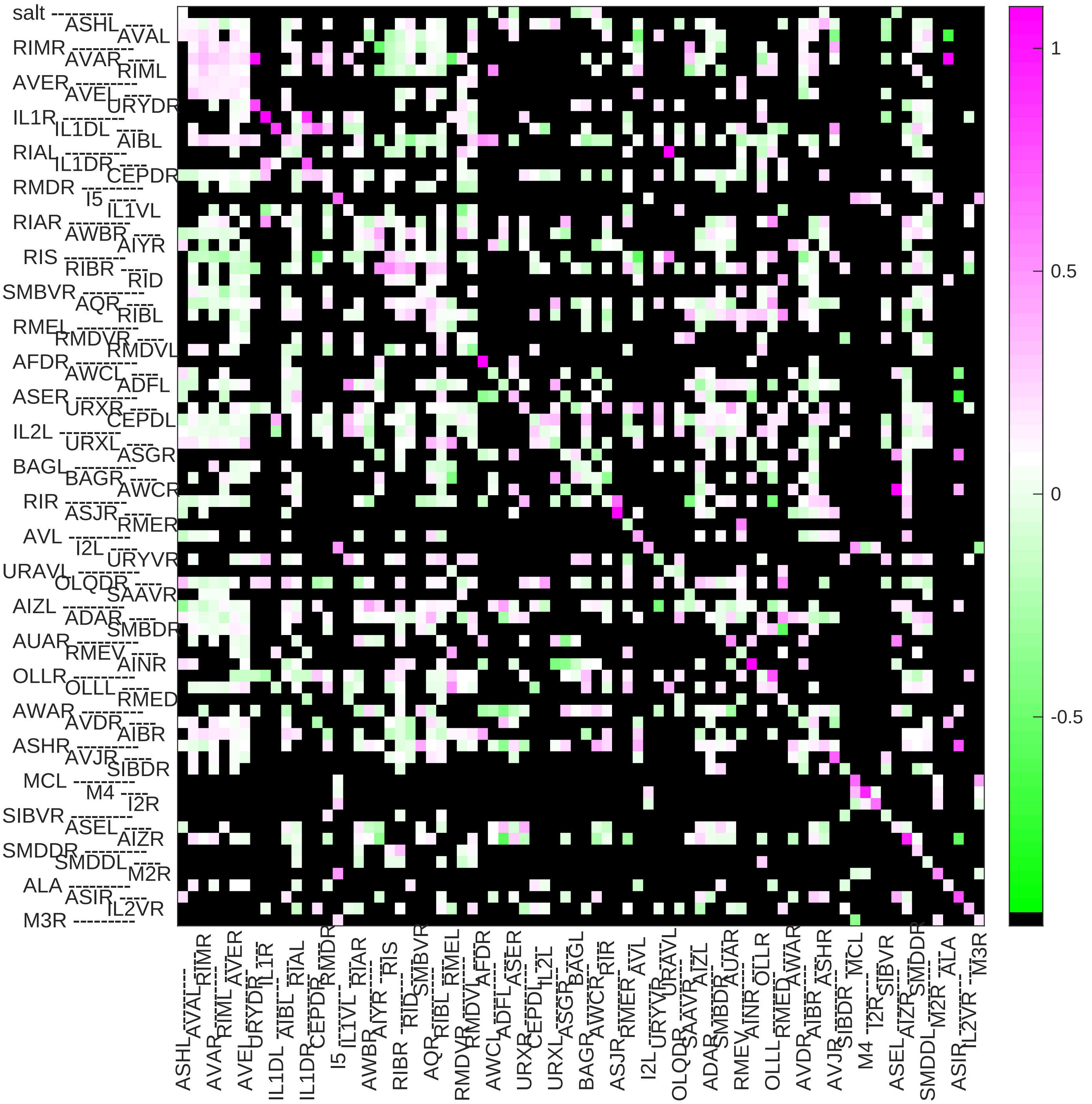

### Sample15 connection weight matrix

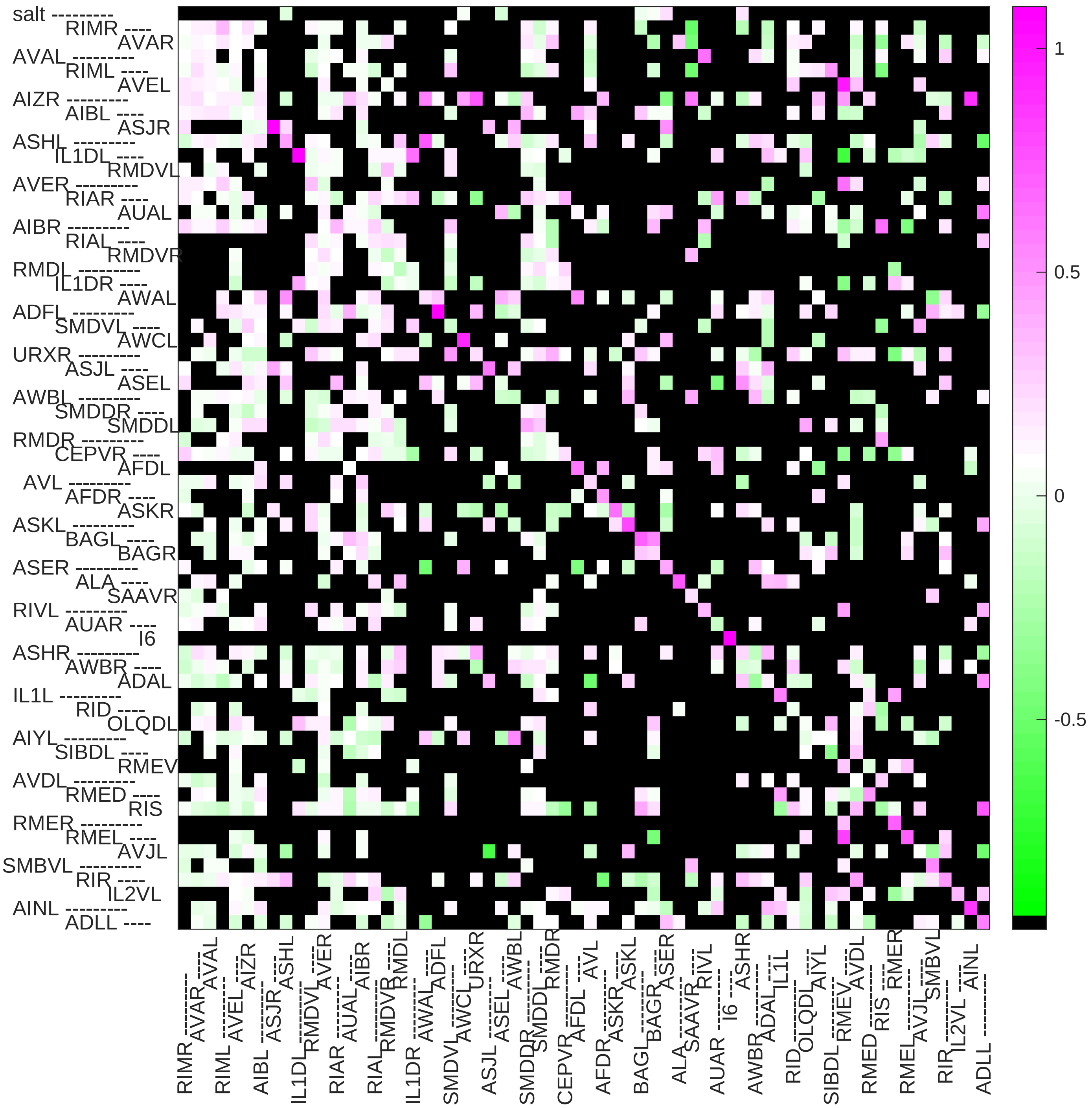

### Sample16 connection weight matrix

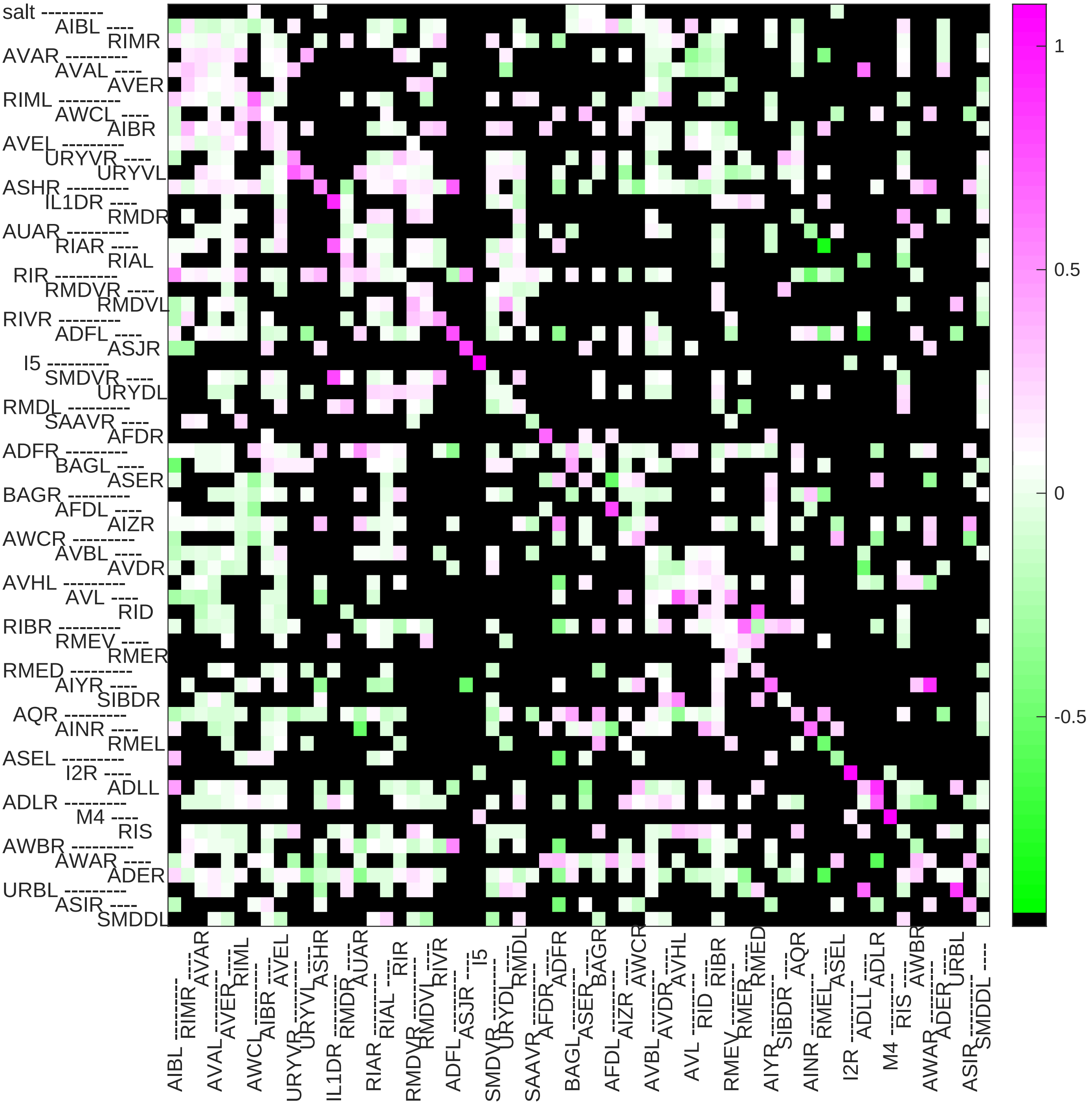

### Sample17 connection weight matrix

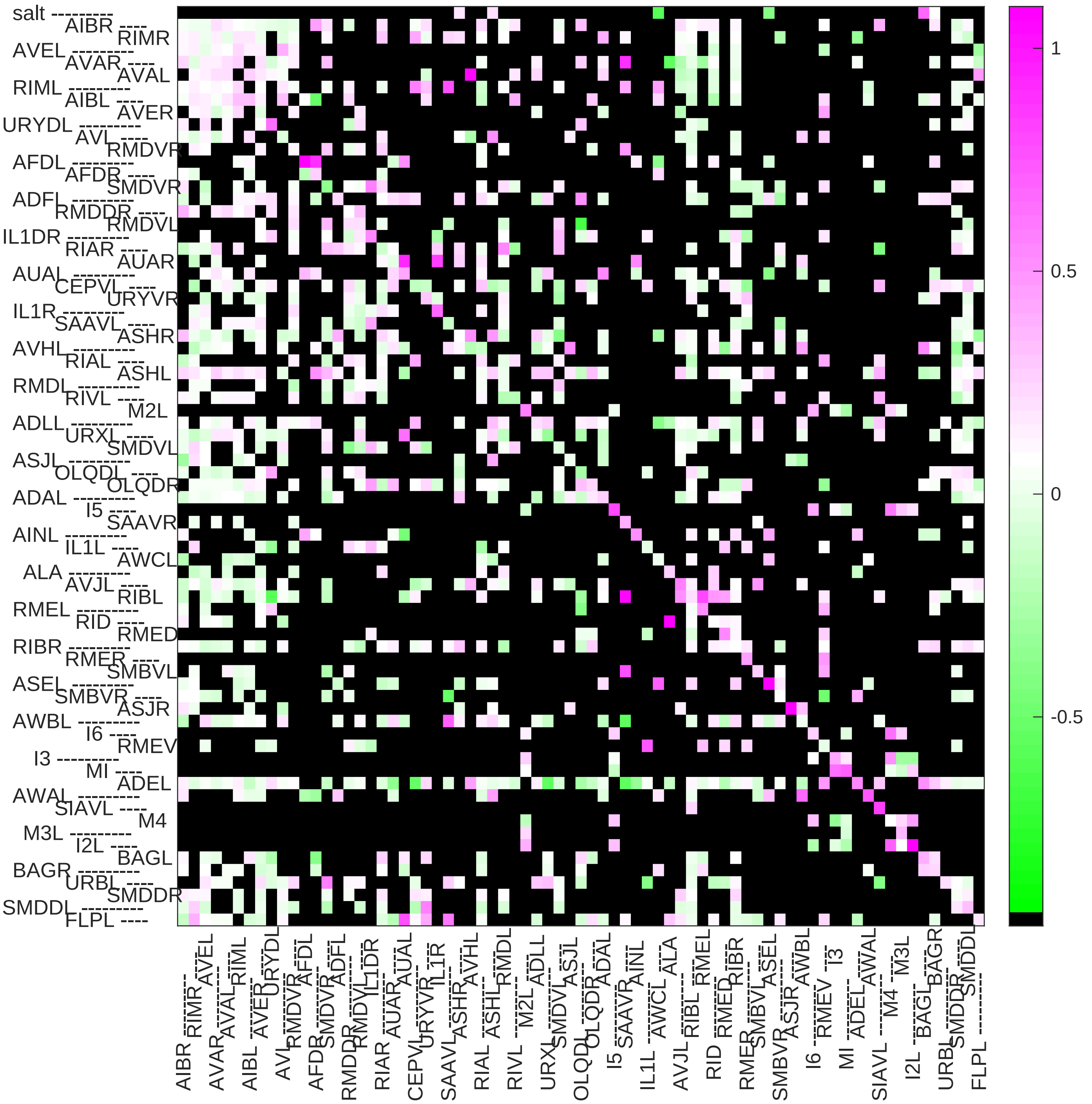

### Sample18 connection weight matrix

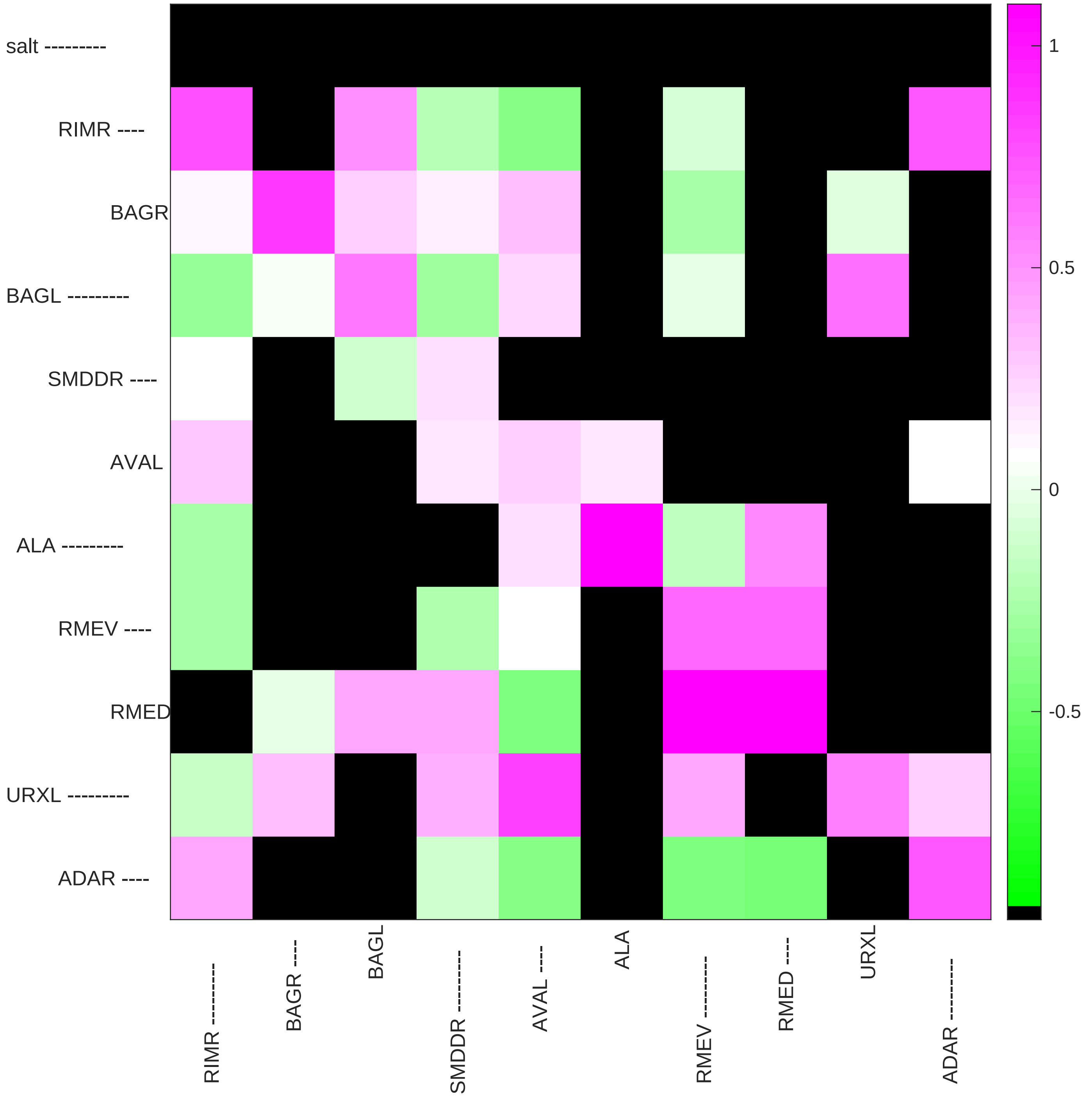

### Sample19 connection weight matrix

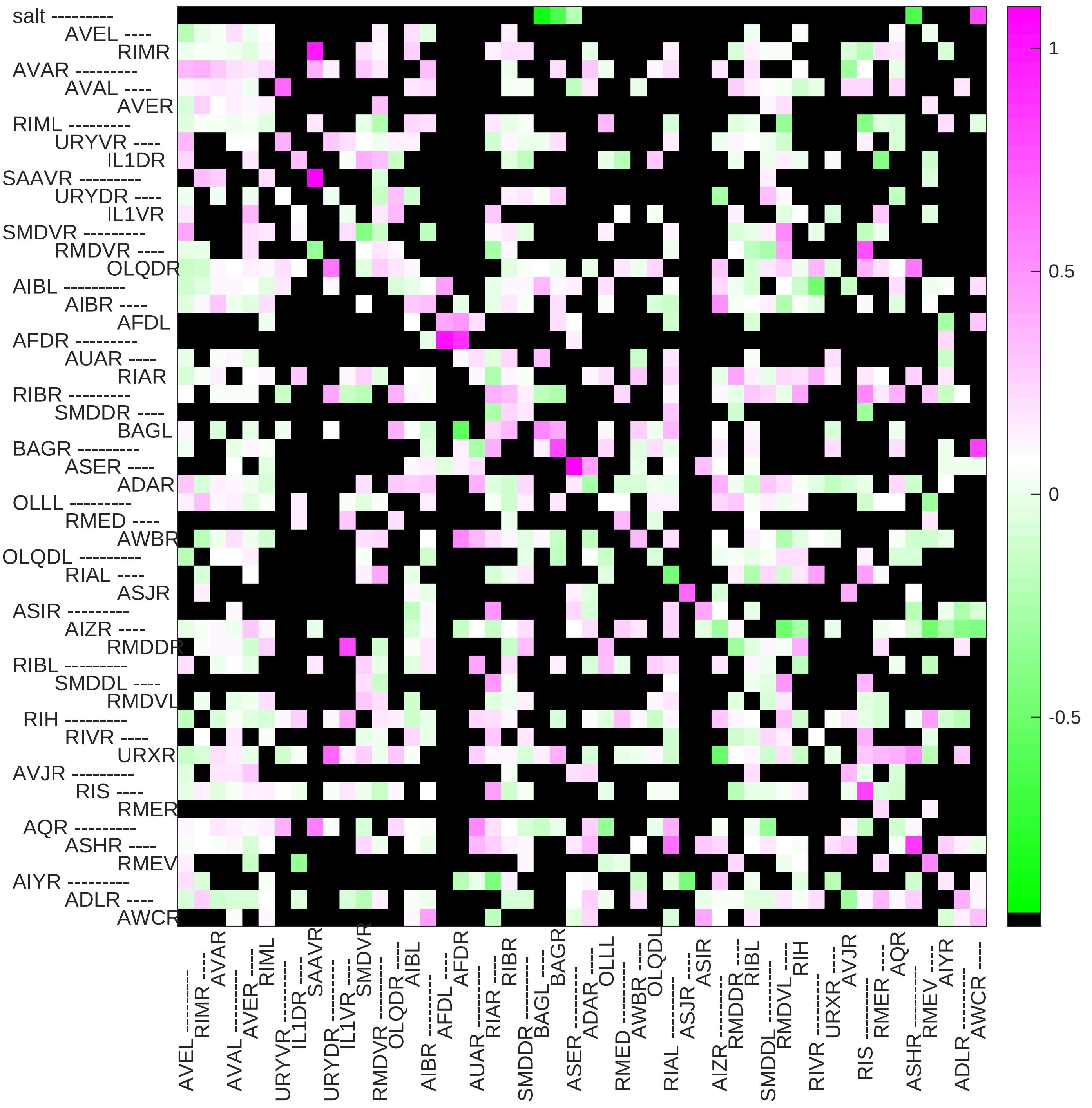

Sample20 connection weight matrix

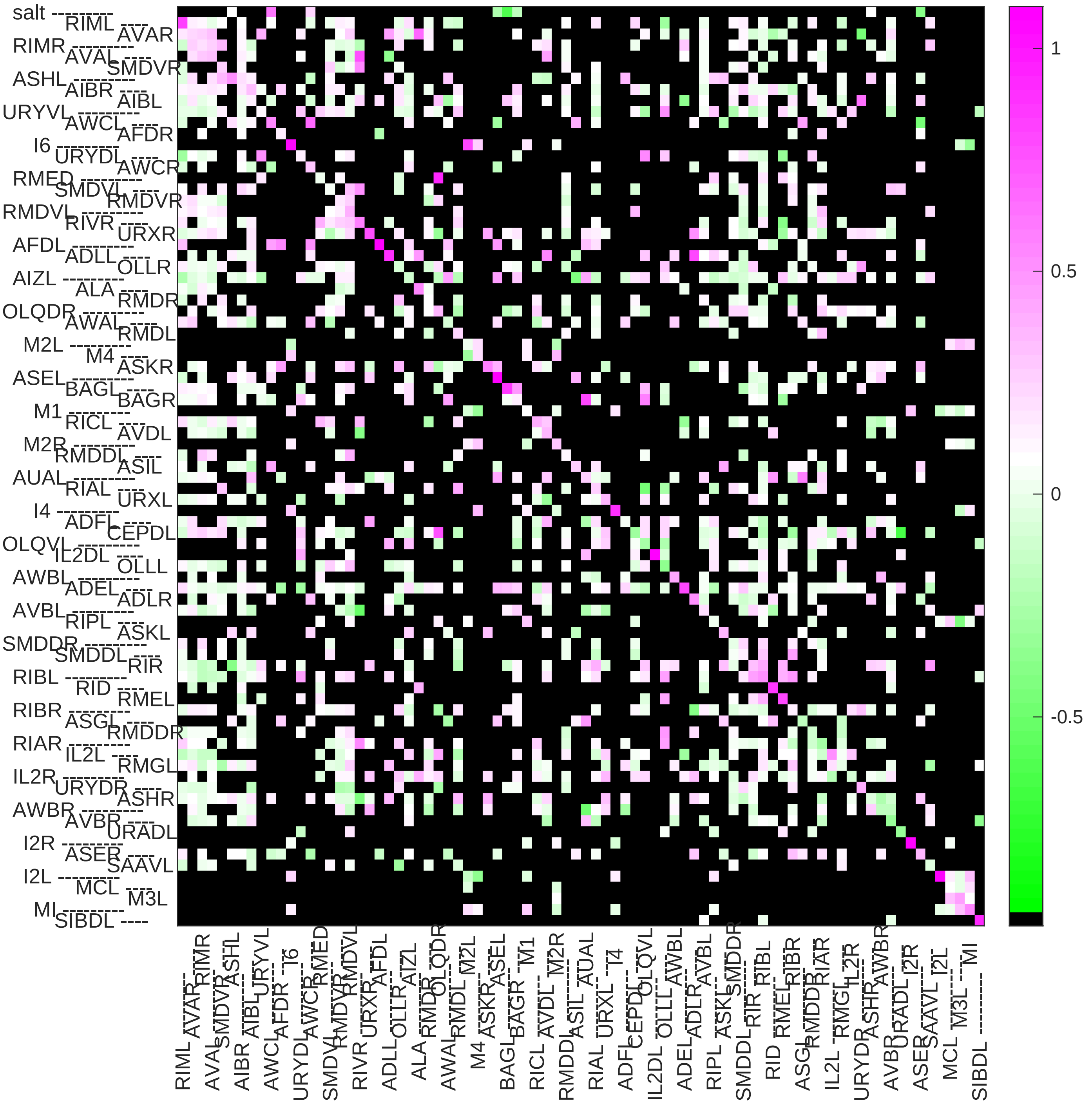

### Sample21 connection weight matrix

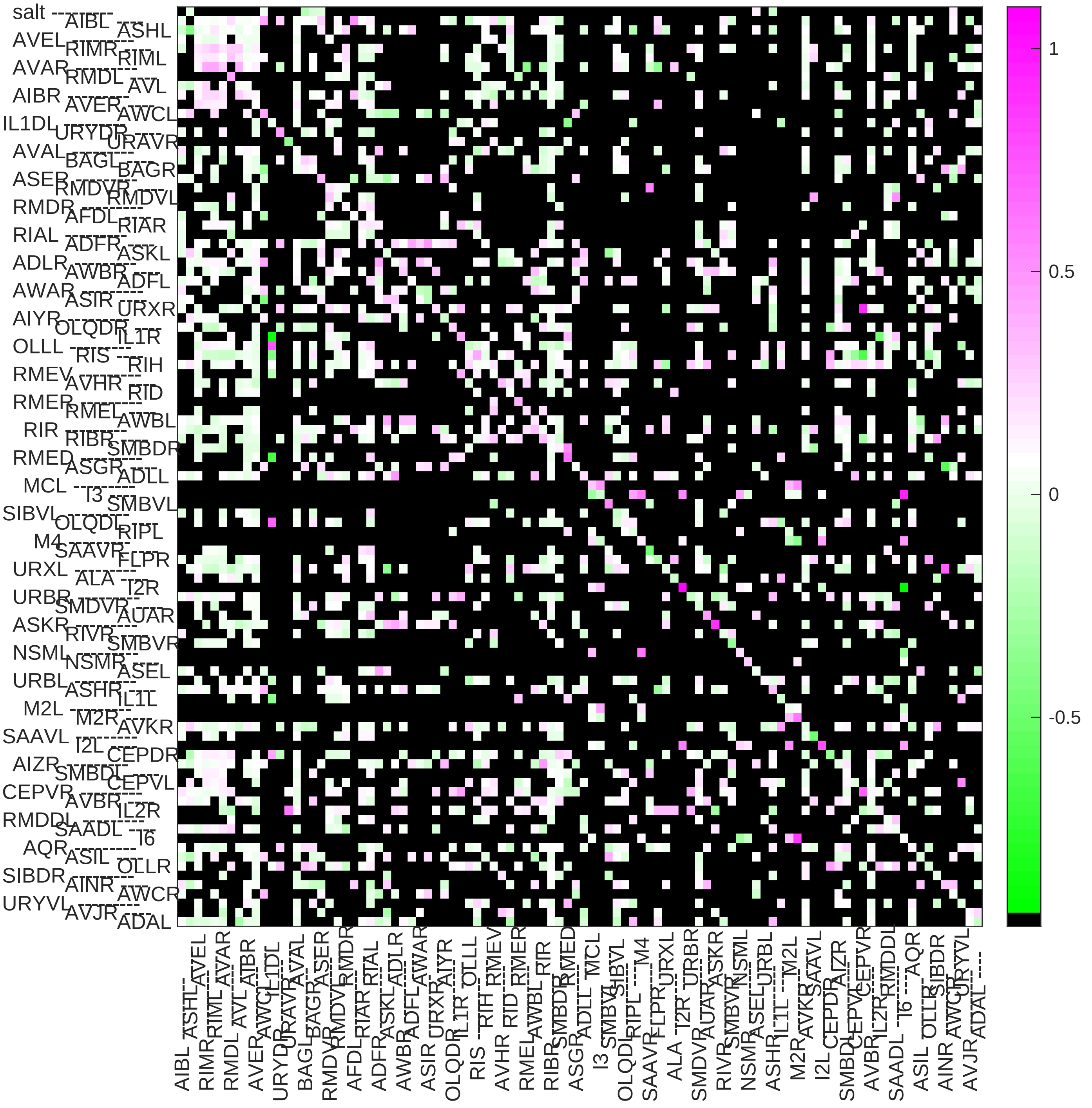

### Sample22 connection weight matrix

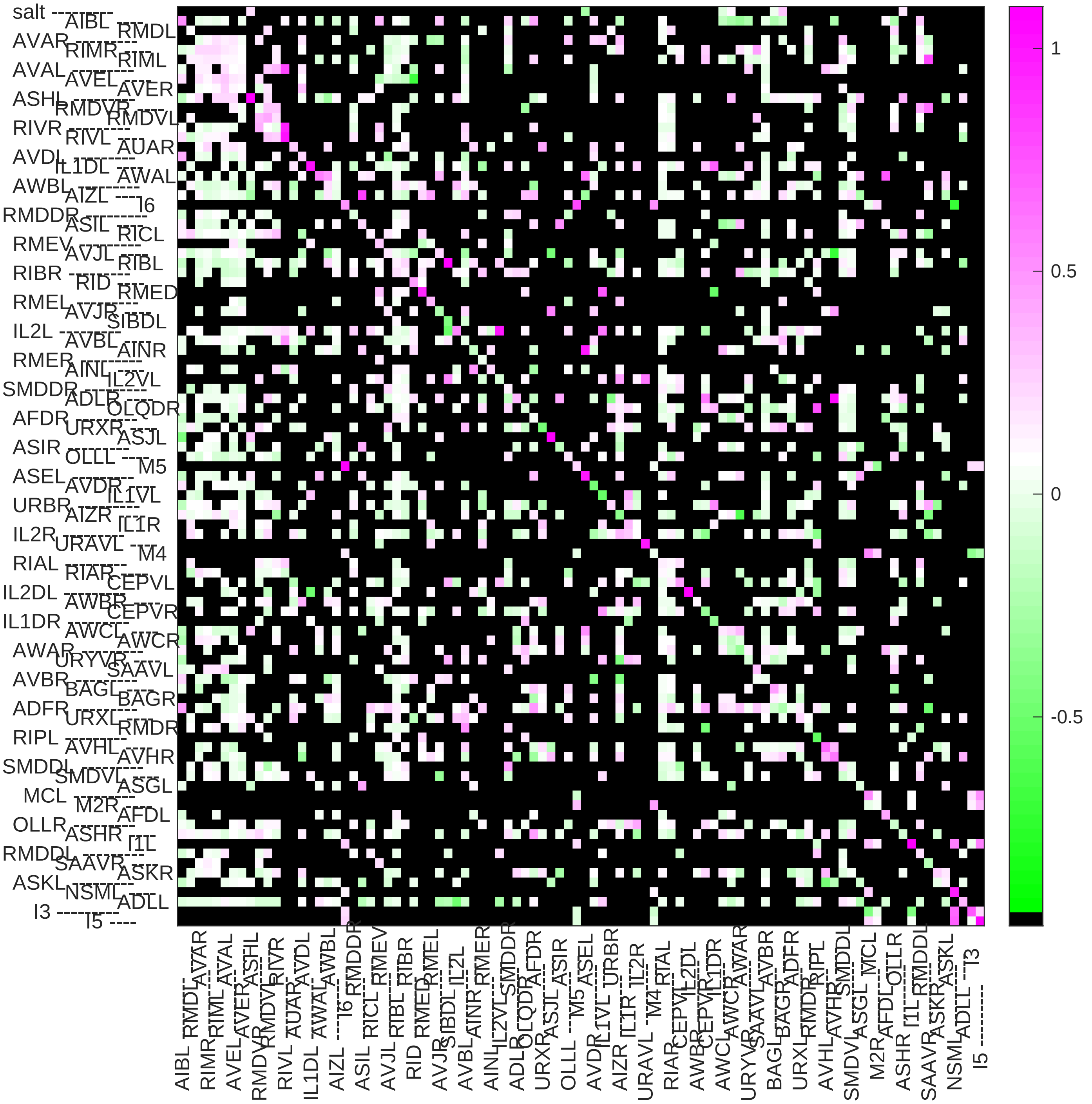

### Sample23 connection weight matrix

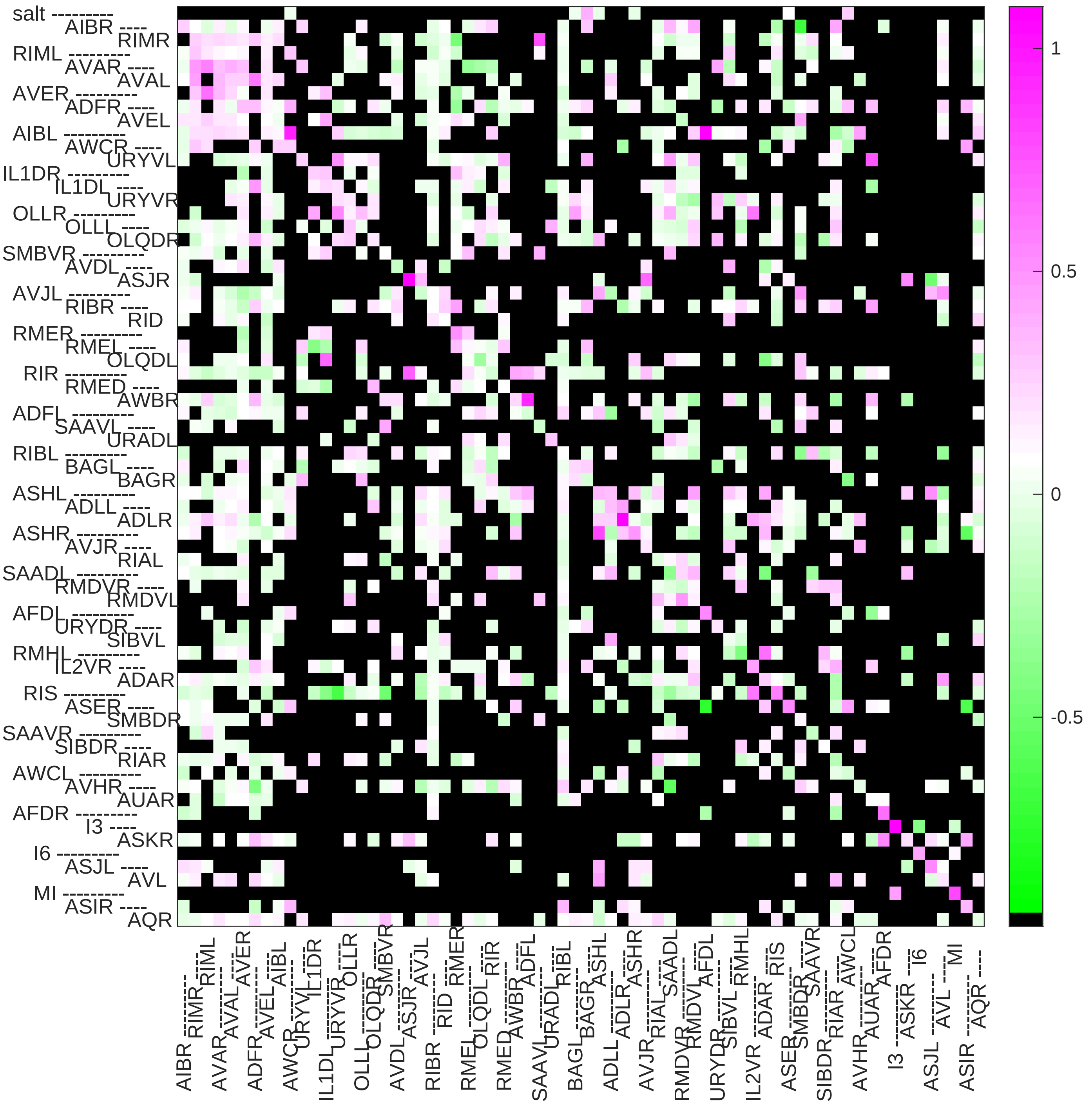

### Sample24 connection weight matrix

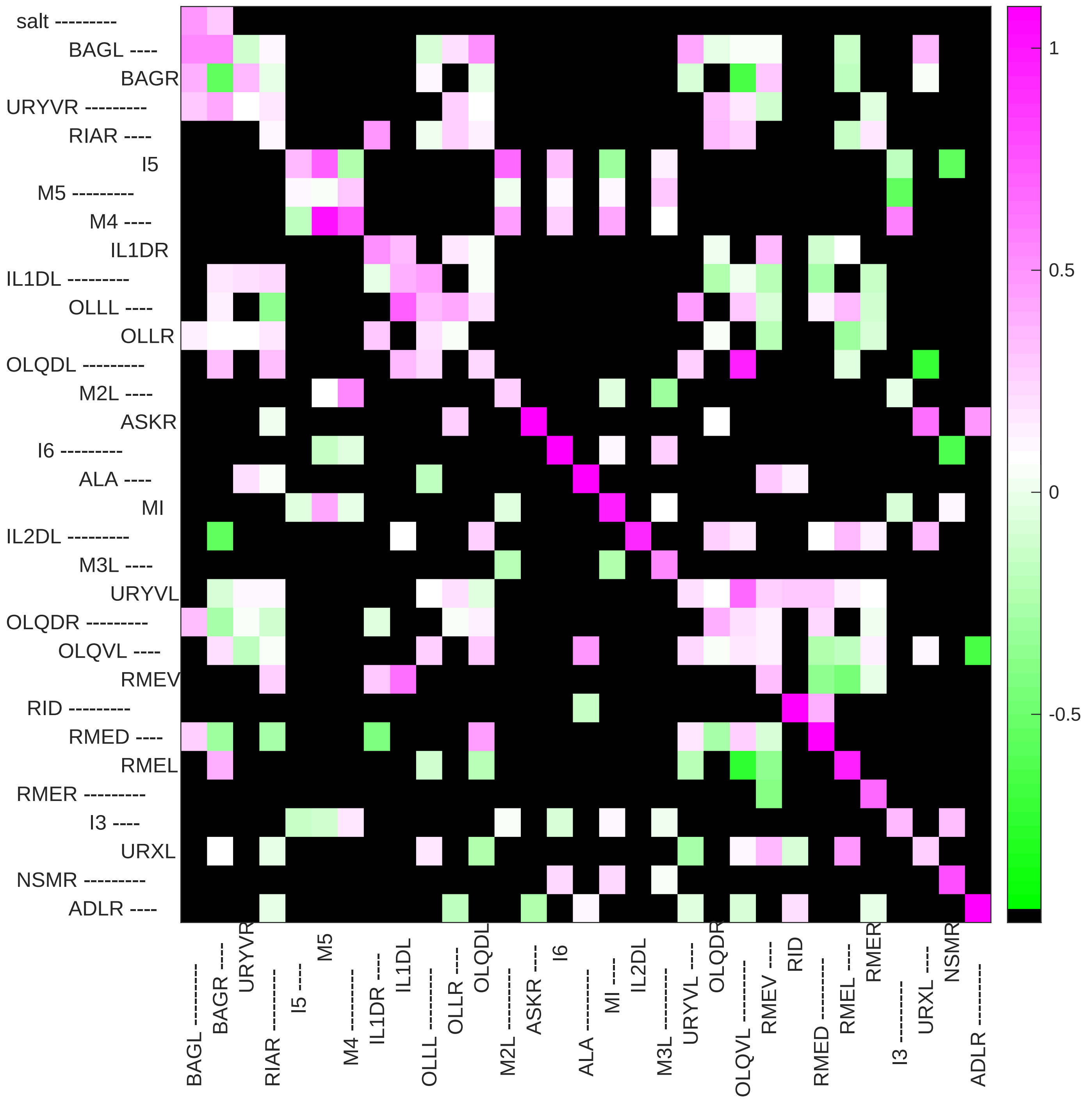
