## Supplementary File 11B for "Deducing ensemble dynamics and information flow from the whole-brain imaging data"

### Sample1 connection weight matrix

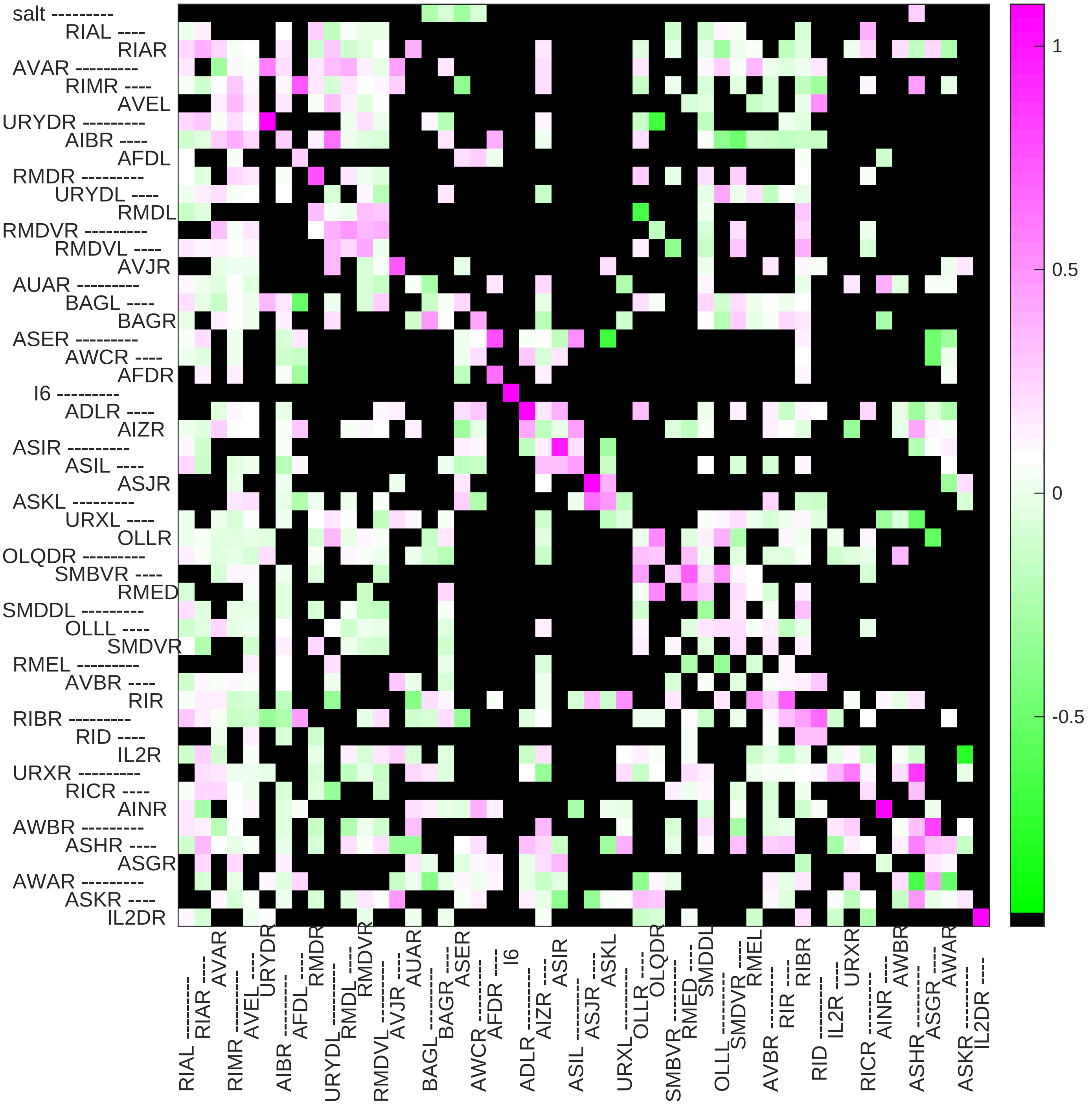

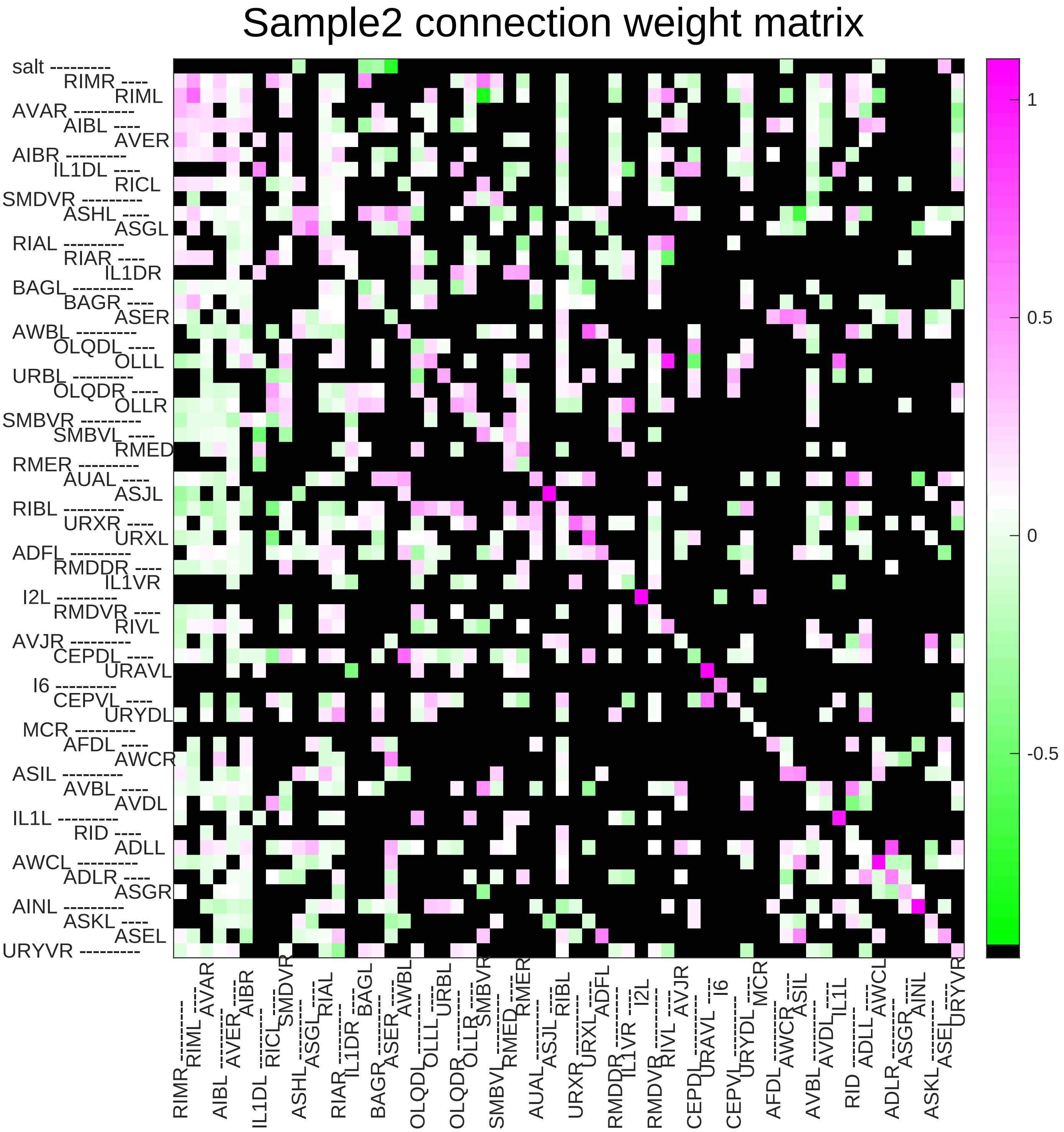
Sample2 connection weight matrix
salt
RIMR
RIML
AVAR
AIBL
AVER
AIBR
IL1DL
RICL
SMDVR
ASHL
ASGL
RIAL
RIAR
IL1DR
BAGL
BAGR
ASER
AWBL
OLQDL
OLLL
URBL
OLQDR
OLLR
SMBVR
SMBVL
RMED
RMER
AUAL
ASJL
RIBL
URXR
URXL
ADFL
RMDDR
IL1VR
I2L
RMDVR
RIVL
AVJR
CEPDL
URAVL
I6
CEPVL
URYDL
MCR
AFDL
AWCR
ASIL
AVBL
AVDL
IL1L
RID
ADLL
AWCL
ADLR
ASGR
AINL
ASKL
ASEL
URYVR
1
0.5
0
-0.5

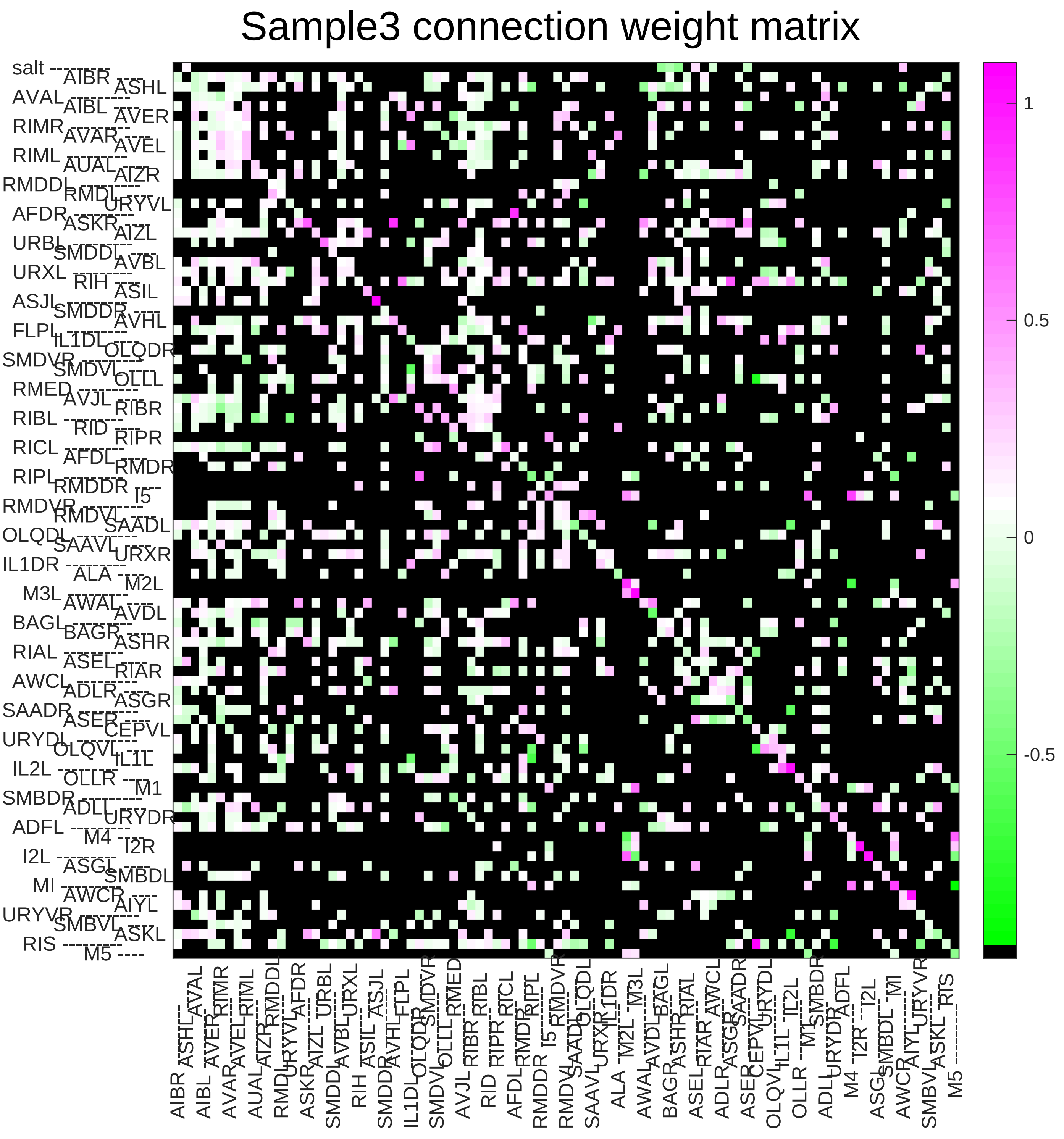
Sample3 connection weight matrix
salt
AIBR
ASHL
AVAL
AIBL
AVER
RIMR
AVAR
AVEL
RIML
AUAL
AIZR
RMDDL
RMDL
URYVL
AFDR
ASKR
AIZL
URBL
SMDDL
AVBL
URXL
RIH
ASIL
ASJL
SMDDR
AVHL
FLPL
IL1DL
OLQDR
SMDVR
SMDVL
OLLL
RMED
AVJL
RIBR
RIBL
RID
RIPR
RICL
AFDL
RMDR
RIPL
RMDDR
I5
RMDVR
RMDVL
SAADL
OLQDL
SAAVL
URXR
IL1DR
ALA
M2L
M3L
AWAL
AVDL
BAGL
BAGR
ASHR
RIAL
ASEL
RIAR
AWCL
ADLR
ASGR
SAADR
ASER
CEPVL
URYDL
OLQVL
IL1L
IL2L
OLLR
M1
SMBDR
ADLL
URYDR
ADFL
M4
I2R
I2L
ASGL
SMBDL
MI
AWCR
AIYL
URYVR
SMBVL
ASKL
RIS
M5
1
0.5
0
-0.5

### Sample4 connection weight matrix

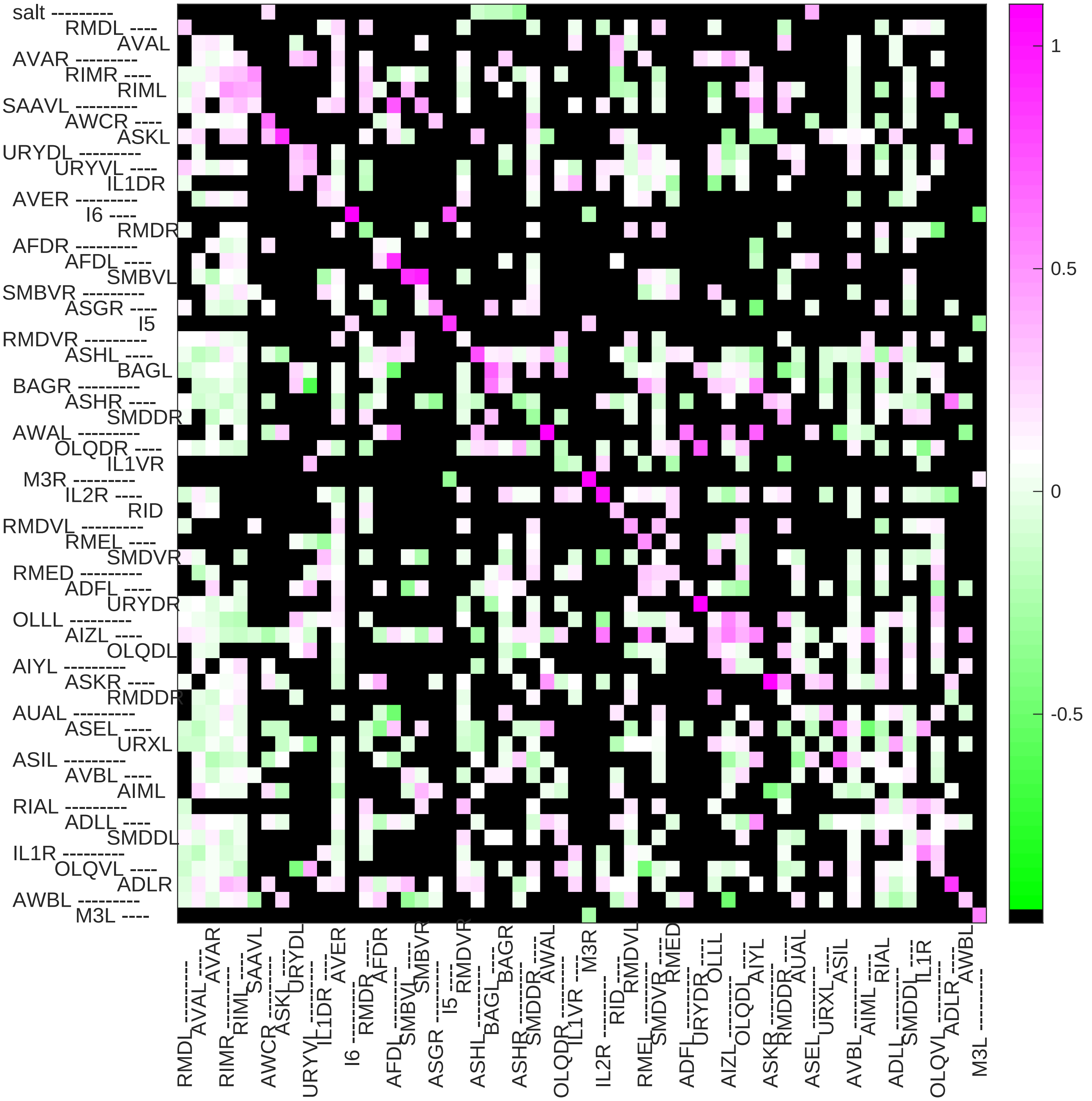

#### Sample5 connection weight matrix

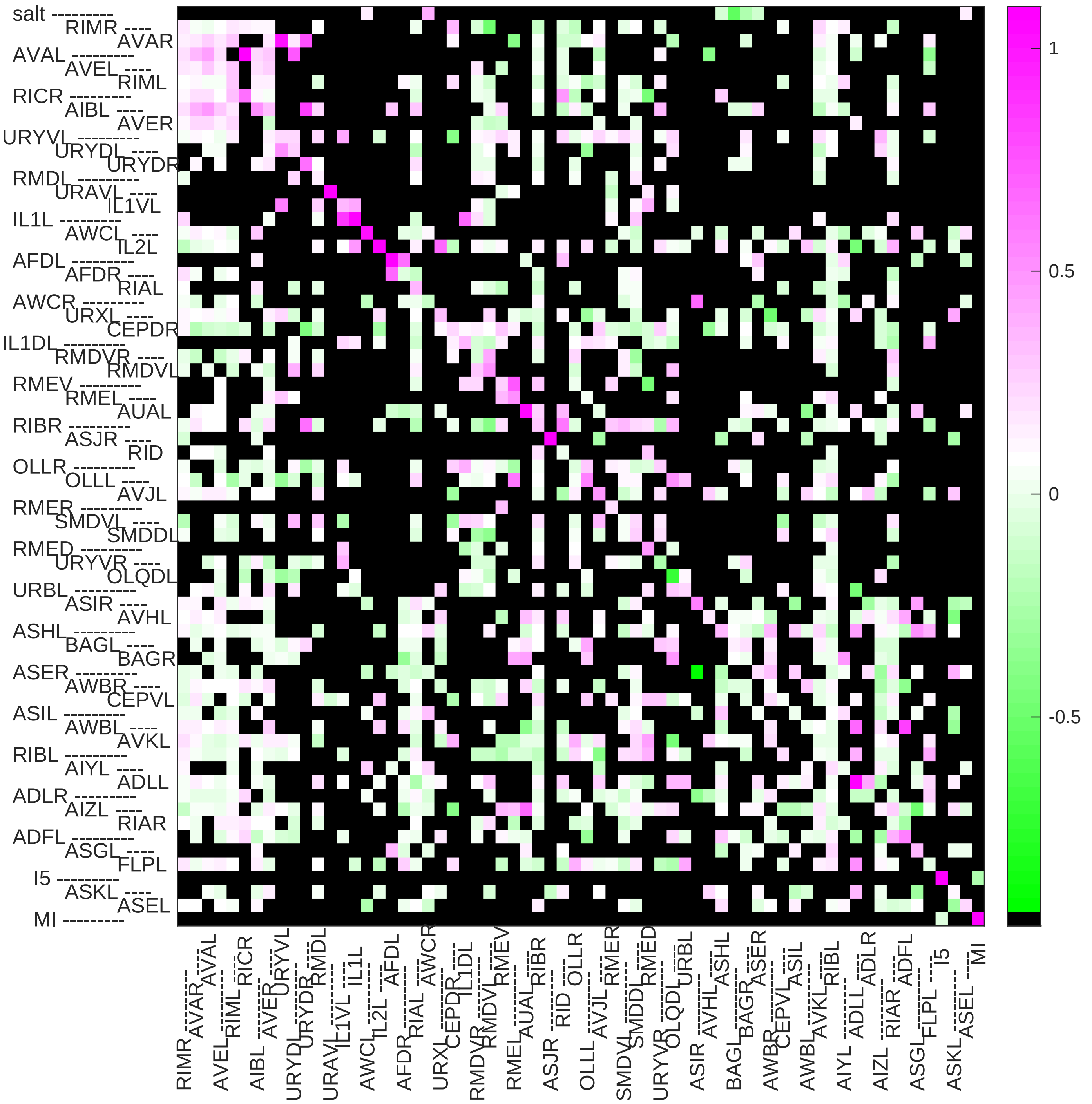

Sample6 connection weight matrix

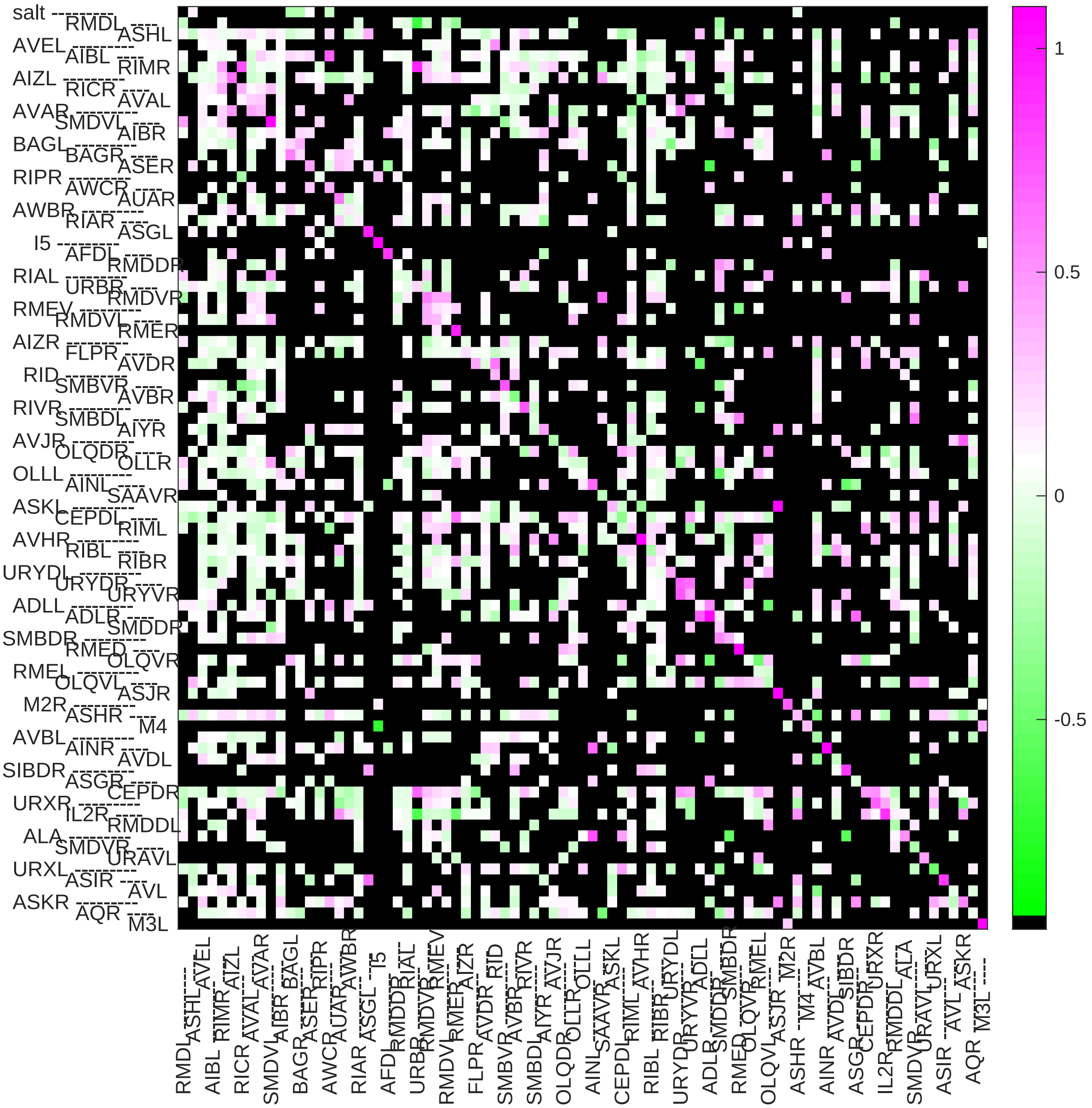

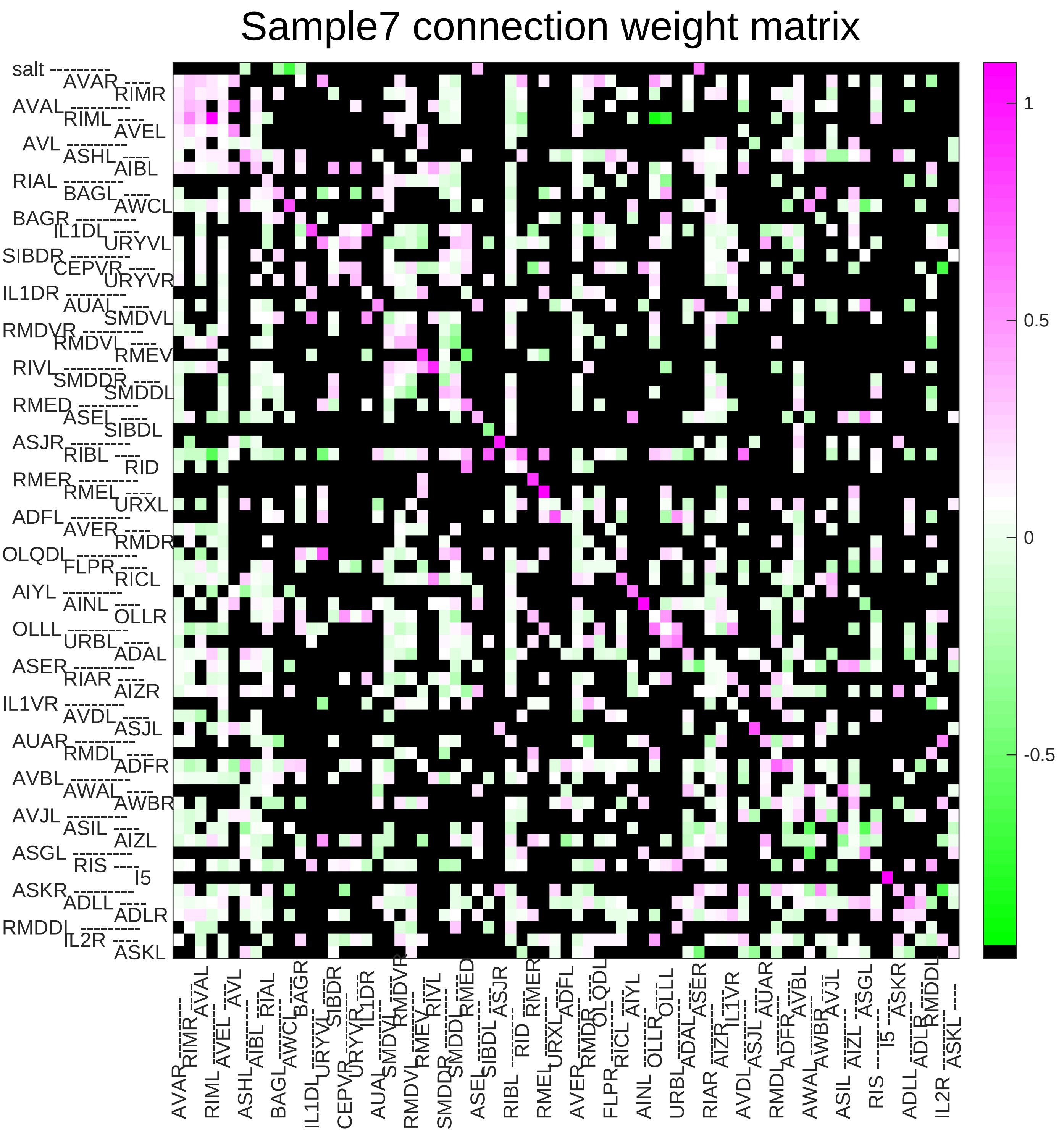
Sample7 connection weight matrix
salt
AVAR
RIMR
AVAL
RIML
AVEL
AVL
ASHL
AIBL
RIAL
BAGL
AWCL
BAGR
IL1DL
URYVL
SIBDR
CEPVR
URYVR
IL1DR
AUAL
SMDVL
RMDVR
RMDVL
RMEV
RIVL
SMDDR
SMDDL
RMED
ASEL
SIBDL
ASJR
RIBL
RID
RMER
RMEL
URXL
ADFL
AVER
RMDR
OLQDL
FLPR
RICL
AIYL
AINL
OLLR
OLLL
URBL
ADAL
ASER
RIAR
AIZR
IL1VR
AVDL
ASJL
AUAR
RMDL
ADFR
AVBL
AWAL
AWBR
AVJL
ASIL
AIZL
ASGL
RIS
I5
ASKR
ADLL
ADLR
RMDDL
IL2R
ASKL
1
0.5
0
-0.5

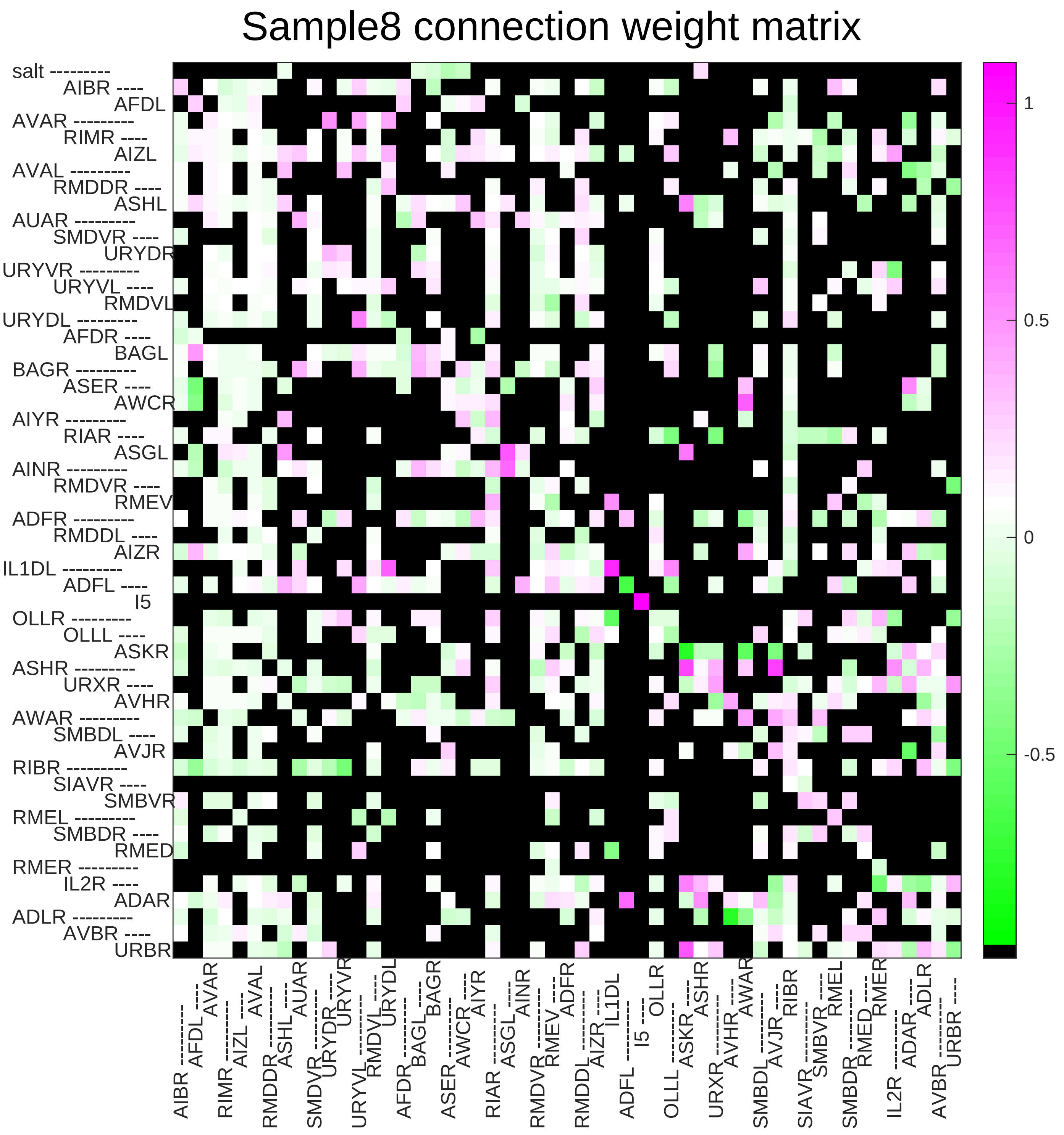
Sample8 connection weight matrix
salt
AIBR
AFDL
AVAR
RIMR
AIZL
AVAL
RMDDR
ASHL
AUAR
SMDVR
URYDR
URYVR
URYVL
RMDVL
URYDL
AFDR
BAGL
BAGR
ASER
AWCR
AIYR
RIAR
ASGL
AINR
RMDVR
RMEV
ADFR
RMDDL
AIZR
IL1DL
ADFL
I5
OLLR
OLLL
ASKR
ASHR
URXR
AVHR
AWAR
SMBDL
AVJR
RIBR
SIAVR
SMBVR
RMEL
SMBDR
RMED
RMER
IL2R
ADAR
ADLR
AVBR
URBR
1
0.5
0
-0.5

### Sample9 connection weight matrix

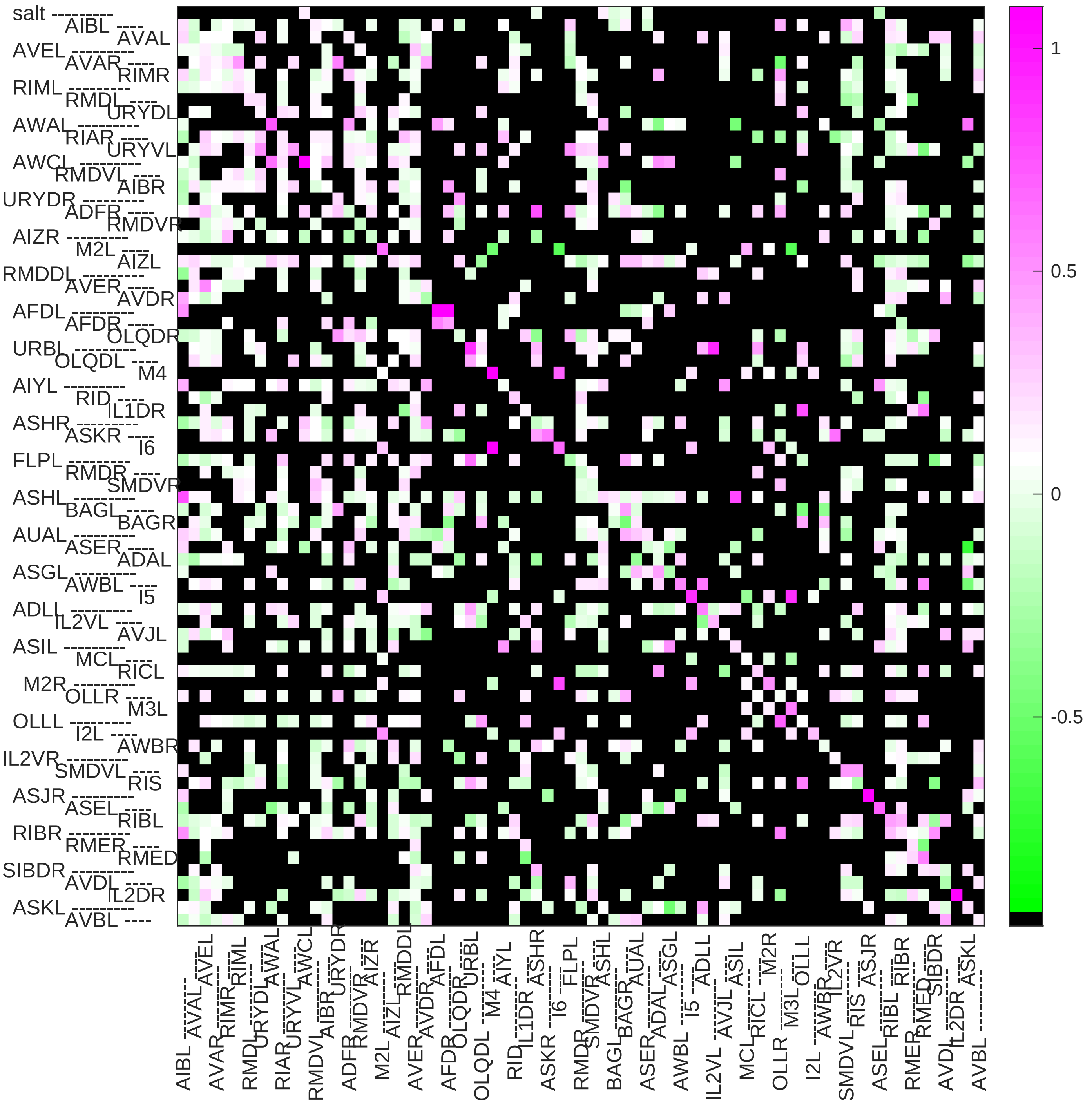

### Sample10 connection weight matrix

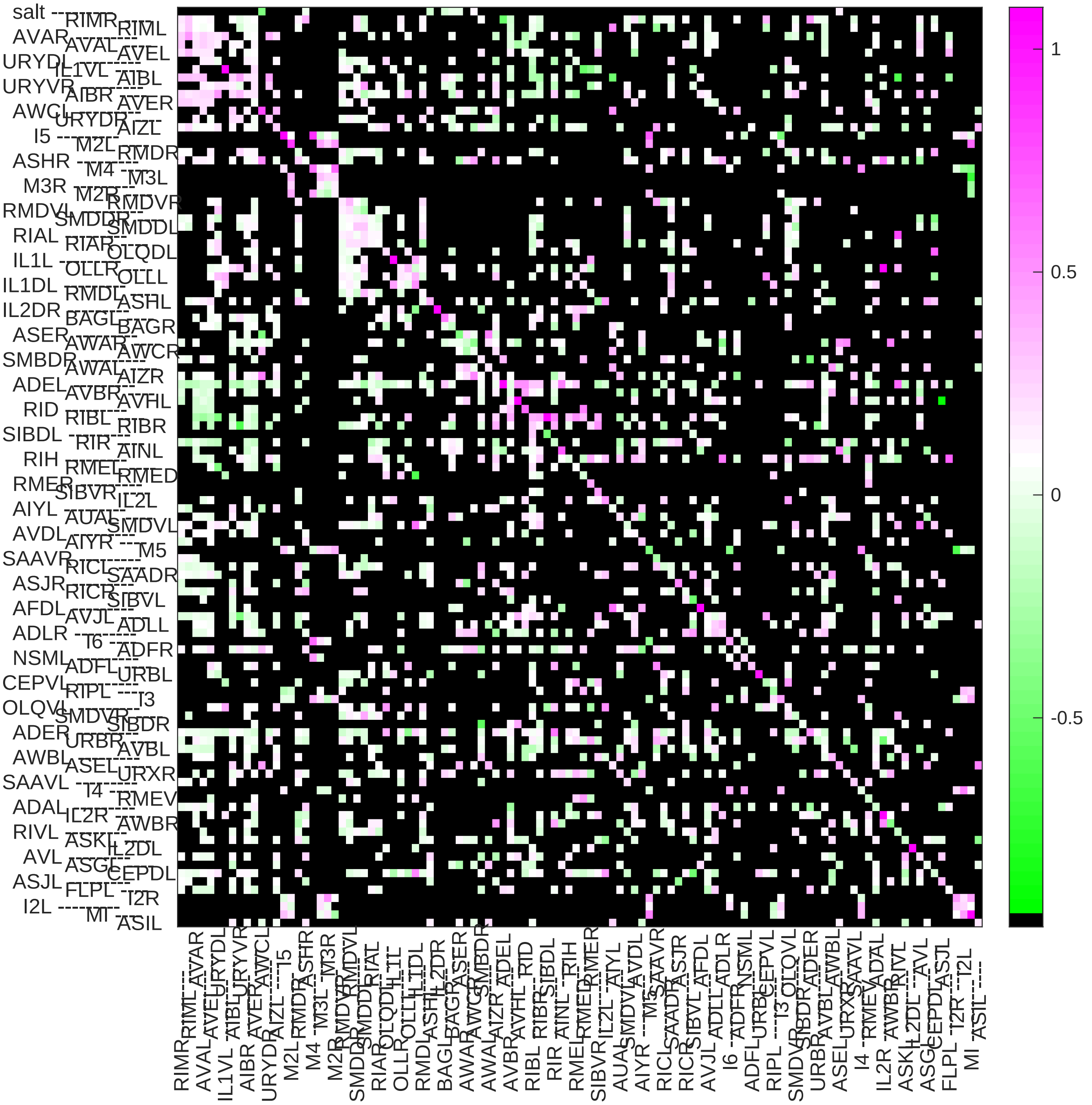

Sample11 connection weight matrix

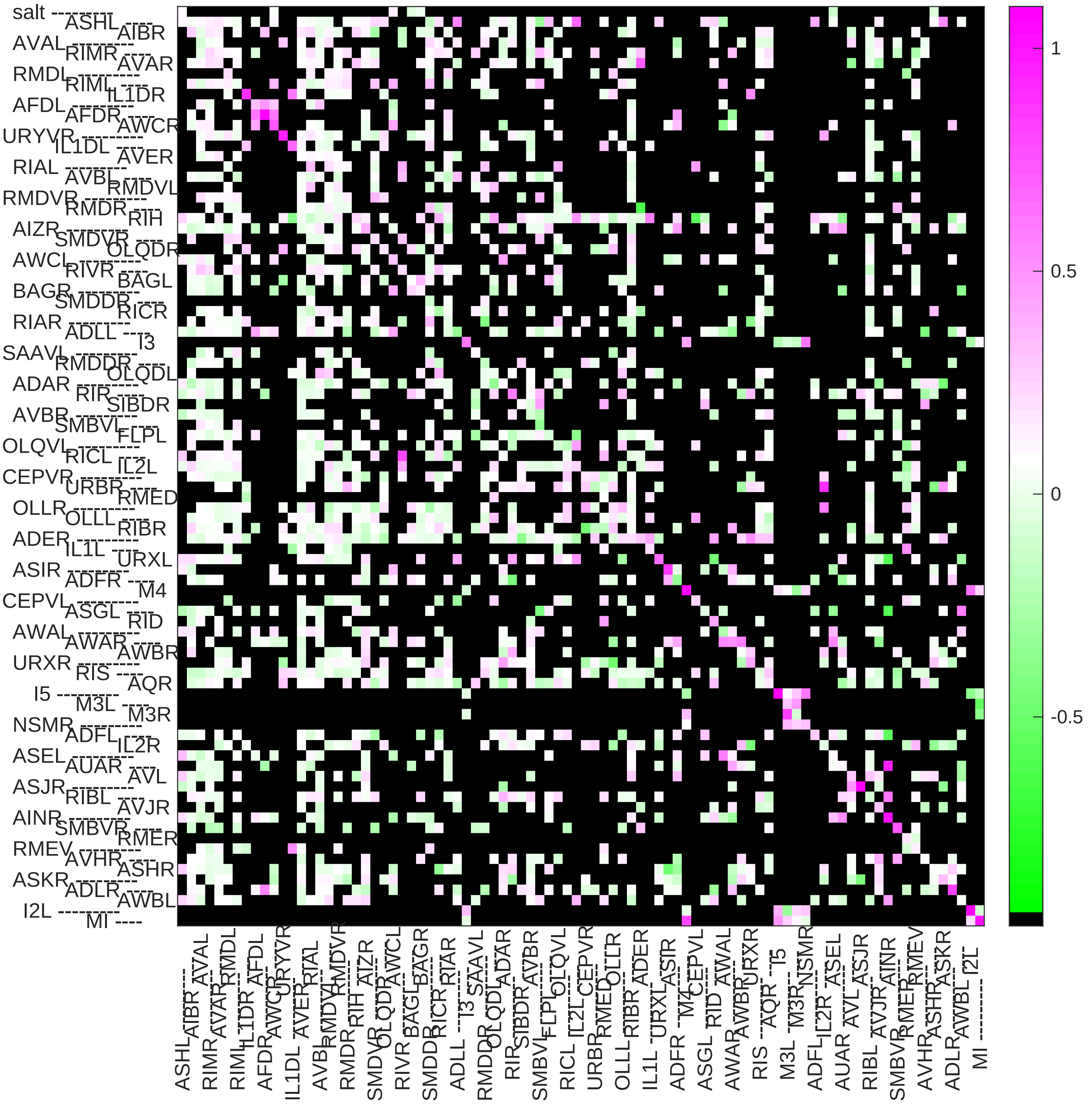

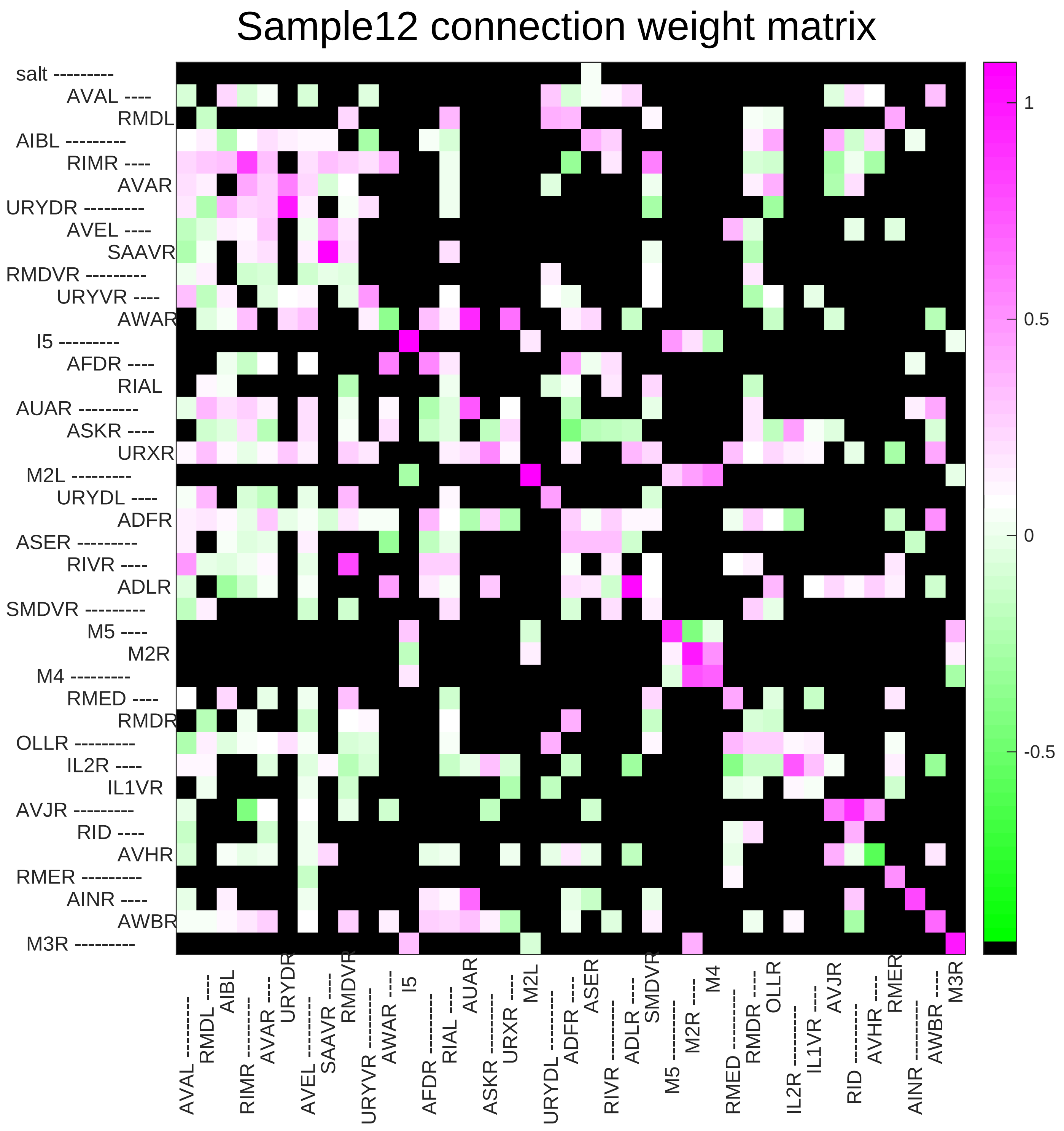
Sample12 connection weight matrix
salt
AVAL
RMDL
AIBL
RIMR
AVAR
URYDR
AVEL
SAAVR
RMDVR
URYVR
AWAR
I5
AFDR
RIAL
AUAR
ASKR
URXR
M2L
URYDL
ADFR
ASER
RIVR
ADLR
SMDVR
M5
M2R
M4
RMED
RMDR
OLLR
IL2R
IL1VR
AVJR
RID
AVHR
RMER
AINR
AWBR
M3R
1
0.5
0
-0.5

### Sample13 connection weight matrix

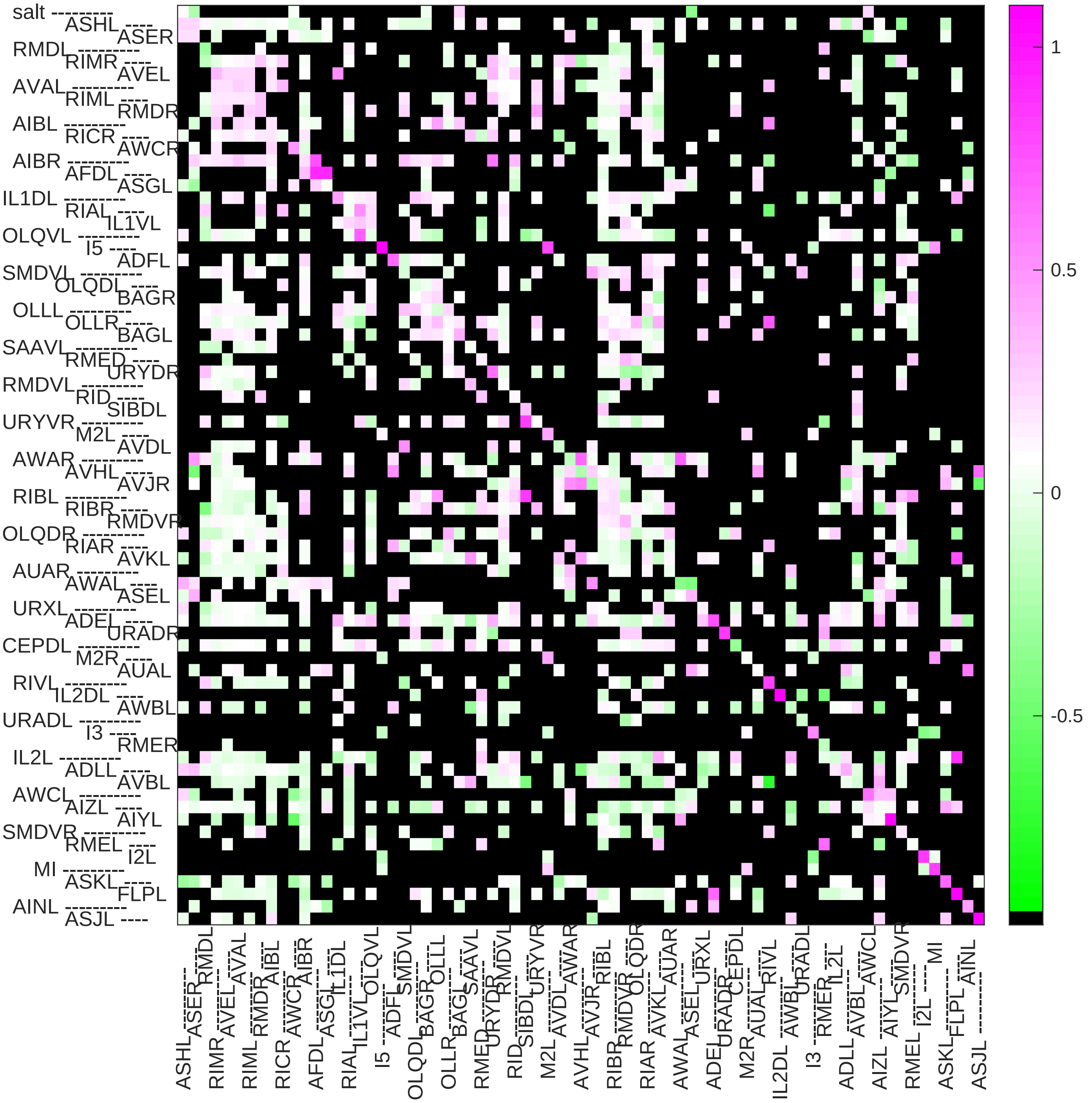

### Sample14 connection weight matrix

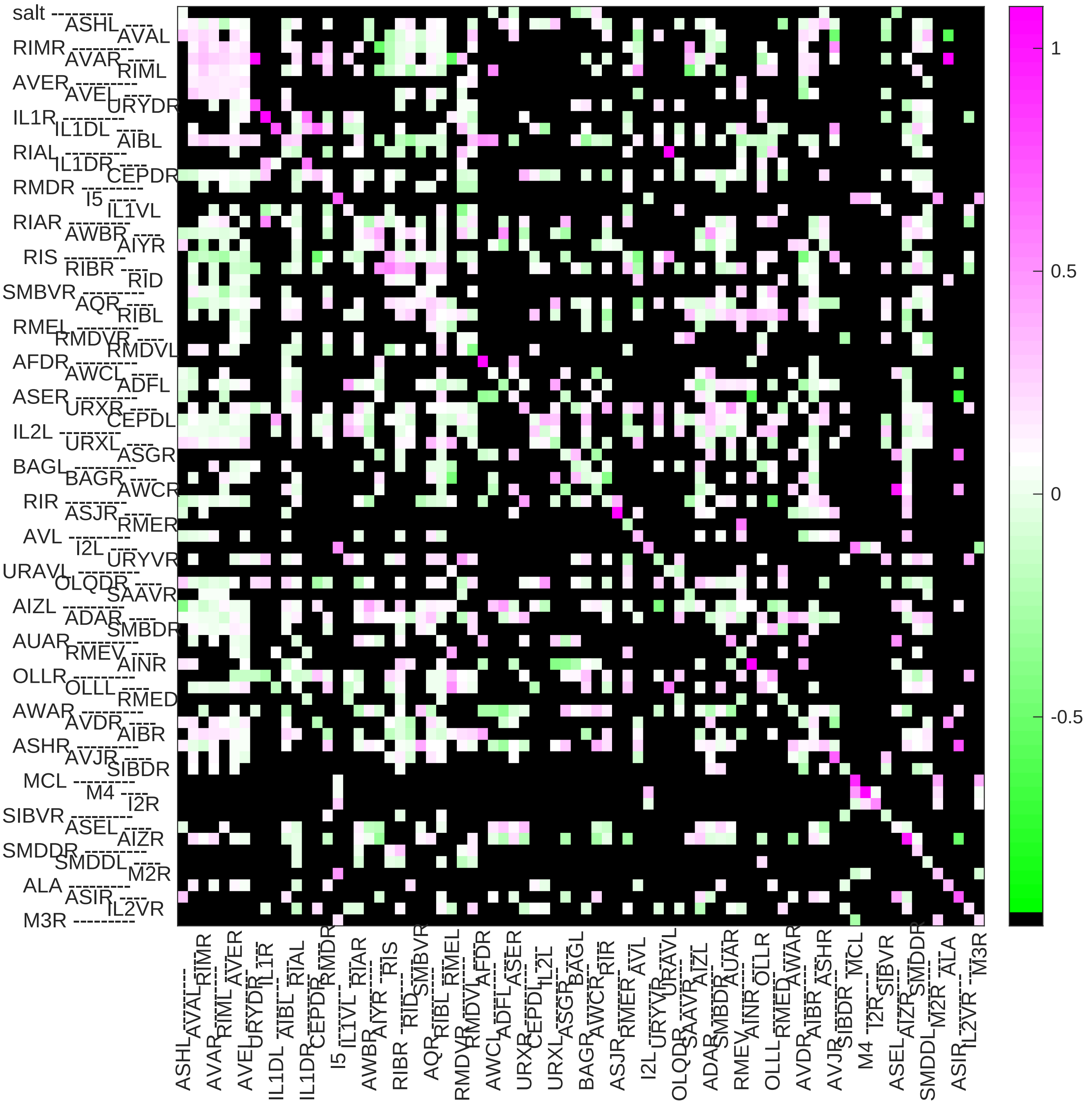

Sample15 connection weight matrix

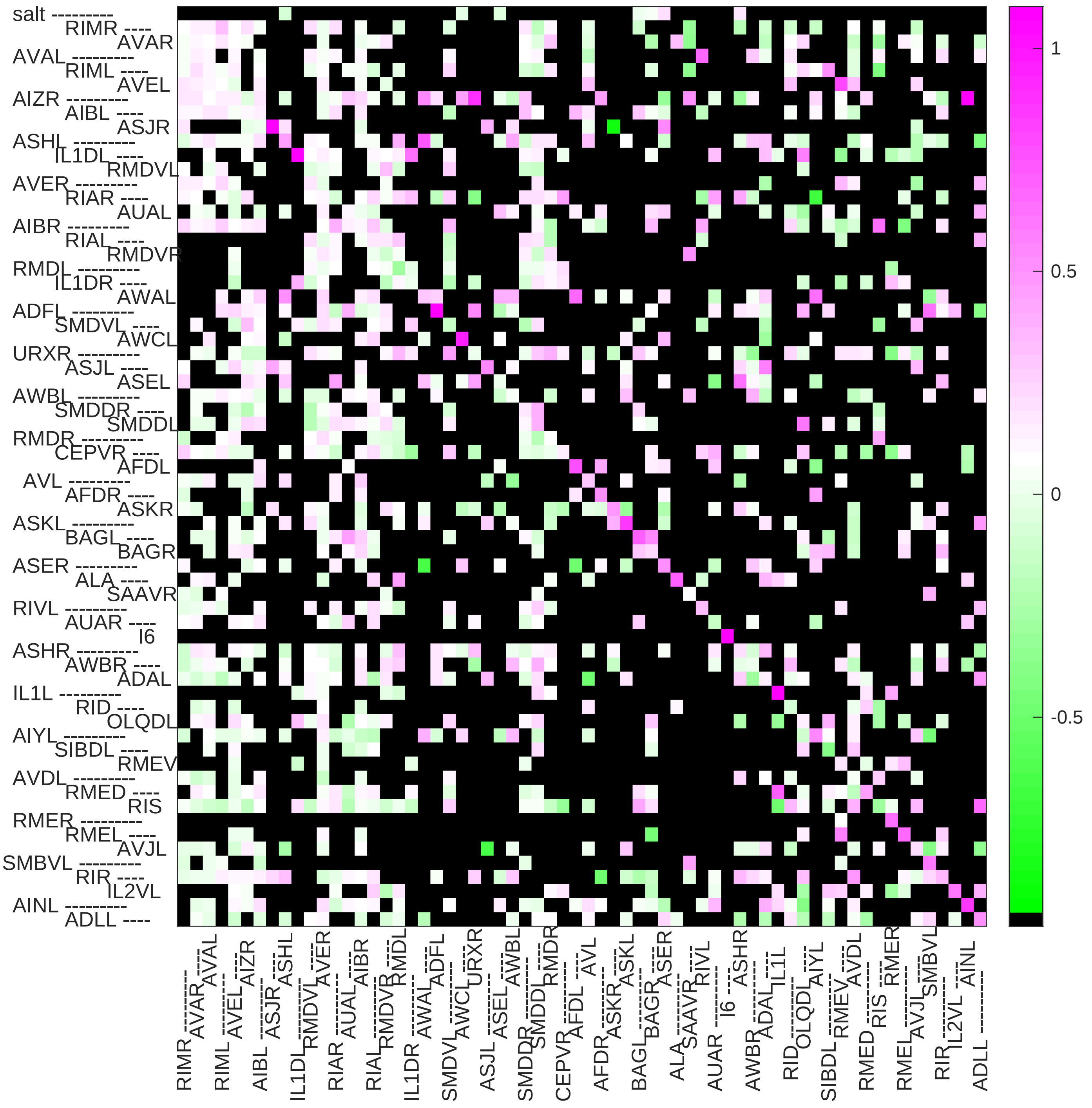

Sample16 connection weight matrix

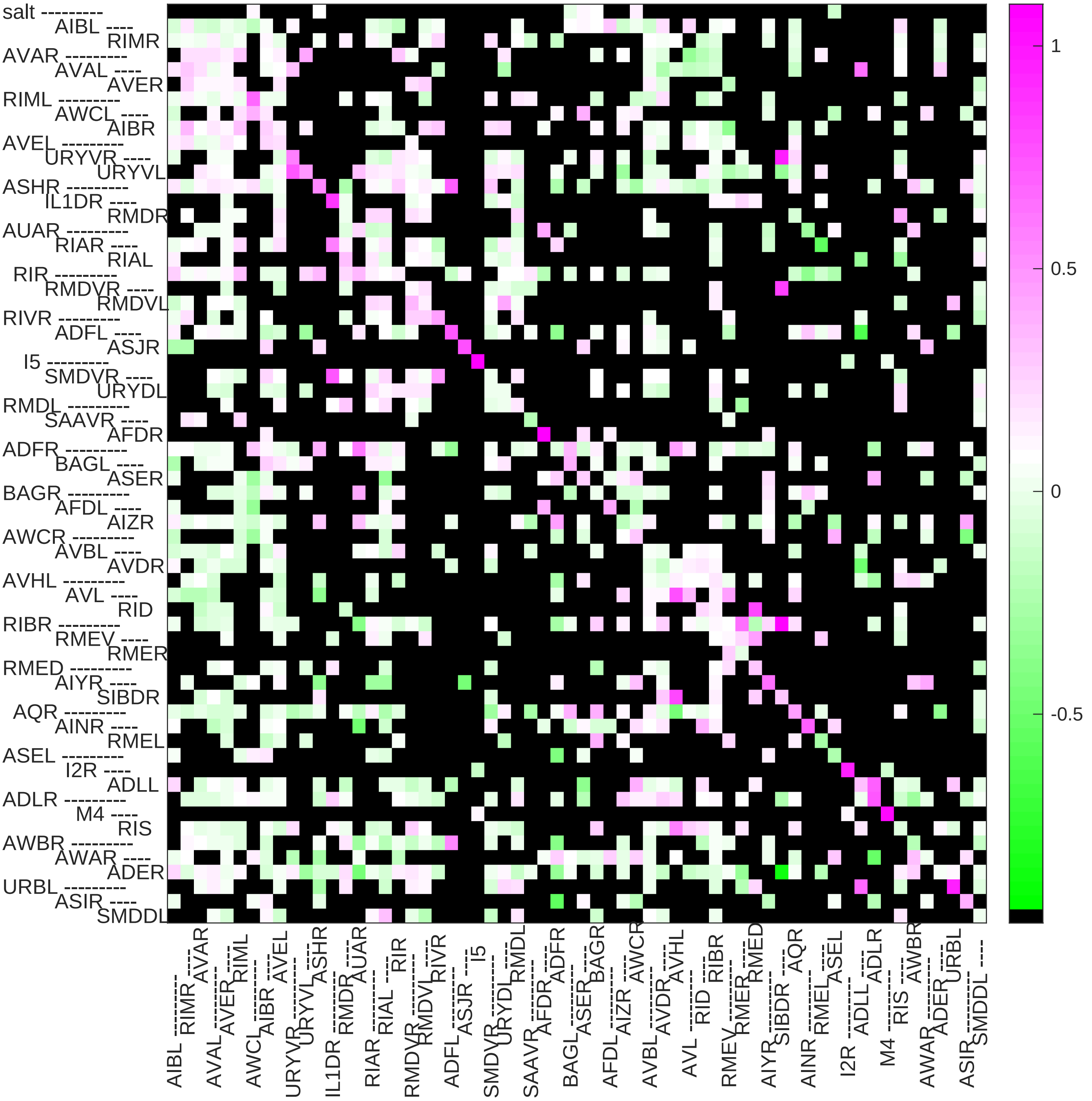

### Sample17 connection weight matrix

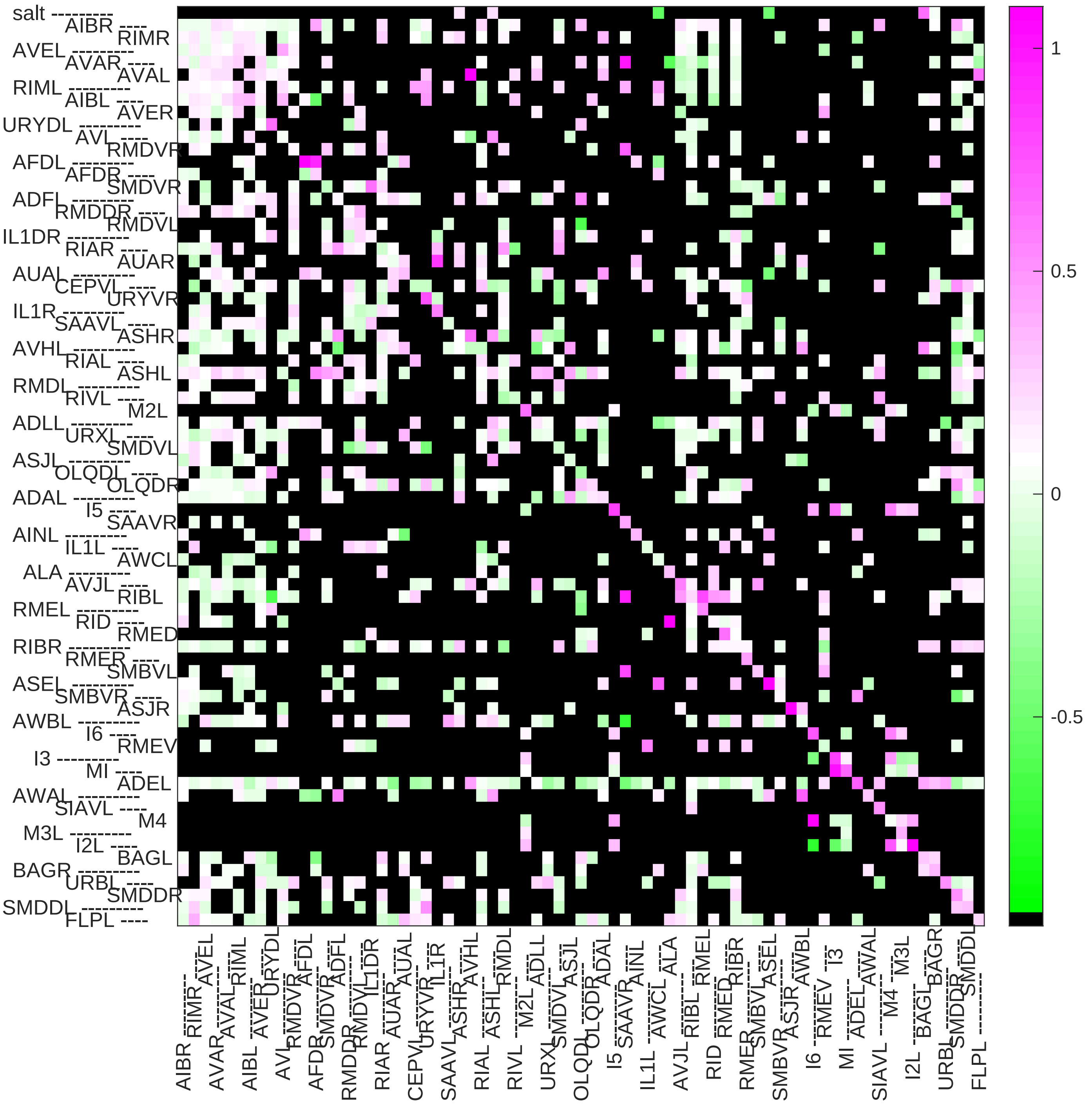

### Sample18 connection weight matrix

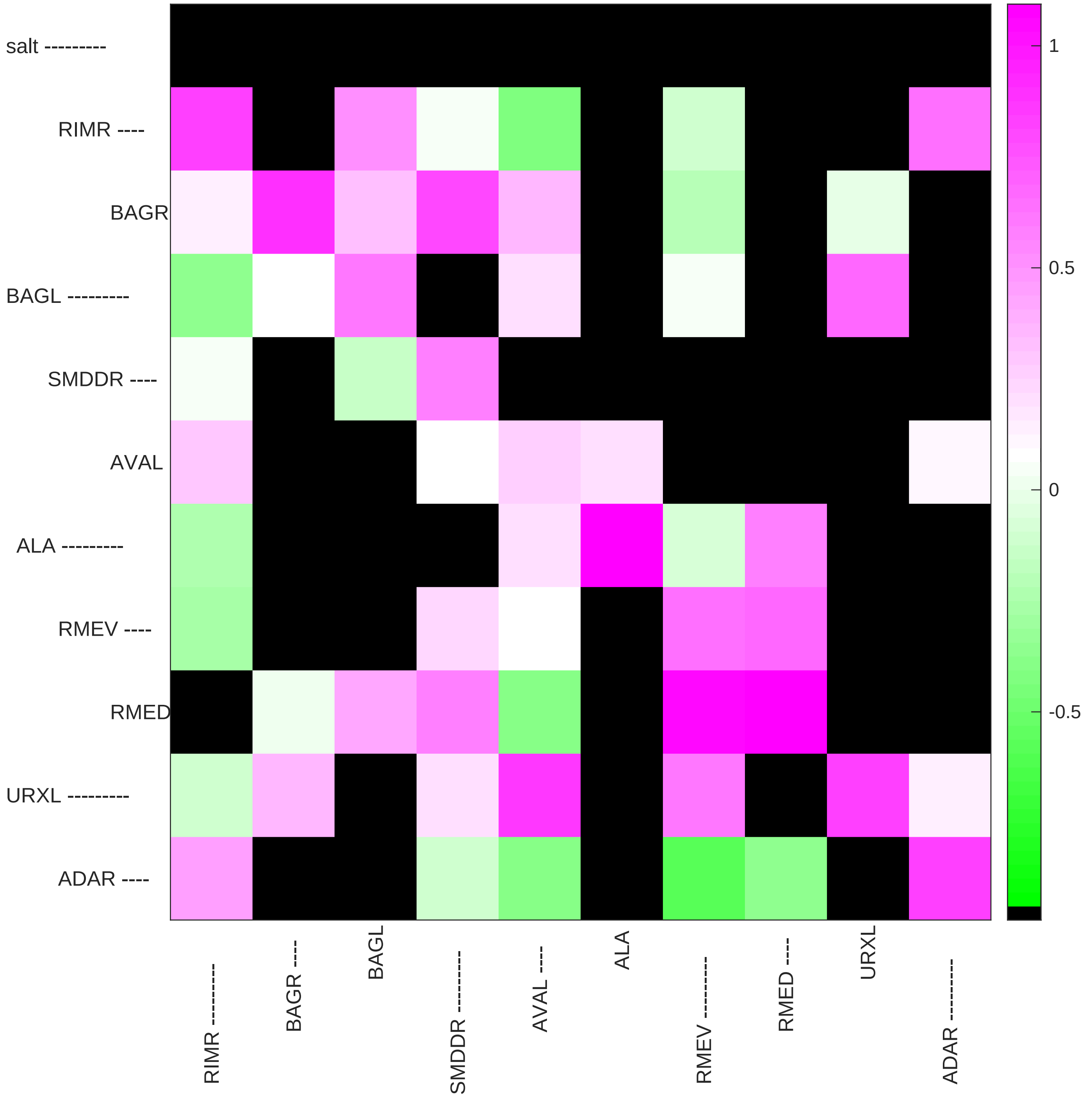

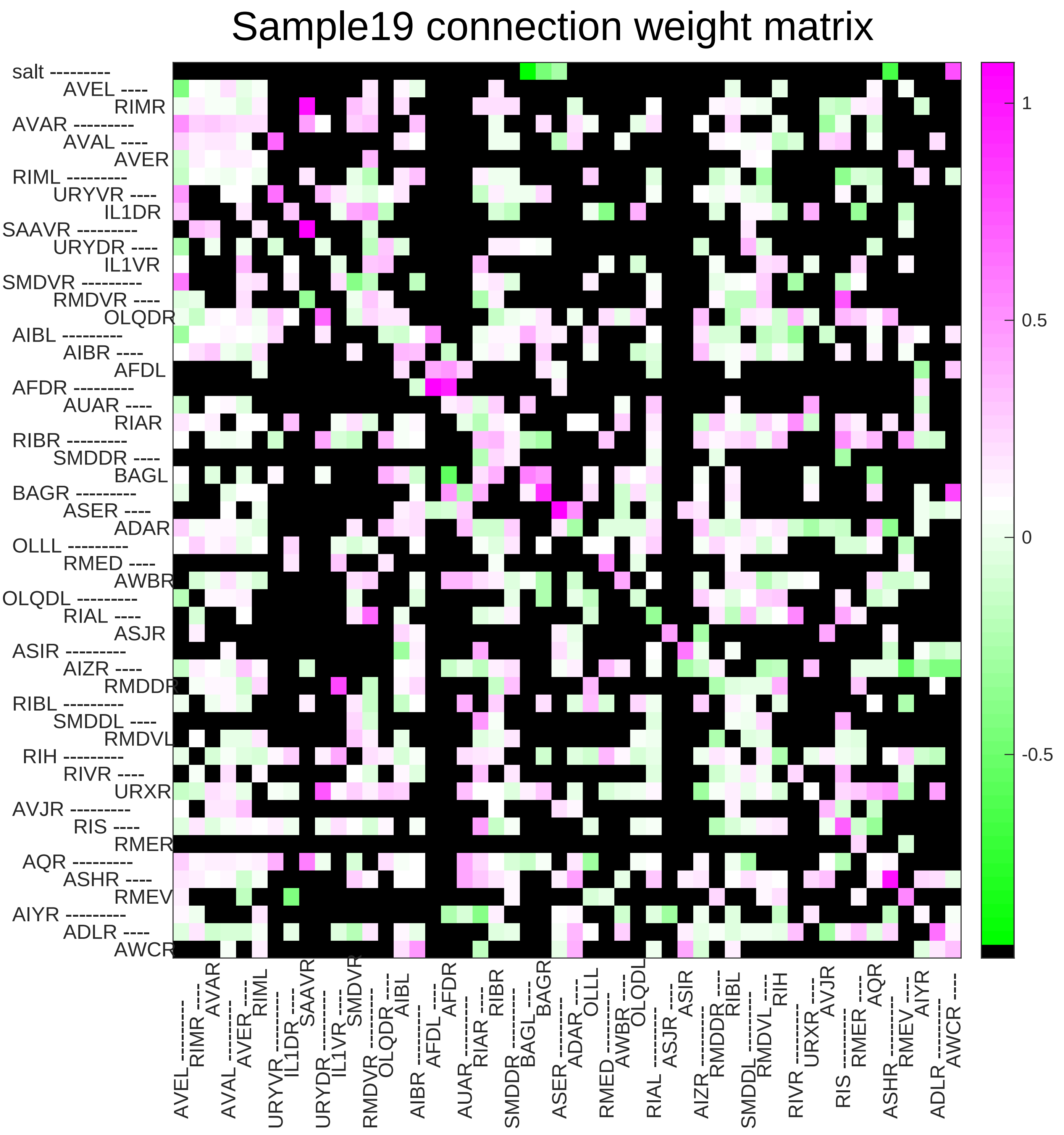

Sample19 connection weight matrix
salt
AVEL
RIMR
AVAR
AVAL
AVER
RIML
URYVR
IL1DR
SAAVR
URYDR
IL1VR
SMDVR
RMDVR
OLQDR
AIBL
AIBR
AFDL
AFDR
AUAR
RIAR
RIBR
SMDDR
BAGL
BAGR
ASER
ADAR
OLLL
RMED
AWBR
OLQDL
RIAL
ASJR
ASIR
AIZR
RMDDR
RIBL
SMDDL
RMDVL
RIH
RIVR
URXR
AVJR
RIS
RMER
AQR
ASHR
RMEV
AIYR
ADLR
AWCR
1
0.5
0
-0.5

### Sample20 connection weight matrix

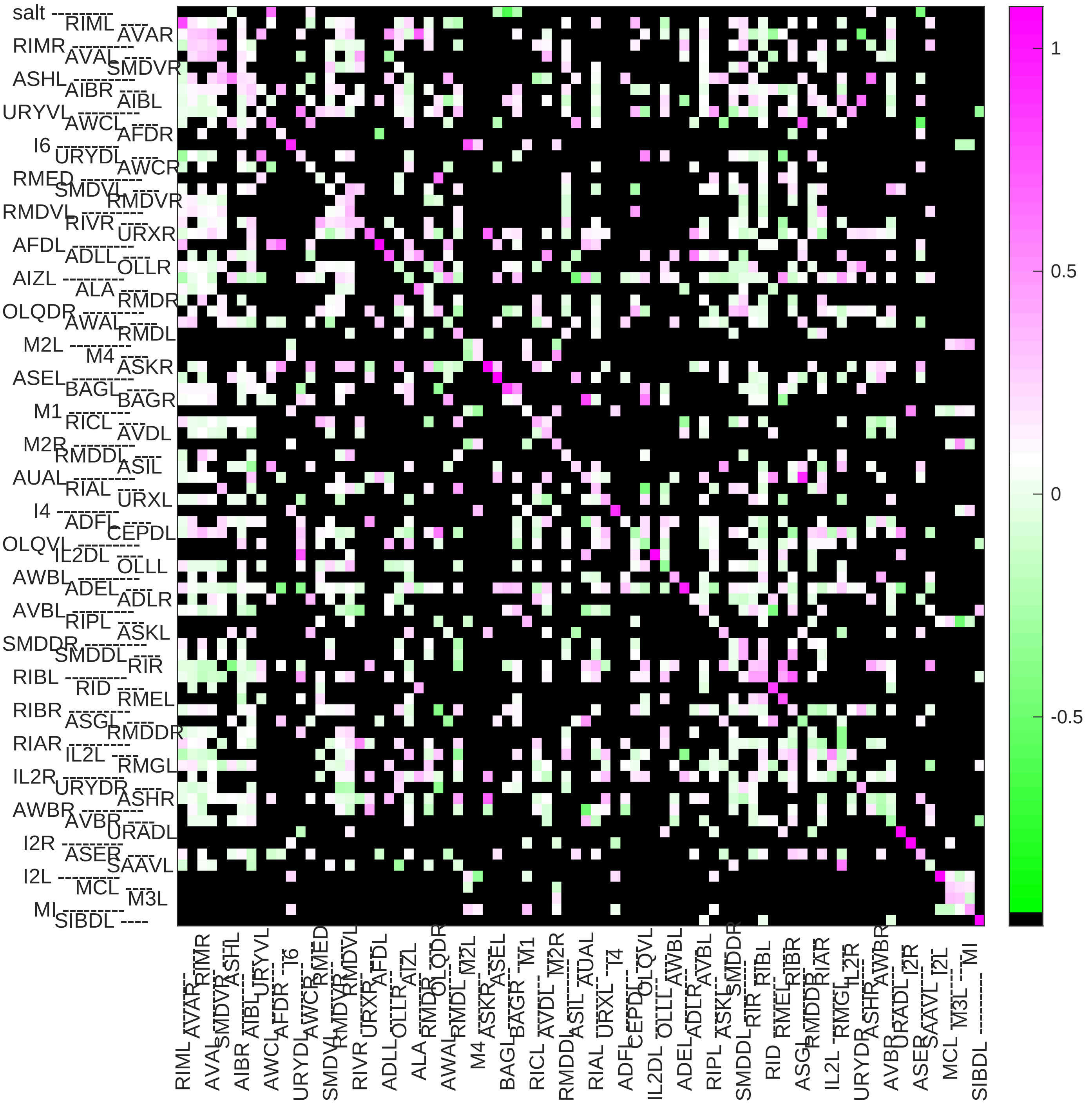

### Sample21 connection weight matrix

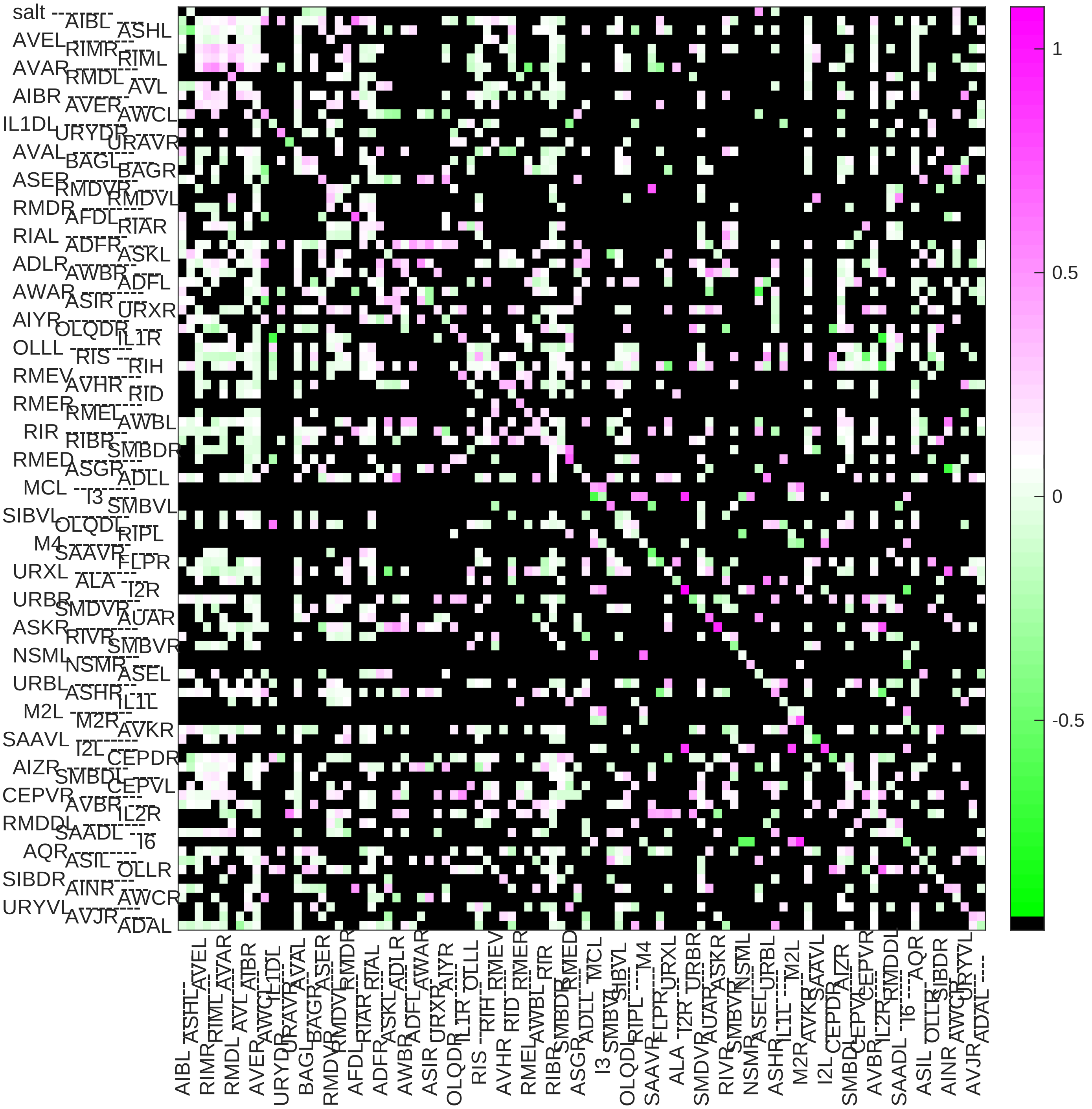

### Sample22 connection weight matrix

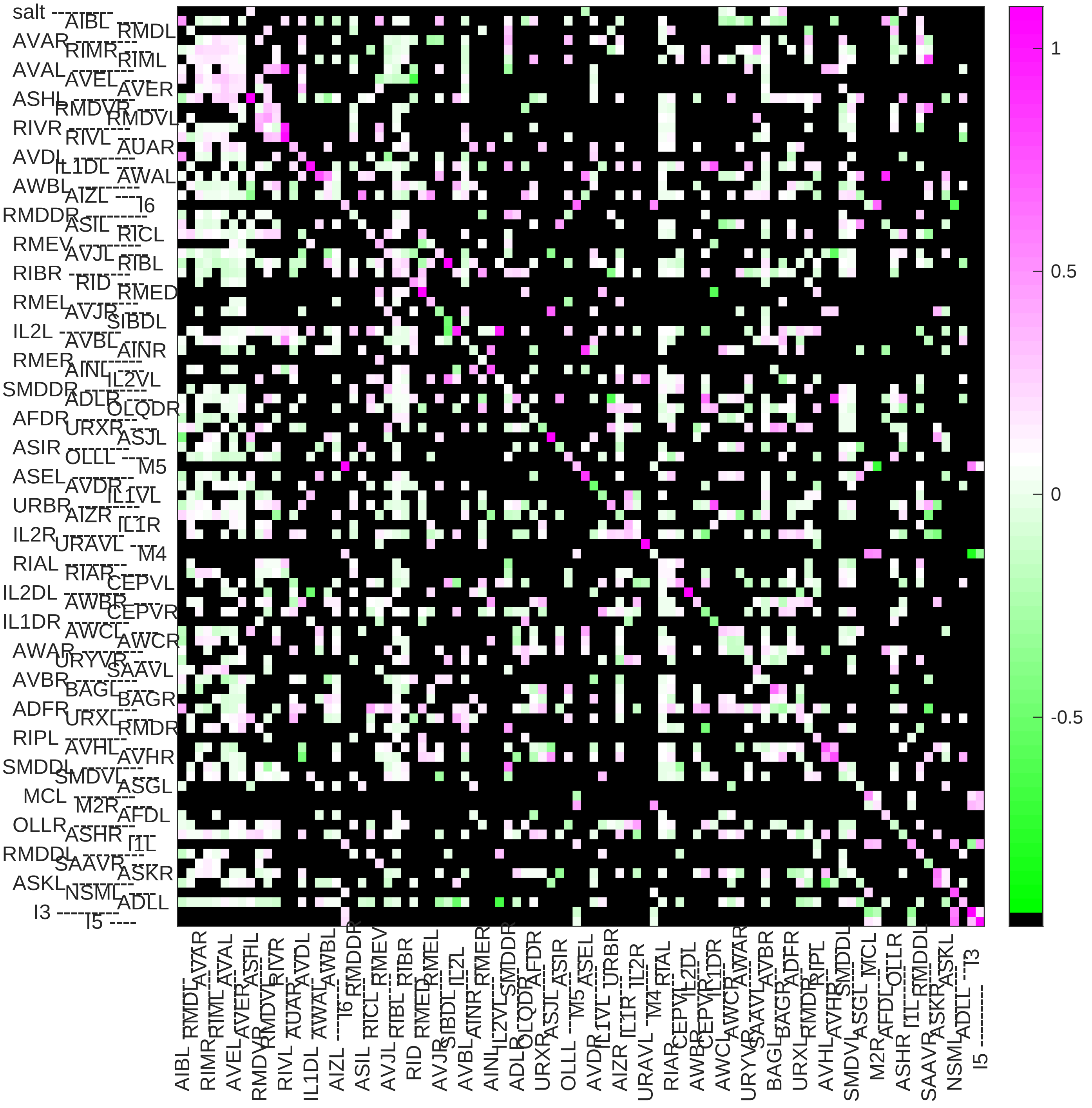

### Sample23 connection weight matrix

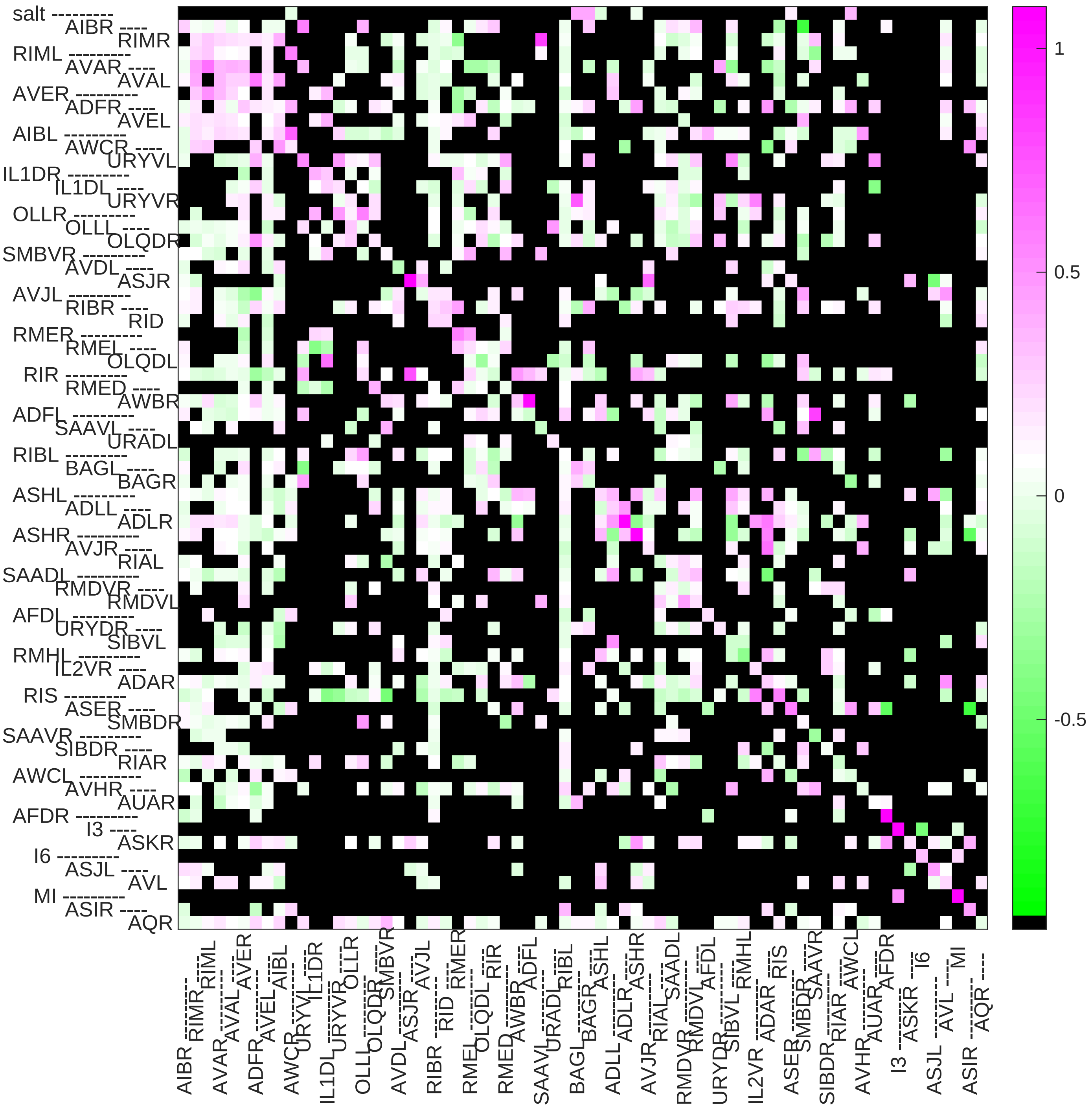

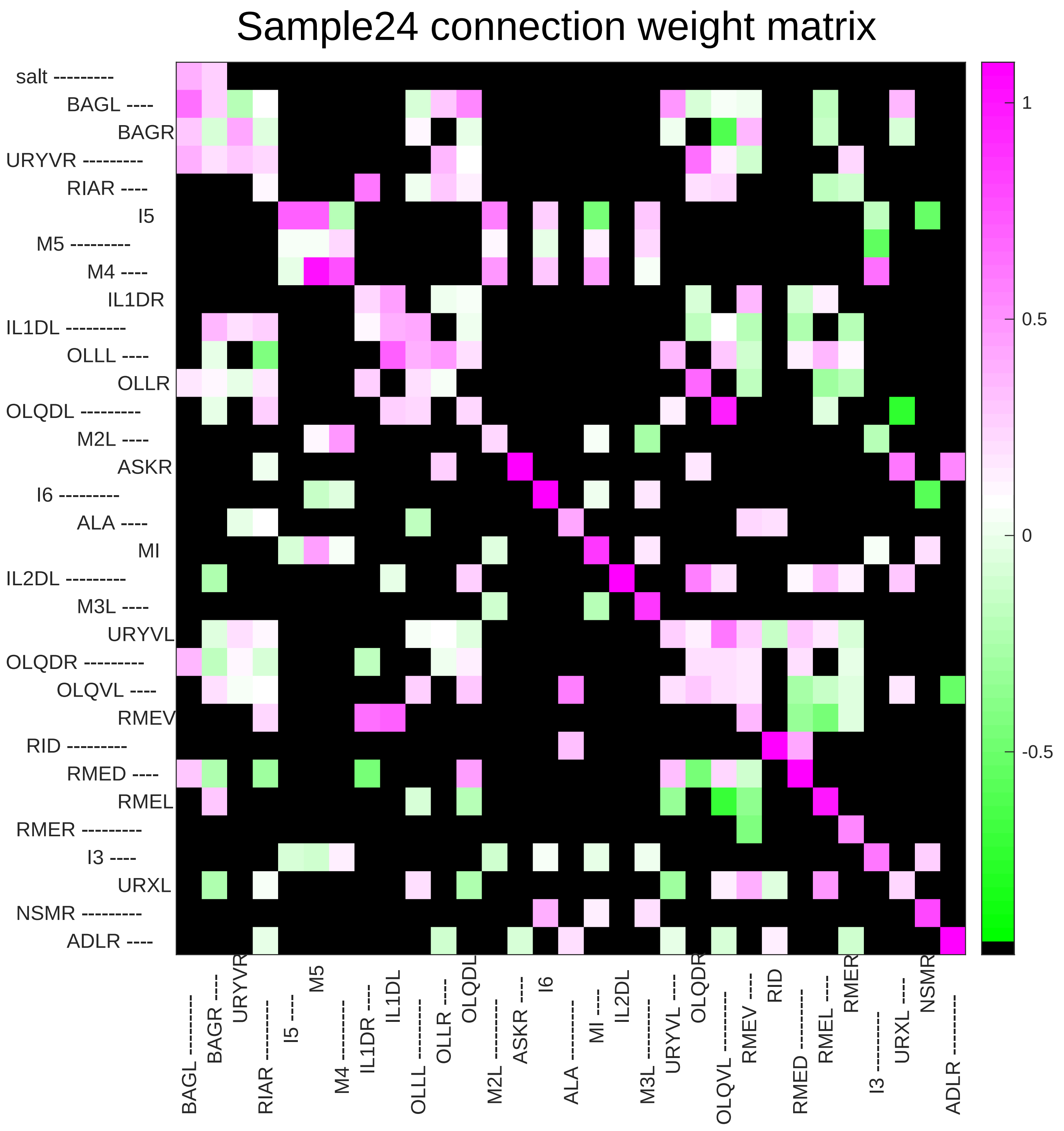
Sample24 connection weight matrix
salt
BAGL
BAGR
URYVR
RIAR
I5
M5
M4
IL1DR
IL1DL
OLLL
OLLR
OLQDL
M2L
ASKR
I6
ALA
MI
IL2DL
M3L
URYVL
OLQDR
OLQVL
RMEV
RID
RMED
RMEL
RMER
I3
URXL
NSMR
ADLR
1
0.5
0
-0.5
