## Supplementary File 11C for "Deducing ensemble dynamics and information flow from the whole-brain imaging data"

Sample1 connection weight matrix

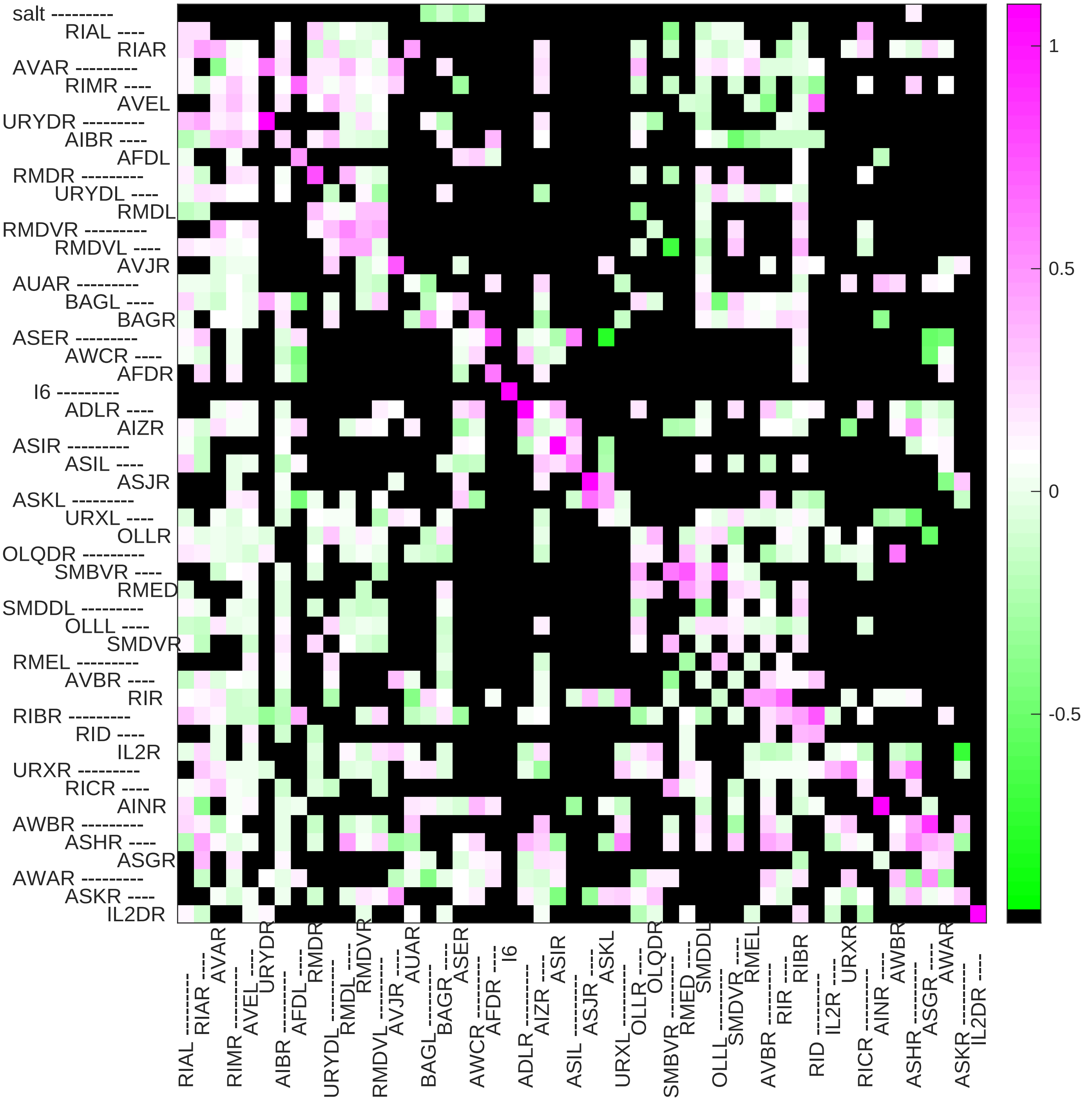

### Sample2 connection weight matrix

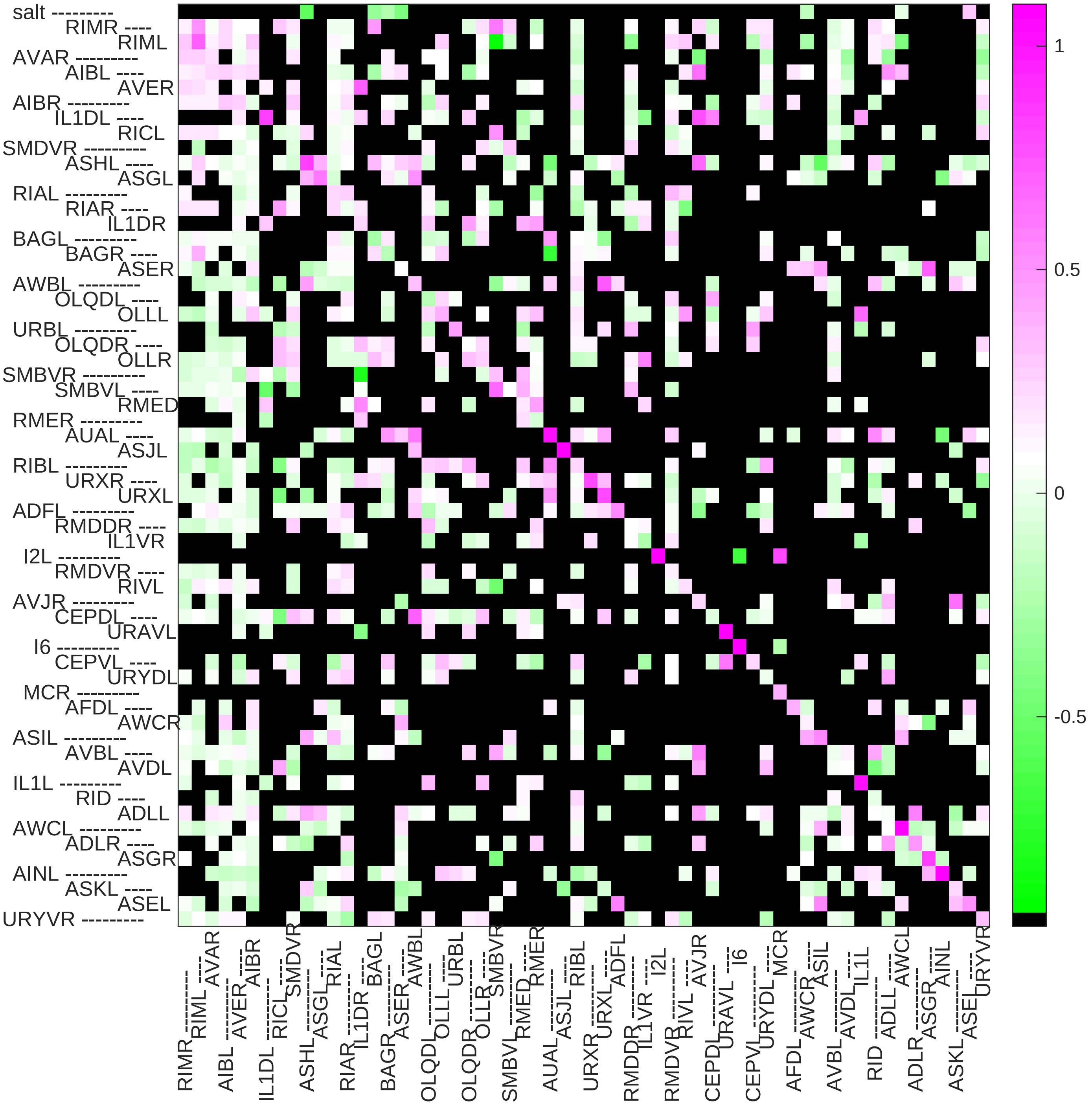

### Sample3 connection weight matrix

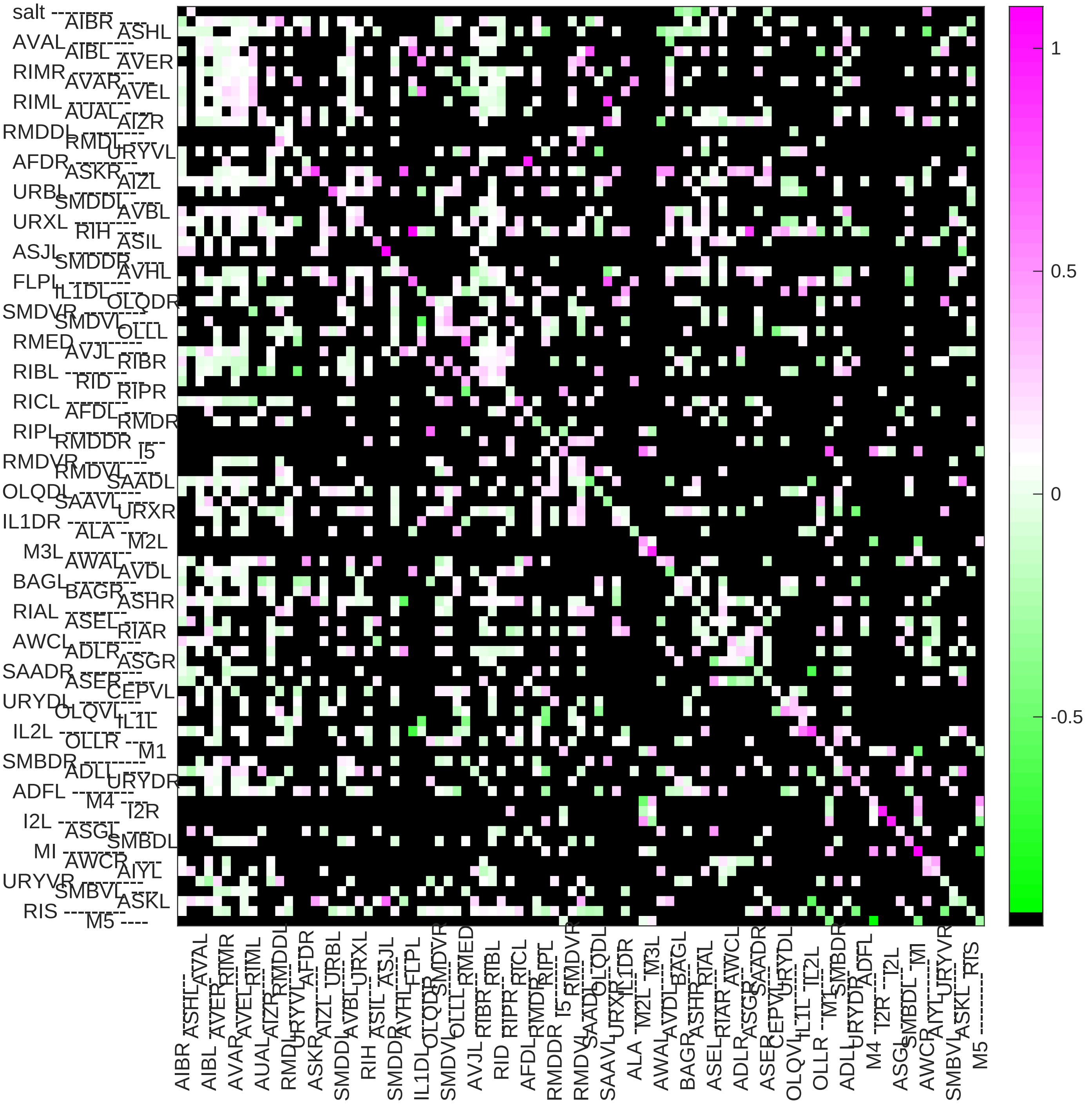

### Sample4 connection weight matrix

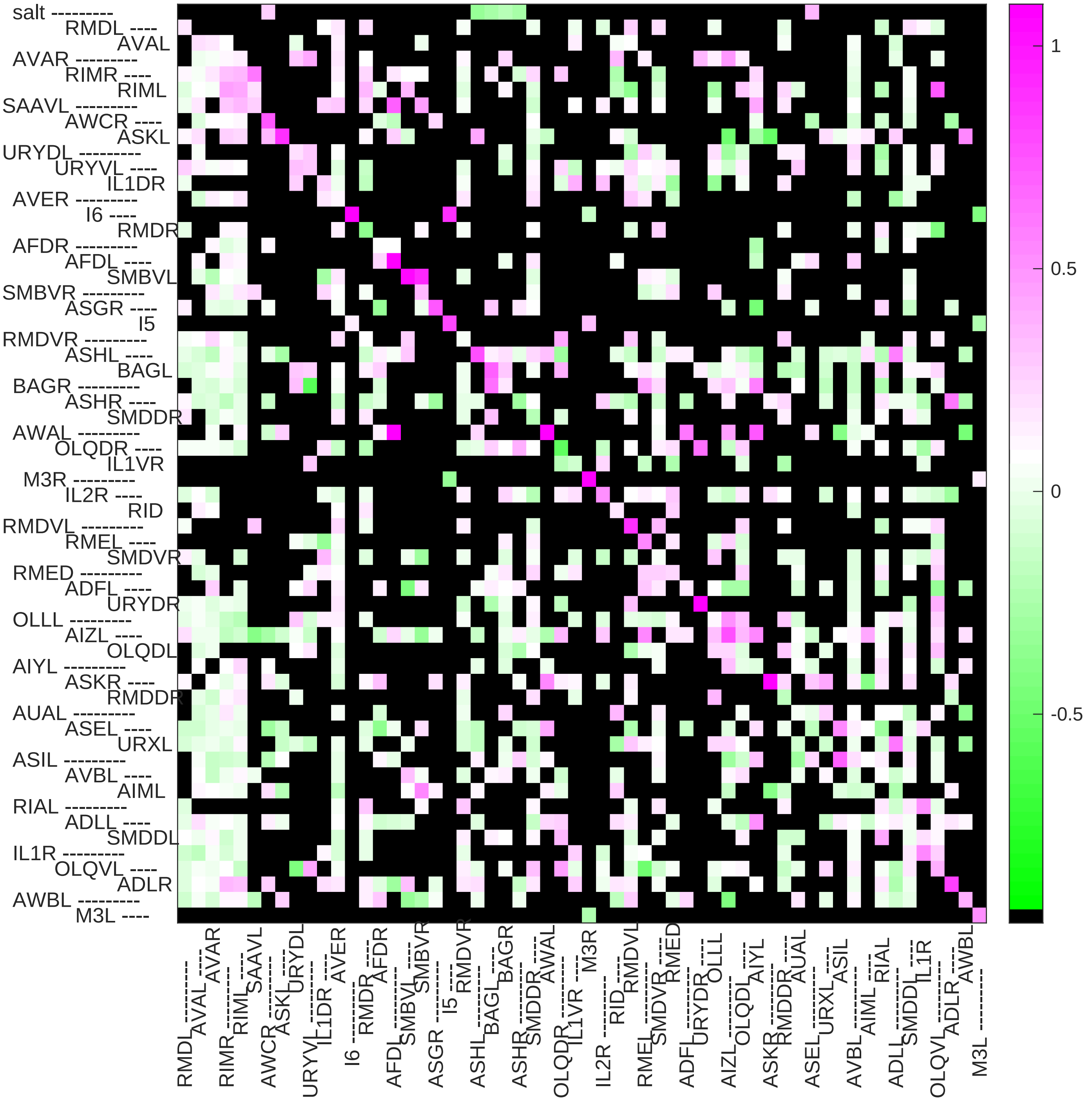

### Sample5 connection weight matrix

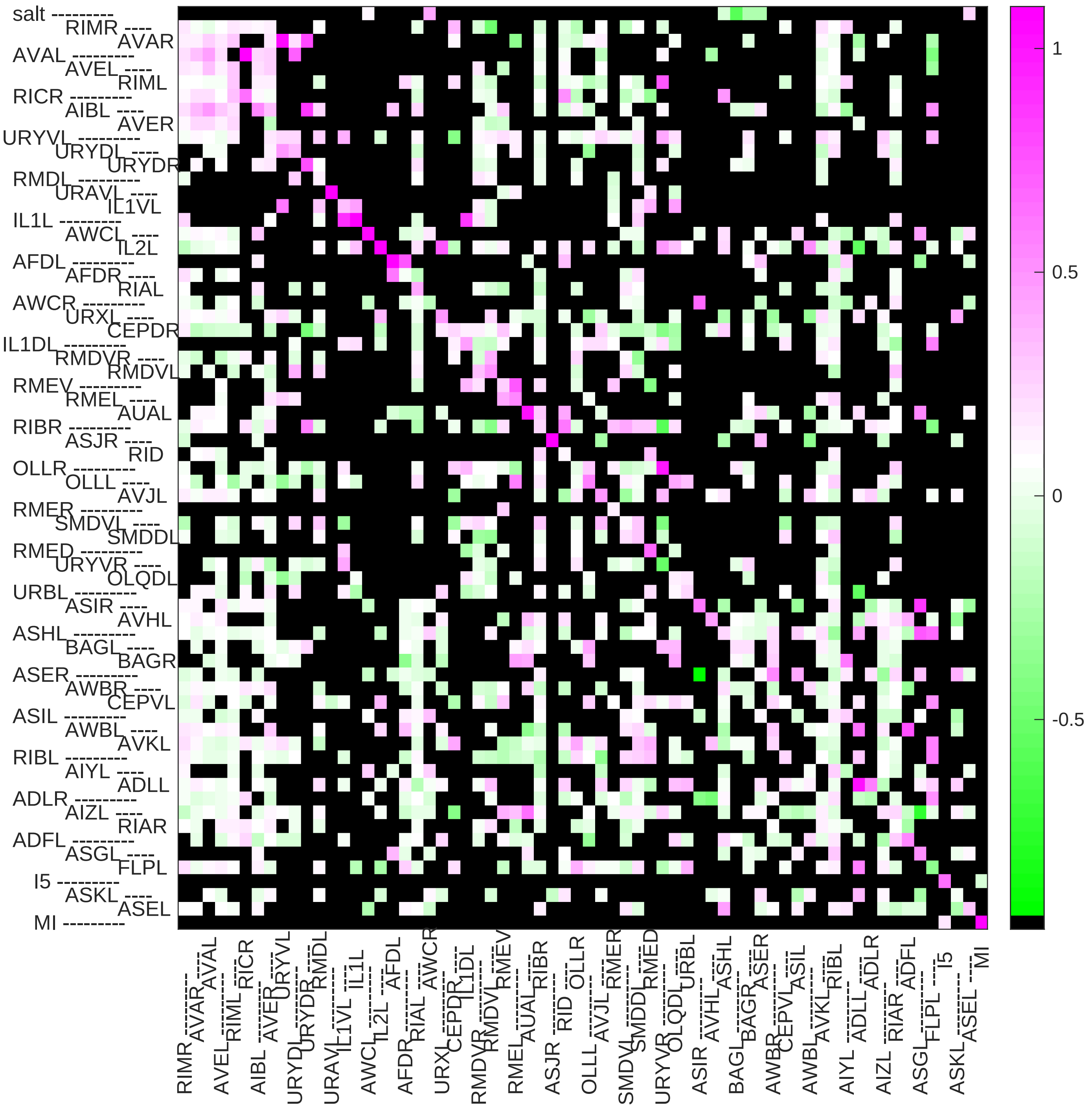

### Sample6 connection weight matrix

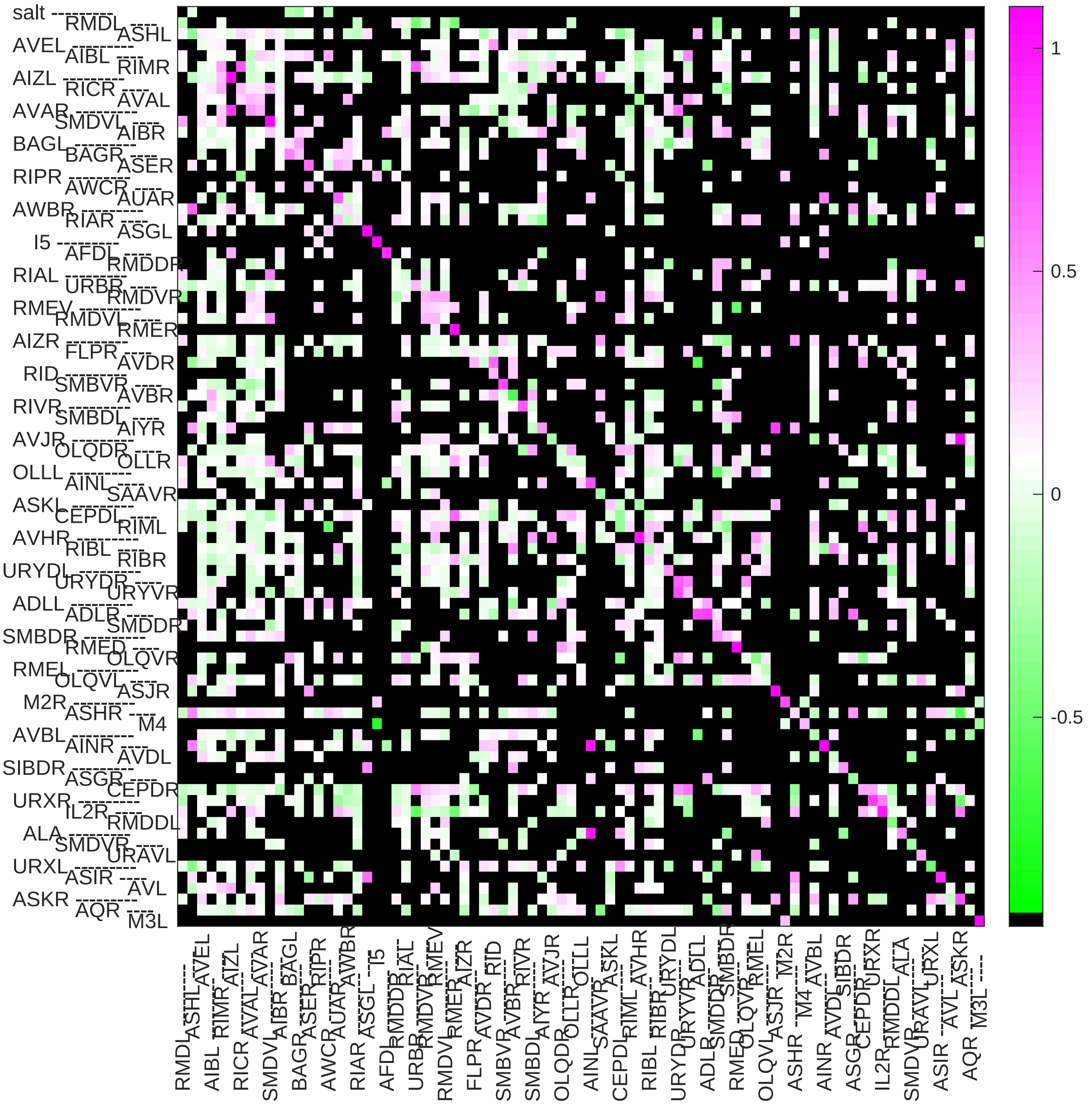

### Sample7 connection weight matrix

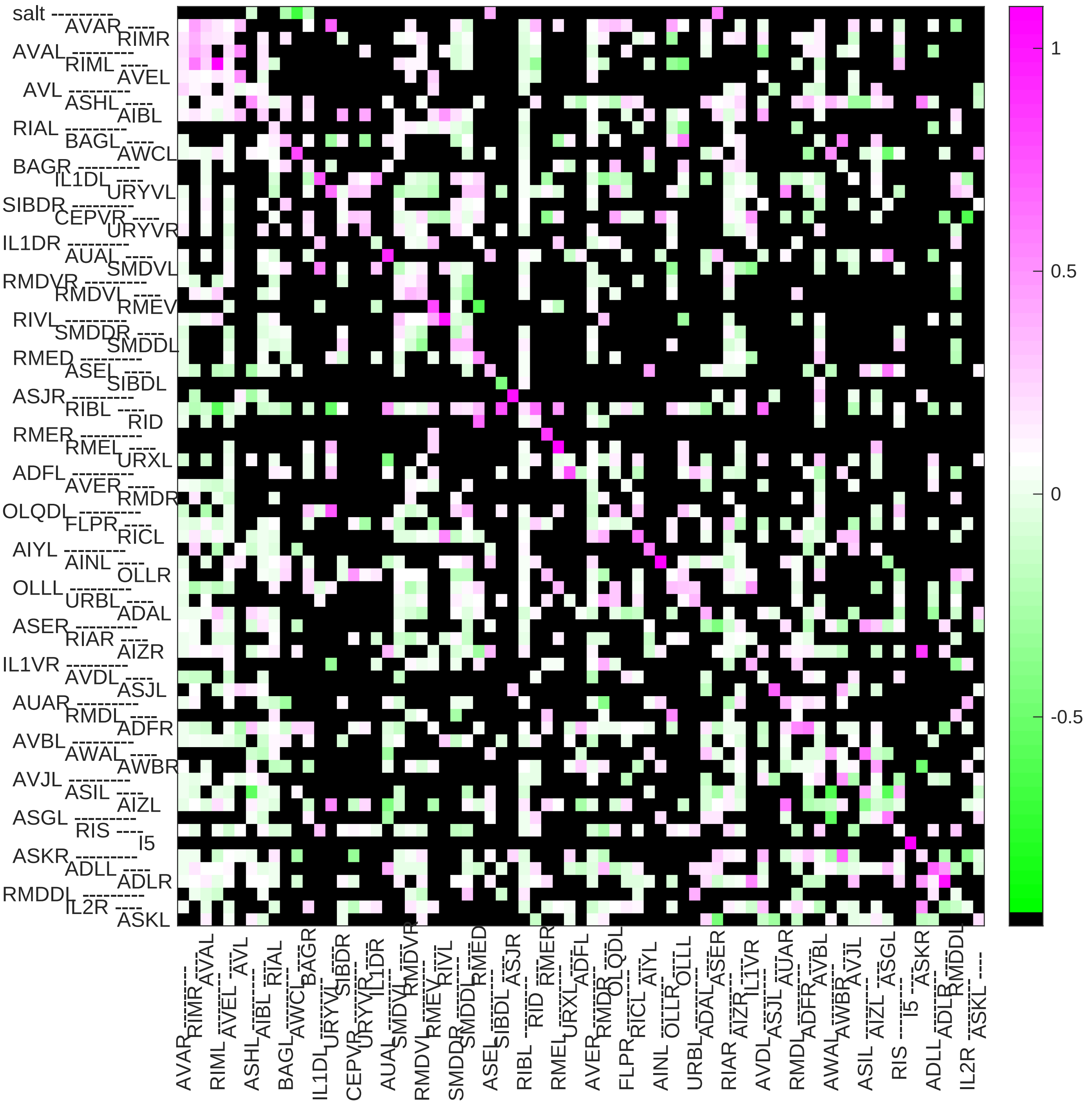

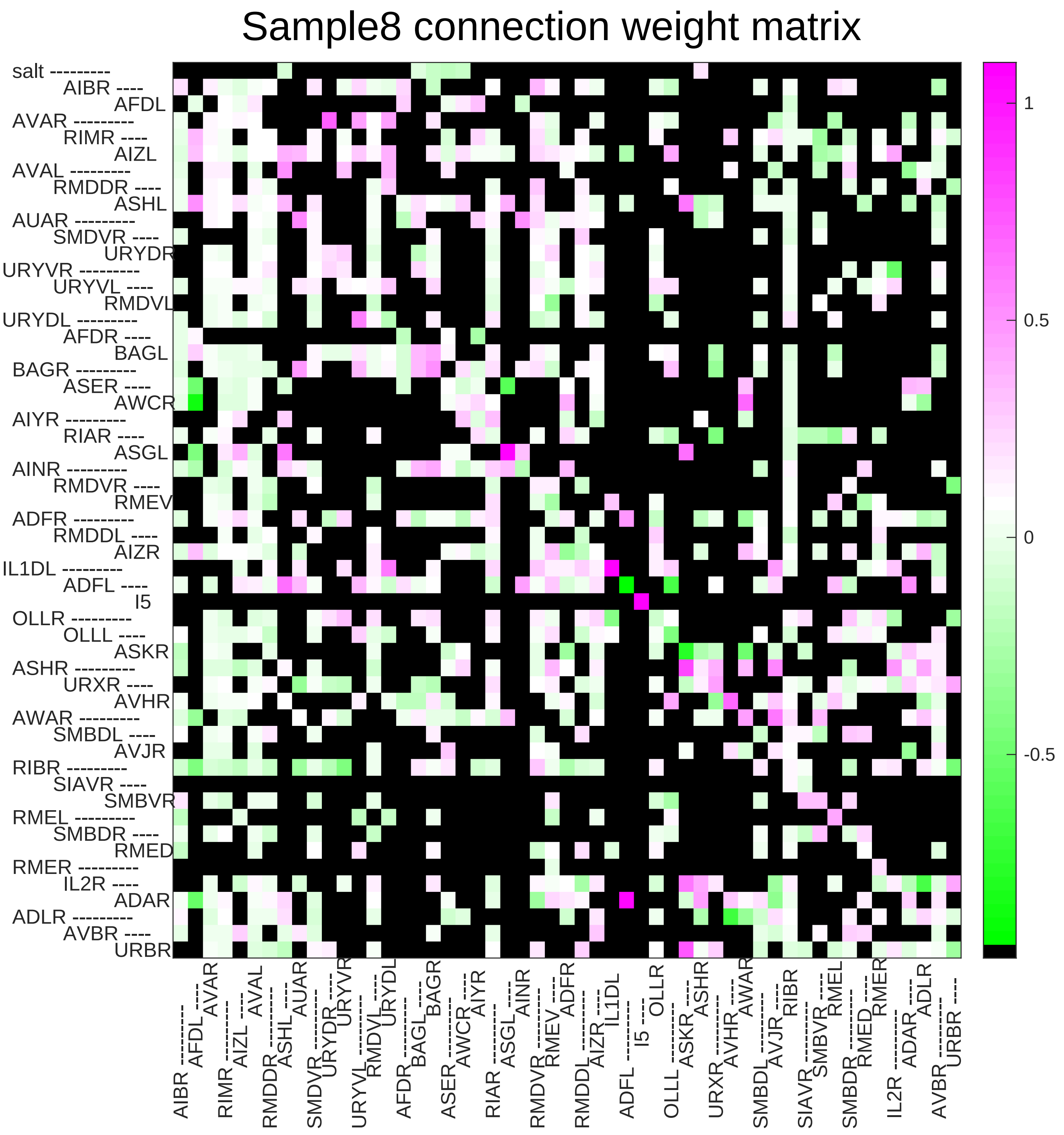

Sample8 connection weight matrix
salt
AIBR
AFDL
AVAR
RIMR
AIZL
AVAL
RMDDR
ASHL
AUAR
SMDVR
URYDR
URYVR
URYVL
RMDVL
URYDL
AFDR
BAGL
BAGR
ASER
AWCR
AIYR
RIAR
ASGL
AINR
RMDVR
RMEV
ADFR
RMDDL
AIZR
IL1DL
ADFL
I5
OLLR
OLLL
ASKR
ASHR
URXR
AVHR
AWAR
SMBDL
AVJR
RIBR
SIAVR
SMBVR
RMEL
SMBDR
RMED
RMER
IL2R
ADAR
ADLR
AVBR
URBR
1
0.5
0
-0.5

### Sample9 connection weight matrix

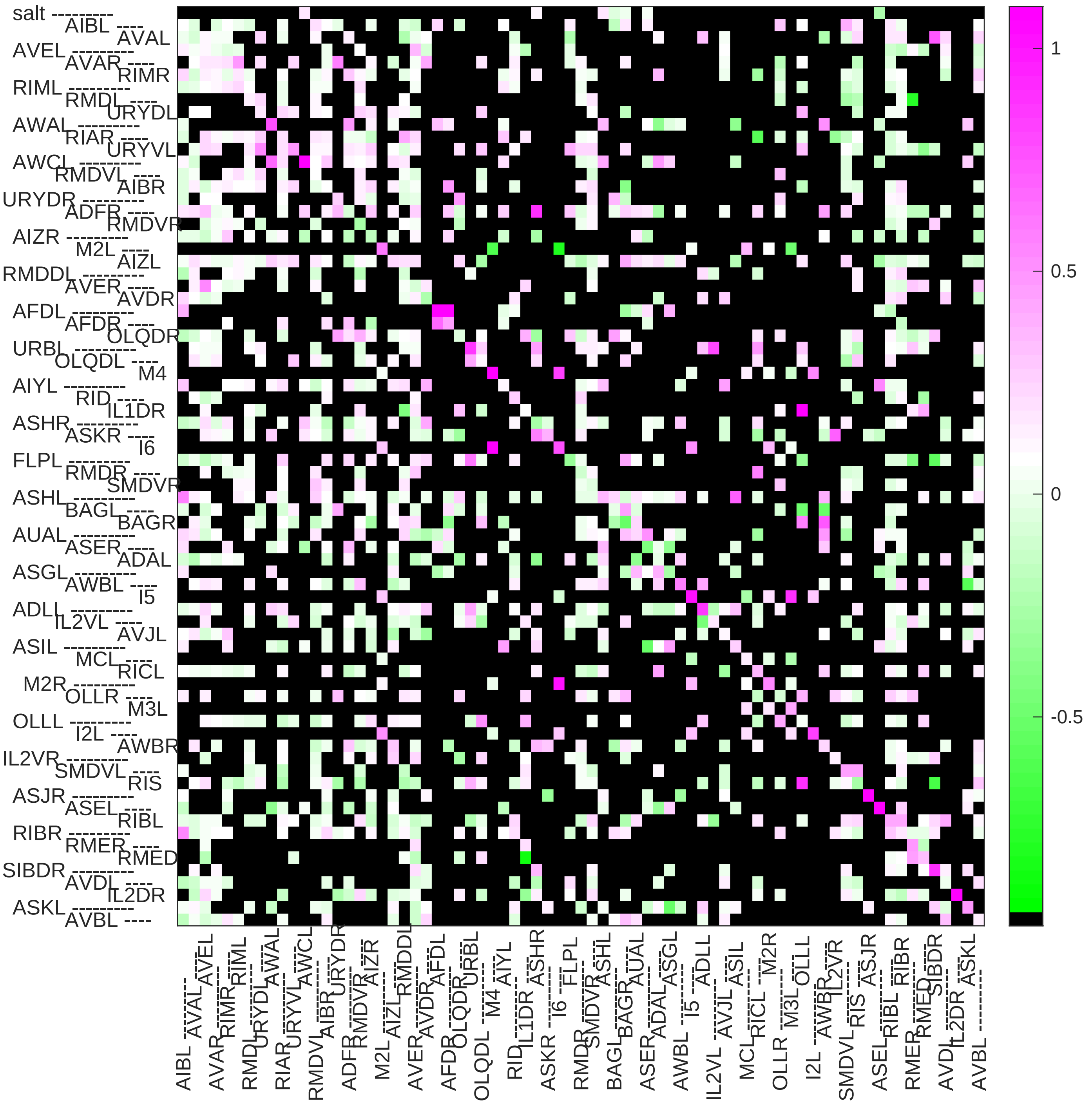

### Sample10 connection weight matrix

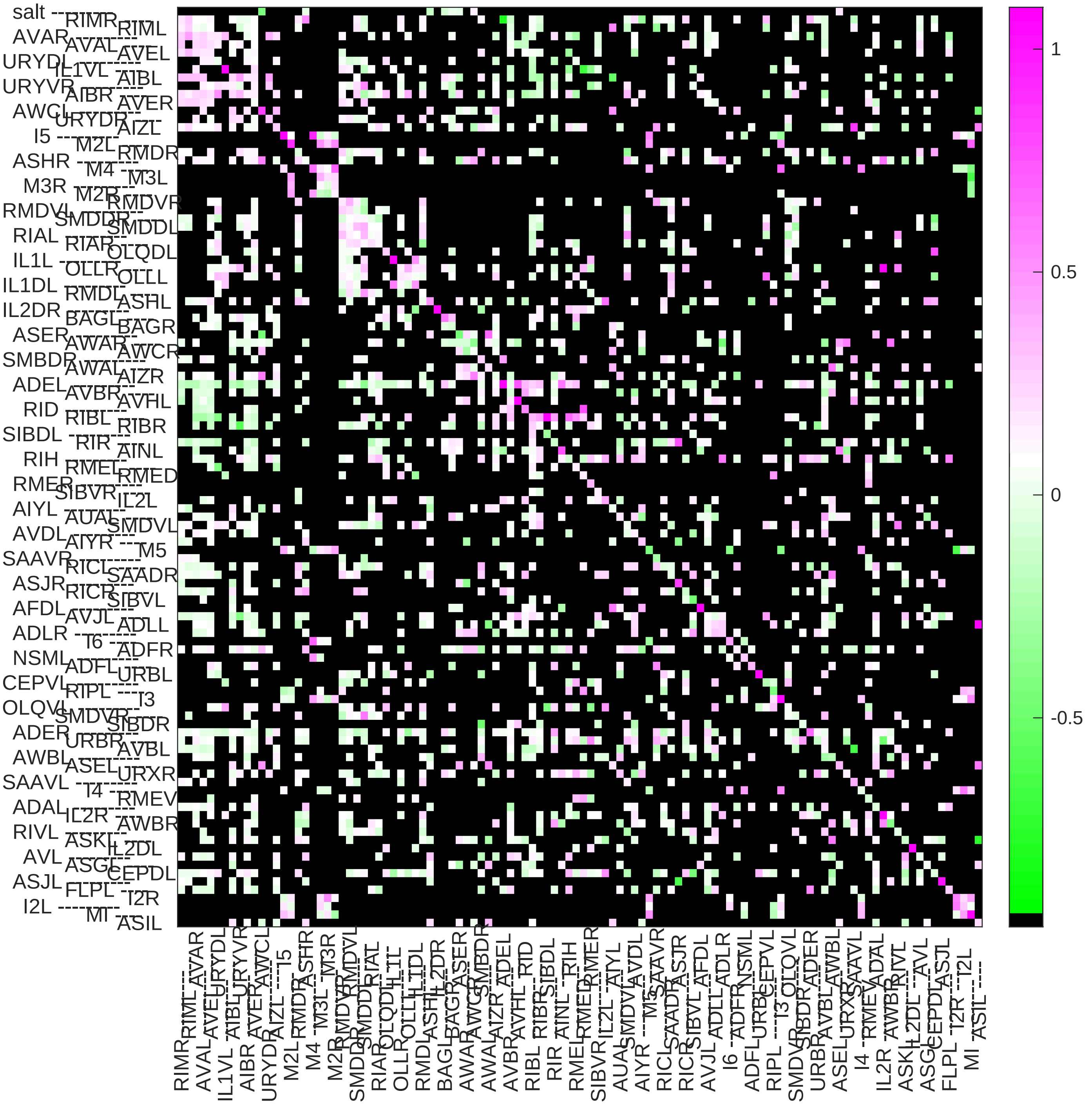

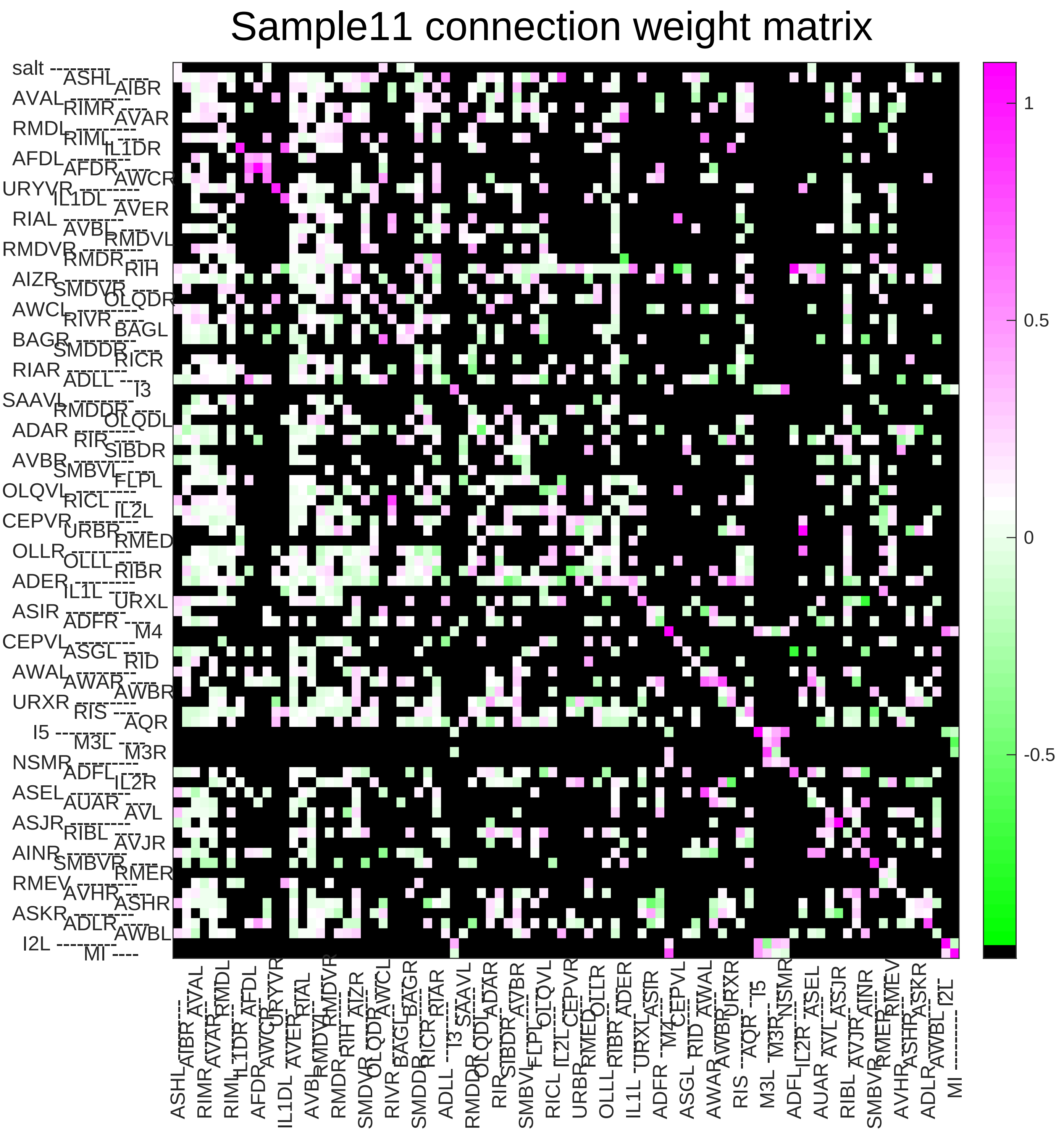
Sample11 connection weight matrix
1
0.5
0
-0.5

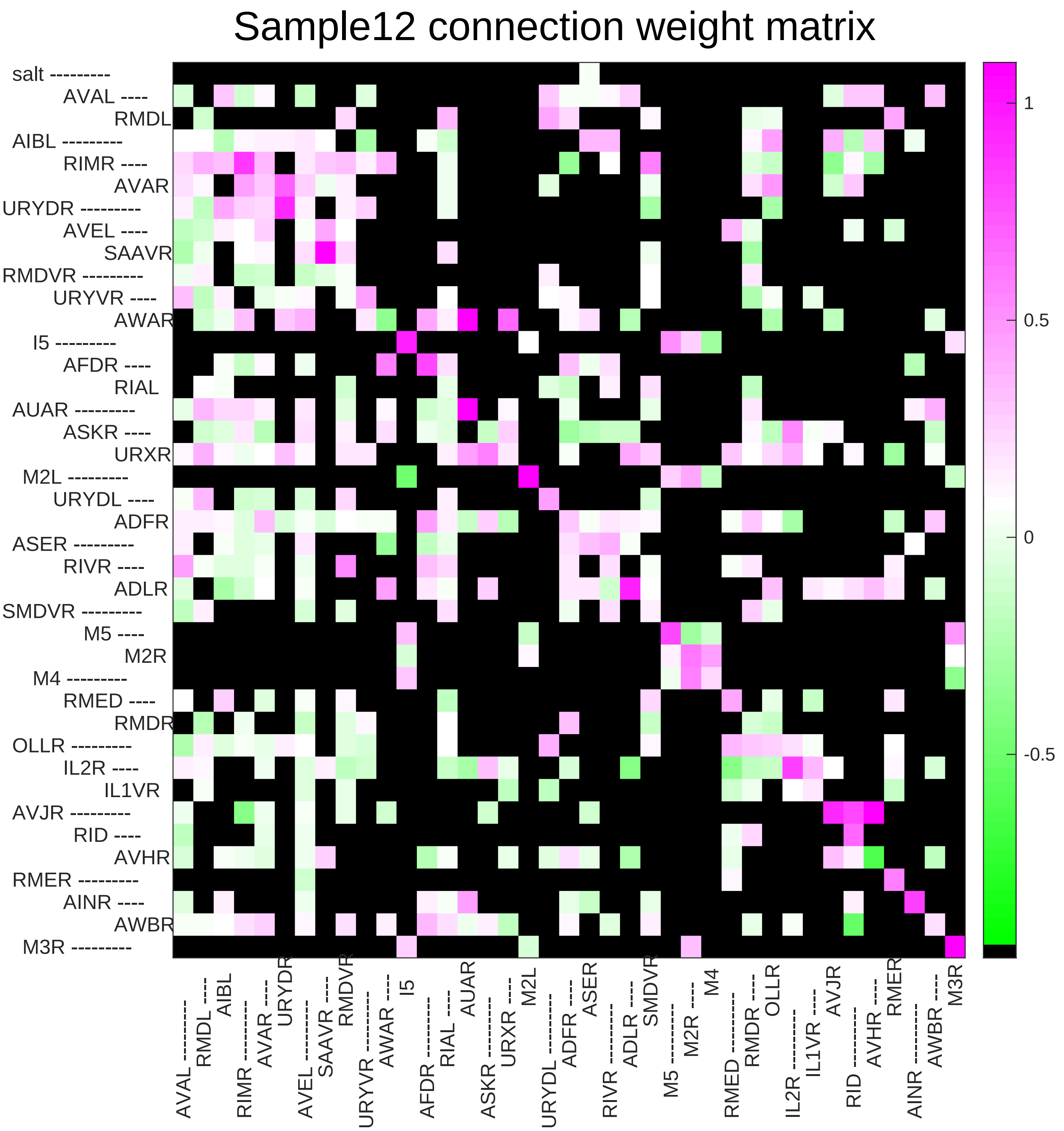

Sample12 connection weight matrix
salt
AVAL
RMDL
AIBL
RIMR
AVAR
URYDR
AVEL
SAAVR
RMDVR
URYVR
AWAR
I5
AFDR
RIAL
AUAR
ASKR
URXR
M2L
URYDL
ADFR
ASER
RIVR
ADLR
SMDVR
M5
M2R
M4
RMED
RMDR
OLLR
IL2R
IL1VR
AVJR
RID
AVHR
RMER
AINR
AWBR
M3R
1
0.5
0
-0.5

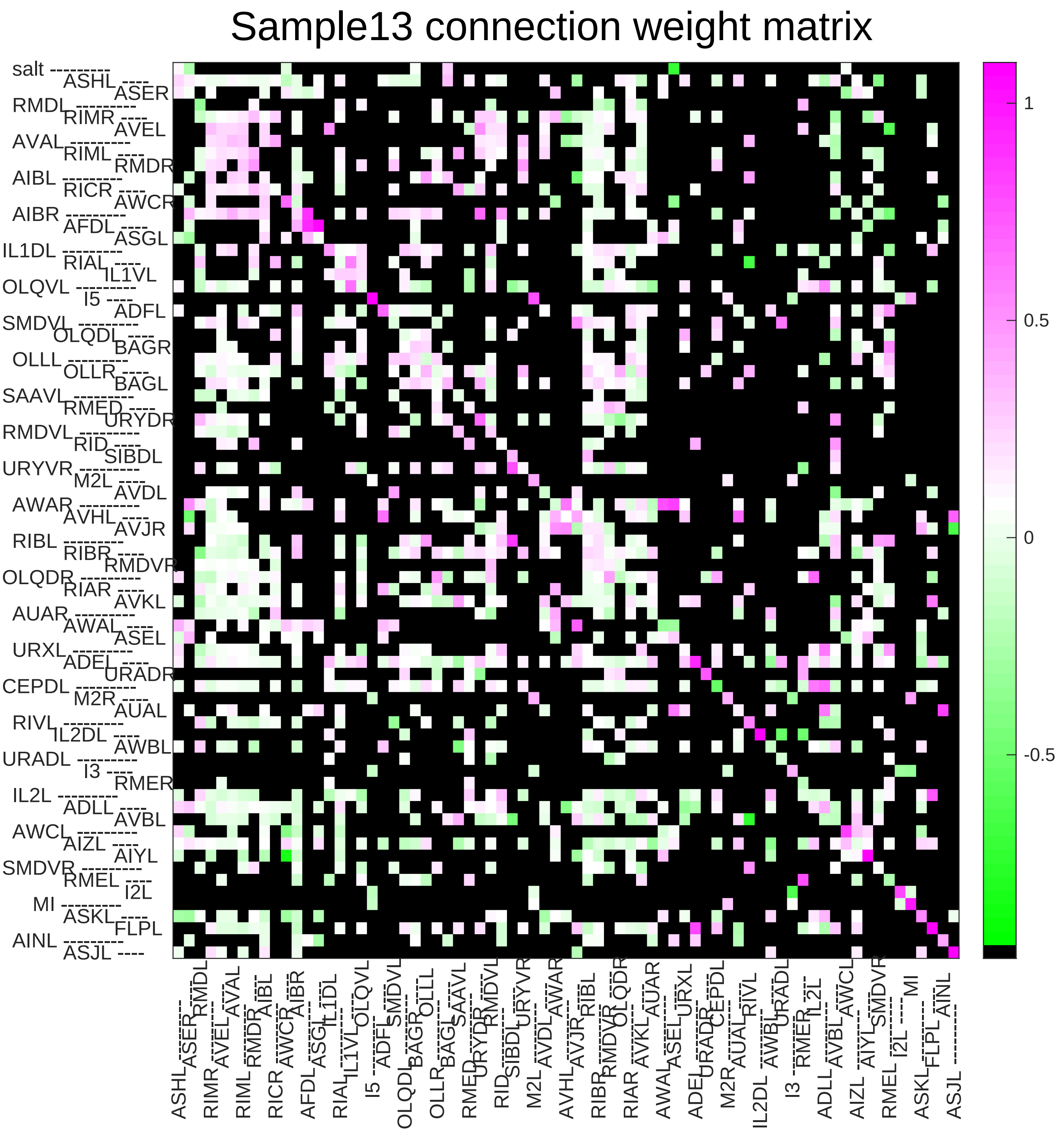
Sample13 connection weight matrix
1
0.5
0
-0.5

### Sample14 connection weight matrix

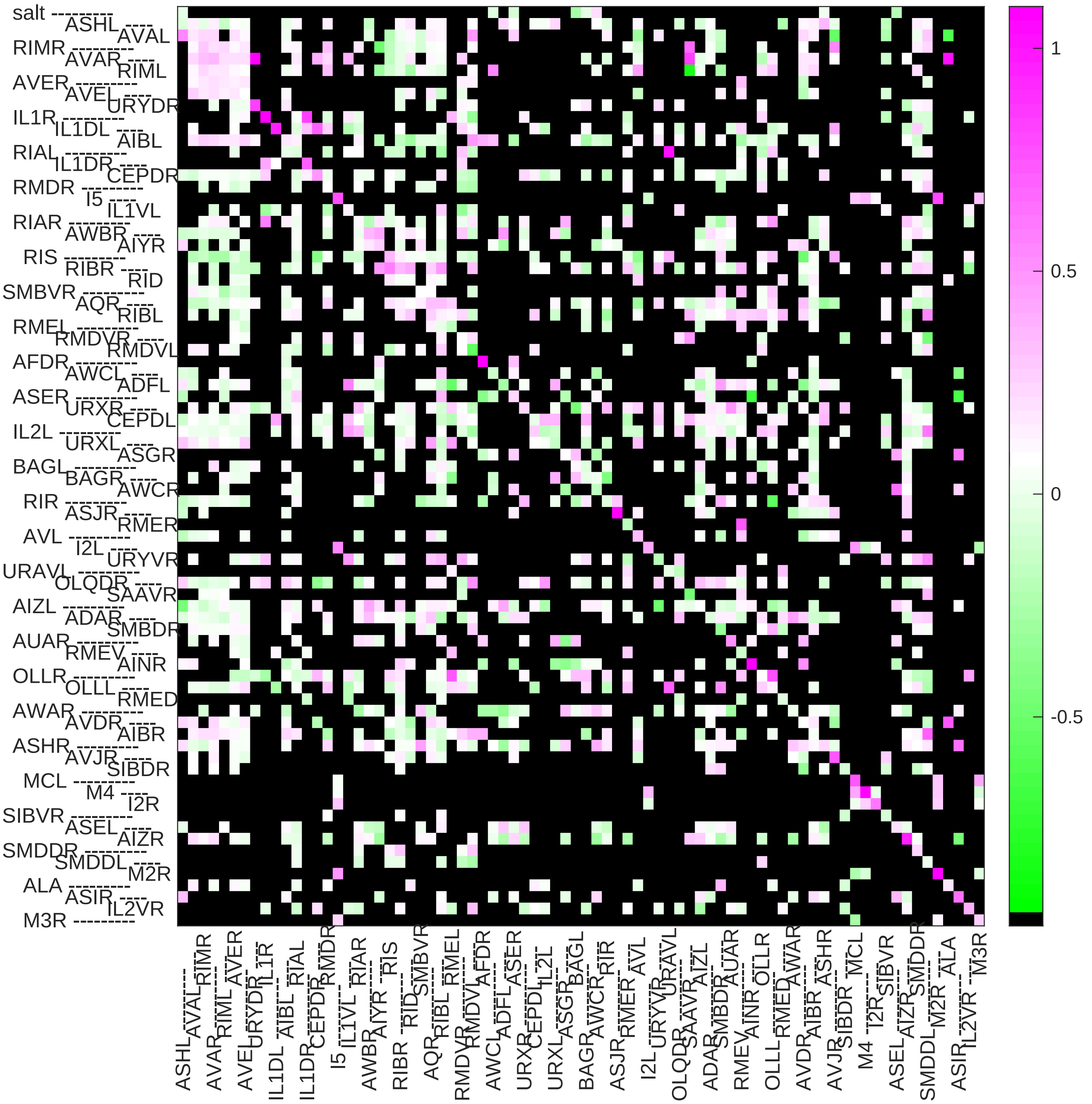

### Sample15 connection weight matrix

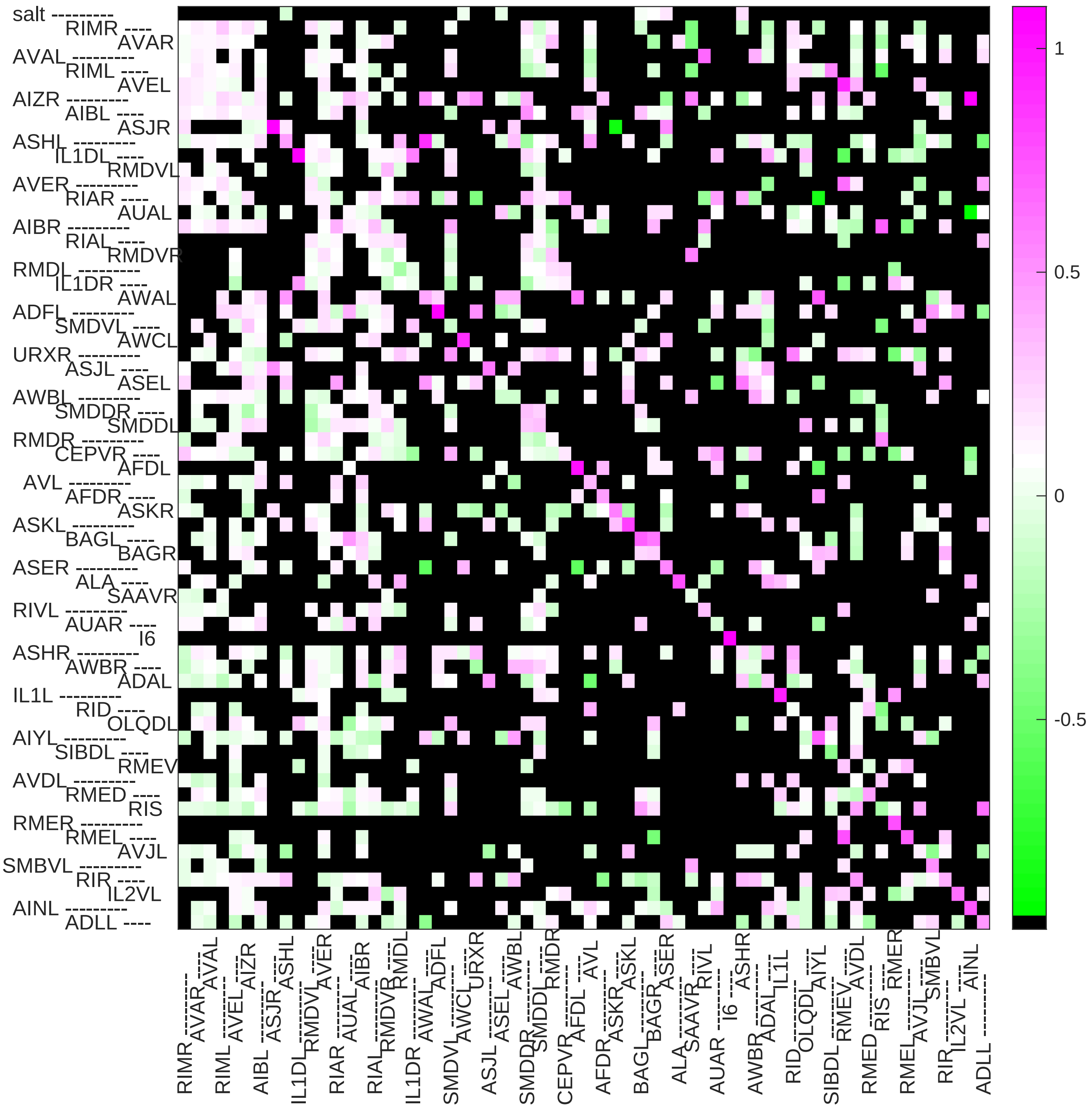

### Sample16 connection weight matrix

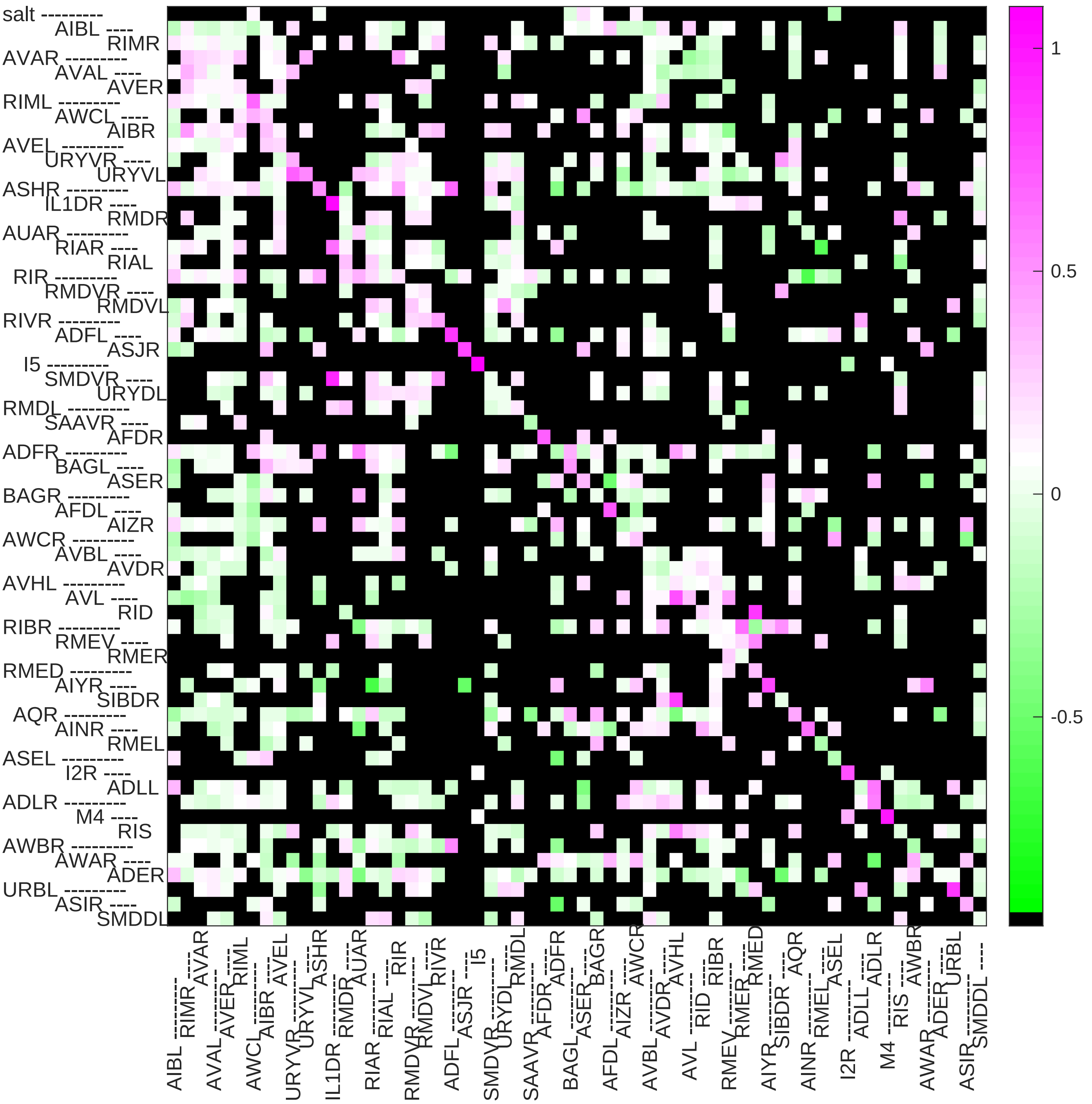

### Sample17 connection weight matrix

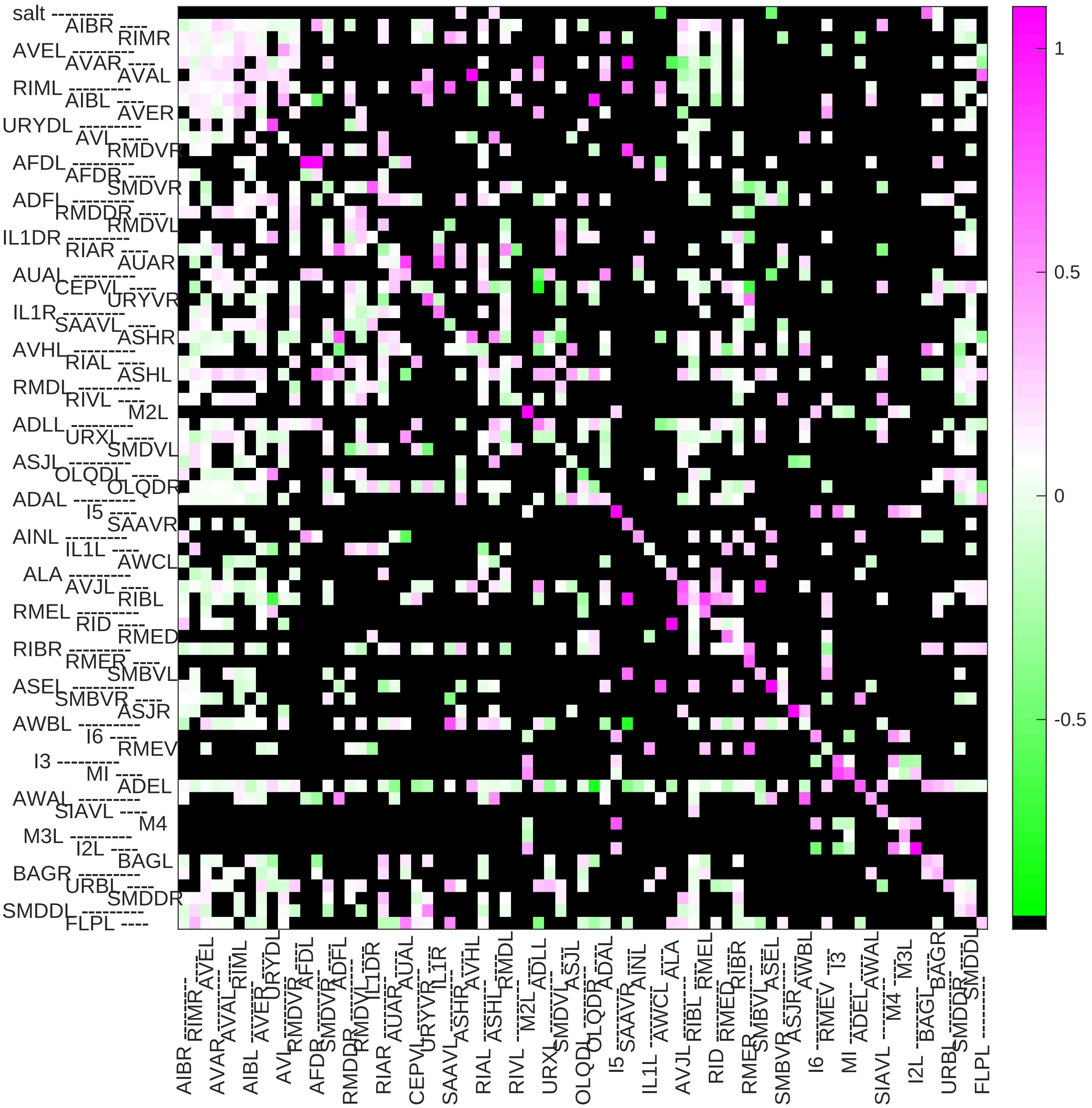

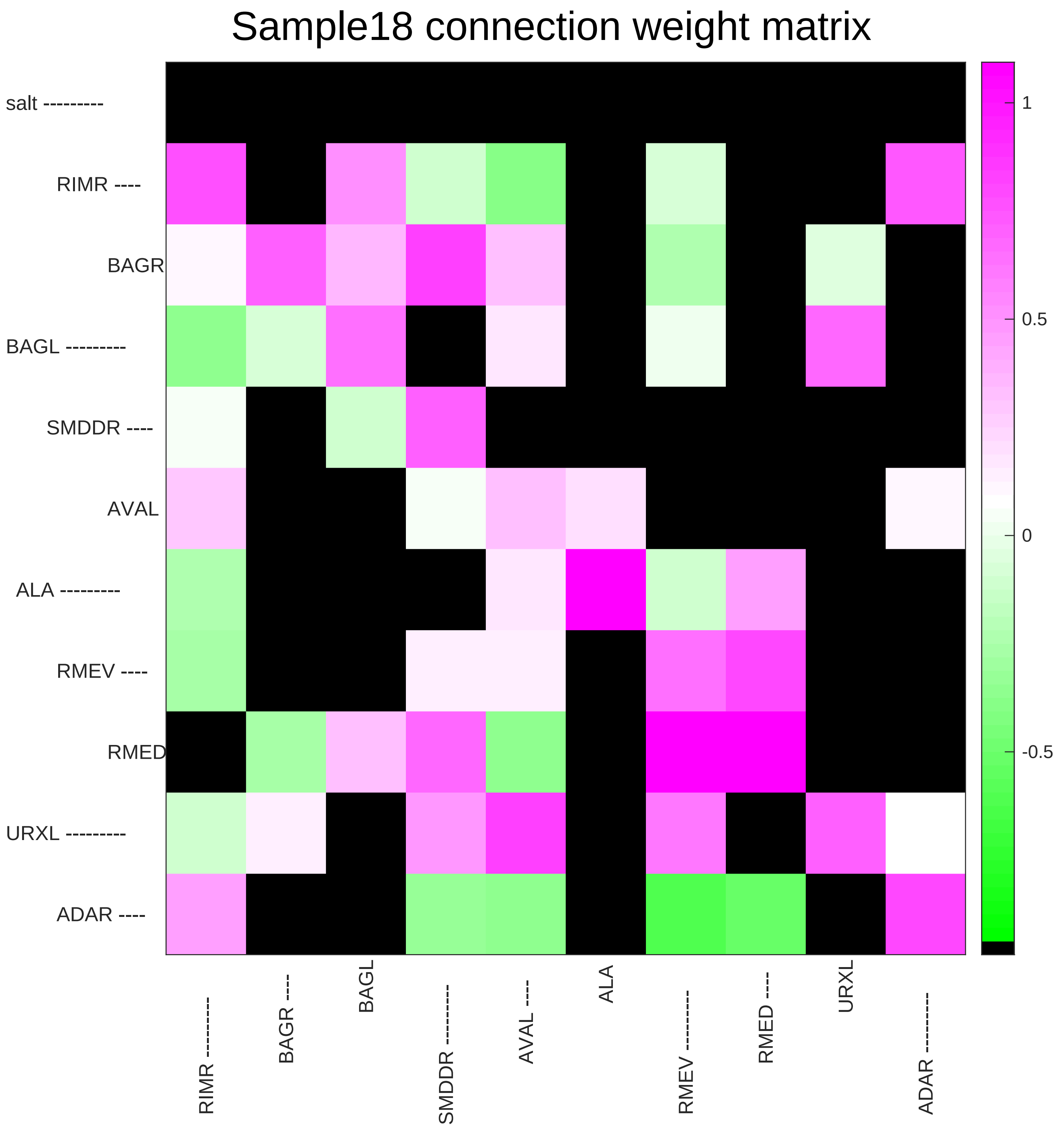

Sample18 connection weight matrix
salt ---------
RIMR ----
BAGR
BAGL ---------
SMDDR ----
AVAL
ALA ---------
RMEV ----
RMED
URXL ---------
ADAR ----
RIMR ---------
BAGR ----
BAGL
SMDDR ---------
AVAL ----
ALA
RMEV ---------
RMED ----
URXL
ADAR ---------
1
0.5
0
-0.5

### Sample19 connection weight matrix

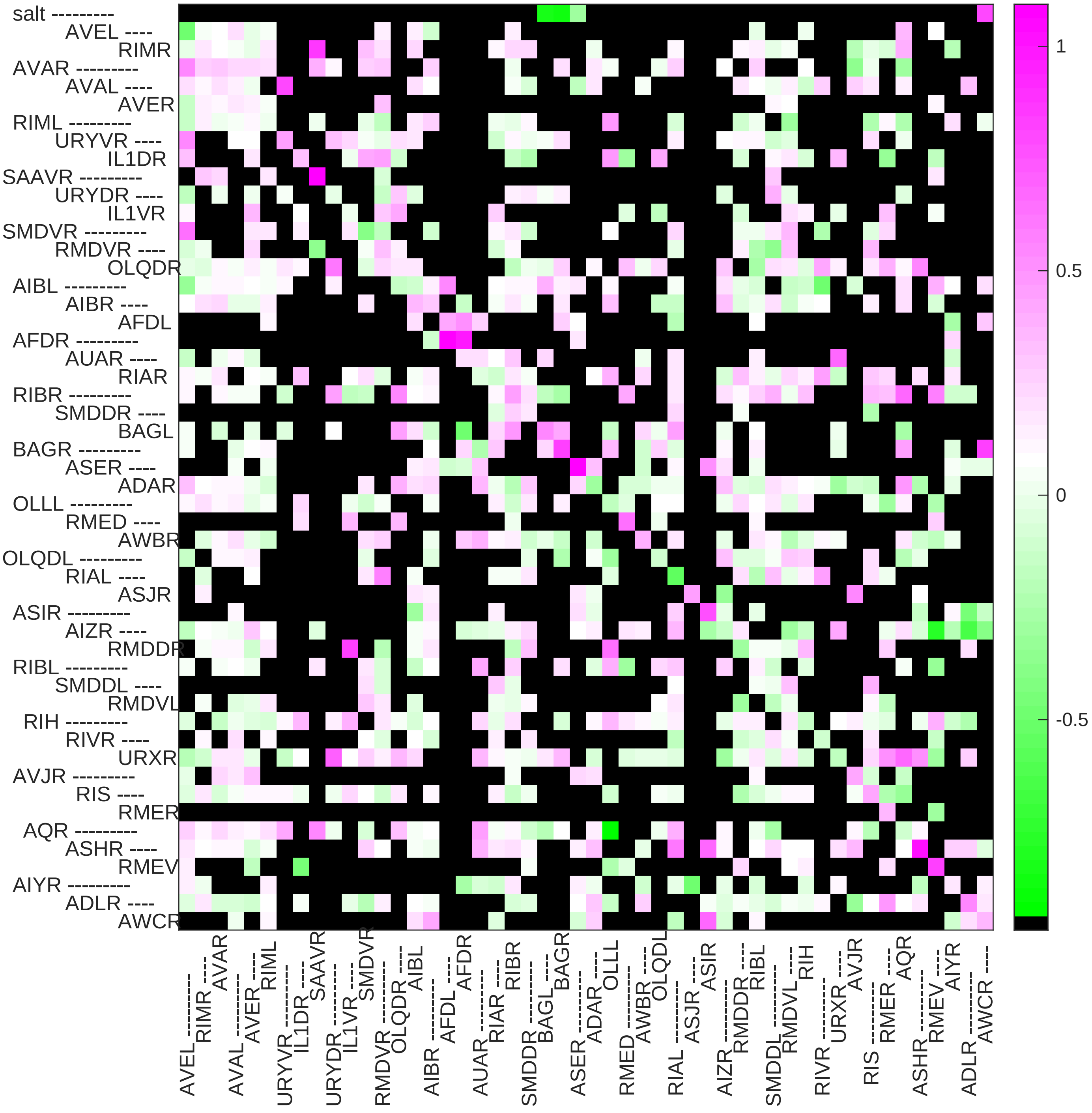

### Sample20 connection weight matrix

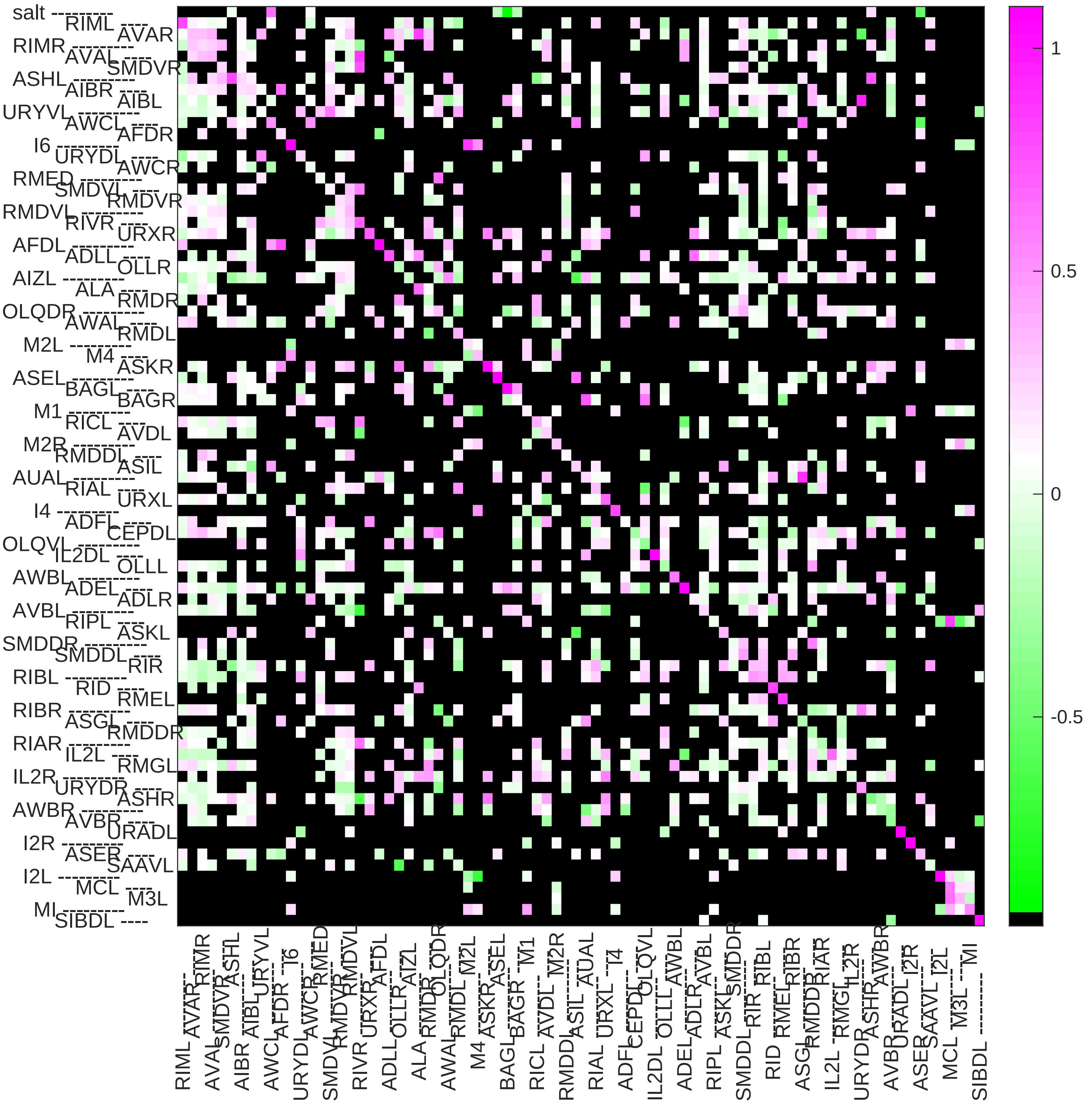

### Sample21 connection weight matrix

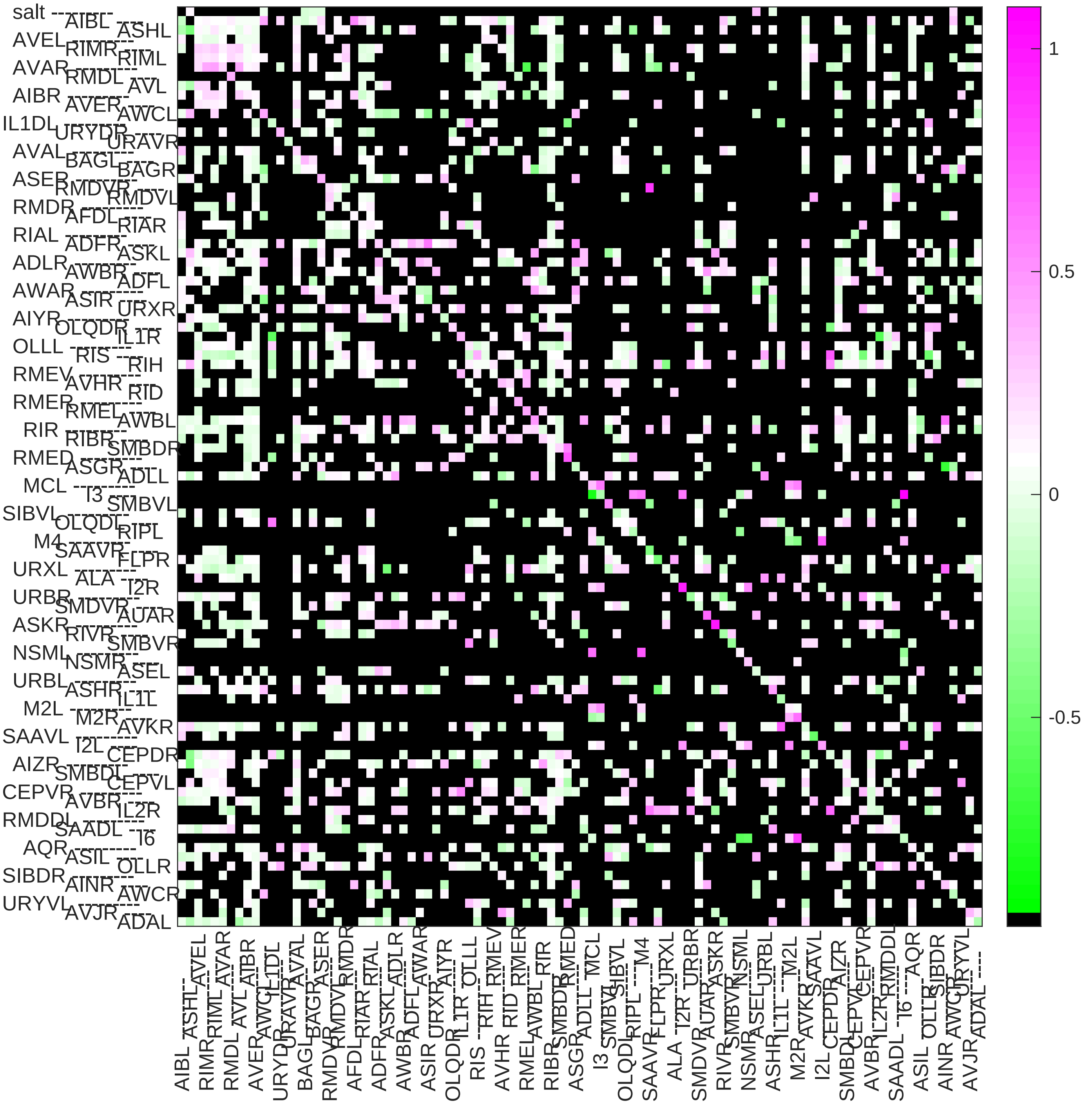

### Sample22 connection weight matrix

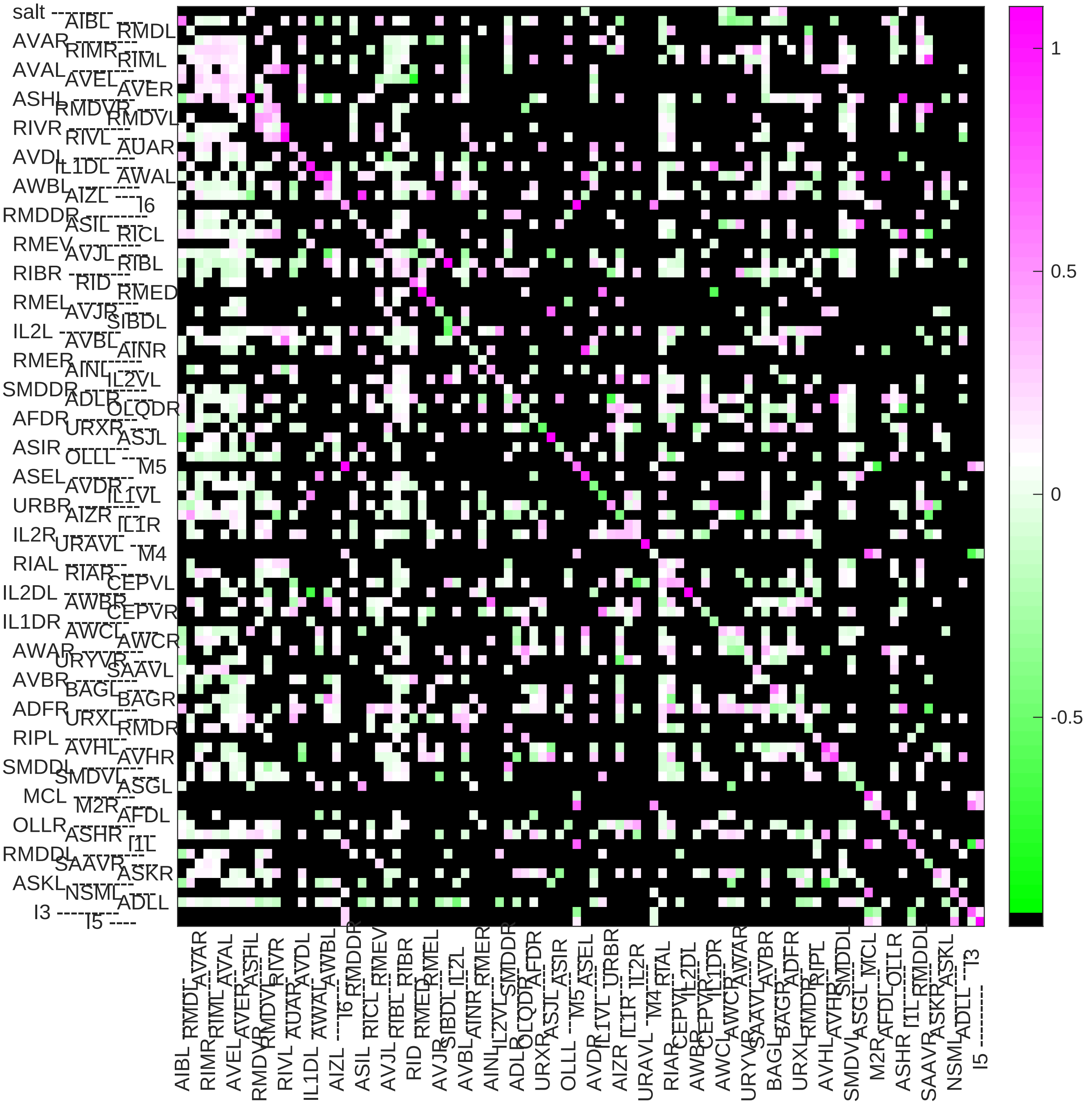

#### Sample23 connection weight matrix

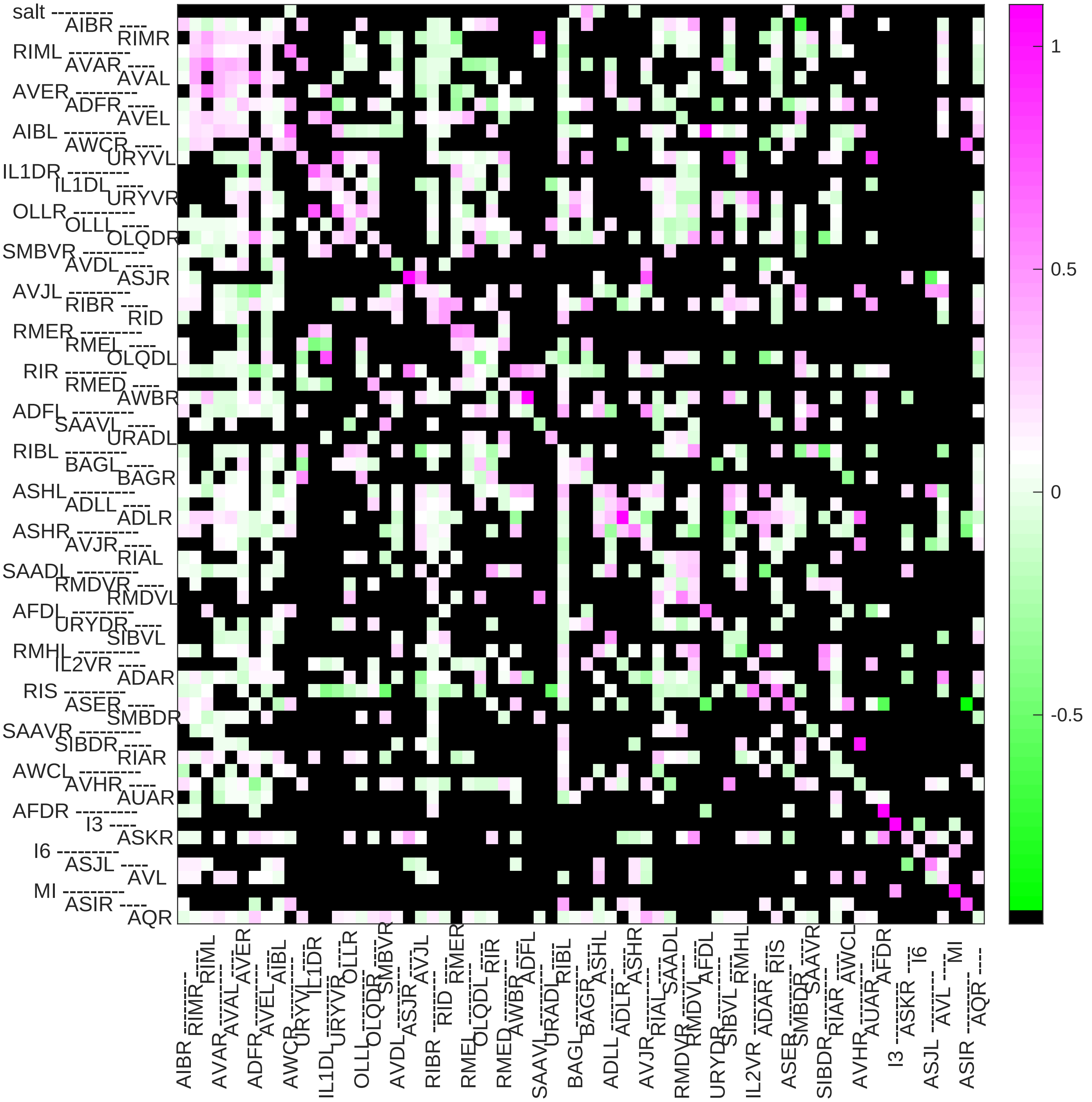

### Sample24 connection weight matrix

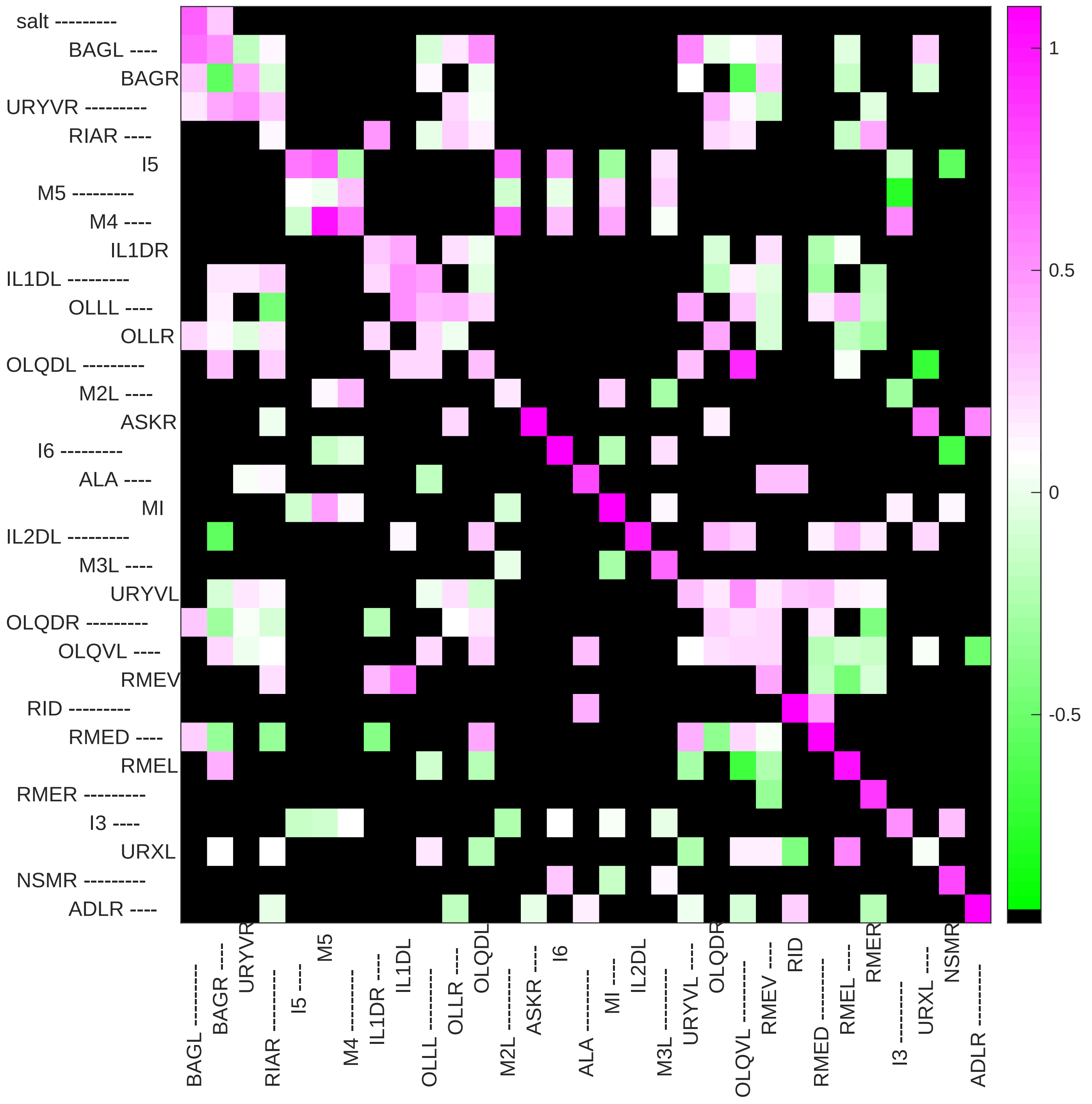
