## Supplementary File 12B for "Deducing ensemble dynamics and information flow from the whole-brain imaging data"

### sample 1

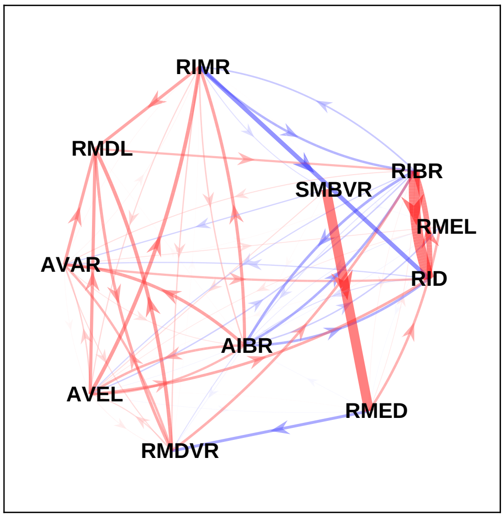


### sample 2

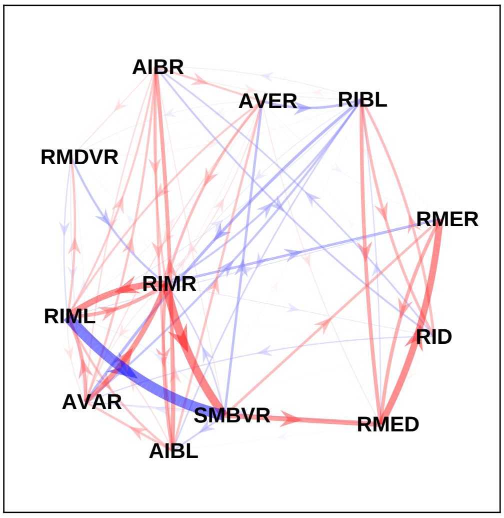


### sample 3

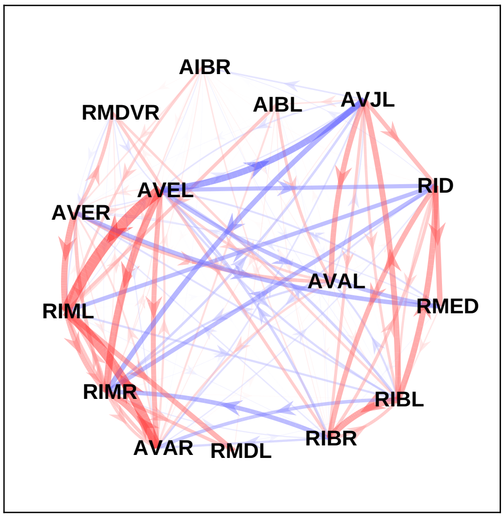


### sample 4

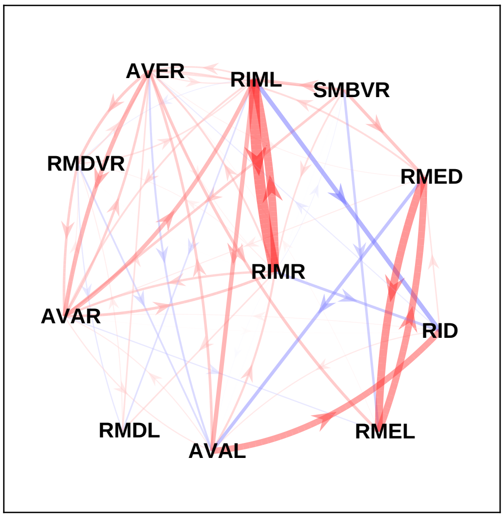


### sample 5

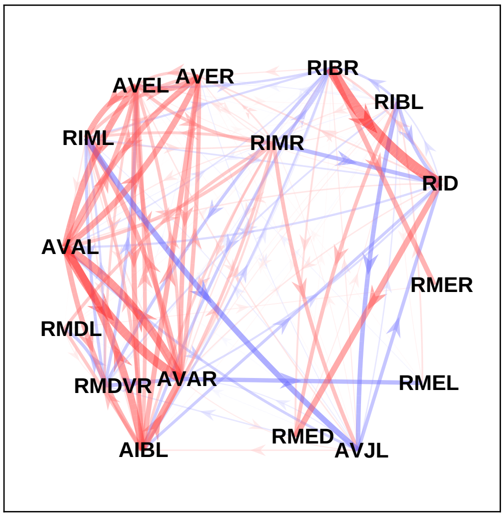


### sample 6

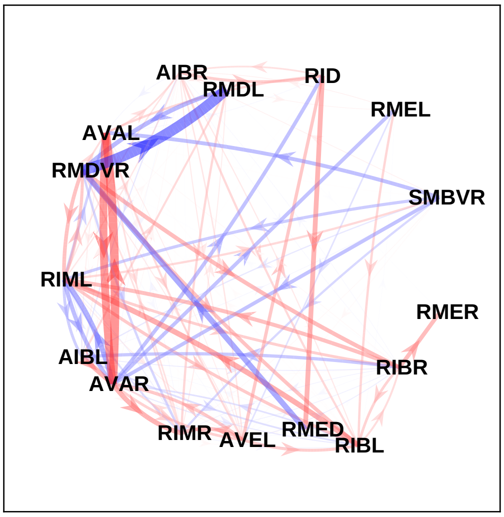

sample 7

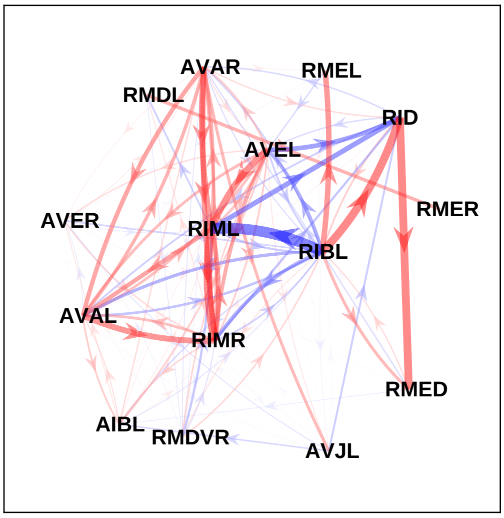


sample 8

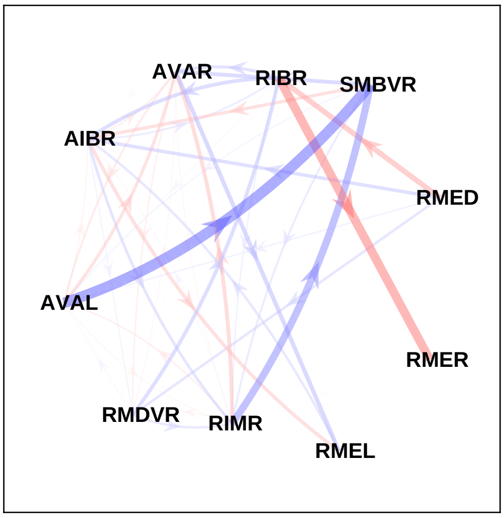


sample 9

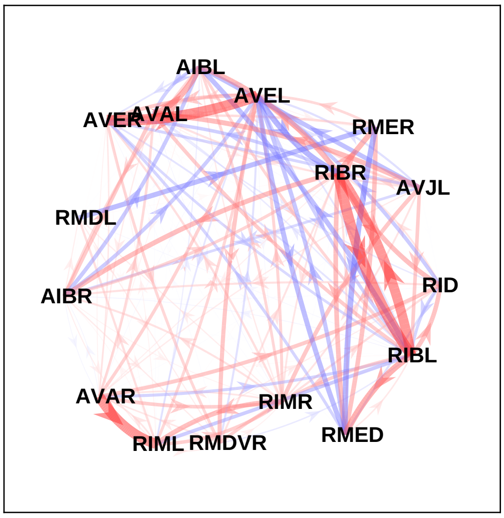


sample 10

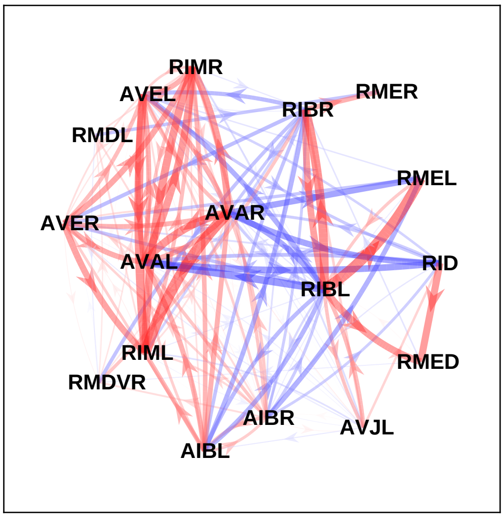


sample 11

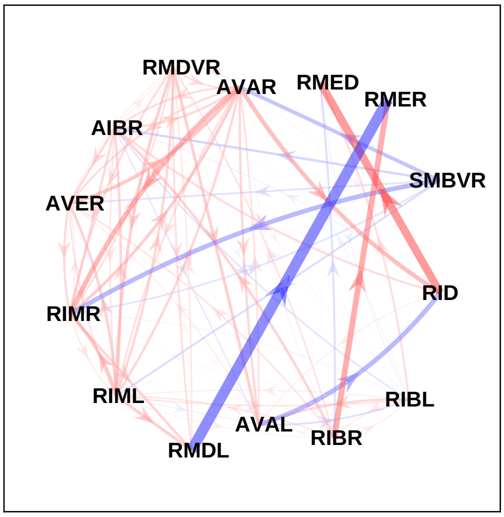


sample 12

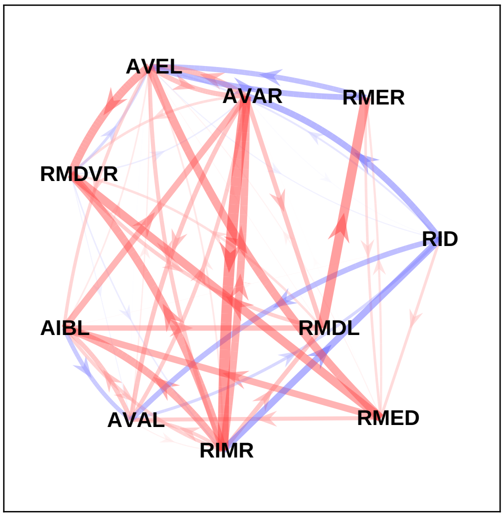

sample 13

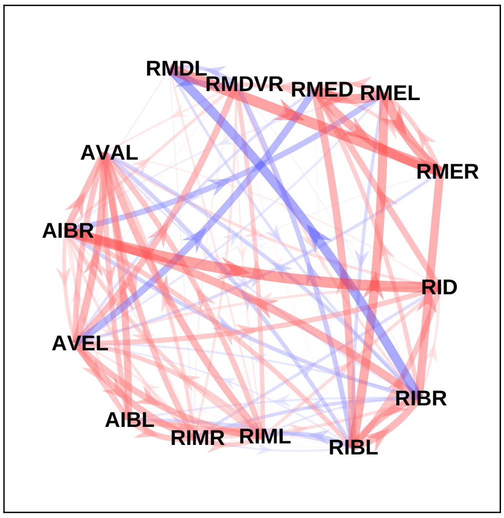


sample 14

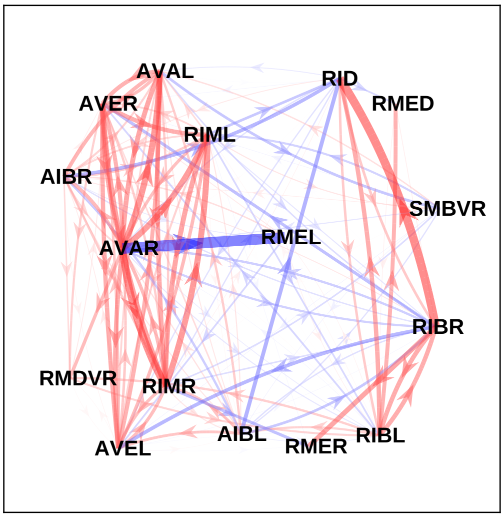


sample 15

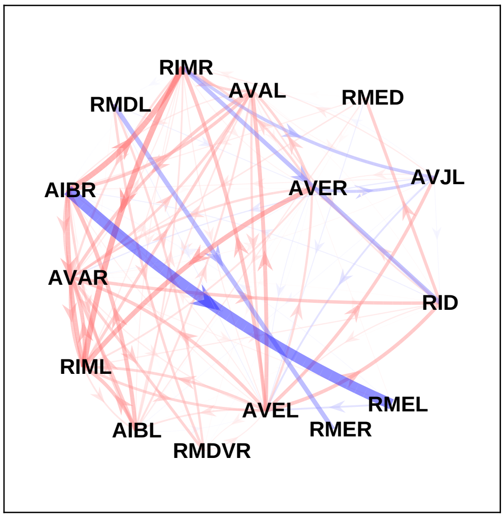


sample 16

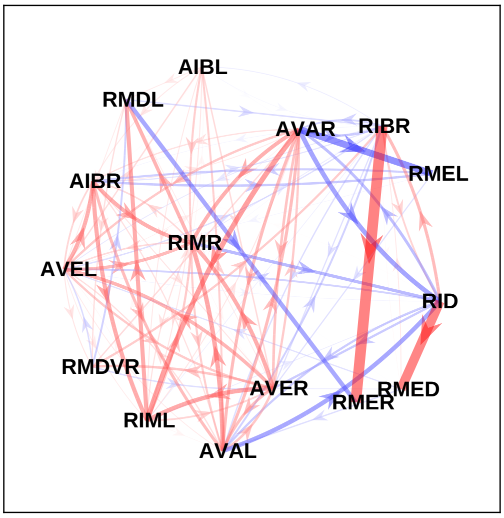


sample 17

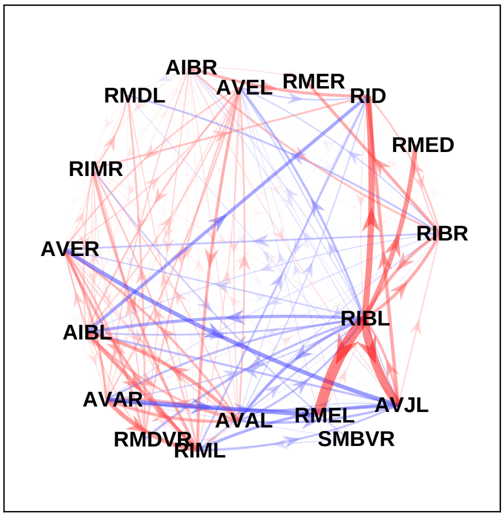


sample 18

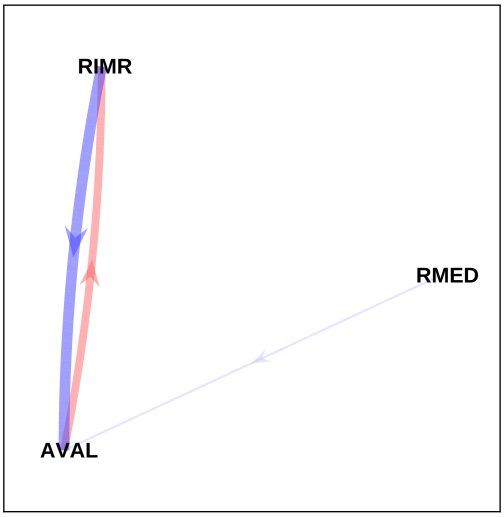

sample 19

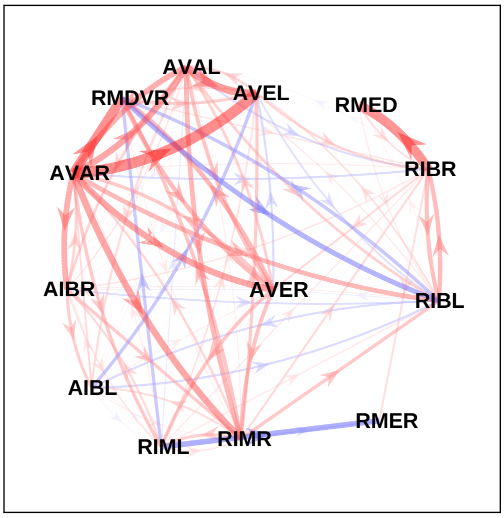


sample 20

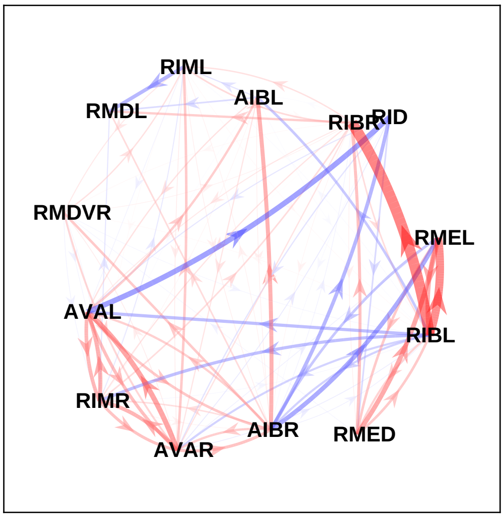


sample 21

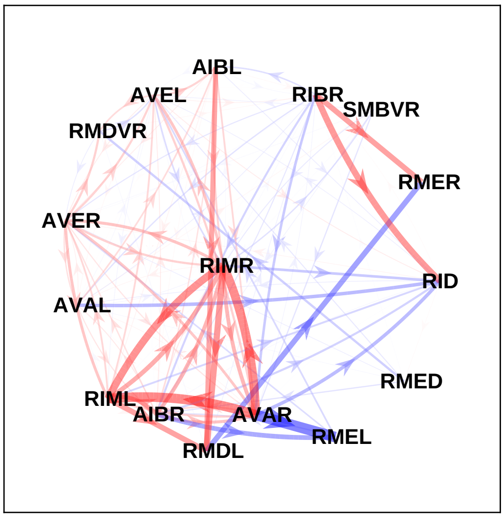


sample 22

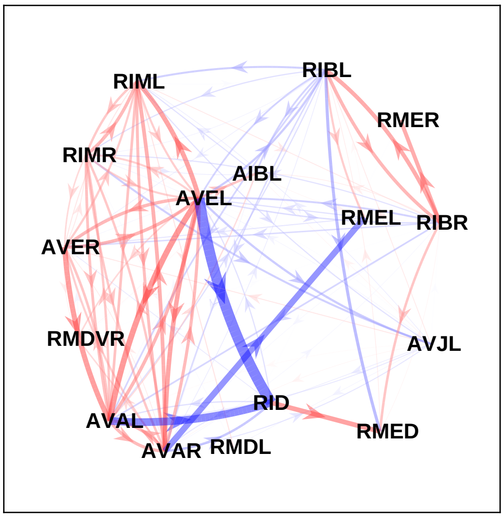


sample 23

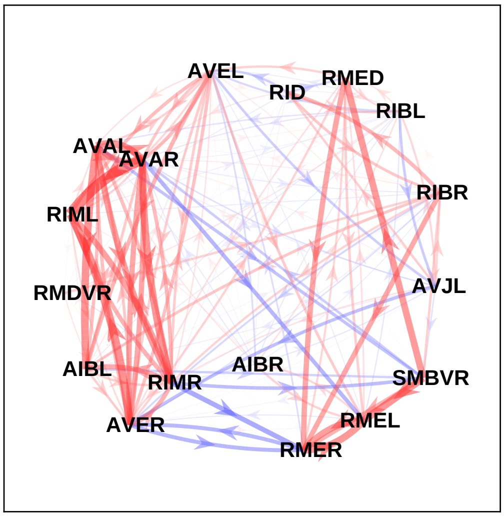


sample 24

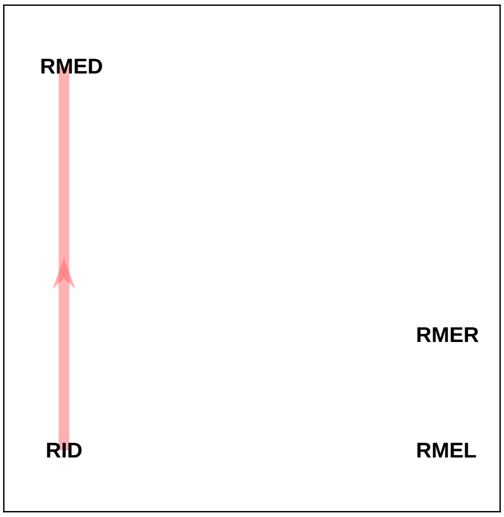
