## Supplementary File 14B for "Deducing ensemble dynamics and information flow from the whole-brain imaging data"

sample 1

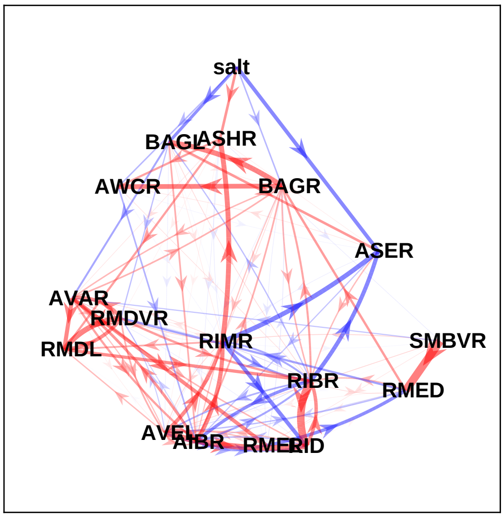


sample 2

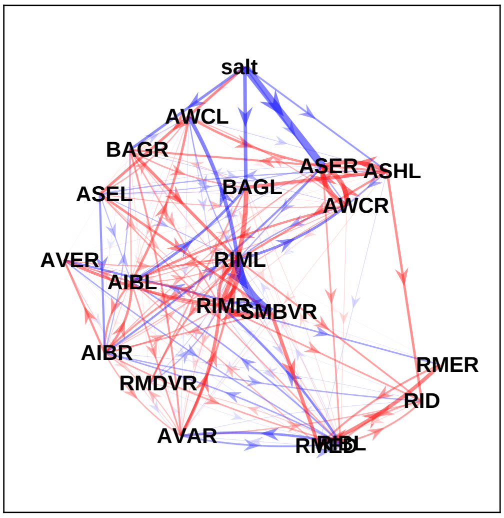


sample 3

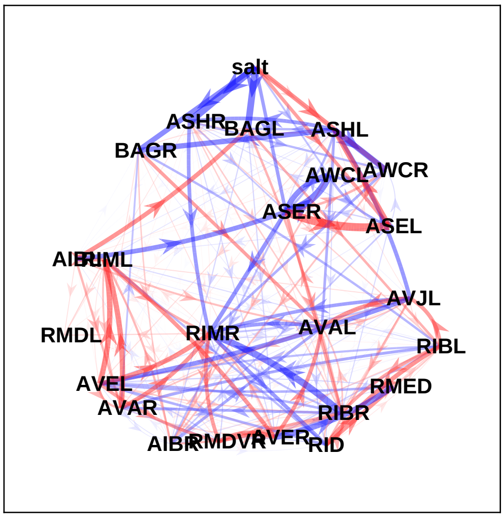


sample 4

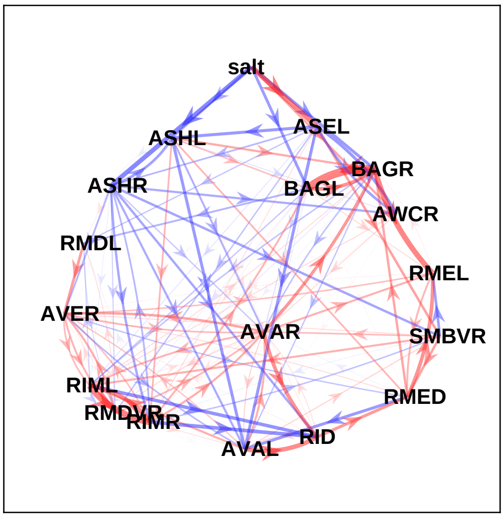


sample 5

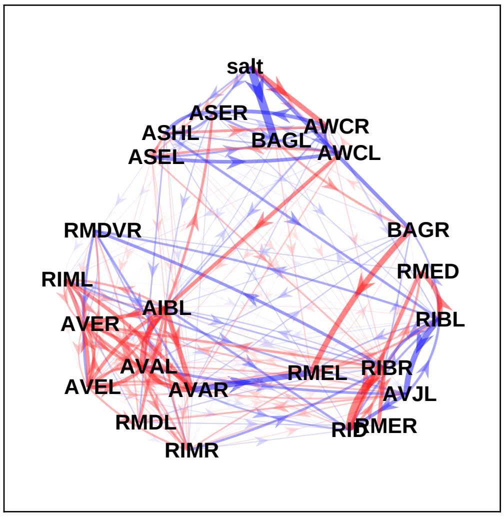


sample 6

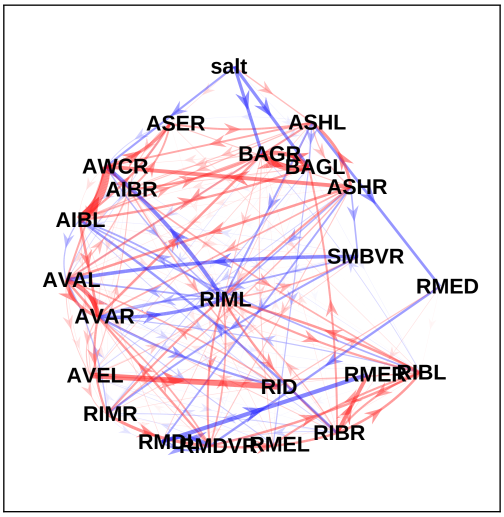

### sample 7

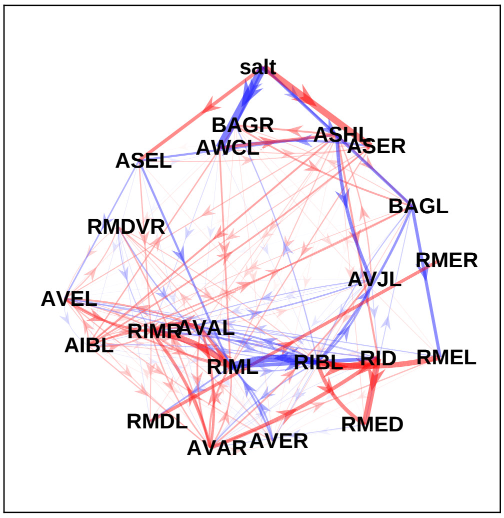


### sample 8

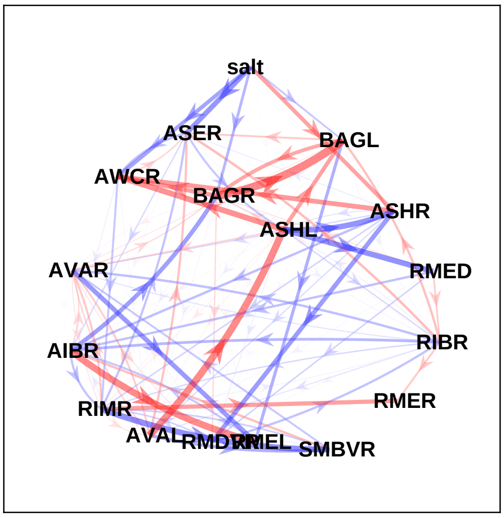


### sample 9

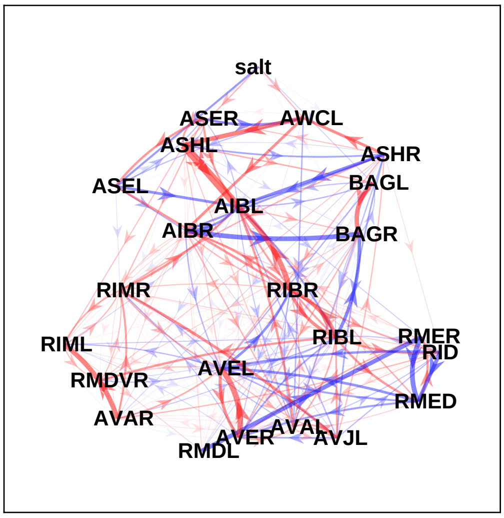


### sample 10

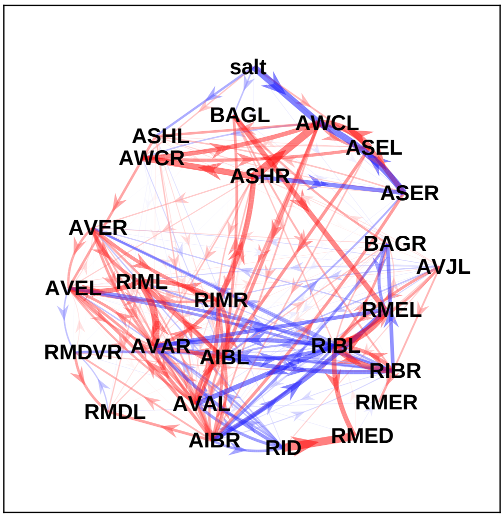


### sample 11

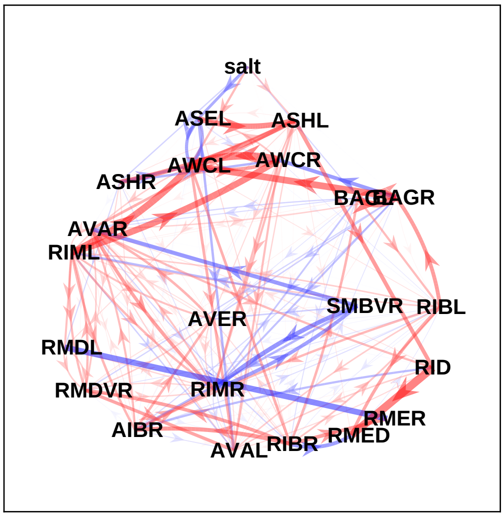


### sample 12

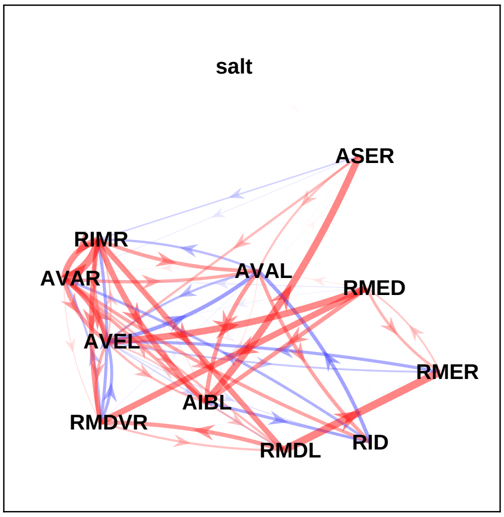

sample 13

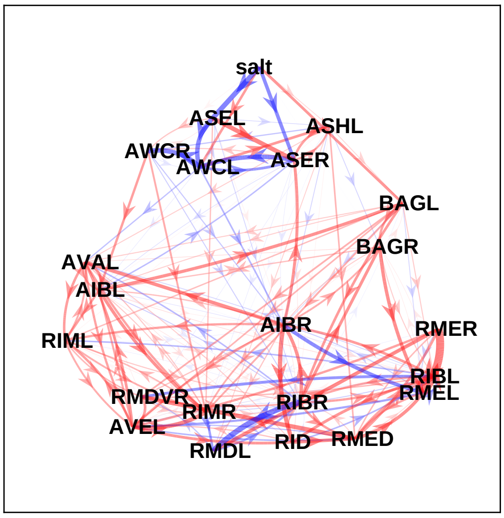


sample 14

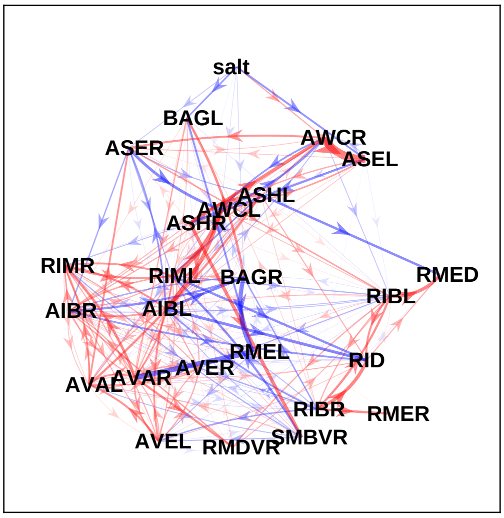


sample 15

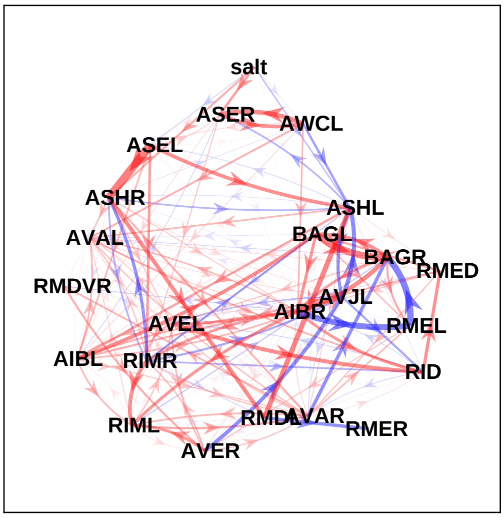


sample 16

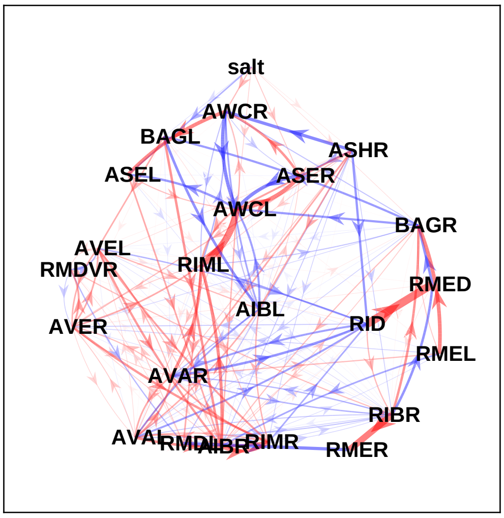


sample 17

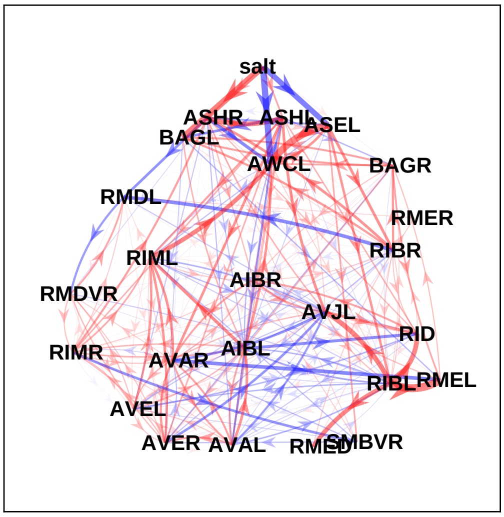


sample 18

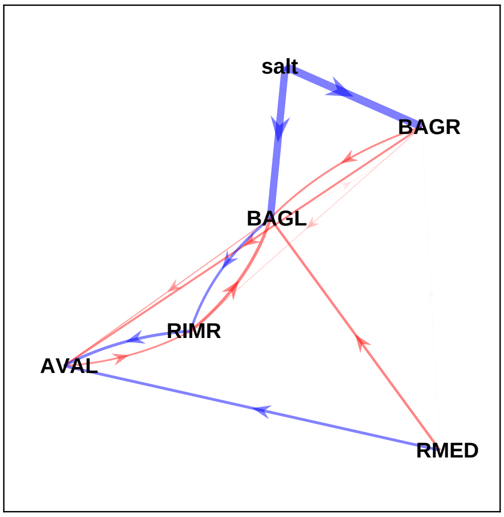

### sample 19

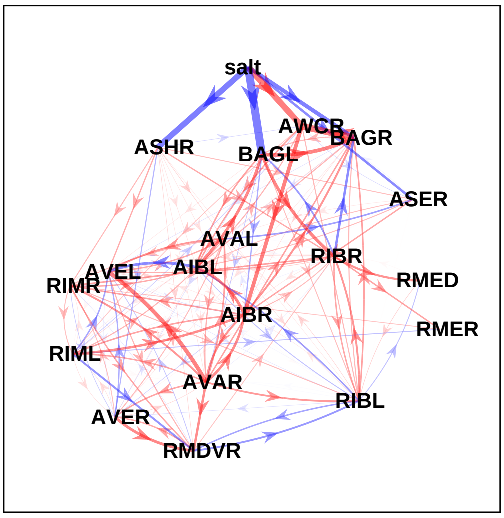


### sample 20

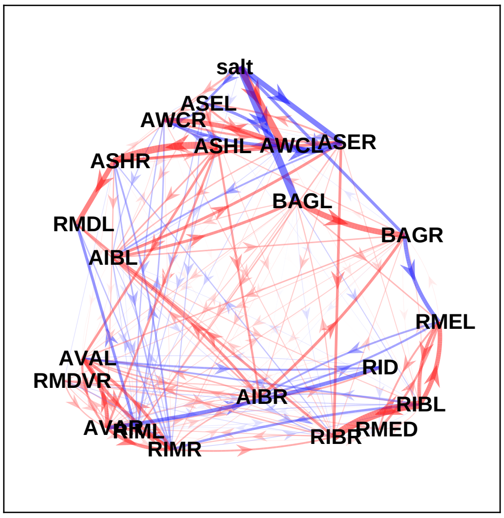


### sample 21

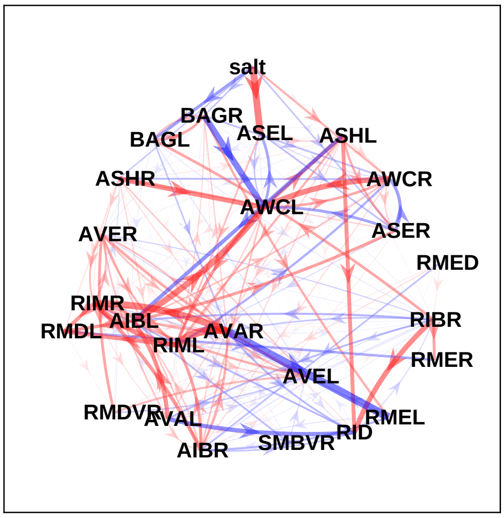


### sample 22

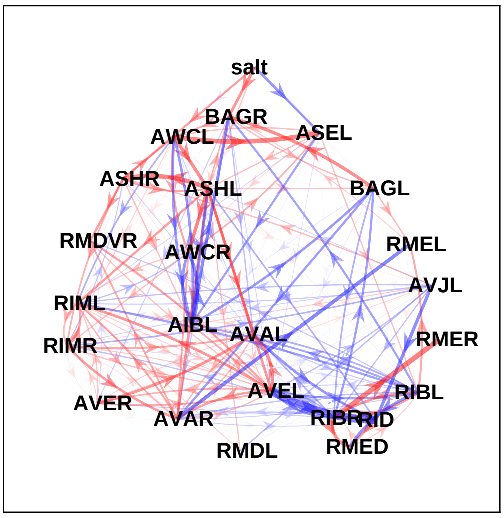


### sample 23

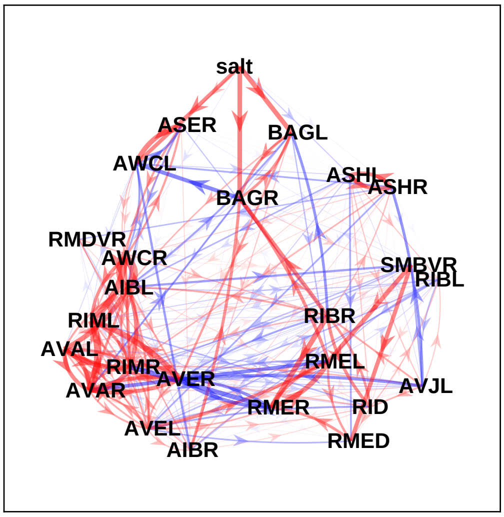


### sample 24

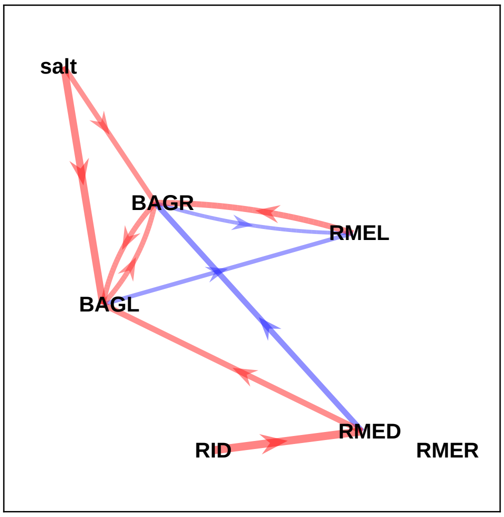
