## Supplementary File 15 for "Deducing ensemble dynamics and information flow from the whole-brain imaging data"

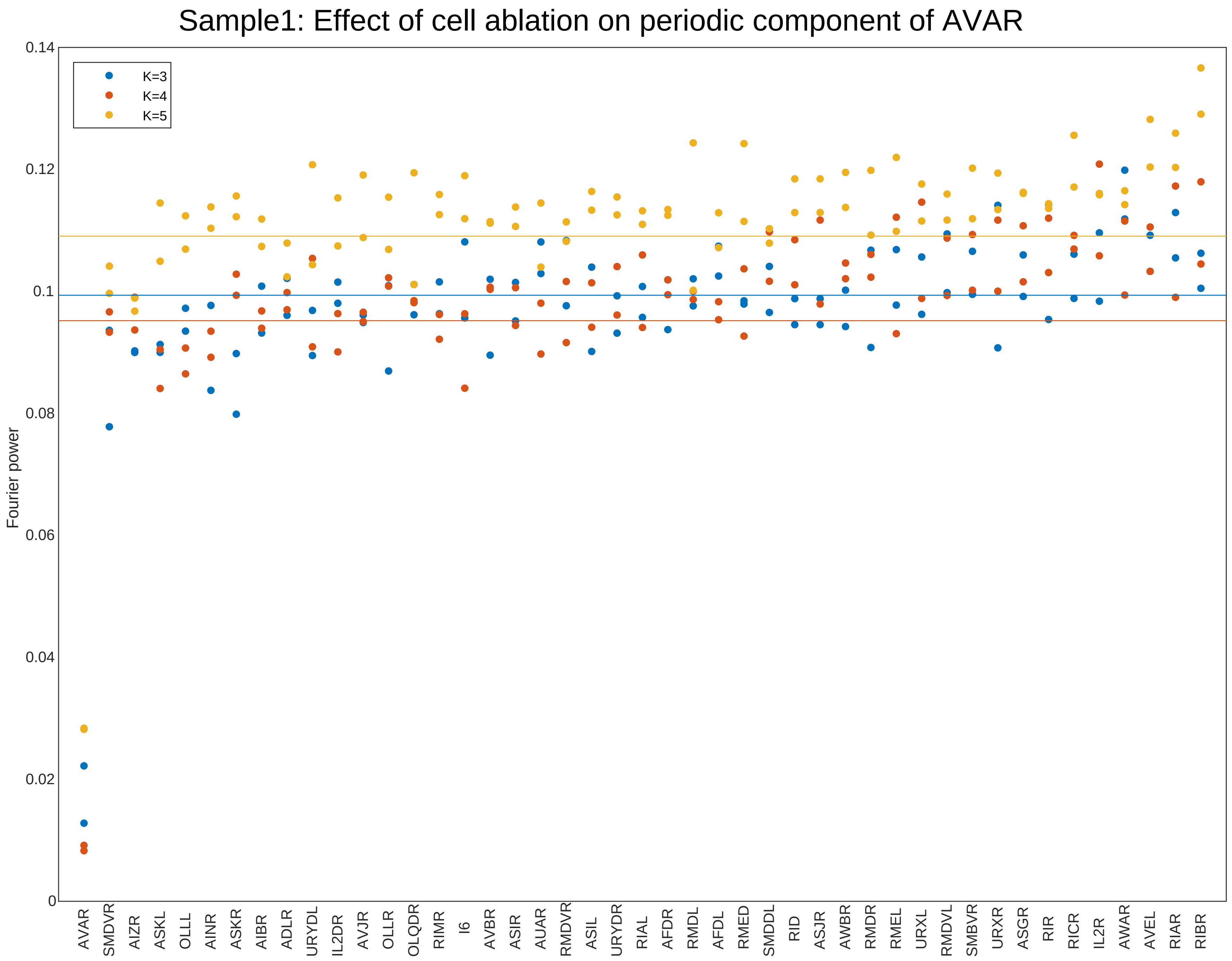

Sample1: Effect of cell ablation on periodic component of AVAR
Fourier power
0.14
0.12
0.1
0.08
0.06
0.04
0.02
0
K=3
K=4
K=5
AVAR SMDVR AIZR ASKL OLLL AINR ASKR AIBR ADLR URYDL IL2DR AVJR OLLR OLQDR RIMR I6 AVBR ASIR AUAR RMDVR ASIL URYDR RIAL AFDR RMDL AFDL RMED SMDDL RID ASJR AWBR RMDR RMEL URXL RMDVL SMBVR URXR ASGR RIR RICR IL2R AWAR AVEL RIAR RIBR

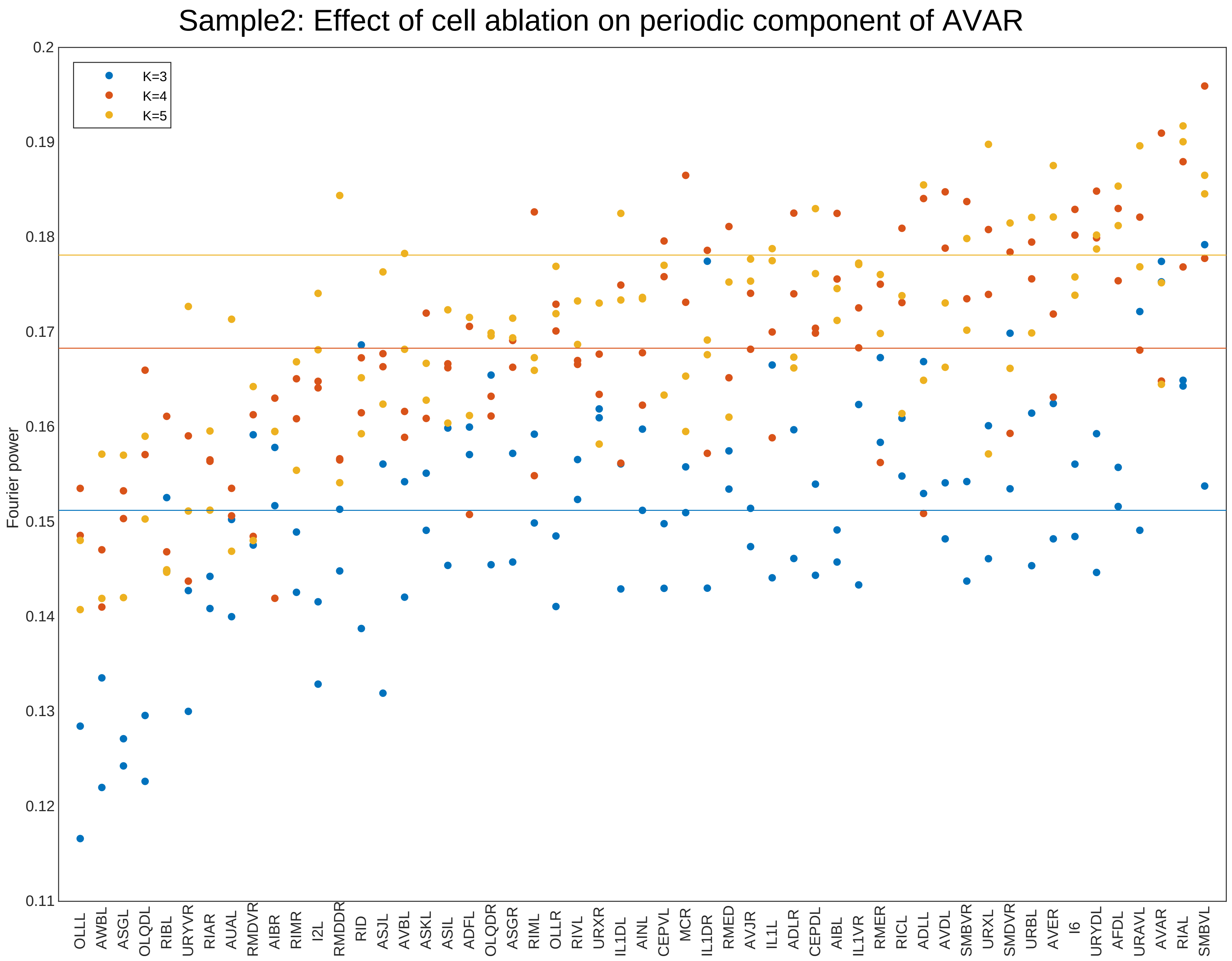
Sample2: Effect of cell ablation on periodic component of AVAR
Fourier power
0.2
0.19
0.18
0.17
0.16
0.15
0.14
0.13
0.12
0.11
K=3
K=4
K=5
OLLL
AWBL
ASGL
OLQDL
RIBL
URYVR
RIAR
AUAL
RMDVR
AIBR
RIMR
I2L
RMDDR
RID
ASJL
AVBL
ASKL
ASIL
ADFL
OLQDR
ASGR
RIML
OLLR
RIVL
URXR
IL1DL
AINL
CEPVL
MCR
IL1DR
RMED
AVJR
IL1L
ADLR
CEPDL
AIBL
IL1VR
RMER
RICL
ADLL
AVDL
SMBVR
URXL
SMDVR
URBL
AVER
I6
URYDL
AFDL
URAVL
AVAR
RIAL
SMBVL

Sample3: Effect of cell ablation on periodic component of AVAR

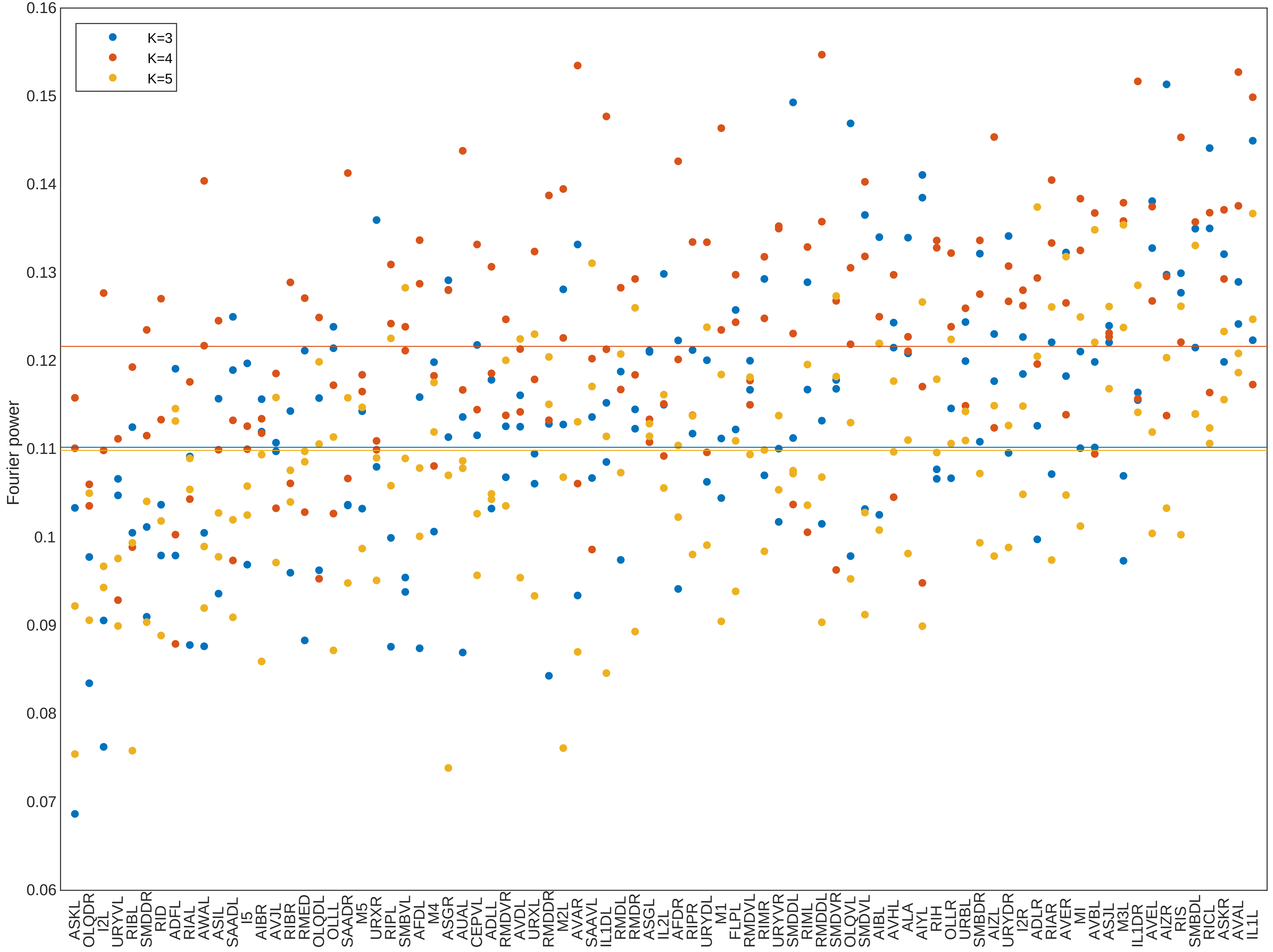

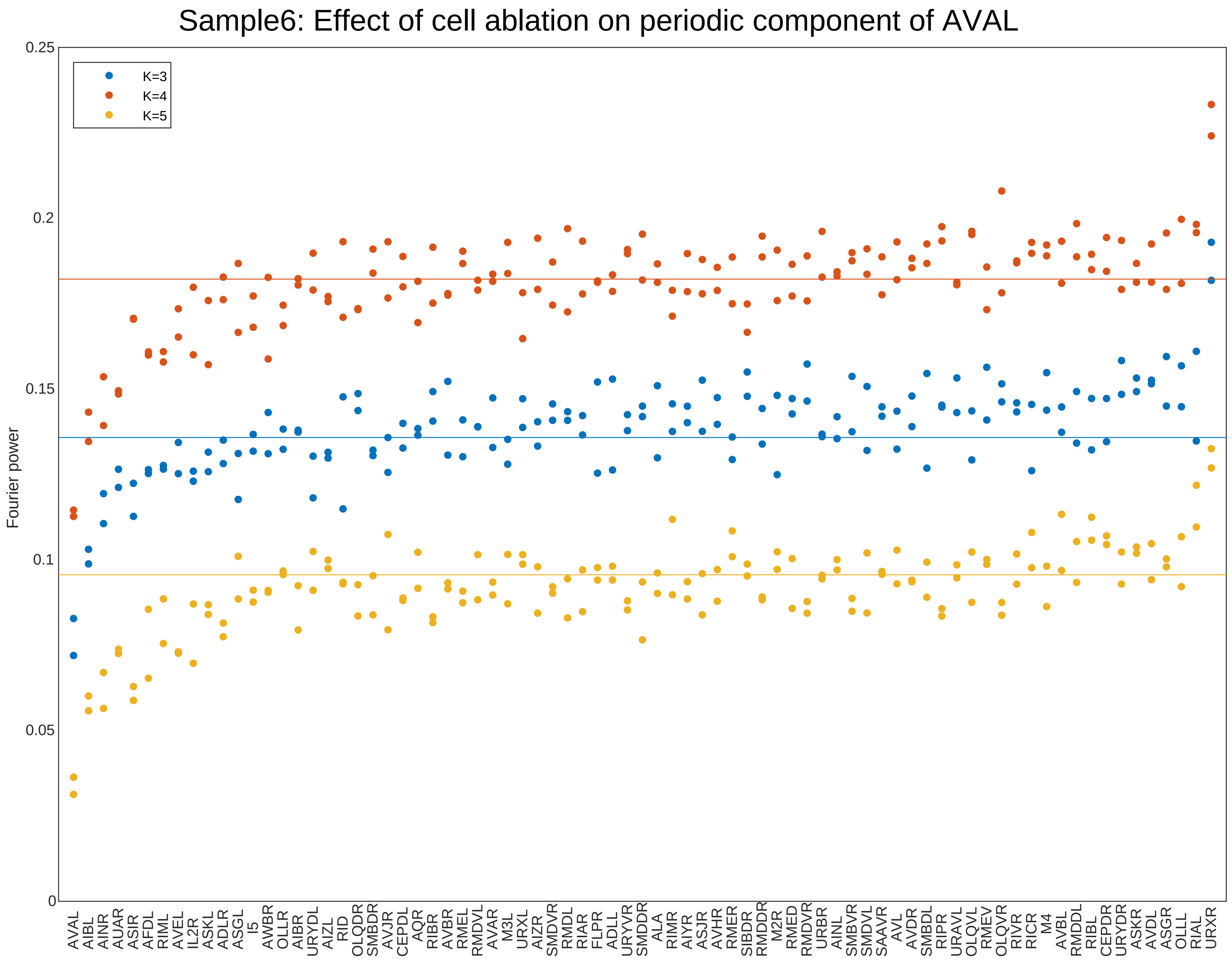

Sample6: Effect of cell ablation on periodic component of AVAL
Fourier power
K=3
K=4
K=5
0
0.05
0.1
0.15
0.2
0.25
AVAL AIBL AINR AUAR ASIR AFDL RIML AVEL IL2R ASKL ADLR ASGL I5 AWBR OLLR AIBR URYDL AIZL RID OLQDR SMBDR AVJR CEPDL AQR RIBR AVBR RMEL RMDVL AVAR M3L URXL AIZR SMDVR RMDL RIAR FLPR ADLL URYVR SMDDR ALA RIMR AIYR ASJR AVHR RMER SIBDR RMDDR M2R RMED RMDVR URBR AINL SMBVR SMDVL SAAVR AVL AVDR SMBDL RIPR URAVL OLQVL RMEV OLQVR RIVR RICR M4 AVBL RMDDL RIBL CEPDR URYDR ASKR AVDL ASGR OLLL RIAL URXR

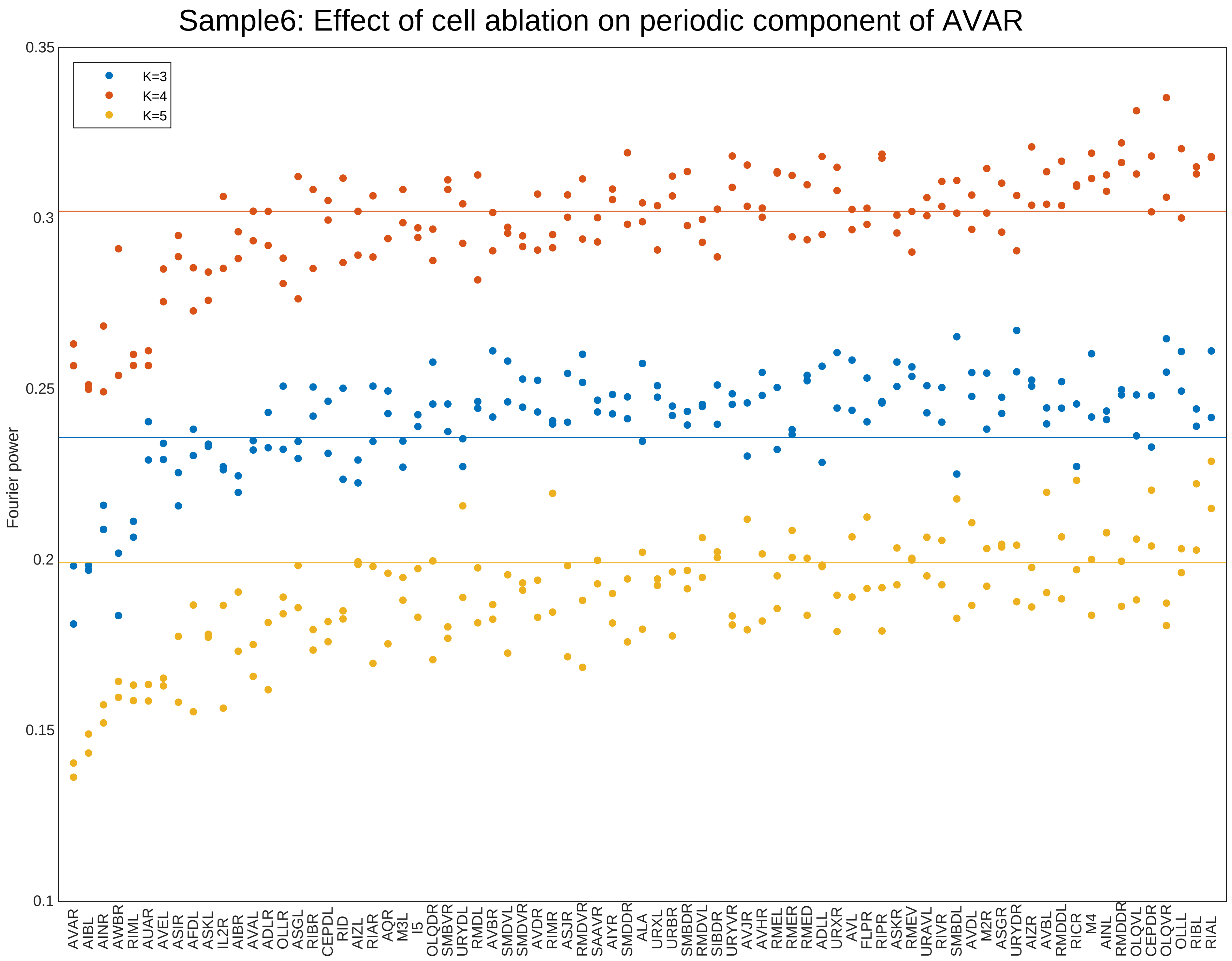

Sample6: Effect of cell ablation on periodic component of AVAR
K=3
K=4
K=5
Fourier power
0.35
0.3
0.25
0.2
0.15
0.1
AVAR
AIBL
AINR
AWBR
RIML
AUAR
AVEL
ASIR
AFDL
ASKL
IL2R
AIBR
AVAL
ADLR
OLLR
ASGL
RIBR
CEPDL
RID
AIZL
RIAR
AQR
M3L
I5
OLQDR
SMBVR
URYDL
RMDL
AVBR
SMDVL
SMDVR
AVDR
RIMR
ASJR
RMDVR
SAAVR
AIYR
SMDDR
ALA
URXL
URBR
SMBDR
RMDVL
SIBDR
URYVR
AVJR
AVHR
RMEL
RMER
RMED
ADLL
URXR
AVL
FLPR
RIPR
ASKR
RMEV
URAVL
RIVR
SMBDL
AVDL
M2R
ASGR
URYDR
AIZR
AVBL
RMDDL
RICR
M4
AINL
RMDDR
OLQVL
CEPDR
OLQVR
OLLL
RIBL
RIAL

### Sample7: Effect of cell ablation on periodic component of AVAL

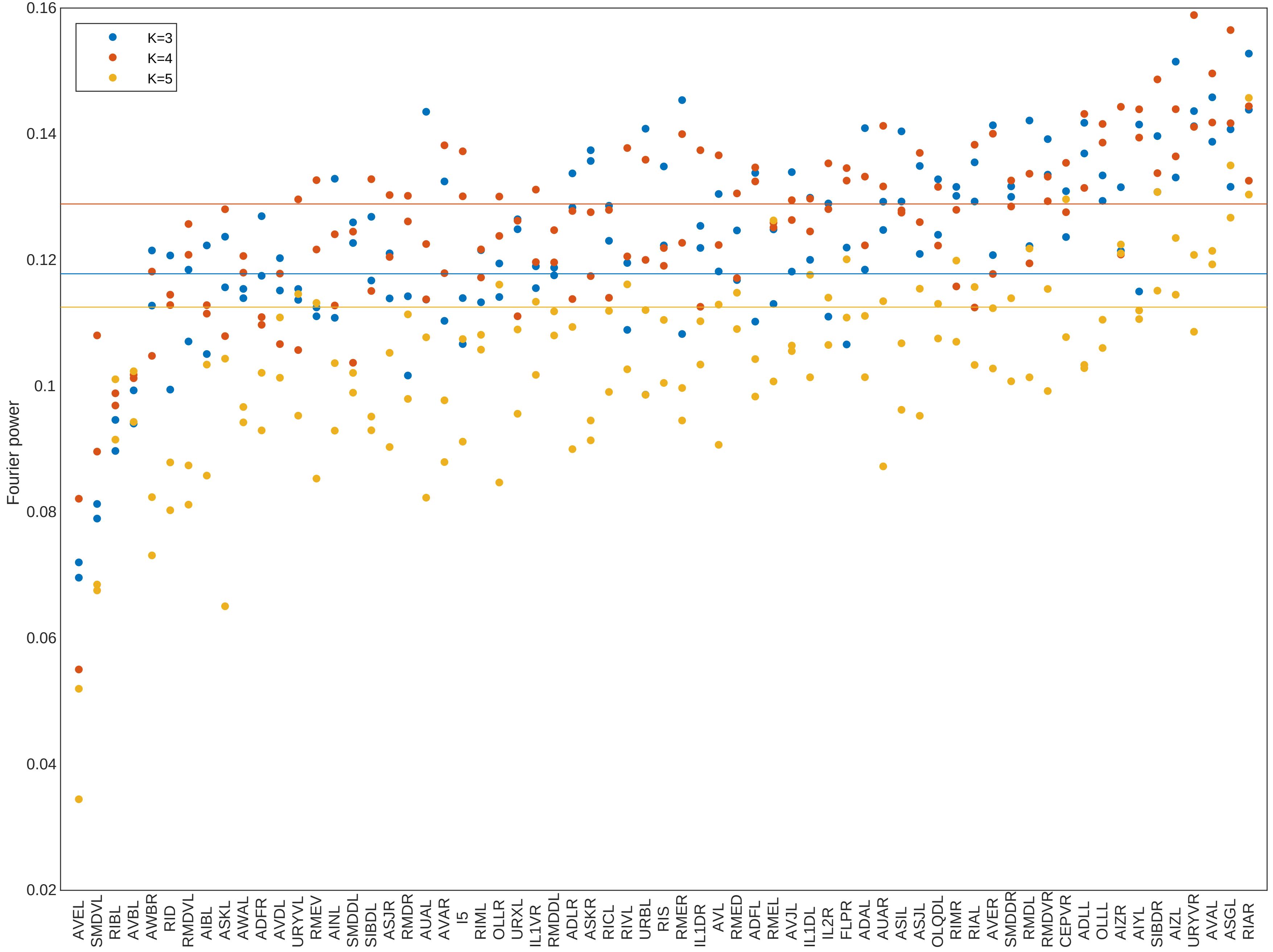

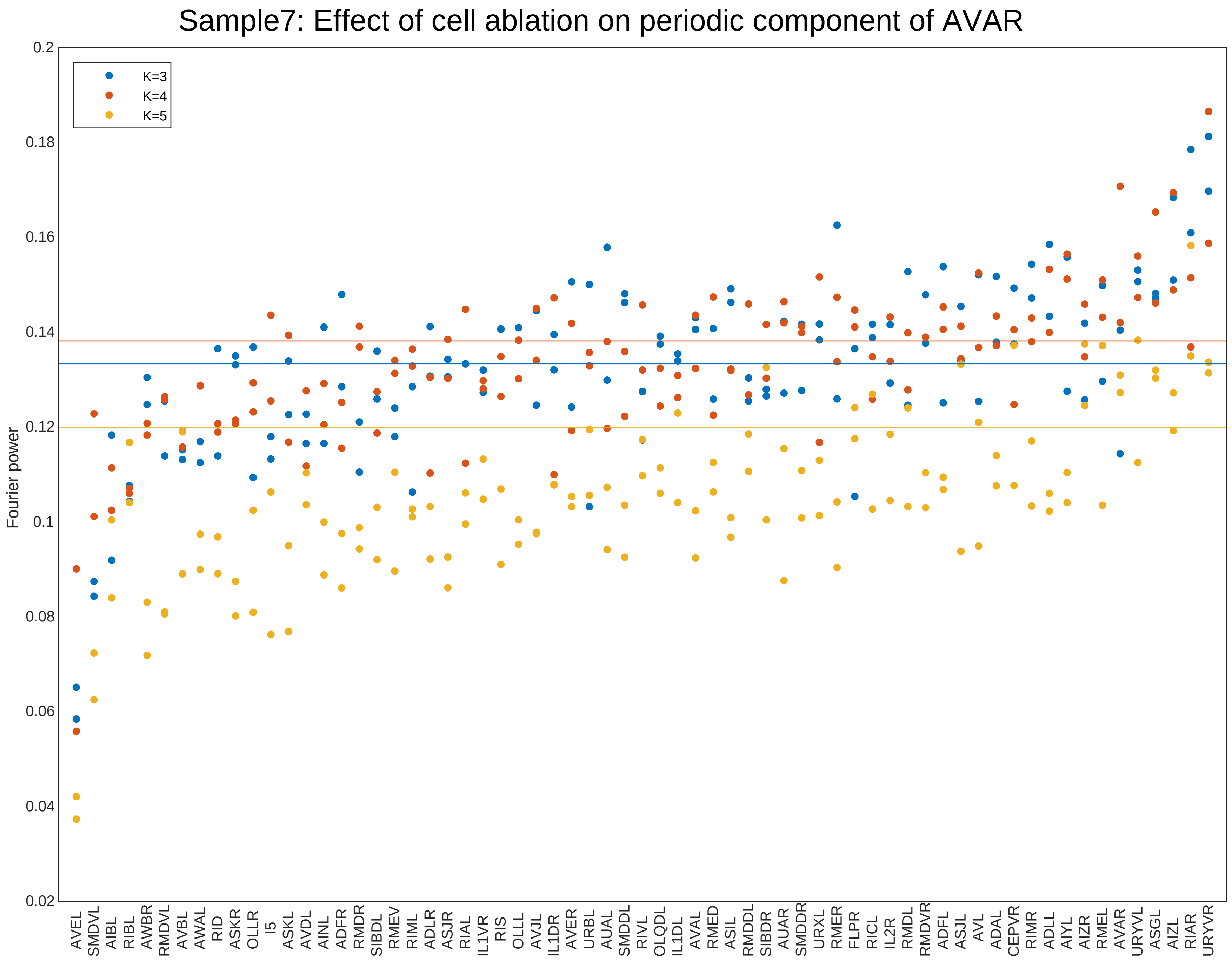
Sample7: Effect of cell ablation on periodic component of AVAR
Fourier power
0.2
0.18
0.16
0.14
0.12
0.1
0.08
0.06
0.04
0.02
K=3
K=4
K=5
AVEL
SMDVL
AIBL
RIBL
AWBR
RMDVL
AVBL
AWAL
RID
ASKR
OLLR
I5
ASKL
AVDL
AINL
ADFR
RMDR
SIBDL
RMEV
RIML
ADLR
ASJR
RIAL
IL1VR
RIS
OLLL
AVJL
IL1DR
AVER
URBL
AUAL
SMDDL
RIVL
OLQDL
IL1DL
AVAL
RMED
ASIL
RMDDL
SIBDR
AUAR
SMDDR
URXL
RMER
FLPR
RICL
IL2R
RMDL
RMDVR
ADFL
ASJL
AVL
ADAL
CEPVR
RIMR
ADLL
AIYL
AIZR
RMEL
AVAR
URYVL
ASGL
AIZL
RIAR
URYVR

Sample21: Effect of cell ablation on periodic component of AVAR

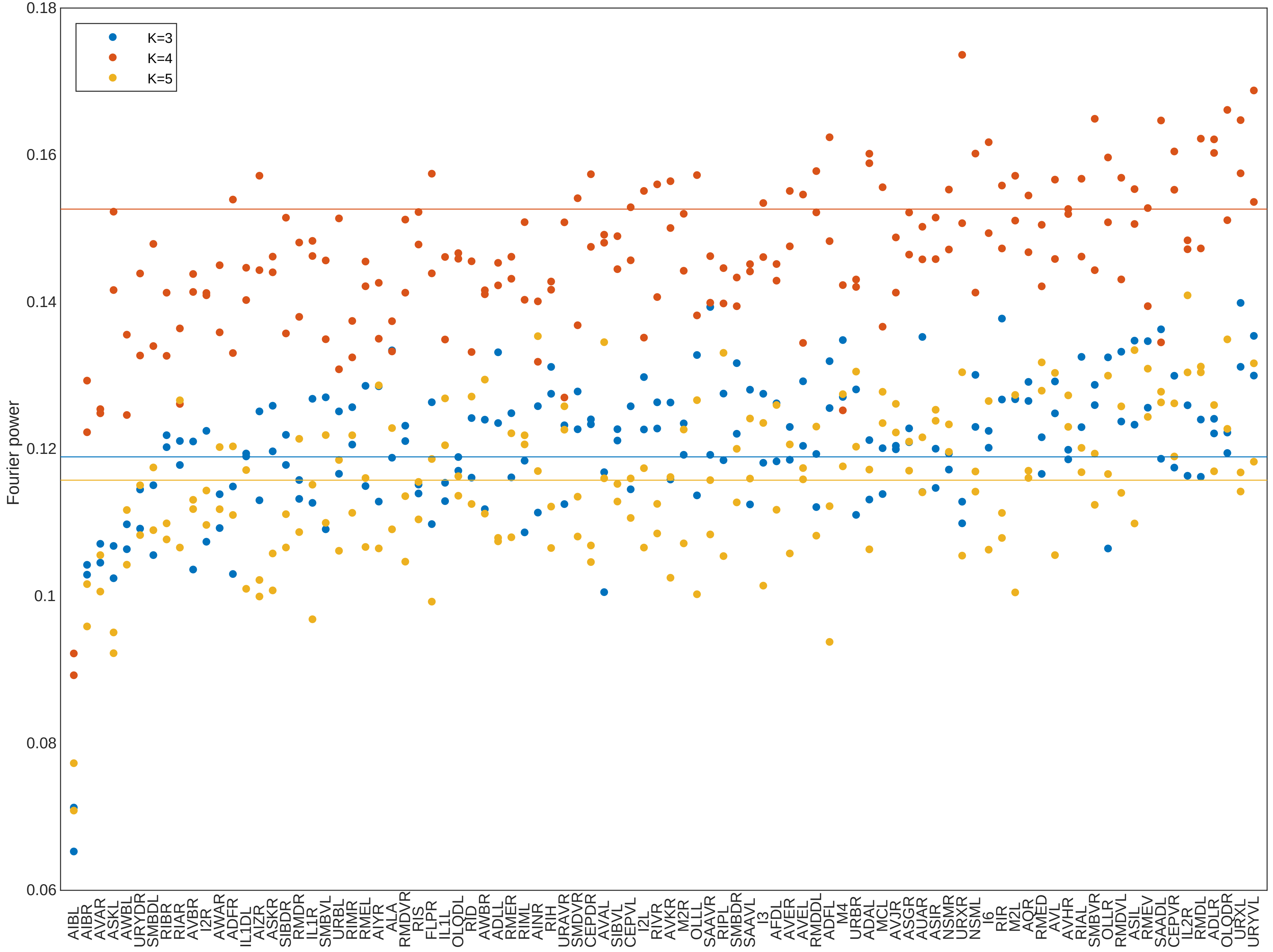
