## Supplementary File 16 for "Deducing ensemble dynamics and information flow from the whole-brain imaging data"

Figure 1

A Example of time series neural activities

Sample 1 Sample 2 Sample 3 Sample 4

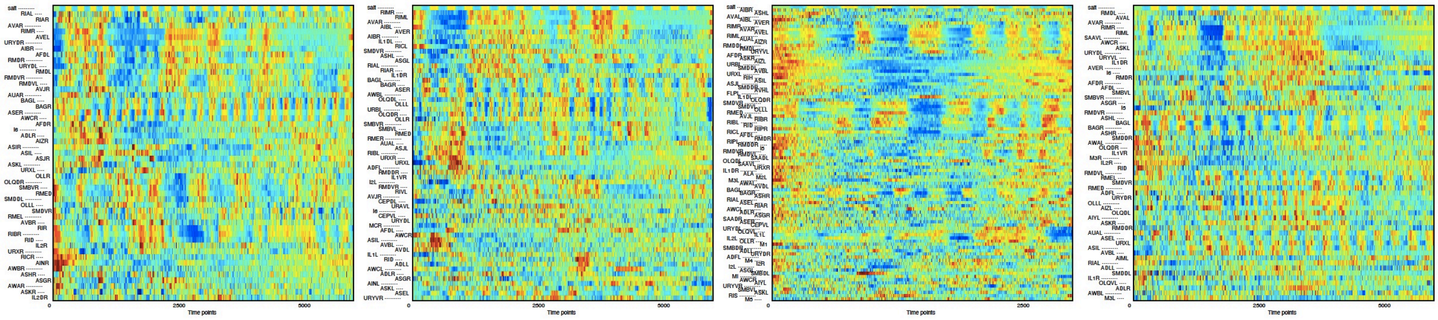

B Example of pairwise cross correlations

Sample 1 Sample 2 Sample 3 Sample 4

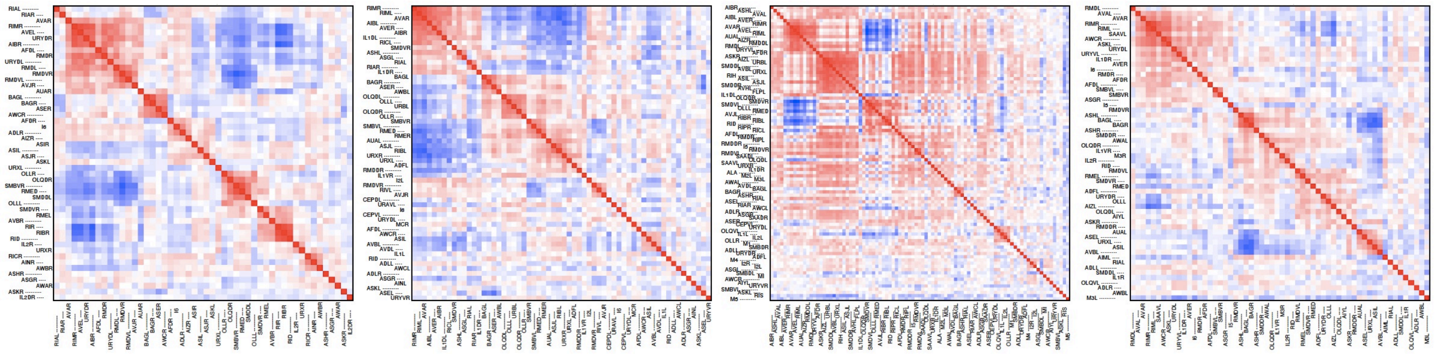

### Figure1-figure supplement 1

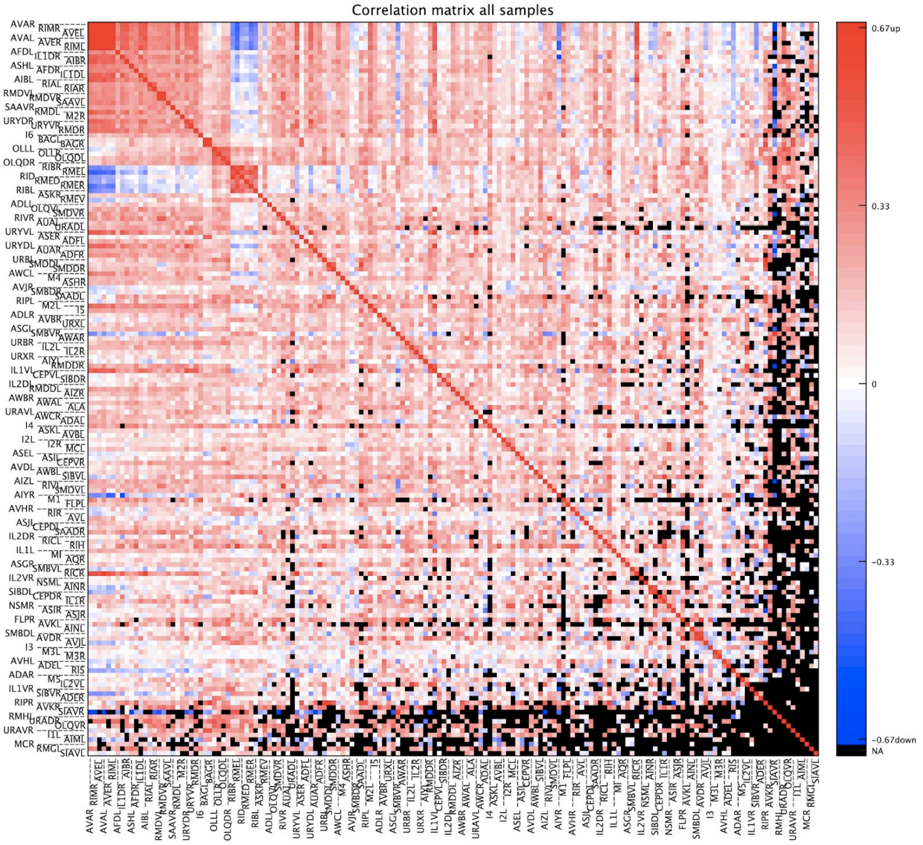

### Figure 7

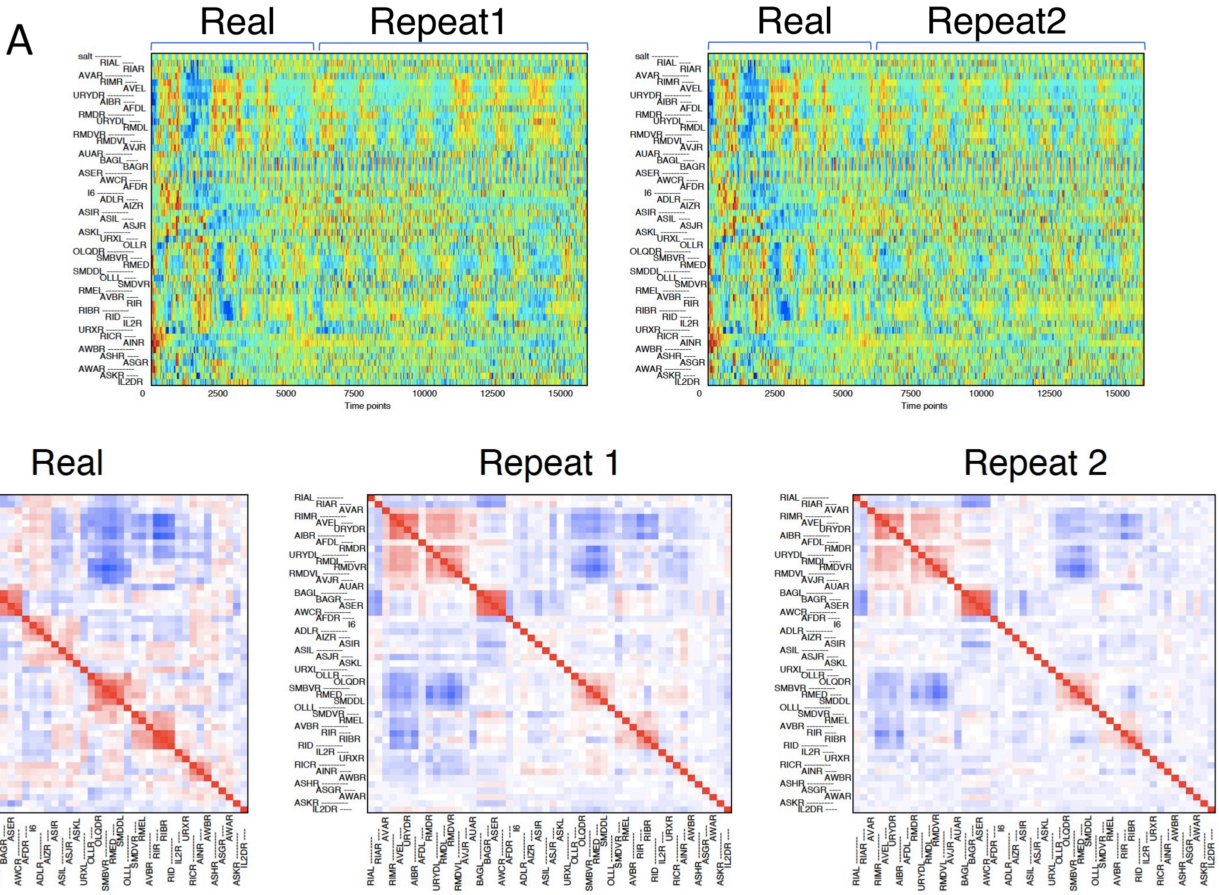

A
Real
Repeat1
Real
Repeat2
Time points
B
Real
Repeat 1
Repeat 2

### Figure 7-Figure supplement 1

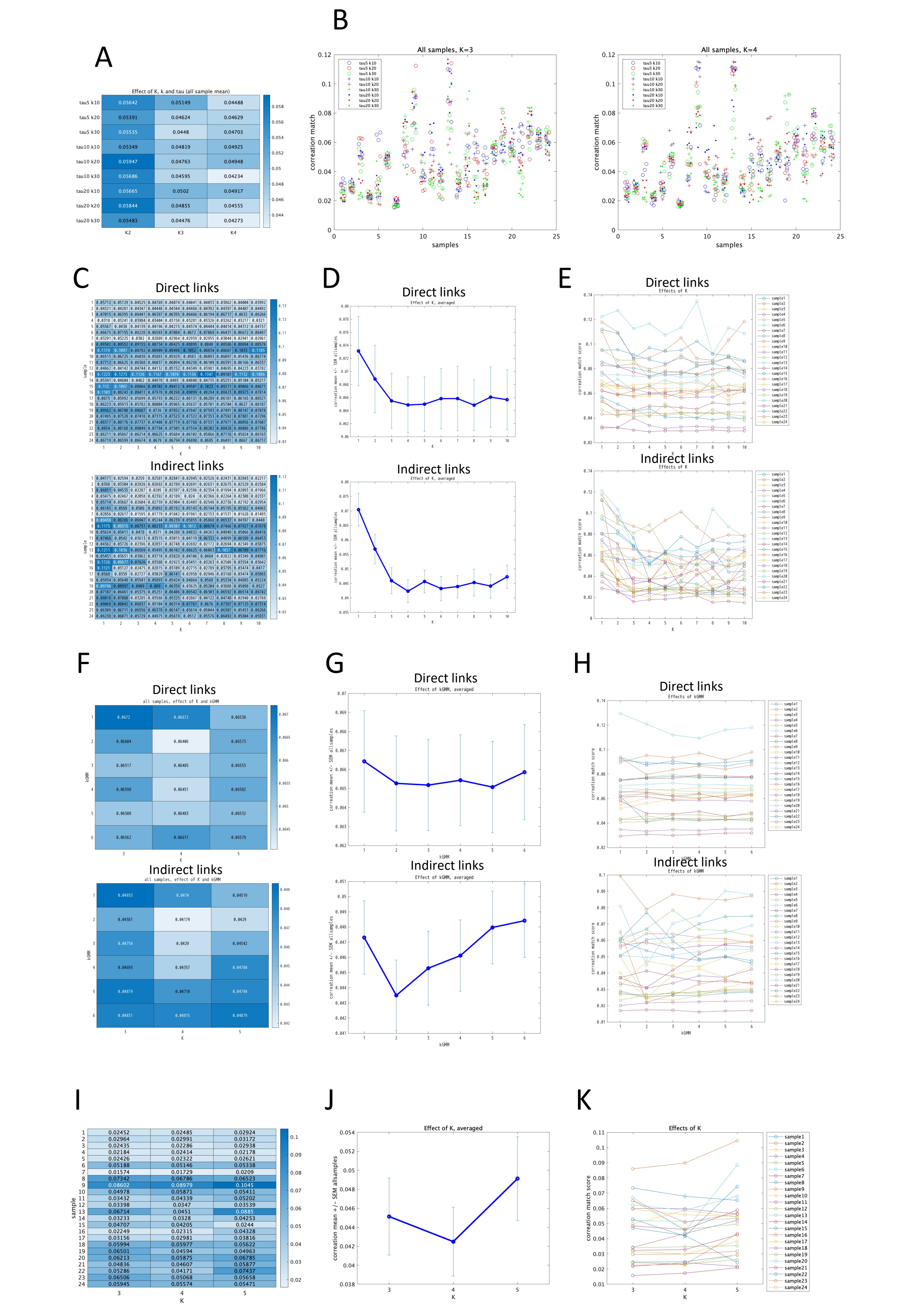

### Figure 7-Figure supplement 2

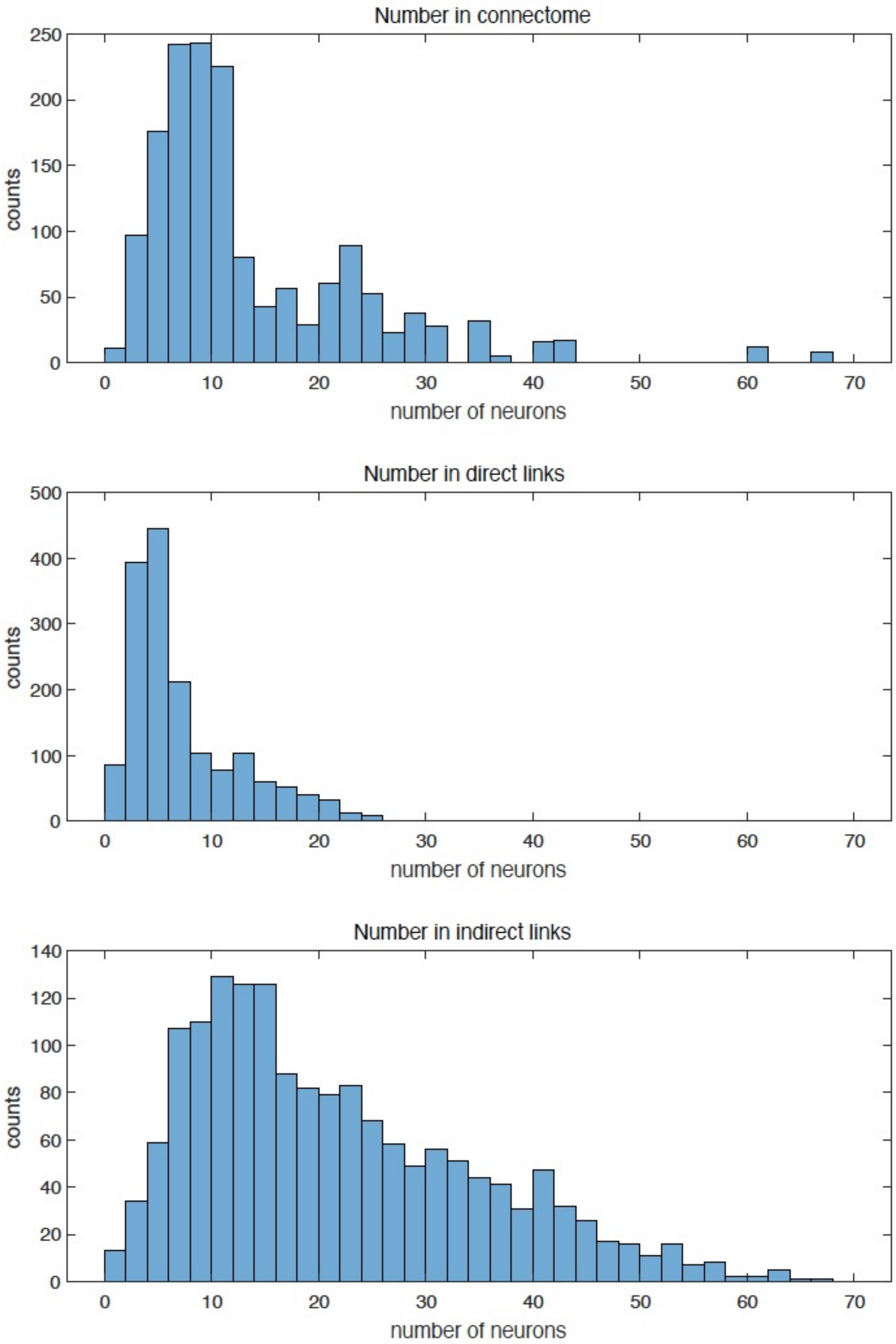

### Figure 8

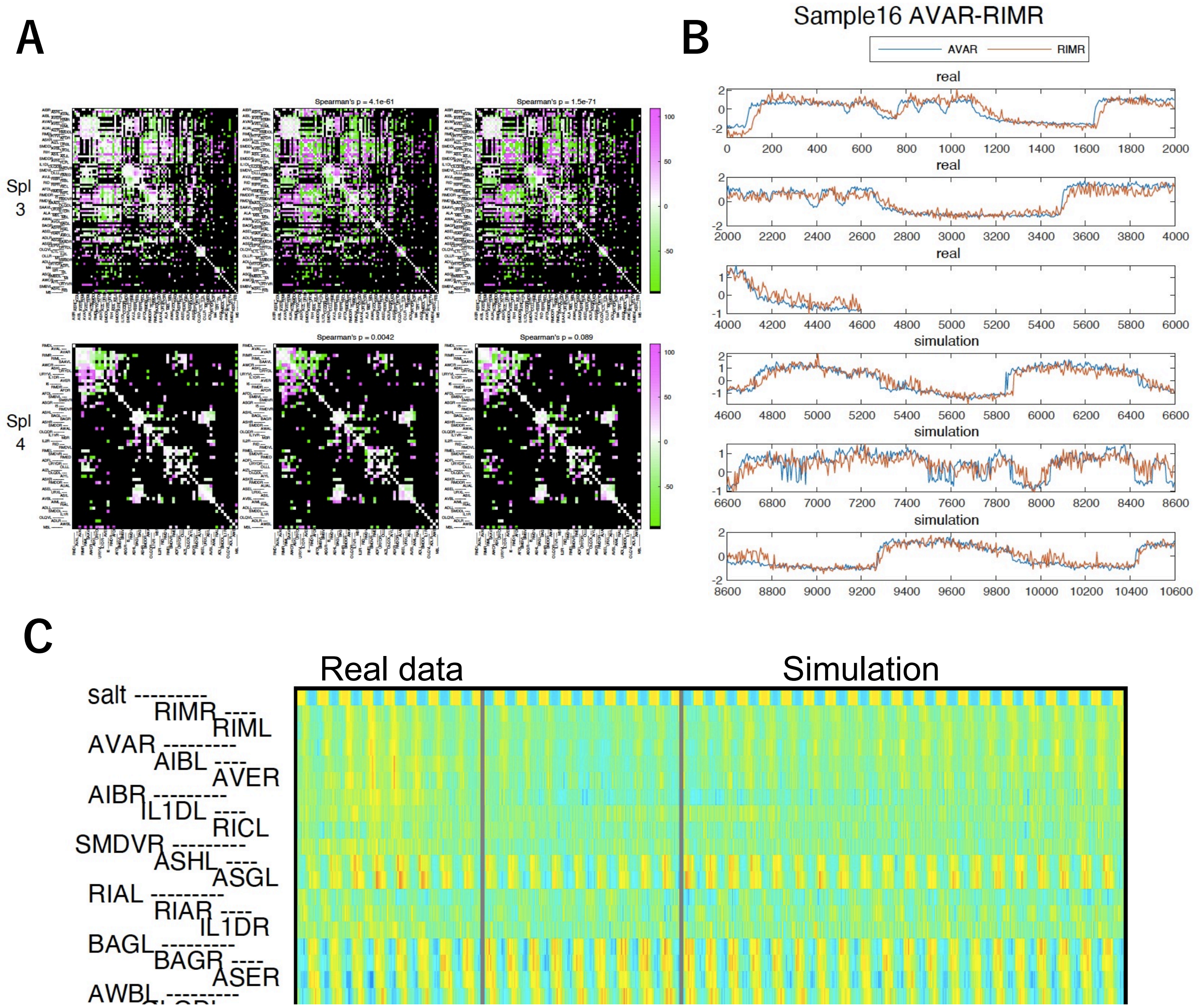
A
B
C
Spl 3
Spl 4
Spearman's p = 4.1e-61
Spearman's p = 1.5e-71
Spearman's p = 0.0042
Spearman's p = 0.089
Sample16 AVAR-RIMR
AVAR
RIMR
real
simulation
Real data
Simulation
salt
RIMR
RIML
AVAR
AIBL
AVER
AIBR
IL1DL
RICL
SMDVR
ASHL
ASGL
RIAL
RIAR
IL1DR
BAGL
BAGR
ASER
AWBL

### Figure 8-Figure Supplement 1

A K=3

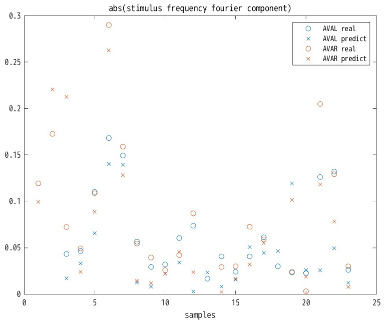


B K=4

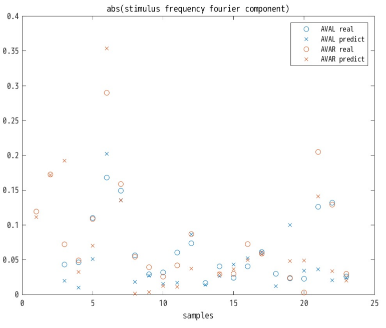


C K=5

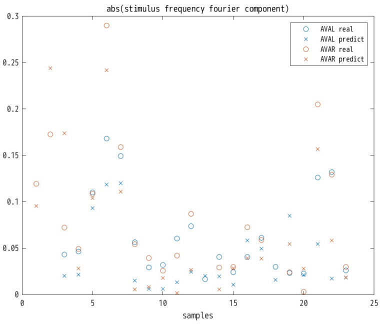

### Figure 9

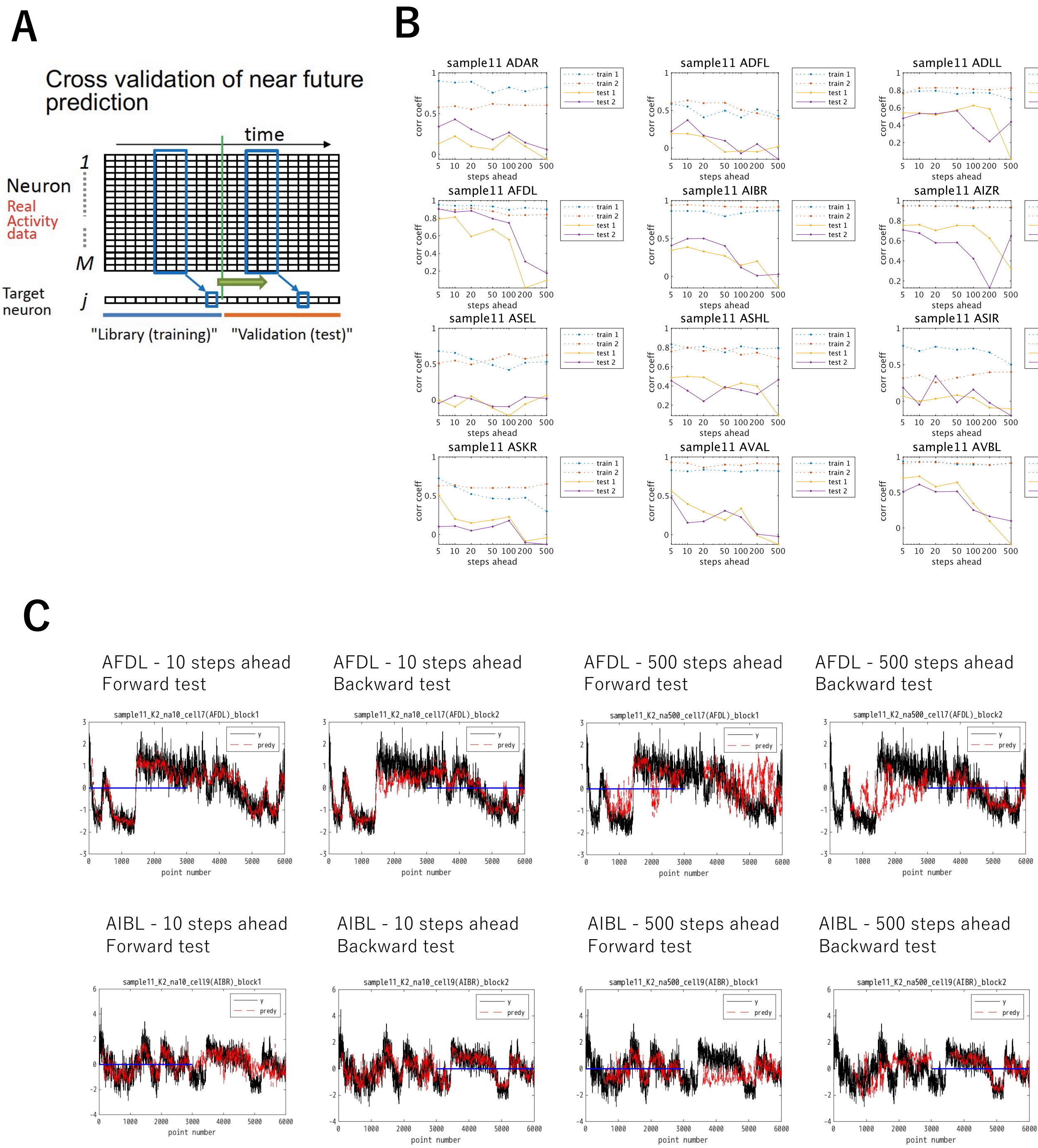
A
B
C
Cross validation of near future prediction
time
Neuron
Real Activity data
1
M
Target neuron
j
"Library (training)"
"Validation (test)"
sample11 ADAR
sample11 ADFL
sample11 ADLL
sample11 AFDL
sample11 AIBR
sample11 AIZR
sample11 ASEL
sample11 ASHL
sample11 ASIR
sample11 ASKR
sample11 AVAL
sample11 AVBL
train 1
train 2
test 1
test 2
corr coeff
steps ahead
AFDL - 10 steps ahead
Forward test
AFDL - 10 steps ahead
Backward test
AFDL - 500 steps ahead
Forward test
AFDL - 500 steps ahead
Backward test
sample11_K2_na10_cell7(AFDL)_block1
sample11_K2_na10_cell7(AFDL)_block2
sample11_K2_na500_cell7(AFDL)_block1
sample11_K2_na500_cell7(AFDL)_block2
AIBL - 10 steps ahead
Forward test
AIBL - 10 steps ahead
Backward test
AIBL - 500 steps ahead
Forward test
AIBL - 500 steps ahead
Backward test
sample11_K2_na10_cell9(AIBR)_block1
sample11_K2_na10_cell9(AIBR)_block2
sample11_K2_na500_cell9(AIBR)_block1
sample11_K2_na500_cell9(AIBR)_block2
y
predy
point number

### Comparison of probabilistic and deterministic models Figure 10

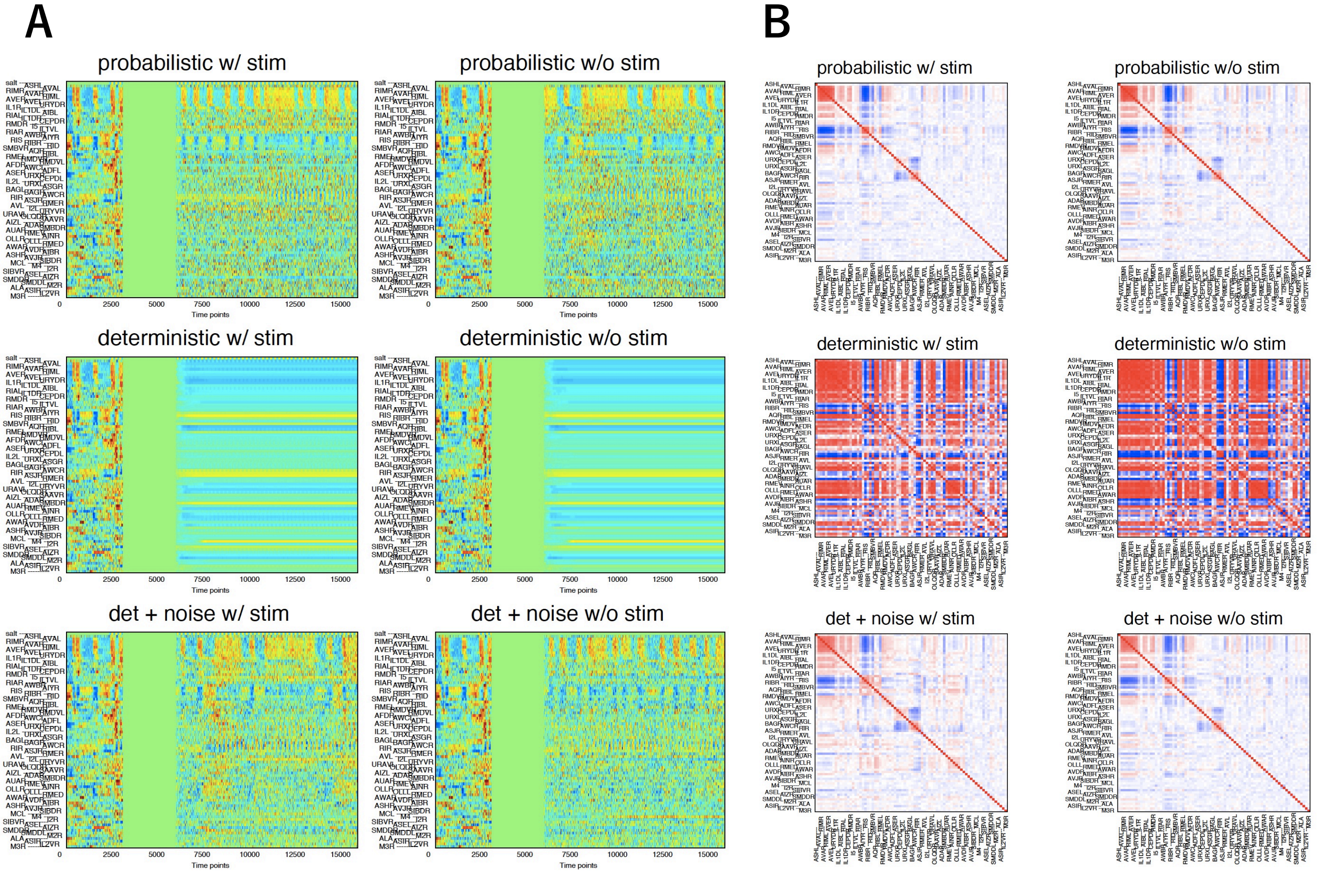

### Figure 11

**A** sample 1 sample 2

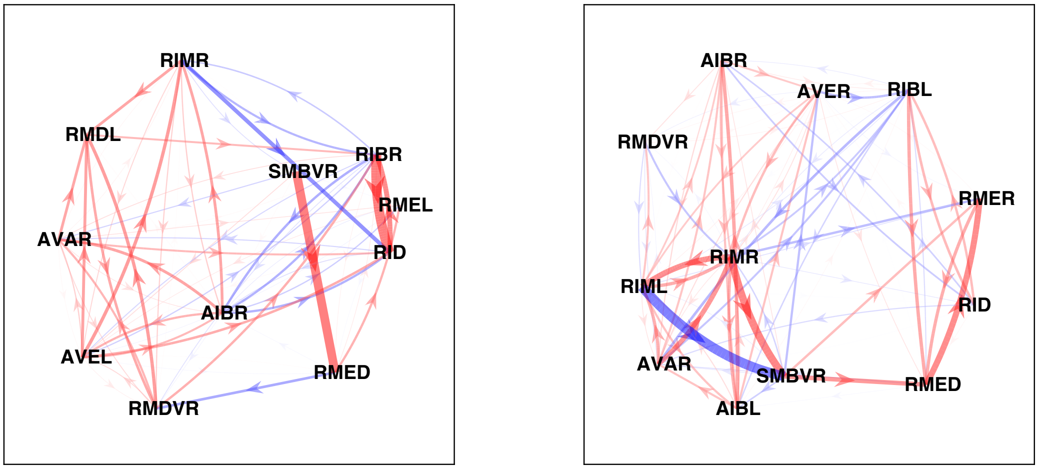


**B** sample 1 sample 2

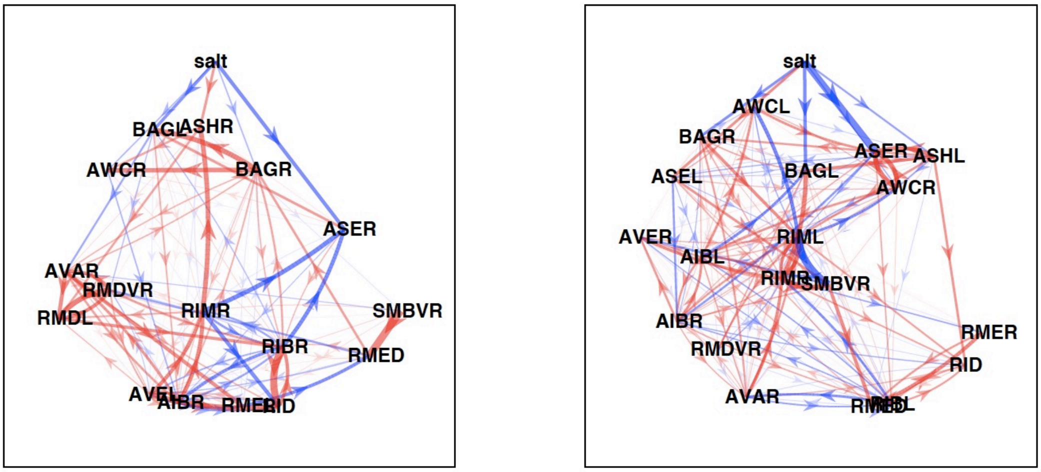

Table 1

**A**

### Sample 1, positive correlation

| intact_r | corr1 | corr1kill_r | corr2 | corr2kill_r | kill1 | kill1_r | kill2 | kill2_r | kill3 | kill3_r | kill4 | kill4_r | kill5 | kill5_r |
|---|---|---|---|---|---|---|---|---|---|---|---|---|---|---|
| 0.661 | AVEL | 0.448 | RIMR | 0.540 | AVEL | 0.448 | RMDL | 0.470 | AIBR | 0.479 | RMDVL | 0.491 | RIR | 0.536 |
| 0.521 | AIBR | 0.362 | RIMR | 0.438 | RMDVL | 0.331 | RMDL | 0.358 | AIBR | 0.362 | AVEL | 0.378 | RIR | 0.423 |
| 0.510 | RMDVL | 0.337 | RMDVR | 0.276 | RMDVR | 0.276 | RMDVL | 0.337 | SMBVR | 0.376 | RMDL | 0.404 | RMED | 0.429 |
| 0.491 | RMED | 0.462 | SMBVR | 0.180 | SMBVR | 0.180 | RMDVL | 0.369 | RIMR | 0.392 | RMDL | 0.396 | RMDR | 0.403 |
| 0.479 | RMDL | 0.292 | RMDR | 0.276 | RMDR | 0.276 | RMDL | 0.292 | RMDVR | 0.333 | AVEL | 0.361 | RMDVL | 0.374 |
| 0.478 | RMDL | 0.288 | RMDVR | 0.218 | RMDVR | 0.218 | RMDVL | 0.238 | RMDL | 0.288 | RMDR | 0.376 | SMBVR | 0.385 |
| 0.466 | RIMR | 0.413 | URYDR | 0.335 | RMDL | 0.258 | RMDVL | 0.267 | RIBR | 0.269 | AIBR | 0.291 | AVEL | 0.292 |
| 0.465 | RIBR | 0.204 | RIR | 0.388 | RIBR | 0.204 | RMDVL | 0.364 | RMDL | 0.365 | RIR | 0.388 | AIBR | 0.407 |
| 0.422 | AIBR | 0.263 | AVEL | 0.244 | RMDVL | 0.217 | RMDL | 0.233 | AVEL | 0.244 | AIBR | 0.263 | RIR | 0.297 |
| 0.408 | RIBR | 0.126 | RID | 0.371 | RIBR | 0.126 | RMDVL | 0.313 | RMDL | 0.359 | RIR | 0.370 | RIAL | 0.371 |
| 0.397 | RMDR | 0.289 | RMDVR | 0.283 | RMDL | 0.074 | RMDVL | 0.232 | AVEL | 0.233 | RIBR | 0.252 | RIMR | 0.267 |
| 0.383 | RIMR | 0.166 | RMDVR | 0.257 | RMDL | 0.112 | RIMR | 0.166 | OLLR | 0.197 | AVEL | 0.200 | RMDVL | 0.209 |
| 0.372 | RIMR | 0.215 | URYDL | 0.334 | AIBR | 0.136 | RMDVL | 0.158 | RMDL | 0.175 | AVEL | 0.187 | RIMR | 0.215 |
| 0.372 | RIMR | 0.143 | RMDL | 0.194 | RIMR | 0.143 | AVEL | 0.173 | RMDL | 0.194 | RMDVL | 0.222 | OLLR | 0.228 |
| 0.362 | RIMR | 0.184 | RMDR | 0.233 | AVEL | 0.156 | RIMR | 0.184 | SMDDL | 0.216 | RMDL | 0.224 | RMDR | 0.233 |
| 0.352 | RMDL | 0.201 | RMDVL | 0.181 | RMDVR | 0.110 | RMDVL | 0.181 | RMDL | 0.201 | SMBVR | 0.237 | RMDR | 0.276 |
| 0.335 | AVEL | 0.187 | URYDR | 0.236 | RMDVL | 0.147 | RMDL | 0.162 | RIBR | 0.173 | AVEL | 0.187 | AIBR | 0.201 |
| 0.324 | BAGL | 0.230 | BAGR | 0.076 | BAGR | 0.076 | BAGL | 0.230 | RIR | 0.241 | AWAR | 0.260 | RIBR | 0.280 |
| 0.316 | AVEL | 0.178 | RMDVR | 0.204 | RMDL | 0.152 | AVEL | 0.178 | OLQDR | 0.184 | RIMR | 0.193 | RMDVL | 0.194 |
| 0.312 | RMED | 0.202 | SMDDL | 0.336 | SMBVR | 0.142 | RMDVL | 0.193 | RMED | 0.202 | RMDVR | 0.224 | RMDL | 0.230 |

### Sample 3, positive correlation

| intact_r | corr1 | corr1kill_r | corr2 | corr2kill_r | kill1 | kill1_r | kill2 | kill2_r | kill3 | kill3_r | kill4 | kill4_r | kill5 | kill5_r |
|---|---|---|---|---|---|---|---|---|---|---|---|---|---|---|
| 0.841 | AVEL | 0.741 | RIML | 0.826 | AVEL | 0.741 | AVAR | 0.792 | RIBL | 0.803 | AUAL | 0.820 | AVER | 0.820 |
| 0.825 | SMDDL | 0.835 | URBL | 0.760 | RIS | 0.393 | OLQVL | 0.530 | RICL | 0.754 | URBL | 0.760 | RIH | 0.769 |
| 0.816 | AVAR | 0.767 | RIML | 0.775 | AVEL | 0.727 | AUAL | 0.767 | AVAR | 0.767 | RIML | 0.775 | RIBL | 0.786 |
| 0.813 | AVBL | 0.793 | SMDDR | 0.816 | RIS | 0.370 | OLQVL | 0.634 | RIH | 0.747 | URBL | 0.751 | RICL | 0.752 |
| 0.800 | AVBL | 0.782 | URBL | 0.821 | RIS | 0.269 | OLQVL | 0.579 | RICL | 0.717 | RIH | 0.740 | ASKR | 0.766 |
| 0.796 | AVAR | 0.736 | AVEL | 0.714 | AVEL | 0.714 | AVAR | 0.736 | RIBL | 0.742 | RIBR | 0.762 | AUAL | 0.764 |
| 0.793 | RIH | 0.705 | URBL | 0.783 | RIS | 0.327 | OLQVL | 0.523 | RIH | 0.705 | FLPL | 0.748 | SAADL | 0.749 |
| 0.783 | AIZL | 0.679 | AVBL | 0.750 | RIS | 0.562 | OLQVL | 0.666 | RIH | 0.678 | AIZL | 0.679 | ASIL | 0.711 |
| 0.773 | AVBL | 0.743 | RIH | 0.723 | RIS | 0.358 | OLQVL | 0.609 | RICL | 0.712 | SAADL | 0.714 | ASKR | 0.718 |
| 0.764 | SMDDR | 0.776 | URBL | 0.730 | RIS | 0.277 | OLQVL | 0.526 | RIH | 0.658 | RICL | 0.707 | FLPL | 0.708 |
| 0.761 | AVBL | 0.740 | SMDDL | 0.767 | RIS | 0.370 | OLQVL | 0.571 | SMBDL | 0.697 | RIH | 0.711 | SAADL | 0.713 |
| 0.749 | AIZL | 0.641 | SMDDR | 0.757 | RIS | 0.359 | OLQVL | 0.496 | RIH | 0.639 | AIZL | 0.641 | ASIL | 0.670 |
| 0.741 | RIH | 0.670 | SMDDL | 0.754 | RIS | 0.313 | OLQVL | 0.489 | RIH | 0.670 | URBL | 0.679 | ASKR | 0.685 |
| 0.726 | AIZL | 0.365 | ASIL | 0.764 | AIZL | 0.365 | RIS | 0.558 | RIH | 0.633 | AVBL | 0.640 | SAADL | 0.649 |
| 0.722 | AIZL | 0.739 | URXL | 0.710 | RIS | 0.441 | RIH | 0.535 | SAADL | 0.599 | AVBL | 0.599 | ASKR | 0.626 |
| 0.722 | RICL | 0.452 | SMDVL | 0.710 | RIS | 0.359 | RICL | 0.452 | ASIL | 0.652 | ASKR | 0.663 | ADLR | 0.667 |
| 0.721 | SMDDL | 0.719 | SMDDR | 0.729 | RIS | 0.290 | OLQVL | 0.478 | RIH | 0.633 | URBL | 0.636 | SMBDL | 0.644 |
| 0.719 | RIH | 0.671 | SMDDR | 0.731 | RIS | 0.240 | OLQVL | 0.494 | URBL | 0.660 | RICL | 0.663 | ASKR | 0.663 |
| 0.717 | RIBL | 0.618 | RID | 0.701 | RIBL | 0.618 | RIS | 0.668 | RIBR | 0.681 | OLQVL | 0.690 | ASIL | 0.693 |
| 0.704 | AIZL | 0.669 | URBL | 0.750 | RIS | 0.305 | OLQVL | 0.475 | RIH | 0.500 | FLPL | 0.550 | ASIL | 0.583 |

**B**

### Sample 1, negative correlation

| intact_r | corr1 | corr1kill_r | corr2 | corr2kill_r | kill1 | kill1_r | kill2 | kill2_r | kill3 | kill3_r | kill4 | kill4_r | kill5 | kill5_r |
|---|---|---|---|---|---|---|---|---|---|---|---|---|---|---|
| -0.492 | RMDVR | -0.403 | RMED | -0.379 | SMBVR | -0.275 | RMDVL | -0.347 | RMDL | -0.352 | RMED | -0.379 | RMDVR | -0.403 |
| -0.449 | RMDVR | -0.376 | SMBVR | -0.322 | RMDVL | -0.234 | SMBVR | -0.322 | RMDL | -0.345 | ADLR | -0.373 | RMDVR | -0.376 |
| -0.391 | RIMR | -0.389 | RIR | -0.195 | RIR | -0.195 | AIBR | -0.208 | RMDVL | -0.230 | RIAL | -0.306 | RMDL | -0.309 |
| -0.377 | AIBR | -0.237 | RIR | -0.193 | RIR | -0.193 | RMDVL | -0.223 | AIBR | -0.237 | RMDL | -0.243 | AVEL | -0.266 |
| -0.361 | AVEL | -0.245 | RIR | -0.184 | AIBR | -0.154 | RIR | -0.184 | RMDVL | -0.187 | RMDL | -0.230 | AVEL | -0.245 |
| -0.351 | RMDR | -0.211 | RMED | -0.319 | RMDL | -0.149 | AVEL | -0.155 | RMDVL | -0.197 | RMDVR | -0.208 | RMDR | -0.211 |
| -0.341 | RMDVL | -0.149 | SMBVR | -0.186 | RMDVL | -0.149 | RMDVR | -0.176 | SMBVR | -0.186 | RMDL | -0.223 | RIBR | -0.274 |
| -0.328 | RMDVL | -0.175 | RMED | -0.274 | SMBVR | -0.089 | RMDVR | -0.163 | RMDVL | -0.175 | RMDL | -0.194 | AVBR | -0.258 |
| -0.323 | RMDL | -0.189 | RMED | -0.272 | RMDVR | -0.136 | RMDVL | -0.139 | SMBVR | -0.152 | RMDL | -0.189 | AVEL | -0.225 |
| -0.318 | RMDVR | -0.188 | SMDDL | -0.340 | RMDVL | -0.151 | RMDVR | -0.188 | SMBVR | -0.194 | RMED | -0.214 | RMDL | -0.223 |
| -0.309 | OLQDR | -0.255 | RMDVR | -0.204 | RMED | -0.170 | RMDVR | -0.204 | RMDVL | -0.224 | SMBVR | -0.227 | OLQDR | -0.255 |
| -0.300 | OLLR | -0.267 | RMDVR | -0.229 | SMBVR | -0.148 | RMDVL | -0.208 | OLQDR | -0.210 | RMDL | -0.229 | RMDVR | -0.229 |
| -0.292 | RMDL | -0.199 | SMBVR | -0.195 | RMDVL | -0.102 | RMDVR | -0.142 | SMBVR | -0.195 | RMDL | -0.199 | RMDR | -0.214 |
| -0.274 | RMDR | -0.135 | SMBVR | -0.197 | RMDL | -0.124 | RMDVL | -0.129 | RMDR | -0.135 | AVEL | -0.140 | RIMR | -0.149 |
| -0.258 | RIAR | -0.259 | RMDR | -0.163 | RMDVL | -0.150 | RMDL | -0.160 | RMDR | -0.163 | BAGR | -0.180 | AVEL | -0.181 |
| -0.251 | RIMR | -0.095 | RMED | -0.212 | RIMR | -0.095 | AVEL | -0.113 | RMDVL | -0.130 | RMDL | -0.138 | OLQDR | -0.143 |
| -0.247 | AUAR | -0.243 | RIR | -0.072 | RIR | -0.072 | RMDL | -0.117 | RMDVL | -0.131 | RIBR | -0.148 | AVEL | -0.193 |
| -0.245 | OLLR | -0.171 | RIMR | -0.159 | RMDL | -0.065 | OLQDR | -0.148 | RIMR | -0.159 | RMDVL | -0.165 | RIBR | -0.167 |
| -0.243 | AIBR | -0.101 | RIBR | 0.008 | RIBR | 0.008 | AIBR | -0.101 | RIR | -0.132 | AVEL | -0.192 | RMEL | -0.201 |
| -0.230 | RIMR | -0.022 | SMBVR | -0.188 | RIMR | -0.022 | RMDVL | -0.104 | RMDL | -0.107 | OLLR | -0.122 | AVEL | -0.125 |

### Sample 3, negative correlation

| intact_r | corr1 | corr1kill_r | corr2 | corr2kill_r | kill1 | kill1_r | kill2 | kill2_r | kill3 | kill3_r | kill4 | kill4_r | kill5 | kill5_r |
|---|---|---|---|---|---|---|---|---|---|---|---|---|---|---|
| -0.719 | RIML | -0.698 | SMDVR | -0.684 | RIBL | -0.574 | OLLL | -0.653 | AVEL | -0.657 | AVAR | -0.663 | RIMR | -0.681 |
| -0.692 | AVEL | -0.657 | SMDVR | -0.682 | RIBL | -0.535 | AVAR | -0.619 | OLLL | -0.622 | AVEL | -0.657 | RIML | -0.664 |
| -0.661 | RIS | -0.298 | URBL | -0.583 | RIS | -0.298 | OLQVL | -0.330 | SMDDL | -0.563 | RICL | -0.567 | URBL | -0.583 |
| -0.641 | RIML | -0.598 | RMED | -0.630 | AVEL | -0.419 | OLLL | -0.450 | AVAR | -0.529 | AVER | -0.560 | AVBL | -0.563 |
| -0.613 | OLQVL | -0.474 | RICL | -0.602 | RIS | -0.131 | OLQVL | -0.474 | ASKR | -0.478 | AVBL | -0.495 | URXL | -0.498 |
| -0.611 | AVBL | -0.657 | RIS | -0.243 | RIS | -0.243 | OLQVL | -0.416 | SMDDL | -0.488 | RICL | -0.541 | SMBDL | -0.557 |
| -0.610 | AVEL | -0.567 | RIBL | -0.517 | RIBL | -0.517 | AVEL | -0.567 | RIBR | -0.578 | RICL | -0.581 | SAADR | -0.581 |
| -0.606 | RIS | -0.235 | SMDDL | -0.516 | RIS | -0.235 | OLQVL | -0.388 | RICL | -0.475 | SMDDL | -0.516 | SMBDL | -0.524 |
| -0.602 | RIML | -0.517 | SMDVL | -0.658 | SMDVR | -0.460 | RIBL | -0.472 | AVEL | -0.490 | OLLL | -0.516 | RIML | -0.517 |
| -0.599 | RMDVL | -0.583 | RMED | -0.580 | SMBDL | -0.431 | OLLL | -0.463 | AVEL | -0.512 | SMDVL | -0.526 | AVAR | -0.531 |
| -0.597 | AVAR | -0.515 | SMDVR | -0.591 | RIBL | -0.438 | AVAR | -0.515 | AVEL | -0.516 | RIBR | -0.541 | RIMR | -0.542 |
| -0.592 | AVEL | -0.552 | SMDVL | -0.621 | RIBL | -0.486 | SMDVR | -0.494 | OLLL | -0.513 | RIBR | -0.521 | RIML | -0.530 |
| -0.590 | AVAR | -0.534 | RMED | -0.593 | AVEL | -0.419 | OLLL | -0.444 | AUAL | -0.503 | AVBL | -0.521 | RIML | -0.531 |
| -0.589 | SMDVL | -0.620 | URYVL | -0.592 | RIBL | -0.440 | AVAR | -0.456 | RIS | -0.456 | AVEL | -0.517 | SMDVR | -0.525 |
| -0.572 | RIH | -0.584 | RIS | -0.132 | RIS | -0.132 | OLQVL | -0.247 | SMDDL | -0.479 | RICL | -0.494 | ASIL | -0.495 |
| -0.557 | AVEL | -0.357 | RMED | -0.545 | AVEL | -0.357 | OLLL | -0.408 | AVAR | -0.457 | AVER | -0.484 | AVBL | -0.505 |
| -0.546 | RIS | -0.195 | SMDDR | -0.506 | RIS | -0.195 | OLQVL | -0.337 | SMDDL | -0.419 | SMBDL | -0.439 | URBL | -0.474 |
| -0.535 | AVER | -0.546 | RIBL | -0.467 | RIS | -0.434 | RIBL | -0.467 | AVBL | -0.484 | OLQVL | -0.485 | AVAR | -0.490 |
| -0.526 | RIBL | -0.476 | RIML | -0.502 | RICL | -0.430 | AVEL | -0.443 | RIBL | -0.476 | RIS | -0.486 | AVER | -0.487 |
| -0.520 | RMED | -0.400 | URYVL | -0.556 | AVAR | -0.280 | AVEL | -0.381 | RMED | -0.400 | OLLL | -0.415 | SMBDL | -0.434 |

### Figure 12

**A**

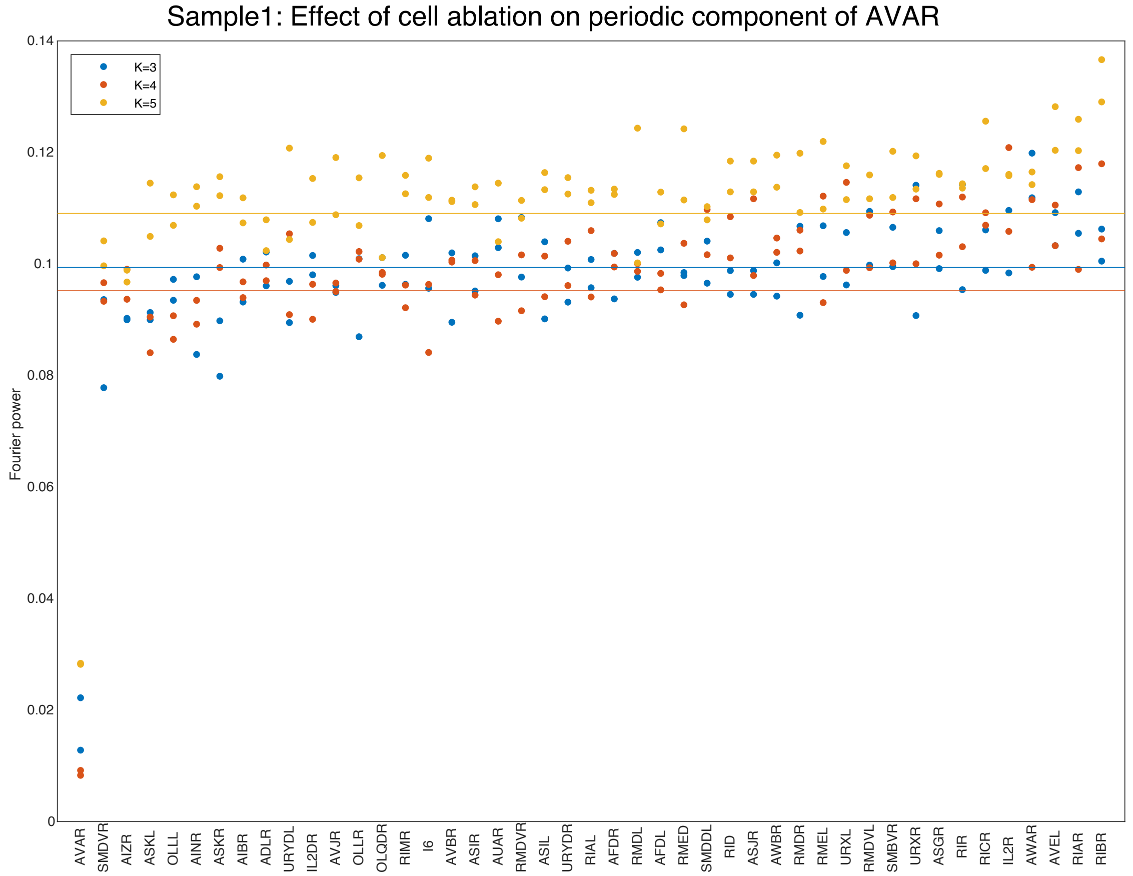

Sample1: Effect of cell ablation on periodic component of AVAR
K=3
K=4
K=5
Fourier power
0.14
0.12
0.1
0.08
0.06
0.04
0.02
0
AVAR
SMDVR
AIZR
ASKL
OLLL
AINR
ASKR
AIBR
ADLR
URYDL
IL2DR
AVJR
OLLR
OLQDR
RIMR
I6
AVBR
ASIR
AUAR
RMDVR
ASIL
URYDR
RIAL
AFDR
RMDL
AFDL
RMED
SMDDL
RID
ASJR
AWBR
RMDR
RMEL
URXL
RMDVL
SMBVR
URXR
ASGR
RIR
RICR
IL2R
AWAR
AVEL
RIAR
RIBR

B

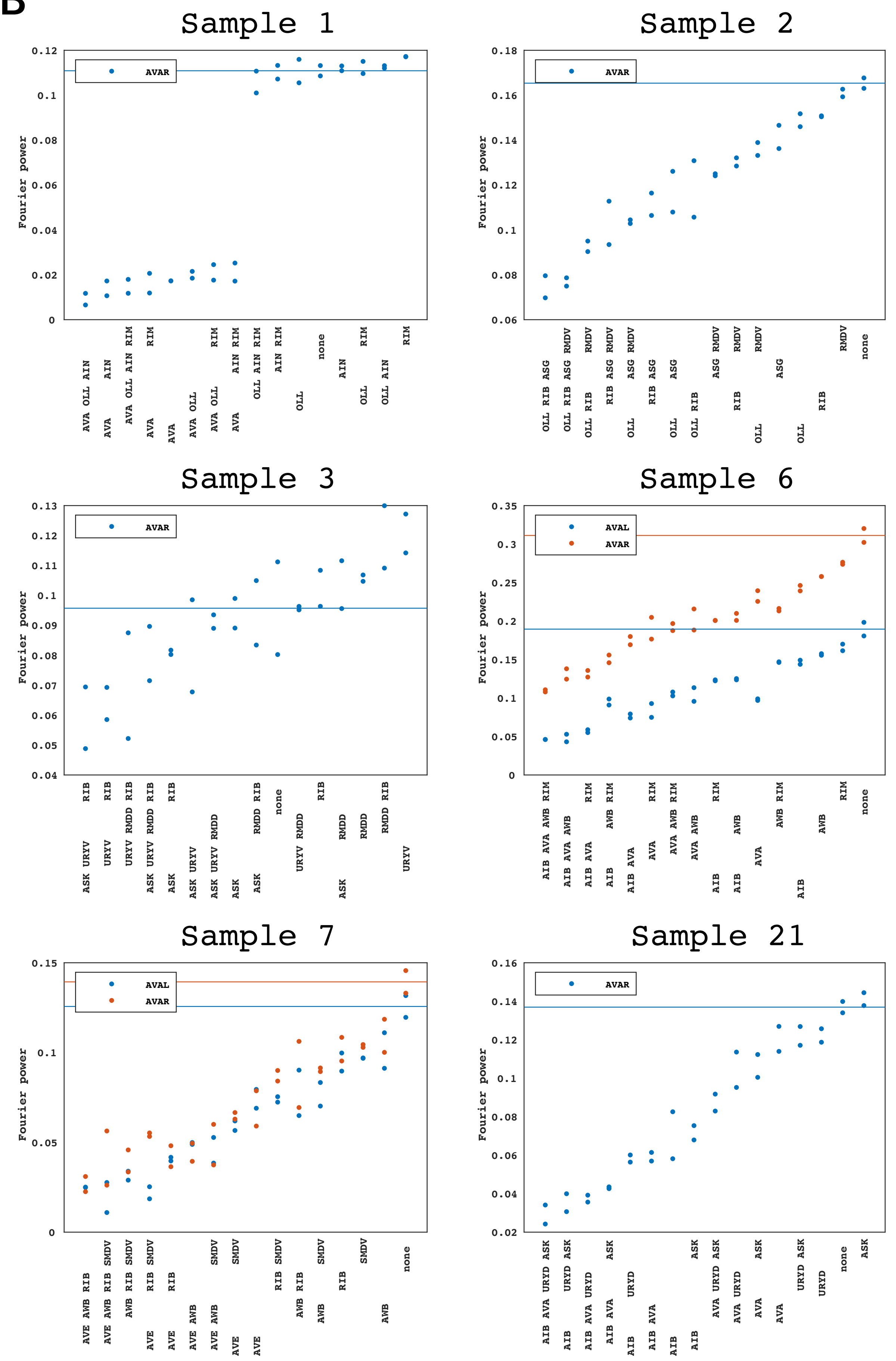


Neurons to which input from salt sensors were blocked
