## Supplementary File 17 for "Deducing ensemble dynamics and information flow from the whole-brain imaging data"

Figure 2: TDE-RICA captures neural motifs

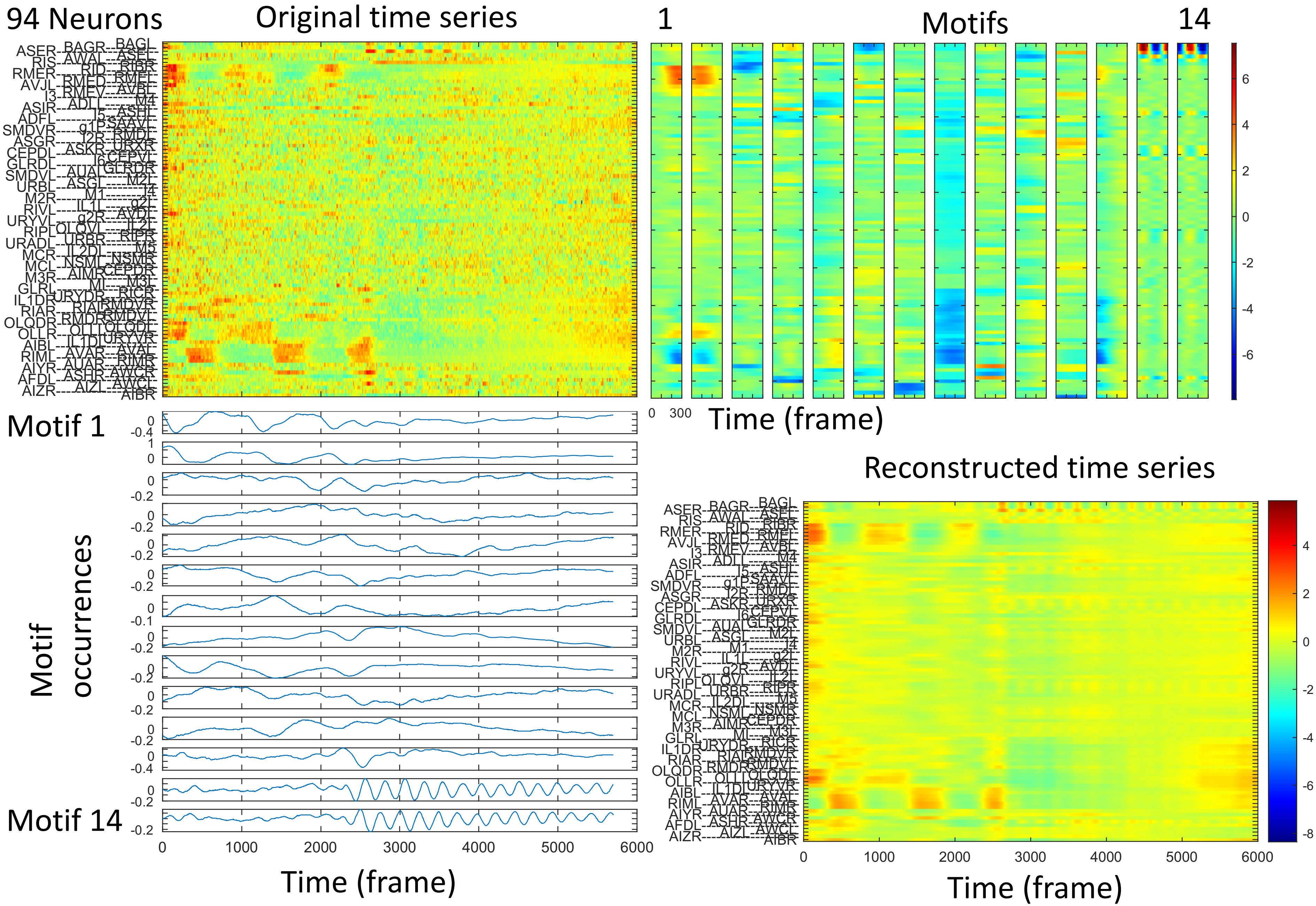

Figure 2-figure supplement 1: completing missing value with TDE-RICA

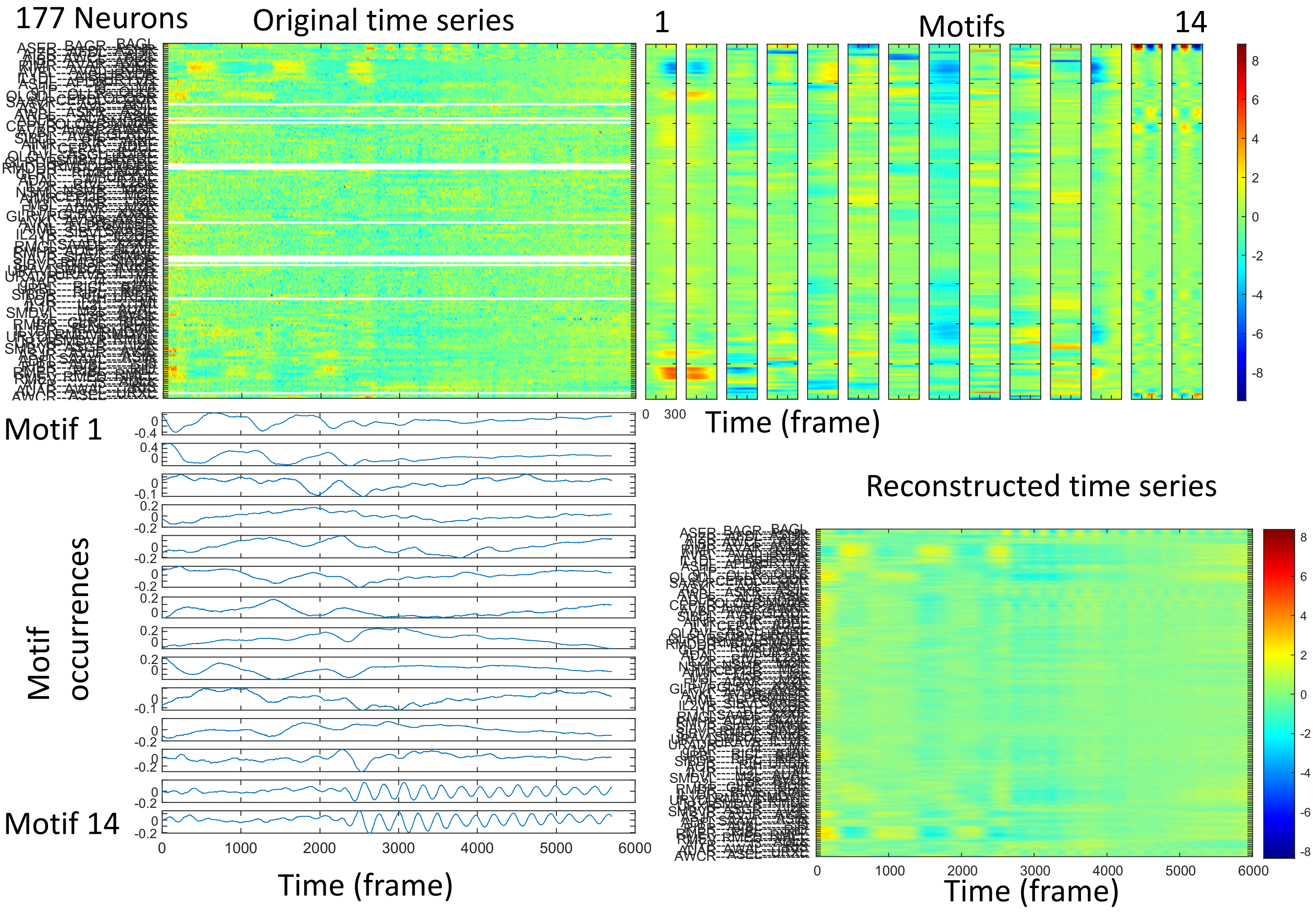

### Figure 3: Neural motifs obtained by TDE-RICA (completed missing value)

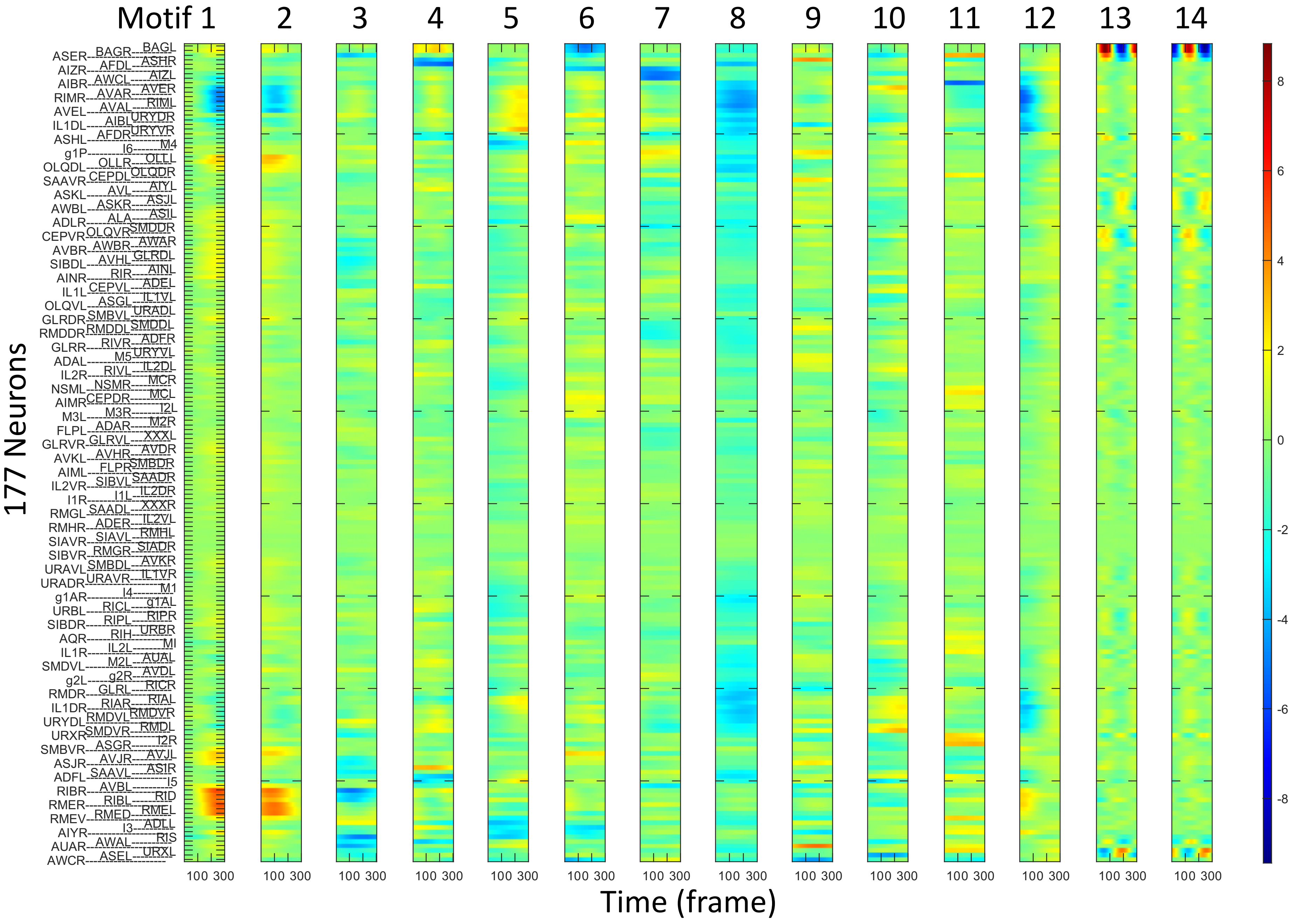

### Figure 4: Cross correlation of motif occurrences

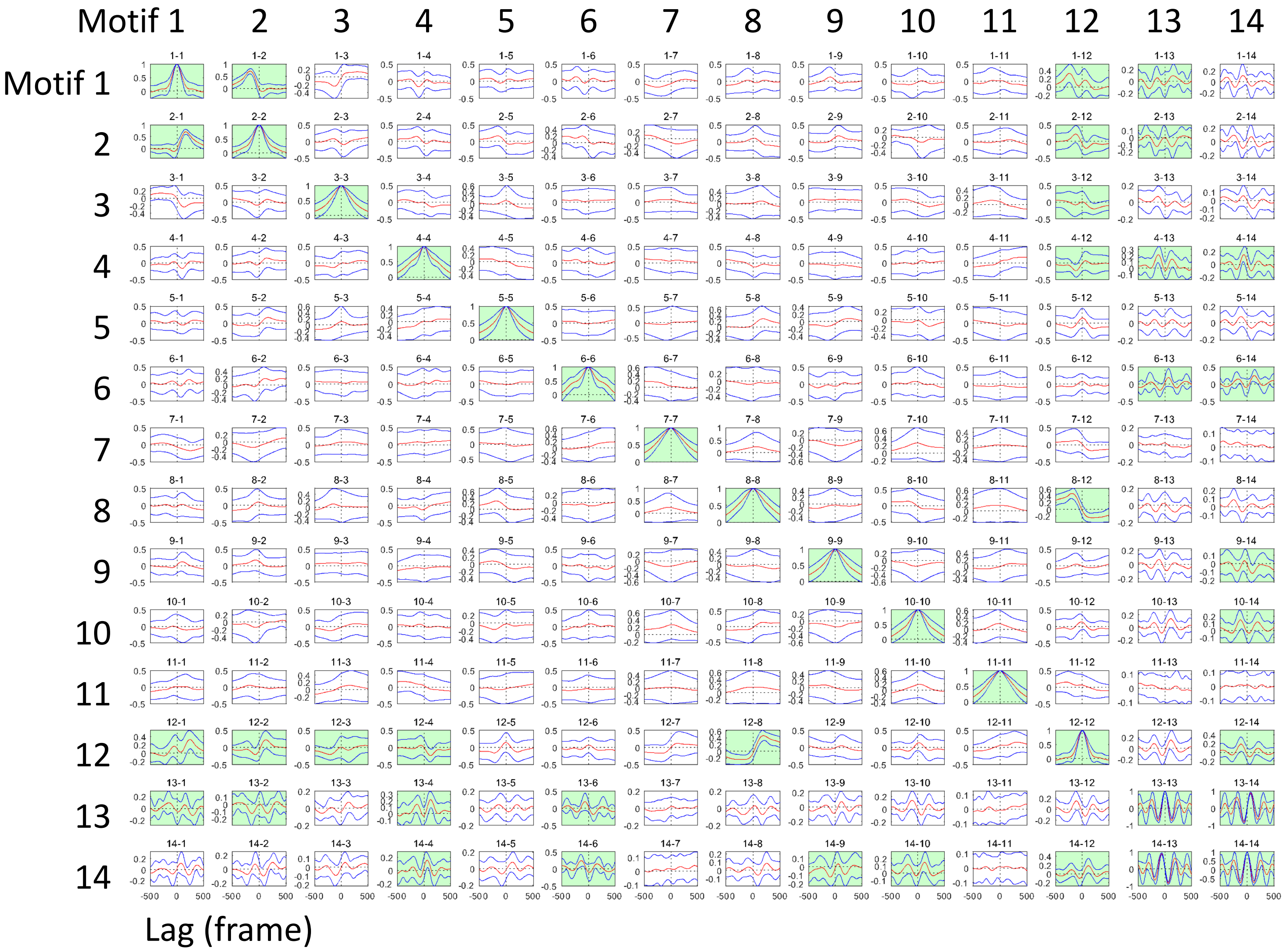

### Figure 5: Common features and individual differences of motif occurrences in the latent space

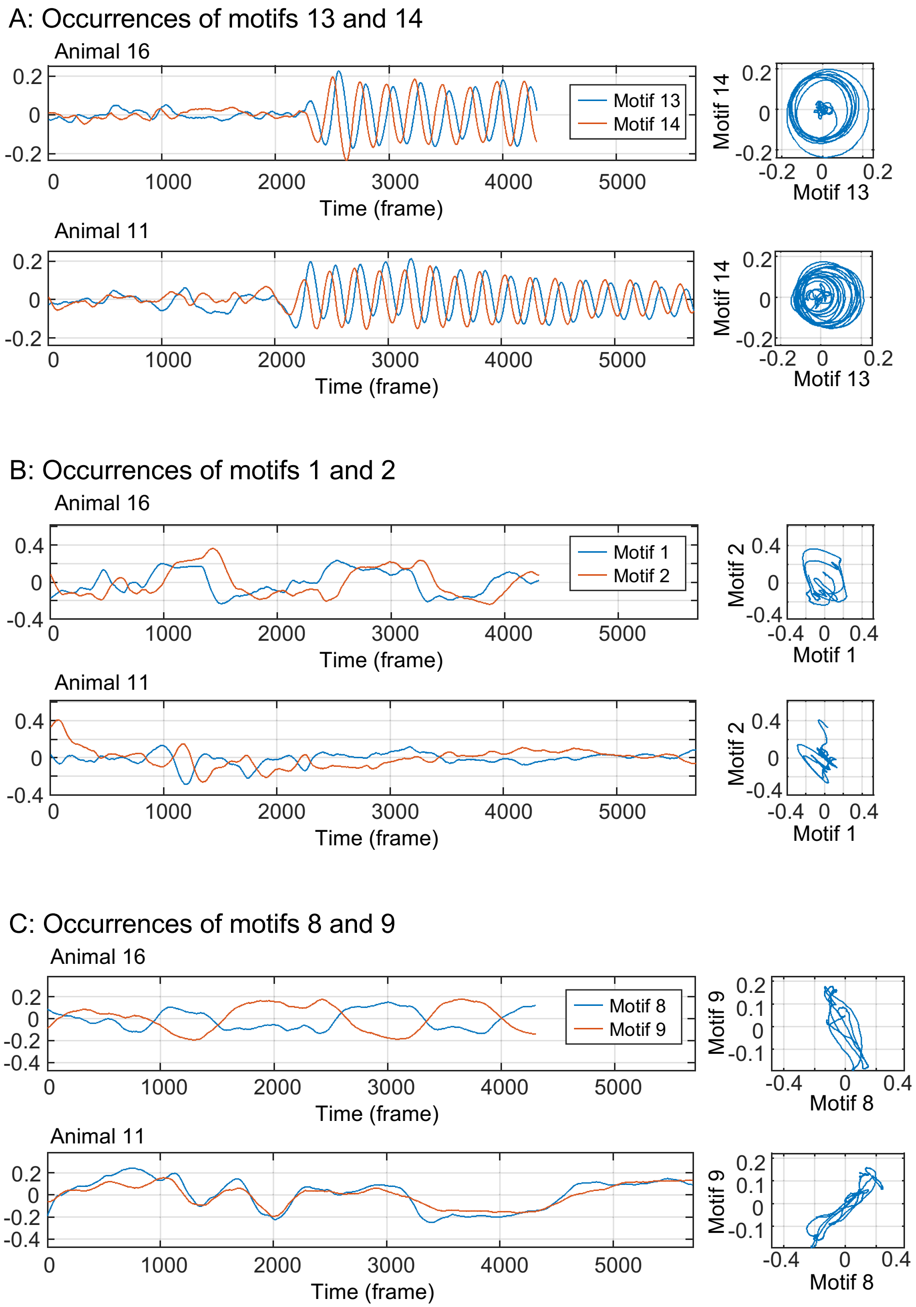

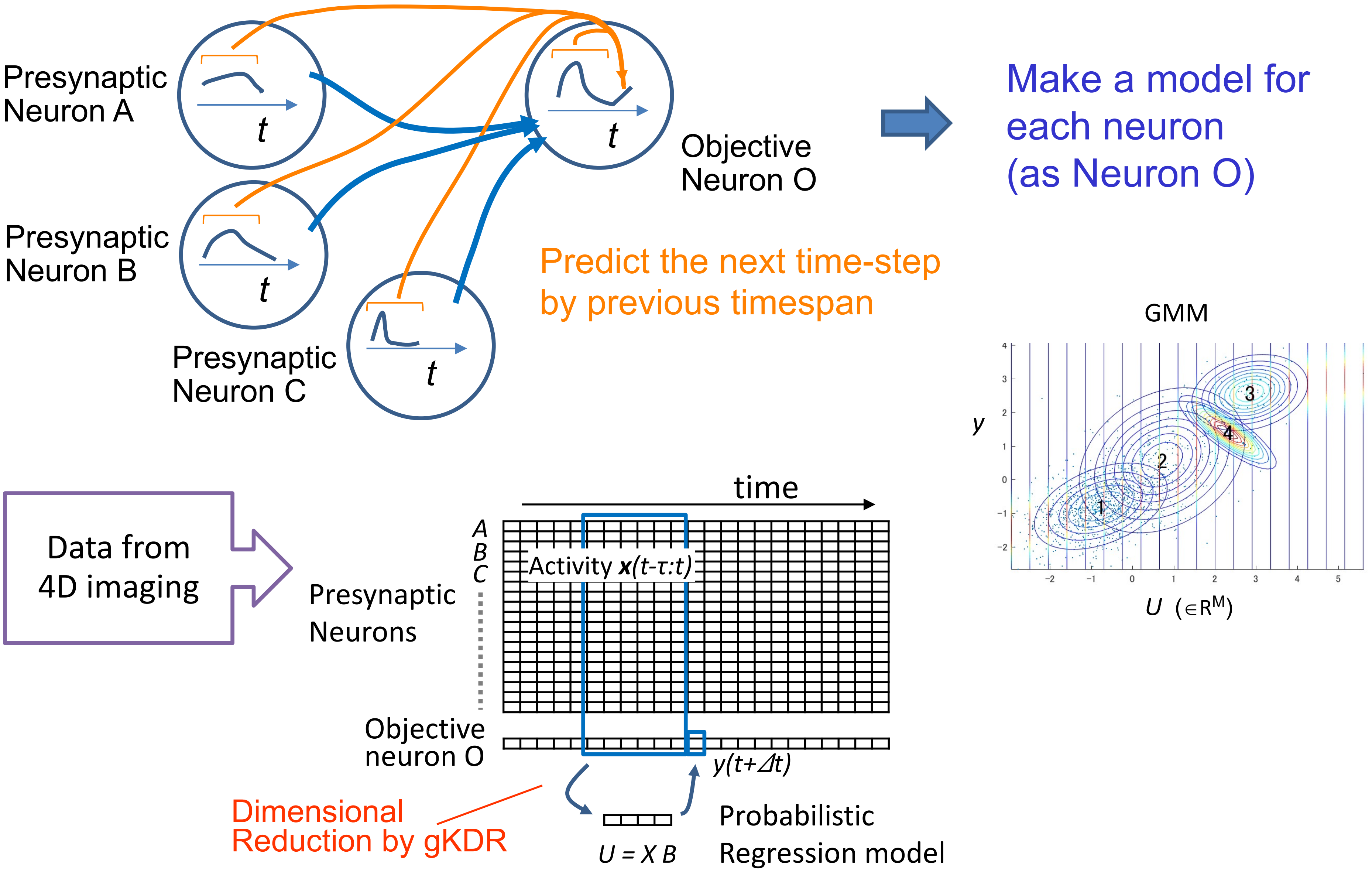

Presynaptic Neuron A
Presynaptic Neuron B
Presynaptic Neuron C
Objective Neuron O
t
Make a model for each neuron (as Neuron O)
Predict the next time-step by previous timespan
GMM
y
U (∈R^M)
Data from 4D imaging
time
A
B
C
Activity x(t-τ:t)
Presynaptic Neurons
Objective neuron O
y(t+Δt)
Dimensional Reduction by gKDR
U = X B
Probabilistic Regression model

Figure 13: Outline of gKDR-GMM

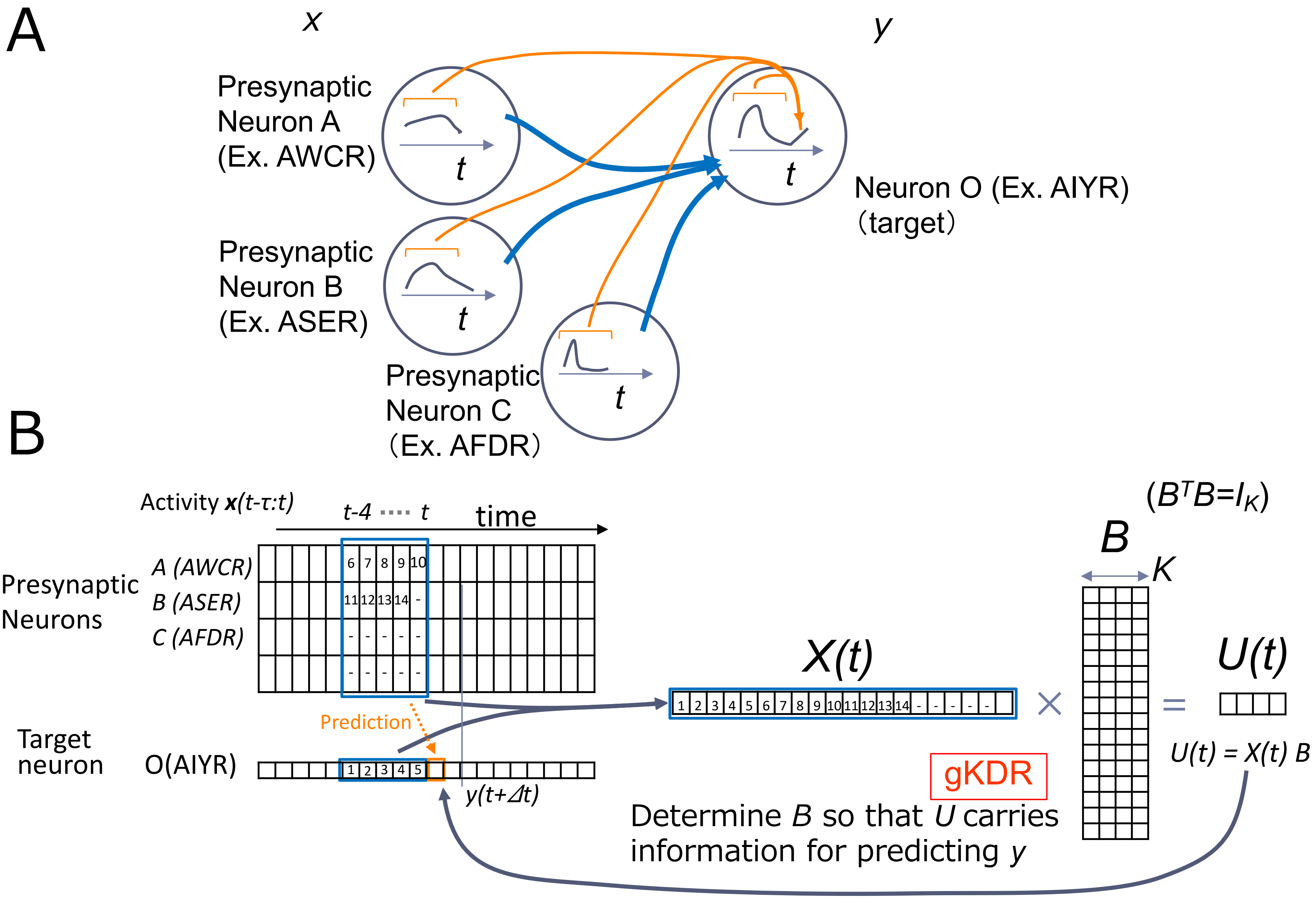

Figure 13: Outline of gKDR-GMM

C

$$y = f(x_1, x_2, x_3)$$

($b_1$, $b_2$: basis vectors)

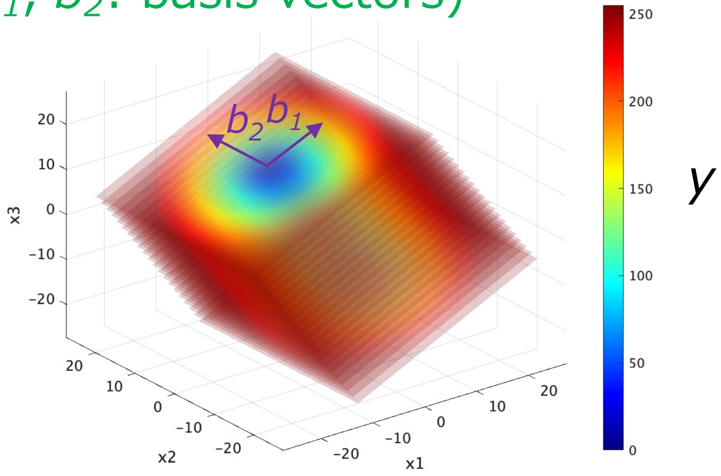


$$y = f(x_1, x_2, x_3) + \xi$$

$$\rightarrow \quad y = g(u_1, u_2) + \xi$$

$$(u_1, u_2) = (x_1, x_2, x_3)\,B$$

$$B = \begin{bmatrix} & b_1 & b_2 \\ & 0.8 & 0 \\ & 7 & \\ & 0 & 1 \\ & 0.5 & 0 \end{bmatrix}$$
